# The Aorta-Gonad-Mesonephros niche shapes the functions of yolk sac-derived macrophages involved in hematopoietic stem and progenitor cell generation *ex vivo*

**DOI:** 10.64898/2026.07.02.736005

**Authors:** Rebecca L. Belmonte, Marianna Romano, Anna Popravko, Amanda MacCallum, Samidha Kulkarni, Malgorzata Rumowska, Cristiana Barone, Alessandro Muratore, Ella Blanks, Hena Modha, Subhankar Mukhopadhyay, Emanuele Azzoni, Siamon Gordon, Samanta A. Mariani

## Abstract

Hematopoietic stem cells (HSCs) generated from induced pluripotent stem cells (iPSCs) offer a promising patient-specific alternative to allogeneic transplantation, yet current differentiation protocols fail to fully recapitulate *in vivo* HSC maturation. During mouse development, yolk sac (YS)-derived macrophages populate the aorta-gonad-mesonephros (AGM) region at the time of HSC emergence, but the mechanisms by which they support *ex vivo* hematopoietic stem and progenitor cell (HSPC) generation remain poorly defined. Bulk RNA sequencing revealed that mature AGM CD206⁺ macrophages upregulate pro-inflammatory cytokines and the adhesion molecule F4/80. Using F4/80 knockout embryos, we identify a previously unreported, niche-specific role for F4/80 in restraining the frequency and colony-forming activity of HSPC subsets in the AGM, while supporting endothelial cell maintenance; this effect was absent in the YS. Lineage-tracing with a Cdh5-CreER^T2^;Rosa26^LSL-tdTomato^ pulse-chase system confirmed that both CD206⁺ and CD206⁻ AGM cells originate from early YS-derived endothelial precursors, with no evidence of local macrophage generation within the AGM. Functional co-culture assays further demonstrated that the ability of CD206⁺ macrophages to enhance the progenitor potential of hemogenic endothelium is AGM-specific and not an intrinsic, ontogeny-determined property, as YS macrophages failed to confer the same benefit even when paired with AGM endothelial cells, and AGM macrophages were ineffective with YS endothelium. Differential expression and NicheNet ligand-receptor interaction analyses identified a small set of AGM-restricted macrophage genes - including *Mmp2, Nrep, Ccl2,* and *Cxcl16* - which are predicted to interact with both endothelial and cluster cells during endothelial-to-hematopoietic transition. Together, these findings establish that AGM macrophages acquire niche-specific transcriptional and functional properties upon entry into the aortic microenvironment, independent of their YS origin, and identify candidate macrophage-derived factors and a novel regulatory role for F4/80 in shaping HSPC output. These insights may guide the refinement of iPSC-based HSC differentiation protocols through the targeted, temporally controlled addition of macrophage-associated signals.

## Introduction

Hematopoietic stem cells (HSCs) are vital components of the hematopoietic system, responsible for the production of all blood cell types throughout the lifetime on an individual. HSCs can be used for hematopoietic stem cell transplantation, a curative therapy for various blood disorders.

HSCs are unique in their ability to self-renew, differentiate, and reconstitute the entire blood system in a recipient after transplantation^1^.

For over six decades, allogeneic HSC transplantation has been employed to treat a range of malignant and non-malignant blood diseases. However, several challenges limit the widespread application of this therapeutic approach. Two major obstacles are the variability in patient response to treatment, and the scarcity of human leukocyte antigen (HLA)-matched donors, particularly for individuals from ethnically diverse backgrounds^2, 3^. The latter can lead to serious consequences such as graft-versus-host disease, a life-threatening complication in which T cells transplanted together with the HSCs recognize the recipient’s body as non-self and mount an immune response against it^4^. The possibility of mitigating transplantation-related side effects by expanding patient-derived functional HSCs *in vitro* is hindered by the difficulty of maintaining and expanding HSCs outside a living organism^5, 6^. As a result, many patients face significant barriers in accessing curative treatments for blood disorders, highlighting the need for innovative solutions to address these challenges.

An alternative approach is to generate HSCs from induced pluripotent stem cells (iPSCs) in a patient-specific manner^2^. Years of research have significantly advanced our knowledge on the topic, with several scientific breakthroughs moving the field from suboptimal early attempts to highly promising iPSC differentiation protocols^7–9^. Recent work from by Ng et al. described a method for generating repopulating iPSC-derived HSCs with high-level multipotent engraftment^9^. However, they reported a 20-fold lower frequency of primary engraftment compared to mobilized CD34^+^ cells, and low levels of secondary engraftment^9^, suggesting that some aspects of *in vivo* HSC generation and maturation remain difficult to recapitulate *in vitro*.

HSCs are generated during embryonic development in both humans and mice^10–13^. They are produced in the aorta-gonad-mesonephros (AGM) region of the embryo during the third definitive wave of developmental hematopoiesis, occurring at embryonic day (E)10.5 in mouse and Carnegie stages (CS)13-17 in humans, through a trans-differentiation process called endothelial-to-hematopoietic transition (EHT)^14–17^. The first two waves preceding HSC generation occur in the extra embryonic yolk sac (YS) at E7.5 and E8.5 in mice, and CS6-CS7 and CS8-CS10 in humans^13, 18^. Moreover, an additional wave of fetal-restricted hematopoietic stem and progenitor cells (HSPCs) has recently been described, emerging from vitelline and umbilical arteries between E8.5 and E9.5 in the mouse embryo^19^.

Both waves of YS hematopoiesis see the production of various progenitors, such as early and late erythroid-myeloid-progenitors (EMPs) which give rise to differentiated cells including macrophages^20–23^.

YS-derived macrophages have well-established functions in tissue specification and remodeling during development^12, 24–26^ Moreover, we and others previously demonstrated that YS-derived macrophages are present in the AGM of mouse and human embryos at the time of HSC generation^27, 28^. However, the mechanism through which macrophages contribute to HSPC generation and/or maturation remains an open question. Understanding which macrophage-related factors can promote HSPC generation and/or maturation is key to informing human iPSC-based differentiation protocols and to more fully recapitulating HSC generation in vitro. Here we demonstrate that macrophages expressing the mannose receptors in the mouse AGM at E10.5 have specific HSPC-promoting functions in an *in vitro* model of endothelial-to-hematopoietic transition. These macrophages, while YS-derived, remodel their transcriptome upon entry into the AGM and acquire niche-specific functions not shared by macrophages present in the YS at the same developmental stage. Transcriptomic analyses revealed that AGM macrophages express a combination of pro-inflammatory genes and metalloproteinases that may mediate their niche-specific functions. Furthermore, we describe a previously unreported role for F4/80 in controlling the frequency of specific HSPC subsets in the mouse AGM.

## Material and Methods

### Timed matings

Wild type C57BL/6 from the University of Edinburgh’s animal facility, as well as *MacGreen (B6N.Cg-Tg(Csf1r-EGFP))*^29^, and *F4/80 knock out (KO)*^30^ mice were used for timed mating experiments. Embryos were generated from *MacGreen* males crossed with wild type females, and *F4/80 KO* males crossed with *F4/80 KO* females. Embryos were staged accurately by counting somite pairs (sp), with 30-34sp corresponding to embryonic day (E) 10 and 35-39sp corresponding to E10.5. The day of vaginal plug detection was designated as E0.5, and plugged females were sacrificed at E10.5 for embryo collection. All mice were housed and bred at the University of Edinburgh’s animal facilities in accordance with Home Office regulations, and all animal experiments were performed under the authority of a UK Home Office Project License and were approved by the University of Edinburgh’s Animal Welfare and Ethical Review Body (AWERB). All procedures were conducted in accordance with the Animals (Scientific Procedures) Act 1986 and the EU Directive 2010/63/EU on the protection of animals used for scientific purposes. For lineage-tracing experiments using the *Cdh5-CreER^T^*^2^ system, pregnant females received a single intraperitoneal (i.p.) injection of 4-hydroxytamoxifen (4-OHT; 37.5 mg/kg body weight) dissolved in corn oil at either E7.5 or E8.5. Embryos were then harvested at E10.5 for analysis. *Cdh5-CreER^T2^* and *Rosa26^LSL-tdTomato^*mice were maintained at the San Raffaele Scientific Institute under standard housing conditions with ad libitum access to food and water. All animal procedures were conducted in accordance with protocols approved by the Institutional Animal Care and Use Committee (IACUC) of the San Raffaele Scientific Institute and by the Italian Ministry of Health.

### Embryo dissection and tissue digestion

E10 and E10.5 embryos were collected in Phosphate-Buffered Saline (PBS, Gibco) supplemented with 10% fetal bovine serum (FBS, Gibco) prior to dissection. The aorta-gonad-mesonephros (AGM) region and the yolk sac (YS) were dissected using a previously described method^31^. Briefly, an incision was made in the lower abdomen of the pregnant females to expose the uterus, and embryos were subsequently collected and dissected with fine-point insulin needles (BD Microfine U-100). The yolk-sac was carefully detached from umbilical and vitelline arteries. To isolate the AGM region, tissues including the head, limbs, intestines, heart, tail, and somites were removed. To obtain a single-cell suspension for use in flow cytometry and functional assays, the dissected YS and AGM tissues were then digested in 0.125% type I collagenase (C0130; Sigma-Aldrich) in PBS + 10% FBS at 37°C for 45 minutes.

### Ex vivo co-culture system and CFU-C assay

Cells from YS and AGM were incubated with TruStain FcX (anti-mouse CD16/32) antibody (BioLegend) in PBS+10% FBS to block non-specific binding, followed by staining with relevant antibodies (Supplementary Table 1) for 30 minutes on ice. Endothelial cells were identified and sorted by fluorescent activated cell sorting (FACS) with FACS Aria II (BD) as CD45^-^CD41^-^CD31^+^. Macrophages were sorted as CD45^+^ CD11b^+^F4/80^+^ CD206^+/-^ and those from *MacGreen* embryos were additionally identified by their expression of Csf1r-GFP. Sorted endothelial cells and macrophages were plated on OP9 stromal cells at a 3:1 ratio in αMEM medium supplemented with cytokines (SCF100ng/ml, IL3 100ng/ml and Flt3-ligand 100ng/ml; Peprotech). After seven days of co-culture, CD45^+^ cells were FACS sorted and plated in methylcellulose (M3434; Stem Cell Technologies) to assess their colony-forming unit (CFU) potential. Colonies were counted and scored after 10 days.

### Immunostaining

E10.5 embryos were fixed in 2% paraformaldehyde (ThermoFisher Scientific) in PBS solution for 30 minutes at 4°C in the dark. This was followed by an overnight incubation in 20% sucrose (Sigma-Aldrich) in PBS solution at 4°C in the dark. The embryos were then embedded in Cryomatrix (Epredia) and frozen at -80°C, with the yolk sac positioned on the side but still attached to the embryo. Frozen embryos were sectioned into 10 μm thick sections using a cryostat (Bright) and collected on Superfrost Plus slides (ThermoFisher Scientific). After thawing, the sections were treated with absolute acetone for 10 minutes and allowed to air dry. The sections were then washed three times with 0.05% Tween 20 (Sigma-Aldrich) in PBS by gentle shaking for 5 minutes. The sections were subsequently incubated with primary antibodies (Supplementary Table 1) in PBS-block buffer (PBS + 0.05% Tween 20 + 1% bovine serum albumin (Sigma-Aldrich)) for 1 hour at room temperature in a humid chamber. This was followed by three washes with PBS-Tween 20 before incubation with secondary antibodies (Supplementary Table 1) in PBS-block for 30 minutes at room temperature. However, for the F4/80 antibody, which was directly conjugated to the BV421 fluorophore, an overnight incubation at -20°C in the humid chamber was performed, omitting the secondary antibody step. After washing, the slides were mounted in Slowfade Diamond mountant (ThermoFisher Scientific) and the coverslips were sealed with clear nail polish and allowed to dry. The slides were stored at 4°C until visualization under a confocal microscope.

To control for non-specific binding, single antibody-stained and secondary antibody-only stained sections were included in each experiment. Immunostained slides were visualized using a Leica SP8 microscope.

### RNA extraction and real time PCR

E10.5 AGM and YS cells were FACS sorted, and RNA was extracted using the RNeasy Plus Micro kit (Qiagen) according to the manufacturer’s instructions. Complementary DNA (cDNA) was synthesized using the SuperScript III Reverse Transcriptase protocol (ThermoFisher Scientific) following the manufacturer’s instructions. Real-time PCR was performed using the SybrGreen master mix (Applied Biosystems), with β-actin serving as the housekeeping gene for ΔCt normalization. The list of primers used is provided in Supplementary Table 2.

### Bulk RNA sequencing preparation

CD206^-^ cells and CD206^+^ macrophages were FACS sorted from E10.5 AGM and YS, and RNA extracted using with the RNeasy Micro kit (Qiagen), according to manufacturer’s instructions. The RNA samples were sent to the BGI (Honk Kong) for library preparation and sequencing using the low-input protocol. The sequencing was performed on three biological replicates, with each sample representing a pool of 4 or 5 embryos.

### RNA sequencing processing

Fastq files of 51 bp single-end reads were quality controlled using FASTQC. Reads were then aligned to the GENCODE M14^32^ mouse reference genome (mm10) using STAR aligner (version 2.3) for Linux Ubuntu, with default options, and ran over 8 threads per sample. For improved accuracy, the exon junction coordinates from the reference annotation were used. Resulting BAM files from different lanes were merged into one sample. Gene quantification on the merged samples was performed using the function “featureCounts” from the subRead package^33^.

### Statistical analysis for differentially expressed genes

All statistical calculations were performed in R programming language (version 4.4.1)^34^. Differential expression analysis was performed using DeSeq2 package (version 1.46.0)^35^ to identify significantly differentially expressed genes with a false discovery rate (likelihood ratio test) of 5% (adjusted p-value > 0.05). Principal component analysis was performed on the variance stabilized matrix of all genes. AGM CD206^+^ enriched genes were identified as genes with a log_2_ fold change greater than |1.5| in comparison to all other cell populations. Genes expressed by endothelial or cluster cells were identified as genes with a log_2_ fold change greater than |0.5|. Normalized counts were calculated using a median of ratios method using the DeSeq2 package and plotted with ggplot2 (version 4.0.0)^36^.

### Interaction analysis

Potential ligands expressed by AGM CD206^+^ cells were identified as genes with a maximum log_2_ fold change greater than 0 and compared with known ligands from the NicheNet (version 2.2.0) network^37^. Potential ligands expressed by AGM CD206^+^ cells interacting with receptors expressed by endothelial or cluster cells were identified by the NicheNet network. Using the genes expressed by endothelial or cluster cells, predicted ligands were determined using the “predict_ligand_activities” function and ligands expressed in AGM CD206^+^ were input as potential ligands. The predicted ligands were ranked according to the area under the precision-recall curve (AUPR). Prior ligand-receptor interaction potential was determined using the “get_weighted_ligand_receptor_links” function.

### Data analysis and statistics

Flow cytometry data were analyzed with FlowJo 10. Graphs were made and statistics calculated with GraphPad Prism 11 (Dotmatics). The statistical tests used are specified in the figure legends.

## Results

### CD206 expression defines mature macrophages in the AGM

We previously identified two main populations of myeloid cells in the mouse AGM at the time of HSC generation, distinguished by expression of CD206 (mannose receptor)^28^. CD206^+^ macrophages were found to be important for the correct *ex vivo* generation and/or maturation of mouse HSCs, while the role of CD206^-^ cells in supporting hematopoiesis was confounded by their residual multipotent progenitor potential in colony-forming unit (CFU-C) assays^28^. To determine whether a more refined sorting strategy would help define the composition and function of AGM CD206⁻ myeloid cells, additional flow cytometry analyses were performed. Using the *MacGreen* mouse model^29^, in which GFP expression is driven by the colony-stimulating factor 1 receptor (Csf1r) promoter, flow cytometry revealed that 9±1.2% of E10.5 AGM CD45⁺GFP⁺CD206⁻ cells express low levels of CD11b and F4/80 and retain cKit expression (Fig. 1A), indicating the presence of myeloid progenitors and explaining the residual CFU-C activity previously observed^28^. In contrast, the CD206⁺ population was confirmed to consist of terminally differentiated macrophages, with only 1±0.4% of cells retaining cKit expression at E10.5 (Fig. 1A). These findings suggested that a more comprehensive sorting strategy incorporating CD11b, F4/80, and cKit is needed to discriminate AGM CD206⁻ myeloid progenitors from CD206⁻ macrophages (Fig. 1A) and to assess the role of the latter in embryonic hematopoiesis.

**Figure 1:**
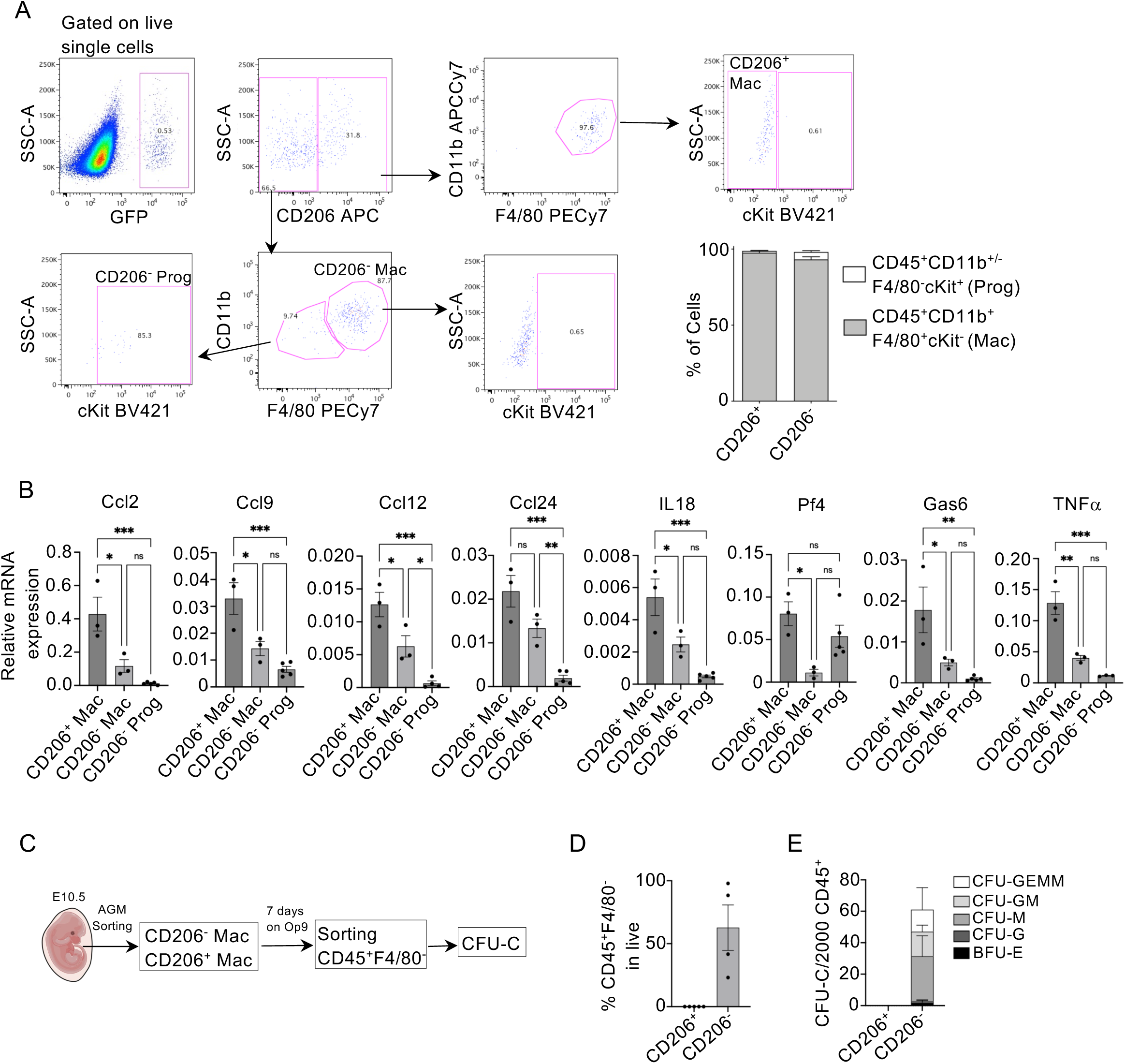
A) Representative pseudocolor dotplots showing the fluorescent activated cell sorting gating strategy for CD206^+^ macrophages, and CD206^-^ macrophages and progenitors in the Aorta-Gonad-Mesonephros (AGM) of embryonic day (E)10.5 *MacGreen* embryos. The improved sorting strategy shows how CD206^+^ and CD206^-^ populations are resolved according to CD45, F4/80 and cKit expression. Arrows are used to show gate dependency. Bottom right, bar chart of percentage of cells shows proportion of progenitors (CD45^+^CD11b^+/-^F4/80^-^cKit^+^) and macrophages (CD45^+^CD11b^+^ F4/80^+^cKit^-^) in CD206^+^ or CD206^-^ AGM populations. B) Real time qPCR analysis of *Ccl2*, *Ccl9*, *Ccl12*, *Ccl24*, interleukin 18 (*Il18)*, platelet factor 4 (*Pf4)*, growth arrest-specific 6 (*Gas6)*, and tumor necrosis factor α (*Tnfα)* expression in CD206^+^ macrophages, CD206^-^ macrophages, and CD206^-^ progenitors, normalized over β *Actin* expression. N = 3, with each N representing 4-to-6 pulled AGM from E10.5 *MacGreen* embryos. Statistical test: Ordinary One Way Anova with Tukey’s multiple comparison test. ns= not significant, **=p<0.01, ***=p<0.001. C) Schematic representation of co-culture experiments on OP9 stromal cells. CD206^+^ (CD45^+^cKit^-^CD11b^+^F4/80^+^CD206^+^) and CD206^-^ (CD45^+^cKit^-^CD11b^+^F4/80^+^CD206^-^) macrophages were sorted from the AGM of E10.5 *MacGreen* embryos and plated on OP9 cells for seven days. Nascent CD45^+^F4/80^-^ hematopoietic cells were subsequently sorted and plated in methylcellulose for a CFU-C (colony forming unit in culture) assay. D) Bar graph showing the frequency of hematopoietic CD45^+^F4/80^-^ cells produced by AGM CD206^+^ and CD206^-^ cells after one week of culture on OP9 stromal cells. Each dot represents an independent biological replicate, with 4-to-7 embryos pulled for each experiment. E) Bar graph showing the number of hematopoietic colonies produced by AGM CD45^+^F4/80^-^ cells retrieved after OP9 culture. N=4, with each N representing 4-to-7 pulled embryos. CFU-GEMM= colony forming unit-granulocyte, erythroid, macrophage, megakaryocyte; CFU-GM= colony forming unit-granulocyte, macrophage; CFU-M= colony forming unit-macrophage; CFU-G= colony forming unit-granulocyte; BFU-E= burst forming unit-erythroid.

We previously observed that *MacGreen*-derived CD206^+^ macrophages express a higher level of pro-inflammatory cytokines as compared to CD45^+^GFP^+^CD206^-^ cells^28^. This was confirmed when CD45^+^GFP^+^CD11b^-^F4/80^-^cKit^+^CD206^-^ progenitors (CD206^-^ Prog) were sorted separately from CD45^+^GFP^+^CD11b^+^F4/80^+^cKit^-^CD206^-^ cells (hereafter referred to as CD206^-^ Mac) from E10.5 *MacGreen* AGMs. CD206+ Mac expressed significantly higher levels of *CCl2, Ccl9, Ccl12, IL18, Pf4, Gas6* and *TNF*α as compared to CD206^-^ Mac and CD206^-^ Prog (Fig. 1B). This analysis further revealed that CD206⁻ Mac more closely resemble myeloid progenitors than terminally differentiated macrophages with respect to pro-inflammatory cytokine production (Fig. 1B), suggesting that they may represent a differentiation intermediate between CD206⁻ Prog and CD206⁺ macrophages, rather than a distinct AGM macrophage subset. To test our hypothesis, AGM CD206^-^ and CD206^+^ Mac were sorted according to the refined strategy shown in Fig. 1A, and plated on OP9 stromal cells^38^ for seven days. CD45^+^F4/80^-^ cells were subsequently sorted and plated in methylcellulose to assess progenitor potential (Fig 1C). Notably, 62.7±35% of cells recovered from wells containing CD206^-^ Mac were not mature macrophages (CD45^+^F4/80^-^) (Fig. 1D), and retained the capacity to generate multiple types of progenitor colonies in methylcellulose (Fig. 1E). As a control, the CD206^+^ subset did not give rise non-macrophage cells and failed to generate CFU-Cs (Fig. 1D-E), consistent with previous findings^28^. Together, these results show that CD206 expression defines mature macrophages in the mouse AGM, while CD206^-^ myeloid cells retain progenitor potential even when expressing CD11b and F4/80 and in the absence of cKit.

### F4/80 expression on AGM macrophages regulates the frequency and the colony forming potential of hematopoietic progenitors

To identify the key features of HSPC-supporting mature AGM macrophages, bulk RNA sequencing (RNAseq) was performed on CD206⁺ Mac and CD206⁻ Mac isolated from E10.5 *MacGreen* embryos according to the refined sorting strategy shown in Figure 1A (Supplementary Fig. 1A). Principal component analysis revealed marked differences between CD206⁺ Mac and CD206⁻ cells, with one CD206⁺ biological replicate behaving as an outlier (Supplementary Fig. 1B). A total of 667 genes were found to be differentially expressed between the two cell subsets (Supplementary Table 3), with CD206⁺ Mac preferentially expressing genes associated with chemotaxis, migration, and endocytosis, as shown by gene ontology analysis (Supplementary Fig. 1C). Notably, RNAseq data also revealed increased *Adgre1* expression (encoding F4/80) in CD206⁺ Mac (Supplementary Fig. 1D). This was confirmed by flow cytometry, which showed higher mean fluorescence intensity for F4/80 in the CD206⁺ Mac subset (Fig. 2A–B). F4/80 is an adhesion molecule used as a pan-macrophage marker in mice, but its role during embryonic development remains poorly understood^39^. To determine whether F4/80 mediates the HSPC-supporting functions of CD206⁺ Mac in the mouse AGM, we characterized the effect of its absence on the endothelial and hematopoietic compartments of the AGM in E10.5 F4/80 knockout (KO) embryos^30^, compared to C57BL/6J wild-type (WT) controls, by flow cytometry. To avoid confounding effects from progenitor and endothelial cell populations that might express F4/80, analysis was restricted to F4/80⁻ cells (Supplementary Fig. 2A–B). The frequency of endothelial cells, defined as CD31⁺cKit⁻CD41⁻CD43⁻CD45⁻, was lower in F4/80 KO embryos compared to WT controls (Fig. 2C and Supplementary Fig. 2A), suggesting a role for F4/80 in the generation and/or maintenance of AGM endothelial cells. Conversely, the frequency of cKit⁺CD31⁻ HSPCs was higher in F4/80 KO embryos (Fig. 2D and Supplementary Fig. 2A), whereas no differences were observed in the frequency of cKit⁺CD31⁺ cluster cells, cKit⁺CD31⁺CD41ˡᵒ cells, or cKit⁺CD31⁺CD45⁺ cells (Fig. 2E). Furthermore, while the frequency of the CD31⁻cKit⁺/⁻CD41ˡᵒ population was unchanged in F4/80 KO embryos (Fig. 2F and Supplementary Fig. 2B), the more differentiated CD31⁻cKit⁺/⁻CD41ˡᵒCD43⁺ and CD31⁻cKit⁺/⁻CD41ˡᵒCD43⁺CD45⁺ progenitor populations were increased compared to WT controls (Fig. 2G and Supplementary Fig. 2B). To confirm that the observed changes in progenitor population frequency were attributable to the absence of F4/80 rather than differences in cell viability or CD206 expression, we measured the percentage of live cells and the frequency and mean fluorescence intensity of CD206 in CD45⁺CD11b⁺ cells isolated from the AGM of C57BL/6 and F4/80 KO embryos. No differences were observed in live cell frequency or CD206 expression compared to C57BL/6 controls (Supplementary Fig. 2C–F). To determine whether the higher hematopoietic progenitor frequency in F4/80 KO embryos reflects a difference in progenitor potential, a CFU-C assay was performed on cells isolated from the AGM of E10.5 F4/80 KO and C57BL/6 embryos. Colonies were counted and scored after ten days; a significant increase in total colony number was observed in F4/80 KO embryos, with no differences in the relative representation of individual colony types (Fig. 2H). Taken together, these data suggest that F4/80 expression on AGM macrophages suppresses the frequency and colony-forming potential of hematopoietic progenitors in the AGM, while promoting an increase in endothelial cells. Transplantation experiments will be needed to determine whether the frequency and/or potency of HSCs is also altered in F4/80 KO embryos.

**Figure 2:**
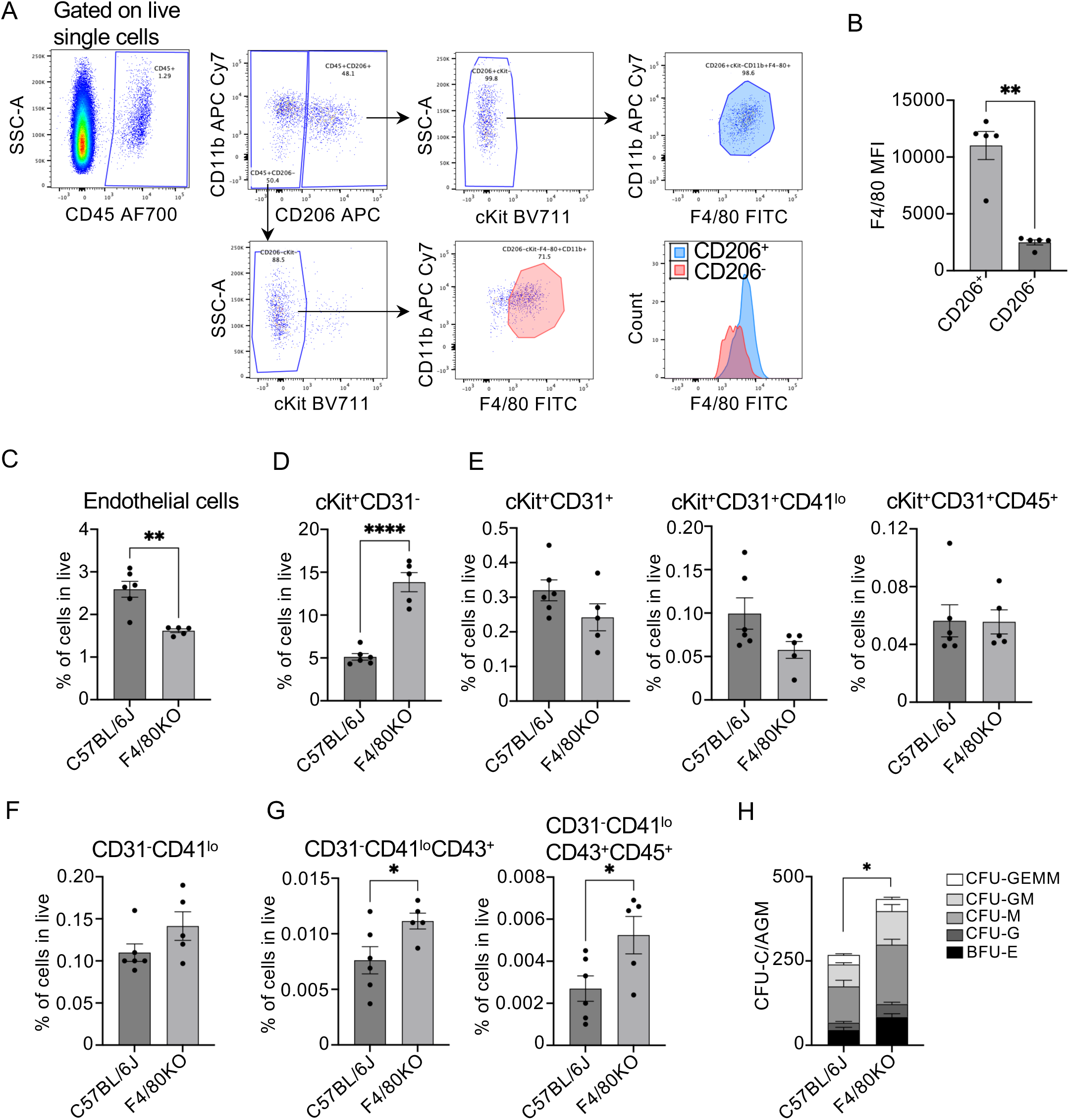
A) Representative flow cytometry plots showing F4/80 expression in CD45^+^GFP^+^cKit^-^CD11b^+^F4/80^+^CD206^+^ macrophages (blue) and CD45^+^GFP^+^cKit^-^CD11b^+^F4/80^+^ CD206^-^ cells (red) from the Aorta-Gonad-Mesonephros (AGM) of embryonic day 10.5 *MacGreen* embryos. B) Bar graph showing the quantification of F4/80 mean fluorescent intensity (MFI) in AGM CD45^+^cKit^-^CD11b^+^F4/80^+^CD206^+^ macrophages and CD45^+^cKit^-^CD11b^+^F4/80^+^ CD206^-^ cells from E10.5 *MacGreen* embryos. Each dot represents one individual embryo. C-G) Bar graphs showing the quantification of the frequency of C) endothelial cells (CD31^+^cKit^-^CD41^-^CD43^-^CD45^-^), D) cKit^+^CD31^-^ cells, E) cKit^+^CD31^+^ cluster cells, cKit^+^CD31^+^CD41^lo^ and cKit^+^CD31^+^CD45^+^ hematopoietic progenitors, F) CD31^-^CD41^lo^ and G) CD31^-^CD41^lo^CD43^+^ and CD31^-^CD41^lo^CD43^+^CD45^+^ hematopoietic progenitors in the AGM of E10.5 C57BL/6J and F4/80 KO embryos. Each dot represents one individual embryo. H) Bar graph showing the number of hematopoietic colonies produced by AGM cells from E10.5 C57BL/6J and F4/80 KO embryos. N=5. CFU-GEMM= colony forming unit-granulocyte, erythroid, macrophage, megakaryocyte; CFU-GM= colony forming unit-granulocyte, macrophage; CFU-M= colony forming unit-macrophage; CFU-G= colony forming unit-granulocyte; BFU-E= burst forming unit-erythroid. B,D,G,H) Statistical analysis: Student’s t-test. *=p<0.05, **=p<0.01, ****=p<0.0001. Only statistically significant differences are indicated.

### AGM E10.5 CD206^+^ macrophages and CD206^-^ cells are yolk sac-derived

It has been previously reported that macrophages in the mouse embryos are yolk sac (YS)-derived^28, 40^. However, the possibility of *de novo* macrophage generation in the AGM has never been formally excluded. To address this, we employed a well-validated pulse-chase approach in which *Cdh5-CreER^T2^*mice were crossed with *Rosa26^LSL-tdTomato^* mice to generate embryos in which tdTomato labelling was induced by administering 4-hydroxytamoxifen (4-OHT) to pregnant dams at E7.5 and E8.5^19, 41, 42^. At these time points, YS progenitors derived from Cdh5⁺ hemogenic endothelium are preferentially labelled. Pulsed embryos were then collected at E10.5 to assess tdTomato expression in AGM CD206⁺ Mac and CD206⁻ cells. Between 96–100% of cells in both populations were tdTomato⁺ following pulse labelling at E7.5, with more variable tdTomato expression (36–100%) when pulse-labelled at E8.5 (Supplementary Fig. 3A–B). Normalization against tdTomato expression in YS erythro-myeloid progenitors (EMPs), used as a control to account for variability in labelling efficiency across experiments, consistently showed equivalent labelling levels in AGM CD206⁺ macrophages, CD206⁻ cells, and YS EMPs upon pulse labelling at E7.5 (Supplementary Fig. 3C). The ratios were still high but more variable (between 0.8 to 1) when cells were pulse-labelled at E8.5 (Supplementary Fig. 3C). These results suggest that both AGM CD206⁺ macrophages and CD206⁻ cells present at E10.5 derive from early YS EMPs, with no evidence of *de novo* macrophage generation within the AGM, up until this developmental stage. Furthermore, the absence of any difference in labelling frequencies between CD206⁺ macrophages and CD206⁻ cells corroborates our hypothesis that these two cell subsets represent distinct maturation states of the same myeloid population.

### CD206^+^ and CD206^-^ myeloid cells are present also in the yolk sac at E10.5 in the mouse embryo

To determine whether the phenotype and function of YS-derived AGM CD206⁺ Mac are ontogeny-related or niche-specific, we first investigated whether macrophages in the YS have similar spatial distribution and phenotype to those in the AGM. Immunofluorescence experiments revealed that F4/80⁺ macrophages localize in proximity to cKit⁺ cells in the YS (Fig. 3A), as previously reported for the AGM^28^. The distribution of CD206⁺ and CD206⁻ cells within the CD45⁺CD11b⁺F4/80⁺GFP⁺ compartment of the YS was similar to that observed in the AGM, but with a higher proportion of immature cKit⁺ progenitors (7.5 ± 3.2% in the CD206⁺ population and 27.3 ± 7.0% in the CD206⁻ population) compared to the AGM (Fig. 3B and Supplementary Fig. 3D). An increase in F4/80 mean fluorescence intensity was also observed in YS CD206⁺ cells compared to their CD206⁻ counterparts (Fig. 3C). To determine whether CD206⁺ and CD206⁻ cells differ in their capacity to generate hematopoietic colonies, both populations were cultured on stromal cells (experimental schematic as in Fig. 1C), and the frequency of CD45⁺F4/80⁻ cells arising after one week was assessed. Unlike AGM cells, YS CD206⁺ Mac were also able to produce CD45⁺F4/80⁻ cells (Fig. 3D), suggesting that residual progenitor potential is retained in this population in the YS. However, no hematopoietic colonies were generated by CD206⁺ Mac (Fig. 3E). In contrast, YS CD206⁻ cells produced both CD45⁺F4/80⁻ cells and a variety of hematopoietic colonies after one week on OP9 cells (Fig. 3D–E). Notably, unlike CD206⁻ cells in the AGM, YS CD206⁻ cells did not give rise to CFU-GEMM (granulocyte, erythroid, megakaryocyte, macrophage) colonies, suggesting that CD206⁻ cells in the YS are less multipotent than their AGM counterparts (compare Fig. 1E with Fig. 3E). When assessing pro-inflammatory cytokine expression, YS CD206⁺ Mac were found to express lower levels of *Ccl2, Ccl9, Ccl12, Pf4,* and *Tnfα* compared to AGM CD206⁺ Mac (Fig. 3F). Notably, while *Ccl*2 and *Tnfα* expression was upregulated also in AGM CD206⁻ cells compared to YS CD206⁻ cells — albeit to a lesser extent than in CD206⁺ macrophages — other mediators such as *Ccl24, Il18, Pf4,* and *Gas6* were expressed at lower levels in AGM CD206⁻ cells relative to YS CD206⁻ cells (Supplementary Fig. 3E), suggesting that macrophages maturing in the AGM acquire a more pro-inflammatory phenotype than those maturing in the YS.

**Figure 3:**
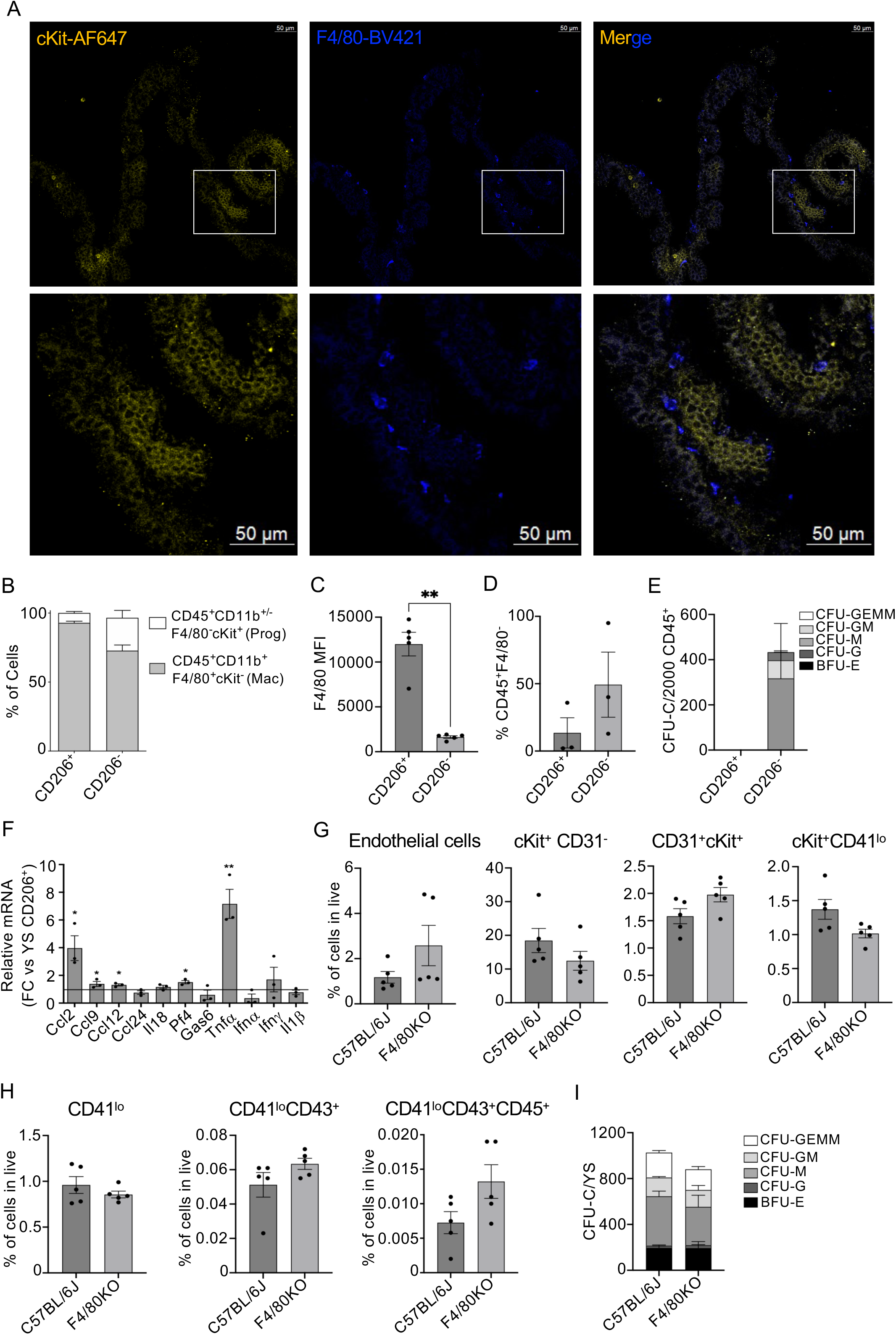
A) Representative immunofluorescence images showing localization of F4/80^+^ (blue) and cKit^+^ (yellow) cells in the yolk sac (YS) of embryonic day (E)10.5 C57BL/6J embryos. B) bar chart of percentage of cells shows proportion of progenitors (CD45^+^CD11b^+/-^F4/80^-^cKit^+^) and macrophages (CD45^+^CD11b^+^F4/80^+^cKit^-^) in YS CD206^+^ or CD206^-^ populations. C) Bar graph showing the quantification of F4/80 mean fluorescent intensity (MFI) in YS CD45^+^cKit^-^CD11b^+^F4/80^+^CD206^+^ macrophages and CD45^+^cKit^-^CD11b^+^F4/80^+^ CD206^-^ cells from E10.5 *MacGreen* embryos. Each dot represents one individual embryo. D) Bar graph showing the frequency of hematopoietic CD45^+^F4/80^-^ cells produced by YS CD45^+^cKit^-^CD11b^+^F4/80^+^CD206^+^ macrophages and CD45^+^cKit^-^CD11b^+^F4/80^+^ CD206^-^ after one week of culture on OP9 stromal cells (see experiment schematics in Fig. 2C). Each dot represents an independent biological replicate, with 4-to-7 embryos pulled for each experiment. E) Bar graph showing the number of hematopoietic colonies produced by yolk sac CD45^+^F4/80^-^ cells retrieved after OP9 culture. N=3. CFU-GEMM= colony forming unit-granulocyte, erythroid, macrophage, megakaryocyte; CFU-GM= colony forming unit-granulocyte, macrophage; CFU-M= colony forming unit-macrophage; CFU-G= colony forming unit-granulocyte; BFU-E= burst forming unit-erythroid. F) Real time qPCR analysis of *Ccl2*, *Ccl9*, *Ccl12*, *Ccl24*, interleukin 18 (*Il18)*, platelet factor 4 (*Pf4)*, growth arrest-specific 6 (*Gas6)*, tumor necrosis factor α (*Tnfα)*, interferon α (*Ifnα*), interferon ψ (*Ifnψ*) and interleukin 1β (*Il1* β) expression in AGM CD206^+^ macrophages (columns) normalized over β *Actin* expression and then plotted as fold change over the expression in YS CD206^+^ macrophages (horizontal line). Each dot represents an independent biological replicate, with 4-to-6 embryos per replicate. G-H) Bar graphs showing the quantification of the frequency of G) endothelial cells (CD31^+^cKit^-^CD41^-^CD43^-^CD45^-^), cKit^+^CD31^-^cells, cKit^+^CD31^+^ cluster cells, and cKit^+^CD31^+^CD41^lo^ hematopoietic progenitors, H) CD31^-^CD41^lo^, CD31^-^CD41^lo^CD43^+^ and CD31^-^CD41^lo^CD43^+^CD45^+^ hematopoietic progenitors in the YS of E10.5 C57BL/6J and F4/80 KO embryos. Each dot represents one individual embryo. I) Bar graph showing the number of hematopoietic colonies produced by YS cells from E10.5 C57BL/6J and F4/80 KO embryos. CFU-GEMM= colony forming unit-granulocyte, erythroid, macrophage, megakaryocyte; CFU-GM= colony forming unit-granulocyte, macrophage; CFU-M= colony forming unit-macrophage; CFU-G= colony forming unit-granulocyte; BFU-E= burst forming unit-erythroid. C,F) Statistical test: Student’s t-test. *=p<0.05, **=p<0.01. Only statistically significant differences are indicated.

### F4/80 expression does not influence the activity of mature CD206^+^ macrophages in the yolk sac

Since, similarly to the AGM, we observed an increase in F4/80 MFI in CD206⁺ YS macrophages compared to CD206⁻ cells (Fig. 3C), we assessed whether its absence would affect the frequencies of endothelial cells and hematopoietic progenitors in the YS, as previously observed in the AGM. The frequencies of CD31⁺cKit⁻CD41⁻CD43⁻CD45⁻ endothelial cells, cKit⁺ and CD41ˡᵒ hematopoietic progenitors were determined by flow cytometry in the YS of F4/80 KO and C57BL/6J control embryos. No statistically significant differences were observed in any endothelial or progenitor populations in the YS of F4/80 KO embryos compared to age-matched controls (Fig. 3G–H). Furthermore, unlike in the AGM, the number of hematopoietic colonies produced by F4/80 KO YS cells was comparable to that of C57BL/6J controls (Fig. 3I). Taken together, these data indicate that the effect of F4/80 on endothelial cells and hematopoietic progenitors is AGM-specific.

### The ability of CD206⁺ macrophages to promote haemogenic endothelial cell progenitor potential is AGM-specific

Having established that the CD45^+^CD11b^+^F4/80^+^GFP^+^cKit^-^CD206^-^ population in both E10.5 YS and AGM is heterogenous and not enriched for macrophages, we proceeded to assess only the potential of YS and AGM CD206^+^ Mac to enhance HSPC generation *in vitro*, in order to determine whether their functions are ontogeny-related or niche-specific. We previously established an *in vitro* co-culture protocol to measure the effect of macrophages on the progenitor potential of hemogenic endothelial cells^28^. Briefly, CD31⁺CD41⁻CD45⁻ endothelial cells were sorted from the AGM and YS of E10.5 *MacGreen* embryos and cultured on OP9 stromal cells in the presence or absence of CD206⁺ macrophages sorted from the same tissues (Fig. 4A). After 7 days, CD45⁺F4/80⁻ hematopoietic cells were collected as a readout of *in vitro* endothelial-to-hematopoietic transition (Fig. 4A). The presence of CD206⁺ Mac did not affect the frequency of CD45⁺F4/80⁻ cells produced by AGM or YS endothelial cells, with AGM endothelial cells consistently generating more hematopoietic cells than YS endothelial cells (Fig. 4B). To assess how the presence of CD206⁺ macrophages affects the progenitor potential of hematopoietic cells produced during co-culture, CD45⁺F4/80⁻ cells were plated in methylcellulose for a CFU-C assay (Fig. 4A). Using the refined macrophage gating strategy (CD45⁺CD11b⁺F4/80⁺GFP⁺cKit⁻CD206⁺), we were able to reproduce previously published data demonstrating an increase in CFU-Cs in the presence of AGM CD206⁺ macrophages (Fig. 4C left and^28^). However, no such increase was observed in the number of colonies produced by YS CD45⁺F4/80⁻ cells following co-culture with YS CD206⁺ macrophages (Fig. 4C, right), suggesting that the pro-hematopoietic function of CD206⁺ macrophages is AGM-specific and not ontogeny-related.

**Figure 4:**
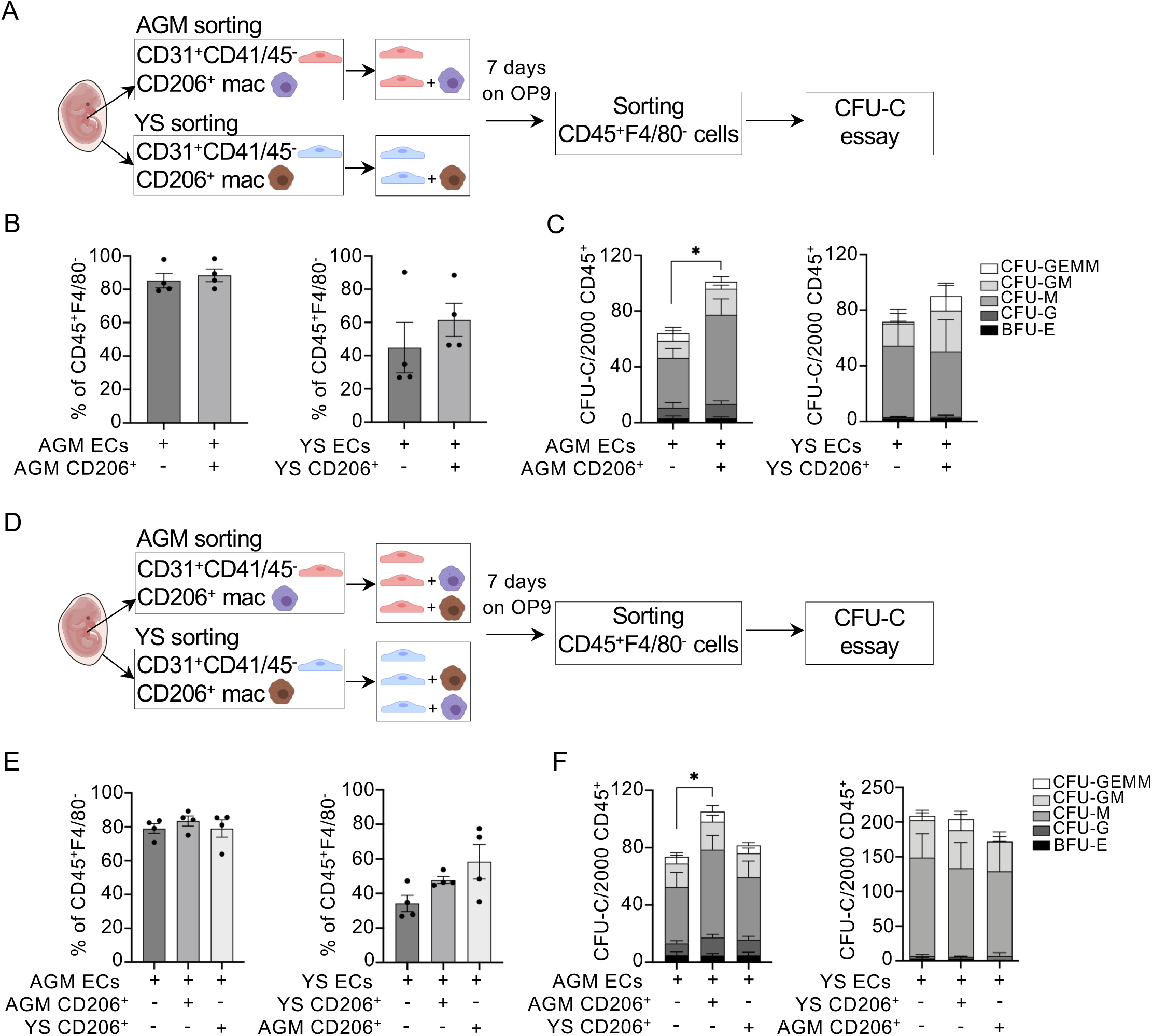
A) Schematic representation of co-culture experiments on OP9 stromal cells. Endothelial cells (CD31^+^CD41^-^CD45^-^) and CD206^+^ macrophages (CD45^+^GFP^+^cKit^-^CD11b^+^F4/80^+^CD206^+^) were sorted from the aorta-gonad-mesonephros (AGM) and yolk sac (YS) of embryonic day (E)10.5 *MacGreen* embryos and mixed and plated on OP9 cells for seven days. Nascent CD45^+^F4/80^-^ hematopoietic cells were subsequently sorted and plated in methylcellulose for a CFU-C (colony forming unit in culture) assay. B). Bar graph showing the frequency of hematopoietic CD45^+^F4/80^-^ cells produced by AGM (left) and YS (right) cells after one week of culture on OP9 stromal cells. Each dot represents an independent biological replicate, with 4-to-7 embryos pulled for each experiment C) Bar graph showing the number of hematopoietic colonies produced by AGM (left) and YS (right) CD45^+^F4/80^-^ cells retrieved after OP9 culture. N=4, with each N representing 4-to-7 pulled embryos. CFU-GEMM= colony forming unit-granulocyte, erythroid, macrophage, megakaryocyte; CFU-GM= colony forming unit-granulocyte, macrophage; CFU-M= colony forming unit-macrophage; CFU-G= colony forming unit-granulocyte; BFU-E= burst forming unit-erythroid. D) Schematic representation of co-culture experiments on OP9 stromal cells. Endothelial cells (CD31^+^CD41^-^CD45^-^) and CD206^+^ macrophages (CD45^+^cKit^-^CD11b^+^F4/80^+^CD206^+^) were sorted from the aorta-gonad-mesonephros (AGM) and yolk sac (YS) of embryonic day (E)10.5 *MacGreen* embryos and then mixed and plated on OP9 cells for seven days. Nascent CD45^+^F4/80^-^hematopoietic cells were subsequently sorted and plated in methylcellulose for a CFU-C (colony forming unit in culture) assay. E) Bar graph showing the frequency of hematopoietic CD45^+^F4/80^-^ cells produced by AGM endothelial (left) and YS enfothelial (right) cells after one week of culture on OP9 stromal cells in the presence of absence of AGM or YS CD206^+^ macrophages. Each dot represents an independent biological replicate, with 4-to-7 embryos pulled for each experiment. F) Bar graph showing the number of hematopoietic colonies produced by AGM and YS CD45^+^F4/80^-^ cells retrieved after OP9 culture. N=4, with each N representing 4-to-7 pulled embryos. CFU-GEMM= colony forming unit-granulocyte, erythroid, macrophage, megakaryocyte; CFU-GM= colony forming unit-granulocyte, macrophage; CFU-M= colony forming unit-macrophage; CFU-G= colony forming unit-granulocyte; BFU-E= burst forming unit-erythroid. C,F) Statistical test: Student’s t-test. *=p<0.05, **=p<0.01. Only statistically significant differences are indicated.

### The functions of AGM CD206^+^ mac are not cell intrinsic but tissue specific

To determine whether AGM CD206⁺ Mac functions are cell-intrinsic, we tested whether they could enhance the progenitor potential of YS hemogenic endothelial cells *in vitro*. Macrophages and endothelial cells from the AGM and YS of E10.5 *MacGreen* embryos were sorted and a mixed co-culture experiment was performed by culturing YS endothelial cells with AGM CD206⁺ Mac, with the reciprocal condition used as a control (Fig. 4D). No differences were observed in the frequency of CD45⁺F4/80⁻ cells after 7 days of culture on OP9 cells in any of the conditions tested (Fig. 4E). Furthermore, when colonies were counted and scored after 10 days in methylcellulose, AGM CD206⁺ Mac retained the ability to increase the progenitor potential of AGM endothelial cells but had no effect on YS endothelial cells (Fig. 4F), suggesting that their pro-hematopoietic function is not cell-intrinsic but dependent on crosstalk with AGM endothelial cells. As expected, YS CD206^+^ Mac had no effect on colony formation in any of the conditions tested (Fig. 4F, right).

### Only eight genes are preferentially upregulated in AGM CD206^+^ Mac

To understand which pathways and genes mediate the cross-talk between CD206⁺ Mac and hematopoietic progenitors in the AGM, we first determined the gene signature uniquely identifying AGM CD206⁺ Mac. Bulk RNA-seq was performed on AGM and yolk sac (YS) CD206⁺ Mac and CD206⁻ progenitors. Principal component analysis (PCA) showed AGM and YS samples clustering separately, with greater variability observed among YS samples (Fig. 5A). To identify genes preferentially upregulated in AGM CD206⁺ Mac, differential expression analysis was performed comparing AGM CD206⁺ Mac against all other cell populations (YS CD206⁺ Mac, and YS and AGM CD206⁻ cells). More than twenty genes were significantly downregulated in AGM CD206⁺ Mac relative to the other populations, whereas only eight genes were upregulated (Supplementary Fig. 4A and Fig. 5B). Two of these genes are linked to extracellular matrix remodeling — A Disintegrin-Like and Metalloprotease with Thrombospondin Type 1 Motifs 1 (*Adamts1*) and Matrix Metalloproteinase-2 (*Mmp2*)^43, 44^, two encode chemokines with known chemoattractant functions (*Ccl2* and *Cxcl16*)^45, 46^, while the remaining four (*Nrep, Tgm2, Lpar4*, and *Dapk1*) are involved in diverse biological processes including tissue remodeling, tissue regeneration, and regulation of autophagy and apoptosis^47–50^. To validate the RNA-seq results, RT-qPCR was performed on CD206⁺ Mac and CD206⁻ cells sorted from the AGM and YS of E10.5 *MacGreen* embryos. Expression of *Lpar4* (lysophosphatidic acid receptor 4), *Tgm2* (transglutaminase 2), *Adamts1*, and *Dapk1* (death-associated protein kinase 1) was not detected by RT-qPCR in any of the populations tested. In contrast, *Nrep* (neuronal regeneration-related protein), *Ccl2*, *Mmp2*, and *Cxcl16* were preferentially expressed in AGM CD206⁺ Mac, consistent with the RNA-seq data (Fig. 5C).

**Figure 5.**
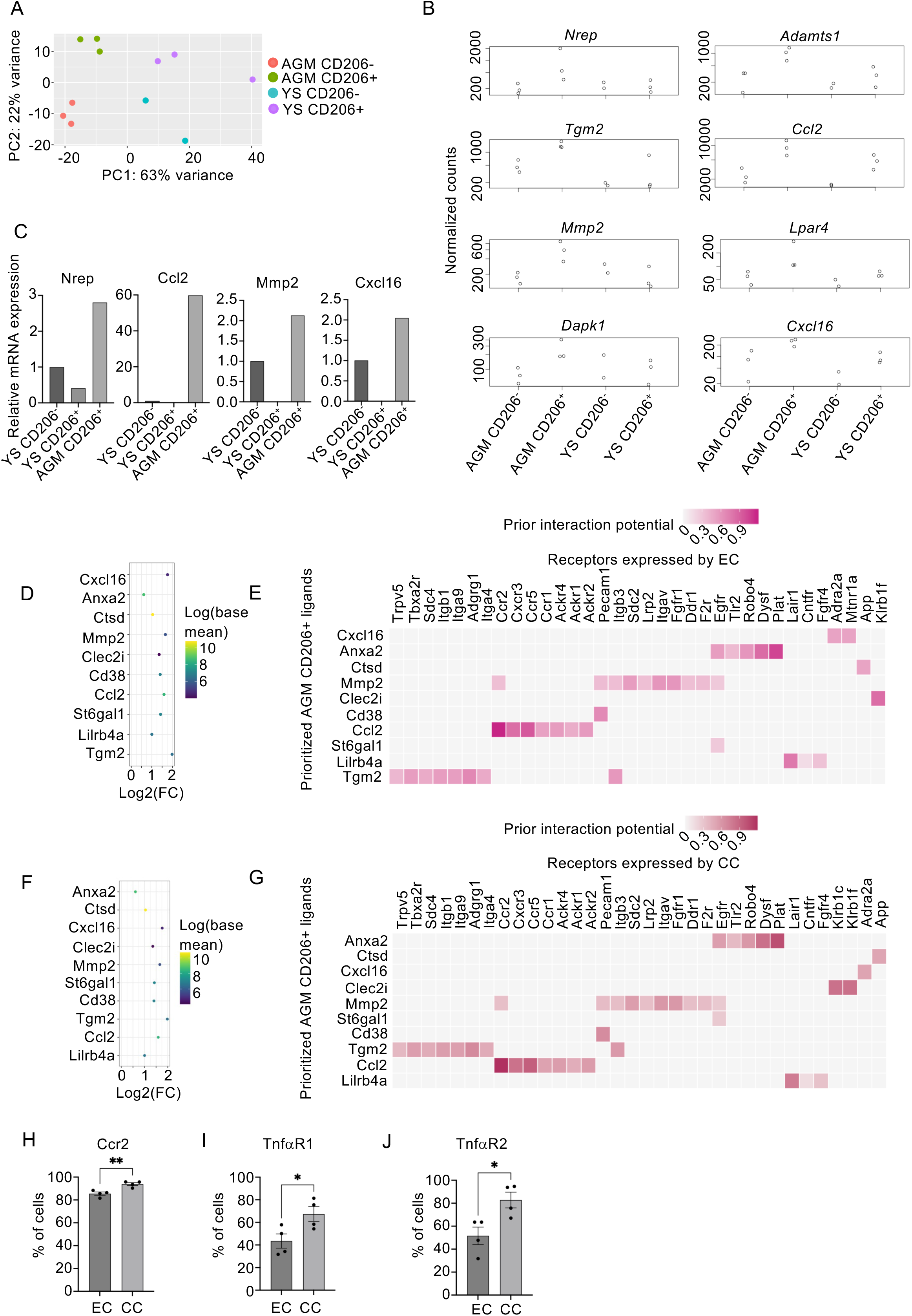
A) Principal component (PC) analysis plot indicated by color of aorta-gonad-mesonephros (AGM) CD45^+^GFP^+^cKit^-^CD11b^+^F4/80^+^CD206^-^ (red), AGM CD45^+^GFP^+^cKit^-^CD11b^+^F4/80^+^CD206^+^ (green), yolk sac (YS) CD45^+^GFP^+^cKit^-^ CD11b^+^F4/80^+^CD206^-^ (blue), and YS CD45^+^GFP^+^cKit^-^CD11b^+^F4/80^+^CD206^+^ (purple) cells based on gene expression of 35727 genes. N = 2-3, with each N representing 5-to-7 pulled *MacGreen* embryos. B) RNA sequencing normalized read count plots for *Adamts1*, *Ccl2*, *Tgm2*, *Nrep*, *Lpar4*, *Cxcl16*, *Mmp2*, and *Dapk1* in AGM CD45^+^GFP^+^cKit^-^CD11b^+^F4/80^+^CD206^-^, AGM CD45^+^GFP^+^cKit^-^CD11b^+^F4/80^+^CD206^+^, YS CD45^+^GFP^+^cKit^-^CD11b^+^F4/80^+^CD206^-^, and YS CD45^+^GFP^+^cKit^-^CD11b^+^F4/80^+^CD206^+^ cells N = 2-3, with each N representing 5-to-7 pulled *MacGreen* embryos. C) Real time qPCR analysis of *Nrep, Ccl2*, *Mmp2*, and *Cxcl16* in YS CD45^+^GFP^+^cKit^-^CD11b^+^F4/80^+^CD206^-^, YS CD45^+^GFP^+^cKit^-^CD11b^+^F4/80^+^CD206^+^, and AGM CD45^+^GFP^+^cKit^-^CD11b^+^F4/80^+^CD206^+^ cells. mRNA expression is relative to YS CD206^-^ cells. N=1 representing 9 E10.5 *MacGreen* embryos from 2 different litters. D) RNA sequencing analysis of potential AGM CD45^+^GFP^+^cKit^-^CD11b^+^F4/80^+^CD206^+^ ligands colored by the log of the mean base expression (from the lowest as purple to the highest as yellow). Log2(FC) is the log2 of the fold change in AGM CD45^+^GFP^+^cKit^-^CD11b^+^F4/80^+^CD206^+^ cells compared to all other macrophages. N = 2-3, with each N representing 5-to-7 pulled *MacGreen* embryos. E) Matrix showing known interactions of ligands expressed by AGM CD45^+^GFP^+^cKit^-^CD11b^+^F4/80^+^CD206^+^ macrophages and receptors expressed by endothelial cells (EC). The squares are colored according to the prior interaction potential from the lowest as gray to the highest as pink. F) RNA sequencing analysis of potential AGM CD206+ ligands colored by the log of the mean base expression (from the lowest as purple to the highest as yellow). Log2(FC) is the log2 of the fold change in AGM CD206+ cells compared to all other macrophages. N = 2-3, with each N representing 5-to-7 pulled *MacGreen* embryos. G) Matrix showing known interactions of ligands expressed by AGM CD45^+^GFP^+^cKit^-^CD11b^+^F4/80^+^CD206^+^ macrophages and receptors expressed by cluster cells (CC). The squares are colored according to the prior interaction potential from the lowest as gray to the highest as red. H) Flow cytometry analysis of Ccr2 surface expression on endothelial cells (ECs) and cluster cells (CCs). I) Flow cytometry analysis of TnfaR1 surface expression on endothelial cells (ECs) and cluster cells (CCs). J) Flow cytometry analysis of TNfaR2 surface expression on endothelial cells (ECs) and cluster cells (CCs). H-J: Statistical analysis: Student’s t-test. *=p<0.05, **=p<0.01.

### Macrophages interact with both endothelial and cluster cells in the mouse AGM

To determine whether AGM CD206⁺ Mac promote HSPC generation *in vitro* by interacting with hemogenic endothelial cells (ECs) or with CD31⁺cKit⁺ cluster cells (CCs), we sorted CD206⁺ Mac, ECs (CD31⁺cKit⁻CD41⁻CD45⁻), and early CCs (CD31⁺cKit⁺CD41⁻CD45⁻) from the AGM of E10.5 MacGreen embryos for co-culture experiments. EC and CC populations were each plated on stromal cells in the presence or absence of CD206⁺ Mac. After one week on OP9 stromal cells, no CD45⁺ cells were recovered under any of the conditions tested (data not shown). These results suggest that the *in vitro* co-culture system is only productive when ECs and CCs are sorted together as CD31⁺cKit⁺^/^⁻CD41⁻CD45⁻, as performed in previous experiments (Fig. 4). To circumvent this experimental limitation, ECs and CCs were sorted from the AGM of E10.5 *MacGreen* embryos and subjected to bulk RNA-seq, followed by NicheNet analysis to predict the intercellular interactions between AGM CD206⁺ Mac and ECs or CCs. The top-ranked predicted ligands were shared between ECs and CCs, with the same base mean expression values (Fig. 5D and 5F). Bioinformatic analysis did not identify differential predicted interactions between CD206⁺ Mac and ECs or CCs, with the sole exception of the CC-specific *Clec2i–Klrb1c* interaction, that we were unable to validate by real time PCR. The near-complete concordance of predicted interactions suggests that ECs and CCs share the same potential of interacting with CD206^+^ macrophages at E10.5 (Fig. 5E and 5G). Among the predicted interactions, *Ccl2* expressed by CD206⁺ Mac showed the strongest predicted interaction with *Ccr2* expressed on both ECs and CCs (Fig. 5E and 5G). To assess whether Ccr2 surface expression differed between these populations at the protein level, flow cytometry was performed on E10.5 AGM ECs and CCs (Supplementary Fig. 4B). Both populations expressed high levels of Ccr2, with a statistically significant increase observed in the CC population (Fig. 5H). NicheNet analysis did not predict any interactions mediated via *Tnfα–TnfR* signaling between CD206⁺ Mac and ECs or CCs (Fig. 5E and 5G). However, given that *Tnf* was the most highly upregulated cytokine in AGM CD206^+^ subset compared to YS CD206⁺ Mac (Fig. 3F), we assessed the protein expression of TnfR1 and TnfR2 on E10.5 AGM ECs and CCs by flow cytometry (Supplementary Fig. 4B). Both receptors were expressed at significantly higher levels on CCs than on ECs (Fig. 5I), suggesting that CD206⁺ Mac can interact with both populations, but with greater receptor-mediated engagement with CCs.

## Discussion

In this study, we employed a combination of *in vivo* and *in vitro* approaches to identify the mediators of AGM macrophage functions that facilitate HSPC generation *in vitro*. We first established that CD206 expression is a hallmark of terminally differentiated macrophages in the mouse AGM, defined by the absence of *in vitro* progenitor potential — a capacity that is retained by CD45⁺CD11b⁺F4/80⁺cKit⁻CD206⁻ cells. CD206⁺ Mac (CD45⁺CD11b⁺F4/80⁺cKit⁻CD206⁺) enhance HSPC generation when co-cultured with endothelial cells *in vitro*. CD206, also known as the mannose receptor, is a carbohydrate-binding endocytic receptor with diverse functions including clearance of endogenous and exogenous molecules, collagen internalization, modulation of cellular activation, and promotion of antigen presentation^51^. CD206 is typically expressed by mature monocytes, macrophages, and certain subsets of vascular and lymphatic endothelial cells^52^. Historically, mannose receptor expression has been associated with alternatively activated macrophages exhibiting predominantly anti-inflammatory and pro-tumoral functions^53, 54^. However, this association has been recently challenged by studies demonstrating a role for CD206⁺ Mac in recruiting anti-tumoral T cells^55^, pointing to a more nuanced role for CD206⁺ Mac in cancer biology. A similar complexity appears to apply to AGM macrophages, in which CD206 is expressed on the surface of macrophages that also express pro-inflammatory cytokines. As previously discussed^28^, in the embryonic AGM — an environment not exposed to extracellular pathogens — a pro-inflammatory milieu has been reported to be necessary for correct HSC generation in vertebrates^56, 57^. Although we consider CD206 to be a marker of macrophage maturity in the AGM rather than a molecule directly involved in HSC generation, additional experiments are needed to determine whether AGM CD206⁺ Mac actively contribute to extracellular matrix remodeling through collagen internalization. That said, the viability of CD206 KO mice, with no reported defects in embryonic HSC generation, argues against a direct functional role for CD206 in embryonic HSPC generation^58, 59^.

We also observed that CD206⁺ Mac are characterized by high F4/80 expression. F4/80 is one of the most specific cell-surface markers for murine macrophages ^39, 60^. It belongs to the epidermal growth factor–seven transmembrane receptor family and is thought to mediate cell adhesion; however, its precise function(s) remain to be fully elucidated^61^. F4/80 knockout mouse models have not revealed a requirement for F4/80 in normal macrophage development or function^62^. Nevertheless, F4/80 has been shown to be required for the induction of antigen-specific CD8⁺ regulatory T cells in models of peripheral tolerance in adult mice, a process dependent on direct cellular interaction between macrophages and NKT cells^30^. This demonstrates that F4/80 can mediate cell-to-cell interactions in specific tissue contexts. Having previously observed direct interactions between macrophages and hematopoietic cells in the mouse AGM at E10.5 ^28^, we used the F4/80 KO mouse model^30^ to investigate whether F4/80 plays a previously unrecognized role in mediating macrophage–HSPC interactions in the mouse embryo. Notably, we found a reduction in the frequency of endothelial cells (CD31⁺cKit⁻CD41⁻CD45⁻) in F4/80 KO embryos, suggesting that F4/80 may contribute to maintenance of the endothelial cell pool in the AGM. Since CD31⁺F4/80⁺ endothelial cells have been reported in both embryos and adult mice^63, 64^, but our aim is to investigate the effect of the absence of macrophage-expressed F4/80, flow cytometry analyses were performed on F4/80⁻ cells to exclude any confounding factors. While endothelial cell frequency was reduced in the absence of F4/80, the frequency of cKit⁺CD31⁻ cells was increased. cKit is a receptor tyrosine kinase whose expression is a canonical marker of HSPCs^65^. An increased frequency of cKit⁺ cells in the AGM of F4/80 KO embryos could indicate a role for F4/80 in HSPC generation or proliferation. However, we cannot exclude the possibility that other non-hematopoietic cell types within the AGM also express cKit. The AGM encompasses not only the aorta — the site of HSPC generation — but also the gonad and mesonephros, which give rise to the gonads and kidneys later in development. Although no studies have specifically examined cKit expression in the gonad–mesonephros region of the AGM at E10.5, cKit expression has been documented during gametogenesis in primordial germ cells, spermatogonia, primordial and growing oocytes, and in the developing kidney^66, 67^. Therefore, the overall increase in cKit⁺ cell frequency observed in F4/80 KO embryos may not be restricted to the hematopoietic compartment. Nevertheless, an expansion of early hematopoietic progenitors expressing CD41^lo^, CD43, and CD45 is evident in F4/80 KO embryos at E10.5 and it is reflected by increased progenitor output from AGM cells isolated from these embryos. This may result from defective interactions between macrophages and HSPCs, leading to an expansion of specific HSPC subsets — suggesting that macrophages contribute to regulating the number of HSPCs generated in the AGM at E10.5. A similar regulatory role has been reported in the caudal hematopoietic tissue of zebrafish (the functional equivalent of the mouse fetal liver), where macrophages influence the number of HSC clones generated in the embryo that subsequently contribute to adult hematopoiesis^68^. In zebrafish, however, fewer HSC clones were observed in the absence of macrophages^68^. It is important to note that in F4/80 KO embryos, macrophages are still present, and macrophage–HSPC interactions may be impaired but not entirely abolished. We do not consider F4/80 to be the primary mediator of the HSPC-promoting functions of AGM macrophages, as its complete absence would be expected to result in fewer — not more — HSPCs. Interestingly, preliminary analysis of phenotypic HSC frequency in the fetal liver of E13.5 F4/80 KO embryos revealed a reduction compared to wild-type controls (unpublished data). Together with the absence of any hematopoietic defect in the yolk sac, this suggests that the progenitor expansion observed in the AGM of F4/80 KO embryos is niche-dependent, and that macrophage–HSPC interactions may yield distinct outcomes in different niches during development. This interpretation is further supported by the inability of YS macrophages to enhance the progenitor potential of YS endothelial cells, indicating that the HSPC-promoting activity of AGM macrophages is niche-specific rather than ontogeny-dependent.

We subsequently identified the key genes expressed by AGM CD206⁺ macrophages. No specific pathways or Gene Ontology (GO) terms were enriched, but eight genes with diverse functions were found to be upregulated specifically in AGM CD206⁺ Mac compared to AGM CD206⁻ cells and YS CD206⁺ and CD206⁻ cells. Upregulation of four of the eight genes (*Dapk1*, *Tgm2*, *Lpar4*, and *Adamts1*) could not be validated by real time PCR (RT-qPCR), possibly due to their low expression levels combined with the limited number of CD206⁺ and CD206⁻ cells obtainable from the AGM, where CD206⁺ Mac represent approximately 0.2% of total live cells at E10.5. Despite this limitation, upregulation of *Nrep* (neuronal regeneration-related protein), *Mmp2* (matrix metallopeptidase 2), *Ccl2*, and *Cxcl16* in AGM CD206⁺ Mac was validated by RT-qPCR. *Nrep* encodes a highly conserved 8 kDa protein associated with the TGF-β pathway, wound healing, nerve and lung regeneration, and kidney fibrosis^48^. Although no role has been described for *Nrep* in the AGM specifically, it plays important roles in nervous system development, and a functional cross-talk between the sympathetic nervous system and HSC generation in the developing embryo has previously been reported^69^. *Mmp2* encodes a metallopeptidase involved in the degradation and remodeling of extracellular matrix components, including type IV and V collagen, fibronectin, laminin, and elastin^44^. *Mmp2* has been shown to be essential for the endothelial-to-hematopoietic transition in zebrafish, where it is upregulated in response to inflammation and contributes to HSC emergence and subsequent egress from the dorsal aorta^70^. *Cxcl16* is an atypical CXC chemokine that exists in both transmembrane and soluble forms, functioning as a chemoattractant and scavenger receptor in its soluble form and as an adhesion molecule when membrane-bound^71^. Although there is currently no published evidence for expression of the *Cxcl16* receptor (*Cxcr6*) on endothelial or hematopoietic cells in the AGM, the Cxcl16–Cxcr6 axis has been shown to promote Tnfα secretion^72^ and may thus contribute to the pro-inflammatory environment in the AGM that is required for HSC generation. Finally, NicheNet analysis predicted broadly similar interactions between CD206⁺ Mac and both endothelial cells (ECs) and cluster cells (CCs) in the AGM. Flow cytometry confirmed expression of receptors for Ccl2 and Tnfα on both ECs and CCs, with significantly higher levels on CCs. This suggests that interactions between macrophages and/or macrophage-secreted factors and AGM cells may be initiated with ECs and subsequently amplified with nascent CCs.

Taken together, our data support a multi-functional role for CD206⁺ Mac in the AGM at E10.5 and may inform improvements to iPSC differentiation protocols for the *in vitro* generation of HSCs, for example through the timed addition of key factors such as Ccl2, Tnfα, and Cxcl16 during differentiation. In addition, our findings suggest a previously unrecognized role for F4/80 during mammalian embryonic development.

## Limitations

While the transcriptomic differences between embryonic macrophage subsets are representative of the *in vivo* situation in the mouse embryo, the functional assays were performed in an *in vitro* 2D model of endothelial-to-hematopoietic transition. This was specifically designed to inform iPSC protocols for *in vitro* HSC/HSPC generation; however, the results may not fully recapitulate what occurs *in vivo*. In particular, the compensatory mechanisms that may operate in a living embryo are absent in our model. We previously reported a decrease in functional HSCs in the absence of macrophages^28^ and here we confirmed impaired HSPC generation *in vitro*. However, Perçin et al. recently reported no deficit in AGM HSC numbers in an *in vivo* model of embryonic macrophage depletion, suggesting that compensatory mechanisms operate *in vivo* that are not recapitulated in our system^74^. A further limitation of this study is the absence of transplantation data. With the current experimental setup, it is not possible to determine whether the loss of macrophages and/or F4/80 also impairs the generation of functional HSCs. Future experiments using human iPSCs will be necessary to evaluate the effects of macrophages on HSC generation and/or maturation, and to assess the translational relevance of our findings. Human iPSC experiments will also be critical for determining whether supplementation with macrophage-related factors — such as CCL2, CXCL16, and TNFα — during differentiation is beneficial for HSC generation, particularly given previous reports that an *in vitro* inflammatory environment may favour HSC differentiation over self-renewal^75^. It should be noted, however, that this effect was observed in fetal bovine serum-containing cultures^75^, and may therefore not directly apply to defined, serum-free iPSC differentiation systems. In conclusion, our data reveal a previously unreported role for F4/80 in regulating HSPC frequency in the E10.5 AGM, and define a gene signature for CD206⁺ Mac that may be used to inform iPSC protocols for *in vitro* HSC differentiation. Validation in human cellular models will nonetheless be essential to establish the translational relevance of these findings.

## Author contributions

R.L.B. performed the bioinformatic analyses and contributed to manuscript preparation. M.R. performed validation experiments and contributed to manuscript preparation. A.P. contributed to RNA-seq analysis, alignment, and initial bioinformatic processing. A.M., S.K., M.R., H.M., and E.B. performed immunofluorescence, co-culture, and RT-qPCR experiments. C.B. and E.A. performed, analyzed, and interpreted lineage-tracing experiments. A.M. provided support for lineage-tracing mouse work. S.M. and S.G. provided mice and scientific discussion. S.A.M. conceived the project, designed the study, performed experiments, and wrote the manuscript. All authors reviewed, edited, and approved the final version of the manuscript

## Acknowledgements

R.L..B. and M.R. are supported by Leukaemia UK 2024/FuF/004 and Blood Cancer UK (grant #25031), respectively. The experiments on F4/80 KO animals were supported by The Carnegie Trust (RIG012518).

S.A.M.’s research is supported by Leukaemia UK (2021/JGF/004), Medical Research Scotland (ECG-1797-2022), Wellcome Trust Institutional Translational Partnership Award (WT iTPA PIII063), the Little Princess Trust through the Children’s Cancer and Leukaemia Group (CCLGA 2023 13 Mariani) and Medical Research Scotland (PHD-50551-2022). E.A. is supported by Worldwide Cancer Research (Grant ref. 24-0083). E.A. and A.M. are supported by Italian Ministry of University and Research (Grant n. P20223HEZC). C.B. is supported by Fondazione Umberto Veronesi.

The authors would like to acknowledge the helpful contributions of the flow cytometry and sorting facility of the Institute for Regeneration and Repair of the University of Edinburgh, and Sarah Coupland, Colin Dick and the rest of the staff at the University of Edinburgh Bioresearch & Veterinary Services (BVS) for their professional animal care, technical assistance, and veterinary support during this study.

