## Supplemental Figures for "The Aorta-Gonad-Mesonephros niche shapes the functions of yolk sac-derived macrophages involved in hematopoietic stem and progenitor cell generation *ex vivo*"

Supplementary Figure 1 – related to Figure 1

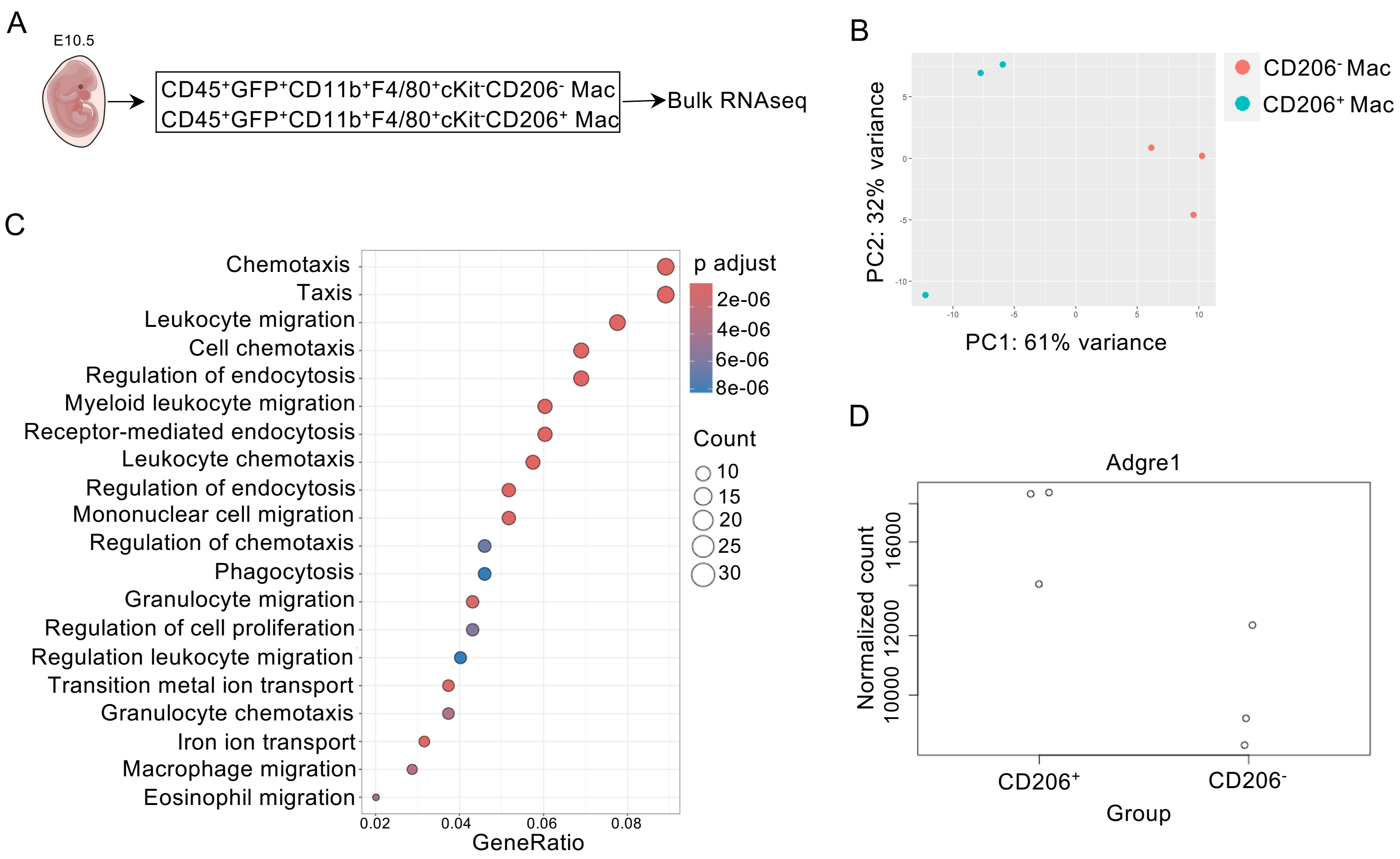


**Supplementary Figure 1:**

A) Experiment schematics of bulk RNA sequencing (RNAseq) analysis of CD206$^+$ macrophages and CD206$^-$ cells with corresponding surface markers used for sorting strategy. B)Principal component analysis plot from RNA sequencing analysis of CD45$^+$GFP$^+$CD11b$^+$F4/80$^+$cKit$^-$CD206$^-$ cells (red) and CD45$^+$GFP$^+$CD11b$^+$F4/80$^+$cKit$^-$CD206$^+$ macrophages (blue). N = 3, with each N representing 5-to-7 embryos pulled together. C) KEGG pathway analysis of genes expressed by CD45$^+$GFP$^+$CD11b$^+$F4/80$^+$cKit$^-$ CD206$^+$ macrophages. Adjusted p-value is indicated by color with red indicating a smaller adjusted p-value and blue indicating a higher adjusted p-value. The number of genes associated with each term is indicated by size of the dot. D) Normalized read count plot for *Adgre1* gene (encoding for F4/80) illustrating differences between CD45$^+$GFP$^+$CD11b$^+$F4/80$^+$cKit$^-$CD206$^+$ macrophages and CD45$^+$GFP$^+$CD11b$^+$F4/80$^+$cKit$^-$ CD206$^-$ cells. Data from bulk RNAseq on N=3, with each N representing 5-to-7 embryos pulled together.

Supplementary Figure 2 – related to Figure 2

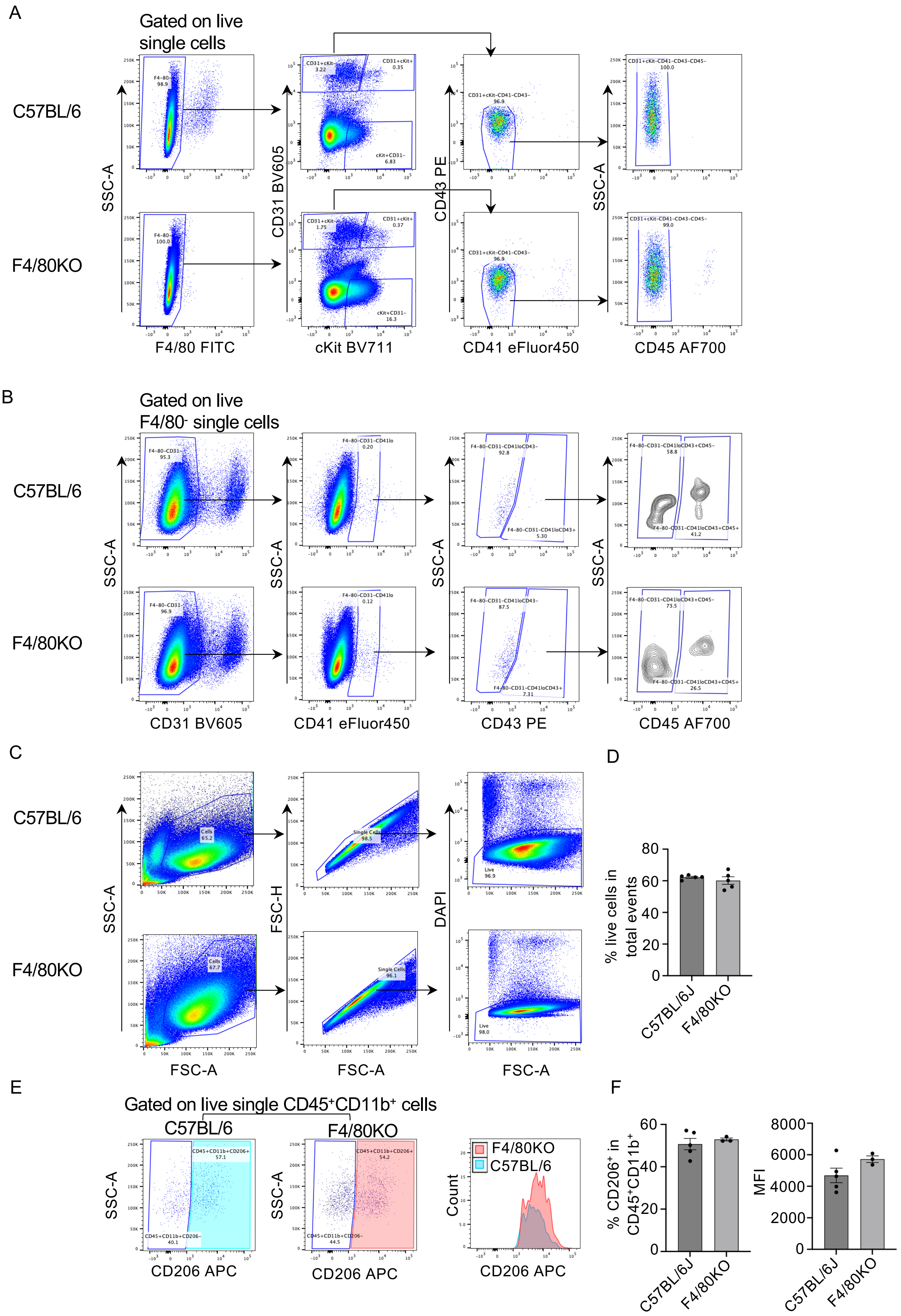

**Supplementary Figure 2:**

A) Representative flow cytometry gating strategy of F4/80$^{-}$, CD31$^{+}$cKit$^{-}$, CD31$^{+}$cKit$^{+}$, CD31$^{-}$cKit$^{+}$, CD31$^{+}$cKit$^{-}$CD41$^{-}$CD43$^{-}$CD45$^{-}$ endothelial cells in the aorta-gonad-mesonephros (AGM) of C57BL/6 and F4/80 knock out (KO) embryos at embryonic day (E)10.5. Arrows indicate gate dependency. B) Representative flow cytometry gating strategy of CD31$^{-}$CD41$^{lo}$CD43$^{+}$CD45$^{+}$ hematopoietic progenitors in the AGM of E10.5 C57BL/6 and F4/80 KO embryos. Arrows indicate gate dependency. C) Left, representative flow cytometry gating strategy of single live cells in the AGM of E10.5 C57BL/6 and F4/80 KO embryos and corresponding quantification on the right. Each dot represents an individual embryo. E) Representative flow cytometry gating strategy of CD206$^{+}$ frequency and mean fluorescent intensity (MFI) in E10.5 C57BL/6 (cyan) and F4/80 KO (red) embryos. F) Bar graphs showing the quantification of the data in E. Each dot represents a different embryo.

Supplementary Figure 3 – related to Figure 3

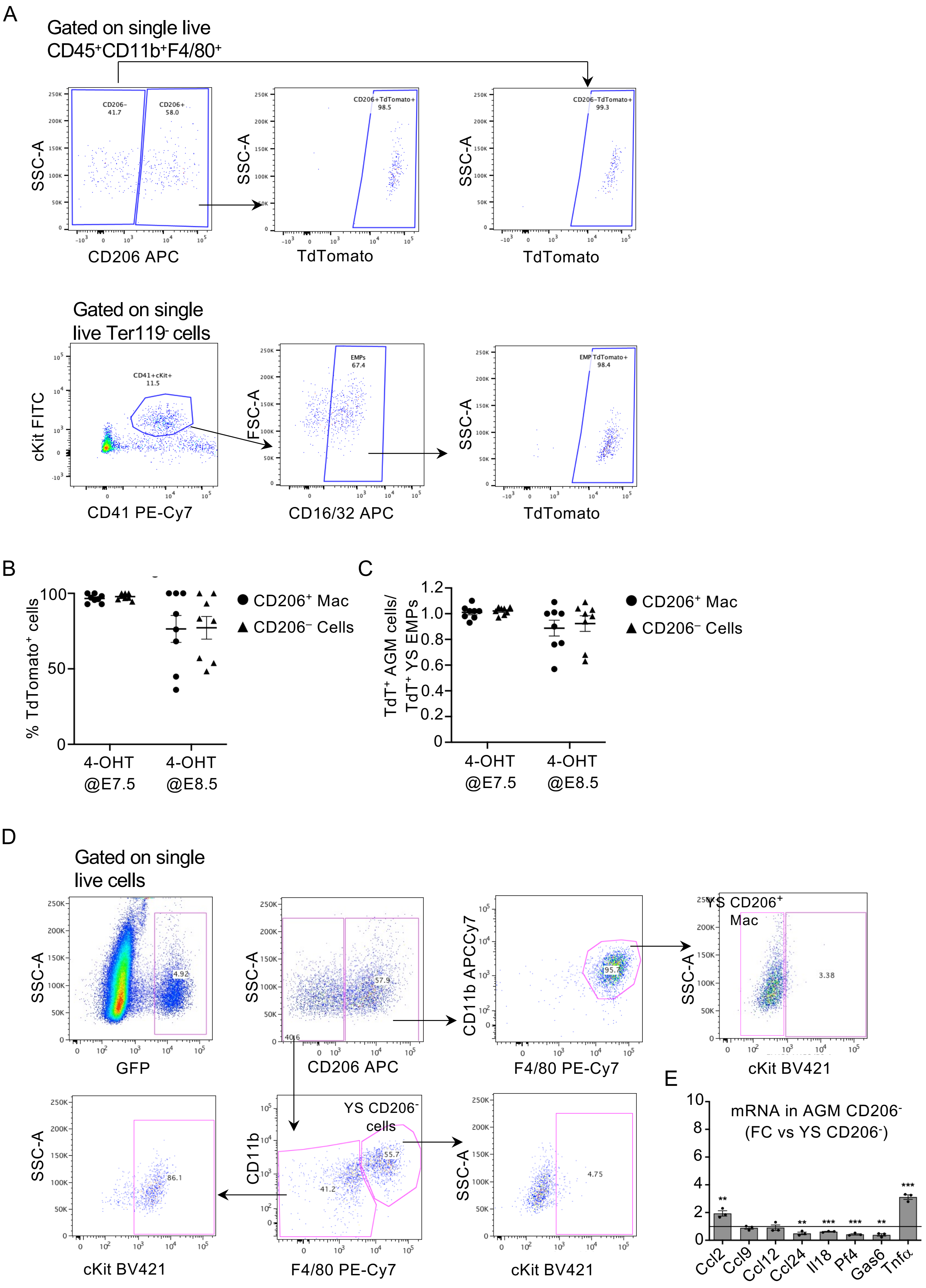

**Supplementary Figure 3:**

A) Top, representative flow cytometry gating strategy of TdTomato$^+$ CD206$^+$ and CD206$^-$ cells in the aorta-gonad-mesonephros (AGM) of Cdh5-Cre$^{ERT2}$::Rosa26$^{tdTomato}$ embryos at embryonic day (E)10.5 after a pulse-chase with 4-hydroxytamoxifen at E7.5. Bottom, representative flow cytometry gating strategy of TdTomato$^+$ cKit$^+$CD41$^+$CD16/32$^+$ erythroid myeloid progenitors (EMPs) in the yolk sac (YS) of E10.5 Cdh5-Cre$^{ERT2}$::Rosa26$^{tdTomato}$ embryos after a pulse-chase with 4-hydroxytamoxifen at E7.5. Arrows indicate gate dependency. B) Interleaved scatter plot showing the quantification of TdTomato$^+$ AGM CD206$^+$ macrophages (black circle) and CD206$^-$ cells (black triangle) after a pulse-chase with 4-hydroxytamoxifen at E7.5. or E8.5. C) Interleaved scatter plot showing the ratio of TdTomato$^+$ AGM CD206$^+$ macrophages (black circle) and CD206$^-$ cells (black triangle) over YS EMPs after a pulse-chase with 4-hydroxytamoxifen at E7.5. or E8.5. D) Representative flow cytometry gating strategy of CD45$^+$GFP$^+$CD11b$^+$F4/80$^+$cKit$^-$ CD206$^+$ macrophages and CD45$^+$GFP$^+$CD11b$^+$F4/80$^+$cKit$^-$ CD206$^-$ cells in the YS of E10.5 *MacGreen* embryos. E) Real time qPCR analysis of *Ccl2*, *Ccl9*, *Ccl12*, *Ccl24*, interleukin 18 (*Il18)*, platelet factor 4 (*Pf4)*, growth arrest-specific 6 (*Gas6)*, tumor necrosis factor $\alpha$ (*Tnf*$\alpha$) expression in AGM CD206$^-$ cells (columns) normalized over $\beta$*Actin* expression and then plotted as fold change over the expression in YS CD206$^-$ cells (horizontal line). Each dot represents an independent biological replicate, with 4-to-6 embryos per replicate. Statistical test: Student's t-test. **=p<0.01, ***=p<0.001. Only statistically significant differences are indicated.

Supplementary Figure 4 – related to Figure 5

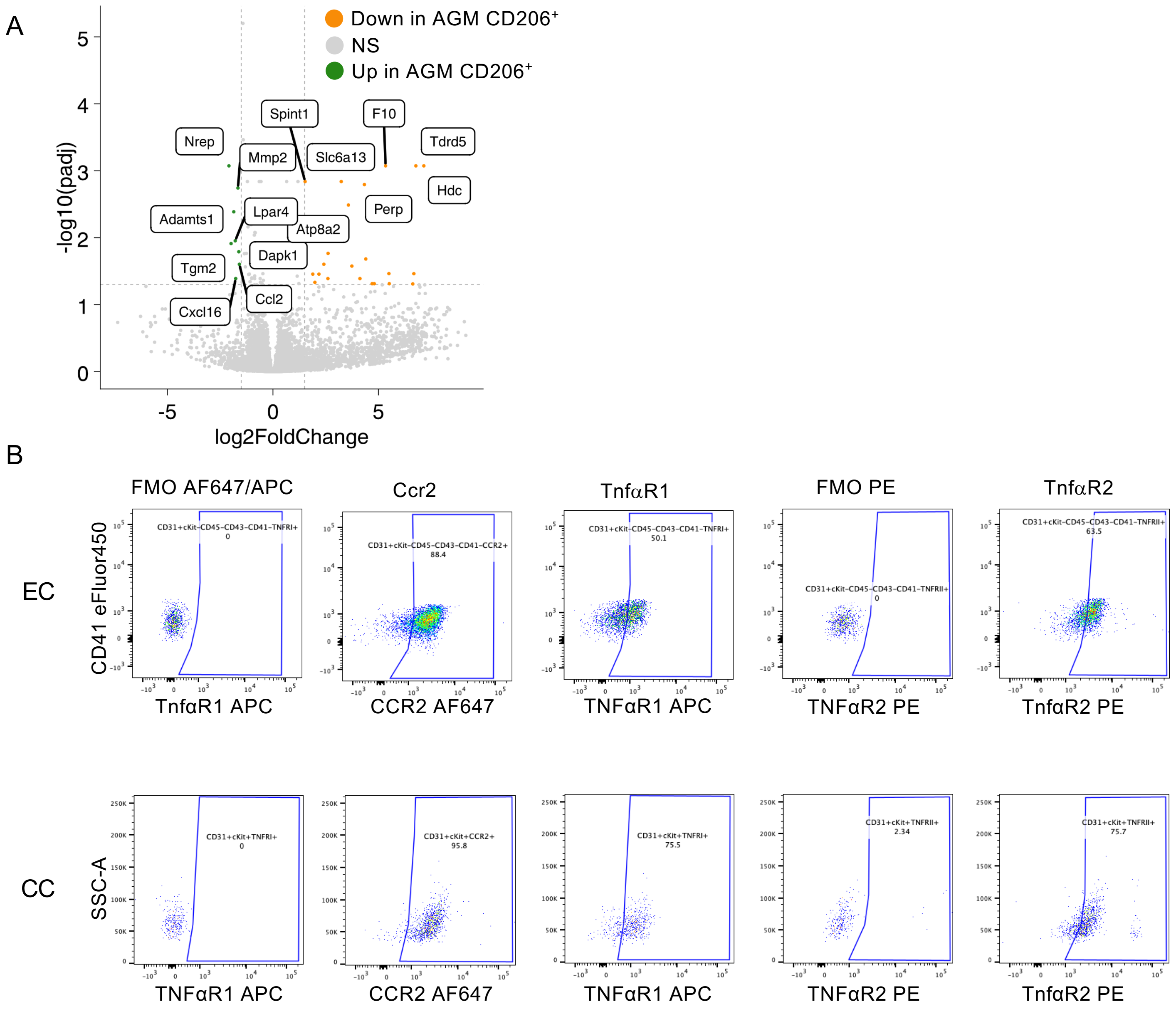


**Supplementary Figure 4:**

A) Volcano plot of differential gene expression in AGM $CD45^+GFP^+CD11b^+F4/80^+cKit^-CD206^+$ ($CD206^+$) macrophages compared YS $CD45^+GFP^+CD11b^+F4/80^+cKit^-CD206^+$ macrophages and AGM and YS $CD45^+GFP^+CD11b^+F4/80^+cKit^-CD206^-$ cells. Genes that are significantly upregulated in AGM $CD206^+$ macrophages (log2(fold change) < -1.5 and adjusted p-value < 0.05) are labeled and show in green. Genes that are significantly downregulated in AGM $CD206^+$ macrophages (log2(fold change) > 1.5 and adjusted p-value < 0.05) are labeled and show in orange.

B-C) Flow cytometry gating strategy in endothelial cells (EC) and cluster cells (CC) of Ccr2, TnfaR1, and TnfaR2 based on their respective fluorescence minus one (FMO).
