## Supplemental Table 1 for "The Aorta-Gonad-Mesonephros niche shapes the functions of yolk sac-derived macrophages involved in hematopoietic stem and progenitor cell generation *ex vivo*"

Supplementary Table 1
Antibody list

| Antibody | Clone | Company |
|---|---|---|
| AF647 α Mouse CD192 (CCR2) | SA203G11 | BioLegend |
| FITC α Mouse F4/80 | BM8 | BioLegend |
| PECy7 α Mouse F4/80 | BM8 | BioLegend |
| BV421 α Mouse F4/80 (immunofluorescence) | BM8 | BioLegend |
| BV605 α Mouse CD31 | 390 | BioLegend |
| AF700 α Mouse CD45 | 30-F11 | BioLegend |
| APC/Cy7 α Mouse/human CD11b (Mac1) | M1/70 | BioLegend |
| APC α Mouse CD206 (mannose receptor) | C068C2 | BioLegend |
| BV421 α Mouse CD117 (cKit) | 2B8 | BD Biosciences |
| BV711 α Mouse CD117 (cKit) | 2B8 | BioLegend |
| FITC α Mouse CD117 (cKit) | 2B8 | BioLegend |
| Rat α Mouse CD117 (cKit) (immunofluorescence) | 2B8 | eBioscience |
| AF647 α Mouse CD192 (Ccr2) | SA203G11 | BioLegend |
| APC α Mouse CD120a (TnfaR1) | 55R-286 | BioLegend |
| PE α Mouse CD120b (TnfaR2) | TR75-89 | BioLegend |
| Efluor450 α Mouse CD41 | MWReg30 | eBiosciences |
| PECy7 α Mouse CD41 | MWReg30 | BioLegend |
| PE α Mouse CD43 | eBioR2/60 | eBiosciences |
| APC α Mouse CD16-32 | 93 | BioLegend |
| AF647 Goat α Rat IgG H+L (Immunofluorescence) | Polyclonal | ThermoFisher Sc. |
