## Supplemental Table 2 for "The Aorta-Gonad-Mesonephros niche shapes the functions of yolk sac-derived macrophages involved in hematopoietic stem and progenitor cell generation *ex vivo*"

Supplementary Table 2
Real Time qPCR primer list

| **Gene** | **Primer sequence 5'-3'** |
|---|---|
| *Ccl2 Fw* | TTAAAAACCTGGATCGGAACCAA |
| *Ccl2 Rev* | GCATTAGCTTCAGATTTACGGGT |
| *Ccl9 Fw* | CCCTCTCCTTCCTCATTCTTACA |
| *Ccl9 Rev* | AGTCTTGAAAGCCCATGTGAAA |
| *Ccl12 Fw* | ATTTCCACACTTCTATGCCTCCT |
| *Ccl12 Rev* | ATCCAGTATGGTCCTGAAGATCA |
| *Ccl24 Fw* | AGGCAGGGGTCATCTTCATC |
| *Ccl24 Rev* | TTGGCCCCTTTAGAAGGCTG |
| *Gas6 Fw* | GACCCCGAGACGGAGTATTTC |
| *Gas6 Rev* | TGCACTGGTCAGGCAAGTTC |
| *Pf4 Fw* | AGAGCCCTAGACCCATTTCCT |
| *Pf4 Rev* | CATTCTTCAGGGTGGCTATGA |
| *Il18 Fw* | GTGAACCCCAGACCAGACTG |
| *Il18 Rev* | CCTGGAACACGTTTCTGAAAGA |
| *Ifnα1 Fw* | TGCCCAGCAGATCAAGAAGG |
| *Ifnα1 Rev* | TCAGGGGAAATTCCTGCACC |
| *Ifnγ Fw* | AGGAACTGGCAAAAGGATGGT |
| *Ifnγ Rev* | TCATTGAATGCTTGGCGCTG |
| *Tnfa Fw* | GGTGCCTATGTCTCAGCCTCTT |
| *Tnfa Rev* | CTCCCTCTCATCAGTTCTATGGC |
| *Il1β Fw* | TGGACCTTCCAGGATGAGGACA |
| *Il1β Rev* | CACTACAGGCTCCGAGATGAAC |
| *Nrep Fw* | GCGGGGCTTTTGTCTGTTG |
| *Nrep Rev* | TGGTTCTTGACTGACCCAGA |
| *Adamts1 Fw* | CTCTCACCCTTCGGAATTTCTG |
| *Adamts1 Rev* | GGAGCCACATAAATCCTGTCTG |
| *Tgm2 Fw* | GAGCGAGATGATCTGGAACTT |
| *Tgm2 Rev* | TGGGCCACAACAGTATGTCC |
| *Mmp2 Fw* | TGCAGGAGACAAGTTCTGGA |
| *Mmp2 Rev* | TTGAAGAAGTAGCTATGACCACCA |
| *Lpar4 Fw* | ACACTCTTTCTTGGGCACTCAAT |
| *Lpar4 Rev* | CTGAGGACCAGTAGAGAATGCTT |
| *Dapk1 Fw* | CGCCAATGTGGAGGCTCTAA |
| *Dapk1 Rev* | GTCCTCGGTGTGTGTCCTTC |
| *Cxcl16 Fw* | TGTCCATTCTTTATCAGGTTCCA |
| *Cxcl16 Rev* | CCCATGACCAGTTCCACACT |
| *β Actin Fw* | CACCACACCTTCTACAATGAG |
| *β Actin Rev* | GACCCAGATCATGTTTGAGAC |
