## Supplemental Table 3 for "The Aorta-Gonad-Mesonephros niche shapes the functions of yolk sac-derived macrophages involved in hematopoietic stem and progenitor cell generation *ex vivo*"

| Gene name | baseMean | log2FoldCha | lfcSE | stat | pvalue | padj |
|---|---|---|---|---|---|---|
| Gnai3 | 8989.06071 | 0.032587 | 0.14684229 | 0.22191839 | 0.82437742 | 0.962942 |
| Cdc45 | 3056.66549 | -0.6406405 | 0.21152378 | -3.0286928 | 0.00245614 | 0.04636729 |
| Scml2 | 30.3412488 | 1.41773835 | 0.84341498 | 1.68094992 | 0.09277264 | 0.46606436 |
| Narf | 2095.28069 | 0.22565706 | 0.1904831 | 1.18465658 | 0.23615326 | 0.67182855 |
| Cav2 | 608.320272 | 0.78820814 | 0.36718422 | 2.14662856 | 0.03182286 | 0.26554509 |
| Klf6 | 25845.2923 | 0.362959 | 0.24106859 | 1.5056254 | 0.13216336 | 0.55179332 |
| Scmh1 | 581.615652 | -0.5021714 | 0.31080934 | -1.6156896 | 0.10616143 | 0.49777747 |
| Cox5a | 7519.93258 | 0.07483182 | 0.23252333 | 0.32182499 | 0.74758528 | 0.9399191 |
| Fer | 1054.07795 | 0.9470587 | 0.38614203 | 2.4526175 | 0.01418211 | 0.16198851 |
| Xpo6 | 1019.22114 | -0.434768 | 0.37544109 | -1.1580191 | 0.24685622 | 0.68182258 |
| Tfe3 | 3349.50648 | 0.19405016 | 0.32374771 | 0.59938698 | 0.54891486 | 0.87082325 |
| Brat1 | 1137.10275 | -0.0992581 | 0.39494805 | -0.2513194 | 0.80156716 | 0.95629571 |
| Gna12 | 988.954794 | 0.2399268 | 0.31844213 | 0.75343926 | 0.451186 | 0.82550977 |
| Slc22a18 | 22.1394756 | 0.87628473 | 1.37224531 | 0.63857731 | 0.52309794 | 0.85870332 |
| Pih1d2 | 34.5497818 | 0.25426726 | 0.82021789 | 0.30999965 | 0.75656122 | 0.94102069 |
| Dlat | 2743.88943 | -0.3972529 | 0.27923758 | -1.4226342 | 0.15484224 | 0.58390415 |
| Sdhd | 15976.2061 | 0.52424847 | 0.36494473 | 1.43651471 | 0.15085593 | 0.57859667 |
| Ccnd2 | 4802.11202 | -1.4810003 | 0.26173223 | -5.6584561 | 1.527407834 | 1.962073271 |
| Gpr107 | 1766.96923 | 0.25512853 | 0.19757338 | 1.2913103 | 0.1965961 | 0.62907667 |
| Tspan32 | 1634.00429 | 0.40107352 | 0.29291103 | 1.36926739 | 0.17091571 | 0.60430173 |
| Gmpr | 1245.32855 | 0.18847297 | 0.23916149 | 0.78805736 | 0.43066315 | 0.81254302 |
| Trim25 | 767.182243 | 0.05206362 | 0.34936391 | 0.14902404 | 0.88153467 | 0.97582756 |
| Dgke | 109.689816 | 0.75837799 | 0.56753764 | 1.33626025 | 0.18146422 | 0.61271097 |
| Scpep1 | 5382.53078 | 0.70588522 | 0.16318583 | 4.32565261 | 1.520809357 | 0.00075182 |
| Mnt | 119.019909 | -0.4529756 | 0.65685979 | -0.6896077 | 0.4904409 | 0.84472295 |
| Itgb2 | 5898.55663 | -0.783884 | 0.1585656 | -4.9435943 | 7.669522839 | 5.791436598 |
| Hddc2 | 1612.53835 | 0.31692104 | 0.46007589 | 0.68884515 | 0.49092072 | 0.84477968 |
| Tpd52l1 | 18.2865598 | -1.3034954 | 1.51279198 | -0.8616488 | 0.38888081 | 0.78942657 |
| Pemt | 205.119793 | -0.4525819 | 0.40624759 | -1.1140544 | 0.26525586 | 0.7011398 |
| Bcl6b | 39.0870426 | 4.31251514 | 1.68514209 | 2.55914036 | 0.01049314 | 0.1323572 |
| Clec10a | 191.072612 | 2.15455384 | 0.52527977 | 4.10172628 | 4.100792153 | 0.00171798 |
| Arvcf | 45.3079522 | 1.61054537 | 1.35906648 | 1.18503796 | 0.23600244 | 0.67182855 |
| Comt | 3970.34858 | 0.68790223 | 0.2098129 | 3.27864601 | 0.00104306 | 0.02416629 |
| Rtca | 2035.48702 | -0.4054009 | 0.19991403 | -2.0278764 | 0.04257286 | 0.31638902 |
| Dbt | 837.532946 | 0.62418747 | 0.28166503 | 2.21606301 | 0.02668719 | 0.24040085 |
| Dazap2 | 18484.2836 | -0.0794818 | 0.17391499 | -0.4570152 | 0.64766012 | 0.9097332 |
| Mcts1 | 2507.16619 | -0.1778756 | 0.26213353 | -0.6785687 | 0.49741116 | 0.84818511 |
| Rem1 | 8.48335144 | -0.867681 | 1.60938553 | -0.5391381 | 0.5897916 | 0.88722166 |
| Rnf17 | 242.139742 | 1.03185693 | 0.41016291 | 2.51572465 | 0.01187879 | 0.14416002 |
| Trappc10 | 64.9177977 | -1.0927212 | 1.12801206 | -0.9687141 | 0.33268784 | 0.75185117 |
| Ccm2 | 2032.86471 | 0.25951407 | 0.20477981 | 1.26728348 | 0.20505394 | 0.63913241 |
| Tbrg4 | 2694.58369 | -0.2713189 | 0.18368113 | -1.4771192 | 0.13964369 | 0.56329174 |
| Mx1 | 196.704188 | -1.8494426 | 0.48903586 | -3.7818138 | 0.00015569 | 0.0052467 |
| Fap | 10.353117 | 4.75287058 | 2.01543151 | 2.35823969 | 0.01836183 | 0.19092573 |
| Ndufa9 | 9519.75729 | 0.15525373 | 0.2607842 | 0.59533413 | 0.55162015 | 0.87093412 |

| | | | | | | |
|---|---|---|---|---|---|---|
| Lck | 67.354001 | 4.19492252 | 1.56751277 | 2.67616482 | NA | NA |
| Galnt1 | 1384.65657 | -0.1187259 | 0.23558101 | -0.5039707 | 0.61428194 | 0.89843508 |
| Mkrn2 | 1317.81926 | -0.0828418 | 0.34053423 | -0.2432703 | 0.80779604 | 0.95809279 |
| Pparg | 563.057196 | -0.7406644 | 0.50484889 | -1.4671013 | 0.14234851 | 0.56795476 |
| Raf1 | 4302.34979 | -0.0517104 | 0.18226519 | -0.2837099 | 0.77663269 | 0.94800948 |
| Sept1 | 17.0178547 | -0.6000208 | 0.9457072 | -0.6344678 | 0.52577557 | 0.86056156 |
| Pdgfb | 11.5564216 | 1.77095549 | 1.97146194 | 0.89829555 | 0.369028 | 0.77480275 |
| Acvrl1 | 707.467658 | 0.4225364 | 0.25196931 | 1.67693599 | 0.093555 | 0.46750793 |
| Grasp | 35.6579307 | -0.9001099 | 1.30346235 | -0.6905531 | 0.48984646 | 0.84448327 |
| Acvr1b | 64.958083 | -0.4330266 | 0.69486198 | -0.6231836 | 0.5331639 | 0.86489775 |
| Tom1l2 | 271.183962 | 0.19880805 | 0.42081385 | 0.47243706 | 0.63661486 | 0.90542378 |
| Zfp385a | 452.757038 | -0.1860154 | 0.24621963 | -0.7554856 | 0.44995769 | 0.82484234 |
| Itga5 | 1778.6453 | 0.67341819 | 0.4364745 | 1.54285804 | 0.12286522 | 0.53283544 |
| Wdr77 | 7184.66132 | -0.6225789 | 0.20058752 | -3.1037769 | 0.00191067 | 0.03885434 |
| Adora3 | 78.1107875 | -2.3824148 | 0.62844908 | -3.7909433 | 0.00015008 | 0.00508555 |
| Atp5f1 | 21884.8674 | 0.09282544 | 0.31398875 | 0.29563301 | 0.76751036 | 0.94401697 |
| Sox9 | 17.8213155 | 1.89659244 | 1.61245844 | 1.17621167 | 0.2395103 | 0.67562272 |
| Hnrnpd | 814.147306 | -0.6599432 | 0.26355322 | -2.5040226 | 0.01227902 | 0.14755327 |
| Dynlt1c | 423.126426 | 0.72756782 | 0.3030899 | 2.40050171 | 0.01637261 | 0.17834923 |
| C1d | 4193.88822 | 0.42933912 | 0.32348027 | 1.32724981 | 0.18442606 | 0.61658486 |
| Gm2a | 5634.80528 | 0.09404243 | 0.17064973 | 0.55108457 | 0.5815757 | 0.88321857 |
| Krit1 | 974.027078 | -0.1711021 | 0.28125212 | -0.6083584 | 0.54294982 | 0.86919453 |
| Clcn4 | 2294.24334 | 0.14197677 | 0.18961571 | 0.74876062 | 0.4540015 | 0.82695062 |
| Hk2 | 2091.29501 | -0.4179547 | 0.34103482 | -1.2255485 | 0.22036863 | 0.65381019 |
| Myo18a | 559.027532 | -0.1953938 | 0.30087586 | -0.6494168 | 0.51606904 | 0.85627177 |
| Haao | 242.056027 | -2.3885448 | 0.48595906 | -4.9151154 | 8.873018504 | 6.538773214 |
| Cd52 | 5768.40107 | -0.6341903 | 0.32372305 | -1.9590519 | 0.05010671 | 0.34262459 |
| Abhd15 | 495.036694 | -0.7716678 | 0.30885506 | -2.4984787 | 0.01247277 | 0.14871281 |
| Loxl3 | 437.211563 | 1.66612731 | 0.5181955 | 3.21524851 | 0.00130332 | 0.02831877 |
| Kat2b | 159.514277 | -0.198225 | 0.8764373 | -0.2261714 | 0.82106814 | 0.96205421 |
| Rab5b | 1945.88247 | 0.23279293 | 0.15762421 | 1.47688563 | 0.1397063 | 0.56329174 |
| Icosl | 118.215561 | 0.91745846 | 0.53097831 | 1.72786429 | 0.08401256 | 0.44471119 |
| Spg7 | 292.026983 | 0.15932569 | 0.36016152 | 0.44237288 | 0.6582194 | 0.91258818 |
| Rpl13 | 39032.5152 | 0.00835606 | 0.30120881 | 0.02774174 | 0.97786813 | 0.99483019 |
| Chmp1a | 4621.8206 | 0.01512247 | 0.19727469 | 0.07665693 | 0.93889647 | 0.98910786 |
| Rpa1 | 12384.8513 | -0.2034479 | 0.14132768 | -1.4395477 | 0.14999541 | 0.57784353 |
| Serpinf1 | 240.37605 | -0.775754 | 0.44424211 | -1.7462415 | 0.080769 | 0.43854735 |
| Tubgcp3 | 1444.20212 | -0.4658555 | 0.23082023 | -2.0182613 | 0.04356406 | 0.31968755 |
| Polr3d | 666.476044 | 0.03952449 | 0.36569942 | 0.10807916 | 0.9139329 | 0.9832153 |
| Ddx3x | 5454.63664 | -0.3783899 | 0.22626079 | -1.6723618 | 0.09445301 | 0.47064914 |
| Il12rb1 | 28.4763285 | -1.1467196 | 0.78336766 | -1.4638332 | 0.14323953 | 0.56846902 |
| Usp32 | 104.755318 | -0.6132447 | 0.47169129 | -1.3000976 | 0.19356753 | 0.62506533 |
| Txnrd3 | 197.603325 | 0.34203506 | 0.42402465 | 0.80663959 | 0.41987416 | 0.80569735 |
| Zfp512b | 620.552076 | 0.13221503 | 0.28474642 | 0.46432551 | 0.64241455 | 0.90715193 |
| Dnajc5 | 3910.3234 | 0.15028983 | 0.23590444 | 0.63707927 | 0.52407321 | 0.8594073 |
| Tpd52l2 | 4924.34076 | 0.42947862 | 0.33106204 | 1.29727536 | 0.19453646 | 0.62651174 |

| | | | | | | |
|---|---|---|---|---|---|---|
| Fmr1 | 512.659797 | -0.1159137 | 0.38332079 | -0.3023934 | 0.76235215 | 0.94298237 |
| Bcl11a | 101.011365 | -1.8511919 | 0.70691993 | -2.6186726 | 0.00882726 | 0.11905611 |
| Pxmp4 | 764.303914 | 0.04106389 | 0.25470636 | 0.16122051 | 0.87191972 | 0.97298927 |
| Gnb1l | 396.655981 | -0.126613 | 0.31259405 | -0.4050396 | 0.6854484 | 0.92314423 |
| Mmp11 | 10.2277065 | 0.02452648 | 2.68612097 | 0.00913082 | 0.99271476 | 0.99800348 |
| Smarcb1 | 4432.93737 | -0.0808551 | 0.16234472 | -0.4980456 | 0.61845193 | 0.89878056 |
| Hip1r | 39.6691247 | -1.3728022 | 0.92572944 | -1.4829411 | 0.13809007 | 0.56068131 |
| Nsun5 | 1106.79116 | -0.3399603 | 0.25858859 | -1.3146764 | 0.18861866 | 0.61935434 |
| Top1mt | 589.17149 | -0.6641826 | 0.28536618 | -2.3274748 | 0.01994001 | 0.19988038 |
| Mmp14 | 40.7387941 | 0.28267478 | 0.86006016 | 0.32866862 | 0.74240618 | 0.93840202 |
| Slc7a7 | 1786.85742 | 0.90831998 | 0.21616234 | 4.20202693 | 2.645356574 | 0.00115162 |
| Oxa1l | 1777.91155 | -0.2227953 | 0.16497425 | -1.3504855 | 0.17686031 | 0.60784615 |
| Heatr6 | 909.063783 | -0.3520394 | 0.30651031 | -1.1485402 | 0.25074563 | 0.6842534 |
| Ccl3 | 10669.025 | 0.38053371 | 0.40018429 | 0.95089617 | 0.34165709 | 0.75866343 |
| Ilf2 | 6107.90451 | 0.06050292 | 0.31843443 | 0.19000118 | 0.84930821 | 0.96765585 |
| Chtop | 5078.25738 | -0.0667116 | 0.18959974 | -0.3518552 | 0.72494689 | 0.93542036 |
| Snapin | 2338.8902 | 0.39354655 | 0.21945706 | 1.79327362 | 0.07292918 | 0.41525386 |
| S100a4 | 301.999254 | 1.48224611 | 0.62365679 | 2.37670164 | 0.01746821 | 0.18560042 |
| S100a6 | 548.811882 | 0.84668501 | 0.3937924 | 2.15007959 | 0.03154892 | 0.26539272 |
| Icam2 | 1275.60629 | -0.2969996 | 0.3030882 | -0.9799115 | 0.3271298 | 0.74811039 |
| Mapk7 | 1308.64038 | -0.2363547 | 0.24534617 | -0.9633519 | 0.33537096 | 0.75359284 |
| Epn2 | 114.179069 | 0.89948506 | 0.65818861 | 1.36660685 | 0.17174857 | 0.60430173 |
| B9d1 | 131.454622 | 1.24242952 | 0.64735271 | 1.91924666 | 0.05495312 | 0.36084578 |
| Sec24b | 2024.27874 | -0.2027586 | 0.30361545 | -0.6678139 | 0.50425239 | 0.85012672 |
| N4bp3 | 72.2602772 | 0.12814904 | 0.9964042 | 0.12861151 | 0.89766506 | 0.97984623 |
| Rmnd5b | 1214.23898 | -0.402795 | 0.19400022 | -2.0762605 | 0.03786987 | 0.29387688 |
| Nhp2 | 8182.91825 | -0.0040951 | 0.32646229 | -0.0125438 | 0.9899918 | 0.99650865 |
| Vps9d1 | 279.186722 | -0.0915932 | 0.32485284 | -0.281953 | 0.77797957 | 0.94837927 |
| Zfp276 | 250.888987 | -0.3024599 | 0.38006923 | -0.7958021 | 0.42614707 | 0.81021713 |
| Mfsd10 | 2703.24547 | -0.4377566 | 0.28742912 | -1.5230071 | 0.12775692 | 0.54227156 |
| Luzp1 | 653.85884 | 0.27183432 | 0.35616572 | 0.76322428 | 0.44532965 | 0.82118815 |
| Slc13a2 | 23.39254 | 2.33190117 | 1.25159121 | 1.8631492 | 0.06244128 | 0.38558513 |
| Kctd10 | 4243.95754 | 0.24662541 | 0.22156933 | 1.11308459 | 0.2656721 | 0.70163359 |
| Poldip2 | 4448.00461 | 0.28308463 | 0.24759007 | 1.1433602 | 0.25288907 | 0.68647781 |
| Ift20 | 845.030004 | 0.35274767 | 0.33575452 | 1.05061184 | 0.2934369 | 0.72646828 |
| Pcbp3 | 19.2752777 | 0.82054956 | 1.2462268 | 0.65842715 | 0.51026369 | 0.85335831 |
| Lgals9 | 5022.64554 | 0.07540021 | 0.18906008 | 0.39881614 | 0.69002868 | 0.92470267 |
| Araf | 2368.54148 | -0.0095641 | 0.18361555 | -0.0520876 | 0.95845885 | 0.99249408 |
| Cfp | 5363.41388 | 0.01566462 | 0.17449748 | 0.08976986 | 0.9284701 | 0.98666423 |
| Uxt | 821.957529 | 0.35463328 | 0.43229584 | 0.82034859 | 0.41201741 | 0.80245328 |
| Cnnm3 | 176.123422 | -0.0593766 | 0.59230353 | -0.1002469 | 0.92014829 | 0.9839023 |
| Lman2l | 1977.37317 | -0.3077962 | 0.21022892 | -1.4641002 | 0.14316657 | 0.56846902 |
| Mcm3ap | 408.637724 | -0.313753 | 0.30965958 | -1.0132192 | 0.31095547 | 0.74011268 |
| Pcnt | 861.919843 | 0.09283178 | 0.44036281 | 0.21080749 | 0.83303749 | 0.96565375 |
| Mxd1 | 994.982803 | 0.19714299 | 0.4644542 | 0.42446164 | 0.67122918 | 0.91711586 |
| Gmcl1 | 221.64562 | 0.56176872 | 0.54479127 | 1.03116321 | 0.30246429 | 0.73372823 |

| | | | | | | |
|---|---|---|---|---|---|---|
| Snrnp27 | 1168.37679 | 0.28933303 | 0.32766205 | 0.88302271 | 0.377224 | 0.7799581 |
| Ocrl | 493.159292 | -0.0909193 | 0.31667418 | -0.2871067 | 0.77403061 | 0.94684176 |
| Calm1 | 41272.6671 | 0.26853478 | 0.23169825 | 1.15898493 | 0.24646233 | 0.68165807 |
| Agpat3 | 388.351347 | 1.73680972 | 0.35601057 | 4.8785341 | 1.068771874 | 7.645781482 |
| Sema6b | 30.4106636 | -0.0904879 | 1.09681662 | -0.0825005 | 0.93424872 | 0.98756396 |
| Uhrf1 | 14997.0499 | -0.2367569 | 0.26265582 | -0.9013959 | 0.36737786 | 0.77325076 |
| Dpp9 | 2515.72621 | -0.3780553 | 0.31555883 | -1.1980501 | 0.23089753 | 0.6653874 |
| Ramp2 | 147.007843 | 1.46901253 | 0.80043652 | 1.83526425 | 0.06646655 | 0.39699582 |
| Lsr | 20.2716308 | -1.0205184 | 1.41666378 | -0.7203674 | 0.47129882 | 0.83393588 |
| Gramd1a | 1042.50261 | 0.5386854 | 0.22327628 | 2.41264057 | 0.01583743 | 0.17485494 |
| Hpn | 315.676696 | 1.14920357 | 0.31105331 | 3.69455496 | 0.00022027 | 0.0070539 |
| Ckb | 30704.8977 | 0.30561383 | 0.21179593 | 1.44296364 | 0.14903071 | 0.57638088 |
| Sp1 | 360.364028 | -0.2298999 | 0.30955089 | -0.7426885 | 0.45767027 | 0.82968569 |
| Itgb7 | 164.688852 | -2.9244942 | 0.40925739 | -7.1458555 | 8.943709424 | 2.668497497 |
| Myg1 | 2234.58277 | 0.0012469 | 0.25292572 | 0.00492992 | 0.99606651 | 0.99884675 |
| Rarg | 47.7740684 | -2.3301675 | 0.85257804 | -2.7330841 | 0.00627443 | 0.09487653 |
| Pfdn5 | 3251.81403 | 0.74400898 | 0.5305012 | 1.40246428 | 0.16077665 | 0.59115743 |
| Rrp15 | 2686.21022 | -0.5975338 | 0.2099244 | -2.8464237 | 0.00442133 | 0.07285835 |
| Rnd2 | 20.034271 | -0.5618882 | 1.4814473 | -0.3792833 | 0.70447752 | 0.9285501 |
| Srr | 411.526692 | 0.16322819 | 0.34439725 | 0.47395322 | 0.63553327 | 0.90531997 |
| Acp5 | 413.043241 | -0.5587532 | 0.34860732 | -1.6028154 | 0.10897542 | 0.50470898 |
| Fbxo9 | 845.857547 | -0.3828745 | 0.20182701 | -1.8970431 | 0.05782225 | 0.36898259 |
| Vps50 | 2512.40055 | 0.0128053 | 0.17913853 | 0.07148266 | 0.94301362 | 0.98983253 |
| Hars | 5627.91599 | -0.2754459 | 0.15885791 | -1.7339135 | 0.08293344 | 0.4429983 |
| Zmat2 | 2178.77692 | -0.0235742 | 0.16200962 | -0.1455111 | 0.88430737 | 0.97721157 |
| Ube2c | 9007.79592 | -0.3024063 | 0.15224484 | -1.9863158 | 0.04699827 | 0.3347601 |
| Smg5 | 1757.47438 | 0.1474791 | 0.31142264 | 0.47356577 | 0.63580959 | 0.90531997 |
| Cct3 | 31613.401 | 0.05704139 | 0.2805875 | 0.20329271 | 0.83890626 | 0.96750639 |
| Glmp | 6447.76391 | 0.46775979 | 0.25290827 | 1.84952347 | 0.06438226 | 0.39202997 |
| Mef2d | 608.241258 | -0.2727549 | 0.62268779 | -0.4380284 | 0.66136572 | 0.91376631 |
| Tmem79 | 19.6588557 | 0.72100393 | 0.96488928 | 0.74724006 | 0.45491867 | 0.82763534 |
| Snd1 | 6184.89125 | -0.2032511 | 0.1763737 | -1.1523891 | 0.24916122 | 0.68371226 |
| Col18a1 | 174.705635 | 1.64429748 | 1.07741812 | 1.52614612 | 0.12697348 | 0.54111158 |
| Slc19a1 | 2047.6313 | -0.4723764 | 0.23778396 | -1.9865783 | 0.04696915 | 0.3347601 |
| Kpnb1 | 7306.89113 | -0.4171886 | 0.23628704 | -1.765601 | 0.07746282 | 0.43008322 |
| Npepps | 1265.56227 | -0.0505402 | 0.26522912 | -0.1905531 | 0.84887574 | 0.96765585 |
| Mrpl10 | 3470.66928 | -0.0574568 | 0.19257274 | -0.298364 | 0.76542538 | 0.94347227 |
| Cyp51 | 4629.37477 | 0.50902692 | 0.25313313 | 2.01090597 | 0.04433539 | 0.32302252 |
| Tcf25 | 6627.28044 | -0.0273755 | 0.2093473 | -0.130766 | 0.89596042 | 0.97984623 |
| Tubb6 | 14401.7473 | 0.18725102 | 0.22336817 | 0.83830665 | 0.4018585 | 0.79752354 |
| Def8 | 1180.44049 | -0.1721743 | 0.23735739 | -0.7253801 | 0.4682189 | 0.83381643 |
| Meox1 | 27.948891 | 1.97997114 | 1.06477616 | 1.85951866 | 0.06295366 | 0.38709283 |
| Col1a1 | 52.8602243 | 1.04915385 | 1.26224428 | 0.83118131 | 0.40587121 | 0.79990697 |
| Foxm1 | 881.261523 | -0.2516845 | 0.35285 | -0.7132903 | 0.47566615 | 0.83639059 |
| Itfg2 | 496.079925 | 0.01450224 | 0.2390674 | 0.06066173 | 0.95162861 | 0.99116839 |
| Nrip2 | 28.3840335 | 0.75965251 | 1.1407347 | 0.66593268 | 0.50545413 | 0.85039477 |

| | | | | | | |
|---|---|---|---|---|---|---|
| Tulp3 | 73.0417658 | -0.8500904 | 0.56724517 | -1.4986296 | 0.13396975 | 0.55441542 |
| Gtf2h4 | 1494.59842 | -0.1326762 | 0.21908758 | -0.6055852 | 0.54479026 | 0.86963259 |
| Tubb5 | 89236.8208 | -0.2213626 | 0.15055341 | -1.4703261 | 0.14147347 | 0.56628837 |
| Ell2 | 386.119989 | 0.1481202 | 0.55034776 | 0.26913927 | 0.78782251 | 0.95225386 |
| Jup | 1262.45059 | 0.34450781 | 0.27090849 | 1.27167595 | 0.20348828 | 0.63809363 |
| Fkbp10 | 50.1738006 | 1.02242467 | 0.90864656 | 1.12521713 | 0.26049704 | 0.69666911 |
| Nom1 | 1884.43916 | -0.1936816 | 0.29489819 | -0.6567744 | 0.51132597 | 0.85373917 |
| Ergic1 | 3328.72241 | -0.2266849 | 0.15851054 | -1.4300937 | 0.15269011 | 0.58090075 |
| Acap1 | 100.455018 | -3.3258444 | 0.828159 | -4.0159491 | 5.920702869 | 0.00232141 |
| Tcea3 | 25.536979 | -0.3129515 | 0.83476816 | -0.3748963 | 0.70773761 | 0.92950127 |
| Ifrd1 | 3821.93533 | -0.0814368 | 0.33225041 | -0.2451068 | 0.80637376 | 0.95741157 |
| Stk38l | 678.24834 | 0.37475987 | 0.64988542 | 0.57665531 | 0.56417233 | 0.87694029 |
| Brpf1 | 996.771058 | -0.2880055 | 0.33329632 | -0.8641123 | 0.38752621 | 0.78918063 |
| Akr1b3 | 10965.8984 | 0.13320826 | 0.29717281 | 0.44825185 | 0.65397144 | 0.91076467 |
| Gstt3 | 40.1120629 | -0.2417849 | 0.77772215 | -0.3108885 | 0.75588535 | 0.94102069 |
| Ddt | 228.000187 | 0.31616726 | 0.40944235 | 0.77218991 | 0.44000196 | 0.81963526 |
| Ddx18 | 2902.5378 | -0.3684341 | 0.24547823 | -1.5008832 | 0.13338579 | 0.55359482 |
| Ubl3 | 2918.92152 | 0.34383693 | 0.22026701 | 1.56100054 | 0.11852362 | 0.52515149 |
| Gramd3 | 237.725291 | 0.612502 | 0.30875848 | 1.98375764 | 0.04728286 | 0.33531089 |
| Eef1e1 | 3024.43121 | -0.0374378 | 0.36279563 | -0.1031926 | 0.91781014 | 0.98390186 |
| Akt1 | 1442.8395 | -0.1594671 | 0.1871874 | -0.8519116 | 0.39426315 | 0.79233901 |
| Cldn15 | 98.8655981 | -7.7580196 | 1.04005225 | -7.4592594 | 8.701014422 | 2.956653039 |
| Il16 | 1965.04208 | -0.2302092 | 0.26834463 | -0.8578864 | 0.39095518 | 0.7899826 |
| Tcirg1 | 2886.8753 | -0.0058819 | 0.26355647 | -0.0223176 | 0.98219462 | 0.99523682 |
| Naglu | 1129.06136 | 0.83931636 | 0.19097207 | 4.39496918 | 1.107884729 | 0.00056706 |
| Coasy | 1593.8731 | 0.19051392 | 0.20721018 | 0.91942357 | 0.35787406 | 0.76674184 |
| Smo | 433.593927 | 0.43503378 | 0.3027951 | 1.43672663 | 0.15079569 | 0.57859667 |
| Tspan33 | 13.3206147 | 0.08441642 | 1.24274579 | 0.06792734 | 0.94584347 | 0.99019648 |
| Crnkl1 | 2112.95429 | -0.0197355 | 0.19173782 | -0.1029294 | 0.91801901 | 0.98390186 |
| Rin2 | 7575.82531 | 0.64367071 | 0.18795492 | 3.42460155 | 0.0006157 | 0.01595737 |
| Chordc1 | 4860.24294 | 0.05730844 | 0.17385124 | 0.32964068 | 0.74167148 | 0.93800224 |
| Rtcb | 9288.96116 | -0.2317951 | 0.16205389 | -1.430358 | 0.15261428 | 0.58090075 |
| Pwp1 | 6494.36996 | -0.0823335 | 0.19571054 | -0.4206901 | 0.6739814 | 0.91822108 |
| Fbxo7 | 1398.25297 | 0.03917669 | 0.196324 | 0.1995512 | 0.8418316 | 0.96765585 |
| Capns1 | 496.762649 | 0.33519592 | 0.27243667 | 1.23036274 | 0.2185613 | 0.65195327 |
| Clpb | 2642.67003 | 0.04598379 | 0.19050642 | 0.24137659 | 0.80926325 | 0.95844313 |
| Sept7 | 10409.2317 | 0.28561184 | 0.24988597 | 1.14296872 | 0.25305158 | 0.68647781 |
| Zdhhc4 | 917.808768 | -0.2163182 | 0.22171557 | -0.9756564 | 0.32923477 | 0.74874249 |
| Rac1 | 25842.0536 | -0.0064139 | 0.1907475 | -0.0336253 | 0.97317596 | 0.99443087 |
| Nup214 | 1156.1476 | -0.0261047 | 0.2627428 | -0.0993547 | 0.92085664 | 0.9839023 |
| Aif1l | 43.5414519 | 1.21404936 | 1.19717051 | 1.01409895 | 0.31053554 | 0.7399039 |
| Ugp2 | 3130.27929 | -0.0158844 | 0.19700344 | -0.0806303 | 0.93573598 | 0.98794094 |
| Trmt1 | 2508.57772 | -0.3382686 | 0.19908767 | -1.6990937 | 0.08930153 | 0.4566019 |
| Nacc1 | 186.609107 | 0.32496452 | 0.44602372 | 0.72858126 | 0.46625785 | 0.83345004 |
| Slc1a5 | 2940.96145 | -0.5762593 | 0.21719957 | -2.6531328 | 0.00797485 | 0.11048282 |
| Uba1 | 13907.9139 | -0.2064828 | 0.17310654 | -1.1928074 | 0.2329448 | 0.66845266 |

| | | | | | | |
|---|---|---|---|---|---|---|
| Vwf | 14.8753114 | 0.5572029 | 1.79453559 | 0.31049978 | 0.75618093 | 0.94102069 |
| Siae | 892.937936 | 0.37806641 | 0.32180036 | 1.17484772 | 0.24005565 | 0.67563574 |
| Esam | 150.744059 | 3.12047121 | 2.26905034 | 1.37523225 | 0.16905945 | 0.60312169 |
| Spa17 | 33.2550999 | 0.27439931 | 0.80192082 | 0.34217756 | 0.73221727 | 0.93704508 |
| Fam50a | 1617.88682 | 0.40763157 | 0.22471745 | 1.81397385 | 0.06968175 | 0.40760152 |
| Emd | 549.144972 | 0.29303274 | 0.30790503 | 0.95169847 | 0.34124992 | 0.75846782 |
| Taco1 | 382.586192 | 0.87361887 | 0.4608377 | 1.8957192 | 0.05799719 | 0.36990593 |
| Gria3 | 488.141551 | -5.3177187 | 0.38184774 | -13.92628 | 4.386105066 | 1.788507442 |
| Sipa1l2 | 292.90554 | 0.13610545 | 0.45453256 | 0.29944049 | 0.76460398 | 0.94307414 |
| Ap4e1 | 359.283104 | 0.02990897 | 0.47954925 | 0.06236892 | 0.95026904 | 0.99069447 |
| Blvra | 4544.17376 | 1.27115194 | 0.34434397 | 3.69151799 | 0.00022292 | 0.00710571 |
| Srpk3 | 19.8492991 | 2.07631904 | 1.79796341 | 1.15481718 | 0.2481653 | 0.68235696 |
| Idh3g | 9869.93672 | -0.2685944 | 0.23300289 | -1.1527514 | 0.24901246 | 0.68345734 |
| Pnck | 9.86197436 | -6.7041493 | 1.97255001 | -3.3987221 | 0.00067701 | 0.01729002 |
| Ssr4 | 4515.36053 | 0.16525321 | 0.41242998 | 0.40068185 | 0.68865437 | 0.92470267 |
| Bcap31 | 8661.68161 | 0.12787725 | 0.27801075 | 0.45997231 | 0.64553609 | 0.90847224 |
| Fam98a | 1195.59847 | -0.3984026 | 0.21795785 | -1.8278883 | 0.06756631 | 0.40045479 |
| Kmt2a | 179.556636 | 0.13299838 | 0.57521673 | 0.23121437 | 0.81714826 | 0.96079676 |
| Ift46 | 788.858866 | 0.3064351 | 0.25916126 | 1.18241091 | 0.2370427 | 0.6732629 |
| Trappc6a | 1383.14367 | 0.36567679 | 0.3337414 | 1.09568902 | 0.27321489 | 0.71005689 |
| Supt6 | 1380.94229 | -0.0606234 | 0.20183225 | -0.3003653 | 0.76389851 | 0.94300309 |
| Spag5 | 1642.69715 | -0.2428779 | 0.17424446 | -1.3938916 | 0.16335035 | 0.59475807 |
| Unc119 | 299.473707 | -0.5293759 | 0.56391409 | -0.9387528 | 0.34785766 | 0.76101142 |
| Rab34 | 339.860424 | 0.77775519 | 0.33337246 | 2.33299174 | 0.01964858 | 0.19864551 |
| Sdf2 | 1618.55431 | 0.41272609 | 0.3699497 | 1.11562757 | 0.26458159 | 0.70112124 |
| Ccne1 | 2053.85081 | -0.3051704 | 0.21455524 | -1.4223393 | 0.15492777 | 0.58404664 |
| Bbc3 | 238.663987 | 0.21942975 | 0.44724784 | 0.49062228 | 0.62369362 | 0.90017327 |
| Psmc3 | 14014.5425 | 0.31424924 | 0.28985473 | 1.08416115 | 0.27829335 | 0.71360814 |
| Acp2 | 3837.86175 | 0.24047504 | 0.17200086 | 1.39810375 | 0.16208191 | 0.59315429 |
| Rapsn | 216.241173 | 0.95638113 | 0.54318539 | 1.76069009 | 0.07829087 | 0.43258005 |
| Slc39a13 | 470.06656 | 0.36372989 | 0.31407152 | 1.15811165 | 0.24681848 | 0.68182258 |
| Celf2 | 7499.17267 | -0.2018346 | 0.26118403 | -0.7727676 | 0.43965994 | 0.81963526 |
| Nr1h3 | 266.730349 | 2.91288283 | 0.54240032 | 5.37035602 | 7.858134982 | 8.078030692 |
| Ddb2 | 102.361134 | 0.08034846 | 0.80003864 | 0.10043072 | 0.92000238 | 0.9839023 |
| Spi1 | 14774.0655 | -0.0754691 | 0.16502706 | -0.4573136 | 0.64744569 | 0.9097332 |
| Sf3a1 | 4620.29303 | -0.2174847 | 0.20811537 | -1.04502 | 0.29601372 | 0.72859877 |
| Stat6 | 6974.88467 | -0.0531316 | 0.2446642 | -0.2171611 | 0.82808277 | 0.96434151 |
| Clgn | 88.6356989 | 2.25282676 | 1.33369555 | 1.68916119 | 0.09118854 | 0.46133559 |
| Napsa | 708.68274 | -4.424255 | 0.39991875 | -11.062885 | 1.898905360 | 2.903663659 |
| Vrk3 | 2209.96773 | -0.2278985 | 0.20392521 | -1.1175592 | 0.26375531 | 0.70008189 |
| Smg9 | 657.805616 | 0.15452467 | 0.35496783 | 0.43532021 | 0.66333002 | 0.91395288 |
| Paxip1 | 958.92354 | -0.5602575 | 0.3885694 | -1.4418466 | 0.14934564 | 0.57685486 |
| Rmnd5a | 405.282958 | 0.18118078 | 0.3971384 | 0.45621573 | 0.64823486 | 0.90973357 |
| Mov10 | 1872.31824 | -0.1723865 | 0.21974031 | -0.7845009 | 0.43274624 | 0.81435873 |
| Ppm1j | 87.5562001 | -0.2051971 | 0.59888884 | -0.3426297 | 0.73187707 | 0.9367077 |
| Rhoc | 1619.1362 | 1.22649641 | 0.22808384 | 5.37739278 | 7.557219064 | 7.834530577 |

| | | | | | | |
|---|---|---|---|---|---|---|
| Ppard | 23.2644652 | 1.00281759 | 0.93575065 | 1.07167181 | 0.28386747 | 0.71951357 |
| Def6 | 1834.50839 | -0.759971 | 0.19125158 | -3.973672 | 7.077300776 | 0.002714 |
| Peg3 | 397.751732 | 1.18166838 | 0.53592695 | 2.2049057 | 0.02746071 | 0.24574023 |
| Metrn | 143.41259 | -0.7143071 | 0.37151627 | -1.9226805 | 0.05452019 | 0.36033644 |
| Lmf1 | 1054.33069 | 0.01244656 | 0.18483394 | 0.06733912 | 0.94631173 | 0.99019648 |
| Narfl | 2604.80412 | 0.30527841 | 0.2470711 | 1.23558932 | 0.21661126 | 0.65009949 |
| Angptl4 | 1360.53616 | -0.0648646 | 0.22347786 | -0.2902507 | 0.77162449 | 0.94526087 |
| Dbf4 | 2096.68669 | -0.0725065 | 0.28869352 | -0.2511537 | 0.80169525 | 0.95629571 |
| Daxx | 1629.12057 | -0.5026261 | 0.2136474 | -2.3525966 | 0.01864285 | 0.19261651 |
| Cd320 | 954.351961 | 0.05181715 | 0.31280452 | 0.16565346 | 0.86842967 | 0.97186901 |
| Ipo4 | 3647.15502 | -0.3076442 | 0.21487482 | -1.4317369 | 0.15221914 | 0.58009243 |
| Tm9sf1 | 3060.81571 | 0.04520494 | 0.18143284 | 0.24915525 | 0.80324069 | 0.95672466 |
| Irf9 | 937.525191 | -0.2252587 | 0.19829618 | -1.135971 | 0.25596869 | 0.69061865 |
| Gmpr2 | 6645.88612 | 0.38178919 | 0.31097686 | 1.22770932 | 0.2195561 | 0.65289787 |
| Mdp1 | 1052.77189 | 0.64166884 | 0.47108478 | 1.36210905 | 0.17316347 | 0.60592355 |
| Dhrs1 | 2375.33633 | 0.04654926 | 0.16994802 | 0.27390293 | 0.78415922 | 0.95094799 |
| Tmem161a | 1345.56037 | -0.1402229 | 0.19853804 | -0.7062772 | 0.48001574 | 0.83935746 |
| Armc6 | 1275.45621 | -0.2861209 | 0.24997741 | -1.1445869 | 0.25238032 | 0.68582922 |
| Borcs8 | 485.190425 | -0.1557721 | 0.31500867 | -0.4945009 | 0.62095246 | 0.89896588 |
| Slc25a42 | 29.5115353 | 0.27510275 | 0.70042702 | 0.39276432 | 0.69449356 | 0.92609254 |
| Snx9 | 1043.8167 | 0.49483093 | 0.37096225 | 1.33391182 | 0.18223276 | 0.61344341 |
| Ranbp3 | 5021.46973 | -0.2821304 | 0.18257323 | -1.5452999 | 0.12227374 | 0.53143108 |
| Ndufa11 | 3189.47851 | 0.45967989 | 0.38061542 | 1.20772796 | 0.22715192 | 0.66050617 |
| Nr2f6 | 134.740782 | 0.46964619 | 0.42489803 | 1.10531506 | 0.26902309 | 0.70592381 |
| Use1 | 2597.37447 | 0.38974 | 0.34963277 | 1.11471244 | 0.26497367 | 0.70112124 |
| Ocel1 | 261.871995 | 0.10931099 | 0.37201544 | 0.29383455 | 0.76888433 | 0.94401697 |
| Dyrk1b | 23.9720027 | -0.2647824 | 1.37902526 | -0.1920069 | 0.8477368 | 0.96765585 |
| Braf | 407.123192 | 0.07768881 | 0.34204798 | 0.2271284 | 0.8203239 | 0.96185395 |
| Ndufb2 | 2165.46635 | 0.64038769 | 0.478196 | 1.33917409 | 0.180514 | 0.61122544 |
| Hltf | 858.666874 | -0.4982435 | 0.32718098 | -1.5228377 | 0.12779931 | 0.54227156 |
| Prpf6 | 2933.73691 | -0.1879423 | 0.21687206 | -0.8666046 | 0.38615869 | 0.7886896 |
| Rgs19 | 5299.76359 | -0.399647 | 0.16715634 | -2.3908573 | 0.01680908 | 0.18128333 |
| Snrpd1 | 6223.37285 | 0.17352586 | 0.34063796 | 0.50941435 | 0.61046182 | 0.89508835 |
| Tchp | 148.727796 | 0.04934467 | 0.37519068 | 0.13151891 | 0.89536483 | 0.97952048 |
| Tiam1 | 45.2658642 | -1.7091954 | 1.01081745 | -1.6909042 | 0.09085511 | 0.46044203 |
| Tsc2 | 1065.00017 | -0.3333953 | 0.29333272 | -1.1365773 | 0.25571502 | 0.69039104 |
| Slc9a3r2 | 283.244885 | 1.44522521 | 0.40123909 | 3.60190533 | 0.00031589 | 0.00922273 |
| Puf60 | 10499.9049 | 0.0002358 | 0.1633054 | 0.00144394 | 0.9988479 | 0.99969186 |
| Golga2 | 917.819821 | -0.3136631 | 0.33731126 | -0.9298922 | 0.35242688 | 0.76358405 |
| Uck1 | 2880.15368 | 0.12789862 | 0.2163144 | 0.59126261 | 0.55434448 | 0.87165957 |
| Mien1 | 2008.05624 | 0.46943146 | 0.3750088 | 1.25178786 | 0.21064717 | 0.64401792 |
| Axl | 445.342034 | -0.1222234 | 0.34598696 | -0.3532601 | 0.72389347 | 0.93466537 |
| Tgfb1 | 52.9618364 | 1.2657051 | 1.09114166 | 1.15998239 | 0.24605598 | 0.68117517 |
| Ccdc97 | 889.379569 | -0.2220847 | 0.22106451 | -1.004615 | 0.31508229 | 0.74123109 |
| Zfp40 | 149.689938 | 1.01490155 | 0.56624696 | 1.79233026 | 0.07308008 | 0.41561534 |
| Akap8l | 762.229714 | -0.0248581 | 0.33863754 | -0.0734062 | 0.94148291 | 0.98948137 |

| | | | | | | |
|---|---|---|---|---|---|---|
| Pdcd2l | 1876.01377 | 0.19524317 | 0.34134931 | 0.57197469 | 0.56733912 | 0.87929304 |
| Gtf2f1 | 3946.73556 | -0.1478374 | 0.18002781 | -0.8211921 | 0.41153688 | 0.80231592 |
| Clpp | 2933.95032 | -0.0464636 | 0.26202593 | -0.1773243 | 0.85925368 | 0.96954641 |
| Alkbh7 | 802.801828 | 0.36389519 | 0.41476371 | 0.87735542 | 0.38029362 | 0.78292357 |
| Dennd1c | 567.416847 | -0.0368341 | 0.30031556 | -0.1226512 | 0.90238333 | 0.98043702 |
| Med6 | 1376.47994 | 0.48056834 | 0.29938717 | 1.60517347 | 0.10845563 | 0.50301538 |
| Lcp2 | 8918.45971 | -0.1993024 | 0.24696066 | -0.8070209 | 0.41965446 | 0.80563709 |
| Cse1l | 13615.7164 | -0.1871784 | 0.1450279 | -1.290637 | 0.19682958 | 0.62932233 |
| Naa20 | 8988.78638 | 0.48619328 | 0.36748696 | 1.32302188 | 0.1858281 | 0.61790851 |
| Prkra | 923.681446 | 0.93597973 | 0.36847417 | 2.5401502 | 0.01108049 | 0.13664072 |
| Fkbp7 | 76.2868021 | 1.4700368 | 0.81266116 | 1.80891725 | 0.07046386 | 0.41006309 |
| Plekha3 | 71.5785045 | -0.5965929 | 0.57616626 | -1.0354527 | 0.30045756 | 0.73158785 |
| Ykt6 | 1110.87718 | 0.50179784 | 0.32990543 | 1.52103542 | 0.12825095 | 0.54327949 |
| Baz1b | 2376.02811 | -0.4158877 | 0.31008815 | -1.341192 | 0.17985814 | 0.61065906 |
| Pex6 | 337.483263 | 0.15968757 | 0.26888243 | 0.59389364 | 0.55258325 | 0.87098968 |
| Mrpl2 | 3576.08132 | -0.0367536 | 0.21090803 | -0.1742635 | 0.86165835 | 0.97020895 |
| Mea1 | 2689.00514 | 0.45554427 | 0.42229771 | 1.07872779 | 0.28070909 | 0.7155865 |
| Kdelr1 | 700.476875 | -0.1143425 | 0.20990787 | -0.5447269 | 0.58594136 | 0.88513468 |
| Tmem143 | 64.8263204 | 0.61576976 | 0.49949342 | 1.23278853 | 0.21765467 | 0.65150232 |
| Ggct | 193.801551 | -0.7760122 | 0.3675404 | -2.1113657 | 0.03474089 | 0.27929255 |
| Jag2 | 10.374613 | -0.1269664 | 1.70213076 | -0.0745926 | 0.94053886 | 0.98935054 |
| Btbd6 | 285.032556 | 0.11532591 | 0.31022166 | 0.37175323 | 0.71007659 | 0.93077435 |
| Nudt14 | 499.263545 | 0.39185848 | 0.51465454 | 0.76140098 | 0.4464176 | 0.8219486 |
| Flii | 5036.60045 | -0.2846045 | 0.29787246 | -0.9554577 | 0.33934631 | 0.75683748 |
| Top3a | 963.882223 | 0.23147003 | 0.19509128 | 1.18647044 | 0.23543657 | 0.67164041 |
| Atg4d | 636.979847 | 0.16292482 | 0.27557152 | 0.59122519 | 0.55436955 | 0.87165957 |
| Qtrt1 | 1573.29317 | -0.044523 | 0.33635206 | -0.1323703 | 0.89469144 | 0.97941957 |
| Hdgfl2 | 1879.12033 | -0.0782344 | 0.17047788 | -0.4589124 | 0.64629705 | 0.90885755 |
| Chaf1a | 3824.73916 | -0.0004905 | 0.28611695 | -0.0017143 | 0.99863217 | 0.99967651 |
| Adprh | 4426.71131 | 0.10756981 | 0.16169121 | 0.66527928 | 0.50587188 | 0.85074333 |
| Tmem39a | 1224.68097 | -0.3056465 | 0.26090503 | -1.1714858 | 0.24140355 | 0.67708722 |
| Timmdc1 | 531.809571 | -0.0018734 | 0.36965678 | -0.0050679 | 0.99595638 | 0.99884675 |
| Pla1a | 14.5398603 | -0.2326477 | 1.18607814 | -0.1961487 | 0.84449375 | 0.96765585 |
| Mcm2 | 15796.0034 | -0.3460523 | 0.1908765 | -1.8129643 | 0.06983733 | 0.40804748 |
| Tpra1 | 1373.39634 | 0.18521668 | 0.34652532 | 0.5344968 | 0.59299786 | 0.88797037 |
| Nab1 | 439.851878 | -0.5095529 | 0.31709384 | -1.6069468 | 0.10806603 | 0.50301538 |
| Adgre5 | 2655.08937 | -0.2240237 | 0.26501438 | -0.8453267 | 0.39792846 | 0.79531492 |
| Il17ra | 5818.595 | -0.5807136 | 0.27095661 | -2.1431977 | 0.03209723 | 0.26624356 |
| Lamb1 | 27.7824992 | 2.79087397 | 1.41317485 | 1.97489644 | 0.0482799 | 0.3386514 |
| Arrdc2 | 181.996315 | 0.67687521 | 0.41147068 | 1.64501444 | 0.09996683 | 0.48325682 |
| Cd36 | 2730.97828 | 1.09042991 | 0.26792058 | 4.06997444 | 4.701829682 | 0.00191725 |
| Map2k7 | 439.268666 | -0.6369173 | 0.33171061 | -1.9200994 | 0.05484534 | 0.36084578 |
| Timm44 | 3261.72461 | -0.2061941 | 0.17774877 | -1.1600312 | 0.24603611 | 0.68117517 |
| Ap2a2 | 9590.48667 | 0.72385076 | 0.27208226 | 2.66041148 | 0.00780452 | 0.10907646 |
| Pnkp | 2598.99084 | -0.0103456 | 0.2096938 | -0.0493366 | 0.96065102 | 0.99325523 |
| Med25 | 1940.59354 | -0.1106876 | 0.3280412 | -0.3374197 | 0.73580054 | 0.9377627 |

| | | | | | | |
|---|---|---|---|---|---|---|
| Clptm1 | 8333.79753 | 0.01149482 | 0.20407624 | 0.05632609 | 0.95508204 | 0.99215259 |
| Relb | 345.929809 | 0.63599302 | 0.640129 | 0.99353884 | 0.32044743 | 0.74413737 |
| Tomm40 | 7462.58689 | -0.4127708 | 0.19612275 | -2.1046552 | 0.03532134 | 0.28154973 |
| Apoe | 74797.7084 | 0.67717278 | 0.19812817 | 3.41785215 | 0.00063117 | 0.01632378 |
| Apoc2 | 221.366914 | -0.4899885 | 0.51699395 | -0.9477644 | 0.34324938 | 0.75947281 |
| Hbp1 | 4253.99003 | 0.59251969 | 0.29643067 | 1.99884743 | 0.04562486 | 0.32966861 |
| Prkar2b | 303.206513 | -0.0869178 | 0.43426232 | -0.2001505 | 0.84136289 | 0.96765585 |
| Cdkn1b | 375.424099 | 0.75325304 | 0.31719824 | 2.37470749 | 0.01756286 | 0.18569273 |
| Klf4 | 152.058606 | -0.8383687 | 0.50600383 | -1.6568426 | 0.09755129 | 0.47887037 |
| Ap1m1 | 8387.67043 | -0.2101227 | 0.15819319 | -1.3282663 | 0.18409015 | 0.61619551 |
| Rab8a | 8019.82727 | 0.05656437 | 0.15531391 | 0.36419384 | 0.71571326 | 0.93235217 |
| Hmgn2 | 60859.5289 | -0.1930676 | 0.21755305 | -0.8874506 | 0.37483633 | 0.77815049 |
| Fam32a | 4573.85071 | 0.0202165 | 0.17167096 | 0.1177631 | 0.90625537 | 0.98159944 |
| Stard3nl | 4626.10542 | 0.70519766 | 0.35977403 | 1.96011276 | 0.04998261 | 0.34252555 |
| Stk11 | 420.831008 | -0.2129138 | 0.40594875 | -0.5244844 | 0.59994165 | 0.8902352 |
| Atp5d | 7141.94776 | -0.3193847 | 0.15066399 | -2.1198474 | 0.03401891 | 0.27569109 |
| Ppp5c | 5576.03186 | -0.2386676 | 0.18145923 | -1.3152683 | 0.18841974 | 0.61935434 |
| Cdk12 | 223.61691 | 0.0720769 | 0.41874676 | 0.17212528 | 0.86333904 | 0.9705375 |
| Lipe | 546.147804 | -0.4309786 | 0.2720145 | -1.5843956 | 0.11310371 | 0.51263346 |
| Pafah1b2 | 6261.59128 | 0.12118314 | 0.23450807 | 0.51675468 | 0.60532742 | 0.89258167 |
| Tbc1d8 | 823.62426 | -0.3208739 | 0.25193273 | -1.273649 | 0.20278782 | 0.63804101 |
| Cnot11 | 127.789737 | -0.6930493 | 0.47343605 | -1.463871 | 0.1432292 | 0.56846902 |
| Slc2a3 | 3671.3088 | -0.1347494 | 0.28022309 | -0.4808646 | 0.63061276 | 0.9036296 |
| Foxj2 | 159.292033 | -1.2586044 | 0.45660139 | -2.7564621 | 0.00584304 | 0.09002253 |
| Sri | 5426.65962 | 0.36740097 | 0.26329924 | 1.3953742 | 0.16290303 | 0.59424967 |
| Dgcr2 | 3406.54369 | -0.1754953 | 0.26610239 | -0.6595029 | 0.50957285 | 0.85335831 |
| Irf3 | 2662.41547 | -0.049453 | 0.24439023 | -0.2023527 | 0.83964101 | 0.96758168 |
| Bcl2l12 | 405.098325 | -0.0266305 | 0.31802445 | -0.0837373 | 0.93326527 | 0.98751425 |
| Zfp959 | 440.288713 | 0.35707816 | 0.45603071 | 0.7830134 | 0.43361925 | 0.81483708 |
| Mpnd | 185.571824 | -0.531268 | 0.36939963 | -1.438193 | 0.15037929 | 0.57837915 |
| Sh3gl1 | 6358.75252 | 0.8897416 | 0.16607284 | 5.35753831 | 8.436348336 | 8.507181661 |
| Ebi3 | 1404.25251 | -0.3564805 | 0.22241587 | -1.6027658 | 0.10898638 | 0.50470898 |
| Yju2 | 696.10041 | -0.0490536 | 0.210619 | -0.232902 | 0.81583753 | 0.96063519 |
| Ranbp2 | 1074.08926 | 0.09391282 | 0.35037669 | 0.26803387 | 0.78867325 | 0.95225386 |
| Grk5 | 44.5011287 | 1.17426687 | 0.99597265 | 1.17901518 | 0.23839213 | 0.67446139 |
| Dvl3 | 958.198103 | 0.24394822 | 0.29517305 | 0.8264583 | 0.40854416 | 0.80100711 |
| Abcf3 | 2494.86698 | 0.37143928 | 0.25144153 | 1.47723917 | 0.13961154 | 0.56329174 |
| Eif2b5 | 3672.66991 | -0.0164177 | 0.26271388 | -0.0624927 | 0.95017048 | 0.99069447 |
| Cyth2 | 1598.75774 | 0.2662263 | 0.21764452 | 1.22321618 | 0.22124806 | 0.65469944 |
| Plag1 | 69.6516022 | -0.117526 | 0.65478675 | -0.1794875 | 0.85755497 | 0.9694296 |
| Hck | 5676.75406 | -0.3062403 | 0.15042569 | -2.0358243 | 0.041768 | 0.31168889 |
| Mrpl4 | 4741.09101 | -0.2772519 | 0.15991761 | -1.7337174 | 0.08296824 | 0.4429983 |
| Keap1 | 1685.98537 | -0.2114387 | 0.1935892 | -1.0922031 | 0.27474382 | 0.71147204 |
| Ap1m2 | 14.2877353 | -0.3147193 | 2.12466979 | -0.1481262 | 0.88224315 | 0.97598447 |
| Glg1 | 1911.00243 | -0.0232025 | 0.49096281 | -0.0472593 | 0.96230659 | 0.99400521 |
| Btbd2 | 114.570963 | -0.3991625 | 0.52160513 | -0.765258 | 0.44411795 | 0.82068545 |

| | | | | | | |
|---|---|---|---|---|---|---|
| Csnk1g2 | 277.503805 | -0.6320748 | 0.53519469 | -1.1810185 | 0.23759537 | 0.67389384 |
| Abhd17a | 195.400648 | -0.4902953 | 0.39031274 | -1.2561602 | 0.2090579 | 0.64228937 |
| Mob3a | 5153.68067 | -0.2798639 | 0.21664863 | -1.291787 | 0.19643093 | 0.62887714 |
| Fkbp11 | 366.320666 | 0.81566773 | 0.48480005 | 1.68248275 | 0.09247527 | 0.46597694 |
| Ddx23 | 211.484226 | 0.56687363 | 0.49011106 | 1.15662281 | 0.2474265 | 0.68235696 |
| Pld3 | 2336.8818 | 0.68336219 | 0.28010491 | 2.4396652 | 0.01470088 | 0.16670141 |
| Rabac1 | 1032.5933 | 0.17703028 | 0.26498897 | 0.6680666 | 0.50409108 | 0.85012672 |
| Etv3 | 346.881886 | 0.68540408 | 0.39508394 | 1.73483155 | 0.08277065 | 0.44254079 |
| Prkcsh | 2844.97626 | -0.2664636 | 0.1987418 | -1.3407528 | 0.18000071 | 0.61080409 |
| St8sia6 | 29.5819352 | -0.3369695 | 1.28228296 | -0.2627887 | 0.79271341 | 0.95297175 |
| Fcgrt | 7236.37763 | 1.85504936 | 0.24132598 | 7.68690281 | 1.507395588 | 5.762490698 |
| Nosip | 2386.83108 | -0.0068806 | 0.21461403 | -0.0320605 | 0.97442384 | 0.99446549 |
| Pih1d1 | 2187.42546 | 0.3497546 | 0.33888193 | 1.03208394 | 0.30203279 | 0.73361866 |
| Rps11 | 39538.9545 | 0.36293295 | 0.36930521 | 0.98274527 | 0.32573282 | 0.74723652 |
| Supt5 | 3235.32439 | -0.255243 | 0.27769494 | -0.9191491 | 0.35801761 | 0.76674184 |
| Paf1 | 3022.39311 | -0.2649178 | 0.19788198 | -1.3387666 | 0.18064668 | 0.61122544 |
| Timm50 | 6648.15292 | -0.127135 | 0.20033193 | -0.6346218 | 0.52567511 | 0.86056156 |
| Med29 | 1187.81514 | -0.0304517 | 0.21255331 | -0.1432661 | 0.88608006 | 0.97793139 |
| Ncstn | 5620.36792 | 0.04198945 | 0.15728597 | 0.26696249 | 0.78949804 | 0.95255415 |
| Pex19 | 3578.57747 | 0.18343224 | 0.19874071 | 0.92297266 | 0.35602145 | 0.76514589 |
| Cyp4f18 | 550.224734 | -2.9004769 | 0.28999705 | -10.001746 | 1.497328977 | 1.526402115 |
| Impdh1 | 508.069941 | -0.056097 | 0.33314768 | -0.1683848 | 0.86628057 | 0.97060021 |
| Dusp3 | 5327.94184 | 0.56593815 | 0.27135002 | 2.08563886 | 0.03701135 | 0.28924449 |
| Prodh | 1209.97215 | -0.5902628 | 0.27531815 | -2.1439298 | 0.03203851 | 0.26624356 |
| Ess2 | 1321.5337 | 0.07124601 | 0.21425161 | 0.33253432 | 0.73948583 | 0.93800224 |
| Slc25a1 | 2158.37588 | 0.00388038 | 0.20086144 | 0.01931869 | 0.98458688 | 0.99564261 |
| Dgcr6 | 965.116752 | 0.37908939 | 0.3856091 | 0.98309245 | 0.32556194 | 0.74706419 |
| Ier3 | 3218.98943 | 0.99414861 | 0.27000581 | 3.68195266 | 0.00023145 | 0.00727862 |
| Fosb | 775.345486 | -0.0072888 | 0.54059857 | -0.0134828 | 0.98924261 | 0.99650865 |
| Klc4 | 1079.31485 | 0.29370768 | 0.36651166 | 0.80135973 | 0.42292342 | 0.80844471 |
| Ercc1 | 850.819352 | -0.0383262 | 0.28278 | -0.1355335 | 0.89219008 | 0.97937556 |
| As3mt | 1781.53866 | 0.18641512 | 0.17883364 | 1.04239402 | 0.29722904 | 0.72954201 |
| Homer3 | 298.940366 | 0.46770994 | 0.26780755 | 1.74644042 | 0.08073445 | 0.43854735 |
| Crtc1 | 229.288391 | 0.48404751 | 0.38609839 | 1.25368952 | 0.20995488 | 0.64350283 |
| Rnf215 | 112.898532 | 0.67250187 | 0.52233794 | 1.28748425 | 0.19792556 | 0.63076766 |
| Aven | 54.5716756 | -0.9417029 | 0.72099845 | -1.3061094 | 0.19151534 | 0.62259956 |
| Crot | 2259.33214 | 0.84559534 | 0.32070267 | 2.6366957 | 0.00837179 | 0.11493265 |
| Rps6ka1 | 11386.346 | -0.6720969 | 0.20973765 | -3.2044644 | 0.00135314 | 0.02909132 |
| Snrnp200 | 1472.88097 | -0.2099149 | 0.26294791 | -0.7983138 | 0.42468843 | 0.80982401 |
| Ciao1 | 3209.53238 | 0.14528695 | 0.25475664 | 0.57029703 | 0.56847626 | 0.87936678 |
| Taf6l | 1671.27308 | 0.25623974 | 0.17261405 | 1.48446628 | 0.13768526 | 0.56012764 |
| Insig2 | 856.49685 | 0.52524754 | 0.2389225 | 2.19840134 | 0.02792052 | 0.24804045 |
| Kpna6 | 858.998718 | 0.11867388 | 0.36006044 | 0.32959433 | 0.74170651 | 0.93800224 |
| Man1a | 713.824298 | 1.1405367 | 0.42302292 | 2.6961582 | 0.00701443 | 0.10251801 |
| Itpkc | 58.2219614 | -0.0077076 | 0.87935569 | -0.008765 | 0.99300662 | 0.99806507 |
| Coq8b | 914.295076 | 0.2283723 | 0.21551694 | 1.05964893 | 0.28930434 | 0.72395811 |

| | | | | | | |
|---|---|---|---|---|---|---|
| Brd8 | 2777.93112 | -0.1131479 | 0.30142081 | -0.3753818 | 0.70737652 | 0.92950127 |
| Kif20a | 8481.30006 | -0.2750171 | 0.1468317 | -1.8730089 | 0.06106715 | 0.38152934 |
| Farsa | 5347.51008 | -0.3399631 | 0.18283466 | -1.8594017 | 0.06297022 | 0.38709283 |
| Gcdh | 1355.31354 | 0.49284299 | 0.24527244 | 2.00936965 | 0.04449795 | 0.32382117 |
| Mast2 | 391.482262 | -0.2713713 | 0.49841165 | -0.5444722 | 0.58611655 | 0.88518071 |
| Dnase2a | 2352.03676 | 0.5885356 | 0.3198494 | 1.84003969 | 0.06576241 | 0.39558985 |
| Rad23a | 3307.37136 | -0.178832 | 0.1762819 | -1.0144662 | 0.31036034 | 0.7399039 |
| Calr | 25331.5018 | 0.011927 | 0.17475239 | 0.06825086 | 0.94558594 | 0.99019648 |
| Syce2 | 1890.85614 | 0.17831356 | 0.34287804 | 0.52004952 | 0.60302906 | 0.89168472 |
| Nfat5 | 83.9152011 | 0.34287106 | 0.88504307 | 0.38740607 | 0.69845561 | 0.92669304 |
| Nob1 | 3098.84812 | -0.3206653 | 0.1967076 | -1.6301625 | 0.10306716 | 0.49059168 |
| Nqo1 | 34.4359413 | 0.88300065 | 1.22012599 | 0.72369629 | 0.46925221 | 0.83381643 |
| Gys1 | 985.360692 | -0.4674176 | 0.37111402 | -1.2594987 | 0.20785025 | 0.6418276 |
| Ruvbl2 | 8628.59894 | -0.4145682 | 0.23837884 | -1.7391149 | 0.08201456 | 0.44108302 |
| Bax | 4337.04903 | 0.04205957 | 0.1958986 | 0.21470073 | 0.83000066 | 0.96464475 |
| Tfam | 1675.08654 | -0.1471095 | 0.23759415 | -0.6191628 | 0.53580911 | 0.86539693 |
| Zfp81 | 301.67239 | -0.2333195 | 0.29508075 | -0.790697 | 0.4291208 | 0.81140481 |
| Mmd | 1281.84124 | 0.76568374 | 0.19882053 | 3.85113022 | 0.00011757 | 0.00413299 |
| Fam162a | 3343.09592 | 0.00606438 | 0.37037701 | 0.01637354 | 0.98693639 | 0.99616298 |
| Rpl8 | 20777.8936 | -0.0934837 | 0.27933832 | -0.3346611 | 0.73788075 | 0.93800224 |
| Ssbp2 | 1690.98705 | -0.9148279 | 0.22576959 | -4.052042 | 5.077253853 | 0.00204309 |
| Fancl | 1148.67844 | -0.0095085 | 0.28684722 | -0.0331482 | 0.97355639 | 0.99443087 |
| Gstm5 | 427.296797 | 0.56057069 | 0.39440686 | 1.42130056 | 0.1552294 | 0.58428346 |
| Gstm7 | 46.2650982 | 0.94006138 | 1.08997939 | 0.86245794 | 0.38843557 | 0.78942657 |
| Stat3 | 4696.98899 | -0.0708741 | 0.39547393 | -0.179213 | 0.85777045 | 0.9694296 |
| Stat5a | 591.349718 | -0.5361523 | 0.27401016 | -1.9566878 | 0.05038418 | 0.3433703 |
| Cavin1 | 80.3943783 | 1.17437327 | 0.87386638 | 1.34388196 | 0.17898656 | 0.60992118 |
| Map3k11 | 2072.25831 | 0.23649179 | 0.3092797 | 0.76465343 | 0.44447794 | 0.82091127 |
| Akt2 | 2188.42978 | 0.16220779 | 0.21201788 | 0.76506657 | 0.44423191 | 0.82068545 |
| Dnaja3 | 3585.9279 | -0.3494543 | 0.17579839 | -1.9878129 | 0.04683239 | 0.33444287 |
| Hmox2 | 7140.80998 | 0.09066754 | 0.18321716 | 0.49486379 | 0.62069627 | 0.89896588 |
| Cdip1 | 1541.26528 | 0.23245408 | 0.19452713 | 1.19496998 | 0.23209876 | 0.66733333 |
| Map3k20 | 809.676393 | -0.4535167 | 0.22940069 | -1.9769632 | 0.04804579 | 0.33778401 |
| Cwc15 | 2999.9253 | -0.1204264 | 0.16484861 | -0.7305275 | 0.46506783 | 0.83339036 |
| Dnmt1 | 5584.76209 | -0.2945341 | 0.37067679 | -0.7945846 | 0.4268552 | 0.81021713 |
| Ppan | 2826.70132 | -0.5348819 | 0.20305798 | -2.6341337 | 0.00843523 | 0.11542295 |
| Angptl2 | 80.1006903 | -1.388903 | 0.61386896 | -2.2625399 | 0.02366406 | 0.22185905 |
| Trmt10a | 643.117109 | 0.48566847 | 0.19907634 | 2.43960924 | 0.01470316 | 0.16670141 |
| Etv1 | 587.449596 | 1.31387099 | 0.40955013 | 3.20808342 | 0.00133623 | 0.02882905 |
| Kifc2 | 14.7670572 | 0.90919878 | 1.97624534 | 0.46006372 | 0.64547048 | 0.90847224 |
| Psap | 13998.4907 | -0.4849349 | 0.1669032 | -2.9054858 | 0.00366683 | 0.06317374 |
| Ikbkg | 1350.71422 | 0.17616213 | 0.25866291 | 0.68104904 | 0.49584046 | 0.84750822 |
| Wars2 | 462.432396 | 0.08616295 | 0.27660689 | 0.31149964 | 0.75542081 | 0.94087513 |
| Atn1 | 38.5902065 | -0.1699383 | 1.39717462 | -0.1216299 | 0.9031921 | 0.98043702 |
| Phb2 | 16650.3401 | -0.1160945 | 0.21966974 | -0.5284959 | 0.59715522 | 0.88891958 |
| Ptpn6 | 19187.2839 | -0.1657704 | 0.14578274 | -1.137106 | 0.255494 | 0.69025133 |

| | | | | | | |
|---|---|---|---|---|---|---|
| Emg1 | 7520.82087 | -0.1062313 | 0.29593679 | -0.3589661 | 0.71962045 | 0.93314808 |
| Lpcat3 | 3647.33133 | -0.2234248 | 0.200327 | -1.1153006 | 0.26472164 | 0.70112124 |
| Atp6v1f | 7252.05037 | 0.58473408 | 0.38876086 | 1.50409708 | 0.13255637 | 0.55219606 |
| Clcn5 | 1029.29342 | 0.8854239 | 0.34885973 | 2.53805133 | 0.01114716 | 0.13718637 |
| Clcn3 | 1141.403 | -0.1500154 | 0.20303181 | -0.7388765 | 0.45998201 | 0.82987212 |
| Hif3a | 162.618084 | 0.72804993 | 0.66998534 | 1.08666547 | 0.27718468 | 0.71247429 |
| Utp20 | 592.427544 | -0.3903338 | 0.28869138 | -1.3520797 | 0.17634983 | 0.60764153 |
| Spic | 12.8631827 | 2.08959488 | 1.62670644 | 1.2845556 | 0.19894762 | 0.63148061 |
| Cul3 | 739.016494 | -0.8122161 | 0.18987378 | -4.2776632 | 1.888655193 | 0.00086532 |
| Large1 | 30.1353381 | 0.37140619 | 1.43023726 | 0.25968152 | 0.79510945 | 0.95367917 |
| Ddx56 | 6363.23856 | -0.2289231 | 0.15644757 | -1.4632576 | 0.1433969 | 0.56846902 |
| Tmed4 | 2682.60474 | 0.16295647 | 0.21907444 | 0.74384066 | 0.45697288 | 0.82915295 |
| Bid | 6509.33785 | -0.2792337 | 0.19756641 | -1.4133665 | 0.157548 | 0.58620126 |
| Ralb | 13384.2426 | 0.31177411 | 0.15071815 | 2.06859029 | 0.03858455 | 0.29684673 |
| Ppp1cc | 1395.07094 | 0.02583625 | 0.23060571 | 0.11203648 | 0.91079448 | 0.98288717 |
| Dnajb11 | 7131.96085 | -0.2479635 | 0.19740477 | -1.2561171 | 0.20907351 | 0.64228937 |
| Tbccd1 | 1424.05651 | -0.1100534 | 0.18259909 | -0.6027053 | 0.54670475 | 0.86994632 |
| Zfp324 | 315.166714 | -0.1338851 | 0.27138566 | -0.4933388 | 0.62177325 | 0.89912494 |
| Gab2 | 649.01011 | 0.44047127 | 0.45123398 | 0.97614829 | 0.32899098 | 0.7486136 |
| Coro1c | 12456.3997 | -0.1284308 | 0.23261093 | -0.5521273 | 0.58086117 | 0.88314588 |
| Tax1bp1 | 4138.10544 | -0.0248099 | 0.1766406 | -0.1404542 | 0.88830116 | 0.97832097 |
| Ctse | 22.6697974 | -4.4795629 | 1.25087753 | -3.5811363 | 0.0003421 | 0.00987016 |
| Ndrg2 | 48.064194 | -2.953766 | 1.23493627 | -2.3918368 | 0.01676429 | 0.18100406 |
| Mettl17 | 623.697974 | -0.5502575 | 0.2915406 | -1.8874129 | 0.05910481 | 0.37462651 |
| Arhgef40 | 131.330061 | 0.30566897 | 0.63295986 | 0.48291999 | 0.62915255 | 0.90280623 |
| Pnpla6 | 379.789345 | 0.20110426 | 0.34234411 | 0.58743308 | 0.55691288 | 0.87276537 |
| Mcoln1 | 1139.78454 | 0.30721551 | 0.22397895 | 1.3716267 | 0.17017968 | 0.60430173 |
| Arhgef18 | 139.184346 | -0.2496625 | 0.5737168 | -0.4351669 | 0.66344131 | 0.91395288 |
| Pkn2 | 526.013621 | -0.1047533 | 0.42564881 | -0.2461026 | 0.80560282 | 0.95737871 |
| Cd33 | 3201.50074 | -0.6739 | 0.16101616 | -4.1852941 | 2.847970644 | 0.00122996 |
| Etfb | 9532.15136 | 0.30183292 | 0.32161845 | 0.93848137 | 0.34799709 | 0.76101142 |
| Nkg7 | 38.5030181 | -8.6673541 | 1.7012406 | -5.0947257 | 3.492466030 | 2.893961448 |
| Stxbp2 | 8048.10333 | -0.3187937 | 0.17476176 | -1.8241619 | 0.0681276 | 0.40222245 |
| Sgce | 274.542022 | 1.83832636 | 0.4320615 | 4.25477941 | 2.092552088 | 0.00094808 |
| Chn2 | 1111.93665 | -1.4736269 | 0.26159488 | -5.6332405 | 1.768545337 | 2.142041100 |
| Wwox | 214.088273 | 0.11980676 | 0.32628381 | 0.36718572 | 0.71348049 | 0.93235217 |
| Slbp | 10630.8172 | -0.2533272 | 0.18084439 | -1.4008019 | 0.16127332 | 0.59141542 |
| Arid3b | 1255.96453 | -0.0169099 | 0.19946903 | -0.0847745 | 0.93244063 | 0.98732332 |
| Cnn2 | 7868.41536 | -0.917665 | 0.19529964 | -4.6987541 | 2.617534119 | 0.00016591 |
| Polr2e | 6343.7069 | 0.14589644 | 0.34719397 | 0.42021595 | 0.67432771 | 0.91822108 |
| Myo9b | 822.885189 | -0.3965233 | 0.49228614 | -0.8054732 | 0.42054666 | 0.80673472 |
| Hdac9 | 2284.13781 | 0.43055631 | 0.23731051 | 1.81431625 | 0.06962905 | 0.40760152 |
| Ly9 | 227.642421 | -0.6434717 | 0.50376967 | -1.2773134 | 0.20149163 | 0.63561051 |
| Cd244a | 23.3646753 | -1.9536084 | 1.0375909 | -1.8828311 | 0.05972327 | 0.37717848 |
| Adgre1 | 13584.9523 | 0.75855289 | 0.22749064 | 3.33443562 | 0.00085473 | 0.02066378 |
| Mtfp1 | 441.550751 | 0.26055886 | 0.38998811 | 0.66812001 | 0.50405699 | 0.85012672 |

| | | | | | | |
|---|---|---|---|---|---|---|
| Rab23 | 219.571146 | -0.1175686 | 0.53786606 | -0.2185834 | 0.82697459 | 0.96379955 |
| Rab11a | 8249.88024 | 0.9123511 | 0.282238 | 3.23255944 | 0.00122687 | 0.0271397 |
| Eif2b2 | 2955.06685 | -0.2288401 | 0.21015862 | -1.0888921 | 0.27620147 | 0.7118544 |
| Dlst | 8006.05077 | -0.0893025 | 0.17028036 | -0.524444 | 0.59996979 | 0.8902352 |
| Pgf | 14.3293265 | -0.0773732 | 1.60124584 | -0.0483206 | 0.96146073 | 0.9935419 |
| Ulk2 | 289.744052 | 1.24830752 | 0.41985374 | 2.97319616 | 0.00294716 | 0.05333227 |
| Ccl24 | 977.39852 | 1.44172156 | 0.40632994 | 3.54815487 | 0.00038794 | 0.01095998 |
| Dgkq | 249.8105 | 0.21767653 | 0.42094453 | 0.51711452 | 0.60507622 | 0.89258167 |
| Grap | 1905.62875 | 0.94426801 | 0.20575825 | 4.58921088 | 4.449247786 | 0.00026167 |
| Chmp2b | 2391.60584 | 0.10183074 | 0.18926555 | 0.53803103 | 0.59055563 | 0.88748446 |
| Plod3 | 1800.03288 | -0.6749189 | 0.21666107 | -3.1150909 | 0.00183888 | 0.03768014 |
| Ap1s1 | 5654.44053 | 0.13069546 | 0.16700595 | 0.78257967 | 0.43387398 | 0.81492099 |
| Mapk13 | 67.6847603 | -1.0019477 | 0.54168339 | -1.8496925 | 0.06435788 | 0.39202997 |
| Srpk1 | 4246.61263 | -0.4168279 | 0.20843517 | -1.9997963 | 0.04552226 | 0.3293163 |
| Lbr | 15121.351 | -0.5398956 | 0.17639651 | -3.0606932 | 0.00220825 | 0.04301521 |
| Crabp2 | 48.6060244 | 0.66972373 | 1.02602234 | 0.65273796 | 0.51392522 | 0.85500439 |
| Nes | 785.601687 | -0.0769687 | 0.33270674 | -0.231341 | 0.81704988 | 0.96079676 |
| Prcc | 96.1754711 | -0.6257983 | 0.56185255 | -1.1138124 | 0.2653597 | 0.70125149 |
| Rrnad1 | 313.970511 | 0.09460998 | 0.3509561 | 0.2695778 | 0.78748508 | 0.95225386 |
| Hdgf | 2777.97864 | -0.4956482 | 0.22590705 | -2.194036 | 0.02823282 | 0.24951635 |
| Slc25a18 | 57.2588502 | -0.5933071 | 0.92392599 | -0.6421587 | 0.52077017 | 0.8576517 |
| Thop1 | 3017.04556 | -0.6260904 | 0.1853744 | -3.3774371 | 0.00073165 | 0.01830314 |
| Apba3 | 1112.99581 | 0.05014169 | 0.20126279 | 0.24913544 | 0.80325602 | 0.95672466 |
| Matk | 210.950517 | -0.1279044 | 0.42520528 | -0.3008063 | 0.76356218 | 0.94300309 |
| Pias4 | 1809.29415 | -0.072779 | 0.25232794 | -0.2884302 | 0.77301746 | 0.94625874 |
| Map2k1 | 6793.28733 | -0.2461865 | 0.1631197 | -1.509238 | 0.13123797 | 0.54970237 |
| Sgta | 5880.69852 | -0.0046542 | 0.20987058 | -0.0221763 | 0.9823073 | 0.9952431 |
| Tmem242 | 1379.58917 | 0.52430287 | 0.39053641 | 1.34251979 | 0.17942752 | 0.60992118 |
| Dtx2 | 858.014398 | -0.1148824 | 0.31201281 | -0.3681977 | 0.71272581 | 0.93235217 |
| Hspb1 | 983.406927 | 0.01942768 | 0.67222766 | 0.02890044 | 0.97694399 | 0.99483019 |
| Rasa4 | 197.400427 | 0.42567791 | 0.5367537 | 0.79306005 | 0.42774284 | 0.81062404 |
| Hnrnpa2b1 | 37388.5962 | -0.3521131 | 0.17835544 | -1.9742215 | 0.04835656 | 0.33880059 |
| Fxyd4 | 11.2881852 | 1.12003244 | 1.21631533 | 0.92084052 | 0.3571337 | 0.76632459 |
| Ccdc130 | 527.488238 | 0.25618559 | 0.30590744 | 0.83746114 | 0.40233341 | 0.79794822 |
| Mri1 | 1597.656 | -0.3043025 | 0.20265157 | -1.5016044 | 0.1331993 | 0.55309812 |
| Prkacb | 1414.29856 | 0.29637331 | 0.33808477 | 0.87662427 | 0.38069075 | 0.78323054 |
| Sgsh | 688.379874 | -0.0538759 | 0.39918982 | -0.1349631 | 0.89264106 | 0.97941957 |
| Cstb | 8380.26452 | 1.05296132 | 0.31349673 | 3.35876331 | 0.00078292 | 0.01942634 |
| Sh2b2 | 704.500823 | -1.0325948 | 0.23951818 | -4.3111333 | 1.624199691 | 0.00077917 |
| Pex5 | 1624.14232 | 0.50354237 | 0.29094289 | 1.73072578 | 0.08350068 | 0.44386868 |
| Jkamp | 1661.41715 | 0.61342887 | 0.32563429 | 1.88379691 | 0.05959245 | 0.37674133 |
| Cd44 | 7797.72849 | -0.6064454 | 0.30907148 | -1.9621527 | 0.04974471 | 0.34225366 |
| Eif2ak4 | 484.945277 | 0.19674663 | 0.24293768 | 0.80986459 | 0.418018 | 0.8051099 |
| Wdr1 | 36851.7699 | -0.1604691 | 0.16210708 | -0.9898959 | 0.322225 | 0.7445747 |
| Slc2a9 | 91.9106027 | -0.398104 | 0.6078082 | -0.6549829 | 0.51247876 | 0.85434078 |
| Ndrg1 | 172.759839 | -0.8703305 | 0.6310423 | -1.3791952 | 0.16783457 | 0.60218592 |

| | | | | | | |
|---|---|---|---|---|---|---|
| Man2b1 | 9789.9855 | -0.0120731 | 0.21370427 | -0.0564942 | 0.95494809 | 0.99215259 |
| Klf5 | 15.5523162 | 0.44724153 | 1.56354292 | 0.28604365 | 0.77484469 | 0.94710982 |
| Wdr83 | 1154.96818 | 0.00588922 | 0.17911206 | 0.03288011 | 0.9737702 | 0.99443087 |
| Prdx2 | 9919.83085 | 0.76840263 | 0.42354483 | 1.81421797 | 0.06964417 | 0.40760152 |
| Polr2a | 825.584258 | -0.2030657 | 0.2432272 | -0.8348808 | 0.40378481 | 0.79875478 |
| Senp3 | 2741.24496 | 0.0916002 | 0.25247271 | 0.3628123 | 0.7167451 | 0.93236313 |
| Plekha8 | 592.533804 | 1.99662002 | 0.52555386 | 3.79907787 | 0.00014524 | 0.00494893 |
| Spc25 | 2771.06544 | 0.13613816 | 0.3041259 | 0.44763751 | 0.65441483 | 0.9108895 |
| Dnah2 | 14.3103597 | 0.2746737 | 1.51771338 | 0.18097864 | 0.85638434 | 0.96924897 |
| Ufd1 | 1904.08968 | 0.17701683 | 0.23672002 | 0.74778985 | 0.45458692 | 0.82758903 |
| Zfp287 | 23.2205563 | 0.3757776 | 1.30571046 | 0.2877955 | 0.77350328 | 0.94651052 |
| Letm1 | 3663.34381 | -0.2637697 | 0.18111949 | -1.4563298 | 0.14530148 | 0.57153471 |
| Ubqln1 | 5827.39445 | 0.01107225 | 0.36717964 | 0.03015485 | 0.97594356 | 0.99472734 |
| Txn2 | 8798.80485 | -0.1865763 | 0.16624323 | -1.1223094 | 0.2617309 | 0.6974044 |
| Slc1a3 | 114.408909 | -0.6183595 | 0.80143701 | -0.7715634 | 0.44037306 | 0.81963526 |
| Crbn | 1786.23071 | 0.41475784 | 0.30263249 | 1.37050008 | 0.17053085 | 0.60430173 |
| Msh6 | 3053.39494 | -0.1459107 | 0.20804103 | -0.7013554 | 0.48308124 | 0.84106286 |
| Fbxo11 | 376.968939 | -0.3846518 | 0.2783573 | -1.3818634 | 0.16701365 | 0.60073616 |
| Mlxipl | 24.2464279 | 1.57364836 | 1.26826892 | 1.24078445 | 0.21468539 | 0.64854028 |
| Tbl2 | 1209.10218 | -0.3214433 | 0.20583684 | -1.5616413 | 0.11837252 | 0.52515149 |
| Bud23 | 3683.87256 | -0.1510191 | 0.17772436 | -0.8497378 | 0.39547089 | 0.79334132 |
| Nid1 | 28.9977906 | 1.43754171 | 1.90684612 | 0.75388449 | 0.45091858 | 0.82549763 |
| Mcm5 | 22191.2205 | -0.1869458 | 0.13370534 | -1.3981921 | 0.16205538 | 0.59315429 |
| Hmox1 | 16878.5477 | 1.09558201 | 0.25353235 | 4.32127114 | 1.551328990 | 0.00075307 |
| Mprip | 552.764088 | -0.1366321 | 0.20657119 | -0.6614284 | 0.50833761 | 0.85238774 |
| Cic | 191.003999 | -0.1976854 | 0.43837929 | -0.450946 | 0.65202845 | 0.91037383 |
| Pafah1b3 | 616.600288 | 0.74553481 | 0.39012889 | 1.91099616 | 0.05600507 | 0.3642729 |
| Prkaca | 506.753545 | 0.15437425 | 0.31461213 | 0.49068118 | 0.62365196 | 0.90017327 |
| Asf1b | 6584.43617 | -0.3836281 | 0.18077508 | -2.1221293 | 0.03382688 | 0.27458805 |
| Ddx39 | 21428.0609 | 0.0082775 | 0.29520843 | 0.02803951 | 0.97763064 | 0.99483019 |
| Dnajb1 | 27264.6092 | 0.10741502 | 0.25472115 | 0.4216965 | 0.67324656 | 0.91786342 |
| Usp40 | 314.921193 | -0.0207196 | 0.49752719 | -0.0416452 | 0.96678154 | 0.99443087 |
| Kbtbd4 | 1462.95906 | -0.1659149 | 0.20493649 | -0.8095917 | 0.41817485 | 0.80521532 |
| Celf1 | 4197.41283 | 0.05953595 | 0.36718827 | 0.16214011 | 0.87119552 | 0.97268995 |
| Ndufs3 | 3933.29905 | 0.01732865 | 0.19196598 | 0.09026939 | 0.92807315 | 0.98666423 |
| Por | 4663.32119 | -0.1285057 | 0.26284925 | -0.4888949 | 0.62491611 | 0.90095448 |
| Igf1r | 188.315249 | -0.5235894 | 0.54190095 | -0.9662087 | 0.33393975 | 0.75320272 |
| Insr | 249.744375 | -0.2890465 | 0.5038168 | -0.5737136 | 0.56616163 | 0.87858116 |
| Atp4a | 10.4985029 | -3.1679896 | 2.47188918 | -1.2816067 | 0.19998067 | 0.63247751 |
| Trim28 | 3611.66837 | -0.0944125 | 0.20224868 | -0.4668139 | 0.64063302 | 0.90685007 |
| Ube2m | 239.706902 | -0.1329913 | 0.28821872 | -0.461425 | 0.64449372 | 0.90830549 |
| Adcy9 | 22.2861671 | 1.41029611 | 1.39514935 | 1.01085673 | 0.31208501 | 0.74050472 |
| Mef2c | 11115.768 | -0.2712615 | 0.23624162 | -1.1482374 | 0.25087056 | 0.6842534 |
| Ctr9 | 3876.42119 | -0.0854338 | 0.20232972 | -0.4222501 | 0.67284244 | 0.91780571 |
| Eif4g2 | 27068.968 | -0.0909156 | 0.17323139 | -0.5248217 | 0.59970712 | 0.8902352 |
| Mrvi1 | 429.152819 | 2.00655123 | 0.34885374 | 5.75184088 | 8.827686885 | 1.316607998 |

| | | | | | | |
|---|---|---|---|---|---|---|
| Pcyt1a | 1266.64085 | 0.23403673 | 0.27981001 | 0.83641301 | 0.4029226 | 0.79795242 |
| Zfp592 | 933.093189 | 0.19138523 | 0.30341013 | 0.6307806 | 0.52818399 | 0.86198732 |
| Psmd4 | 8283.40851 | 0.02596913 | 0.21372511 | 0.12150715 | 0.90328935 | 0.98043702 |
| Snx6 | 14763.304 | 0.93292064 | 0.19989932 | 4.6669525 | 3.056999615 | 0.00018983 |
| Mthfd2 | 4155.408 | -0.9325381 | 0.18184245 | -5.1282752 | 2.924087439 | 2.484052892 |
| Kit | 76.6932076 | 0.47835078 | 1.13583756 | 0.42114365 | 0.67365019 | 0.91822108 |
| Tomm40l | 1573.13703 | 0.30906742 | 0.34914458 | 0.88521328 | 0.37604161 | 0.77836994 |
| Apoa2 | 15.5970877 | 0.94477995 | 1.0707641 | 0.88234182 | 0.37759199 | 0.78033065 |
| Pan2 | 1008.29519 | 0.00745034 | 0.2023794 | 0.03681372 | 0.97063353 | 0.99443087 |
| Cs | 15903.2016 | -0.2912696 | 0.15939047 | -1.8273966 | 0.06764016 | 0.40069836 |
| Ampd3 | 510.453608 | 0.43591868 | 0.37337965 | 1.1674945 | 0.24301073 | 0.67814218 |
| Bcas2 | 4466.85111 | 0.18542791 | 0.20025391 | 0.92596397 | 0.35446471 | 0.76419569 |
| Ctcf | 602.706792 | -0.5063589 | 0.31928394 | -1.5859204 | 0.11275738 | 0.51163242 |
| Pard6a | 249.028836 | 0.79467887 | 0.36989863 | 2.14836931 | 0.03168443 | 0.26539272 |
| Tfap4 | 75.0887867 | -0.2737339 | 0.61534132 | -0.4448488 | 0.656429 | 0.91188916 |
| Ranbp1 | 18221.4017 | -0.1365528 | 0.19024104 | -0.7177884 | 0.47288779 | 0.83483892 |
| Rfx5 | 285.899658 | -0.0419962 | 0.46410856 | -0.0904879 | 0.92789952 | 0.98666423 |
| Psmb4 | 11729.7003 | 0.10690464 | 0.29781717 | 0.35896062 | 0.71962455 | 0.93314808 |
| Mmp8 | 208.513143 | -6.9196767 | 0.66531708 | -10.40057 | 2.464549538 | 2.740803137 |
| Slc30a4 | 44.1511071 | 1.87641075 | 1.15956517 | 1.61820207 | 0.10561906 | 0.49732022 |
| Sqor | 399.464707 | 1.88966842 | 0.39079007 | 4.8355078 | 1.328061693 | 9.336884310 |
| Bloc1s6 | 1413.17796 | 0.02997768 | 0.19272156 | 0.15554915 | 0.87638841 | 0.9745677 |
| Metap1 | 3847.9059 | -0.0252981 | 0.17300904 | -0.1462242 | 0.88374443 | 0.97685422 |
| Gpr108 | 1837.34666 | -0.3969032 | 0.28120489 | -1.4114378 | 0.15811556 | 0.58701903 |
| Tnfsf14 | 65.6695651 | -1.6361463 | 0.697623 | -2.3453159 | 0.01901097 | 0.19493809 |
| Rsl1d1 | 10692.6272 | -0.1250549 | 0.21273302 | -0.587849 | 0.55663367 | 0.87276537 |
| Apc | 240.380727 | 0.32413869 | 0.51103163 | 0.63428302 | 0.52589614 | 0.86056156 |
| Reep5 | 7265.84353 | 0.27706629 | 0.29294968 | 0.94578116 | 0.34426022 | 0.75962035 |
| Ergic3 | 4038.23878 | 0.0907737 | 0.23118742 | 0.39264119 | 0.69458451 | 0.92609254 |
| Uqcc1 | 3254.94476 | 0.16310628 | 0.30906592 | 0.52773943 | 0.5976802 | 0.88928406 |
| Ncoa2 | 247.550989 | -0.2701243 | 0.42140564 | -0.6410078 | 0.52151765 | 0.85808179 |
| Nr2c2 | 104.740011 | 0.57377909 | 0.6479561 | 0.88552155 | 0.3758754 | 0.77828094 |
| Nr2c1 | 445.088794 | 0.22284083 | 0.30362912 | 0.73392443 | 0.4629948 | 0.83156884 |
| Smpd4 | 2855.40106 | -0.0229872 | 0.18714876 | -0.1228284 | 0.90224298 | 0.98043702 |
| Pex1 | 412.684575 | -0.323234 | 0.44388961 | -0.7281857 | 0.46649994 | 0.83345004 |
| Kctd20 | 5147.34667 | -0.1783285 | 0.22044661 | -0.8089421 | 0.41854848 | 0.8052381 |
| Ctns | 1239.59945 | 0.57742238 | 0.24783856 | 2.32983267 | 0.019815 | 0.19950359 |
| Shpk | 97.7762198 | 0.6099308 | 0.72239427 | 0.84431843 | 0.39849149 | 0.79557554 |
| Tuft1 | 517.706767 | 0.3915093 | 0.33687675 | 1.16217369 | 0.24516492 | 0.68039867 |
| Rcn1 | 780.470481 | 0.03777282 | 0.23641636 | 0.15977244 | 0.87306034 | 0.97344599 |
| Dnase1 | 19.6815108 | -1.4130994 | 1.32153065 | -1.0692899 | 0.28493905 | 0.72045503 |
| Trap1 | 9460.11815 | -0.1700048 | 0.168432 | -1.0093377 | 0.31281269 | 0.74065408 |
| Naa60 | 3948.57973 | -0.0529705 | 0.24033133 | -0.2204063 | 0.82555476 | 0.96326388 |
| 1700037C18 | 100.647368 | -0.3375066 | 0.44066596 | -0.7659013 | 0.44373502 | 0.82068545 |
| Ankrd13d | 155.825678 | -0.3789478 | 0.52965652 | -0.7154595 | 0.47432518 | 0.83575195 |
| Tpr | 2882.18759 | -0.2556773 | 0.30180743 | -0.8471537 | 0.39690943 | 0.79492355 |

| | | | | | | |
|---|---|---|---|---|---|---|
| Odr4 | 1500.38448 | 0.4430125 | 0.27978326 | 1.5834132 | 0.11332732 | 0.51285577 |
| Dhx34 | 682.11906 | 0.06573696 | 0.35228229 | 0.18660308 | 0.85197185 | 0.96798445 |
| Kptn | 652.385349 | -0.0313714 | 0.25815437 | -0.1215221 | 0.90327755 | 0.98043702 |
| Napa | 8162.17671 | 0.20487931 | 0.14425496 | 1.42025831 | 0.15553249 | 0.58452503 |
| Sra1 | 3312.71635 | -0.0152603 | 0.26996692 | -0.0565265 | 0.95492236 | 0.99215259 |
| Atp5g1 | 10929.5926 | -0.196999 | 0.20235982 | -0.9735086 | 0.33030059 | 0.74978051 |
| Snf8 | 294.69443 | -0.3417059 | 0.27029521 | -1.2641951 | 0.20616001 | 0.64074069 |
| Tbcb | 4080.5732 | 0.34443453 | 0.275183 | 1.25165627 | 0.21069514 | 0.64401792 |
| Inpp5k | 1538.37885 | -0.0819348 | 0.1932641 | -0.4239523 | 0.67160062 | 0.91719387 |
| Crkl | 1327.20693 | 0.22429285 | 0.31695553 | 0.70764772 | 0.47916405 | 0.83869135 |
| Eps8l1 | 7.21696622 | 0.62427271 | 2.32757728 | 0.26820708 | 0.78853993 | 0.95225386 |
| Clint1 | 782.223773 | -0.7706088 | 0.26629134 | -2.8938559 | 0.00380543 | 0.06465527 |
| Prss16 | 39.7760911 | 0.10254529 | 1.2397045 | 0.08271753 | 0.93407615 | 0.98756396 |
| Cdkal1 | 629.81303 | 0.11688383 | 0.22622675 | 0.5166667 | 0.60538885 | 0.89258167 |
| Zbtb17 | 742.217103 | 0.17497638 | 0.24209602 | 0.72275611 | 0.46982973 | 0.83381643 |
| Fblim1 | 110.187864 | 1.63859711 | 1.04558175 | 1.56716307 | 0.11707658 | 0.52289076 |
| Epor | 152.957342 | 3.75662586 | 0.70460234 | 5.33155461 | 9.737553112 | 9.379487182 |
| Ccdc159 | 13.5803904 | -1.4183533 | 2.00952774 | -0.7058142 | 0.48030366 | 0.83935746 |
| Mob1b | 399.61122 | 0.41731877 | 0.4269338 | 0.97747887 | 0.32833213 | 0.74843736 |
| Atp6v1b2 | 22493.2298 | 0.14079068 | 0.14948978 | 0.94180804 | 0.34629093 | 0.7602916 |
| Eps15l1 | 1082.29828 | -0.1044484 | 0.2082923 | -0.5014511 | 0.61605368 | 0.89878056 |
| Tep1 | 1208.27004 | 0.20495124 | 0.43555139 | 0.47055582 | 0.63795796 | 0.90599527 |
| Ttc5 | 2708.82289 | 0.08993371 | 0.25840586 | 0.34803278 | 0.72781556 | 0.93576253 |
| Osgep | 4173.80817 | 0.28606701 | 0.32952376 | 0.86812256 | 0.38532724 | 0.78798197 |
| Aamp | 8388.78047 | -0.0010191 | 0.16741602 | -0.0060875 | 0.99514293 | 0.99884675 |
| Tmbim1 | 2789.34709 | 0.18283431 | 0.21404923 | 0.85416946 | 0.3930111 | 0.79137753 |
| Arpc2 | 34291.1448 | -0.0118172 | 0.14217553 | -0.0831172 | 0.93375836 | 0.98756396 |
| Kmt2b | 144.405538 | 0.12138802 | 0.36421003 | 0.33329126 | 0.73891443 | 0.93800224 |
| Tmem147 | 2653.50923 | 0.17078502 | 0.35237081 | 0.48467415 | 0.62790751 | 0.90239574 |
| Rps9 | 32771.7385 | 0.12911245 | 0.34989928 | 0.3689989 | 0.71212855 | 0.93235217 |
| Tfpt | 466.699411 | 0.0389255 | 0.35005566 | 0.11119803 | 0.91145931 | 0.98292561 |
| Ggt5 | 49.6168628 | 1.7726955 | 0.73294301 | 2.41859935 | 0.01558039 | 0.1732681 |
| Ggt1 | 12.3772927 | -0.4299023 | 1.83421221 | -0.2343798 | 0.81469014 | 0.95994071 |
| Crip2 | 174.958993 | 2.06025038 | 1.05408225 | 1.95454422 | 0.05063689 | 0.34442343 |
| Crip1 | 652.458368 | 1.29710352 | 0.37208265 | 3.48606292 | 0.00049019 | 0.01323718 |
| Cbfa2t3 | 168.034997 | -0.4236662 | 0.42054422 | -1.0074237 | 0.31373122 | 0.74065408 |
| Fbln1 | 145.028991 | 0.24691046 | 0.56946192 | 0.43358555 | 0.66458943 | 0.91431173 |
| Pgrmc1 | 5707.08093 | 0.50447874 | 0.26853668 | 1.87862139 | 0.06029621 | 0.37881203 |
| Gcat | 1183.62538 | 0.11118431 | 0.26544334 | 0.41886269 | 0.67531648 | 0.91848602 |
| Tek | 103.614118 | 2.23835395 | 1.05950391 | 2.11264341 | 0.0346313 | 0.2788971 |
| Elovl1 | 7478.76911 | -0.2674634 | 0.24624229 | -1.08618 | 0.27739938 | 0.71260013 |
| Med8 | 3645.84144 | 0.04749697 | 0.23172777 | 0.20496882 | 0.83759648 | 0.96683245 |
| Hyi | 71.4854318 | 0.81687189 | 0.76063086 | 1.07393999 | 0.28284958 | 0.71857502 |
| Cdc20 | 11019.3038 | -0.2491531 | 0.17125904 | -1.4548317 | 0.14571586 | 0.57242843 |
| Nectin4 | 79.5831231 | 0.91167262 | 0.68708084 | 1.32687824 | 0.18454896 | 0.61665869 |
| Pfdn2 | 1329.221 | -0.2839712 | 0.22760649 | -1.2476411 | 0.21216254 | 0.64580271 |

| | | | | | | |
|---|---|---|---|---|---|---|
| Rnf114 | 4397.6157 | -0.1593084 | 0.17299595 | -0.9208791 | 0.35711355 | 0.76632459 |
| C330007P06 | 1857.37467 | 0.34082547 | 0.23451929 | 1.45329395 | 0.14614216 | 0.5731306 |
| Neurl1a | 825.212943 | 0.37155718 | 0.36670285 | 1.01323777 | 0.31094661 | 0.74011268 |
| Srm | 12947.6283 | -0.6260985 | 0.16720361 | -3.7445274 | 0.00018073 | 0.00602429 |
| Epha2 | 19.7027886 | -0.6009606 | 2.44391841 | -0.2459005 | 0.80575929 | 0.95741157 |
| Rbm14 | 3611.41634 | -0.2572992 | 0.24931354 | -1.0320306 | 0.30205776 | 0.73361866 |
| Zdhhc24 | 329.172414 | -0.0412215 | 0.39300869 | -0.1048871 | 0.91646542 | 0.98386323 |
| Bbs1 | 35.2486877 | -1.4265105 | 1.01111947 | -1.4108229 | 0.15829683 | 0.5871574 |
| Ndor1 | 752.623304 | -0.2056627 | 0.22607004 | -0.90973 | 0.36296492 | 0.76990376 |
| Nsmf | 337.575478 | 0.78350946 | 0.47868268 | 1.63680344 | 0.10167155 | 0.48717118 |
| Pdk1 | 7335.52309 | 0.39936028 | 0.17996912 | 2.21904884 | 0.0264834 | 0.23919285 |
| Ptbp1 | 13540.9008 | -0.4162147 | 0.24201308 | -1.7198024 | 0.08546837 | 0.44669728 |
| Mvd | 2525.04491 | 0.64632716 | 0.22263568 | 2.90307093 | 0.00369523 | 0.06331056 |
| Cyba | 14646.6324 | 0.50343081 | 0.37696845 | 1.33547199 | 0.18172192 | 0.61290438 |
| Stimate | 304.465944 | 0.02748429 | 0.47507914 | 0.05785203 | 0.95386649 | 0.99171765 |
| Sfmbt1 | 1079.00568 | 0.33050555 | 0.2876248 | 1.14908574 | 0.25052063 | 0.6842534 |
| Rundc3a | 15.5653172 | -0.7447948 | 0.98103966 | -0.7591893 | 0.4477393 | 0.82332359 |
| Cdt1 | 3918.67813 | -0.2014172 | 0.2949853 | -0.6828042 | 0.49473058 | 0.84678075 |
| Runx1t1 | 12.8889051 | 2.78053584 | 1.41411774 | 1.96626897 | 0.04926755 | 0.34139551 |
| Aprt | 8156.77952 | -0.2300557 | 0.27579719 | -0.834148 | 0.40419761 | 0.79917613 |
| Gtf2h1 | 5242.16592 | -0.1750333 | 0.21324684 | -0.8208015 | 0.41175935 | 0.80245328 |
| Hfe | 3430.37785 | 1.27465435 | 0.29493214 | 4.32185641 | 1.547218752 | 0.00075307 |
| Abhd1 | 30.8700698 | -0.4080808 | 0.86892237 | -0.4696401 | 0.63861215 | 0.90615794 |
| Slc5a6 | 1360.45301 | 0.20540767 | 0.27717612 | 0.74107277 | 0.45864932 | 0.82986925 |
| Nphs1 | 35.010269 | 0.11365893 | 0.82138467 | 0.13837479 | 0.88994422 | 0.97871887 |
| Qrich1 | 6068.44262 | -0.0021009 | 0.23436898 | -0.0089642 | 0.99284768 | 0.99800348 |
| P4htm | 17.3440594 | 0.95477703 | 1.47964132 | 0.645276 | 0.51874834 | 0.8570838 |
| Usp19 | 1353.93442 | 0.0734426 | 0.25067313 | 0.29298152 | 0.76953627 | 0.94401697 |
| Pola1 | 1088.91794 | -0.2944047 | 0.20802799 | -1.4152167 | 0.15700496 | 0.58620126 |
| Cdc42 | 46438.3044 | 0.13879925 | 0.17248044 | 0.80472458 | 0.42097863 | 0.80708017 |
| Pknox1 | 1268.09724 | 0.00121108 | 0.23588133 | 0.00513429 | 0.99590345 | 0.99884675 |
| Gmnn | 4794.97504 | 0.24932414 | 0.3504202 | 0.71150047 | 0.47677416 | 0.83666308 |
| Acot13 | 1706.98653 | 1.02643224 | 0.406613 | 2.52434684 | 0.01159135 | 0.14165533 |
| Zfp184 | 92.5961947 | 1.55132365 | 0.6784979 | 2.28640892 | 0.02223035 | 0.21429774 |
| Cdk4 | 21332.831 | 0.100654 | 0.23369108 | 0.4307139 | 0.66667641 | 0.91552004 |
| B4galnt1 | 3838.50818 | -1.9511629 | 0.22344925 | -8.7320183 | 2.501678177 | 1.610685744 |
| Mettl1 | 715.45789 | -0.2758677 | 0.29356883 | -0.9397036 | 0.34736964 | 0.76101142 |
| Tspan31 | 6043.38346 | 0.32720099 | 0.22356957 | 1.46353101 | 0.14332213 | 0.56846902 |
| Kif5b | 5125.21966 | -0.0092809 | 0.20666355 | -0.0449085 | 0.96418024 | 0.99406401 |
| Saal1 | 1544.77044 | -0.1032706 | 0.30061718 | -0.3435287 | 0.73120078 | 0.93653187 |
| Cnp | 2257.73 | -0.1100339 | 0.16743211 | -0.6571852 | 0.51106181 | 0.85372377 |
| Ttc25 | 44.0792732 | 0.5069414 | 1.10042638 | 0.46067726 | 0.64503018 | 0.90835328 |
| Sulf2 | 21.8180711 | 0.26245307 | 1.86555527 | 0.14068362 | 0.88811989 | 0.97832097 |
| Sod2 | 4413.20802 | -0.1063826 | 0.18659462 | -0.570127 | 0.56859159 | 0.87936678 |
| Tmco6 | 291.955827 | -0.1948464 | 0.29040964 | -0.6709365 | 0.50226098 | 0.85012672 |
| Stambp | 963.283108 | -0.1222621 | 0.25404408 | -0.4812635 | 0.63032924 | 0.9036296 |

| | | | | | | |
|---|---|---|---|---|---|---|
| Ezh1 | 460.887342 | 0.37108923 | 0.3276844 | 1.13245925 | 0.25744141 | 0.69274795 |
| P3h4 | 49.5301133 | 1.36558692 | 0.77182349 | 1.76929951 | 0.07684391 | 0.42792174 |
| Ctnnb1 | 8126.06838 | 0.24899025 | 0.24619265 | 1.01136346 | 0.3118425 | 0.74030066 |
| Eif1b | 2321.75336 | 0.17873741 | 0.25459369 | 0.70204966 | 0.48264819 | 0.84106286 |
| Psmd2 | 21944.3679 | -0.0799605 | 0.1562603 | -0.5117136 | 0.60885145 | 0.89437372 |
| Vars | 8893.70635 | -0.3309413 | 0.25304379 | -1.3078421 | 0.19092686 | 0.62216522 |
| Vwa7 | 10.4709384 | 0.48655661 | 1.32557566 | 0.36705307 | 0.71357944 | 0.93235217 |
| Hspa1l | 199.958976 | 0.02485571 | 0.59892998 | 0.04150019 | 0.96689714 | 0.99443087 |
| Slc44a4 | 25.0142886 | -0.9241124 | 0.94339298 | -0.9795625 | 0.32730212 | 0.74811039 |
| Msh5 | 89.9933845 | 1.25993528 | 0.79002597 | 1.59480235 | 0.11075644 | 0.50858992 |
| Abhd16a | 1832.08478 | -0.2386175 | 0.15968185 | -1.4943307 | 0.13508921 | 0.55654288 |
| Neu1 | 2899.10654 | 0.58082467 | 0.24156101 | 2.40446369 | 0.01619621 | 0.17737538 |
| Ddah2 | 663.870592 | 0.99756856 | 0.331649 | 3.00790465 | 0.00263056 | 0.04860967 |
| Clic1 | 21413.9151 | 0.22828375 | 0.23024888 | 0.99146519 | 0.32145848 | 0.74447686 |
| Lsm2 | 3070.99885 | -0.2095128 | 0.15362943 | -1.3637542 | 0.17264493 | 0.60492813 |
| Pole | 645.205863 | 0.02334185 | 0.37877308 | 0.06162489 | 0.95086155 | 0.99093022 |
| Stx1a | 73.861734 | 0.52160077 | 0.52452778 | 0.99441973 | 0.32001857 | 0.74413737 |
| Zfp775 | 149.79713 | 0.71927669 | 0.63521605 | 1.13233394 | 0.25749407 | 0.69274795 |
| Mrpl49 | 3116.47437 | -0.0201944 | 0.24869703 | -0.0812009 | 0.93528221 | 0.98794094 |
| Dennd2c | 427.431075 | 0.37703783 | 0.29943181 | 1.25917762 | 0.20796618 | 0.6418276 |
| Mark3 | 213.915054 | -0.7787047 | 0.51781624 | -1.5038245 | 0.13262657 | 0.55221948 |
| Gatad1 | 2189.59073 | 0.11105231 | 0.21516741 | 0.51612047 | 0.60577027 | 0.89258167 |
| M6pr | 13177.0913 | -0.0297263 | 0.19280584 | -0.1541776 | 0.87746974 | 0.97458574 |
| Lrrc8a | 1417.87345 | 0.38596803 | 0.331027 | 1.16597146 | 0.24362598 | 0.67897148 |
| Ppp2r1a | 11380.8518 | -0.211656 | 0.19083963 | -1.1090776 | 0.26739669 | 0.70441538 |
| Fance | 714.506135 | -0.4628713 | 0.29874189 | -1.5494021 | 0.12128508 | 0.5291774 |
| Tinf2 | 1677.89502 | 0.04617751 | 0.2393296 | 0.19294526 | 0.84700184 | 0.96765585 |
| Dus3l | 1437.4168 | -0.0985603 | 0.2342577 | -0.4207346 | 0.67394889 | 0.91822108 |
| Gtpbp3 | 637.760272 | -0.0144207 | 0.29168081 | -0.0494401 | 0.96056857 | 0.99325523 |
| Tgfbr1 | 727.29931 | 0.69412851 | 0.27718158 | 2.50423753 | 0.01227156 | 0.14755327 |
| Homer1 | 127.757721 | -0.3411121 | 0.35139695 | -0.9707315 | 0.33168199 | 0.75068746 |
| Rad51c | 315.797097 | 0.34964779 | 0.37314429 | 0.93703107 | 0.34874258 | 0.76163771 |
| Cav1 | 28.4416708 | 1.27054217 | 1.87846889 | 0.67637115 | 0.49880503 | 0.84872332 |
| Arpp19 | 5002.64484 | 0.16978392 | 0.26804866 | 0.63340708 | 0.52646784 | 0.86111527 |
| Bcl2l1 | 1929.38275 | 0.17671274 | 0.21683157 | 0.81497695 | 0.4150855 | 0.80395688 |
| Khsrp | 265.684852 | -0.26506 | 0.38484161 | -0.6887509 | 0.49098004 | 0.84477968 |
| Ccdc124 | 2364.92326 | -0.1094079 | 0.16422653 | -0.6662011 | 0.50528259 | 0.85034006 |
| Cct4 | 28791.2616 | 0.10112141 | 0.29905492 | 0.33813657 | 0.73526027 | 0.93750666 |
| 0610009B22I | 756.753266 | 0.45484909 | 0.3649471 | 1.24634252 | 0.21263867 | 0.64596443 |
| Cpt1c | 33.4752581 | 2.42733829 | 1.20236882 | 2.01879678 | 0.04350835 | 0.31959733 |
| Zfp655 | 2990.48844 | 0.30565584 | 0.36399937 | 0.8397153 | 0.40106804 | 0.79698916 |
| Rhoa | 16380.6281 | 0.10748642 | 0.20850363 | 0.51551341 | 0.6061943 | 0.89258167 |
| Zmiz1 | 1521.56276 | -0.0036794 | 0.35973222 | -0.0102283 | 0.99183916 | 0.99738334 |
| Ankrd26 | 368.596995 | -0.2003477 | 0.6170059 | -0.3247095 | 0.74540095 | 0.93917909 |
| Aldh16a1 | 2891.78436 | -0.0670318 | 0.19402391 | -0.3454823 | 0.72973181 | 0.93649408 |
| Hnrnpa0 | 1037.23271 | -0.3347849 | 0.20655219 | -1.6208248 | 0.10505523 | 0.49581042 |

| | | | | | | |
|---|---|---|---|---|---|---|
| Prrg2 | 146.952072 | 0.7505186 | 0.39927858 | 1.87968659 | 0.0601508 | 0.37831609 |
| Hnrnph1 | 5767.96804 | -0.549057 | 0.22209509 | -2.4721709 | 0.01342953 | 0.1567552 |
| Ift43 | 99.5719595 | 0.25767494 | 0.63651803 | 0.40481955 | 0.68561016 | 0.92314423 |
| Id3 | 4096.79404 | 0.61055624 | 0.45046796 | 1.35538218 | 0.17529582 | 0.607477 |
| Arid1a | 1897.08756 | -0.3434735 | 0.35100896 | -0.978532 | 0.32781127 | 0.74843736 |
| Ctsd | 56356.7388 | 0.89180297 | 0.18370924 | 4.85442638 | 1.207357148 | 8.586976744 |
| Rplp1 | 13579.4244 | -0.1232347 | 0.31546108 | -0.3906495 | 0.69605632 | 0.92623267 |
| Abhd8 | 241.699713 | 0.13911166 | 0.33682422 | 0.41300969 | 0.67959952 | 0.92051478 |
| Ift22 | 531.080688 | 0.79889447 | 0.30492179 | 2.61999798 | 0.00879303 | 0.11898797 |
| Fzd3 | 24.2618625 | 1.69849122 | 2.12512756 | 0.79924201 | 0.4241501 | 0.80933211 |
| Mid1ip1 | 3127.42204 | -0.0096519 | 0.17492691 | -0.055177 | 0.95599747 | 0.99215259 |
| Ap2s1 | 5752.02258 | 0.59622579 | 0.21315712 | 2.79711878 | 0.00515606 | 0.08191435 |
| Fgfrl1 | 11.7645741 | 1.04892191 | 1.69515991 | 0.61877461 | 0.53606485 | 0.8654666 |
| 4930432K21I | 28.8374579 | 3.00780502 | 1.15397219 | 2.60647965 | 0.00914783 | 0.12230094 |
| Fhl2 | 93.6219057 | -0.702887 | 0.48152904 | -1.459698 | 0.1443731 | 0.57027792 |
| Emc10 | 1740.49384 | -0.442883 | 0.27027651 | -1.6386294 | 0.10129047 | 0.48631215 |
| Clstn3 | 16.6954464 | -0.1213562 | 2.10925327 | -0.0575352 | 0.9541189 | 0.99173562 |
| Fbxw9 | 300.297988 | 0.98599753 | 0.52534637 | 1.87685226 | 0.06053834 | 0.3793881 |
| Fnbp4 | 802.684992 | -0.4382967 | 0.34073687 | -1.2863203 | 0.19833131 | 0.63122167 |
| Cers4 | 178.688399 | -0.2249019 | 0.541048 | -0.4156782 | 0.67764551 | 0.91958179 |
| Scrn3 | 666.464985 | 0.04701604 | 0.30213478 | 0.15561281 | 0.87633823 | 0.9745677 |
| Phax | 1902.19714 | -0.2798556 | 0.15906911 | -1.7593332 | 0.07852094 | 0.43353356 |
| Tle1 | 45.6142579 | -0.12667 | 0.98606666 | -0.1284599 | 0.89778505 | 0.97988977 |
| Relt | 1078.96023 | -0.4507426 | 0.22935274 | -1.9652812 | 0.0493817 | 0.34139551 |
| Snrpb2 | 3485.84355 | -0.0718926 | 0.22503397 | -0.3194746 | 0.74936663 | 0.9399191 |
| Ubc | 111511.326 | 0.10456822 | 0.24840566 | 0.42095746 | 0.67378614 | 0.91822108 |
| Prpf31 | 8402.98299 | -0.312876 | 0.16736146 | -1.8694624 | 0.06155852 | 0.38288591 |
| Sertad1 | 1256.03389 | 0.42877723 | 0.3030401 | 1.41491911 | 0.15709222 | 0.58620126 |
| Carhsp1 | 3501.12854 | 0.77759628 | 0.17178318 | 4.5266149 | 5.993601689 | 0.00033944 |
| Elk3 | 633.653135 | 0.11128573 | 0.22925017 | 0.48543356 | 0.62736883 | 0.9022352 |
| Herpud2 | 525.686044 | -0.2492755 | 0.31932528 | -0.780632 | 0.43501894 | 0.81606911 |
| Rdh13 | 1444.01268 | -0.0672698 | 0.18891548 | -0.3560844 | 0.72177738 | 0.93398714 |
| Nutf2 | 3074.37763 | -0.009565 | 0.23749868 | -0.0402738 | 0.96787481 | 0.99443087 |
| Arpc5 | 34728.0286 | 0.11895864 | 0.27023007 | 0.44021244 | 0.65978325 | 0.91280998 |
| Pou2f2 | 430.418312 | -0.8065409 | 0.39448929 | -2.0445192 | 0.04090228 | 0.30810197 |
| Mgst1 | 123.903608 | -0.5313117 | 0.84905251 | -0.6257702 | 0.5314657 | 0.86432064 |
| Ubqln4 | 993.504349 | -0.1799859 | 0.19749973 | -0.9113223 | 0.36212558 | 0.76947754 |
| Rps18 | 33975.5978 | 0.52073035 | 0.48633375 | 1.07072634 | 0.2842925 | 0.71974495 |
| Rpl10 | 8623.63734 | 0.10944481 | 0.27129975 | 0.40340916 | 0.68664725 | 0.92416721 |
| Rps15a | 10209.7102 | 0.20475024 | 0.40169727 | 0.5097128 | 0.61025268 | 0.89508835 |
| Ncaph2 | 9182.81656 | 0.13296048 | 0.16897574 | 0.78686137 | 0.43136303 | 0.81332674 |
| Hipk1 | 1314.93252 | -0.4269188 | 0.34079059 | -1.2527307 | 0.21030372 | 0.64378984 |
| Man1a2 | 1786.52574 | 0.15783761 | 0.36845401 | 0.42837807 | 0.66837589 | 0.91604147 |
| Acyp1 | 300.232368 | 0.13365877 | 0.5331466 | 0.25069797 | 0.80204763 | 0.95632784 |
| Hdac5 | 1199.56349 | 0.02703713 | 0.23189702 | 0.11659112 | 0.90718408 | 0.98176897 |
| Rala | 1455.83793 | 0.00226549 | 0.17066147 | 0.01327477 | 0.98940858 | 0.99650865 |

| | | | | | | |
|---|---|---|---|---|---|---|
| Vdac3 | 12430.9246 | -0.0643037 | 0.18445231 | -0.3486195 | 0.72737502 | 0.93576253 |
| Vps72 | 1605.26363 | 0.07437705 | 0.19575828 | 0.37994332 | 0.70398749 | 0.92850227 |
| Gabpa | 1230.40741 | -0.0003447 | 0.2612612 | -0.0013194 | 0.99894728 | 0.99969186 |
| Dynll1 | 8597.74939 | 0.05226115 | 0.28889717 | 0.18089879 | 0.85644702 | 0.96924897 |
| Pdcl | 2023.70502 | 0.44750776 | 0.23729144 | 1.88589932 | 0.05930852 | 0.37572303 |
| Tmem184b | 1672.53682 | 0.17799479 | 0.29917238 | 0.5949573 | 0.55187201 | 0.87093412 |
| Nf2 | 1196.3446 | -0.0121969 | 0.31655236 | -0.0385304 | 0.96926477 | 0.99443087 |
| Zmat5 | 1000.02514 | -0.0154934 | 0.29630767 | -0.0522883 | 0.95829897 | 0.99249408 |
| Ewsr1 | 19959.6202 | -0.2885151 | 0.19606687 | -1.471514 | 0.14115216 | 0.56557954 |
| Ap1b1 | 9214.16991 | 0.79850197 | 0.28768043 | 2.77565615 | 0.00550904 | 0.08595935 |
| Gnat2 | 53.7808962 | 2.05269712 | 0.94889477 | 2.16325052 | 0.03052192 | 0.26073645 |
| Bcl2l13 | 2443.37881 | -0.3848539 | 0.2707731 | -1.4213151 | 0.15522517 | 0.58428346 |
| Dqx1 | 12.0630596 | -0.5238296 | 1.20586854 | -0.4344003 | 0.66399779 | 0.91399629 |
| Lnpk | 1063.76189 | -0.0037334 | 0.20401114 | -0.0182998 | 0.98539972 | 0.99564261 |
| Ascl2 | 122.003714 | 0.78465229 | 0.47876776 | 1.63889961 | 0.10123417 | 0.48631215 |
| Pttg1ip | 4414.18503 | 0.30569168 | 0.17578274 | 1.73903132 | 0.08202925 | 0.44108302 |
| Trpm2 | 253.662204 | 0.47962858 | 0.36491058 | 1.3143729 | 0.18872074 | 0.61935434 |
| Ube2g2 | 5153.524 | -0.0327813 | 0.17803831 | -0.184125 | 0.85391538 | 0.96839628 |
| Mpo | 20639.6532 | -6.5175074 | 0.31192021 | -20.89479 | 5.971842262 | 7.305354640 |
| Met | 75.4335515 | -3.5677909 | 0.96887208 | -3.6824169 | 0.00023103 | 0.00727862 |
| Slc16a12 | 45.8255416 | 3.1162517 | 1.61822716 | 1.92571957 | 0.0541394 | 0.35989907 |
| Elk1 | 129.998512 | 0.26234758 | 0.69762442 | 0.37605849 | 0.70687341 | 0.92935286 |
| Nav1 | 169.607112 | 0.7413171 | 0.36718084 | 2.01894277 | 0.04349317 | 0.31959733 |
| Tnpo1 | 4746.68998 | -0.199802 | 0.18174251 | -1.0993685 | 0.27160736 | 0.70934517 |
| Rnmt | 2714.51421 | -0.2177426 | 0.22325834 | -0.9752943 | 0.32941429 | 0.74874249 |
| Srp14 | 4126.33218 | 0.45209128 | 0.3785476 | 1.19427854 | 0.23236903 | 0.6674267 |
| Cdk9 | 672.010145 | -0.446696 | 0.33613578 | -1.3289153 | 0.18387591 | 0.61583 |
| Tor2a | 1433.47385 | 0.30202354 | 0.21279229 | 1.41933499 | 0.15580138 | 0.58481688 |
| Fpgs | 1304.49024 | -0.0777795 | 0.23441657 | -0.3318003 | 0.74004007 | 0.93800224 |
| Mrtfb | 393.571398 | 0.57577546 | 0.3438265 | 1.67461048 | 0.09401068 | 0.46882701 |
| Cbx5 | 7314.91492 | -0.12107 | 0.15206483 | -0.7961735 | 0.42593119 | 0.81021713 |
| Apobec3 | 4960.01501 | -0.2934583 | 0.14041059 | -2.0900014 | 0.03661767 | 0.28732779 |
| Vav2 | 572.299667 | 0.10045009 | 0.36429276 | 0.27574001 | 0.78274776 | 0.95075701 |
| Ppp2cb | 204.772638 | -0.7823839 | 0.40957191 | -1.910248 | 0.05610129 | 0.36429573 |
| G0s2 | 67.6198681 | -1.13949 | 0.84201994 | -1.3532814 | 0.17596574 | 0.60763832 |
| Dmap1 | 1529.87637 | -0.1898677 | 0.20401817 | -0.9306413 | 0.35203912 | 0.76348958 |
| Mcu | 412.783896 | -0.2530684 | 0.28302553 | -0.894154 | 0.37123951 | 0.7765614 |
| Oit3 | 27.2739516 | 0.76475442 | 1.34075747 | 0.57038982 | 0.56841333 | 0.87936678 |
| Bcr | 73.067143 | -2.5017377 | 1.06485957 | -2.3493593 | 0.01880575 | 0.19348256 |
| Fxyd5 | 2386.80996 | -0.8435615 | 0.18802124 | -4.4865221 | 7.239520074 | 0.00039536 |
| Kcnd1 | 258.583193 | -0.6816277 | 0.39306666 | -1.7341274 | 0.08289548 | 0.4429983 |
| Tfcp2 | 179.647115 | 0.72939354 | 0.62203451 | 1.17259337 | 0.24095891 | 0.67637686 |
| Ubp1 | 437.511946 | -0.3851207 | 0.3059011 | -1.2589712 | 0.20804075 | 0.6418276 |
| Nuak2 | 640.307125 | -1.614901 | 0.38710471 | -4.171742 | 3.022798073 | 0.00129747 |
| Ick | 268.687072 | 0.88217073 | 0.30935722 | 2.8516248 | 0.00434964 | 0.07200157 |
| Sdhb | 8959.6163 | 0.11651596 | 0.22649311 | 0.51443488 | 0.60694798 | 0.89307142 |

| | | | | | | |
|---|---|---|---|---|---|---|
| Cox4i2 | 515.9582 | -1.0947347 | 0.35848258 | -3.0538017 | 0.00225961 | 0.04359914 |
| Snap47 | 1008.2539 | 0.52544109 | 0.21050576 | 2.49608892 | 0.01255711 | 0.14928197 |
| Kdsr | 537.28564 | 0.03499291 | 0.25221946 | 0.13873994 | 0.88965566 | 0.97870754 |
| Vps4b | 5963.2732 | 0.072487 | 0.21102568 | 0.34349847 | 0.73122349 | 0.93653187 |
| Rps25 | 20232.6824 | 0.43331337 | 0.44074756 | 0.98313278 | 0.32554209 | 0.74706419 |
| Nxf2 | 25.4382768 | -0.5522638 | 1.99291888 | -0.277113 | 0.78169331 | 0.95004728 |
| Taz | 1574.73435 | 0.24885144 | 0.24333666 | 1.0226632 | 0.30646712 | 0.73582655 |
| Kif19a | 35.8741395 | 2.21189587 | 1.26332852 | 1.75084773 | 0.07997214 | 0.43723638 |
| Aldh3a2 | 2229.98977 | -0.2536113 | 0.26357179 | -0.9622094 | 0.3359444 | 0.75405649 |
| Tmem115 | 1057.58214 | 0.09490816 | 0.19049118 | 0.49822865 | 0.61832289 | 0.89878056 |
| Hyal2 | 839.649067 | 0.31802385 | 0.32025921 | 0.99302016 | 0.32070013 | 0.74413737 |
| Ifrd2 | 1470.73131 | -0.5766853 | 0.1966292 | -2.9328567 | 0.00335859 | 0.05886192 |
| Hyal1 | 29.5952994 | 1.78175342 | 0.9613132 | 1.85345778 | 0.06381677 | 0.39033529 |
| Tusc2 | 1820.4907 | 0.0892847 | 0.26227131 | 0.34042878 | 0.73353365 | 0.93735685 |
| Nprl2 | 720.577001 | 0.47816495 | 0.39376853 | 1.21433002 | 0.22462173 | 0.65815511 |
| Rassf1 | 1978.90879 | 0.08907531 | 0.19404063 | 0.45905495 | 0.64619471 | 0.9088181 |
| Slc3a2 | 17021.7541 | 0.14999047 | 0.24291317 | 0.61746537 | 0.53692781 | 0.86596304 |
| Nxf1 | 4961.05096 | -0.1464959 | 0.17947031 | -0.8162679 | 0.41434693 | 0.80395688 |
| Stx5a | 1896.40389 | 0.29184829 | 0.19484739 | 1.49783009 | 0.1341774 | 0.55489931 |
| Tnfrsf13b | 3862.38306 | -0.347138 | 0.18356422 | -1.8910982 | 0.05861123 | 0.37246298 |
| Prox1 | 65.3439254 | 0.79951533 | 1.42731768 | 0.56015234 | 0.57537553 | 0.88070182 |
| Raver1 | 1914.63507 | -0.5035209 | 0.28877892 | -1.7436205 | 0.08122528 | 0.43985341 |
| 2610507B11I | 3224.66364 | 0.21982605 | 0.28733706 | 0.76504593 | 0.44424419 | 0.82068545 |
| AI597479 | 920.41739 | -0.1919315 | 0.2356156 | -0.814596 | 0.4153036 | 0.80395688 |
| Tmem86a | 1381.65677 | 1.31547098 | 0.23203564 | 5.66926268 | 1.434133499 | 1.866356925 |
| Ifi35 | 1466.0902 | -0.0583922 | 0.2913177 | -0.2004416 | 0.84113522 | 0.96765585 |
| Rdm1 | 323.113735 | 0.109251 | 0.39664588 | 0.27543713 | 0.78298042 | 0.95078415 |
| Nedd8 | 7755.8862 | 0.04272588 | 0.23785008 | 0.17963368 | 0.85744016 | 0.96941456 |
| Gosr1 | 1586.35397 | -0.1363046 | 0.19486579 | -0.6994795 | 0.48425241 | 0.84128398 |
| Mrpl52 | 395.864318 | 0.32925135 | 0.3896848 | 0.84491709 | 0.39815713 | 0.79546892 |
| Kansl3 | 849.51976 | -0.1562757 | 0.30110507 | -0.5190071 | 0.60375578 | 0.89232143 |
| Faf1 | 3113.42814 | 0.0018355 | 0.25752524 | 0.00712744 | 0.99431317 | 0.99880383 |
| Tsacc | 64.7025464 | 0.29355665 | 0.73528805 | 0.39924033 | 0.68971613 | 0.92470267 |
| Mettl16 | 2864.00211 | -0.2351511 | 0.17392998 | -1.3519869 | 0.1763795 | 0.60764153 |
| Pigyl | 668.927621 | 0.72122399 | 0.3767202 | 1.91448186 | 0.05555861 | 0.3627292 |
| Rbm25 | 3054.8974 | -0.2768458 | 0.27329731 | -1.0129839 | 0.31106784 | 0.74011268 |
| Psen2 | 556.409299 | -0.5072933 | 0.40331616 | -1.2578054 | 0.20846213 | 0.6418276 |
| Acaa1b | 15.1641994 | 1.7218883 | 1.56845875 | 1.09782185 | 0.27228229 | 0.71005689 |
| Plcd1 | 365.276393 | 1.2813747 | 0.51478722 | 2.48913461 | 0.01280545 | 0.15120563 |
| Fads1 | 2564.49544 | 0.78735641 | 0.2318604 | 3.39582094 | 0.00068423 | 0.01743791 |
| Lmbr1 | 102.555043 | 0.19587276 | 0.62369776 | 0.31405076 | 0.75348248 | 0.94018119 |
| Tnfrsf22 | 91.3656232 | 0.56077992 | 0.46231342 | 1.21298648 | 0.22513499 | 0.65870021 |
| Cars | 3059.15005 | -0.2085942 | 0.20044044 | -1.0406791 | 0.29802451 | 0.730169 |
| Apip | 1688.83334 | 0.60277163 | 0.39074064 | 1.54263868 | 0.12291847 | 0.53283544 |
| Pdhx | 2471.26349 | 0.02456892 | 0.20647815 | 0.11899043 | 0.90528293 | 0.9814201 |
| Vac14 | 1062.77588 | 0.00713144 | 0.23212563 | 0.03072232 | 0.97549099 | 0.99450266 |

| | | | | | | |
|---|---|---|---|---|---|---|
| Mcoln2 | 1764.11222 | 1.07899527 | 0.22295744 | 4.83946734 | 1.301875809 | 9.205691781 |
| Akt1s1 | 728.398138 | 0.25939413 | 0.23872183 | 1.08659579 | 0.27721549 | 0.71247429 |
| Tbrg1 | 2463.51551 | 0.50002968 | 0.30485674 | 1.64021197 | 0.1009611 | 0.48566935 |
| Adssl1 | 907.678206 | -1.2796396 | 0.29981438 | -4.2681061 | 1.971395365 | 0.00089985 |
| Brf1 | 773.951626 | -0.3993024 | 0.23042161 | -1.732921 | 0.08310971 | 0.4429983 |
| Vipr2 | 47.5694728 | -1.3556087 | 0.89755879 | -1.5103286 | 0.13095961 | 0.54940637 |
| Odc1 | 1390.66771 | -0.2149026 | 0.3169728 | -0.6779844 | 0.49778156 | 0.84857327 |
| Thg1l | 744.263953 | -0.0303718 | 0.21732829 | -0.1397508 | 0.88885691 | 0.97852652 |
| Adam19 | 241.731744 | 0.41812054 | 0.53836715 | 0.77664571 | 0.43736782 | 0.8185065 |
| Pabpc4 | 711.823217 | -1.0430196 | 0.30838985 | -3.3821463 | 0.00071922 | 0.01810316 |
| Exoc3l2 | 64.3855882 | -1.4651343 | 0.79807446 | -1.8358366 | 0.06638182 | 0.39699582 |
| Zfp296 | 63.8332312 | -1.9329625 | 0.70618354 | -2.7371957 | 0.00619654 | 0.09399815 |
| Sugp1 | 1512.80299 | -0.0928141 | 0.22187562 | -0.418316 | 0.67571608 | 0.91855038 |
| Dhdh | 570.120423 | 0.03207322 | 0.20789223 | 0.15427809 | 0.87739047 | 0.97458574 |
| Zfp790 | 411.824011 | -0.0754955 | 0.34665228 | -0.2177844 | 0.82759706 | 0.96423051 |
| Fuz | 112.411362 | 0.20879612 | 0.45835072 | 0.4555379 | 0.64872231 | 0.90973357 |
| Sptbn4 | 15.5708663 | 1.14060184 | 1.13806832 | 1.00222616 | 0.31623438 | 0.74186126 |
| Pgam1 | 48507.2585 | -0.273946 | 0.15833454 | -1.7301719 | 0.08359956 | 0.44386868 |
| Evi5 | 220.115577 | 0.79685877 | 0.44618191 | 1.78595042 | 0.07410729 | 0.41931287 |
| Evi5l | 209.785403 | 0.0637946 | 0.47397561 | 0.13459469 | 0.89293234 | 0.97941957 |
| Snapc2 | 1082.04406 | -0.3190103 | 0.19145494 | -1.6662421 | 0.09566521 | 0.47437068 |
| Git1 | 359.301069 | -0.0006311 | 0.37944825 | -0.0016631 | 0.99867301 | 0.99967651 |
| Gltp | 4244.85868 | 0.22443328 | 0.20739876 | 1.08213411 | 0.27919294 | 0.71377266 |
| Bnip2 | 8508.83024 | 0.15624776 | 0.16818762 | 0.92900873 | 0.35288456 | 0.76376537 |
| Ccnt1 | 2056.19915 | -0.0066686 | 0.49869685 | -0.0133721 | 0.98933094 | 0.99650865 |
| Brms1l | 620.978855 | -0.0123249 | 0.45388131 | -0.0271545 | 0.97833647 | 0.99483019 |
| Med15 | 1112.29546 | -0.0223533 | 0.34586367 | -0.0646305 | 0.94846822 | 0.99019648 |
| Dhdds | 1433.30109 | -0.1358684 | 0.18680201 | -0.7273393 | 0.46701812 | 0.8334534 |
| Crybg2 | 41.7327314 | -0.2826821 | 1.11315855 | -0.2539459 | 0.79953735 | 0.95515043 |
| Ubxn11 | 90.2494434 | -0.1950375 | 0.75055727 | -0.2598569 | 0.79497416 | 0.95367917 |
| Tjap1 | 561.586794 | -0.3653354 | 0.42224226 | -0.8652269 | 0.38691425 | 0.78911204 |
| Rpl15 | 28828.8651 | 0.16757912 | 0.33904 | 0.49427537 | 0.62111172 | 0.89896588 |
| Tmem167 | 5141.04922 | 0.05346755 | 0.21723926 | 0.24612289 | 0.80558711 | 0.95737871 |
| Steap4 | 62.754459 | 1.46411993 | 1.81365979 | 0.80727375 | 0.41950879 | 0.80563709 |
| Mplkip | 664.499243 | 0.40678412 | 0.30755302 | 1.32264712 | 0.18595276 | 0.61790851 |
| Kif11 | 3541.46589 | -0.3351564 | 0.20480696 | -1.6364504 | 0.10174537 | 0.48733401 |
| Rpa3 | 1483.36741 | 0.49387097 | 0.50651026 | 0.97504632 | 0.32953728 | 0.74874249 |
| Mlkl | 213.112316 | 1.78388377 | 0.36075086 | 4.94491896 | 7.617554641 | 5.787922107 |
| Tnpo3 | 5908.87401 | -0.264109 | 0.25056595 | -1.0540499 | 0.29186005 | 0.72575203 |
| Ttll5 | 298.794187 | -0.7024332 | 0.51690872 | -1.3589115 | 0.17417464 | 0.60653633 |
| Zfp715 | 658.335686 | 0.61990379 | 0.25568536 | 2.424479 | 0.01533037 | 0.17119251 |
| Retn | 15.2720067 | -0.6364472 | 1.29474068 | -0.4915635 | 0.62302798 | 0.90017327 |
| Cdh23 | 67.6990247 | -1.2289132 | 0.93718675 | -1.3112789 | 0.18976354 | 0.62019167 |
| Rps5 | 33456.819 | 0.1487308 | 0.36082584 | 0.41219553 | 0.68019611 | 0.92061965 |
| Adgrl1 | 45.4070708 | -0.1026106 | 0.74516798 | -0.1377013 | 0.89047647 | 0.97871887 |
| Amotl1 | 64.5191194 | 0.25032822 | 0.79597436 | 0.31449282 | 0.75314677 | 0.94018119 |

| | | | | | | |
|---|---|---|---|---|---|---|
| Etv5 | 6356.755 | -0.0112454 | 0.27496617 | -0.0408974 | 0.9673777 | 0.99443087 |
| Gfod2 | 195.064893 | -0.0649067 | 0.40751241 | -0.1592753 | 0.87345196 | 0.97344599 |
| Enkd1 | 251.182552 | -0.1537995 | 0.30341305 | -0.506898 | 0.61222641 | 0.89650055 |
| Atp6v0d1 | 4043.42517 | 0.34768997 | 0.16261055 | 2.13817594 | 0.03250247 | 0.2677459 |
| Ptprs | 62.1954556 | -0.1596018 | 0.70772308 | -0.2255145 | 0.82157903 | 0.96231102 |
| Slc41a1 | 195.778335 | -0.7098789 | 0.44747328 | -1.5864163 | 0.11264492 | 0.51139203 |
| Igf2bp1 | 8909.62699 | -0.2517976 | 0.21156601 | -1.190161 | 0.23398314 | 0.66970419 |
| Nelfb | 4371.97146 | 0.04835331 | 0.13808349 | 0.35017446 | 0.72620777 | 0.93561871 |
| Tmem175 | 1823.16889 | 0.05393432 | 0.20391339 | 0.26449621 | 0.79139759 | 0.95268772 |
| Tango2 | 2292.19954 | 0.82495843 | 0.32309244 | 2.5533201 | 0.01067014 | 0.13344172 |
| Aldh1a2 | 30.8377677 | 0.83626228 | 0.77895016 | 1.07357611 | 0.28301271 | 0.71857502 |
| Ndufs2 | 12520.9342 | 0.03441858 | 0.17454315 | 0.19719239 | 0.84367699 | 0.96765585 |
| Atraid | 2953.94165 | 0.40560233 | 0.29428958 | 1.37824222 | 0.16812852 | 0.60261241 |
| Cad | 1497.88088 | -0.8843834 | 0.27654496 | -3.1979734 | 0.00138397 | 0.029592 |
| Sh3bp5l | 578.072931 | -0.0047713 | 0.24646779 | -0.0193587 | 0.98455493 | 0.99564261 |
| Atad1 | 1964.09199 | -0.0813276 | 0.18457969 | -0.4406095 | 0.65949575 | 0.91280998 |
| Pten | 593.931603 | -0.3925324 | 0.23751435 | -1.6526682 | 0.09839841 | 0.48011435 |
| Pea15a | 5811.75922 | 0.8497868 | 0.19333142 | 4.39549244 | 1.105219126 | 0.00056706 |
| Timm23 | 15182.5594 | 0.0435214 | 0.33756942 | 0.12892577 | 0.89741639 | 0.97984623 |
| Tnfaip8l2 | 2431.9829 | 0.71279541 | 0.39366698 | 1.81065582 | 0.07019415 | 0.40928741 |
| Trnt1 | 2888.04136 | 0.05272715 | 0.28379776 | 0.18579128 | 0.85260845 | 0.96798445 |
| Ehmt2 | 1579.26984 | -0.1043915 | 0.20606172 | -0.5066033 | 0.61243319 | 0.89669602 |
| Elof1 | 4134.92868 | 0.63938278 | 0.41785954 | 1.53013805 | 0.12598256 | 0.53789768 |
| Med16 | 1410.19149 | -0.2019824 | 0.3111845 | -0.6490759 | 0.5162893 | 0.85627177 |
| St3gal1 | 245.818987 | 0.38900186 | 0.50566115 | 0.76929354 | 0.44171907 | 0.81967485 |
| Tmem259 | 186.030451 | -0.6973647 | 0.40236578 | -1.7331612 | 0.08306703 | 0.4429983 |
| Rnf170 | 531.171217 | 0.27593818 | 0.34507806 | 0.79963987 | 0.42391948 | 0.80917688 |
| Thap8 | 10.8246067 | 2.99637593 | 1.56643979 | 1.91285738 | 0.05576631 | 0.36325309 |
| Dedd | 1070.27769 | -0.0255424 | 0.24556035 | -0.104017 | 0.91715587 | 0.98390186 |
| Mcemp1 | 495.98358 | -3.3609443 | 0.36523191 | -9.2022199 | 3.506298350 | 2.859503181 |
| Nit1 | 952.199991 | -0.0589989 | 0.31437833 | -0.1876685 | 0.85113654 | 0.9679302 |
| Prdm15 | 102.796958 | 0.10137464 | 0.59672232 | 0.16988579 | 0.86509996 | 0.9705375 |
| Rnf168 | 1295.21969 | -0.2326752 | 0.20510999 | -1.1343924 | 0.25663 | 0.6916575 |
| Tctex1d2 | 260.274507 | -0.4914133 | 0.4840872 | -1.0151339 | 0.31004201 | 0.73989254 |
| Chp1 | 2023.58201 | -0.4662629 | 0.3215414 | -1.4500866 | 0.14703438 | 0.57373895 |
| Trpv4 | 95.1647271 | 0.97129516 | 0.73771715 | 1.31662271 | 0.18796512 | 0.61935434 |
| Klhl3 | 74.8706256 | 2.21276745 | 0.93327734 | 2.37096451 | 0.01774173 | 0.18742197 |
| Tvp23b | 1451.42484 | 0.20897603 | 0.25228351 | 0.82833804 | 0.40747909 | 0.8005951 |
| Dnajc7 | 3748.245 | -0.2662452 | 0.22734614 | -1.1711008 | 0.24155828 | 0.67708722 |
| Cacybp | 17429.152 | -0.1762782 | 0.28645588 | -0.6153764 | 0.5383062 | 0.86663962 |
| Cluap1 | 516.193021 | -0.409635 | 0.33688048 | -1.2159654 | 0.22399811 | 0.65774578 |
| Zswim7 | 385.212343 | 0.07882241 | 0.34869327 | 0.22605084 | 0.82116188 | 0.96205421 |
| Pigl | 555.535827 | 0.412422 | 0.23923811 | 1.72389761 | 0.08472633 | 0.44578804 |
| Ndufa2 | 4727.6669 | 0.29383332 | 0.34191996 | 0.85936286 | 0.39014035 | 0.78985201 |
| Pam16 | 535.251491 | -0.3457969 | 0.38173931 | -0.9058458 | 0.36501749 | 0.77118172 |
| Glis2 | 24.1087166 | 1.02265032 | 0.84974063 | 1.20348526 | 0.22878859 | 0.66258779 |

| | | | | | | |
|---|---|---|---|---|---|---|
| Cox6c | 4680.75692 | 0.20001473 | 0.36557703 | 0.54712063 | 0.58429586 | 0.88404713 |
| Ube2z | 471.11469 | -0.205368 | 0.38677501 | -0.5309754 | 0.59543581 | 0.88830085 |
| Tmem87b | 747.916674 | 1.24724138 | 0.29939478 | 4.16587545 | 3.10160007€ | 0.00132664 |
| Anapc1 | 1493.77 | -0.4952294 | 0.29889307 | -1.6568783 | 0.09754409 | 0.47887037 |
| Mertk | 1081.70995 | 0.71720723 | 0.34419884 | 2.08370028 | 0.03718744 | 0.28956966 |
| Tsg101 | 3920.56848 | 0.21641468 | 0.17272148 | 1.2529691 | 0.21021695 | 0.64378984 |
| Hps5 | 688.051419 | 0.02727362 | 0.45085775 | 0.06049275 | 0.95176319 | 0.99122417 |
| Map3k4 | 757.269225 | 0.03882616 | 0.29337285 | 0.13234407 | 0.89471215 | 0.97941957 |
| Piezo1 | 609.147308 | -0.1832617 | 0.38728818 | -0.4731922 | 0.63607608 | 0.90531997 |
| Rnf166 | 664.493968 | 0.03144411 | 0.30132717 | 0.10435205 | 0.91688997 | 0.98389485 |
| Ankrd28 | 221.392539 | 0.37363106 | 0.36772308 | 1.0160664 | 0.30959777 | 0.73898722 |
| Ankrd52 | 199.199034 | -0.6302818 | 0.52604022 | -1.1981628 | 0.23085366 | 0.6653874 |
| Pkd2l2 | 15.6858877 | 1.92383795 | 1.08687156 | 1.77006928 | 0.0767156 | 0.42754531 |
| Srp19 | 3361.39022 | 0.04666582 | 0.26308879 | 0.1773767 | 0.85921251 | 0.96954641 |
| Wdfy2 | 1027.72011 | 0.31064 | 0.25268595 | 1.22935206 | 0.21893984 | 0.65247231 |
| Rbsn | 973.842763 | 0.13078942 | 0.30022821 | 0.43563335 | 0.66310277 | 0.91395288 |
| Mrps25 | 2094.46928 | 0.22583703 | 0.27032822 | 0.83541788 | 0.40348247 | 0.79841492 |
| Dguok | 656.832441 | -0.035519 | 0.21661252 | -0.1639748 | 0.86975101 | 0.97239152 |
| Camta1 | 301.832351 | 0.32874394 | 0.39182499 | 0.83900707 | 0.40146534 | 0.79737342 |
| Csf1 | 105.300664 | 0.53663899 | 0.59104639 | 0.90794732 | 0.36390605 | 0.77045439 |
| Strip1 | 2711.68849 | 0.16557863 | 0.27012513 | 0.61297012 | 0.5398961 | 0.86719392 |
| Slc25a11 | 6293.60662 | 0.0973613 | 0.19092639 | 0.50994158 | 0.61009239 | 0.89508835 |
| Cmc2 | 1151.51212 | -0.1469298 | 0.28598086 | -0.5137749 | 0.60740939 | 0.89332558 |
| Chfr | 2800.33699 | -0.171197 | 0.19062924 | -0.8980624 | 0.36915226 | 0.77480275 |
| Tex261 | 3299.89413 | -0.0167787 | 0.17376613 | -0.0965589 | 0.92307672 | 0.98465273 |
| Fam120b | 1347.14311 | 0.29483776 | 0.23201851 | 1.27075102 | 0.20381724 | 0.63824694 |
| Tbp | 1448.58874 | -0.0575806 | 0.25836108 | -0.2228688 | 0.82363764 | 0.9628698 |
| Psmb1 | 13763.4513 | 0.28109375 | 0.34680406 | 0.81052612 | 0.41763786 | 0.80506837 |
| Pdcd2 | 1223.73981 | 0.25410902 | 0.33215704 | 0.76502673 | 0.44425563 | 0.82068545 |
| Fhod1 | 51.73607 | -0.8311941 | 0.63687731 | -1.3051087 | 0.19185585 | 0.62286958 |
| Elmo3 | 20.2440016 | 2.02003909 | 1.9080021 | 1.05871953 | 0.28972753 | 0.72395811 |
| Stc1 | 14.7982245 | 1.6824359 | 1.69958973 | 0.98990708 | 0.32221954 | 0.7445747 |
| 4931428F04l | 168.300232 | 1.02947565 | 0.68501546 | 1.50285024 | 0.13287767 | 0.55270061 |
| Tppp3 | 42.9430412 | 0.89278456 | 0.80551205 | 1.10834414 | 0.26771322 | 0.70441538 |
| Msh3 | 538.781449 | -0.0999886 | 0.26519622 | -0.3770364 | 0.70614657 | 0.92896305 |
| Tmem208 | 2136.37273 | 0.60460781 | 0.44457405 | 1.359971 | 0.1738391 | 0.60653633 |
| E2f4 | 1140.28805 | -0.3229108 | 0.25704351 | -1.2562494 | 0.20902557 | 0.64228937 |
| Surf4 | 14601.8519 | 0.10563921 | 0.13772229 | 0.76704514 | 0.44305468 | 0.82038676 |
| Surf2 | 966.087584 | -0.3424279 | 0.25490604 | -1.3433494 | 0.17915885 | 0.60992118 |
| Dnajb9 | 1717.56973 | 0.48378751 | 0.22508182 | 2.14938512 | 0.03160388 | 0.26539272 |
| Naf1 | 75.9482799 | -1.3229737 | 0.52601684 | -2.5150786 | 0.01190058 | 0.14428131 |
| Yes1 | 221.800544 | 0.51738951 | 0.63521524 | 0.81451055 | 0.41535253 | 0.80395688 |
| Ppp1cb | 3445.52634 | -0.0420245 | 0.19231587 | -0.2185179 | 0.82702558 | 0.96379955 |
| Gorasp2 | 6600.14693 | -0.2774699 | 0.23949287 | -1.1585727 | 0.24663039 | 0.68166056 |
| Tsen15 | 2247.4673 | 0.60275228 | 0.34837583 | 1.73017825 | 0.08359843 | 0.44386868 |
| Efr3a | 4090.22363 | 0.16700943 | 0.22582983 | 0.73953664 | 0.4595812 | 0.82987212 |

| | | | | | | |
|---|---|---|---|---|---|---|
| Trappc2l | 2502.22216 | 0.24822586 | 0.38377962 | 0.64679271 | 0.51776611 | 0.85708158 |
| Acsf3 | 455.813797 | -0.2269432 | 0.25820563 | -0.8789245 | 0.37944224 | 0.78211296 |
| Ddx19a | 574.951124 | -0.2352147 | 0.32614771 | -0.7211907 | 0.47079219 | 0.83381643 |
| Galns | 1592.53399 | -0.4028609 | 0.19031158 | -2.1168493 | 0.03427264 | 0.27692023 |
| Gata2 | 85.4608143 | -0.3017444 | 0.70870158 | -0.4257707 | 0.67027495 | 0.91711586 |
| C8g | 31.187341 | -1.042492 | 0.87321085 | -1.1938606 | 0.2325325 | 0.66773946 |
| Rabl6 | 1961.5425 | 0.24163788 | 0.20730515 | 1.16561447 | 0.24377036 | 0.67897148 |
| Edf1 | 3514.94646 | 0.09026841 | 0.2464993 | 0.36620147 | 0.71421475 | 0.93235217 |
| Fbxw5 | 1424.97767 | -0.1345539 | 0.24103455 | -0.5582347 | 0.57668412 | 0.88105119 |
| Slc25a13 | 3010.00578 | -0.2861381 | 0.18547035 | -1.5427698 | 0.12288663 | 0.53283544 |
| Ube2i | 9094.44049 | -0.096988 | 0.13736746 | -0.7060479 | 0.48015836 | 0.83935746 |
| Tsr3 | 1038.46922 | 0.09639981 | 0.23366728 | 0.41255161 | 0.67993516 | 0.92054299 |
| Unkl | 114.33494 | 0.09299698 | 0.5475086 | 0.16985484 | 0.8651243 | 0.9705375 |
| Lrrk1 | 360.122658 | -0.3001109 | 0.36540882 | -0.8213018 | 0.41147439 | 0.80231592 |
| Aldh1a3 | 67.3253839 | -0.1859772 | 0.64076881 | -0.2902408 | 0.77163206 | 0.94526087 |
| Actn1 | 90.6451472 | 0.30151851 | 0.45805329 | 0.65826078 | 0.51037057 | 0.85335831 |
| Sirt2 | 1868.47836 | 0.49895367 | 0.28661118 | 1.74087305 | 0.08170583 | 0.44050571 |
| Hnrnpl | 736.342499 | 0.11063471 | 0.25625091 | 0.43174368 | 0.66592772 | 0.91520501 |
| Nolc1 | 6007.26677 | -0.5285408 | 0.26579322 | -1.9885414 | 0.04675184 | 0.33444287 |
| Casd1 | 278.504873 | -0.266913 | 0.4175093 | -0.6392984 | 0.52262882 | 0.85816354 |
| Cnksr3 | 112.182158 | 1.09076464 | 0.66825043 | 1.63226927 | 0.10262277 | 0.48942862 |
| Mtmr1 | 194.368481 | -0.4771905 | 0.47619366 | -1.0020933 | 0.31629855 | 0.74186126 |
| Hmgb3 | 4045.11457 | 0.51237605 | 0.27417615 | 1.86878416 | 0.06165285 | 0.38323136 |
| Abca1 | 1311.75948 | 1.1233299 | 0.38579735 | 2.91170977 | 0.00359456 | 0.06219563 |
| Nipsnap3b | 3997.08133 | 0.3328938 | 0.29377306 | 1.13316654 | 0.25714432 | 0.6927211 |
| Lage3 | 1588.374 | 0.46685642 | 0.45672749 | 1.02217718 | 0.30669705 | 0.73594057 |
| Ubl4a | 2729.38179 | 0.63422084 | 0.39754756 | 1.59533325 | 0.11063773 | 0.50858841 |
| Gdi1 | 1201.44197 | -0.1913419 | 0.23929382 | -0.7996106 | 0.42393645 | 0.80917688 |
| Sash1 | 123.734133 | 0.39415511 | 0.49197578 | 0.80116772 | 0.42303456 | 0.80846457 |
| Gadd45b | 535.13132 | 0.44201113 | 0.47844578 | 0.9238479 | 0.35556551 | 0.7647925 |
| Slamf6 | 362.367314 | -0.4720983 | 0.32695706 | -1.4439154 | 0.14876278 | 0.57589084 |
| Slamf1 | 38.1227287 | 2.38190628 | 1.11554807 | 2.13518928 | 0.03274555 | 0.26920453 |
| Zdhhc12 | 1092.83554 | -0.3734518 | 0.28944159 | -1.2902494 | 0.19696408 | 0.62932233 |
| Endog | 128.500129 | -0.2953258 | 0.44150784 | -0.6689027 | 0.50355756 | 0.85012672 |
| Cybb | 8187.66668 | -1.7014497 | 0.14327769 | -11.875189 | 1.592730202 | 3.247311427 |
| Golga7 | 6695.46562 | 0.03012242 | 0.18544208 | 0.16243575 | 0.87096272 | 0.97268995 |
| Xk | 98.9935844 | 0.45552441 | 0.57589876 | 0.79098001 | 0.42895566 | 0.81140481 |
| Cd48 | 1319.81348 | -0.1384432 | 0.35032264 | -0.3951878 | 0.69270433 | 0.92549717 |
| Clpx | 2437.75594 | 0.1240434 | 0.23168416 | 0.53539871 | 0.59237418 | 0.88797037 |
| Trabd | 1300.88165 | 0.62509395 | 0.30645385 | 2.03976533 | 0.0413737 | 0.30955629 |
| Dennd6b | 57.5493545 | 0.17126309 | 0.68328197 | 0.25064775 | 0.80208646 | 0.95632784 |
| Cd83 | 23721.9088 | 0.91941399 | 0.4697117 | 1.95740065 | 0.05030038 | 0.34299029 |
| Atf6b | 1478.81959 | -0.6213224 | 0.29473152 | -2.1080963 | 0.03502266 | 0.28075507 |
| Notch4 | 18.8455375 | 2.29840925 | 1.55477884 | 1.47828694 | 0.13933099 | 0.56326372 |
| Ppt2 | 116.067371 | -0.4277552 | 0.38793478 | -1.1026472 | 0.2701804 | 0.70727945 |
| Rnf5 | 2450.41074 | 0.14494646 | 0.15903217 | 0.91142856 | 0.36206961 | 0.76947754 |

| | | | | | | |
|---|---|---|---|---|---|---|
| Cacfd1 | 546.666489 | -0.2422547 | 0.28453722 | -0.8513991 | 0.39454769 | 0.79263117 |
| Hivep2 | 221.866692 | 0.35236901 | 0.48880818 | 0.7208738 | 0.47098717 | 0.83381643 |
| Arnt | 1522.38233 | 0.06757075 | 0.36715431 | 0.1840391 | 0.85398278 | 0.96839628 |
| Mocs2 | 775.115925 | 0.61289208 | 0.33285598 | 1.84131309 | 0.06557569 | 0.39558985 |
| Nat9 | 502.893244 | 0.45141902 | 0.33185115 | 1.3603057 | 0.1737332 | 0.60653633 |
| Lpl | 22090.1124 | 0.12270067 | 0.14404821 | 0.85180282 | 0.39432354 | 0.79233901 |
| Atp6v0e | 6419.03614 | 0.13654649 | 0.26593787 | 0.51345258 | 0.60763477 | 0.89332558 |
| Zfp318 | 108.373854 | -0.3667117 | 0.48726421 | -0.7525932 | 0.45169439 | 0.82570905 |
| Srf | 178.281741 | -0.9822114 | 0.4170726 | -2.3550131 | 0.01852205 | 0.19153023 |
| Gata3 | 38.8897115 | -1.905472 | 0.71007321 | -2.6834867 | 0.00728589 | 0.10438985 |
| Steap1 | 36.2024508 | 2.1890854 | 0.6682178 | 3.2760058 | 0.00105286 | 0.02425859 |
| Hspa8 | 288520.306 | 0.00065078 | 0.1716215 | 0.00379193 | 0.99697448 | 0.99942546 |
| Serac1 | 27.3484314 | -0.7184917 | 0.92969973 | -0.7728212 | 0.43962819 | 0.81963526 |
| Pdzd11 | 2799.10056 | 0.07305756 | 0.21433595 | 0.34085536 | 0.73321247 | 0.93728136 |
| Psma2 | 16912.6894 | 0.15946399 | 0.32303012 | 0.49365052 | 0.62155301 | 0.89912494 |
| Mrpl32 | 2481.10108 | 0.23446295 | 0.37200862 | 0.63026215 | 0.52852308 | 0.86217134 |
| Setdb1 | 2065.45929 | 0.28554916 | 0.30629737 | 0.93226123 | 0.35120154 | 0.76309919 |
| Anxa9 | 108.098682 | -0.4371396 | 0.69899645 | -0.6253817 | 0.5317206 | 0.8643494 |
| Prune1 | 417.301735 | -0.3333614 | 0.27957095 | -1.1924036 | 0.23310303 | 0.66870814 |
| Cers2 | 7703.4605 | -0.1548715 | 0.18296012 | -0.8464768 | 0.3972868 | 0.79515861 |
| Capza2 | 46564.4458 | -0.0102865 | 0.23190492 | -0.0443566 | 0.96462019 | 0.99406401 |
| Plekho1 | 676.696082 | 0.1973727 | 0.3074649 | 0.64193573 | 0.52091491 | 0.8576517 |
| Vps45 | 2040.42518 | 0.52544774 | 0.2698304 | 1.94732594 | 0.05149567 | 0.34938799 |
| Prpf3 | 2232.374 | 0.02652371 | 0.17939237 | 0.14785305 | 0.88245874 | 0.97605043 |
| Anp32e | 6652.49738 | -0.2869203 | 0.15433809 | -1.8590375 | 0.06302183 | 0.38721549 |
| Aph1a | 1242.01609 | -0.2357767 | 0.21789607 | -1.0820605 | 0.27922564 | 0.71377266 |
| Tab2 | 995.990502 | -0.0284508 | 0.23057493 | -0.1233908 | 0.90179767 | 0.98043702 |
| Ppil4 | 2131.24416 | -0.2054221 | 0.23785473 | -0.8636451 | 0.38778287 | 0.7893659 |
| Cnih1 | 5774.1827 | 0.37472753 | 0.36825793 | 1.01756812 | 0.30888324 | 0.73867239 |
| Eps8 | 2139.97704 | 1.25859531 | 0.26530053 | 4.7440362 | 2.095013898 | 0.00013705 |
| Med22 | 921.8053 | -0.097589 | 0.29053665 | -0.3358921 | 0.73695223 | 0.93800224 |
| Surf1 | 1330.40011 | -0.1637839 | 0.16571592 | -0.9883416 | 0.32298537 | 0.74562749 |
| Med28 | 4807.5669 | 0.33592728 | 0.30110693 | 1.11564115 | 0.26457577 | 0.70112124 |
| Qdpr | 4556.91755 | 0.2449739 | 0.25087965 | 0.97645984 | 0.32883664 | 0.74854087 |
| Sqstm1 | 14367.2177 | 0.34845695 | 0.3230753 | 1.07856266 | 0.28078274 | 0.7155865 |
| Nfe2l2 | 11285.2328 | 0.23074047 | 0.38664365 | 0.59677813 | 0.55065552 | 0.87093412 |
| Rxra | 226.357625 | 1.43110532 | 0.5886729 | 2.43107051 | 0.01505428 | 0.16957556 |
| Adamtsl4 | 12.3783379 | -3.5595382 | 1.39972558 | -2.5430258 | 0.01098971 | 0.13620785 |
| Fcrls | 20803.5681 | 1.85163205 | 0.28898349 | 6.40739731 | 1.480247222 | 3.084209888 |
| Cd5l | 60.5856552 | 3.25606493 | 1.52812675 | 2.13075579 | 0.03310927 | 0.27019723 |
| Prpsap1 | 4375.29208 | -0.2302058 | 0.16849859 | -1.3662182 | 0.1718705 | 0.60433798 |
| Ncapg | 3963.57279 | -0.1515565 | 0.18077822 | -0.8383557 | 0.40183095 | 0.79752354 |
| Lcorl | 366.916146 | 0.94243206 | 0.51805271 | 1.81918179 | 0.0688837 | 0.40473309 |
| Lta4h | 5445.20754 | -0.483567 | 0.16365142 | -2.9548597 | 0.00312811 | 0.05561949 |
| Dstn | 8816.50378 | 0.00719909 | 0.28675714 | 0.02510519 | 0.97997106 | 0.99486654 |
| H2afy | 4339.31963 | -0.7222617 | 0.1860423 | -3.8822445 | 0.0001035 | 0.00369118 |

| | | | | | | |
|---|---|---|---|---|---|---|
| Gtf2ird2 | 239.399663 | 0.85846934 | 0.38889714 | 2.20744575 | 0.02728293 | 0.24474748 |
| Bola1 | 747.574835 | 0.25811501 | 0.39302659 | 0.65673676 | 0.51135019 | 0.85373917 |
| Castor2 | 160.683014 | 1.14125543 | 0.7259291 | 1.57213071 | 0.11592024 | 0.52081758 |
| Fcgr1 | 7484.60944 | 0.18311502 | 0.26806302 | 0.68310436 | 0.4945409 | 0.84678075 |
| Ncf1 | 10759.9529 | -0.270889 | 0.21280622 | -1.2729375 | 0.20304021 | 0.63809363 |
| Adss | 4057.07141 | -0.3410625 | 0.19637396 | -1.7368008 | 0.08242233 | 0.44183004 |
| Cacna1d | 28.3091051 | -0.7965452 | 1.87155767 | -0.4256055 | 0.67039535 | 0.91711586 |
| Actr8 | 1776.60118 | -0.011829 | 0.18864273 | -0.062706 | 0.95000065 | 0.99065364 |
| Lrrc27 | 52.7939688 | 0.11414047 | 0.67496478 | 0.16910582 | 0.86571341 | 0.9705375 |
| Fnta | 3921.39645 | 0.35475581 | 0.27616567 | 1.28457607 | 0.19894047 | 0.63148061 |
| Mtrex | 4023.91707 | -0.4279388 | 0.26729752 | -1.6009831 | 0.10938067 | 0.5054982 |
| Lbp | 155.157585 | -1.1665175 | 0.45326233 | -2.5736035 | 0.01006455 | 0.12973625 |
| Fli1 | 5469.29614 | 0.17417122 | 0.16766671 | 1.03879427 | 0.29890042 | 0.73093388 |
| Diexf | 231.321293 | -0.0352391 | 0.44499727 | -0.0791895 | 0.93688186 | 0.98843258 |
| Hsd11b1 | 230.562395 | -4.7077413 | 0.62103268 | -7.580505 | 3.442124129 | 1.275984984 |
| Syt14 | 40.3666966 | 0.46215403 | 1.01783272 | 0.45405696 | 0.64978783 | 0.90973357 |
| H2-M3 | 685.518591 | 1.29546786 | 0.41741334 | 3.10356125 | 0.00191207 | 0.03885434 |
| Lonrf3 | 39.3598155 | 2.41130979 | 1.33124201 | 1.81132339 | 0.07009081 | 0.40907483 |
| Atp5e | 3387.37124 | 0.72650106 | 0.54037464 | 1.3444396 | 0.17880628 | 0.60992118 |
| Nelfcd | 2529.12177 | -0.2229181 | 0.18092983 | -1.2320694 | 0.21792315 | 0.65150232 |
| Ctsz | 53653.9153 | -0.2961788 | 0.18846261 | -1.5715519 | 0.11605451 | 0.52084358 |
| Prelid3b | 5116.01962 | -0.3871276 | 0.19744309 | -1.9607047 | 0.04991348 | 0.34225988 |
| Ube2a | 3537.67985 | 0.23835251 | 0.31746424 | 0.75080115 | 0.45277234 | 0.82610332 |
| Slc25a5 | 71626.97 | 0.05376443 | 0.28609952 | 0.18792214 | 0.85093769 | 0.9679302 |
| Ppdpf | 1828.93451 | 0.30357115 | 0.19583786 | 1.55011471 | 0.12111399 | 0.52912326 |
| Pls3 | 849.62586 | 0.600457 | 0.28523298 | 2.10514579 | 0.03527862 | 0.28154973 |
| Mpped2 | 23.9578912 | 3.10596583 | 1.47982139 | 2.09887885 | 0.03582758 | 0.28424306 |
| Nkap | 709.555981 | 0.34280337 | 0.22032784 | 1.55587864 | 0.11973695 | 0.52718991 |
| Ndufa1 | 2122.65985 | 0.14506073 | 0.33820836 | 0.42890937 | 0.66798918 | 0.91595755 |
| E2f3 | 441.457155 | -0.2141299 | 0.43309837 | -0.4944141 | 0.62101378 | 0.89896588 |
| Cr1l | 3124.96202 | 0.45553054 | 0.33520009 | 1.35898095 | 0.17415263 | 0.60653633 |
| Ppfibp1 | 640.751387 | 0.25946738 | 0.41016905 | 0.63258643 | 0.52700375 | 0.86130085 |
| Cd34 | 5627.3847 | -2.6109526 | 0.31827824 | -8.2033653 | 2.337496042 | 1.143783563 |
| Plgrkt | 1146.89917 | 0.44538311 | 0.33426484 | 1.33242581 | 0.1827203 | 0.61407074 |
| Cd274 | 111.647307 | -0.3840688 | 0.66231269 | -0.5798904 | 0.56198853 | 0.87543686 |
| Gtf3a | 4240.64556 | 0.03775202 | 0.31858447 | 0.11849925 | 0.90567209 | 0.98144617 |
| Mtif3 | 728.945827 | 0.68425126 | 0.35267865 | 1.94015504 | 0.05236085 | 0.35368873 |
| Lnx2 | 501.707017 | -0.2711368 | 0.45427197 | -0.5968601 | 0.55060082 | 0.87093412 |
| Dyrk3 | 256.395168 | -1.3594474 | 0.37508679 | -3.6243542 | 0.00028968 | 0.00866432 |
| Mapkapk2 | 1719.74848 | -0.1408259 | 0.26288475 | -0.5356945 | 0.59216968 | 0.88797037 |
| Il10 | 33.8386046 | 0.98527642 | 1.36325084 | 0.72274037 | 0.46983941 | 0.83381643 |
| Lamp2 | 12017.7972 | 0.56942016 | 0.22528318 | 2.52757512 | 0.01148533 | 0.14064062 |
| Atxn10 | 13288.9819 | -0.0454153 | 0.16543815 | -0.2745151 | 0.78368884 | 0.95094799 |
| Foxred2 | 76.0361929 | 0.43142526 | 0.67206617 | 0.64193866 | 0.52091301 | 0.8576517 |
| Eif3d | 9867.03476 | -0.1776623 | 0.18823426 | -0.9438363 | 0.3452533 | 0.76016623 |
| H3f3b | 35654.7729 | 0.03705237 | 0.18472886 | 0.20057707 | 0.84102929 | 0.96765585 |

| | | | | | | |
|---|---|---|---|---|---|---|
| Nup50 | 4074.10998 | -0.1659698 | 0.21451318 | -0.7737044 | 0.43910564 | 0.81963526 |
| Ift27 | 713.278606 | -0.0232698 | 0.23708163 | -0.0981511 | 0.92181234 | 0.98437722 |
| Pacsin2 | 8522.44015 | 0.16430038 | 0.24378731 | 0.67394966 | 0.50034332 | 0.84938937 |
| Cmah | 859.846581 | 0.79957453 | 0.31104806 | 2.570582 | 0.01015278 | 0.13017076 |
| Ttll12 | 1352.65384 | -0.2298162 | 0.27237467 | -0.8437504 | 0.39880888 | 0.7957029 |
| Tox4 | 6178.32745 | -0.0819888 | 0.20603651 | -0.3979334 | 0.6906793 | 0.92470267 |
| Mrps18c | 2257.59519 | 0.38148464 | 0.36427916 | 1.04723158 | 0.29499278 | 0.72771732 |
| Srsf6 | 12707.0241 | -0.1817114 | 0.18676748 | -0.9729286 | 0.33058881 | 0.75003046 |
| Plcg1 | 295.692373 | 0.02781523 | 0.36527148 | 0.07614946 | 0.93930019 | 0.98913801 |
| Kctd2 | 26.7875933 | 0.03310809 | 1.02616706 | 0.03226384 | 0.97426164 | 0.99446549 |
| Kctd5 | 3371.37914 | -0.0684795 | 0.18978825 | -0.3608207 | 0.71823346 | 0.93286094 |
| Etaa1 | 566.197313 | 0.29627508 | 0.38719195 | 0.76518916 | 0.44415892 | 0.82068545 |
| Slpi | 126.212981 | -3.1517291 | 0.45052222 | -6.9957239 | 2.638928150 | 7.173779569 |
| Sdc4 | 2114.06147 | 2.61582673 | 0.36534169 | 7.15994594 | 8.070888504 | 2.468279476 |
| Il13ra1 | 3289.67709 | 0.05860625 | 0.20506689 | 0.28579089 | 0.77503829 | 0.94725181 |
| Nbr1 | 1617.86687 | 0.54055437 | 0.31381809 | 1.72250865 | 0.08497742 | 0.44637738 |
| Cyth1 | 435.958325 | 0.41691339 | 0.26671434 | 1.56314576 | 0.11801831 | 0.52479754 |
| Rnd3 | 167.715517 | -0.5498938 | 0.37695735 | -1.4587693 | 0.14462862 | 0.57032823 |
| Brca1 | 414.050319 | 0.07027136 | 0.57136936 | 0.12298762 | 0.9021169 | 0.98043702 |
| Nt5c3b | 1371.67411 | 0.59272328 | 0.34514105 | 1.7173364 | 0.08591772 | 0.44801003 |
| Coa3 | 3719.3746 | 0.43706837 | 0.40640409 | 1.07545269 | 0.2821721 | 0.71778152 |
| Med24 | 1914.58035 | 0.10657804 | 0.20494071 | 0.52004329 | 0.60303341 | 0.89168472 |
| Psmd3 | 7897.2705 | -0.3039089 | 0.21958664 | -1.3840044 | 0.16635711 | 0.59960123 |
| Exosc10 | 3872.06999 | -0.293122 | 0.23993714 | -1.2216618 | 0.22183554 | 0.65525654 |
| Glod4 | 4829.32711 | 0.08489451 | 0.20240909 | 0.41942043 | 0.67490889 | 0.91847374 |
| Vps53 | 1769.17887 | 0.07861536 | 0.20726394 | 0.37930072 | 0.70446457 | 0.9285501 |
| Taok1 | 972.013852 | 0.3076729 | 0.32545614 | 0.94535904 | 0.34447561 | 0.75972469 |
| Dnttip1 | 78.2002548 | -0.282037 | 0.90987468 | -0.3099735 | 0.75658112 | 0.94102069 |
| Acot8 | 813.482214 | 0.06251546 | 0.2659116 | 0.23509867 | 0.81413215 | 0.95988497 |
| Cd300lg | 231.003563 | -0.2401531 | 0.59948186 | -0.400601 | 0.68871388 | 0.92470267 |
| Nlk | 189.953751 | -0.7221923 | 0.42372006 | -1.704409 | 0.08830471 | 0.45403754 |
| Traf4 | 213.910559 | -0.4624132 | 0.3444052 | -1.342643 | 0.17938761 | 0.60992118 |
| Aldoc | 157.69137 | 0.20567736 | 0.48985194 | 0.41987659 | 0.67457561 | 0.91832667 |
| Rpl19 | 37320.4391 | 0.24010009 | 0.39918913 | 0.60146952 | 0.54752731 | 0.87008334 |
| Nek8 | 134.044712 | 0.31591814 | 0.485803 | 0.65030093 | 0.51549786 | 0.85587477 |
| Cacnb4 | 22.7904259 | 0.04330565 | 0.93156024 | 0.04648723 | 0.96292191 | 0.99401755 |
| Arl5b | 792.985954 | 0.49224004 | 0.42624397 | 1.15483168 | 0.24815936 | 0.68235696 |
| Zfp207 | 13558.7207 | 0.07913537 | 0.24886046 | 0.31799094 | 0.75049181 | 0.9399191 |
| Psmd11 | 8812.88749 | -0.1864597 | 0.16125606 | -1.1562954 | 0.24756033 | 0.68235696 |
| Timp2 | 209.983598 | 2.24159935 | 0.61859326 | 3.62370474 | 0.00029041 | 0.00866494 |
| Zc3h18 | 1048.64182 | -0.382846 | 0.27328338 | -1.4009123 | 0.16124031 | 0.59141542 |
| Top2b | 1206.5325 | -0.8049401 | 0.43582828 | -1.8469204 | 0.06475868 | 0.39236898 |
| Rarb | 18.2727471 | 1.30086228 | 1.59527362 | 0.81544775 | 0.41481606 | 0.80395688 |
| Igfbp4 | 6452.08323 | 1.25643453 | 0.29096631 | 4.31814429 | 1.573465577 | 0.0007608 |
| Cdc6 | 5186.39162 | -0.0827245 | 0.25510486 | -0.3242763 | 0.74572885 | 0.93939872 |
| Suz12 | 585.151061 | -0.5098408 | 0.33215519 | -1.5349474 | 0.12479675 | 0.53626132 |

| | | | | | | |
|---|---|---|---|---|---|---|
| Atad5 | 926.319533 | -0.4154434 | 0.25163357 | -1.6509858 | 0.09874148 | 0.48025164 |
| Crlf3 | 5294.47346 | -0.3140759 | 0.17857022 | -1.758837 | 0.0786052 | 0.43353356 |
| Tns4 | 280.976374 | 0.65360787 | 0.6276306 | 1.04138943 | 0.29769484 | 0.72979999 |
| Tnfaip1 | 4522.51678 | 0.23942213 | 0.2160883 | 1.10798284 | 0.26786923 | 0.70441538 |
| Abr | 1131.80971 | -0.6181701 | 0.28787617 | -2.1473473 | 0.03176564 | 0.26554509 |
| Cd40 | 25.2181336 | 4.50989844 | 1.5197481 | 2.96753023 | 0.00300203 | 0.05400559 |
| Slc35c2 | 4519.36695 | -0.5591772 | 0.24316931 | -2.2995386 | 0.02147437 | 0.21049361 |
| Zfp334 | 64.3825172 | 1.30177262 | 1.25165386 | 1.04004203 | 0.29832038 | 0.73074753 |
| Elmo2 | 4901.40846 | -0.2189767 | 0.22119647 | -0.9899647 | 0.32219136 | 0.7445747 |
| Wsb1 | 11770.1739 | 0.26847809 | 0.26620395 | 1.00854284 | 0.31319393 | 0.74065408 |
| Ttpal | 907.524163 | -0.3508719 | 0.33808647 | -1.0378171 | 0.29935521 | 0.73113229 |
| Rhot1 | 3669.60346 | 0.16935109 | 0.16116575 | 1.05078835 | 0.29335581 | 0.72646828 |
| Ada | 1743.18777 | 0.35033693 | 0.17804554 | 1.96768154 | 0.04910469 | 0.34139551 |
| Serinc3 | 24164.0691 | -0.0514326 | 0.17964299 | -0.2863047 | 0.77464479 | 0.94709254 |
| Tha1 | 104.094605 | 0.70194835 | 0.471127 | 1.48993447 | 0.13624147 | 0.55833898 |
| Pgs1 | 2709.50163 | -0.5731217 | 0.28267546 | -2.0274902 | 0.0426123 | 0.31638902 |
| Birc5 | 9159.99118 | 0.0737295 | 0.22507454 | 0.32757815 | 0.74323065 | 0.93856621 |
| Afmid | 67.5391099 | 0.16008424 | 0.74033543 | 0.21623204 | 0.82880688 | 0.96462215 |
| Pigt | 761.972501 | -0.5348153 | 0.29723969 | -1.7992729 | 0.07197553 | 0.41298154 |
| Dbndd2 | 194.704626 | 1.05708203 | 0.56535507 | 1.8697666 | 0.06151624 | 0.38288591 |
| Mmp9 | 300.189502 | 2.04996354 | 0.47126709 | 4.34989749 | 1.362011894 | 0.00068849 |
| Ghdc | 459.056146 | 0.3508002 | 0.26132974 | 1.34236616 | 0.1794773 | 0.60992118 |
| Pltp | 128.844698 | 1.71536021 | 0.81476675 | 2.10533898 | 0.03526181 | 0.28154973 |
| Slc12a7 | 1202.36891 | 1.05081605 | 0.38564379 | 2.72483594 | 0.00643334 | 0.09680087 |
| Ctsa | 23511.6606 | 0.26767871 | 0.14928905 | 1.79302305 | 0.07296924 | 0.41525386 |
| Zswim1 | 587.392854 | -0.0872542 | 0.31834947 | -0.274083 | 0.7840208 | 0.95094799 |
| Slc12a4 | 584.835399 | -0.3799071 | 0.29456361 | -1.2897284 | 0.19714496 | 0.62935134 |
| Myo1c | 1530.96235 | 0.56871171 | 0.31647912 | 1.79699598 | 0.07233626 | 0.41349974 |
| Crk | 2046.04563 | 0.30260469 | 0.30066121 | 1.00646403 | 0.31419241 | 0.74065408 |
| Cox7c | 6840.58068 | 0.40607796 | 0.42826842 | 0.94818561 | 0.34303497 | 0.75947281 |
| Pitpna | 728.523333 | -0.4665949 | 0.23103818 | -2.0195577 | 0.04342928 | 0.31959733 |
| Mlx | 2490.51208 | 0.3793589 | 0.32309537 | 1.17413909 | 0.24033932 | 0.67564175 |
| Retreg3 | 538.009713 | 0.15273253 | 0.30386069 | 0.50264 | 0.6152174 | 0.8987425 |
| Dhx58 | 378.329959 | -0.0193359 | 0.3313682 | -0.0583516 | 0.95346855 | 0.99149988 |
| Rab5a | 1655.99584 | 0.3832131 | 0.31062387 | 1.23368851 | 0.217319 | 0.65150232 |
| Nkiras2 | 4107.53311 | 0.35384019 | 0.16341605 | 2.16527197 | 0.03036686 | 0.26008374 |
| Ppp2r5c | 6683.5957 | -0.0595504 | 0.15460962 | -0.3851663 | 0.7001142 | 0.92669304 |
| Ift52 | 2089.35095 | 0.12657306 | 0.25571834 | 0.4949706 | 0.62062087 | 0.89896588 |
| Mybl2 | 1637.66829 | -0.1634183 | 0.46101004 | -0.3544789 | 0.72298003 | 0.93433544 |
| B4galt5 | 245.612103 | 0.19754021 | 0.33241715 | 0.59425397 | 0.55234226 | 0.87093412 |
| Gdap1l1 | 22.8144547 | -1.1972187 | 1.02513812 | -1.1678608 | 0.24286291 | 0.67814218 |
| Ddx27 | 3570.18927 | -0.6583293 | 0.23963703 | -2.7471937 | 0.00601076 | 0.09179733 |
| Cyth3 | 557.219019 | 0.18252544 | 0.42964767 | 0.42482584 | 0.67096364 | 0.91711586 |
| Cyth4 | 18044.7626 | 0.16498883 | 0.16471848 | 1.00164127 | 0.31651688 | 0.74186126 |
| Rac3 | 25.9940074 | 0.53681278 | 0.83746984 | 0.64099356 | 0.52152686 | 0.85808179 |
| Rrp7a | 6787.88532 | -0.2325082 | 0.18446405 | -1.2604528 | 0.20750607 | 0.6418276 |

| | | | | | | |
|---|---|---|---|---|---|---|
| Cyb5r3 | 4294.70357 | 0.29517601 | 0.19685126 | 1.4994875 | 0.13374721 | 0.55435014 |
| Ints2 | 1268.15256 | 0.07549434 | 0.2030248 | 0.37184785 | 0.71000613 | 0.93077435 |
| Med13l | 432.704781 | -0.3521138 | 0.59177802 | -0.5950099 | 0.55183686 | 0.87093412 |
| Hist1h2bc | 543.133081 | 0.78523589 | 0.2819845 | 2.78467753 | 0.0053581 | 0.08414071 |
| Mafk | 1275.23658 | -0.1805815 | 0.35083594 | -0.5147177 | 0.60675032 | 0.89307142 |
| Med1 | 2874.98345 | -0.3698862 | 0.20542199 | -1.8006162 | 0.0717634 | 0.4119576 |
| Erbb3 | 92.8037383 | 0.85488338 | 0.70457441 | 1.213333 | 0.22500253 | 0.65870021 |
| Stard3 | 2272.25124 | 0.13605241 | 0.200029 | 0.68016343 | 0.49640098 | 0.84787395 |
| Mfng | 1389.1311 | -0.2357153 | 0.23791522 | -0.9907532 | 0.32180612 | 0.74447686 |
| Vmp1 | 5915.0037 | 0.40650958 | 0.28747579 | 1.41406545 | 0.15734269 | 0.58620126 |
| Uchl5 | 7211.64165 | -0.2712906 | 0.15713328 | -1.7264999 | 0.08425751 | 0.44526833 |
| Glrx2 | 875.869514 | 0.12802443 | 0.33439042 | 0.38285914 | 0.70182421 | 0.92745118 |
| Ro60 | 245.890407 | -0.1046272 | 0.60675389 | -0.1724377 | 0.86309344 | 0.9705375 |
| Stk4 | 1694.85676 | -0.231079 | 0.24625781 | -0.938362 | 0.34805842 | 0.76101142 |
| Pmp22 | 814.16455 | 2.73847499 | 0.63937593 | 4.28304364 | 1.843539974 | 0.00085424 |
| Gdf9 | 49.6527373 | 0.18436254 | 0.8233328 | 0.22392226 | 0.82281779 | 0.9626559 |
| Zcchc10 | 789.112429 | -0.4329138 | 0.3062992 | -1.4133689 | 0.15754728 | 0.58620126 |
| Psmb6 | 14659.8329 | 0.47652982 | 0.35860379 | 1.32884769 | 0.18389822 | 0.61583 |
| Spag7 | 4036.06489 | 0.1149299 | 0.14461211 | 0.79474608 | 0.42676122 | 0.81021713 |
| Pfn1 | 63058.2285 | -0.0407703 | 0.14504617 | -0.2810848 | 0.77864537 | 0.94853305 |
| Tomm34 | 3613.72035 | -0.5619392 | 0.21209464 | -2.6494737 | 0.00806173 | 0.11156005 |
| Ywhab | 18629.2482 | -0.0513726 | 0.1369959 | -0.3749936 | 0.70766526 | 0.92950127 |
| Ksr1 | 12.8438212 | 0.84935421 | 1.56928073 | 0.5412379 | 0.58834362 | 0.88690478 |
| Gpx3 | 7839.76253 | 0.45940307 | 0.13981211 | 3.28586029 | 0.00101671 | 0.02370428 |
| Anxa6 | 4941.66312 | 0.19170591 | 0.23250297 | 0.82453101 | 0.4096379 | 0.80100711 |
| Il12rb2 | 456.150905 | 0.14570034 | 0.31531798 | 0.46207432 | 0.64402802 | 0.90823885 |
| Zkscan6 | 887.225087 | -0.1810078 | 0.3319026 | -0.5453643 | 0.58550302 | 0.8848945 |
| Kpna2 | 27666.8666 | -0.3540501 | 0.21901017 | -1.616592 | 0.10596639 | 0.49777747 |
| Smurf2 | 193.730037 | 0.25787844 | 0.40114761 | 0.64285173 | 0.52032031 | 0.8576517 |
| Cep95 | 278.871842 | -0.0691964 | 0.4796902 | -0.1442523 | 0.88530123 | 0.97751512 |
| Vezf1 | 743.035423 | -0.2208394 | 0.22592494 | -0.9774903 | 0.3283265 | 0.74843736 |
| Srsf1 | 2624.48481 | -0.2690016 | 0.22880098 | -1.1757015 | 0.2397142 | 0.67563574 |
| Abi3 | 1816.58502 | -0.6634978 | 0.21264765 | -3.1201748 | 0.00180744 | 0.03728563 |
| Shroom1 | 26.7809302 | -0.6613037 | 1.35583003 | -0.4877483 | 0.62572816 | 0.90151518 |
| Kif3a | 544.078992 | 0.98340941 | 0.25956156 | 3.78873289 | 0.00015142 | 0.00511683 |
| Sept8 | 392.538311 | 1.30599904 | 0.38940855 | 3.3538017 | 0.00079709 | 0.01965899 |
| Mtmr4 | 1204.23037 | -0.0131932 | 0.28194267 | -0.046794 | 0.96267739 | 0.99401755 |
| Mrm1 | 705.314371 | -0.1752831 | 0.2322675 | -0.7546606 | 0.45045263 | 0.8251553 |
| Kansl1 | 820.326724 | -0.155844 | 0.3541428 | -0.4400599 | 0.65989373 | 0.91280998 |
| Gid4 | 803.008433 | -0.3832872 | 0.21837729 | -1.7551606 | 0.07923184 | 0.4360068 |
| Myo1b | 261.024496 | 0.4495038 | 0.3647332 | 1.23241811 | 0.21779294 | 0.65150232 |
| Dhx40 | 998.508123 | 0.12651863 | 0.18040315 | 0.70131051 | 0.48310926 | 0.84106286 |
| Ypel2 | 38.4151069 | 0.63029354 | 0.69078598 | 0.91242955 | 0.36154264 | 0.76887042 |
| Akap1 | 253.795517 | -0.6519567 | 0.34681932 | -1.8798167 | 0.06013307 | 0.37831609 |
| Nol11 | 4709.16505 | -0.4156016 | 0.21573879 | -1.926411 | 0.05405308 | 0.35989907 |
| Derl2 | 2536.76988 | 0.44826872 | 0.2431877 | 1.8433034 | 0.06528472 | 0.39438421 |

| | | | | | | |
|---|---|---|---|---|---|---|
| C1qbp | 13689.7498 | -0.486402 | 0.1448396 | -3.3582116 | 0.00078449 | 0.01942634 |
| Rpain | 1047.74317 | 1.10742199 | 0.4154236 | 2.66576568 | 0.00768132 | 0.10775872 |
| 6330403K07I | 35.1815521 | 3.28142192 | 1.48423951 | 2.21084393 | 0.02704665 | 0.24328061 |
| Slc13a3 | 13.921986 | -1.1030767 | 1.99020378 | -0.5542531 | 0.57940561 | 0.88286755 |
| Chd3 | 34.9439714 | 1.70839122 | 1.57412459 | 1.08529606 | 0.27779055 | 0.71286172 |
| Kdm6b | 564.505725 | 0.46445821 | 0.4650862 | 0.99864974 | 0.3179644 | 0.74329418 |
| Appbp2 | 3921.93013 | 0.2916147 | 0.27385971 | 1.06483244 | 0.28695176 | 0.72260415 |
| Adora2b | 434.975103 | -0.2594727 | 0.45876152 | -0.5655939 | 0.57166986 | 0.88035336 |
| Ncor1 | 1019.18471 | 0.01226967 | 0.20393739 | 0.0601639 | 0.9520251 | 0.99132814 |
| Trpv2 | 2972.91628 | -0.2101302 | 0.20063475 | -1.0473272 | 0.29494869 | 0.72771732 |
| Cenpv | 14.4318604 | 0.25540681 | 1.34693577 | 0.18962063 | 0.84960642 | 0.96766052 |
| Pcgf2 | 59.3976544 | 0.01021037 | 0.67994957 | 0.01501637 | 0.98801912 | 0.9962454 |
| Cwc25 | 965.197108 | -0.1597018 | 0.36834133 | -0.4335701 | 0.66460066 | 0.91431173 |
| Pip4k2b | 61.404569 | -0.7398002 | 0.8342023 | -0.8868355 | 0.37516749 | 0.77815049 |
| Trim37 | 1366.95647 | 0.01031373 | 0.26636484 | 0.03872031 | 0.96911338 | 0.99443087 |
| Ctdnep1 | 130.280414 | -0.2934276 | 0.38376812 | -0.7645961 | 0.44451208 | 0.82091127 |
| Elp5 | 3485.83417 | 0.17108097 | 0.21886159 | 0.78168568 | 0.43439932 | 0.81542071 |
| Gabarap | 7981.49776 | 0.45937436 | 0.21918085 | 2.09586904 | 0.03609381 | 0.28486166 |
| 2810408A11I | 9.78799224 | 0.52118451 | 2.65708652 | 0.19614887 | 0.84449365 | 0.96765585 |
| Phf23 | 1462.33241 | -0.1112725 | 0.23987045 | -0.4638856 | 0.64272969 | 0.90743513 |
| Acadvl | 2484.64382 | 0.00554668 | 0.18105631 | 0.03063513 | 0.97556053 | 0.99450266 |
| G3bp1 | 22471.5752 | -0.3753604 | 0.1503439 | -2.4966784 | 0.01253626 | 0.1492686 |
| Atox1 | 1919.4943 | 0.24678859 | 0.38224027 | 0.64563735 | 0.51851424 | 0.8570838 |
| Sparc | 462.520476 | 0.99827251 | 0.7549799 | 1.32225045 | 0.18608477 | 0.61790851 |
| Mief2 | 213.469608 | -0.2397431 | 0.41413382 | -0.5789024 | 0.56265501 | 0.8755831 |
| Tbx3 | 26.1798192 | 2.35454241 | 1.98183145 | 1.1880639 | 0.23480824 | 0.670497 |
| Dusp14 | 31.565991 | -0.0401619 | 0.72568942 | -0.0553431 | 0.95586514 | 0.99215259 |
| Tada2a | 567.372849 | -0.2795429 | 0.26411521 | -1.0584127 | 0.28986731 | 0.72395811 |
| Ikzf1 | 2372.86045 | -0.024338 | 0.37090185 | -0.0656186 | 0.94768152 | 0.99019648 |
| Pnpo | 2445.53366 | -0.3750803 | 0.18196442 | -2.0612837 | 0.03927599 | 0.29991457 |
| Cog1 | 1171.33129 | -0.4671872 | 0.24326065 | -1.9205209 | 0.05479213 | 0.36084578 |
| Cbx1 | 1695.0031 | 0.0930575 | 0.26927729 | 0.34558245 | 0.72965654 | 0.93649408 |
| Cdk5rap3 | 2808.62095 | -0.2464969 | 0.21766808 | -1.1324439 | 0.25744785 | 0.69274795 |
| Copz2 | 250.249077 | 1.36343253 | 0.40818636 | 3.3402207 | 0.00083712 | 0.02031839 |
| Slc25a39 | 4771.73679 | -0.0598465 | 0.16219552 | -0.3689773 | 0.71214463 | 0.93235217 |
| Sp2 | 589.120223 | -0.1978933 | 0.28768532 | -0.6878811 | 0.49152769 | 0.84497726 |
| Aatf | 2575.01056 | -0.4091428 | 0.16372587 | -2.4989505 | 0.01245617 | 0.14871281 |
| Dync1h1 | 1282.84597 | 0.77887586 | 0.31741568 | 2.453804 | 0.0141354 | 0.16169868 |
| Pex12 | 411.082993 | -0.1474052 | 0.33508654 | -0.4399019 | 0.66000816 | 0.91281852 |
| Ndel1 | 4025.15423 | 0.27735 | 0.23041996 | 1.20367173 | 0.22871648 | 0.66258779 |
| Slc25a35 | 48.1325661 | -0.5799486 | 0.87229911 | -0.6648507 | 0.50614601 | 0.85085669 |
| Zbtb4 | 27.9147979 | 2.13353389 | 1.73636604 | 1.2287351 | 0.21917114 | 0.65257977 |
| Mpdu1 | 3637.93132 | 0.56874424 | 0.29005325 | 1.96082702 | 0.04989921 | 0.34225988 |
| Fxr2 | 178.430959 | -0.5295423 | 0.51685715 | -1.0245429 | 0.30557894 | 0.73530116 |
| Atp5g3 | 5477.17773 | 0.1909292 | 0.29687024 | 0.64314024 | 0.52013311 | 0.8576517 |
| Cd68 | 10562.8181 | 0.43894427 | 0.16952937 | 2.58919312 | 0.00962011 | 0.12626914 |

| | | | | | | |
|---|---|---|---|---|---|---|
| Acsl1 | 1630.62388 | -0.5274811 | 0.23314264 | -2.2624822 | 0.02366763 | 0.22185905 |
| Smyd4 | 395.854421 | 0.31131221 | 0.41484875 | 0.7504234 | 0.45299974 | 0.82610332 |
| Lsp1 | 13010.6599 | -0.6947734 | 0.18063412 | -3.8463022 | 0.00011991 | 0.00420317 |
| Zfyve27 | 710.946965 | -0.1983524 | 0.26233937 | -0.756091 | 0.44959465 | 0.82484234 |
| Avpi1 | 339.070741 | 0.24829097 | 0.51997894 | 0.47750198 | 0.63300471 | 0.90430301 |
| Rad51d | 792.713962 | -0.3337544 | 0.21183805 | -1.5755165 | 0.11513727 | 0.51884494 |
| Pank3 | 354.96037 | 0.25189962 | 0.42878693 | 0.58747037 | 0.55688784 | 0.87276537 |
| Rars | 11495.3475 | -0.2283718 | 0.18582952 | -1.2289316 | 0.21909745 | 0.65257977 |
| Mrpl58 | 1987.34712 | 0.37919004 | 0.35886552 | 1.0566355 | 0.29067797 | 0.72461526 |
| Fdxr | 649.395599 | -0.1261555 | 0.26379839 | -0.4782268 | 0.6324888 | 0.90423307 |
| Mrpl45 | 3454.39733 | 0.13414911 | 0.16893195 | 0.7941015 | 0.42713635 | 0.81021713 |
| Irf1 | 2377.15551 | 0.40908904 | 0.27302391 | 1.49836343 | 0.13403885 | 0.5545138 |
| Slc22a5 | 479.562372 | 0.17491887 | 0.24993825 | 0.69984834 | 0.48402202 | 0.84128398 |
| P4ha2 | 159.950728 | 2.09698877 | 0.51875753 | 4.04232932 | 5.292283364 | 0.00210881 |
| Arrb1 | 626.129079 | 0.47141761 | 0.21248432 | 2.21859956 | 0.02651398 | 0.23919285 |
| Cxcl16 | 157.777896 | 1.55637232 | 0.66001798 | 2.35807566 | 0.01836995 | 0.19092573 |
| Pelp1 | 433.017111 | -0.0013464 | 0.22862837 | -0.0058891 | 0.99530124 | 0.99884675 |
| Med11 | 1229.0409 | 0.28876845 | 0.34534853 | 0.83616529 | 0.40306192 | 0.79806139 |
| Ccl6 | 4771.75201 | 1.55610552 | 0.32985493 | 4.71754512 | 2.387074682 | 0.00015209 |
| Ccl4 | 11106.1831 | 0.75294121 | 0.53044131 | 1.41946186 | 0.15576441 | 0.58481688 |
| Natd1 | 487.24364 | 1.09635076 | 0.36839895 | 2.97598773 | 0.00292047 | 0.05300601 |
| Map2k3 | 6683.25946 | -0.2161661 | 0.19235099 | -1.1238107 | 0.26109336 | 0.69691361 |
| Ywhah | 19861.6827 | 0.13009599 | 0.23152238 | 0.56191538 | 0.57417367 | 0.88059206 |
| Sart3 | 2479.6482 | -0.2247979 | 0.30590417 | -0.7348637 | 0.46242248 | 0.83108016 |
| E2f2 | 223.250186 | 0.24945913 | 0.57951482 | 0.43046204 | 0.66685957 | 0.91565144 |
| Slfn3 | 12.4405694 | 0.2059476 | 1.59533092 | 0.12909397 | 0.89728329 | 0.97984623 |
| Nars2 | 851.444073 | 0.60611845 | 0.19991283 | 3.03191369 | 0.00243009 | 0.0460372 |
| Slc35b4 | 2439.72359 | 0.02576249 | 0.22339545 | 0.11532236 | 0.90818961 | 0.9817854 |
| Dalrd3 | 612.889018 | -0.4901004 | 0.269595 | -1.8179135 | 0.06907736 | 0.40548143 |
| Fis1 | 3671.18976 | 0.01989208 | 0.34499749 | 0.05765863 | 0.95402054 | 0.99171765 |
| Plod1 | 5610.54932 | 1.04424142 | 0.24675651 | 4.23186982 | 2.317565403 | 0.0010272 |
| Rab3d | 1144.07964 | -0.3109111 | 0.23801499 | -1.3062668 | 0.19146185 | 0.62259956 |
| Mfsd3 | 341.386215 | -0.0559758 | 0.32654709 | -0.1714171 | 0.86389582 | 0.9705375 |
| Slc25a22 | 857.503874 | -0.5809465 | 0.26796528 | -2.1679917 | 0.03015932 | 0.25945074 |
| Atp6ap1 | 10688.1945 | 0.17700891 | 0.13404631 | 1.32050565 | 0.18666625 | 0.61862792 |
| Dnase1l1 | 663.04254 | 1.24648607 | 0.4032629 | 3.09100108 | 0.00199483 | 0.0400045 |
| Ccl9 | 11820.1293 | 1.89409794 | 0.40139425 | 4.71879691 | 2.372435146 | 0.00015209 |
| Scrn1 | 21.4834089 | 1.10515623 | 1.52690571 | 0.72378813 | 0.46919581 | 0.83381643 |
| BC005537 | 2334.89344 | -0.3775653 | 0.29003605 | -1.3017876 | 0.19298902 | 0.624687 |
| Isyna1 | 10778.3751 | -0.1758757 | 0.18668767 | -0.9420854 | 0.34614891 | 0.7602916 |
| Hars2 | 1193.74494 | 0.23896384 | 0.28456187 | 0.83976056 | 0.40104265 | 0.79698916 |
| Tmem160 | 1005.0134 | 0.05962772 | 0.29134111 | 0.20466634 | 0.83783282 | 0.96683245 |
| Rab5c | 1109.78382 | 0.2526855 | 0.28182572 | 0.89660199 | 0.36993133 | 0.77529039 |
| Mdh2 | 20566.6943 | -0.2703542 | 0.16823043 | -1.6070472 | 0.10804402 | 0.50301538 |
| H13 | 3729.68894 | -0.3814121 | 0.1920348 | -1.9861613 | 0.04701542 | 0.3347601 |
| Rnf145 | 1265.91781 | 0.50798759 | 0.31703257 | 1.60231987 | 0.10908489 | 0.50489425 |

| | | | | | | |
|---|---|---|---|---|---|---|
| Scn1b | 6.2423805 | 6.12202848 | 2.28987493 | 2.6735209 | 0.00750596 | 0.10639676 |
| Atp6v1e1 | 6305.09691 | 0.13231899 | 0.17519139 | 0.75528248 | 0.45007951 | 0.82484234 |
| Chtf18 | 1399.78713 | -0.5682332 | 0.29235253 | -1.9436574 | 0.05193678 | 0.3516008 |
| Rps6kl1 | 25.6244156 | -0.4491295 | 1.08158466 | -0.4152513 | 0.67795793 | 0.91965617 |
| Ppp1r12c | 498.791547 | -0.0459935 | 0.24512279 | -0.1876346 | 0.85116307 | 0.9679302 |
| Ahr | 44.6526188 | -1.347443 | 0.91788954 | -1.4679795 | 0.14210979 | 0.5673724 |
| Map1s | 1607.17382 | -0.1216489 | 0.24491603 | -0.4966963 | 0.61940325 | 0.89896588 |
| Tmem129 | 1161.94887 | 0.1084635 | 0.23599322 | 0.45960432 | 0.64580026 | 0.90847224 |
| Nop9 | 5010.6164 | -0.3562801 | 0.20493162 | -1.7385318 | 0.08211716 | 0.44123582 |
| Atp6v0a1 | 1878.61659 | 0.76867133 | 0.31685554 | 2.42593621 | 0.01526895 | 0.17073593 |
| Psmc3ip | 777.308725 | 0.03707343 | 0.34740886 | 0.10671412 | 0.91501578 | 0.98347559 |
| Zfp687 | 355.694663 | -0.0260681 | 0.47733319 | -0.054612 | 0.95644762 | 0.99215259 |
| Gdpd2 | 35.2724475 | -2.0509318 | 1.16599041 | -1.7589612 | 0.07858411 | 0.43353356 |
| D8Ertd738e | 4066.70249 | 0.09813676 | 0.28808573 | 0.34065123 | 0.73336616 | 0.93728136 |
| Calm3 | 2522.75003 | 0.22875347 | 0.23979181 | 0.953967 | 0.34010035 | 0.75756302 |
| Cops3 | 7827.78934 | 0.20297057 | 0.3686931 | 0.55051361 | 0.58196714 | 0.88321857 |
| Fkbp8 | 4558.87562 | -0.091158 | 0.1649332 | -0.5526966 | 0.58047116 | 0.88286755 |
| Ddx39b | 9703.04724 | -0.3967333 | 0.1389597 | -2.8550245 | 0.00430335 | 0.07142862 |
| Gipc1 | 4371.15278 | 0.47782936 | 0.16324118 | 2.92713736 | 0.00342098 | 0.05967533 |
| Tlcd1 | 149.613113 | -0.1628723 | 0.34009 | -0.4789095 | 0.63200298 | 0.90405469 |
| Plscr3 | 1002.46971 | -0.0879853 | 0.22027514 | -0.3994338 | 0.68957358 | 0.92470267 |
| Arhgef25 | 35.6381588 | 0.85323436 | 0.65313009 | 1.30637736 | 0.19142426 | 0.62259956 |
| Xab2 | 4505.01563 | -0.3487677 | 0.20012148 | -1.7427802 | 0.08137202 | 0.43986918 |
| Cdc37 | 8893.45617 | -0.2135882 | 0.13407748 | -1.593021 | 0.11115549 | 0.50888686 |
| Rab4a | 620.598204 | 0.52439308 | 0.20824669 | 2.51813406 | 0.01179784 | 0.14331975 |
| Trip10 | 887.627974 | -0.0831289 | 0.3070509 | -0.2707331 | 0.7865963 | 0.95205625 |
| Cops6 | 9802.69971 | 0.01065816 | 0.2484622 | 0.0428965 | 0.96578404 | 0.99443087 |
| Ubb | 71903.1562 | 0.04585676 | 0.1779634 | 0.25767525 | 0.79665755 | 0.95380086 |
| Ap4m1 | 444.159122 | -0.0401158 | 0.36281907 | -0.1105669 | 0.91195977 | 0.98292561 |
| Gyg | 4630.73554 | -0.3993315 | 0.16172526 | -2.4691966 | 0.01354168 | 0.1574243 |
| Rcn3 | 4535.48682 | 1.77742005 | 0.20929585 | 8.49238075 | 2.024464877 | 1.179298992 |
| Arid3a | 1609.09422 | -0.588404 | 0.19938557 | -2.9510861 | 0.00316659 | 0.05605911 |
| Pdk4 | 37.1235877 | 5.75260236 | 1.1292343 | 5.09425045 | 3.501236772 | 2.893961448 |
| Ubxn6 | 1331.24121 | 0.14555101 | 0.24101725 | 0.60390288 | 0.54590822 | 0.86990153 |
| Mydgf | 6786.76967 | 0.19398972 | 0.237592 | 0.81648252 | 0.41422422 | 0.80395688 |
| Sema6a | 46.5245753 | 3.73860302 | 1.27742976 | 2.92666035 | 0.00342623 | 0.05967533 |
| Ccdc12 | 2433.53512 | -0.1001715 | 0.24197154 | -0.4139804 | 0.67888845 | 0.92012773 |
| Fmc1 | 764.194082 | 0.60030414 | 0.5571162 | 1.07752052 | 0.28124779 | 0.71634017 |
| Akt3 | 80.3136676 | -1.4508699 | 1.19038293 | -1.2188262 | 0.22291016 | 0.65613364 |
| Mrpl24 | 2400.09425 | 0.10993023 | 0.35673348 | 0.30815786 | 0.75796221 | 0.94129085 |
| Gle1 | 3729.79506 | -0.077746 | 0.20746993 | -0.3747337 | 0.70785856 | 0.92950127 |
| L3hypdh | 86.9803451 | 0.77318577 | 0.61223679 | 1.26288682 | 0.20662986 | 0.64143381 |
| Lyst | 134.157281 | -1.2940968 | 0.53250017 | -2.430228 | 0.01508933 | 0.16981393 |
| Slc35e1 | 502.913637 | -0.6606252 | 0.53442758 | -1.2361361 | 0.21640797 | 0.64982959 |
| Calr3 | 82.7084701 | 0.35884052 | 0.65807501 | 0.54528817 | 0.58555534 | 0.8848945 |
| Tmc4 | 14.5131411 | 0.327704 | 1.43579153 | 0.22823926 | 0.81946024 | 0.96120981 |

| | | | | | | |
|---|---|---|---|---|---|---|
| Polr2i | 970.556439 | 0.20315182 | 0.36459685 | 0.55719576 | 0.57739368 | 0.88147472 |
| Krt10 | 69.7466986 | 0.80837738 | 0.62317513 | 1.29719132 | 0.19456537 | 0.62651174 |
| Rmnd1 | 1904.52485 | 0.14825169 | 0.21276795 | 0.6967764 | 0.48594274 | 0.84183502 |
| Esr1 | 19.6770359 | -1.1974176 | 1.43764009 | -0.832905 | 0.40489832 | 0.79917613 |
| Fbxo5 | 1757.57013 | -0.2095658 | 0.28638127 | -0.731772 | 0.46430772 | 0.83257898 |
| Mtrf1l | 490.9987 | 0.11572895 | 0.2904613 | 0.39843157 | 0.69031209 | 0.92470267 |
| Hdac2 | 1544.84716 | -0.0352786 | 0.4650214 | -0.0758644 | 0.93952698 | 0.98913801 |
| Frk | 19.8405145 | 1.66070196 | 1.51391824 | 1.09695617 | 0.27266055 | 0.71005689 |
| Rwdd1 | 3713.62448 | 0.0474556 | 0.33416659 | 0.1420118 | 0.88707069 | 0.97827644 |
| Stxbp5 | 262.916726 | 0.68793665 | 0.4939129 | 1.39282991 | 0.16367123 | 0.59518139 |
| Hint3 | 262.916895 | 0.57426027 | 0.5107793 | 1.12428258 | 0.26089317 | 0.69691361 |
| Trmt11 | 1370.42561 | -0.1481457 | 0.247708 | -0.5980659 | 0.54979599 | 0.87093412 |
| Katna1 | 2563.58264 | 0.0095309 | 0.20835775 | 0.04574298 | 0.96351511 | 0.99401755 |
| Pcmt1 | 5871.45859 | 0.01900654 | 0.20346478 | 0.09341439 | 0.92557436 | 0.98580325 |
| Lrp11 | 145.040756 | 1.11425568 | 0.55873498 | 1.99424721 | 0.04612504 | 0.3319104 |
| 1700021F05I | 600.446169 | -0.0775528 | 0.28578863 | -0.2713641 | 0.78611101 | 0.95184068 |
| Sec63 | 482.842003 | -0.2635127 | 0.27699798 | -0.9513163 | 0.34144382 | 0.75846782 |
| Snx3 | 14485.9762 | 0.86941223 | 0.16497361 | 5.27000797 | 1.364178317 | 1.265524296 |
| Aig1 | 234.948086 | -0.06206 | 0.51227006 | -0.121147 | 0.90357457 | 0.98060928 |
| Adat2 | 212.70581 | -0.2510859 | 0.48129195 | -0.5216915 | 0.60188512 | 0.89103677 |
| Pex3 | 861.345177 | 0.38212448 | 0.33613089 | 1.13683238 | 0.25560836 | 0.69025543 |
| Fuca2 | 5164.81888 | 0.09865775 | 0.15584617 | 0.63304571 | 0.52670379 | 0.86115578 |
| Cep57l1 | 903.313735 | -0.1885734 | 0.27842161 | -0.6772946 | 0.49821907 | 0.84860011 |
| Ltv1 | 3739.14971 | -0.4315327 | 0.20087899 | -2.1482219 | 0.03169613 | 0.26539272 |
| Plagl1 | 27.9780663 | 3.61064211 | 1.41778417 | 2.54667966 | 0.01087532 | 0.13520102 |
| Cd164 | 20324.6053 | -0.2133266 | 0.1909624 | -1.1171133 | 0.2639459 | 0.70024945 |
| Utrn | 114.202141 | 0.05293882 | 0.6342876 | 0.08346186 | 0.9334843 | 0.98756396 |
| Smpd2 | 1023.21102 | 0.06847707 | 0.2048614 | 0.33426048 | 0.73818301 | 0.93800224 |
| Mical1 | 951.693566 | -0.6792278 | 0.23008221 | -2.9521093 | 0.00315611 | 0.05595465 |
| Zbtb24 | 489.110373 | -0.4598484 | 0.36089197 | -1.2741996 | 0.20259269 | 0.63759103 |
| Wasf1 | 25.0590078 | 1.38742473 | 1.02634417 | 1.35181236 | 0.17643535 | 0.60764153 |
| Rab32 | 16356.0812 | 0.36781839 | 0.23639056 | 1.55597746 | 0.11971345 | 0.52718991 |
| Amd-ps4 | 8.89724531 | -0.4017965 | 1.398074 | -0.2873929 | 0.7738115 | 0.94669828 |
| Gtf3c6 | 2308.68519 | -0.0307531 | 0.17090659 | -0.1799408 | 0.85719908 | 0.96936093 |
| Slc16a10 | 2934.16707 | 0.09200829 | 0.26057547 | 0.35309649 | 0.7240161 | 0.93466537 |
| Rev3l | 172.830676 | 0.12848109 | 0.77280822 | 0.16625223 | 0.86795845 | 0.97169724 |
| Traf3ip2 | 70.4947325 | 0.49063512 | 0.82963821 | 0.59138443 | 0.55426287 | 0.87165957 |
| Fyn | 334.889674 | 0.6996829 | 0.32801034 | 2.1331123 | 0.03291551 | 0.26976292 |
| Tube1 | 754.702656 | -0.2615763 | 0.21973077 | -1.19044 | 0.23387351 | 0.66968409 |
| Prep | 7979.31099 | -0.4297341 | 0.2003903 | -2.1444856 | 0.031994 | 0.26606568 |
| Tnfaip3 | 4373.52716 | 0.89137935 | 0.58055451 | 1.53539303 | 0.12468731 | 0.53626132 |
| Perp | 227.347349 | -4.4800377 | 0.61817301 | -7.2472231 | 4.254030041 | 1.369461828 |
| Hebp2 | 171.106342 | 0.42146426 | 0.50055036 | 0.84200171 | 0.399787 | 0.79612475 |
| Reps1 | 255.361154 | -0.17104 | 0.41713522 | -0.410035 | 0.68178026 | 0.92061965 |
| Asf1a | 2436.89464 | -0.2753427 | 0.26849278 | -1.0255125 | 0.30512142 | 0.73518175 |
| Gopc | 321.04563 | -0.5620462 | 0.48006148 | -1.1707796 | 0.2416874 | 0.67708722 |

| | | | | | | |
|---|---|---|---|---|---|---|
| Qrsl1 | 611.48516 | -0.2321984 | 0.23267623 | -0.9979461 | 0.31830547 | 0.74336577 |
| Rtn4ip1 | 933.916909 | -0.4169791 | 0.22305403 | -1.8694087 | 0.06156598 | 0.38288591 |
| Crybg1 | 9.16587736 | -4.2663677 | 1.82072393 | -2.343226 | 0.0191178 | 0.19537847 |
| Vta1 | 3484.73246 | 0.26948129 | 0.23243519 | 1.15938249 | 0.24630031 | 0.6814256 |
| Smpdl3a | 567.689068 | 1.29134363 | 0.39403601 | 3.2772224 | 0.00104834 | 0.02424256 |
| Reep3 | 1228.72141 | 0.44458555 | 0.32230151 | 1.37940884 | 0.16776874 | 0.60218592 |
| Pkib | 590.810351 | 0.78344125 | 0.32688663 | 2.39667572 | 0.01654456 | 0.17972015 |
| Serinc1 | 6567.82144 | 0.36184674 | 0.15057016 | 2.40317699 | 0.01625332 | 0.17782865 |
| Hsf2 | 514.965986 | -0.194304 | 0.30163657 | -0.6441658 | 0.51946791 | 0.8576517 |
| Echdc1 | 1587.19535 | 0.00514342 | 0.21465671 | 0.02396113 | 0.98088361 | 0.99522729 |
| Ptprk | 56.0438568 | 2.75999505 | 0.64365056 | 4.28803331 | 1.802620682 | 0.00084488 |
| Dcbld1 | 363.949241 | 0.06920582 | 0.40385811 | 0.17136173 | 0.86393935 | 0.9705375 |
| Ccdc59 | 826.931049 | 0.0871825 | 0.25538033 | 0.34138298 | 0.73281528 | 0.93728136 |
| Ppp1r12a | 1332.2467 | -0.3786911 | 0.33103248 | -1.1439695 | 0.25263627 | 0.68616774 |
| P4ha1 | 6489.27766 | -0.0898855 | 0.17867891 | -0.5030561 | 0.61492486 | 0.89851598 |
| Sept10 | 513.69979 | 0.47626706 | 0.28022831 | 1.69956796 | 0.08921222 | 0.4566019 |
| Lims1 | 3649.08092 | -0.4186719 | 0.18070831 | -2.316838 | 0.02051256 | 0.20351188 |
| Zwint | 7692.697 | 0.09876074 | 0.15909212 | 0.62077707 | 0.53474631 | 0.8652846 |
| Ube2d1 | 579.910031 | 0.03856903 | 0.23670625 | 0.16294047 | 0.8705653 | 0.97265735 |
| Cdk1 | 10365.2127 | 0.12609138 | 0.23724005 | 0.5314928 | 0.59507733 | 0.88830085 |
| Atp2b1 | 4816.2873 | 0.0280306 | 0.36460271 | 0.07687984 | 0.93871913 | 0.9890914 |
| Rhobtb1 | 471.867342 | 1.05211907 | 0.52407072 | 2.00758987 | 0.04468689 | 0.32480971 |
| Arid5b | 193.635199 | 0.27955879 | 0.48178055 | 0.58026167 | 0.56173817 | 0.87540592 |
| Actr6 | 708.552145 | 0.18839772 | 0.40796259 | 0.46180147 | 0.6442237 | 0.90823885 |
| Uhrf1bp1l | 685.743458 | -0.1034831 | 0.33547577 | -0.3084667 | 0.75772724 | 0.94129085 |
| Poc1b | 1106.97181 | -0.2871111 | 0.28851416 | -0.9951368 | 0.31966973 | 0.7440831 |
| Dusp6 | 10771.787 | 0.30500328 | 0.19153019 | 1.59245534 | 0.11128243 | 0.50888686 |
| Tmpo | 1695.30216 | -0.2307399 | 0.17594803 | -1.3114095 | 0.18971945 | 0.62019167 |
| Kitl | 107.100398 | 1.66693981 | 0.56080805 | 2.97238923 | 0.00295492 | 0.05339367 |
| Psen1 | 7253.55742 | -0.3613892 | 0.25710441 | -1.4056127 | 0.15983918 | 0.58912704 |
| Sgk1 | 919.015238 | -0.0193342 | 0.43179216 | -0.0447767 | 0.96428528 | 0.99406401 |
| Cep290 | 82.8640083 | 0.11304305 | 0.4628484 | 0.24423343 | 0.80705004 | 0.95778548 |
| Ikbip | 468.3005 | 0.52071644 | 0.31009211 | 1.67923149 | 0.09310694 | 0.46660269 |
| Hbs1l | 3682.62001 | -0.0929697 | 0.19007443 | -0.4891227 | 0.6247548 | 0.90093428 |
| Epb41l2 | 3413.74805 | 0.23552309 | 0.2315224 | 1.01727994 | 0.30902028 | 0.73867239 |
| Apaf1 | 653.706093 | 0.40710372 | 0.30727378 | 1.32488925 | 0.18520789 | 0.61717463 |
| Myb | 117.488415 | -2.1468633 | 0.91510593 | -2.3460271 | 0.01897473 | 0.19472974 |
| Med23 | 593.743817 | -0.0868506 | 0.24965521 | -0.3478821 | 0.72792874 | 0.93576253 |
| Ahi1 | 79.3539981 | -0.1866432 | 0.61913139 | -0.3014597 | 0.76306395 | 0.94298237 |
| Arg1 | 97.8273389 | -0.2703218 | 0.67330878 | -0.4014827 | 0.68806475 | 0.92470267 |
| Nedd1 | 2500.00586 | 0.25935177 | 0.23796552 | 1.08987123 | 0.27576987 | 0.7118544 |
| Mtfr2 | 69.5929865 | -0.7741691 | 0.67575856 | -1.1456297 | 0.25194841 | 0.68505999 |
| Map7 | 214.009195 | 0.22652589 | 0.52026096 | 0.43540821 | 0.66326615 | 0.91395288 |
| Stx7 | 7304.97695 | 0.38246247 | 0.17350387 | 2.20434542 | 0.02750006 | 0.24591246 |
| Pex7 | 1920.9842 | 0.25257491 | 0.36288526 | 0.69601864 | 0.48641715 | 0.84183502 |
| Ifngr1 | 15526.6125 | -0.2520836 | 0.14356069 | -1.7559374 | 0.0790991 | 0.43547225 |

| | | | | | | |
|---|---|---|---|---|---|---|
| Cdk17 | 307.199004 | 0.16552511 | 0.28233715 | 0.58626756 | 0.55769573 | 0.87288227 |
| Hal | 12.5747917 | -2.6086531 | 1.84282537 | -1.4155726 | 0.15690068 | 0.58620126 |
| Snrpf | 3185.02439 | -0.4811635 | 0.18930458 | -2.5417423 | 0.01103015 | 0.13649787 |
| Usp44 | 15.1882668 | -2.6487713 | 1.81224037 | -1.4616005 | 0.14385073 | 0.56966439 |
| Fgd6 | 102.025876 | 1.99740525 | 0.54180086 | 3.68660406 | 0.00022727 | 0.00718385 |
| Ndufa12 | 3971.84055 | 0.121436 | 0.28300117 | 0.42910069 | 0.66784995 | 0.91595755 |
| Tmcc3 | 328.018859 | 1.09017077 | 0.40869244 | 2.66746009 | 0.0076427 | 0.10754172 |
| Cep83 | 915.983571 | -0.2872005 | 0.25562487 | -1.1235234 | 0.26121525 | 0.69693483 |
| Socs2 | 30.6642824 | 1.10779482 | 0.72197729 | 1.53439013 | 0.1249337 | 0.53626132 |
| Nudt4 | 1392.30723 | -0.23381 | 0.18741197 | -1.2475723 | 0.21218772 | 0.64580271 |
| Nuak1 | 52.2870841 | -0.0968586 | 0.61311918 | -0.1579768 | 0.87447506 | 0.97364644 |
| Tcp11l2 | 597.293577 | -0.0927595 | 0.42601088 | -0.2177398 | 0.82763185 | 0.96423051 |
| Cry1 | 1454.30826 | -0.6254457 | 0.21178209 | -2.953251 | 0.00314446 | 0.05582903 |
| Btbd11 | 15.6150048 | -0.1360082 | 1.75778375 | -0.0773748 | 0.93832539 | 0.98906962 |
| Timp3 | 82.2596081 | 0.9373929 | 1.45116814 | 0.64595747 | 0.5183069 | 0.8570838 |
| Hsp90b1 | 56045.213 | 0.11233546 | 0.16138104 | 0.69608834 | 0.48637351 | 0.84183502 |
| Igf1 | 1009.22432 | 1.86748089 | 0.30755996 | 6.07192471 | 1.263861879 | 2.208688910 |
| Washc3 | 1672.76017 | 0.43401083 | 0.32813319 | 1.32266667 | 0.18594625 | 0.61790851 |
| Dram1 | 311.024501 | -0.3169573 | 0.28161773 | -1.1254876 | 0.26038246 | 0.6965359 |
| Sirt1 | 200.735689 | -0.8738406 | 0.58833211 | -1.4852846 | 0.13746846 | 0.55999057 |
| Herc4 | 1602.72442 | 0.27514828 | 0.23816257 | 1.15529607 | 0.2479692 | 0.68235696 |
| Hnrnph3 | 798.389241 | -0.3744214 | 0.23036369 | -1.6253492 | 0.10408817 | 0.49391412 |
| Rufy2 | 61.1173458 | 0.06558171 | 0.87596378 | 0.07486806 | 0.94031969 | 0.98934006 |
| Pbld2 | 8.94506294 | -0.7665759 | 2.12392733 | -0.3609238 | 0.71815642 | 0.93286094 |
| Ccar1 | 4721.5501 | -0.1910652 | 0.25675889 | -0.7441424 | 0.45679032 | 0.82915295 |
| Ddx21 | 3565.38088 | -0.6047295 | 0.27100108 | -2.2314652 | 0.02565033 | 0.23347396 |
| Ddx50 | 2096.70069 | -0.1612481 | 0.23246688 | -0.6936392 | 0.48790852 | 0.84313956 |
| Srgn | 4267.89475 | -1.1197647 | 0.23230665 | -4.8202008 | 1.434138073 | 9.801011760 |
| Vps26a | 11341.8673 | 0.408304 | 0.17761331 | 2.29883672 | 0.02151421 | 0.21054669 |
| Supv3l1 | 1552.75013 | -0.2361526 | 0.21282358 | -1.1096168 | 0.26716418 | 0.70405417 |
| Aifm2 | 244.901293 | -0.3530919 | 0.40548787 | -0.8707828 | 0.38387277 | 0.78704643 |
| H2afy2 | 86.5005837 | 1.10932628 | 0.48616158 | 2.28180572 | 0.02250081 | 0.21507833 |
| Tysnd1 | 1857.42841 | -0.0674217 | 0.17494167 | -0.3853955 | 0.69994439 | 0.92669304 |
| Sar1a | 7982.92107 | 0.2402546 | 0.20416682 | 1.17675636 | 0.23929277 | 0.67541957 |
| Ppa1 | 5245.98293 | -0.493009 | 0.23916614 | -2.0613664 | 0.0392681 | 0.29991457 |
| Eif4ebp2 | 813.792585 | -0.3491853 | 0.48060403 | -0.726555 | 0.46749861 | 0.83381643 |
| Pald1 | 1520.21695 | 0.55967169 | 0.22803588 | 2.45431411 | 0.01411536 | 0.16167905 |
| Sgpl1 | 7243.28096 | 0.60154015 | 0.20760085 | 2.8975803 | 0.00376053 | 0.06410023 |
| Pcbd1 | 120.959792 | 0.19883247 | 0.50868863 | 0.39087264 | 0.69589138 | 0.92623267 |
| Slc29a3 | 1355.34682 | -0.086797 | 0.25957784 | -0.3343776 | 0.73809461 | 0.93800224 |
| Vsir | 7521.71315 | -0.4966456 | 0.203767 | -2.4373209 | 0.01479654 | 0.16744321 |
| Slc16a7 | 1410.64048 | 0.03607007 | 0.20211511 | 0.17846299 | 0.85835938 | 0.96947659 |
| Anapc16 | 2118.97478 | 0.54909163 | 0.35462673 | 1.54836502 | 0.12153444 | 0.52946253 |
| Ddit4 | 25.3590176 | 0.62875166 | 0.90777022 | 0.69263306 | 0.48853986 | 0.84363468 |
| Dnajb12 | 571.682273 | -0.4063761 | 0.38734585 | -1.0491299 | 0.29411835 | 0.72700541 |
| Micu1 | 2133.92505 | 0.18115756 | 0.21404928 | 0.84633575 | 0.39736547 | 0.79518596 |

| | | | | | | |
|---|---|---|---|---|---|---|
| Cand1 | 4447.27543 | -0.1558403 | 0.21811317 | -0.714493 | 0.47492237 | 0.83578199 |
| Tbk1 | 3033.67933 | 0.14655972 | 0.16488738 | 0.88884738 | 0.37408511 | 0.77748705 |
| Pno1 | 4301.52337 | -0.1780582 | 0.22404463 | -0.7947442 | 0.42676232 | 0.81021713 |
| Plek | 17839.1904 | 0.09768653 | 0.30913014 | 0.31600454 | 0.75199906 | 0.94018119 |
| Usp15 | 4119.99563 | -0.3660134 | 0.28866552 | -1.26795 | 0.20481581 | 0.63901202 |
| Elane | 233.843428 | -7.4327671 | 0.70843216 | -10.491854 | 9.416191001 | 1.151882645 |
| Vps54 | 375.133109 | -0.3762041 | 0.54821949 | -0.686229 | 0.49256869 | 0.84581594 |
| Tbc1d15 | 3466.06334 | 0.17555076 | 0.18936695 | 0.92704009 | 0.35390572 | 0.76401657 |
| Pcsk4 | 51.4249763 | -2.0866101 | 0.94517274 | -2.2076495 | 0.02726871 | 0.24474748 |
| Rab21 | 234.936536 | -0.4350742 | 0.49233956 | -0.8836873 | 0.37686502 | 0.77940657 |
| 2310011J03F | 2942.32452 | 0.36466504 | 0.31498293 | 1.15772954 | 0.24697443 | 0.68185412 |
| Peli1 | 2690.92721 | 0.13982855 | 0.29508454 | 0.47385929 | 0.63560025 | 0.90531997 |
| Thap2 | 320.726027 | -0.1556835 | 0.43574068 | -0.3572847 | 0.72087867 | 0.93398714 |
| Slc1a4 | 143.09165 | -0.1190531 | 0.61463803 | -0.1936963 | 0.84641372 | 0.96765585 |
| Dock2 | 832.051526 | -0.1453391 | 0.25963492 | -0.5597827 | 0.57562764 | 0.88080372 |
| Rab1a | 19491.5674 | 0.34935219 | 0.13395981 | 2.607888 | 0.00911028 | 0.12206572 |
| Gamt | 589.21188 | 0.48197524 | 0.26861475 | 1.79429926 | 0.07276542 | 0.41489174 |
| Actr2 | 14087.0082 | -0.1895431 | 0.17727032 | -1.0692322 | 0.28496504 | 0.72045503 |
| Ndufs7 | 5213.56128 | -0.0334171 | 0.23754674 | -0.1406758 | 0.88812605 | 0.97832097 |
| Pwwp3a | 1388.52247 | 0.02049542 | 0.22313296 | 0.09185297 | 0.92681486 | 0.98630274 |
| Meis1 | 784.317336 | -0.5199657 | 0.28285441 | -1.8382805 | 0.06602108 | 0.39562181 |
| Uqcr11 | 6534.428 | 0.32464049 | 0.42805472 | 0.75840885 | 0.44820626 | 0.82382295 |
| Cnot2 | 3455.17283 | -0.2485285 | 0.26860339 | -0.9252619 | 0.35482968 | 0.76419569 |
| Tcf3 | 1258.4812 | 0.21521479 | 0.46307916 | 0.46474731 | 0.64211243 | 0.90705802 |
| Frs2 | 150.661364 | -0.1066949 | 0.50213272 | -0.2124834 | 0.83172994 | 0.96514441 |
| Yeats4 | 1623.42629 | -0.3326175 | 0.22713264 | -1.4644195 | 0.14307937 | 0.56846902 |
| Rab36 | 17.6710556 | 3.67979181 | 1.71545545 | 2.14508154 | 0.03194633 | 0.26603092 |
| Grb10 | 80.0437041 | 1.00570757 | 0.8654839 | 1.16201765 | 0.24522829 | 0.68039867 |
| Snrpd3 | 2136.09478 | -0.0631083 | 0.21579763 | -0.2924419 | 0.76994875 | 0.9442389 |
| Cpm | 15.3273796 | 0.99164894 | 2.21748691 | 0.44719495 | 0.65473431 | 0.91108261 |
| Mdm2 | 3905.76705 | 0.45377016 | 0.3342044 | 1.35776238 | 0.1745391 | 0.60656845 |
| E2f7 | 232.74418 | -0.05193 | 0.40979191 | -0.1267229 | 0.89915972 | 0.98010294 |
| Csrp2 | 415.86593 | 0.68304576 | 0.47160274 | 1.44834984 | 0.14751924 | 0.5745313 |
| Osbpl8 | 1096.49016 | 0.37759458 | 0.27246535 | 1.38584443 | 0.16579441 | 0.5992759 |
| Mknk2 | 1506.19001 | -0.1428841 | 0.27448519 | -0.5205532 | 0.60267804 | 0.89159034 |
| Zpbp | 14.1680413 | -0.6049114 | 1.18470293 | -0.5106018 | 0.60962995 | 0.89508835 |
| Cabin1 | 1204.54181 | -0.1370064 | 0.27347524 | -0.5009829 | 0.61638316 | 0.89878056 |
| Ap3d1 | 4774.25517 | 0.02459769 | 0.32737203 | 0.07513682 | 0.94010585 | 0.98934006 |
| Phlda1 | 400.987849 | 0.46936283 | 0.40388873 | 1.16210925 | 0.24519109 | 0.68039867 |
| Sf3a2 | 359.112718 | -0.5730035 | 0.35070809 | -1.6338473 | 0.10229092 | 0.4886081 |
| Mdm1 | 1516.43084 | 0.08006821 | 0.20399288 | 0.39250494 | 0.69468516 | 0.92609254 |
| Timm13 | 5669.23363 | 0.34756335 | 0.37352617 | 0.93049263 | 0.35211608 | 0.76348958 |
| Vps13d | 238.734176 | -0.2409589 | 0.79034771 | -0.3048771 | 0.76045979 | 0.94215089 |
| Llph | 2498.06673 | -0.1881267 | 0.22930914 | -0.8204065 | 0.41198442 | 0.80245328 |
| Tmbim4 | 2174.54612 | -0.0085073 | 0.14956999 | -0.0568787 | 0.95464181 | 0.99215259 |
| Irak3 | 199.056332 | -0.9645658 | 0.47599208 | -2.0264324 | 0.0427205 | 0.31653532 |

| | | | | | | |
|---|---|---|---|---|---|---|
| Helb | 432.686547 | 0.11155114 | 0.28867219 | 0.38642843 | 0.6991794 | 0.92669304 |
| Prmt2 | 555.044968 | 0.96217585 | 0.26683798 | 3.60584302 | 0.00031114 | 0.00917154 |
| Dip2a | 100.940812 | -0.2311181 | 0.76646326 | -0.3015384 | 0.76300394 | 0.94298237 |
| Hmg20b | 501.56509 | 0.19879742 | 0.21866296 | 0.90914995 | 0.36327098 | 0.77000799 |
| Fzr1 | 4644.22823 | -0.1102072 | 0.2310201 | -0.4770459 | 0.63332943 | 0.90434445 |
| Ncln | 6145.83702 | -0.3947735 | 0.21544025 | -1.8324038 | 0.06689128 | 0.39799658 |
| Hcfc2 | 867.366847 | 0.12293214 | 0.2991917 | 0.41088086 | 0.6811599 | 0.92061965 |
| Nfyb | 1476.1621 | 0.03262246 | 0.16899829 | 0.19303425 | 0.84693214 | 0.96765585 |
| Txnrd1 | 5701.5305 | -0.0698021 | 0.31409537 | -0.2222321 | 0.82413318 | 0.96290555 |
| Ppm1m | 1901.343 | -0.6163546 | 0.22487237 | -2.7409087 | 0.00612695 | 0.09327527 |
| D10Wsu102e | 1565.35358 | -0.4485914 | 0.22065334 | -2.0330143 | 0.04205108 | 0.31328312 |
| Aldh1l2 | 38.2925363 | -0.4186292 | 1.21546522 | -0.3444189 | 0.73053128 | 0.93653187 |
| Wdr82 | 998.926771 | -0.1784216 | 0.30723958 | -0.5807248 | 0.56142594 | 0.8752292 |
| Glyctk | 58.8595255 | -2.1202302 | 1.19322469 | -1.776891 | 0.07558618 | 0.42337259 |
| Pofut2 | 2544.14271 | 0.25583458 | 0.20642402 | 1.23936441 | 0.21521058 | 0.64854028 |
| Slc36a1 | 216.121341 | 0.07766245 | 0.49354401 | 0.15735668 | 0.87496375 | 0.97366476 |
| Appl2 | 977.27586 | 0.87818813 | 0.18795406 | 4.67235523 | 2.977653813 | 0.00018585 |
| Sumo3 | 9238.01654 | 0.06680052 | 0.22159185 | 0.30145746 | 0.76306568 | 0.94298237 |
| Hint1 | 9828.79754 | 0.46771313 | 0.42639276 | 1.09690682 | 0.27268213 | 0.71005689 |
| Lyrm7 | 122.672619 | -0.2840428 | 0.46290168 | -0.6136137 | 0.53947061 | 0.86719392 |
| Fbxw11 | 2750.61882 | 0.25066608 | 0.30100488 | 0.83276416 | 0.40497775 | 0.79917613 |
| Stk10 | 1394.96762 | -0.2781481 | 0.33074392 | -0.8409772 | 0.40036071 | 0.79668215 |
| Papolg | 506.558029 | 0.07414713 | 0.25000888 | 0.29657798 | 0.76678872 | 0.94381438 |
| Rel | 1363.48492 | 0.02805241 | 0.51793173 | 0.05416238 | 0.95680579 | 0.99216795 |
| Pfkl | 15513.8892 | -0.3544125 | 0.21055034 | -1.6832672 | 0.09232337 | 0.46597694 |
| Pus10 | 1845.37595 | -0.2030307 | 0.19468706 | -1.0428568 | 0.29701461 | 0.72954201 |
| Rhbdf1 | 12.4522917 | -0.13998 | 1.59461059 | -0.0877832 | 0.93004901 | 0.9870122 |
| Pex13 | 1665.99283 | 0.53104198 | 0.28873258 | 1.83921736 | 0.06588322 | 0.39558985 |
| 1810043G02 | 452.315336 | 0.29917313 | 0.24285423 | 1.23190411 | 0.21798491 | 0.65150232 |
| Mpg | 592.675698 | 0.22843441 | 0.32071769 | 0.71226011 | 0.47630372 | 0.83644936 |
| Ahsa2 | 1483.85531 | -0.0895289 | 0.20588191 | -0.4348554 | 0.66366741 | 0.91395288 |
| Nprl3 | 571.930107 | -0.1917609 | 0.22402419 | -0.8559829 | 0.39200723 | 0.79031213 |
| Xpo1 | 4056.33722 | -0.1863022 | 0.25144103 | -0.740938 | 0.45873103 | 0.82986925 |
| Cpeb4 | 139.493862 | 0.54305719 | 0.60653116 | 0.89534919 | 0.37060045 | 0.7762831 |
| Stc2 | 209.33241 | -0.2114048 | 0.40640115 | -0.5201875 | 0.60293292 | 0.89168472 |
| Asb3 | 1003.33438 | 0.18810506 | 0.20605323 | 0.91289545 | 0.36129753 | 0.76887042 |
| Cdc34 | 2560.55704 | 0.01574178 | 0.30271746 | 0.05200157 | 0.95852744 | 0.99249408 |
| Tpgs1 | 1002.62132 | 0.04126262 | 0.22589878 | 0.18265978 | 0.85506498 | 0.96887828 |
| Chac2 | 152.760125 | 1.72442704 | 0.71596593 | 2.40853226 | 0.01601681 | 0.17615771 |
| Erlec1 | 207.290837 | 0.01747336 | 0.47855342 | 0.03651286 | 0.97087342 | 0.99443087 |
| Sptbn1 | 1069.60451 | -0.0769402 | 0.29341544 | -0.2622226 | 0.79314985 | 0.95310033 |
| Wdpcp | 495.684459 | 0.3459173 | 0.22327025 | 1.54932108 | 0.12130455 | 0.5291774 |
| Mdh1 | 10519.0985 | 0.23731219 | 0.18668618 | 1.27118238 | 0.20366377 | 0.63809363 |
| Prss57 | 84.0694938 | -9.7934309 | 1.43564189 | -6.8216391 | 9.000754366 | 2.247065880 |
| Ccng1 | 4360.58424 | 0.26122981 | 0.2036411 | 1.28279514 | 0.19956386 | 0.6323909 |
| Nudcd2 | 4524.87321 | 0.18427301 | 0.31381913 | 0.58719494 | 0.55707279 | 0.87276537 |

| | | | | | | |
|---|---|---|---|---|---|---|
| Polrmt | 1115.41742 | -0.0422707 | 0.18243133 | -0.2317075 | 0.81676523 | 0.96079676 |
| Hmmr | 3148.02823 | -0.3564739 | 0.21330857 | -1.6711652 | 0.09468904 | 0.47133886 |
| Slc22a4 | 27.4598076 | 0.58645619 | 1.15129643 | 0.50938765 | 0.61048053 | 0.89508835 |
| Cyfip2 | 170.355255 | -1.0632582 | 0.44991201 | -2.363258 | 0.01811505 | 0.19054293 |
| Mgat1 | 2598.21318 | 0.12562177 | 0.17119431 | 0.73379639 | 0.46307284 | 0.83158692 |
| Ppp2ca | 806.991183 | -0.2018913 | 0.30660261 | -0.6584789 | 0.51023046 | 0.85335831 |
| Flt4 | 30.8300428 | 2.52453955 | 1.95691901 | 1.29005827 | 0.19703043 | 0.62932233 |
| Hnrnpab | 2672.92254 | -0.2234407 | 0.40985636 | -0.5451684 | 0.58563769 | 0.8848945 |
| Phykpl | 724.348838 | 0.0495374 | 0.257991 | 0.19201214 | 0.8477327 | 0.96765585 |
| Hspa4 | 13624.2449 | -0.2226138 | 0.21013984 | -1.0593602 | 0.28943577 | 0.72395811 |
| Cnot6 | 845.649985 | -0.4086288 | 0.27818278 | -1.4689221 | 0.14185393 | 0.56696251 |
| Gfpt2 | 34.9526676 | 0.71958797 | 1.61195877 | 0.44640594 | 0.65530404 | 0.91136134 |
| Zfp354a | 59.0139612 | 0.32274452 | 1.3470236 | 0.23959827 | 0.81064171 | 0.95905029 |
| Mapk9 | 1308.61731 | 0.12414858 | 0.36934789 | 0.33612911 | 0.73677351 | 0.93800224 |
| Canx | 38691.0419 | -0.0228474 | 0.19672665 | -0.116138 | 0.90754317 | 0.98176897 |
| Rack1 | 34802.3019 | 0.17121225 | 0.37696663 | 0.45418411 | 0.64969632 | 0.90973357 |
| Rufy1 | 995.989615 | 0.40909843 | 0.41215765 | 0.99257754 | 0.32091587 | 0.74413737 |
| Rnf130 | 1621.37443 | 0.3487117 | 0.20461377 | 1.70424355 | 0.0883356 | 0.45403754 |
| Ltc4s | 410.669665 | 2.39080625 | 0.35986737 | 6.64357612 | 3.061622534 | 6.809605176 |
| Rad50 | 1510.6271 | -0.0336282 | 0.22307523 | -0.1507481 | 0.88017442 | 0.97549733 |
| Mrnip | 458.395566 | 0.15818294 | 0.22976896 | 0.68844347 | 0.49117355 | 0.84491231 |
| Clk4 | 1650.70066 | -0.3173152 | 0.28914709 | -1.0974179 | 0.27245876 | 0.71005689 |
| Sar1b | 6222.54384 | 0.51789256 | 0.41010014 | 1.26284416 | 0.20664519 | 0.64143381 |
| Pdlim4 | 1019.7225 | 1.15507209 | 0.23917249 | 4.82945212 | 1.369092259 | 9.462206557 |
| Cdkl3 | 70.4409141 | 1.13883719 | 0.79136601 | 1.43907771 | 0.15012851 | 0.5778861 |
| Ube2b | 2809.99243 | 0.1682405 | 0.28922177 | 0.58170068 | 0.56076831 | 0.87469256 |
| Cdkn2aipnl | 3669.19557 | 0.24568696 | 0.25869657 | 0.94971092 | 0.34225916 | 0.75889838 |
| Kremen1 | 234.815723 | -0.0297844 | 0.38729375 | -0.0769039 | 0.93869997 | 0.9890914 |
| Med7 | 1671.34509 | 0.34765167 | 0.24593846 | 1.41357178 | 0.15748767 | 0.58620126 |
| Havcr2 | 243.202501 | -0.2684786 | 0.52173288 | -0.5145902 | 0.6068394 | 0.89307142 |
| Tnip1 | 388.83571 | -0.1877696 | 0.30016317 | -0.6255584 | 0.53160462 | 0.8643494 |
| Vdac1 | 7278.99292 | -0.1693721 | 0.15605536 | -1.0853334 | 0.27777401 | 0.71286172 |
| Upp1 | 34.4235947 | 1.87123859 | 1.69851554 | 1.10169059 | 0.27059621 | 0.70756376 |
| Slu7 | 2464.14973 | -0.3365643 | 0.30389265 | -1.1075104 | 0.26807333 | 0.70459677 |
| Ascc2 | 1818.76768 | -0.0332804 | 0.26694852 | -0.1246696 | 0.90078515 | 0.98043702 |
| Hus1 | 638.44529 | -0.1884794 | 0.25829866 | -0.7296958 | 0.46557615 | 0.83339036 |
| Pttg1 | 994.488125 | 0.20624654 | 0.19815482 | 1.04083535 | 0.29795197 | 0.73013751 |
| Hormad2 | 67.7154224 | 0.05673545 | 0.49262878 | 0.11516877 | 0.90831135 | 0.98183024 |
| Zfp607a | 164.394443 | 0.25372545 | 0.40590715 | 0.62508249 | 0.53191692 | 0.8643494 |
| Tns3 | 454.074154 | 0.37527982 | 0.32624238 | 1.15030986 | 0.25001627 | 0.6842534 |
| Btg2 | 10850.5786 | 0.26750361 | 0.33557463 | 0.79715088 | 0.42536341 | 0.81013087 |
| Castor1 | 243.743863 | -0.1862678 | 0.46736965 | -0.398545 | 0.69022848 | 0.92470267 |
| Igfbp3 | 39.8015737 | 1.94343454 | 1.99071555 | 0.97624924 | 0.32894097 | 0.7486136 |
| Pes1 | 8479.81014 | -0.2349897 | 0.17445598 | -1.3469857 | 0.17798485 | 0.60920221 |
| Tcn2 | 3707.98632 | 0.77711208 | 0.19262545 | 4.03431678 | 5.476139928 | 0.00216096 |
| Myo1g | 6835.60192 | -0.88692 | 0.21592669 | -4.1075052 | 3.999557125 | 0.00168133 |

| | | | | | | |
|---|---|---|---|---|---|---|
| Smtn | 499.765525 | 0.1817567 | 0.33782966 | 0.53801287 | 0.59056816 | 0.88748446 |
| Arf5 | 1506.19098 | 0.30609767 | 0.18894428 | 1.62004202 | 0.10522325 | 0.49640504 |
| 2310033P09I | 460.82076 | -0.387066 | 0.28388056 | -1.363482 | 0.17273064 | 0.60492813 |
| Guk1 | 1797.89913 | 0.10076821 | 0.35426743 | 0.28444109 | 0.77607238 | 0.94776024 |
| Rnf185 | 2070.81593 | 0.44279531 | 0.19574947 | 2.2620512 | 0.02369424 | 0.22193847 |
| Limk2 | 846.666141 | -0.7247882 | 0.32835249 | -2.207348 | 0.02728975 | 0.24474748 |
| Patz1 | 919.109533 | -0.3929972 | 0.21903324 | -1.7942353 | 0.07277562 | 0.41489174 |
| Eif4enif1 | 577.707773 | -0.5174418 | 0.49373702 | -1.048011 | 0.29463354 | 0.72739699 |
| Trim11 | 473.887191 | -0.2213693 | 0.41710238 | -0.5307313 | 0.59560501 | 0.88830374 |
| Ogdh | 8970.50816 | -0.2342735 | 0.18717994 | -1.251595 | 0.21071748 | 0.64401792 |
| Drg1 | 11344.1806 | 0.21800462 | 0.33725135 | 0.64641587 | 0.51801006 | 0.8570838 |
| Rtn4 | 803.721784 | 0.16322673 | 0.27499843 | 0.59355514 | 0.55280969 | 0.8711362 |
| Mtif2 | 1107.04964 | 0.10397071 | 0.2233511 | 0.46550348 | 0.64157095 | 0.90700632 |
| Rps27a | 27251.1028 | 0.37793664 | 0.42417072 | 0.89100125 | 0.3729285 | 0.77748705 |
| Cfap36 | 689.531318 | 0.16420858 | 0.36794658 | 0.44628375 | 0.6553923 | 0.91138047 |
| Ppp4r3b | 3500.07999 | -0.1665685 | 0.18245564 | -0.912926 | 0.36128145 | 0.76887042 |
| Pnpt1 | 3505.44137 | -0.156479 | 0.19448308 | -0.8045895 | 0.42105662 | 0.80708017 |
| Pold2 | 3671.43375 | -0.4444475 | 0.20123014 | -2.2086529 | 0.02719879 | 0.24446937 |
| Zkscan17 | 631.647846 | -0.1638921 | 0.41206808 | -0.3977307 | 0.69082869 | 0.92470267 |
| Polm | 283.567478 | 0.39135136 | 0.54354259 | 0.72000127 | 0.47152421 | 0.83393588 |
| Dbnl | 8116.13412 | 0.14579978 | 0.18858285 | 0.77313381 | 0.43944319 | 0.81963526 |
| Mrps24 | 2018.54873 | 0.3642436 | 0.38539566 | 0.945116 | 0.34459966 | 0.75972469 |
| Ccdc117 | 2748.06809 | -0.0980336 | 0.21102595 | -0.464557 | 0.64224874 | 0.90705802 |
| Dynll2 | 160.076628 | 0.59917083 | 0.42362134 | 1.4144019 | 0.15724393 | 0.58620126 |
| Xbp1 | 11119.5408 | 0.17752714 | 0.13907839 | 1.2764538 | 0.20179515 | 0.63606288 |
| Supt4a | 4009.33293 | 0.58219888 | 0.44681612 | 1.30299435 | 0.19257669 | 0.6243813 |
| 2810021J22F | 152.759395 | 1.64100517 | 1.15948721 | 1.41528527 | 0.15698487 | 0.58620126 |
| Ska2 | 2010.67233 | -0.0696597 | 0.15259726 | -0.4564935 | 0.64803514 | 0.90973357 |
| Prr11 | 348.506952 | -0.5205161 | 0.2949426 | -1.7648047 | 0.07759659 | 0.43049394 |
| Smg8 | 1041.87972 | 0.10151434 | 0.25182973 | 0.40310707 | 0.68686946 | 0.92426981 |
| Rnf187 | 7934.17181 | -0.1034257 | 0.14745535 | -0.7014035 | 0.48305124 | 0.84106286 |
| Tubd1 | 1060.80334 | 0.22185571 | 0.39196806 | 0.56600456 | 0.57139068 | 0.88035336 |
| Mrpl22 | 1950.58616 | 0.00766912 | 0.33943792 | 0.02259359 | 0.98197445 | 0.99523682 |
| Cnot8 | 3040.13352 | -0.1151088 | 0.19777978 | -0.5820051 | 0.56056323 | 0.87469256 |
| Rps6kb1 | 2419.72693 | -0.2384029 | 0.19654128 | -1.2129913 | 0.22513314 | 0.65870021 |
| Sap30l | 73.3669985 | 0.52642955 | 0.59377097 | 0.88658688 | 0.37530136 | 0.77815049 |
| Galnt10 | 528.438741 | 0.04573498 | 0.45322259 | 0.10091064 | 0.9196214 | 0.9839023 |
| Rnft1 | 2500.59768 | 0.32536139 | 0.26578692 | 1.22414369 | 0.22089803 | 0.65452328 |
| Mfap3 | 2694.62226 | 0.32460864 | 0.17571681 | 1.84733965 | 0.06469793 | 0.39236898 |
| Fam114a2 | 2500.33117 | 0.08054233 | 0.16039169 | 0.50216022 | 0.61555483 | 0.89876301 |
| Ppm1d | 833.772265 | -0.1833323 | 0.31008564 | -0.5912311 | 0.55436556 | 0.87165957 |
| Znhit3 | 2314.59437 | 0.7474013 | 0.51199396 | 1.45978538 | 0.14434907 | 0.57027792 |
| Myo19 | 295.556777 | 0.12518314 | 0.41044208 | 0.30499587 | 0.7603693 | 0.94215089 |
| Prpsap2 | 1574.87415 | 0.44459028 | 0.20300587 | 2.19003655 | 0.02852159 | 0.25141958 |
| Ggnbp2 | 3065.3746 | 0.24898648 | 0.17471776 | 1.42507827 | 0.15413458 | 0.58303286 |
| Acaca | 539.161417 | -0.3864556 | 0.2931961 | -1.3180789 | 0.18747725 | 0.61935434 |

| | | | | | | |
|---|---|---|---|---|---|---|
| Shmt1 | 4718.04422 | -0.347451 | 0.18468867 | -1.8812795 | 0.05993391 | 0.37827102 |
| Llgl1 | 431.932281 | -0.1482106 | 0.255997 | -0.5789545 | 0.56261985 | 0.8755831 |
| Drg2 | 4942.3297 | -0.2201204 | 0.16628455 | -1.3237573 | 0.18558368 | 0.61775377 |
| Srebf1 | 1936.24475 | -0.4931691 | 0.32507786 | -1.5170799 | 0.12924651 | 0.54538551 |
| Tom1l1 | 455.114297 | -0.0562594 | 0.28946961 | -0.1943534 | 0.84589915 | 0.96765585 |
| Cox11 | 403.807368 | 0.17111762 | 0.27949949 | 0.61222875 | 0.54038642 | 0.86741192 |
| Stxbp4 | 272.404789 | -0.07525 | 0.28790952 | -0.261367 | 0.79380953 | 0.95310033 |
| Bzw2 | 5385.01267 | -0.3886627 | 0.18452632 | -2.1062727 | 0.03518067 | 0.28128442 |
| Elac2 | 2994.79487 | -0.4953276 | 0.16621798 | -2.9799883 | 0.00288259 | 0.05255257 |
| Pctp | 2148.49852 | 0.32608876 | 0.25356119 | 1.28603576 | 0.19843058 | 0.63122167 |
| Twistnb | 1774.53287 | -0.0366581 | 0.31689985 | -0.1156773 | 0.90790832 | 0.9817854 |
| Atxn7l1 | 899.821937 | -0.0241251 | 0.28549638 | -0.0845022 | 0.93265717 | 0.98738166 |
| Sypl | 4010.94839 | -0.0333484 | 0.18909492 | -0.1763577 | 0.86001291 | 0.96954641 |
| Pdia6 | 41714.3026 | 0.46635654 | 0.18692934 | 2.49482796 | 0.01260182 | 0.14937799 |
| Nampt | 3223.87321 | 0.16574622 | 0.23114969 | 0.71705148 | 0.47334233 | 0.83483892 |
| Pik3cg | 2367.24952 | 0.22274749 | 0.30326095 | 0.73450767 | 0.46263939 | 0.83117457 |
| Nbas | 222.653714 | 0.20976879 | 0.37217159 | 0.5636346 | 0.57300283 | 0.88035336 |
| Tspan13 | 1603.88121 | 0.53436053 | 0.35057643 | 1.52423404 | 0.12745026 | 0.54145382 |
| Rock2 | 336.041891 | 0.11679485 | 0.38928982 | 0.30002031 | 0.76416166 | 0.94300309 |
| Matn3 | 47.3843888 | 1.02845417 | 1.2788651 | 0.80419285 | 0.42128561 | 0.80727887 |
| Laptm4a | 1929.34774 | 0.4636399 | 0.24129242 | 1.9214856 | 0.05467052 | 0.36033644 |
| Fam49a | 613.318494 | 0.67806456 | 0.26096039 | 2.59834289 | 0.00936749 | 0.12401786 |
| Snx13 | 1184.13865 | 0.52730498 | 0.19150863 | 2.75342678 | 0.0058975 | 0.0904061 |
| Sdc1 | 199.697176 | -0.2788566 | 0.41048041 | -0.6793419 | 0.49692122 | 0.84814011 |
| Lpin1 | 108.390706 | 0.72867417 | 0.44639704 | 1.63234545 | 0.10260673 | 0.48942862 |
| Pum2 | 2405.37349 | -0.0482071 | 0.28566391 | -0.1687546 | 0.86598969 | 0.97055903 |
| Nrcam | 10.3342743 | -1.6618967 | 2.08384353 | -0.7975151 | 0.42515192 | 0.80988459 |
| Trib2 | 14.7003558 | 0.77074072 | 1.69359451 | 0.45509165 | 0.6490433 | 0.90973357 |
| Arsg | 33.1444588 | 0.04553223 | 1.52659451 | 0.02982602 | 0.97620581 | 0.994746 |
| Hs1bp3 | 1056.43594 | 0.70647381 | 0.3256674 | 2.16931083 | 0.03005909 | 0.25895274 |
| Fam84a | 151.821379 | 0.05927109 | 0.53003222 | 0.11182545 | 0.91096181 | 0.98288717 |
| Smc6 | 2599.72585 | -0.3999278 | 0.21478297 | -1.862009 | 0.06260182 | 0.38618662 |
| Amz2 | 975.179652 | -0.2401538 | 0.2904915 | -0.8267154 | 0.40839837 | 0.80088767 |
| Gna13 | 1363.86745 | 0.27261239 | 0.20784315 | 1.31162554 | 0.18964651 | 0.62019167 |
| Prkar1a | 10891.5323 | 0.1257983 | 0.14486558 | 0.8683795 | 0.38518662 | 0.78782609 |
| Rdh14 | 580.698962 | 0.39362935 | 0.24123365 | 1.63173486 | 0.10273535 | 0.48958378 |
| Map2k6 | 443.158101 | 1.00207233 | 0.26555548 | 3.77349516 | 0.00016098 | 0.00540996 |
| Trappc12 | 1825.64835 | -0.1879946 | 0.19488196 | -0.9646589 | 0.33471568 | 0.75351065 |
| Adi1 | 2289.54511 | -0.1179701 | 0.17288805 | -0.6823494 | 0.49501805 | 0.84681245 |
| Rnaseh1 | 775.092169 | -0.1500878 | 0.24142579 | -0.6216725 | 0.53415723 | 0.86520872 |
| Ubxn2a | 786.106101 | -0.1389896 | 0.2845742 | -0.4884125 | 0.62525771 | 0.9011746 |
| Fkbp1b | 578.743098 | -0.25446 | 0.3180778 | -0.799993 | 0.42371486 | 0.80917688 |
| Cmpk2 | 146.291822 | -0.922015 | 0.42381545 | -2.1755105 | 0.02959188 | 0.25669202 |
| Pfn4 | 35.2839576 | -0.777431 | 1.04504264 | -0.7439228 | 0.4569232 | 0.82915295 |
| Itsn2 | 1682.80632 | 0.02926526 | 0.2680475 | 0.10917938 | 0.91306021 | 0.98292561 |
| Rsad2 | 23.8395237 | 4.54337752 | 1.52767343 | 2.97405024 | 0.00293897 | 0.05326284 |

| | | | | | | |
|---|---|---|---|---|---|---|
| Rnf144a | 21.1624722 | 0.60155015 | 2.00892025 | 0.29943954 | 0.7646047 | 0.94307414 |
| Id2 | 2794.47958 | 0.91184254 | 0.32337254 | 2.81978965 | 0.00480551 | 0.07714679 |
| Mboat2 | 120.00581 | 0.80339397 | 0.75253274 | 1.06758674 | 0.28570697 | 0.721488 |
| Ncoa1 | 42.9262836 | -0.0348991 | 1.69066971 | -0.0206422 | 0.98353111 | 0.99564261 |
| Dus4l | 93.403673 | -0.6226001 | 0.45194547 | -1.3776001 | 0.16832678 | 0.60296969 |
| Rrm2 | 16404.6104 | -0.2759588 | 0.18274945 | -1.5100389 | 0.1310335 | 0.54951416 |
| Bcap29 | 3777.92935 | -0.1645156 | 0.34727367 | -0.4737348 | 0.63568903 | 0.90531997 |
| Cenpo | 1292.76758 | -0.1316787 | 0.20131408 | -0.6540957 | 0.51305013 | 0.85482732 |
| Klf11 | 717.960574 | 0.27524155 | 0.49740112 | 0.55335933 | 0.58001738 | 0.88286755 |
| Adcy3 | 36.03117 | -0.6554127 | 1.13173668 | -0.5791212 | 0.56250743 | 0.8755831 |
| Grhl1 | 19.5628602 | -1.2781281 | 1.94173207 | -0.6582412 | 0.51038314 | 0.85335831 |
| Dnajc27 | 120.609079 | -0.5774239 | 0.59795171 | -0.9656697 | 0.33420946 | 0.75320272 |
| Efr3b | 24.0118488 | 2.45484707 | 2.30959863 | 1.06288904 | 0.28783227 | 0.72300867 |
| Cbll1 | 1024.26592 | -0.2906814 | 0.28600715 | -1.0163432 | 0.30946598 | 0.73896103 |
| Pomc | 20.804535 | 0.4238842 | 0.80934269 | 0.52373884 | 0.60046019 | 0.8903551 |
| Dnmt3a | 3177.86054 | 0.03105183 | 0.22650196 | 0.13709301 | 0.89095728 | 0.97889088 |
| Dld | 6058.98841 | -0.0514778 | 0.14089007 | -0.3653758 | 0.71483093 | 0.93235217 |
| Kif3c | 416.257565 | -0.5953768 | 0.36179295 | -1.6456285 | 0.09984026 | 0.48298772 |
| Sh3yl1 | 77.5793791 | -0.2411769 | 0.59009228 | -0.4087104 | 0.68275217 | 0.92125604 |
| Rab10 | 1956.82203 | 0.3865045 | 0.43463541 | 0.88926143 | 0.3738626 | 0.77748705 |
| Pxdn | 47.7089388 | 2.30437826 | 1.43958982 | 1.60071865 | 0.10943925 | 0.50553519 |
| Ddx52 | 2687.04191 | -0.3927532 | 0.21953763 | -1.7890015 | 0.07361457 | 0.41763833 |
| Hnf1b | 13.6078538 | 2.30002718 | 2.00103058 | 1.1494213 | 0.2503823 | 0.6842534 |
| Taf15 | 32.1742156 | 0.18765121 | 0.7230714 | 0.25951961 | 0.79523435 | 0.9537323 |
| Mmp28 | 11.5540027 | -0.3916818 | 1.94782795 | -0.2010865 | 0.84063097 | 0.96765585 |
| Cdc27 | 1669.29386 | -0.1131596 | 0.18870842 | -0.599653 | 0.54873749 | 0.87082325 |
| Itgb3 | 50.9219213 | -1.7715217 | 1.03517263 | -1.7113297 | 0.08702026 | 0.45109601 |
| Mettl2 | 1418.52669 | 0.04936892 | 0.2751933 | 0.17939724 | 0.8576258 | 0.9694296 |
| Nle1 | 513.073901 | -0.5386448 | 0.37376385 | -1.4411367 | 0.14954608 | 0.57685486 |
| Tlk2 | 396.418502 | -0.4372768 | 0.2926475 | -1.4942099 | 0.13512077 | 0.55654288 |
| Rffl | 1760.477 | -0.294129 | 0.28442214 | -1.0341284 | 0.30107613 | 0.73265651 |
| Lig3 | 1473.54631 | -0.0162735 | 0.25005899 | -0.0650788 | 0.94811129 | 0.99019648 |
| Map3k3 | 555.003929 | -0.3833967 | 0.31111554 | -1.2323288 | 0.21782627 | 0.65150232 |
| Ddx42 | 3134.78483 | -0.1508786 | 0.22372587 | -0.6743907 | 0.50006297 | 0.84914912 |
| Ftsj3 | 6858.30494 | -0.6641633 | 0.22664066 | -2.9304685 | 0.00338451 | 0.0592314 |
| Rnf135 | 1423.15864 | 0.06164063 | 0.21980887 | 0.2804283 | 0.77914892 | 0.9488468 |
| Psmc5 | 12194.8388 | -0.0622837 | 0.20928565 | -0.2976016 | 0.76600724 | 0.94347227 |
| Adap2 | 1276.41176 | 0.3407384 | 0.24828868 | 1.37234771 | 0.16995522 | 0.60430173 |
| Tcam1 | 10.0546823 | -1.7285939 | 2.2195849 | -0.7787915 | 0.43610253 | 0.81725842 |
| Ern1 | 115.214014 | -0.4493513 | 0.45103811 | -0.9962602 | 0.31912374 | 0.74398524 |
| Nf1 | 188.536187 | 0.47825122 | 0.38830804 | 1.23162844 | 0.21808791 | 0.65150232 |
| Pecam1 | 1125.11848 | 0.21151561 | 0.30231943 | 0.6996428 | 0.4841504 | 0.84128398 |
| Polg2 | 331.193223 | -0.2075644 | 0.34929215 | -0.5942429 | 0.55234968 | 0.87093412 |
| Ddx5 | 57140.2397 | -0.1438733 | 0.18933044 | -0.759906 | 0.44731078 | 0.82297186 |
| Psmd12 | 9097.63763 | 0.03647452 | 0.1938062 | 0.18820101 | 0.85071908 | 0.9679302 |
| Helz | 824.19532 | 0.08771083 | 0.27951257 | 0.31379925 | 0.75367352 | 0.94018119 |

| | | | | | | |
|---|---|---|---|---|---|---|
| Cep112 | 61.1047156 | 0.71447352 | 0.56994629 | 1.25358043 | 0.20999455 | 0.64350283 |
| Slc9a3r1 | 1771.43027 | -0.3112651 | 0.19191332 | -1.6219047 | 0.10482376 | 0.49510003 |
| Nt5c | 2538.77131 | 0.07332877 | 0.39440772 | 0.18592123 | 0.85250653 | 0.96798445 |
| Jpt1 | 13571.447 | -0.075552 | 0.1876851 | -0.4025464 | 0.68728198 | 0.92427909 |
| Sumo2 | 12220.0615 | -0.2760827 | 0.15281804 | -1.8066104 | 0.07082304 | 0.41080052 |
| Nup85 | 5880.49156 | -0.1801327 | 0.16010357 | -1.1251012 | 0.26054616 | 0.69666911 |
| Gga3 | 642.752048 | -0.2534308 | 0.27117972 | -0.9345491 | 0.35002071 | 0.76216093 |
| Cluh | 2275.43991 | -0.5942055 | 0.22960434 | -2.5879542 | 0.00965478 | 0.12658836 |
| Mif4gd | 638.606404 | 0.0609995 | 0.31076715 | 0.19628683 | 0.84438567 | 0.96765585 |
| Slc25a19 | 1058.88848 | -0.0892545 | 0.21197982 | -0.4210521 | 0.67371706 | 0.91822108 |
| Pafah1b1 | 1518.04656 | -0.0931712 | 0.34889054 | -0.26705 | 0.78943063 | 0.95255415 |
| Tmem94 | 501.227058 | -0.7206673 | 0.39991215 | -1.802064 | 0.07153534 | 0.41122734 |
| Recql5 | 478.293921 | -0.5103033 | 0.23770156 | -2.1468234 | 0.03180734 | 0.26554509 |
| Sap30bp | 2355.94844 | 0.35408966 | 0.28070369 | 1.26143572 | 0.20715191 | 0.6418276 |
| Galk1 | 6443.77541 | 0.04131935 | 0.29252243 | 0.1412519 | 0.88767095 | 0.97832097 |
| Unk | 1041.30071 | 0.45586904 | 0.19331677 | 2.35814539 | 0.0183665 | 0.19092573 |
| Trim47 | 681.46894 | 2.16142284 | 0.42103357 | 5.13361159 | 2.842343765 | 2.448619104 |
| Aspa | 59.4146206 | 3.68539106 | 1.0430637 | 3.5332368 | 0.0004105 | 0.01149131 |
| Mrpl38 | 1680.7106 | -0.191915 | 0.19914849 | -0.9636778 | 0.33520751 | 0.75359284 |
| Fbf1 | 274.251508 | -0.2761223 | 0.60759394 | -0.454452 | 0.6495035 | 0.90973357 |
| Acox1 | 1759.2091 | 0.15771501 | 0.34574944 | 0.45615405 | 0.64827921 | 0.90973357 |
| Ten1 | 1707.98408 | 0.21487186 | 0.29516479 | 0.72797253 | 0.4666304 | 0.83345004 |
| Srp68 | 3805.66455 | -0.3112651 | 0.24540398 | -1.2683783 | 0.2046629 | 0.63901002 |
| Tsen54 | 965.805043 | -0.4885976 | 0.28319046 | -1.7253323 | 0.08446761 | 0.44526833 |
| Llgl2 | 52.3344194 | -0.6586157 | 0.60318503 | -1.0918966 | 0.27487857 | 0.71147204 |
| Ncbp3 | 1997.82505 | -0.1840663 | 0.22446204 | -0.8200329 | 0.41219735 | 0.80251302 |
| Camkk1 | 834.119655 | -1.0808321 | 0.34835712 | -3.1026553 | 0.00191793 | 0.03890883 |
| P2rx1 | 1763.30251 | -1.0429915 | 0.26263321 | -3.9712858 | 7.148577139 | 0.00272425 |
| Atp2a3 | 1045.13212 | -1.6088021 | 0.4128075 | -3.8972212 | 9.730273630 | 0.00352161 |
| Ankfy1 | 1650.24542 | -0.1231157 | 0.24494677 | -0.5026221 | 0.61522997 | 0.8987425 |
| Exoc7 | 1813.02485 | 0.30226777 | 0.20339974 | 1.48607747 | 0.13725863 | 0.55981285 |
| Ube2g1 | 2870.55459 | 0.02341552 | 0.1509631 | 0.1551076 | 0.87673649 | 0.9745677 |
| Spns3 | 83.3793933 | -5.0463091 | 0.93631415 | -5.389547 | 7.063548962 | 7.448999521 |
| Med31 | 696.317704 | 0.38866525 | 0.44273912 | 0.87786515 | 0.3800169 | 0.78250296 |
| Ube2o | 211.641581 | -0.3935158 | 0.42391529 | -0.9282889 | 0.35325774 | 0.76376846 |
| Txndc17 | 4315.26196 | 0.33943812 | 0.28768118 | 1.17991077 | 0.23803571 | 0.67403588 |
| Rhbdf2 | 791.162904 | 0.00239435 | 0.43144026 | 0.00554966 | 0.99557203 | 0.99884675 |
| 4933427D14 | 230.190772 | 0.12361148 | 0.54694386 | 0.22600397 | 0.82119834 | 0.96205421 |
| Pimreg | 2984.4011 | 0.4183767 | 0.31965042 | 1.30885706 | 0.19058277 | 0.62154068 |
| Mxra7 | 31.2022843 | 2.8654288 | 0.78943317 | 3.62972942 | 0.00028372 | 0.00854234 |
| Rabep1 | 980.13096 | -0.0029135 | 0.18877959 | -0.0154332 | 0.98768656 | 0.9962376 |
| Mfsd11 | 4512.69601 | 1.15746432 | 0.19175309 | 6.03622252 | 1.577636635 | 2.718201261 |
| Sec14l1 | 1430.34023 | 0.56634017 | 0.27136899 | 2.08697455 | 0.03689044 | 0.28869994 |
| Mink1 | 61.4049168 | 0.31217108 | 0.57698753 | 0.5410361 | 0.5884827 | 0.88690478 |
| Pld2 | 212.26953 | 1.78717374 | 0.54437947 | 3.28295579 | 0.00102725 | 0.02384501 |
| Slc46a1 | 588.196478 | 1.31353416 | 0.3253047 | 4.03785796 | 5.394149578 | 0.00213549 |

| | | | | | | |
|---|---|---|---|---|---|---|
| 0610010K14l | 2506.28023 | 0.27052651 | 0.2607431 | 1.03752126 | 0.299493 | 0.73113229 |
| Eral1 | 1418.77707 | -0.2071352 | 0.22529533 | -0.9193943 | 0.35788936 | 0.76674184 |
| Dhrs13 | 380.029253 | -0.073293 | 0.34023954 | -0.2154159 | 0.82944308 | 0.96462215 |
| Coro6 | 23.7492647 | 1.39803566 | 1.83785569 | 0.76068848 | 0.44684316 | 0.82248455 |
| Blmh | 11273.8189 | 0.15064136 | 0.16767388 | 0.89841875 | 0.36896234 | 0.77480275 |
| Cpd | 360.183668 | -0.381677 | 0.38873324 | -0.981848 | 0.32617472 | 0.74761857 |
| Timm22 | 2446.3161 | 0.04692807 | 0.25827322 | 0.18169934 | 0.85581868 | 0.96915653 |
| Nxn | 96.2754428 | -0.5429732 | 0.54414073 | -0.9978544 | 0.31834997 | 0.74336577 |
| Rflnb | 173.742652 | -2.9653996 | 1.00157858 | -2.9607259 | 0.00306915 | 0.05497058 |
| Ywhae | 21329.1328 | 0.05765944 | 0.18342899 | 0.31434202 | 0.75326128 | 0.94018119 |
| Prpf8 | 4150.35049 | -0.1922069 | 0.22324704 | -0.8609608 | 0.38925963 | 0.78942657 |
| Nme2 | 13807.9954 | 0.81303076 | 0.52402763 | 1.55150362 | 0.12078104 | 0.52910763 |
| Spag9 | 267.509159 | 0.56476842 | 0.38093815 | 1.4825725 | 0.13818803 | 0.56068131 |
| Luc7l3 | 1257.01009 | -0.1506642 | 0.23366584 | -0.6447851 | 0.51906647 | 0.8574936 |
| Ankrd40 | 446.096674 | -0.2292451 | 0.38188447 | -0.6002995 | 0.54830664 | 0.87059427 |
| Abcc3 | 562.769197 | 1.47220209 | 0.58121076 | 2.53299181 | 0.01130936 | 0.13890808 |
| Xylt2 | 453.178928 | 0.84863232 | 0.53292115 | 1.59241628 | 0.1112912 | 0.50888686 |
| Lrrc59 | 7543.29081 | -0.212004 | 0.14714765 | -1.440757 | 0.14965334 | 0.57685486 |
| Slc35b1 | 2504.50611 | -0.4825762 | 0.16302662 | -2.9601065 | 0.00307533 | 0.0550007 |
| Snx11 | 628.423935 | -0.26646 | 0.26475422 | -1.006443 | 0.31420251 | 0.74065408 |
| Scrn2 | 212.54803 | -0.2101992 | 0.32569504 | -0.6453865 | 0.51867673 | 0.8570838 |
| Fbxl20 | 542.814787 | 0.31614019 | 0.29838916 | 1.05948951 | 0.2893769 | 0.72395811 |
| Dlg4 | 174.989739 | -0.4930884 | 0.39403443 | -1.2513841 | 0.21079439 | 0.64401792 |
| Dvl2 | 571.655082 | 0.01218767 | 0.33476481 | 0.03640665 | 0.97095811 | 0.99443087 |
| Nr1d1 | 98.1781422 | 0.46143304 | 0.75382464 | 0.61212253 | 0.54045669 | 0.86741192 |
| Per1 | 590.162624 | -0.2951758 | 0.48648248 | -0.6067552 | 0.54401341 | 0.86947682 |
| Vamp2 | 166.881443 | 0.22198148 | 0.47144616 | 0.47085224 | 0.63774625 | 0.90599527 |
| Tmem107 | 184.791145 | 0.41607037 | 0.44065658 | 0.94420549 | 0.34506465 | 0.76002446 |
| Aurkb | 7728.42673 | -0.2469643 | 0.17641168 | -1.3999319 | 0.16153372 | 0.5919838 |
| Ctc1 | 2193.71956 | -0.209216 | 0.23602647 | -0.886409 | 0.37539717 | 0.77815049 |
| Pfas | 4490.60937 | -0.6855737 | 0.30695662 | -2.2334545 | 0.02551899 | 0.2326183 |
| Myh10 | 257.444789 | 0.91670798 | 0.51424024 | 1.78264537 | 0.07464405 | 0.42093241 |
| Pik3r5 | 759.841313 | -0.4163992 | 0.31172912 | -1.3357724 | 0.18162367 | 0.61290438 |
| Stx8 | 854.968074 | 0.26458279 | 0.23215131 | 1.13969977 | 0.2544114 | 0.68839078 |
| Adprm | 673.07033 | 0.14858104 | 0.32578228 | 0.45607465 | 0.6483363 | 0.90973357 |
| Top2a | 8777.72589 | -0.1841072 | 0.13124761 | -1.4027473 | 0.16069222 | 0.59102462 |
| Acly | 12179.6392 | 0.01517963 | 0.25719011 | 0.05902104 | 0.95293535 | 0.99149988 |
| Kat2a | 1897.90282 | -0.4138299 | 0.22865349 | -1.809856 | 0.07031812 | 0.40971651 |
| Stat5b | 757.063595 | 0.36038108 | 0.29589592 | 1.21793189 | 0.22324986 | 0.65665196 |
| Tmem101 | 599.246258 | 0.42397515 | 0.26397806 | 1.60609992 | 0.10825196 | 0.50301538 |
| Lsm12 | 1698.3104 | -0.2954752 | 0.20257106 | -1.4586248 | 0.1446684 | 0.57032823 |
| Ubtf | 3337.7527 | -0.4621202 | 0.18196899 | -2.5395548 | 0.01109937 | 0.13673569 |
| Ccdc43 | 2054.2356 | 0.02821489 | 0.26242074 | 0.10751777 | 0.91437823 | 0.98347559 |
| Eftud2 | 8248.68482 | -0.4044019 | 0.23198838 | -1.7431991 | 0.08129884 | 0.43986231 |
| Dcakd | 3119.59895 | 0.03137485 | 0.2040801 | 0.15373792 | 0.87781638 | 0.97488223 |
| Nmt1 | 6578.64302 | 0.12811142 | 0.18966698 | 0.6754545 | 0.49938705 | 0.84872332 |

| | | | | | | |
|---|---|---|---|---|---|---|
| Plcd3 | 19.1051241 | 1.12304384 | 1.36978284 | 0.81986999 | 0.41229023 | 0.80251302 |
| Map3k14 | 183.808031 | 0.11081564 | 0.32683371 | 0.33905817 | 0.73456591 | 0.93750666 |
| Gosr2 | 4958.45512 | -0.125692 | 0.17732835 | -0.7088095 | 0.47844269 | 0.83826833 |
| Klhl28 | 379.558738 | 0.3968399 | 0.38471747 | 1.03150994 | 0.30230175 | 0.73372823 |
| Fkbp3 | 6974.65728 | 0.03345319 | 0.2958484 | 0.11307546 | 0.90997073 | 0.98262295 |
| Scfd1 | 3716.68304 | 0.13636364 | 0.2073859 | 0.65753576 | 0.51083648 | 0.85359566 |
| Strn3 | 335.77528 | 0.21024623 | 0.30215033 | 0.69583319 | 0.4865333 | 0.84183502 |
| Ap4s1 | 647.561378 | 0.11162353 | 0.3253675 | 0.34306907 | 0.7315465 | 0.93653187 |
| Dtd2 | 961.537973 | 0.22958468 | 0.22295196 | 1.02974954 | 0.30312759 | 0.73428908 |
| Ston2 | 119.831172 | 1.11828186 | 0.65629758 | 1.70392501 | 0.0883951 | 0.45415255 |
| Gtf2a1 | 4062.12146 | -0.172138 | 0.2978705 | -0.5778954 | 0.56333476 | 0.8760902 |
| Sel1l | 4439.05273 | 0.30961026 | 0.24242193 | 1.27715448 | 0.20154771 | 0.63561051 |
| Dnaaf2 | 793.756067 | -0.2810132 | 0.23773312 | -1.1820531 | 0.23718464 | 0.67338667 |
| Pole2 | 1250.71368 | 0.30254706 | 0.34586494 | 0.87475493 | 0.38170727 | 0.78451361 |
| Klhdc2 | 1762.52123 | 0.34902461 | 0.29842979 | 1.16953676 | 0.24218744 | 0.67749347 |
| Nemf | 2376.10716 | -0.276321 | 0.23532887 | -1.174191 | 0.24031852 | 0.67564175 |
| Sec23a | 1627.49721 | 0.53850575 | 0.25122182 | 2.1435469 | 0.03206921 | 0.26624356 |
| L2hgdh | 468.708314 | -0.089469 | 0.35855078 | -0.2495296 | 0.80295112 | 0.95672466 |
| Trappc6b | 1432.95556 | 0.10821073 | 0.1715567 | 0.63075782 | 0.52819889 | 0.86198732 |
| Pnn | 1702.39724 | -0.4083046 | 0.21977727 | -1.8578109 | 0.06319587 | 0.38764352 |
| Mia2 | 1365.14242 | 0.09131869 | 0.38548492 | 0.23689304 | 0.81273977 | 0.95967622 |
| Galc | 288.718632 | 0.51922516 | 0.46886654 | 1.10740502 | 0.26811886 | 0.70459677 |
| Spata7 | 202.662042 | -0.4081497 | 0.38734907 | -1.0537 | 0.29202028 | 0.72584701 |
| Ptpn21 | 55.1297911 | 0.67750475 | 0.62586428 | 1.08251065 | 0.27902568 | 0.71377266 |
| Zc3h14 | 1476.22152 | -0.1407615 | 0.22948352 | -0.6133838 | 0.53962262 | 0.86719392 |
| Ttc8 | 274.588259 | 0.59869927 | 0.49636042 | 1.2061785 | 0.22774867 | 0.66161232 |
| Polr2h | 2257.4348 | -0.0804684 | 0.3412979 | -0.2357717 | 0.81360981 | 0.95988497 |
| Ppp2r3c | 2052.14322 | 0.14687998 | 0.27464566 | 0.5347981 | 0.59278947 | 0.88797037 |
| 1110008L16R | 820.016053 | -0.9372509 | 0.52366179 | -1.7898019 | 0.07348576 | 0.41734043 |
| Psma6 | 13625.7948 | 0.52822433 | 0.42629874 | 1.23909427 | 0.21531059 | 0.64854028 |
| Nfkbia | 9498.41278 | 0.76415893 | 0.96330533 | 0.79326762 | 0.42762192 | 0.81062404 |
| Ralgapa1 | 125.066833 | 0.83231735 | 0.49557264 | 1.67950625 | 0.09305342 | 0.46652562 |
| Mbip | 616.889777 | 0.69452919 | 0.42811392 | 1.62229995 | 0.10473914 | 0.49501875 |
| Gstz1 | 1209.50447 | 0.36290037 | 0.34066048 | 1.06528462 | 0.28674715 | 0.72260415 |
| Sptlc2 | 5412.67643 | 0.07376824 | 0.2325355 | 0.3172343 | 0.75106583 | 0.94003547 |
| Ahsa1 | 12229.8493 | -0.2610907 | 0.18168501 | -1.4370514 | 0.1507034 | 0.57859667 |
| Vipas39 | 1828.38619 | 0.08575294 | 0.23846545 | 0.35960319 | 0.7191439 | 0.93310883 |
| Snw1 | 4505.60937 | -0.1372196 | 0.15588319 | -0.8802722 | 0.37871189 | 0.78126394 |
| Slirp | 2380.50914 | 0.26648633 | 0.36938678 | 0.72142899 | 0.47064561 | 0.83381643 |
| Adck1 | 1349.35233 | -0.2922392 | 0.26794068 | -1.0906863 | 0.27541095 | 0.71181528 |
| Nova1 | 46.5495411 | 1.5981474 | 0.87478879 | 1.82689515 | 0.06771554 | 0.4009507 |
| Mthfd1 | 6055.9003 | -0.5888699 | 0.20258394 | -2.9067945 | 0.00365153 | 0.06309203 |
| Ppp2r5e | 402.088606 | 0.18629245 | 0.43306152 | 0.43017548 | 0.667068 | 0.91574939 |
| Sgpp1 | 210.350237 | -0.180721 | 0.4634655 | -0.3899341 | 0.69658527 | 0.92646244 |
| Akap5 | 94.405723 | -0.8947682 | 0.52455571 | -1.7057639 | 0.08805205 | 0.45353293 |
| Sptb | 9.32110781 | 1.92912413 | 2.16737974 | 0.89007205 | 0.3734272 | 0.77748705 |

| | | | | | | |
|---|---|---|---|---|---|---|
| Fut8 | 986.421491 | -0.2103917 | 0.30452694 | -0.6908805 | 0.48964061 | 0.84434362 |
| Atl1 | 33.2504155 | 1.31762017 | 1.17521751 | 1.12117132 | 0.26221495 | 0.69762406 |
| Sav1 | 32.6414682 | -0.9995518 | 1.12332617 | -0.8898144 | 0.37356556 | 0.77748705 |
| Nin | 490.648561 | -0.8511469 | 0.3713643 | -2.291946 | 0.02190876 | 0.21320672 |
| Pygl | 2769.60016 | -0.3179638 | 0.23398793 | -1.3588899 | 0.17418149 | 0.60653633 |
| Tmx1 | 6423.99794 | 0.124933 | 0.17723124 | 0.70491523 | 0.48086299 | 0.83962275 |
| Actr10 | 3549.74074 | 0.20549593 | 0.30269908 | 0.67887862 | 0.49721478 | 0.84814011 |
| Timm9 | 1542.79846 | -0.0675753 | 0.33956504 | -0.1990056 | 0.84225838 | 0.96765585 |
| Rtn1 | 466.602239 | 0.55138676 | 0.35282569 | 1.56277382 | 0.1181058 | 0.52499575 |
| Lrrc9 | 28.6212597 | -0.7314708 | 1.06207023 | -0.6887217 | 0.49099841 | 0.84477968 |
| Dhrs7 | 1177.80684 | 0.22730048 | 0.30452509 | 0.74640968 | 0.45541997 | 0.82793172 |
| Ppm1a | 268.493418 | -0.2375714 | 0.40280177 | -0.5897972 | 0.5553266 | 0.8722354 |
| 4930447C04 | 30.7363672 | -0.6308738 | 1.22375725 | -0.515522 | 0.60618831 | 0.89258167 |
| Glrx5 | 1363.84702 | 0.12562165 | 0.32708373 | 0.38406572 | 0.70092973 | 0.92722415 |
| Mnat1 | 1122.715 | -0.0219384 | 0.3297553 | -0.0665294 | 0.94695635 | 0.99019648 |
| Prkch | 1071.01204 | 0.3120999 | 0.42387904 | 0.73629472 | 0.46155136 | 0.83025001 |
| Hif1a | 5104.12483 | 0.05148839 | 0.24121903 | 0.2134508 | 0.83097536 | 0.96466657 |
| Papola | 6559.5161 | -0.2171705 | 0.20481678 | -1.0603157 | 0.28900098 | 0.72395811 |
| Mpp5 | 428.007975 | 0.48419746 | 0.28780573 | 1.68237601 | 0.09249595 | 0.46597694 |
| Snapc1 | 630.96064 | -0.1755618 | 0.4029453 | -0.4356965 | 0.66305698 | 0.91395288 |
| Atp6v1d | 3145.86141 | 0.17817222 | 0.17938158 | 0.99325816 | 0.32058416 | 0.74413737 |
| Vrk1 | 6698.6065 | -0.23957 | 0.16657949 | -1.4381724 | 0.15038516 | 0.57837915 |
| Eif2s1 | 18381.7245 | 0.15371199 | 0.30935826 | 0.49687372 | 0.6192781 | 0.89896588 |
| Pigh | 988.63379 | 0.45686097 | 0.30036808 | 1.52100372 | 0.1282589 | 0.54327949 |
| Rdh12 | 22.5480903 | -2.8698108 | 1.25778344 | -2.2816415 | 0.02251052 | 0.21507833 |
| Vti1b | 3248.92425 | 0.46817714 | 0.34860882 | 1.34298709 | 0.17927616 | 0.60992118 |
| Arg2 | 20.6017071 | 0.59065748 | 1.86197156 | 0.31722154 | 0.75107551 | 0.94003547 |
| Zfp36l1 | 2925.01686 | 0.32003359 | 0.25644175 | 1.24797771 | 0.21203922 | 0.64580271 |
| Erh | 16556.644 | 0.18744076 | 0.33899247 | 0.5529349 | 0.58030799 | 0.88286755 |
| Susd6 | 1272.88502 | 0.06155358 | 0.31972122 | 0.19252266 | 0.84733282 | 0.96765585 |
| Srsf5 | 5049.41166 | -0.1030227 | 0.18413096 | -0.5595077 | 0.57581528 | 0.88080372 |
| Smoc1 | 20.8214405 | -1.6917758 | 1.15913637 | -1.459514 | 0.14442369 | 0.57027792 |
| Gm20498 | 278.549175 | 0.23293938 | 0.46731003 | 0.4984686 | 0.6181538 | 0.89878056 |
| Pcnx | 212.037156 | -0.6417418 | 0.49613483 | -1.2934827 | 0.19584418 | 0.62779346 |
| Pacs2 | 343.823553 | 0.32168439 | 0.3662155 | 0.87840189 | 0.37972566 | 0.78237978 |
| Mta1 | 118.928559 | -1.2448294 | 0.52419935 | -2.3747251 | 0.01756202 | 0.18569273 |
| Wdr37 | 418.962476 | 0.18076123 | 0.33864974 | 0.53377046 | 0.59350035 | 0.88797037 |
| Gtpbp4 | 4780.17363 | -0.3020327 | 0.22997979 | -1.3133011 | 0.18908151 | 0.61961804 |
| Zmynd11 | 899.989479 | 0.14533952 | 0.27185108 | 0.5346292 | 0.59290629 | 0.88797037 |
| Esyt2 | 210.241069 | -0.66493 | 0.33810173 | -1.9666567 | 0.0492228 | 0.34139551 |
| Cdca7l | 2316.15131 | -0.5417608 | 0.22434901 | -2.4148125 | 0.01574331 | 0.17444227 |
| Efcab11 | 443.872329 | 0.71980368 | 0.29061625 | 2.47681839 | 0.01325593 | 0.15566077 |
| Tdp1 | 2942.1203 | -0.3676376 | 0.19572915 | -1.8782974 | 0.0603405 | 0.37881203 |
| Psmc1 | 12496.9094 | 0.03085206 | 0.18090813 | 0.17053996 | 0.86458551 | 0.9705375 |
| Nrde2 | 721.151106 | -0.430856 | 0.33742194 | -1.2769057 | 0.20163555 | 0.63572364 |
| Rps6ka5 | 104.419409 | 0.04565137 | 0.61538437 | 0.07418351 | 0.94086436 | 0.98935054 |

| | | | | | | |
|---|---|---|---|---|---|---|
| Ccdc88c | 47.6557956 | -1.7664371 | 1.31451041 | -1.3437985 | 0.17901356 | 0.60992118 |
| Dglucy | 62.4130105 | -0.3259927 | 1.33821493 | -0.2436027 | 0.80753856 | 0.95797316 |
| Fbln5 | 8.76916779 | 1.87536685 | 2.16527427 | 0.86611053 | 0.38642955 | 0.78902002 |
| Trip11 | 580.804368 | 0.37457951 | 0.39800453 | 0.94114383 | 0.34663116 | 0.76056509 |
| Atxn3 | 893.932588 | 0.41538435 | 0.24269259 | 1.71156585 | 0.08697671 | 0.45109601 |
| Lgmn | 39926.1737 | 0.94236541 | 0.17123995 | 5.50318672 | 3.729870192 | 4.166133446 |
| Golga5 | 1034.42177 | 0.10062628 | 0.20303098 | 0.49562031 | 0.62016232 | 0.89896588 |
| Pitrm1 | 3531.18175 | -0.1425649 | 0.28773909 | -0.4954657 | 0.62027143 | 0.89896588 |
| Pfkp | 3412.66751 | -0.6117273 | 0.26185798 | -2.3361033 | 0.01948586 | 0.19765383 |
| Asb2 | 1263.88815 | -0.7132959 | 0.2021737 | -3.5281339 | 0.0004185 | 0.01163526 |
| Otub2 | 164.319693 | 0.26109068 | 0.59989632 | 0.43522633 | 0.66339815 | 0.91395288 |
| Akr1c12 | 25.8830985 | 1.96341551 | 1.55831783 | 1.25995832 | 0.2076844 | 0.6418276 |
| Akr1c13 | 90.9817613 | 0.50252805 | 0.78491805 | 0.64022997 | 0.5220231 | 0.85816354 |
| Net1 | 845.307669 | -0.6629274 | 0.29013409 | -2.2849 | 0.02231869 | 0.21464194 |
| Tshz3 | 26.0800312 | 2.18106388 | 1.40073504 | 1.55708526 | 0.11945024 | 0.52676093 |
| Gdi2 | 30685.0329 | 0.02258852 | 0.15345636 | 0.14719832 | 0.88297549 | 0.97644541 |
| Dcaf4 | 879.742541 | -0.4134948 | 0.21818766 | -1.895134 | 0.05807465 | 0.37020699 |
| Numb | 733.481225 | -0.1454516 | 0.32848318 | -0.4427977 | 0.65791203 | 0.91258818 |
| Acot2 | 584.805627 | 0.76018188 | 0.27916859 | 2.72302082 | 0.0064688 | 0.09721475 |
| Fam161b | 28.5905546 | 0.63293857 | 1.38503925 | 0.4569824 | 0.64768369 | 0.9097332 |
| Coq6 | 1421.45697 | 0.07502526 | 0.2872755 | 0.26116137 | 0.79396807 | 0.95310033 |
| Entpd5 | 232.775984 | 0.91081463 | 0.42878936 | 2.12415398 | 0.03365727 | 0.27375624 |
| Aldh6a1 | 802.801735 | 0.50981354 | 0.28352091 | 1.79815146 | 0.07215301 | 0.41303124 |
| Abcd4 | 477.258046 | 0.69598479 | 0.30506354 | 2.28144203 | 0.0225223 | 0.21507833 |
| Isca2 | 1524.91185 | 0.30248562 | 0.35853989 | 0.84365959 | 0.39885966 | 0.7957029 |
| Npc2 | 12286.1769 | 0.37324652 | 0.27585106 | 1.35307265 | 0.17603242 | 0.60763832 |
| Fcf1 | 3326.96415 | -0.0054315 | 0.30105097 | -0.0180418 | 0.98560551 | 0.99564261 |
| Ylpm1 | 463.128901 | -0.3010705 | 0.34344885 | -0.8766094 | 0.38069883 | 0.78323054 |
| Mlh3 | 125.499769 | -0.1000613 | 0.57184414 | -0.17498 | 0.86109539 | 0.96996131 |
| Tmed10 | 11409.9175 | 0.10326961 | 0.16918036 | 0.61041133 | 0.54158936 | 0.86797625 |
| Fos | 43728.172 | 0.13764345 | 0.41930063 | 0.32826912 | 0.74270819 | 0.93850397 |
| Erg28 | 2162.70962 | 0.11876353 | 0.28957265 | 0.41013381 | 0.6817078 | 0.92061965 |
| Tgfb3 | 13.1735188 | -2.2007672 | 2.06598024 | -1.0652412 | 0.28676681 | 0.72260415 |
| Gpatch2l | 847.825618 | -0.2924705 | 0.31803847 | -0.9196074 | 0.35777794 | 0.76674184 |
| Angel1 | 159.828248 | 0.5491586 | 0.47771657 | 1.14954898 | 0.25032968 | 0.6842534 |
| Ccnk | 244.687511 | -0.3328488 | 0.35343754 | -0.9417471 | 0.34632212 | 0.7602916 |
| Evl | 634.044842 | -0.4277614 | 0.22236586 | -1.9236827 | 0.05439437 | 0.36033644 |
| Yy1 | 757.53757 | -0.3716387 | 0.24044434 | -1.5456331 | 0.12219321 | 0.53138624 |
| Slc25a29 | 65.005565 | -0.8677395 | 0.76905958 | -1.1283124 | 0.25918799 | 0.69546977 |
| Wars | 5764.95627 | -0.1445381 | 0.19510105 | -0.740837 | 0.45879226 | 0.82986925 |
| Hsp90aa1 | 68997.5876 | -0.3217405 | 0.14693874 | -2.1896235 | 0.02855155 | 0.25145509 |
| Zfp839 | 105.209736 | 0.53302244 | 0.54873427 | 0.97136715 | 0.33136548 | 0.75038762 |
| Fdft1 | 2629.41596 | 0.44450659 | 0.21588025 | 2.0590424 | 0.03949018 | 0.30082941 |
| Tecpr2 | 121.255365 | 0.02416458 | 0.52264226 | 0.0462354 | 0.96312262 | 0.99401755 |
| Cinp | 3611.51272 | 0.29793425 | 0.31950691 | 0.93248141 | 0.35108779 | 0.76309919 |
| Traf3 | 166.985591 | -0.1635835 | 0.65970109 | -0.2479661 | 0.80416064 | 0.9570372 |

| | | | | | | |
|---|---|---|---|---|---|---|
| Cdc42bpb | 243.728502 | 0.33569266 | 0.33735373 | 0.99507616 | 0.31969922 | 0.7440831 |
| Tnfaip2 | 44.1471901 | 0.44697712 | 0.8333118 | 0.53638641 | 0.59169152 | 0.88797037 |
| Eif5 | 16659.2546 | -0.163601 | 0.16155309 | -1.0126763 | 0.31121482 | 0.74011268 |
| Ppp1r13b | 207.746693 | 0.25617177 | 0.34531847 | 0.74184208 | 0.45818302 | 0.82986925 |
| Zfyve21 | 1704.27696 | 0.88323439 | 0.40642519 | 2.17317828 | 0.0297669 | 0.25715996 |
| Xrcc3 | 938.849969 | -0.222637 | 0.24904526 | -0.8939622 | 0.37134213 | 0.7765614 |
| Klc1 | 1832.55878 | -0.0851509 | 0.20310343 | -0.4192489 | 0.67503421 | 0.91848602 |
| Atp5mpl | 4700.59458 | 0.51526248 | 0.43865921 | 1.17463049 | 0.24014258 | 0.67563574 |
| Gpr132 | 27.4639901 | -1.1154635 | 1.31930259 | -0.8454949 | 0.3978346 | 0.79531492 |
| Ggps1 | 1329.47514 | 0.13780404 | 0.35774496 | 0.38520191 | 0.70008783 | 0.92669304 |
| Gpr137b | 125.744866 | 0.79869436 | 0.56376251 | 1.4167213 | 0.15656443 | 0.58588334 |
| Mtr | 2476.34813 | 0.21961009 | 0.28459489 | 0.7716586 | 0.44031666 | 0.81963526 |
| Gli3 | 27.8204463 | -0.1178746 | 0.94786716 | -0.1243577 | 0.90103207 | 0.98043702 |
| Aoah | 2359.03562 | 0.58659894 | 0.26015532 | 2.2548028 | 0.02414571 | 0.22455944 |
| Trim27 | 7406.61569 | -0.279446 | 0.19505961 | -1.4326186 | 0.15196686 | 0.5800345 |
| Zkscan3 | 1047.12886 | 0.28783505 | 0.20741506 | 1.38772492 | 0.16522082 | 0.59797229 |
| Carmil1 | 97.7843481 | 0.03161038 | 0.67662217 | 0.04671791 | 0.96273805 | 0.99401755 |
| Mrs2 | 751.420878 | -0.1988626 | 0.37426848 | -0.5313369 | 0.59518537 | 0.88830085 |
| Gpld1 | 8.50481026 | -1.9766024 | 2.63524322 | -0.7500645 | 0.45321585 | 0.82610332 |
| Irf4 | 62.1088933 | 0.41500844 | 1.29820292 | 0.31967918 | 0.74921154 | 0.9399191 |
| Exoc2 | 1477.64167 | 0.34815834 | 0.17726012 | 1.96410976 | 0.04951736 | 0.34145768 |
| Gcnt2 | 530.710139 | -0.9727636 | 0.51724945 | -1.880647 | 0.06001995 | 0.37827102 |
| Tmem14c | 7630.57958 | 0.19625081 | 0.37576233 | 0.52227377 | 0.60147973 | 0.89094157 |
| Gcm2 | 14.8794464 | -2.261812 | 2.43509935 | -0.9288377 | 0.35297322 | 0.76376537 |
| Elovl2 | 49.2151824 | 1.50374019 | 1.25849754 | 1.1948694 | 0.23213806 | 0.66733333 |
| Nedd9 | 1257.68161 | -0.8439332 | 0.21727623 | -3.8841487 | 0.00010269 | 0.00368315 |
| Hivep1 | 140.868511 | 0.90904801 | 0.72035815 | 1.26193894 | 0.20697076 | 0.64174209 |
| Edn1 | 23.7506004 | -0.2219234 | 1.59408588 | -0.1392167 | 0.88927891 | 0.97870754 |
| Tbc1d7 | 724.049871 | 0.24716761 | 0.24270551 | 1.0183848 | 0.30849512 | 0.7383723 |
| Mcur1 | 1996.6751 | -0.1035551 | 0.26015133 | -0.3980571 | 0.6905881 | 0.92470267 |
| Nup153 | 699.600519 | -0.4995184 | 0.32574663 | -1.5334567 | 0.12516337 | 0.53667138 |
| Kif13a | 260.503589 | -1.3449932 | 0.80022793 | -1.6807627 | 0.09280902 | 0.46606436 |
| Tpmt | 311.518396 | 0.50118042 | 0.32142293 | 1.55925534 | 0.11893596 | 0.52515149 |
| Dek | 7904.43025 | -0.4197129 | 0.19347107 | -2.1693831 | 0.03005361 | 0.25895274 |
| Susd3 | 1139.5778 | 0.25039507 | 0.38448376 | 0.65125004 | 0.5148851 | 0.85532175 |
| Ippk | 1297.42771 | -0.0405517 | 0.24123199 | -0.1681027 | 0.86650248 | 0.97060021 |
| Cenpp | 471.662615 | -0.115417 | 0.36156654 | -0.3192136 | 0.74956453 | 0.9399191 |
| Nol8 | 755.362268 | -0.1609441 | 0.33678212 | -0.4778878 | 0.63273009 | 0.90430301 |
| Spin1 | 1111.46139 | -0.2359397 | 0.19016547 | -1.2407075 | 0.21471384 | 0.64854028 |
| Wrnip1 | 744.323248 | -0.2507641 | 0.6866552 | -0.3651965 | 0.71496469 | 0.93235217 |
| Ripk1 | 1791.71711 | 0.66966494 | 0.49870904 | 1.34279687 | 0.17933776 | 0.60992118 |
| Pxdc1 | 178.314104 | 1.81764879 | 0.58747587 | 3.09399738 | 0.00197479 | 0.03966771 |
| Prpf4b | 958.512215 | -0.1536579 | 0.41712934 | -0.36837 | 0.71259734 | 0.93235217 |
| Eci2 | 2444.12572 | 0.25940762 | 0.2407961 | 1.07729164 | 0.28135 | 0.71643516 |
| Rpp40 | 1291.56707 | -0.1342629 | 0.37117343 | -0.3617256 | 0.71755712 | 0.93275665 |
| Fars2 | 1198.85587 | 0.06963708 | 0.26049732 | 0.26732359 | 0.78922003 | 0.95255415 |

| | | | | | | |
|---|---|---|---|---|---|---|
| Ly86 | 13763.8128 | 0.03373131 | 0.26037366 | 0.12954964 | 0.89692275 | 0.97984623 |
| Ssr1 | 22726.069 | -0.1100862 | 0.18769821 | -0.5865065 | 0.55753518 | 0.87283439 |
| Riok1 | 1203.62861 | -0.2667607 | 0.22954614 | -1.1621225 | 0.24518572 | 0.68039867 |
| Snrnp48 | 409.541681 | 0.33540488 | 0.32463469 | 1.03317632 | 0.30152139 | 0.73330242 |
| Slc35b3 | 844.61987 | 0.60788751 | 0.38681797 | 1.57150795 | 0.11606471 | 0.52084358 |
| Sema4d | 1583.38622 | -0.9985168 | 0.37367538 | -2.6721504 | 0.00753669 | 0.10658529 |
| Gadd45g | 633.721331 | 0.53536741 | 0.2277585 | 2.35059245 | 0.01874355 | 0.19316753 |
| Fbp2 | 29.7294229 | -2.1296447 | 1.34497454 | -1.5834089 | 0.11332829 | 0.51285577 |
| Syk | 2383.7108 | -0.0188106 | 0.27516535 | -0.0683611 | 0.94549818 | 0.99019648 |
| Aopep | 388.401952 | -0.2266223 | 0.25132163 | -0.9017222 | 0.36720444 | 0.77315179 |
| Auh | 1080.82969 | 0.08180202 | 0.26675361 | 0.30665758 | 0.75910402 | 0.94184564 |
| Fancc | 783.583879 | -0.1480027 | 0.25555809 | -0.5791354 | 0.56249778 | 0.8755831 |
| Ptch1 | 128.544493 | 0.57177609 | 0.67119879 | 0.85187294 | 0.39428462 | 0.79233901 |
| Sptlc1 | 2589.18073 | 0.00176527 | 0.16733575 | 0.01054927 | 0.99158306 | 0.99720779 |
| Ercc6l2 | 496.566902 | -0.0308626 | 0.33052344 | -0.0933748 | 0.92560583 | 0.98580325 |
| Sfxn1 | 5917.20381 | -0.3916857 | 0.15145994 | -2.5860683 | 0.00970777 | 0.12687515 |
| Habp4 | 76.652988 | 0.94621155 | 0.78015128 | 1.2128565 | 0.22518469 | 0.65870021 |
| Ctsl | 21533.3916 | 0.60343868 | 0.18368701 | 3.28514614 | 0.00101929 | 0.02370539 |
| Zfp346 | 479.642336 | -0.0589388 | 0.30186561 | -0.1952485 | 0.84519841 | 0.96765585 |
| Prxl2c | 2206.56653 | 0.30686213 | 0.18615965 | 1.6483815 | 0.09927441 | 0.48095994 |
| Cdk20 | 939.678174 | 0.1006108 | 0.26764791 | 0.37590727 | 0.70698584 | 0.92935286 |
| Lman2 | 13171.0001 | -0.1796923 | 0.16757244 | -1.0723262 | 0.28357356 | 0.71951357 |
| Mxd3 | 71.4711156 | 0.20125493 | 0.57468218 | 0.35020215 | 0.726187 | 0.93561871 |
| Prelid1 | 19009.2466 | -0.0181976 | 0.15830272 | -0.1149541 | 0.90848149 | 0.98192738 |
| Nsd1 | 1816.72881 | 0.01098178 | 0.19670552 | 0.05582851 | 0.95547842 | 0.99215259 |
| Pdlim7 | 403.720604 | 0.52794869 | 0.25707985 | 2.053637 | 0.04001084 | 0.30381911 |
| Ddx41 | 3623.50205 | 0.23152033 | 0.19779639 | 1.17049824 | 0.24180052 | 0.67708722 |
| Fam193b | 113.938762 | -0.0712238 | 0.53415099 | -0.1333401 | 0.8939244 | 0.97941957 |
| Pcbd2 | 537.669221 | 0.5973219 | 0.37304162 | 1.60122051 | 0.10932809 | 0.50544614 |
| Txndc15 | 1785.71234 | 0.21334975 | 0.24240875 | 0.88012396 | 0.37879216 | 0.78127879 |
| Ddx46 | 2397.33706 | -0.520311 | 0.31340502 | -1.6601871 | 0.09687682 | 0.47766794 |
| Caml | 924.015625 | 0.48053492 | 0.31487488 | 1.52611389 | 0.1269815 | 0.54111158 |
| B4galt7 | 752.458244 | 0.38614809 | 0.22040956 | 1.75195709 | 0.07978119 | 0.43686807 |
| Zfp729a | 169.403489 | -0.0504228 | 0.58031922 | -0.0868881 | 0.93076048 | 0.98716777 |
| Zfp369 | 91.7020786 | -0.5308708 | 0.56857143 | -0.9336924 | 0.3504626 | 0.76238338 |
| Ptdss1 | 2868.91568 | -0.1391566 | 0.20496981 | -0.6789125 | 0.49719332 | 0.84814011 |
| Mterf3 | 950.072488 | -0.1264558 | 0.29980314 | -0.421796 | 0.6731739 | 0.91786342 |
| Uqcrb | 7122.6547 | 0.17178192 | 0.26106556 | 0.65800298 | 0.51053622 | 0.85342847 |
| Fastkd3 | 1234.5549 | -0.1645018 | 0.24643482 | -0.6675266 | 0.50443584 | 0.85031882 |
| Cetn3 | 4083.75106 | 0.29188599 | 0.29295095 | 0.99636469 | 0.31907298 | 0.74398524 |
| Smad5 | 584.204957 | -0.0673432 | 0.24343985 | -0.2766317 | 0.78206295 | 0.95004728 |
| Hnrnpk | 68541.9374 | 0.01674516 | 0.18147542 | 0.09227232 | 0.92648168 | 0.98630274 |
| Ccnh | 3794.37115 | 0.12467204 | 0.23345501 | 0.53403025 | 0.5933206 | 0.88797037 |
| Rasa1 | 476.205848 | 0.06831261 | 0.33575023 | 0.20346259 | 0.83877348 | 0.96744446 |
| 2210016F16I | 2601.17699 | -0.0289312 | 0.21498415 | -0.1345738 | 0.89294882 | 0.97941957 |
| Gkap1 | 54.1379731 | 0.12554242 | 0.58293406 | 0.21536298 | 0.82948433 | 0.96462215 |

| | | | | | | |
|---|---|---|---|---|---|---|
| Naa35 | 1811.64682 | -0.2266361 | 0.27821705 | -0.8146016 | 0.41530037 | 0.80395688 |
| Golm1 | 3197.27497 | -0.0195576 | 0.1937631 | -0.1009356 | 0.91960155 | 0.9839023 |
| Agtpbp1 | 502.270064 | -0.8720489 | 1.35063966 | -0.6456562 | 0.51850202 | 0.8570838 |
| Dapk1 | 188.510869 | 1.75115726 | 0.51519327 | 3.3990298 | 0.00067625 | 0.01729002 |
| Trip13 | 4364.6637 | 0.00801956 | 0.23581707 | 0.03400753 | 0.97287114 | 0.99443087 |
| Cep72 | 678.036724 | -0.4829466 | 0.24559294 | -1.9664514 | 0.04924649 | 0.34139551 |
| Tppp | 62.8866639 | 0.10024107 | 0.76120594 | 0.13168719 | 0.89523172 | 0.97949899 |
| Ahrr | 58.6207955 | 0.50070767 | 0.77895648 | 0.64279287 | 0.52035851 | 0.8576517 |
| Pdcd6 | 5779.7121 | 0.32349803 | 0.34819853 | 0.92906202 | 0.35285695 | 0.76376537 |
| Sdha | 21051.7937 | -0.1922048 | 0.22856485 | -0.8409204 | 0.40039253 | 0.79668215 |
| Ccdc127 | 1981.96825 | 0.2547755 | 0.25291109 | 1.00737181 | 0.31375613 | 0.74065408 |
| Erap1 | 3014.2524 | -0.3378506 | 0.20758013 | -1.627567 | 0.10361672 | 0.49263248 |
| Cast | 2315.37724 | 0.04752511 | 0.19632318 | 0.24207589 | 0.80872136 | 0.95844313 |
| Rhobtb3 | 29.9085965 | 0.49162609 | 0.72593751 | 0.67722921 | 0.49826054 | 0.84860011 |
| Glrx | 7999.8713 | -0.045635 | 0.14651182 | -0.3114769 | 0.75543809 | 0.94087513 |
| Arsk | 553.558014 | 0.5626767 | 0.24533189 | 2.29353262 | 0.02181736 | 0.21283234 |
| Srd5a1 | 178.165742 | -0.1329606 | 0.42004251 | -0.3165407 | 0.75159212 | 0.94018119 |
| Nsun2 | 15882.2763 | -0.3068716 | 0.15930804 | -1.926278 | 0.05406967 | 0.35989907 |
| Mctp1 | 640.075198 | -0.4277048 | 0.35917042 | -1.190813 | 0.233727 | 0.66954422 |
| Slf1 | 377.285542 | -0.0099855 | 0.31961297 | -0.0312426 | 0.97507606 | 0.99450266 |
| Med10 | 2937.92881 | 0.49976667 | 0.28782866 | 1.7363339 | 0.08250481 | 0.44183004 |
| Ndufs6 | 2713.22526 | 0.2557143 | 0.40721358 | 0.62796113 | 0.5300294 | 0.86347712 |
| Mrpl36 | 3067.02414 | -0.3027099 | 0.223967 | -1.3515826 | 0.17650888 | 0.60764153 |
| Lpcat1 | 1183.97572 | -0.1055396 | 0.4090564 | -0.2580074 | 0.79640117 | 0.9537323 |
| Clptm1l | 3846.19585 | -0.0267893 | 0.35350757 | -0.0757814 | 0.93959304 | 0.98913801 |
| Tert | 14.4115146 | 1.58882179 | 1.76571442 | 0.89981809 | 0.36821706 | 0.77421783 |
| Hapln1 | 166.191268 | 1.95375935 | 1.08809728 | 1.79557414 | 0.07256227 | 0.41440442 |
| Vcan | 17.9754106 | -7.3070074 | 1.63841981 | -4.4597895 | 8.204019457 | 0.00044017 |
| Xrcc4 | 444.038409 | 0.46001172 | 0.42426227 | 1.08426262 | 0.27824838 | 0.71360814 |
| Atg10 | 570.344647 | 0.42194384 | 0.33103848 | 1.27460663 | 0.20244851 | 0.63746528 |
| Zcchc9 | 870.725663 | 0.020077 | 0.22370136 | 0.08974912 | 0.92848658 | 0.98666423 |
| Cd180 | 838.006606 | 0.29387964 | 0.39737642 | 0.73954976 | 0.45957324 | 0.82987212 |
| Slc30a5 | 5068.59356 | 0.08587181 | 0.2374832 | 0.3615911 | 0.71765762 | 0.93275665 |
| Rad17 | 1224.46679 | -0.2646884 | 0.20851286 | -1.2694104 | 0.20429471 | 0.63848041 |
| Marveld2 | 58.4763348 | 0.48640716 | 1.19293277 | 0.40774063 | 0.68346411 | 0.92201328 |
| Ocln | 33.7426894 | 2.39148869 | 1.75004683 | 1.3665284 | 0.17177317 | 0.60430173 |
| Gtf2h2 | 2030.2186 | -0.0693611 | 0.32282543 | -0.2148565 | 0.82987924 | 0.96464393 |
| Serf1 | 117.72811 | 0.06045187 | 0.57995561 | 0.10423533 | 0.91698259 | 0.98390037 |
| Smn1 | 1311.97013 | -0.4155705 | 0.24723652 | -1.6808623 | 0.09278966 | 0.46606436 |
| Mccc2 | 992.719734 | 0.06524344 | 0.31885724 | 0.20461646 | 0.83787179 | 0.96683245 |
| Ptcd2 | 2959.55366 | -0.1704438 | 0.18331656 | -0.9297784 | 0.35248583 | 0.76358405 |
| Btf3 | 16861.7583 | 0.24093793 | 0.37898579 | 0.63574398 | 0.52494331 | 0.86011673 |
| Ankra2 | 236.863075 | 0.20810093 | 0.32188501 | 0.64650705 | 0.51795103 | 0.8570838 |
| Arhgef28 | 10.5693208 | -1.713403 | 1.99820025 | -0.8574731 | 0.39118345 | 0.79005237 |
| Hexb | 6461.90962 | -0.056487 | 0.19108872 | -0.295606 | 0.76753097 | 0.94401697 |
| Gfm2 | 1819.18822 | -0.1494816 | 0.19032226 | -0.7854132 | 0.43221137 | 0.81379739 |

| | | | | | | |
|---|---|---|---|---|---|---|
| Polk | 289.204337 | 0.30391264 | 0.3478962 | 0.87357276 | 0.38235097 | 0.78530883 |
| Col4a3bp | 559.82191 | -0.1892783 | 0.28151766 | -0.6723496 | 0.50136113 | 0.84968141 |
| Hmgcr | 4074.9193 | 0.48003981 | 0.22433561 | 2.13982883 | 0.0323686 | 0.26700278 |
| Poc5 | 1144.89679 | 0.06785128 | 0.20837047 | 0.3256281 | 0.74470575 | 0.93917909 |
| Iqgap2 | 585.81203 | 0.08076692 | 0.3464337 | 0.23313817 | 0.81565412 | 0.96063519 |
| Aggf1 | 2139.56756 | -0.2938751 | 0.29357627 | -1.0010178 | 0.31681821 | 0.74186126 |
| Pde8b | 45.2483624 | -0.9675522 | 1.22142481 | -0.7921504 | 0.42827296 | 0.81130995 |
| Ap3b1 | 3747.56735 | 0.08236687 | 0.23253089 | 0.35421905 | 0.72317473 | 0.93433544 |
| Scamp1 | 2090.08992 | 0.15804095 | 0.26505808 | 0.59625027 | 0.55100805 | 0.87093412 |
| Jmy | 102.356682 | 0.04455542 | 0.6605336 | 0.06745368 | 0.94622053 | 0.99019648 |
| Dimt1 | 951.105376 | 0.15291259 | 0.34089552 | 0.44856146 | 0.65374804 | 0.91076467 |
| Kif2a | 426.51853 | -0.325141 | 0.35514702 | -0.9155109 | 0.35992346 | 0.76857232 |
| Ercc8 | 859.300058 | -0.4984286 | 0.25743512 | -1.9361331 | 0.0528514 | 0.35543223 |
| Elovl7 | 159.782193 | -2.0334133 | 0.65665724 | -3.0966129 | 0.00195745 | 0.03946709 |
| Depdc1b | 300.666086 | -0.1605313 | 0.46759046 | -0.3433161 | 0.73136067 | 0.93653187 |
| Pde4d | 91.9653542 | 0.09782238 | 0.55903486 | 0.1749844 | 0.86109189 | 0.96996131 |
| Plk2 | 4325.32696 | 0.22026829 | 0.24671396 | 0.89280841 | 0.37195978 | 0.77719221 |
| Serinc5 | 90.0458214 | 1.14122007 | 0.54645731 | 2.08839747 | 0.036762 | 0.2879062 |
| Mtx3 | 45.0697552 | -0.2192475 | 0.69408722 | -0.3158789 | 0.7520944 | 0.94018119 |
| Zfyve16 | 234.772749 | 0.1893755 | 0.3925791 | 0.48238812 | 0.62953026 | 0.90310807 |
| Dhfr | 3845.94482 | 0.03298961 | 0.20172264 | 0.16353944 | 0.87009373 | 0.97248575 |
| Erbin | 558.606358 | -0.1950523 | 0.31657784 | -0.6161274 | 0.53781046 | 0.86655355 |
| Nln | 1472.25144 | 0.18281213 | 0.20362241 | 0.89779962 | 0.36929238 | 0.77488057 |
| Trappc13 | 3383.00034 | 0.2748785 | 0.19173845 | 1.43361176 | 0.15168309 | 0.57985599 |
| Trim23 | 707.991766 | 0.18525128 | 0.35562247 | 0.52092119 | 0.60242168 | 0.89131886 |
| Ppwd1 | 545.040636 | -0.1507531 | 0.38958288 | -0.3869602 | 0.69878571 | 0.92669304 |
| Cenpk | 1473.30486 | 0.03231449 | 0.30761702 | 0.10504781 | 0.91633787 | 0.98386323 |
| Cwc27 | 961.690999 | -0.1628355 | 0.32927892 | -0.4945216 | 0.62093789 | 0.89896588 |
| Srek1ip1 | 428.398594 | 0.31416364 | 0.33727909 | 0.93146493 | 0.35161312 | 0.76348958 |
| Rgs7bp | 439.516675 | 1.82305555 | 0.50186989 | 3.63252623 | 0.00028066 | 0.00847732 |
| Rnf180 | 338.237779 | 1.23686174 | 0.38275046 | 3.23150949 | 0.00123138 | 0.02719043 |
| Parp8 | 819.953032 | -1.1252625 | 0.26092335 | -4.3126172 | 1.613333245 | 0.000777 |
| Emb | 22908.9344 | -0.8423457 | 0.1814595 | -4.642059 | 3.449542568 | 0.00020994 |
| Mrps30 | 3538.96041 | -0.023696 | 0.22398087 | -0.1057945 | 0.91574538 | 0.98369452 |
| Slc4a7 | 646.518792 | -0.5281644 | 0.33992269 | -1.5537781 | 0.12023735 | 0.5281377 |
| Psmd6 | 8756.74248 | -0.0198537 | 0.20282506 | -0.0978856 | 0.92202315 | 0.98438725 |
| Atxn7 | 94.8768911 | -0.2521659 | 0.63525943 | -0.3969495 | 0.69140471 | 0.92487194 |
| Ptprg | 51.4652095 | 1.23698579 | 1.25762978 | 0.983585 | 0.3253196 | 0.74692844 |
| Pdhb | 7727.60167 | -0.1078288 | 0.1995828 | -0.5402709 | 0.58901023 | 0.88691905 |
| Kctd6 | 226.504427 | 0.55729325 | 0.54210501 | 1.02801716 | 0.30394175 | 0.73436097 |
| Map3k1 | 4406.11861 | -0.3772634 | 0.39089136 | -0.9651363 | 0.33447655 | 0.75348848 |
| Il6st | 653.41362 | 0.10047664 | 0.36502855 | 0.27525693 | 0.78311885 | 0.95081654 |
| Ddx4 | 33.2009449 | 0.21391323 | 1.34999259 | 0.15845511 | 0.87409819 | 0.97350615 |
| Plpp1 | 923.597985 | 0.1783325 | 0.22598418 | 0.78913709 | 0.43003189 | 0.81197362 |
| Gpx8 | 32.3642271 | 0.77990611 | 1.10464777 | 0.70602244 | 0.48017418 | 0.83935746 |
| Ndufs4 | 3205.88314 | 0.49128433 | 0.41562991 | 1.18202352 | 0.23719637 | 0.67338667 |

| | | | | | | |
|---|---|---|---|---|---|---|
| Kat6b | 70.8503891 | 1.29647952 | 0.71214441 | 1.82052895 | 0.06867849 | 0.40407556 |
| Samd8 | 811.136096 | 0.06939337 | 0.3644506 | 0.19040543 | 0.84899144 | 0.96765585 |
| Vdac2 | 11234.0575 | -0.1701917 | 0.14956699 | -1.1378965 | 0.25516372 | 0.68966367 |
| Nkiras1 | 116.098796 | 0.1045279 | 0.49492526 | 0.21119937 | 0.8327317 | 0.96565375 |
| Comtd1 | 640.617732 | -0.1877723 | 0.26145075 | -0.7181939 | 0.47263775 | 0.83483892 |
| Ube2e1 | 4899.20253 | 0.07020985 | 0.27431718 | 0.25594406 | 0.79799403 | 0.95433165 |
| Nr1d2 | 46.3832421 | 0.32882363 | 1.30671323 | 0.25164177 | 0.80131797 | 0.95615712 |
| Thrb | 33.4321371 | 2.88719162 | 1.29094628 | 2.23649246 | 0.02531953 | 0.23131726 |
| Ngly1 | 1697.16108 | 0.3277832 | 0.20095521 | 1.63112564 | 0.10286381 | 0.48985567 |
| Oxsm | 260.768923 | 0.8368141 | 0.48014308 | 1.74284319 | 0.08136101 | 0.43986918 |
| Prxl2a | 452.741108 | 0.62738173 | 0.28043687 | 2.23715846 | 0.02527598 | 0.23122295 |
| Glud1 | 11346.5877 | -0.0857153 | 0.16152167 | -0.5306738 | 0.59564483 | 0.88830374 |
| Bmpr1a | 215.62804 | -0.0196709 | 0.50116158 | -0.0392506 | 0.9686906 | 0.99443087 |
| Nid2 | 1443.7109 | 1.80704926 | 0.40082491 | 4.50832569 | 6.534121858 | 0.00036333 |
| Rtraf | 6273.62472 | 0.37183668 | 0.34116805 | 1.08989304 | 0.27576026 | 0.7118544 |
| Nudt13 | 374.367031 | 0.45759393 | 0.31999509 | 1.43000298 | 0.15271616 | 0.58090075 |
| Ecd | 5105.08589 | -0.3016495 | 0.18637855 | -1.6184775 | 0.10555972 | 0.49723223 |
| Dnajc9 | 5662.86744 | -0.1048854 | 0.2175318 | -0.4821612 | 0.62969147 | 0.90315579 |
| Anxa7 | 3942.23542 | 0.13928695 | 0.21642419 | 0.64358312 | 0.5198458 | 0.8576517 |
| Ppp3cb | 411.867022 | -0.5729589 | 0.38391521 | -1.4924101 | 0.13559166 | 0.55758574 |
| Zswim8 | 315.744774 | 0.07144722 | 0.47315432 | 0.15100194 | 0.87997419 | 0.97541902 |
| Camk2g | 1120.61242 | 0.0623444 | 0.25763925 | 0.24198331 | 0.8087931 | 0.95844313 |
| Plau | 604.664776 | 1.45542221 | 0.41557602 | 3.50218049 | 0.00046147 | 0.01267531 |
| Vcl | 433.383962 | -0.7057499 | 0.29454659 | -2.3960554 | 0.01657259 | 0.17972015 |
| Ap3m1 | 5711.47108 | 0.20612895 | 0.20935533 | 0.98458897 | 0.32482601 | 0.74621532 |
| Txndc16 | 758.490846 | -0.1111297 | 0.45505561 | -0.2442113 | 0.80706717 | 0.95778548 |
| Ero1l | 2596.38747 | -0.5713066 | 0.23609751 | -2.4197907 | 0.01552944 | 0.17285867 |
| Psmc6 | 9720.31999 | -0.0547965 | 0.23325796 | -0.2349179 | 0.81427245 | 0.95988497 |
| Bmp4 | 52.1307332 | 4.09240295 | 1.52010088 | 2.69219168 | 0.00709841 | 0.10284238 |
| Samd4 | 23.934566 | 2.2918468 | 1.46826991 | 1.56091655 | 0.11854344 | 0.52515149 |
| Mapk1ip1l | 3646.11748 | -0.0527552 | 0.25374255 | -0.2079083 | 0.83530056 | 0.96641606 |
| Ktn1 | 502.020041 | 0.13436075 | 0.33700843 | 0.39868663 | 0.69012412 | 0.92470267 |
| Peli2 | 89.7967443 | 0.50969161 | 0.59433207 | 0.85758725 | 0.39112042 | 0.79005237 |
| Anxa11 | 102.805231 | -0.3946005 | 0.46031246 | -0.8572448 | 0.39130962 | 0.79017671 |
| Tmem254b | 288.617751 | 0.04263862 | 0.3876875 | 0.10998192 | 0.91242371 | 0.98292561 |
| Ppif | 2750.56905 | -0.5228284 | 0.25313161 | -2.0654408 | 0.03888131 | 0.29821741 |
| Slmap | 431.252587 | 0.39079572 | 0.30142351 | 1.29650046 | 0.19480312 | 0.62694727 |
| Rnase10 | 28.5439857 | 2.04367561 | 1.80267475 | 1.1336907 | 0.25692431 | 0.69228086 |
| Rnase4 | 11617.9441 | 1.31172183 | 0.32065629 | 4.09074102 | 4.299970894 | 0.0017831 |
| Arf4 | 10150.6955 | 0.24372416 | 0.19356369 | 1.25914191 | 0.20797908 | 0.6418276 |
| Rnase6 | 289.751075 | -1.8446527 | 0.40813724 | -4.5196874 | 6.193102062 | 0.00034913 |
| Hacl1 | 405.477316 | 1.24477534 | 0.46486604 | 2.67770764 | 0.00741279 | 0.10532015 |
| Gpr65 | 2661.55509 | -0.0765484 | 0.26569046 | -0.288111 | 0.77326175 | 0.9464043 |
| Eaf1 | 1153.82422 | 0.0147627 | 0.31494282 | 0.04687422 | 0.96261348 | 0.99401755 |
| Mettl6 | 2464.18622 | 0.13703906 | 0.24334448 | 0.56314841 | 0.57333383 | 0.88046468 |
| Sh3bp5 | 1681.33943 | 0.91771023 | 0.29817253 | 3.07778265 | 0.00208547 | 0.04128082 |

| | | | | | | |
|---|---|---|---|---|---|---|
| Capn7 | 1836.56611 | -0.0372926 | 0.29435028 | -0.1266946 | 0.89918216 | 0.98010294 |
| Arhgef3 | 900.898911 | 0.05736211 | 0.27170608 | 0.21111825 | 0.832795 | 0.96565375 |
| Btd | 886.089935 | 0.3184082 | 0.20014777 | 1.59086561 | 0.11163982 | 0.50906326 |
| Bap1 | 4013.38181 | -0.1543563 | 0.18286679 | -0.8440912 | 0.39861843 | 0.7956109 |
| Phf7 | 558.769863 | -0.0547379 | 0.28008416 | -0.1954339 | 0.84505331 | 0.96765585 |
| Dph3 | 2085.59933 | 0.35157742 | 0.45540241 | 0.77201484 | 0.44010564 | 0.81963526 |
| Oxnad1 | 546.655127 | -0.1181718 | 0.32474658 | -0.3638894 | 0.71594059 | 0.93235217 |
| Nisch | 4247.23226 | 0.3429133 | 0.2441365 | 1.40459657 | 0.16014128 | 0.58970748 |
| Parg | 659.88682 | -0.1399413 | 0.32756035 | -0.427223 | 0.6692169 | 0.91643685 |
| Ogdhl | 25.9801689 | 2.4361207 | 1.38710239 | 1.75626595 | 0.07904301 | 0.43547225 |
| Glt8d1 | 534.121817 | 0.79779207 | 0.31617307 | 2.52327647 | 0.0116267 | 0.14194547 |
| Spcs1 | 4986.72828 | 0.43019069 | 0.30596054 | 1.40603326 | 0.15971425 | 0.58902465 |
| Nek4 | 787.994785 | -0.1463153 | 0.27072042 | -0.5404663 | 0.58887552 | 0.88690478 |
| Ebpl | 821.693163 | 0.41503902 | 0.25585357 | 1.62217405 | 0.10476609 | 0.49501875 |
| Kpna3 | 2380.97113 | -0.2272807 | 0.20324735 | -1.1182467 | 0.26346166 | 0.69987547 |
| Spryd7 | 744.536293 | 0.02433238 | 0.40875727 | 0.0595277 | 0.9525318 | 0.9914597 |
| Rnaseh2b | 3039.74878 | 0.24674755 | 0.28577005 | 0.86344788 | 0.38789126 | 0.7893695 |
| Mapk8 | 108.376124 | 0.01843947 | 0.63461774 | 0.02905603 | 0.97681991 | 0.99483019 |
| Pspc1 | 894.895251 | 0.0054953 | 0.35443543 | 0.01550437 | 0.9876298 | 0.9962376 |
| Ctsb | 119827.686 | 1.19628504 | 0.15930779 | 7.50926872 | 5.945856544 | 2.139284208 |
| Zmym2 | 354.139129 | 0.08551182 | 0.323282 | 0.26451155 | 0.79138577 | 0.95268772 |
| Cryl1 | 2232.29458 | 0.50950148 | 0.21313184 | 2.39054606 | 0.01682334 | 0.18128333 |
| Prkcd | 8341.59067 | -0.4331216 | 0.19982353 | -2.1675206 | 0.03019518 | 0.25957667 |
| Eef1akmt1 | 2245.19268 | 0.34124393 | 0.40491933 | 0.84274547 | 0.39937081 | 0.79612475 |
| Xpo4 | 628.946128 | -0.1999458 | 0.34427421 | -0.5807749 | 0.56139217 | 0.8752292 |
| Tkt | 17519.8471 | -0.4302961 | 0.14725932 | -2.9220299 | 0.00347758 | 0.06034221 |
| Pinx1 | 423.633663 | -0.5267672 | 0.27735388 | -1.8992604 | 0.05753025 | 0.36897087 |
| Lats2 | 334.713383 | 0.56538237 | 0.72988972 | 0.77461341 | 0.43856813 | 0.81933475 |
| Dcp1a | 387.74172 | -0.5416246 | 0.4837438 | -1.1196518 | 0.26286216 | 0.69888997 |
| Sap18 | 1446.43153 | -0.0132338 | 0.18886406 | -0.0700704 | 0.94413764 | 0.99005579 |
| Ska3 | 1607.453 | 0.0166531 | 0.17276768 | 0.09639014 | 0.92321073 | 0.98470981 |
| Mrpl57 | 2236.7839 | 0.34754944 | 0.46006523 | 0.75543513 | 0.44998794 | 0.82484234 |
| Zdhhc20 | 2886.32004 | 0.46579015 | 0.25144985 | 1.85241773 | 0.06396586 | 0.39085634 |
| Hmbox1 | 183.293424 | 0.39764838 | 0.44418078 | 0.89523993 | 0.37065884 | 0.7762831 |
| Micu2 | 3092.05986 | 0.11867839 | 0.1706592 | 0.69541165 | 0.48679736 | 0.841933 |
| Ints9 | 2869.38961 | -0.2356372 | 0.22125813 | -1.0649879 | 0.28688143 | 0.72260415 |
| Extl3 | 437.466269 | 0.42175369 | 0.38075213 | 1.10768569 | 0.26799758 | 0.70458078 |
| Cab39l | 4413.89915 | 0.17459144 | 0.26710795 | 0.65363625 | 0.51334618 | 0.85497712 |
| Cdadc1 | 877.255991 | 0.19375413 | 0.32460723 | 0.59688791 | 0.55058222 | 0.87093412 |
| Atp8a2 | 215.014866 | -4.6344629 | 0.82089533 | -5.6456198 | 1.645872553 | 2.031058461 |
| Mtmr6 | 2410.92224 | 0.14148815 | 0.16963598 | 0.83406923 | 0.40424197 | 0.79917613 |
| Spata13 | 268.045598 | -1.2185664 | 0.36573672 | -3.3318131 | 0.00086282 | 0.02081834 |
| Mipep | 1071.22498 | 0.16342158 | 0.41785394 | 0.39109737 | 0.69572527 | 0.92623267 |
| Esd | 5515.95983 | 0.4301669 | 0.33277532 | 1.29266466 | 0.19612707 | 0.62823315 |
| Lcp1 | 66548.1705 | -0.0346064 | 0.18398382 | -0.1880948 | 0.85080233 | 0.9679302 |
| Zc3h13 | 1136.21426 | -0.1794569 | 0.25471874 | -0.7045298 | 0.48110292 | 0.83974159 |

| | | | | | | |
|---|---|---|---|---|---|---|
| Slc25a30 | 359.856901 | -0.3735763 | 0.32168846 | -1.1612986 | 0.24552049 | 0.6807462 |
| Gpalpp1 | 854.270734 | 0.04916402 | 0.26336419 | 0.18667692 | 0.85191395 | 0.96798445 |
| Nufip1 | 2418.9757 | -0.1625897 | 0.2563499 | -0.6342491 | 0.52591827 | 0.86056156 |
| Tsc22d1 | 875.718799 | -0.092603 | 0.4694312 | -0.1972663 | 0.84361913 | 0.96765585 |
| Dnajc15 | 2480.27283 | 0.89519121 | 0.50285073 | 1.78023249 | 0.07503792 | 0.42184691 |
| Epsti1 | 150.43635 | -0.0531497 | 0.37071081 | -0.1433725 | 0.88599603 | 0.97793139 |
| Akap11 | 294.740226 | 0.34910972 | 0.3819679 | 0.9139766 | 0.36072914 | 0.76887042 |
| Rgcc | 136.388863 | -1.6264385 | 0.47946199 | -3.3922158 | 0.0006933 | 0.01763226 |
| Tdrd3 | 588.569874 | -0.0936028 | 0.24957156 | -0.3750539 | 0.70762035 | 0.92950127 |
| Naa16 | 860.664466 | -0.6686559 | 0.24327801 | -2.7485259 | 0.00598639 | 0.0915394 |
| Diaph3 | 198.175678 | 0.11929743 | 0.38589451 | 0.30914517 | 0.7572111 | 0.94116141 |
| Mtrf1 | 611.255075 | 0.14842583 | 0.29578203 | 0.50180813 | 0.61580249 | 0.89878056 |
| Wbp4 | 878.546016 | 0.11108601 | 0.20292106 | 0.54743459 | 0.5840802 | 0.88404713 |
| Sugt1 | 3675.04806 | 0.31333331 | 0.27503803 | 1.1392363 | 0.25460461 | 0.68860893 |
| Olfm4 | 1572.91631 | -4.7753434 | 0.25508738 | -18.720422 | 3.374655644 | 2.064108124 |
| Elp3 | 2695.95504 | -0.0355955 | 0.1654966 | -0.2150828 | 0.8297028 | 0.96462215 |
| Pbk | 3799.56029 | 0.13551467 | 0.24359154 | 0.55631929 | 0.5779926 | 0.8819488 |
| Esco2 | 1013.46559 | 0.12547069 | 0.23538349 | 0.53304796 | 0.59400038 | 0.88797037 |
| Ccdc25 | 1555.05655 | -0.232806 | 0.23716864 | -0.9816054 | 0.32629429 | 0.74762278 |
| Clu | 207.22739 | -1.0926579 | 0.44944011 | -2.4311535 | 0.01505084 | 0.16957556 |
| Trim35 | 4658.71943 | 0.05159265 | 0.22315662 | 0.23119481 | 0.81716346 | 0.96079676 |
| Dpysl2 | 28856.6528 | 0.11579054 | 0.23454457 | 0.49368247 | 0.62153044 | 0.89912494 |
| Bnip3l | 3953.68005 | 0.20020559 | 0.25969412 | 0.7709285 | 0.44074931 | 0.81965436 |
| Ppp2r2a | 5839.79572 | -0.2764457 | 0.21815001 | -1.2672277 | 0.20507388 | 0.63913241 |
| Nefm | 13.9862733 | 0.84725725 | 1.95832437 | 0.43264398 | 0.66527343 | 0.9149286 |
| Nefl | 17.8238243 | 2.64527291 | 1.94075812 | 1.36301009 | 0.17287933 | 0.60527557 |
| Pibf1 | 449.099416 | 0.27650511 | 0.29537258 | 0.93612315 | 0.34920978 | 0.76186861 |
| Bora | 1662.93236 | -0.3699631 | 0.1860948 | -1.9880354 | 0.04680778 | 0.33444287 |
| Tnfrsf10b | 64.8242645 | -0.5726484 | 0.54565307 | -1.0494734 | 0.2939603 | 0.72683372 |
| Rhobtb2 | 137.087371 | -0.1918543 | 0.48931291 | -0.3920891 | 0.69499236 | 0.92609254 |
| Bin3 | 3055.51217 | 0.03420157 | 0.24615723 | 0.13894197 | 0.88949601 | 0.97870754 |
| Pdlim2 | 312.890097 | -0.2585087 | 0.33990839 | -0.7605247 | 0.44694103 | 0.82254094 |
| Sorbs3 | 100.24294 | 0.76997447 | 0.70253116 | 1.09600046 | 0.27307857 | 0.71005689 |
| Ppp3cc | 24.6669493 | -1.7468498 | 1.3553251 | -1.2888788 | 0.19744023 | 0.63012948 |
| Slc39a14 | 249.520539 | 0.08500774 | 0.5337688 | 0.15925948 | 0.87346445 | 0.97344599 |
| Fam160b2 | 473.259084 | 0.02734263 | 0.46649229 | 0.05861324 | 0.95326016 | 0.99149988 |
| Bmp1 | 44.073128 | 0.23411261 | 1.79900614 | 0.13013441 | 0.89646008 | 0.97984623 |
| Xpo7 | 1468.61943 | -0.1965981 | 0.35015242 | -0.5614643 | 0.57448106 | 0.88059206 |
| Dok2 | 11865.2298 | 0.44597797 | 0.17456575 | 2.55478507 | 0.01062534 | 0.13331258 |
| Rb1 | 299.632718 | -1.2293717 | 0.33323275 | -3.6892284 | 0.00022494 | 0.00712858 |
| Rcbtb2 | 5892.39594 | 0.21956246 | 0.31524606 | 0.69647962 | 0.48612852 | 0.84183502 |
| Itm2b | 33378.3024 | 1.16707842 | 0.31206169 | 3.73989647 | 0.0001841 | 0.00608661 |
| Med4 | 2556.89106 | -0.0233773 | 0.18878102 | -0.1238328 | 0.90144764 | 0.98043702 |
| Sucla2 | 4480.06271 | -0.1734518 | 0.17317095 | -1.0016218 | 0.31652628 | 0.74186126 |
| Uchl3 | 3311.95447 | -0.2478073 | 0.19045984 | -1.3010998 | 0.1932243 | 0.624687 |
| Spry2 | 259.713291 | 0.6751651 | 0.382273 | 1.7661857 | 0.0773647 | 0.43008322 |

| | | | | | | |
|---|---|---|---|---|---|---|
| Rbm26 | 286.819575 | 0.30959972 | 0.46551811 | 0.66506483 | 0.50600903 | 0.85074333 |
| Rnf219 | 991.602563 | -0.4915673 | 0.22352991 | -2.1991121 | 0.02786995 | 0.24777117 |
| Ednrb | 13614.8006 | 1.35973617 | 0.29759201 | 4.5691286 | 4.897562127 | 0.00028394 |
| Scel | 135.827687 | -0.1147804 | 0.60697718 | -0.1891017 | 0.85001312 | 0.9679302 |
| Fbxl3 | 199.853264 | 0.22476147 | 0.3935296 | 0.57114247 | 0.56790307 | 0.87929304 |
| Cln5 | 4607.48269 | 0.20431479 | 0.21978953 | 0.92959293 | 0.35258189 | 0.76365692 |
| Acod1 | 31.4379021 | -0.8248902 | 1.64408822 | -0.5017311 | 0.61585668 | 0.89878056 |
| Dct | 33.5414 | -0.3605354 | 0.70513534 | -0.5112996 | 0.6091413 | 0.89466029 |
| Tgds | 1230.5974 | 0.39808423 | 0.35807602 | 1.11173105 | 0.2662538 | 0.70256315 |
| Gpr180 | 1237.83455 | 0.16404189 | 0.25178413 | 0.65151801 | 0.51471216 | 0.85526675 |
| Dnajc3 | 4265.32166 | 0.05570147 | 0.1741704 | 0.31981021 | 0.7491122 | 0.9399191 |
| Mbnl2 | 1821.9555 | -0.2233239 | 0.35868422 | -0.6226199 | 0.53353431 | 0.86492515 |
| Nipbl | 1248.50167 | -0.42536 | 0.3083672 | -1.3793946 | 0.16777314 | 0.60218592 |
| Nup155 | 1379.16946 | -0.0125934 | 0.32578071 | -0.0386561 | 0.96916458 | 0.99443087 |
| Fyb | 3854.40322 | 0.05669403 | 0.26297087 | 0.21559055 | 0.82930692 | 0.96462215 |
| Dab2 | 51422.8524 | 1.60687997 | 0.28199922 | 5.69817167 | 1.210990804 | 1.610222881 |
| Ttc33 | 1061.97798 | -0.0269335 | 0.24243746 | -0.1110946 | 0.91154132 | 0.98292561 |
| Rab2b | 438.070916 | 0.56929339 | 0.41220268 | 1.38110066 | 0.16724801 | 0.60121803 |
| Mettl3 | 1108.22513 | -0.1813501 | 0.18655612 | -0.9720939 | 0.33100382 | 0.75011761 |
| Dad1 | 4011.73342 | -0.0977821 | 0.16798335 | -0.5820942 | 0.56050322 | 0.87469256 |
| Lrp10 | 1692.06863 | -0.1265684 | 0.33818566 | -0.3742572 | 0.70821294 | 0.92967665 |
| Haus4 | 2900.12412 | -0.1273522 | 0.16716064 | -0.7618549 | 0.4461466 | 0.82166369 |
| Ajuba | 145.150495 | 0.90330923 | 0.65196623 | 1.38551537 | 0.16589494 | 0.5992759 |
| 4931414P19I | 98.5407561 | 0.66306757 | 0.53095664 | 1.2488168 | 0.21173209 | 0.64559288 |
| Slc7a8 | 6454.26552 | 0.7809251 | 0.34408431 | 2.26957487 | 0.02323339 | 0.21896307 |
| Fbxo4 | 85.3800581 | 0.58773176 | 0.56816644 | 1.03443588 | 0.30093244 | 0.73259831 |
| Acin1 | 754.997767 | -0.4520053 | 0.30383354 | -1.4876743 | 0.13683681 | 0.55946682 |
| Oxct1 | 2884.83814 | 0.06171768 | 0.29498279 | 0.2092247 | 0.83427283 | 0.96629899 |
| Drosha | 713.504036 | -0.3020662 | 0.27655821 | -1.0922338 | 0.27473033 | 0.71147204 |
| Psmb5 | 10758.7011 | 0.20515782 | 0.3737006 | 0.54898979 | 0.58301246 | 0.88343755 |
| Pabpn1 | 345.821111 | -0.4670856 | 0.30815861 | -1.5157311 | 0.12958736 | 0.54639717 |
| 6030458C11 | 1068.10926 | -0.0524051 | 0.21669432 | -0.2418387 | 0.80890519 | 0.95844313 |
| Pdzd2 | 30.3205309 | 2.96342418 | 1.72555481 | 1.71737471 | 0.08591073 | 0.44801003 |
| Slc22a17 | 70.1446162 | 0.03350057 | 0.59982573 | 0.0558505 | 0.95546091 | 0.99215259 |
| Golph3 | 1804.13903 | 0.26449729 | 0.45116917 | 0.58624858 | 0.55770848 | 0.87288227 |
| Zfr | 2783.03728 | -0.0726804 | 0.22513316 | -0.3228329 | 0.74682176 | 0.9398077 |
| Ngdn | 2908.96767 | -0.1070457 | 0.23161894 | -0.4621628 | 0.64396454 | 0.90823885 |
| Sub1 | 12152.6183 | -0.09659 | 0.28036185 | -0.3445192 | 0.73045588 | 0.93653187 |
| Dhrs4 | 899.909294 | 0.46980402 | 0.31065476 | 1.51230267 | 0.13045688 | 0.54803538 |
| Dcaf11 | 2024.39863 | -0.090087 | 0.18326247 | -0.4915736 | 0.62302079 | 0.90017327 |
| Fitm1 | 8.96189869 | 0.0706826 | 1.84898975 | 0.03822769 | 0.96950614 | 0.99443087 |
| Psme1 | 6793.23397 | -0.3078618 | 0.23689377 | -1.2995773 | 0.19374587 | 0.62518945 |
| Emc9 | 115.156898 | 0.72830249 | 0.49891866 | 1.45976198 | 0.1443555 | 0.57027792 |
| Tgm1 | 320.747418 | 0.40463373 | 0.40644398 | 0.99554612 | 0.31947072 | 0.7440831 |
| Cideb | 11.8286634 | -1.150987 | 1.34029664 | -0.8587554 | 0.39047547 | 0.7899826 |
| Adcy4 | 50.4810992 | 1.50648217 | 1.38166851 | 1.09033546 | 0.2755654 | 0.71181528 |

| | | | | | | |
|---|---|---|---|---|---|---|
| Ripk3 | 3386.1532 | -0.1842636 | 0.19530896 | -0.9434469 | 0.34545235 | 0.7602916 |
| Sdr39u1 | 93.7459149 | -0.3892646 | 0.68018929 | -0.5722887 | 0.56712644 | 0.87929304 |
| Zscan26 | 1438.39953 | 0.20793718 | 0.16868859 | 1.23266892 | 0.21769931 | 0.65150232 |
| Cct5 | 38376.9435 | -0.0398851 | 0.22713445 | -0.1756013 | 0.86060715 | 0.96994723 |
| Cmbl | 66.8853382 | 3.47591352 | 1.10900662 | 3.1342586 | 0.00172289 | 0.03578286 |
| Ropn1l | 40.0653718 | 1.40057766 | 0.82569522 | 1.69624048 | 0.08984035 | 0.45792373 |
| Ankrd33b | 18.5124957 | 1.52769997 | 1.23242662 | 1.23958696 | 0.21512821 | 0.64854028 |
| Tars | 7450.45318 | -0.4645107 | 0.17738389 | -2.6186749 | 0.0088272 | 0.11905611 |
| Amacr | 198.918541 | 0.99655426 | 0.39861754 | 2.50002611 | 0.01241842 | 0.148499 |
| Rai14 | 120.976125 | 0.34055669 | 0.62290795 | 0.54672073 | 0.58457061 | 0.88426515 |
| Brix1 | 4908.17376 | 0.05607753 | 0.27417458 | 0.20453219 | 0.83793764 | 0.96683245 |
| Rad1 | 810.983355 | -0.3493616 | 0.30588472 | -1.1421347 | 0.25339804 | 0.68662948 |
| Nadk2 | 254.211283 | 0.05508533 | 0.34893307 | 0.1578679 | 0.87456089 | 0.97365258 |
| Mtdh | 5355.44099 | -0.3642078 | 0.1960996 | -1.8572593 | 0.06327427 | 0.38779263 |
| Laptm4b | 745.283181 | 0.8717961 | 0.27028929 | 3.22541853 | 0.00125789 | 0.02752723 |
| Sdc2 | 9.0947118 | 1.5692751 | 2.15800075 | 0.72718932 | 0.46710999 | 0.8334534 |
| Trio | 195.969458 | 0.1553722 | 0.58332981 | 0.26635394 | 0.78996663 | 0.95255415 |
| Ank | 161.24213 | -1.7841571 | 0.71349676 | -2.500582 | 0.01239894 | 0.14841123 |
| Retreg1 | 292.337747 | 0.08915165 | 0.3382207 | 0.26359016 | 0.79209574 | 0.95297175 |
| Myo10 | 107.211286 | -0.5015508 | 0.60593339 | -0.8277325 | 0.40782198 | 0.80064439 |
| Rnf19a | 571.505913 | 0.3221858 | 0.39664659 | 0.81227422 | 0.4166343 | 0.8047849 |
| Pabpc1 | 8421.53783 | -0.7857802 | 0.18346118 | -4.2830869 | 1.843181679 | 0.00085424 |
| Ywhaz | 42534.751 | -0.2540851 | 0.14637138 | -1.7358932 | 0.08258272 | 0.44192232 |
| Grhl2 | 28.528024 | -0.0888237 | 1.31844739 | -0.0673699 | 0.94628719 | 0.99019648 |
| Rrm2b | 136.67028 | -0.2765901 | 0.653395 | -0.4233123 | 0.67206745 | 0.91736232 |
| Atp6v1c1 | 7407.92737 | 0.33330083 | 0.18714373 | 1.78098855 | 0.07491432 | 0.4213457 |
| Slc25a32 | 688.936348 | -0.3692966 | 0.22982448 | -1.6068634 | 0.10808433 | 0.50301538 |
| Dcaf13 | 4622.16029 | 0.03924727 | 0.25808796 | 0.15206934 | 0.87913224 | 0.97541065 |
| Dcstamp | 48.4146779 | -2.1237504 | 0.96708578 | -2.1960311 | 0.02808972 | 0.24918167 |
| Lrp12 | 303.027927 | 0.47146652 | 0.33956475 | 1.38844362 | 0.165002 | 0.59753389 |
| Oxr1 | 930.256316 | 0.64600124 | 0.32809187 | 1.96896447 | 0.04895717 | 0.34113306 |
| Angpt1 | 21.8013163 | -0.9346403 | 1.05584521 | -0.8852058 | 0.37604564 | 0.77836994 |
| Eif3h | 12218.5954 | 0.38663376 | 0.41326376 | 0.93556173 | 0.34949889 | 0.76197111 |
| Utp23 | 874.716429 | -0.1256058 | 0.22875349 | -0.549088 | 0.58294509 | 0.88343755 |
| Rad21 | 1371.00018 | 0.02282525 | 0.25713596 | 0.08876725 | 0.92926689 | 0.98669576 |
| Shcbp1 | 5748.42498 | 0.12318692 | 0.21177154 | 0.58169725 | 0.56077062 | 0.87469256 |
| Rida | 541.344168 | 0.39132593 | 0.37496464 | 1.04363421 | 0.29665466 | 0.72954201 |
| Matn2 | 118.039739 | 0.53395043 | 0.62801063 | 0.85022515 | 0.39519992 | 0.79310606 |
| Pop1 | 644.371767 | -0.6181746 | 0.40490512 | -1.5267146 | 0.12683198 | 0.54111158 |
| Stk3 | 1492.94762 | 0.18230211 | 0.17989356 | 1.01338875 | 0.31087451 | 0.74011268 |
| Khdrbs3 | 15.117386 | 0.28150166 | 1.0568692 | 0.2663543 | 0.78996635 | 0.95255415 |
| Zfat | 370.276279 | 0.1162985 | 0.33878439 | 0.34328176 | 0.7313865 | 0.93653187 |
| Eif3e | 12434.0967 | -0.0474442 | 0.28770018 | -0.1649085 | 0.86901601 | 0.97208054 |
| Emc2 | 3172.73236 | 0.21364727 | 0.24196778 | 0.88295752 | 0.37725922 | 0.7799581 |
| Eny2 | 1938.22826 | 0.12413145 | 0.33942374 | 0.36571235 | 0.71457973 | 0.93235217 |
| Ebag9 | 492.538853 | 0.34841499 | 0.36037082 | 0.96682353 | 0.33363226 | 0.75315066 |

| | | | | | | |
|---|---|---|---|---|---|---|
| Myc | 4209.19277 | -0.3090165 | 0.29190608 | -1.0586163 | 0.28977455 | 0.72395811 |
| Washc5 | 3331.00976 | 0.40557934 | 0.34290583 | 1.1827718 | 0.2368996 | 0.6732629 |
| Sqle | 4239.51518 | 0.29458901 | 0.27072366 | 1.08815391 | 0.27652717 | 0.7118544 |
| Mtss1 | 891.897169 | 1.50106137 | 0.40866576 | 3.67307838 | 0.00023965 | 0.00747854 |
| Ndufb9 | 7530.64667 | 0.39956896 | 0.42175291 | 0.9474006 | 0.34343468 | 0.75947281 |
| Fbxo32 | 85.2528622 | 0.5206615 | 0.92122924 | 0.56518126 | 0.57195048 | 0.88035336 |
| Wdyhv1 | 515.320552 | 0.11649562 | 0.29931732 | 0.38920443 | 0.69712493 | 0.92650558 |
| Atad2 | 1906.33261 | -0.1885407 | 0.25925483 | -0.7272408 | 0.46707848 | 0.8334534 |
| Zhx1 | 1789.31146 | 0.04243415 | 0.27851228 | 0.15236008 | 0.87890294 | 0.97535404 |
| Gm29394 | 32.2116552 | 0.97920973 | 0.95739551 | 1.02278496 | 0.30640953 | 0.73582655 |
| Tbc1d31 | 1155.59363 | -0.4700855 | 0.3600223 | -1.3057121 | 0.19165048 | 0.62272506 |
| Derl1 | 8011.36273 | 0.09892401 | 0.14811395 | 0.66789121 | 0.50420304 | 0.85012672 |
| Mtbp | 647.814466 | -0.1792068 | 0.28398055 | -0.6310532 | 0.52800573 | 0.86198732 |
| Mrpl13 | 2528.22781 | 0.47083858 | 0.43059295 | 1.0934656 | 0.27418943 | 0.71138054 |
| Col14a1 | 59.715507 | 1.86342726 | 0.68756311 | 2.71019088 | 0.00672445 | 0.09958861 |
| Sla | 8032.3762 | 0.21247485 | 0.22661479 | 0.93760362 | 0.34844815 | 0.76130849 |
| Asap1 | 1567.47324 | -0.1093037 | 0.18558815 | -0.5889585 | 0.55588913 | 0.87260256 |
| Fam49b | 9583.59052 | -0.1598321 | 0.14769707 | -1.0821614 | 0.27918081 | 0.71377266 |
| Gtse1 | 1154.77988 | -0.4871903 | 0.28773379 | -1.6931982 | 0.09041775 | 0.45983389 |
| Trmu | 547.817598 | -0.4744734 | 0.24260485 | -1.9557458 | 0.05049511 | 0.34374326 |
| Brd1 | 671.405353 | 0.05829172 | 0.29785812 | 0.19570298 | 0.84484266 | 0.96765585 |
| Tef | 141.88676 | 0.77063175 | 0.59462018 | 1.29600673 | 0.19497317 | 0.62699967 |
| Zc3h7b | 139.980206 | 0.0903282 | 0.59758561 | 0.15115525 | 0.87985325 | 0.97541902 |
| Rangap1 | 9750.54678 | -0.2302438 | 0.18304591 | -1.2578474 | 0.20844695 | 0.6418276 |
| L3mbtl2 | 2497.97075 | -0.3273812 | 0.26788213 | -1.2221093 | 0.22166629 | 0.65514465 |
| Rbx1 | 9378.83036 | 0.42405747 | 0.38685381 | 1.09616982 | 0.27300446 | 0.71005689 |
| Xpnpep3 | 751.249502 | -0.2082219 | 0.32420875 | -0.6422465 | 0.52071316 | 0.8576517 |
| St13 | 20802.5718 | 0.03403017 | 0.16600308 | 0.2049972 | 0.83757431 | 0.96683245 |
| Slc25a17 | 2355.87916 | 0.03072868 | 0.24019723 | 0.12793102 | 0.89820356 | 0.97989497 |
| Adsl | 9018.64125 | -0.2641864 | 0.20257258 | -1.3041566 | 0.19218019 | 0.62359158 |
| Fam83f | 22.0011443 | -1.8651452 | 1.89251842 | -0.9855361 | 0.32436082 | 0.74596967 |
| Mief1 | 945.168619 | -0.4004044 | 0.26947478 | -1.4858695 | 0.13731363 | 0.55981285 |
| Tab1 | 1174.51028 | -0.1628182 | 0.27319122 | -0.595986 | 0.55118455 | 0.87093412 |
| Syngr1 | 93.0415473 | 1.14458623 | 0.53272654 | 2.14854366 | 0.03167059 | 0.26539272 |
| Deptor | 450.370746 | 0.57448703 | 0.4231605 | 1.35761025 | 0.17458739 | 0.60656845 |
| Dnal4 | 912.032454 | -0.0053693 | 0.1870669 | -0.0287027 | 0.97710174 | 0.99483019 |
| Dscc1 | 978.003216 | 0.0636753 | 0.32013108 | 0.19890383 | 0.84233797 | 0.96765585 |
| Enpp2 | 98.698279 | 2.27255744 | 1.10188088 | 2.06243478 | 0.03916636 | 0.29945131 |
| Josd1 | 297.260259 | 0.40344897 | 0.41717171 | 0.96710529 | 0.3334914 | 0.75297164 |
| Tomm22 | 5877.48679 | 0.06756085 | 0.29955219 | 0.2255395 | 0.8215596 | 0.96231102 |
| Cby1 | 751.465869 | 0.20966314 | 0.3904867 | 0.53692773 | 0.59131754 | 0.88792087 |
| Dmc1 | 7.76044329 | 1.73916985 | 2.20759511 | 0.78781197 | 0.4308067 | 0.81254302 |
| Csnk1e | 537.856382 | -0.2058311 | 0.3089366 | -0.6662566 | 0.50524709 | 0.85034006 |
| Fam118a | 1252.18692 | 0.56850927 | 0.22237891 | 2.55648921 | 0.01057344 | 0.13320786 |
| Sh3bp1 | 3033.23867 | -0.2046616 | 0.23250254 | -0.8802552 | 0.3787211 | 0.78126394 |
| Samm50 | 6539.22019 | -0.0006249 | 0.17167339 | -0.0036403 | 0.9970955 | 0.9994418 |

| | | | | | | |
|---|---|---|---|---|---|---|
| Parvb | 27.7791776 | -0.2670118 | 0.84937757 | -0.3143618 | 0.75324629 | 0.94018119 |
| Parvg | 6418.0741 | -0.9167947 | 0.1768342 | -5.1844873 | 2.166098865 | 1.948374075 |
| C1qtnf6 | 63.6335113 | 0.79520722 | 0.75583796 | 1.05208691 | 0.29275968 | 0.72584701 |
| Ttll1 | 498.408315 | 0.40405371 | 0.33151236 | 1.21881945 | 0.22291273 | 0.65613364 |
| Myh9 | 992.921932 | -0.6884797 | 0.32568217 | -2.1139618 | 0.03451853 | 0.27856237 |
| Ndufa6 | 2287.27007 | 0.06401928 | 0.33751735 | 0.18967702 | 0.84956223 | 0.96766052 |
| Twf1 | 12440.2384 | 1.05082228 | 0.14642625 | 7.17646121 | 7.153893574 | 2.243937951 |
| Smdt1 | 3999.72474 | 0.18815094 | 0.31603995 | 0.59533911 | 0.55161682 | 0.87093412 |
| Naga | 1525.27549 | -0.0004923 | 0.19327859 | -0.0025473 | 0.99796753 | 0.99967651 |
| Slc38a2 | 3479.24855 | -0.1091986 | 0.40415337 | -0.270191 | 0.78701334 | 0.95223213 |
| Srebf2 | 1236.04902 | 0.09242307 | 0.20878755 | 0.44266563 | 0.65800761 | 0.91258818 |
| Slc38a4 | 298.08057 | 0.93326499 | 0.28736902 | 3.24761866 | 0.00116375 | 0.02636328 |
| Rpap3 | 907.425937 | -0.0207411 | 0.2090863 | -0.0991989 | 0.92098035 | 0.9839023 |
| Rapgef3 | 67.0701268 | -0.2524958 | 0.63323507 | -0.3987395 | 0.69008519 | 0.92470267 |
| Xrcc6 | 3872.18366 | -0.2705539 | 0.20195193 | -1.3396946 | 0.18034466 | 0.61122544 |
| Desi1 | 1292.02492 | -0.442896 | 0.21180052 | -2.0910996 | 0.03651913 | 0.28679157 |
| Pmm1 | 513.443879 | -0.5160679 | 0.21491576 | -2.4012567 | 0.01633887 | 0.17814028 |
| Hdac7 | 43.6144753 | -0.4541979 | 0.80278358 | -0.5657788 | 0.57154415 | 0.88035336 |
| Polr3h | 2583.31209 | -0.1647953 | 0.25303892 | -0.6512647 | 0.51487561 | 0.85532175 |
| Aco2 | 12160.6434 | -0.1172568 | 0.17025123 | -0.6887283 | 0.49099428 | 0.84477968 |
| Vdr | 151.952725 | -1.5194865 | 0.42696424 | -3.5588143 | 0.00037253 | 0.01059812 |
| Nckap1l | 11649.1759 | 0.10775885 | 0.24270343 | 0.44399393 | 0.65704697 | 0.91251438 |
| Pde1b | 23.2168864 | -0.4609718 | 1.43682834 | -0.320826 | 0.74834228 | 0.9399191 |
| Txndc11 | 2013.03326 | 0.01246218 | 0.45295667 | 0.02751295 | 0.97805061 | 0.99483019 |
| Litaf | 18845.7082 | 0.46644828 | 0.25837876 | 1.80528884 | 0.07102949 | 0.41122734 |
| Nubp1 | 3859.77236 | 0.14183036 | 0.29548939 | 0.47998459 | 0.63123835 | 0.90394609 |
| 1810013L24I | 1112.98902 | -0.0367181 | 0.34417942 | -0.1066831 | 0.91504042 | 0.98347559 |
| Bcl6 | 759.29235 | 0.0758538 | 0.66319122 | 0.11437697 | 0.90893897 | 0.98209583 |
| Il1rap | 259.176699 | -0.2956995 | 0.32176555 | -0.9189905 | 0.35810054 | 0.76676242 |
| Anks3 | 633.55875 | 0.07330717 | 0.20006759 | 0.36641205 | 0.71405763 | 0.93235217 |
| Nudt16l1 | 5002.34481 | 0.34797034 | 0.26595602 | 1.30837549 | 0.19074598 | 0.62174143 |
| Mgrn1 | 2682.75154 | 0.11165917 | 0.1827029 | 0.61115161 | 0.54109922 | 0.86776854 |
| Srl | 185.488959 | -0.1945664 | 0.47544226 | -0.4092325 | 0.68236901 | 0.92104707 |
| Crebbp | 1530.00506 | 0.22476098 | 0.28019653 | 0.80215477 | 0.42246343 | 0.80821275 |
| Zfp251 | 435.928738 | -0.3639526 | 0.23681461 | -1.536867 | 0.12432587 | 0.53570919 |
| Hes1 | 225.974726 | 1.33888091 | 0.3953245 | 3.3867896 | 0.00070716 | 0.01787322 |
| Zfp263 | 1403.23309 | 0.08927497 | 0.23170909 | 0.38528902 | 0.7000233 | 0.92669304 |
| Atp13a3 | 439.919732 | 0.33042697 | 0.46338222 | 0.7130765 | 0.47579842 | 0.83639059 |
| Mefv | 18.5097147 | -1.5998061 | 2.4182771 | -0.6615479 | 0.50826102 | 0.85238774 |
| Glyr1 | 6996.06072 | -0.1992853 | 0.20120057 | -0.9904807 | 0.32193923 | 0.74447686 |
| Tmem44 | 37.2646308 | 3.35872496 | 1.00877192 | 3.32951868 | 0.00086996 | 0.02090881 |
| Lsg1 | 2933.79159 | -0.2941469 | 0.22450872 | -1.3101803 | 0.19013486 | 0.62090756 |
| Rogdi | 7545.12463 | -0.5814986 | 0.1643952 | -3.5371995 | 0.00040439 | 0.0113723 |
| 4930451G09 | 15.4775811 | -1.9663106 | 1.95401968 | -1.0062901 | 0.31427604 | 0.74065408 |
| Eef2kmt | 1695.68761 | 0.03785772 | 0.29046547 | 0.13033468 | 0.89630164 | 0.97984623 |
| Ercc4 | 554.661005 | -0.2787031 | 0.35666553 | -0.781413 | 0.43455965 | 0.81552669 |

| | | | | | | |
|---|---|---|---|---|---|---|
| Gpt | 122.882975 | 1.17281842 | 0.71734341 | 1.63494694 | 0.10206017 | 0.48845074 |
| Adck5 | 795.770791 | 0.69661292 | 0.34442895 | 2.0225156 | 0.04312311 | 0.31893896 |
| Cyc1 | 11481.2733 | -0.0176291 | 0.25473089 | -0.0692068 | 0.94482497 | 0.99019648 |
| Sharpin | 1097.4613 | 0.00096468 | 0.20131272 | 0.00479193 | 0.99617661 | 0.99887119 |
| Maf1 | 3706.86022 | 0.21208088 | 0.22644961 | 0.93654781 | 0.34899121 | 0.7617085 |
| Hgh1 | 660.734651 | -0.8235014 | 0.28512044 | -2.888258 | 0.00387382 | 0.06536336 |
| Dgat1 | 557.26091 | -0.0630002 | 0.29526776 | -0.2133663 | 0.83104128 | 0.96466657 |
| Hsf1 | 1920.10179 | -0.3675237 | 0.26560063 | -1.3837454 | 0.16643641 | 0.59971036 |
| Bop1 | 5318.24732 | -0.6364041 | 0.19771847 | -3.2187387 | 0.00128756 | 0.02802615 |
| Mroh1 | 963.434825 | 0.51563436 | 0.37645517 | 1.36970988 | 0.17077748 | 0.60430173 |
| Fbxl6 | 1300.03514 | -0.3179145 | 0.19386789 | -1.6398514 | 0.10103608 | 0.48583896 |
| Slc52a2 | 981.704789 | -0.2530007 | 0.19432042 | -1.301977 | 0.19292425 | 0.62468035 |
| Gpaa1 | 3136.60285 | -0.2169019 | 0.19175557 | -1.1311376 | 0.25799717 | 0.69349141 |
| Oplah | 213.006203 | 1.02597158 | 0.33682562 | 3.04600215 | 0.00231906 | 0.04446565 |
| Grina | 838.338994 | 0.59479521 | 0.33778371 | 1.76087596 | 0.0782594 | 0.43258005 |
| Plec | 1103.25048 | 0.31995134 | 0.46850567 | 0.68291882 | 0.49465814 | 0.84678075 |
| Scrib | 190.203817 | -0.4416249 | 0.36016656 | -1.2261684 | 0.22013529 | 0.65346155 |
| Tsta3 | 3253.62153 | -0.0009865 | 0.20031816 | -0.0049248 | 0.99607059 | 0.99884675 |
| Pycrl | 3145.18908 | 0.0607245 | 0.2568121 | 0.23645496 | 0.81307965 | 0.95979961 |
| Naprt | 94.5061203 | 1.05226933 | 0.7377577 | 1.4263075 | 0.1537796 | 0.58242478 |
| Gsdmd | 985.486766 | -0.6438531 | 0.27065504 | -2.3788699 | 0.01736581 | 0.18521003 |
| Ly6c2 | 32.8049704 | -7.2408902 | 1.64267265 | -4.4079934 | 1.043327290 | 0.00054311 |
| Ly6e | 6180.23394 | -1.0839253 | 0.18845518 | -5.7516344 | 8.838479731 | 1.316607998 |
| Zbtb11 | 681.843893 | -0.5531604 | 0.27985902 | -1.9765681 | 0.04809047 | 0.3379039 |
| Arc | 72.8864645 | 2.1699261 | 1.50170576 | 1.44497421 | 0.14846513 | 0.5754118 |
| Cep97 | 163.83977 | -0.1310163 | 0.59800936 | -0.2190873 | 0.82658205 | 0.96370073 |
| Ptk2 | 487.049429 | 0.65480592 | 0.37081508 | 1.76585569 | 0.07742006 | 0.43008322 |
| Lmf2 | 1912.64267 | -0.0105949 | 0.29761202 | -0.0355998 | 0.97160145 | 0.99443087 |
| Tymp | 49.1877712 | -0.118397 | 0.53766896 | -0.2202043 | 0.82571202 | 0.96326388 |
| Chkb | 892.784359 | 0.6726166 | 0.26073611 | 2.57968334 | 0.00988909 | 0.12828558 |
| Arsa | 932.374521 | 0.44261671 | 0.40627028 | 1.08946366 | 0.27594947 | 0.7118544 |
| Rabl2 | 758.378839 | 0.80472969 | 0.37450171 | 2.14880112 | 0.03165017 | 0.26539272 |
| Kif21a | 19.6010435 | -0.5696167 | 1.19115029 | -0.4782073 | 0.63250268 | 0.90423307 |
| Yaf2 | 1295.76422 | 0.44385206 | 0.25326987 | 1.75248662 | 0.07969017 | 0.43676068 |
| Zcrb1 | 1262.05863 | 0.04488802 | 0.32220682 | 0.1393143 | 0.88920179 | 0.97870754 |
| Alcam | 13.8778729 | -0.6166409 | 1.93851553 | -0.3180995 | 0.75040944 | 0.9399191 |
| Cblb | 216.137277 | -0.0084617 | 0.4641324 | -0.0182312 | 0.98545441 | 0.99564261 |
| Bbx | 905.919412 | 0.54621018 | 0.36919271 | 1.47947173 | 0.13901428 | 0.56272724 |
| Nectin3 | 57.508999 | -0.0278569 | 0.74057607 | -0.0376151 | 0.96999454 | 0.99443087 |
| Atg3 | 4108.2911 | -0.162102 | 0.19835203 | -0.8172441 | 0.41378896 | 0.80395688 |
| Slc35a5 | 919.325168 | 0.60835833 | 0.40315982 | 1.5089756 | 0.13130502 | 0.54970237 |
| Ccdc80 | 32.9303799 | 1.94855531 | 1.45551867 | 1.33873605 | 0.18065662 | 0.61122544 |
| Cd200r1 | 855.520861 | 0.09764184 | 0.28848701 | 0.33846181 | 0.7350152 | 0.93750666 |
| Gtpbp8 | 946.768891 | 0.50900848 | 0.38710253 | 1.31491901 | 0.18853712 | 0.61935434 |
| Mzt2 | 384.70119 | 0.00595567 | 0.23737958 | 0.02508922 | 0.9799838 | 0.99486654 |
| Prkdc | 301.658555 | 0.02733453 | 0.35844047 | 0.0762596 | 0.93921257 | 0.98913801 |

| | | | | | | |
|---|---|---|---|---|---|---|
| Mcm4 | 14509.9452 | -0.1886058 | 0.23020933 | -0.8192795 | 0.41262699 | 0.80261821 |
| Ube2v2 | 5030.46867 | 0.25333831 | 0.36517304 | 0.69374867 | 0.48783984 | 0.84313956 |
| Snai2 | 39.0690072 | -0.3898477 | 1.69979075 | -0.2293504 | 0.81859657 | 0.96105733 |
| Fopnl | 2607.74745 | 0.75837678 | 0.44865193 | 1.69034554 | 0.09096187 | 0.46056976 |
| Nde1 | 3336.42647 | 0.02621053 | 0.14765616 | 0.1775106 | 0.85910734 | 0.96954641 |
| Mpv17l | 66.0967634 | 0.11695613 | 0.74131807 | 0.15776781 | 0.87463977 | 0.97365258 |
| Pdxdc1 | 3248.90779 | -0.119633 | 0.19706908 | -0.6070614 | 0.54381019 | 0.86947682 |
| Ntan1 | 2127.33172 | 0.33144032 | 0.32244469 | 1.02789822 | 0.30399771 | 0.73436097 |
| Rrn3 | 4327.74608 | -0.3798367 | 0.1919677 | -1.978649 | 0.04785554 | 0.33761064 |
| Bfar | 2419.95526 | 0.25358867 | 0.2573712 | 0.98530321 | 0.32447516 | 0.74596967 |
| Parn | 2909.62558 | -0.0046562 | 0.1958736 | -0.0237717 | 0.98103473 | 0.99522729 |
| B3gnt5 | 23.1970125 | 0.11547256 | 1.59676193 | 0.07231671 | 0.94234987 | 0.98976268 |
| Naa50 | 9542.11261 | -0.2160105 | 0.25823342 | -0.8364933 | 0.40287742 | 0.79795242 |
| Ccdc191 | 113.504651 | 0.77092617 | 0.52501457 | 1.46839005 | 0.1419983 | 0.56729759 |
| Hira | 369.275572 | -0.6228893 | 0.35666635 | -1.7464201 | 0.08073797 | 0.43854735 |
| Qtrt2 | 411.47712 | -0.4546472 | 0.27037855 | -1.6815211 | 0.09266174 | 0.46597694 |
| Mrpl40 | 3569.44612 | 0.09216495 | 0.28369664 | 0.32487149 | 0.74527834 | 0.93917909 |
| Gbe1 | 1187.20192 | -0.1056839 | 0.2537236 | -0.4165315 | 0.67702112 | 0.91950697 |
| Zbtb20 | 74.6685391 | 0.24646177 | 0.81062782 | 0.30403814 | 0.76109884 | 0.94245593 |
| Usp7 | 990.38811 | -0.2699897 | 0.2092629 | -1.2901939 | 0.19698333 | 0.62932233 |
| Pmm2 | 3126.5391 | 0.08715263 | 0.18259723 | 0.47729434 | 0.63315254 | 0.90434445 |
| Dgcr8 | 1172.75667 | -0.1541305 | 0.22242037 | -0.6929694 | 0.48832875 | 0.84338918 |
| Trmt2a | 7114.73978 | -0.1168768 | 0.15950041 | -0.7327677 | 0.4637001 | 0.83202414 |
| Arl6 | 297.058637 | 0.67283114 | 0.50015886 | 1.34523488 | 0.1785494 | 0.60992118 |
| Crybg3 | 337.369307 | -0.4477837 | 0.34054651 | -1.3148974 | 0.18854438 | 0.61935434 |
| Riox2 | 2306.22033 | -0.4281813 | 0.18810954 | -2.2762338 | 0.02283202 | 0.2163471 |
| Cpox | 300.564885 | 0.15856668 | 0.45109598 | 0.35151428 | 0.72520256 | 0.93542036 |
| Cldnd1 | 2933.52735 | -0.0234244 | 0.18647932 | -0.125614 | 0.90003751 | 0.98043702 |
| St3gal6 | 2050.12208 | 1.11507017 | 0.28768278 | 3.87604066 | 0.00010617 | 0.00377551 |
| Cmss1 | 809.518496 | 0.05148946 | 0.32456678 | 0.15864056 | 0.87395207 | 0.97350615 |
| Tbc1d23 | 805.966183 | 0.29153862 | 0.25833885 | 1.12851251 | 0.25910353 | 0.69546977 |
| Klhl22 | 579.05043 | 0.38479355 | 0.33705824 | 1.14162331 | 0.25361063 | 0.6869838 |
| Nit2 | 2371.51714 | 0.52264154 | 0.36795983 | 1.4203766 | 0.15549807 | 0.58452503 |
| Tomm70a | 3779.12664 | -0.1261399 | 0.46307029 | -0.272399 | 0.78531523 | 0.95163558 |
| Tfg | 4698.49459 | -0.020503 | 0.16774742 | -0.1222256 | 0.90272035 | 0.98043702 |
| Thap7 | 616.206378 | -0.098635 | 0.3012626 | -0.3274055 | 0.74336117 | 0.93856621 |
| Lztr1 | 1925.0773 | 0.36289618 | 0.27706975 | 1.30976469 | 0.19027545 | 0.6209089 |
| Snap29 | 815.925796 | 0.45564424 | 0.22320857 | 2.04133844 | 0.0412172 | 0.30914162 |
| Sdf2l1 | 9233.10951 | 0.22194984 | 0.26651934 | 0.83277197 | 0.40497335 | 0.79917613 |
| Dlg1 | 180.978567 | -0.1673831 | 0.51698699 | -0.3237665 | 0.74611481 | 0.93959466 |
| Ppil2 | 5522.62315 | 0.40757293 | 0.25496465 | 1.59854679 | 0.10992134 | 0.50665703 |
| Senp5 | 605.635449 | 0.27474404 | 0.22147098 | 1.24054194 | 0.21477501 | 0.64854028 |
| Ypel1 | 42.1431056 | 1.61555687 | 1.38699464 | 1.16478956 | 0.24410419 | 0.67912815 |
| Ncbp2 | 2533.83311 | -0.0855409 | 0.18822101 | -0.4544705 | 0.6494902 | 0.90973357 |
| Top3b | 1465.22309 | -0.3587279 | 0.22753691 | -1.57657 | 0.1148945 | 0.51833667 |
| Pak2 | 4124.47611 | -0.0053592 | 0.2718748 | -0.0197121 | 0.98427301 | 0.99564261 |

| | | | | | | |
|---|---|---|---|---|---|---|
| Wdr53 | 421.126445 | 0.2519362 | 0.29192732 | 0.86301003 | 0.38813195 | 0.78942657 |
| Fgd4 | 1200.68456 | -0.155237 | 0.30386388 | -0.5108767 | 0.60943737 | 0.89498768 |
| Dnm1l | 3022.7798 | -0.1278079 | 0.24935135 | -0.5125615 | 0.60825811 | 0.89389974 |
| Tnk2 | 48.0617049 | -0.2206144 | 0.65964919 | -0.334442 | 0.73804604 | 0.93800224 |
| Yars2 | 1808.14106 | -0.007313 | 0.27758821 | -0.0263448 | 0.97898236 | 0.99483019 |
| B4galt4 | 462.11573 | 0.49865281 | 0.36455243 | 1.36784937 | 0.17135923 | 0.60430173 |
| Tfrc | 1444.0115 | 0.53123694 | 0.20494186 | 2.59213482 | 0.00953824 | 0.12534914 |
| Arhgap31 | 482.78588 | 0.50598688 | 0.41443383 | 1.22091114 | 0.22211967 | 0.65543602 |
| Fyttd1 | 3299.95394 | 0.26513833 | 0.25353924 | 1.04574868 | 0.29567708 | 0.72852118 |
| Lrch3 | 286.521437 | -0.0275553 | 0.3032723 | -0.09086 | 0.92760386 | 0.98666423 |
| Lmln | 135.07356 | -0.4227828 | 0.75364019 | -0.5609875 | 0.57480603 | 0.88059206 |
| Osbpl11 | 3147.27859 | 0.96941364 | 0.29968045 | 3.23482444 | 0.00121718 | 0.02712151 |
| Snx4 | 6346.24779 | -0.160718 | 0.23895167 | -0.6725963 | 0.50120416 | 0.84968141 |
| Zfp148 | 1289.23026 | 0.22754652 | 0.26327533 | 0.86429107 | 0.38742801 | 0.78911204 |
| Gsk3b | 4406.23422 | 0.33480853 | 0.31363815 | 1.06749939 | 0.28574639 | 0.721488 |
| Umps | 9790.2645 | -0.2468731 | 0.18245042 | -1.353097 | 0.17602465 | 0.60763832 |
| Fstl1 | 173.799511 | -0.0602839 | 0.72702844 | -0.0829182 | 0.93391659 | 0.98756396 |
| Itgb5 | 1161.80063 | 0.09359461 | 0.26821474 | 0.34895403 | 0.72712382 | 0.93576253 |
| Ndufb4 | 1414.33734 | 0.66484525 | 0.48313161 | 1.37611622 | 0.16878565 | 0.60312169 |
| Abcc5 | 262.103404 | -0.0707716 | 0.37959031 | -0.1864419 | 0.85209821 | 0.96798445 |
| Rabl3 | 1210.84683 | 0.18543856 | 0.26557715 | 0.69824741 | 0.48502248 | 0.84171939 |
| Gtf2e1 | 1489.01946 | -0.2930816 | 0.23516776 | -1.2462659 | 0.21266679 | 0.64596443 |
| Hcls1 | 5397.78302 | -0.2513621 | 0.1782391 | -1.4102524 | 0.15846516 | 0.58731982 |
| Ccdc14 | 455.186749 | -0.068069 | 0.3483602 | -0.1953983 | 0.84508116 | 0.96765585 |
| Mylk | 38.3246949 | 4.66539152 | 1.64538745 | 2.83543642 | 0.00457631 | 0.07434532 |
| Iqcb1 | 409.202944 | 0.63412423 | 0.34321305 | 1.84761108 | 0.06465863 | 0.39234572 |
| Ap2m1 | 29179.8085 | 0.75793069 | 0.15076119 | 5.02735945 | 4.972797192 | 3.950144678 |
| Pdia5 | 762.937033 | 0.21193508 | 0.30620739 | 0.6921292 | 0.4888562 | 0.84370455 |
| Slc49a4 | 92.6448053 | -0.6112481 | 0.79330091 | -0.7705123 | 0.44099609 | 0.81967485 |
| Hspbap1 | 523.613035 | -0.9390347 | 0.33623774 | -2.79277 | 0.00522588 | 0.0828086 |
| Ehhadh | 233.069645 | 1.13537756 | 0.47459568 | 2.39230488 | 0.01674293 | 0.18093307 |
| Senp2 | 1555.66842 | -0.1520459 | 0.25202823 | -0.6032892 | 0.5463163 | 0.86994632 |
| Tmem41a | 759.172235 | -0.4404574 | 0.20835971 | -2.1139279 | 0.03452142 | 0.27856237 |
| Tra2b | 16914.2398 | -0.108059 | 0.20061925 | -0.538627 | 0.59014422 | 0.88731984 |
| Dgkg | 36.2367512 | -3.0583209 | 1.47427276 | -2.0744607 | 0.03803654 | 0.29468083 |
| Btg3 | 130.352197 | -0.2592273 | 0.44457335 | -0.5830923 | 0.55983116 | 0.87441453 |
| D16Ertd472e | 474.801505 | -0.5152031 | 0.44955784 | -1.1460217 | 0.25178616 | 0.68505999 |
| Usp25 | 469.724581 | -0.0045874 | 0.23978523 | -0.0191312 | 0.98473646 | 0.99564261 |
| Samsn1 | 2565.75123 | -0.36275 | 0.19382388 | -1.8715441 | 0.0612697 | 0.38220919 |
| Rfc4 | 4203.3954 | 0.18130542 | 0.30979232 | 0.58524828 | 0.55838078 | 0.87326414 |
| Eif4a2 | 3975.45656 | -0.0473023 | 0.27518679 | -0.1718915 | 0.86352286 | 0.9705375 |
| St6gal1 | 1571.74082 | 1.20984735 | 0.28684154 | 4.21782479 | 2.466703677 | 0.00108155 |
| Mrpl39 | 2471.13697 | -0.0290594 | 0.22441775 | -0.1294878 | 0.89697168 | 0.97984623 |
| Atp5j | 8493.3294 | 0.2692038 | 0.31834465 | 0.84563631 | 0.39775566 | 0.79531492 |
| App | 3285.87311 | 0.65154737 | 0.32276703 | 2.01863049 | 0.04352564 | 0.31959733 |
| Adamts1 | 608.861393 | 2.0108291 | 0.46972895 | 4.28082852 | 1.861987954 | 0.00085954 |

| | | | | | | |
|---|---|---|---|---|---|---|
| Ets2 | 3226.24406 | 0.14958795 | 0.42858044 | 0.34903122 | 0.72706587 | 0.93576253 |
| Dyrk1a | 918.902017 | -0.0825796 | 0.26186671 | -0.3153496 | 0.75249623 | 0.94018119 |
| Vps26c | 7051.01203 | 0.56332901 | 0.26288051 | 2.14290902 | 0.0321204 | 0.26624356 |
| Ildr1 | 10.0818052 | -0.1998779 | 2.67019714 | -0.0748551 | 0.94033 | 0.98934006 |
| Cd86 | 7122.64009 | 1.61745709 | 0.26982067 | 5.9945633 | 2.040329788 | 3.466576986 |
| Kpna1 | 3530.76115 | -0.0485287 | 0.20085764 | -0.2416074 | 0.80908441 | 0.95844313 |
| Parp9 | 1513.91294 | 0.19089331 | 0.21425899 | 0.89094658 | 0.37295783 | 0.77748705 |
| Arl13b | 306.295122 | 0.2629455 | 0.47546827 | 0.55302429 | 0.58024677 | 0.88286755 |
| Pros1 | 8263.10844 | -0.1289829 | 0.1904667 | -0.677194 | 0.49828289 | 0.84860011 |
| Psmg1 | 3602.09515 | -0.1288492 | 0.3201088 | -0.402517 | 0.68730355 | 0.92427909 |
| Brwd1 | 905.304611 | 0.17640912 | 0.21633344 | 0.81545007 | 0.41481474 | 0.80395688 |
| Pigp | 650.798783 | 0.56316234 | 0.37230305 | 1.512645 | 0.13036986 | 0.54785794 |
| Chaf1b | 3492.93746 | 0.12136737 | 0.18017081 | 0.67362394 | 0.50055043 | 0.84962306 |
| Dop1b | 201.948518 | 0.50946159 | 0.4044581 | 1.25961525 | 0.20780819 | 0.6418276 |
| Cbr3 | 26.5577283 | 1.39504667 | 0.96000037 | 1.45317306 | 0.14617571 | 0.5731306 |
| Setd4 | 122.414332 | -0.2567314 | 0.59290291 | -0.4330075 | 0.66500932 | 0.91477105 |
| Clic6 | 15.2753339 | -0.7849896 | 1.15156207 | -0.6816737 | 0.49544529 | 0.84722895 |
| Rcan1 | 542.693101 | 1.76923194 | 0.43218696 | 4.09367262 | 4.245937059 | 0.00176669 |
| Runx1 | 1499.7123 | -0.2903064 | 0.33916084 | -0.8559549 | 0.39202273 | 0.79031213 |
| Atp5o | 11721.6944 | 0.39871215 | 0.32939158 | 1.21045033 | 0.22610614 | 0.65937941 |
| Itsn1 | 673.446583 | 1.72544372 | 0.51494258 | 3.35074976 | 0.00080593 | 0.0197775 |
| Donson | 1534.38235 | -0.1918225 | 0.32565159 | -0.589042 | 0.55583309 | 0.87260256 |
| Son | 2927.64403 | -0.105208 | 0.19881851 | -0.5291658 | 0.59669042 | 0.88891958 |
| Gart | 11596.1465 | -0.4281096 | 0.16306222 | -2.6254371 | 0.00865378 | 0.11762411 |
| Tmem50b | 2130.31495 | -0.3650145 | 0.16666765 | -2.1900738 | 0.02851888 | 0.25141958 |
| Ifngr2 | 5315.19117 | -0.0580055 | 0.17977598 | -0.3226541 | 0.74695723 | 0.93988149 |
| Ifnar1 | 4276.02606 | 0.20983112 | 0.24561429 | 0.85431154 | 0.3929324 | 0.79137753 |
| Il10rb | 5143.94047 | 0.45599693 | 0.15701006 | 2.90425289 | 0.00368131 | 0.06324919 |
| Ifnar2 | 4370.82812 | -0.5594549 | 0.16000585 | -3.4964655 | 0.00047147 | 0.01284507 |
| Cfap298 | 902.932975 | -0.2588325 | 0.29289508 | -0.8837038 | 0.37685615 | 0.77940657 |
| Synj1 | 495.343016 | 0.51376317 | 0.29945839 | 1.71564128 | 0.08622772 | 0.44886114 |
| Paxbp1 | 219.977076 | -0.1738494 | 0.43716253 | -0.3976769 | 0.69086839 | 0.92470267 |
| Mis18a | 1501.49263 | 0.10372143 | 0.29553715 | 0.35095901 | 0.7256191 | 0.93544211 |
| Sod1 | 12871.3413 | -0.2347145 | 0.22148446 | -1.0597337 | 0.28926579 | 0.72395811 |
| Scaf4 | 1163.21447 | 0.3003885 | 0.35658351 | 0.84240715 | 0.39956009 | 0.79612475 |
| Kansl2 | 3638.73275 | -0.2023296 | 0.19482038 | -1.0385445 | 0.29901661 | 0.73093388 |
| Adcy6 | 21.5972942 | 2.29567464 | 1.94833373 | 1.17827588 | 0.23868664 | 0.67452555 |
| Enah | 158.02508 | 2.1864804 | 0.54607674 | 4.00398012 | 6.228560208 | 0.00242656 |
| Lmbr1l | 77.8186922 | 0.1692766 | 0.49095795 | 0.34478839 | 0.73025346 | 0.93653187 |
| Tuba1b | 127579.677 | -0.1093491 | 0.16303451 | -0.6707116 | 0.50240425 | 0.85012672 |
| Prpf40b | 93.608589 | 0.38320517 | 0.43652714 | 0.8778496 | 0.38002535 | 0.78250296 |
| Fmnl3 | 426.262474 | -0.3765325 | 0.32992037 | -1.1412829 | 0.25375221 | 0.68721513 |
| Nckap5l | 127.48083 | 0.42807108 | 0.6076455 | 0.70447504 | 0.48113699 | 0.83974159 |
| Tmbim6 | 19035.465 | 0.08961699 | 0.21412445 | 0.4185276 | 0.67556141 | 0.91851958 |
| Racgap1 | 8313.61631 | -0.0076899 | 0.13556869 | -0.0567231 | 0.95476576 | 0.99215259 |
| Asic1 | 10.9003214 | -1.7836629 | 1.25055727 | -1.4262944 | 0.15378338 | 0.58242478 |

| | | | | | | |
|---|---|---|---|---|---|---|
| Smarcd1 | 188.769059 | 0.08442756 | 0.41298496 | 0.20443253 | 0.83801551 | 0.96683245 |
| Gpd1 | 7.41073833 | 0.87969169 | 1.76259073 | 0.49909016 | 0.61771587 | 0.89878056 |
| Cox14 | 2704.33053 | 0.60941838 | 0.39717568 | 1.5343799 | 0.12493622 | 0.53626132 |
| Cers5 | 3556.51102 | -0.1678968 | 0.18393366 | -0.9128117 | 0.36134161 | 0.76887042 |
| Lima1 | 892.227085 | 0.87746669 | 0.34340452 | 2.55519843 | 0.01061273 | 0.13331258 |
| Larp4 | 1526.15085 | -0.0690479 | 0.18900725 | -0.3653189 | 0.7148734 | 0.93235217 |
| Dip2b | 1174.758 | -0.1626323 | 0.38319936 | -0.4244065 | 0.67126936 | 0.91711586 |
| Atf1 | 1826.74353 | -0.4986854 | 0.18539419 | -2.6898654 | 0.00714808 | 0.10299472 |
| Slc11a2 | 908.861082 | 0.60899797 | 0.19922666 | 3.05680966 | 0.00223706 | 0.04336924 |
| Cela1 | 885.648182 | 1.90763556 | 0.33798653 | 5.64411704 | 1.660311012 | 2.031058461 |
| Slc4a8 | 10.2919433 | -1.29783 | 1.60265207 | -0.8098015 | 0.41805429 | 0.8051099 |
| Nr4a1 | 9217.90353 | -0.1649765 | 0.47318665 | -0.3486499 | 0.72735219 | 0.93576253 |
| Pcdhgc4 | 28.014632 | 1.71090905 | 2.3184372 | 0.7379579 | 0.46054003 | 0.82987212 |
| Krt18 | 74.2882776 | -0.3077749 | 0.6464834 | -0.4760754 | 0.63402064 | 0.90465549 |
| Csad | 276.844669 | 0.81566688 | 0.48640325 | 1.67693552 | 0.09355509 | 0.46750793 |
| Soat2 | 20.8981145 | 1.37847752 | 1.0686678 | 1.28990274 | 0.19708443 | 0.62932233 |
| Prr13 | 2486.53499 | 0.09657702 | 0.18165497 | 0.53165086 | 0.59496783 | 0.88830085 |
| Map3k12 | 63.5966118 | 0.6746145 | 0.93493483 | 0.72156313 | 0.47056312 | 0.83381643 |
| Tarbp2 | 2822.94358 | 0.08858102 | 0.22588788 | 0.39214597 | 0.69495036 | 0.92609254 |
| Calcoco1 | 180.168049 | 0.74690137 | 0.37828714 | 1.97442974 | 0.0483329 | 0.33880059 |
| Rttn | 143.47428 | 0.06144535 | 0.52709412 | 0.11657378 | 0.90719782 | 0.98176897 |
| Cdkn1a | 1377.52333 | 0.14861507 | 0.31861154 | 0.46644597 | 0.64089631 | 0.90685007 |
| Nus1 | 753.056624 | -0.0491591 | 0.36753539 | -0.1337535 | 0.8935975 | 0.97941957 |
| Cep89 | 720.602621 | -0.2814775 | 0.27413726 | -1.0267758 | 0.30452605 | 0.73467909 |
| Mospd1 | 918.915786 | 0.64286354 | 0.28236577 | 2.27670494 | 0.02280385 | 0.2163471 |
| Akirin1 | 4803.27574 | 0.04850106 | 0.16948741 | 0.2861632 | 0.77475313 | 0.94709254 |
| Gtf2ird1 | 310.234919 | 0.53678813 | 0.45601807 | 1.17712029 | 0.2391475 | 0.67535535 |
| Noct | 765.154216 | -0.0329987 | 0.36253629 | -0.0910218 | 0.92747529 | 0.98666423 |
| Abcc1 | 682.848671 | -0.4339822 | 0.37554516 | -1.1556058 | 0.24784245 | 0.68235696 |
| Ndufa5 | 1964.45123 | 0.18396449 | 0.3035512 | 0.60604107 | 0.54448748 | 0.86960875 |
| Fhl1 | 53.948369 | 0.76360601 | 1.42035843 | 0.53761501 | 0.59084287 | 0.88755516 |
| Msrb2 | 830.048004 | 0.26597083 | 0.33347482 | 0.79757397 | 0.42511776 | 0.80988459 |
| Rfc2 | 10607.5667 | -0.1948413 | 0.17759996 | -1.0970796 | 0.2726066 | 0.71005689 |
| Denr | 6792.85317 | -0.1066364 | 0.2131119 | -0.5003777 | 0.61680913 | 0.89878056 |
| Prmt5 | 7478.79429 | -0.3179393 | 0.13835501 | -2.2979961 | 0.02156201 | 0.21084581 |
| Sympk | 463.854008 | -0.4516304 | 0.36669829 | -1.2316129 | 0.21809371 | 0.65150232 |
| Nagpa | 6216.43934 | -0.065737 | 0.18931321 | -0.3472396 | 0.72841134 | 0.93605089 |
| Wrb | 2437.1664 | -0.0252313 | 0.2071387 | -0.1218085 | 0.90305068 | 0.98043702 |
| Ivns1abp | 11475.4246 | -0.183916 | 0.13715674 | -1.3409185 | 0.17994691 | 0.61079094 |
| Rpp14 | 3125.08068 | 0.02367286 | 0.27865351 | 0.08495447 | 0.93229758 | 0.98730569 |
| Slc38a1 | 4806.35584 | -0.3143795 | 0.28872506 | -1.0888542 | 0.27621819 | 0.7118544 |
| Gps2 | 1218.36965 | -0.1922321 | 0.22399445 | -0.8582002 | 0.39078195 | 0.7899826 |
| Bsg | 12850.9773 | 0.08599375 | 0.23383408 | 0.36775541 | 0.71305562 | 0.93235217 |
| Vwa5a | 4995.93271 | 0.47220219 | 0.1608475 | 2.93571366 | 0.00332781 | 0.05849015 |
| Il15ra | 97.9097861 | -0.0076215 | 0.55306825 | -0.0137804 | 0.98900516 | 0.99650865 |
| Serping1 | 13.4240475 | -0.2326649 | 2.24858796 | -0.1034715 | 0.91758873 | 0.98390186 |

| | | | | | | |
|---|---|---|---|---|---|---|
| Ccl25 | 223.480347 | -0.2040445 | 0.36037122 | -0.5662064 | 0.57125347 | 0.88035336 |
| Kcnk5 | 504.111086 | 0.20567196 | 0.30081537 | 0.68371492 | 0.4941552 | 0.84678075 |
| Parp3 | 233.865439 | -0.7056459 | 0.40902837 | -1.7251759 | 0.08449578 | 0.44526833 |
| Slc26a6 | 172.067051 | 0.76043283 | 0.33732234 | 2.25432099 | 0.02417598 | 0.22455944 |
| Acy1 | 2784.5119 | -0.3772775 | 0.19632762 | -1.921673 | 0.05464692 | 0.36033644 |
| Creld2 | 4402.68504 | -0.0890611 | 0.16110711 | -0.5528069 | 0.58039564 | 0.88286755 |
| Twf2 | 9220.53607 | -0.1481844 | 0.22129847 | -0.6696131 | 0.50310446 | 0.85012672 |
| Zfp605 | 124.118334 | -0.8478456 | 0.68074917 | -1.2454596 | 0.21296285 | 0.64596443 |
| Ube2j2 | 1859.13136 | 0.21130015 | 0.30217519 | 0.69926374 | 0.48438723 | 0.84133309 |
| March5 | 890.393559 | -0.0250659 | 0.26113921 | -0.0959868 | 0.92353109 | 0.98493282 |
| Dtwd1 | 689.238094 | -0.0058933 | 0.30505897 | -0.0193185 | 0.98458699 | 0.99564261 |
| Mx2 | 24.980183 | -1.0150593 | 1.66769463 | -0.6086602 | 0.54274972 | 0.86903891 |
| Poc1a | 2224.80476 | -0.2133044 | 0.16185962 | -1.3178361 | 0.18755852 | 0.61935434 |
| Trip6 | 205.746996 | 0.87653399 | 0.38492682 | 2.2771445 | 0.0227776 | 0.2163471 |
| Clec4n | 695.194846 | 0.25487965 | 0.43770721 | 0.58230626 | 0.56036041 | 0.87469256 |
| Agap3 | 222.043595 | -0.3231309 | 0.38771252 | -0.8334291 | 0.40460274 | 0.79917613 |
| Tmem176a | 1273.7047 | -0.7412471 | 0.21119123 | -3.5098386 | 0.00044838 | 0.01240954 |
| Slc17a9 | 309.921256 | 0.1863462 | 0.362662 | 0.51382887 | 0.60737164 | 0.89332558 |
| Pisd | 4272.74817 | -0.1504017 | 0.20663556 | -0.7278599 | 0.46669932 | 0.83345004 |
| Tpi1 | 56418.8304 | -0.2412626 | 0.1517466 | -1.5899047 | 0.11185629 | 0.50962308 |
| Rab12 | 60.0803054 | 0.59304914 | 0.74702279 | 0.7938836 | 0.4272632 | 0.81021713 |
| Tulp2 | 16.7208714 | -0.1164606 | 1.63215557 | -0.0713539 | 0.94311614 | 0.98983253 |
| Pcbp4 | 519.448542 | -0.9722751 | 0.28544562 | -3.4061657 | 0.00065882 | 0.01693144 |
| Cdca3 | 4762.58553 | -0.147625 | 0.18859322 | -0.7827691 | 0.43376269 | 0.81483708 |
| C1qtnf12 | 478.001276 | -0.4986871 | 0.2701654 | -1.8458586 | 0.06491274 | 0.39268276 |
| Ccndbp1 | 1022.92138 | -0.0980127 | 0.25991306 | -0.3770981 | 0.70610074 | 0.92896305 |
| Ogfod2 | 598.42338 | 0.02920634 | 0.21446692 | 0.1361811 | 0.89167811 | 0.97925293 |
| Mrps23 | 2310.9616 | 0.00655814 | 0.32452701 | 0.0202083 | 0.98387721 | 0.99564261 |
| Rhebl1 | 129.398726 | -0.0776201 | 0.53332322 | -0.1455404 | 0.88428425 | 0.97721157 |
| Sfi1 | 177.270575 | -0.1389183 | 0.4096189 | -0.3391404 | 0.73450397 | 0.93750666 |
| Hes7 | 7.17127901 | 0.39333992 | 1.41576096 | 0.27782933 | 0.78114337 | 0.94996788 |
| Pigx | 1706.54289 | 0.56614256 | 0.41244807 | 1.37263961 | 0.16986441 | 0.60430173 |
| Pisd-ps2 | 14.332392 | 1.06609506 | 0.9581089 | 1.11270761 | 0.26583403 | 0.7019097 |
| Tiam2 | 88.2540339 | -1.4252803 | 0.52372659 | -2.7214205 | 0.0065002 | 0.09744727 |
| Synj2 | 65.6934748 | -1.1845113 | 0.65868242 | -1.7983041 | 0.07212884 | 0.41303124 |
| Rsph3b | 768.923495 | -0.0355956 | 0.22102936 | -0.1610445 | 0.87205833 | 0.97299248 |
| Rps6ka2 | 82.0282529 | 1.12959977 | 0.77215394 | 1.46292042 | 0.14348916 | 0.56861122 |
| Prkn | 95.831421 | 1.50534288 | 1.33328779 | 1.12904573 | 0.25887855 | 0.69546977 |
| Agpat4 | 364.864341 | 0.31597837 | 0.31062574 | 1.01723177 | 0.30904319 | 0.73867239 |
| Igf2r | 153.422608 | 0.77154183 | 0.4268509 | 1.80752071 | 0.07068113 | 0.41036651 |
| Acat2 | 5699.87905 | 0.65889286 | 0.23220493 | 2.83754895 | 0.00454614 | 0.07415053 |
| Lnpep | 818.572098 | -0.2928517 | 0.30379776 | -0.9639692 | 0.33506137 | 0.75359284 |
| Chd1 | 694.842481 | 0.0005077 | 0.70874498 | 0.00071634 | 0.99942845 | 0.99975535 |
| Mpc1 | 2556.85689 | -0.0770016 | 0.32463561 | -0.237194 | 0.81250628 | 0.95967622 |
| 1700010I14R | 148.562299 | 0.42006112 | 0.42536458 | 0.98753197 | 0.32338192 | 0.74583177 |
| Zfp54 | 216.983401 | 1.50918214 | 0.55713203 | 2.70884109 | 0.00675187 | 0.09966081 |

| | | | | | | |
|---|---|---|---|---|---|---|
| Phf10 | 392.300725 | -0.4397048 | 0.29934902 | -1.4688699 | 0.14186808 | 0.56696251 |
| Zfp51 | 633.245382 | 0.28156104 | 0.4358051 | 0.64607102 | 0.51823336 | 0.8570838 |
| Hcfc1r1 | 427.218927 | 0.45503754 | 0.30319832 | 1.50079177 | 0.13340943 | 0.55359482 |
| Tnfrsf12a | 127.887347 | 0.19197403 | 0.56953595 | 0.33707096 | 0.73606341 | 0.9377627 |
| Cldn6 | 30.2065676 | -1.1197708 | 1.43994907 | -0.7776461 | 0.43677769 | 0.81786338 |
| Pkmyt1 | 1442.71707 | -0.0728765 | 0.28237785 | -0.2580814 | 0.79634405 | 0.9537323 |
| Paqr4 | 483.389664 | 1.69935705 | 0.33951938 | 5.00518429 | 5.580861207 | 4.404559687 |
| Pla2g7 | 360.969562 | 0.20881302 | 0.54747371 | 0.38141195 | 0.70289759 | 0.9278673 |
| Tnfrsf21 | 2545.69156 | -0.1074768 | 0.30158638 | -0.3563716 | 0.72156233 | 0.93398714 |
| Cenpq | 1188.42817 | 0.38795815 | 0.2804819 | 1.38318424 | 0.16660838 | 0.59997655 |
| Mmut | 1799.71833 | 0.27820393 | 0.22932614 | 1.21313656 | 0.22507761 | 0.65870021 |
| Tbc1d5 | 728.449173 | -0.0876508 | 0.28098828 | -0.3119374 | 0.75508811 | 0.94084431 |
| Satb1 | 136.118623 | -0.5740257 | 0.52750286 | -1.0881946 | 0.27650922 | 0.7118544 |
| Cdc5l | 3430.56983 | -0.1836017 | 0.18478842 | -0.9935778 | 0.32042845 | 0.74413737 |
| Spats1 | 57.1831589 | -0.7361585 | 0.65287006 | -1.1275727 | 0.2595004 | 0.69591124 |
| Aars2 | 689.762245 | -0.0778219 | 0.28534613 | -0.2727281 | 0.78506221 | 0.95142322 |
| Mrpl14 | 1869.9923 | 0.87762827 | 0.45224146 | 1.94061878 | 0.05230453 | 0.35365015 |
| Sgo1 | 1488.81777 | -0.0596446 | 0.18361854 | -0.3248287 | 0.74531072 | 0.93917909 |
| Slc29a1 | 4146.12372 | 0.28979492 | 0.21986954 | 1.3180312 | 0.1874932 | 0.61935434 |
| Hsp90ab1 | 109298.424 | -0.2386664 | 0.14531195 | -1.6424416 | 0.10049853 | 0.48478567 |
| Nfkbie | 412.005072 | 0.28421174 | 0.47072159 | 0.60377886 | 0.54599068 | 0.86990153 |
| Gtpbp2 | 238.046165 | 0.41530427 | 0.50492061 | 0.822514 | 0.41078442 | 0.80139357 |
| Polh | 811.62258 | -0.0771799 | 0.25723123 | -0.3000409 | 0.76414596 | 0.94300309 |
| Enpp4 | 140.049618 | 0.17951048 | 0.47751475 | 0.37592656 | 0.70697149 | 0.92935286 |
| Cyp39a1 | 67.7223511 | 0.44808482 | 0.75820055 | 0.59098457 | 0.55453076 | 0.87165957 |
| Fbxl17 | 58.5340852 | -0.6442594 | 1.06434804 | -0.605309 | 0.54497371 | 0.86963259 |
| Rsph9 | 10.0837953 | 0.29492274 | 1.46835227 | 0.20085285 | 0.84081364 | 0.96765585 |
| Mrps18a | 2959.90356 | 0.34968574 | 0.32094789 | 1.08954054 | 0.27591559 | 0.7118544 |
| Rrp36 | 1239.36292 | 0.41437207 | 0.27975218 | 1.48121123 | 0.13855029 | 0.56122045 |
| Ptk7 | 712.926563 | -0.6111358 | 0.28946324 | -2.1112725 | 0.0347489 | 0.27929255 |
| Cnpy3 | 3236.82419 | -0.2488991 | 0.19747022 | -1.2604386 | 0.20751117 | 0.6418276 |
| Ubr2 | 978.810719 | 0.07809328 | 0.39350687 | 0.19845468 | 0.84268934 | 0.96765585 |
| Taf8 | 968.498603 | -0.1374201 | 0.25817821 | -0.5322683 | 0.5945402 | 0.88828726 |
| Guca1a | 204.468565 | 1.11974739 | 0.63122728 | 1.77392111 | 0.0760762 | 0.42514288 |
| Gm20517 | 61.509705 | 0.40628716 | 0.74213084 | 0.54746028 | 0.58406255 | 0.88404713 |
| Bysl | 3395.25394 | -0.616825 | 0.23952835 | -2.5751648 | 0.01001923 | 0.12928829 |
| Tfeb | 226.335453 | -0.5928237 | 0.49045373 | -1.208725 | 0.22676852 | 0.66015092 |
| Foxp4 | 26.6069775 | 0.50661635 | 1.03880242 | 0.4876927 | 0.62576753 | 0.90151518 |
| Trem2 | 4030.07212 | 0.71120481 | 0.33421605 | 2.12797922 | 0.03333881 | 0.27170794 |
| Nfya | 296.240932 | -0.2364973 | 0.32673877 | -0.7238116 | 0.46918143 | 0.83381643 |
| Brd4 | 2025.35198 | -0.0500283 | 0.25486485 | -0.1962933 | 0.84438063 | 0.96765585 |
| Stk38 | 2368.30848 | -0.0045406 | 0.24982067 | -0.0181756 | 0.98549875 | 0.99564261 |
| Ppil1 | 7374.18946 | 0.13410114 | 0.22398499 | 0.59870592 | 0.54936901 | 0.87093412 |
| Mtch1 | 460.576592 | -0.1760201 | 0.3319934 | -0.5301914 | 0.59597922 | 0.8886604 |
| Fgd2 | 5714.53845 | 0.09570826 | 0.22600305 | 0.42348218 | 0.6719435 | 0.91729548 |
| Pim1 | 794.440678 | 0.20420062 | 0.38271556 | 0.53355715 | 0.59364796 | 0.88797037 |

| | | | | | | |
|---|---|---|---|---|---|---|
| Ccdc167 | 401.241025 | 1.01081462 | 0.54382314 | 1.8587194 | 0.06306692 | 0.38729802 |
| Cmtr1 | 3621.58127 | -0.2426206 | 0.24751601 | -0.9802218 | 0.32697665 | 0.74811039 |
| Glo1 | 4144.30174 | -0.0694143 | 0.15955795 | -0.4350413 | 0.66353244 | 0.91395288 |
| Abcg1 | 576.232877 | 0.89823925 | 0.35743853 | 2.51298944 | 0.01197129 | 0.14499487 |
| Slc37a1 | 393.726678 | -0.1905616 | 0.31799514 | -0.5992595 | 0.54899986 | 0.87083964 |
| Wdr4 | 3082.13361 | 0.00286013 | 0.28631061 | 0.00998959 | 0.99202959 | 0.99749285 |
| Ndufv3 | 3071.38238 | 0.11575543 | 0.30015673 | 0.38564994 | 0.69975594 | 0.92669304 |
| Cbs | 10.3129732 | -0.3756834 | 2.27175363 | -0.1653715 | 0.86865157 | 0.9719395 |
| Sik1 | 117.669895 | -1.3048951 | 0.56017674 | -2.3294347 | 0.01983605 | 0.19955131 |
| Arhgap28 | 105.750665 | -0.021191 | 0.59148219 | -0.035827 | 0.97142032 | 0.99443087 |
| Epb41l3 | 216.592321 | 3.50715102 | 0.63621773 | 5.51250124 | 3.537696992 | 4.007096972 |
| Akap8 | 3442.65106 | -0.2500301 | 0.24289899 | -1.0293583 | 0.30331131 | 0.73436097 |
| Myl12a | 14007.7786 | 0.05321614 | 0.21320688 | 0.2495986 | 0.80289778 | 0.95672466 |
| Myom1 | 69.3160897 | -0.2939344 | 0.65108435 | -0.4514537 | 0.65166258 | 0.91036394 |
| Wiz | 166.65559 | -0.5111834 | 0.6438406 | -0.7939596 | 0.42721895 | 0.81021713 |
| Lpin2 | 1057.61184 | 0.545892 | 0.50546571 | 1.0799783 | 0.28015184 | 0.71532524 |
| Emilin2 | 5118.14999 | -1.1137163 | 0.29726752 | -3.7465118 | 0.00017931 | 0.00599319 |
| Smchd1 | 1299.14257 | -0.6255856 | 0.27104065 | -2.3080877 | 0.02099426 | 0.20694822 |
| Cyp4f13 | 209.552759 | 0.27993737 | 0.42257737 | 0.66245234 | 0.50768137 | 0.85197408 |
| Ndc80 | 2593.84708 | 0.34283319 | 0.20883175 | 1.64167181 | 0.10065804 | 0.48497431 |
| Lbh | 217.925675 | -0.0578494 | 0.32343711 | -0.1788583 | 0.85804895 | 0.96944425 |
| Xdh | 37.8523441 | -4.5050092 | 0.94806519 | -4.7517926 | 2.016210947 | 0.00013332 |
| Dpy30 | 3174.14195 | 0.17958086 | 0.37034878 | 0.48489659 | 0.6277497 | 0.90236182 |
| Spast | 539.852251 | -0.3005182 | 0.29119914 | -1.0320022 | 0.30207109 | 0.73361866 |
| Slc30a6 | 1282.93263 | 0.32532344 | 0.28158881 | 1.15531381 | 0.24796194 | 0.68235696 |
| Prkd3 | 634.682167 | 0.13896252 | 0.260582 | 0.53327753 | 0.59384148 | 0.88797037 |
| Yipf4 | 2075.02888 | 0.17118009 | 0.20071599 | 0.85284731 | 0.39374398 | 0.79233901 |
| Birc6 | 1798.73046 | 0.10434258 | 0.33141791 | 0.31483688 | 0.75288551 | 0.94018119 |
| Crim1 | 78.2769681 | 0.53542839 | 0.69752573 | 0.76761095 | 0.44271836 | 0.82038676 |
| Strn | 780.841839 | -0.2535305 | 0.4094335 | -0.6192226 | 0.53576971 | 0.86539693 |
| Ttc27 | 1401.30748 | -0.0597857 | 0.1961042 | -0.3048671 | 0.76046741 | 0.94215089 |
| Eif2ak2 | 447.976705 | 0.26948974 | 0.29502585 | 0.91344449 | 0.36100882 | 0.76887042 |
| Cebpz | 1890.91357 | -0.323514 | 0.21974383 | -1.4722324 | 0.14095812 | 0.5653576 |
| Ndufaf7 | 1793.96581 | -0.0065963 | 0.23895565 | -0.0276046 | 0.97797753 | 0.99483019 |
| Pja2 | 1682.90463 | 0.27289561 | 0.3145385 | 0.86760637 | 0.38560986 | 0.78819532 |
| Qpct | 323.720815 | 1.56484182 | 0.45099788 | 3.46973209 | 0.00052098 | 0.01392544 |
| Man2a1 | 277.650654 | -0.2969572 | 0.38708288 | -0.767167 | 0.4429822 | 0.82038676 |
| Vapa | 11978.8068 | -0.0882021 | 0.16451048 | -0.5361486 | 0.59185582 | 0.88797037 |
| Hnrnpll | 1856.1909 | 0.26110542 | 0.48926411 | 0.53366968 | 0.59357009 | 0.88797037 |
| Ralbp1 | 949.721135 | 0.38139831 | 0.40657595 | 0.93807397 | 0.3482064 | 0.76101142 |
| Srsf7 | 16250.9051 | 0.08579937 | 0.31596237 | 0.27154933 | 0.78596856 | 0.95184068 |
| Twsg1 | 332.520462 | 0.35995569 | 0.43498559 | 0.82751175 | 0.40794707 | 0.80064439 |
| Ndufv2 | 7909.49995 | 0.29470584 | 0.36211182 | 0.81385313 | 0.41572909 | 0.80433694 |
| Washc1 | 669.222549 | 0.31557 | 0.25363453 | 1.24419181 | 0.21342898 | 0.64625661 |
| Washc2 | 3525.37398 | 0.09307012 | 0.33990252 | 0.27381416 | 0.78422744 | 0.95094799 |
| Tedc2 | 496.001213 | 0.0050533 | 0.24323163 | 0.02077566 | 0.98342461 | 0.99564261 |

| | | | | | | |
|---|---|---|---|---|---|---|
| Lrpprc | 3058.88394 | -0.227246 | 0.34505881 | -0.6585719 | 0.51017071 | 0.85335831 |
| Atp6v0c | 20539.4786 | 0.21918114 | 0.18622782 | 1.17695165 | 0.23921481 | 0.67535535 |
| Pdpk1 | 988.490907 | -0.1235828 | 0.27454472 | -0.4501372 | 0.65261148 | 0.91051509 |
| Prepl | 591.971522 | 0.17596364 | 0.20977409 | 0.83882449 | 0.4015678 | 0.79737342 |
| Abca3 | 540.094739 | 0.07789651 | 0.32869642 | 0.23698618 | 0.81266751 | 0.95967622 |
| Eci1 | 2017.12978 | 0.11090533 | 0.25452109 | 0.43574123 | 0.66302449 | 0.91395288 |
| Srbd1 | 792.708225 | -0.0604192 | 0.22796547 | -0.2650366 | 0.79098124 | 0.95268772 |
| Dnase1l2 | 11.5015806 | 2.02456025 | 1.99662384 | 1.01399182 | 0.31058665 | 0.7399039 |
| E4f1 | 600.587551 | 0.25374557 | 0.32961157 | 0.76983211 | 0.44139949 | 0.81967485 |
| Mlst8 | 699.701309 | 0.2697112 | 0.30418654 | 0.88666384 | 0.37525991 | 0.77815049 |
| Rhoq | 183.323273 | -0.4866956 | 0.41405577 | -1.1754349 | 0.23982075 | 0.67563574 |
| Pigf | 579.64606 | -0.0111541 | 0.41095306 | -0.027142 | 0.97834645 | 0.99483019 |
| Cript | 3847.61898 | 0.62708438 | 0.43893017 | 1.42866548 | 0.1531004 | 0.58163889 |
| Mcfd2 | 5331.90402 | 0.55932585 | 0.21517116 | 2.59944615 | 0.00933743 | 0.12388808 |
| Msh2 | 3108.83103 | -0.2582435 | 0.18778085 | -1.3752386 | 0.16905749 | 0.60312169 |
| Hagh | 940.257026 | 0.47623272 | 0.35942114 | 1.32499918 | 0.18517142 | 0.61717463 |
| Spsb3 | 1171.13034 | -0.0403436 | 0.22660789 | -0.1780328 | 0.8586972 | 0.96954641 |
| Mapk8ip3 | 204.008517 | 0.38320296 | 0.43066793 | 0.88978757 | 0.37357996 | 0.77748705 |
| C3 | 64.9632895 | -5.1425233 | 0.88763934 | -5.7934828 | 6.894155451 | 1.081233379 |
| Jpt2 | 14959.4148 | 0.00534211 | 0.16185041 | 0.03300643 | 0.97366946 | 0.99443087 |
| Tmem204 | 37.6625466 | 3.38870178 | 1.29242789 | 2.62196584 | 0.00874242 | 0.11869702 |
| Ift140 | 258.45599 | -0.0170575 | 0.65210884 | -0.0261575 | 0.97913171 | 0.99483019 |
| Telo2 | 939.23813 | -0.0308881 | 0.23314296 | -0.1324858 | 0.89460008 | 0.97941957 |
| Pot1b | 119.869721 | -0.2585276 | 0.57189513 | -0.4520542 | 0.65122997 | 0.91021183 |
| Nme4 | 4663.99625 | 1.21133131 | 0.36685704 | 3.30191654 | 0.00096027 | 0.02272135 |
| Tmem8 | 512.344698 | 1.19619939 | 0.45065346 | 2.65436636 | 0.00794575 | 0.1102045 |
| Mrpl28 | 9718.68072 | 0.67996607 | 0.39989229 | 1.70037302 | 0.08906078 | 0.4566019 |
| Axin1 | 470.991992 | -0.0635781 | 0.30983259 | -0.2052014 | 0.83741476 | 0.96683245 |
| Fam234a | 1239.3417 | 1.07688591 | 0.35859186 | 3.0030964 | 0.00267248 | 0.04916153 |
| Luc7l | 1765.67203 | -0.0338296 | 0.16735799 | -0.2021391 | 0.83980802 | 0.96762336 |
| Dusp1 | 28462.583 | 0.66050194 | 0.3099026 | 2.13132108 | 0.0330627 | 0.26999733 |
| Bnip1 | 1393.71001 | 0.3070807 | 0.32161926 | 0.95479574 | 0.339681 | 0.75726534 |
| Phf1 | 41.6496773 | 0.19250714 | 0.75907512 | 0.2536075 | 0.7997988 | 0.95518266 |
| Cuta | 2424.43815 | -0.0138206 | 0.18435619 | -0.0749671 | 0.94024089 | 0.98934006 |
| Plin3 | 3772.02507 | 0.53862808 | 0.19590443 | 2.74944305 | 0.00596966 | 0.09139787 |
| Kdm4b | 557.722832 | -0.3301731 | 0.39569942 | -0.8344039 | 0.40405342 | 0.79915692 |
| Uqcc2 | 1879.17588 | 0.64212127 | 0.5564049 | 1.15405394 | 0.24847806 | 0.68263757 |
| Mllt1 | 225.668391 | -0.1515751 | 0.70629323 | -0.2146065 | 0.83007416 | 0.96464475 |
| Nudt3 | 2843.82537 | -0.3452815 | 0.24342604 | -1.4184248 | 0.15606679 | 0.58521232 |
| Snrpc | 1521.39061 | 0.21477972 | 0.28827989 | 0.74503888 | 0.45624822 | 0.82894467 |
| Taf11 | 874.542248 | 0.14755643 | 0.31529056 | 0.46800143 | 0.63978357 | 0.90663689 |
| Anks1 | 565.582921 | 0.06545647 | 0.31209373 | 0.20973338 | 0.83387577 | 0.96598507 |
| Zfp523 | 358.802071 | -0.0567075 | 0.28372548 | -0.1998674 | 0.84158431 | 0.96765585 |
| Fkbp5 | 447.118381 | -0.4390336 | 0.27175619 | -1.6155424 | 0.10619329 | 0.49777747 |
| Nudt12 | 99.626311 | 0.04317714 | 0.54454914 | 0.0792897 | 0.9368022 | 0.98843258 |
| Cul2 | 4887.12626 | 0.22896814 | 0.19235877 | 1.19031815 | 0.23392137 | 0.66968409 |

| | | | | | | |
|---|---|---|---|---|---|---|
| Bambi | 840.781984 | -0.0724796 | 0.4423575 | -0.1638485 | 0.86985044 | 0.97239152 |
| Mtpap | 1826.56387 | -0.2538767 | 0.24581992 | -1.0327751 | 0.30170917 | 0.73346745 |
| Map3k8 | 1810.27927 | 0.66011898 | 0.40395584 | 1.6341365 | 0.10223019 | 0.4886081 |
| Svil | 355.193871 | -0.5052635 | 0.37359565 | -1.3524341 | 0.17623647 | 0.60763832 |
| Zeb1 | 59.2531974 | 0.30719323 | 0.71557578 | 0.42929518 | 0.66770842 | 0.91595755 |
| Epc1 | 400.317527 | 0.43184029 | 0.39648548 | 1.08917048 | 0.27607871 | 0.7118544 |
| Sos1 | 158.748879 | -0.393998 | 0.4304854 | -0.9152413 | 0.36006496 | 0.76857232 |
| Map4k3 | 67.2234124 | -0.2118851 | 0.57649748 | -0.3675386 | 0.71321733 | 0.93235217 |
| Thumpd2 | 437.806354 | -0.24403 | 0.24481549 | -0.9967916 | 0.31886569 | 0.74398524 |
| Pkdcc | 61.1158909 | -0.3343332 | 0.83165979 | -0.4020072 | 0.68767875 | 0.92463993 |
| Cox7a2l | 6202.48466 | 0.48591131 | 0.32652633 | 1.4881229 | 0.13671848 | 0.55946682 |
| Thada | 436.062161 | 0.05816707 | 0.32185088 | 0.18072676 | 0.85658205 | 0.9692506 |
| Dync2li1 | 87.0134212 | 0.80282256 | 0.60004425 | 1.33793892 | 0.18091635 | 0.61136731 |
| Polr2d | 3164.80699 | 0.17513355 | 0.28667545 | 0.61091225 | 0.54125767 | 0.86776854 |
| Slc25a46 | 2988.26879 | -0.0706465 | 0.21803733 | -0.3240108 | 0.74592985 | 0.93955517 |
| Sap130 | 1299.11864 | -0.1275794 | 0.32737934 | -0.389699 | 0.69675913 | 0.92646244 |
| Tpgs2 | 729.949872 | 0.11515411 | 0.25984257 | 0.44316877 | 0.65764366 | 0.91258818 |
| Slc39a6 | 2009.61136 | -0.3255481 | 0.20191783 | -1.6122802 | 0.10690098 | 0.49970181 |
| Elp2 | 6199.79021 | -0.3372319 | 0.17994546 | -1.8740786 | 0.06091959 | 0.38080193 |
| 2700062C07 | 891.2645 | -0.1214484 | 0.27431549 | -0.4427326 | 0.65795913 | 0.91258818 |
| Zscan30 | 26.434989 | -1.0742052 | 1.37012071 | -0.7840223 | 0.43302701 | 0.81435873 |
| Zfp397 | 385.919877 | -0.0143518 | 0.40069141 | -0.0358175 | 0.97142789 | 0.99443087 |
| Mapre2 | 8040.99177 | -0.2306843 | 0.2206335 | -1.0455543 | 0.29576687 | 0.72857755 |
| Wac | 1176.17086 | -0.2082093 | 0.22644358 | -0.9194756 | 0.35784687 | 0.76674184 |
| Ccny | 1233.08338 | -0.0088777 | 0.25615395 | -0.0346577 | 0.97235273 | 0.99443087 |
| Thoc1 | 2918.114 | 0.23729543 | 0.28699017 | 0.82684165 | 0.40832682 | 0.80087574 |
| Rock1 | 979.688499 | -0.1490149 | 0.3273232 | -0.4552531 | 0.64892715 | 0.90973357 |
| Esco1 | 533.4431 | 0.01986767 | 0.31523815 | 0.06302433 | 0.94974713 | 0.99063619 |
| Mib1 | 505.881075 | 0.22017084 | 0.43232533 | 0.5092712 | 0.61056214 | 0.89508835 |
| Cyp4f41-ps | 46.7095501 | 1.20938268 | 0.76977754 | 1.57108076 | 0.11616389 | 0.52109749 |
| Zfp871 | 622.020222 | 0.20490604 | 0.32031254 | 0.63970659 | 0.52236337 | 0.85816354 |
| Adamts10 | 254.747829 | 0.1618894 | 0.38905497 | 0.41610933 | 0.67733001 | 0.91958179 |
| Myo1f | 12903.5774 | -0.4775135 | 0.19387484 | -2.4629986 | 0.01377805 | 0.15900646 |
| Kifc5b | 22.6080849 | 1.33661062 | 1.13762589 | 1.17491227 | 0.24002982 | 0.67563574 |
| Cdh2 | 70.3484219 | 1.99274068 | 2.74234812 | 0.72665489 | 0.46743739 | 0.83381643 |
| Tapbp | 4171.0542 | -0.3632575 | 0.16768407 | -2.1663208 | 0.03028668 | 0.26001357 |
| Pfdn6 | 2198.48901 | -0.1550413 | 0.26772757 | -0.5791011 | 0.56252096 | 0.8755831 |
| Wdr46 | 2910.8707 | -0.6339671 | 0.16309698 | -3.8870561 | 0.00010147 | 0.00365073 |
| Rnf138 | 1428.95339 | 0.03760549 | 0.28565843 | 0.13164496 | 0.89526512 | 0.97949899 |
| Vps52 | 2478.87604 | -0.0746493 | 0.22400204 | -0.3332529 | 0.73894341 | 0.93800224 |
| Ring1 | 80.2019895 | 0.41961449 | 0.50318366 | 0.83391914 | 0.40432655 | 0.79917613 |
| Slc39a7 | 5610.65595 | -0.1277529 | 0.23892848 | -0.534691 | 0.59286357 | 0.88797037 |
| H2-Oa | 58.9466511 | -1.4206369 | 0.5852716 | -2.4273122 | 0.01521116 | 0.17040121 |
| Brd2 | 2425.30894 | -0.1253279 | 0.28625516 | -0.4378188 | 0.66151762 | 0.91376977 |
| Psmb8 | 3522.48653 | -0.5085475 | 0.27777421 | -1.8307944 | 0.06713123 | 0.39891584 |
| Tap2 | 605.021016 | -0.2938432 | 0.25108144 | -1.1703102 | 0.24187617 | 0.67708722 |

| | | | | | | |
|---|---|---|---|---|---|---|
| Btnl2 | 267.717734 | 1.31868528 | 0.31058575 | 4.24580104 | 2.178138429 | 0.00097352 |
| Pfdn1 | 2680.51026 | 0.44710773 | 0.3641874 | 1.2276859 | 0.2195649 | 0.65289787 |
| Tmem173 | 2928.2862 | -0.3765115 | 0.15963306 | -2.3586059 | 0.01834372 | 0.19092573 |
| Dnajc18 | 690.419389 | 0.43151533 | 0.30783643 | 1.40176825 | 0.16098446 | 0.59141542 |
| Spata24 | 208.581783 | 0.84279233 | 0.54737865 | 1.53968797 | 0.12363643 | 0.53443267 |
| Sil1 | 697.141539 | 0.33488878 | 0.23113225 | 1.44890544 | 0.14736399 | 0.57410946 |
| Hspa9 | 22921.0075 | -0.2797207 | 0.21080104 | -1.3269415 | 0.18452804 | 0.61665869 |
| Etf1 | 8051.13992 | -0.1500759 | 0.15387577 | -0.9753056 | 0.3294087 | 0.74874249 |
| Nelfe | 1948.97521 | 0.02150362 | 0.16903813 | 0.12721165 | 0.89877288 | 0.97991878 |
| Cdc23 | 5579.11439 | -0.1284718 | 0.16381314 | -0.784258 | 0.43288876 | 0.81435873 |
| C2 | 72.6848398 | -0.6160334 | 0.65052239 | -0.9469826 | 0.34364765 | 0.75947281 |
| Epb41l4a | 13.9547971 | 1.23860889 | 2.13856275 | 0.57917819 | 0.56246895 | 0.8755831 |
| Stard4 | 1266.24733 | 0.69503711 | 0.23005063 | 3.02123539 | 0.00251746 | 0.04719108 |
| Bin1 | 3981.84503 | 0.95969903 | 0.24232932 | 3.96030909 | 7.485281447 | 0.00282616 |
| Ercc3 | 1286.12379 | -0.5230595 | 0.23192388 | -2.2553066 | 0.02411409 | 0.22455944 |
| Map3k2 | 131.543361 | 0.04153596 | 0.64314715 | 0.06458236 | 0.94850653 | 0.99019648 |
| Iws1 | 1728.88987 | -0.2387534 | 0.23714778 | -1.0067707 | 0.31404498 | 0.74065408 |
| Csnk2b | 2946.64834 | -0.2134059 | 0.19094171 | -1.1176494 | 0.26371676 | 0.70008189 |
| Bag6 | 563.493546 | -0.7340365 | 0.25553655 | -2.8725305 | 0.00407199 | 0.06805004 |
| Prrc2a | 465.21766 | -0.6441572 | 0.30060605 | -2.1428616 | 0.03212421 | 0.26624356 |
| Aif1 | 4778.72443 | 0.66826238 | 0.24923561 | 2.68124754 | 0.00733482 | 0.1045768 |
| Ltb | 45.4279809 | 0.17362601 | 0.65425513 | 0.26537966 | 0.790717 | 0.95261385 |
| Wdr33 | 2566.10128 | -0.0340119 | 0.18357849 | -0.1852718 | 0.85301585 | 0.96808079 |
| Tnf | 5092.33041 | 1.40677501 | 1.14822916 | 1.22516921 | 0.22051147 | 0.65394348 |
| Atp6v1g2 | 53.9607144 | -0.2481122 | 0.55548866 | -0.4466557 | 0.65512369 | 0.91121411 |
| Riok3 | 2688.27819 | 0.26412861 | 0.24188128 | 1.09197622 | 0.27484355 | 0.71147204 |
| Rmc1 | 1499.52453 | 0.06741973 | 0.20434217 | 0.32993547 | 0.74144872 | 0.93800224 |
| Npc1 | 538.015911 | 0.42596516 | 0.35803666 | 1.18972497 | 0.23415451 | 0.67003791 |
| Mrpl27 | 2986.16628 | 0.45363075 | 0.35670916 | 1.27171039 | 0.20347604 | 0.63809363 |
| Zfp521 | 96.3601417 | 3.0097412 | 1.25481352 | 2.39855655 | NA | NA |
| Dhx16 | 1687.14796 | -0.3723847 | 0.20045973 | -1.8576534 | 0.06321825 | 0.38764352 |
| Impact | 1435.91562 | 0.48253383 | 0.38366639 | 1.25769117 | 0.20850347 | 0.6418276 |
| Ttc39c | 17.0136115 | -2.9499214 | 1.44380011 | -2.0431647 | 0.04103614 | 0.30873004 |
| Ndfip1 | 7522.87201 | 0.58670229 | 0.27578768 | 2.12736944 | 0.0333894 | 0.2719391 |
| Atat1 | 177.716487 | 0.46033958 | 0.41298376 | 1.11466752 | 0.26499293 | 0.70112124 |
| Spry4 | 20.1031695 | 1.26845206 | 1.23956381 | 1.02330517 | 0.30616358 | 0.73554152 |
| Gnl1 | 1071.62888 | -0.4411238 | 0.30718341 | -1.4360273 | 0.15099457 | 0.57867059 |
| Nr3c1 | 820.80342 | -0.1030271 | 0.37621486 | -0.2738517 | 0.78419856 | 0.95094799 |
| Mrps18b | 4287.98813 | -0.0890991 | 0.22391577 | -0.3979136 | 0.69069388 | 0.92470267 |
| Pcdh12 | 9.77028312 | 4.93850593 | 1.7985766 | 2.74578571 | 0.00603662 | 0.09207724 |
| Dele1 | 1063.68496 | 0.21742173 | 0.28595105 | 0.76034595 | 0.44704782 | 0.82261371 |
| Rpp21 | 744.496833 | 0.12310249 | 0.35584917 | 0.34594009 | 0.72938774 | 0.93636271 |
| Arap3 | 433.418702 | 0.40465432 | 0.53027304 | 0.76310559 | 0.44540043 | 0.82118815 |
| Hdac3 | 3881.83272 | 0.01066426 | 0.16736807 | 0.0637174 | 0.94919525 | 0.99028544 |
| Diaph1 | 446.458392 | -0.4243308 | 0.2351856 | -1.8042381 | 0.071194 | 0.41122734 |
| Trim26 | 1644.22402 | 0.1784611 | 0.17343352 | 1.02898853 | 0.30348506 | 0.73436097 |

| | | | | | | |
|---|---|---|---|---|---|---|
| Dcp2 | 790.857865 | -0.0217722 | 0.3576009 | -0.060884 | 0.95145163 | 0.99115284 |
| Ik | 5097.43017 | -0.2480959 | 0.29225181 | -0.8489114 | 0.39593057 | 0.79400306 |
| Pggt1b | 1647.43619 | 0.03256787 | 0.22047234 | 0.14771862 | 0.88256483 | 0.97607952 |
| Ap3s1 | 3205.10567 | -0.1711447 | 0.18718195 | -0.9143226 | 0.36054734 | 0.76881241 |
| Ankhd1 | 572.006003 | -0.0850308 | 0.29999795 | -0.2834381 | 0.77684105 | 0.94800948 |
| Hbegf | 203.384098 | 1.22795884 | 0.5380138 | 2.28239286 | 0.02246616 | 0.21507833 |
| Yipf5 | 3256.17295 | 0.38335972 | 0.30157385 | 1.27119683 | 0.20365863 | 0.63809363 |
| Rbm27 | 1079.44661 | -0.4297721 | 0.25348022 | -1.6954859 | 0.08998328 | 0.45827038 |
| Lars | 5084.57694 | -0.3824958 | 0.24901553 | -1.5360318 | 0.12453059 | 0.53617731 |
| Tcerg1 | 3892.1011 | -0.3614278 | 0.25552591 | -1.4144466 | 0.15723082 | 0.58620126 |
| Dpysl3 | 62.405187 | 1.96231322 | 0.72074894 | 2.722603 | 0.00647698 | 0.09721834 |
| Dtwd2 | 401.800036 | -0.2006864 | 0.4066087 | -0.4935615 | 0.62161591 | 0.89912494 |
| Hsd17b4 | 4181.05736 | 0.17315322 | 0.2305704 | 0.75097768 | 0.45266609 | 0.82610332 |
| Mbd2 | 296.916825 | -0.5309017 | 0.40215136 | -1.320154 | 0.1867836 | 0.61862792 |
| Smad4 | 265.516048 | -0.9599368 | 0.34620227 | -2.772763 | 0.00555826 | 0.0866168 |
| Sec11c | 4476.53065 | 0.26238608 | 0.35433595 | 0.74050086 | 0.45899614 | 0.82987212 |
| Pmaip1 | 8078.99782 | -0.2858357 | 0.37515983 | -0.7619039 | 0.44611738 | 0.82166369 |
| Gnal | 10.1807793 | -0.7220337 | 2.18089856 | -0.3310717 | 0.74059035 | 0.93800224 |
| Impa2 | 1708.48231 | -0.4140351 | 0.23679762 | -1.7484764 | 0.08038156 | 0.43799895 |
| Afg3l2 | 3361.0941 | -0.3317192 | 0.2319246 | -1.430289 | 0.15263409 | 0.58090075 |
| Srfbp1 | 1362.54996 | -0.1022946 | 0.26768987 | -0.3821385 | 0.70235867 | 0.92762537 |
| Prelid3a | 30.816648 | 0.65297789 | 1.16743767 | 0.5593257 | 0.57593946 | 0.88080372 |
| Spire1 | 441.495312 | -0.0990388 | 0.39216509 | -0.2525437 | 0.80062085 | 0.95576786 |
| Sncaip | 9.46663219 | 2.18189441 | 2.57697355 | 0.84668871 | 0.39716865 | 0.79505223 |
| Snx24 | 922.11859 | 1.06783171 | 0.22943805 | 4.6541178 | 3.253707507 | 0.00020001 |
| Psmg2 | 1757.33501 | -0.3932026 | 0.19241413 | -2.0435222 | 0.04100077 | 0.30865381 |
| Ppic | 2647.45357 | 0.97911379 | 0.39371006 | 2.48689044 | 0.01288651 | 0.15201605 |
| Ptpn2 | 2332.04716 | -0.3327633 | 0.18554013 | -1.7934842 | 0.07289553 | 0.41525386 |
| Cep192 | 448.61681 | -0.1047616 | 0.29519194 | -0.354893 | 0.72266973 | 0.93430763 |
| Ldlrad4 | 66.9188096 | -0.145698 | 0.66519259 | -0.2190313 | 0.82662566 | 0.96370073 |
| Me2 | 7850.66151 | -0.6230816 | 0.17103703 | -3.6429633 | 0.00026952 | 0.00820151 |
| Cxxc1 | 1332.98766 | -0.3011468 | 0.26794504 | -1.1239126 | 0.26105012 | 0.69691361 |
| Mbd1 | 813.7646 | 0.19049036 | 0.28757683 | 0.66239813 | 0.5077161 | 0.85197408 |
| Smad2 | 2373.12022 | -0.0426217 | 0.19116842 | -0.2229538 | 0.82357146 | 0.9628698 |
| Sall3 | 153.554871 | -0.8389333 | 0.49565402 | -1.6925783 | 0.09053576 | 0.46012626 |
| Atp9b | 917.560378 | 0.31946293 | 0.25952584 | 1.23094844 | 0.21834215 | 0.65177636 |
| Rbfa | 5260.66972 | 0.20952181 | 0.16544823 | 1.26638891 | 0.20537388 | 0.63913241 |
| Gm16286 | 10716.3483 | -0.0598717 | 0.16241715 | -0.3686294 | 0.71240395 | 0.93235217 |
| Csnk1a1 | 8681.56528 | 0.01675309 | 0.13473738 | 0.12433883 | 0.901047 | 0.98043702 |
| Pcyox1l | 1411.44617 | -0.658391 | 0.21097289 | -3.1207376 | 0.00180399 | 0.03727732 |
| Grpel2 | 1247.96593 | 0.02133608 | 0.27726635 | 0.07695158 | 0.93866207 | 0.9890914 |
| Napg | 854.297664 | -0.2846351 | 0.24984583 | -1.1392429 | 0.25460185 | 0.68860893 |
| Txnl1 | 7328.78427 | 0.21078845 | 0.27203409 | 0.77486041 | 0.43842215 | 0.81918713 |
| Nars | 13755.286 | -0.2853065 | 0.22208742 | -1.2846588 | 0.19891156 | 0.63148061 |
| Fech | 2046.04407 | 0.3030119 | 0.24937618 | 1.21507957 | 0.22433575 | 0.65793926 |
| Nedd4l | 100.197369 | -0.0670283 | 0.57063235 | -0.1174632 | 0.906493 | 0.98159944 |

| | | | | | | |
|---|---|---|---|---|---|---|
| Lmnb1 | 3450.46067 | -0.3131259 | 0.2819354 | -1.11063 | 0.26672764 | 0.70350997 |
| C330018D2C | 675.053322 | 0.29963942 | 0.41702833 | 0.71851097 | 0.47244229 | 0.83483892 |
| Prrc1 | 1415.17846 | 0.12751013 | 0.30472032 | 0.41844972 | 0.67561834 | 0.91851958 |
| Slc12a2 | 86.4939005 | -0.2730644 | 1.15538049 | -0.2363416 | 0.81316765 | 0.95981087 |
| Fbn2 | 18.9343009 | 3.56798154 | 1.35338511 | 2.6363387 | 0.0083806 | 0.11493265 |
| Isoc1 | 1397.42266 | -0.017664 | 0.19504918 | -0.0905617 | 0.92784083 | 0.98666423 |
| Dctn4 | 2929.51507 | 0.14146302 | 0.19035284 | 0.74316208 | 0.45738356 | 0.82946634 |
| Rbm22 | 1024.095 | -0.2906782 | 0.29455294 | -0.9868455 | 0.32371839 | 0.74583177 |
| Rps14 | 31223.3333 | 0.46038267 | 0.46652707 | 0.98682947 | 0.32372624 | 0.74583177 |
| Cd74 | 528.591198 | -0.6581321 | 0.29846481 | -2.2050577 | 0.02745005 | 0.24574023 |
| Tcof1 | 4078.37536 | -0.5122581 | 0.27739826 | -1.8466521 | 0.06479758 | 0.3924103 |
| Tmx3 | 1194.60325 | 0.74253151 | 0.24708479 | 3.00516888 | 0.00265433 | 0.04897507 |
| Camk2a | 149.669341 | 0.69876253 | 0.46847441 | 1.49157035 | 0.13581182 | 0.55758574 |
| Pdgfrb | 16.1450111 | 2.16921267 | 1.17352103 | 1.84846511 | 0.06453509 | 0.39212252 |
| Csf1r | 59682.2695 | 0.02439035 | 0.2706273 | 0.09012523 | 0.9281877 | 0.98666423 |
| Hmgxb3 | 874.123219 | 0.13409805 | 0.3554803 | 0.3772306 | 0.70600224 | 0.92896305 |
| Gnaq | 1657.32114 | -0.1444927 | 0.30074483 | -0.4804496 | 0.6309077 | 0.90384049 |
| Psat1 | 15922.9182 | 0.19479776 | 0.30302654 | 0.6428406 | 0.52032754 | 0.8576517 |
| Tle4 | 921.960083 | 0.26570515 | 0.33738375 | 0.78754576 | 0.43096245 | 0.81269672 |
| Cndp2 | 11269.6426 | 0.91308688 | 0.18559051 | 4.91990073 | 8.658811498 | 6.458734210 |
| Timm21 | 1616.00253 | 0.35896824 | 0.35472068 | 1.01197437 | 0.31155031 | 0.74017637 |
| Cyb5a | 4284.75148 | 0.85564719 | 0.27981785 | 3.05787206 | 0.00222915 | 0.04328438 |
| Asrgl1 | 1067.28894 | -0.4335584 | 0.26288151 | -1.6492541 | 0.0990956 | 0.48073092 |
| Anxa1 | 1991.64686 | -1.545688 | 0.37955558 | -4.0723627 | 4.653862632 | 0.00191043 |
| Incenp | 5266.08614 | -0.1965961 | 0.25211322 | -0.7797929 | 0.4355128 | 0.81649473 |
| Fth1 | 81664.5873 | 0.48490077 | 0.13924415 | 3.48237812 | 0.00049698 | 0.01336039 |
| Rab3il1 | 20993.5915 | 0.28588836 | 0.19213266 | 1.48797381 | 0.1367578 | 0.55946682 |
| Fads3 | 87.810393 | -0.5852118 | 0.78355228 | -0.7468701 | 0.45514198 | 0.8277954 |
| Fads2 | 1044.96422 | 0.60496745 | 0.34992893 | 1.72882947 | 0.08383962 | 0.44417933 |
| Tmem138 | 511.344372 | 0.06587468 | 0.27899812 | 0.23611156 | 0.81334611 | 0.95983628 |
| Tmem216 | 380.290349 | -0.5656505 | 0.34719341 | -1.6292085 | 0.10326888 | 0.49136065 |
| Sdhaf2 | 3562.57439 | 0.31839842 | 0.36118541 | 0.88153732 | 0.37802707 | 0.78087685 |
| Cd5 | 12.347254 | 0.2447399 | 1.31707381 | 0.18582095 | 0.85258518 | 0.96798445 |
| Ms4a7 | 321.711842 | 2.65140408 | 0.53905865 | 4.91858184 | 8.717347030 | 6.462988255 |
| Ms4a4c | 340.605237 | 0.18582149 | 0.52147112 | 0.3563409 | 0.72158528 | 0.93398714 |
| Ms4a6b | 20088.5876 | 1.87439243 | 0.45366349 | 4.13168009 | 3.601213815 | 0.00152964 |
| Ms4a6d | 6835.91028 | 1.52053161 | 0.30731131 | 4.94785442 | 7.503599379 | 5.736970700 |
| Ms4a3 | 267.964644 | -5.3853504 | 0.85677645 | -6.2855958 | 3.265986955 | 6.341717210 |
| Mrpl16 | 1861.03081 | 0.18546535 | 0.31304551 | 0.5924549 | 0.55354601 | 0.87160875 |
| Osbp | 189.520619 | -0.7598939 | 0.52516323 | -1.4469672 | 0.14790612 | 0.57472133 |
| Fam111a | 8869.26609 | -0.2260242 | 0.15759595 | -1.4342009 | 0.15151494 | 0.57975674 |
| Zfp91 | 618.046938 | -0.2888485 | 0.23766901 | -1.2153391 | 0.22423677 | 0.65793926 |
| Lpxn | 2228.87578 | 0.3756817 | 0.16081446 | 2.33611893 | 0.01948504 | 0.19765383 |
| Rfk | 4184.97106 | 0.04669842 | 0.23723196 | 0.19684708 | 0.84394721 | 0.96765585 |
| Ostf1 | 12097.5665 | 0.58280002 | 0.31331712 | 1.86009631 | 0.0628719 | 0.38701669 |
| Carnmt1 | 1070.50823 | 0.05307513 | 0.22611473 | 0.23472655 | 0.81442098 | 0.95988497 |

| | | | | | | |
|---|---|---|---|---|---|---|
| Ccdc86 | 4040.94655 | -0.3051989 | 0.17555707 | -1.7384599 | 0.08212981 | 0.44123582 |
| Prpf19 | 542.534175 | -0.5304988 | 0.32331521 | -1.64081 | 0.10083686 | 0.48545348 |
| Tmem132a | 183.890048 | 0.6446176 | 0.61496076 | 1.04822557 | 0.29453469 | 0.72738877 |
| Slc15a3 | 918.099314 | 0.15736372 | 0.26298404 | 0.59837745 | 0.54958811 | 0.87093412 |
| Ddb1 | 12957.9201 | -0.0495986 | 0.17443561 | -0.2843377 | 0.77615156 | 0.94776024 |
| Fen1 | 11106.6191 | -0.2013514 | 0.16800336 | -1.1984965 | 0.23072379 | 0.6653874 |
| Zfand5 | 9091.80793 | 0.21997035 | 0.34983839 | 0.62877706 | 0.52949502 | 0.86283637 |
| Cemip2 | 852.857608 | 0.60961367 | 0.28256147 | 2.15745507 | 0.03097022 | 0.26283091 |
| Rtn3 | 12731.4893 | -0.168713 | 0.16844018 | -1.0016199 | 0.31652719 | 0.74186126 |
| Atl3 | 1026.23805 | -0.089615 | 0.40170868 | -0.2230845 | 0.82346972 | 0.9628698 |
| Naa40 | 3481.44202 | -0.2014413 | 0.24835497 | -0.8111025 | 0.41730683 | 0.8049685 |
| Lipo3 | 714.570642 | 0.02829854 | 0.19312724 | 0.14652798 | 0.8835046 | 0.97667737 |
| Otub1 | 5404.72389 | -0.0881617 | 0.18364144 | -0.4800755 | 0.63117373 | 0.90394609 |
| Ehd1 | 5850.78338 | 1.04363489 | 0.18283162 | 5.708175 | 1.141939776 | 1.569589807 |
| Atg2a | 221.75094 | -0.315724 | 0.42432064 | -0.7440694 | 0.45683448 | 0.82915295 |
| Stambpl1 | 970.754381 | -0.3488166 | 0.36217636 | -0.9631125 | 0.33549107 | 0.75359284 |
| Ppp2r5b | 209.709324 | 0.77570406 | 0.48984631 | 1.5835662 | 0.11329247 | 0.51285577 |
| Fas | 55.1757459 | 1.1120514 | 0.87713203 | 1.26782669 | 0.20485985 | 0.63901202 |
| Cdc37l1 | 723.564918 | 0.25688358 | 0.25846653 | 0.99387559 | 0.32028344 | 0.74413737 |
| Lipa | 6893.64446 | 0.34968692 | 0.1627518 | 2.14859019 | 0.0316669 | 0.26539272 |
| Ak3 | 3881.06023 | 0.03444015 | 0.1988997 | 0.17315337 | 0.86253088 | 0.9705375 |
| Rcl1 | 3134.93351 | -0.3540747 | 0.25775384 | -1.3736932 | 0.16953694 | 0.60382474 |
| Snx15 | 430.579997 | -0.0727714 | 0.35920515 | -0.20259 | 0.83945552 | 0.96757914 |
| Jak2 | 128.871676 | -0.6124551 | 0.57139092 | -1.0718671 | 0.28377972 | 0.71951357 |
| Sac3d1 | 677.254853 | -0.4943749 | 0.26284624 | -1.8808521 | 0.05999204 | 0.37827102 |
| Cdca5 | 3669.13925 | -0.1674954 | 0.1990248 | -0.8415804 | 0.40002288 | 0.79633522 |
| Zfpl1 | 964.352686 | 0.10045086 | 0.21664467 | 0.46366643 | 0.64288675 | 0.90750446 |
| Tnfrsf25 | 12.968696 | -1.857387 | 1.94398624 | -0.9554528 | 0.33934878 | 0.75683748 |
| Kif20b | 1670.51319 | -0.2274513 | 0.20770944 | -1.0950457 | 0.27349664 | 0.71048723 |
| Vps51 | 2057.99323 | 0.20865617 | 0.20525225 | 1.01658405 | 0.30935135 | 0.73890515 |
| Tm7sf2 | 419.017607 | 0.91889933 | 0.33145913 | 2.77228549 | 0.00556642 | 0.0866336 |
| Rpp30 | 2323.36173 | 0.12457712 | 0.24855026 | 0.50121501 | 0.61621981 | 0.89878056 |
| Pcgf5 | 364.193278 | -0.156113 | 0.45431681 | -0.3436215 | 0.73113092 | 0.93653187 |
| Syvn1 | 980.306817 | -0.0583738 | 0.29184233 | -0.2000183 | 0.84146629 | 0.96765585 |
| Tnks2 | 1040.79618 | -0.0831091 | 0.31011261 | -0.2679966 | 0.78870195 | 0.95225386 |
| Tjp2 | 497.433592 | 0.20463588 | 0.27756297 | 0.73725928 | 0.46096469 | 0.82987212 |
| Frmd8 | 4643.67474 | -0.081889 | 0.30216716 | -0.2710056 | 0.78638676 | 0.9519156 |
| Uhrf2 | 840.044917 | 0.25230985 | 0.2072347 | 1.21750772 | 0.2234111 | 0.65696827 |
| Slc25a45 | 5359.43873 | 0.00037589 | 0.17812957 | 0.0021102 | 0.99831631 | 0.99967651 |
| Rad9a | 1421.38337 | -0.0499875 | 0.20083152 | -0.2489028 | 0.80343599 | 0.95672466 |
| Dpf2 | 2486.72783 | -0.2362012 | 0.27572628 | -0.8566509 | 0.39163786 | 0.79018736 |
| Mrpl21 | 1767.48063 | -0.1960005 | 0.27998399 | -0.7000419 | 0.48390115 | 0.84128398 |
| Rps6kb2 | 1060.17303 | -0.0567656 | 0.21961593 | -0.2584767 | 0.79603906 | 0.9537323 |
| Ighmbp2 | 424.164476 | 0.02562286 | 0.30599776 | 0.08373546 | 0.93326676 | 0.98751425 |
| Pola2 | 4264.17358 | -0.2611837 | 0.19217413 | -1.3590993 | 0.17411512 | 0.60653633 |
| Coro1b | 13592.6575 | 0.26283001 | 0.2142658 | 1.22665405 | 0.21995264 | 0.65307784 |

| | | | | | | |
|---|---|---|---|---|---|---|
| Eif1ad | 4741.39729 | -0.3217644 | 0.19399272 | -1.6586414 | 0.09718807 | 0.47808129 |
| Chka | 1250.75277 | -0.199386 | 0.20955773 | -0.9514611 | 0.34137037 | 0.75846782 |
| Banf1 | 8929.52412 | -0.0845334 | 0.25488491 | -0.3316532 | 0.74015115 | 0.93800224 |
| Aip | 2566.00401 | 0.05450413 | 0.25432318 | 0.21431052 | 0.83030492 | 0.96466657 |
| Pitpnm1 | 135.806191 | 0.34332587 | 0.4365601 | 0.78643437 | 0.43161305 | 0.81342204 |
| Sf3b2 | 8293.97094 | -0.3965905 | 0.21271586 | -1.8644145 | 0.06226352 | 0.38520862 |
| Pold4 | 713.473754 | -0.0594179 | 0.39960488 | -0.1486917 | 0.88179695 | 0.97582756 |
| Pacs1 | 787.471288 | 0.15266631 | 0.42459854 | 0.35955449 | 0.71918032 | 0.93310883 |
| Cdk2ap2 | 2406.45285 | -0.3753906 | 0.18815485 | -1.9951149 | 0.04603035 | 0.33161911 |
| Grk2 | 1995.57047 | -0.4163823 | 0.21654921 | -1.9228067 | 0.05450433 | 0.36033644 |
| Acy3 | 76.2909941 | 0.251857 | 0.48697511 | 0.51718661 | 0.6050259 | 0.89258167 |
| Rab1b | 6336.35993 | -0.188721 | 0.17577759 | -1.0736354 | 0.28298611 | 0.71857502 |
| Yif1a | 3255.50258 | 0.46554721 | 0.28105696 | 1.65641589 | 0.09763762 | 0.47892724 |
| Cbwd1 | 508.311926 | 0.21191478 | 0.33575372 | 0.63116139 | 0.527935 | 0.86198732 |
| Aldh3b1 | 530.876776 | 0.25105114 | 0.26764823 | 0.93798917 | 0.34824997 | 0.76101142 |
| Asah2 | 27.3973818 | 1.76829326 | 1.28020742 | 1.38125528 | 0.16720048 | 0.60121803 |
| Rce1 | 1591.7026 | -0.1214805 | 0.22433232 | -0.5415202 | 0.58814909 | 0.88690478 |
| Slc29a2 | 239.495649 | -0.3645038 | 0.49440004 | -0.7372648 | 0.46096131 | 0.82987212 |
| Pcx | 217.097279 | -0.1976918 | 0.48026911 | -0.4116272 | 0.68061272 | 0.92061965 |
| Minpp1 | 2528.81296 | -0.3203015 | 0.15217042 | -2.1048869 | 0.03530115 | 0.28154973 |
| Apba1 | 15.5058741 | -2.8736759 | 1.73409569 | -1.6571611 | 0.09748692 | 0.47887037 |
| Papss2 | 30.4759477 | -1.2726626 | 1.54720982 | -0.8225534 | 0.41076203 | 0.80139357 |
| Cpt1a | 1242.28472 | 0.55505272 | 0.39788502 | 1.39500785 | 0.16301347 | 0.59424967 |
| Mrpl11 | 2322.40603 | 0.3635861 | 0.38736462 | 0.93861465 | 0.34792863 | 0.76101142 |
| Tesmin | 21.7820343 | -0.3330632 | 1.19864523 | -0.2778664 | 0.78111494 | 0.94996788 |
| Mus81 | 1252.71724 | -0.1793973 | 0.23832864 | -0.7527306 | 0.45161183 | 0.82570905 |
| Ppp6r3 | 1382.58023 | 0.00889118 | 0.26829464 | 0.0331396 | 0.97356327 | 0.99443087 |
| Efemp2 | 494.947028 | 1.01536437 | 0.33344696 | 3.04505513 | 0.00232638 | 0.04446651 |
| Fibp | 996.165326 | -0.0159833 | 0.19853827 | -0.0805049 | 0.93583569 | 0.98794094 |
| Fosl1 | 20.0915915 | -0.0082004 | 1.44604553 | -0.0056709 | 0.99547531 | 0.99884675 |
| Lrp5 | 63.3615539 | -0.5644916 | 0.99238058 | -0.5688257 | 0.56947443 | 0.87961466 |
| Drap1 | 283.970899 | 0.15200557 | 0.40490235 | 0.37541291 | 0.7073534 | 0.92950127 |
| Smarca2 | 219.280713 | 0.8215728 | 0.52374891 | 1.56863869 | 0.11673215 | 0.52252063 |
| Rnaseh2c | 1355.03321 | -0.0277238 | 0.29093767 | -0.0952911 | 0.9240836 | 0.9849488 |
| Kat5 | 1384.21809 | 0.17930724 | 0.21665384 | 0.82762086 | 0.40788525 | 0.80064439 |
| Rela | 2919.19128 | 0.20807164 | 0.47416428 | 0.43881763 | 0.66079369 | 0.91322024 |
| Ehbp1l1 | 5101.18233 | -0.2359241 | 0.28319195 | -0.8330889 | 0.40479458 | 0.79917613 |
| Fam89b | 2154.42091 | 0.34256912 | 0.17046289 | 2.00964048 | 0.04446925 | 0.32380499 |
| Scyl1 | 2295.45507 | 0.04369876 | 0.19145624 | 0.22824411 | 0.81945647 | 0.96120981 |
| Capn1 | 57.5072062 | -0.2225533 | 1.03967 | -0.2140614 | 0.83049914 | 0.96466657 |
| Smc5 | 1057.87106 | -0.150223 | 0.23744246 | -0.632671 | 0.52694851 | 0.86130085 |
| Arl2 | 1177.74209 | 0.32599239 | 0.32349306 | 1.00772606 | 0.31358599 | 0.74065408 |
| Men1 | 1984.57899 | -0.0504436 | 0.20495012 | -0.246126 | 0.80558467 | 0.95737871 |
| Map4k2 | 527.586077 | 0.19082063 | 0.42438909 | 0.44963604 | 0.6529729 | 0.9106891 |
| Sf1 | 4849.93389 | -0.2289724 | 0.24786774 | -0.9237683 | 0.35560694 | 0.7647925 |
| Rps6ka4 | 1431.99168 | 0.12821219 | 0.26422573 | 0.48523735 | 0.62750799 | 0.9022352 |

| | | | | | | |
|---|---|---|---|---|---|---|
| Prdx5 | 4853.12005 | -0.0742723 | 0.30319611 | -0.2449645 | 0.80648392 | 0.95741157 |
| Esrra | 73.4239986 | 0.01757064 | 0.5253862 | 0.03344328 | 0.9733211 | 0.99443087 |
| Gpr137 | 173.791448 | 0.18613745 | 0.37257238 | 0.49960079 | 0.6173562 | 0.89878056 |
| Bad | 1718.17549 | 0.41542592 | 0.3930418 | 1.05695098 | 0.29053396 | 0.72461526 |
| Plcb3 | 1132.05814 | 0.24139075 | 0.33471069 | 0.72119224 | 0.47079125 | 0.83381643 |
| Vegfb | 152.689774 | -0.0244436 | 0.37332954 | -0.0654747 | 0.94779604 | 0.99019648 |
| Dnajc4 | 320.539185 | 0.00755415 | 0.36372789 | 0.02076869 | 0.98343018 | 0.99564261 |
| Fermt3 | 17450.655 | 0.01110697 | 0.15241476 | 0.07287335 | 0.9419069 | 0.9896127 |
| Stip1 | 17698.4866 | -0.0644565 | 0.16117983 | -0.399904 | 0.68922722 | 0.92470267 |
| Rcor2 | 24.2420193 | 0.21956953 | 0.94312135 | 0.23281153 | 0.81590776 | 0.96063519 |
| Mark2 | 970.97036 | -0.2812354 | 0.44415601 | -0.6331905 | 0.52660926 | 0.86115578 |
| Spindoc | 54.6615806 | -0.5938197 | 0.63291862 | -0.9382244 | 0.34812909 | 0.76101142 |
| Smc3 | 3436.93673 | -0.3602734 | 0.27909944 | -1.2908424 | 0.19675835 | 0.62932233 |
| Pdcd4 | 295.255156 | -0.1105019 | 0.31029958 | -0.3561136 | 0.72175548 | 0.93398714 |
| Shoc2 | 1538.57459 | 0.16852807 | 0.20321746 | 0.82929916 | 0.40693515 | 0.79992362 |
| Gpam | 265.40286 | 0.02772713 | 0.34422281 | 0.08054995 | 0.93579987 | 0.98794094 |
| Acsl5 | 12309.3449 | -0.2909544 | 0.16726816 | -1.7394485 | 0.0819559 | 0.44107633 |
| Zdhhc6 | 4243.27023 | -0.2347672 | 0.1981481 | -1.1848068 | 0.23609385 | 0.67182855 |
| Vti1a | 965.629755 | 0.11044396 | 0.2107352 | 0.5240888 | 0.60021678 | 0.8902352 |
| Tcf7l2 | 1517.36723 | 0.30345594 | 0.27590025 | 1.09987553 | 0.27138636 | 0.70898285 |
| Hhex | 2739.16918 | -0.3457015 | 0.17230754 | -2.0063053 | 0.04482369 | 0.32541732 |
| Cep55 | 1940.82557 | -0.1361894 | 0.24678143 | -0.5518626 | 0.58104248 | 0.88314588 |
| Rbp4 | 39.9693736 | 1.2657322 | 1.18488871 | 1.06822876 | 0.28541733 | 0.7212374 |
| Eif3a | 9233.54627 | -0.272101 | 0.30961383 | -0.87884 | 0.37948804 | 0.78211296 |
| Fam45a | 2751.45279 | 0.61807823 | 0.28260389 | 2.18708323 | 0.02873645 | 0.25235676 |
| Prdx3 | 7924.66252 | -0.0532733 | 0.2269557 | -0.23473 | 0.8144183 | 0.95988497 |
| Noc3l | 441.157192 | -0.4524613 | 0.28803656 | -1.5708469 | 0.11621822 | 0.52115009 |
| Hells | 4142.39946 | -0.3334621 | 0.24527248 | -1.3595576 | 0.17396996 | 0.60653633 |
| Sorbs1 | 55.9108141 | 0.85228981 | 1.04289963 | 0.81723091 | 0.41379648 | 0.80395688 |
| Aldh18a1 | 3255.10423 | -0.2818581 | 0.18559263 | -1.5186921 | 0.12884002 | 0.54471351 |
| Tctn3 | 328.072557 | 0.32546044 | 0.37380171 | 0.87067669 | 0.38393071 | 0.78704643 |
| Ccnj | 367.046103 | -0.0030607 | 0.35907432 | -0.0085238 | 0.99319905 | 0.99809447 |
| Tm9sf3 | 2497.32524 | -0.0371569 | 0.25497279 | -0.145729 | 0.88413534 | 0.97719801 |
| Pik3ap1 | 6794.41774 | -0.1118572 | 0.25634529 | -0.4363537 | 0.66258013 | 0.91395288 |
| Lcor | 336.127876 | 0.06423692 | 0.33121819 | 0.19394139 | 0.84622177 | 0.96765585 |
| Smndc1 | 1109.7231 | -0.2187204 | 0.2019864 | -1.082847 | 0.27887633 | 0.71377266 |
| Mxi1 | 238.586438 | 0.50840452 | 0.36433938 | 1.39541466 | 0.16289083 | 0.59424967 |
| Add3 | 2454.41956 | 0.02072489 | 0.3251719 | 0.06373519 | 0.94918109 | 0.99028544 |
| Xpnpep1 | 5735.37388 | -0.0374227 | 0.19102588 | -0.1959036 | 0.8446856 | 0.96765585 |
| Trim8 | 550.187936 | -0.0630155 | 0.33944519 | -0.1856425 | 0.85272512 | 0.96798445 |
| Arl3 | 392.347552 | 0.41761387 | 0.40435397 | 1.03279281 | 0.30170087 | 0.73346745 |
| Sfxn2 | 959.954801 | 0.26909309 | 0.26522903 | 1.01456879 | 0.31031143 | 0.7399039 |
| Maoa | 1717.39548 | 0.31687502 | 0.18678756 | 1.69644603 | 0.08980144 | 0.45791624 |
| Fundc1 | 2659.36884 | 0.70308413 | 0.31916719 | 2.20287097 | 0.02760384 | 0.24636643 |
| Nt5c2 | 3044.24311 | 0.433898 | 0.2089967 | 2.07609977 | 0.03788472 | 0.29387688 |
| Msr1 | 1932.43064 | 1.36256957 | 0.23754134 | 5.73613658 | 9.686056002 | 1.377785152 |

| | | | | | | |
|---|---|---|---|---|---|---|
| Pdcd11 | 805.78245 | -0.8911873 | 0.34879389 | -2.5550543 | 0.01061712 | 0.13331258 |
| Taf5 | 114.993732 | -0.7280269 | 0.48372195 | -1.5050525 | 0.13231057 | 0.55203111 |
| Pcgf6 | 814.155575 | 0.27505736 | 0.28481512 | 0.96574002 | 0.33417429 | 0.75320272 |
| 5430427O19 | 715.393767 | -0.3765568 | 0.20327511 | -1.8524491 | 0.06396136 | 0.39085634 |
| Gk | 616.028594 | -0.2715618 | 0.35279637 | -0.7697409 | 0.44145362 | 0.81967485 |
| Slk | 526.65999 | -0.3613777 | 0.28317715 | -1.2761542 | 0.20190102 | 0.63623266 |
| Sfr1 | 5121.9425 | -0.1435484 | 0.15939056 | -0.9006078 | 0.36779691 | 0.77386646 |
| Gsto1 | 4357.69792 | -0.4532703 | 0.1762077 | -2.5723637 | 0.01010067 | 0.12992799 |
| Gsto2 | 24.7542376 | 0.29203578 | 1.4453652 | 0.20204982 | 0.83987778 | 0.96762336 |
| Casp7 | 1293.00748 | -0.1418495 | 0.23054406 | -0.6152814 | 0.5383689 | 0.86663962 |
| Dclre1a | 619.144007 | 0.0772259 | 0.26349756 | 0.29308013 | 0.7694609 | 0.94401697 |
| Nhlrc2 | 1434.97272 | -0.1055427 | 0.20466578 | -0.5156832 | 0.60607571 | 0.89258167 |
| Ablim1 | 14.0039726 | 1.82827301 | 2.12500771 | 0.86036065 | 0.38959027 | 0.78944141 |
| Trub1 | 953.656755 | -0.2192216 | 0.3066026 | -0.7150024 | 0.47460756 | 0.83578199 |
| Hspa12a | 26.173276 | -0.560086 | 1.67775443 | -0.3338308 | 0.73850728 | 0.93800224 |
| Slc18a2 | 244.241331 | -0.1361435 | 0.36468671 | -0.3733162 | 0.7089131 | 0.92998755 |
| 3110040N11 | 646.225565 | -0.0547813 | 0.27724797 | -0.1975896 | 0.84336619 | 0.96765585 |
| Btbd1 | 717.540808 | -0.0846103 | 0.28366366 | -0.2982768 | 0.76549194 | 0.94347227 |
| Hdgfl3 | 23.6608938 | 1.81785428 | 1.14271556 | 1.59081958 | 0.11165019 | 0.50906326 |
| P4hb | 39034.4763 | 0.30125594 | 0.27470149 | 1.09666658 | 0.27278717 | 0.71005689 |
| Arhgdia | 29237.5933 | -0.1519795 | 0.19725717 | -0.7704639 | 0.44102475 | 0.81967485 |
| Ints4 | 2397.42379 | -0.0261773 | 0.15421713 | -0.1697434 | 0.86521197 | 0.9705375 |
| Alyref | 475.094332 | -0.4170114 | 0.28440036 | -1.466283 | 0.14257121 | 0.56828728 |
| Anapc11 | 1917.68893 | 0.25559674 | 0.33526376 | 0.76237509 | 0.44583617 | 0.82166369 |
| Pcyt2 | 2176.85427 | 0.03253682 | 0.21090357 | 0.15427345 | 0.87739413 | 0.97458574 |
| Sirt7 | 2787.7842 | 0.07069359 | 0.26963111 | 0.26218631 | 0.7931778 | 0.95310033 |
| Tollip | 4553.78931 | 0.06736571 | 0.22892374 | 0.2942714 | 0.76855053 | 0.94401697 |
| Pycr1 | 222.589255 | -0.3125675 | 0.46510147 | -0.6720416 | 0.50155722 | 0.84968141 |
| Myadml2 | 12.9724384 | 0.38963362 | 1.56703797 | 0.24864338 | 0.80363665 | 0.95680296 |
| Aspscr1 | 1999.71518 | -0.1803683 | 0.20865081 | -0.8644505 | 0.38734044 | 0.78911204 |
| Cenpx | 2597.79852 | 0.36860038 | 0.35751717 | 1.03100052 | 0.30254058 | 0.73372823 |
| Lrrc45 | 685.449226 | -0.1753315 | 0.33811051 | -0.5185627 | 0.6040657 | 0.89257579 |
| Mob2 | 1263.35475 | -0.0931587 | 0.18714278 | -0.4977949 | 0.61862859 | 0.89878056 |
| Cbr2 | 19.3306501 | 0.71489476 | 1.14475347 | 0.62449669 | 0.53230144 | 0.86442675 |
| Maged1 | 3643.21065 | 0.91994611 | 0.26269133 | 3.50200412 | 0.00046177 | 0.01267531 |
| Fasn | 1390.02072 | -0.6174874 | 0.40914764 | -1.5092043 | 0.13124657 | 0.54970237 |
| Arhgap19 | 1509.50018 | 0.79706869 | 0.26398561 | 3.01936415 | 0.00253306 | 0.04723614 |
| Dus1l | 3778.63266 | -0.0212318 | 0.22966871 | -0.0924452 | 0.92634429 | 0.98624627 |
| Gps1 | 9691.26033 | -0.1004832 | 0.19222977 | -0.5227242 | 0.6011662 | 0.89086204 |
| Zdhhc16 | 2438.6167 | -0.0461311 | 0.26570776 | -0.1736158 | 0.86216741 | 0.9705375 |
| Rfng | 795.611291 | 0.00900873 | 0.31696948 | 0.02842144 | 0.97732603 | 0.99483019 |
| Mms19 | 1964.44127 | 0.23700718 | 0.25456896 | 0.93101369 | 0.35184648 | 0.76348958 |
| Slc16a3 | 10007.0127 | -0.3444989 | 0.16622131 | -2.0725317 | 0.03821589 | 0.29537366 |
| Csnk1d | 4415.82648 | 0.00353645 | 0.30196205 | 0.01171157 | 0.99065573 | 0.9966603 |
| Ogfod3 | 424.372922 | 0.29285227 | 0.30681174 | 0.95450151 | 0.33982985 | 0.75726534 |
| Ubtd1 | 876.441873 | 0.45687867 | 0.2000495 | 2.28382807 | 0.02238164 | 0.21507822 |

| | | | | | | |
|---|---|---|---|---|---|---|
| Wdr45b | 4854.09435 | 0.54696975 | 0.22715044 | 2.40796251 | 0.01604183 | 0.17615771 |
| Pi4k2a | 383.572962 | -0.1051386 | 0.35884381 | -0.2929927 | 0.7695277 | 0.94401697 |
| R3hcc1l | 1552.30131 | 0.65215511 | 0.28483758 | 2.28956841 | 0.02204635 | 0.21353363 |
| Hps1 | 1104.16551 | -0.2973528 | 0.27119167 | -1.0964671 | 0.27287439 | 0.71005689 |
| Got1 | 2840.67224 | 0.00380238 | 0.18256663 | 0.02082735 | 0.98338338 | 0.99564261 |
| Entpd7 | 410.080094 | -0.035269 | 0.43930991 | -0.0802828 | 0.93601238 | 0.98802652 |
| Cutc | 581.558131 | -0.147049 | 0.29984251 | -0.4904207 | 0.62383623 | 0.90017327 |
| Dnmbp | 111.015046 | 0.78508656 | 0.57989117 | 1.35385155 | 0.17578374 | 0.60763832 |
| Cpn1 | 8.00913644 | 2.38555349 | 1.60945616 | 1.48221092 | 0.13828418 | 0.56085884 |
| Erlin1 | 2330.30608 | -0.0216544 | 0.18285165 | -0.1184261 | 0.90573008 | 0.98144617 |
| Chuk | 1152.59198 | 0.10217809 | 0.35319296 | 0.28929821 | 0.77235318 | 0.94595479 |
| Cwf19l1 | 941.127858 | -0.1853351 | 0.23330049 | -0.7944051 | 0.42695962 | 0.81021713 |
| Scd2 | 2555.34172 | -0.0051662 | 0.18813033 | -0.0274609 | 0.97809211 | 0.99483019 |
| Ndufb8 | 15807.5061 | 0.75817646 | 0.45698622 | 1.65907947 | 0.09709978 | 0.47799662 |
| Mrpl43 | 2400.55548 | -0.085301 | 0.29035521 | -0.2937815 | 0.76892487 | 0.94401697 |
| Twnk | 566.632487 | -0.7892311 | 0.32033739 | -2.4637495 | 0.01374922 | 0.15882359 |
| Sfxn3 | 282.630781 | 0.44167761 | 0.44741284 | 0.98718135 | 0.32355374 | 0.74583177 |
| Btrc | 1262.40103 | 0.04001404 | 0.20638032 | 0.19388498 | 0.84626595 | 0.96765585 |
| Poll | 811.361981 | -0.2372643 | 0.37284024 | -0.6363698 | 0.52453544 | 0.85967874 |
| Oga | 786.902782 | 0.28365251 | 0.22550862 | 1.25783444 | 0.20845164 | 0.6418276 |
| Ldb1 | 211.376949 | -0.4222947 | 0.31864036 | -1.3253021 | 0.18507098 | 0.61717463 |
| Gbf1 | 957.341281 | 0.36163318 | 0.3327549 | 1.08678545 | 0.27713164 | 0.71247429 |
| Nfkb2 | 869.788659 | 0.46073942 | 0.41263295 | 1.11658415 | 0.26417217 | 0.70060794 |
| Fbxl15 | 191.909504 | 0.36256503 | 0.30696498 | 1.18112833 | 0.23755175 | 0.67389384 |
| Mfsd13a | 586.205657 | -0.0495881 | 0.28893887 | -0.1716216 | 0.86373504 | 0.9705375 |
| Actr1a | 5586.66654 | 0.03331159 | 0.14076595 | 0.23664523 | 0.81293203 | 0.95974923 |
| Sufu | 635.449743 | -0.1562938 | 0.36882171 | -0.4237651 | 0.67173714 | 0.91719387 |
| Hexa | 24591.7347 | 0.60785718 | 0.22341572 | 2.72074491 | 0.0065135 | 0.09752711 |
| Arih1 | 399.509567 | -0.2994613 | 0.40760198 | -0.7346904 | 0.46252805 | 0.8310966 |
| Bbs4 | 352.731117 | 1.04346214 | 0.39533353 | 2.63944762 | 0.00830413 | 0.11452578 |
| Adpgk | 2661.44804 | -0.3215036 | 0.16383028 | -1.9624188 | 0.04971374 | 0.34225366 |
| Parp6 | 200.821432 | -0.2752542 | 0.4169301 | -0.6601927 | 0.50913017 | 0.85329351 |
| Limd1 | 1764.29197 | 0.12652933 | 0.25799216 | 0.49043868 | 0.62382351 | 0.90017327 |
| Sacm1l | 2635.83574 | 0.66472351 | 0.20729962 | 3.20658333 | 0.00134321 | 0.02892876 |
| Fyco1 | 287.935835 | -0.0971053 | 0.53887205 | -0.1802011 | 0.85699473 | 0.96936093 |
| Lztfl1 | 483.333997 | 0.5499722 | 0.4579292 | 1.20099834 | 0.22975185 | 0.6643583 |
| Tbl1x | 3151.05414 | -0.1497019 | 0.28014485 | -0.5343733 | 0.59308327 | 0.88797037 |
| Ribc1 | 99.4064889 | -0.8245993 | 0.43951616 | -1.8761523 | 0.06063437 | 0.37940675 |
| Hsd17b10 | 2726.58487 | 0.40710469 | 0.37846758 | 1.07566595 | 0.28207668 | 0.71778152 |
| Huwe1 | 1014.16379 | 0.02634969 | 0.40522133 | 0.06502542 | 0.94815376 | 0.99019648 |
| Fam120c | 42.8735993 | -1.1905111 | 0.62408871 | -1.9075992 | 0.05644304 | 0.36517938 |
| Tsr2 | 1675.20929 | 0.04738053 | 0.24118278 | 0.19645071 | 0.84425741 | 0.96765585 |
| Fgd1 | 10.2472645 | -1.3437056 | 2.07588472 | -0.647293 | 0.51744234 | 0.85694744 |
| Gnl3l | 1690.94674 | -0.5260252 | 0.24220777 | -2.1717931 | 0.02987127 | 0.25769767 |
| Maged2 | 2541.68072 | 1.01847191 | 0.30435278 | 3.3463532 | 0.00081882 | 0.01999327 |
| Apex2 | 739.56555 | 0.16191434 | 0.24162329 | 0.67011064 | 0.50278726 | 0.85012672 |

| | | | | | | |
|---|---|---|---|---|---|---|
| Pfkfb1 | 461.117783 | 0.78118024 | 0.44863304 | 1.74124546 | 0.08164056 | 0.44034788 |
| Abhd6 | 512.926099 | 0.63848484 | 0.3538628 | 1.8043288 | 0.07117978 | 0.41122734 |
| Flnb | 336.341538 | -0.4190775 | 0.30239796 | -1.3858476 | 0.16579344 | 0.5992759 |
| Dnase1l3 | 202.818195 | 1.43313899 | 0.56819065 | 2.52228539 | 0.01165951 | 0.14220414 |
| Polr3a | 649.816469 | -0.6633709 | 0.36718966 | -1.8066166 | 0.07082208 | 0.41080052 |
| Sat1 | 27350.1957 | 0.49916502 | 0.32495215 | 1.53611852 | 0.12450931 | 0.53617731 |
| Acot9 | 4132.27031 | -0.0520777 | 0.31266884 | -0.1665587 | 0.86771729 | 0.97155457 |
| Prdx4 | 4557.38392 | 0.63003001 | 0.3510447 | 1.79472875 | 0.07269693 | 0.41489174 |
| Rps24 | 13415.3804 | 0.46817626 | 0.46731454 | 1.00184398 | 0.31641895 | 0.74186126 |
| Ptprj | 882.012399 | 0.03791536 | 0.33496741 | 0.11319119 | 0.90987898 | 0.98262295 |
| Banp | 1092.66645 | -0.3538098 | 0.19557709 | -1.8090557 | 0.07044236 | 0.41006309 |
| Sp4 | 88.2520579 | 1.06122795 | 0.57957647 | 1.83104041 | 0.06709451 | 0.39891584 |
| Atp10a | 129.455833 | -0.8086925 | 0.67852333 | -1.1918418 | 0.23332327 | 0.66891108 |
| Ube3a | 2613.6606 | -0.0769541 | 0.36557904 | -0.2104993 | 0.83327802 | 0.96565375 |
| Kdm5c | 665.165079 | -9.80E-05 | 0.27658708 | -0.0003542 | 0.9997174 | 0.99979912 |
| Sbds | 2249.1309 | 0.00161425 | 0.21540762 | 0.00749391 | 0.99402078 | 0.99875616 |
| Rabgef1 | 1391.16245 | 0.37535549 | 0.54093735 | 0.69389827 | 0.48774601 | 0.84309692 |
| Rdh5 | 46.4984374 | 1.0274909 | 0.71666746 | 1.43370666 | 0.15165599 | 0.57985599 |
| Cd63 | 7193.60935 | 1.00319848 | 0.34199262 | 2.93339223 | 0.0033528 | 0.05884477 |
| Gdf11 | 32.7264562 | 1.14288002 | 0.89879736 | 1.27156584 | 0.20352742 | 0.63809363 |
| Ormdl2 | 3675.36834 | 0.2294732 | 0.30022854 | 0.76432841 | 0.44467156 | 0.82108184 |
| Dnajc14 | 1937.5919 | 0.16499261 | 0.23008032 | 0.71710874 | 0.473307 | 0.83483892 |
| Mmp19 | 139.018552 | -1.2550102 | 0.5406511 | -2.321294 | 0.02027098 | 0.20128169 |
| Dgka | 207.577223 | -0.1805956 | 0.41099589 | -0.4394098 | 0.66036464 | 0.91302324 |
| Cdk2 | 5070.07008 | -0.2122336 | 0.17896291 | -1.1859086 | 0.23565841 | 0.6716704 |
| Pmel | 56.950983 | -1.3374699 | 0.85931868 | -1.5564306 | 0.11960572 | 0.52706657 |
| Rps26 | 14100.1999 | -0.3325163 | 0.19310231 | -1.7219697 | 0.085075 | 0.44637738 |
| Pa2g4 | 17254.9057 | -0.4259762 | 0.13507653 | -3.1535914 | 0.00161275 | 0.03378208 |
| Esyt1 | 1265.71278 | -0.9193629 | 0.21458605 | -4.2843552 | 1.832699120 | 0.00085424 |
| Smarcc2 | 838.042977 | 0.11537894 | 0.24378507 | 0.47328141 | 0.63601242 | 0.90531997 |
| Chmp6 | 2826.71711 | 0.2883007 | 0.19115372 | 1.50821395 | 0.13149978 | 0.54977334 |
| Baiap2 | 718.185176 | 0.37739545 | 0.3326896 | 1.13437704 | 0.25663643 | 0.6916575 |
| Rnf41 | 1873.52245 | 0.0270368 | 0.2291699 | 0.1179771 | 0.9060858 | 0.98150603 |
| Nabp2 | 1200.96829 | -0.417987 | 0.19333184 | -2.1620185 | 0.03061675 | 0.26095692 |
| Aatk | 32.3558178 | 0.98255532 | 1.33174036 | 0.73779797 | 0.46063723 | 0.82987212 |
| Tepsin | 460.322612 | 0.10238672 | 0.30834907 | 0.33204809 | 0.73985295 | 0.93800224 |
| Cnpy2 | 2826.52357 | 0.73516497 | 0.37748856 | 1.9475159 | 0.05147292 | 0.34938799 |
| Faap100 | 918.112159 | 0.32291972 | 0.21729983 | 1.48605606 | 0.13726429 | 0.55981285 |
| Atp5b | 91503.8555 | -0.0934704 | 0.16569915 | -0.5640972 | 0.57268801 | 0.88035336 |
| Prim1 | 3203.89882 | 0.30302553 | 0.31254853 | 0.96953112 | 0.33228026 | 0.75138937 |
| Nab2 | 250.235683 | 0.4586783 | 0.82929303 | 0.55309557 | 0.58019797 | 0.88286755 |
| Shmt2 | 14139.2727 | -0.583518 | 0.19509134 | -2.9909987 | 0.00278067 | 0.05092199 |
| R3hdm2 | 392.124959 | -0.1483798 | 0.40400425 | -0.3672729 | 0.71341548 | 0.93235217 |
| Ddit3 | 1110.84084 | -0.0451733 | 0.43589947 | -0.1036324 | 0.91746104 | 0.98390186 |
| Mbd6 | 66.636499 | -0.3112243 | 0.72907538 | -0.4268753 | 0.66947017 | 0.91657847 |
| Dctn2 | 3507.0602 | 0.35870957 | 0.30648984 | 1.17037999 | 0.24184808 | 0.67708722 |

| | | | | | | |
|---|---|---|---|---|---|---|
| Ttc4 | 2093.15506 | 0.32929733 | 0.31379284 | 1.04940996 | 0.29398948 | 0.72683372 |
| Pip4k2c | 891.832304 | 0.03629391 | 0.19392049 | 0.18715874 | 0.85153618 | 0.9679302 |
| Katnal2 | 41.6004296 | 0.73233488 | 1.2804437 | 0.57193837 | 0.56736373 | 0.87929304 |
| Hdhd2 | 1904.06875 | 0.14731553 | 0.20779064 | 0.70896134 | 0.47834847 | 0.83822329 |
| Pias2 | 831.557628 | -0.1905523 | 0.38144614 | -0.4995522 | 0.61739043 | 0.89878056 |
| Atp5a1 | 46994.2466 | -0.2239181 | 0.13155994 | -1.7020236 | 0.08875094 | 0.45521604 |
| Pstpip2 | 1630.81518 | -0.2757913 | 0.21025351 | -1.3117087 | 0.18961845 | 0.62019167 |
| Atp23 | 644.086746 | 0.12816813 | 0.4251295 | 0.30148021 | 0.76304834 | 0.94298237 |
| Usp33 | 1305.38797 | 0.3683419 | 0.21357387 | 1.72465808 | 0.08458911 | 0.44544923 |
| Clns1a | 9290.20729 | -0.0136661 | 0.24873425 | -0.0549426 | 0.95618417 | 0.99215259 |
| Paip1 | 406.394603 | -0.1503148 | 0.27187357 | -0.5528849 | 0.58034224 | 0.88286755 |
| Nnt | 955.136653 | -0.0513142 | 0.22704096 | -0.2260131 | 0.82119124 | 0.96205421 |
| Paox | 1787.02284 | 0.96503053 | 0.21183521 | 4.5555719 | 5.224324091 | 0.00030004 |
| Echs1 | 3778.20489 | 0.47347304 | 0.38704801 | 1.22329279 | 0.22121913 | 0.65469944 |
| Fuom | 66.6865602 | 0.62497425 | 0.59427672 | 1.05165528 | 0.29295774 | 0.72604376 |
| Zfp511 | 635.309304 | 0.00790619 | 0.23799328 | 0.03322024 | 0.97349895 | 0.99443087 |
| Adam8 | 630.122054 | -1.4649817 | 0.36205908 | -4.0462503 | 5.204458191 | 0.00208059 |
| Tubgcp2 | 3430.08516 | -0.0827562 | 0.21128758 | -0.3916756 | 0.69529789 | 0.92616369 |
| Inpp5a | 63.1691857 | -0.3078429 | 0.79917004 | -0.3852032 | 0.70008686 | 0.92669304 |
| Bet1l | 663.689021 | 0.18515847 | 0.2654249 | 0.6975927 | 0.48543195 | 0.84183502 |
| Ric8a | 2621.0612 | -0.0842149 | 0.20465804 | -0.4114909 | 0.68071262 | 0.92061965 |
| Sirt3 | 423.074303 | -0.0419405 | 0.3551831 | -0.1180813 | 0.90600326 | 0.98150353 |
| Psmd13 | 9492.83226 | 0.19226758 | 0.35001805 | 0.54930761 | 0.58279436 | 0.88343755 |
| Ifitm5 | 16.544195 | 1.29291453 | 1.42332142 | 0.90837847 | 0.3636783 | 0.77036824 |
| Ifitm1 | 259.868638 | -1.7148179 | 0.42660011 | -4.0197316 | 5.826449039 | 0.0022918 |
| Ifitm3 | 3534.26932 | -1.0561084 | 0.19620858 | -5.3825803 | 7.342561540 | 7.677056010 |
| Sigirr | 394.665368 | -0.0881664 | 0.43060152 | -0.2047517 | 0.83776611 | 0.96683245 |
| Ptdss2 | 1173.07702 | 0.11218649 | 0.21766468 | 0.51540971 | 0.60626675 | 0.89258167 |
| Irf7 | 284.040948 | -0.6504492 | 0.3251213 | -2.0006354 | 0.0454317 | 0.32924523 |
| Hras | 338.803976 | -0.37893 | 0.36731415 | -1.0316239 | 0.30224836 | 0.73372823 |
| Taldo1 | 13796.5016 | -0.4450636 | 0.19571572 | -2.2740312 | 0.02296411 | 0.21709424 |
| Tmem80 | 268.475672 | 0.06512747 | 0.52170972 | 0.12483469 | 0.90065442 | 0.98043702 |
| Pidd1 | 570.36441 | -0.227637 | 0.40400163 | -0.5634555 | 0.57312472 | 0.88035336 |
| Rplp2 | 7392.31526 | -0.0208081 | 0.31982383 | -0.0650612 | 0.94812525 | 0.99019648 |
| Pnpla2 | 1024.06147 | -0.1769646 | 0.31299363 | -0.5653935 | 0.57180611 | 0.88035336 |
| Cd151 | 159.80747 | 1.04292127 | 0.46999829 | 2.21898951 | 0.02648744 | 0.23919285 |
| Tspan4 | 3238.76354 | 0.32322735 | 0.25459061 | 1.26959651 | 0.20422839 | 0.63846817 |
| Chid1 | 3313.5842 | 0.47354217 | 0.22203038 | 2.13278098 | 0.0329427 | 0.26976292 |
| Tmem192 | 2017.39516 | 0.29959614 | 0.18137731 | 1.651784 | 0.09857859 | 0.48025164 |
| Apool | 1577.18783 | 0.38571316 | 0.25075238 | 1.53822331 | 0.12399402 | 0.53550701 |
| Zfp711 | 199.604839 | 1.25538771 | 0.55684849 | 2.25445114 | 0.0241678 | 0.22455944 |
| Chm | 896.412866 | 0.626224 | 0.23640014 | 2.64900008 | 0.00807303 | 0.11159027 |
| Crcp | 2359.58272 | 0.12912054 | 0.26525956 | 0.48677054 | 0.62642095 | 0.90195474 |
| Asl | 2504.29869 | 0.38795388 | 0.17913221 | 2.16574053 | 0.03033102 | 0.26001357 |
| Gusb | 26928.5827 | 0.13349629 | 0.18620187 | 0.71694387 | 0.47340873 | 0.83483892 |
| Sumf2 | 1377.89271 | 0.08347668 | 0.19645212 | 0.42492125 | 0.67089408 | 0.91711586 |

| | | | | | | |
|---|---|---|---|---|---|---|
| Tm9sf2 | 12566.0535 | 0.07143971 | 0.19218402 | 0.37172553 | 0.71009721 | 0.93077435 |
| Clybl | 1129.85268 | 0.56338263 | 0.4161179 | 1.35390146 | 0.17576782 | 0.60763832 |
| Farp1 | 94.5715452 | 1.66022083 | 0.70926521 | 2.34076169 | 0.01924445 | 0.19585467 |
| Dock9 | 43.0126895 | -0.037408 | 1.43789767 | -0.0260157 | 0.97924478 | 0.99483019 |
| Tnrc6c | 207.817503 | -0.7097641 | 0.48038301 | -1.4774963 | 0.13954266 | 0.56329174 |
| Tmc6 | 1439.94895 | -0.3770888 | 0.28678194 | -1.3148973 | 0.18854443 | 0.61935434 |
| Tk1 | 5303.83152 | 0.36625884 | 0.25357325 | 1.44439069 | 0.14862911 | 0.57573777 |
| Cant1 | 1470.44041 | -0.1993852 | 0.25045295 | -0.7960984 | 0.42597487 | 0.81021713 |
| Rbfox3 | 11.6176715 | 4.07441567 | 1.79719251 | 2.2671003 | 0.0233841 | 0.22004437 |
| Cbx2 | 110.472335 | -0.2177896 | 0.40803573 | -0.5337513 | 0.5935136 | 0.88797037 |
| Cbx8 | 262.245407 | 0.18189638 | 0.46316677 | 0.3927233 | 0.69452386 | 0.92609254 |
| Gaa | 3143.15214 | 0.13998326 | 0.25434566 | 0.55036624 | 0.5820682 | 0.88321857 |
| Eif4a3 | 11706.5266 | -0.2453841 | 0.16913886 | -1.4507852 | 0.14683969 | 0.57365079 |
| Rptor | 898.939219 | 0.18543413 | 0.3246511 | 0.57117973 | 0.56787782 | 0.87929304 |
| Tma16 | 931.993826 | -0.3137979 | 0.23905935 | -1.3126359 | 0.18930565 | 0.61985441 |
| Dach2 | 17.157848 | 1.04809084 | 1.56941377 | 0.66782314 | 0.50424649 | 0.85012672 |
| Klhl4 | 132.11558 | 4.44262677 | 3.03989602 | 1.46144037 | 0.14389463 | 0.56966439 |
| Zfp202 | 255.455177 | -0.6874541 | 0.66848509 | -1.0283761 | 0.30377294 | 0.73436097 |
| Copg2 | 294.572842 | 0.29575644 | 0.43135735 | 0.68564135 | 0.4929393 | 0.84609645 |
| Mkln1 | 3799.14498 | 0.30780763 | 0.18871348 | 1.63108449 | 0.10287249 | 0.48985567 |
| Bach1 | 747.587372 | 0.46085238 | 0.3885785 | 1.18599556 | 0.23562405 | 0.6716704 |
| Cct8 | 41945.0715 | -0.1759785 | 0.14360546 | -1.2254304 | 0.22041308 | 0.65381019 |
| Usp16 | 966.949439 | -0.1303844 | 0.28936531 | -0.4505874 | 0.65228698 | 0.91037383 |
| Phf6 | 895.709937 | 0.18494738 | 0.28347677 | 0.65242515 | 0.51412694 | 0.8551074 |
| Hprt | 13064.3664 | 0.34276565 | 0.34980329 | 0.97988113 | 0.3271448 | 0.74811039 |
| Ccdc51 | 519.551787 | -0.3040338 | 0.28711191 | -1.0589383 | 0.28962788 | 0.72395811 |
| Atrip | 618.601982 | -0.2272182 | 0.25333496 | -0.8969082 | 0.36976793 | 0.77508072 |
| Shisa5 | 4400.21264 | -0.2560939 | 0.18831923 | -1.3598924 | 0.17386398 | 0.60653633 |
| Pfkfb4 | 809.552592 | 0.43530831 | 0.25448642 | 1.71053648 | 0.08716671 | 0.45109601 |
| Uqcrc1 | 24587.8384 | -0.0588832 | 0.18131316 | -0.3247597 | 0.74536296 | 0.93917909 |
| Arhgef9 | 41.6213643 | 0.24073198 | 0.87519007 | 0.27506251 | 0.78326821 | 0.95081654 |
| Rps6ka6 | 120.038441 | -0.0454157 | 0.43509483 | -0.1043813 | 0.9168668 | 0.98389485 |
| Tmem47 | 27.7226463 | 0.25348177 | 1.1004012 | 0.23035396 | 0.81781674 | 0.96079676 |
| Alox5 | 692.581707 | 0.1833271 | 0.21297545 | 0.86078983 | 0.38935381 | 0.78942657 |
| March8 | 702.110194 | 0.44399147 | 0.27788858 | 1.59773197 | 0.11010264 | 0.50692212 |
| Wdr73 | 1502.61315 | -0.0734856 | 0.25208903 | -0.2915066 | 0.77066387 | 0.94454775 |
| Nmb | 51.6133872 | 0.13488232 | 0.6069074 | 0.2222453 | 0.82412293 | 0.96290555 |
| Sec11a | 4758.79682 | 0.31195579 | 0.31247393 | 0.99834182 | 0.31811363 | 0.74336577 |
| Pigq | 2421.16205 | -0.4162201 | 0.21624464 | -1.924765 | 0.05425877 | 0.35994983 |
| Rab40c | 306.43371 | 0.63278159 | 0.33001954 | 1.91740644 | 0.05518632 | 0.36178683 |
| Mettl26 | 567.206663 | -0.0791544 | 0.35918059 | -0.220375 | 0.82557911 | 0.96326388 |
| Mcrip2 | 707.164118 | 0.25334259 | 0.39992111 | 0.63348141 | 0.52641932 | 0.86111527 |
| Rhot2 | 624.093919 | -0.442018 | 0.26809089 | -1.6487619 | 0.09919643 | 0.48078421 |
| Jmjd8 | 1267.01098 | -0.3448368 | 0.25490342 | -1.3528135 | 0.17611522 | 0.60763832 |
| Wdr24 | 977.858896 | 0.16311995 | 0.31758464 | 0.51362671 | 0.607513 | 0.89332558 |
| Prps2 | 5621.74928 | 0.32855931 | 0.15623818 | 2.10293873 | 0.03547113 | 0.28231508 |

| | | | | | | |
|---|---|---|---|---|---|---|
| Sdc3 | 287.561668 | 0.87469244 | 0.39521319 | 2.21321674 | 0.02688271 | 0.24198392 |
| Hadha | 5001.25505 | 0.03054396 | 0.23668246 | 0.12905036 | 0.8973178 | 0.97984623 |
| Il6 | 36.3509708 | -1.1009384 | 1.3486861 | -0.8163044 | 0.41432608 | 0.80395688 |
| Tyms | 6579.76225 | 0.4061796 | 0.34932543 | 1.16275415 | 0.24492926 | 0.68039867 |
| Hspa4l | 712.170383 | -0.4189249 | 0.28624902 | -1.4634982 | 0.1433311 | 0.56846902 |
| Plk4 | 3095.50726 | 0.0273604 | 0.17166012 | 0.15938706 | 0.87336393 | 0.97344599 |
| Mfsd8 | 639.874494 | 0.23194156 | 0.49439777 | 0.46913958 | 0.63896987 | 0.90615794 |
| Larp1b | 24.4422509 | -0.4468702 | 0.94229553 | -0.4742357 | 0.63533183 | 0.90531997 |
| Jade1 | 430.48735 | -0.3035051 | 0.40395618 | -0.7513318 | 0.45245301 | 0.82609816 |
| D3Ertd751e | 397.565623 | 0.31170434 | 0.30032059 | 1.03790532 | 0.29931414 | 0.73113229 |
| Ly96 | 462.54006 | 0.67424202 | 0.43003319 | 1.5678837 | 0.11690828 | 0.52252063 |
| Itih5 | 15.9391672 | -4.4358771 | 1.3038106 | -3.4022404 | 0.00066836 | 0.01714051 |
| Atp5c1 | 23639.9912 | 0.11451509 | 0.32641838 | 0.35082304 | 0.72572111 | 0.93544211 |
| Taf3 | 921.213763 | -0.1849271 | 0.31010163 | -0.5963436 | 0.55094573 | 0.87093412 |
| Exosc7 | 3433.67241 | -0.3303622 | 0.26709443 | -1.236874 | 0.21613387 | 0.64982959 |
| Zdhhc3 | 3563.95982 | -0.4907779 | 0.25027555 | -1.9609502 | 0.04988483 | 0.34225988 |
| Slco3a1 | 20.9002186 | -0.7380476 | 1.24922938 | -0.5908023 | 0.55465291 | 0.87166868 |
| Pgm1 | 5278.08984 | -0.2542469 | 0.2157544 | -1.1784087 | 0.23863369 | 0.67452555 |
| Slc25a10 | 2488.22207 | 0.13893814 | 0.18306252 | 0.75896551 | 0.44787319 | 0.82338935 |
| Hgs | 2491.06449 | 0.08276319 | 0.18356923 | 0.45085544 | 0.65209374 | 0.91037383 |
| Rpl14 | 20892.4922 | 0.01762201 | 0.32613178 | 0.0540334 | 0.95690855 | 0.99217217 |
| Rassf3 | 126.897544 | 0.12365963 | 0.4390198 | 0.28167209 | 0.77819496 | 0.94837927 |
| Ccr1 | 6622.91024 | 0.10542927 | 0.44276139 | 0.23811758 | 0.8117899 | 0.95967622 |
| Itgb1 | 12302.8686 | 0.35063854 | 0.2072306 | 1.69202108 | 0.09064195 | 0.46044203 |
| Nrp1 | 5041.03913 | 0.58900288 | 0.32678121 | 1.80243803 | 0.07147652 | 0.41122734 |
| Pard3 | 35.7536116 | 1.51538828 | 1.45884502 | 1.03875892 | 0.29891687 | 0.73093388 |
| Dhtkd1 | 19.5439971 | 0.62294744 | 0.89339781 | 0.6972789 | 0.48562827 | 0.84183502 |
| Sec61a2 | 482.127459 | 0.12735503 | 0.3434276 | 0.37083515 | 0.71076032 | 0.93141199 |
| Nudt5 | 7170.10519 | 0.12998584 | 0.35089322 | 0.37044271 | 0.71105266 | 0.93149573 |
| Zfp282 | 356.146276 | -0.0913432 | 0.48421009 | -0.1886438 | 0.85037198 | 0.9679302 |
| Pdia4 | 14673.5257 | 0.07699314 | 0.15297615 | 0.50330158 | 0.61475225 | 0.89844745 |
| Iscu | 3025.2232 | -0.1337325 | 0.18826174 | -0.7103543 | 0.47748444 | 0.836949 |
| Fam20c | 316.612911 | -0.7812672 | 0.42329536 | -1.8456787 | 0.06493887 | 0.39268276 |
| Prkar1b | 83.747324 | -0.3662035 | 0.85924586 | -0.4261917 | 0.66996815 | 0.91689368 |
| Pdgfa | 12.050705 | -1.2722576 | 1.68368871 | -0.7556371 | 0.44986682 | 0.82484234 |
| Dnaaf5 | 973.012046 | -0.0525466 | 0.25563175 | -0.2055558 | 0.83713787 | 0.96683245 |
| Get4 | 4797.36054 | 0.27996245 | 0.24391878 | 1.14776912 | 0.25106389 | 0.6842534 |
| Xiap | 856.654085 | 0.55059365 | 0.24367941 | 2.2595001 | 0.02385229 | 0.2230773 |
| Stag2 | 1123.08638 | -0.0234068 | 0.19616002 | -0.1193249 | 0.90501792 | 0.9814201 |
| Higd2a | 2020.63296 | 0.06358094 | 0.26470791 | 0.24019284 | 0.81018077 | 0.95884376 |
| Nop16 | 6432.60019 | -0.168169 | 0.27318752 | -0.615581 | 0.53817109 | 0.86663962 |
| Arl10 | 310.677862 | -0.509035 | 0.30616801 | -1.6626002 | 0.09639249 | 0.47604735 |
| 4833439L19R | 3551.156 | -0.198092 | 0.18118479 | -1.0933145 | 0.27425574 | 0.7114017 |
| Thoc3 | 4891.85478 | -0.3030283 | 0.22710783 | -1.3342928 | 0.18210793 | 0.61344341 |
| Faf2 | 2122.53026 | 0.25733009 | 0.20805043 | 1.23686399 | 0.21613758 | 0.64982959 |
| Tspan17 | 24.4384977 | -0.0372555 | 1.08544566 | -0.0343228 | 0.97261978 | 0.99443087 |

| | | | | | | |
|---|---|---|---|---|---|---|
| Hk3 | 3664.28812 | 0.77320093 | 0.23241613 | 3.32679542 | 0.00087851 | 0.02107214 |
| Uimc1 | 911.260605 | -0.163895 | 0.19539462 | -0.8387897 | 0.40158732 | 0.79737342 |
| Smad7 | 34.0592411 | -0.8601744 | 0.83036951 | -1.0358936 | 0.30025179 | 0.73152364 |
| Casp1 | 2703.99442 | 0.31888436 | 0.33915241 | 0.94023912 | 0.34709492 | 0.76101142 |
| Kbtbd3 | 92.1076365 | 1.4646296 | 0.68165635 | 2.14863339 | 0.03166347 | 0.26539272 |
| Aasdhppt | 1417.88477 | 0.16568402 | 0.334864 | 0.49478002 | 0.62075541 | 0.89896588 |
| Cwf19l2 | 303.55302 | -0.2591846 | 0.3953159 | -0.6556392 | 0.5120563 | 0.85398564 |
| Alkbh8 | 694.881607 | -0.08148 | 0.24974385 | -0.3262542 | 0.74423204 | 0.93906039 |
| Sox17 | 11.8244965 | 0.7836992 | 2.63035479 | 0.29794429 | 0.76574568 | 0.94347227 |
| Lypla1 | 4395.73793 | -0.1148931 | 0.2161132 | -0.5316339 | 0.59497961 | 0.88830085 |
| Rb1cc1 | 919.814793 | 0.53272672 | 0.51254329 | 1.03937898 | 0.29862852 | 0.73093388 |
| Adhfe1 | 56.6989913 | 0.25959321 | 0.52548714 | 0.49400487 | 0.62130274 | 0.89912494 |
| Sgk3 | 2124.90936 | -0.0342499 | 0.21922618 | -0.1562307 | 0.8758512 | 0.97420329 |
| Cops5 | 7251.13471 | 0.26487312 | 0.32201821 | 0.82254081 | 0.41076917 | 0.80139357 |
| Stau2 | 419.383028 | -0.1770375 | 0.37314883 | -0.4744422 | 0.63518463 | 0.90531997 |
| Rdh10 | 301.040304 | 0.99423825 | 0.38228398 | 2.60078453 | 0.00930109 | 0.12353982 |
| Terf1 | 1438.30075 | 0.07160231 | 0.21062565 | 0.33995056 | 0.73389376 | 0.93750666 |
| Paqr8 | 23.0504789 | -0.6899177 | 1.90963688 | -0.3612821 | 0.71788854 | 0.93285856 |
| Tmem14a | 50.289404 | -0.705078 | 0.74281397 | -0.9491987 | 0.34251958 | 0.75920312 |
| Tram1 | 11557.7391 | 0.17156704 | 0.2393802 | 0.7167136 | 0.47355083 | 0.83483892 |
| Lactb2 | 1374.5643 | 0.58230672 | 0.27630685 | 2.10746394 | 0.03507738 | 0.28086853 |
| Ube2w | 1478.49163 | 0.37963467 | 0.28868107 | 1.31506601 | 0.18848772 | 0.61935434 |
| Tmem70 | 3606.20468 | 0.2783181 | 0.32333738 | 0.86076686 | 0.38936646 | 0.78942657 |
| Pikfyve | 321.692664 | -0.2586866 | 0.53245192 | -0.4858403 | 0.62708036 | 0.90205481 |
| Idh1 | 8505.28177 | 0.57540513 | 0.1546061 | 3.72174931 | 0.00019785 | 0.00645763 |
| Mettl21a | 747.970274 | 0.16819946 | 0.27406845 | 0.61371333 | 0.53940478 | 0.86719392 |
| Creb1 | 1284.23133 | 0.05725395 | 0.23515918 | 0.24346891 | 0.80764216 | 0.95800316 |
| Klf7 | 747.020639 | 0.92907362 | 0.35805207 | 2.59480034 | 0.00946459 | 0.12489785 |
| Fastkd2 | 3608.17813 | 0.45716934 | 0.34827117 | 1.31268212 | 0.18929008 | 0.61985441 |
| Eef1b2 | 15849.2217 | 0.07714398 | 0.34328393 | 0.22472354 | 0.82219435 | 0.96238671 |
| Ndufs1 | 7643.33165 | -0.1595122 | 0.17127298 | -0.9313333 | 0.35168119 | 0.76348958 |
| Nrp2 | 353.629399 | 2.19618414 | 0.60194724 | 3.64846615 | 0.00026381 | 0.00806799 |
| Maip1 | 1560.50562 | 0.42447439 | 0.35104538 | 1.20917243 | 0.22659661 | 0.66015092 |
| Rftn2 | 37.5340388 | 0.29992645 | 0.88173119 | 0.34015634 | 0.7337388 | 0.93750666 |
| Mob4 | 3008.69672 | 0.14430128 | 0.26819726 | 0.5380416 | 0.59054833 | 0.88748446 |
| Hspd1 | 35821.5334 | -0.4940929 | 0.23574384 | -2.095889 | 0.03609204 | 0.28486166 |
| Coq10b | 1265.84711 | -0.1556386 | 0.24285254 | -0.6408771 | 0.52160254 | 0.8580909 |
| Sf3b1 | 16092.6955 | -0.2887024 | 0.17432214 | -1.656143 | 0.09769286 | 0.47892724 |
| Slc39a10 | 998.2259 | -0.0427076 | 0.3516512 | -0.1214486 | 0.9033357 | 0.98043702 |
| Slc40a1 | 9623.49179 | 2.61656624 | 0.29313322 | 8.92620168 | 4.408863600 | 3.172566378 |
| Wdr75 | 3194.86366 | -0.310641 | 0.24577554 | -1.2639214 | 0.20625824 | 0.64088316 |
| Ikzf2 | 54.4082262 | -0.2282319 | 0.56262889 | -0.4056526 | 0.6849979 | 0.92314423 |
| Lancl1 | 859.268579 | 0.5688334 | 0.36476546 | 1.55944977 | 0.11888997 | 0.52515149 |
| Acadl | 3476.55317 | 0.38761254 | 0.1805663 | 2.14664948 | 0.0318212 | 0.26554509 |
| Kansl1l | 83.8133892 | -0.1587264 | 0.52110195 | -0.3045976 | 0.76067265 | 0.94230972 |
| Rpe | 3830.51706 | -0.2547232 | 0.17456609 | -1.4591792 | 0.14451578 | 0.57027792 |

| | | | | | | |
|---|---|---|---|---|---|---|
| Icos | 431.682772 | 1.06317933 | 0.53542209 | 1.98568448 | 0.04706837 | 0.3347601 |
| Cd28 | 443.575746 | -0.085189 | 0.23838863 | -0.3573535 | 0.72082717 | 0.93398714 |
| Raph1 | 1939.95205 | 0.11393508 | 0.20649219 | 0.5517646 | 0.58110964 | 0.88314588 |
| Carf | 199.577033 | -0.4655091 | 0.48521202 | -0.9593931 | 0.33736074 | 0.75598717 |
| Ica1l | 43.4061936 | 0.12780622 | 0.66719543 | 0.19155739 | 0.84808892 | 0.96765585 |
| Wdr12 | 6064.68011 | -0.2787841 | 0.23877411 | -1.1675641 | 0.24298263 | 0.67814218 |
| Nop58 | 6540.60021 | -0.4262465 | 0.17461607 | -2.4410497 | 0.01464464 | 0.16649431 |
| Sumo1 | 10136.3598 | 0.29348798 | 0.3324385 | 0.8828339 | 0.37732602 | 0.77996438 |
| Als2 | 316.161916 | 0.20711759 | 0.46205696 | 0.4482512 | 0.65397191 | 0.91076467 |
| Stradb | 1050.54329 | 0.41512347 | 0.21677217 | 1.91502195 | 0.0554897 | 0.3627292 |
| Trak2 | 217.466582 | -0.1779091 | 0.68596928 | -0.2593544 | 0.79536182 | 0.9537323 |
| Casp8 | 6314.15667 | 0.29518909 | 0.14480671 | 2.03850419 | 0.04149954 | 0.31027743 |
| Cflar | 1395.86668 | 0.80396838 | 0.29969584 | 2.68261438 | 0.00730492 | 0.10439375 |
| Ndufb3 | 2491.48747 | -0.003598 | 0.30981395 | -0.0116133 | 0.99073415 | 0.9966603 |
| Clk1 | 4370.50631 | 0.29041818 | 0.32660047 | 0.88921543 | 0.37388731 | 0.77748705 |
| Ppil3 | 1350.52523 | 0.03630402 | 0.36603835 | 0.09918091 | 0.92099463 | 0.9839023 |
| Nif3l1 | 2113.633 | 0.31886208 | 0.29870375 | 1.06748602 | 0.28575242 | 0.721488 |
| Orc2 | 3268.35625 | -0.2837306 | 0.22086061 | -1.284659 | 0.19891146 | 0.63148061 |
| Sgo2a | 1249.41361 | -0.3148739 | 0.29952321 | -1.0512505 | 0.29314357 | 0.72635716 |
| Col5a2 | 27.2340226 | 4.13216805 | 1.55183093 | 2.66276948 | 0.00775005 | 0.10847408 |
| Col3a1 | 68.1794264 | 2.41911753 | 0.84770823 | 2.85371479 | 0.00432113 | 0.07162655 |
| Kdelc1 | 1489.59312 | -0.1414432 | 0.2569476 | -0.5504749 | 0.58199367 | 0.88321857 |
| Ercc5 | 417.405158 | 0.09616269 | 0.53269145 | 0.18052232 | 0.85674253 | 0.96934253 |
| Tex30 | 3655.79531 | 0.02816628 | 0.35964621 | 0.07831663 | 0.93757619 | 0.98856831 |
| Ptp4a1 | 52.6896157 | 0.20182367 | 1.11576678 | 0.18088339 | 0.85645911 | 0.96924897 |
| Il18rap | 696.100042 | -0.9292334 | 0.25328086 | -3.6687864 | 0.0002437 | 0.00756659 |
| Il1rl1 | 87.4942152 | 0.16236746 | 0.56732633 | 0.28619764 | 0.77472675 | 0.94709254 |
| Il1r2 | 91.8422553 | -2.127308 | 0.59359845 | -3.5837492 | 0.0003387 | 0.00981304 |
| Map4k4 | 374.491179 | -0.3823655 | 0.35411843 | -1.0797674 | 0.28024579 | 0.71532524 |
| Pdcl3 | 2634.63675 | 0.18257334 | 0.19014144 | 0.96019753 | 0.33695581 | 0.7553565 |
| Chst10 | 68.0049467 | -0.5587364 | 1.41153718 | -0.3958354 | 0.69222644 | 0.92539651 |
| Rev1 | 323.007899 | -0.10713 | 0.34883683 | -0.3071064 | 0.75876241 | 0.94184564 |
| Eif5b | 2138.65766 | -0.2771081 | 0.33464221 | -0.8280728 | 0.40762926 | 0.8005951 |
| Mrpl30 | 3355.9455 | 0.12575377 | 0.30500791 | 0.41229674 | 0.68012194 | 0.92061965 |
| Mitd1 | 653.135694 | 0.31421347 | 0.32222002 | 0.97515191 | 0.32948491 | 0.74874249 |
| Stk17b | 7005.02751 | 0.0208335 | 0.2235766 | 0.09318283 | 0.92575831 | 0.98587982 |
| Asnsd1 | 4296.57698 | -0.1735763 | 0.15293117 | -1.1349962 | 0.25637691 | 0.69126266 |
| Osgepl1 | 617.846258 | 0.04056657 | 0.34882285 | 0.11629561 | 0.90741826 | 0.98176897 |
| Ormdl1 | 3793.67564 | 0.66500649 | 0.39538565 | 1.68191863 | 0.09258462 | 0.46597694 |
| Pms1 | 689.748331 | 0.08782977 | 0.26183066 | 0.33544494 | 0.73728947 | 0.93800224 |
| Inpp1 | 437.544714 | -0.0531646 | 0.26938134 | -0.1973581 | 0.8435473 | 0.96765585 |
| Gls | 523.675863 | -0.6381312 | 0.34110877 | -1.8707559 | 0.06137893 | 0.38244534 |
| Stat1 | 2619.07842 | -0.4398949 | 0.18469334 | -2.3817582 | 0.01723021 | 0.18456841 |
| Nabp1 | 1426.17383 | 0.74619449 | 0.21535466 | 3.46495632 | 0.00053032 | 0.01408534 |
| Mgat4a | 2962.59893 | 0.51648396 | 0.2407666 | 2.14516449 | 0.0319397 | 0.26603092 |
| Unc50 | 3288.53666 | 0.62089579 | 0.3267666 | 1.90012012 | 0.05741736 | 0.36870684 |

| | | | | | | |
|---|---|---|---|---|---|---|
| Coa5 | 7888.87551 | 0.11965159 | 0.25429406 | 0.47052452 | 0.63798032 | 0.90599527 |
| Inpp4a | 985.893786 | 0.40500362 | 0.34824932 | 1.16297032 | 0.24484154 | 0.68039867 |
| Tmem131 | 377.428461 | 0.02407252 | 0.52036589 | 0.04626075 | 0.96310242 | 0.99401755 |
| Sema4c | 157.058741 | 0.23712718 | 0.42240448 | 0.56137467 | 0.57454215 | 0.88059206 |
| Plekhb2 | 4065.86962 | 0.626469 | 0.20451214 | 3.06323617 | 0.00218957 | 0.04271935 |
| Ptpn18 | 3789.42954 | 0.964245 | 0.4277248 | 2.2543584 | 0.02417363 | 0.22455944 |
| Imp4 | 8037.90489 | 0.04734357 | 0.27384459 | 0.17288481 | 0.86274198 | 0.9705375 |
| Dst | 76.8368958 | 3.99940327 | 1.31729025 | 3.03608356 | 0.00239673 | 0.0455975 |
| Prim2 | 2338.05145 | 0.1250116 | 0.27137897 | 0.46065324 | 0.64504741 | 0.90835328 |
| Zfp142 | 234.669128 | 0.21276678 | 0.65555147 | 0.32456152 | 0.74551296 | 0.93922348 |
| Rhbdd1 | 652.323978 | -0.0446193 | 0.24929725 | -0.1789803 | 0.85795314 | 0.96944425 |
| Col9a1 | 10.5232463 | 2.12228562 | 2.04819203 | 1.03617512 | 0.30012045 | 0.73145694 |
| Mff | 4956.69623 | 0.57866699 | 0.31803149 | 1.81952735 | 0.06883101 | 0.40461788 |
| Fam135a | 57.4399874 | -0.668473 | 1.239076 | -0.5394931 | 0.58954665 | 0.88722166 |
| Sdhaf4 | 715.000748 | 0.1620232 | 0.37095912 | 0.43676835 | 0.66227935 | 0.91395288 |
| Smap1 | 458.64207 | -0.0686834 | 0.30209583 | -0.2273565 | 0.82014655 | 0.96176948 |
| Ogfrl1 | 456.293645 | -0.8152799 | 0.26886689 | -3.0322807 | 0.00242713 | 0.0460372 |
| Agfg1 | 261.902799 | -0.1976138 | 0.29613045 | -0.6673201 | 0.50456772 | 0.85034006 |
| Nhej1 | 322.969162 | 0.55338332 | 0.37504527 | 1.47551071 | 0.14007529 | 0.56396008 |
| Cyp27a1 | 358.052005 | 0.55257668 | 0.48910214 | 1.12977768 | 0.25856992 | 0.69487825 |
| Rnf25 | 559.865173 | 0.33668385 | 0.36768451 | 0.91568679 | 0.3598312 | 0.76857232 |
| Bcs1l | 990.070097 | 0.01300651 | 0.23958669 | 0.05428728 | 0.95670628 | 0.99215259 |
| Cnot9 | 9121.60906 | -0.0319947 | 0.18935885 | -0.1689632 | 0.86582557 | 0.9705375 |
| Ctdsp1 | 444.025883 | -0.2246058 | 0.2496735 | -0.899598 | 0.36833421 | 0.77433105 |
| Slc11a1 | 8609.54492 | 0.18155978 | 0.1421489 | 1.27725065 | 0.20151377 | 0.63561051 |
| Pnkd | 846.286795 | 0.45241959 | 0.35949551 | 1.25848466 | 0.20821653 | 0.6418276 |
| Cxcr2 | 501.438931 | -1.6840137 | 0.35553173 | -4.736606 | 2.173270983 | 0.00014066 |
| Ppm1f | 1433.29682 | -0.233481 | 0.33532766 | -0.6962771 | 0.48625534 | 0.84183502 |
| Igfbp5 | 19.6126795 | -1.0745968 | 1.62960108 | -0.6594232 | 0.50962404 | 0.85335831 |
| Xrcc5 | 212.540507 | 0.2095677 | 0.39829336 | 0.52616418 | 0.59877414 | 0.88979641 |
| Pecr | 432.425192 | 0.37764923 | 0.32574843 | 1.15932786 | 0.24632257 | 0.6814256 |
| Atic | 2131.43191 | -0.7332389 | 0.20496828 | -3.5773289 | 0.00034712 | 0.00995083 |
| Fn1 | 501.837382 | -0.9399459 | 0.35754728 | -2.6288717 | 0.00856687 | 0.11670212 |
| Bard1 | 269.394883 | -0.5868319 | 0.44514435 | -1.3182956 | 0.1874047 | 0.61935434 |
| Zfand2b | 756.713357 | 0.67038557 | 0.37229033 | 1.80070637 | 0.07174917 | 0.4119576 |
| Abcb6 | 686.444462 | 0.57005177 | 0.24202972 | 2.3552966 | 0.01850793 | 0.19153023 |
| Ankzf1 | 630.961204 | -0.0734272 | 0.20933468 | -0.3507648 | 0.72576482 | 0.93544211 |
| Glb1l | 68.0656936 | 0.34049967 | 0.64475778 | 0.52810478 | 0.59742661 | 0.88908999 |
| Stk16 | 1663.12951 | -0.2046299 | 0.34021616 | -0.6014703 | 0.5475268 | 0.87008334 |
| Tuba4a | 1815.38897 | -2.9482908 | 0.24452187 | -12.057371 | 1.773547090 | 4.339160311 |
| Dnajb2 | 92.53424 | 0.05253338 | 0.60247328 | 0.0871962 | 0.93051556 | 0.98716777 |
| Speg | 1342.67298 | 2.14449839 | 0.30853957 | 6.95048079 | 3.640433652 | 9.475196781 |
| Dnpep | 2327.32398 | 0.58938107 | 0.2632112 | 2.2391945 | 0.02514326 | 0.23084234 |
| Obsl1 | 68.3958029 | 0.95154015 | 0.64890938 | 1.46636831 | 0.14254798 | 0.56828728 |
| Stk11ip | 973.930553 | -0.1133166 | 0.27008192 | -0.4195638 | 0.67480415 | 0.91843337 |
| Trip12 | 1343.26531 | -0.1513407 | 0.3039913 | -0.4978455 | 0.61859292 | 0.89878056 |

| | | | | | | |
|---|---|---|---|---|---|---|
| Sp100 | 377.878129 | -1.4911559 | 0.31890502 | -4.6758623 | 2.927210504 | 0.00018458 |
| Itm2c | 4871.11271 | 0.3326197 | 0.1548758 | 2.14765443 | 0.03174122 | 0.26554509 |
| Psmd1 | 16596.8318 | -0.0298014 | 0.15175799 | -0.1963743 | 0.84431724 | 0.96765585 |
| Ncl | 29614.0826 | -0.5183688 | 0.21664851 | -2.3926717 | 0.0167262 | 0.1809121 |
| Epha4 | 12.742713 | -0.122039 | 2.07714231 | -0.0587533 | 0.9531486 | 0.99149988 |
| Ptma | 37462.8102 | -0.3983218 | 0.14990184 | -2.6572172 | 0.00787887 | 0.10963695 |
| Pde6d | 1112.40396 | 0.10211064 | 0.26756193 | 0.38163368 | 0.7027331 | 0.92775027 |
| Cops7b | 1272.90394 | 0.02796203 | 0.22442523 | 0.12459398 | 0.900845 | 0.98043702 |
| Farsb | 9203.34811 | -0.6095963 | 0.18431033 | -3.3074448 | 0.00094151 | 0.02232078 |
| Mrpl44 | 2161.87527 | -0.2970759 | 0.2004475 | -1.4820632 | 0.13832348 | 0.56085884 |
| Serpine2 | 527.476822 | -0.0828733 | 0.36456312 | -0.2273222 | 0.82017323 | 0.96176948 |
| Eif4e2 | 5392.57638 | 0.16131351 | 0.2681261 | 0.60163299 | 0.54741846 | 0.87008334 |
| Ndufa10 | 7002.74503 | -0.1390343 | 0.15481717 | -0.8980547 | 0.36915639 | 0.77480275 |
| Rnpepl1 | 387.411591 | 0.64777278 | 0.50478132 | 1.28327408 | 0.19939607 | 0.63224783 |
| Capn10 | 407.235288 | 0.34057589 | 0.33971341 | 1.00253885 | 0.31608341 | 0.74186126 |
| Gpr35 | 61.4454304 | -0.4663871 | 0.90074029 | -0.517782 | 0.60461038 | 0.89258167 |
| Mterf4 | 575.38199 | 0.08755917 | 0.29392387 | 0.29789745 | 0.76578143 | 0.94347227 |
| Pask | 405.908031 | 0.20704355 | 0.30290513 | 0.68352608 | 0.49427448 | 0.84678075 |
| Ppp1r7 | 3342.24198 | 0.06665326 | 0.16716196 | 0.39873465 | 0.69008873 | 0.92470267 |
| Sept2 | 1302.86497 | 0.12819442 | 0.21425357 | 0.59833035 | 0.54961953 | 0.87093412 |
| Stk25 | 2152.84217 | 0.45216055 | 0.25920571 | 1.74440816 | 0.08108794 | 0.43949881 |
| Bok | 31.8549765 | 3.62832957 | 1.46786668 | 2.47183863 | 0.01344202 | 0.1567552 |
| Thap4 | 866.499043 | -0.0985277 | 0.22391979 | -0.4400135 | 0.65992737 | 0.91280998 |
| Atg4b | 2612.05865 | -0.0974276 | 0.18653924 | -0.5222903 | 0.60146823 | 0.89094157 |
| Dtymk | 4891.43862 | 0.31706154 | 0.36637199 | 0.86540881 | 0.38681445 | 0.78911204 |
| Ing5 | 1734.65719 | 0.0697833 | 0.24675891 | 0.28279953 | 0.77733051 | 0.94834788 |
| Pdcd1 | 20.3861635 | -2.2827982 | 1.90244334 | -1.1999296 | 0.23016667 | 0.66484742 |
| Inpp5d | 6569.51512 | -0.1214238 | 0.31438729 | -0.3862235 | 0.69933116 | 0.92669304 |
| Atg16l1 | 949.669836 | 0.39018709 | 0.25278379 | 1.54356059 | 0.12269482 | 0.53280998 |
| Lrrfip1 | 5491.16378 | -0.5153967 | 0.23616151 | -2.1823908 | 0.02908071 | 0.25392168 |
| Scly | 117.083447 | 0.64268274 | 0.49075523 | 1.309579 | 0.1903383 | 0.6209089 |
| Ilkap | 1997.83936 | -0.2288202 | 0.17818288 | -1.2841876 | 0.19907632 | 0.63169391 |
| Asb1 | 132.66099 | 0.52282279 | 0.65362497 | 0.79988192 | 0.42377921 | 0.80917688 |
| Hdac4 | 666.096226 | 0.20668127 | 0.40579063 | 0.50932982 | 0.61052106 | 0.89508835 |
| Serpinb8 | 1443.0458 | 0.6076514 | 0.19730079 | 3.07982237 | 0.00207124 | 0.041123 |
| Cln8 | 4572.4249 | 0.49458987 | 0.16717555 | 2.95850604 | 0.00309134 | 0.0552064 |
| Relch | 493.384134 | -0.1350829 | 0.27285273 | -0.4950762 | 0.62054631 | 0.89896588 |
| Tnfrsf11a | 1635.45277 | 0.18379752 | 0.22985158 | 0.79963567 | 0.42392191 | 0.80917688 |
| Gin1 | 491.317284 | 0.19915495 | 0.41974685 | 0.47446443 | 0.63516876 | 0.90531997 |
| Pam | 232.769639 | -0.170185 | 0.41703302 | -0.4080853 | 0.68321106 | 0.92177356 |
| Ccdc93 | 864.008243 | 0.81493556 | 0.3169518 | 2.57116559 | 0.01013569 | 0.13017076 |
| Actr3 | 52027.4076 | -0.2119494 | 0.18423454 | -1.1504325 | 0.24996579 | 0.6842534 |
| Slc35f5 | 1861.09573 | 0.29959842 | 0.20850236 | 1.43690661 | 0.15074453 | 0.57859667 |
| Ccnt2 | 826.475589 | 0.51062534 | 0.36036764 | 1.41695671 | 0.15649558 | 0.58588334 |
| Ubxn4 | 3186.90243 | -0.0395974 | 0.23403404 | -0.1691949 | 0.86564334 | 0.9705375 |
| Mcm6 | 26984.3202 | -0.1675976 | 0.21528196 | -0.7785028 | 0.43627267 | 0.81729305 |

| | | | | | | |
|---|---|---|---|---|---|---|
| Dars | 10828.2353 | -0.0037253 | 0.26496502 | -0.0140595 | 0.9887825 | 0.99650865 |
| Rgs18 | 1438.08565 | -1.0671056 | 0.30756463 | -3.4695328 | 0.00052136 | 0.01392544 |
| Rgs1 | 2704.10053 | -0.2624054 | 0.2783605 | -0.942682 | 0.34584358 | 0.7602916 |
| Rgs2 | 13025.4735 | -0.2694154 | 0.19842861 | -1.357745 | 0.17454462 | 0.60656845 |
| Cdc73 | 276.408265 | -0.2450493 | 0.51120346 | -0.4793577 | 0.63168417 | 0.90399255 |
| Cfh | 7822.6718 | 0.36916604 | 0.21382674 | 1.72647278 | 0.08426239 | 0.44526833 |
| Tsn | 5207.75376 | 0.06516149 | 0.1841589 | 0.35383296 | 0.72346407 | 0.93444578 |
| Nifk | 6438.02701 | -0.3718982 | 0.14819779 | -2.5094718 | 0.01209119 | 0.1460133 |
| Epb41l5 | 475.247288 | 0.17785899 | 0.29067316 | 0.61188651 | 0.54061285 | 0.86741237 |
| Ptpn4 | 176.976487 | -1.3728707 | 0.80814193 | -1.698799 | 0.08935706 | 0.4566019 |
| Dbi | 6488.73574 | 0.60374446 | 0.34686287 | 1.74058542 | 0.08175628 | 0.44058349 |
| 3110009E18I | 74.5166556 | 0.16014739 | 0.60411007 | 0.26509637 | 0.79093522 | 0.95268772 |
| Steap3 | 490.315463 | -0.1511021 | 0.39650852 | -0.3810816 | 0.7031427 | 0.92804961 |
| Nek7 | 3094.58801 | 0.27391259 | 0.31529532 | 0.86874929 | 0.38498427 | 0.78754751 |
| Ptprc | 8729.47375 | 0.20949212 | 0.18892405 | 1.10886952 | 0.26748648 | 0.70441538 |
| Cd55 | 22.1072691 | -0.1387011 | 2.09167662 | -0.066311 | 0.94713025 | 0.99019648 |
| Ddx59 | 462.686942 | 0.30599524 | 0.32444016 | 0.94314846 | 0.34560497 | 0.7602916 |
| Pfkfb2 | 48.710901 | -0.0421469 | 1.4692744 | -0.0286855 | 0.97711542 | 0.99483019 |
| Tmem9 | 1613.48308 | 0.8848097 | 0.29677168 | 2.98144925 | 0.00286888 | 0.05245883 |
| Tnni1 | 42.8963384 | 0.07098032 | 1.03442716 | 0.06861799 | 0.9452937 | 0.99019648 |
| Csrp1 | 1406.25058 | 0.52287529 | 0.29418948 | 1.77734192 | 0.075512 | 0.42315085 |
| Srgap2 | 1098.62198 | 0.17181415 | 0.29749011 | 0.57754577 | 0.56357083 | 0.87626321 |
| Arl8a | 157.193062 | 0.00962508 | 0.368941 | 0.02608841 | 0.97918682 | 0.99483019 |
| Eif2d | 1635.74885 | -0.3887618 | 0.20857309 | -1.8639115 | 0.06233414 | 0.38520862 |
| Ube2t | 872.614168 | 0.3183439 | 0.34666106 | 0.91831457 | 0.3584542 | 0.76713922 |
| Rassf5 | 3302.64468 | -0.2286218 | 0.19572418 | -1.1680813 | 0.24277398 | 0.67814218 |
| Rab29 | 1229.94082 | 0.38750502 | 0.37228188 | 1.04089144 | 0.29792594 | 0.73013751 |
| Nucks1 | 4732.57119 | -0.0929752 | 0.27998642 | -0.3320704 | 0.73983612 | 0.93800224 |
| Slc45a3 | 10.6529824 | -0.5853667 | 2.15336222 | -0.2718385 | 0.78574623 | 0.95168899 |
| Elk4 | 237.466203 | -0.6785771 | 0.55435553 | -1.2240828 | 0.220921 | 0.65452328 |
| Cdk18 | 39.2334715 | 0.3689127 | 0.97122334 | 0.37984332 | 0.70406172 | 0.92850227 |
| Rbbp5 | 1840.78027 | -0.024652 | 0.32438802 | -0.0759955 | 0.93942269 | 0.98913801 |
| Pik3c2b | 63.7960556 | -0.5083887 | 0.7445667 | -0.682798 | 0.49473449 | 0.84678075 |
| Klhl12 | 383.495213 | -0.1252628 | 0.35129582 | -0.3565737 | 0.721411 | 0.93398714 |
| Cyb5r1 | 859.778319 | -0.2845456 | 0.23740559 | -1.198563 | 0.23069791 | 0.6653874 |
| Adipor1 | 5630.27144 | 0.09556525 | 0.16008525 | 0.59696476 | 0.55053091 | 0.87093412 |
| Ppfia4 | 2622.4015 | -0.5024705 | 0.20354356 | -2.4686141 | 0.01356374 | 0.1574243 |
| Myog | 29.7318469 | -1.1616612 | 1.52154054 | -0.763477 | 0.44517898 | 0.82118815 |
| Atp2b4 | 17.810425 | -0.6921259 | 1.56634124 | -0.4418743 | 0.65858019 | 0.91265586 |
| Tor1aip1 | 2336.17905 | -0.4071548 | 0.21513778 | -1.8925307 | 0.05842031 | 0.37144262 |
| Xpr1 | 257.080742 | 0.13801343 | 0.28634504 | 0.48198296 | 0.62981805 | 0.90321876 |
| Stx6 | 1980.19565 | 0.30866163 | 0.24373767 | 1.26636816 | 0.2053813 | 0.63913241 |
| Mr1 | 53.1517125 | 0.85197383 | 0.89675155 | 0.95006674 | 0.34207834 | 0.75878732 |
| Glul | 4402.5322 | 0.5235127 | 0.23780673 | 2.20142087 | 0.02770624 | 0.24685395 |
| Rgs16 | 12.7897691 | 1.9541696 | 1.97223277 | 0.99084126 | 0.3217631 | 0.74447686 |
| Lamc1 | 310.935583 | -0.6446676 | 0.4366152 | -1.4765121 | 0.13980646 | 0.56350984 |

| | | | | | | |
|---|---|---|---|---|---|---|
| Ncf2 | 9908.05005 | 0.06906187 | 0.17202591 | 0.40146201 | 0.68808 | 0.92470267 |
| Rgl1 | 3320.01301 | 0.53409076 | 0.37164739 | 1.43709001 | 0.15069242 | 0.57859667 |
| Fam129a | 651.046055 | -0.3968069 | 0.37293898 | -1.0639994 | 0.28732897 | 0.72278333 |
| Rnf2 | 4744.08893 | 0.21766138 | 0.15641417 | 1.3915707 | 0.16405244 | 0.59585062 |
| Coq8a | 128.647272 | 0.07803482 | 0.53837904 | 0.14494401 | 0.88475508 | 0.97742337 |
| Cdc42bpa | 265.922413 | 0.56451208 | 0.60076135 | 0.9396611 | 0.34739142 | 0.76101142 |
| Ahctf1 | 1769.55471 | 0.04282778 | 0.47335716 | 0.09047667 | 0.92790844 | 0.98666423 |
| Tfb2m | 1008.34957 | -0.1276062 | 0.25005555 | -0.5103113 | 0.60983342 | 0.89508835 |
| Efcab2 | 228.059793 | -0.04902 | 0.38351274 | -0.1278186 | 0.89829256 | 0.97989497 |
| Parp1 | 10818.7043 | -0.0796433 | 0.20290317 | -0.3925186 | 0.69467506 | 0.92609254 |
| Acbd3 | 460.677682 | -0.1690401 | 0.36150199 | -0.467605 | 0.64006708 | 0.90671662 |
| Cox20 | 572.807436 | 0.1189216 | 0.26520393 | 0.44841567 | 0.65385324 | 0.91076467 |
| Desi2 | 550.268936 | 0.45090973 | 0.41908913 | 1.075928 | 0.28195946 | 0.71768831 |
| Sdccag8 | 738.590114 | -0.4239484 | 0.2105968 | -2.013081 | 0.04410611 | 0.32226735 |
| Capn2 | 1862.58455 | 0.8358824 | 0.31051361 | 2.69193478 | 0.00710388 | 0.10284238 |
| Trp53bp2 | 424.362294 | 0.09927487 | 0.42028781 | 0.23620687 | 0.81327215 | 0.95983628 |
| Srp9 | 3742.1205 | 0.34551348 | 0.33671674 | 1.02612505 | 0.30483264 | 0.7350715 |
| Nvl | 1791.10364 | -0.5317587 | 0.38979159 | -1.3642129 | 0.17250056 | 0.60492813 |
| Tmem63a | 1825.43377 | 0.04714779 | 0.36571395 | 0.12891987 | 0.89742105 | 0.97984623 |
| Pycr2 | 3685.79224 | -0.3184727 | 0.2079012 | -1.5318466 | 0.1255603 | 0.53724874 |
| Opn3 | 608.954834 | -0.0332157 | 0.28432493 | -0.1168229 | 0.90700037 | 0.98176897 |
| Fh1 | 9009.53196 | 0.18865227 | 0.28365066 | 0.66508665 | 0.50599507 | 0.85074333 |
| Ifi211 | 882.325304 | -2.3309155 | 0.31345583 | -7.4361849 | 1.036347671 | 3.426389477 |
| Dusp23 | 133.266115 | 0.16465069 | 0.39812849 | 0.41356169 | 0.67919514 | 0.92031392 |
| Cfap45 | 56.2820022 | 1.79208622 | 1.04127917 | 1.721043 | 0.08524302 | 0.44656163 |
| Tagln2 | 16640.8611 | -0.451257 | 0.17448167 | -2.5862715 | 0.00970205 | 0.12687515 |
| Slamf9 | 420.305075 | 1.30351403 | 0.42541487 | 3.06410077 | 0.00218325 | 0.04271935 |
| Copa | 12288.9505 | 0.09029833 | 0.20944461 | 0.43113228 | 0.66637219 | 0.91520501 |
| Dcaf8 | 3423.56798 | 0.06009582 | 0.17110186 | 0.35122834 | 0.72541705 | 0.93544211 |
| Vangl2 | 193.171293 | -0.2495755 | 0.41237875 | -0.6052094 | 0.5450399 | 0.86963259 |
| Uck2 | 3365.56172 | -0.5964972 | 0.24086239 | -2.4765062 | 0.01326753 | 0.15566077 |
| Tada1 | 3231.01856 | 0.12453748 | 0.21550409 | 0.57788915 | 0.56333896 | 0.8760902 |
| Dusp27 | 160.773696 | 0.31299816 | 0.37567473 | 0.83316266 | 0.404753 | 0.79917613 |
| Pou2f1 | 343.740328 | 0.06166939 | 0.49905164 | 0.12357317 | 0.90165324 | 0.98043702 |
| Mpzl1 | 993.318116 | 0.72981379 | 0.34997388 | 2.08533789 | 0.03703864 | 0.28924449 |
| Mpc2 | 2014.84497 | 0.15151561 | 0.41595086 | 0.36426323 | 0.71566145 | 0.93235217 |
| Dcaf6 | 54.3928407 | -0.079133 | 0.78512062 | -0.1007909 | 0.91971643 | 0.9839023 |
| Nme7 | 894.437327 | 0.30921707 | 0.29487223 | 1.04864766 | 0.29434031 | 0.72726016 |
| Atp1b1 | 31.3598834 | 0.89538729 | 0.96642151 | 0.92649769 | 0.3541874 | 0.76419569 |
| Blzf1 | 882.924767 | 0.22394517 | 0.22601159 | 0.99085705 | 0.32175539 | 0.74447686 |
| Ccdc181 | 159.303303 | -0.3949173 | 0.38012095 | -1.0389254 | 0.29883942 | 0.73093388 |
| F5 | 24.0068954 | -4.6409086 | 1.30063793 | -3.5681787 | 0.00035947 | 0.01025038 |
| Sell | 247.634558 | -5.7889201 | 0.91380957 | -6.3349304 | 2.374483856 | 4.761813280 |
| Scyl3 | 1004.35361 | -0.1870816 | 0.24449897 | -0.7651632 | 0.44417436 | 0.82068545 |
| Kifap3 | 625.412379 | 0.14859957 | 0.39172622 | 0.37934548 | 0.70443133 | 0.9285501 |
| Prrx1 | 10.7856752 | -0.1212112 | 2.15200634 | -0.0563247 | 0.95508313 | 0.99215259 |

| | | | | | | |
|---|---|---|---|---|---|---|
| Ralgps2 | 1561.87337 | 0.4611511 | 0.25305174 | 1.82235893 | 0.06840054 | 0.40325004 |
| Abl2 | 448.424287 | -0.205609 | 0.56012003 | -0.3670803 | 0.71355913 | 0.93235217 |
| Soat1 | 4981.36588 | -0.0225377 | 0.20458414 | -0.1101635 | 0.91227973 | 0.98292561 |
| Smyd2 | 3103.10459 | -0.1806922 | 0.26182395 | -0.6901286 | 0.49011332 | 0.84448327 |
| Cenpf | 1058.9504 | 0.07765406 | 0.28334311 | 0.2740637 | 0.78403567 | 0.95094799 |
| Kctd3 | 741.378851 | 0.02391303 | 0.53952712 | 0.04432219 | 0.96464758 | 0.99406401 |
| Eprs | 10726.2222 | -0.4867226 | 0.28657405 | -1.6984184 | 0.08942882 | 0.45677777 |
| Bpnt1 | 2074.0923 | -0.1300712 | 0.20867018 | -0.6233342 | 0.53306495 | 0.86485193 |
| Iars2 | 2926.17074 | 0.04440975 | 0.16468149 | 0.26967058 | 0.78741369 | 0.95225386 |
| Nek2 | 2965.9139 | -0.2132587 | 0.22787209 | -0.9358703 | 0.34933996 | 0.76189618 |
| Lpgat1 | 568.149579 | 0.15763058 | 0.40869848 | 0.38568917 | 0.69972688 | 0.92669304 |
| Ppp2r5a | 902.806531 | -0.5104723 | 0.28326457 | -1.8021042 | 0.07152901 | 0.41122734 |
| Tmem206 | 3770.13522 | 0.69365947 | 0.18180859 | 3.81532828 | 0.000136 | 0.00469975 |
| Atf3 | 18976.9014 | 0.68795829 | 0.28317082 | 2.4294816 | 0.01512043 | 0.17000758 |
| Batf3 | 263.386899 | 0.07404374 | 0.46230911 | 0.16016067 | 0.87275451 | 0.97344599 |
| Tatdn3 | 240.206469 | -0.2969724 | 0.41032433 | -0.7237504 | 0.46921899 | 0.83381643 |
| Angel2 | 1428.69625 | 0.04286855 | 0.26235392 | 0.16339969 | 0.87020375 | 0.97251987 |
| Traf5 | 134.301381 | 0.7108374 | 0.50585471 | 1.40522048 | 0.15995573 | 0.58927692 |
| Plxna2 | 13.8175257 | 0.76105666 | 2.09263247 | 0.36368386 | 0.71609409 | 0.93235217 |
| Usf1 | 2538.35861 | 0.1257454 | 0.28060905 | 0.44811599 | 0.65406949 | 0.91078339 |
| Nmt2 | 3911.54124 | -0.0489571 | 0.22602922 | -0.2165965 | 0.82852279 | 0.96462215 |
| Suv39h2 | 314.621142 | -0.0303722 | 0.5650911 | -0.0537475 | 0.95713632 | 0.99217427 |
| Dclre1c | 1929.7688 | 0.57813948 | 0.19836338 | 2.91454742 | 0.00356205 | 0.06172028 |
| Cfap126 | 82.049119 | 0.58397566 | 0.67673312 | 0.86293348 | 0.38817404 | 0.78942657 |
| Meig1 | 22.241061 | 1.38858913 | 0.90744271 | 1.53022237 | 0.1259617 | 0.53789768 |
| Fam107b | 7431.71873 | -1.0233208 | 0.19423614 | -5.2684364 | 1.375907230 | 1.265524296 |
| Fcgr2b | 257.907998 | -1.5780464 | 0.49352059 | -3.197529 | 0.0013861 | 0.029592 |
| Frmd4a | 461.60664 | -0.0081178 | 0.3684034 | -0.022035 | 0.98242006 | 0.99527491 |
| Dusp12 | 1429.60029 | 0.3008349 | 0.36330823 | 0.82804315 | 0.40764606 | 0.8005951 |
| Sephs1 | 3684.14922 | -0.1488503 | 0.21027328 | -0.7078898 | 0.47901367 | 0.83869135 |
| Atf6 | 1923.50942 | -0.2270236 | 0.25167903 | -0.9020362 | 0.36703766 | 0.77293367 |
| Phyh | 1630.69694 | 0.17440325 | 0.2016581 | 0.86484622 | 0.3871232 | 0.78911204 |
| Uhmk1 | 409.868021 | -0.6438016 | 0.82947366 | -0.7761568 | 0.43765642 | 0.8185065 |
| Mcm10 | 4511.70261 | -0.1001401 | 0.168584 | -0.594007 | 0.55250743 | 0.8709824 |
| Uap1 | 932.981116 | -0.0724917 | 0.37651964 | -0.1925309 | 0.84732636 | 0.96765585 |
| Optn | 32.5058469 | 1.46769647 | 1.01111858 | 1.45155721 | 0.14662476 | 0.57365079 |
| Hsd17b7 | 409.988286 | 0.67169696 | 0.3367295 | 1.99476717 | 0.04606828 | 0.33169704 |
| Enkur | 12.8589423 | -0.7611252 | 1.2129198 | -0.6275148 | 0.53032181 | 0.86349351 |
| Nuf2 | 3271.25009 | 0.40064451 | 0.24121465 | 1.66094604 | 0.09672429 | 0.4773006 |
| Aldh9a1 | 7791.35718 | 0.14687916 | 0.21246263 | 0.6913176 | 0.48936597 | 0.84401986 |
| Mgst3 | 683.464437 | 0.94436271 | 0.44290803 | 2.13218691 | 0.03299148 | 0.26977595 |
| Eef1aknmt | 1039.82125 | 0.00480996 | 0.23181222 | 0.02074939 | 0.98344557 | 0.99564261 |
| Vamp4 | 1405.92264 | 0.37440506 | 0.34696918 | 1.07907296 | 0.2805552 | 0.7155865 |
| Pigc | 1251.09879 | 0.61053306 | 0.43036537 | 1.41863892 | 0.15600431 | 0.58521232 |
| Prdx6 | 8112.51762 | -0.5032168 | 0.15044877 | -3.3447718 | 0.0008235 | 0.02006755 |
| Klhl20 | 233.114303 | -0.1245981 | 0.3428819 | -0.3633848 | 0.71631746 | 0.93235217 |

| | | | | | | |
|---|---|---|---|---|---|---|
| Nsun6 | 400.383871 | 0.53084978 | 0.47731787 | 1.11215149 | 0.26607302 | 0.7023537 |
| Cenpl | 1417.31295 | -0.1342295 | 0.25497464 | -0.5264425 | 0.59858083 | 0.889626 |
| Dars2 | 2134.40458 | -0.5157016 | 0.2520753 | -2.0458236 | 0.04077373 | 0.30763217 |
| Mrc1 | 12582.995 | 2.18195705 | 0.38570423 | 5.65707314 | 1.539761579 | 1.962073271 |
| Serpinc1 | 16.973867 | 0.85701112 | 1.62914088 | 0.52605096 | 0.5988528 | 0.88980521 |
| Stam | 1024.83435 | -0.0900705 | 0.2671957 | -0.3370955 | 0.73604494 | 0.9377627 |
| Rabgap1l | 132.516167 | 0.06077154 | 0.37471269 | 0.1621817 | 0.87116277 | 0.97268995 |
| Trdmt1 | 572.492192 | -0.0235275 | 0.38584269 | -0.0609769 | 0.95137761 | 0.99115284 |
| Rsu1 | 4089.43432 | -0.0590736 | 0.17337269 | -0.3407321 | 0.73330525 | 0.93728136 |
| Vim | 9331.94935 | -0.4456048 | 0.18067372 | -2.4663508 | 0.01364976 | 0.15812263 |
| Pter | 1141.63388 | 0.17305207 | 0.23879987 | 0.72467404 | 0.46865202 | 0.83381643 |
| Pip4k2a | 1180.98042 | 0.02175841 | 0.20304616 | 0.10715992 | 0.91466211 | 0.98347559 |
| Bmi1 | 225.918195 | 0.17915672 | 0.28834696 | 0.6213234 | 0.53438686 | 0.8652846 |
| Dnajc1 | 561.385991 | -0.2465292 | 0.27234987 | -0.9051928 | 0.36536323 | 0.77139945 |
| Mllt10 | 483.73131 | 0.01112611 | 0.48743486 | 0.02282584 | 0.9817892 | 0.99523682 |
| Plxdc2 | 260.836113 | -0.6896324 | 0.34858336 | -1.978386 | 0.04788518 | 0.33762502 |
| Nek6 | 6211.72099 | 0.4694762 | 0.15489003 | 3.03102915 | 0.00243722 | 0.04608111 |
| Psmb7 | 9301.79569 | 0.09930524 | 0.26497208 | 0.37477622 | 0.7078269 | 0.92950127 |
| Ppp6c | 3212.37224 | 0.19505936 | 0.20638908 | 0.94510501 | 0.34460527 | 0.75972469 |
| Golga1 | 284.255519 | 0.43770033 | 0.34553615 | 1.26672806 | 0.20525254 | 0.63913241 |
| Arpc5l | 3797.67084 | -0.2289907 | 0.35529729 | -0.6445046 | 0.51924828 | 0.8576517 |
| Orc4 | 2420.54497 | 0.19800951 | 0.33035626 | 0.59938175 | 0.54891834 | 0.87082325 |
| Kif5c | 53.8522111 | 2.78055586 | 1.40373528 | 1.98082637 | 0.04761075 | 0.33685499 |
| Mmadhc | 3525.16071 | 0.09219502 | 0.22763432 | 0.40501372 | 0.68546742 | 0.92314423 |
| Mindy3 | 1416.29116 | -0.1501761 | 0.30842825 | -0.4869078 | 0.6263237 | 0.90192087 |
| Itga8 | 22.8069893 | -2.5884932 | 1.17991329 | -2.1937995 | 0.02824983 | 0.24951635 |
| Spopl | 88.7855308 | 2.11667791 | 0.94856415 | 2.23145468 | 0.02565103 | 0.23347396 |
| Pfkfb3 | 1032.88656 | 0.34489592 | 0.53774259 | 0.64137737 | 0.52127753 | 0.85801777 |
| Yme1l1 | 5167.25544 | 0.2924766 | 0.26854973 | 1.08909658 | 0.2761113 | 0.7118544 |
| Prkcq | 49.8203394 | -1.2092637 | 0.98140853 | -1.2321716 | 0.21788498 | 0.65150232 |
| Mastl | 532.742124 | -0.0634726 | 0.24598447 | -0.2580349 | 0.79637995 | 0.9537323 |
| Acbd5 | 931.318282 | 0.41581694 | 0.32417251 | 1.28270263 | 0.19959628 | 0.6323909 |
| Abi2 | 1165.20804 | 0.06818341 | 0.23619055 | 0.28867967 | 0.77282653 | 0.94625031 |
| Pdss1 | 1595.104 | -0.4123587 | 0.2050302 | -2.0112097 | 0.04430332 | 0.32302252 |
| Pkn3 | 344.9237 | 0.34702725 | 0.32836662 | 1.05682865 | 0.2905898 | 0.72461526 |
| Apbb1ip | 614.208479 | -0.5532596 | 0.25852818 | -2.1400361 | 0.03235185 | 0.26700278 |
| Zbtb43 | 371.68231 | -0.4039621 | 0.49185082 | -0.8213102 | 0.41146958 | 0.80231592 |
| Odf2 | 2573.43398 | -0.2938352 | 0.28191088 | -1.0422982 | 0.29727344 | 0.72954201 |
| Slc2a8 | 882.799867 | 0.9855248 | 0.30247072 | 3.25824859 | 0.00112102 | 0.02561249 |
| Lrsam1 | 246.393503 | 0.68501774 | 0.30034965 | 2.28073427 | 0.02256418 | 0.21531012 |
| Fam129b | 385.935005 | -0.2895995 | 0.35168286 | -0.8234677 | 0.4102421 | 0.80117888 |
| Stxbp1 | 196.394147 | 1.16871482 | 0.40230226 | 2.9050665 | 0.00367175 | 0.06317374 |
| Coq4 | 1340.63378 | -0.0610613 | 0.28079604 | -0.2174578 | 0.82785162 | 0.96432112 |
| Med27 | 1330.20019 | -0.3223922 | 0.19873258 | -1.6222415 | 0.10475166 | 0.49501875 |
| Ttf1 | 745.003622 | 0.06216085 | 0.28838331 | 0.21554942 | 0.82933899 | 0.96462215 |
| Ddx31 | 1527.69403 | -0.1127061 | 0.24171833 | -0.4662702 | 0.64102211 | 0.90685007 |

| | | | | | | |
|---|---|---|---|---|---|---|
| Ak8 | 243.403851 | 0.0103277 | 0.43321232 | 0.0238398 | 0.98098039 | 0.99522729 |
| Spaca9 | 494.804822 | -0.1730609 | 0.25087916 | -0.6898178 | 0.49030878 | 0.84466234 |
| Dpm2 | 1976.34899 | 0.23847927 | 0.17026239 | 1.40065737 | 0.16131656 | 0.59141542 |
| St6galnac6 | 290.94874 | -0.472858 | 0.40545697 | -1.1662347 | 0.24351957 | 0.67889128 |
| Tsc1 | 375.716517 | 0.27842694 | 0.44158454 | 0.63051786 | 0.52835582 | 0.86212842 |
| Eng | 988.688849 | 0.25248398 | 0.31770665 | 0.79470788 | 0.42678345 | 0.81021713 |
| Gfi1b | 21.7794816 | -2.1895498 | 1.34940889 | -1.6225992 | 0.1046751 | 0.49501875 |
| Gtf3c5 | 1962.85257 | -0.1639246 | 0.20203792 | -0.8113556 | 0.4171615 | 0.8049685 |
| Ak1 | 179.480075 | 0.23167227 | 0.40604808 | 0.57055379 | 0.56830215 | 0.87933724 |
| Slc25a25 | 1175.16249 | -0.1203016 | 0.23176753 | -0.5190613 | 0.60371798 | 0.89232143 |
| Ptges2 | 466.308173 | 0.4342089 | 0.34580775 | 1.25563669 | 0.20924772 | 0.64266316 |
| Ralgds | 357.634569 | 0.67881173 | 0.41425948 | 1.63861485 | 0.1012935 | 0.48631215 |
| Nr4a2 | 415.581055 | -1.1338963 | 0.54676042 | -2.0738449 | 0.03809372 | 0.29493701 |
| Gpd2 | 2471.64037 | -0.8108185 | 0.25144373 | -3.224652 | 0.00126126 | 0.02755175 |
| Cytip | 1668.59031 | -1.1540219 | 0.26196085 | -4.4053221 | 1.056268043 | 0.00054751 |
| Fcnb | 125.501797 | -6.6548642 | 0.82112437 | -8.1045752 | 5.293012959 | 2.398126945 |
| Acvr1 | 132.228526 | 0.16491568 | 0.57299352 | 0.28781421 | 0.77348896 | 0.94651052 |
| Abl1 | 809.839091 | 0.04918697 | 0.2728669 | 0.18025995 | 0.8569485 | 0.96936093 |
| Fubp3 | 1237.42368 | 0.53010086 | 0.25281669 | 2.09677954 | 0.0360131 | 0.28477582 |
| Tor1b | 1131.90446 | -0.3296872 | 0.20308297 | -1.6234113 | 0.1045015 | 0.49501875 |
| Tor1a | 3146.9759 | 0.12249434 | 0.33619528 | 0.36435474 | 0.71559313 | 0.93235217 |
| BC005624 | 1707.52478 | -0.1781471 | 0.19948219 | -0.8930477 | 0.37183164 | 0.77714275 |
| Crat | 2430.83693 | 0.30090858 | 0.24343051 | 1.23611697 | 0.21641509 | 0.64982959 |
| Usp20 | 356.410481 | 0.35495038 | 0.36232053 | 0.97965848 | 0.32725473 | 0.74811039 |
| Dolpp1 | 1818.65292 | 0.05515118 | 0.20170845 | 0.27342025 | 0.78453019 | 0.95094799 |
| Ntmt1 | 1594.52559 | 0.30130727 | 0.30546117 | 0.98640123 | 0.32393626 | 0.74583177 |
| Miga2 | 452.299345 | -0.4881032 | 0.35471361 | -1.3760487 | 0.16880657 | 0.60312169 |
| Sh3glb2 | 700.738838 | 0.06984162 | 0.26540634 | 0.26314977 | 0.79243515 | 0.95297175 |
| Hspa5 | 86966.0247 | 0.26555686 | 0.16520099 | 1.60747736 | 0.10794969 | 0.50301538 |
| Kynu | 217.464093 | -0.5337498 | 0.3232068 | -1.6514189 | 0.09865307 | 0.48025164 |
| Gapvd1 | 2776.06312 | 0.34223315 | 0.21607277 | 1.5838791 | 0.11322124 | 0.51285577 |
| Psmd5 | 4710.31175 | -0.1022662 | 0.2085676 | -0.4903264 | 0.62390296 | 0.90017327 |
| Zeb2 | 2606.36629 | 0.02890434 | 0.27644554 | 0.10455709 | 0.91672727 | 0.98389485 |
| Phf19 | 100.55414 | -0.8518517 | 0.91373958 | -0.9322696 | 0.35119719 | 0.76309919 |
| Traf1 | 25.865145 | -0.2920992 | 1.88095691 | -0.1552929 | 0.87659043 | 0.9745677 |
| Rab14 | 9985.53253 | 0.16201812 | 0.18579256 | 0.87203774 | 0.38318779 | 0.78636743 |
| Gsn | 7773.87878 | -0.0420132 | 0.17936086 | -0.2342383 | 0.81479998 | 0.95996209 |
| Stom | 1437.27462 | 0.87467499 | 0.24030729 | 3.63981876 | 0.00027283 | 0.00826121 |
| Dab2ip | 55.5112464 | -0.3662287 | 0.62507479 | -0.5858958 | 0.55794557 | 0.87291829 |
| Mrrf | 1556.03581 | 0.3745363 | 0.3102765 | 1.20710496 | 0.22739173 | 0.66104634 |
| Rbm18 | 2123.18337 | 0.31099017 | 0.22903961 | 1.35780086 | 0.17452688 | 0.60656845 |
| Gca | 108.767499 | -0.5023372 | 0.61467282 | -0.8172432 | 0.41378943 | 0.80395688 |
| Morn5 | 12.4622654 | 0.33206653 | 1.4255848 | 0.23293355 | 0.815813 | 0.96063519 |
| Ndufa8 | 5332.48184 | -0.0162585 | 0.30773451 | -0.052833 | 0.95786497 | 0.99242502 |
| Ifih1 | 449.454193 | 0.2729668 | 0.29679721 | 0.91970811 | 0.35772531 | 0.76674184 |
| Psmd14 | 13455.35 | 0.31864674 | 0.37139704 | 0.85796791 | 0.39091019 | 0.7899826 |

| | | | | | | |
|---|---|---|---|---|---|---|
| Strbp | 176.995206 | -0.3292567 | 0.7000053 | -0.4703632 | 0.63809555 | 0.90599527 |
| Wdr5 | 3610.68867 | -0.2978987 | 0.17584408 | -1.694107 | 0.09024496 | 0.45941183 |
| Brd3 | 1924.00394 | -0.178526 | 0.33787785 | -0.5283745 | 0.59723943 | 0.88891958 |
| Egfl7 | 1718.65194 | 0.55502591 | 0.33442365 | 1.65964907 | 0.09698507 | 0.47799662 |
| Agpat2 | 404.383179 | -0.3075923 | 0.38124585 | -0.8068084 | 0.4197769 | 0.80563709 |
| Notch1 | 384.755856 | -0.1604454 | 0.33706086 | -0.4760133 | 0.63406492 | 0.90465549 |
| Sec16a | 658.261882 | 0.24713329 | 0.55988151 | 0.44140284 | 0.65892139 | 0.91280998 |
| Inpp5e | 331.185607 | 0.12827233 | 0.54654216 | 0.23469795 | 0.81444318 | 0.95988497 |
| Pmpca | 10530.7462 | -0.0820708 | 0.16889058 | -0.4859404 | 0.62700943 | 0.90205481 |
| Entr1 | 2138.97011 | -0.4366918 | 0.29618636 | -1.474382 | 0.14037878 | 0.56414376 |
| Card9 | 640.803378 | -0.4421732 | 0.20806453 | -2.1251735 | 0.03357214 | 0.27324551 |
| Gpsm1 | 285.332465 | -0.0289444 | 0.41026685 | -0.0705502 | 0.94375577 | 0.98992676 |
| Nacc2 | 98.246444 | 1.33320664 | 0.85448982 | 1.56023701 | 0.11870388 | 0.52515149 |
| Camsap1 | 80.2552031 | 1.71448463 | 0.64465976 | 2.65951863 | 0.00782524 | 0.10907646 |
| Fcna | 2464.9436 | 3.99672498 | 0.48232819 | 8.28631852 | 1.168077018 | 6.212646159 |
| Tmem141 | 864.897629 | 1.75669874 | 0.41832983 | 4.19931498 | 2.677236888 | 0.00116137 |
| Mamdc4 | 13.7151548 | -0.8685522 | 1.86699178 | -0.4652148 | 0.64177764 | 0.90702012 |
| Traf2 | 2117.42228 | 0.04012014 | 0.27288988 | 0.14701953 | 0.88311661 | 0.97651319 |
| Abca2 | 77.7708572 | -1.094908 | 1.17703662 | -0.9302243 | 0.35225498 | 0.76348958 |
| Nmi | 932.803067 | 0.07805318 | 0.33699998 | 0.23161181 | 0.81683953 | 0.96079676 |
| Sapcd2 | 386.331736 | 0.09915529 | 0.35970444 | 0.27565768 | 0.78281101 | 0.95075701 |
| Uap1l1 | 5323.24047 | 0.80328358 | 0.17275061 | 4.64996088 | 3.319979990 | 0.00020307 |
| Dpp7 | 2794.48675 | 1.227304 | 0.29553436 | 4.15283021 | 3.283884001 | 0.00139971 |
| Arl6ip6 | 1848.16892 | -0.2788452 | 0.25200785 | -1.1064942 | 0.26851265 | 0.70532185 |
| Anapc2 | 5984.50817 | 0.20872764 | 0.17986626 | 1.16046023 | 0.24586148 | 0.68117517 |
| Ssna1 | 4303.63606 | 0.28281966 | 0.34094704 | 0.82951198 | 0.40681476 | 0.79992362 |
| Rbms1 | 6908.27037 | 0.16289339 | 0.22512482 | 0.72356922 | 0.46933024 | 0.83381643 |
| Arrdc1 | 2119.52465 | -0.0513471 | 0.18728645 | -0.2741632 | 0.78395919 | 0.95094799 |
| Zmynd19 | 129.389785 | -0.7730625 | 0.60414752 | -1.2795922 | 0.20068858 | 0.63371798 |
| Dph7 | 897.378264 | 0.16179159 | 0.30215728 | 0.53545488 | 0.59233535 | 0.88797037 |
| Pax8 | 10.1665745 | -1.2386797 | 2.08234065 | -0.5948497 | 0.55194395 | 0.87093412 |
| March7 | 1355.65108 | -0.0587987 | 0.20444263 | -0.2876048 | 0.77364925 | 0.94659445 |
| Psd4 | 2032.80827 | -0.0764902 | 0.28740426 | -0.2661415 | 0.79013027 | 0.95261385 |
| Ly75 | 26.3143896 | -2.2935525 | 1.19486272 | -1.9195113 | 0.05491966 | 0.36084578 |
| Il1rn | 204.517434 | -0.0985588 | 0.51634113 | -0.1908793 | 0.8486202 | 0.96765585 |
| Hnmt | 179.703469 | 1.78225924 | 0.39262914 | 4.53929435 | 5.644279846 | 0.00032115 |
| Baz2b | 542.107073 | 0.1812089 | 0.40671216 | 0.44554581 | 0.65592536 | 0.91165371 |
| Wdsub1 | 426.700421 | 0.02886371 | 0.47201732 | 0.06114968 | 0.95124 | 0.99115284 |
| Pkp4 | 378.92986 | 0.99250698 | 0.36553565 | 2.71521257 | 0.00662332 | 0.09868834 |
| Galnt3 | 52.0056268 | -0.3317333 | 1.10114398 | -0.3012624 | 0.76321437 | 0.94300309 |
| Nup35 | 2932.94069 | -0.1268684 | 0.29321995 | -0.4326731 | 0.66525229 | 0.9149286 |
| Dusp19 | 618.120407 | 0.45855946 | 0.4213543 | 1.08829898 | 0.27646314 | 0.7118544 |
| Nckap1 | 22.1042711 | 0.68020647 | 2.13075337 | 0.31923285 | 0.74954995 | 0.9399191 |
| Dnajc10 | 2975.66556 | 0.36400978 | 0.22024717 | 1.65273306 | 0.0983852 | 0.48011435 |
| Itprid2 | 827.0851 | 0.05930141 | 0.42980005 | 0.13797442 | 0.89026064 | 0.97871887 |
| Itga4 | 1049.07715 | 0.04385319 | 0.40922219 | 0.1071623 | 0.91466022 | 0.98347559 |

| | | | | | | |
|---|---|---|---|---|---|---|
| Slc25a12 | 6245.79324 | -1.0796901 | 0.22882124 | -4.7184871 | 2.376049837 | 0.00015209 |
| Ube2e3 | 3529.58725 | 0.38474511 | 0.19981238 | 1.92553187 | 0.05416285 | 0.35989907 |
| Dync1i2 | 3466.08378 | 0.38153145 | 0.15112855 | 2.52454923 | 0.01158468 | 0.14165533 |
| Cwc22 | 1542.56871 | 0.10922833 | 0.29670133 | 0.36814238 | 0.71276707 | 0.93235217 |
| Hat1 | 12966.6059 | -0.0197985 | 0.16971503 | -0.1166571 | 0.90713175 | 0.98176897 |
| Stk39 | 54.4823583 | 2.38163663 | 0.60425719 | 3.94142873 | 8.099770357 | 0.00301169 |
| Cers6 | 679.945706 | -0.1058503 | 0.20186426 | -0.5243639 | 0.60002546 | 0.8902352 |
| Ssrp1 | 8832.19043 | -0.3868443 | 0.19140833 | -2.0210422 | 0.0432754 | 0.31923815 |
| Slc43a3 | 2367.19505 | 0.30223374 | 0.17608964 | 1.71636303 | 0.08609562 | 0.44836429 |
| Slc43a1 | 586.774746 | -0.3277327 | 0.31357839 | -1.0451381 | 0.29595913 | 0.72859877 |
| Timm10 | 1712.02305 | 0.27119093 | 0.38760295 | 0.69966168 | 0.48413861 | 0.84128398 |
| Ube2l6 | 304.274277 | 1.10476129 | 0.56111517 | 1.96886726 | 0.04896834 | 0.34113306 |
| Clp1 | 2526.79115 | 0.14042597 | 0.23729406 | 0.5917804 | 0.55399765 | 0.87165957 |
| Med19 | 975.967737 | 0.19015202 | 0.19393239 | 0.98050678 | 0.32683602 | 0.74811039 |
| Tfpi | 2571.67498 | 0.34729611 | 0.34824651 | 0.9972709 | 0.31863304 | 0.7438622 |
| Fastkd1 | 238.907995 | -0.4337218 | 0.34322852 | -1.2636533 | 0.2063545 | 0.64101946 |
| Itgav | 340.697947 | 0.09545916 | 0.56446926 | 0.16911312 | 0.86570767 | 0.9705375 |
| Phospho2 | 1095.1383 | 0.26024905 | 0.37167268 | 0.70021033 | 0.48379596 | 0.84128398 |
| Zc3h15 | 2562.05234 | -0.2559647 | 0.20565208 | -1.2446492 | 0.21326071 | 0.64606694 |
| Mtx2 | 3205.27318 | 0.12792273 | 0.27951938 | 0.45765244 | 0.64720216 | 0.90950305 |
| Atf2 | 1704.83742 | 0.26774832 | 0.2661501 | 1.00600494 | 0.3144132 | 0.74065408 |
| Chrna1 | 23.941429 | 0.5890644 | 1.11897117 | 0.52643394 | 0.59858674 | 0.889626 |
| Ola1 | 3713.52504 | -0.2514365 | 0.19189849 | -1.310258 | 0.19010858 | 0.62090756 |
| Sp3 | 1886.17381 | -0.1388239 | 0.33856364 | -0.4100377 | 0.68177827 | 0.92061965 |
| Itga6 | 1876.28872 | 0.03415935 | 0.22932259 | 0.14895763 | 0.88158706 | 0.97582756 |
| Kif18a | 943.950953 | -0.0151905 | 0.30048411 | -0.0505533 | 0.95968145 | 0.99287747 |
| Arl14ep | 1453.53371 | 0.05553738 | 0.18866146 | 0.29437588 | 0.76847069 | 0.94401697 |
| Slc12a6 | 534.426754 | 0.01655458 | 0.23791875 | 0.0695808 | 0.94452732 | 0.99017934 |
| Emc4 | 3214.87374 | 0.26597875 | 0.33615041 | 0.79124921 | 0.42879858 | 0.81136783 |
| Katnbl1 | 2457.95319 | 0.42623875 | 0.19899865 | 2.14191783 | 0.03220009 | 0.2665769 |
| Nop10 | 6120.70264 | 0.09199476 | 0.40977067 | 0.22450303 | 0.82236591 | 0.96247351 |
| Lpcat4 | 444.320892 | -1.764537 | 0.41236045 | -4.279113 | 1.876395500 | 0.00086293 |
| Ccdc34 | 508.331829 | -0.0497791 | 0.22762367 | -0.2186905 | 0.82689114 | 0.96379955 |
| Lin7c | 1290.11271 | 0.24650106 | 0.24751591 | 0.99589985 | 0.31929881 | 0.7440831 |
| Commd9 | 2633.34757 | 0.41022435 | 0.20080512 | 2.04289792 | 0.04106254 | 0.30873884 |
| Traf6 | 671.131041 | -0.2549794 | 0.30476897 | -0.8366317 | 0.4027996 | 0.79795242 |
| B230118H07 | 683.099423 | -0.0106837 | 0.2613071 | -0.0408857 | 0.96738702 | 0.99443087 |
| Dnajc24 | 705.687364 | 0.59036194 | 0.35953087 | 1.64203405 | 0.10058295 | 0.48480347 |
| Elp4 | 1094.87576 | -0.1281034 | 0.23989994 | -0.5339868 | 0.59335064 | 0.88797037 |
| Eif3m | 12740.5913 | 0.27955136 | 0.37196109 | 0.75156075 | 0.45231525 | 0.82603347 |
| Depdc7 | 553.40817 | -0.1104588 | 0.24236581 | -0.4557525 | 0.64856796 | 0.90973357 |
| Tcp11l1 | 298.640355 | 0.65411949 | 0.27303383 | 2.39574525 | 0.01658661 | 0.17972015 |
| Cstf3 | 2285.41397 | 0.23341652 | 0.16605159 | 1.40568673 | 0.15981717 | 0.58912704 |
| Hipk3 | 616.764105 | -0.2578421 | 0.29663446 | -0.8692252 | 0.38472397 | 0.78748265 |
| Fbxo3 | 2974.26518 | 0.40273472 | 0.25749956 | 1.564021 | 0.11781264 | 0.52437333 |
| Caprin1 | 19701.5649 | -0.2202744 | 0.21952963 | -1.0033924 | 0.31567155 | 0.74186126 |

| | | | | | | |
|---|---|---|---|---|---|---|
| Nat10 | 1994.98705 | -0.4328986 | 0.33493694 | -1.2924779 | 0.19619168 | 0.62827561 |
| Cat | 4693.33255 | 0.10857793 | 0.15842748 | 0.68534783 | 0.49312446 | 0.8462951 |
| Trim44 | 436.621257 | -1.2137811 | 0.36256773 | -3.3477361 | 0.00081475 | 0.01993356 |
| Api5 | 13574.115 | -0.173743 | 0.16972283 | -1.0236866 | 0.30598335 | 0.73554152 |
| Ttc17 | 503.891747 | -0.5091801 | 0.34030628 | -1.4962406 | 0.13459096 | 0.55585235 |
| Hsd17b12 | 7379.41288 | 0.16937222 | 0.16973979 | 0.99783449 | 0.31835962 | 0.74336577 |
| Ext2 | 3225.51304 | -0.0960355 | 0.21134495 | -0.4544018 | 0.64953964 | 0.90973357 |
| Gatm | 2062.17188 | 1.63719236 | 0.33385673 | 4.90387713 | 9.396327985 | 6.841980967 |
| Sema6d | 63.8474681 | 4.6393897 | 1.06281783 | 4.36517865 | 1.270189072 | 0.00064743 |
| Myef2 | 2094.1678 | 0.11210299 | 0.1811993 | 0.61867231 | 0.53613225 | 0.8654666 |
| Dut | 3915.57495 | -0.4298544 | 0.27742841 | -1.5494247 | 0.12127965 | 0.5291774 |
| Cops2 | 4310.44123 | -0.1920103 | 0.16638333 | -1.1540234 | 0.24849056 | 0.68263757 |
| Galk2 | 4170.50949 | 0.4460196 | 0.19471559 | 2.29062083 | 0.02198535 | 0.21347378 |
| Meis2 | 26.0738035 | 1.6587429 | 1.34296365 | 1.23513611 | 0.21677985 | 0.6504459 |
| Cd82 | 135.528366 | -0.1480968 | 0.48493855 | -0.3053928 | 0.76006698 | 0.94215089 |
| Tspan18 | 411.155835 | 1.77200957 | 0.48316083 | 3.66753562 | 0.0002449 | 0.00758444 |
| Chst1 | 11.626969 | -0.0666958 | 1.61620636 | -0.0412669 | 0.96708312 | 0.99443087 |
| Pex16 | 757.397814 | 0.01963983 | 0.36441531 | 0.05389409 | 0.95701954 | 0.99217217 |
| Sord | 2850.93718 | 0.71387505 | 0.31086887 | 2.29638639 | 0.0216538 | 0.21157421 |
| Creb3l1 | 155.446276 | -0.3697928 | 0.42770519 | -0.8645974 | 0.38725982 | 0.78911204 |
| Eif3j1 | 2275.62844 | -0.3118968 | 0.18868621 | -1.6529922 | 0.09833246 | 0.48011435 |
| Mdk | 154.702747 | 0.51780958 | 0.7528954 | 0.68775766 | 0.49160541 | 0.84499212 |
| Wdr76 | 670.369979 | -0.2263072 | 0.30981882 | -0.7304501 | 0.46511509 | 0.83339036 |
| Harbi1 | 362.172615 | 0.0299332 | 0.35246673 | 0.08492491 | 0.93232109 | 0.98730569 |
| Atg13 | 1210.29087 | -0.1598621 | 0.28845305 | -0.5542049 | 0.57943862 | 0.88286755 |
| Hypk | 2413.31955 | -0.2814078 | 0.22760618 | -1.2363805 | 0.21631716 | 0.64982959 |
| Ell3 | 27.9825565 | 0.49989388 | 1.66875401 | 0.29956116 | 0.76451192 | 0.94307414 |
| Arhgap1 | 6009.72401 | -0.0692459 | 0.23051955 | -0.3003905 | 0.7638793 | 0.94300309 |
| Pdia3 | 56376.1 | 0.32036639 | 0.19803271 | 1.61774476 | 0.10571761 | 0.49740138 |
| Lrp4 | 90.0421553 | 0.3062872 | 0.60841536 | 0.50341793 | 0.61467047 | 0.89844745 |
| Arfgap2 | 4810.27918 | -0.0022945 | 0.19171449 | -0.0119684 | 0.9904508 | 0.9966603 |
| Pacsin3 | 51.1041269 | 0.53249642 | 0.66619582 | 0.79930917 | 0.42411116 | 0.80933211 |
| Adal | 615.596092 | -0.1415745 | 0.2464876 | -0.5743678 | 0.56571892 | 0.87845133 |
| Tubgcp4 | 1549.05016 | -0.0124208 | 0.3435106 | -0.0361585 | 0.97115594 | 0.99443087 |
| Ubr1 | 489.506839 | -0.3253102 | 0.2483875 | -1.3096884 | 0.19030127 | 0.6209089 |
| Mkks | 525.555836 | 0.16177425 | 0.53261506 | 0.30373578 | 0.7613292 | 0.94247845 |
| Jag1 | 29.2888854 | 1.14648067 | 0.9927125 | 1.15489698 | 0.24813261 | 0.68235696 |
| Slx4ip | 200.723432 | -0.3699892 | 0.40106292 | -0.9225216 | 0.35625656 | 0.76538225 |
| Mtch2 | 12402.4746 | 0.40601317 | 0.36953571 | 1.09871161 | 0.27189388 | 0.70940613 |
| Cdan1 | 193.55879 | -0.9964793 | 0.50558996 | -1.9709237 | 0.0487326 | 0.34046898 |
| Haus2 | 1537.83727 | -0.0105015 | 0.25476768 | -0.04122 | 0.96712053 | 0.99443087 |
| Lrrc57 | 913.725576 | 0.3985368 | 0.19238767 | 2.07152978 | 0.03830931 | 0.29553145 |
| Snap23 | 5116.74921 | 0.72024453 | 0.29830094 | 2.41448962 | 0.01575727 | 0.17444227 |
| Zfp106 | 2422.84267 | -0.1290981 | 0.42967417 | -0.3004557 | 0.76382958 | 0.94300309 |
| Vps39 | 1539.98046 | 0.22831288 | 0.26341488 | 0.86674255 | 0.38608308 | 0.7886896 |
| Ehd4 | 17031.9886 | 0.50582561 | 0.22414744 | 2.25666461 | 0.02402904 | 0.22438721 |

| | | | | | | |
|---|---|---|---|---|---|---|
| Tyro3 | 9.88719582 | 2.334726 | 2.20260038 | 1.05998619 | 0.28915088 | 0.72395811 |
| Ubox5 | 876.901487 | -0.5563299 | 0.21229683 | -2.6205285 | 0.00877936 | 0.11893454 |
| Ptpra | 1513.15534 | -0.0612931 | 0.24759655 | -0.2475523 | 0.80448079 | 0.9570372 |
| Rtf1 | 239.788779 | -0.2102965 | 0.60442722 | -0.3479269 | 0.72789506 | 0.93576253 |
| Ndufaf1 | 1117.42438 | -0.065809 | 0.22195658 | -0.2964947 | 0.76685228 | 0.94381438 |
| Nusap1 | 2955.52532 | -0.0209878 | 0.19229403 | -0.1091444 | 0.91308796 | 0.98292561 |
| 4930402H24 | 245.414478 | 0.5049106 | 0.47939701 | 1.05322017 | 0.2922401 | 0.72584701 |
| Atrn | 309.495854 | -1.1437857 | 0.35863241 | -3.1892982 | 0.00142619 | 0.03028914 |
| Chac1 | 164.826354 | -1.2579881 | 0.57606137 | -2.1837745 | 0.02897882 | 0.25346382 |
| Spint1 | 109.927397 | -1.3935413 | 0.4878825 | -2.8563051 | 0.00428603 | 0.07123778 |
| Siglec1 | 36.8627458 | 0.70460312 | 0.88659663 | 0.79472794 | 0.42677177 | 0.81021713 |
| Rad51 | 4547.39633 | 0.02088107 | 0.29058596 | 0.0718585 | 0.94271452 | 0.98983253 |
| Rpusd2 | 244.871226 | -0.8776092 | 0.55843292 | -1.571557 | 0.11605333 | 0.52084358 |
| Knl1 | 1425.39733 | -0.4213898 | 0.24081466 | -1.7498512 | 0.08014399 | 0.43723638 |
| 1700037H04 | 744.448778 | -0.1266334 | 0.23623171 | -0.5360558 | 0.59191998 | 0.88797037 |
| Cdc25b | 368.966823 | -0.4907328 | 0.56840118 | -0.8633563 | 0.38794159 | 0.7893695 |
| Knstrn | 4421.21113 | 0.12484702 | 0.25049599 | 0.49839928 | 0.61820264 | 0.89878056 |
| Ivd | 2024.31859 | -0.2621737 | 0.2080399 | -1.2602088 | 0.20759407 | 0.6418276 |
| Smox | 818.857338 | 0.30334429 | 0.24797046 | 1.22330818 | 0.22121332 | 0.65469944 |
| Rassf2 | 1927.30318 | -0.2788252 | 0.26390471 | -1.0565374 | 0.29072277 | 0.72461526 |
| Slc23a2 | 621.210924 | 0.26253945 | 0.44396028 | 0.59135798 | 0.55428059 | 0.87165957 |
| Tmem230 | 1137.40001 | -0.0707924 | 0.20580571 | -0.3439768 | 0.73086377 | 0.93653187 |
| Pcna | 14209.3639 | 0.1647352 | 0.23037111 | 0.71508617 | 0.4745558 | 0.83578199 |
| Gpcpd1 | 1072.52272 | 0.53732178 | 0.28322074 | 1.89718375 | 0.05780369 | 0.36898259 |
| Fam98b | 293.863996 | -0.341757 | 0.26808123 | -1.2748262 | 0.20237078 | 0.63738458 |
| Spred1 | 688.021578 | 0.17124897 | 0.25271603 | 0.67763399 | 0.49800379 | 0.84860011 |
| Mcm8 | 980.822268 | -0.027236 | 0.23469133 | -0.1160504 | 0.90761263 | 0.98176897 |
| Crls1 | 135.559848 | 0.27480644 | 0.44761878 | 0.61392963 | 0.53926183 | 0.86719392 |
| Bmp2 | 533.629828 | 1.47694721 | 0.64729893 | 2.28170811 | 0.02250658 | 0.21507833 |
| Hdc | 288.031944 | -7.6170496 | 0.84851932 | -8.9768724 | 2.785727647 | 2.129862894 |
| Gabpb1 | 2337.9133 | -0.0285634 | 0.21176105 | -0.1348849 | 0.89270291 | 0.97941957 |
| Usp8 | 3253.00116 | -0.017837 | 0.27926044 | -0.0638724 | 0.9490718 | 0.99028544 |
| Usp50 | 503.871515 | -0.2635658 | 0.28518938 | -0.9241782 | 0.35539354 | 0.7647925 |
| Trpm7 | 1233.26265 | -0.2948407 | 0.33070834 | -0.8915429 | 0.37263798 | 0.77748705 |
| Sppl2a | 3948.49182 | 0.26008261 | 0.21603355 | 1.20389918 | 0.22862855 | 0.66258779 |
| Stard7 | 5349.90506 | 0.18686315 | 0.25898695 | 0.72151569 | 0.47059229 | 0.83381643 |
| Dusp2 | 3656.14687 | 0.59368609 | 0.56327872 | 1.05398281 | 0.29189079 | 0.72575203 |
| Fahd2a | 327.537923 | 0.63672542 | 0.55152235 | 1.15448706 | 0.24830054 | 0.68257539 |
| Mrps5 | 2191.68818 | -0.2129886 | 0.19525539 | -1.0908207 | 0.2753518 | 0.71181528 |
| Nphp1 | 153.183596 | 0.23864043 | 0.35813865 | 0.66633531 | 0.5051968 | 0.85034006 |
| Bub1 | 4706.15507 | -0.2330843 | 0.19359919 | -1.2039531 | 0.2286077 | 0.66258779 |
| Bcl2l11 | 804.414267 | -0.1602919 | 0.46717239 | -0.3431108 | 0.73151511 | 0.93653187 |
| Ndufaf5 | 788.649136 | 0.48061253 | 0.51827912 | 0.92732374 | 0.35375847 | 0.76401657 |
| Zc3h8 | 143.170765 | -0.2201329 | 0.3423995 | -0.6429123 | 0.520281 | 0.8576517 |
| Ttl | 210.264699 | 0.04620191 | 0.49316556 | 0.09368439 | 0.92535987 | 0.98571293 |
| Polr1b | 1951.40782 | -0.6627855 | 0.25080817 | -2.6425994 | 0.00822723 | 0.11359337 |

| | | | | | | |
|---|---|---|---|---|---|---|
| Slc20a1 | 1614.13915 | 0.05786432 | 0.41004242 | 0.14111788 | 0.88777682 | 0.97832097 |
| Il1b | 4251.02785 | 0.13992515 | 0.44022319 | 0.31785048 | 0.75059835 | 0.9399191 |
| Il1a | 836.155834 | 1.44270344 | 1.05388491 | 1.36893832 | 0.17101856 | 0.60430173 |
| Snrpb | 23119.3862 | -0.2578745 | 0.18747312 | -1.3755278 | 0.16896787 | 0.60312169 |
| Nop56 | 11176.1326 | -0.3109833 | 0.22805758 | -1.3636175 | 0.17268798 | 0.60492813 |
| Idh3b | 8048.26161 | 0.08592258 | 0.24331499 | 0.35313309 | 0.72398867 | 0.93466537 |
| Cpxm1 | 88.3636627 | 1.69135091 | 0.77075583 | 2.1944056 | 0.02820626 | 0.24951635 |
| Vps16 | 2466.1918 | -0.0521115 | 0.15922453 | -0.3272831 | 0.74345376 | 0.93856621 |
| Rrbp1 | 2657.05251 | -0.2446107 | 0.28217246 | -0.8668837 | 0.38600575 | 0.7886896 |
| Snx5 | 24805.488 | 0.5542014 | 0.24660728 | 2.24730353 | 0.02462064 | 0.22749037 |
| Mgme1 | 1315.86501 | 0.00024999 | 0.27060848 | 0.00092379 | 0.99926292 | 0.99973522 |
| Kat14 | 994.040785 | -0.1851859 | 0.20164419 | -0.9183795 | 0.35842023 | 0.76713922 |
| Polr3f | 592.055179 | 0.06683966 | 0.26127083 | 0.25582517 | 0.79808584 | 0.95433165 |
| Rbbp9 | 909.072302 | 0.6152889 | 0.29315433 | 2.09885661 | 0.03582954 | 0.28424306 |
| Sec23b | 5657.91731 | -0.0038483 | 0.20241123 | -0.0190121 | 0.98483142 | 0.99564261 |
| Dtd1 | 1389.34779 | -0.2470579 | 0.25078209 | -0.9851497 | 0.32455052 | 0.74600273 |
| Xrn2 | 6147.04669 | -0.2052147 | 0.16820828 | -1.2200036 | 0.22246351 | 0.65591616 |
| Cd93 | 183.747899 | -1.5292608 | 0.43253608 | -3.5355683 | 0.0004069 | 0.01141651 |
| Napb | 70.5756122 | -0.7325113 | 0.88809996 | -0.8248073 | 0.40948101 | 0.80100711 |
| Gzf1 | 684.215838 | 0.32621169 | 0.22500735 | 1.4497824 | 0.14711921 | 0.57388689 |
| Cst3 | 23321.2926 | -0.1808346 | 0.27291864 | -0.6625954 | 0.50758969 | 0.85197408 |
| Acss1 | 4765.25683 | 0.90131094 | 0.25153317 | 3.58326881 | 0.00033932 | 0.00981304 |
| Gins1 | 2171.65322 | 0.32200526 | 0.2970219 | 1.08411287 | 0.27831476 | 0.71360814 |
| Nsfl1c | 5183.39587 | -0.0710699 | 0.16030025 | -0.4433548 | 0.65750915 | 0.91258818 |
| Fam110a | 806.974801 | 0.1799608 | 0.24032473 | 0.74882347 | 0.45396361 | 0.82695062 |
| Angpt4 | 35.933727 | 3.09372131 | 1.58628166 | 1.9502976 | 0.05114066 | 0.3475576 |
| Slc52a3 | 22.4268934 | 0.29498678 | 1.43358629 | 0.20576842 | 0.83697181 | 0.96683245 |
| Tbc1d20 | 1803.13004 | 0.30104592 | 0.36883947 | 0.81619766 | 0.4143871 | 0.80395688 |
| Rbck1 | 3426.02661 | -0.3898987 | 0.18461215 | -2.1119884 | 0.03468744 | 0.27916547 |
| Tpx2 | 7275.38849 | -0.0917513 | 0.22304618 | -0.4113557 | 0.68081174 | 0.92061965 |
| Pdrg1 | 1990.85522 | 0.14264615 | 0.276473 | 0.51594965 | 0.60588958 | 0.89258167 |
| Kif3b | 262.534301 | 0.93050225 | 0.47488783 | 1.95941482 | 0.05006423 | 0.34252555 |
| Dnmt3b | 1248.34116 | -0.318875 | 0.25357895 | -1.2574979 | 0.20857341 | 0.6418814 |
| Mapre1 | 14636.5201 | -0.0323401 | 0.14410143 | -0.224426 | 0.82242586 | 0.96247351 |
| Cdk5rap1 | 766.730899 | 0.29991137 | 0.32205157 | 0.93125261 | 0.35172291 | 0.76348958 |
| Snta1 | 204.114598 | 1.42405637 | 0.46244508 | 3.07940645 | 0.00207414 | 0.041123 |
| E2f1 | 633.081624 | -0.1349811 | 0.29778601 | -0.4532821 | 0.65034561 | 0.9101573 |
| Fam210b | 316.02979 | 0.86977564 | 0.37640615 | 2.31073705 | 0.02084738 | 0.20583216 |
| Aurka | 9366.21375 | -0.3135807 | 0.18670154 | -1.6795829 | 0.09303849 | 0.46652562 |
| Cstf1 | 2268.93186 | -0.0781261 | 0.19453104 | -0.4016127 | 0.68796907 | 0.92470267 |
| Pkia | 118.454396 | 1.00361851 | 0.4353309 | 2.30541529 | 0.02114333 | 0.20774804 |
| Rtf2 | 4330.53685 | 0.05157561 | 0.16434753 | 0.31382039 | 0.75365746 | 0.94018119 |
| Tpd52 | 7066.63927 | 0.07110683 | 0.17091215 | 0.41604314 | 0.67737843 | 0.91958179 |
| Pag1 | 292.378689 | 0.14345441 | 0.34776047 | 0.41250927 | 0.67996619 | 0.92054299 |
| Rae1 | 6076.83198 | -0.4136437 | 0.17552812 | -2.3565663 | 0.01844478 | 0.19120986 |
| Rbm38 | 2428.01064 | -0.6244073 | 0.26385661 | -2.3664644 | 0.0179589 | 0.18931452 |

| | | | | | | |
|---|---|---|---|---|---|---|
| Zbp1 | 50.112424 | -2.1299895 | 0.90882265 | -2.34368 | 0.01909454 | 0.19530399 |
| Rab22a | 302.356499 | -0.1808604 | 0.64314706 | -0.2812115 | 0.77854815 | 0.94853305 |
| Stx16 | 192.262289 | 0.0103124 | 0.44424739 | 0.02321319 | 0.98148022 | 0.99523682 |
| Gnas | 4695.39873 | -0.269742 | 0.22211948 | -1.2144007 | 0.22459475 | 0.65815511 |
| Impa1 | 4398.02824 | -0.1871714 | 0.17577937 | -1.064809 | 0.28696239 | 0.72260415 |
| Fabp5 | 4788.63807 | 0.25475326 | 0.32392566 | 0.78645595 | 0.43160042 | 0.81342204 |
| Snx16 | 588.724921 | 0.63648306 | 0.24974803 | 2.54850083 | 0.0108187 | 0.13490845 |
| Chmp4c | 10.4293688 | -0.2428943 | 2.08779846 | -0.1163399 | 0.90738316 | 0.98176897 |
| Ptpn1 | 7195.72503 | -0.3517732 | 0.18492125 | -1.9022867 | 0.05713368 | 0.3676572 |
| Nfatc2 | 193.743374 | 1.71368897 | 0.43655987 | 3.92543864 | 8.657183847 | 0.00318986 |
| Atp9a | 167.245791 | -0.5703698 | 0.4779769 | -1.1933 | 0.23275191 | 0.66805784 |
| Sall4 | 24.6048005 | 0.84398586 | 0.99238363 | 0.85046331 | 0.39506755 | 0.79305241 |
| Lrrcc1 | 688.762124 | 0.24139108 | 0.28743962 | 0.8397975 | 0.40102193 | 0.79698916 |
| Zfp64 | 1188.73052 | 0.01192932 | 0.18208747 | 0.06551423 | 0.94776457 | 0.99019648 |
| E2f5 | 306.530984 | 0.37286568 | 0.30503041 | 1.22238858 | 0.22156072 | 0.65514465 |
| Car13 | 490.131998 | 1.40719932 | 0.35831249 | 3.92729631 | 8.590612119 | 0.00317489 |
| Car3 | 8.44402703 | 0.32307711 | 1.81456642 | 0.17804645 | 0.85868649 | 0.96954641 |
| Car2 | 64.3180208 | -1.3195479 | 0.80146663 | -1.6464166 | 0.09967802 | 0.48253314 |
| Psma7 | 18316.0945 | 0.3410336 | 0.44533185 | 0.76579657 | 0.44379736 | 0.82068545 |
| Mrgbp | 475.209013 | -0.3345495 | 0.32411899 | -1.0321812 | 0.30198724 | 0.73361866 |
| Gid8 | 592.58061 | -0.2710148 | 0.26333943 | -1.0291462 | 0.303411 | 0.73436097 |
| Arfgap1 | 1584.10676 | -0.0989136 | 0.20888484 | -0.4735319 | 0.63583376 | 0.90531997 |
| Helz2 | 85.751271 | 0.70853554 | 0.49483279 | 1.43186861 | 0.15218143 | 0.58009243 |
| Zgpat | 711.585375 | -0.3058238 | 0.24666796 | -1.2398196 | 0.21504212 | 0.64854028 |
| Zbtb46 | 138.235579 | 1.08249133 | 0.52043711 | 2.07996566 | 0.03752868 | 0.29185529 |
| Pcmtd2 | 416.909187 | -0.1503856 | 0.34207278 | -0.4396304 | 0.66020483 | 0.91298731 |
| Raly | 957.019659 | 0.08423274 | 0.32055698 | 0.26276993 | 0.79272791 | 0.95297175 |
| Ahcy | 5782.39045 | -0.4445141 | 0.22600967 | -1.9667923 | 0.04920716 | 0.34139551 |
| Itch | 901.725066 | -0.094467 | 0.24059063 | -0.3926463 | 0.69458073 | 0.92609254 |
| Armc1 | 3097.73079 | -0.1900834 | 0.19833181 | -0.9584109 | 0.33785563 | 0.75611716 |
| Mtfr1 | 1910.46097 | 0.11681625 | 0.23296265 | 0.50143766 | 0.61606314 | 0.89878056 |
| Map1lc3a | 205.733724 | 0.36122924 | 0.38145656 | 0.94697347 | 0.3436523 | 0.75947281 |
| Acss2 | 201.645231 | 0.54955948 | 0.47553747 | 1.15565968 | 0.24782039 | 0.68235696 |
| Gss | 1114.16463 | 0.32319832 | 0.28034591 | 1.15285546 | 0.24896972 | 0.68345734 |
| Procr | 43.4201108 | 1.8819491 | 1.29331015 | 1.45514137 | 0.14563013 | 0.57242843 |
| Eif6 | 9208.60727 | -0.2718608 | 0.21176539 | -1.2837831 | 0.19921785 | 0.6318465 |
| Hps3 | 1827.80378 | -0.1431847 | 0.20855 | -0.6865726 | 0.49235208 | 0.84568141 |
| Nfs1 | 2854.94631 | -0.2190249 | 0.16798946 | -1.3038015 | 0.19230128 | 0.62381902 |
| Rbm39 | 7686.09218 | 0.11906932 | 0.16470867 | 0.72290865 | 0.46973601 | 0.83381643 |
| Epb41l1 | 203.454539 | 2.11999907 | 0.78429706 | 2.70305625 | 0.00687051 | 0.10086772 |
| Aar2 | 2295.14538 | 0.28733998 | 0.18483405 | 1.55458358 | 0.12004528 | 0.52767297 |
| Tbl1xr1 | 673.421472 | -0.75915 | 0.51550598 | -1.4726308 | 0.14085061 | 0.56517498 |
| Ndrg3 | 1413.06757 | -0.4032671 | 0.2107968 | -1.9130607 | 0.05574028 | 0.36325309 |
| Dsn1 | 1508.79766 | 0.18892753 | 0.16366801 | 1.15433388 | 0.24836331 | 0.68259456 |
| Rab5if | 9385.75562 | 0.18285764 | 0.33419488 | 0.54715869 | 0.58426971 | 0.88404713 |
| Samhd1 | 6154.798 | -0.0499055 | 0.2121047 | -0.2352871 | 0.81398594 | 0.95988497 |

| | | | | | | |
|---|---|---|---|---|---|---|
| Rbl1 | 1144.19027 | -0.2398673 | 0.20802999 | -1.1530418 | 0.24889324 | 0.68345734 |
| Rpn2 | 19833.3136 | -0.2415904 | 0.17526807 | -1.3784051 | 0.16807826 | 0.60260883 |
| Src | 17.4992336 | 1.39475123 | 1.52013626 | 0.91751724 | 0.35887166 | 0.76764331 |
| Ctnnbl1 | 3191.25835 | -0.1767288 | 0.18149389 | -0.9737454 | 0.33018299 | 0.74978051 |
| Tti1 | 1517.07288 | -0.0868709 | 0.29526016 | -0.294218 | 0.76859134 | 0.94401697 |
| Rprd1b | 1864.12737 | -0.5019075 | 0.25512844 | -1.9672738 | 0.04915166 | 0.34139551 |
| Ralgapb | 172.544449 | 0.24401244 | 0.78084387 | 0.31249837 | 0.7546618 | 0.94048266 |
| Fam83d | 519.890581 | 0.06671633 | 0.32054551 | 0.20813372 | 0.83512456 | 0.96641606 |
| Dhx35 | 682.576697 | 0.01707752 | 0.27389012 | 0.06235171 | 0.95028275 | 0.99069447 |
| Skil | 1684.69156 | 0.02237109 | 0.45990242 | 0.04864313 | 0.9612037 | 0.99338983 |
| Zmat3 | 82.4822971 | 1.12386365 | 0.67952385 | 1.65389876 | 0.09814809 | 0.47978865 |
| Pik3ca | 254.917528 | -0.7282711 | 0.54362609 | -1.3396544 | 0.18035774 | 0.61122544 |
| Zfp639 | 885.945576 | -0.0847598 | 0.30987297 | -0.2735309 | 0.78444512 | 0.95094799 |
| Mfn1 | 417.027951 | 0.48989365 | 0.39048246 | 1.25458556 | 0.20962925 | 0.64347165 |
| Gnb4 | 919.162437 | 0.41023767 | 0.39359885 | 1.04227356 | 0.29728487 | 0.72954201 |
| Ocstamp | 42.1866529 | 1.20205587 | 1.26070115 | 0.95348201 | 0.34034591 | 0.75795585 |
| Actl6a | 6652.18239 | -0.0287875 | 0.21507008 | -0.1338517 | 0.89351982 | 0.97941957 |
| Ndufb5 | 4512.55699 | 0.07667696 | 0.32608946 | 0.23514087 | 0.8140994 | 0.95988497 |
| Ttc14 | 699.451012 | 0.13451473 | 0.28819944 | 0.46674181 | 0.64068461 | 0.90685007 |
| Ncoa3 | 4573.14095 | 0.14957436 | 0.30039172 | 0.49793103 | 0.61853266 | 0.89878056 |
| Dnajc19 | 2627.70281 | 0.43673637 | 0.31981571 | 1.36558759 | 0.17206845 | 0.60451272 |
| Fxr1 | 2488.78346 | -0.2698201 | 0.23497334 | -1.1483008 | 0.2508444 | 0.6842534 |
| Tnik | 59.4282553 | -0.3344981 | 1.28801555 | -0.2597003 | 0.79509492 | 0.95367917 |
| Pld1 | 670.421579 | 0.3721155 | 0.340135 | 1.09402295 | 0.27394491 | 0.7109191 |
| Nceh1 | 1024.28977 | -0.1546691 | 0.26136291 | -0.5917789 | 0.55399864 | 0.87165957 |
| Ect2 | 3196.38237 | 0.22875289 | 0.26753363 | 0.85504348 | 0.39252708 | 0.79093786 |
| Lrrc34 | 179.660849 | 0.55888069 | 0.36771776 | 1.51986319 | 0.12854536 | 0.54411607 |
| Sec62 | 2086.76905 | -0.0389333 | 0.26069618 | -0.1493437 | 0.88128247 | 0.97582756 |
| Dcun1d1 | 2611.99786 | -0.0071954 | 0.22663684 | -0.0317484 | 0.97467268 | 0.9944856 |
| Mccc1 | 1126.66897 | 0.20926776 | 0.19840724 | 1.05473852 | 0.29154493 | 0.72554658 |
| Acad9 | 3435.75768 | -0.1115747 | 0.19853178 | -0.5619994 | 0.57411645 | 0.88059206 |
| Anxa5 | 6985.8916 | 0.82482946 | 0.17087562 | 4.82707525 | 1.385527803 | 9.522000908 |
| Exosc9 | 2941.7876 | -0.0995059 | 0.21778635 | -0.4568969 | 0.64774518 | 0.9097332 |
| Ccna2 | 12105.159 | -0.354408 | 0.14373356 | -2.465729 | 0.01367347 | 0.15824751 |
| Spata5 | 1570.99602 | -0.4227533 | 0.2722417 | -1.5528603 | 0.12045651 | 0.52834152 |
| Slc7a11 | 25.8333585 | -0.2891343 | 1.31353641 | -0.220119 | 0.82577849 | 0.96326388 |
| Rab33b | 460.212343 | 0.19554912 | 0.40186857 | 0.48659968 | 0.62654205 | 0.90198854 |
| Cog6 | 1628.29797 | 0.23402735 | 0.17076169 | 1.37049096 | 0.1705337 | 0.60430173 |
| Ufm1 | 2271.1271 | 0.54374571 | 0.35642436 | 1.52555711 | 0.1271202 | 0.54111158 |
| Postn | 51.1327377 | 2.09647843 | 1.70260419 | 1.23133635 | 0.21819709 | 0.65150232 |
| Supt20 | 2299.72512 | -0.0928419 | 0.18708348 | -0.4962593 | 0.61971144 | 0.89896588 |
| Exosc8 | 4589.85919 | -0.2114163 | 0.27686995 | -0.7635942 | 0.44510913 | 0.82118815 |
| Mbnl1 | 7880.77074 | -0.259818 | 0.3191015 | -0.8142174 | 0.41552041 | 0.80415459 |
| P2ry1 | 52.9624917 | -0.6189415 | 0.94027568 | -0.6582554 | 0.51037403 | 0.85335831 |
| Dhx36 | 407.716748 | -0.3017995 | 0.24764951 | -1.2186557 | 0.2229749 | 0.65615877 |
| Gfm1 | 3087.75001 | -0.2870497 | 0.26295281 | -1.0916397 | 0.27499152 | 0.71147204 |

| | | | | | | |
|---|---|---|---|---|---|---|
| Mfsd1 | 20538.1844 | 0.48540154 | 0.23430767 | 2.07164171 | 0.03829887 | 0.29553145 |
| Schip1 | 23.0180168 | 1.11604572 | 1.20395298 | 0.92698447 | 0.3539346 | 0.76401657 |
| Ift80 | 157.65403 | -0.7894267 | 0.41489706 | -1.902705 | 0.05707905 | 0.36749894 |
| Kpna4 | 1005.69374 | -0.3306023 | 0.23167534 | -1.4270071 | 0.15357786 | 0.58242478 |
| Nmd3 | 5459.30769 | 0.10215141 | 0.25500781 | 0.40058148 | 0.68872829 | 0.92470267 |
| Nbea | 34.6695247 | -0.1310793 | 0.86281038 | -0.1519213 | 0.87924902 | 0.97541065 |
| Wwtr1 | 71.4585815 | 0.65568764 | 0.695982 | 0.94210431 | 0.34613924 | 0.7602916 |
| Ppid | 6780.08621 | -0.6048382 | 0.17068818 | -3.5435272 | 0.00039481 | 0.01112844 |
| Pfn2 | 42.3928415 | 1.01533259 | 0.98730711 | 1.02838577 | 0.3037684 | 0.73436097 |
| Tsc22d2 | 777.967185 | -0.098921 | 0.37453241 | -0.2641187 | 0.79168843 | 0.95275204 |
| Serp1 | 45308.208 | 0.41522281 | 0.27999591 | 1.48296028 | 0.13808496 | 0.56068131 |
| Etfdh | 2609.08773 | 0.15302494 | 0.20645589 | 0.7411992 | 0.45857267 | 0.82986925 |
| Eif2a | 3449.49753 | -0.0091477 | 0.23676914 | -0.0386355 | 0.969181 | 0.99443087 |
| 4930579G24 | 1272.54278 | -0.0378086 | 0.19846857 | -0.1905018 | 0.84891593 | 0.96765585 |
| Slc33a1 | 867.311834 | -0.0645642 | 0.39176205 | -0.1648046 | 0.86909779 | 0.97208313 |
| Gmps | 7013.97391 | -0.1842552 | 0.1798364 | -1.0245712 | 0.30556557 | 0.73530116 |
| Kcnab1 | 9.92327808 | -0.1468993 | 1.44742647 | -0.10149 | 0.91916151 | 0.9839023 |
| Ssr3 | 13537.6288 | 0.38063845 | 0.17818195 | 2.13623462 | 0.03266029 | 0.26868419 |
| Ccnl1 | 3162.16488 | -0.1181344 | 0.316741 | -0.3729684 | 0.70917195 | 0.93012763 |
| Serpini1 | 15.9886088 | -0.9240063 | 2.03128615 | -0.4548873 | 0.64919031 | 0.90973357 |
| Pdcd10 | 3567.07398 | 0.72046904 | 0.42181825 | 1.70800822 | 0.08763482 | 0.45214539 |
| Ptpn22 | 911.242052 | -0.679821 | 0.25410161 | -2.6753902 | 0.00746423 | 0.105928 |
| Dclre1b | 534.204219 | -0.4856362 | 0.37161739 | -1.3068177 | 0.19127463 | 0.62246941 |
| Olfml3 | 326.660589 | -0.1893839 | 0.40348365 | -0.4693719 | 0.63880385 | 0.90615794 |
| Nras | 12824.4748 | 0.23706441 | 0.15350165 | 1.54437699 | 0.12249703 | 0.53213998 |
| Sike1 | 2085.40378 | 0.41273933 | 0.26815767 | 1.53916661 | 0.12376363 | 0.53479352 |
| Tspan2 | 46.6816675 | -0.5539352 | 1.10009409 | -0.5035344 | 0.61458862 | 0.89844745 |
| Vangl1 | 167.720406 | 0.14117909 | 0.66113962 | 0.213539 | 0.83090657 | 0.96466657 |
| Gdap2 | 2193.70222 | -0.1043073 | 0.22173688 | -0.4704101 | 0.63806208 | 0.90599527 |
| Notch2 | 1320.63747 | -0.0117745 | 0.38051297 | -0.0309437 | 0.97531442 | 0.99450266 |
| Sec22b | 5214.1998 | 0.34834811 | 0.26495269 | 1.3147559 | 0.18859195 | 0.61935434 |
| Prpf38b | 2296.6554 | -0.3755297 | 0.25045515 | -1.4993891 | 0.13377272 | 0.55435014 |
| Stxbp3 | 1566.00869 | 0.63669236 | 0.24934686 | 2.55344047 | 0.01066645 | 0.13344172 |
| Gpsm2 | 1015.63406 | -0.0260395 | 0.25210816 | -0.1032872 | 0.91773507 | 0.98390186 |
| Clcc1 | 2802.76753 | -0.3137331 | 0.19645608 | -1.5969631 | 0.11027393 | 0.50751729 |
| Ampd2 | 2621.79906 | -0.1953368 | 0.30545129 | -0.6395024 | 0.52249614 | 0.85816354 |
| Gstm4 | 642.534118 | -0.2331887 | 0.31936913 | -0.7301543 | 0.46529589 | 0.83339036 |
| Ahcyl1 | 433.389902 | -0.3097461 | 0.29692442 | -1.0431817 | 0.29686414 | 0.72954201 |
| Dram2 | 4344.57034 | 0.89295435 | 0.35603262 | 2.50806891 | 0.0121393 | 0.14630546 |
| Dennd2d | 272.122024 | 0.35202823 | 0.47736929 | 0.73743375 | 0.46085861 | 0.82987212 |
| Ddx20 | 2572.31684 | -0.2608198 | 0.15689329 | -1.6624027 | 0.09643206 | 0.47605061 |
| S100a11 | 402.110761 | -1.339431 | 0.33092162 | -4.0475777 | 5.175041624 | 0.00207562 |
| Npr1 | 84.8882715 | 0.87022534 | 0.48843805 | 1.78164935 | 0.07480643 | 0.42093241 |
| Slc27a3 | 99.4706158 | -0.0615002 | 0.53325427 | -0.11533 | 0.90818354 | 0.9817854 |
| Ints3 | 1942.00009 | 0.05379233 | 0.17128253 | 0.31405616 | 0.75347838 | 0.94018119 |
| Rab13 | 138.448494 | 1.31925853 | 0.53705914 | 2.45644927 | 0.01403176 | 0.16117421 |

| | | | | | | |
|---|---|---|---|---|---|---|
| Crtc2 | 708.950476 | -0.3161288 | 0.31295545 | -1.0101398 | 0.3124283 | 0.74051791 |
| Jtb | 2441.76288 | 0.23580352 | 0.29358924 | 0.80317495 | 0.42187362 | 0.80821275 |
| Tpm3 | 32167.283 | -0.0811798 | 0.15385334 | -0.5276443 | 0.59774623 | 0.88928406 |
| 4933434E20I | 7729.62413 | 0.38569155 | 0.28531501 | 1.35180955 | 0.17643625 | 0.60764153 |
| Hax1 | 3402.83905 | 0.23236068 | 0.35738029 | 0.65017767 | 0.51557746 | 0.85589077 |
| Il6ra | 3002.94109 | -0.3104524 | 0.3425212 | -0.9063741 | 0.36473785 | 0.77108653 |
| Chrnb2 | 10.2857035 | -0.2616987 | 1.72256976 | -0.1519234 | 0.87924733 | 0.97541065 |
| Adar | 839.56373 | -0.6577872 | 0.24622886 | -2.6714462 | 0.00755252 | 0.10668585 |
| Pmvk | 1684.9203 | 0.46879884 | 0.24112398 | 1.94422321 | 0.05186854 | 0.35133325 |
| Slc50a1 | 790.937007 | 0.04995418 | 0.32184129 | 0.15521371 | 0.87665284 | 0.9745677 |
| Efna1 | 63.5269918 | 2.34512992 | 1.46701749 | 1.59856984 | 0.10991622 | 0.50665703 |
| Gask1b | 138.484135 | 0.93724932 | 0.68947016 | 1.35937618 | 0.17402742 | 0.60653633 |
| Tmem144 | 1200.00194 | 1.46121783 | 0.32486501 | 4.4979231 | 6.862050709 | 0.00037812 |
| Slc35a3 | 662.749446 | 0.35273747 | 0.45445753 | 0.77617257 | 0.4376471 | 0.8185065 |
| Sass6 | 1156.10453 | 0.12313078 | 0.18804224 | 0.65480384 | 0.51259406 | 0.85441655 |
| Lrrc39 | 13.4560068 | -0.2954076 | 1.66916461 | -0.1769793 | 0.85952462 | 0.96954641 |
| Vcam1 | 30.9439872 | 0.95887348 | 1.86740677 | 0.51347863 | 0.60761655 | 0.89332558 |
| Extl2 | 947.884629 | 0.34718757 | 0.31997844 | 1.08503426 | 0.27790648 | 0.71300965 |
| Larp7 | 2323.68854 | -0.3323676 | 0.21044659 | -1.5793441 | 0.11425715 | 0.51614023 |
| Rnpc3 | 511.392453 | -0.2307238 | 0.26282052 | -0.877876 | 0.380011 | 0.78250296 |
| Hadh | 2421.11208 | 0.29367379 | 0.25574625 | 1.14830149 | 0.25084413 | 0.6842534 |
| Trim2 | 38.4554761 | 0.56474932 | 1.26338238 | 0.44701377 | 0.65486512 | 0.91108261 |
| Mcub | 344.6973 | 0.08443442 | 0.3156614 | 0.26748414 | 0.78909642 | 0.95255415 |
| Tlr2 | 10689.5235 | -0.0538073 | 0.38533028 | -0.1396395 | 0.88894483 | 0.97853524 |
| Sfrp2 | 20.7854329 | 1.06121522 | 1.37988007 | 0.76906337 | 0.44185569 | 0.81971803 |
| Casp6 | 819.526177 | 0.46298171 | 0.27380447 | 1.69092091 | 0.09085191 | 0.46044203 |
| Plrg1 | 3504.5045 | -0.0059174 | 0.15797596 | -0.0374574 | 0.97012032 | 0.99443087 |
| Pla2g12a | 145.063301 | -1.0239861 | 0.4277128 | -2.3940974 | 0.01666132 | 0.1803699 |
| Snx7 | 850.736723 | 0.2912631 | 0.25942005 | 1.12274709 | 0.26154492 | 0.69720616 |
| Gar1 | 642.23718 | -0.5129079 | 0.42781132 | -1.1989115 | 0.23056236 | 0.6653874 |
| Ppa2 | 2130.54694 | 0.05556335 | 0.25040696 | 0.22189218 | 0.82439782 | 0.962942 |
| Ctso | 1240.34359 | 0.94041917 | 0.20835022 | 4.5136461 | 6.372251749 | 0.00035758 |
| Ints12 | 1484.22326 | 0.23561858 | 0.2133197 | 1.10453267 | 0.26936214 | 0.70619525 |
| Egf | 32.0505461 | -0.3910409 | 1.56560194 | -0.2497703 | 0.80276497 | 0.95672466 |
| Gstcd | 325.530149 | -0.3422462 | 0.2541437 | -1.3466642 | 0.1780884 | 0.60928307 |
| Alpk1 | 135.131416 | 0.12867157 | 0.53397023 | 0.24097143 | 0.80957726 | 0.95844313 |
| Aimp1 | 2169.70795 | 0.02178858 | 0.34623311 | 0.06293037 | 0.94982195 | 0.99063619 |
| Tbck | 1894.39061 | 0.08207192 | 0.25725728 | 0.31902664 | 0.74970632 | 0.9399191 |
| Papss1 | 1336.80062 | 0.0468606 | 0.26756363 | 0.17513815 | 0.86097108 | 0.96996131 |
| Fubp1 | 6341.60908 | -0.5245922 | 0.15562591 | -3.370854 | 0.00074936 | 0.01866979 |
| Dnajb4 | 916.327539 | -0.1497046 | 0.45508561 | -0.3289593 | 0.74218648 | 0.93840202 |
| Efna4 | 12.3280511 | 2.69838812 | 2.09597328 | 1.28741533 | 0.19794957 | 0.63076766 |
| Adam15 | 7051.45984 | 0.27542671 | 0.23042271 | 1.1953106 | 0.2319657 | 0.66733333 |
| Zbtb7b | 218.209841 | -0.1535608 | 0.36221528 | -0.4239489 | 0.67160311 | 0.91719387 |
| Cks1b | 8609.85713 | 0.00871347 | 0.30364465 | 0.02869627 | 0.97710683 | 0.99483019 |
| Gba | 4043.69929 | 0.24305866 | 0.17237912 | 1.41002378 | 0.15853266 | 0.58731982 |

| | | | | | | |
|---|---|---|---|---|---|---|
| Scamp3 | 2714.27513 | -0.083365 | 0.23231679 | -0.3588421 | 0.71971325 | 0.93314808 |
| Ash1l | 443.162015 | 0.05720013 | 0.4726831 | 0.12101158 | 0.90368186 | 0.98063872 |
| Rit1 | 1917.03043 | 0.69781434 | 0.3327315 | 2.09722955 | 0.03597326 | 0.28477582 |
| Arhgef2 | 3179.8684 | 0.08033592 | 0.27188021 | 0.29548278 | 0.76762511 | 0.94401697 |
| Khdc4 | 1592.71234 | 0.1073484 | 0.3510144 | 0.30582335 | 0.75973915 | 0.94210735 |
| Lamtor2 | 3271.93951 | 0.0887898 | 0.22365985 | 0.39698589 | 0.69137787 | 0.92487194 |
| Lmna | 1467.11092 | 0.86594475 | 0.39607534 | 2.18631323 | 0.0287927 | 0.25266934 |
| Sema4a | 580.461886 | -0.6943828 | 0.21390314 | -3.2462488 | 0.00116937 | 0.0264415 |
| Pmf1 | 3668.0548 | 0.08198371 | 0.1818301 | 0.45088087 | 0.65207541 | 0.91037383 |
| Iqgap3 | 591.621562 | 0.43205795 | 0.25150932 | 1.71786062 | 0.08582204 | 0.44801003 |
| Gpatch4 | 1387.44376 | -0.3170022 | 0.16495618 | -1.9217361 | 0.05463898 | 0.36033644 |
| Naxe | 3690.84771 | 0.78351689 | 0.37942941 | 2.06498723 | 0.03892421 | 0.29821741 |
| Pear1 | 82.1176236 | 1.01618848 | 0.55278482 | 1.8383075 | 0.06601711 | 0.39562181 |
| Cd1d1 | 648.858997 | -0.6861396 | 0.38050847 | -1.8032177 | 0.07135402 | 0.41122734 |
| Lrba | 38.3451723 | -2.598929 | 1.40660546 | -1.8476602 | 0.06465151 | 0.39234572 |
| Rps3a1 | 34292.6776 | -0.077927 | 0.26584193 | -0.2931328 | 0.76942064 | 0.94401697 |
| Sh3d19 | 49.9893205 | 0.31544145 | 0.99898646 | 0.31576149 | 0.75218355 | 0.94018119 |
| Gatb | 2255.07875 | -0.0626257 | 0.2470375 | -0.2535067 | 0.79987666 | 0.95518266 |
| Fbxw7 | 189.964726 | -0.2188356 | 0.4323254 | -0.5061825 | 0.61272851 | 0.8969137 |
| Fmo5 | 84.9776207 | -0.0220607 | 0.70097171 | -0.0314716 | 0.97489343 | 0.9944856 |
| Chd1l | 1079.64604 | -0.0484855 | 0.28694061 | -0.1689741 | 0.86581705 | 0.9705375 |
| Acp6 | 1190.76947 | -0.0827753 | 0.202337 | -0.4090963 | 0.68246896 | 0.92104707 |
| Gpr89 | 1570.18466 | 0.18985506 | 0.21060003 | 0.90149589 | 0.36732472 | 0.77325076 |
| Rnf115 | 2405.79452 | 0.05542256 | 0.24889338 | 0.22267593 | 0.82378773 | 0.9628698 |
| Polr3c | 2877.59856 | -0.2871419 | 0.16662838 | -1.7232475 | 0.08484377 | 0.44602228 |
| Nudt17 | 17.8150089 | 0.36904837 | 0.95714324 | 0.38557277 | 0.6998131 | 0.92669304 |
| Pias3 | 525.63601 | 0.17617439 | 0.36259472 | 0.48587135 | 0.62705838 | 0.90205481 |
| Pex11b | 1393.637 | -0.1341388 | 0.17064645 | -0.7860629 | 0.43183067 | 0.81344118 |
| Polr3gl | 489.301 | 0.29782962 | 0.22066046 | 1.34971901 | 0.17710613 | 0.60806604 |
| Rprd2 | 515.40554 | -0.0312059 | 0.31322195 | -0.0996287 | 0.92063912 | 0.9839023 |
| Tars2 | 3284.51932 | -0.1922233 | 0.2005489 | -0.9584859 | 0.33781782 | 0.75611716 |
| Ecm1 | 149.451436 | -0.0899659 | 0.42517145 | -0.2115992 | 0.83241972 | 0.96556977 |
| Mettl14 | 636.953066 | 0.00678605 | 0.25094524 | 0.02704197 | 0.97842626 | 0.99483019 |
| Bcar3 | 66.2700249 | 0.57258005 | 0.6366775 | 0.8993251 | 0.36847952 | 0.77450344 |
| Gclm | 674.752435 | 0.3306218 | 0.47263893 | 0.69952301 | 0.48422524 | 0.84128398 |
| Pip5k1a | 926.513353 | -0.127601 | 0.22262462 | -0.5731664 | 0.56653201 | 0.87883863 |
| Abcd3 | 1448.12346 | 0.20983302 | 0.21168669 | 0.99124336 | 0.32156676 | 0.74447686 |
| F3 | 23.7383772 | 1.65540858 | 1.34343744 | 1.23221859 | 0.21786744 | 0.65150232 |
| Rwdd3 | 43.7781789 | 0.4313909 | 0.84633972 | 0.50971364 | 0.61025209 | 0.89508835 |
| Ptbp2 | 153.25411 | 0.05617531 | 0.42286892 | 0.13284333 | 0.89431729 | 0.97941957 |
| Snx27 | 1372.64555 | 0.22582067 | 0.47486339 | 0.4755487 | 0.63439593 | 0.90491667 |
| Adh5 | 6726.34966 | -0.1339099 | 0.17530554 | -0.7638659 | 0.44494714 | 0.82118815 |
| Mrpl9 | 3292.74598 | -0.0110299 | 0.1791764 | -0.0615591 | 0.95091391 | 0.99093022 |
| Them4 | 278.41725 | 0.3231538 | 0.42897971 | 0.75330788 | 0.45126493 | 0.82553071 |
| Rap1gds1 | 4212.76672 | 0.29668419 | 0.2229051 | 1.33098876 | 0.18319271 | 0.61498255 |
| Tspan5 | 436.775887 | 0.08503848 | 0.46277892 | 0.18375616 | 0.85420475 | 0.96844177 |

| | | | | | | |
|---|---|---|---|---|---|---|
| Eif4e | 15852.9373 | 0.04199618 | 0.21547073 | 0.19490435 | 0.84546783 | 0.96765585 |
| Mttp | 141.803977 | 0.52585604 | 0.59124583 | 0.88940338 | 0.37378633 | 0.77748705 |
| Dapp1 | 3401.223 | 0.2093654 | 0.16094408 | 1.30085803 | 0.19330705 | 0.62476227 |
| Ppp3ca | 1250.639 | 0.02359685 | 0.23439908 | 0.10066955 | 0.91981278 | 0.9839023 |
| Nfkb1 | 1604.69027 | -0.3882571 | 0.27605772 | -1.4064345 | 0.15959514 | 0.58893735 |
| Manba | 2833.51089 | 0.11008021 | 0.22466061 | 0.48998447 | 0.62414488 | 0.90037316 |
| Cisd2 | 3971.66903 | 0.34634667 | 0.31572845 | 1.09697643 | 0.27265169 | 0.71005689 |
| Wls | 5000.05758 | 1.00881306 | 0.24852979 | 4.05912335 | 4.925729051 | 0.00199525 |
| Depdc1a | 1085.80797 | 0.0631646 | 0.35141514 | 0.17974353 | 0.85735392 | 0.96940664 |
| Cth | 45.739427 | 0.36221751 | 0.73523281 | 0.49265688 | 0.62225505 | 0.89955637 |
| Zranb2 | 2957.31869 | 0.0783608 | 0.25406527 | 0.30842783 | 0.7577568 | 0.94129085 |
| Adgrl2 | 345.760004 | 0.5964034 | 0.58085052 | 1.02677604 | 0.30452593 | 0.73467909 |
| Rpf1 | 3376.07709 | 0.03744283 | 0.23532039 | 0.15911427 | 0.87357885 | 0.97344599 |
| Spata1 | 24.8847956 | 1.26342769 | 0.99485582 | 1.2699606 | 0.20409866 | 0.63846817 |
| Ctbs | 937.409999 | 0.2670692 | 0.30335876 | 0.88037412 | 0.37865667 | 0.78126394 |
| Bcl10 | 2875.81392 | 0.32702244 | 0.1640787 | 1.99308285 | 0.04625238 | 0.33263099 |
| Ddah1 | 269.763624 | 1.97244178 | 0.52010827 | 3.79236768 | 0.00014922 | 0.0050705 |
| Ccn1 | 2794.91777 | 1.30675243 | 0.37386977 | 3.49520752 | 0.00047369 | 0.01287709 |
| Cryz | 672.133461 | 1.02365003 | 0.45680249 | 2.24090292 | 0.02503236 | 0.23024129 |
| Asph | 2986.92108 | 0.19675471 | 0.2128991 | 0.92416881 | 0.35539843 | 0.7647925 |
| Trp53inp1 | 19.7946012 | -0.4973597 | 1.3703761 | -0.3629366 | 0.71665221 | 0.93235217 |
| Ccne2 | 3445.36797 | 0.50551226 | 0.33994286 | 1.48705067 | 0.13700143 | 0.55976569 |
| Gem | 1811.92449 | 0.77295365 | 0.31883532 | 2.42430368 | 0.01533777 | 0.17119251 |
| Fam92a | 754.782843 | 0.38587617 | 0.32620075 | 1.18294078 | 0.23683262 | 0.6732629 |
| Pip4p2 | 2162.8591 | 0.12266635 | 0.18636951 | 0.65818896 | 0.51041672 | 0.85335831 |
| Decr1 | 1871.73146 | 0.5010583 | 0.29219112 | 1.71483068 | 0.08637627 | 0.44906118 |
| Nbn | 1635.05994 | -0.2522214 | 0.19332026 | -1.3046815 | 0.19200134 | 0.62317653 |
| Cpne3 | 1926.24535 | 0.01047256 | 0.18780713 | 0.0557623 | 0.95553117 | 0.99215259 |
| Rmdn1 | 900.801424 | 1.30934147 | 0.46263745 | 2.83016752 | 0.00465236 | 0.07538062 |
| Tmem68 | 1569.81255 | 0.52878656 | 0.31126084 | 1.69885348 | 0.08934679 | 0.4566019 |
| Tgs1 | 2214.56809 | -0.1652381 | 0.19422584 | -0.8507522 | 0.39490701 | 0.7929904 |
| Rps20 | 10227.1975 | -0.0539118 | 0.31923152 | -0.1688799 | 0.86589113 | 0.9705375 |
| Atp6v0d2 | 10.9093726 | 0.82551602 | 1.79535353 | 0.45980695 | 0.6456548 | 0.90847224 |
| Ubxn2b | 763.738234 | -0.2934438 | 0.24315532 | -1.2068162 | 0.22750292 | 0.66121245 |
| Nsmaf | 892.506538 | -0.2756178 | 0.36665858 | -0.7517016 | 0.45223054 | 0.82603347 |
| Coq3 | 1442.98522 | 0.46930337 | 0.457353 | 1.02612942 | 0.30483058 | 0.7350715 |
| Pnisr | 990.167126 | 0.16912669 | 0.20934012 | 0.80790386 | 0.41914593 | 0.80529433 |
| Sdcbp | 49547.1609 | 0.53955574 | 0.16721066 | 3.22680233 | 0.00125182 | 0.02749282 |
| Tstd3 | 1156.57625 | 0.32659809 | 0.28506775 | 1.14568586 | 0.25192516 | 0.68505999 |
| Ccnc | 1615.4962 | 0.33151937 | 0.22512943 | 1.47257234 | 0.14086639 | 0.56517498 |
| Odf2l | 254.043774 | 0.21512549 | 0.30483938 | 0.7057011 | 0.48037404 | 0.83935746 |
| Ndufaf4 | 1517.54379 | -0.4407423 | 0.30294091 | -1.4548786 | 0.14570287 | 0.57242843 |
| Lmo4 | 834.6487 | -0.5436061 | 0.30728056 | -1.7690872 | 0.07687933 | 0.42792174 |
| Gbp3 | 1272.95169 | -0.3181461 | 0.16542147 | -1.9232457 | 0.0544492 | 0.36033644 |
| Gbp2 | 342.284416 | -0.2021398 | 0.38009909 | -0.5318081 | 0.59485892 | 0.88830085 |
| Gtf2b | 2172.50729 | -0.0402446 | 0.21172233 | -0.1900818 | 0.84924504 | 0.96765585 |

| | | | | | | |
|---|---|---|---|---|---|---|
| Pdlim5 | 854.205499 | 0.67634179 | 0.26263513 | 2.57521449 | 0.01001779 | 0.12928829 |
| Rngtt | 1765.37691 | 0.15784484 | 0.19189315 | 0.82256632 | 0.41075466 | 0.80139357 |
| Ube2j1 | 2224.99063 | 0.32703588 | 0.25232516 | 1.29608906 | 0.19494481 | 0.62699967 |
| Casp8ap2 | 1431.96252 | 0.13512454 | 0.33191538 | 0.40710537 | 0.6839306 | 0.92254085 |
| Map3k7 | 449.874684 | -0.1089464 | 0.31542416 | -0.3453965 | 0.72979629 | 0.93649408 |
| Akirin2 | 769.437245 | -0.0140549 | 0.21316708 | -0.0659338 | 0.94743055 | 0.99019648 |
| Rars2 | 2917.63674 | -0.0195474 | 0.20787471 | -0.0940345 | 0.92508174 | 0.98569821 |
| Slc35a1 | 1099.48112 | 0.09739098 | 0.24747089 | 0.39354521 | 0.69391684 | 0.92609254 |
| Smim8 | 496.672118 | 0.45890089 | 0.50743708 | 0.90435033 | 0.36580967 | 0.77194233 |
| C9orf72 | 2705.55943 | 0.63310149 | 0.2054776 | 3.08112165 | 0.00206222 | 0.04112033 |
| Rnf20 | 1309.23151 | -0.6268409 | 0.2290281 | -2.7369605 | 0.00620097 | 0.09399815 |
| Smc2 | 4311.13326 | -0.1752316 | 0.1946636 | -0.9001764 | 0.36802638 | 0.77410534 |
| Polr1e | 1221.61123 | -0.4122988 | 0.34778024 | -1.1855153 | 0.23581378 | 0.67182855 |
| Exosc3 | 2881.84379 | -0.1903546 | 0.25666529 | -0.7416451 | 0.45830236 | 0.82986925 |
| Tmod1 | 43.04273 | 2.87353785 | 1.21911289 | 2.35707281 | 0.01841963 | 0.19120986 |
| Xpa | 370.346493 | -0.2822554 | 0.32276483 | -0.8744925 | 0.38185009 | 0.78454353 |
| Ncbp1 | 5472.51815 | -0.3858914 | 0.13661873 | -2.8245865 | 0.00473417 | 0.07631593 |
| Trmo | 844.904679 | -0.0442887 | 0.31618631 | -0.1400716 | 0.8886034 | 0.97846539 |
| Anp32b | 13194.7881 | -0.4402542 | 0.22093025 | -1.9927296 | 0.04629106 | 0.33271362 |
| Nans | 4116.89287 | 0.25972825 | 0.25878762 | 1.00363475 | 0.3155547 | 0.74186126 |
| Coro2a | 1010.49001 | -2.3395428 | 0.33562412 | -6.9707229 | 3.153164132 | 8.385360180 |
| Col15a1 | 18.7353227 | 0.80999181 | 1.02547847 | 0.7898672 | 0.42960533 | 0.81155572 |
| Nr4a3 | 21.1678667 | -0.6535639 | 1.25296649 | -0.5216132 | 0.60193966 | 0.89103677 |
| Erp44 | 4499.21014 | 0.05224582 | 0.16410747 | 0.31836346 | 0.75020925 | 0.9399191 |
| Invs | 168.599925 | 0.28695861 | 0.39657152 | 0.72359862 | 0.46931219 | 0.83381643 |
| Tex10 | 3038.15227 | -0.3856467 | 0.23007626 | -1.6761692 | 0.09370506 | 0.46787508 |
| Tmeff1 | 31.7048304 | -0.0888331 | 1.03419395 | -0.085896 | 0.9315491 | 0.98730569 |
| Txn1 | 9613.37645 | 0.11252894 | 0.29442203 | 0.38220284 | 0.70231092 | 0.92762537 |
| Ptgr1 | 1864.15749 | 0.72244187 | 0.32324169 | 2.23498979 | 0.02541802 | 0.23204375 |
| Ugcg | 390.930175 | -0.0679346 | 0.26918978 | -0.2523671 | 0.80075736 | 0.95576786 |
| Ptbp3 | 4930.16154 | 0.05704585 | 0.25793251 | 0.2211658 | 0.82496334 | 0.96312579 |
| Hsdl2 | 1061.44688 | 0.06911104 | 0.20906062 | 0.33057893 | 0.74096256 | 0.93800224 |
| Snx30 | 774.919677 | 0.57706623 | 0.50970609 | 1.13215485 | 0.25756934 | 0.69279809 |
| Zfp37 | 51.9244427 | 3.27379099 | 1.4136448 | 2.31585118 | 0.0205664 | 0.20384393 |
| Wdr31 | 90.7386603 | 0.38600133 | 0.52147305 | 0.74021337 | 0.45917053 | 0.82987212 |
| Alad | 1457.71612 | -0.0267576 | 0.20664141 | -0.1294879 | 0.89697164 | 0.97984623 |
| Pole3 | 3694.67152 | 0.26467228 | 0.23872392 | 1.1086961 | 0.26756131 | 0.70441538 |
| Kdm4c | 618.784206 | 0.44924908 | 0.2390007 | 1.8796978 | 0.06014928 | 0.37831609 |
| Dmac1 | 2639.60811 | 0.59628953 | 0.35265606 | 1.69085291 | 0.0908649 | 0.46044203 |
| Mpdz | 82.2418537 | 0.16994473 | 0.99427465 | 0.17092333 | 0.86428405 | 0.9705375 |
| Zdhhc21 | 651.388731 | -0.4470625 | 0.43088385 | -1.0375475 | 0.29948076 | 0.73113229 |
| Aco1 | 2697.04604 | 0.47159024 | 0.17680998 | 2.66721509 | 0.00764827 | 0.10754172 |
| Smim27 | 136.191172 | -0.5765895 | 0.58543749 | -0.9848865 | 0.32467983 | 0.74601961 |
| Smu1 | 10921.2222 | 0.17662206 | 0.25197331 | 0.70095542 | 0.48333084 | 0.84107543 |
| Dnaja1 | 18571.7316 | -0.1050248 | 0.20900985 | -0.5024874 | 0.6153247 | 0.8987425 |
| Aptx | 968.68551 | 0.22765429 | 0.33644066 | 0.6766551 | 0.49862481 | 0.84870979 |

| | | | | | | |
|---|---|---|---|---|---|---|
| Slc44a1 | 55.3964438 | 0.5616919 | 0.58246856 | 0.96432999 | 0.33488051 | 0.75359284 |
| B4galt1 | 6752.33235 | -0.1648245 | 0.18603157 | -0.8860027 | 0.37561608 | 0.77819605 |
| Fktn | 388.105794 | 0.54441114 | 0.32823409 | 1.65860635 | 0.09719514 | 0.47808129 |
| Bag1 | 1348.78036 | 0.24872432 | 0.4097344 | 0.60703793 | 0.54382574 | 0.86947682 |
| Chmp5 | 2462.36164 | 0.35796951 | 0.18820147 | 1.90205485 | 0.05716398 | 0.36765879 |
| Tmem38b | 401.924349 | -0.0772998 | 0.34203667 | -0.2259986 | 0.82120249 | 0.96205421 |
| Nfx1 | 3258.60094 | -0.2350702 | 0.18610022 | -1.2631377 | 0.2065397 | 0.64143186 |
| Rad23b | 1209.35124 | 0.08977962 | 0.4137108 | 0.21701058 | 0.8282001 | 0.96434151 |
| Nol6 | 1495.23376 | -0.5248855 | 0.27988815 | -1.8753401 | 0.06074595 | 0.37991065 |
| Elp1 | 1718.37848 | -0.2004345 | 0.2453273 | -0.8170086 | 0.41392352 | 0.80395688 |
| Ubap2 | 2570.62111 | -0.3346724 | 0.28911349 | -1.1575815 | 0.24703486 | 0.68185412 |
| Dcaf12 | 3862.84435 | -0.1228987 | 0.23428544 | -0.524568 | 0.59988351 | 0.8902352 |
| Ubap1 | 2292.63872 | -0.1157826 | 0.18924917 | -0.6117997 | 0.54067026 | 0.86741237 |
| Kif24 | 308.073056 | -0.0302649 | 0.31139116 | -0.0971927 | 0.92257337 | 0.98465273 |
| Fam219a | 77.1941508 | 0.84107293 | 0.54687812 | 1.53795314 | 0.12406007 | 0.53550701 |
| Nudt2 | 751.428441 | 0.31859581 | 0.27494019 | 1.15878225 | 0.24654495 | 0.68166056 |
| Enho | 13.7849774 | 1.97769412 | 1.94374776 | 1.01746438 | 0.30893257 | 0.73867239 |
| Dctn3 | 3184.29994 | 0.29906095 | 0.40152491 | 0.74481296 | 0.4563848 | 0.82906968 |
| Vcp | 33782.6723 | -0.1651077 | 0.23984967 | -0.6883798 | 0.49121363 | 0.84491231 |
| Fancg | 893.415154 | 0.38350576 | 0.2804559 | 1.36743694 | 0.17148839 | 0.60430173 |
| Pigo | 349.484063 | -0.3250462 | 0.28191872 | -1.1529784 | 0.24891926 | 0.68345734 |
| Stoml2 | 6567.97165 | -0.0037138 | 0.29118488 | -0.0127542 | 0.98982393 | 0.99650865 |
| Unc13b | 27.1699419 | -0.4727557 | 1.03712618 | -0.4558324 | 0.64851051 | 0.90973357 |
| Tesk1 | 60.4418657 | 0.93923592 | 0.83343151 | 1.12695034 | 0.25976348 | 0.69596146 |
| Cd72 | 76.3439099 | 0.52652066 | 0.58278308 | 0.90345907 | 0.3662823 | 0.77230417 |
| Ccdc107 | 766.379947 | 0.38544036 | 0.42703996 | 0.90258618 | 0.36674557 | 0.7726996 |
| Car9 | 220.594433 | 0.20861087 | 0.42905561 | 0.4862094 | 0.6268187 | 0.90205481 |
| Tpm2 | 3143.96888 | 0.59950725 | 0.23868344 | 2.51172535 | 0.01201426 | 0.1453713 |
| Tln1 | 6946.37875 | -0.1175247 | 0.17957391 | -0.6544644 | 0.51281266 | 0.85459923 |
| Creb3 | 2378.57084 | 0.35372658 | 0.17892098 | 1.97699893 | 0.04804175 | 0.33778401 |
| Gba2 | 793.220973 | -0.0528001 | 0.22491314 | -0.2347578 | 0.81439676 | 0.95988497 |
| Rgp1 | 564.252038 | -0.7297782 | 0.37135755 | -1.9651632 | 0.04939535 | 0.34139551 |
| Hint2 | 676.100856 | 0.52176667 | 0.35381206 | 1.47470009 | 0.14029319 | 0.56406756 |
| Reck | 16.4758715 | 1.24050102 | 2.00384635 | 0.61905995 | 0.53587687 | 0.86539693 |
| Clta | 21534.4149 | 0.6977779 | 0.23266772 | 2.99903185 | 0.00270839 | 0.04974735 |
| Gne | 885.718358 | -0.4706252 | 0.38836822 | -1.2118014 | 0.2255884 | 0.65900564 |
| Glipr2 | 2070.20043 | -0.8557366 | 0.30095101 | -2.8434415 | 0.00446292 | 0.07328174 |
| Snapc3 | 729.229261 | -0.2590564 | 0.29363327 | -0.8822448 | 0.37764443 | 0.78033065 |
| Psip1 | 528.340463 | 0.05906451 | 0.26256935 | 0.22494821 | 0.82201956 | 0.96235404 |
| Plin2 | 26574.5141 | 0.5051176 | 0.24052379 | 2.10007333 | 0.03572239 | 0.28378802 |
| Rps6 | 44330.7542 | -0.0494361 | 0.29367833 | -0.1683342 | 0.86632039 | 0.97060021 |
| Mllt3 | 76.2879121 | 0.92183887 | 0.85378298 | 1.07971099 | 0.2802709 | 0.71532524 |
| Hacd4 | 3665.38018 | 1.00415935 | 0.30159375 | 3.32950983 | 0.00086999 | 0.02090881 |
| Usp24 | 572.381602 | 0.28762698 | 0.33731511 | 0.85269523 | 0.39382834 | 0.79233901 |
| Plpp3 | 31.3388923 | 1.4786669 | 1.193722 | 1.23870289 | 0.21545555 | 0.64854028 |
| Slc35d1 | 494.503222 | -1.1034679 | 0.45994937 | -2.3991073 | 0.0164351 | 0.1787116 |

| | | | | | | |
|---|---|---|---|---|---|---|
| Mier1 | 1194.75109 | 0.13514652 | 0.21549555 | 0.62714298 | 0.53056552 | 0.86356925 |
| Sgip1 | 149.598796 | 0.09962655 | 0.35929122 | 0.27728635 | 0.78156024 | 0.95004728 |
| Pde4b | 1840.0862 | -0.3203329 | 0.54197104 | -0.5910518 | 0.55448574 | 0.87165957 |
| Ak4 | 520.147662 | 0.04317803 | 0.35205947 | 0.12264413 | 0.90238891 | 0.98043702 |
| Jak1 | 5848.97004 | 0.06179003 | 0.21043343 | 0.29363222 | 0.76903896 | 0.94401697 |
| Cachd1 | 20.4130841 | -1.4940547 | 1.62853939 | -0.9174201 | 0.35892256 | 0.76764331 |
| St3gal3 | 64.5901247 | 1.37916354 | 0.70378498 | 1.95963765 | 0.05003816 | 0.34252555 |
| Dph2 | 1017.74634 | -0.144195 | 0.26731 | -0.5394299 | 0.58959023 | 0.88722166 |
| B4galt2 | 22.2377586 | -0.1555673 | 1.33930896 | -0.1161549 | 0.90752977 | 0.98176897 |
| Bend5 | 7.39977853 | -0.0955052 | 2.23512957 | -0.0427292 | 0.96591744 | 0.99443087 |
| Elavl4 | 235.073164 | 0.30521023 | 0.36241955 | 0.84214616 | 0.39970614 | 0.79612475 |
| Itgb3bp | 482.19637 | 0.60172783 | 0.50363051 | 1.19478034 | 0.23217286 | 0.66733333 |
| Atg4c | 419.433613 | 0.21948401 | 0.32485531 | 0.67563621 | 0.49927165 | 0.84872332 |
| Cdkn2c | 194.382869 | 1.47630922 | 0.71700273 | 2.05900084 | 0.03949416 | 0.30082941 |
| Eps15 | 2610.94571 | 0.56500959 | 0.23665034 | 2.38752918 | 0.01696206 | 0.18249502 |
| Ttc39a | 76.1561728 | 0.93607728 | 0.51279087 | 1.8254562 | 0.06793222 | 0.40145642 |
| Dock7 | 309.107983 | -0.2450078 | 0.43031105 | -0.5693736 | 0.56910262 | 0.87961466 |
| Rnf11 | 79.8917553 | 0.25594089 | 0.51720646 | 0.49485246 | 0.62070427 | 0.89896588 |
| Osbpl9 | 5399.22925 | 0.89617702 | 0.20882068 | 4.29161042 | 1.773819581 | 0.00083458 |
| Usp1 | 1712.47807 | 0.28203677 | 0.39065495 | 0.7219588 | 0.47031981 | 0.83381643 |
| Tm2d1 | 1011.89272 | 0.58681215 | 0.36614253 | 1.60268776 | 0.10900361 | 0.50470898 |
| Nfia | 50.6863747 | -0.6884357 | 1.02307037 | -0.6729114 | 0.50100365 | 0.84968141 |
| Txndc12 | 1485.66176 | 0.10542761 | 0.26523102 | 0.39749354 | 0.69100355 | 0.92470267 |
| Btf3l4 | 2691.93937 | 0.3204647 | 0.32490847 | 0.986323 | 0.32397463 | 0.74583177 |
| Fggy | 349.633407 | 0.57050416 | 0.3875859 | 1.4719425 | 0.14103641 | 0.5654142 |
| Ift74 | 275.352196 | 0.47814293 | 0.38566559 | 1.23978635 | 0.21505443 | 0.64854028 |
| Plaa | 3410.97714 | -0.2965232 | 0.2153386 | -1.3770091 | 0.16850943 | 0.60312169 |
| Caap1 | 97.7061774 | -0.4009946 | 0.42874945 | -0.9352656 | 0.34965144 | 0.76203275 |
| Pum1 | 2492.62333 | -0.2603591 | 0.31866517 | -0.8170304 | 0.41391107 | 0.80395688 |
| Laptm5 | 41731.9141 | -0.3447408 | 0.23757668 | -1.4510716 | 0.14675992 | 0.57365079 |
| Cc2d1b | 1234.04995 | 0.43159003 | 0.22585214 | 1.91094066 | 0.05601221 | 0.3642729 |
| Pdpn | 18.3668183 | 0.13495405 | 1.04795873 | 0.12877802 | 0.8975333 | 0.97984623 |
| Orc1 | 1419.79782 | 0.34468868 | 0.3887577 | 0.88664142 | 0.37527199 | 0.77815049 |
| Gpx7 | 29.0371716 | 1.39495236 | 1.00295163 | 1.39084709 | 0.1642718 | 0.59585062 |
| Tnfrsf1b | 6256.23038 | -0.0645114 | 0.30067952 | -0.2145521 | 0.83011651 | 0.96464475 |
| Echdc2 | 39.4037362 | 2.36023838 | 1.04145365 | 2.2662923 | 0.02343349 | 0.22033967 |
| Tnfrsf8 | 34.5623883 | -0.7851142 | 0.67380973 | -1.1651867 | 0.24394341 | 0.67911981 |
| Scp2 | 4491.94578 | 0.24044015 | 0.16492675 | 1.45786022 | 0.14487908 | 0.57067981 |
| Cpt2 | 1354.45204 | -0.0713223 | 0.28187829 | -0.2530251 | 0.80024881 | 0.95544053 |
| Czib | 1616.35633 | -0.0963526 | 0.2244014 | -0.4293759 | 0.66764966 | 0.91595755 |
| Magoh | 2596.85833 | 0.02650519 | 0.36516699 | 0.07258375 | 0.94213735 | 0.98962444 |
| Lrp8 | 34.9477467 | -0.7154323 | 1.14662505 | -0.6239462 | 0.53266294 | 0.8646427 |
| Ndc1 | 5290.33953 | -0.1671042 | 0.18318346 | -0.9122231 | 0.3616513 | 0.76887042 |
| Lrrc42 | 383.356495 | 0.60003127 | 0.43481648 | 1.37996443 | 0.1675976 | 0.60194404 |
| Tmem59 | 10145.8243 | 0.29986906 | 0.22966361 | 1.30568818 | 0.19165862 | 0.62272506 |
| Tceanc2 | 974.168928 | 0.43140422 | 0.265126 | 1.6271668 | 0.10370168 | 0.49284483 |

| | | | | | | |
|---|---|---|---|---|---|---|
| Cyb5rl | 283.913286 | -0.6472591 | 0.54617018 | -1.1850869 | 0.23598308 | 0.67182855 |
| Mrpl37 | 4111.12937 | -0.0010046 | 0.18572888 | -0.0054088 | 0.99568442 | 0.99884675 |
| Exo5 | 599.089261 | 0.19637147 | 0.22345718 | 0.87878794 | 0.37951627 | 0.78211296 |
| Dyrk2 | 304.133632 | -0.1565912 | 0.51081679 | -0.3065507 | 0.75918539 | 0.94184564 |
| Ctps | 3539.30345 | -0.4957684 | 0.18552352 | -2.6722671 | 0.00753406 | 0.10658529 |
| Hivep3 | 316.633826 | 0.40120006 | 0.40914056 | 0.98059226 | 0.32679385 | 0.74811039 |
| Ppcs | 687.349752 | 0.75780784 | 0.46383386 | 1.63379156 | 0.10230262 | 0.4886081 |
| Ybx1 | 19223.0233 | 0.25760126 | 0.46024041 | 0.55971021 | 0.57567711 | 0.88080372 |
| P3h1 | 900.851171 | 0.19839025 | 0.24521756 | 0.80903773 | 0.41849345 | 0.8052381 |
| Svbp | 761.228542 | -0.1419936 | 0.25925666 | -0.5476951 | 0.58390126 | 0.88404713 |
| Slc2a1 | 9585.65177 | -0.4350152 | 0.18501896 | -2.3511927 | 0.01871334 | 0.1930188 |
| Rragc | 623.061831 | 0.02290472 | 0.30254734 | 0.07570624 | 0.93965281 | 0.98913801 |
| Mycbp | 1755.15208 | 0.19626212 | 0.34680808 | 0.56590989 | 0.57145504 | 0.88035336 |
| Ndufs5 | 3077.98128 | 0.14554524 | 0.30508928 | 0.47705787 | 0.63332091 | 0.90434445 |
| Macf1 | 669.166103 | -0.074663 | 0.32418592 | -0.2303093 | 0.81785144 | 0.96079676 |
| Ppie | 2619.58011 | -0.3123196 | 0.2082155 | -1.4999827 | 0.13361889 | 0.55425547 |
| Trit1 | 781.418371 | -0.1065961 | 0.26538272 | -0.4016693 | 0.68792742 | 0.92470267 |
| Mycl | 89.1665135 | -1.1702687 | 0.51102325 | -2.2900498 | 0.02201843 | 0.21347378 |
| Mfsd2a | 15.4227249 | -0.3869111 | 2.08598826 | -0.1854809 | 0.85285183 | 0.96798445 |
| Cap1 | 2220.16439 | -0.2927673 | 0.21141824 | -1.384778 | 0.16612037 | 0.59942225 |
| Ppt1 | 6131.66803 | 0.43968124 | 0.20127774 | 2.18445041 | 0.02892917 | 0.25332176 |
| Eloa | 5512.18214 | 0.0781269 | 0.16499175 | 0.47352007 | 0.63584219 | 0.90531997 |
| Pithd1 | 2491.89163 | 0.06639929 | 0.21468284 | 0.30929014 | 0.75710083 | 0.94112534 |
| Lypla2 | 1631.36898 | -0.3134945 | 0.24210056 | -1.2948939 | 0.19535683 | 0.62707955 |
| Gale | 1434.90325 | 0.17456792 | 0.24250879 | 0.7198416 | 0.47162253 | 0.83396334 |
| Hmgcl | 4582.78316 | 0.46921256 | 0.29481554 | 1.59154623 | 0.1114867 | 0.50888686 |
| Fuca1 | 3924.45653 | 0.33600408 | 0.24496459 | 1.37164343 | 0.17017447 | 0.60430173 |
| Pnrc2 | 5306.85776 | 0.1189035 | 0.27335773 | 0.43497397 | 0.66358132 | 0.91395288 |
| Srsf10 | 6771.52227 | -0.175365 | 0.15960258 | -1.0987601 | 0.2718727 | 0.70940613 |
| Rnf220 | 637.122329 | -0.0546878 | 0.31065343 | -0.1760412 | 0.86026156 | 0.96973642 |
| Kif2c | 3461.79841 | 0.05691077 | 0.19412543 | 0.29316494 | 0.76939608 | 0.94401697 |
| Plk3 | 492.78281 | 0.00208694 | 0.5851148 | 0.00356671 | 0.99715418 | 0.9994418 |
| Eif2b3 | 2070.30966 | -0.4352669 | 0.20176423 | -2.1573045 | 0.03098195 | 0.26283091 |
| Urod | 1979.65514 | 0.29635722 | 0.32381784 | 0.91519731 | 0.36008806 | 0.76857232 |
| Mutyh | 445.054569 | 0.14636648 | 0.31763679 | 0.46079827 | 0.64494334 | 0.90835328 |
| Toe1 | 1206.42831 | -0.0716831 | 0.24439883 | -0.2933037 | 0.76928999 | 0.94401697 |
| Ccdc163 | 414.634401 | -0.0972516 | 0.29275482 | -0.3321948 | 0.73974215 | 0.93800224 |
| Mmachc | 1152.58669 | -0.3175998 | 0.25051424 | -1.2677913 | 0.20487249 | 0.63901202 |
| Prdx1 | 44887.3587 | 0.53180755 | 0.32718039 | 1.6254261 | 0.1040718 | 0.49391412 |
| Akr1a1 | 32746.4738 | 0.38562338 | 0.20318035 | 1.89793642 | 0.05770446 | 0.36898259 |
| Nasp | 5454.10082 | -0.3013136 | 0.14952776 | -2.0151013 | 0.04389403 | 0.32095378 |
| Ipp | 618.821151 | -0.2189199 | 0.25341529 | -0.8638779 | 0.38765494 | 0.7893114 |
| Pik3r3 | 28.6260283 | -0.6742748 | 0.93927228 | -0.7178694 | 0.47283782 | 0.83483892 |
| Pomgnt1 | 282.389911 | 0.40052764 | 0.44200008 | 0.90617097 | 0.36484536 | 0.77108653 |
| Rad54l | 2677.86642 | -0.2381027 | 0.17187265 | -1.3853437 | 0.16594739 | 0.5992759 |
| Lrrc41 | 4068.60154 | 0.1120912 | 0.17175932 | 0.6526062 | 0.51401019 | 0.85502946 |

| | | | | | | |
|---|---|---|---|---|---|---|
| Nsun4 | 1089.49905 | 0.10417262 | 0.1912304 | 0.54474926 | 0.58592598 | 0.88513468 |
| Mknk1 | 3680.19844 | -0.0848305 | 0.21575339 | -0.3931829 | 0.69418443 | 0.92609254 |
| Mob3c | 1791.31912 | 0.31416739 | 0.36554346 | 0.8594529 | 0.39009069 | 0.78985201 |
| Atpaf1 | 2067.58511 | 0.5861229 | 0.29738652 | 1.97091281 | 0.04873385 | 0.34046898 |
| Tal1 | 343.965178 | 0.22613243 | 0.36928523 | 0.61235167 | 0.54030511 | 0.86741192 |
| Stil | 498.247695 | 0.28765887 | 0.43991694 | 0.65389361 | 0.51318035 | 0.85492786 |
| Cmpk1 | 9459.62509 | 0.0865406 | 0.25098479 | 0.34480416 | 0.7302416 | 0.93653187 |
| Ebna1bp2 | 3508.18273 | -0.3934209 | 0.20244356 | -1.9433609 | 0.05197256 | 0.35164843 |
| Aldh4a1 | 864.061079 | -0.4119219 | 0.29681539 | -1.387805 | 0.16519644 | 0.59797229 |
| Mrto4 | 3677.65439 | -0.2176543 | 0.31166326 | -0.6983636 | 0.48494981 | 0.84171939 |
| Akr7a5 | 1404.48156 | 0.20661876 | 0.2096576 | 0.98550572 | 0.32437572 | 0.74596967 |
| Pqlc2 | 2117.41176 | 0.92915744 | 0.2188963 | 4.24473791 | 2.188490598 | 0.00097352 |
| Capzb | 26257.3763 | 0.0520128 | 0.14637018 | 0.35535106 | 0.72232661 | 0.93423486 |
| Pink1 | 403.516224 | 0.58899707 | 0.59172919 | 0.99538281 | 0.31955012 | 0.7440831 |
| Ddost | 20071.9899 | 0.08135888 | 0.24486968 | 0.33225381 | 0.73969762 | 0.93800224 |
| Hp1bp3 | 2647.95843 | -0.1896564 | 0.25285458 | -0.750061 | 0.45321796 | 0.82610332 |
| Eif4g3 | 398.212598 | -0.5133838 | 0.56401099 | -0.9102372 | 0.36269741 | 0.76990376 |
| Hspg2 | 14.779762 | 2.71137547 | 2.014146 | 1.3461663 | 0.17824889 | 0.60928307 |
| Ptpn12 | 267.244968 | 0.45987457 | 0.43257501 | 1.06310941 | 0.28773234 | 0.72300867 |
| Zcchc17 | 2407.61053 | -0.1053196 | 0.20699827 | -0.5087948 | 0.61089606 | 0.89526044 |
| Fabp3 | 72.9643592 | 0.88474117 | 0.76953424 | 1.14970995 | 0.25026336 | 0.6842534 |
| Tinagl1 | 8.46357704 | 0.85930702 | 1.49678862 | 0.57410045 | 0.56589983 | 0.878492 |
| Pef1 | 2015.87271 | 0.08976226 | 0.23261244 | 0.38588762 | 0.6995799 | 0.92669304 |
| Ptp4a2 | 22158.4017 | -0.132885 | 0.15168034 | -0.8760858 | 0.38098342 | 0.7832891 |
| Khdrbs1 | 4661.89443 | -0.1927641 | 0.2983186 | -0.6461685 | 0.51817021 | 0.8570838 |
| Ak2 | 7778.72899 | 0.00754875 | 0.15261371 | 0.04946311 | 0.96055023 | 0.99325523 |
| Rnf19b | 83.7470672 | 0.70624508 | 0.59842725 | 1.18016866 | 0.23793314 | 0.67403588 |
| Ccdc28b | 1090.99758 | 0.08652638 | 0.2787278 | 0.31043325 | 0.75623151 | 0.94102069 |
| Phc2 | 291.446741 | 0.23354637 | 0.31295145 | 0.7462703 | 0.45550415 | 0.8279617 |
| Tmem234 | 5068.34633 | 0.24878862 | 0.24519203 | 1.01466845 | 0.3102639 | 0.7399039 |
| Eif3i | 8834.11749 | 0.08327363 | 0.35489515 | 0.23464292 | 0.8144859 | 0.95988497 |
| Zfp362 | 122.389234 | 0.41509241 | 0.57965845 | 0.71609826 | 0.47393068 | 0.83526783 |
| Hdac1 | 8772.25509 | -0.1600016 | 0.21212343 | -0.7542851 | 0.45067802 | 0.825321 |
| Nipal3 | 58.8385882 | 1.46866689 | 0.55226173 | 2.65936747 | 0.00782875 | 0.10907646 |
| Zbtb8a | 40.6233083 | 0.00974202 | 0.86662985 | 0.01124128 | 0.99103095 | 0.99681644 |
| Srrm1 | 2691.63388 | -0.4031115 | 0.26209249 | -1.5380505 | 0.12403628 | 0.53550701 |
| Yars | 6283.5116 | -0.4464066 | 0.17408529 | -2.5642981 | 0.01033847 | 0.1312161 |
| Sfpq | 2374.64731 | -0.2423301 | 0.25088088 | -0.9659169 | 0.33408578 | 0.75320272 |
| Syf2 | 1927.96273 | -0.2551646 | 0.21344931 | -1.1954341 | 0.23191748 | 0.66733333 |
| Tmem50a | 7096.85467 | 0.4444541 | 0.3034448 | 1.46469509 | 0.14300412 | 0.56846902 |
| Maco1 | 237.332369 | 0.14968528 | 0.33394169 | 0.44823778 | 0.6539816 | 0.91076467 |
| AU040320 | 726.686285 | -0.0435699 | 0.31765424 | -0.1371615 | 0.89090316 | 0.97889088 |
| Stmn1 | 12811.6241 | 0.33981454 | 0.27848345 | 1.22023243 | 0.22237678 | 0.65586668 |
| Ncdn | 953.64533 | -0.5197057 | 0.25692161 | -2.022818 | 0.04309191 | 0.31890102 |
| Psmb2 | 14793.2074 | 0.06250942 | 0.23764412 | 0.26303792 | 0.79252135 | 0.95297175 |
| Zfp593 | 1125.71197 | -0.3839312 | 0.2709955 | -1.4167439 | 0.15655781 | 0.58588334 |

| | | | | | | |
|---|---|---|---|---|---|---|
| Ago3 | 207.352511 | 0.79461167 | 0.34161552 | 2.32604086 | 0.02001637 | 0.20003355 |
| Sh3bgrl3 | 15719.5679 | -0.3298386 | 0.14156272 | -2.329982 | 0.0198071 | 0.19950359 |
| Trappc3 | 2883.42062 | 0.47950701 | 0.26913554 | 1.78165618 | 0.07480532 | 0.42093241 |
| Gpn2 | 553.804689 | -0.5913916 | 0.22945793 | -2.5773422 | 0.00995633 | 0.129021 |
| Map7d1 | 1983.44244 | 0.28462656 | 0.34732187 | 0.81948929 | 0.41250731 | 0.80251302 |
| Gpatch3 | 847.23619 | -0.0821835 | 0.31304385 | -0.2625304 | 0.79291256 | 0.95303665 |
| Nudc | 10712.4712 | -0.4271248 | 0.19952569 | -2.1407009 | 0.03229817 | 0.26700278 |
| Slc9a1 | 330.541226 | -0.3440567 | 0.46663295 | -0.7373175 | 0.46092927 | 0.82987212 |
| Tmem222 | 2475.44707 | 0.1545174 | 0.20526431 | 0.75277282 | 0.45158644 | 0.82570905 |
| Csf3r | 3316.46198 | -1.3456958 | 0.21787577 | -6.1764361 | 6.556474805 | 1.215232671 |
| Mrps15 | 2551.63938 | 0.03518479 | 0.15353407 | 0.22916603 | 0.81873987 | 0.96105733 |
| Map3k6 | 55.7608004 | -1.3756884 | 1.34247787 | -1.0247383 | 0.30548671 | 0.73530116 |
| Meaf6 | 870.852504 | -0.189633 | 0.20234027 | -0.9371984 | 0.34865652 | 0.76162771 |
| Hgf | 112.017773 | -0.4226771 | 0.71904877 | -0.5878282 | 0.55664761 | 0.87276537 |
| Wasf2 | 3397.61803 | 0.237686 | 0.31803078 | 0.74736791 | 0.45484151 | 0.82761805 |
| Gnl2 | 3604.53525 | -0.2246726 | 0.16362625 | -1.3730838 | 0.1697263 | 0.6042671 |
| Rspo1 | 29.1829336 | -0.1811302 | 1.06910274 | -0.1694226 | 0.86546425 | 0.9705375 |
| Cdca8 | 7889.2151 | -0.2430929 | 0.17263837 | -1.4081048 | 0.15910004 | 0.58838861 |
| Fgr | 141.628147 | -2.7651233 | 0.50519441 | -5.4733846 | 4.415204805 | 4.822428606 |
| Fam76a | 1292.61183 | 0.24256635 | 0.22418407 | 1.08199636 | 0.27925414 | 0.71377266 |
| Stx12 | 2945.2161 | 0.30026606 | 0.21042973 | 1.42691841 | 0.15360341 | 0.58242478 |
| Ppp1r8 | 1403.96952 | -0.2336274 | 0.31582435 | -0.7397385 | 0.45945867 | 0.82987212 |
| Sema3a | 20.8199579 | -1.2199129 | 2.12131912 | -0.5750728 | 0.56524207 | 0.87804524 |
| Rpa2 | 4999.57998 | -0.1326956 | 0.21138746 | -0.6277364 | 0.53017662 | 0.86348696 |
| Smpdl3b | 262.798319 | 0.86815434 | 0.44127495 | 1.96737736 | 0.04913972 | 0.34139551 |
| Eya3 | 1653.15088 | 0.11511031 | 0.20539555 | 0.56043234 | 0.57518458 | 0.88059206 |
| Yrdc | 501.767807 | 0.76823574 | 0.4907877 | 1.56531171 | 0.11750984 | 0.52372617 |
| Mtf1 | 531.795338 | -0.0386795 | 0.38213635 | -0.1012192 | 0.91937644 | 0.9839023 |
| Sesn2 | 922.877974 | -0.0744853 | 0.26442506 | -0.2816876 | 0.77818306 | 0.94837927 |
| Inpp5b | 2225.81331 | -0.6404224 | 0.22520436 | -2.8437388 | 0.00445876 | 0.07328174 |
| Rcc1 | 5181.72619 | -0.5688659 | 0.17155834 | -3.3158742 | 0.00091357 | 0.02178498 |
| Trnau1ap | 2238.15105 | 0.2668356 | 0.33497046 | 0.79659443 | 0.42568662 | 0.81021713 |
| Taf12 | 2530.50534 | 0.07757582 | 0.23512481 | 0.32993464 | 0.74144935 | 0.93800224 |
| Gmeb1 | 848.482149 | -0.1858097 | 0.30721507 | -0.6048198 | 0.54529877 | 0.86970532 |
| Sf3a3 | 7158.17543 | -0.2005329 | 0.14942134 | -1.3420631 | 0.17957552 | 0.60992118 |
| Epb41 | 1712.65087 | 0.25646065 | 0.33953233 | 0.75533498 | 0.45004801 | 0.82484234 |
| Utp11 | 2836.42466 | 0.30213033 | 0.28763252 | 1.05040392 | 0.29353244 | 0.72646828 |
| Mecr | 1634.85812 | -0.1868503 | 0.28264226 | -0.6610841 | 0.50855839 | 0.85251363 |
| Srsf4 | 2326.35903 | -0.3474488 | 0.24407099 | -1.4235562 | 0.15457497 | 0.58343244 |
| Casp9 | 1652.65308 | 0.30662158 | 0.21939584 | 1.39757245 | 0.1622415 | 0.59320305 |
| Plekhm2 | 136.736986 | 0.30061165 | 0.48106961 | 0.6248818 | 0.53204864 | 0.8643494 |
| Fbxo42 | 1460.04 | -0.2338443 | 0.28890091 | -0.8094273 | 0.4182694 | 0.8052381 |
| Necap2 | 6869.7739 | 0.3986204 | 0.14825865 | 2.68868215 | 0.00717347 | 0.10311756 |
| Cdk14 | 381.486625 | 1.62447524 | 0.28473989 | 5.70511995 | 1.162611833 | 1.580247839 |
| Padi2 | 37.2711049 | -3.4394982 | 1.32227526 | -2.6011968 | 0.00928991 | 0.12352557 |
| Kcnab2 | 888.248745 | 0.14356042 | 0.21028467 | 0.68269562 | 0.4947992 | 0.84678075 |

| | | | | | | |
|---|---|---|---|---|---|---|
| Psmc2 | 10552.5753 | -0.0591304 | 0.19469306 | -0.3037109 | 0.76134816 | 0.94247845 |
| Xrcc2 | 690.11788 | 0.44503374 | 0.38951989 | 1.14251865 | 0.2532385 | 0.68647781 |
| Rpl22 | 4100.41359 | 0.21993251 | 0.37215924 | 0.59096346 | 0.5545449 | 0.87165957 |
| Acot7 | 1830.31573 | -1.1941041 | 0.2772192 | -4.3074366 | 1.651574357 | 0.00078921 |
| Prkag2 | 208.776676 | -1.007048 | 0.39010945 | -2.5814498 | 0.00983863 | 0.12803827 |
| Rheb | 5049.0323 | 0.09831896 | 0.20822584 | 0.47217464 | 0.63680214 | 0.90549815 |
| Nol9 | 3112.77355 | -0.4738386 | 0.17031886 | -2.7820678 | 0.00540138 | 0.08471157 |
| Zbtb48 | 250.047026 | 0.41347308 | 0.50359188 | 0.82104795 | 0.41161896 | 0.80231592 |
| Abcf2 | 6871.58876 | -0.177852 | 0.17142033 | -1.0375202 | 0.29949349 | 0.73113229 |
| Nub1 | 1923.35363 | 0.07784708 | 0.25286714 | 0.30785763 | 0.75819067 | 0.94131116 |
| Vamp3 | 12299.4334 | 0.87219457 | 0.20474483 | 4.25991019 | 2.045090622 | 0.00093002 |
| Per3 | 322.183314 | 1.01791777 | 0.39203326 | 2.59650868 | 0.00941765 | 0.12454718 |
| Tmub1 | 358.106395 | -0.5065339 | 0.3052966 | -1.6591533 | 0.09708491 | 0.47799662 |
| Fastk | 1236.0113 | -0.1961583 | 0.23565073 | -0.8324111 | 0.40517695 | 0.79931134 |
| Ube4b | 1024.90716 | -0.019338 | 0.28187392 | -0.0686051 | 0.94530398 | 0.99019648 |
| Pgd | 13044.5667 | -0.3051889 | 0.18169005 | -1.6797228 | 0.09301126 | 0.46652562 |
| Slc4a2 | 595.934037 | 0.20056095 | 0.4339694 | 0.4621546 | 0.64397046 | 0.90823885 |
| Park7 | 10711.5321 | -0.0254417 | 0.23195671 | -0.1096832 | 0.91266066 | 0.98292561 |
| Errfi1 | 2378.18965 | 0.18481504 | 0.38301021 | 0.48253295 | 0.6294274 | 0.90309469 |
| Cdk5 | 595.201707 | 0.16180635 | 0.24584022 | 0.65817687 | 0.51042449 | 0.85335831 |
| Abcb1b | 373.890793 | 0.20655207 | 0.41045487 | 0.50322725 | 0.61480451 | 0.89844745 |
| Car6 | 27.6976527 | 0.55962191 | 1.05570752 | 0.53009181 | 0.59604828 | 0.8886604 |
| Abcb8 | 2484.28863 | -0.3358335 | 0.23681065 | -1.4181519 | 0.15614642 | 0.58521421 |
| Dffa | 1331.57553 | 0.08261173 | 0.26821328 | 0.30800761 | 0.75807654 | 0.94129085 |
| Pex14 | 1574.26021 | -0.1582849 | 0.21359095 | -0.7410656 | 0.45865367 | 0.82986925 |
| Slc2a5 | 816.93557 | 0.41489227 | 0.21897287 | 1.89471995 | 0.05812951 | 0.37027169 |
| Casz1 | 20.6585459 | -1.257079 | 1.35177437 | -0.9299474 | 0.35239834 | 0.76358405 |
| Nos3 | 22.8862461 | 0.89101839 | 1.09581041 | 0.81311363 | 0.41615291 | 0.80440685 |
| H6pd | 1179.86774 | 0.27362146 | 0.28598568 | 0.9567663 | 0.33868523 | 0.7564609 |
| Slc25a33 | 413.62944 | 0.49090875 | 0.60615844 | 0.80986871 | 0.41801564 | 0.8051099 |
| Klhl7 | 1018.89478 | 0.31365857 | 0.23801269 | 1.31782287 | 0.18756295 | 0.61935434 |
| Ctnnbip1 | 459.601438 | 0.58366495 | 0.41454734 | 1.40795728 | 0.15914372 | 0.58838861 |
| Lzic | 1781.80102 | 0.027106 | 0.19958733 | 0.1358102 | 0.89197132 | 0.97931118 |
| Mtor | 591.920647 | 0.0374892 | 0.43984944 | 0.08523188 | 0.93207704 | 0.98730569 |
| Nmnat1 | 380.262518 | 0.62006002 | 0.32849432 | 1.88758219 | 0.05908206 | 0.37462651 |
| Fam126a | 481.865878 | 0.8836588 | 0.25687521 | 3.44003137 | 0.00058165 | 0.0152036 |
| Tomm7 | 3232.31643 | 0.23639091 | 0.40276002 | 0.58692743 | 0.55725245 | 0.87276537 |
| Rint1 | 658.331628 | 0.18363888 | 0.22405758 | 0.81960573 | 0.41244091 | 0.80251302 |
| Mad2l2 | 851.225131 | -0.1045372 | 0.20486448 | -0.5102748 | 0.60985898 | 0.89508835 |
| Kmt2e | 1489.85249 | 0.45129107 | 0.21926375 | 2.058211 | 0.03956988 | 0.30121865 |
| Agtrap | 1580.67625 | 0.53194533 | 0.29509303 | 1.802636 | 0.0714454 | 0.41122734 |
| Mthfr | 661.765326 | 0.22251066 | 0.28374289 | 0.78419818 | 0.43292383 | 0.81435873 |
| Orc5 | 2163.75624 | 0.02685431 | 0.17121604 | 0.15684461 | 0.87536731 | 0.97384215 |
| Dnajc2 | 3043.21343 | -0.5372098 | 0.19019458 | -2.8245271 | 0.00473504 | 0.07631593 |
| Clcn6 | 91.2243738 | -1.0467602 | 1.15842762 | -0.9036043 | 0.36620526 | 0.77230417 |
| Pmpcb | 3508.75171 | 0.02391503 | 0.23390757 | 0.10224139 | 0.91856508 | 0.98390186 |

| | | | | | | |
|---|---|---|---|---|---|---|
| Mfn2 | 2249.22142 | -0.1606839 | 0.28779873 | -0.5583204 | 0.57662563 | 0.88105119 |
| Miip | 1343.83146 | -0.0565815 | 0.19273115 | -0.2935771 | 0.76908108 | 0.94401697 |
| Dffb | 840.31066 | -0.0285528 | 0.29288677 | -0.0974874 | 0.92233932 | 0.98463888 |
| Lrrc47 | 323.959576 | -0.040087 | 0.33487624 | -0.1197069 | 0.90471535 | 0.98132496 |
| Wrap73 | 843.694135 | -0.1518124 | 0.2590459 | -0.5860443 | 0.55784571 | 0.87291829 |
| Tprgl | 451.571697 | 0.40776664 | 0.37176616 | 1.09683636 | 0.27271293 | 0.71005689 |
| Acap3 | 553.180669 | 0.11718039 | 0.38526867 | 0.30415239 | 0.7610118 | 0.94244354 |
| Ints11 | 4303.42023 | -0.200448 | 0.17417398 | -1.1508494 | 0.24979419 | 0.6842534 |
| Atad3a | 5102.16798 | -0.3866907 | 0.21072307 | -1.835066 | 0.06649591 | 0.39699582 |
| Ssu72 | 6719.13024 | 0.19093269 | 0.27605736 | 0.69164137 | 0.48916257 | 0.84401986 |
| Pex10 | 434.935154 | -0.1497765 | 0.33164191 | -0.4516211 | 0.65154199 | 0.91036394 |
| Rer1 | 7897.38089 | 0.1171897 | 0.25228028 | 0.46452186 | 0.6422739 | 0.90705802 |
| Morn1 | 29.6194276 | -1.2312887 | 1.3570556 | -0.9073237 | 0.36423566 | 0.77081383 |
| Ski | 467.752703 | 0.13256457 | 0.31643274 | 0.41893445 | 0.67526404 | 0.91848602 |
| Pank4 | 1226.91401 | -0.1732458 | 0.25921434 | -0.6683496 | 0.50391044 | 0.85012672 |
| Prxl2b | 1580.09892 | 1.04889059 | 0.36121826 | 2.90375852 | 0.00368712 | 0.0632603 |
| Mib2 | 222.57408 | -0.4494218 | 0.60642684 | -0.7410981 | 0.45863393 | 0.82986925 |
| Cdk11b | 1095.67818 | -0.3391764 | 0.30431459 | -1.1145586 | 0.26503964 | 0.70112124 |
| Nadk | 6210.88054 | -0.3726547 | 0.18924728 | -1.9691416 | 0.04893683 | 0.34113306 |
| Gnb1 | 12118.5696 | -0.0640048 | 0.18926202 | -0.3381808 | 0.73522693 | 0.93750666 |
| Mrpl20 | 4810.26696 | 0.162481 | 0.38931148 | 0.41735475 | 0.67641894 | 0.91889316 |
| Ccnl2 | 2372.0387 | -0.1229646 | 0.19384464 | -0.6343464 | 0.52585481 | 0.86056156 |
| Mxra8 | 159.87906 | 3.03287237 | 0.38951992 | 7.78618037 | 6.906533316 | 2.725407163 |
| Dvl1 | 65.874524 | 0.05303807 | 1.10014669 | 0.04821 | 0.96154888 | 0.99354908 |
| Cptp | 1214.8864 | 0.00313593 | 0.18451897 | 0.01699517 | 0.98644047 | 0.99611549 |
| Sdf4 | 6046.38088 | 0.19873006 | 0.26124986 | 0.76068964 | 0.44684246 | 0.82248455 |
| Bst1 | 157.478489 | -0.7991282 | 0.48248301 | -1.6562826 | 0.0976646 | 0.47892724 |
| Cd38 | 1452.93275 | 1.08933063 | 0.33814525 | 3.22148729 | 0.00127527 | 0.02780819 |
| Prom1 | 45.5050777 | -1.4558588 | 0.77244192 | -1.8847485 | 0.0594638 | 0.3765117 |
| 5730480H06 | 423.260648 | 0.10605262 | 0.42782862 | 0.24788576 | 0.8042228 | 0.9570372 |
| Adgra3 | 102.944743 | 0.15268341 | 0.81294821 | 0.18781443 | 0.85102212 | 0.9679302 |
| Afap1 | 170.034839 | 0.95838097 | 0.48466797 | 1.97739694 | 0.04799678 | 0.33778401 |
| Htra3 | 13.6228441 | -4.0927295 | 1.65785764 | -2.4686858 | 0.01356102 | 0.1574243 |
| Trmt44 | 485.047954 | 0.12506333 | 0.27355556 | 0.45717706 | 0.64754379 | 0.9097332 |
| Acox3 | 759.154023 | -0.3301907 | 0.53005769 | -0.6229335 | 0.53332821 | 0.86492515 |
| Rgs12 | 77.0731137 | -0.1612249 | 0.67339989 | -0.2394192 | 0.81078052 | 0.95912175 |
| Lrpap1 | 6016.61073 | -0.1855083 | 0.17444457 | -1.0634224 | 0.28759043 | 0.72300867 |
| Htt | 524.680691 | 0.43945887 | 0.52583517 | 0.83573503 | 0.40330399 | 0.79832001 |
| Add1 | 3277.07794 | 0.33238404 | 0.19280174 | 1.72396805 | 0.08471361 | 0.44578804 |
| Pcdh7 | 19.7073201 | -0.9335534 | 2.07532213 | -0.4498354 | 0.6528291 | 0.9106891 |
| Rnf4 | 8392.3875 | 0.04310655 | 0.15440708 | 0.27917471 | 0.78011075 | 0.94918389 |
| Nelfa | 2206.4259 | -0.438911 | 0.2261915 | -1.9404401 | 0.05232623 | 0.35365015 |
| Man2b2 | 2286.47034 | -0.0276594 | 0.27295217 | -0.1013343 | 0.91928507 | 0.9839023 |
| Crmp1 | 40.9639746 | 1.03700569 | 1.24959467 | 0.82987365 | 0.40661022 | 0.79992362 |
| Evc | 40.7773164 | -0.6670123 | 1.31274274 | -0.5081059 | 0.61137907 | 0.89568865 |
| Stx18 | 1535.85977 | 0.32327874 | 0.31198696 | 1.03619312 | 0.30011206 | 0.73145694 |

| | | | | | | |
|---|---|---|---|---|---|---|
| Nsg1 | 19.692516 | -0.3625182 | 1.15715389 | -0.3132843 | 0.75406466 | 0.94018119 |
| Zbtb49 | 102.308396 | 0.6514084 | 0.52442311 | 1.2421428 | 0.21418388 | 0.64776608 |
| Rab28 | 1509.11024 | -0.1419329 | 0.18550303 | -0.7651242 | 0.44419759 | 0.82068545 |
| Dnajb6 | 5960.00496 | -0.0867623 | 0.18010428 | -0.4817337 | 0.62999513 | 0.90327361 |
| Fosl2 | 257.073002 | -0.7749093 | 0.75178867 | -1.0307541 | 0.30265615 | 0.73372823 |
| Rbks | 582.730338 | 0.25866387 | 0.35290824 | 0.73294937 | 0.46358931 | 0.83202414 |
| Slc4a1ap | 743.538671 | -0.0755069 | 0.23776673 | -0.3175671 | 0.75081337 | 0.94003261 |
| Eif2b4 | 2229.53089 | -0.0391799 | 0.17545384 | -0.2233062 | 0.8232972 | 0.9628698 |
| Snx17 | 6907.10341 | 0.09002573 | 0.14306706 | 0.62925549 | 0.52918181 | 0.86278569 |
| Ppm1g | 7638.65044 | -0.5617151 | 0.21714907 | -2.5867717 | 0.00968798 | 0.12687515 |
| Nrbp1 | 5078.5131 | 0.21263335 | 0.2021879 | 1.05166206 | 0.29295462 | 0.72604376 |
| Ociad1 | 6575.00259 | 0.1198493 | 0.27916507 | 0.42931338 | 0.66769518 | 0.91595755 |
| Ociad2 | 95.6391922 | 1.43337109 | 0.72544084 | 1.97586214 | 0.04817039 | 0.33815639 |
| Sgcb | 783.111219 | 0.15611543 | 0.41093091 | 0.37990677 | 0.70401462 | 0.92850227 |
| Khk | 1385.18081 | 0.17959786 | 0.36017287 | 0.49864351 | 0.61803055 | 0.89878056 |
| Emilin1 | 4678.19452 | 0.08923615 | 0.26357389 | 0.33856218 | 0.73493958 | 0.93750666 |
| Agbl5 | 64.2478616 | -0.2053713 | 0.68369718 | -0.3003834 | 0.76388469 | 0.94300309 |
| Mapre3 | 28.4840019 | 2.26097165 | 1.71964452 | 1.31479014 | 0.18858044 | 0.61935434 |
| Dhx15 | 5870.68596 | -0.3588796 | 0.18239583 | -1.9675866 | 0.04911563 | 0.34139551 |
| Pgm2 | 2085.7235 | -0.214746 | 0.20048419 | -1.0711366 | 0.28410799 | 0.71971279 |
| Sepsecs | 809.697945 | 0.05056949 | 0.32529902 | 0.15545539 | 0.87646232 | 0.9745677 |
| Tbc1d1 | 1207.25204 | 0.0137605 | 0.38150317 | 0.03606916 | 0.97122721 | 0.99443087 |
| Slc35f6 | 1806.38861 | 0.74315939 | 0.26628637 | 2.79082773 | 0.00525734 | 0.08309185 |
| Anapc4 | 3623.00861 | 0.01641723 | 0.16315367 | 0.10062432 | 0.91984869 | 0.9839023 |
| Cenpa | 6030.28678 | 0.31504258 | 0.3240566 | 0.9721838 | 0.33095912 | 0.75011761 |
| Klf3 | 1054.68684 | 0.57865677 | 0.58411454 | 0.99065633 | 0.32185342 | 0.74447686 |
| Zcchc4 | 435.349331 | 0.01414286 | 0.2760759 | 0.05122816 | 0.95914372 | 0.99271349 |
| Fam114a1 | 401.054668 | 0.72746384 | 0.29375971 | 2.47639082 | 0.01327182 | 0.15566077 |
| Pi4k2b | 336.782913 | -0.0840509 | 0.46721929 | -0.179896 | 0.85723424 | 0.96936093 |
| D5Ertd579e | 408.719941 | -0.011887 | 0.25158551 | -0.0472483 | 0.96231536 | 0.99400521 |
| Rfc1 | 2358.68735 | -0.0468977 | 0.263577 | -0.1779277 | 0.85877972 | 0.96954641 |
| Tbc1d14 | 4737.05857 | 0.58180949 | 0.2139784 | 2.71901044 | 0.00654775 | 0.09792014 |
| Tada2b | 101.837338 | -0.3829723 | 0.41388904 | -0.925302 | 0.35480886 | 0.76419569 |
| Grpel1 | 8151.40945 | 0.3396654 | 0.37069445 | 0.91629482 | 0.35951229 | 0.76857232 |
| Lias | 1718.31343 | 0.12135247 | 0.25532794 | 0.47528082 | 0.63458683 | 0.9050692 |
| Ugdh | 1998.44288 | -0.1216095 | 0.19887304 | -0.6114933 | 0.54087306 | 0.86762394 |
| Pds5a | 882.6061 | -0.3651167 | 0.34072956 | -1.0715733 | 0.28391172 | 0.71951357 |
| Ube2k | 3759.30442 | -0.0524799 | 0.15571011 | -0.3370357 | 0.73609001 | 0.9377627 |
| Rhoh | 1436.46155 | 0.61837381 | 0.37857 | 1.6334464 | 0.10237515 | 0.48862862 |
| Apbb2 | 112.845018 | -0.4471471 | 0.98363551 | -0.4545862 | 0.64940695 | 0.90973357 |
| Guf1 | 781.417244 | -0.2209096 | 0.34289459 | -0.6442494 | 0.51941374 | 0.8576517 |
| Gnpda2 | 962.423324 | 0.2620838 | 0.27743607 | 0.94466374 | 0.34483057 | 0.75991937 |
| Commd8 | 6287.0823 | 0.20293527 | 0.28181412 | 0.72010328 | 0.47146141 | 0.83393588 |
| Tec | 1487.49034 | 0.47582892 | 0.20909122 | 2.2757002 | 0.02286396 | 0.21648205 |
| Slc30a9 | 1931.12599 | 0.21554558 | 0.22489737 | 0.95841753 | 0.33785226 | 0.75611716 |
| Uchl1 | 12.7505425 | 1.55808342 | 1.5685937 | 0.99329955 | 0.320564 | 0.74413737 |

| | | | | | | |
|---|---|---|---|---|---|---|
| Fip1l1 | 2270.87638 | -0.1796866 | 0.21914598 | -0.8199404 | 0.4122501 | 0.80251302 |
| Chic2 | 191.94814 | 0.57569309 | 0.42730634 | 1.34726083 | 0.17789624 | 0.60906934 |
| Srd5a3 | 1846.36187 | 0.4434536 | 0.17086607 | 2.59532861 | 0.00945005 | 0.1248407 |
| Tmem165 | 941.532151 | -0.1215756 | 0.2811441 | -0.4324315 | 0.6654278 | 0.91503803 |
| Clock | 156.087671 | 0.20493472 | 0.50865579 | 0.40289469 | 0.6870257 | 0.92426981 |
| Ppat | 4575.65572 | -0.422576 | 0.17514801 | -2.4126794 | 0.01583574 | 0.17485494 |
| Paics | 19612.6824 | -0.0090992 | 0.23918891 | -0.0380419 | 0.9696543 | 0.99443087 |
| Rest | 1337.84144 | -0.1019259 | 0.39463178 | -0.258281 | 0.79619007 | 0.9537323 |
| Polr2b | 5532.02719 | -0.3582997 | 0.21725833 | -1.6491874 | 0.09910924 | 0.48073092 |
| Cenpc1 | 915.957847 | 0.11695257 | 0.21202944 | 0.55158647 | 0.58123171 | 0.88314588 |
| Stap1 | 300.925703 | -1.0737953 | 0.47679016 | -2.2521339 | 0.02431381 | 0.22532643 |
| Pigg | 399.959183 | 0.13036757 | 0.37408575 | 0.34849649 | 0.72746735 | 0.93576253 |
| Dr1 | 2433.25748 | -0.1411298 | 0.19332367 | -0.7300183 | 0.46537899 | 0.83339036 |
| Mtf2 | 2907.71842 | -0.266685 | 0.16138278 | -1.6525 | 0.09843266 | 0.48011435 |
| Dipk1a | 1874.23936 | -0.1216361 | 0.17194199 | -0.7074254 | 0.47930214 | 0.8387067 |
| Glmn | 865.156448 | 0.32148486 | 0.30554464 | 1.05216986 | 0.29272163 | 0.72584701 |
| Cdc7 | 1070.07081 | -0.1008442 | 0.24157574 | -0.4174433 | 0.67635415 | 0.91889316 |
| Tgfbr3 | 34.8214446 | 0.83659002 | 1.81685923 | 0.46045946 | 0.64518647 | 0.90835953 |
| Zfp326 | 492.81164 | -0.2403181 | 0.25612787 | -0.938274 | 0.34810362 | 0.76101142 |
| Rufy3 | 154.7926 | 0.28872397 | 0.46448448 | 0.62160091 | 0.53420432 | 0.86520872 |
| Gbp9 | 363.946523 | 0.26894584 | 0.33931754 | 0.79260813 | 0.42800617 | 0.81099744 |
| Abcg3 | 1934.01843 | 0.58005937 | 0.27395914 | 2.11732075 | 0.03423263 | 0.27677979 |
| Spp1 | 1184.71637 | 0.35379386 | 0.43228598 | 0.81842548 | 0.41311428 | 0.80343831 |
| Sparcl1 | 40.0176166 | 3.65392262 | 3.60845435 | 1.01260048 | 0.31125103 | 0.74011268 |
| Nudt9 | 1938.51382 | 0.29375725 | 0.18994789 | 1.54651495 | 0.12198025 | 0.5306488 |
| Hsd17b11 | 43.3466397 | 0.26418075 | 0.68702079 | 0.38453094 | 0.70058497 | 0.92706828 |
| Klhl8 | 145.505436 | 0.06488776 | 0.50444977 | 0.12863076 | 0.89764983 | 0.97984623 |
| Aff1 | 447.122383 | -0.1503681 | 0.40941624 | -0.3672743 | 0.71341441 | 0.93235217 |
| Gpat3 | 39.3207788 | 1.58734314 | 1.05185584 | 1.50908811 | 0.13127627 | 0.54970237 |
| Coq2 | 700.525596 | 0.19946379 | 0.26983446 | 0.739208 | 0.4597807 | 0.82987212 |
| Plac8 | 4271.86543 | -2.1064567 | 0.53202636 | -3.959309 | 7.516693951 | 0.00282928 |
| Enoph1 | 3488.85452 | -0.2877064 | 0.22345554 | -1.2875333 | 0.19790847 | 0.63076766 |
| Hnrnpdl | 3879.3583 | -0.1846697 | 0.21058732 | -0.8769267 | 0.38052643 | 0.78323054 |
| Cds1 | 29.1181802 | -0.3467937 | 1.62076872 | -0.2139687 | 0.8305715 | 0.96466657 |
| Antxr2 | 482.386117 | 0.19668425 | 0.44001651 | 0.44699289 | 0.65488019 | 0.91108261 |
| Crybb1 | 154.438269 | 2.04519439 | 0.45832843 | 4.46229001 | 8.108837821 | 0.00043892 |
| Tpst2 | 5820.8063 | -0.2003485 | 0.16943888 | -1.1824233 | 0.23703777 | 0.6732629 |
| Tfip11 | 1938.63955 | 0.06113035 | 0.22794098 | 0.26818498 | 0.78855693 | 0.95225386 |
| Srrd | 120.255271 | -0.2556714 | 0.46016366 | -0.5556097 | 0.5784777 | 0.88230592 |
| Rfc5 | 7508.62035 | -0.1691181 | 0.2333801 | -0.7246465 | 0.46866889 | 0.83381643 |
| Wsb2 | 238.086905 | 0.60968018 | 0.48916829 | 1.24636078 | 0.21263197 | 0.64596443 |
| Dck | 2592.00556 | -0.170983 | 0.20570613 | -0.8312004 | 0.40586041 | 0.79990697 |
| Ppbp | 103.345955 | 1.19690694 | 0.58377498 | 2.05028817 | 0.04033632 | 0.3051541 |
| Pf4 | 2825.45901 | 3.616741 | 0.38142705 | 9.48213044 | 2.491435628 | 2.176980860 |
| Mthfd2l | 122.150392 | 1.09777952 | 0.46188365 | 2.37674469 | 0.01746617 | 0.18560042 |
| Cxcl3 | 22.3086353 | 1.19396315 | 1.92800692 | 0.61927327 | 0.53573635 | 0.86539693 |

| | | | | | | |
|---|---|---|---|---|---|---|
| Cxcl1 | 84.0654252 | 4.03966703 | 1.48802273 | 2.71478853 | 0.00663181 | 0.09869457 |
| Ccng2 | 498.060014 | 0.62944602 | 0.40635719 | 1.54899687 | 0.12138247 | 0.5291774 |
| Gtf2h3 | 1343.24921 | -0.2377415 | 0.21334694 | -1.1143423 | 0.26513237 | 0.70112124 |
| Eif2b1 | 6903.90137 | -0.0648974 | 0.18323448 | -0.3541768 | 0.72320638 | 0.93433544 |
| Ddx55 | 451.414011 | -0.5309926 | 0.29274784 | -1.8138223 | 0.06970509 | 0.40760152 |
| Tmed2 | 31108.1612 | 0.2118168 | 0.21109367 | 1.00342561 | 0.31565555 | 0.74186126 |
| Rilpl1 | 31.2061838 | 0.92996066 | 0.83405443 | 1.11498797 | 0.26485558 | 0.70112124 |
| Cdk2ap1 | 474.645942 | -0.5210744 | 0.33496876 | -1.5555909 | 0.11980541 | 0.52718991 |
| Rchy1 | 1919.57585 | 0.0255318 | 0.23043312 | 0.11079918 | 0.9117756 | 0.98292561 |
| Rilpl2 | 2133.5963 | 0.35649316 | 0.33927633 | 1.05074574 | 0.29337538 | 0.72646828 |
| Snrnp35 | 433.250891 | -0.2428658 | 0.39152004 | -0.6203151 | 0.53505034 | 0.86539693 |
| Cdkl2 | 15.5254601 | 2.11236196 | 1.40623608 | 1.50213893 | 0.13306123 | 0.55290835 |
| Arl6ip4 | 3262.84707 | 0.07693859 | 0.24902269 | 0.30896216 | 0.75735031 | 0.94116141 |
| G3bp2 | 9013.90979 | -0.1965136 | 0.15337393 | -1.281271 | 0.20009849 | 0.63253708 |
| Pitpnm2 | 43.5872826 | -2.9919228 | 0.78163254 | -3.8277869 | 0.0001293 | 0.00449356 |
| Uso1 | 524.065003 | -0.8189745 | 0.4337732 | -1.8880246 | 0.05902265 | 0.3744938 |
| Abcb9 | 27.8008997 | 0.31786744 | 1.867916 | 0.17017223 | 0.86487469 | 0.9705375 |
| Naaa | 1325.18209 | 0.06165952 | 0.24551287 | 0.25114576 | 0.80170142 | 0.95629571 |
| Kntc1 | 333.063235 | -0.4016804 | 0.41955664 | -0.9573924 | 0.33836921 | 0.75644564 |
| Sdad1 | 1436.34107 | -0.5435525 | 0.29408107 | -1.8483085 | 0.06455773 | 0.39212252 |
| Slc15a4 | 1782.50465 | -0.2111025 | 0.30266714 | -0.697474 | 0.48550619 | 0.84183502 |
| Rsrc2 | 2014.16372 | -0.2101005 | 0.31428601 | -0.668501 | 0.50381387 | 0.85012672 |
| Scarb2 | 2178.46514 | 0.2188566 | 0.17769473 | 1.23164378 | 0.21808218 | 0.65150232 |
| Zcchc8 | 1458.15808 | -0.1417 | 0.29595762 | -0.4787846 | 0.63209187 | 0.90405469 |
| Stx2 | 620.512196 | 0.07312541 | 0.38063057 | 0.1921165 | 0.84765095 | 0.96765585 |
| Ran | 72853.5125 | 0.23997124 | 0.37455335 | 0.64068642 | 0.52172644 | 0.85816354 |
| Nipsnap2 | 1589.87638 | 0.40446095 | 0.20021284 | 2.02015492 | 0.04336732 | 0.31946485 |
| Diablo | 2301.48768 | -0.232327 | 0.24242615 | -0.9583412 | 0.33789074 | 0.75611716 |
| Vps33a | 4162.18134 | 0.37736136 | 0.18513613 | 2.03829123 | 0.04152082 | 0.31027743 |
| Bcl7a | 103.685336 | -0.3587567 | 0.50683338 | -0.7078396 | 0.4790449 | 0.83869135 |
| Sfswap | 733.191407 | -0.2496331 | 0.39974146 | -0.6244863 | 0.53230824 | 0.86442675 |
| Psmd9 | 1964.58286 | 0.17389928 | 0.24451837 | 0.71119106 | 0.47696585 | 0.83675398 |
| Psph | 1097.34645 | -0.12802 | 0.34931333 | -0.3664903 | 0.71399924 | 0.93235217 |
| Cct6a | 35117.7599 | 0.05541047 | 0.27335956 | 0.20270177 | 0.83936814 | 0.96757914 |
| Rhof | 317.463068 | -0.6921531 | 0.29928908 | -2.3126574 | 0.02074148 | 0.20511769 |
| Tmem116 | 20.3607455 | 0.26396931 | 1.24803671 | 0.21150765 | 0.83249116 | 0.96556977 |
| Mapkapk5 | 235.1463 | -0.4385772 | 0.29841711 | -1.4696786 | 0.14164883 | 0.5664564 |
| Aldh2 | 3647.62542 | -0.0043822 | 0.21016243 | -0.0208514 | 0.98336421 | 0.99564261 |
| Acad10 | 69.3439352 | -0.2839466 | 1.4596855 | -0.1945259 | 0.84576413 | 0.96765585 |
| Brap | 1563.22681 | -0.0390688 | 0.26960789 | -0.1449096 | 0.88478222 | 0.97742337 |
| Fam168a | 518.56993 | 0.20859041 | 0.33408575 | 0.62436187 | 0.53238996 | 0.8644447 |
| Vps29 | 8831.10416 | 0.81210965 | 0.39769191 | 2.04205726 | 0.04114585 | 0.30904863 |
| Fam216a | 531.679765 | 0.3223471 | 0.44470851 | 0.7248503 | 0.46854388 | 0.83381643 |
| Gpn3 | 1593.69309 | 0.04054222 | 0.25474602 | 0.15914763 | 0.87355256 | 0.97344599 |
| Arpc3 | 1869.1098 | 0.23104277 | 0.31695898 | 0.72893589 | 0.46604088 | 0.83345004 |
| Anapc7 | 2501.56951 | 0.1376098 | 0.18022838 | 0.76353017 | 0.44514728 | 0.82118815 |

| | | | | | | |
|---|---|---|---|---|---|---|
| Atp2a2 | 1637.00396 | -0.6170797 | 0.3129715 | -1.9716802 | 0.04864613 | 0.34046898 |
| P2rx7 | 920.480257 | 0.0506531 | 0.25859964 | 0.1958746 | 0.84470832 | 0.96765585 |
| Ift81 | 83.3334481 | -0.27239 | 0.63548206 | -0.4286352 | 0.66818875 | 0.91597175 |
| P2rx4 | 2812.32416 | 0.83126212 | 0.25878184 | 3.21221194 | 0.00131717 | 0.02856908 |
| Camkk2 | 393.039981 | -0.115481 | 0.44543103 | -0.2592567 | 0.7954372 | 0.9537323 |
| Anapc5 | 16155.9703 | -0.1844029 | 0.15466137 | -1.1923012 | 0.23314316 | 0.66870814 |
| Rnf34 | 1823.53036 | -0.2823324 | 0.22639995 | -1.2470514 | 0.21237866 | 0.64580271 |
| Kdm2b | 2672.9876 | -0.1149266 | 0.31752436 | -0.3619458 | 0.71739255 | 0.93270943 |
| Ncor2 | 246.895542 | -0.9691768 | 0.40085017 | -2.4178031 | 0.01561452 | 0.17348998 |
| Dhx37 | 934.013619 | -0.8426028 | 0.27981614 | -3.0112731 | 0.00260155 | 0.04821929 |
| Aacs | 1875.77477 | 0.20528233 | 0.22654097 | 0.90615982 | 0.36485126 | 0.77108653 |
| Anxa3 | 13312.7618 | 0.00881669 | 0.23709877 | 0.03718571 | 0.97033693 | 0.99443087 |
| Mrpl1 | 2097.68587 | 0.06572787 | 0.35862001 | 0.18327997 | 0.85457835 | 0.96859981 |
| Pxmp2 | 129.329281 | 0.98302225 | 0.53578573 | 1.83473018 | 0.06654568 | 0.39709916 |
| Pgam5 | 3458.49149 | -0.0663695 | 0.17577745 | -0.3775769 | 0.70574493 | 0.92896305 |
| Ankle2 | 1465.93686 | -0.0363672 | 0.24470937 | -0.1486138 | 0.88185839 | 0.97582756 |
| Golga3 | 731.628856 | -0.1781154 | 0.49446338 | -0.3602197 | 0.71868286 | 0.93310883 |
| Ddx51 | 1214.76672 | -0.1791202 | 0.46716432 | -0.38342 | 0.70140834 | 0.92738786 |
| Ep400 | 1511.80698 | 0.235036 | 0.41546184 | 0.56572223 | 0.57158262 | 0.88035336 |
| Pus1 | 1266.58506 | -0.4614889 | 0.26472452 | -1.7432797 | 0.08128477 | 0.43986231 |
| Ulk1 | 196.542833 | 0.02891198 | 0.38999714 | 0.07413382 | 0.94090391 | 0.98935054 |
| Prkab1 | 3712.00063 | 0.093388 | 0.21537376 | 0.43360899 | 0.66457241 | 0.91431173 |
| Cit | 500.373964 | 0.34563109 | 0.47751575 | 0.72381086 | 0.46918186 | 0.83381643 |
| Rab35 | 305.473216 | -0.3727239 | 0.34202164 | -1.089767 | 0.2758158 | 0.7118544 |
| Chek2 | 1489.30791 | 0.18305521 | 0.30356364 | 0.60302086 | 0.54649481 | 0.86994632 |
| Sirt4 | 89.6035726 | 0.656859 | 0.50721347 | 1.29503461 | 0.19530829 | 0.62707955 |
| Pxn | 2005.41391 | -0.150863 | 0.27793721 | -0.5427952 | 0.58727082 | 0.88632735 |
| St7 | 160.544727 | -0.0135419 | 0.36242248 | -0.037365 | 0.97019399 | 0.99443087 |
| Triap1 | 2188.75991 | 0.42862094 | 0.34408249 | 1.24569241 | 0.21287734 | 0.64596443 |
| Gatc | 1179.69621 | -0.1998196 | 0.20712408 | -0.9647339 | 0.3346781 | 0.75351065 |
| Srsf9 | 348.806673 | -0.4979189 | 0.37409636 | -1.330991 | 0.18319196 | 0.61498255 |
| Acads | 2062.34497 | 0.27047543 | 0.26119416 | 1.035534 | 0.3004196 | 0.73158785 |
| Ints1 | 508.228015 | -0.63069 | 0.32425527 | -1.9450416 | 0.05176997 | 0.35085985 |
| Sppl3 | 222.658265 | 0.10298126 | 0.41994634 | 0.24522481 | 0.80628237 | 0.95741157 |
| Psmg3 | 1840.66291 | 0.15916047 | 0.38779693 | 0.4104222 | 0.68149627 | 0.92061965 |
| Tes | 2056.84432 | -0.5195767 | 0.27155704 | -1.9133244 | 0.05570654 | 0.36325056 |
| Tfec | 2811.63428 | -0.0966148 | 0.23869329 | -0.4047656 | 0.68564985 | 0.92314423 |
| Mad1l1 | 1949.86004 | -0.2589415 | 0.29131478 | -0.8888718 | 0.37407196 | 0.77748705 |
| Mrm2 | 490.892158 | 0.56085968 | 0.39362545 | 1.42485623 | 0.15419876 | 0.58309536 |
| 2210016L21F | 1008.82486 | 0.13724288 | 0.27726041 | 0.49499629 | 0.62060274 | 0.89896588 |
| Snx8 | 2315.49422 | 0.63265992 | 0.1743209 | 3.62928317 | 0.00028421 | 0.00854234 |
| Oasl2 | 126.731222 | -0.166022 | 0.76925938 | -0.2158206 | 0.82912762 | 0.96462215 |
| Foxp2 | 31.1664452 | -3.3318799 | 1.64046928 | -2.0310529 | 0.04224963 | 0.31437938 |
| Tmem168 | 1738.91676 | 0.23733557 | 0.23888277 | 0.9935232 | 0.32045505 | 0.74413737 |
| Lfng | 98.6613434 | -0.2949776 | 0.49571525 | -0.5950546 | 0.551807 | 0.87093412 |
| Tmem106b | 688.251843 | 0.78984973 | 0.30202436 | 2.61518549 | 0.0089179 | 0.1197646 |

| | | | | | | |
|---|---|---|---|---|---|---|
| Mmab | 746.729437 | 0.30240179 | 0.21103621 | 1.4329379 | 0.15187558 | 0.57999887 |
| Ube3b | 1668.40052 | -0.0367769 | 0.29875039 | -0.1231024 | 0.90202604 | 0.98043702 |
| Wipi2 | 2367.57404 | 0.12502686 | 0.22903387 | 0.54588808 | 0.58514287 | 0.8848945 |
| Actb | 525046.3 | -0.083354 | 0.19133745 | -0.4356389 | 0.66309877 | 0.91395288 |
| Fscn1 | 882.767045 | 0.97620284 | 0.27051365 | 3.60870081 | 0.00030773 | 0.00909303 |
| Zfp12 | 125.848279 | -0.1623508 | 0.62743564 | -0.258753 | 0.79582583 | 0.9537323 |
| Ung | 1761.16323 | -0.2067265 | 0.17857272 | -1.1576599 | 0.24700285 | 0.68185412 |
| Usp30 | 150.755317 | -0.4598044 | 0.46154497 | -0.9962288 | 0.31913899 | 0.74398524 |
| Rbm19 | 1130.12571 | -0.2317053 | 0.34133399 | -0.6788228 | 0.49725013 | 0.84814011 |
| Plbd2 | 2390.04132 | 0.03078971 | 0.26857811 | 0.11463967 | 0.90873073 | 0.98202324 |
| Ddx54 | 4348.9151 | -0.4477717 | 0.21605805 | -2.0724601 | 0.03822255 | 0.29537366 |
| Rita1 | 270.658486 | -0.1027961 | 0.29557437 | -0.3477842 | 0.72800229 | 0.93576253 |
| Rasal1 | 29.3207364 | -1.8739946 | 1.27458017 | -1.4702838 | 0.1414849 | 0.56628837 |
| Aimp2 | 4667.26925 | 0.07451318 | 0.32351465 | 0.23032398 | 0.81784003 | 0.96079676 |
| Eif2ak1 | 2890.88123 | -0.465767 | 0.24011441 | -1.939771 | 0.05240752 | 0.35380862 |
| Rpl6 | 37792.984 | -0.0099856 | 0.28737998 | -0.0347471 | 0.97228142 | 0.99443087 |
| Erp29 | 15960.8366 | 0.03682078 | 0.27344011 | 0.13465758 | 0.89288262 | 0.97941957 |
| Ccz1 | 1185.76511 | -0.0971254 | 0.2870878 | -0.3383127 | 0.73512756 | 0.93750666 |
| Arpc1a | 3019.2212 | -0.0115267 | 0.23713909 | -0.0486072 | 0.96123236 | 0.99338983 |
| Arpc1b | 43729.8839 | -0.0414794 | 0.18446716 | -0.2248605 | 0.82208777 | 0.96235404 |
| Pdap1 | 740.741233 | -0.0457324 | 0.19967541 | -0.2290335 | 0.81884288 | 0.96105733 |
| Ptcd1 | 877.263942 | -0.2550518 | 0.20254167 | -1.2592559 | 0.20793791 | 0.6418276 |
| Cpsf4 | 1364.9621 | 0.09407313 | 0.2008531 | 0.4683678 | 0.63952159 | 0.90646657 |
| Zkscan14 | 765.362497 | 0.27156627 | 0.38102107 | 0.71273295 | 0.47601102 | 0.83644936 |
| Phf14 | 351.85013 | 0.17568625 | 0.29667172 | 0.59219077 | 0.55372285 | 0.87165957 |
| Ndufa4 | 7804.53753 | 0.38981453 | 0.38584619 | 1.01028478 | 0.31235887 | 0.74051791 |
| Rnf6 | 1297.1551 | -0.1182578 | 0.40648231 | -0.2909299 | 0.77110497 | 0.94488554 |
| Cdk8 | 255.449861 | -0.4483978 | 0.33605739 | -1.3342893 | 0.18210907 | 0.61344341 |
| Glcci1 | 52.8415615 | -0.3916848 | 1.24229746 | -0.3152907 | 0.75254095 | 0.94018119 |
| Usp12 | 696.732283 | -0.0969462 | 0.51776529 | -0.1872397 | 0.85147272 | 0.9679302 |
| Rasl11a | 23.8072741 | 1.23778818 | 1.3842667 | 0.89418331 | 0.37122382 | 0.7765614 |
| Polr1d | 5766.72727 | 0.04099255 | 0.34074335 | 0.12030332 | 0.90424288 | 0.98098644 |
| Pan3 | 706.847252 | -0.0019657 | 0.27135115 | -0.007244 | 0.99422014 | 0.99880383 |
| Flt1 | 212.3516 | -0.9793194 | 0.47872975 | -2.0456622 | 0.04078962 | 0.30763217 |
| Pomp | 9763.59947 | 0.13439126 | 0.29648533 | 0.4532813 | 0.65034619 | 0.9101573 |
| Slc46a3 | 2956.17241 | 0.46692325 | 0.23594992 | 1.97890829 | 0.04782633 | 0.33759927 |
| Mtus2 | 60.1411457 | 0.14265591 | 0.54451238 | 0.26198836 | 0.79333042 | 0.95310033 |
| N4bp2l2 | 1503.67285 | 0.11557003 | 0.21754254 | 0.53125255 | 0.59524378 | 0.88830085 |
| Hsph1 | 11408.3101 | -0.1746867 | 0.36502033 | -0.4785671 | 0.63224661 | 0.90417031 |
| Col1a2 | 37.2116118 | 2.31657665 | 1.38040299 | 1.67818867 | 0.09331027 | 0.46713342 |
| Tspan12 | 121.911617 | -0.5743019 | 0.55049517 | -1.0432461 | 0.29683432 | 0.72954201 |
| Ing3 | 1030.75697 | -0.2296496 | 0.23184592 | -0.9905269 | 0.32191665 | 0.74447686 |
| Fam3c | 833.343619 | -0.2852961 | 0.33151198 | -0.8605905 | 0.38946363 | 0.78942657 |
| Auts2 | 25.7275278 | -1.1892847 | 1.32410748 | -0.8981784 | 0.36909044 | 0.77480275 |
| Limk1 | 146.734612 | 0.08319232 | 0.5035408 | 0.16521465 | 0.86877504 | 0.97198875 |
| Pot1a | 1507.71234 | 0.00090908 | 0.27575687 | 0.00329666 | 0.99736965 | 0.99957586 |

| | | | | | | |
|---|---|---|---|---|---|---|
| Bcl7b | 1338.3067 | 0.36404159 | 0.2690615 | 1.35300517 | 0.17605398 | 0.60763832 |
| Wasl | 284.753396 | 0.71772348 | 0.39797838 | 1.80342331 | 0.07132176 | 0.41122734 |
| Cul1 | 6835.97657 | -0.139821 | 0.17192623 | -0.8132617 | 0.41606804 | 0.80440685 |
| Ezh2 | 1650.35981 | -0.23761 | 0.28611867 | -0.8304597 | 0.4062789 | 0.79992362 |
| Ssc4d | 119.37909 | 0.10161189 | 0.56155379 | 0.18094775 | 0.85640859 | 0.96924897 |
| Rbm28 | 1497.16285 | -0.7495918 | 0.25231169 | -2.9708961 | 0.00296932 | 0.0534959 |
| Lrwd1 | 2252.51896 | -0.2825475 | 0.20160883 | -1.4014638 | 0.16107541 | 0.59141542 |
| Cux1 | 1177.6738 | 0.43722338 | 0.31908626 | 1.37023569 | 0.17061334 | 0.60430173 |
| Gcc1 | 358.778982 | -0.3500618 | 0.49627417 | -0.7053799 | 0.48057383 | 0.83935746 |
| Ephb4 | 73.1450307 | 3.10816396 | 3.00799634 | 1.03330045 | 0.30146332 | 0.73330242 |
| Gnb2 | 1823.38236 | -0.2415439 | 0.22786699 | -1.0600215 | 0.28913482 | 0.72395811 |
| Gigyf1 | 100.733779 | -0.0976894 | 0.6568243 | -0.1487299 | 0.88176675 | 0.97582756 |
| Pop7 | 1015.33924 | 0.04008329 | 0.22636416 | 0.17707436 | 0.85944998 | 0.96954641 |
| Pcolce | 42.4789599 | 2.18932403 | 1.51238399 | 1.44759799 | 0.14772951 | 0.57472133 |
| Gm20605 | 279.357003 | -0.1793027 | 0.38906602 | -0.4608542 | 0.64490319 | 0.90835328 |
| Agfg2 | 584.139915 | -0.0772841 | 0.37382074 | -0.206741 | 0.83621215 | 0.96683245 |
| Tsc22d4 | 2487.83861 | -0.2128937 | 0.22867628 | -0.9309828 | 0.35186244 | 0.76348958 |
| Ppp1r35 | 224.223749 | 0.63124761 | 0.59385242 | 1.06297052 | 0.28779532 | 0.72300867 |
| Mepce | 204.249957 | 0.07957521 | 0.5515696 | 0.14427047 | 0.88528689 | 0.97751512 |
| Zkscan1 | 377.335912 | -0.3695099 | 0.39545688 | -0.9343874 | 0.35010409 | 0.76216093 |
| Mcm7 | 15094.9785 | -0.2373581 | 0.14249847 | -1.6656891 | 0.09577535 | 0.47453216 |
| Tpk1 | 297.921303 | 0.38229405 | 0.58879665 | 0.64928028 | 0.51615723 | 0.85627177 |
| Asns | 4706.1962 | -0.1603684 | 0.16274556 | -0.9853936 | 0.32443076 | 0.74596967 |
| Pon3 | 1899.77334 | 1.05148909 | 0.32499175 | 3.23543322 | 0.00121458 | 0.02711313 |
| Cald1 | 136.358408 | 0.898909 | 1.11361817 | 0.80719678 | 0.41955312 | 0.80563709 |
| Akr1b8 | 142.817831 | 1.45090565 | 0.44304992 | 3.27481305 | 0.00105732 | 0.02426677 |
| Exoc4 | 1222.81647 | 0.17214776 | 0.25157867 | 0.68427011 | 0.49380462 | 0.84678075 |
| Calu | 10497.0748 | 0.13844206 | 0.17244157 | 0.80283461 | 0.42207033 | 0.80821275 |
| Irf5 | 8790.3905 | -0.6387072 | 0.2361102 | -2.7051233 | 0.0068279 | 0.10051233 |
| Ahcyl2 | 152.461503 | -0.1098256 | 0.58371295 | -0.1881501 | 0.85075902 | 0.9679302 |
| Klhdc10 | 277.650597 | -0.6074186 | 0.38770145 | -1.5667174 | 0.11718077 | 0.5231651 |
| Hibadh | 3072.13148 | 0.00234197 | 0.17042657 | 0.01374179 | 0.98903598 | 0.99650865 |
| Gars | 11287.7971 | -0.4608264 | 0.14590817 | -3.1583318 | 0.00158675 | 0.0332945 |
| Nt5c3 | 1674.28076 | -0.1293981 | 0.2506661 | -0.5162168 | 0.60570297 | 0.89258167 |
| Fkbp9 | 1699.50024 | 0.66538972 | 0.21781349 | 3.05485998 | 0.00225166 | 0.04351423 |
| Tmem209 | 2033.85086 | 0.17465552 | 0.16139247 | 1.08217891 | 0.27917304 | 0.71377266 |
| Avl9 | 353.424593 | -0.3179708 | 0.48295452 | -0.6583866 | 0.51028972 | 0.85335831 |
| Cep41 | 235.926978 | -0.0174739 | 0.34077232 | -0.0512775 | 0.95910441 | 0.99271349 |
| Herc6 | 526.813572 | -0.0917895 | 0.31367778 | -0.2926237 | 0.76980983 | 0.9442118 |
| Abcg2 | 3352.99531 | -0.2832446 | 0.29795666 | -0.9506234 | 0.3417956 | 0.75869817 |
| Herc3 | 45.5044793 | -0.0991606 | 0.83794166 | -0.1183383 | 0.90579963 | 0.98144617 |
| Tmem176b | 2282.98053 | -0.9439457 | 0.15104154 | -6.249577 | 4.115657102 | 7.866692709 |
| Igf2bp3 | 6349.18106 | -0.0663813 | 0.19867172 | -0.3341257 | 0.73828474 | 0.93800224 |
| Malsu1 | 1119.81275 | 0.13309139 | 0.30195677 | 0.44076307 | 0.65938453 | 0.91280998 |
| Gpnmb | 55.9937356 | 0.19308525 | 0.74089583 | 0.26061053 | 0.79439287 | 0.95338055 |
| Tra2a | 1559.43695 | -0.0804249 | 0.16162665 | -0.497597 | 0.61876811 | 0.89878056 |

| | | | | | | |
|---|---|---|---|---|---|---|
| Npy | 10.3194676 | -0.3441863 | 1.6453774 | -0.2091838 | 0.83430474 | 0.96629899 |
| Gsdme | 2660.02598 | -0.3538025 | 0.20748331 | -1.7052097 | 0.08815532 | 0.45368281 |
| Osbpl3 | 78.3062518 | 0.0323909 | 0.79840477 | 0.04056952 | 0.96763909 | 0.99443087 |
| Luc7l2 | 256.201643 | -0.0269585 | 0.31493871 | -0.0855993 | 0.93178498 | 0.98730569 |
| Zc3hav1 | 4757.10293 | -0.3315603 | 0.31488013 | -1.0529731 | 0.29235333 | 0.72584701 |
| Trim24 | 116.511499 | 0.8801108 | 0.61148051 | 1.43931128 | 0.15006235 | 0.57784353 |
| Cbx3 | 19912.7194 | -0.3369848 | 0.18320919 | -1.8393442 | 0.06586458 | 0.39558985 |
| Ptn | 33.9406681 | 2.34317947 | 1.30952222 | 1.78933923 | 0.0735602 | 0.41756934 |
| Mtpn | 16529.5667 | -0.104045 | 0.14607945 | -0.7122492 | 0.47631049 | 0.83644936 |
| Zyx | 6091.82408 | -0.2475942 | 0.25482381 | -0.9716289 | 0.33123521 | 0.75037043 |
| Casp2 | 1435.40789 | -0.396622 | 0.18411179 | -2.1542454 | 0.03122094 | 0.26439131 |
| Gstk1 | 95.4963595 | 0.95398696 | 0.60254626 | 1.58325928 | 0.11336238 | 0.51285577 |
| Ephb6 | 20.2194066 | 3.21920501 | 1.48621415 | 2.16604384 | 0.03030784 | 0.26001357 |
| Mad2l1 | 6255.63948 | 0.47974099 | 0.37429882 | 1.28170587 | 0.19994585 | 0.63247751 |
| Ssbp1 | 2978.63326 | -0.0810144 | 0.32095746 | -0.2524148 | 0.80072047 | 0.95576786 |
| Prdm5 | 273.538159 | -0.0151043 | 0.32206881 | -0.0468978 | 0.96259471 | 0.99401755 |
| Clec5a | 2196.2884 | -0.9055376 | 0.21322265 | -4.2469111 | 2.167378984 | 0.00097352 |
| Agk | 1985.24721 | -0.1599917 | 0.20815505 | -0.7686177 | 0.44212027 | 0.81996017 |
| Mrps33 | 4096.54433 | 0.2131007 | 0.34421727 | 0.61908777 | 0.53585854 | 0.86539693 |
| Hpgds | 11533.3248 | 0.88329441 | 0.24150956 | 3.657389 | 0.0002548 | 0.0078315 |
| Smarcad1 | 652.551409 | 0.13322103 | 0.3352863 | 0.39733513 | 0.69112035 | 0.92470267 |
| Mkrn1 | 1691.87043 | 0.1488142 | 0.24682346 | 0.60291756 | 0.54656353 | 0.86994632 |
| Rab19 | 20.6748174 | 0.8125945 | 1.45900537 | 0.556951 | 0.57756091 | 0.88150999 |
| Slc37a3 | 266.379938 | 0.97298902 | 0.38896862 | 2.50145895 | 0.01236828 | 0.14818917 |
| Tbxas1 | 1213.20525 | 0.84675044 | 0.2636822 | 3.21125366 | 0.00132157 | 0.02861379 |
| Gfpt1 | 450.227455 | -0.1859278 | 0.36250109 | -0.5129027 | 0.6080194 | 0.8936563 |
| Nfu1 | 2728.22309 | -0.0362602 | 0.20815821 | -0.1741955 | 0.86171178 | 0.97020895 |
| Anxa4 | 720.438531 | 0.67036137 | 0.28601342 | 2.34381089 | 0.01908785 | 0.19530399 |
| Pcyox1 | 5557.27946 | 0.58445042 | 0.20225831 | 2.88962376 | 0.00385703 | 0.06516998 |
| Dusp11 | 1497.68595 | 0.48682394 | 0.34603289 | 1.4068719 | 0.15946538 | 0.5886662 |
| Cct7 | 35525.5375 | -0.1717736 | 0.20502114 | -0.8378336 | 0.40212419 | 0.79779195 |
| Pradc1 | 1066.39927 | 0.22844793 | 0.31499635 | 0.72523992 | 0.46830486 | 0.83381643 |
| Zfp638 | 1071.49664 | -0.1423431 | 0.28741013 | -0.4952612 | 0.62041573 | 0.89896588 |
| Fbxl14 | 72.3452291 | -0.7188899 | 0.52263517 | -1.3755101 | 0.16897336 | 0.60312169 |
| Kbtbd8 | 251.816493 | 0.40545164 | 0.47366923 | 0.85598054 | 0.39200854 | 0.79031213 |
| Wdr54 | 45.6491392 | 1.57087221 | 1.59245108 | 0.98644927 | 0.3239127 | 0.74583177 |
| Ino80b | 74.5846005 | 0.66434101 | 0.64290757 | 1.03333829 | 0.30144561 | 0.73330242 |
| Wbp1 | 245.589632 | -0.2870994 | 0.34106347 | -0.8417771 | 0.39991274 | 0.79624553 |
| Mogs | 696.92365 | -0.4676472 | 0.23627522 | -1.9792478 | 0.04778812 | 0.33752427 |
| Mrpl53 | 1053.23392 | 0.56624601 | 0.44624335 | 1.26891753 | 0.20447048 | 0.63873528 |
| M1ap | 159.404945 | -0.5059228 | 0.42574993 | -1.1883097 | 0.2347114 | 0.67038085 |
| Pole4 | 2616.32494 | 0.24673722 | 0.31196577 | 0.7909112 | 0.42899581 | 0.81140481 |
| Mrpl19 | 2898.84159 | 0.07419426 | 0.33680161 | 0.22029068 | 0.82564478 | 0.96326388 |
| Arhgap25 | 3810.91256 | 0.25367985 | 0.17660468 | 1.43642768 | 0.15088068 | 0.57859667 |
| Aplf | 660.634454 | -0.1297923 | 0.39552012 | -0.328156 | 0.74279371 | 0.93850397 |
| Gp9 | 9.82255933 | -1.0777984 | 1.52933293 | -0.7047506 | 0.48096543 | 0.83968176 |

| | | | | | | |
|---|---|---|---|---|---|---|
| Rab43 | 161.61598 | 0.83061718 | 0.58206994 | 1.4270058 | 0.15357822 | 0.58242478 |
| Isy1 | 2615.32776 | -0.4437714 | 0.17048716 | -2.602961 | 0.00924225 | 0.12315948 |
| Cnbp | 61717.4105 | 0.20266041 | 0.32827652 | 0.61734665 | 0.5370061 | 0.86596304 |
| Copg1 | 8606.04841 | -0.077948 | 0.15748749 | -0.4949473 | 0.62063735 | 0.89896588 |
| Tmf1 | 2041.49161 | 0.29557965 | 0.20276044 | 1.45777769 | 0.14490183 | 0.57067981 |
| Hmces | 982.92685 | 0.59657684 | 0.2052802 | 2.90615869 | 0.00365896 | 0.06313122 |
| Uba3 | 3202.9209 | 0.4353961 | 0.25746806 | 1.69106838 | 0.09082374 | 0.46044203 |
| Rpn1 | 17452.4062 | -0.0450193 | 0.14399881 | -0.3126366 | 0.75455678 | 0.94044759 |
| Frmd4b | 3027.7914 | 0.78956189 | 0.37785654 | 2.0895811 | 0.03665545 | 0.28743979 |
| Foxp1 | 1734.02365 | -0.4757834 | 0.28578891 | -1.6648071 | 0.09595124 | 0.47482666 |
| Gm20696 | 16.5282198 | -2.2498189 | 1.40160763 | -1.6051703 | 0.10845634 | 0.50301538 |
| Ruvbl1 | 8345.49862 | -0.4267236 | 0.17963076 | -2.3755596 | 0.01752236 | 0.18560042 |
| Sec61a1 | 13958.5846 | -0.1604211 | 0.19613983 | -0.8178913 | 0.41341924 | 0.80364811 |
| Abtb1 | 1015.77676 | -0.1520915 | 0.23568751 | -0.6453099 | 0.51872635 | 0.8570838 |
| Plxna1 | 70.3867042 | -0.2722673 | 0.78580831 | -0.3464805 | 0.72898163 | 0.93617556 |
| Chchd6 | 1529.68246 | 0.34325044 | 0.39249936 | 0.87452485 | 0.38183249 | 0.78454353 |
| Klf15 | 19.201257 | 1.64707145 | 1.8933691 | 0.86991567 | 0.38434649 | 0.78704643 |
| Aldh1l1 | 145.316597 | 1.09319696 | 0.37215661 | 2.93746489 | 0.00330908 | 0.05824451 |
| Slc41a3 | 176.680784 | 1.01096845 | 0.55733955 | 1.81391836 | 0.0696903 | 0.40760152 |
| Nup210 | 1523.84828 | -1.3358816 | 0.22536978 | -5.9275097 | 3.075631373 | 5.084351161 |
| Xpc | 438.766256 | -0.338991 | 0.32467454 | -1.0440947 | 0.29644157 | 0.72935836 |
| Tmem43 | 2451.60795 | -0.1178399 | 0.25203594 | -0.467552 | 0.64010502 | 0.90671662 |
| Slc6a6 | 7802.68885 | 0.28241428 | 0.22515114 | 1.25433197 | 0.20972137 | 0.64347165 |
| Sumf1 | 2680.28247 | 0.22583236 | 0.22054918 | 1.02395464 | 0.3058567 | 0.73554152 |
| Itpr1 | 183.664994 | -2.4226652 | 0.65097253 | -3.721609 | 0.00019796 | 0.00645763 |
| Bhlhe40 | 174.125526 | -0.4748265 | 0.70734829 | -0.6712769 | 0.50204418 | 0.85006796 |
| Edem1 | 2560.86592 | 0.34825967 | 0.48809921 | 0.7135018 | 0.4755353 | 0.83639059 |
| Arl8b | 9293.16194 | 0.1857251 | 0.20345637 | 0.91284978 | 0.36132156 | 0.76887042 |
| Usp18 | 130.807328 | -0.4154233 | 0.59918702 | -0.6933116 | 0.488114 | 0.84329777 |
| Slc6a13 | 27.8551844 | -2.6617432 | 1.15774491 | -2.2990757 | 0.02150064 | 0.21054669 |
| Gdf3 | 111.584606 | 0.36385341 | 0.47365607 | 0.7681806 | 0.4423799 | 0.82019298 |
| Mlf2 | 5059.8725 | 0.09076968 | 0.19870609 | 0.45680371 | 0.64781214 | 0.9097332 |
| Ptms | 47.4413765 | 1.17562064 | 0.74397266 | 1.58019334 | 0.1140626 | 0.51545169 |
| Plxnd1 | 220.945308 | 1.54434164 | 0.39508722 | 3.90886253 | 9.273171119 | 0.00336613 |
| Lag3 | 14.4355162 | 1.50029259 | 2.04868878 | 0.73231845 | 0.46397422 | 0.83222825 |
| Tmcc1 | 182.800154 | -0.0157721 | 0.43143873 | -0.0365569 | 0.97083828 | 0.99443087 |
| Cops7a | 5447.82841 | 2.069368641 | 0.16106425 | 0.00012848 | 0.99989749 | 0.99989749 |
| Tuba8 | 20.7170039 | -3.1354628 | 1.0215607 | -3.0692868 | 0.0021457 | 0.04220001 |
| Bms1 | 2261.69072 | -0.4701961 | 0.28973354 | -1.6228572 | 0.10461994 | 0.49501875 |
| Clec4e | 40.8888519 | -0.1133239 | 1.37927345 | -0.082162 | 0.93451787 | 0.98756396 |
| Clec4d | 106.119242 | 0.02070731 | 0.51106285 | 0.04051812 | 0.96768006 | 0.99443087 |
| Zfp248 | 120.056729 | 0.11506482 | 0.62179354 | 0.1850531 | 0.85318738 | 0.96811477 |
| Clec4a2 | 1630.79181 | 0.33003245 | 0.28446419 | 1.16018977 | 0.24597155 | 0.68117517 |
| Klrk1 | 34.5494351 | -8.5127068 | 1.54276131 | -5.5178378 | 3.431958722 | 3.923658977 |
| Klrb1f | 78.6967871 | -2.966445 | 1.05159548 | -2.8208993 | 0.00478892 | 0.07698147 |
| Cd69 | 707.941044 | -1.353584 | 0.35570637 | -3.805341 | 0.00014161 | 0.00485545 |

| | | | | | | |
|---|---|---|---|---|---|---|
| Clec2d | 20.0323511 | 0.58748344 | 0.87295803 | 0.67298016 | 0.50095991 | 0.84968141 |
| Clec1b | 114.844459 | -1.2041077 | 0.39450505 | -3.0521986 | 0.00227172 | 0.04376365 |
| Gabarapl1 | 1511.25033 | -0.3439164 | 0.22179976 | -1.5505714 | 0.12100442 | 0.52910763 |
| Olr1 | 84.6984848 | -3.1456882 | 0.72710935 | -4.3262931 | 1.516396059 | 0.00075182 |
| Rad52 | 337.485768 | 0.39338556 | 0.42735429 | 0.92051389 | 0.35730428 | 0.7664509 |
| Adipor2 | 827.026875 | -0.523343 | 0.28009008 | -1.868481 | 0.06169506 | 0.38329897 |
| Erc1 | 208.582989 | 0.86239529 | 0.56991976 | 1.51318719 | 0.13023212 | 0.54746717 |
| Ccdc77 | 600.259463 | -0.0774748 | 0.24287603 | -0.318989 | 0.74973482 | 0.9399191 |
| Kdm5a | 1792.86737 | -0.2652234 | 0.21305465 | -1.2448611 | 0.21318283 | 0.64599096 |
| Magohb | 293.383439 | -0.8204607 | 0.3075395 | -2.6678223 | 0.00763446 | 0.10754172 |
| Ybx3 | 1444.70875 | -0.7057707 | 0.58000005 | -1.2168459 | 0.22366284 | 0.65739248 |
| Etv6 | 808.243821 | -0.9135328 | 0.28512869 | -3.2039314 | 0.00135565 | 0.0290941 |
| Bcl2l14 | 18.4444722 | -3.296882 | 1.6584294 | -1.9879544 | 0.04681674 | 0.33444287 |
| Lrp6 | 234.286891 | 0.76467255 | 0.38814846 | 1.97005174 | 0.04883244 | 0.34076854 |
| Dusp16 | 413.159349 | 0.21687103 | 0.44181212 | 0.49086709 | 0.62352045 | 0.90017327 |
| Ddx47 | 5750.08045 | -0.2277617 | 0.15451726 | -1.4740211 | 0.1404759 | 0.56425905 |
| Gsg1 | 25.9492837 | 1.70431478 | 1.45063487 | 1.1748751 | 0.24004469 | 0.67563574 |
| Fam234b | 433.656793 | 0.17443518 | 0.25232626 | 0.69130807 | 0.48937196 | 0.84401986 |
| Emp1 | 12.3241741 | 1.90323221 | 2.01168538 | 0.94608841 | 0.3441035 | 0.75962035 |
| Atf7ip | 2605.72053 | -0.096774 | 0.32227402 | -0.3002849 | 0.76395987 | 0.94300309 |
| Plbd1 | 85.873778 | -2.6050352 | 0.85542011 | -3.0453285 | 0.00232426 | 0.04446651 |
| Wbp11 | 2483.01461 | -0.4151141 | 0.2870045 | -1.446368 | 0.14807402 | 0.57486179 |
| Erp27 | 9.10348198 | -5.6933128 | 1.88734616 | -3.0165705 | 0.00255652 | 0.04760103 |
| Arhgdib | 42532.1673 | -0.1067601 | 0.27864514 | -0.3831401 | 0.70161592 | 0.92738786 |
| Ptpro | 1195.98979 | 0.40979856 | 0.23406389 | 1.75079787 | 0.07998073 | 0.43723638 |
| Strap | 12856.5618 | -0.2468317 | 0.18073886 | -1.3656815 | 0.17203896 | 0.60451272 |
| Dera | 5964.56173 | 0.36745906 | 0.27720127 | 1.32560381 | 0.18497096 | 0.61705748 |
| Plekha5 | 160.519549 | -1.3103089 | 0.51444182 | -2.5470497 | 0.0108638 | 0.13520102 |
| Aebp2 | 262.220676 | -0.2778768 | 0.44761854 | -0.6207893 | 0.53473828 | 0.8652846 |
| Recql | 1823.7078 | -0.3797338 | 0.19704039 | -1.9271874 | 0.05395628 | 0.35989907 |
| Golt1b | 7890.67065 | 0.09105222 | 0.17645097 | 0.51601995 | 0.60584048 | 0.89258167 |
| Ldhb | 489.482337 | -1.0345921 | 0.40641248 | -2.54567 | 0.01090682 | 0.13545501 |
| Abcc9 | 30.6577317 | 4.48395274 | 1.88062897 | 2.38428357 | 0.01711241 | 0.18378939 |
| Rad18 | 1696.99676 | 0.07345247 | 0.18658362 | 0.39367052 | 0.69382431 | 0.92609254 |
| Bhlhe41 | 291.455714 | -0.1168079 | 0.38977181 | -0.2996829 | 0.76441907 | 0.94307414 |
| Srgap3 | 40.3302466 | -0.9842127 | 0.99227059 | -0.9918793 | 0.32125639 | 0.7444458 |
| Rassf8 | 27.4880437 | 0.67913478 | 1.44457034 | 0.47012926 | 0.63826267 | 0.90599527 |
| Lrmp | 3834.16495 | -0.1622413 | 0.17283617 | -0.9387 | 0.34788479 | 0.76101142 |
| Thumpd3 | 2587.11191 | 0.27221517 | 0.28252794 | 0.9634982 | 0.33529757 | 0.75359284 |
| Kras | 862.483672 | 0.59759718 | 0.44124296 | 1.3543495 | 0.1756249 | 0.60763832 |
| Bcat1 | 454.741833 | -1.6085983 | 0.41055532 | -3.9181037 | 8.924831911 | 0.00324933 |
| Mtmr14 | 2148.17026 | 0.05649753 | 0.16793864 | 0.33641768 | 0.73655592 | 0.93798548 |
| Ogg1 | 203.094275 | 0.30268438 | 0.4502918 | 0.67219609 | 0.50145887 | 0.84968141 |
| Camk1 | 6624.73175 | 1.28670543 | 0.28960467 | 4.44297188 | 8.872476614 | 0.00047173 |
| Etnk1 | 476.642887 | -0.2494666 | 0.38852686 | -0.6420833 | 0.52081911 | 0.8576517 |
| Ttll3 | 9.71625685 | -0.6135831 | 1.5369192 | -0.3992293 | 0.68972427 | 0.92470267 |

| | | | | | | |
|---|---|---|---|---|---|---|
| C2cd5 | 473.287719 | -0.4195242 | 0.31696994 | -1.3235457 | 0.18565398 | 0.61781966 |
| Il17rc | 19.4122301 | 0.79584199 | 1.22584615 | 0.64921849 | 0.51619716 | 0.85627177 |
| Cmas | 276.776196 | -0.2210598 | 0.417809 | -0.5290929 | 0.59674099 | 0.88891958 |
| Creld1 | 415.171996 | 0.13965914 | 0.44483099 | 0.31396001 | 0.75355141 | 0.94018119 |
| Emc3 | 3743.34502 | 0.20650859 | 0.15637585 | 1.32059133 | 0.18663767 | 0.61862792 |
| Itpr2 | 133.465503 | 0.23097487 | 0.40778927 | 0.56640742 | 0.57111685 | 0.88035336 |
| Med21 | 3451.31095 | 0.58530611 | 0.4182964 | 1.39926166 | 0.16173453 | 0.59254223 |
| Sec13 | 11463.496 | 0.3696917 | 0.35074751 | 1.05401088 | 0.29187793 | 0.72575203 |
| Ccdc91 | 574.108637 | 0.0085364 | 0.20833706 | 0.04097398 | 0.96731664 | 0.99443087 |
| Far2 | 14.1748866 | -1.2043126 | 1.9848652 | -0.6067478 | 0.54401828 | 0.86947682 |
| Ergic2 | 3775.5184 | 0.17930468 | 0.26267586 | 0.68260811 | 0.49485451 | 0.84678075 |
| Caprin2 | 154.620514 | 1.60306378 | 0.45910338 | 3.49172722 | 0.00047991 | 0.01301711 |
| Dennd5b | 402.370049 | 0.09287217 | 0.46668146 | 0.19900549 | 0.84225845 | 0.96765585 |
| Atg7 | 2545.92311 | -0.1598846 | 0.23598678 | -0.6775152 | 0.49807913 | 0.84860011 |
| Vgll4 | 825.186797 | 0.17366289 | 0.33030072 | 0.52577205 | 0.5990466 | 0.88987699 |
| Tamm41 | 1356.81787 | -0.2333505 | 0.22327742 | -1.0451148 | 0.29596993 | 0.72859877 |
| Cand2 | 545.402766 | -0.055861 | 0.30513326 | -0.1830709 | 0.85474242 | 0.96862197 |
| Efcab12 | 9.74721452 | 1.94985155 | 2.59087546 | 0.75258405 | 0.4516999 | 0.82570905 |
| Mbd4 | 708.258698 | -0.014043 | 0.27504193 | -0.0510578 | 0.9592795 | 0.99271349 |
| Ift122 | 231.281311 | 0.43423501 | 0.41001401 | 1.05907361 | 0.28956626 | 0.72395811 |
| Necap1 | 3821.7953 | 0.12678421 | 0.234063 | 0.54166703 | 0.5880479 | 0.88690478 |
| Ing4 | 1911.346 | 0.70864577 | 0.29004996 | 2.44318516 | 0.01455827 | 0.16582055 |
| Mrpl51 | 3625.96762 | 0.05473222 | 0.20606829 | 0.26560234 | 0.79054548 | 0.95261385 |
| Vamp1 | 123.102004 | 1.40574423 | 0.65414376 | 2.14898364 | 0.0316357 | 0.26539272 |
| Ltbr | 1431.15022 | 0.09735951 | 0.20331044 | 0.47887115 | 0.6320303 | 0.90405469 |
| Tnfrsf1a | 2996.65883 | -0.0158542 | 0.27228296 | -0.0582269 | 0.9535679 | 0.99149988 |
| Cd9 | 2995.04069 | -0.0358317 | 0.36132831 | -0.0991667 | 0.92100591 | 0.9839023 |
| Dyrk4 | 5.41706902 | 0.99390413 | 2.91251867 | 0.34125245 | 0.73291354 | 0.93728136 |
| Rad51ap1 | 2020.90681 | 0.21857815 | 0.22020683 | 0.99260387 | 0.32090304 | 0.74413737 |
| D6Wsu163e | 2897.08269 | 0.02544156 | 0.19286867 | 0.13191129 | 0.89505446 | 0.9794437 |
| Tspan9 | 140.213264 | 0.16481675 | 0.63066711 | 0.26133716 | 0.79383251 | 0.95310033 |
| Fkbp4 | 18991.3566 | -0.3358967 | 0.16230141 | -2.0695861 | 0.03849112 | 0.29651254 |
| Clec2i | 72.4253816 | 1.48403059 | 0.58398314 | 2.54122165 | 0.01104659 | 0.13649787 |
| Strn4 | 651.163766 | -0.0391355 | 0.25912164 | -0.1510312 | 0.87995109 | 0.97541902 |
| Mzf1 | 12.3633301 | 1.26752886 | 2.11082832 | 0.60048885 | 0.54818049 | 0.87059427 |
| Zfp606 | 271.247893 | -0.0191152 | 0.47892463 | -0.0399127 | 0.96816272 | 0.99443087 |
| Zik1 | 129.645122 | 0.59656305 | 0.72229684 | 0.82592504 | 0.40884661 | 0.80100711 |
| Mark4 | 234.773347 | 0.10493911 | 0.42833903 | 0.24499077 | 0.80646357 | 0.95741157 |
| Ckm | 10.0072068 | -0.4998249 | 1.19410439 | -0.4185772 | 0.67552516 | 0.91851958 |
| Ercc2 | 1448.05814 | -0.3229189 | 0.32983122 | -0.9790427 | 0.32755887 | 0.74829972 |
| Vasp | 1292.04341 | -0.6983906 | 0.19402699 | -3.5994506 | 0.00031889 | 0.00926599 |
| Qpctl | 772.974102 | 0.12443603 | 0.3220976 | 0.38633019 | 0.69925215 | 0.92669304 |
| Dmpk | 60.9091852 | 0.6674795 | 0.72254813 | 0.92378553 | 0.35559799 | 0.7647925 |
| Dmwd | 29.5580196 | 1.08773044 | 0.88332295 | 1.23140743 | 0.21817052 | 0.65150232 |
| Pglyrp1 | 45.6630994 | -6.1087173 | 1.14108903 | -5.3534099 | 8.631203558 | 8.584188059 |
| Pdcd5 | 3312.81861 | 0.18472649 | 0.3037845 | 0.60808397 | 0.54313178 | 0.86919453 |

| | | | | | | |
|---|---|---|---|---|---|---|
| Uri1 | 1492.54675 | -0.0197706 | 0.3308037 | -0.0597654 | 0.95234251 | 0.9914597 |
| Pop4 | 2169.89874 | -0.0074192 | 0.23995384 | -0.0309193 | 0.97533392 | 0.99450266 |
| Zfp939 | 52.8430453 | -0.8186849 | 0.85724217 | -0.9550218 | 0.33956669 | 0.75718544 |
| Lilra6 | 36.0662577 | -2.4882445 | 1.48300295 | -1.6778419 | 0.09337797 | 0.46719538 |
| Tmem238 | 68.2029448 | -0.4868649 | 0.62378054 | -0.7805068 | 0.43509265 | 0.81608225 |
| Rpl28 | 3109.81205 | -0.2217386 | 0.18379066 | -1.2064738 | 0.22763484 | 0.66143871 |
| U2af2 | 1387.95614 | -0.3378908 | 0.18605317 | -1.8160983 | 0.06935529 | 0.40672258 |
| Zfp583 | 149.284097 | -0.0271739 | 0.55561317 | -0.048908 | 0.96099259 | 0.99338983 |
| Zfp273 | 147.030909 | 0.08029322 | 0.56797067 | 0.14136861 | 0.88757875 | 0.97832097 |
| Cyfip1 | 19212.3308 | 0.34810457 | 0.24908563 | 1.39752969 | 0.16225435 | 0.59320305 |
| Herc2 | 397.829572 | 0.19811334 | 0.38290772 | 0.51739186 | 0.60488264 | 0.89258167 |
| Nipa2 | 4321.3428 | 0.31287074 | 0.20177884 | 1.55056269 | 0.12100652 | 0.52910763 |
| Psd3 | 199.25164 | 0.98786267 | 0.42164523 | 2.3428764 | 0.01913572 | 0.1953984 |
| Siglecg | 95.683331 | -1.6003074 | 0.71465032 | -2.2392874 | 0.02513722 | 0.23084234 |
| Zfp719 | 451.367273 | 0.09840903 | 0.30704057 | 0.32050824 | 0.74858309 | 0.9399191 |
| Zdhhc13 | 1584.07131 | -0.4263593 | 0.24791115 | -1.7198067 | 0.08546758 | 0.44669728 |
| Siglece | 589.417955 | 0.21607762 | 0.2747348 | 0.78649527 | 0.43157739 | 0.81342204 |
| Zfp108 | 6.05009976 | 1.24983353 | 2.83916166 | 0.44021217 | 0.65978344 | 0.91280998 |
| Faap24 | 1200.13682 | -0.2020316 | 0.25856727 | -0.7813504 | 0.43459646 | 0.81552669 |
| Gas2 | 391.734078 | 0.30236083 | 0.48628152 | 0.62178144 | 0.53408559 | 0.86520872 |
| Prmt3 | 2538.45571 | -0.2682946 | 0.1717713 | -1.5619292 | 0.11830466 | 0.52515149 |
| Asb7 | 256.966326 | 0.51992459 | 0.38353396 | 1.35561553 | 0.17522153 | 0.60743838 |
| Cers3 | 17.3784377 | 1.08567371 | 1.69301443 | 0.64126666 | 0.52134945 | 0.85802069 |
| Snrpa1 | 5825.65687 | -0.1279896 | 0.2639988 | -0.4848114 | 0.62781014 | 0.90236182 |
| Tarsl2 | 143.375416 | 0.2819084 | 0.69355623 | 0.40646798 | 0.68439878 | 0.92286709 |
| Tjp1 | 35.2655824 | 1.93217837 | 1.69801129 | 1.13790667 | 0.25515946 | 0.68966367 |
| Mphosph10 | 2638.08452 | -0.2798686 | 0.25639986 | -1.0915318 | 0.27503893 | 0.71147204 |
| Mtmr10 | 386.603175 | 0.59359045 | 0.39645481 | 1.49724616 | 0.13432922 | 0.5551518 |
| Crtc3 | 154.15194 | -0.1280577 | 0.49259094 | -0.2599676 | 0.79488878 | 0.95367917 |
| Blm | 1039.17043 | 0.15650541 | 0.28483177 | 0.54946613 | 0.5826856 | 0.88343755 |
| Furin | 194.850099 | 0.04253075 | 0.34665769 | 0.12268803 | 0.90235414 | 0.98043702 |
| Hddc3 | 102.211889 | 0.57995956 | 0.64120396 | 0.90448531 | 0.36573812 | 0.7719245 |
| Unc45a | 790.049809 | -0.5075384 | 0.22885858 | -2.2176944 | 0.02657567 | 0.23957275 |
| Vps33b | 981.244397 | -0.3426854 | 0.3525611 | -0.9719887 | 0.33105615 | 0.75011761 |
| Iqgap1 | 7406.80266 | -0.1488639 | 0.27863382 | -0.5342634 | 0.5931593 | 0.88797037 |
| Cib1 | 4514.80933 | 0.17285157 | 0.33018637 | 0.52349699 | 0.60062845 | 0.89046491 |
| Sema4b | 649.554612 | 0.29822961 | 0.2707243 | 1.10159898 | 0.27063605 | 0.70756376 |
| Idh2 | 19590.1247 | 0.32095292 | 0.14466374 | 2.21861341 | 0.02651304 | 0.23919285 |
| Pex11a | 65.4150172 | -0.2808568 | 0.62066895 | -0.4525066 | 0.65090405 | 0.91021183 |
| Nr2f2 | 36.8985297 | 2.44756133 | 0.91190825 | 2.68399954 | 0.00727472 | 0.10438985 |
| Ttc23 | 14.1606811 | 1.10804655 | 1.80010127 | 0.61554678 | 0.53819368 | 0.86663962 |
| Lrrc28 | 290.983306 | 0.04372585 | 0.38928719 | 0.11232285 | 0.91056742 | 0.98288717 |
| Mef2a | 1906.50094 | 0.43717483 | 0.24896608 | 1.75596143 | 0.079095 | 0.43547225 |
| Rab38 | 103.295502 | 0.7640602 | 0.62285947 | 1.22669757 | 0.21993628 | 0.65307784 |
| Ctsc | 105765.09 | -0.7181774 | 0.17603633 | -4.0797109 | 4.509173876 | 0.00186354 |
| Tyrobp | 20855.4605 | 0.31160833 | 0.39048382 | 0.79800575 | 0.42486716 | 0.80988459 |

| | | | | | | |
|---|---|---|---|---|---|---|
| Sipa1l3 | 40.0844614 | -0.4539242 | 1.10955413 | -0.409105 | 0.6824626 | 0.92104707 |
| Dpf1 | 18.8662422 | -0.0363921 | 1.34706707 | -0.0270158 | 0.97844715 | 0.99483019 |
| Yif1b | 3474.45199 | 0.20336786 | 0.2201166 | 0.92390967 | 0.35553335 | 0.7647925 |
| Rasgrp4 | 62.7701957 | -0.5031345 | 0.4916824 | -1.0232916 | 0.30616999 | 0.73554152 |
| Fam98c | 572.099472 | 0.08350002 | 0.26352993 | 0.31685215 | 0.75135579 | 0.94018119 |
| Psmd8 | 10003.449 | 0.17424872 | 0.20611355 | 0.8454016 | 0.39788665 | 0.79531492 |
| Ryr1 | 48.4270357 | 0.98844945 | 0.8852874 | 1.11652945 | 0.26419557 | 0.70060794 |
| Nfkbib | 1585.62569 | -0.0306966 | 0.26615827 | -0.115332 | 0.90818194 | 0.9817854 |
| Fbxo17 | 68.5017975 | 1.03636458 | 0.63786196 | 1.62474744 | 0.10421638 | 0.49431405 |
| Pak4 | 819.890574 | 0.03664848 | 0.38369065 | 0.0955157 | 0.92390522 | 0.9849488 |
| Psmc4 | 10038.4768 | -0.0497522 | 0.24205235 | -0.2055433 | 0.8371477 | 0.96683245 |
| Zfp626 | 57.8779926 | 0.69869347 | 0.73732991 | 0.94759953 | 0.34333337 | 0.75947281 |
| Mfge8 | 583.922985 | 0.59935099 | 0.3609886 | 1.66030447 | 0.09685322 | 0.47766794 |
| Aen | 2362.58 | -0.2658184 | 0.21995578 | -1.2085082 | 0.22685185 | 0.66015092 |
| Det1 | 426.553229 | -0.1812769 | 0.3079431 | -0.58867 | 0.55608269 | 0.87268244 |
| Mrps11 | 1608.82954 | 0.01160562 | 0.32321905 | 0.03590635 | 0.97135704 | 0.99443087 |
| Mrpl46 | 2949.57955 | 0.36529441 | 0.36294746 | 1.00646637 | 0.31419128 | 0.74065408 |
| Ccdc90b | 1276.83105 | 0.73264456 | 0.45352385 | 1.61544881 | 0.10621354 | 0.49777747 |
| Tmem126b | 1893.60594 | 0.58416 | 0.24744341 | 2.36078222 | 0.01823644 | 0.19092573 |
| Tmem126a | 2622.56705 | 0.66828071 | 0.35305317 | 1.89286138 | 0.05837631 | 0.37135588 |
| Eed | 3081.56568 | 0.13426064 | 0.25891905 | 0.51854291 | 0.60407953 | 0.89257579 |
| Zfand6 | 3811.41013 | 0.53239098 | 0.2257048 | 2.35879332 | 0.01833446 | 0.19092573 |
| Fah | 69.3564708 | 1.4205596 | 0.73707273 | 1.92729909 | 0.05394237 | 0.35989907 |
| Ddias | 1075.65015 | -0.0466455 | 0.19644817 | -0.2374443 | 0.81231213 | 0.95967622 |
| Rab30 | 7.45978493 | 0.0096446 | 1.89614184 | 0.00508643 | 0.99594163 | 0.99884675 |
| Ndufc2 | 3568.5944 | 0.22463778 | 0.31736834 | 0.70781407 | 0.47906073 | 0.83869135 |
| Anapc15 | 3058.26088 | 0.10800234 | 0.22864819 | 0.47235163 | 0.63667583 | 0.90542378 |
| Coq7 | 2607.78605 | 0.0646477 | 0.20073211 | 0.32205958 | 0.74740756 | 0.9399191 |
| Arl6ip1 | 21351.8104 | 0.22699423 | 0.16291073 | 1.39336574 | 0.16350922 | 0.59494596 |
| Smg1 | 557.101659 | 0.76740796 | 0.49078177 | 1.56364398 | 0.1179012 | 0.52446741 |
| Nucb2 | 744.747868 | 0.12352068 | 0.20732439 | 0.5957846 | 0.55131913 | 0.87093412 |
| Pik3c2a | 296.784183 | -0.189711 | 0.26979783 | -0.7031599 | 0.4819561 | 0.84038934 |
| Ipo5 | 10732.9193 | -0.3400692 | 0.19081773 | -1.7821678 | 0.07472187 | 0.42093241 |
| 1110004F10I | 8394.17023 | 0.25526828 | 0.29754296 | 0.85792073 | 0.39093623 | 0.7899826 |
| Cyp2r1 | 70.0832954 | 1.84170479 | 0.71475211 | 2.57670423 | 0.00997473 | 0.12912257 |
| Pde3b | 548.927107 | 0.20160183 | 0.43151651 | 0.46719378 | 0.64036125 | 0.90677142 |
| Mylpf | 64.4268072 | 0.15122495 | 0.66081093 | 0.22884754 | 0.81898742 | 0.96105733 |
| Qprt | 153.17956 | 0.05686613 | 0.49105685 | 0.11580356 | 0.90780823 | 0.9817854 |
| Kif22 | 5764.09083 | -0.2388729 | 0.15297228 | -1.5615437 | 0.11839553 | 0.52515149 |
| Maz | 1114.63572 | -0.0792113 | 0.2377526 | -0.3331671 | 0.73900815 | 0.93800224 |
| Pagr1a | 186.548946 | -0.1842661 | 0.46064047 | -0.4000216 | 0.68914061 | 0.92470267 |
| Mvp | 2742.67668 | 0.34957449 | 0.25555812 | 1.36788645 | 0.17134762 | 0.60430173 |
| Cdipt | 3344.66475 | -0.3177175 | 0.23241852 | -1.3670058 | 0.1716235 | 0.60430173 |
| Kctd13 | 505.009125 | -0.1641016 | 0.34350664 | -0.4777246 | 0.63284624 | 0.90430301 |
| Stard10 | 10.0183533 | -1.0280879 | 1.1961344 | -0.8595087 | 0.39005995 | 0.78985201 |
| Ino80e | 2982.05102 | -0.5691199 | 0.24458087 | -2.3269192 | 0.01996957 | 0.19990812 |

| | | | | | | |
|---|---|---|---|---|---|---|
| Fchsd2 | 1609.59856 | 0.51297905 | 0.3253931 | 1.57649025 | 0.11491286 | 0.51833667 |
| Aldoa | 89663.188 | -0.1535468 | 0.15891994 | -0.9661899 | 0.33394919 | 0.75320272 |
| Ppp4c | 10568.6018 | -0.0733801 | 0.16723686 | -0.4387797 | 0.66082119 | 0.91322024 |
| Gdpd3 | 21.5084101 | 1.54295572 | 1.12665998 | 1.36949545 | 0.17084446 | 0.60430173 |
| Rab6a | 2455.71461 | -0.1425011 | 0.22955149 | -0.6207807 | 0.53474392 | 0.8652846 |
| Mrpl48 | 2780.54406 | 0.45932184 | 0.40835851 | 1.12480046 | 0.2606736 | 0.69685755 |
| Coro1a | 64461.034 | -0.5186559 | 0.19680466 | -2.6353839 | 0.00840422 | 0.11512741 |
| Dnajb13 | 30.3679634 | 0.45038923 | 0.79120568 | 0.56924419 | 0.56919044 | 0.87961466 |
| Sult1a1 | 10.4557188 | -0.8491792 | 1.70314544 | -0.4985947 | 0.61806497 | 0.89878056 |
| Sgf29 | 1452.99343 | -0.4054852 | 0.26106521 | -1.553195 | 0.12037656 | 0.52825513 |
| Nupr1 | 1426.28891 | 0.16664242 | 0.19709455 | 0.84549481 | 0.39783463 | 0.79531492 |
| Ppme1 | 1890.54807 | -0.4748377 | 0.20339398 | -2.3345711 | 0.01956584 | 0.1981365 |
| Cln3 | 2245.48119 | 0.78635232 | 0.20408931 | 3.85298146 | 0.00011669 | 0.00411368 |
| Nfatc2ip | 70.1375846 | -0.1235517 | 0.72788699 | -0.1697402 | 0.86521446 | 0.9705375 |
| Lipt2 | 413.167945 | 0.28327766 | 0.41690795 | 0.67947292 | 0.49683826 | 0.84814011 |
| Pold3 | 4182.90395 | -0.0876971 | 0.16555108 | -0.5297282 | 0.59630039 | 0.88873045 |
| Rabep2 | 661.894093 | -0.0073677 | 0.25311132 | -0.0291086 | 0.976778 | 0.99483019 |
| Pgm2l1 | 169.875015 | 0.2900084 | 0.50806649 | 0.57080796 | 0.56812983 | 0.87929304 |
| Sh2b1 | 541.233166 | -0.1846358 | 0.55243779 | -0.33422 | 0.73821356 | 0.93800224 |
| Slco2b1 | 6795.40878 | 0.38270937 | 0.22017941 | 1.73817059 | 0.08218076 | 0.44131573 |
| Eif3c | 11688.2752 | -0.3609779 | 0.20688435 | -1.7448297 | 0.08101452 | 0.4392955 |
| Spns1 | 3014.76037 | 0.09798085 | 0.18536918 | 0.52857138 | 0.59710282 | 0.88891958 |
| Lat | 20.5084382 | 0.60707979 | 1.41564871 | 0.42883505 | 0.66804327 | 0.91595755 |
| Rps3 | 44396.9077 | 0.43705466 | 0.43524666 | 1.00415396 | 0.31530441 | 0.741611 |
| Il21r | 3411.58796 | 0.44601007 | 0.21431207 | 2.08112438 | 0.03742252 | 0.29121483 |
| Dgat2 | 359.706452 | -1.3422037 | 0.34235276 | -3.9205284 | 8.835502585 | 0.00322641 |
| Il4ra | 11714.0503 | 0.65524253 | 0.24275317 | 2.69921303 | 0.00695037 | 0.10170316 |
| Nsmce1 | 5472.29628 | 0.69774775 | 0.40790208 | 1.71057661 | 0.0871593 | 0.45109601 |
| Psma1 | 21828.443 | 0.07377345 | 0.26580507 | 0.27754719 | 0.78135998 | 0.9500424 |
| Kdm8 | 617.214924 | -0.4234871 | 0.22269928 | -1.9016098 | 0.05722219 | 0.36783974 |
| Thap12 | 424.95666 | -0.2246578 | 0.31762619 | -0.7073025 | 0.47937851 | 0.8387067 |
| Copb1 | 8555.32268 | 0.27447339 | 0.18029632 | 1.52234604 | 0.1279224 | 0.54260565 |
| Far1 | 4831.4535 | -0.0970185 | 0.14825187 | -0.6544167 | 0.51284337 | 0.85459923 |
| Acer3 | 3111.1061 | -0.0275472 | 0.27729661 | -0.0993421 | 0.92086667 | 0.9839023 |
| Myo7a | 3244.36465 | 0.15789281 | 0.28613489 | 0.55181249 | 0.58107683 | 0.88314588 |
| Lcmt1 | 963.398194 | -0.0024543 | 0.21271925 | -0.011538 | 0.99079424 | 0.9966603 |
| Arhgap17 | 1733.94336 | -0.0763936 | 0.27613878 | -0.2766492 | 0.78204951 | 0.95004728 |
| Disp1 | 220.774276 | 0.25698961 | 0.52480614 | 0.48968483 | 0.62435694 | 0.90046668 |
| Slc5a11 | 26.8727855 | 0.44846026 | 1.9016511 | 0.23582678 | 0.81356709 | 0.95988497 |
| Parva | 31.4379082 | 2.14361851 | 1.77076876 | 1.21055812 | 0.2260648 | 0.65937941 |
| Pak1 | 3335.88831 | -0.6254804 | 0.21053244 | -2.9709456 | 0.00296884 | 0.0534959 |
| Rbbp6 | 506.577132 | -0.3192201 | 0.50869641 | -0.6275257 | 0.53031471 | 0.86349351 |
| BC017158 | 1085.33673 | -0.0093904 | 0.23708954 | -0.0396068 | 0.96840661 | 0.99443087 |
| Itgam | 2988.41647 | -1.3161281 | 0.26503964 | -4.9657782 | 6.842612243 | 5.264507897 |
| Lyve1 | 2830.05099 | 4.85210617 | 0.61339312 | 7.91027163 | 2.568278960 | 1.083370914 |
| Rnf141 | 5626.33664 | 1.47831873 | 0.25163883 | 5.87476407 | 4.234459898 | 6.815808939 |

| | | | | | | |
|---|---|---|---|---|---|---|
| Adm | 39.9299478 | 1.56065421 | 1.60922912 | 0.9698148 | 0.33213882 | 0.75130439 |
| Pycard | 7510.59175 | 0.16782956 | 0.43184072 | 0.38863765 | 0.69754421 | 0.92667877 |
| Fus | 2404.02805 | -0.2413317 | 0.26403107 | -0.9140277 | 0.36070231 | 0.76887042 |
| Tead2 | 42.2501275 | 1.59827095 | 0.81327205 | 1.96523531 | 0.04938701 | 0.34139551 |
| Cd37 | 353.487412 | -1.1497687 | 0.3390935 | -3.390713 | 0.00069711 | 0.01769244 |
| Kat8 | 859.452466 | 0.14735783 | 0.34771972 | 0.42378336 | 0.67172381 | 0.91719387 |
| Bckdk | 5320.8452 | 0.03660645 | 0.16542426 | 0.22128829 | 0.82486797 | 0.96312579 |
| Stx4a | 293.087234 | 0.47031689 | 0.40142537 | 1.17161724 | 0.24135076 | 0.67708722 |
| Fbxl19 | 42.5405097 | -1.9464508 | 0.79726103 | -2.4414222 | 0.01462954 | 0.16647734 |
| Bcl7c | 123.581023 | 0.71959218 | 0.41684476 | 1.72628338 | 0.08429644 | 0.44526833 |
| Phkg2 | 1241.21986 | -0.1772645 | 0.19964069 | -0.8879179 | 0.37458493 | 0.77797919 |
| Rnf40 | 1612.58789 | -0.2072109 | 0.23101081 | -0.896975 | 0.36973225 | 0.77508072 |
| Prr14 | 1247.081 | -0.0768313 | 0.29189976 | -0.2632111 | 0.79238787 | 0.95297175 |
| 9130019O22 | 161.047664 | 0.23510614 | 0.60872545 | 0.38622689 | 0.69932864 | 0.92669304 |
| Nucb1 | 4166.41842 | 0.33134054 | 0.22797395 | 1.45341405 | 0.14610883 | 0.5731306 |
| Bcat2 | 3206.41049 | 0.17385202 | 0.21104916 | 0.8237513 | 0.41008088 | 0.80117888 |
| Itgal | 2027.83989 | -0.3177621 | 0.28634088 | -1.1097337 | 0.26711378 | 0.70405417 |
| Nomo1 | 1977.75602 | -0.7678177 | 0.19244588 | -3.9897851 | 6.613318800 | 0.00255207 |
| Sergef | 771.167928 | -0.0139064 | 0.20376217 | -0.0682484 | 0.94558789 | 0.99019648 |
| Lamtor1 | 7854.7771 | 0.4741023 | 0.26354427 | 1.79894746 | 0.07202699 | 0.41303124 |
| Rgs10 | 2321.05978 | 0.70389448 | 0.35520234 | 1.98167184 | 0.04751598 | 0.33637905 |
| Tial1 | 694.797699 | 0.07276698 | 0.33286446 | 0.21860844 | 0.82695508 | 0.96379955 |
| Bag3 | 5222.59417 | 0.10813686 | 0.34893376 | 0.30990655 | 0.75663202 | 0.94102069 |
| Ate1 | 1036.20573 | -0.18481 | 0.2862599 | -0.6456022 | 0.51853704 | 0.8570838 |
| Tacc2 | 136.123529 | 0.7403589 | 0.49513625 | 1.49526297 | 0.13484583 | 0.55634704 |
| Acadsb | 1136.11259 | 0.2819972 | 0.31819915 | 0.88622862 | 0.37549435 | 0.77815049 |
| Plk1 | 7497.89024 | -0.4025742 | 0.17537916 | -2.2954505 | 0.02170732 | 0.21192788 |
| Dctn5 | 6022.76754 | 0.10309471 | 0.16310834 | 0.6320628 | 0.52734584 | 0.86151465 |
| Ndufab1 | 5757.65338 | 0.05908389 | 0.27438436 | 0.21533258 | 0.82950803 | 0.96462215 |
| Ubfd1 | 1949.13675 | 0.4086034 | 0.50207871 | 0.81382339 | 0.41574613 | 0.80433694 |
| Ears2 | 589.315087 | -0.4816237 | 0.27027497 | -1.7819765 | 0.07475306 | 0.42093241 |
| Gga2 | 3244.24262 | 0.65500536 | 0.33871776 | 1.93377918 | 0.05314028 | 0.35678655 |
| Mettl9 | 2943.03404 | -0.0276387 | 0.17456533 | -0.1583286 | 0.87419784 | 0.97351499 |
| Cdr2 | 48.6549498 | 0.03588743 | 0.65110954 | 0.05511735 | 0.95604497 | 0.99215259 |
| Mrpl17 | 6801.56414 | 0.34752855 | 0.45967719 | 0.7560274 | 0.44963277 | 0.82484234 |
| Polr3e | 2671.58203 | -0.2002241 | 0.23135454 | -0.8654425 | 0.38679595 | 0.78911204 |
| Arfip2 | 835.660927 | -0.4507169 | 0.30482223 | -1.4786222 | 0.13924132 | 0.56308728 |
| Uqcrc2 | 11803.6024 | 0.28509343 | 0.34745136 | 0.82052762 | 0.41191539 | 0.80245328 |
| Rrp8 | 1946.2278 | -0.5318216 | 0.25432147 | -2.0911391 | 0.0365156 | 0.28679157 |
| Ilk | 9962.81528 | 0.48124508 | 0.23275967 | 2.06756211 | 0.03868122 | 0.29722823 |
| Tpp1 | 7658.75764 | 0.8332678 | 0.231338 | 3.60194951 | 0.00031584 | 0.00922273 |
| Tmem159 | 792.906256 | 0.3210523 | 0.31420885 | 1.02177994 | 0.30688507 | 0.73595594 |
| Trim30a | 2091.32907 | 0.17996179 | 0.30822237 | 0.58386997 | 0.5593078 | 0.87397022 |
| Lyrm1 | 203.199022 | 0.37005605 | 0.53049257 | 0.69757066 | 0.48544574 | 0.84183502 |
| Rexo5 | 231.358655 | 0.29972952 | 0.40601462 | 0.73822346 | 0.46037867 | 0.82987212 |
| Eri2 | 774.348874 | -0.2693659 | 0.29830686 | -0.9029824 | 0.36653522 | 0.77259343 |

| | | | | | | |
|---|---|---|---|---|---|---|
| Chst15 | 188.436384 | -1.1464532 | 0.3809833 | -3.0091955 | 0.00261941 | 0.04847683 |
| Oat | 10197.8671 | 0.20954692 | 0.15865891 | 1.32073841 | 0.1865886 | 0.61862792 |
| Thumpd1 | 5130.06568 | -0.2170423 | 0.20098121 | -1.0799135 | 0.28018069 | 0.71532524 |
| Lhpp | 716.517524 | -0.1047051 | 0.28953716 | -0.3616291 | 0.71762919 | 0.93275665 |
| Fam53b | 597.271109 | 0.66561275 | 0.31182984 | 2.13453836 | 0.03279874 | 0.26946068 |
| Eef1akmt2 | 950.042755 | -0.0717377 | 0.23907739 | -0.3000606 | 0.76413094 | 0.94300309 |
| Abraxas2 | 2580.27239 | 0.02685373 | 0.20356308 | 0.13191849 | 0.89504876 | 0.9794437 |
| Zranb1 | 236.330472 | -0.238391 | 0.54875077 | -0.434425 | 0.66397988 | 0.91399629 |
| Ctbp2 | 516.974777 | -0.262422 | 0.27308077 | -0.9609686 | 0.33656797 | 0.75494233 |
| Rrm1 | 28718.2103 | -0.1988746 | 0.14808806 | -1.342948 | 0.17928882 | 0.60992118 |
| Uros | 890.55092 | 0.2981076 | 0.39407171 | 0.75648059 | 0.4493611 | 0.82484234 |
| Knop1 | 2379.93757 | -0.1267014 | 0.17184004 | -0.7373217 | 0.46092673 | 0.82987212 |
| Vps35l | 5542.10853 | 0.28795195 | 0.21346322 | 1.34895347 | 0.17735191 | 0.60839763 |
| Bccip | 3792.20261 | -0.3045645 | 0.1939732 | -1.5701368 | 0.11638329 | 0.521508 |
| Dhx32 | 2010.48934 | 0.61415153 | 0.15926767 | 3.85609676 | 0.00011521 | 0.00407337 |
| Stim1 | 1470.41701 | 0.25712794 | 0.26289879 | 0.97804916 | 0.32805001 | 0.74843736 |
| Pgap2 | 1485.85494 | -0.3573144 | 0.19466424 | -1.835542 | 0.06642542 | 0.39699582 |
| Mki67 | 2828.09644 | -0.222674 | 0.39578465 | -0.562614 | 0.57369777 | 0.8805577 |
| Atp6ap2 | 11020.0823 | 0.53111288 | 0.13676197 | 3.88348367 | 0.00010297 | 0.00368315 |
| Usp9x | 1775.70459 | 0.28307619 | 0.28099561 | 1.00740433 | 0.31374051 | 0.74065408 |
| Cask | 313.860914 | -0.4274535 | 0.37734898 | -1.1327804 | 0.25730647 | 0.69274795 |
| Swap70 | 4559.32105 | 0.16287498 | 0.22696881 | 0.71760951 | 0.47299809 | 0.83483892 |
| Wee1 | 1212.51263 | -0.0626281 | 0.48430417 | -0.1293157 | 0.89710787 | 0.97984623 |
| Tmem9b | 3765.45648 | 0.22144708 | 0.15896693 | 1.39303864 | 0.1636081 | 0.59512874 |
| Akip1 | 1289.23956 | 0.19849307 | 0.2236791 | 0.88740109 | 0.374863 | 0.77815049 |
| St5 | 457.36207 | 0.59322442 | 0.3875379 | 1.53075199 | 0.1258307 | 0.5378361 |
| Eif3f | 7318.80246 | -0.3367128 | 0.16093155 | -2.0922731 | 0.0364141 | 0.28646536 |
| Ndufb11 | 3522.48305 | 0.1863592 | 0.31862057 | 0.58489382 | 0.55861912 | 0.87330194 |
| Rbm10 | 2829.09231 | -0.3514478 | 0.15420423 | -2.2791061 | 0.02266076 | 0.21589491 |
| Cdk16 | 2184.14554 | 0.20271304 | 0.26238042 | 0.77259212 | 0.43976381 | 0.81963526 |
| Usp11 | 745.820538 | -0.4092544 | 0.38243978 | -1.0701146 | 0.28456771 | 0.72013173 |
| Glrx3 | 11762.8881 | 0.31040681 | 0.37628158 | 0.82493224 | 0.40941006 | 0.80100711 |
| LTO1 | 737.478859 | 0.46649018 | 0.21198538 | 2.20057715 | 0.02776597 | 0.24720607 |
| Fadd | 860.556228 | 0.12998478 | 0.21488044 | 0.60491676 | 0.54523431 | 0.86970532 |
| Cttn | 609.039705 | 0.95374856 | 0.28443024 | 3.35318971 | 0.00079886 | 0.01966287 |
| Nadsyn1 | 360.66435 | -0.1883348 | 0.37781581 | -0.498483 | 0.61814362 | 0.89878056 |
| Dock11 | 1184.58953 | -0.0850788 | 0.26543502 | -0.320526 | 0.74856966 | 0.9399191 |
| Cul4b | 1301.77559 | -0.394582 | 0.22585692 | -1.7470442 | 0.08062968 | 0.43854735 |
| Tnni2 | 802.946463 | 0.01135809 | 0.42449081 | 0.02675697 | 0.97865357 | 0.99483019 |
| Sash3 | 8620.81259 | -0.1953292 | 0.14194551 | -1.3760861 | 0.16879497 | 0.60312169 |
| Elf4 | 914.048823 | -0.2931024 | 0.48697014 | -0.60189 | 0.54724739 | 0.87008334 |
| Rab33a | 8.27886633 | 0.746065 | 2.16747466 | 0.34420933 | 0.73068887 | 0.93653187 |
| Slc25a14 | 333.959928 | 1.10456217 | 0.50161777 | 2.20199971 | 0.02766533 | 0.24666908 |
| Rbmx2 | 394.507123 | -0.2538674 | 0.28560596 | -0.888873 | 0.37407134 | 0.77748705 |
| Enox2 | 625.352151 | 0.59137661 | 0.38038491 | 1.55467949 | 0.12002242 | 0.52767297 |
| Stk26 | 92.6235491 | -2.1859094 | 0.64641593 | -3.3815835 | 0.00072069 | 0.01810316 |

| | | | | | | |
|---|---|---|---|---|---|---|
| Gpc4 | 14.8914579 | 0.80262385 | 2.56330888 | 0.31312022 | 0.75418932 | 0.94018119 |
| 3830403N18 | 228.459001 | 0.31252931 | 0.50109601 | 0.62369147 | 0.53283023 | 0.8646427 |
| Slc9a9 | 1998.91338 | 1.40299097 | 0.29153073 | 4.81249768 | 1.490556002 | 0.00010074 |
| Arhgef6 | 4035.78178 | 0.04261314 | 0.27803717 | 0.15326419 | 0.87818994 | 0.97503155 |
| Rbmx | 1234.56192 | -0.2555575 | 0.19491657 | -1.3111122 | 0.18981985 | 0.62020999 |
| Fgf13 | 59.1923093 | 0.60032622 | 0.94528887 | 0.63507171 | 0.52538165 | 0.86048918 |
| F9 | 21.2515163 | 0.57575469 | 1.42181274 | 0.4049441 | 0.6855186 | 0.92314423 |
| Cacna1f | 200.622385 | 0.409958 | 0.38943356 | 1.05270333 | 0.29247698 | 0.72584701 |
| Ccdc22 | 3263.60225 | 0.12532197 | 0.20223022 | 0.61969949 | 0.53545565 | 0.86539693 |
| Syp | 20.2718421 | 1.14696684 | 1.27110125 | 0.90234105 | 0.36687572 | 0.77272568 |
| Prickle3 | 76.2017089 | 1.15372032 | 0.60243846 | 1.9150841 | 0.05548178 | 0.3627292 |
| Plp2 | 744.827866 | -0.3074617 | 0.22964678 | -1.3388459 | 0.18062084 | 0.61122544 |
| Gpkow | 2007.00614 | 0.09968538 | 0.20184961 | 0.49385967 | 0.62140529 | 0.89912494 |
| Praf2 | 445.822055 | -0.0635192 | 0.32710169 | -0.1941881 | 0.84602862 | 0.96765585 |
| Ccdc120 | 17.3567918 | 0.12104085 | 1.53736447 | 0.0787327 | 0.93724524 | 0.98845735 |
| Gripap1 | 889.939846 | 0.27969265 | 0.26259203 | 1.0651224 | 0.28682055 | 0.72260415 |
| Otud5 | 749.036329 | -0.1855807 | 0.19264405 | -0.9633349 | 0.33537947 | 0.75359284 |
| Pim2 | 407.550046 | -0.0116384 | 0.27456934 | -0.0423878 | 0.96618951 | 0.99443087 |
| Slc35a2 | 551.529193 | -0.1955904 | 0.26462959 | -0.7391099 | 0.45984026 | 0.82987212 |
| Pqbp1 | 2082.5577 | -0.2006038 | 0.17092575 | -1.1736314 | 0.24054269 | 0.67567365 |
| Timm17b | 1620.9866 | 0.35659077 | 0.29350341 | 1.21494595 | 0.22438671 | 0.65793926 |
| Hdac6 | 1002.19158 | -0.2451852 | 0.24853795 | -0.9865101 | 0.32388286 | 0.74583177 |
| Gata1 | 17.3907689 | -2.7731032 | 1.56114194 | -1.77633 | 0.07567854 | 0.42369594 |
| Was | 1688.98338 | -0.4693661 | 0.20373835 | -2.3037691 | 0.02123561 | 0.20848729 |
| Wdr13 | 477.340731 | 0.0499098 | 0.45366861 | 0.11001378 | 0.91239845 | 0.98292561 |
| Rbm3 | 17100.3446 | 0.22443768 | 0.34621615 | 0.64825885 | 0.51681754 | 0.85673255 |
| Ebp | 3399.20258 | 0.12526345 | 0.24022445 | 0.52144338 | 0.60205794 | 0.89110403 |
| Ftsj1 | 1421.98971 | 0.03473779 | 0.32398745 | 0.10721957 | 0.91461479 | 0.98347559 |
| Rpgr | 312.524756 | 0.3076379 | 0.33227695 | 0.92584785 | 0.35452505 | 0.76419569 |
| Dynlt3 | 2292.10481 | 0.63991097 | 0.36557501 | 1.75042315 | 0.08004532 | 0.43723638 |
| Vbp1 | 4759.4833 | 0.03718963 | 0.30589944 | 0.12157469 | 0.90323586 | 0.98043702 |
| Fundc2 | 2402.97696 | 0.60680215 | 0.43638072 | 1.39053381 | 0.16436684 | 0.5959394 |
| Mtcp1 | 67.9886927 | -0.0917871 | 0.69095802 | -0.1328403 | 0.89431969 | 0.97941957 |
| Brcc3 | 3319.32284 | 0.13514899 | 0.31473711 | 0.42940278 | 0.66763013 | 0.91595755 |
| Msn | 20120.6331 | -0.3396724 | 0.20423126 | -1.6631752 | 0.09627737 | 0.47586307 |
| Gpr165 | 24.7519456 | 1.62955951 | 1.83107869 | 0.8899451 | 0.37349537 | 0.77748705 |
| Ophn1 | 895.334041 | 1.02474353 | 0.32732917 | 3.13062083 | 0.00174437 | 0.03615604 |
| Stard8 | 5767.6235 | 1.82213404 | 0.36936575 | 4.93314288 | 8.091692026 | 6.072740402 |
| Efnb1 | 28.1186211 | 0.46681721 | 1.42493611 | 0.32760571 | 0.7432098 | 0.93856621 |
| Awat2 | 50.8977368 | -0.9091835 | 0.84292984 | -1.0785993 | 0.2807664 | 0.7155865 |
| Igbp1 | 2507.65032 | 0.20604947 | 0.24126073 | 0.85405308 | 0.39307558 | 0.79137753 |
| Magee2 | 30.3738094 | 0.99629754 | 1.76822938 | 0.56344361 | 0.57313285 | 0.88035336 |
| Pbdc1 | 3411.81261 | -0.1428727 | 0.20449674 | -0.6986553 | 0.48476748 | 0.84163505 |
| Magee1 | 152.383118 | 0.84723443 | 0.49542689 | 1.7101099 | 0.08724555 | 0.45127901 |
| Atrx | 2404.15001 | -0.3303935 | 0.24700389 | -1.3376045 | 0.18102541 | 0.61156693 |
| Cox7b | 7412.08798 | 0.49684683 | 0.44114339 | 1.12627059 | 0.260051 | 0.69625824 |

| | | | | | | |
|---|---|---|---|---|---|---|
| Magt1 | 6272.37092 | 0.12081092 | 0.14506651 | 0.83279681 | 0.40495934 | 0.79917613 |
| Itm2a | 398.730006 | 0.5352447 | 0.59307988 | 0.90248332 | 0.36680018 | 0.7726996 |
| 2610002M06 | 631.166307 | 0.67801815 | 0.29813393 | 2.27420659 | 0.02295357 | 0.21709424 |
| Hmgn5 | 403.924799 | -0.5809038 | 0.27351022 | -2.1238833 | 0.0336799 | 0.27375828 |
| Sh3bgrl | 7584.95706 | 0.5992211 | 0.25430987 | 2.35626368 | 0.01845981 | 0.19120986 |
| Sytl4 | 15.685398 | 2.93072256 | 1.8918204 | 1.54915475 | 0.12134452 | 0.5291774 |
| Cstf2 | 3143.91751 | -0.0725155 | 0.18004357 | -0.4027664 | 0.68712007 | 0.92427909 |
| Cenpi | 1099.18517 | 0.17377754 | 0.27062623 | 0.64213118 | 0.52078801 | 0.8576517 |
| Btk | 4153.05087 | -0.142789 | 0.16820132 | -0.8489171 | 0.39592741 | 0.79400306 |
| Gla | 1686.90245 | 0.12658228 | 0.26902372 | 0.47052461 | 0.63798026 | 0.90599527 |
| Col4a5 | 16.5230357 | 0.16177012 | 1.15208897 | 0.14041461 | 0.88833242 | 0.97832097 |
| Acsl4 | 1468.70555 | -0.2120136 | 0.22696239 | -0.9341355 | 0.35023398 | 0.76221532 |
| Cdkl5 | 24.1169865 | 1.00987001 | 1.90149637 | 0.53109226 | 0.59535484 | 0.88830085 |
| Phka2 | 2063.64988 | -0.0487458 | 0.28941714 | -0.1684275 | 0.866247 | 0.97060021 |
| Slc7a3 | 11.0423275 | -0.9127812 | 2.09618207 | -0.4354494 | 0.66323625 | 0.91395288 |
| Pdha1 | 8849.62181 | -0.1789098 | 0.15161789 | -1.1800045 | 0.23799841 | 0.67403588 |
| Map3k15 | 34.9288643 | -0.4973407 | 1.64588293 | -0.3021726 | 0.7625205 | 0.94298237 |
| Il2rg | 2104.40316 | -0.1443408 | 0.19782423 | -0.7296419 | 0.46560912 | 0.83339036 |
| Rps6ka3 | 351.410762 | 0.2114599 | 0.44375085 | 0.47652844 | 0.63369795 | 0.90455392 |
| Zmym3 | 351.623368 | 0.12454055 | 0.31342857 | 0.39734907 | 0.69111007 | 0.92470267 |
| Nono | 17288.1132 | -0.2794516 | 0.16883753 | -1.6551508 | 0.09789392 | 0.47939805 |
| Taf1 | 534.398032 | -0.6447801 | 0.56521401 | -1.1407717 | 0.25396496 | 0.68763907 |
| Rps4x | 29499.9966 | -0.2040183 | 0.21602749 | -0.9444089 | 0.34496074 | 0.75993243 |
| Flna | 7461.97099 | -0.449008 | 0.27296761 | -1.644913 | 0.09998775 | 0.48325682 |
| Abcb7 | 787.167337 | -0.3497257 | 0.32635328 | -1.071617 | 0.2838921 | 0.71951357 |
| Mtm1 | 286.4062 | 0.12175858 | 0.35942048 | 0.33876363 | 0.7347878 | 0.93750666 |
| Gpm6b | 57.2844199 | 0.36938341 | 0.55400539 | 0.66675057 | 0.50493147 | 0.85034006 |
| Cetn2 | 2033.51998 | 0.18351082 | 0.30440479 | 0.6028513 | 0.54660762 | 0.86994632 |
| Nsdhl | 1490.184 | 0.13471974 | 0.24086519 | 0.55931593 | 0.57594613 | 0.88080372 |
| Hccs | 2052.16944 | -0.2497417 | 0.19719739 | -1.2664552 | 0.20535016 | 0.63913241 |
| Rbbp7 | 27446.7321 | -0.0696796 | 0.14267865 | -0.4883672 | 0.62528979 | 0.9011746 |
| Arhgap6 | 64.4948433 | -2.7852905 | 0.61074141 | -4.560507 | 5.103025620 | 0.00029446 |
| Syap1 | 897.273132 | 0.31192239 | 0.26720919 | 1.16733408 | 0.24307548 | 0.67814218 |
| Msl3 | 1840.45101 | -0.2394056 | 0.21989909 | -1.0887069 | 0.27628317 | 0.7118544 |
| Ctps2 | 3349.17195 | -0.4227268 | 0.18841495 | -2.2435946 | 0.0248585 | 0.2288851 |
| Xlr4c | 20.7463879 | 0.70956788 | 1.20186594 | 0.59038855 | 0.55493019 | 0.87189251 |
| Zfp275 | 204.92791 | -0.5047196 | 0.37232932 | -1.3555731 | 0.17523502 | 0.60743838 |
| Ap1s2 | 1873.39358 | -0.0856577 | 0.2121908 | -0.4036826 | 0.68644615 | 0.92399822 |
| Zrsr2 | 362.944201 | -0.1765041 | 0.37342693 | -0.4726604 | 0.63645547 | 0.90542378 |
| Haus7 | 1730.16927 | 0.24884507 | 0.20587569 | 1.20871519 | 0.22677229 | 0.66015092 |
| Car5b | 9.91327553 | 0.97374448 | 1.762478 | 0.55248604 | 0.5806154 | 0.88297715 |
| Bgn | 29.4147973 | 2.76891122 | 2.03459655 | 1.36091415 | 0.17354082 | 0.60653633 |
| Abcd1 | 3172.83305 | -0.2837279 | 0.22129548 | -1.2821223 | 0.19979976 | 0.63247751 |
| Pir | 42.5224888 | 1.69446 | 1.25999586 | 1.34481394 | 0.17868533 | 0.60992118 |
| Piga | 1018.12143 | 0.11734785 | 0.21169933 | 0.55431376 | 0.57936414 | 0.88286755 |
| Dusp9 | 19.3991099 | 0.62920184 | 1.02619073 | 0.61314318 | 0.53978167 | 0.86719392 |

| | | | | | | |
|---|---|---|---|---|---|---|
| Plxnb3 | 116.50152 | -0.6814369 | 0.67485113 | -1.0097588 | 0.31261084 | 0.74065408 |
| Hcfc1 | 2323.01843 | -0.4408266 | 0.38364768 | -1.1490402 | 0.25053939 | 0.6842534 |
| Renbp | 4631.66802 | 1.10758991 | 0.20129156 | 5.50241603 | 3.746216619 | 4.166133446 |
| Naa10 | 3771.71723 | -0.3444385 | 0.22249519 | -1.5480715 | 0.12160508 | 0.52958169 |
| Arhgap4 | 2928.15946 | -0.2538728 | 0.20044468 | -1.2665482 | 0.20531689 | 0.63913241 |
| Irak1 | 523.835081 | -0.442031 | 0.31055032 | -1.4233796 | 0.15462614 | 0.58343244 |
| Mecp2 | 287.469179 | -0.6777973 | 0.31155242 | -2.1755482 | 0.02958906 | 0.25669202 |
| Plxna3 | 22.996644 | -0.0273551 | 1.54084339 | -0.0177533 | 0.98583565 | 0.99568424 |
| Fam3a | 615.509209 | 0.05889683 | 0.2180238 | 0.27013944 | 0.78705298 | 0.95223213 |
| G6pdx | 13079.6286 | -0.6334391 | 0.19236016 | -3.292985 | 0.0009913 | 0.02319222 |
| Mpp1 | 8452.39244 | 1.20480442 | 0.23447132 | 5.13838711 | 2.771065818 | 2.404145259 |
| Dkc1 | 9473.80473 | -0.5226001 | 0.18799611 | -2.7798453 | 0.00543848 | 0.08518429 |
| Morf4l2 | 15932.5608 | 0.27970039 | 0.18281277 | 1.52998279 | 0.12602099 | 0.53789768 |
| Psmd10 | 2869.21042 | 0.59625041 | 0.41047648 | 1.4525812 | 0.14634007 | 0.57340745 |
| Tsc22d3 | 5020.94279 | 0.32708412 | 0.2132678 | 1.53367795 | 0.1251089 | 0.53662594 |
| Prps1 | 7678.97442 | -0.0198893 | 0.28694988 | -0.0693129 | 0.94474057 | 0.99019648 |
| Rbm41 | 77.8411644 | -0.2121753 | 0.75294978 | -0.2817922 | 0.77810288 | 0.94837927 |
| Morc4 | 212.625844 | 1.09924383 | 0.50691895 | 2.16848044 | 0.03012215 | 0.25931335 |
| Rnf128 | 343.846814 | 1.41319008 | 0.31681646 | 4.46059547 | 8.173223990 | 0.00044017 |
| Atp11a | 74.4297545 | -1.2736858 | 1.11022653 | -1.1472306 | 0.25128634 | 0.6844769 |
| F10 | 120.235875 | -5.3559262 | 0.83599486 | -6.4066497 | 1.487520505 | 3.084209888 |
| Proz | 19.4183958 | -1.6435645 | 1.24025255 | -1.3251853 | 0.18510969 | 0.61717463 |
| Cul4a | 3140.57707 | 0.22815957 | 0.21952198 | 1.03934727 | 0.29864326 | 0.73093388 |
| Lamp1 | 43501.2294 | 0.33409972 | 0.16264741 | 2.05413498 | 0.03996263 | 0.30364152 |
| Gas6 | 6953.14122 | 2.66431066 | 0.37581376 | 7.08944421 | 1.346516707 | 3.830683461 |
| Rasa3 | 5731.3148 | -0.1496037 | 0.25009878 | -0.5981785 | 0.54972085 | 0.87093412 |
| Coprs | 525.254484 | 0.92153638 | 0.30503932 | 3.02104135 | 0.00251907 | 0.04719108 |
| Angpt2 | 93.8770289 | 0.70692019 | 0.59882682 | 1.18050856 | 0.23779801 | 0.67403588 |
| Agpat5 | 3558.05675 | -0.1874328 | 0.16496374 | -1.1362058 | 0.25587042 | 0.6905302 |
| Nek3 | 87.6684303 | -0.7705297 | 0.54261842 | -1.4200213 | 0.15560148 | 0.58460471 |
| Vps36 | 3415.84045 | 0.02069868 | 0.20411615 | 0.10140636 | 0.91922788 | 0.9839023 |
| Thsd1 | 248.897748 | 0.61198176 | 0.48314122 | 1.26667264 | 0.20527237 | 0.63913241 |
| Slc25a15 | 818.462664 | -0.040751 | 0.25110848 | -0.1622845 | 0.87108183 | 0.97268995 |
| Erlin2 | 3047.88697 | 0.35339166 | 0.24206271 | 1.4599178 | 0.14431267 | 0.57027792 |
| Plpbp | 2430.77736 | 0.08337231 | 0.18718882 | 0.44539148 | 0.65603687 | 0.91165371 |
| Adgra2 | 11.3950222 | 0.52513754 | 2.11629646 | 0.24813988 | 0.80402618 | 0.9570372 |
| Brf2 | 196.801441 | -0.442014 | 0.32223448 | -1.3717154 | 0.17015205 | 0.60430173 |
| Rab11fip1 | 144.979728 | -1.5243587 | 0.74006959 | -2.0597505 | 0.03942239 | 0.3006572 |
| Eif4ebp1 | 4942.79143 | 0.01322762 | 0.17880704 | 0.07397705 | 0.94102865 | 0.98935054 |
| Ggn | 12.2433509 | 0.44283111 | 1.31838017 | 0.3358903 | 0.73695359 | 0.93800224 |
| Cd209a | 25.2586995 | -5.3115675 | 1.30484364 | -4.0706544 | 4.688126053 | 0.00191725 |
| Tnfsf13b | 20.383545 | -1.1307041 | 1.43009901 | -0.7906474 | 0.42914977 | 0.81140481 |
| Col4a1 | 57.4443792 | 1.99660335 | 1.29894368 | 1.5370977 | 0.12426939 | 0.53565448 |
| Col4a2 | 36.3269445 | 1.79587652 | 1.65436076 | 1.08554105 | 0.27768209 | 0.71286172 |
| Rab20 | 555.487836 | 0.93150398 | 0.49056891 | 1.89882391 | 0.05758763 | 0.36897087 |
| Naxd | 1675.5254 | 0.06516221 | 0.31217193 | 0.2087382 | 0.83465262 | 0.96641606 |

| | | | | | | |
|---|---|---|---|---|---|---|
| Ptpn7 | 7153.74794 | -0.5241964 | 0.20727811 | -2.5289519 | 0.01144037 | 0.14023051 |
| Ankrd10 | 2475.64045 | -0.4232708 | 0.22269349 | -1.9006877 | 0.05734293 | 0.36842228 |
| Arhgef7 | 1232.54558 | -0.0475193 | 0.39865815 | -0.1191982 | 0.9051183 | 0.9814201 |
| Leprotl1 | 1717.58246 | -0.1280371 | 0.18000828 | -0.7112842 | 0.47690811 | 0.83675398 |
| Dctn6 | 1636.84952 | -0.1307593 | 0.23027249 | -0.5678462 | 0.57013944 | 0.88006507 |
| Aga | 898.169211 | 0.32342921 | 0.32763859 | 0.98715236 | 0.32356795 | 0.74583177 |
| Dlc1 | 54.410532 | 0.17123673 | 1.36457395 | 0.12548732 | 0.90013776 | 0.98043702 |
| Eri1 | 5385.02647 | -0.3277614 | 0.17097556 | -1.9170072 | 0.05523702 | 0.36192523 |
| Tnks | 244.381859 | -0.1342979 | 0.47991323 | -0.2798378 | 0.77960194 | 0.9488468 |
| Dusp4 | 49.0219203 | 0.9044992 | 1.31173853 | 0.68954229 | 0.49048207 | 0.84472295 |
| Saraf | 5621.66686 | 0.39663649 | 0.189123 | 2.09724085 | 0.03597226 | 0.28477582 |
| Mrps31 | 939.796411 | 0.01003833 | 0.24840454 | 0.04041123 | 0.96776528 | 0.99443087 |
| Smim19 | 140.040334 | 0.17096828 | 0.4859934 | 0.35179136 | 0.72499474 | 0.93542036 |
| Polb | 1134.87692 | -0.0550842 | 0.30055356 | -0.1832757 | 0.85458169 | 0.96859981 |
| Ikbkb | 2569.11372 | -0.2108632 | 0.17980438 | -1.1727369 | 0.24090134 | 0.67637047 |
| Plat | 14.7464686 | 0.00291026 | 2.54683227 | 0.0011427 | 0.99908826 | 0.99969186 |
| Ap3m2 | 540.760386 | 0.42761918 | 0.27582741 | 1.5503143 | 0.1210661 | 0.5291181 |
| Kat6a | 499.355904 | 0.08314845 | 0.40958571 | 0.20300623 | 0.83913016 | 0.96757914 |
| Gpat4 | 4063.48549 | 0.18480057 | 0.24065702 | 0.76790018 | 0.44254649 | 0.82037752 |
| Gins4 | 3209.56798 | 0.47885033 | 0.36679506 | 1.30549829 | 0.19172323 | 0.62276959 |
| Sfrp1 | 37.0327244 | 1.8595712 | 1.51487748 | 1.22753901 | 0.21962007 | 0.65289787 |
| Adam3 | 275.435911 | 1.16172586 | 0.47122154 | 2.46534966 | 0.01368796 | 0.15826542 |
| Adam9 | 1829.8305 | 0.63321201 | 0.28384561 | 2.23083249 | 0.02569223 | 0.2336751 |
| Tm2d2 | 4238.28641 | 0.40834924 | 0.2525532 | 1.61688405 | 0.10590331 | 0.49777747 |
| Plekha2 | 4573.0828 | 0.27439493 | 0.18060218 | 1.51933348 | 0.12867858 | 0.54430326 |
| Dctd | 1429.43089 | -0.5892084 | 0.1948891 | -3.0233011 | 0.00250033 | 0.04705627 |
| Wwc2 | 284.958146 | 0.5865086 | 0.42075195 | 1.39395338 | 0.16333168 | 0.59475807 |
| Fgfr1 | 20.0633683 | 1.42774621 | 1.17543647 | 1.21465195 | 0.22449886 | 0.65811038 |
| Rwdd4a | 1645.69914 | -0.1089858 | 0.18143466 | -0.6006889 | 0.54804722 | 0.87059427 |
| Plpp5 | 821.791769 | 0.11695472 | 0.22955685 | 0.50948043 | 0.61041551 | 0.89508835 |
| Star | 36.5765086 | 0.24159774 | 0.72980553 | 0.33104399 | 0.74061126 | 0.93800224 |
| Ash2l | 4355.18869 | 0.06686774 | 0.35578468 | 0.18794442 | 0.85092022 | 0.9679302 |
| Tti2 | 635.38787 | -0.1874848 | 0.24360498 | -0.7696265 | 0.44152148 | 0.81967485 |
| Mak16 | 2015.04561 | -0.1695818 | 0.16659826 | -1.0179088 | 0.30872129 | 0.73867239 |
| Wrn | 294.667679 | -0.6295683 | 0.43025289 | -1.4632517 | 0.14339851 | 0.56846902 |
| Gsr | 2579.83165 | -1.2709212 | 0.40527159 | -3.1359741 | 0.00171284 | 0.03569542 |
| Gtf2e2 | 3803.67513 | 0.12416013 | 0.22056665 | 0.56291433 | 0.57349322 | 0.88046468 |
| Rbpms | 369.096417 | -0.5907926 | 0.4832172 | -1.2226233 | 0.221472 | 0.65504522 |
| Frg1 | 2123.96626 | -0.0547127 | 0.23848667 | -0.2294164 | 0.81854533 | 0.96105733 |
| Asah1 | 14715.9547 | 0.38846423 | 0.14446373 | 2.68900865 | 0.00716646 | 0.10311756 |
| Pcm1 | 1080.77134 | -0.5767422 | 0.32896878 | -1.7531823 | 0.07957071 | 0.4364971 |
| Fgl1 | 14.4937218 | 1.00633752 | 1.4830435 | 0.67856238 | 0.49741519 | 0.84818511 |
| Pdgfrl | 27.193862 | 1.07533477 | 1.12505454 | 0.9558068 | 0.33916985 | 0.75683748 |
| Slc7a2 | 89.9608232 | -1.2845682 | 0.58701488 | -2.1883061 | 0.02864732 | 0.25211699 |
| Vps37a | 505.498122 | 0.18448427 | 0.32826604 | 0.56199622 | 0.57411859 | 0.88059206 |
| Cnot7 | 2476.04133 | -0.1857109 | 0.21620417 | -0.8589609 | 0.39036211 | 0.7899826 |

| | | | | | | |
|---|---|---|---|---|---|---|
| Msmo1 | 4955.56441 | 0.97397379 | 0.27416421 | 3.55251976 | 0.00038156 | 0.01080469 |
| Klhl2 | 103.417066 | -0.6468343 | 0.62922171 | -1.027991 | 0.30395405 | 0.73436097 |
| Galnt7 | 1145.43564 | 0.15797429 | 0.24892503 | 0.63462597 | 0.52567238 | 0.86056156 |
| Sap30 | 1607.43278 | 0.29469638 | 0.31920327 | 0.92322482 | 0.35589006 | 0.76499791 |
| Hpgd | 3727.57679 | 0.32220023 | 0.24600242 | 1.30974413 | 0.19028241 | 0.6209089 |
| Tmem184c | 868.479821 | -0.1735852 | 0.28561303 | -0.6077636 | 0.54334428 | 0.86919453 |
| Sin3b | 3636.03601 | 0.38701167 | 0.28946589 | 1.33698538 | 0.18122741 | 0.61208031 |
| Irf2 | 423.261459 | -0.595395 | 0.46407993 | -1.2829578 | 0.19950686 | 0.6323909 |
| Casp3 | 2372.33824 | 0.55223201 | 0.31039536 | 1.77912454 | 0.07521935 | 0.42247854 |
| Cenpu | 671.343746 | -0.1121534 | 0.23866969 | -0.4699106 | 0.63841887 | 0.90611185 |
| Cfap97 | 372.200693 | -0.207001 | 0.34405507 | -0.6016508 | 0.54740659 | 0.87008334 |
| Slc25a4 | 15536.3599 | -0.0515951 | 0.24187308 | -0.2133146 | 0.83108159 | 0.96466657 |
| Ufsp2 | 4458.45792 | -0.0114114 | 0.14919096 | -0.0764888 | 0.93903019 | 0.98913801 |
| Pdlim3 | 11.4150477 | -1.3027082 | 2.29059785 | -0.5687197 | 0.56954635 | 0.87961466 |
| Lrp2bp | 8.05288659 | -1.521088 | 1.81808905 | -0.8366411 | 0.40279436 | 0.79795242 |
| Tlr3 | 100.444458 | 0.98321302 | 0.77377273 | 1.27067418 | 0.20384459 | 0.63824694 |
| Cbr4 | 614.435036 | 0.44740872 | 0.34577887 | 1.29391574 | 0.19569453 | 0.62750489 |
| Sh3rf1 | 63.8875182 | -0.1054238 | 0.85125055 | -0.1238458 | 0.90143739 | 0.98043702 |
| Nek1 | 385.413277 | -0.4672074 | 0.58823651 | -0.794251 | 0.42704935 | 0.81021713 |
| N4bp1 | 484.741951 | -0.3858785 | 0.35992565 | -1.0721062 | 0.28367232 | 0.71951357 |
| Heatr3 | 4156.50471 | -0.5680302 | 0.17234864 | -3.2958204 | 0.00098135 | 0.02304188 |
| Adcy7 | 1849.33219 | -0.053956 | 0.17518035 | -0.3080025 | 0.75808039 | 0.94129085 |
| Brd7 | 2025.82559 | -0.4295435 | 0.18840156 | -2.2799359 | 0.02261149 | 0.21559342 |
| Nkd1 | 14.8051638 | 2.20387363 | 1.51078759 | 1.4587581 | 0.1446317 | 0.57032823 |
| Snx20 | 1311.33901 | 0.02214561 | 0.22866923 | 0.09684562 | 0.922849 | 0.98465273 |
| Rbl2 | 362.687458 | 0.36226632 | 0.3208807 | 1.12897509 | 0.25890834 | 0.69546977 |
| Aktip | 1103.81642 | 0.65946579 | 0.29354665 | 2.24654509 | 0.02466911 | 0.22775644 |
| Eif2ak3 | 383.3317 | 0.40602845 | 0.60472769 | 0.6714236 | 0.50195071 | 0.85006796 |
| Gins3 | 773.811288 | -0.1249722 | 0.26896424 | -0.4646424 | 0.64218759 | 0.90705802 |
| Setd6 | 558.518816 | -0.5897922 | 0.34417221 | -1.7136544 | 0.08659222 | 0.44976392 |
| Got2 | 7739.33877 | -0.3341725 | 0.19023785 | -1.7566038 | 0.07898536 | 0.43543393 |
| Cdh11 | 37.9969081 | 1.18292825 | 1.43383881 | 0.82500783 | 0.40936714 | 0.80100711 |
| Smad1 | 207.090461 | 0.8511127 | 0.43474688 | 1.95772009 | 0.05026287 | 0.34299029 |
| Lsm6 | 3730.85987 | -0.3256928 | 0.24313382 | -1.3395617 | 0.18038787 | 0.61122544 |
| Slc10a7 | 1034.95477 | -0.0256205 | 0.23081506 | -0.1110002 | 0.91161617 | 0.98292561 |
| Tnpo2 | 567.261064 | -0.3011699 | 0.32190105 | -0.9355979 | 0.34948027 | 0.76197111 |
| Vps35 | 23070.4909 | 0.68470276 | 0.23160472 | 2.95634194 | 0.00311312 | 0.05543343 |
| Orc6 | 4473.60605 | 0.08901429 | 0.26645451 | 0.33406938 | 0.73832721 | 0.93800224 |
| Gpt2 | 754.695956 | 1.68581326 | 0.26037922 | 6.47445384 | 9.515549292 | 2.078637758 |
| Dnaja2 | 7917.0498 | -0.3431706 | 0.25070504 | -1.3688221 | 0.17105488 | 0.60430173 |
| Itfg1 | 4036.72007 | 0.22962868 | 0.18526342 | 1.23947123 | 0.21517104 | 0.64854028 |
| Rfx1 | 182.961945 | -0.0831227 | 0.54909376 | -0.1513816 | 0.87967469 | 0.97541902 |
| Tecr | 8062.50005 | -0.2764674 | 0.16836454 | -1.6420762 | 0.10057421 | 0.48480347 |
| Tbc1d9 | 120.933194 | 0.37450302 | 0.60513863 | 0.61887145 | 0.53600105 | 0.8654666 |
| Zfp330 | 2712.5843 | -0.0045191 | 0.19215646 | -0.0235176 | 0.98123744 | 0.99523071 |
| Il15 | 453.37503 | 1.12246326 | 0.44525431 | 2.5209487 | 0.01170389 | 0.1424614 |

| | | | | | | |
|---|---|---|---|---|---|---|
| Gab1 | 529.127026 | 1.5478644 | 0.50264682 | 3.0794274 | 0.00207399 | 0.041123 |
| Smarca5 | 1619.15071 | -0.4093472 | 0.25576414 | -1.600487 | 0.1094906 | 0.50553519 |
| Hp | 3547.72478 | -4.7686071 | 0.34475607 | -13.831829 | 1.638015979 | 5.009462367 |
| Txnl4b | 1438.29715 | -0.1012233 | 0.17384663 | -0.5822563 | 0.56039403 | 0.87469256 |
| Zfp821 | 64.782017 | 0.32920818 | 0.60737179 | 0.54202085 | 0.58780414 | 0.88685349 |
| Ist1 | 4816.93697 | 0.16779816 | 0.15166592 | 1.10636691 | 0.26856774 | 0.70532185 |
| Dhodh | 2295.76704 | -0.099258 | 0.27348007 | -0.3629443 | 0.71664646 | 0.93235217 |
| Ap1g1 | 1253.49616 | -0.1653176 | 0.35869164 | -0.4608906 | 0.64487711 | 0.90835328 |
| Phlpp2 | 88.4702789 | -0.6656916 | 0.72697755 | -0.9156975 | 0.35982556 | 0.76857232 |
| Mmp2 | 400.992923 | 1.81684994 | 0.42042471 | 4.32146326 | 1.549978610 | 0.00075307 |
| St3gal2 | 99.1517626 | 0.14386987 | 0.59376758 | 0.24229997 | 0.80854774 | 0.95844313 |
| Amfr | 1696.54646 | -0.0219745 | 0.2547626 | -0.0862547 | 0.93126398 | 0.98730569 |
| Cog4 | 2725.13057 | 0.10587193 | 0.19839897 | 0.53363143 | 0.59359656 | 0.88797037 |
| Nudt21 | 3407.95998 | -0.3344269 | 0.20978331 | -1.5941539 | 0.11090156 | 0.50887427 |
| Bbs2 | 198.945417 | -0.0469467 | 0.44168292 | -0.1062906 | 0.91535179 | 0.98353083 |
| Cenpn | 1560.12626 | -0.1825055 | 0.21006303 | -0.8688129 | 0.38494949 | 0.78754751 |
| Cdyl2 | 116.217511 | 0.53397806 | 0.78486291 | 0.68034563 | 0.49628564 | 0.84787395 |
| Mt2 | 205.572522 | 0.8623791 | 0.37699003 | 2.28753819 | 0.02216443 | 0.21416861 |
| Mt1 | 4433.7009 | 0.42557217 | 0.18905236 | 2.25108097 | 0.02438041 | 0.22560178 |
| Nudt7 | 451.641833 | 0.85716192 | 0.53739129 | 1.59504245 | 0.11070274 | 0.50858841 |
| Herpud1 | 8985.60296 | 0.20732193 | 0.29129608 | 0.71172236 | 0.47663672 | 0.83666192 |
| Fam192a | 2426.17059 | -0.0448616 | 0.2132935 | -0.2103282 | 0.83341154 | 0.96565375 |
| Arl2bp | 772.776989 | 0.181052 | 0.29848968 | 0.60656035 | 0.54414272 | 0.8695007 |
| Cx3cl1 | 85.4836308 | 0.416398 | 0.58086882 | 0.71685376 | 0.47346434 | 0.83483892 |
| Ciapin1 | 2570.68981 | -0.4961677 | 0.23413257 | -2.1191743 | 0.03407574 | 0.27569457 |
| Coq9 | 2083.13805 | -0.1302056 | 0.21418485 | -0.6079125 | 0.54324552 | 0.86919453 |
| Polr2c | 6058.2794 | 0.06186729 | 0.29997055 | 0.20624453 | 0.83659991 | 0.96683245 |
| Adgrg1 | 178.181235 | -2.8128324 | 0.40123326 | -7.0104667 | 2.375244880 | 6.603720596 |
| Katnb1 | 940.862033 | -0.13346 | 0.33761822 | -0.3952986 | 0.69262253 | 0.92549717 |
| Kifc3 | 234.575238 | 1.39812658 | 0.47396525 | 2.94985037 | 0.00317928 | 0.05620248 |
| Tmem38a | 476.357362 | 1.06966298 | 0.26026705 | 4.10986711 | 3.958868807 | 0.00166996 |
| Usb1 | 1251.11992 | -0.1130383 | 0.2286615 | -0.4943477 | 0.62106068 | 0.89896588 |
| Cfap20 | 1591.40322 | -0.11827 | 0.16750403 | -0.7060726 | 0.48014298 | 0.83935746 |
| Tpm4 | 17367.8533 | 0.08725375 | 0.16881987 | 0.51684525 | 0.60526419 | 0.89258167 |
| B3gnt3 | 60.1775926 | 2.6683411 | 1.44347699 | 1.84855119 | 0.06452265 | 0.39212252 |
| Jak3 | 1881.45605 | -0.2536624 | 0.30761601 | -0.8246073 | 0.40959456 | 0.80100711 |
| Pgls | 2065.29101 | -0.1712364 | 0.17662472 | -0.9694931 | 0.33229923 | 0.75138937 |
| Slc27a1 | 275.304564 | 0.67253212 | 0.39353203 | 1.70896414 | 0.08745759 | 0.45207544 |
| Map1lc3b | 10807.7447 | 0.84323271 | 0.3203739 | 2.63202682 | 0.00848772 | 0.11601143 |
| Mvb12a | 1870.56327 | 0.04475683 | 0.31996736 | 0.13987935 | 0.88875532 | 0.97850273 |
| Mthfsd | 914.683149 | 0.05236436 | 0.27347766 | 0.19147583 | 0.84815281 | 0.96765585 |
| Cox4i1 | 27879.6813 | 0.39658588 | 0.38698812 | 1.02480117 | 0.30545701 | 0.73530116 |
| Emc8 | 3333.72452 | 0.32564897 | 0.40108107 | 0.81192805 | 0.41683292 | 0.8047849 |
| Babam1 | 2653.25259 | 0.12812295 | 0.1863766 | 0.68744122 | 0.49180473 | 0.84516159 |
| Gins2 | 4665.8044 | 0.59666341 | 0.33634033 | 1.77398712 | 0.07606528 | 0.42514288 |
| Gse1 | 40.7412058 | -1.006943 | 0.95708486 | -1.0520937 | 0.29275656 | 0.72584701 |

| | | | | | | |
|---|---|---|---|---|---|---|
| Zdhhc7 | 2066.19508 | -0.1119984 | 0.2636949 | -0.4247271 | 0.67103561 | 0.91711586 |
| 6430548M08 | 255.558925 | 0.21645756 | 0.5404006 | 0.40055018 | 0.68875133 | 0.92470267 |
| Crispld2 | 459.687871 | 0.55018765 | 0.4029589 | 1.36536914 | 0.17213706 | 0.60458014 |
| Usp10 | 4709.89135 | -0.566694 | 0.26190718 | -2.1637208 | 0.03048579 | 0.26060979 |
| Cotl1 | 6913.46387 | -0.061359 | 0.22764285 | -0.2695406 | 0.78751372 | 0.95225386 |
| Klhl36 | 454.667948 | -0.1151795 | 0.30553543 | -0.376976 | 0.70619144 | 0.92896305 |
| Taf1c | 275.400799 | -0.4417948 | 0.57829456 | -0.7639616 | 0.44489014 | 0.82118815 |
| Mast3 | 358.159641 | 0.19824786 | 0.30628972 | 0.64725601 | 0.51746627 | 0.85694744 |
| Pik3r2 | 1492.30064 | -0.2250842 | 0.2059383 | -1.0929691 | 0.27440736 | 0.71147204 |
| Mbtps1 | 3120.86016 | -0.0042408 | 0.18895724 | -0.0224433 | 0.98209435 | 0.99523682 |
| Hsbp1 | 4558.282 | 0.87919707 | 0.31673153 | 2.77584323 | 0.00550587 | 0.08595935 |
| Rab3a | 36.3866441 | 0.68019903 | 1.30243851 | 0.5222504 | 0.601496 | 0.89094157 |
| Mphosph6 | 1158.92969 | 0.04461646 | 0.30060742 | 0.14842102 | 0.88201051 | 0.97590762 |
| 1700030J22F | 17.0500825 | -0.6927118 | 1.4117251 | -0.4906847 | 0.6236495 | 0.90017327 |
| Lsm4 | 5941.10516 | 0.12677804 | 0.2831639 | 0.44771963 | 0.65435555 | 0.9108895 |
| Ntpcr | 3439.68562 | 1.32063662 | 0.30533786 | 4.32516505 | 1.524177026 | 0.00075182 |
| Mau2 | 1857.29216 | 0.38650626 | 0.20936013 | 1.84613116 | 0.06487316 | 0.39267363 |
| Atp13a1 | 2804.31586 | -0.2029383 | 0.19717819 | -1.0292127 | 0.30337976 | 0.73436097 |
| Ints10 | 2711.92228 | -0.0039157 | 0.17855516 | -0.0219299 | 0.98250384 | 0.99527736 |
| Dctn1 | 2167.83384 | 0.15574811 | 0.34267194 | 0.45451084 | 0.64946119 | 0.90973357 |
| Cdh5 | 54.5567386 | 1.2534438 | 1.30910977 | 0.95747799 | 0.33832605 | 0.75644564 |
| Cmtm3 | 3375.48536 | 0.82800999 | 0.20748276 | 3.99074117 | 6.586713326 | 0.00255207 |
| Nae1 | 2161.19141 | 0.2998196 | 0.28869253 | 1.03854297 | 0.29901734 | 0.73093388 |
| Ciao2b | 1812.28737 | 0.34324505 | 0.32548895 | 1.05455209 | 0.29163023 | 0.72554658 |
| Rrad | 1350.96517 | 0.51146838 | 0.35396859 | 1.44495411 | 0.14847077 | 0.5754118 |
| Car7 | 22.1982206 | 1.41547195 | 1.19432573 | 1.18516408 | 0.23595258 | 0.67182855 |
| Cbfb | 1074.80901 | -0.5606029 | 0.39301608 | -1.426412 | 0.15374944 | 0.58242478 |
| Tradd | 1013.41774 | 0.17083169 | 0.23584105 | 0.72435095 | 0.46885031 | 0.83381643 |
| D230025D16 | 1133.85987 | 0.16707082 | 0.27009907 | 0.61855384 | 0.53621032 | 0.86547841 |
| Psmb10 | 3553.20637 | 0.06889022 | 0.27655741 | 0.24909918 | 0.80328406 | 0.95672466 |
| Dus2 | 726.8738 | 0.35775843 | 0.26092308 | 1.37112604 | 0.17033567 | 0.60430173 |
| Nfatc3 | 274.854946 | -0.4680972 | 0.4085608 | -1.1457223 | 0.25191009 | 0.68505999 |
| Pla2g15 | 4309.72501 | 0.36113792 | 0.244172 | 1.47903085 | 0.13913207 | 0.56286245 |
| Slc7a6 | 1940.56204 | -0.3636521 | 0.22699896 | -1.6019991 | 0.1091558 | 0.50503137 |
| Smpd3 | 22.9773764 | 3.03284502 | 1.21833748 | 2.48933082 | 0.01279838 | 0.15120563 |
| Zfp90 | 557.846135 | 0.79815156 | 0.34854148 | 2.28997582 | 0.02202272 | 0.21347378 |
| Vps4a | 2172.14853 | -0.0288326 | 0.21656532 | -0.133136 | 0.89408583 | 0.97941957 |
| Cog8 | 2596.34235 | -0.17661 | 0.17457849 | -1.0116369 | 0.3117117 | 0.74027747 |
| Nip7 | 5174.64472 | 0.09172877 | 0.37185009 | 0.24668213 | 0.80515424 | 0.95727979 |
| Mtmr2 | 3512.89907 | 0.06837067 | 0.20985879 | 0.3257937 | 0.74458045 | 0.93917909 |
| Terf2 | 146.514556 | -0.6770187 | 0.54157558 | -1.2500909 | 0.21126634 | 0.64504405 |
| Cep57 | 2423.07239 | 0.15019745 | 0.27649618 | 0.54321708 | 0.58698035 | 0.88615705 |
| Cyb5b | 9370.10448 | 0.02420529 | 0.1417349 | 0.17077865 | 0.86439782 | 0.9705375 |
| Maml2 | 420.329651 | 1.56733038 | 0.33046808 | 4.74275876 | 2.108272747 | 0.00013718 |
| Mre11a | 1593.3675 | -0.1348365 | 0.20839232 | -0.6470321 | 0.51761116 | 0.85695893 |
| Wwp2 | 3314.51891 | -0.0297121 | 0.23581808 | -0.1259957 | 0.89973531 | 0.98043702 |

| | | | | | | |
|---|---|---|---|---|---|---|
| Ankrd49 | 1213.29817 | 0.03325616 | 0.27975267 | 0.11887702 | 0.90537279 | 0.98142891 |
| Panx1 | 971.994317 | -0.1878665 | 0.28185162 | -0.6665441 | 0.50506339 | 0.85034006 |
| Med17 | 2560.08641 | -0.2157815 | 0.19366761 | -1.1141847 | 0.26519997 | 0.7011398 |
| 4931406C07 | 386.793435 | -0.2614141 | 0.28238794 | -0.9257268 | 0.35458798 | 0.76419569 |
| Taf1d | 1304.34749 | -0.1778945 | 0.27815757 | -0.6395459 | 0.52246785 | 0.85816354 |
| Kars | 11259.7924 | -0.3703808 | 0.14045209 | -2.6370614 | 0.00836277 | 0.11493265 |
| Adat1 | 215.059471 | 0.00506072 | 0.3116114 | 0.01624047 | 0.98704255 | 0.99616298 |
| Gabarapl2 | 2634.44606 | 0.51244642 | 0.40502171 | 1.26523197 | 0.20578818 | 0.63991024 |
| Tmem231 | 22.824453 | 0.96756909 | 1.35783847 | 0.71258041 | 0.47610544 | 0.83644936 |
| Tmem170 | 149.390396 | 1.17603213 | 0.44106084 | 2.66637168 | 0.00766749 | 0.10768814 |
| Cfdp1 | 3514.29991 | 0.13470096 | 0.20367229 | 0.66136123 | 0.50838069 | 0.85238774 |
| Bcar1 | 65.7510363 | 1.82446093 | 0.68487206 | 2.66394416 | 0.00772304 | 0.10821985 |
| Wdr59 | 101.624053 | 0.56181599 | 0.42433256 | 1.32399925 | 0.18550329 | 0.61775377 |
| Aars | 7367.34162 | -0.2162903 | 0.24708101 | -0.875382 | 0.38136609 | 0.7839441 |
| Afg3l1 | 2205.55269 | -0.2823155 | 0.28282325 | -0.9982046 | 0.31818017 | 0.74336577 |
| Acad8 | 1710.59735 | -0.041701 | 0.20307456 | -0.2053482 | 0.83730012 | 0.96683245 |
| Ccsap | 52.191335 | 0.31167403 | 0.66819735 | 0.46644009 | 0.64090052 | 0.90685007 |
| Abcb10 | 815.795771 | 0.08935512 | 0.26361394 | 0.33896205 | 0.73463832 | 0.93750666 |
| Urb2 | 300.253958 | -0.3509675 | 0.47011082 | -0.7465633 | 0.45532723 | 0.82788614 |
| Cog2 | 1991.72572 | 0.27879548 | 0.26265471 | 1.06145242 | 0.28848435 | 0.72375492 |
| Arv1 | 182.262469 | 0.00532394 | 0.44193838 | 0.0120468 | 0.99038828 | 0.9966603 |
| 2310022B05I | 190.640223 | 0.11626721 | 0.31904373 | 0.36442408 | 0.71554136 | 0.93235217 |
| 2810004N23 | 2430.93125 | 0.16797761 | 0.23745587 | 0.7074056 | 0.47931446 | 0.8387067 |
| Gnpat | 4713.30615 | -0.1572929 | 0.18546647 | -0.8480934 | 0.39638598 | 0.79439542 |
| Sprtn | 332.026621 | -0.0468126 | 0.35021605 | -0.1336678 | 0.89366525 | 0.97941957 |
| Egln1 | 156.897273 | -0.5127037 | 0.5259755 | -0.9747672 | 0.32967573 | 0.74891795 |
| Vps26b | 1415.48056 | -0.1986366 | 0.26544959 | -0.7483026 | 0.45427767 | 0.82733047 |
| Jam3 | 154.070315 | 0.12345924 | 0.43033727 | 0.28688948 | 0.77419694 | 0.94688575 |
| Snx19 | 616.407139 | 0.1904214 | 0.37214618 | 0.51168442 | 0.60887189 | 0.89437372 |
| St14 | 66.5624569 | -0.1253429 | 0.54449311 | -0.2302011 | 0.8179355 | 0.96080324 |
| Aplp2 | 4110.09248 | 0.54791173 | 0.23754405 | 2.30656896 | 0.02107886 | 0.20744788 |
| Birc3 | 1280.71096 | -0.1685977 | 0.44565096 | -0.3783177 | 0.70519458 | 0.92879471 |
| Dcun1d5 | 1227.99243 | 0.42077384 | 0.29106795 | 1.44562066 | 0.14828363 | 0.57512797 |
| Sesn3 | 32.9872984 | 1.49091311 | 1.41216824 | 1.05576168 | 0.29107711 | 0.72510985 |
| Usp2 | 768.059203 | 0.01031825 | 0.42579761 | 0.02423275 | 0.98066695 | 0.99516906 |
| Oaf | 595.423002 | -0.694153 | 0.31922228 | -2.174513 | 0.02966663 | 0.25669202 |
| Sc5d | 2038.33764 | 0.0740683 | 0.26435099 | 0.28018922 | 0.77933234 | 0.9488468 |
| Ubash3b | 2845.23533 | 0.00812355 | 0.29092382 | 0.02792328 | 0.97772334 | 0.99483019 |
| Clmp | 109.814189 | 1.23707501 | 0.57224118 | 2.16180704 | 0.03063305 | 0.26095692 |
| Rexo2 | 2596.20806 | -0.0662378 | 0.26720225 | -0.2478939 | 0.80421654 | 0.9570372 |
| Cul5 | 2465.29781 | 0.25214961 | 0.20620487 | 1.22281114 | 0.22140104 | 0.65499369 |
| Ets1 | 565.011848 | 0.61409725 | 0.28075599 | 2.18729879 | 0.02872072 | 0.25235676 |
| St3gal4 | 781.550599 | 0.26607383 | 0.33916351 | 0.78450017 | 0.43274668 | 0.81435873 |
| Dcps | 3454.84892 | -0.3498587 | 0.16021936 | -2.1836231 | 0.02898995 | 0.25346382 |
| Tirap | 1608.47547 | 0.18580547 | 0.33421302 | 0.55594922 | 0.57824557 | 0.88211473 |
| Srpr | 6044.21726 | -0.0956084 | 0.16555209 | -0.5775122 | 0.56359347 | 0.87626321 |

| | | | | | | |
|---|---|---|---|---|---|---|
| Rpusd4 | 1388.11905 | -0.1783348 | 0.2353874 | -0.7576224 | 0.4486771 | 0.82417561 |
| Abhd12 | 8883.02464 | 1.20066443 | 0.25289637 | 4.74765377 | 2.057899190 | 0.00013535 |
| Acat1 | 7477.32782 | 0.64196716 | 0.24638111 | 2.60558599 | 0.00917172 | 0.12248658 |
| Rdx | 5598.32018 | 0.01810283 | 0.15859456 | 0.11414532 | 0.9091226 | 0.98218641 |
| Fdx1 | 627.159858 | 0.43823945 | 0.29586559 | 1.48121126 | 0.13855029 | 0.56122045 |
| Ppp2r1b | 6266.27926 | -0.1404707 | 0.22512837 | -0.6239583 | 0.53265499 | 0.8646427 |
| Alg9 | 470.571403 | 0.57220276 | 0.52374775 | 1.09251594 | 0.27460639 | 0.71147204 |
| Cryab | 142.373607 | 0.38304308 | 0.54943607 | 0.69715677 | 0.48570469 | 0.84183502 |
| Dixdc1 | 26.5302268 | 1.15699409 | 1.3851417 | 0.83528933 | 0.40355482 | 0.79842894 |
| Bco2 | 301.759034 | 1.00986756 | 0.31017837 | 3.25576397 | 0.00113088 | 0.02575347 |
| Pts | 2769.3207 | 0.8132842 | 0.43551301 | 1.86741653 | 0.06184344 | 0.38383097 |
| Cadm1 | 1182.87295 | 1.47313864 | 0.27031849 | 5.44964068 | 5.047168675 | 5.415966176 |
| Bud13 | 854.269556 | -0.280406 | 0.27228665 | -1.0298193 | 0.30309483 | 0.73428908 |
| Zpr1 | 3676.05134 | -0.5157988 | 0.19230291 | -2.6822204 | 0.00731353 | 0.10439482 |
| Apoa1 | 66.486242 | 2.38078446 | 0.90491078 | 2.63096044 | 0.00851439 | 0.11624619 |
| Tagln | 18.885559 | -0.961864 | 1.64252773 | -0.5855999 | 0.55814443 | 0.87311776 |
| Bace1 | 238.348682 | 0.58336627 | 0.39818035 | 1.4650805 | 0.14289896 | 0.56846902 |
| Il10ra | 2214.0673 | 0.02393208 | 0.29130943 | 0.08215346 | 0.93452468 | 0.98756396 |
| Arcn1 | 16400.0996 | -0.021994 | 0.19607261 | -0.1121728 | 0.91068639 | 0.98288717 |
| Ddx6 | 2117.68964 | -0.0183372 | 0.28036645 | -0.0654043 | 0.94785207 | 0.99019648 |
| Ddx25 | 13.7191512 | 1.48278352 | 2.12856536 | 0.69661169 | 0.48604583 | 0.84183502 |
| Pus3 | 549.846753 | 0.1713293 | 0.29164861 | 0.58745112 | 0.55690077 | 0.87276537 |
| Nlrx1 | 686.107957 | -0.3998099 | 0.32870208 | -1.2163292 | 0.22385955 | 0.65774578 |
| Trappc4 | 2479.77027 | 0.06360749 | 0.22434739 | 0.28352233 | 0.77677647 | 0.94800948 |
| Chek1 | 1274.49764 | 0.02123337 | 0.2880096 | 0.07372451 | 0.94122959 | 0.98935054 |
| Slc37a4 | 916.597014 | -0.2570608 | 0.2989153 | -0.8599786 | 0.38980082 | 0.78973724 |
| Hyou1 | 1087.72212 | -0.0979818 | 0.26511021 | -0.3695888 | 0.71168891 | 0.9321296 |
| Stt3a | 14671.4192 | 0.0540747 | 0.14797655 | 0.36542748 | 0.71479233 | 0.93235217 |
| Hinfp | 605.402944 | -0.1183637 | 0.27828727 | -0.4253291 | 0.67059679 | 0.91711586 |
| C2cd2l | 251.689142 | 0.28431859 | 0.38238509 | 0.74353994 | 0.45715485 | 0.82936012 |
| Tmem218 | 251.525998 | 0.30812196 | 0.49517878 | 0.62224386 | 0.53378153 | 0.86500711 |
| Slc37a2 | 1703.73836 | 0.86016511 | 0.27606698 | 3.11578414 | 0.00183457 | 0.03765476 |
| Dpagt1 | 2463.49032 | -0.0390809 | 0.22845169 | -0.1710687 | 0.86416976 | 0.9705375 |
| Robo4 | 38.9250961 | 2.96952527 | 3.66153816 | 0.81100487 | 0.41736288 | 0.8049685 |
| Hmbs | 7361.18805 | 0.06901094 | 0.24937897 | 0.27673118 | 0.78198653 | 0.95004728 |
| Vps11 | 2986.818 | 0.01713331 | 0.24023793 | 0.07131808 | 0.9431446 | 0.98983253 |
| Mcam | 43.0037591 | 2.38114691 | 1.27990849 | 1.86040402 | 0.06282839 | 0.38699881 |
| Pin1 | 6618.92188 | 0.0350626 | 0.29841201 | 0.11749729 | 0.90646599 | 0.98159944 |
| Tyk2 | 1192.6381 | -0.3252563 | 0.27421064 | -1.186155 | 0.23556108 | 0.6716704 |
| Pde4a | 371.857714 | 0.05435251 | 0.30352324 | 0.179072 | 0.85788116 | 0.96944425 |
| Ilf3 | 9197.91681 | -0.3162095 | 0.25349732 | -1.247388 | 0.21225525 | 0.64580271 |
| Tmed1 | 1791.3606 | 0.26053454 | 0.36368672 | 0.7163708 | 0.47376242 | 0.8350916 |
| Yipf2 | 593.537216 | -0.0449368 | 0.25216408 | -0.1782047 | 0.8585622 | 0.96954641 |
| Lysmd2 | 222.439964 | -0.1820928 | 0.35855211 | -0.507856 | 0.61155435 | 0.89583815 |
| Carm1 | 447.170408 | -0.9372194 | 0.56131125 | -1.6696965 | 0.09497943 | 0.47206881 |
| Tmod2 | 21.8669606 | 0.26543805 | 1.51824811 | 0.1748318 | 0.8612118 | 0.97000313 |

| | | | | | | |
|---|---|---|---|---|---|---|
| Smarca4 | 4247.02839 | -0.1833202 | 0.27404063 | -0.6689525 | 0.5035258 | 0.85012672 |
| Gnb5 | 127.915537 | 0.37816894 | 0.72167972 | 0.52401214 | 0.60027009 | 0.8902352 |
| Ldlr | 1941.43142 | 0.50960967 | 0.34092205 | 1.49479819 | 0.13496712 | 0.55648241 |
| Kank2 | 334.098506 | 1.09532406 | 0.39993403 | 2.73876182 | 0.0061671 | 0.09371699 |
| Dock6 | 117.204909 | 0.45174705 | 0.72553115 | 0.62264322 | 0.53351901 | 0.86492515 |
| Polr2m | 13351.7674 | -0.1408725 | 0.14368151 | -0.9804494 | 0.32686433 | 0.74811039 |
| Rab27a | 1518.11348 | -1.8667033 | 0.22917415 | -8.1453485 | 3.781909833 | 1.779388576 |
| Aqp9 | 72.1252377 | -3.0275431 | 1.0631702 | -2.8476562 | 0.00440425 | 0.07270873 |
| Sltm | 1142.78906 | -0.5132333 | 0.31057835 | -1.6525084 | 0.09843096 | 0.48011435 |
| Rsl24d1 | 4454.08998 | -0.1502661 | 0.27883093 | -0.5389147 | 0.58994569 | 0.88723944 |
| Nedd4 | 1711.52195 | -0.6907342 | 0.35315754 | -1.9558813 | 0.05047914 | 0.34374326 |
| Rnf111 | 525.77328 | -0.0454533 | 0.50578463 | -0.089867 | 0.92839291 | 0.98666423 |
| Ccnb2 | 8753.51413 | -0.0504628 | 0.19480647 | -0.2590409 | 0.7956037 | 0.9537323 |
| Myo1e | 45.2687898 | 0.84686366 | 0.80254177 | 1.0552269 | 0.29132157 | 0.72536876 |
| Mns1 | 484.177206 | -0.6379844 | 0.3578683 | -1.7827353 | 0.07462941 | 0.42093241 |
| Tcf12 | 714.60806 | 0.10179987 | 0.38199081 | 0.26649821 | 0.78985554 | 0.95255415 |
| Anxa2 | 6964.26976 | 0.5331494 | 0.16096668 | 3.31217251 | 0.00092574 | 0.02201314 |
| Cgnl1 | 13.807841 | 0.30429405 | 1.57409041 | 0.19331422 | 0.84671289 | 0.96765585 |
| Ice2 | 453.739246 | 0.49549778 | 0.32576221 | 1.52104133 | 0.12824947 | 0.54327949 |
| Rora | 54.2400252 | 0.96793284 | 1.26815139 | 0.76326285 | 0.44530666 | 0.82118815 |
| Rp9 | 932.205651 | -0.258484 | 0.22936877 | -1.1269361 | 0.25976948 | 0.69596146 |
| Fem1b | 1034.52853 | 0.47795147 | 0.2407527 | 1.98523823 | 0.04711797 | 0.33491816 |
| Cln6 | 1913.33628 | -0.0037619 | 0.25279188 | -0.0148816 | 0.98812664 | 0.9962454 |
| Calml4 | 51.9412926 | 1.13417089 | 0.68706829 | 1.65073968 | 0.09879174 | 0.48025164 |
| Anp32a | 1487.9181 | -0.2249645 | 0.25020325 | -0.8991272 | 0.36858492 | 0.77456461 |
| Irak1bp1 | 123.51649 | 0.06605282 | 0.51234059 | 0.12892366 | 0.89741806 | 0.97984623 |
| Glce | 234.978507 | 0.31547717 | 0.42403147 | 0.74399471 | 0.45687968 | 0.82915295 |
| Phip | 1591.394 | 0.06997941 | 0.23502814 | 0.29774908 | 0.76589468 | 0.94347227 |
| Kif23 | 2331.71042 | -0.47012 | 0.22310602 | -2.1071594 | 0.03510376 | 0.28086853 |
| Lca5 | 77.1104192 | 0.04872256 | 0.6400738 | 0.07612023 | 0.93932345 | 0.98913801 |
| Sh3bgrl2 | 215.337276 | 0.43345763 | 0.51514897 | 0.84142191 | 0.40011162 | 0.79638227 |
| Bckdhb | 306.505787 | -0.8601845 | 0.31927536 | -2.6941777 | 0.00705625 | 0.10256275 |
| Zw10 | 1892.68679 | -0.080951 | 0.25888819 | -0.3126873 | 0.75451825 | 0.94044759 |
| Tent5a | 438.538294 | -0.4436647 | 0.35260907 | -1.2582339 | 0.20830716 | 0.6418276 |
| Usp28 | 767.405978 | 0.52066537 | 0.23275269 | 2.23698969 | 0.02528701 | 0.23122295 |
| Idh3a | 8135.08714 | -0.4573172 | 0.19377428 | -2.360051 | 0.01827242 | 0.19092573 |
| Tle3 | 409.4234 | -0.4682349 | 0.27082044 | -1.7289495 | 0.08381814 | 0.44417933 |
| Dnaja4 | 31.2308215 | -0.4835557 | 1.55313726 | -0.3113412 | 0.75554121 | 0.94090763 |
| Imp3 | 1583.4648 | -0.4443525 | 0.21189522 | -2.0970387 | 0.03599015 | 0.28477582 |
| Ptpn9 | 465.8564 | 0.65842369 | 0.51229854 | 1.28523435 | 0.1987104 | 0.63148061 |
| Crabp1 | 31.4457047 | 1.15052863 | 0.93362974 | 1.2323179 | 0.21783036 | 0.65150232 |
| Ireb2 | 542.172267 | -0.0598195 | 0.2837819 | -0.2107938 | 0.83304816 | 0.96565375 |
| Pkm | 100425.906 | -0.1824084 | 0.17036118 | -1.0707159 | 0.2842972 | 0.71974495 |
| Man2c1 | 1007.13053 | -0.1291032 | 0.25366233 | -0.5089568 | 0.61078251 | 0.89524352 |
| Neil1 | 190.266972 | 0.18310995 | 0.47210568 | 0.38785797 | 0.69812115 | 0.92669304 |
| Commd4 | 2500.6606 | 0.30080174 | 0.28131922 | 1.06925411 | 0.28495518 | 0.72045503 |

| | | | | | | |
|---|---|---|---|---|---|---|
| 1700017B05l | 1590.77455 | -0.1969482 | 0.22035543 | -0.893775 | 0.37144227 | 0.7765614 |
| Psma4 | 13121.8088 | 0.02605311 | 0.28317723 | 0.09200284 | 0.92669578 | 0.98630274 |
| Fam219b | 1015.09211 | 0.00533288 | 0.21188428 | 0.02516881 | 0.97992031 | 0.99486654 |
| Mpi | 2418.34195 | 0.34091236 | 0.17881189 | 1.90654186 | 0.05657994 | 0.36524665 |
| Ube2q2 | 672.837812 | 0.32391622 | 0.23466093 | 1.38035855 | 0.16747628 | 0.60168496 |
| Ulk3 | 221.710996 | -0.1447443 | 0.54222017 | -0.2669474 | 0.78950963 | 0.95255415 |
| Fbxo22 | 3348.95201 | -0.0648078 | 0.15985976 | -0.405404 | 0.68518056 | 0.92314423 |
| Nrg4 | 148.652121 | 0.71913615 | 0.72158806 | 0.99660205 | 0.31895771 | 0.74398524 |
| Csk | 1738.63175 | -0.237256 | 0.19157298 | -1.238463 | 0.21554445 | 0.64864826 |
| Etfa | 6480.67811 | 0.2011308 | 0.26056384 | 0.77190605 | 0.44017008 | 0.81963526 |
| Clk3 | 2712.03465 | -0.020659 | 0.19035512 | -0.1085288 | 0.91357625 | 0.98309098 |
| Rcn2 | 3845.44622 | 0.42337854 | 0.247839 | 1.70828051 | 0.08758431 | 0.45207544 |
| Pstpip1 | 1246.77741 | -0.4544171 | 0.23822096 | -1.9075445 | 0.05645011 | 0.36517938 |
| Tspan3 | 4054.65061 | 0.50662551 | 0.17916457 | 2.82771032 | 0.00468822 | 0.0757609 |
| Tmem30a | 4569.43228 | 0.64184636 | 0.21925885 | 2.92734534 | 0.00341869 | 0.05967533 |
| Hmg20a | 1265.97116 | 0.19740339 | 0.20512089 | 0.96237583 | 0.33586085 | 0.7540073 |
| Cox7a2 | 6320.43017 | 0.37022481 | 0.390183 | 0.94884914 | 0.34269735 | 0.75932198 |
| Stoml1 | 486.385724 | 0.09095288 | 0.26473332 | 0.34356415 | 0.73117409 | 0.93653187 |
| Nptn | 4348.02548 | -0.0177907 | 0.17770812 | -0.1001121 | 0.92025531 | 0.9839023 |
| Neo1 | 63.9514441 | -0.0562857 | 0.65941031 | -0.0853577 | 0.93197704 | 0.98730569 |
| Mto1 | 2094.23322 | -0.0660523 | 0.2048815 | -0.3223928 | 0.74715517 | 0.9399191 |
| Cgas | 849.165015 | -0.2564435 | 0.27791489 | -0.9227411 | 0.35614215 | 0.76527084 |
| Gsta4 | 27.3771356 | 1.62273386 | 0.89899204 | 1.8050592 | 0.07106542 | 0.41122734 |
| Elovl5 | 2238.02799 | -0.5219441 | 0.22353455 | -2.3349592 | 0.01954555 | 0.19809504 |
| Gclc | 253.865212 | 0.00079349 | 0.37747747 | 0.00210208 | 0.99832279 | 0.99967651 |
| Lrrc1 | 10.6315477 | 2.07597868 | 1.99180644 | 1.04225925 | 0.2972915 | 0.72954201 |
| Tmed3 | 2576.33916 | -0.1440457 | 0.33109338 | -0.4350606 | 0.66351846 | 0.91395288 |
| Ctsh | 2689.40666 | -0.0847326 | 0.38970816 | -0.2174258 | 0.82787656 | 0.96432112 |
| Tpm1 | 1614.49129 | -0.2778024 | 0.22547757 | -1.2320621 | 0.2179259 | 0.65150232 |
| Plscr1 | 846.213579 | 0.34729968 | 0.2626061 | 1.32251188 | 0.18599776 | 0.61790851 |
| Lactb | 290.247168 | -0.5307025 | 0.58263423 | -0.9108674 | 0.36236523 | 0.76969598 |
| Car12 | 43.8789186 | -0.2404778 | 0.75734264 | -0.3175284 | 0.75084269 | 0.94003261 |
| Plod2 | 175.623598 | 0.51422029 | 0.4242413 | 1.2120939 | 0.22547644 | 0.65900564 |
| Aph1b | 139.321497 | 1.05332514 | 0.57593229 | 1.82890447 | 0.06741392 | 0.40006856 |
| Usp3 | 1719.26008 | -0.2029392 | 0.20767546 | -0.977194 | 0.32847313 | 0.74843736 |
| Dapk2 | 14.0761911 | -0.1183379 | 1.94184563 | -0.0609409 | 0.95140625 | 0.99115284 |
| Ciao2a | 16481.4366 | 0.36368608 | 0.40684917 | 0.89390886 | 0.37137066 | 0.7765614 |
| Snx1 | 8557.76291 | 0.23689535 | 0.29563883 | 0.80129985 | 0.42295808 | 0.80844471 |
| Ppib | 9157.63786 | 0.27386178 | 0.31481984 | 0.8699 | 0.38435506 | 0.78704643 |
| Csnk1g1 | 506.934839 | -0.0846102 | 0.26998682 | -0.3133865 | 0.75398701 | 0.94018119 |
| Trip4 | 1311.79881 | 0.34903751 | 0.18159353 | 1.9220812 | 0.05459554 | 0.36033644 |
| Rbpms2 | 59.5953242 | 0.45779943 | 0.80163158 | 0.57108457 | 0.56794232 | 0.87929304 |
| Spg21 | 5430.23304 | 0.17613436 | 0.2257199 | 0.78032267 | 0.43520097 | 0.81616028 |
| Parp16 | 220.106001 | 1.50160332 | 0.40420382 | 3.71496566 | 0.00020323 | 0.00661204 |
| Dpp8 | 1244.96916 | -0.0204102 | 0.41416832 | -0.04928 | 0.96069614 | 0.99325523 |
| Dis3l | 1301.5032 | -0.0076985 | 0.23411751 | -0.0328829 | 0.97376795 | 0.99443087 |

| | | | | | | |
|---|---|---|---|---|---|---|
| Tipin | 6604.12017 | -0.2252319 | 0.25596079 | -0.8799468 | 0.37888811 | 0.78134494 |
| Snapc5 | 1395.12382 | 0.0292973 | 0.23321757 | 0.12562217 | 0.90003101 | 0.98043702 |
| Rpl4 | 73716.5016 | -0.1812491 | 0.18682448 | -0.9701572 | 0.33196812 | 0.75105716 |
| Zwilch | 1851.48137 | -0.0176556 | 0.21382488 | -0.0825703 | 0.93419321 | 0.98756396 |
| Smad3 | 1155.40334 | -0.0876251 | 0.33194974 | -0.263971 | 0.79180229 | 0.95279534 |
| 2300009A05I | 145.042617 | 0.15253512 | 0.38968527 | 0.39143157 | 0.69547826 | 0.92623267 |
| Pias1 | 3596.58814 | -0.008216 | 0.2234961 | -0.0367613 | 0.97067534 | 0.99443087 |
| U2surp | 4063.79019 | -0.1638478 | 0.2548903 | -0.6428169 | 0.52034289 | 0.8576517 |
| Atr | 176.319781 | -0.2137595 | 0.44051789 | -0.485246 | 0.62750189 | 0.9022352 |
| Xrn1 | 435.080367 | -0.2560597 | 0.30793426 | -0.8315402 | 0.40566855 | 0.79982698 |
| Tfdp2 | 271.709225 | -0.3309231 | 0.29467056 | -1.1230274 | 0.26142587 | 0.69719264 |
| Atp1b3 | 13906.9702 | 0.04017751 | 0.16890774 | 0.23786659 | 0.81198457 | 0.95967622 |
| Rasa2 | 622.55913 | -0.5953104 | 0.39299019 | -1.5148226 | 0.12981732 | 0.54647951 |
| Ube2cbp | 123.782607 | -0.0891646 | 0.46762851 | -0.190674 | 0.84878103 | 0.96765585 |
| Rwdd2a | 124.792524 | 0.58227066 | 0.52067189 | 1.11830632 | 0.26343619 | 0.69987547 |
| Me1 | 54.6400034 | 0.84430532 | 1.41427176 | 0.59698945 | 0.55051443 | 0.87093412 |
| Nt5e | 30.6169109 | -5.2761397 | 1.52684847 | -3.4555752 | 0.00054912 | 0.01441499 |
| Snx14 | 1535.50604 | 0.13426978 | 0.22287531 | 0.6024435 | 0.54687896 | 0.86994632 |
| Syncrip | 5791.06075 | -0.3677575 | 0.20881304 | -1.7611809 | 0.07820779 | 0.43258005 |
| Zfp949 | 91.0117058 | 0.33852088 | 0.59647561 | 0.56753516 | 0.57035065 | 0.88027118 |
| Crtap | 1157.16676 | 0.30678111 | 0.17772885 | 1.72611876 | 0.08432605 | 0.44526833 |
| Cmtm6 | 5965.6084 | -0.0494306 | 0.16244831 | -0.3042852 | 0.76091061 | 0.94244354 |
| Dync1li1 | 3457.59447 | -0.2309527 | 0.213964 | -1.0793998 | 0.28040952 | 0.71552977 |
| Cmtm7 | 6975.95964 | -0.1783608 | 0.17371816 | -1.0267254 | 0.3045498 | 0.73467909 |
| Stt3b | 3184.92726 | 0.15563288 | 0.44935627 | 0.3463463 | 0.72908248 | 0.93617556 |
| Tgfbr2 | 3529.11307 | 0.29987325 | 0.29137251 | 1.02917484 | 0.30339753 | 0.73436097 |
| Slc25a36 | 1976.98871 | 0.09861036 | 0.18068161 | 0.54576864 | 0.58522498 | 0.8848945 |
| Nmnat3 | 482.290827 | 0.53822466 | 0.32063202 | 1.67863666 | 0.09322288 | 0.4669924 |
| Copb2 | 8262.22457 | -0.1165506 | 0.1616937 | -0.7208108 | 0.47102592 | 0.83381643 |
| Mrps22 | 2816.98579 | 0.13780383 | 0.31135764 | 0.44259017 | 0.6580622 | 0.91258818 |
| Pik3cb | 95.3062751 | -0.9898175 | 0.59971329 | -1.6504844 | 0.09884389 | 0.48025164 |
| Faim | 1531.54133 | -0.3548295 | 0.22097546 | -1.6057415 | 0.10833072 | 0.50301538 |
| Armc8 | 1625.36966 | 0.15659326 | 0.27166137 | 0.57642815 | 0.56432582 | 0.87706744 |
| Dbr1 | 1350.20722 | -0.2033664 | 0.22374048 | -0.908939 | 0.36338234 | 0.77000799 |
| Nck1 | 1633.03572 | 0.47124767 | 0.32971189 | 1.42927108 | 0.15292634 | 0.58128098 |
| Cdc25a | 626.270014 | -0.6116021 | 0.43120302 | -1.4183623 | 0.15608503 | 0.58521232 |
| Nme6 | 2293.96038 | 0.33454952 | 0.23779079 | 1.40690697 | 0.15945498 | 0.5886662 |
| Map4 | 2644.13163 | 0.3795254 | 0.33801671 | 1.12280069 | 0.26152215 | 0.69720616 |
| Dhx30 | 2574.75071 | -0.4899941 | 0.25647146 | -1.9105211 | 0.05606615 | 0.36429573 |
| Smarcc1 | 2180.85344 | -0.1220191 | 0.44387289 | -0.2748965 | 0.78339572 | 0.95081654 |
| Scap | 957.570738 | -0.4486548 | 0.28769514 | -1.55948 | 0.11888282 | 0.52515149 |
| Ptgs2 | 591.417047 | -0.0147108 | 0.55359142 | -0.0265734 | 0.97879999 | 0.99483019 |
| Kif9 | 57.4574464 | -0.0887731 | 1.26834624 | -0.0699913 | 0.94420062 | 0.99005579 |
| Nradd | 48.8071285 | 0.89715849 | 0.91578561 | 0.97965996 | 0.327254 | 0.74811039 |
| Lrrfip2 | 815.869155 | -0.2380906 | 0.27020615 | -0.8811443 | 0.37823975 | 0.78106125 |
| Mlh1 | 2117.43758 | -0.1050304 | 0.234648 | -0.4476085 | 0.65443575 | 0.9108895 |

| | | | | | | |
|---|---|---|---|---|---|---|
| Trib1 | 1367.46396 | 0.55990599 | 0.53989915 | 1.03705661 | 0.29970947 | 0.73128092 |
| Pdcd6ip | 7309.43556 | -0.0285153 | 0.25211863 | -0.1131028 | 0.90994902 | 0.98262295 |
| Fbxl2 | 33.2550743 | 1.12498649 | 1.72301373 | 0.65291789 | 0.51380922 | 0.85497712 |
| Myd88 | 6395.14265 | 0.21968393 | 0.15245666 | 1.44095986 | 0.14959602 | 0.57685486 |
| Wdr48 | 2817.09551 | -0.1773468 | 0.24940293 | -0.7110856 | 0.47703117 | 0.83675398 |
| Gorasp1 | 1554.34002 | 0.15942937 | 0.16812902 | 0.94825614 | 0.34299907 | 0.75947281 |
| Csrnp1 | 237.839027 | 0.35627224 | 0.63731629 | 0.55901952 | 0.5761484 | 0.88091263 |
| Rpsa | 40498.2054 | 0.0664638 | 0.34476384 | 0.19278065 | 0.84713075 | 0.96765585 |
| Slc25a38 | 1091.57007 | 0.02743583 | 0.28699731 | 0.09559613 | 0.92384134 | 0.9849488 |
| Nktr | 901.037021 | -0.0926102 | 0.29243953 | -0.3166814 | 0.75148535 | 0.94018119 |
| Ss18l2 | 819.777563 | 0.32247559 | 0.44620151 | 0.72271292 | 0.46985628 | 0.83381643 |
| Pccb | 520.251163 | 0.37485645 | 0.26151322 | 1.43341299 | 0.15173985 | 0.57988151 |
| Amotl2 | 91.8927582 | -0.3889759 | 0.47445831 | -0.8198316 | 0.41231213 | 0.80251302 |
| Cep63 | 1148.64757 | -0.0984553 | 0.27789637 | -0.3542878 | 0.72312323 | 0.93433544 |
| Trak1 | 270.756613 | -0.1992578 | 0.51191603 | -0.3892392 | 0.69709917 | 0.92650558 |
| Abhd5 | 909.140694 | 0.17129429 | 0.31130105 | 0.55025287 | 0.58214595 | 0.88321857 |
| Ryk | 30.8712956 | 2.49226589 | 1.44702214 | 1.72234123 | 0.08500772 | 0.44637738 |
| Rab6b | 8.79388296 | -1.0504195 | 2.30106346 | -0.4564931 | 0.64803545 | 0.90973357 |
| 1110059G10 | 603.805173 | 0.29253468 | 0.33989734 | 0.86065598 | 0.38942755 | 0.78942657 |
| Srprb | 3557.95772 | -0.2610648 | 0.1638676 | -1.5931448 | 0.11112772 | 0.50888686 |
| Trf | 8045.04542 | 1.3313995 | 0.2303458 | 5.78000343 | 7.469910099 | 1.156701395 |
| Topbp1 | 770.512858 | 0.23485664 | 0.31251552 | 0.75150393 | 0.45234944 | 0.82603347 |
| Uba5 | 2188.28188 | 0.32079663 | 0.27225341 | 1.17830158 | 0.2386764 | 0.67452555 |
| Nphp3 | 68.8098148 | 2.23442454 | 0.97619555 | 2.28891079 | 0.02208454 | 0.21356532 |
| Dnajc13 | 734.312 | -0.3350288 | 0.41419549 | -0.8088663 | 0.41859204 | 0.8052381 |
| Acpp | 15.0183339 | -0.9749341 | 1.50509181 | -0.6477573 | 0.51714197 | 0.85694744 |
| Gnai2 | 2360.24376 | -0.3995776 | 0.19821161 | -2.0159144 | 0.04380893 | 0.32052311 |
| Mrpl3 | 4788.65941 | 0.2713496 | 0.28977947 | 0.93640035 | 0.3490671 | 0.7617085 |
| Nudt16 | 587.354952 | 0.81250873 | 0.34169534 | 2.37787481 | 0.01741274 | 0.18554879 |
| Aste1 | 471.984084 | 0.30847025 | 0.31022705 | 0.99433705 | 0.32005881 | 0.74413737 |
| Atp2c1 | 3668.78864 | 0.20997755 | 0.27643836 | 0.7595818 | 0.4475046 | 0.82308282 |
| Pik3r4 | 757.353721 | 0.15908751 | 0.29580273 | 0.53781625 | 0.59070391 | 0.88750688 |
| Manf | 7415.98551 | 0.07696714 | 0.18778611 | 0.409866 | 0.68190425 | 0.92061965 |
| Mapkapk3 | 2850.4378 | -1.1901165 | 0.17549303 | -6.7815598 | 1.188852510 | 2.796775530 |
| Cish | 56.1217966 | -0.8555556 | 0.74869826 | -1.1427242 | 0.25315311 | 0.68647781 |
| Hemk1 | 196.825994 | -0.3751433 | 0.51650535 | -0.7263106 | 0.46764837 | 0.83381643 |
| Rbm5 | 3833.93305 | -0.0137229 | 0.14646725 | -0.0936929 | 0.9253531 | 0.98571293 |
| Rbm6 | 1914.49116 | -0.3688972 | 0.26997576 | -1.3664085 | 0.17181079 | 0.60430173 |
| Mon1a | 289.514956 | -0.7754635 | 0.49704131 | -1.5601591 | 0.11872229 | 0.52515149 |
| Traip | 1471.81013 | -0.0487728 | 0.29403168 | -0.1658759 | 0.86825458 | 0.97185088 |
| Bsn | 116.651424 | -1.460678 | 0.45252278 | -3.2278552 | 0.00124722 | 0.02744111 |
| Apeh | 2787.60029 | -0.1357948 | 0.16648183 | -0.8156737 | 0.41468681 | 0.80395688 |
| Amigo3 | 41.5149178 | 1.00780011 | 1.80673916 | 0.55780056 | 0.57698058 | 0.88121027 |
| Ip6k1 | 655.320879 | 0.35861516 | 0.31246419 | 1.14770004 | 0.25109241 | 0.6842534 |
| Cdhr4 | 62.0583226 | 1.10495068 | 0.69517388 | 1.58945942 | 0.11195671 | 0.50987629 |
| Uba7 | 826.652683 | 0.33112404 | 0.3838328 | 0.86267782 | 0.38831463 | 0.78942657 |

| | | | | | | |
|---|---|---|---|---|---|---|
| Nckipsd | 615.87821 | -0.2126317 | 0.31716456 | -0.6704145 | 0.50259359 | 0.85012672 |
| Ip6k2 | 111.496544 | 0.15127243 | 0.54661333 | 0.27674485 | 0.78197603 | 0.95004728 |
| Prkar2a | 1151.50559 | -0.3236382 | 0.32116375 | -1.0077046 | 0.31359629 | 0.74065408 |
| Slc25a20 | 986.84904 | 0.03653677 | 0.29836691 | 0.12245582 | 0.90253803 | 0.98043702 |
| Qars | 4462.36975 | -0.4546445 | 0.19132592 | -2.3762827 | 0.01748806 | 0.18560042 |
| Nicn1 | 268.459276 | 0.7578821 | 0.60841293 | 1.2456706 | 0.21288535 | 0.64596443 |
| Amt | 123.803879 | 0.84981883 | 0.6348548 | 1.33860346 | 0.1806998 | 0.61122544 |
| Klhdc8b | 13.0067347 | 0.7416112 | 1.02677175 | 0.72227464 | 0.47012564 | 0.83381643 |
| Usp4 | 5650.07275 | -0.1997882 | 0.24097357 | -0.8290877 | 0.40705476 | 0.79992362 |
| Nt5m | 925.990956 | 0.40520377 | 0.22061398 | 1.83670937 | 0.06625281 | 0.39670614 |
| Srek1 | 1660.78127 | -0.4165943 | 0.23626074 | -1.763282 | 0.07785293 | 0.4314473 |
| Eml4 | 1075.72673 | -0.7020892 | 0.29749245 | -2.3600234 | 0.01827378 | 0.19092573 |
| Flcn | 4995.14401 | 0.34168095 | 0.38370249 | 0.89048404 | 0.37320604 | 0.77748705 |
| Atxn2l | 1666.05456 | -0.222447 | 0.41695905 | -0.5334984 | 0.59368864 | 0.88797037 |
| Chsy1 | 550.383072 | -0.6650592 | 0.30809505 | -2.1586168 | 0.0308799 | 0.26251138 |
| Gpr19 | 173.618805 | 0.08964001 | 0.39823795 | 0.22509157 | 0.82190804 | 0.96232782 |
| Fhl3 | 130.540119 | 0.60775492 | 0.39972304 | 1.52044003 | 0.12840042 | 0.54369067 |
| Pygm | 152.970486 | 1.16409407 | 0.63242772 | 1.84067527 | 0.06566916 | 0.39558985 |
| Crebl2 | 27.8914675 | 0.80600845 | 0.7772656 | 1.03697945 | 0.29974544 | 0.73128092 |
| Fam189b | 58.2354618 | 0.21338465 | 0.75246872 | 0.28357942 | 0.77673271 | 0.94800948 |
| 1700025G04 | 128.894745 | -1.0621518 | 0.45317891 | -2.3437803 | 0.01908941 | 0.19530399 |
| Pon2 | 3849.03939 | 0.65048163 | 0.20867779 | 3.11715799 | 0.00182604 | 0.0375427 |
| Prorsd1 | 1252.46178 | 0.40964144 | 0.35662092 | 1.14867472 | 0.25069014 | 0.6842534 |
| Cd59a | 139.1959 | 2.38322509 | 0.79589167 | 2.99440888 | 0.00274977 | 0.05043172 |
| 6820408C15 | 5.58945267 | 0.93456081 | 2.0516853 | 0.45550885 | 0.6487432 | 0.90973357 |
| Malt1 | 247.357172 | -0.1361709 | 0.41278716 | -0.3298817 | 0.74148935 | 0.93800224 |
| Oas2 | 62.8897315 | 1.23143767 | 0.80941533 | 1.52139159 | 0.1281616 | 0.54327949 |
| Nlrp3 | 542.976354 | 0.44459479 | 0.57522995 | 0.77289923 | 0.43958202 | 0.81963526 |
| Lmo2 | 804.634458 | -0.1580481 | 0.34797585 | -0.4541929 | 0.64969002 | 0.90973357 |
| Exd2 | 648.672512 | 0.42855852 | 0.31725527 | 1.35083184 | 0.17674931 | 0.60784615 |
| Resf1 | 2079.53206 | 0.15206707 | 0.33563648 | 0.45307075 | 0.6504978 | 0.91021183 |
| Syde1 | 27.6730651 | 2.03710668 | 1.63099096 | 1.24899937 | 0.2116653 | 0.64559288 |
| Trib3 | 72.2400454 | 0.83155507 | 0.83231444 | 0.99908764 | 0.31775224 | 0.74307991 |
| Mdfi | 12.4024618 | 0.7876075 | 1.57399766 | 0.5003867 | 0.61680281 | 0.89878056 |
| Mansc1 | 41.2400541 | -0.1947938 | 0.69553395 | -0.2800637 | 0.77942864 | 0.9488468 |
| Abtb2 | 57.9641379 | -0.6407862 | 1.2207162 | -0.5249264 | 0.59963434 | 0.8902352 |
| Folr2 | 462.606435 | 3.28215537 | 0.39394503 | 8.33150606 | 7.982035349 | 4.438374473 |
| Mier3 | 527.586499 | -0.870292 | 0.45559809 | -1.9102187 | 0.05610506 | 0.36429573 |
| Snx33 | 67.5792471 | -0.4867912 | 0.76631033 | -0.6352403 | 0.52527168 | 0.86045516 |
| Inppl1 | 454.120581 | 0.59133838 | 0.48317853 | 1.22385071 | 0.22100855 | 0.65462412 |
| Pram1 | 343.71411 | -1.3717458 | 0.36049386 | -3.8051849 | 0.0001417 | 0.00485545 |
| Ccdc88a | 597.86065 | 0.21685025 | 0.40683112 | 0.53302277 | 0.59401782 | 0.88797037 |
| Tpcn1 | 421.268363 | 0.16731153 | 0.3391988 | 0.49325507 | 0.62183237 | 0.89912494 |
| D430042O09 | 131.516869 | -0.1266071 | 0.5315036 | -0.2382055 | 0.81172171 | 0.95967622 |
| Gpbp1 | 3244.73925 | -0.0975627 | 0.22894093 | -0.4261477 | 0.67000021 | 0.91689368 |
| Gab3 | 711.971973 | 1.07854632 | 0.38849195 | 2.77623855 | 0.00549918 | 0.08595935 |

| | | | | | | |
|---|---|---|---|---|---|---|
| Slc8b1 | 1227.03832 | -0.324577 | 0.25764691 | -1.2597746 | 0.20775067 | 0.6418276 |
| Bet1 | 1455.75519 | 0.41883092 | 0.28967859 | 1.44584697 | 0.14822013 | 0.57506401 |
| Ilvbl | 1181.84549 | 0.23770632 | 0.26715537 | 0.88976808 | 0.37359043 | 0.77748705 |
| Gng11 | 109.302269 | 1.82640524 | 0.73198244 | 2.49514897 | 0.01259043 | 0.14937799 |
| Trpa1 | 10.416194 | 5.81362175 | 1.87516455 | 3.10032618 | 0.00193308 | 0.03908648 |
| Gtf3c1 | 603.646388 | -0.2038733 | 0.20840915 | -0.978236 | 0.32795762 | 0.74843736 |
| Cntrob | 119.128581 | -0.4706594 | 0.59886912 | -0.7859136 | 0.4319181 | 0.81344118 |
| Troap | 1035.10543 | -0.1535861 | 0.257059 | -0.597474 | 0.55019096 | 0.87093412 |
| Alas1 | 3599.4325 | -0.4406792 | 0.21148862 | -2.0837019 | 0.0371873 | 0.28956966 |
| Pdxk | 2455.91464 | -0.1459244 | 0.171639 | -0.8501819 | 0.39522395 | 0.79310606 |
| Srxn1 | 179.661605 | 0.52611304 | 0.44417512 | 1.18447211 | 0.23622623 | 0.6718799 |
| Cdv3 | 10190.3415 | -0.3832585 | 0.2668476 | -1.4362449 | 0.15093266 | 0.57861461 |
| Slc10a3 | 626.831668 | -0.3327236 | 0.23850273 | -1.3950516 | 0.16300027 | 0.59424967 |
| Arap1 | 3718.76396 | 0.4087973 | 0.27569032 | 1.48281341 | 0.13812399 | 0.56068131 |
| Fanca | 673.176107 | -0.9711168 | 0.36100158 | -2.6900624 | 0.00714387 | 0.10299472 |
| Ank2 | 24.3733195 | 0.31251882 | 1.48588851 | 0.21032454 | 0.83341438 | 0.96565375 |
| Ppp1r9a | 272.054234 | 1.01836005 | 0.70136779 | 1.45196296 | 0.14651191 | 0.57365079 |
| Pwp2 | 1697.10971 | -0.6656732 | 0.25900926 | -2.5700747 | 0.01016766 | 0.13017076 |
| 2410131K14R | 606.397962 | 0.30552592 | 0.26339979 | 1.1599323 | 0.24607637 | 0.68117517 |
| Prr5l | 235.982828 | -0.0756361 | 0.39799446 | -0.190043 | 0.84927546 | 0.96765585 |
| Abcc10 | 89.4973068 | -0.5291745 | 0.51588437 | -1.0257619 | 0.30500383 | 0.73510332 |
| Abcc4 | 136.770634 | -0.5391746 | 0.61755378 | -0.8730812 | 0.3826188 | 0.78559512 |
| Pkd1 | 140.098339 | -0.8697334 | 0.67196663 | -1.2943104 | 0.19555823 | 0.62739675 |
| P2ry2 | 574.372352 | -0.5600896 | 0.38531703 | -1.4535811 | 0.14606247 | 0.5731306 |
| Fbxw8 | 607.263122 | -0.3492198 | 0.28002327 | -1.24711 | 0.21235715 | 0.64580271 |
| Psmf1 | 1920.46239 | -0.117515 | 0.18801556 | -0.6250281 | 0.53195264 | 0.8643494 |
| Smap2 | 421.648971 | -0.6353032 | 0.3686174 | -1.723476 | 0.08480249 | 0.44599693 |
| Cyb5r4 | 3926.84105 | -0.1017728 | 0.20977883 | -0.4851435 | 0.62757453 | 0.9022352 |
| Acsl3 | 935.285107 | -0.1275105 | 0.21982526 | -0.5800537 | 0.56187838 | 0.87540592 |
| Rims3 | 5.90662398 | 0.66678686 | 2.87977275 | 0.23154149 | 0.81689416 | 0.96079676 |
| Rangrf | 746.472155 | -0.3783934 | 0.34434319 | -1.0988844 | 0.27181848 | 0.70940613 |
| Nfyc | 3298.97741 | -0.4342175 | 0.25805229 | -1.6826727 | 0.09243847 | 0.46597694 |
| Fbxo21 | 1026.94061 | -0.2325829 | 0.20648718 | -1.1263793 | 0.26000501 | 0.69625824 |
| Slc16a1 | 7206.25362 | -0.2532939 | 0.18144493 | -1.395982 | 0.16271991 | 0.59419484 |
| Atg12 | 2452.58166 | 0.3233802 | 0.2387169 | 1.35465989 | 0.17552594 | 0.60763832 |
| Lrig2 | 350.542479 | 0.46608424 | 0.28258595 | 1.64935391 | 0.09907515 | 0.48073092 |
| Adgre4 | 658.288016 | -1.5256479 | 0.33216946 | -4.5929805 | 4.369599025 | 0.00025948 |
| Hspa13 | 890.177251 | 0.31687263 | 0.21981365 | 1.44155122 | 0.14942902 | 0.57685486 |
| Nup93 | 6212.27791 | -0.3037291 | 0.20989959 | -1.4470211 | 0.14789102 | 0.57472133 |
| Rasgrp2 | 489.483549 | -0.4309905 | 0.41376179 | -1.0416391 | 0.29757901 | 0.72979999 |
| Ap4b1 | 1240.53438 | -0.106607 | 0.26924563 | -0.3959469 | 0.69214422 | 0.92539651 |
| Pebp1 | 13801.134 | 0.15042099 | 0.31842783 | 0.47238646 | 0.63665097 | 0.90542378 |
| Ift57 | 324.196294 | -0.3136973 | 0.47648833 | -0.6583524 | 0.51031169 | 0.85335831 |
| Fkbp1a | 21978.5865 | -0.0453658 | 0.13880849 | -0.3268228 | 0.74380192 | 0.93880767 |
| Fam207a | 2402.12791 | -0.1566351 | 0.20333573 | -0.7703277 | 0.44110555 | 0.81967485 |
| Chpf | 11.8323689 | -1.5947313 | 1.46228928 | -1.0905717 | 0.27546138 | 0.71181528 |

| | | | | | | |
|---|---|---|---|---|---|---|
| Foxj3 | 345.095926 | 0.44095088 | 0.36687396 | 1.20191383 | 0.22939692 | 0.66356408 |
| Mycbp2 | 755.625759 | -0.4464908 | 0.24358689 | -1.8329837 | 0.06680499 | 0.39767663 |
| Ogfod1 | 307.704304 | -0.7383374 | 0.7263388 | -1.0165193 | 0.30938218 | 0.73890515 |
| Trim33 | 185.293597 | -0.2870676 | 0.60667747 | -0.47318 | 0.63608478 | 0.90531997 |
| Nfatc1 | 885.998939 | -0.1608092 | 0.33628465 | -0.4781938 | 0.63251225 | 0.90423307 |
| Polr2f | 2322.24325 | -0.109936 | 0.26489534 | -0.4150166 | 0.67812975 | 0.91968528 |
| Gmppa | 1638.63351 | 0.23386382 | 0.22378012 | 1.04506072 | 0.29599491 | 0.72859877 |
| Cdo1 | 29.1996026 | 0.29837523 | 0.77140349 | 0.38679528 | 0.69890777 | 0.92669304 |
| Cip2a | 1795.98858 | 0.20463727 | 0.19126297 | 1.06992622 | 0.28465252 | 0.72019737 |
| Afap1l1 | 1377.03484 | 0.49438358 | 0.35582472 | 1.38940202 | 0.16471054 | 0.59700859 |
| Calhm2 | 274.073121 | -0.2275848 | 0.28537722 | -0.7974876 | 0.42516789 | 0.80988459 |
| Gm7879 | 20.0177847 | 0.0992421 | 1.19248002 | 0.08322329 | 0.933674 | 0.98756396 |
| Micall1 | 100.418969 | 0.14929393 | 0.53928832 | 0.27683509 | 0.78190673 | 0.95004728 |
| Eif3l | 17768.5032 | -0.2444376 | 0.16131853 | -1.5152481 | 0.12970959 | 0.54639717 |
| Npat | 922.977925 | -0.0803067 | 0.28917237 | -0.2777123 | 0.78123323 | 0.94998271 |
| Ankrd54 | 1745.68288 | -0.0743903 | 0.20381635 | -0.364987 | 0.71512108 | 0.93235217 |
| Pygb | 4043.07837 | 0.16562628 | 0.26946578 | 0.6146468 | 0.538788 | 0.86689381 |
| Pfkm | 378.109787 | -0.0200041 | 0.28238907 | -0.0708389 | 0.94352597 | 0.98989307 |
| Gas7 | 11724.7627 | 0.1494103 | 0.2821562 | 0.52953045 | 0.59643753 | 0.88880744 |
| Entpd6 | 2070.01925 | 0.0248807 | 0.20009196 | 0.12434635 | 0.90104105 | 0.98043702 |
| Senp1 | 1307.3333 | -0.2015407 | 0.24820357 | -0.8119975 | 0.41679306 | 0.8047849 |
| Tbc1d4 | 390.639226 | 0.67038623 | 0.28692616 | 2.33644167 | 0.01946823 | 0.19765383 |
| Triobp | 1459.61446 | 0.10529305 | 0.17405051 | 0.60495685 | 0.54520767 | 0.86970532 |
| Apmap | 3770.99099 | -0.398622 | 0.17107753 | -2.3300664 | 0.01980264 | 0.19950359 |
| Nol12 | 1545.67301 | -0.1424088 | 0.26725602 | -0.5328555 | 0.5941336 | 0.88797037 |
| Cdc14b | 492.861908 | 0.32398003 | 0.30966907 | 1.04621372 | 0.29546237 | 0.72852118 |
| Lss | 1424.38778 | 0.78879777 | 0.3420314 | 2.30621449 | 0.02109865 | 0.20747571 |
| Slc7a6os | 601.867781 | 0.35856998 | 0.31046423 | 1.15494779 | 0.24811181 | 0.68235696 |
| Rnf125 | 78.7978188 | -0.2067286 | 0.70326232 | -0.2939567 | 0.76879102 | 0.94401697 |
| 3830406C13 | 798.913816 | -0.1101198 | 0.26380324 | -0.4174315 | 0.67636282 | 0.91889316 |
| Slc35d2 | 695.22723 | 0.82051908 | 0.45067273 | 1.82065392 | 0.06865948 | 0.40407556 |
| Atg9a | 473.957061 | 0.19893311 | 0.26541566 | 0.74951536 | 0.45354665 | 0.8263682 |
| Ybey | 323.245947 | -0.0511016 | 0.30956122 | -0.1650776 | 0.86888291 | 0.97202055 |
| Gga1 | 914.790552 | -0.0366439 | 0.21777395 | -0.1682658 | 0.86637421 | 0.97060021 |
| Slc22a15 | 85.4322647 | -0.5730793 | 0.55814332 | -1.0267601 | 0.30453345 | 0.73467909 |
| Phldb2 | 17.409645 | 3.34092734 | 1.99089774 | 1.67810092 | 0.0933274 | 0.46713342 |
| Podxl2 | 11.7778704 | 0.53917117 | 1.59269243 | 0.33852812 | 0.73496524 | 0.93750666 |
| Abhd10 | 1067.87118 | -0.0073929 | 0.24667794 | -0.0299698 | 0.97609113 | 0.994746 |
| Cnppd1 | 1269.35051 | 0.07772136 | 0.26721411 | 0.29085799 | 0.77115993 | 0.94488554 |
| Atp1a1 | 2072.93986 | -0.2025354 | 0.34360529 | -0.5894422 | 0.5555647 | 0.87235279 |
| Dis3 | 2343.2983 | -0.4171039 | 0.18321464 | -2.2765862 | 0.02281095 | 0.2163471 |
| Card10 | 15.3426142 | -0.6510579 | 1.98493868 | -0.327999 | 0.74291241 | 0.938557 |
| Mgll | 10.1345713 | -1.3808678 | 2.02989562 | -0.6802654 | 0.49633641 | 0.84787395 |
| Tmed7 | 1056.91544 | 0.16160041 | 0.40400305 | 0.399998 | 0.68915799 | 0.92470267 |
| Mzt1 | 1429.83253 | 0.14339409 | 0.26656004 | 0.53794294 | 0.59061644 | 0.88748446 |
| Tie1 | 322.336597 | 0.52792625 | 0.55236462 | 0.95575682 | 0.33919511 | 0.75683748 |

| | | | | | | |
|---|---|---|---|---|---|---|
| Lpcat2 | 2956.69959 | -0.4081122 | 0.19517334 | -2.0910242 | 0.03652589 | 0.28679157 |
| Ttc28 | 60.8796609 | 0.69563196 | 0.82795406 | 0.84018183 | 0.40080644 | 0.79698916 |
| Eefsec | 2782.64126 | -0.3899344 | 0.21606467 | -1.8047115 | 0.07111984 | 0.41122734 |
| Rac2 | 19482.4875 | -0.4016495 | 0.20900386 | -1.9217326 | 0.05463942 | 0.36033644 |
| Ttf2 | 940.490186 | -0.1240618 | 0.32910003 | -0.3769728 | 0.70619381 | 0.92896305 |
| Wnt6 | 14.34038 | -1.1216819 | 1.22069089 | -0.918891 | 0.35815257 | 0.76676242 |
| Scaf11 | 3816.54263 | -0.2166347 | 0.3573078 | -0.606297 | 0.54431753 | 0.8695007 |
| Trim45 | 44.1559774 | -0.7593497 | 0.70819034 | -1.0722396 | 0.28361244 | 0.71951357 |
| Arid2 | 565.511768 | -0.0749222 | 0.25406125 | -0.2948981 | 0.76807173 | 0.94401697 |
| Szt2 | 109.587712 | -0.2974079 | 0.56851936 | -0.5231272 | 0.60088576 | 0.89066224 |
| Gm5134 | 14.5721157 | 0.03996621 | 1.65110925 | 0.02420567 | 0.98068855 | 0.99516906 |
| Ttll4 | 673.654126 | -0.3015272 | 0.27550401 | -1.0944567 | 0.27375474 | 0.71078816 |
| Slc35a4 | 12767.8296 | -0.361264 | 0.2451021 | -1.4739327 | 0.14049972 | 0.56425905 |
| Stk36 | 23.4582357 | -1.1621702 | 1.37848602 | -0.8430773 | 0.39918523 | 0.79611427 |
| Ptprm | 23.448386 | 0.26025147 | 1.56293006 | 0.16651511 | 0.86775159 | 0.97155457 |
| Rpgrip1l | 124.871892 | 1.27689959 | 0.87058289 | 1.46671801 | 0.14245278 | 0.56818549 |
| Wdr3 | 3758.02247 | -0.6192893 | 0.20376926 | -3.0391695 | 0.00237231 | 0.04520328 |
| Noc4l | 4644.46746 | -0.5228447 | 0.21590714 | -2.4216185 | 0.01545156 | 0.17214842 |
| Ptprf | 36.1356272 | -0.0403491 | 1.1585618 | -0.0348269 | 0.97221778 | 0.99443087 |
| Lpp | 1016.7537 | 0.65961465 | 0.34491652 | 1.91238928 | 0.05582628 | 0.36345019 |
| Mif | 19491.3417 | -0.3345676 | 0.14168021 | -2.3614281 | 0.0182047 | 0.19092573 |
| Fbxl8 | 97.1635474 | -0.0254965 | 0.65511945 | -0.0389189 | 0.96895506 | 0.99443087 |
| Gstt2 | 182.514573 | 0.63320497 | 0.58305829 | 1.08600628 | 0.27747621 | 0.71260013 |
| Fem1c | 1328.48235 | -0.009634 | 0.24528245 | -0.039277 | 0.96866955 | 0.99443087 |
| Ctdp1 | 1626.39179 | -0.0821519 | 0.40681624 | -0.2019387 | 0.83996468 | 0.96763235 |
| Kdm4a | 49.6529565 | 0.62212239 | 0.88984962 | 0.69913205 | 0.48446951 | 0.84135655 |
| Dnm2 | 7361.52899 | 0.17723764 | 0.21903303 | 0.80918223 | 0.41841034 | 0.8052381 |
| Chst2 | 10.9247479 | 2.34379593 | 2.015265 | 1.1630212 | 0.2448209 | 0.68039867 |
| Map2k4 | 2035.37468 | -0.332697 | 0.20235216 | -1.6441486 | 0.1001455 | 0.48364783 |
| Rtp4 | 233.820801 | 1.35289949 | 0.68474805 | 1.97576246 | 0.04818169 | 0.33815639 |
| Pus7l | 691.316973 | -0.1624738 | 0.27061432 | -0.6003888 | 0.54824717 | 0.87059427 |
| Usp37 | 2739.70508 | -0.069225 | 0.33672113 | -0.2055855 | 0.83711468 | 0.96683245 |
| Ipo13 | 2656.4824 | 0.01426981 | 0.17997188 | 0.07928913 | 0.93680265 | 0.98843258 |
| Fntb | 825.006886 | 0.24029014 | 0.27880578 | 0.86185493 | 0.38876735 | 0.78942657 |
| Atp6v0b | 10353.6403 | 0.74097608 | 0.39750596 | 1.8640628 | 0.06231289 | 0.38520862 |
| Trappc8 | 434.669205 | -0.3112847 | 0.36923283 | -0.843058 | 0.39919602 | 0.79611427 |
| Frrs1 | 2386.98563 | -0.4041945 | 0.20253149 | -1.9957118 | 0.0459653 | 0.33134561 |
| Clasp2 | 2193.2017 | -0.084882 | 0.31984765 | -0.2653826 | 0.79071477 | 0.95261385 |
| Spg11 | 471.475883 | 0.05868742 | 0.27228239 | 0.21553881 | 0.82934725 | 0.96462215 |
| Agl | 298.197621 | -0.2618178 | 0.44608355 | -0.5869254 | 0.5572538 | 0.87276537 |
| Nudt15 | 611.742084 | -0.1411391 | 0.30788455 | -0.4584157 | 0.64665381 | 0.90925224 |
| Ctdspl2 | 1460.30889 | -0.0023929 | 0.21680151 | -0.0110373 | 0.99119365 | 0.99689813 |
| Gucd1 | 1108.74483 | 0.33231445 | 0.23036453 | 1.44255913 | 0.1491447 | 0.57645724 |
| Cacul1 | 2136.67251 | 0.4428001 | 0.17809167 | 2.48636056 | 0.01290571 | 0.15209595 |
| Eri3 | 1471.70804 | 0.07955219 | 0.22619174 | 0.35170244 | 0.72506143 | 0.93542036 |
| Mcee | 826.526351 | 0.46029726 | 0.36254227 | 1.2696375 | 0.20421379 | 0.63846817 |

| | | | | | | |
|---|---|---|---|---|---|---|
| Terf2ip | 858.952544 | -0.2407439 | 0.24540201 | -0.9810183 | 0.32658374 | 0.74811039 |
| Gtpbp6 | 303.038551 | 0.06070012 | 0.30616331 | 0.19826058 | 0.84284119 | 0.96765585 |
| Armcx2 | 508.745443 | 0.1040963 | 0.28988021 | 0.3591011 | 0.71951946 | 0.93314808 |
| Trmt13 | 236.523142 | 0.16209915 | 0.29632281 | 0.54703569 | 0.58435422 | 0.88404713 |
| Specc1l | 810.500373 | 0.15668066 | 0.39258784 | 0.39909707 | 0.68982168 | 0.92470267 |
| Lpar6 | 1519.83748 | -0.1606347 | 0.17619355 | -0.9116946 | 0.36192953 | 0.76932823 |
| Tagap | 1048.12881 | -0.7776219 | 0.32966136 | -2.3588505 | 0.01833164 | 0.19092573 |
| Zbtb1 | 1255.94197 | 0.19505262 | 0.25805351 | 0.75586114 | 0.44973245 | 0.82484234 |
| Fan1 | 12.4041723 | -0.4787771 | 2.08915363 | -0.2291727 | 0.81873466 | 0.96105733 |
| Armcx1 | 34.778179 | 3.0585387 | 1.58733081 | 1.9268439 | 0.05399909 | 0.35989907 |
| Crlf2 | 91.195863 | -0.7704996 | 0.54612286 | -1.410854 | 0.15828768 | 0.5871574 |
| Tomm6 | 1717.2771 | -0.3824845 | 0.1879751 | -2.0347617 | 0.04187485 | 0.31216033 |
| Fam160b1 | 206.214833 | -0.4459755 | 0.86103288 | -0.5179541 | 0.6044903 | 0.89258167 |
| Catsper2 | 19.8352939 | 1.42389455 | 1.48941299 | 0.95601056 | 0.3390669 | 0.75683748 |
| Fndc3a | 1001.40145 | 0.18150193 | 0.32690833 | 0.55520741 | 0.5787528 | 0.88255834 |
| Cryzl2 | 611.676529 | 0.2452592 | 0.29442706 | 0.83300496 | 0.40484193 | 0.79917613 |
| Larp4b | 2435.1843 | -0.3614176 | 0.37170465 | -0.9723247 | 0.33088905 | 0.75011761 |
| Cdc14a | 204.062068 | -0.3974281 | 0.41775214 | -0.951349 | 0.34142724 | 0.75846782 |
| Ppip5k1 | 168.244089 | 0.24944403 | 0.42981694 | 0.58034945 | 0.56167899 | 0.87540592 |
| Ttc7b | 139.21912 | -0.3619286 | 0.49762086 | -0.7273179 | 0.46703124 | 0.8334534 |
| Casp4 | 107.031597 | 0.37178652 | 0.50059898 | 0.74268333 | 0.45767343 | 0.82968569 |
| Idua | 1325.17862 | 0.53155211 | 0.25850581 | 2.05624818 | 0.0397586 | 0.30227903 |
| Arhgef5 | 8.63580253 | 2.04150979 | 2.09779745 | 0.97316821 | 0.33046971 | 0.75002522 |
| Gtf2a2 | 2820.90813 | 0.31953831 | 0.37339109 | 0.85577379 | 0.39212292 | 0.79038386 |
| Znrf1 | 86.7676688 | -0.2164866 | 0.44496097 | -0.4865294 | 0.62659189 | 0.90198854 |
| Dph5 | 1776.68921 | 0.11881437 | 0.23504262 | 0.50550141 | 0.61320671 | 0.89739894 |
| Fam20b | 529.477826 | 0.15364127 | 0.27300477 | 0.56277869 | 0.57358559 | 0.88049599 |
| Rbfox2 | 51.1784329 | -2.9586436 | 1.09076648 | -2.7124445 | 0.0066789 | 0.09915404 |
| Myo6 | 184.716938 | 0.08680046 | 0.53595221 | 0.16195559 | 0.87134083 | 0.97272425 |
| Igf2bp2 | 750.728068 | 0.26426276 | 0.22746751 | 1.16176047 | 0.24533278 | 0.6805342 |
| Ndn | 269.856575 | 0.90249687 | 0.40015202 | 2.25538503 | 0.02410917 | 0.22455944 |
| Reep4 | 2169.97602 | -0.1598444 | 0.19785486 | -0.8078873 | 0.41915546 | 0.80529433 |
| Spata2l | 100.673973 | 0.07178073 | 0.51740318 | 0.13873268 | 0.88966139 | 0.97870754 |
| Rfwd3 | 5035.38561 | -0.1499271 | 0.2685549 | -0.5582735 | 0.57665765 | 0.88105119 |
| Pank1 | 145.065067 | 0.18053359 | 0.54066633 | 0.33390943 | 0.7384479 | 0.93800224 |
| Map3k13 | 35.8081146 | 0.78772179 | 0.63723251 | 1.23616069 | 0.21639884 | 0.64982959 |
| Pcgf3 | 575.380916 | -0.1866669 | 0.33096214 | -0.5640128 | 0.57274544 | 0.88035336 |
| Pdpr | 114.988335 | 0.41830511 | 0.52655275 | 0.79442205 | 0.42694977 | 0.81021713 |
| Pik3c3 | 1404.14351 | 0.02710629 | 0.17537987 | 0.15455762 | 0.87717009 | 0.97458574 |
| Hacd3 | 4162.98567 | 0.24407925 | 0.1954304 | 1.24893184 | 0.21169001 | 0.64559288 |
| AW554918 | 108.38637 | 1.70140363 | 0.49519699 | 3.43581176 | 0.00059078 | 0.01540944 |
| Piwil2 | 147.757904 | 0.14386626 | 0.52325629 | 0.27494417 | 0.78335913 | 0.95081654 |
| Vps8 | 869.620978 | 0.32374873 | 0.31260646 | 1.0356431 | 0.30036868 | 0.73158785 |
| Ddx19b | 728.405973 | -0.1516107 | 0.2163478 | -0.7007731 | 0.4834446 | 0.84112897 |
| Zfp7 | 125.81605 | 0.1729228 | 0.51951989 | 0.33285117 | 0.73924663 | 0.93800224 |
| Cep350 | 335.911738 | 0.03302133 | 0.51133983 | 0.06457806 | 0.94850996 | 0.99019648 |

| | | | | | | |
|---|---|---|---|---|---|---|
| Qsox1 | 439.536455 | 0.43458557 | 0.32891587 | 1.32126667 | 0.18641246 | 0.61862792 |
| Ucp2 | 13602.9639 | -0.5948857 | 0.21235406 | -2.8013863 | 0.00508836 | 0.08094393 |
| Arhgap39 | 196.007251 | 0.40336361 | 0.58512172 | 0.68936701 | 0.49059234 | 0.84477968 |
| Acbd6 | 1250.7582 | -0.3312129 | 0.22238448 | -1.4893705 | 0.13638983 | 0.5587598 |
| Fuk | 135.741571 | -0.0964347 | 0.4184239 | -0.2304713 | 0.81772555 | 0.96079676 |
| Stard9 | 100.936648 | 0.36656521 | 0.67776649 | 0.54084293 | 0.58861585 | 0.88690478 |
| Smyd5 | 3707.6461 | -0.5049143 | 0.2801232 | -1.8024721 | 0.07147116 | 0.41122734 |
| Ccar2 | 1887.20401 | -0.2961224 | 0.26903649 | -1.1006775 | 0.27103706 | 0.70830941 |
| Foxn3 | 1123.51203 | 0.39191541 | 0.39226897 | 0.99909868 | 0.31774689 | 0.74307991 |
| Sfxn5 | 1241.25205 | 0.54204507 | 0.20311933 | 2.66860411 | 0.00761672 | 0.10746862 |
| Vav3 | 416.566877 | 0.32538398 | 0.3934758 | 0.82694789 | 0.4082666 | 0.80087574 |
| Lrrc14 | 916.330866 | -0.1126314 | 0.2653625 | -0.4244434 | 0.67124247 | 0.91711586 |
| Egr3 | 112.851688 | -0.3949354 | 0.83188041 | -0.4747502 | 0.63496504 | 0.90531997 |
| Sf3b3 | 11309.4306 | -0.2472213 | 0.22574445 | -1.0951378 | 0.27345625 | 0.71048723 |
| Spr | 857.21005 | 0.17082826 | 0.28641426 | 0.59643769 | 0.55088287 | 0.87093412 |
| Fndc3c1 | 45.6161322 | 1.24989392 | 0.65553989 | 1.90666342 | 0.05656419 | 0.36524665 |
| Fkbpl | 155.61874 | -0.3544045 | 0.4689357 | -0.7557635 | 0.44979101 | 0.82484234 |
| Gadd45gip1 | 2358.06107 | 0.06349786 | 0.34954912 | 0.18165646 | 0.85585234 | 0.96915653 |
| Mnd1 | 155.460139 | -0.3311844 | 0.36014867 | -0.9195768 | 0.35779396 | 0.76674184 |
| Rbm4b | 771.340137 | 0.00075232 | 0.30406553 | 0.00247419 | 0.99802588 | 0.99967651 |
| Recql4 | 324.61899 | -0.5077502 | 0.34049068 | -1.4912309 | 0.13590088 | 0.55769054 |
| Mtss1l | 11.769789 | 0.14745233 | 1.69689497 | 0.08689538 | 0.93075467 | 0.98716777 |
| Tmem131l | 930.255038 | 0.11281053 | 0.29028034 | 0.38862615 | 0.69755272 | 0.92667877 |
| Exoc6b | 196.27604 | 0.74042434 | 0.33143637 | 2.23398642 | 0.02548397 | 0.2324723 |
| Rpap2 | 753.327853 | 0.38226955 | 0.30060093 | 1.27168451 | 0.20348524 | 0.63809363 |
| Tlr13 | 2193.27097 | 0.49065213 | 0.20041697 | 2.44815665 | 0.01435892 | 0.16370241 |
| Asb13 | 163.398195 | 0.35469022 | 0.68066144 | 0.5210964 | 0.60229962 | 0.89130082 |
| Tubgcp5 | 2087.79419 | 0.64905191 | 0.27322409 | 2.37552961 | 0.01752379 | 0.18560042 |
| Atp7a | 1085.79497 | 0.42282249 | 0.36887732 | 1.1462415 | 0.25169524 | 0.68505999 |
| Atp6v1h | 6534.83994 | 0.19037569 | 0.14053643 | 1.3546359 | 0.17553359 | 0.60763832 |
| Tasor2 | 91.370606 | 0.42758699 | 0.58988011 | 0.72487101 | 0.46853117 | 0.83381643 |
| Tmem87a | 118.345675 | -1.1031946 | 0.42992667 | -2.5660065 | 0.01028769 | 0.13082049 |
| Alg3 | 1511.60533 | -0.3692975 | 0.27139551 | -1.3607356 | 0.17359727 | 0.60653633 |
| Tcea1 | 4977.74257 | 0.23969359 | 0.31246724 | 0.76709992 | 0.44302211 | 0.82038676 |
| Ppp1r16a | 504.065685 | 0.03170894 | 0.28440353 | 0.11149276 | 0.91122561 | 0.98289594 |
| Dnah8 | 16.6406881 | -3.6560434 | 1.30806248 | -2.7950067 | 0.00518986 | 0.0823445 |
| Mrpl15 | 4920.91975 | 0.08857055 | 0.3392365 | 0.26108791 | 0.79402471 | 0.95310033 |
| Pla2g4c | 28.8508983 | 0.34246417 | 0.93918836 | 0.36463843 | 0.71538133 | 0.93235217 |
| Gm28042 | 21.9248647 | 0.71250078 | 1.28628743 | 0.55392035 | 0.57963336 | 0.88286755 |
| Engase | 453.197023 | 0.52119131 | 0.39065599 | 1.33414389 | 0.18215671 | 0.61344341 |
| Cdk10 | 893.431804 | -0.1123505 | 0.2523089 | -0.4452896 | 0.65611046 | 0.91165371 |
| Klf9 | 19.5497203 | -0.8455765 | 1.51655106 | -0.5575655 | 0.57714113 | 0.88121027 |
| Ppargc1b | 63.9081793 | -0.4725715 | 1.0411163 | -0.4539085 | 0.64989469 | 0.90973357 |
| Lgals3bp | 543.771702 | 0.53795973 | 0.39837157 | 1.35039689 | 0.17688871 | 0.60784615 |
| D3Ertd254e | 169.969003 | -0.2822613 | 0.47468633 | -0.5946269 | 0.5520929 | 0.87093412 |
| Pxk | 1327.48184 | 0.03523071 | 0.18099989 | 0.19464494 | 0.84567092 | 0.96765585 |

| | | | | | | |
|---|---|---|---|---|---|---|
| Map9 | 14.0893372 | 0.3673983 | 1.63183116 | 0.2251448 | 0.82186663 | 0.96232782 |
| Mapkbp1 | 76.114827 | 0.43504679 | 0.68328583 | 0.63669809 | 0.52432152 | 0.85953586 |
| Ccp110 | 223.664303 | -0.4113397 | 0.35746207 | -1.1507225 | 0.24984641 | 0.6842534 |
| Zdhhc15 | 144.754069 | -0.1878288 | 0.57219699 | -0.3282589 | 0.74271588 | 0.93850397 |
| Usp36 | 796.219161 | -0.2589789 | 0.36352117 | -0.7124177 | 0.47620618 | 0.83644936 |
| Chmp2a | 3877.8633 | 0.31731015 | 0.32406973 | 0.9791416 | 0.32751002 | 0.74829972 |
| Gde1 | 1814.69266 | -0.3308285 | 0.20336487 | -1.6267731 | 0.10378529 | 0.49305066 |
| Parl | 4675.69518 | 0.02135752 | 0.28176762 | 0.07579834 | 0.93957953 | 0.98913801 |
| Rbm34 | 1238.43393 | -0.1952118 | 0.23794781 | -0.8203974 | 0.4119896 | 0.80245328 |
| Vhl | 1745.32154 | 0.37476043 | 0.35572325 | 1.05351683 | 0.29210418 | 0.72584701 |
| Ndufb7 | 3142.62022 | 0.32561737 | 0.39262986 | 0.82932401 | 0.40692109 | 0.79992362 |
| Brk1 | 7739.42331 | 0.27335266 | 0.31743179 | 0.86113827 | 0.38916189 | 0.78942657 |
| Mga | 482.91094 | 0.44209284 | 0.30468412 | 1.45098746 | 0.14678335 | 0.57365079 |
| Zswim5 | 11.1459078 | -1.7804143 | 2.10493358 | -0.8458292 | 0.39764803 | 0.79531492 |
| Trim36 | 531.811141 | 0.0294866 | 0.34997218 | 0.08425412 | 0.93285439 | 0.98748011 |
| Aspm | 648.495443 | 0.29406751 | 0.44340439 | 0.66320388 | 0.50719999 | 0.85169216 |
| Ppp3r1 | 6027.07004 | -0.2583204 | 0.17660145 | -1.4627308 | 0.14354104 | 0.56863263 |
| Jcad | 176.446282 | 0.612878 | 0.40173998 | 1.52555888 | 0.12711976 | 0.54111158 |
| Zfp446 | 264.244526 | 0.46933266 | 0.47210563 | 0.99412639 | 0.32016134 | 0.74413737 |
| Zbtb41 | 207.685734 | -0.080674 | 0.67485284 | -0.1195431 | 0.90484511 | 0.98137869 |
| Slc16a2 | 242.016025 | -1.036138 | 0.58906698 | -1.7589478 | 0.07858638 | 0.43353356 |
| Rnf225 | 9.25560048 | -0.4274768 | 1.60313479 | -0.2666505 | 0.78973824 | 0.95255415 |
| Rfc3 | 3747.30895 | 0.35259214 | 0.33147486 | 1.06370705 | 0.28746143 | 0.72296786 |
| Zfp944 | 360.044707 | 0.01123545 | 0.2528456 | 0.04443602 | 0.96455685 | 0.99406401 |
| Coil | 950.924438 | -0.0699696 | 0.28342209 | -0.2468742 | 0.80500561 | 0.95725854 |
| Tesk2 | 999.611639 | 0.88895097 | 0.29053895 | 3.05966195 | 0.00221587 | 0.04309496 |
| Ttc37 | 280.952272 | 0.26502848 | 0.7690056 | 0.3446379 | 0.73036661 | 0.93653187 |
| Pqlc1 | 712.249474 | 0.04886441 | 0.25666534 | 0.19038182 | 0.84900994 | 0.96765585 |
| Scaper | 163.895766 | -0.0792317 | 0.44188667 | -0.1793032 | 0.8576996 | 0.9694296 |
| Pds5b | 811.670063 | 0.03384658 | 0.28007191 | 0.1208496 | 0.90381016 | 0.98069095 |
| Cpsf1 | 3212.42471 | -0.4383503 | 0.32591485 | -1.3449841 | 0.17863037 | 0.60992118 |
| Fancd2 | 458.009537 | -0.2657667 | 0.48577418 | -0.5470994 | 0.58431046 | 0.88404713 |
| Cct2 | 12375.7865 | -0.4883079 | 0.15886364 | -3.0737548 | 0.00211383 | 0.04170725 |
| Rpap1 | 1382.38052 | -0.1841657 | 0.25246394 | -0.7294734 | 0.46571213 | 0.83339036 |
| Ccdc17 | 35.1052077 | 1.15441501 | 1.12423017 | 1.02684934 | 0.30449141 | 0.73467909 |
| Fgd5 | 62.0910297 | 1.40372156 | 1.54624752 | 0.90782461 | 0.36397089 | 0.77045439 |
| Lyl1 | 2154.57499 | -0.2905173 | 0.17982083 | -1.6155933 | 0.10618228 | 0.49777747 |
| Gpbp1l1 | 166.482274 | -0.2550032 | 0.33842858 | -0.753492 | 0.45115434 | 0.82550977 |
| Phka1 | 114.710999 | 0.86825276 | 0.59577067 | 1.45736069 | 0.14501684 | 0.57067981 |
| Poglut1 | 1805.53557 | -0.2435152 | 0.17086849 | -1.4251614 | 0.15411056 | 0.58303286 |
| Farp2 | 51.197241 | -0.0702931 | 0.95290348 | -0.0737673 | 0.94119556 | 0.98935054 |
| Zfp551 | 18.1495208 | -0.4446893 | 1.26879634 | -0.3504813 | 0.72597755 | 0.93558387 |
| Zdhhc5 | 261.042316 | -0.1122791 | 0.37197282 | -0.3018476 | 0.76276826 | 0.94298237 |
| Ccdc174 | 452.83194 | -0.132588 | 0.30386912 | -0.4363326 | 0.66259543 | 0.91395288 |
| Hdlbp | 6956.86899 | 0.06844454 | 0.23033231 | 0.29715561 | 0.76634771 | 0.94360659 |
| Ctnnd1 | 822.920355 | 0.46586962 | 0.38456059 | 1.21143359 | 0.22572927 | 0.65900564 |

| | | | | | | |
|---|---|---|---|---|---|---|
| Meak7 | 518.22531 | 0.57663923 | 0.43080111 | 1.33852773 | 0.18072447 | 0.61122544 |
| Ccs | 1189.40279 | 0.02665513 | 0.23656777 | 0.1126744 | 0.9102887 | 0.98275189 |
| Golim4 | 383.815158 | -0.1078307 | 0.35719845 | -0.3018789 | 0.76274442 | 0.94298237 |
| Kctd7 | 202.163466 | 0.60851087 | 0.35482262 | 1.71497201 | 0.08635036 | 0.44906118 |
| Tmed8 | 98.0358773 | -0.9236203 | 0.52816232 | -1.7487433 | 0.0803354 | 0.4379425 |
| Vav1 | 9542.46965 | -0.2770581 | 0.2147809 | -1.2899571 | 0.19706557 | 0.62932233 |
| Tpst1 | 393.397616 | 0.6253077 | 0.29821569 | 2.0968303 | 0.0360086 | 0.28477582 |
| Srsf2 | 8109.93776 | 0.21651536 | 0.28419755 | 0.7618481 | 0.44615067 | 0.82166369 |
| Mks1 | 326.288631 | 0.42433359 | 0.33749947 | 1.25728671 | 0.20864984 | 0.64195509 |
| Pomt2 | 446.887675 | 0.0124531 | 0.34301336 | 0.03630499 | 0.97103917 | 0.99443087 |
| Tspan8 | 72.8316305 | 0.66995354 | 0.63008624 | 1.06327276 | 0.28765827 | 0.72300867 |
| Sik3 | 142.006101 | -0.8799189 | 0.45962832 | -1.914414 | 0.05556728 | 0.3627292 |
| Exoc3 | 3514.06671 | 0.11011433 | 0.28416592 | 0.38750014 | 0.69838598 | 0.92669304 |
| Ino80 | 243.452921 | -0.3986031 | 0.45639571 | -0.8733717 | 0.3824605 | 0.78540194 |
| Tspoap1 | 93.4842968 | 0.83733266 | 0.63768175 | 1.31308864 | 0.18915308 | 0.61968656 |
| Cipc | 655.403727 | -0.2125154 | 0.33677954 | -0.6310222 | 0.52802603 | 0.86198732 |
| Lrrc58 | 3627.8729 | 0.04124855 | 0.33479859 | 0.12320407 | 0.9019455 | 0.98043702 |
| Ogt | 920.216484 | -0.2400348 | 0.23413571 | -1.0251952 | 0.30527109 | 0.73525914 |
| Zfc3h1 | 238.516832 | -0.1785431 | 0.46276294 | -0.3858199 | 0.69963006 | 0.92669304 |
| Emid1 | 241.189644 | -1.4256843 | 0.39575853 | -3.6024097 | 0.00031528 | 0.00922273 |
| Ccnd3 | 11085.5913 | -0.7201916 | 0.19900052 | -3.6190437 | 0.00029569 | 0.00880102 |
| Irf2bpl | 732.363485 | 0.2615339 | 0.36418788 | 0.71812906 | 0.47267771 | 0.83483892 |
| Zbed5 | 108.009286 | -0.2710562 | 0.67347415 | -0.4024745 | 0.68733482 | 0.92427909 |
| Rhbdd3 | 522.084354 | -0.0796236 | 0.24879352 | -0.320039 | 0.74893879 | 0.9399191 |
| Rnf43 | 31.4813767 | -4.9024346 | 1.38946677 | -3.5282848 | 0.00041826 | 0.01163526 |
| Nsf | 5756.18056 | -0.2724924 | 0.20061546 | -1.3582821 | 0.17437418 | 0.60656845 |
| Hsdl1 | 3500.42374 | 0.38044884 | 0.18636215 | 2.04144907 | 0.04120621 | 0.30914162 |
| Chmp7 | 1915.61943 | -0.0618131 | 0.18344615 | -0.3369548 | 0.73615101 | 0.9377627 |
| Lsm3 | 2725.80001 | 0.15854974 | 0.36303147 | 0.43673828 | 0.66230117 | 0.91395288 |
| R3hcc1 | 623.387703 | -0.0738381 | 0.26427031 | -0.2794036 | 0.77993514 | 0.94906462 |
| Gas2l1 | 251.0778 | 1.18837329 | 0.57021861 | 2.08406614 | 0.03715415 | 0.28956966 |
| Chchd4 | 4283.07232 | -0.0470569 | 0.31340204 | -0.1501486 | 0.88064734 | 0.97572311 |
| Loxl2 | 29.0033312 | 0.80181276 | 0.81651126 | 0.9819984 | 0.32610063 | 0.74759915 |
| Polq | 192.831762 | -0.6359301 | 0.40883347 | -1.5554746 | 0.11983308 | 0.52718991 |
| Efcab14 | 1033.89435 | 0.00592781 | 0.24268424 | 0.02442601 | 0.98051281 | 0.99515582 |
| Mrps17 | 3102.49633 | 0.16550803 | 0.34971012 | 0.47327207 | 0.63601908 | 0.90531997 |
| Vps18 | 2927.16182 | 0.4350264 | 0.34891777 | 1.24678774 | 0.21247534 | 0.64592714 |
| Atm | 228.018337 | 0.37647616 | 0.52660309 | 0.71491446 | 0.4746619 | 0.83578199 |
| Gpc1 | 20.0559873 | -1.3816743 | 1.61123472 | -0.8575252 | 0.39115472 | 0.79005237 |
| Rhov | 32.2567713 | 0.31793189 | 1.02582265 | 0.30992871 | 0.75661517 | 0.94102069 |
| Usp54 | 77.6475739 | -0.434036 | 0.75548343 | -0.5745143 | 0.56561985 | 0.87840899 |
| Golgb1 | 422.989262 | -0.0657476 | 0.3863631 | -0.1701704 | 0.86487615 | 0.9705375 |
| Hdac11 | 16.5876809 | 3.02987501 | 1.67515033 | 1.80871827 | 0.07049478 | 0.41006309 |
| Plekhm1 | 418.739997 | 0.31218709 | 0.41604729 | 0.75036444 | 0.45303524 | 0.82610332 |
| Slc25a37 | 368.347651 | -0.5120938 | 0.51498576 | -0.9943844 | 0.32003578 | 0.74413737 |
| Senp6 | 359.690978 | -0.7354814 | 0.46156095 | -1.5934655 | 0.11105581 | 0.50888686 |

| | | | | | | |
|---|---|---|---|---|---|---|
| Agpat1 | 576.940965 | -0.5931403 | 0.33825211 | -1.7535451 | 0.07950848 | 0.4364971 |
| Arhgap27 | 1020.73162 | -0.244747 | 0.24071079 | -1.0167679 | 0.30926387 | 0.73890515 |
| Exosc4 | 2498.64744 | -0.0861281 | 0.22514205 | -0.3825502 | 0.70205327 | 0.92762537 |
| Ints14 | 4021.74547 | -0.1179025 | 0.17353331 | -0.679423 | 0.49686986 | 0.84814011 |
| Zdhhc14 | 42.2060625 | 2.05199891 | 0.95614945 | 2.14610688 | 0.03186445 | 0.26571087 |
| Batf | 1548.01961 | 0.22107729 | 0.32554999 | 0.67908861 | 0.49708172 | 0.84814011 |
| Setd5 | 1053.26009 | -0.120023 | 0.21561233 | -0.5566611 | 0.57775899 | 0.88170231 |
| Thoc5 | 1797.14174 | -0.3133996 | 0.17624434 | -1.7782109 | 0.07536922 | 0.42273804 |
| Igsf9b | 19.1121711 | -0.1376948 | 1.95930882 | -0.0702772 | 0.943973 | 0.99002244 |
| Dnajc17 | 393.911693 | -0.3080798 | 0.30604729 | -1.0066412 | 0.31410725 | 0.74065408 |
| Nipsnap1 | 762.982855 | 0.06369345 | 0.36489843 | 0.17455117 | 0.86143232 | 0.97007287 |
| Nek9 | 739.744042 | -0.1164411 | 0.29369922 | -0.3964639 | 0.69176286 | 0.92504756 |
| Traf3ip1 | 60.0506504 | 1.73742734 | 0.8612244 | 2.01739215 | 0.04365461 | 0.32015995 |
| Med13 | 293.765691 | -0.9606029 | 0.46683895 | -2.0576751 | 0.03962134 | 0.30142277 |
| Fam53c | 637.799289 | 0.1994638 | 0.28456915 | 0.70093261 | 0.48334507 | 0.84107543 |
| Ccdc15 | 142.00574 | 0.05315051 | 0.50620783 | 0.1049974 | 0.91637787 | 0.98386323 |
| Sdr42e1 | 189.076083 | 1.07102861 | 0.51363338 | 2.08520053 | 0.0370511 | 0.28924449 |
| Kif4 | 1692.29439 | -0.1217392 | 0.33791531 | -0.3602655 | 0.71864858 | 0.93310883 |
| Iqsec1 | 124.95123 | -0.1445533 | 0.62622818 | -0.2308316 | 0.81744562 | 0.96079676 |
| Trim59 | 2203.89435 | -0.0964315 | 0.25484774 | -0.3783889 | 0.70514173 | 0.92879471 |
| Slc26a2 | 942.102904 | 0.0930353 | 0.44980783 | 0.20683343 | 0.83613994 | 0.96683245 |
| Exosc1 | 1792.57114 | -0.4276866 | 0.27873175 | -1.534402 | 0.12493078 | 0.53626132 |
| Kctd9 | 321.770622 | -0.3967723 | 0.45671832 | -0.8687461 | 0.38498603 | 0.78754751 |
| Brip1 | 1227.11709 | -0.0694876 | 0.30003673 | -0.2315971 | 0.81685099 | 0.96079676 |
| Plcg2 | 8764.0257 | -0.8108139 | 0.2938026 | -2.7597236 | 0.00578503 | 0.08924119 |
| Zbed4 | 491.627546 | -0.119357 | 0.36026764 | -0.3313009 | 0.7404172 | 0.93800224 |
| Wbp2 | 3328.75364 | 0.15039633 | 0.2681344 | 0.56089904 | 0.57486637 | 0.88059206 |
| Cbl | 1023.41827 | -0.2032422 | 0.44026064 | -0.4616407 | 0.64433898 | 0.90829026 |
| Ube2f | 3444.67388 | -0.1561133 | 0.17610295 | -0.8864891 | 0.37535404 | 0.77815049 |
| Gtf2h5 | 2399.29541 | 0.31827688 | 0.32347838 | 0.9839201 | 0.3251548 | 0.7466902 |
| Smc4 | 6170.59369 | -0.2927912 | 0.14558628 | -2.011118 | 0.04431299 | 0.32302252 |
| Ramp1 | 488.392672 | -2.6373654 | 0.38753614 | -6.8054695 | 1.007202185 | 2.415902810 |
| Mtmr3 | 522.091954 | -0.2427559 | 0.40314319 | -0.602158 | 0.54706899 | 0.87003314 |
| Cpne2 | 8.36849395 | -0.1507517 | 1.77570087 | -0.084897 | 0.93234328 | 0.98730569 |
| Tkfc | 100.021233 | -0.5767922 | 0.50161271 | -1.1498756 | 0.25019512 | 0.6842534 |
| Tulp4 | 864.428265 | -0.1563549 | 0.22727053 | -0.6879684 | 0.49147268 | 0.84497726 |
| Wdr5b | 135.181036 | 0.20731215 | 0.50396359 | 0.41136335 | 0.68080613 | 0.92061965 |
| Cmip | 396.650218 | -0.1929054 | 0.25798782 | -0.7477307 | 0.4546226 | 0.82758903 |
| Lif | 15.6611763 | 0.44010571 | 2.03440898 | 0.21633099 | 0.82872975 | 0.96462215 |
| Spata6 | 1098.8493 | 0.74311438 | 0.24150468 | 3.07701854 | 0.00209082 | 0.04131993 |
| Pja1 | 2732.40325 | -0.1872471 | 0.15864274 | -1.180307 | 0.23787814 | 0.67403588 |
| Tbc1d10a | 700.183294 | -0.1965206 | 0.26592265 | -0.7390142 | 0.45989835 | 0.82987212 |
| Pkd1l2 | 50.342958 | -1.3779337 | 1.00897231 | -1.3656804 | 0.17203931 | 0.60451272 |
| Parp14 | 288.877973 | 0.04529598 | 0.416852 | 0.10866202 | 0.91347057 | 0.98309098 |
| Gcsh | 4747.23509 | 0.22153887 | 0.3023129 | 0.73281317 | 0.46367239 | 0.83202414 |
| Myo15b | 22.4067702 | -0.6975817 | 1.16771148 | -0.5973921 | 0.5502456 | 0.87093412 |

| | | | | | | |
|---|---|---|---|---|---|---|
| Zfp707 | 96.6197736 | 0.14071446 | 0.76081036 | 0.1849534 | 0.85326559 | 0.96811477 |
| Zxdc | 142.332119 | -0.1445676 | 0.53122709 | -0.2721391 | 0.78551509 | 0.95168899 |
| Cops8 | 4278.58515 | -0.2242033 | 0.19812482 | -1.1316266 | 0.25779144 | 0.69309071 |
| Gbp8 | 56.3302784 | 2.54663418 | 1.12494991 | 2.26377562 | 0.02358792 | 0.22145127 |
| Trmt5 | 725.923947 | 0.13420444 | 0.2752249 | 0.48761738 | 0.62582089 | 0.90151518 |
| Cyb561a3 | 1767.10641 | -0.6188101 | 0.25063748 | -2.4689449 | 0.01355121 | 0.1574243 |
| Dhrs11 | 549.384283 | -0.6520487 | 0.26739442 | -2.4385275 | 0.01474724 | 0.16703976 |
| Polr3b | 1055.57858 | 0.12541057 | 0.23095672 | 0.54300463 | 0.58712661 | 0.88626849 |
| Eda2r | 11.8296353 | 0.4342079 | 1.75847218 | 0.24692338 | 0.80496752 | 0.95725854 |
| Ifit1 | 57.4970708 | 1.70077258 | 0.99526438 | 1.70886512 | 0.08747594 | 0.45207544 |
| Pkd2 | 43.7432352 | 0.26797558 | 0.8392544 | 0.31930197 | 0.74949754 | 0.9399191 |
| Caskin2 | 26.4561775 | -0.4727699 | 1.1679568 | -0.4047838 | 0.68563647 | 0.92314423 |
| Sec22a | 1139.66929 | 0.94020471 | 0.4102729 | 2.29165685 | 0.02192545 | 0.21320672 |
| Diaph2 | 159.604469 | -0.0084728 | 0.53804972 | -0.0157472 | 0.98743608 | 0.9962376 |
| Snx2 | 50183.7671 | 0.71656212 | 0.20422296 | 3.50872463 | 0.00045026 | 0.0124335 |
| Uaca | 519.520204 | 0.56736474 | 0.40501867 | 1.40083601 | 0.16126312 | 0.59141542 |
| Kdelc2 | 1534.59982 | 0.9530625 | 0.22774942 | 4.18469778 | 2.855457360 | 0.00122996 |
| Pcnx4 | 477.262506 | 0.77365771 | 0.41824511 | 1.84977109 | 0.06434655 | 0.39202997 |
| Mad2l1bp | 1295.81797 | 0.3067464 | 0.2901974 | 1.05702671 | 0.2904994 | 0.72461526 |
| Hmgxb4 | 617.7302 | 0.06268953 | 0.31792348 | 0.19718435 | 0.84368328 | 0.96765585 |
| Zfp395 | 101.626342 | -1.0951075 | 0.66295446 | -1.6518592 | 0.09856325 | 0.48025164 |
| Ice1 | 236.985926 | -0.671514 | 0.51805421 | -1.2962234 | 0.19489853 | 0.62699967 |
| Zfp418 | 60.4349205 | -0.7422685 | 1.35163617 | -0.5491629 | 0.58289364 | 0.88343755 |
| Morc2a | 478.008929 | -0.3429233 | 0.27942615 | -1.2272413 | 0.21973192 | 0.65289787 |
| Rsrc1 | 1541.6184 | -0.3188346 | 0.16709015 | -1.9081594 | 0.05637062 | 0.36517938 |
| Hdx | 16.3324091 | -0.9569894 | 1.7736152 | -0.5395699 | 0.58949366 | 0.88722166 |
| Zfyve9 | 114.019131 | -0.277378 | 0.51537214 | -0.5382091 | 0.59043267 | 0.88748446 |
| Washc4 | 3128.48443 | 0.46920657 | 0.21950314 | 2.13758479 | 0.03255046 | 0.2679608 |
| Ccpg1 | 982.224486 | 0.11061332 | 0.25510409 | 0.43360074 | 0.6645784 | 0.91431173 |
| Atp5h | 7527.54534 | 0.41920946 | 0.41641289 | 1.00671585 | 0.31407134 | 0.74065408 |
| Ptpn13 | 20.8734327 | -1.6615327 | 1.35337417 | -1.2276964 | 0.21956094 | 0.65289787 |
| Daam1 | 86.6513752 | -1.1993102 | 0.83085044 | -1.443473 | 0.14888727 | 0.57600823 |
| Tent4a | 134.825084 | -0.488087 | 0.46062849 | -1.0596111 | 0.28932158 | 0.72395811 |
| Hid1 | 29.6672141 | 0.74161919 | 1.8977254 | 0.39079373 | 0.69594971 | 0.92623267 |
| Myo5a | 2604.23357 | 0.46038768 | 0.33948712 | 1.3561271 | 0.17505873 | 0.60743838 |
| Ppp1r18 | 1899.17906 | 0.23218605 | 0.17699499 | 1.31182276 | 0.18957994 | 0.62019167 |
| 2700049A03I | 429.778743 | -0.2201443 | 0.32312049 | -0.6813071 | 0.49567717 | 0.84734751 |
| Mon2 | 380.933901 | 0.19334821 | 0.55772675 | 0.34667193 | 0.7288378 | 0.93617556 |
| Tut4 | 242.892487 | 0.06379546 | 0.41231834 | 0.15472381 | 0.87703906 | 0.97458574 |
| Chst11 | 30.6251946 | -3.5751034 | 1.38282373 | -2.5853645 | 0.00972761 | 0.12699877 |
| Ppm1h | 1745.53588 | 0.24144754 | 0.30561052 | 0.79004983 | 0.42949867 | 0.81155572 |
| Pik3ip1 | 34.8280375 | -0.0262142 | 1.06483245 | -0.0246181 | 0.98035956 | 0.99508285 |
| Ssh3 | 65.3872466 | -0.0537504 | 0.76248764 | -0.0704935 | 0.94380085 | 0.98992676 |
| Mtrr | 938.786715 | 0.06819723 | 0.21491799 | 0.31731745 | 0.75100274 | 0.94003547 |
| Rxylt1 | 1458.19359 | 0.45061511 | 0.18315503 | 2.46029339 | 0.01388235 | 0.15990844 |
| Gpatch8 | 738.287935 | -0.0591615 | 0.33799225 | -0.175038 | 0.86104979 | 0.96996131 |

| | | | | | | |
|---|---|---|---|---|---|---|
| Zyg11b | 389.249916 | 0.100236 | 0.31416954 | 0.31905066 | 0.7496881 | 0.9399191 |
| Setmar | 134.921947 | 0.88334 | 0.53815706 | 1.6414167 | 0.10071094 | 0.48503818 |
| Tiparp | 1872.96099 | 0.27096546 | 0.19381556 | 1.39805836 | 0.16209554 | 0.59315429 |
| Cd300ld | 1015.95362 | 1.21422929 | 0.22816715 | 5.32166564 | 1.028214292 | 9.826676122 |
| Ankrd12 | 290.83551 | 1.08803702 | 0.42346895 | 2.56934306 | 0.01018915 | 0.13024442 |
| Cd300a | 6127.80945 | 1.20263852 | 0.33373031 | 3.60362392 | 0.00031381 | 0.00922273 |
| Ythdc2 | 117.659024 | -0.4432981 | 0.54494158 | -0.8134783 | 0.41594389 | 0.80440685 |
| Cacna1a | 20.9086708 | -0.1248476 | 0.91833581 | -0.1359499 | 0.89186092 | 0.97927786 |
| Tmem109 | 4019.90394 | 0.01310548 | 0.20207584 | 0.06485428 | 0.94829002 | 0.99019648 |
| Bmp2k | 3141.0328 | 0.17205339 | 0.33446655 | 0.51441136 | 0.60696442 | 0.89307142 |
| Itga2b | 167.996835 | -1.570768 | 0.51986136 | -3.0215133 | 0.00251515 | 0.04719108 |
| Xpot | 2965.50363 | -0.1997501 | 0.21893223 | -0.9123832 | 0.36156705 | 0.76887042 |
| Pbx2 | 132.169413 | -0.4286844 | 0.39925396 | -1.0737137 | 0.28295102 | 0.71857502 |
| Tdg | 6834.12096 | -0.0209962 | 0.20506573 | -0.1023875 | 0.91844913 | 0.98390186 |
| Dbn1 | 106.960088 | 1.05155445 | 0.60732088 | 1.73146434 | 0.08336898 | 0.44341423 |
| Rnps1 | 8182.60161 | -0.5248922 | 0.20501377 | -2.560278 | 0.01045885 | 0.1321726 |
| Gns | 4434.23058 | 0.69760827 | 0.33051996 | 2.11063884 | 0.03480337 | 0.27954668 |
| Grn | 22030.539 | 0.85754938 | 0.27834583 | 3.08087746 | 0.00206392 | 0.04112033 |
| Ppp1r21 | 1733.77258 | 0.0837965 | 0.2205399 | 0.37996071 | 0.70397458 | 0.92850227 |
| Ttyh2 | 1404.96787 | 0.36604708 | 0.32916448 | 1.11204916 | 0.26611701 | 0.7023537 |
| Tmx4 | 162.136388 | 0.34685599 | 0.5287384 | 0.6560068 | 0.51181975 | 0.85393154 |
| Cnot6l | 733.344452 | -0.1547461 | 0.32713527 | -0.4730341 | 0.63618885 | 0.90536276 |
| Mrps10 | 2715.83071 | 0.03199749 | 0.31599743 | 0.10125869 | 0.91934511 | 0.9839023 |
| Dgkh | 132.966152 | 0.80974021 | 0.93009933 | 0.87059541 | 0.3839751 | 0.78704643 |
| Nostrin | 247.228338 | -0.1226364 | 0.31703459 | -0.3868235 | 0.69888686 | 0.92669304 |
| Nagk | 1407.83037 | 0.50870628 | 0.24662663 | 2.06265754 | 0.03914518 | 0.29945131 |
| Sirt6 | 499.846703 | 0.32514288 | 0.22038726 | 1.47532524 | 0.14012512 | 0.56396008 |
| Mast4 | 95.7011715 | 1.34830226 | 0.84388639 | 1.59772959 | 0.11010317 | 0.50692212 |
| Tmub2 | 1310.09472 | 0.71647803 | 0.30109281 | 2.37959196 | 0.01733182 | 0.18521003 |
| Tle6 | 67.8156951 | 1.82147174 | 1.1771779 | 1.54732071 | 0.12178593 | 0.53018053 |
| Map4k5 | 350.701386 | 0.40226536 | 0.54615145 | 0.73654544 | 0.46139882 | 0.83025001 |
| Dusp5 | 784.588252 | 0.5687116 | 1.21178982 | 0.46931538 | 0.63884422 | 0.90615794 |
| BC030867 | 856.763037 | -0.0575374 | 0.21714671 | -0.2649702 | 0.79103242 | 0.95268772 |
| Gna11 | 205.525377 | -0.8077722 | 0.35386442 | -2.2827168 | 0.02244706 | 0.21507833 |
| Gpsm3 | 354.805476 | -0.5350111 | 0.28486104 | -1.8781477 | 0.06036097 | 0.37881203 |
| Rab24 | 1059.94275 | 0.54715291 | 0.54926193 | 0.99616026 | 0.31917228 | 0.74398524 |
| Gna15 | 2624.78751 | 0.06500194 | 0.17636225 | 0.36857061 | 0.71244781 | 0.93235217 |
| G6pc3 | 1075.00968 | -0.4270159 | 0.2653575 | -1.6092098 | 0.10757048 | 0.50187249 |
| Ccdc122 | 44.2997304 | 1.01602911 | 0.85736947 | 1.18505399 | 0.2359961 | 0.67182855 |
| Unc13a | 59.6517975 | 0.07865424 | 0.95467284 | 0.08238868 | 0.93433764 | 0.98756396 |
| Zfp661 | 123.361757 | 0.40289827 | 0.68864965 | 0.58505551 | 0.5585104 | 0.87327051 |
| Sos2 | 146.896852 | 0.17711442 | 0.38535283 | 0.45961626 | 0.64579169 | 0.90847224 |
| Colgalt1 | 4863.82603 | -0.0648396 | 0.18730578 | -0.3461696 | 0.72921525 | 0.93623951 |
| Cpsf7 | 469.559826 | -0.1298219 | 0.32062816 | -0.4048987 | 0.68555197 | 0.92314423 |
| Nup54 | 3416.01192 | -0.2877203 | 0.21895534 | -1.3140592 | 0.18882629 | 0.61935434 |
| Tet3 | 789.43649 | -0.5681689 | 0.47026615 | -1.2081858 | 0.22697579 | 0.66015092 |

| | | | | | | |
|---|---|---|---|---|---|---|
| Tespa1 | 37.3366421 | -8.6262299 | 1.62855733 | -5.2968537 | 1.178150061 | 1.117233309 |
| Gnat1 | 12.5203458 | 0.41492247 | 1.09200364 | 0.37996437 | 0.70397186 | 0.92850227 |
| Plvap | 236.257263 | 2.33981569 | 1.56318476 | 1.49682606 | 0.13443852 | 0.55541589 |
| Ttc21b | 342.989397 | 0.29639543 | 0.40190032 | 0.73748493 | 0.46082751 | 0.82987212 |
| Tmem127 | 1621.75322 | 0.15405699 | 0.22515266 | 0.6842335 | 0.49382773 | 0.84678075 |
| Acot11 | 107.012287 | -0.762715 | 0.71066732 | -1.0732377 | 0.28316447 | 0.71881115 |
| Mfsd12 | 1144.67465 | 0.39326216 | 0.27177597 | 1.44700855 | 0.14789453 | 0.57472133 |
| Cxcl10 | 1418.10521 | 0.94975799 | 0.65067202 | 1.45965702 | 0.14438436 | 0.57027792 |
| Fam214a | 101.19051 | 0.08596979 | 0.61145253 | 0.14059929 | 0.88818651 | 0.97832097 |
| Ano8 | 8.63562063 | -1.0277376 | 1.46814195 | -0.700026 | 0.48391105 | 0.84128398 |
| Ankrd27 | 1617.18189 | -0.5393578 | 0.19330262 | -2.790225 | 0.00526714 | 0.08313929 |
| Myl12b | 6071.65932 | -0.5197529 | 0.16013585 | -3.2456999 | 0.00117162 | 0.02644365 |
| Nudt19 | 2420.66177 | 0.26292926 | 0.33218151 | 0.79152287 | 0.42863893 | 0.81136783 |
| Mrpl34 | 1515.50281 | 0.16588162 | 0.28952811 | 0.57293788 | 0.56668676 | 0.87883863 |
| Lrr1 | 729.032172 | 0.3047577 | 0.36082962 | 0.84460277 | 0.39833266 | 0.79555974 |
| Cactin | 1770.39285 | -0.2083402 | 0.21293328 | -0.9784293 | 0.32786203 | 0.74843736 |
| Rps29 | 8879.67128 | -0.0443796 | 0.2697408 | -0.1645269 | 0.86931637 | 0.97223872 |
| Cog3 | 722.511257 | 0.09295058 | 0.27359883 | 0.33973309 | 0.73405754 | 0.93750666 |
| Pip5k1c | 1131.73746 | -0.2913969 | 0.24068071 | -1.21072 | 0.22600274 | 0.65937941 |
| Cobll1 | 32.6873824 | 0.3056122 | 0.86444931 | 0.35353398 | 0.72368816 | 0.93453787 |
| Ncaph | 3523.36665 | 0.04288053 | 0.27964941 | 0.15333676 | 0.87813271 | 0.97503155 |
| Sidt2 | 352.029441 | -0.3474484 | 0.69751393 | -0.498124 | 0.61839662 | 0.89878056 |
| Ushbp1 | 9.20531343 | 1.98134146 | 2.24603164 | 0.88215207 | 0.37769458 | 0.78033065 |
| Tjp3 | 65.4827039 | -0.5568527 | 0.68934121 | -0.8078042 | 0.41920333 | 0.80529433 |
| Dhcr24 | 227.997688 | -0.6099573 | 0.38845226 | -1.5702247 | 0.11636286 | 0.521508 |
| Rnf44 | 506.222984 | -0.0050102 | 0.42807556 | -0.0117041 | 0.99066172 | 0.9966603 |
| Rtkn | 140.591987 | 0.23237277 | 0.49353744 | 0.47083109 | 0.63776136 | 0.90599527 |
| Dhx8 | 2423.70858 | -0.3050743 | 0.25019434 | -1.2193493 | 0.22271166 | 0.65613364 |
| Mrpl54 | 3307.22534 | 0.48917793 | 0.3970657 | 1.23198233 | 0.21795569 | 0.65150232 |
| Arl4d | 125.919465 | 1.10163961 | 0.8909496 | 1.23647803 | 0.21628093 | 0.64982959 |
| Synrg | 280.618966 | 0.04525738 | 0.44052203 | 0.10273579 | 0.91817267 | 0.98390186 |
| Tmem106a | 1244.21718 | 1.36026931 | 0.26362704 | 5.1598247 | 2.471811795 | 2.175372208 |
| Cog7 | 1366.54367 | -0.1783455 | 0.39565128 | -0.4507643 | 0.65215945 | 0.91037383 |
| Cebpa | 83.5650233 | -0.1320661 | 0.67458056 | -0.1957752 | 0.84478613 | 0.96765585 |
| Rubcnl | 516.481638 | 0.00358718 | 0.25670398 | 0.01397401 | 0.98885071 | 0.99650865 |
| Lbx2 | 19.8106352 | 1.10435396 | 0.94263357 | 1.1715623 | 0.24137283 | 0.67708722 |
| Dop1a | 93.7389175 | 0.64663063 | 0.5047039 | 1.28120792 | 0.20012065 | 0.63253708 |
| Dapk3 | 66.3280982 | 0.40120625 | 1.19143842 | 0.33674107 | 0.7363121 | 0.93787026 |
| Hrh2 | 29.6270048 | -2.0888094 | 1.2079719 | -1.7291871 | 0.08377562 | 0.44417933 |
| Vat1 | 83.269104 | 0.84906117 | 0.69025009 | 1.2300776 | 0.21866805 | 0.65211268 |
| Eef2 | 33777.8459 | -0.4897076 | 0.18726328 | -2.6150756 | 0.00892077 | 0.1197646 |
| Foxn2 | 860.743599 | -0.6797864 | 0.35341394 | -1.923485 | 0.05441917 | 0.36033644 |
| Igsf6 | 7084.64681 | -0.1161575 | 0.25357384 | -0.4580814 | 0.64689396 | 0.90927884 |
| Rundc1 | 726.317855 | -0.2640631 | 0.27960211 | -0.9444245 | 0.34495275 | 0.75993243 |
| Zbtb7a | 460.123367 | -0.1855188 | 0.38022934 | -0.4879129 | 0.62561156 | 0.90151518 |
| Baz1a | 862.844206 | -0.3253781 | 0.35587402 | -0.9143069 | 0.36055561 | 0.76881241 |

| | | | | | | |
|---|---|---|---|---|---|---|
| Ncapd3 | 864.785314 | 0.11146467 | 0.3341925 | 0.33353432 | 0.73873099 | 0.93800224 |
| Map2k2 | 4214.77205 | -0.3151096 | 0.23606984 | -1.3348152 | 0.18193682 | 0.61344341 |
| Creb3l3 | 21.8604905 | 1.61965925 | 1.00828447 | 1.60635148 | 0.10819671 | 0.50301538 |
| Zc3h12b | 12.1608111 | 3.47492661 | 2.40174791 | 1.44683237 | 0.14794388 | 0.57472133 |
| Kri1 | 265.630207 | -0.6701952 | 0.41538228 | -1.6134421 | 0.1066485 | 0.49890292 |
| Anapc13 | 666.471453 | 0.24417701 | 0.36732776 | 0.6647388 | 0.50621757 | 0.85086007 |
| Rrp12 | 2035.69509 | -0.7405245 | 0.32387303 | -2.2864655 | 0.02222704 | 0.21429774 |
| Dhx57 | 310.979179 | 0.36356441 | 0.28251971 | 1.28686389 | 0.19814173 | 0.630887 |
| Zc4h2 | 442.478742 | -0.0551041 | 0.3566334 | -0.154512 | 0.87720603 | 0.97458574 |
| Eef2k | 970.078949 | 0.77231015 | 0.30152402 | 2.56135533 | 0.01042647 | 0.13203618 |
| Oma1 | 1668.31224 | -0.0898185 | 0.23943068 | -0.3751337 | 0.70756101 | 0.92950127 |
| Mtmr9 | 809.129223 | 0.40539741 | 0.29139729 | 1.39121888 | 0.16415906 | 0.59585062 |
| Becn1 | 4766.48478 | -0.1079071 | 0.17868632 | -0.603891 | 0.54591613 | 0.86990153 |
| Secisbp2l | 398.314582 | 0.56609955 | 0.277484 | 2.04011604 | 0.04133877 | 0.30948419 |
| Eva1a | 24.5201173 | 2.08760827 | 1.29870791 | 1.60745018 | 0.10795565 | 0.50301538 |
| Egln3 | 146.350668 | -1.8601531 | 0.53183625 | -3.497605 | 0.00046946 | 0.01281887 |
| Dcbld2 | 74.9690679 | 0.72219935 | 0.68268812 | 1.05787596 | 0.290112 | 0.72418639 |
| Shc4 | 62.4644487 | -1.4699693 | 1.20275479 | -1.2221687 | 0.22164382 | 0.65514465 |
| Gcfc2 | 48.5457982 | -0.0172579 | 0.96050844 | -0.0179674 | 0.98566483 | 0.99564261 |
| Wdr78 | 58.9694384 | 0.97159382 | 0.74791753 | 1.29906544 | 0.19392147 | 0.62559108 |
| Arhgap5 | 606.053791 | 0.04568868 | 0.3169878 | 0.14413387 | 0.88539476 | 0.97753016 |
| Secisbp2 | 753.490096 | -0.3855427 | 0.41722901 | -0.9240553 | 0.35545754 | 0.7647925 |
| Nubpl | 694.440004 | -0.3160788 | 0.2810702 | -1.1245547 | 0.26077779 | 0.69691361 |
| Eif2s3x | 11095.7345 | -0.1640233 | 0.14986648 | -1.094463 | 0.27375197 | 0.71078816 |
| Elmod2 | 350.992732 | 0.13655986 | 0.43447291 | 0.31431156 | 0.75328442 | 0.94018119 |
| Ap2b1 | 1735.86324 | 0.01065189 | 0.21887486 | 0.04866659 | 0.961185 | 0.99338983 |
| Mitf | 113.655531 | 0.59756384 | 1.2501318 | 0.47800067 | 0.63264973 | 0.90430301 |
| Ints6 | 326.653235 | -0.6929876 | 0.54630717 | -1.2684944 | 0.20462144 | 0.63901002 |
| Zc3h12c | 74.4674842 | 0.34941838 | 0.8510838 | 0.41055696 | 0.68139743 | 0.92061965 |
| Kcne3 | 18.1499169 | 0.22588137 | 1.49276742 | 0.15131719 | 0.87972551 | 0.97541902 |
| Tanc1 | 15.5527549 | 3.68538611 | 1.71841508 | 2.14464255 | 0.03198144 | 0.26606568 |
| 1110059E24I | 3515.8259 | 0.34663318 | 0.35539499 | 0.97534626 | 0.32938853 | 0.74874249 |
| Ccdc186 | 335.539839 | 0.03593117 | 0.32799572 | 0.10954767 | 0.91276812 | 0.98292561 |
| Heatr5a | 679.141229 | -0.1492214 | 0.40042728 | -0.3726554 | 0.70940491 | 0.93033343 |
| Slc24a5 | 408.790484 | 0.20929164 | 0.58935493 | 0.35511986 | 0.7224998 | 0.93423486 |
| Tubg1 | 3478.52444 | -0.397109 | 0.17621943 | -2.2534915 | 0.02422817 | 0.22470299 |
| Arl6ip5 | 2514.56849 | 0.18097685 | 0.26991395 | 0.6704983 | 0.50254019 | 0.85012672 |
| Chrnb4 | 19.1480027 | 0.51514387 | 1.45507622 | 0.35403222 | 0.72331474 | 0.93435155 |
| Lars2 | 11100.8554 | -0.0726316 | 0.24365968 | -0.2980863 | 0.76563728 | 0.94347227 |
| Epn1 | 206.247523 | -0.439048 | 0.57606794 | -0.7621462 | 0.44597275 | 0.82166369 |
| Sppl2b | 907.398064 | -0.48237 | 0.31100916 | -1.5509833 | 0.1209057 | 0.52910763 |
| Slfn8 | 222.95984 | -0.1133236 | 0.5907277 | -0.1918372 | 0.84786971 | 0.96765585 |
| Leprot | 4732.65588 | 0.31364115 | 0.14699245 | 2.13372293 | 0.03286547 | 0.26964673 |
| Lsm7 | 2322.78828 | 0.28105362 | 0.43680424 | 0.64343153 | 0.51994413 | 0.8576517 |
| Spcs2 | 15413.9351 | 0.3996396 | 0.26023138 | 1.53570872 | 0.12460983 | 0.53617731 |
| Ccdc106 | 66.9756728 | 1.88852589 | 0.68561534 | 2.75449774 | 0.00587823 | 0.09033717 |

| | | | | | | |
|---|---|---|---|---|---|---|
| Pdk3 | 3826.12649 | -0.3909558 | 0.18410757 | -2.1235183 | 0.03371044 | 0.27382458 |
| Abraxas1 | 964.539511 | 0.52127691 | 0.39017195 | 1.33601842 | 0.18154325 | 0.61280867 |
| Trim13 | 337.550938 | 0.27446259 | 0.27640714 | 0.99296491 | 0.32072706 | 0.74413737 |
| Scai | 86.1433276 | -2.3623122 | 1.10777569 | -2.1324825 | 0.03296721 | 0.26976292 |
| Neu3 | 49.6790377 | 0.9820049 | 1.60454515 | 0.6120145 | 0.54052817 | 0.86741192 |
| Oaz1 | 27168.0508 | 0.42540024 | 0.32032528 | 1.32802582 | 0.18416958 | 0.61623261 |
| Eogt | 595.743672 | 0.03526217 | 0.32991261 | 0.10688336 | 0.91488151 | 0.98347559 |
| Pcyt1b | 9.5994769 | 0.63101499 | 2.10486173 | 0.29978928 | 0.76433789 | 0.94307414 |
| Hectd1 | 1663.49905 | 0.00904784 | 0.26159153 | 0.03458765 | 0.97240855 | 0.99443087 |
| Tut7 | 1222.50138 | 0.48330732 | 0.37764687 | 1.27978636 | 0.20062028 | 0.63366587 |
| Helq | 362.735048 | -0.0563367 | 0.42452214 | -0.1327061 | 0.89442583 | 0.97941957 |
| Pkig | 3117.2031 | -0.1869647 | 0.18693747 | -1.0001459 | 0.31723992 | 0.74230987 |
| Hpse | 472.420483 | -1.3647873 | 0.25479015 | -5.3565153 | 8.484232508 | 8.507181661 |
| Plekhj1 | 3401.49551 | 0.07892942 | 0.29699211 | 0.26576267 | 0.79042199 | 0.95261385 |
| Adrb1 | 10.305598 | 0.27543725 | 1.84340487 | 0.14941766 | 0.88122407 | 0.97582756 |
| Vps13c | 349.032214 | -0.1187354 | 0.55616117 | -0.213491 | 0.83094401 | 0.96466657 |
| G2e3 | 745.257352 | -0.246106 | 0.21080604 | -1.1674523 | 0.24302775 | 0.67814218 |
| Wdr38 | 10.0697012 | -0.9984711 | 1.6655925 | -0.599469 | 0.54886017 | 0.87082325 |
| Cops4 | 12506.1059 | 0.24861789 | 0.33580761 | 0.74035811 | 0.45908273 | 0.82987212 |
| Mid1 | 86.2682329 | 0.05540124 | 0.8352443 | 0.06632938 | 0.94711559 | 0.99019648 |
| Lin54 | 608.522986 | 0.42839912 | 0.45535332 | 0.94080597 | 0.3468043 | 0.76070594 |
| Gnptab | 340.377343 | -0.9222571 | 0.52945477 | -1.7418997 | 0.08152599 | 0.44011802 |
| Gdpd5 | 169.130232 | 1.21717764 | 0.5686746 | 2.1403763 | 0.03232437 | 0.26700278 |
| Sec31a | 2463.61497 | -0.4645051 | 0.32750429 | -1.4183175 | 0.15609808 | 0.58521232 |
| Fbxo33 | 239.390234 | -0.0248239 | 0.484336 | -0.0512534 | 0.95912359 | 0.99271349 |
| Lzts2 | 138.656903 | 0.61016099 | 0.55913767 | 1.0912536 | 0.27516131 | 0.71163812 |
| Nup37 | 2883.19737 | 0.06322238 | 0.3685571 | 0.17154024 | 0.86379899 | 0.9705375 |
| Ccl12 | 914.749096 | 2.87954225 | 0.45722909 | 6.29781067 | 3.018789052 | 5.956265561 |
| Uvrag | 2205.7194 | 0.19937165 | 0.23168992 | 0.86051069 | 0.38950759 | 0.78942657 |
| Nfkbiz | 6378.06203 | 0.99316613 | 0.94553284 | 1.05037718 | 0.29354473 | 0.72646828 |
| Parpbp | 988.528458 | -0.0472671 | 0.29105518 | -0.1623991 | 0.87099156 | 0.97268995 |
| Rmi1 | 1508.5182 | 0.31177907 | 0.17243978 | 1.80804607 | 0.07059933 | 0.41021047 |
| Gm49322 | 54.1464065 | -0.1583283 | 1.191138 | -0.1329219 | 0.89425517 | 0.97941957 |
| 1810055G02 | 610.121549 | -0.0970853 | 0.27832691 | -0.3488175 | 0.72722629 | 0.93576253 |
| Ccl7 | 580.923573 | 3.21222948 | 0.42802137 | 7.5048344 | 6.150630715 | 2.149733301 |
| Hacd2 | 3627.11933 | 0.2148457 | 0.32749856 | 0.65602029 | 0.51181107 | 0.85393154 |
| Shq1 | 1247.78704 | 0.04638946 | 0.22931197 | 0.20229845 | 0.83968341 | 0.96758168 |
| Pcsk7 | 1272.98649 | 0.33714762 | 0.1868106 | 1.80475632 | 0.07111282 | 0.41122734 |
| Ccl2 | 5852.50326 | 1.93982181 | 0.34610673 | 5.60469252 | 2.086249907 | 2.477776225 |
| Dennd1a | 307.91775 | 0.53686607 | 0.34254542 | 1.56728434 | 0.11704824 | 0.52289076 |
| Klf16 | 246.641527 | -0.3056842 | 0.38598306 | -0.7919627 | 0.42838245 | 0.81133341 |
| Oser1 | 9059.01455 | 0.30603662 | 0.25283497 | 1.21042044 | 0.2261176 | 0.65937941 |
| Emsy | 768.109119 | -0.0668943 | 0.26367382 | -0.2537009 | 0.79972666 | 0.95518266 |
| Tmem98 | 98.3821941 | 1.82215039 | 0.64133526 | 2.84118232 | 0.00449466 | 0.07360533 |
| Rabgap1 | 455.418288 | 0.28806687 | 0.33639417 | 0.85633728 | 0.39181125 | 0.7902765 |
| Haus8 | 2799.58734 | -0.2874958 | 0.26040447 | -1.1040356 | 0.26957769 | 0.70660893 |

| | | | | | | |
|---|---|---|---|---|---|---|
| Myo1d | 27.456902 | 1.09993037 | 1.02712929 | 1.07087821 | 0.2842242 | 0.71974495 |
| Thyn1 | 2879.22851 | 0.19472502 | 0.36223896 | 0.53755956 | 0.59088116 | 0.88755516 |
| Ccr3 | 38.6593438 | 2.4327722 | 1.42552061 | 1.70658508 | 0.0878992 | 0.45312723 |
| Fignl1 | 4802.93275 | 0.07225312 | 0.15643656 | 0.46186848 | 0.64417564 | 0.90823885 |
| Rcbtb1 | 1173.39772 | 0.39006806 | 0.30163098 | 1.29319628 | 0.19594318 | 0.62780852 |
| Galm | 266.251407 | 0.83976623 | 0.5274298 | 1.59218579 | 0.11134297 | 0.50888686 |
| Tab3 | 63.5298505 | 0.42668405 | 0.62413428 | 0.68364142 | 0.49420162 | 0.84678075 |
| Mbd3 | 1175.51636 | -0.4309704 | 0.31668557 | -1.3608779 | 0.17355227 | 0.60653633 |
| Tgfbi | 4007.39774 | 0.77308381 | 0.24088611 | 3.20933335 | 0.00133043 | 0.02875471 |
| Tstd2 | 2375.50514 | -0.0039407 | 0.2022348 | -0.019486 | 0.98445342 | 0.99564261 |
| Reep6 | 21.4160815 | 0.23636877 | 1.42298177 | 0.16610808 | 0.8680719 | 0.97173531 |
| Cox18 | 527.569084 | -0.4168205 | 0.25858497 | -1.6119285 | 0.10697751 | 0.49986855 |
| Fbxl21 | 17.0572976 | 0.52315182 | 1.5739284 | 0.33238604 | 0.73959778 | 0.93800224 |
| Tdrd7 | 524.244581 | 0.3586822 | 0.37767709 | 0.94970601 | 0.34226166 | 0.75889838 |
| Gnptg | 809.620616 | 0.26560279 | 0.29788681 | 0.8916232 | 0.37259493 | 0.77748705 |
| Prdm4 | 402.293424 | 0.08121937 | 0.47718877 | 0.17020385 | 0.86484982 | 0.9705375 |
| Eif1 | 16381.0173 | 0.11640973 | 0.23063329 | 0.50473949 | 0.61374182 | 0.89776035 |
| Ccdc180 | 24.1773706 | -1.7982457 | 0.85775802 | -2.0964487 | 0.0360424 | 0.28482346 |
| Leng8 | 570.109207 | 0.07780982 | 0.3507897 | 0.2218133 | 0.82445923 | 0.962942 |
| Capn5 | 22.3300374 | -0.7392597 | 1.48231889 | -0.4987184 | 0.61797778 | 0.89878056 |
| Mpv17l2 | 1191.17263 | 0.20077887 | 0.24647137 | 0.81461337 | 0.41529365 | 0.80395688 |
| Aldh1b1 | 198.052778 | -0.4590793 | 0.40733529 | -1.1270305 | 0.25972957 | 0.69596146 |
| Ankrd11 | 765.974539 | -0.1959874 | 0.33329379 | -0.5880318 | 0.55651093 | 0.87276537 |
| Dcaf10 | 165.13659 | 0.77659625 | 0.59055424 | 1.31502951 | 0.18849998 | 0.61935434 |
| Utp6 | 2298.44524 | -0.2307394 | 0.16059035 | -1.4368198 | 0.15076921 | 0.57859667 |
| Iqcg | 90.2690766 | 0.62136337 | 0.63592895 | 0.97709559 | 0.32852183 | 0.74843736 |
| Tsen34 | 2239.43622 | 0.24236096 | 0.14918427 | 1.62457443 | 0.10425327 | 0.49431405 |
| 1600002K03I | 256.750261 | 0.20949682 | 0.47391286 | 0.44205768 | 0.65844747 | 0.91265586 |
| Mboat7 | 1326.32196 | -0.4048567 | 0.18197453 | -2.2247985 | 0.02609477 | 0.23668568 |
| Prpf39 | 1202.63919 | -0.1335752 | 0.28915633 | -0.4619482 | 0.64411848 | 0.90823885 |
| Trmt10b | 421.843564 | 0.63633039 | 0.50738332 | 1.25414133 | 0.20979064 | 0.64350283 |
| Togaram1 | 360.536802 | -0.2470626 | 0.45119595 | -0.5475727 | 0.58398533 | 0.88404713 |
| Ric8b | 462.187844 | 0.18254836 | 0.29467197 | 0.61949686 | 0.53558909 | 0.86539693 |
| Midn | 123.94493 | 0.0581234 | 0.73973632 | 0.07857314 | 0.93737215 | 0.98845735 |
| Rsf1 | 475.53469 | 0.21500739 | 0.28609582 | 0.75152233 | 0.45233837 | 0.82603347 |
| Rubcn | 373.955296 | 0.67000821 | 0.48384652 | 1.38475359 | 0.16612783 | 0.59942225 |
| Cnot3 | 639.539392 | -0.2251665 | 0.25944704 | -0.8678708 | 0.38546504 | 0.78813202 |
| Grhpr | 1569.57591 | -0.2150463 | 0.18095488 | -1.1883973 | 0.2346769 | 0.67038085 |
| Aamdc | 279.882565 | 0.33240035 | 0.47467707 | 0.70026629 | 0.48376102 | 0.84128398 |
| Zcchc7 | 463.655119 | -0.0690155 | 0.35717584 | -0.1932255 | 0.84678234 | 0.96765585 |
| Gtf3c4 | 425.533785 | -0.3718927 | 0.42957413 | -0.8657242 | 0.38664142 | 0.78911204 |
| Zswim4 | 172.702438 | -0.7581665 | 0.51414231 | -1.4746238 | 0.14031371 | 0.56406756 |
| Sbno2 | 820.729901 | -0.2920224 | 0.37945947 | -0.7695746 | 0.44155228 | 0.81967485 |
| Ndufa3 | 2158.97287 | 0.26506265 | 0.37306024 | 0.71050899 | 0.47738856 | 0.836949 |
| Tnfsf9 | 1837.17561 | 0.20514514 | 0.51245891 | 0.4003153 | 0.6889243 | 0.92470267 |
| Melk | 2690.22589 | -0.1556836 | 0.18348143 | -0.848498 | 0.39616071 | 0.79433436 |

| | | | | | | |
|---|---|---|---|---|---|---|
| Isg15 | 45.1079141 | -1.3981926 | 0.69468498 | -2.0127002 | 0.04414618 | 0.32226735 |
| Rnf38 | 263.985858 | -0.1099097 | 0.33011837 | -0.3329403 | 0.73917932 | 0.93800224 |
| Arhgap45 | 2774.5893 | -0.5133787 | 0.28188487 | -1.8212354 | 0.06857108 | 0.40386619 |
| Alg8 | 2879.52655 | -0.1703539 | 0.20210886 | -0.8428822 | 0.39929435 | 0.79612475 |
| Dok3 | 1407.50937 | -0.0453043 | 0.17132706 | -0.2644317 | 0.79144732 | 0.95268772 |
| Usp35 | 9.94436636 | 0.839276 | 1.7109093 | 0.49054383 | 0.62374912 | 0.90017327 |
| Abca7 | 150.835574 | -0.8216202 | 0.54467626 | -1.508456 | 0.13143787 | 0.54970237 |
| Prkx | 1305.5651 | -0.2428546 | 0.27108972 | -0.895846 | 0.37033501 | 0.77600345 |
| Supt16 | 3643.78094 | -0.350001 | 0.28066465 | -1.2470433 | 0.21238161 | 0.64580271 |
| Dagla | 8.36719661 | 3.5814676 | 1.60668376 | 2.22910549 | 0.02580689 | 0.23454357 |
| Wdr18 | 8782.57224 | -0.1133374 | 0.20186402 | -0.561454 | 0.57448809 | 0.88059206 |
| Selenoo | 599.464145 | 0.43515768 | 0.27795098 | 1.56559147 | 0.11744429 | 0.52372617 |
| Bbs10 | 95.9575574 | 0.32377737 | 0.85771127 | 0.37748993 | 0.70580954 | 0.92896305 |
| Tmem161b | 429.429637 | 0.25891816 | 0.28865019 | 0.89699631 | 0.36972089 | 0.77508072 |
| Fbxo45 | 187.976909 | -0.3625322 | 0.55484263 | -0.6533965 | 0.51350072 | 0.85497712 |
| Dym | 1389.86906 | 0.03850154 | 0.24451784 | 0.15745904 | 0.87488309 | 0.97366476 |
| Xylb | 282.023277 | 0.04983218 | 0.41301676 | 0.12065414 | 0.90396499 | 0.98077195 |
| Dync1li2 | 1330.50838 | 0.47414702 | 0.29843463 | 1.58878018 | 0.11211003 | 0.51001933 |
| Mrps2 | 2296.74022 | -0.1847547 | 0.22260263 | -0.8299753 | 0.40655274 | 0.79992362 |
| Cd99l2 | 720.718573 | 0.53668658 | 0.35634378 | 1.50609217 | 0.13204352 | 0.55148116 |
| Ggta1 | 3106.79562 | 0.31582457 | 0.24244454 | 1.30266727 | 0.19268838 | 0.62446288 |
| R3hdm4 | 1823.27607 | -0.339752 | 0.18992872 | -1.7888397 | 0.07364063 | 0.41763833 |
| Acta2 | 82.3108465 | 0.75500339 | 0.48540571 | 1.5554069 | 0.11984919 | 0.52718991 |
| Cep19 | 377.082497 | -0.2173214 | 0.2380023 | -0.9131065 | 0.36118655 | 0.76887042 |
| Zdhhc17 | 319.32792 | 0.08818506 | 0.32793448 | 0.26891062 | 0.78799846 | 0.95225386 |
| Tk2 | 1167.89618 | 0.60641106 | 0.21432295 | 2.82942664 | 0.00466315 | 0.07545542 |
| Pim3 | 746.793009 | -0.1834069 | 0.32083993 | -0.571646 | 0.56756183 | 0.87929304 |
| Polr3g | 554.181057 | 0.17341501 | 0.41233848 | 0.42056471 | 0.67407297 | 0.91822108 |
| Plppr3 | 614.140875 | -1.7593621 | 0.28737031 | -6.122282 | 9.224455328 | 1.659452382 |
| Lysmd3 | 652.774615 | 0.6781519 | 0.31685409 | 2.14026557 | 0.03233331 | 0.26700278 |
| Ddx11 | 1419.96823 | -0.2339558 | 0.27135651 | -0.8621713 | 0.38859323 | 0.78942657 |
| Alg12 | 741.956102 | 0.01205085 | 0.22767882 | 0.05292919 | 0.95778833 | 0.99242502 |
| Ids | 54.7959449 | 0.00178663 | 0.64827362 | 0.00275598 | 0.99780105 | 0.99967651 |
| Ythdc1 | 824.299297 | 0.28958752 | 0.42329111 | 0.68413325 | 0.49389103 | 0.84678075 |
| Palm | 182.302403 | -1.6915349 | 0.45996873 | -3.6774998 | 0.00023553 | 0.00737525 |
| Zfp983 | 161.784909 | 0.33159165 | 0.5914976 | 0.56059678 | 0.57507245 | 0.88059206 |
| Pawr | 6.22500507 | 0.78149836 | 2.41592224 | 0.32347828 | 0.74633306 | 0.93960567 |
| Zhx3 | 278.329864 | 1.18608506 | 0.52361731 | 2.26517541 | 0.02350192 | 0.22081333 |
| Hykk | 38.9078556 | 1.23153261 | 1.11726971 | 1.10226975 | 0.27034442 | 0.70733226 |
| Cox8a | 8532.26232 | -0.1244116 | 0.21004185 | -0.5923182 | 0.55363753 | 0.87164066 |
| Rnf126 | 181.015143 | -0.334918 | 0.37749906 | -0.8872023 | 0.37496997 | 0.77815049 |
| Cerk | 544.046985 | -0.1798029 | 0.35715362 | -0.5034329 | 0.61465995 | 0.89844745 |
| Uba6 | 666.323134 | 0.19949087 | 0.30542885 | 0.65315005 | 0.51365955 | 0.85497712 |
| Gramd4 | 64.5809496 | -1.115382 | 1.0377767 | -1.0747803 | 0.2824731 | 0.71787592 |
| Dennd5a | 511.855581 | 0.11832572 | 0.34436828 | 0.34360227 | 0.73114542 | 0.93653187 |
| Cd276 | 492.304984 | 0.48949619 | 0.30839207 | 1.58725283 | 0.11245539 | 0.51121027 |

| | | | | | | |
|---|---|---|---|---|---|---|
| Bbs9 | 198.967103 | -0.3369776 | 0.51151489 | -0.6587836 | 0.51003476 | 0.85335831 |
| Cog5 | 901.70076 | 0.02133933 | 0.20451106 | 0.10434317 | 0.91689702 | 0.98389485 |
| Pknox2 | 15.6714346 | 1.03830024 | 2.50011895 | 0.41530034 | 0.67792206 | 0.91965617 |
| Aldh5a1 | 80.2105138 | 0.47006549 | 0.88740205 | 0.52970971 | 0.59631321 | 0.88873045 |
| Ibtk | 762.155343 | -0.7433382 | 0.4519509 | -1.6447323 | 0.10002504 | 0.48325682 |
| Ttc38 | 224.324546 | -0.3338032 | 0.36438227 | -0.9160797 | 0.35962509 | 0.76857232 |
| Fbxw2 | 1203.74026 | 0.05844519 | 0.24638701 | 0.23720889 | 0.81249474 | 0.95967622 |
| Pip4p1 | 1403.28242 | 0.6583373 | 0.39153167 | 1.68144074 | 0.09267734 | 0.46597694 |
| Dock4 | 129.83078 | 0.49044545 | 0.41667554 | 1.17704404 | 0.23917793 | 0.67535535 |
| Tdp2 | 2204.30248 | 0.10186929 | 0.20577448 | 0.49505307 | 0.62056266 | 0.89896588 |
| Apex1 | 10118.1738 | -0.1732613 | 0.18620813 | -0.930471 | 0.35212725 | 0.76348958 |
| Ints6l | 2750.90932 | 0.66393452 | 0.16364184 | 4.05724179 | 4.965566782 | 0.00200475 |
| Fnip1 | 354.610682 | 0.07935957 | 0.53604953 | 0.14804521 | 0.88230709 | 0.97598447 |
| Fam214b | 245.655379 | 0.29475887 | 0.42523708 | 0.69316361 | 0.48820688 | 0.84329777 |
| Ripor2 | 2046.97679 | 0.03803041 | 0.27067231 | 0.1405035 | 0.88826219 | 0.97832097 |
| Mettl25 | 455.993294 | 0.62775029 | 0.28664889 | 2.18996237 | 0.02852697 | 0.25141958 |
| Tmtc2 | 20.4773314 | -0.7558717 | 1.42088812 | -0.5319713 | 0.59474585 | 0.88830085 |
| Fam122b | 315.350695 | -0.0789122 | 0.41038557 | -0.1922879 | 0.84751669 | 0.96765585 |
| Parp2 | 3120.82697 | -0.079622 | 0.16899935 | -0.4711379 | 0.63754227 | 0.90599527 |
| Tmem63b | 106.008932 | 0.64369854 | 0.69549113 | 0.92553092 | 0.35468981 | 0.76419569 |
| Prtg | 145.861056 | -1.7798974 | 1.00093066 | -1.7782425 | 0.07536404 | 0.42273804 |
| Zfp57 | 22.8653474 | 3.24205169 | 1.76375193 | 1.83815628 | 0.06603938 | 0.39562181 |
| 5031439G07 | 1404.28758 | -0.0934877 | 0.30125208 | -0.3103303 | 0.75630978 | 0.94102069 |
| Dnajb5 | 38.6834046 | 1.19419665 | 1.29332757 | 0.92335204 | 0.35582378 | 0.76499791 |
| Fmnl2 | 34.8572946 | 1.19101822 | 1.12500581 | 1.0586774 | 0.28974673 | 0.72395811 |
| Sugp2 | 462.1261 | 0.03590378 | 0.42753231 | 0.0839791 | 0.93307305 | 0.98748011 |
| Ptpn23 | 202.022736 | 0.92488129 | 0.47246907 | 1.95754884 | 0.05028297 | 0.34299029 |
| Smug1 | 582.562514 | 0.17455909 | 0.30095047 | 0.58002597 | 0.5618971 | 0.87540592 |
| Slc2a6 | 250.416269 | -1.8136326 | 0.41959524 | -4.3223382 | 1.543842653 | 0.00075307 |
| Galt | 811.337959 | 0.3709722 | 0.19893217 | 1.86481756 | 0.06220698 | 0.38511032 |
| Sigmar1 | 1428.89599 | -0.0889298 | 0.20181517 | -0.4406497 | 0.65946664 | 0.91280998 |
| Zranb3 | 396.017335 | -0.1246891 | 0.34105339 | -0.3656 | 0.71466354 | 0.93235217 |
| Slain2 | 413.498699 | 0.31573605 | 0.48798845 | 0.64701542 | 0.51762197 | 0.85695893 |
| Hyal3 | 12.9920148 | -0.9820162 | 1.50400409 | -0.6529345 | 0.51379849 | 0.85497712 |
| Arl5a | 454.761513 | -0.4150523 | 0.29364934 | -1.4134282 | 0.15752986 | 0.58620126 |
| Slf2 | 1231.63992 | 0.02330265 | 0.33321764 | 0.06993223 | 0.9442476 | 0.99005579 |
| Vezt | 820.221864 | 0.61920932 | 0.27878747 | 2.22108013 | 0.02634553 | 0.23837642 |
| Colec12 | 1535.00212 | 1.84275789 | 0.34507099 | 5.34022832 | 9.282959409 | 9.043841866 |
| Rab3gap1 | 1441.88676 | 0.20565543 | 0.29557357 | 0.69578423 | 0.48656397 | 0.84183502 |
| Prr5 | 811.566745 | -0.2672455 | 0.55556545 | -0.4810334 | 0.63049275 | 0.9036296 |
| Mbnl3 | 227.008609 | -1.5239254 | 0.48134523 | -3.1659717 | 0.00154566 | 0.03254395 |
| Lmo1 | 32.1747426 | -0.4737349 | 0.72526204 | -0.6531914 | 0.51363289 | 0.85497712 |
| Metap2 | 9621.90483 | -0.2684575 | 0.18812418 | -1.4270229 | 0.15357328 | 0.58242478 |
| Rpp25l | 889.12004 | -0.0080075 | 0.25913781 | -0.0309007 | 0.97534876 | 0.99450266 |
| Rfxank | 437.324039 | 0.35216604 | 0.37924961 | 0.92858642 | 0.35310346 | 0.76376846 |
| Acaa1a | 5012.02347 | 0.49544902 | 0.23992292 | 2.06503412 | 0.03891977 | 0.29821741 |

| | | | | | | |
|---|---|---|---|---|---|---|
| Mgat5 | 269.872122 | -0.3221803 | 0.33179075 | -0.9710346 | 0.33153103 | 0.75062356 |
| Prickle1 | 227.440277 | 0.16786389 | 0.87180569 | 0.19254737 | 0.84731347 | 0.96765585 |
| Surf6 | 1138.93649 | -0.3404786 | 0.23724493 | -1.4351352 | 0.15124856 | 0.57926708 |
| Pphln1 | 3460.63736 | -0.210438 | 0.18031821 | -1.167037 | 0.24319544 | 0.67814218 |
| Ccdc38 | 12.7112378 | -0.1692165 | 1.08216898 | -0.1563679 | 0.87574307 | 0.97417159 |
| Gatad2a | 3277.36984 | -0.2500551 | 0.24386318 | -1.0253907 | 0.30517885 | 0.73518175 |
| Hist1h1c | 421.760183 | 0.67573219 | 0.32802225 | 2.06001934 | 0.03939669 | 0.30064861 |
| Ankmy2 | 847.318982 | -0.0031708 | 0.21271225 | -0.0149064 | 0.98810687 | 0.9962454 |
| Gxylt1 | 369.376234 | -0.0729812 | 0.47719732 | -0.1529371 | 0.87844792 | 0.97518697 |
| Ndufa13 | 3885.39073 | 0.05923348 | 0.26960677 | 0.21970323 | 0.82610229 | 0.96345784 |
| Rif1 | 582.954134 | -0.522509 | 0.42267868 | -1.2361849 | 0.21638986 | 0.64982959 |
| Sh3bp4 | 128.8428 | -0.4454446 | 0.49630894 | -0.8975148 | 0.3694443 | 0.77502139 |
| Nepro | 511.128364 | -0.0156468 | 0.36553802 | -0.0428049 | 0.96585703 | 0.99443087 |
| Znrd1as | 149.024657 | 1.20281365 | 0.53613867 | 2.24347491 | 0.02486621 | 0.2288851 |
| Ska1 | 1200.18188 | 0.39185998 | 0.43978162 | 0.8910331 | 0.37291141 | 0.77748705 |
| Ube2r2 | 1465.08823 | 0.19047301 | 0.19266811 | 0.98860684 | 0.32285554 | 0.74560918 |
| Armh4 | 18.439623 | 0.52950308 | 2.20941972 | 0.23965708 | 0.81059611 | 0.95905029 |
| Gmip | 2519.85212 | -0.219803 | 0.27446982 | -0.8008274 | 0.42323156 | 0.80870984 |
| Rbm43 | 1124.00064 | 0.15723275 | 0.38327877 | 0.41023078 | 0.68163667 | 0.92061965 |
| Pnpla8 | 1656.48722 | 0.30350884 | 0.21854057 | 1.38879862 | 0.164894 | 0.59731959 |
| Edc4 | 1504.76197 | -0.2608768 | 0.27535421 | -0.9474225 | 0.34342351 | 0.75947281 |
| Lrrk2 | 34.8144514 | -0.3137066 | 0.94822141 | -0.3308369 | 0.74076768 | 0.93800224 |
| 9530068E07I | 8758.60778 | 0.13574071 | 0.16650638 | 0.81522829 | 0.41494165 | 0.80395688 |
| Snapc4 | 393.671419 | -0.1157039 | 0.41356053 | -0.2797749 | 0.77965019 | 0.9488468 |
| Naa30 | 154.993852 | -0.4328747 | 0.34507081 | -1.2544517 | 0.20967786 | 0.64347165 |
| Noa1 | 1202.09007 | -0.3102887 | 0.17026559 | -1.8223806 | 0.06839726 | 0.40325004 |
| Ap5m1 | 1167.0574 | 0.03043211 | 0.33232326 | 0.09157382 | 0.92703665 | 0.98638248 |
| Gramd1c | 14.003369 | -0.96286 | 1.62490362 | -0.5925644 | 0.55347271 | 0.87160552 |
| BC031181 | 2613.07365 | 0.08698349 | 0.27474576 | 0.31659629 | 0.75154995 | 0.94018119 |
| Zdhhc23 | 12.9710083 | 1.84468327 | 1.35968981 | 1.3566942 | 0.1748784 | 0.60708046 |
| Lzts1 | 32.9401321 | 0.19592877 | 1.10297797 | 0.17763616 | 0.85900873 | 0.96954641 |
| Skp1a | 7029.58797 | 0.2845198 | 0.31836669 | 0.89368583 | 0.37149001 | 0.7765614 |
| Znrd1 | 774.216835 | -0.0532648 | 0.21042615 | -0.2531285 | 0.80016894 | 0.95543842 |
| Srp72 | 4805.64351 | -0.2351653 | 0.18074689 | -1.3010752 | 0.1932327 | 0.624687 |
| Qsox2 | 786.360557 | -0.0265575 | 0.19402182 | -0.136879 | 0.89112641 | 0.97889088 |
| Slc18a1 | 28.6780765 | 2.21713094 | 1.81703848 | 1.22018932 | 0.22239312 | 0.65586668 |
| Kidins220 | 599.129838 | 0.0534537 | 0.35538421 | 0.15041101 | 0.88044036 | 0.97567052 |
| Igsf10 | 444.220691 | 1.09983461 | 0.44443497 | 2.47468063 | 0.01333554 | 0.15610873 |
| Tmem260 | 421.720417 | 0.38582036 | 0.35643662 | 1.08243749 | 0.27905818 | 0.71377266 |
| Ubac1 | 526.612344 | -0.0646116 | 0.25905014 | -0.2494174 | 0.80303794 | 0.95672466 |
| P2ry12 | 2673.20078 | 0.89550862 | 0.20841596 | 4.29673729 | 1.733304083 | 0.00081867 |
| P2ry13 | 2679.61588 | 0.49381238 | 0.20293567 | 2.43334445 | 0.01496006 | 0.16898101 |
| Rmdn2 | 143.663985 | 0.38983819 | 0.80576711 | 0.48381001 | 0.62852071 | 0.90276005 |
| Serbp1 | 17487.4068 | -0.3704746 | 0.17372989 | -2.1324747 | 0.03296784 | 0.26976292 |
| Tmem258 | 2329.16909 | 0.15880458 | 0.2865295 | 0.55423468 | 0.57941825 | 0.88286755 |
| Abt1 | 1234.71938 | -0.1737919 | 0.19251916 | -0.9027251 | 0.3666718 | 0.77269528 |

| | | | | | | |
|---|---|---|---|---|---|---|
| C530008M17 | 17.7994713 | -1.9262431 | 1.32538969 | -1.4533409 | 0.14612912 | 0.5731306 |
| P2ry14 | 212.884023 | -1.848664 | 0.34174321 | -5.4095119 | 6.319675538 | 6.722486161 |
| Gadd45a | 43.5095087 | -0.0982471 | 0.74058014 | -0.1326624 | 0.89446037 | 0.97941957 |
| Sec24a | 562.470943 | 0.1241987 | 0.40993572 | 0.30297116 | 0.76191184 | 0.94298237 |
| Glb1l2 | 23.304312 | -0.1110741 | 1.39190088 | -0.0798003 | 0.93639607 | 0.98826099 |
| Ppp1r11 | 1645.70016 | 0.35455965 | 0.37422307 | 0.94745536 | 0.34340679 | 0.75947281 |
| Gng12 | 3072.85607 | 0.6146657 | 0.16430992 | 3.74089221 | 0.00018337 | 0.00607898 |
| Cep135 | 305.575211 | -0.3736158 | 0.45997674 | -0.8122493 | 0.41664862 | 0.8047849 |
| 9530077C05 | 78.5450281 | 0.81088473 | 0.84382403 | 0.96096426 | 0.33657014 | 0.75494233 |
| Gpi1 | 34824.8402 | -0.2533715 | 0.16771068 | -1.5107655 | 0.13084822 | 0.54930207 |
| Tbcc | 822.941928 | -0.1503516 | 0.19234584 | -0.7816732 | 0.43440667 | 0.81542071 |
| Siah2 | 68.3477775 | -0.5615348 | 0.68838188 | -0.8157315 | 0.41465374 | 0.80395688 |
| Exoc1 | 2220.22797 | 0.03869231 | 0.2059835 | 0.18784178 | 0.85100068 | 0.9679302 |
| Calm2 | 26547.6459 | 0.38634112 | 0.19063452 | 2.02660626 | 0.0427027 | 0.31653532 |
| Thap11 | 1734.94433 | -0.1313039 | 0.21276724 | -0.6171244 | 0.53715269 | 0.86608526 |
| Lum | 21.8318842 | 1.37821399 | 1.40005047 | 0.98440308 | 0.32491737 | 0.74628504 |
| Hif1an | 566.759569 | -0.401388 | 0.33608007 | -1.1943225 | 0.23235185 | 0.6674267 |
| Arhgap26 | 156.990788 | -1.7151085 | 0.38949281 | -4.4034407 | 1.065474259 | 0.00054996 |
| Elf1 | 2714.34837 | -0.3748799 | 0.22373146 | -1.6755798 | 0.09382053 | 0.46826055 |
| March1 | 1089.78387 | 0.29535832 | 0.28928284 | 1.02100185 | 0.30725356 | 0.73640925 |
| Tbc1d24 | 146.413542 | 0.20531195 | 0.54017849 | 0.38008168 | 0.70388478 | 0.92850227 |
| Btg1 | 3161.64614 | -0.0393562 | 0.28577084 | -0.1377195 | 0.89046209 | 0.97871887 |
| Eea1 | 896.316918 | 0.14494726 | 0.27818661 | 0.52104328 | 0.60233662 | 0.89130082 |
| Fam13b | 294.383888 | 0.099786 | 0.28529539 | 0.34976379 | 0.72651598 | 0.93576253 |
| Tmem255a | 110.764637 | 1.39280908 | 0.54539583 | 2.5537582 | 0.01065672 | 0.13344172 |
| Rnf13 | 6111.98409 | 0.54129623 | 0.21990304 | 2.46152227 | 0.01383488 | 0.15951187 |
| Phpt1 | 2057.87998 | 0.55147621 | 0.3777498 | 1.4598981 | 0.14431808 | 0.57027792 |
| Commd2 | 2940.58229 | 0.99670364 | 0.5013691 | 1.98796382 | 0.04681569 | 0.33444287 |
| Greb1 | 18.1816429 | -2.715515 | 1.64701425 | -1.6487501 | 0.09919883 | 0.48078421 |
| Card11 | 46.1503278 | 0.05886084 | 0.82274545 | 0.07154199 | 0.94296641 | 0.98983253 |
| Ppfibp2 | 291.797074 | 0.71118442 | 0.43167995 | 1.64748079 | 0.09945926 | 0.48166474 |
| Sbf1 | 416.42444 | -0.5049945 | 0.49983469 | -1.0103231 | 0.31234053 | 0.74051791 |
| Cdc42ep3 | 269.209338 | -1.3235386 | 0.43376779 | -3.0512606 | 0.00227883 | 0.04383158 |
| Slc38a7 | 2346.31013 | 0.44202875 | 0.27386415 | 1.61404389 | 0.10651791 | 0.49867338 |
| Rnf113a1 | 104.381464 | -1.1361088 | 0.63177338 | -1.7982853 | 0.0721318 | 0.41303124 |
| Cnot1 | 1906.11194 | -0.1183508 | 0.38738978 | -0.3055083 | 0.75997907 | 0.94215089 |
| Ermard | 588.091247 | 0.10349911 | 0.29224442 | 0.35415256 | 0.72322455 | 0.93433544 |
| Sh3tc1 | 1352.45719 | -0.1469299 | 0.18557089 | -0.7917726 | 0.42849331 | 0.81136783 |
| Iqce | 181.533025 | 0.30761699 | 0.38691057 | 0.79505968 | 0.42657879 | 0.81021713 |
| Lgi4 | 15.7734337 | -0.1279864 | 1.42402231 | -0.0898767 | 0.9283852 | 0.98666423 |
| Ppp6r2 | 245.373206 | 0.58591818 | 0.34254189 | 1.71050083 | 0.0871733 | 0.45109601 |
| Ndrg4 | 22.7489868 | 0.3531997 | 0.85391187 | 0.41362547 | 0.67914842 | 0.92031392 |
| Ttyh3 | 108.024828 | -0.4696941 | 0.46000668 | -1.0210592 | 0.30722637 | 0.73640925 |
| Bicral | 139.590569 | -0.7654874 | 0.69276198 | -1.104979 | 0.26916869 | 0.7061421 |
| Upf3b | 275.975515 | -0.6384862 | 0.36867732 | -1.7318294 | 0.08330394 | 0.44341423 |
| Spg20 | 1299.87098 | 0.03555242 | 0.31209395 | 0.11391576 | 0.90930457 | 0.98229626 |

| | | | | | | |
|---|---|---|---|---|---|---|
| Arhgap21 | 212.936052 | -0.5178862 | 0.53488321 | -0.9682229 | 0.33293304 | 0.7521274 |
| Chst12 | 2273.01757 | -0.6590953 | 0.23648901 | -2.7870017 | 0.00531982 | 0.08375465 |
| Plxnb2 | 5010.86929 | 0.29655501 | 0.17536619 | 1.6910615 | 0.09082506 | 0.46044203 |
| Eepd1 | 189.767433 | 0.02419233 | 0.53478034 | 0.04523787 | 0.96391771 | 0.99406401 |
| Eipr1 | 2088.15533 | -0.2055852 | 0.22215746 | -0.9254032 | 0.3547562 | 0.76419569 |
| Rfxap | 24.6497836 | -1.3189404 | 1.30797677 | -1.0083821 | 0.31327105 | 0.74065408 |
| Mgat4b | 308.294187 | 0.27160473 | 0.29580284 | 0.91819514 | 0.35851671 | 0.76713922 |
| Atp13a2 | 6790.03493 | 0.37725379 | 0.29674788 | 1.27129397 | 0.20362409 | 0.63809363 |
| Alg5 | 3093.50819 | 0.4920196 | 0.39719748 | 1.23872792 | 0.21544627 | 0.64854028 |
| Mag | 51.272785 | 1.3026773 | 0.8545769 | 1.52435351 | 0.12742043 | 0.54145382 |
| Clcn7 | 1384.26658 | 0.26826776 | 0.26258149 | 1.02165524 | 0.3069441 | 0.73595594 |
| Nudt1 | 974.879415 | -0.0761913 | 0.39890544 | -0.191001 | 0.8485248 | 0.96765585 |
| Tbc1d9b | 2020.41505 | 0.41095373 | 0.20379673 | 2.01648838 | 0.04374893 | 0.32046747 |
| Man1b1 | 714.520908 | 0.30689348 | 0.2206357 | 1.39095115 | 0.16424024 | 0.59585062 |
| Dennd3 | 89.921733 | -0.7516758 | 0.91548982 | -0.8210641 | 0.41160975 | 0.80231592 |
| Tcaf1 | 512.234527 | 1.06887552 | 0.29067006 | 3.67728111 | 0.00023573 | 0.00737525 |
| Cenpt | 1815.55716 | 0.07511098 | 0.23775213 | 0.31592139 | 0.75206218 | 0.94018119 |
| Tmtc3 | 707.988254 | 0.65873379 | 0.2833038 | 2.32518515 | 0.02006206 | 0.2001788 |
| Aaas | 3657.06955 | -0.1838673 | 0.15053839 | -1.2213979 | 0.22193539 | 0.65525654 |
| Cc2d1a | 228.178565 | 0.02211769 | 0.34057073 | 0.06494303 | 0.94821936 | 0.99019648 |
| Nop14 | 2076.86685 | -0.3862103 | 0.25939814 | -1.4888708 | 0.1365214 | 0.55911158 |
| Ago2 | 1131.64436 | -0.1652236 | 0.36693461 | -0.4502808 | 0.65250797 | 0.91051509 |
| Cab39 | 1251.96114 | -0.1625849 | 0.22556252 | -0.7207974 | 0.47103415 | 0.83381643 |
| Cyld | 688.991002 | -0.2025601 | 0.43287483 | -0.4679415 | 0.63982639 | 0.90663689 |
| Micall2 | 21.059251 | -0.7184879 | 0.93226389 | -0.7706916 | 0.44088977 | 0.81967485 |
| Zscan12 | 292.114751 | -0.0105895 | 0.36515574 | -0.0289999 | 0.97686469 | 0.99483019 |
| Rbm42 | 3640.11912 | -0.1820272 | 0.158471 | -1.148647 | 0.25070159 | 0.6842534 |
| Oxsr1 | 551.476456 | 0.278766 | 0.24398493 | 1.14255417 | 0.25322374 | 0.68647781 |
| Psma8 | 413.800385 | 0.6570725 | 0.4861803 | 1.35149964 | 0.17653544 | 0.60764153 |
| Cuedc2 | 3984.90347 | 0.22745954 | 0.30314866 | 0.75032343 | 0.45305994 | 0.82610332 |
| Cox6b1 | 11360.681 | 0.57656578 | 0.43288008 | 1.33192957 | 0.18288333 | 0.61444982 |
| Tubb4b | 22128.1966 | 0.09810465 | 0.3161337 | 0.31032644 | 0.75631273 | 0.94102069 |
| Dnajc12 | 363.928572 | 1.01623439 | 0.37516759 | 2.70874783 | 0.00675377 | 0.09966081 |
| Dner | 20.7883354 | 0.06197797 | 1.3800577 | 0.0449097 | 0.96417929 | 0.99406401 |
| Kif15 | 2495.35255 | -0.2180238 | 0.17744798 | -1.2286632 | 0.21919809 | 0.65257977 |
| Wdr44 | 180.65607 | 0.29014277 | 0.45443572 | 0.63846823 | 0.52316893 | 0.85870462 |
| Decr2 | 825.89348 | 0.58523414 | 0.32469602 | 1.80240628 | 0.07148151 | 0.41122734 |
| Anln | 1874.88921 | 0.05218005 | 0.25749605 | 0.20264406 | 0.83941325 | 0.96757914 |
| Tent4b | 261.901666 | 0.09533826 | 0.44229311 | 0.21555448 | 0.82933504 | 0.96462215 |
| Rps27l | 9139.9655 | 0.41180032 | 0.42489747 | 0.96917575 | 0.33245751 | 0.75146946 |
| Klhl13 | 554.571652 | 1.60457161 | 0.4442384 | 3.61196062 | 0.00030389 | 0.0090012 |
| Mbd5 | 78.6075506 | 0.78535702 | 0.63331877 | 1.24006592 | 0.21495101 | 0.64854028 |
| Cnep1r1 | 739.719915 | 0.35806559 | 0.30996455 | 1.15518239 | 0.24801574 | 0.68235696 |
| Sun1 | 1061.12671 | -0.0890531 | 0.39977358 | -0.2227589 | 0.82372319 | 0.9628698 |
| Jmjd4 | 395.347138 | -0.0069894 | 0.30913175 | -0.0226099 | 0.98196147 | 0.99523682 |
| Amdhd2 | 1848.05172 | 0.39965416 | 0.20944715 | 1.90813847 | 0.05637332 | 0.36517938 |

| | | | | | | |
|---|---|---|---|---|---|---|
| Topors | 3195.54474 | 0.07896333 | 0.20278754 | 0.38938945 | 0.69698808 | 0.92650558 |
| Ssx2ip | 69.5794796 | 0.51675468 | 0.52605169 | 0.98232682 | 0.32593887 | 0.74750846 |
| Pnpla7 | 1151.78756 | 0.17039862 | 0.32975072 | 0.51674979 | 0.60533084 | 0.89258167 |
| Psenen | 1560.46362 | 0.06411574 | 0.22696567 | 0.2824909 | 0.77756712 | 0.94837927 |
| Siah1a | 27.129961 | -0.1831105 | 1.06900362 | -0.1712908 | 0.86399513 | 0.9705375 |
| Lin37 | 475.495921 | -0.1240241 | 0.22912914 | -0.5412846 | 0.58831144 | 0.88690478 |
| Mrpl41 | 2426.73119 | 0.7021797 | 0.50382504 | 1.39369752 | 0.16340897 | 0.59475807 |
| Mcoln3 | 10.336857 | 0.70512314 | 2.20613488 | 0.31961923 | 0.74925699 | 0.9399191 |
| Hspb6 | 58.1160648 | 0.40332611 | 0.55573482 | 0.72575282 | 0.46799031 | 0.83381643 |
| Ptcra | 28.7487193 | -0.4152146 | 0.87132296 | -0.4765335 | 0.63369432 | 0.90455392 |
| Mrpl55 | 1849.88383 | 0.02684879 | 0.30033531 | 0.08939605 | 0.92876716 | 0.986679 |
| Dchs1 | 28.9723867 | 3.0027051 | 1.38939148 | 2.16116562 | 0.03068255 | 0.26119667 |
| Proser3 | 35.1505976 | 0.34289752 | 0.91599219 | 0.37434547 | 0.70814731 | 0.92967665 |
| 2410004B18I | 1333.45636 | 0.12677284 | 0.20992939 | 0.60388323 | 0.54592128 | 0.86990153 |
| Dna2 | 1659.94745 | -0.8424427 | 0.21977697 | -3.8331709 | 0.0001265 | 0.00442143 |
| Phkb | 914.750617 | -0.0714762 | 0.30137953 | -0.2371634 | 0.81253 | 0.95967622 |
| Acaa2 | 4196.82002 | 0.01019463 | 0.2233988 | 0.0456342 | 0.96360181 | 0.99401755 |
| Arhgap33 | 14.7530852 | 1.07371546 | 0.98344503 | 1.09179002 | 0.27492541 | 0.71147204 |
| C1qa | 14320.3283 | 1.33182053 | 0.26675742 | 4.99262779 | 5.956327806 | 4.670753721 |
| Gtdc1 | 119.105994 | -0.0217783 | 0.43176611 | -0.05044 | 0.95977174 | 0.99288691 |
| Ehmt1 | 1552.1937 | -0.0134004 | 0.25176635 | -0.0532257 | 0.95755208 | 0.99242502 |
| Rap2b | 2777.86572 | 0.18532917 | 0.18915386 | 0.97978001 | 0.32719472 | 0.74811039 |
| C1qc | 29036.4087 | 1.00022777 | 0.18732259 | 5.33960045 | 9.315164515 | 9.043841866 |
| Zfp157 | 392.839832 | 0.17359676 | 0.47032304 | 0.36910111 | 0.71205236 | 0.93235217 |
| C1qb | 64184.5674 | 0.96182779 | 0.244565 | 3.9328104 | 8.395846052 | 0.00311231 |
| Unc93b1 | 12831.3798 | -0.0779439 | 0.1961092 | -0.3974513 | 0.69103466 | 0.92470267 |
| Zfp280c | 330.320878 | -0.2172026 | 0.41788174 | -0.5197705 | 0.60322355 | 0.8917503 |
| Ttc7 | 589.880551 | -0.4374914 | 0.27946313 | -1.5654709 | 0.11747254 | 0.52372617 |
| Stag3 | 94.6041973 | 0.27968737 | 0.70333335 | 0.39765976 | 0.690881 | 0.92470267 |
| Nfkbid | 2296.77893 | 0.52504345 | 0.62308771 | 0.84264773 | 0.39942549 | 0.79612475 |
| Aifm1 | 2781.77582 | -0.484471 | 0.22409499 | -2.1618999 | 0.03062589 | 0.26095692 |
| 4921524J17F | 611.518972 | 0.00381486 | 0.29494476 | 0.01293415 | 0.98968033 | 0.99650865 |
| Kdm1a | 600.885598 | -0.1119646 | 0.29315388 | -0.3819312 | 0.70251239 | 0.92762537 |
| Elac1 | 408.166306 | -0.238509 | 0.23562715 | -1.0122304 | 0.31142792 | 0.74017637 |
| Rab8b | 2654.48332 | 0.63184847 | 0.24164587 | 2.61477047 | 0.00892874 | 0.1197646 |
| Tmem71 | 222.037954 | 0.81021435 | 0.50316327 | 1.61024144 | 0.10734516 | 0.50120355 |
| Map11 | 36.4594258 | -1.0684897 | 0.88866407 | -1.202355 | 0.22922603 | 0.6635423 |
| Slc39a12 | 9.67792172 | 1.01762093 | 2.17522984 | 0.46782225 | 0.63991171 | 0.90665276 |
| Kif1bp | 337.677254 | 0.68823661 | 0.47342235 | 1.4537476 | 0.1460163 | 0.5731306 |
| Bcorl1 | 36.1080806 | 0.40856984 | 0.93951121 | 0.43487489 | 0.66365324 | 0.91395288 |
| Trim17 | 11.368034 | -0.9928838 | 1.27900126 | -0.7762962 | 0.4375741 | 0.8185065 |
| Spryd3 | 528.063004 | -0.2746409 | 0.24433088 | -1.1240531 | 0.26099049 | 0.69691361 |
| Cnpy4 | 1577.73777 | -0.0504877 | 0.18743477 | -0.2693616 | 0.7876514 | 0.95225386 |
| Tmem177 | 785.953654 | 0.12474244 | 0.28616707 | 0.43590773 | 0.66290367 | 0.91395288 |
| Anapc10 | 1326.33675 | 0.29060253 | 0.38280602 | 0.75913782 | 0.44777012 | 0.82332359 |
| Taf6 | 3143.23575 | -0.1801595 | 0.16057037 | -1.1219974 | 0.26186353 | 0.69744756 |

| | | | | | | |
|---|---|---|---|---|---|---|
| Tfb1m | 675.737839 | 0.23871316 | 0.41428412 | 0.57620639 | 0.56447569 | 0.87718891 |
| Zdhhc9 | 261.800274 | 0.46576548 | 0.40672297 | 1.14516641 | 0.25214023 | 0.68542921 |
| Pml | 954.78885 | 0.44034454 | 0.37608279 | 1.17087129 | 0.24165052 | 0.67708722 |
| Trim3 | 59.0072151 | -0.3393865 | 0.6083581 | -0.5578728 | 0.57693122 | 0.88121027 |
| Otud4 | 494.200303 | -0.0047128 | 0.36975615 | -0.0127457 | 0.98983071 | 0.99650865 |
| Nxt1 | 2727.76208 | -0.0476764 | 0.33959495 | -0.140392 | 0.88835031 | 0.97832097 |
| Zfp39 | 231.212976 | 0.24395947 | 0.4693072 | 0.51982895 | 0.6031828 | 0.8917503 |
| Zfp113 | 161.163791 | 0.66864664 | 0.53371529 | 1.25281523 | 0.21027296 | 0.64378984 |
| Hk1 | 886.30217 | -0.5450905 | 0.30331885 | -1.7970873 | 0.07232176 | 0.41349974 |
| Ss18 | 4196.07348 | -0.1024422 | 0.20610176 | -0.4970468 | 0.61915603 | 0.89896588 |
| Zscan21 | 1279.16505 | 0.4335999 | 0.3080008 | 1.40778824 | 0.15919378 | 0.58838861 |
| Wdr62 | 463.731202 | 0.44054183 | 0.42688083 | 1.03200193 | 0.30207122 | 0.73361866 |
| Mmaa | 261.555979 | 0.46538275 | 0.50889086 | 0.91450405 | 0.36045205 | 0.76881241 |
| Zfp146 | 118.897854 | 0.68305378 | 0.42746968 | 1.59789994 | 0.11006525 | 0.50692212 |
| Tspan15 | 43.9149248 | 4.77519592 | 1.44875066 | 3.29607852 | 0.00098045 | 0.02304188 |
| Apbb1 | 48.5663761 | 1.49207997 | 1.10042676 | 1.35591029 | 0.17512771 | 0.60743838 |
| Smpd1 | 732.340906 | 0.24515319 | 0.24498965 | 1.00066755 | 0.31698756 | 0.74186126 |
| Paip2 | 4396.55575 | 0.33995895 | 0.29379384 | 1.1571344 | 0.24721746 | 0.68220418 |
| Cavin3 | 54.0365052 | 2.55645973 | 1.29377928 | 1.9759628 | NA | NA |
| Sh3glb1 | 5248.80504 | 0.56237111 | 0.21901997 | 2.56767051 | 0.01023844 | 0.13050277 |
| Rbmxl1 | 10665.0471 | 0.03443752 | 0.25799972 | 0.1334789 | 0.89381465 | 0.97941957 |
| Scd1 | 50.0705216 | 0.83168226 | 0.95452395 | 0.87130582 | 0.3835872 | 0.78679111 |
| Selenof | 18983.0259 | 0.67355803 | 0.38024415 | 1.77138304 | 0.07649703 | 0.42671597 |
| Rnf139 | 542.866404 | 0.17230624 | 0.26781037 | 0.64338897 | 0.51997174 | 0.8576517 |
| Trmt12 | 495.125156 | 0.23454826 | 0.52655695 | 0.4454376 | 0.65600354 | 0.91165371 |
| Slc35b2 | 5877.7299 | 0.4487778 | 0.32805947 | 1.36797699 | 0.17131928 | 0.60430173 |
| Gm9762 | 136.858338 | 0.38726046 | 0.37755185 | 1.02571463 | 0.30502611 | 0.73510332 |
| Rab11fip3 | 27.0711177 | 1.2432945 | 1.08178874 | 1.14929511 | 0.25043432 | 0.6842534 |
| Dcaf15 | 137.954695 | -0.4100222 | 0.63296601 | -0.6477792 | 0.5171278 | 0.85694744 |
| Socs5 | 179.467882 | 0.55018596 | 0.45418564 | 1.21136801 | 0.2257544 | 0.65900564 |
| Zcwpw1 | 94.9365204 | -0.0639015 | 0.68133531 | -0.0937886 | 0.92527707 | 0.98571293 |
| Ralgapa2 | 152.995258 | -0.407172 | 0.57710407 | -0.7055435 | 0.48047206 | 0.83935746 |
| Setd7 | 143.123125 | -0.7510912 | 0.74178824 | -1.0125413 | 0.3112793 | 0.74011268 |
| Sik2 | 191.358 | -0.0707368 | 0.384699 | -0.1838758 | 0.85411088 | 0.9684251 |
| Fam91a1 | 560.71156 | -0.2387105 | 0.38998782 | -0.6120974 | 0.54047334 | 0.86741192 |
| Psd | 23.5168988 | 1.74750123 | 1.86035786 | 0.93933607 | 0.34755823 | 0.76101142 |
| Prmt9 | 736.204989 | 0.04412187 | 0.25585269 | 0.17245028 | 0.86308354 | 0.9705375 |
| Arhgap10 | 385.761925 | 0.20706145 | 0.28523043 | 0.72594445 | 0.46787283 | 0.83381643 |
| Ddx1 | 9838.94616 | -0.2218113 | 0.1768945 | -1.2539184 | 0.20987167 | 0.64350283 |
| Lrrc20 | 230.710951 | 0.12665696 | 0.34496716 | 0.36715657 | 0.71350223 | 0.93235217 |
| Ndufc1 | 2290.42625 | 0.23989158 | 0.26993172 | 0.88871209 | 0.37415783 | 0.77748815 |
| Mgarp | 88.5492154 | 0.02116111 | 0.63536371 | 0.03330551 | 0.97343096 | 0.99443087 |
| Mycn | 52.2599798 | 0.22088991 | 1.30642662 | 0.16907946 | 0.86573414 | 0.9705375 |
| Nodal | 18.9260442 | -0.2028657 | 1.62754251 | -0.1246454 | 0.90080426 | 0.98043702 |
| E330009J07F | 69.1188862 | -0.8041522 | 0.68535043 | -1.1733446 | 0.24065765 | 0.67584138 |
| Elf2 | 1325.92048 | 0.51506705 | 0.24656029 | 2.08901054 | 0.03670677 | 0.28765789 |

| | | | | | | |
|---|---|---|---|---|---|---|
| Krt80 | 44.4831826 | -0.9042186 | 0.8070644 | -1.1203797 | 0.262552 | 0.69836891 |
| Cyb561d2 | 1404.12917 | 0.07420745 | 0.32190426 | 0.23052646 | 0.81768271 | 0.96079676 |
| Rbm17 | 4432.03231 | -0.4989701 | 0.23943234 | -2.083971 | 0.0371628 | 0.28956966 |
| Atg101 | 2434.13971 | 0.11641641 | 0.2463005 | 0.47266006 | 0.63645572 | 0.90542378 |
| Fam193a | 360.235132 | 0.19310479 | 0.29869795 | 0.64648848 | 0.51796306 | 0.8570838 |
| Spry1 | 42.0669509 | 1.10774254 | 1.26432025 | 0.87615661 | 0.3809449 | 0.7832891 |
| Thap1 | 526.460301 | -0.0633482 | 0.23571648 | -0.2687474 | 0.78812409 | 0.95225386 |
| Lipt1 | 72.4009113 | 0.2982781 | 0.56210329 | 0.53064642 | 0.59566382 | 0.88830374 |
| Mospd3 | 650.80305 | -0.3726234 | 0.33765922 | -1.1035487 | 0.26978897 | 0.7068598 |
| Zfyve28 | 9.03425976 | 0.97310979 | 1.63696792 | 0.59445868 | 0.55220537 | 0.87093412 |
| Hook3 | 506.140954 | -0.3902836 | 0.38793819 | -1.0060458 | 0.31439354 | 0.74065408 |
| Mxd4 | 465.099001 | 0.16929557 | 0.37404691 | 0.45260518 | 0.65083307 | 0.91021183 |
| Matr3 | 5643.10722 | -0.009314 | 0.18281933 | -0.0509463 | 0.95936827 | 0.99272137 |
| Clic4 | 5495.35254 | 0.46096937 | 0.28819386 | 1.59951142 | 0.10970701 | 0.50605049 |
| Zfp692 | 198.606542 | -0.4969238 | 0.36344884 | -1.3672454 | 0.1715484 | 0.60430173 |
| Pomk | 762.723678 | 0.71134152 | 0.24307956 | 2.92637324 | 0.00342939 | 0.05967533 |
| Mex3c | 338.130272 | -0.4906538 | 0.62710959 | -0.7824053 | 0.43397644 | 0.81498831 |
| Aagab | 5166.23067 | 0.10535863 | 0.20386477 | 0.51680646 | 0.60529127 | 0.89258167 |
| Hgsnat | 7891.52397 | 1.44765526 | 0.28416774 | 5.09436878 | 3.499050883 | 2.893961448 |
| Kin | 545.058239 | -0.4599501 | 0.3122856 | -1.4728509 | 0.14079125 | 0.56517498 |
| Rsrp1 | 5995.3885 | 0.24049841 | 0.25902183 | 0.92848701 | 0.353155 | 0.76376846 |
| 4932438A13I | 249.430551 | 0.17209358 | 0.32302846 | 0.53275053 | 0.59420628 | 0.88797037 |
| Gemin5 | 1442.53557 | -0.4742517 | 0.31946444 | -1.4845212 | 0.13767072 | 0.56012764 |
| Tmem97 | 5776.01222 | -0.323661 | 0.15389297 | -2.1031564 | 0.0354521 | 0.28231508 |
| Galnt6 | 367.9624 | 1.57708623 | 0.25826733 | 6.10641014 | 1.018969641 | 1.806529801 |
| Stag1 | 843.875452 | -0.2085522 | 0.24481823 | -0.8518657 | 0.39428865 | 0.79233901 |
| Tbcel | 62.2532528 | 0.57729174 | 0.74635346 | 0.77348304 | 0.43923656 | 0.81963526 |
| Ldlrap1 | 1344.23132 | -0.1255289 | 0.31319582 | -0.4008001 | 0.68856727 | 0.92470267 |
| Lsm1 | 1435.82088 | 0.23800171 | 0.34592295 | 0.68801943 | 0.49144055 | 0.84497726 |
| Ttc13 | 1860.8695 | -0.1081534 | 0.21921376 | -0.4933694 | 0.62175163 | 0.89912494 |
| Man1c1 | 508.343052 | 0.25092601 | 0.26340016 | 0.95264185 | 0.34077157 | 0.75821364 |
| Tacc3 | 11500.4728 | 0.01588093 | 0.17269249 | 0.09196076 | 0.92672922 | 0.98630274 |
| Jade3 | 223.629051 | 0.24560825 | 0.50493855 | 0.48641216 | 0.62667497 | 0.90200199 |
| Bag4 | 852.509299 | 0.24373983 | 0.33581423 | 0.72581746 | 0.46795068 | 0.83381643 |
| Traf3ip3 | 4604.32659 | -0.2462912 | 0.18482147 | -1.3325895 | 0.18266654 | 0.61405874 |
| Tap1 | 893.364531 | -0.1606446 | 0.25590781 | -0.6277441 | 0.53017161 | 0.86348696 |
| Bbs7 | 345.875259 | 0.25432152 | 0.36687128 | 0.69321731 | 0.48817319 | 0.84329777 |
| Capn15 | 425.462084 | -0.2298118 | 0.43706372 | -0.5258084 | 0.59902134 | 0.88987699 |
| Larp1 | 932.066828 | -0.5575637 | 0.37799231 | -1.4750661 | 0.14019477 | 0.56396008 |
| Hand1 | 10.7072778 | -0.0970355 | 1.24880771 | -0.0777025 | 0.93806472 | 0.98891293 |
| Map4k1 | 1919.17533 | -1.1748307 | 0.2652056 | -4.4298863 | 9.428276798 | 0.00049289 |
| Fam53a | 1428.70573 | -0.156672 | 0.27760891 | -0.5643624 | 0.57250752 | 0.88035336 |
| Slc9a7 | 15.103941 | 0.22318584 | 2.04517975 | 0.10912774 | 0.91310117 | 0.98292561 |
| Taf2 | 745.450265 | -0.3704595 | 0.33012789 | -1.1221696 | 0.26179032 | 0.6974044 |
| Slc12a9 | 1641.46549 | 0.26652772 | 0.23810026 | 1.1193928 | 0.2629726 | 0.69903169 |
| Chst7 | 94.4098838 | 0.57367216 | 0.51667584 | 1.11031351 | 0.26686395 | 0.7035729 |

| | | | | | | |
|---|---|---|---|---|---|---|
| Paqr7 | 1219.80481 | 0.14511896 | 0.28991446 | 0.50055784 | 0.61668234 | 0.89878056 |
| Nudt22 | 382.413202 | 0.5898028 | 0.38370107 | 1.53714141 | 0.12425868 | 0.53565448 |
| Actr1b | 3795.18082 | -0.0530394 | 0.15998535 | -0.3315267 | 0.74024667 | 0.93800224 |
| Letmd1 | 1145.66394 | -0.1099488 | 0.24854215 | -0.4423748 | 0.65821804 | 0.91258818 |
| Uvssa | 148.78688 | 0.74250794 | 0.48405205 | 1.53394235 | 0.12504383 | 0.53653498 |
| Sf3b6 | 4518.96976 | 0.26851087 | 0.42079774 | 0.63809958 | 0.52340886 | 0.85898317 |
| Letm2 | 34.1128242 | 0.58509932 | 0.88095835 | 0.66416229 | 0.50658644 | 0.85123041 |
| Srrt | 4196.17513 | -0.3846869 | 0.19927672 | -1.9304156 | 0.05355536 | 0.35852954 |
| Pafah2 | 654.25429 | 0.52449019 | 0.37552357 | 1.39669047 | 0.16250667 | 0.59359334 |
| Kdm6a | 347.552952 | 0.11615209 | 0.47619735 | 0.24391586 | 0.80729598 | 0.95787117 |
| Enpp1 | 225.783882 | 1.54442397 | 0.64784178 | 2.38395241 | 0.01712782 | 0.1837935 |
| Ctbp1 | 647.568754 | -0.2454972 | 0.38233945 | -0.6420922 | 0.52081331 | 0.8576517 |
| Hhat | 198.596708 | 0.50663963 | 0.3585388 | 1.41306779 | 0.15763579 | 0.58620126 |
| Trmt6 | 1983.70675 | -0.344247 | 0.22011852 | -1.5639165 | 0.11783719 | 0.52437333 |
| Rims2 | 14.1824229 | 1.6301818 | 1.94716221 | 0.83720904 | 0.40247508 | 0.79795242 |
| Rcor3 | 239.245078 | 0.01126911 | 0.32242851 | 0.03495072 | 0.97211903 | 0.99443087 |
| Atp11b | 401.158 | -0.673412 | 0.30570828 | -2.2027927 | 0.02760937 | 0.24636643 |
| Icam1 | 5461.88678 | 0.4172476 | 0.33320122 | 1.25223908 | 0.21048276 | 0.64401792 |
| Htra4 | 9.09676929 | 2.19294629 | 1.99999798 | 1.09647425 | 0.27287129 | 0.71005689 |
| Cnnm4 | 2558.63541 | -0.7627912 | 0.27603127 | -2.763423 | 0.00571986 | 0.08847273 |
| Tbc1d2b | 469.060123 | -0.1063825 | 0.57072294 | -0.1863995 | 0.85213147 | 0.96798445 |
| Serpine1 | 8.29363525 | 0.904509 | 1.69642017 | 0.5331869 | 0.59390421 | 0.88797037 |
| Ranbp10 | 176.234025 | 0.55979974 | 0.46786355 | 1.19650212 | 0.23150067 | 0.66689373 |
| Dmxl1 | 129.334486 | 0.22838635 | 0.54353337 | 0.42018827 | 0.67434792 | 0.91822108 |
| Best1 | 64.0538516 | 0.08344208 | 0.54703639 | 0.15253478 | 0.87876516 | 0.97535404 |
| Endod1 | 150.743925 | 0.39443812 | 0.48886727 | 0.80684091 | 0.41975815 | 0.80563709 |
| Depdc5 | 433.030998 | 0.00216663 | 0.35030262 | 0.00618502 | 0.9950651 | 0.99884675 |
| Fer1l5 | 19.9624571 | -0.3669922 | 0.81411579 | -0.4507863 | 0.65214358 | 0.91037383 |
| Slc30a1 | 75.4719766 | -1.4855614 | 0.92756392 | -1.6015731 | 0.10925004 | 0.50527626 |
| Cep85 | 3224.6289 | -0.5047997 | 0.195593 | -2.5808681 | 0.00985522 | 0.12811791 |
| Arid5a | 2388.09226 | -0.2218088 | 0.19073365 | -1.1629243 | 0.24486023 | 0.68039867 |
| Slc18b1 | 60.3134339 | 1.44108849 | 0.78801449 | 1.82875888 | 0.06743574 | 0.40006856 |
| Azin1 | 5984.35341 | -0.0364016 | 0.16860345 | -0.2159005 | 0.82906532 | 0.96462215 |
| Ints7 | 3200.87706 | -0.2108628 | 0.25917115 | -0.8136046 | 0.41587152 | 0.80440685 |
| Klf10 | 5465.96796 | 0.3137358 | 0.49876805 | 0.62902145 | 0.52933501 | 0.86280549 |
| Tedc1 | 650.177459 | 0.18411748 | 0.26006199 | 0.70797537 | 0.47896055 | 0.83869135 |
| Uggt1 | 1235.91596 | -0.3021308 | 0.30243222 | -0.9990034 | 0.31779303 | 0.74307991 |
| Dtl | 2428.04566 | -0.0276103 | 0.3359123 | -0.082195 | 0.93449164 | 0.98756396 |
| Thoc2 | 2221.96183 | -0.220521 | 0.20817754 | -1.0592931 | 0.28946633 | 0.72395811 |
| Asxl2 | 743.514647 | -0.1444175 | 0.53671709 | -0.2690756 | 0.78787149 | 0.95225386 |
| Ubr5 | 448.054021 | 0.19446968 | 0.38398762 | 0.50644778 | 0.61254234 | 0.8967485 |
| Cib2 | 53.5768877 | 1.14768214 | 0.74113431 | 1.54854812 | 0.12149039 | 0.52945917 |
| Nenf | 651.23894 | 0.73567452 | 0.40580103 | 1.81289467 | 0.06984807 | 0.40804748 |
| Fam168b | 1453.98577 | -0.2130459 | 0.30460411 | -0.6994191 | 0.48429018 | 0.84128398 |
| Arhgef4 | 41.6868423 | 0.42911653 | 0.79146875 | 0.54217747 | 0.58769625 | 0.88680008 |
| Pank2 | 136.811616 | 0.5627056 | 0.65575176 | 0.85810765 | 0.39083303 | 0.7899826 |

| | | | | | | |
|---|---|---|---|---|---|---|
| Ppfia1 | 262.030907 | -0.1595861 | 0.44790711 | -0.3562929 | 0.72162121 | 0.93398714 |
| Mavs | 1939.66113 | -0.1346181 | 0.2840967 | -0.4738461 | 0.63560964 | 0.90531997 |
| Bcdin3d | 207.488844 | 0.22597187 | 0.33926888 | 0.66605539 | 0.5053757 | 0.85037976 |
| Atg14 | 791.039294 | 0.26470184 | 0.37761556 | 0.70098236 | 0.48331402 | 0.84107543 |
| Mrpl47 | 1520.96841 | 0.02639514 | 0.29864701 | 0.08838239 | 0.92957276 | 0.98676358 |
| Rapgef6 | 754.931977 | 0.33743911 | 0.24643539 | 1.36928024 | 0.17091169 | 0.60430173 |
| Fbxo34 | 1478.68392 | -0.1991609 | 0.33401249 | -0.5962678 | 0.55099635 | 0.87093412 |
| Dlgap5 | 2457.55781 | 0.16244081 | 0.17405718 | 0.93326118 | 0.35068514 | 0.76238338 |
| H2-DMb2 | 87.8974073 | -1.5122257 | 0.87069217 | -1.7368086 | 0.08242095 | 0.44183004 |
| Plekhg2 | 177.605152 | 0.11775595 | 0.47634943 | 0.24720497 | 0.8047496 | 0.95725854 |
| Zdhhc18 | 84.8977438 | -1.0733694 | 0.70058994 | -1.5320937 | 0.12549932 | 0.53724874 |
| Rps16 | 16855.1803 | 0.1947762 | 0.36372421 | 0.53550518 | 0.59230058 | 0.88797037 |
| Vash2 | 44.3772331 | 1.30662031 | 1.42757718 | 0.91527122 | 0.36004926 | 0.76857232 |
| Mcrs1 | 3149.42593 | 0.04038999 | 0.15386793 | 0.26249776 | 0.79293771 | 0.95303665 |
| Wdhd1 | 2161.11107 | -0.0815122 | 0.25526016 | -0.3193298 | 0.74947645 | 0.9399191 |
| Tob1 | 216.087466 | -0.1768823 | 0.39855105 | -0.4438135 | 0.65717745 | 0.91251438 |
| Gch1 | 372.12712 | -0.1419351 | 0.32181024 | -0.4410522 | 0.65917524 | 0.91280998 |
| Nr0b2 | 69.7288455 | -0.4184164 | 0.55968955 | -0.7475865 | 0.45470958 | 0.82760373 |
| Clba1 | 43.6045255 | -0.2526183 | 0.59907848 | -0.4216782 | 0.67325991 | 0.91786342 |
| Nme1 | 15216.3306 | 0.00573052 | 0.31045984 | 0.01845818 | 0.98527334 | 0.99564261 |
| Adgrl3 | 10.3036052 | 1.70099068 | 2.09234949 | 0.81295725 | 0.41624257 | 0.80440685 |
| Bclaf1 | 5563.25465 | -0.3486869 | 0.21611073 | -1.6134642 | 0.1066437 | 0.49890292 |
| Tnfrsf23 | 68.6196104 | 0.26120228 | 0.71481784 | 0.36541097 | 0.71480466 | 0.93235217 |
| Spag1 | 37.0284027 | 1.28366927 | 0.86396112 | 1.48579517 | 0.13733331 | 0.55981285 |
| Wdtc1 | 160.834452 | 0.07441823 | 0.38555227 | 0.19301724 | 0.84694547 | 0.96765585 |
| Cdkn3 | 1554.09018 | 0.34428344 | 0.26748576 | 1.28710943 | 0.19805615 | 0.63077867 |
| Slc25a43 | 69.8837746 | 0.84270478 | 0.63446561 | 1.3282119 | 0.18410812 | 0.61619551 |
| Zbtb42 | 14.4854148 | 0.26994558 | 2.03152093 | 0.13287856 | 0.89428942 | 0.97941957 |
| Zfp60 | 179.778957 | -0.1672111 | 0.4280086 | -0.3906722 | 0.69603957 | 0.92623267 |
| Prkci | 350.038296 | 0.42587541 | 0.3384217 | 1.25841638 | 0.20824121 | 0.6418276 |
| Vps13b | 454.805085 | -0.4389692 | 0.44699426 | -0.9820466 | 0.32607691 | 0.74759915 |
| H2-DMa | 343.500271 | -1.7346676 | 0.34275087 | -5.0610158 | 4.170286309 | 3.378484266 |
| Phc3 | 530.771673 | -0.1184469 | 0.28017352 | -0.4227627 | 0.67246837 | 0.91770477 |
| Slc20a2 | 395.780935 | 0.33158665 | 0.2724578 | 1.21702024 | 0.22359652 | 0.65735549 |
| Gpr160 | 1567.66415 | -0.544773 | 0.20643421 | -2.6389665 | 0.00831592 | 0.1145593 |
| Cdkn1c | 61.4471266 | 2.09101529 | 0.98097585 | 2.13156653 | 0.0330425 | 0.26999733 |
| Ldah | 2737.01259 | -0.2434176 | 0.17299981 | -1.4070397 | 0.15941561 | 0.5886662 |
| Rfx7 | 351.609675 | -0.7535511 | 0.60833746 | -1.2387058 | 0.21545448 | 0.64854028 |
| Inf2 | 137.196374 | -0.7337397 | 0.76611956 | -0.9577352 | 0.33819628 | 0.75644564 |
| Armc3 | 7.81083549 | -0.2062096 | 1.67195162 | -0.1233347 | 0.9018421 | 0.98043702 |
| Atp8a1 | 959.144844 | -0.0197995 | 0.254199 | -0.0778898 | 0.93791571 | 0.98884106 |
| Ahdc1 | 17.46897 | -1.2161665 | 1.40702615 | -0.8643524 | 0.38739431 | 0.78911204 |
| Ddhd1 | 58.9213794 | -0.5333983 | 0.71756875 | -0.743341 | 0.45727524 | 0.82945552 |
| Cd81 | 10942.3694 | 0.41708703 | 0.1506786 | 2.76805743 | 0.00563915 | 0.08754282 |
| Fam13a | 96.0225044 | 1.81368362 | 0.79602048 | 2.27843839 | 0.02270047 | 0.21610495 |
| Cisd1 | 2381.19981 | -0.4845952 | 0.17835077 | -2.7170909 | 0.00658585 | 0.09836961 |

| | | | | | | |
|---|---|---|---|---|---|---|
| Fermt2 | 105.712157 | 2.36836331 | 1.33675915 | 1.77172028 | 0.076441 | 0.42659798 |
| Tmem33 | 4627.07801 | 0.01107345 | 0.20226274 | 0.05474783 | 0.95633936 | 0.99215259 |
| Gnpnat1 | 1924.03715 | -0.1565402 | 0.35865585 | -0.4364636 | 0.66250041 | 0.91395288 |
| Ckap2 | 1763.88733 | -0.0732095 | 0.21969458 | -0.3332331 | 0.73895833 | 0.93800224 |
| Mynn | 1171.05919 | 0.47656283 | 0.24887806 | 1.91484468 | 0.05551231 | 0.3627292 |
| Themis2 | 5846.59715 | 0.17921449 | 0.22707814 | 0.78921948 | 0.42998374 | 0.81197362 |
| Limch1 | 31.549429 | 1.69343772 | 1.960918 | 0.86359436 | 0.38781076 | 0.7893659 |
| Mrps26 | 2847.3917 | -0.3636017 | 0.22256921 | -1.6336569 | 0.1023309 | 0.4886081 |
| Eef1a1 | 225709.983 | 0.1578773 | 0.34100927 | 0.46297069 | 0.64338538 | 0.90810354 |
| Fam222b | 554.471987 | -0.1519686 | 0.34516855 | -0.4402735 | 0.65973903 | 0.91280998 |
| Xkr8 | 483.223497 | -0.4847711 | 0.34044643 | -1.4239277 | 0.1544674 | 0.58343244 |
| Ppp1r16b | 72.1277709 | 0.07882527 | 0.79450875 | 0.09921259 | 0.92096948 | 0.9839023 |
| Ptger2 | 97.5538512 | -2.1431861 | 0.57080916 | -3.7546456 | 0.00017359 | 0.00581778 |
| Actr5 | 62.8330319 | -0.1673645 | 0.62111381 | -0.2694587 | 0.78757674 | 0.95225386 |
| Mrpl23 | 572.403046 | 0.3500517 | 0.38509513 | 0.90900058 | 0.36334982 | 0.77000799 |
| Pced1a | 401.398423 | 0.0299681 | 0.2288565 | 0.13094713 | 0.89581713 | 0.97984623 |
| Dzip1l | 14.4112028 | -0.8377238 | 2.06070068 | -0.4065238 | 0.6843578 | 0.92286709 |
| Apopt1 | 1416.60275 | 0.12351006 | 0.34418429 | 0.35884863 | 0.71970833 | 0.93314808 |
| Vopp1 | 140.555865 | -0.3913501 | 0.48772916 | -0.8023922 | 0.4223261 | 0.80821275 |
| Phf12 | 251.485278 | -0.5589117 | 0.38527765 | -1.4506724 | 0.14687111 | 0.57365079 |
| N4bp2 | 397.832583 | 0.13201261 | 0.29189952 | 0.45225362 | 0.65108629 | 0.91021183 |
| Rpl10a | 15787.0275 | 0.22884634 | 0.38480222 | 0.59471159 | 0.55203628 | 0.87093412 |
| Fam76b | 355.274448 | 0.18299182 | 0.35328853 | 0.51796707 | 0.60448125 | 0.89258167 |
| Ctnna1 | 2984.40614 | 0.3689141 | 0.29857262 | 1.23559253 | 0.21661006 | 0.65009949 |
| Fbxw17 | 692.739919 | 0.10159591 | 0.37832374 | 0.26854224 | 0.78828196 | 0.95225386 |
| Abhd18 | 380.171633 | -0.1049925 | 0.3971202 | -0.2643848 | 0.79148342 | 0.95268772 |
| Tgm2 | 995.309525 | 1.61925782 | 0.31564133 | 5.13005636 | 2.896554461 | 2.477870679 |
| Smim14 | 927.040598 | 0.50686713 | 0.25505272 | 1.9873034 | 0.04688879 | 0.3345146 |
| Tspan14 | 5201.90161 | -0.0128938 | 0.19605396 | -0.0657667 | 0.94756355 | 0.99019648 |
| Ppm1k | 294.32348 | 0.12601157 | 0.43900727 | 0.28703754 | 0.77408358 | 0.94684176 |
| Fdxacb1 | 502.613576 | 0.57454238 | 0.41991391 | 1.36823848 | 0.17123744 | 0.60430173 |
| Rtkn2 | 552.526204 | 0.67139104 | 0.24950178 | 2.6909269 | 0.00712538 | 0.10299472 |
| Nmrk1 | 565.113899 | 1.10992968 | 0.36755155 | 3.01979319 | 0.00252947 | 0.04723614 |
| Iars | 8313.55027 | -0.4221507 | 0.26524464 | -1.5915521 | 0.11148537 | 0.50888686 |
| Cpe | 35.7847441 | 3.25421233 | 1.69572568 | 1.91906767 | 0.05497577 | 0.36084578 |
| Nufip2 | 206.06032 | 0.14436257 | 0.48827841 | 0.29565626 | 0.7674926 | 0.94401697 |
| Aim2 | 631.782566 | 0.10972498 | 0.29377164 | 0.37350433 | 0.70877312 | 0.92997777 |
| Egr2 | 1303.80519 | 0.52243819 | 0.41149665 | 1.26960497 | 0.20422538 | 0.63846817 |
| Jmjd1c | 992.739863 | 0.49814947 | 0.35238412 | 1.4136547 | 0.15746331 | 0.58620126 |
| Stk35 | 267.163531 | -0.3757157 | 0.34065041 | -1.1029362 | 0.27005487 | 0.70725351 |
| Dusp8 | 310.075078 | 1.0012776 | 0.71031405 | 1.40962662 | 0.15864996 | 0.58757644 |
| Wdr19 | 86.5377475 | 0.66714772 | 0.85103654 | 0.78392371 | 0.43308488 | 0.81435873 |
| Pcdh18 | 15.633942 | 1.51981872 | 2.52660492 | 0.60152607 | 0.54748965 | 0.87008334 |
| H2afz | 65412.7055 | 0.02153536 | 0.34314607 | 0.06275859 | 0.94995874 | 0.99065364 |
| Rcor1 | 422.350829 | -0.3036508 | 0.25864303 | -1.1740152 | 0.24038894 | 0.67564175 |
| Sirpa | 20952.036 | 0.28712113 | 0.22746367 | 1.26227248 | 0.20685076 | 0.64174209 |

| | | | | | | |
|---|---|---|---|---|---|---|
| Ankrd9 | 59.6015843 | 0.31254641 | 0.57952699 | 0.53931295 | 0.58967094 | 0.88722166 |
| Bri3bp | 2747.72136 | -1.0855576 | 0.22687477 | -4.784832 | 1.711304499 | 0.0001144 |
| Ankrd13b | 35.4715787 | -1.2681987 | 0.8835622 | -1.4353248 | 0.15119457 | 0.5792556 |
| Tmem156 | 388.049454 | -0.1686056 | 0.40260009 | -0.4187916 | 0.67536841 | 0.91848602 |
| Ndufv1 | 5705.88894 | 0.00464801 | 0.18494014 | 0.02513252 | 0.97994926 | 0.99486654 |
| Ddx60 | 74.1536004 | 1.43394866 | 1.10686883 | 1.29550008 | 0.19514777 | 0.62707955 |
| Bank1 | 1861.69989 | 0.48547983 | 0.21170928 | 2.29314388 | 0.02183972 | 0.21288074 |
| Ssh2 | 2272.07978 | 0.64700534 | 0.30391984 | 2.12886838 | 0.03326515 | 0.2712884 |
| Bicd2 | 806.454952 | -0.2776503 | 0.43426333 | -0.6393593 | 0.52258925 | 0.85816354 |
| Smarce1 | 4438.18671 | -0.1475707 | 0.22059305 | -0.6689726 | 0.50351297 | 0.85012672 |
| Scarb1 | 7842.91195 | -0.0311645 | 0.17157282 | -0.1816402 | 0.85586512 | 0.96915653 |
| Chchd5 | 720.489033 | 0.08056207 | 0.26335401 | 0.30590789 | 0.75967478 | 0.94210735 |
| Inpp4b | 466.316018 | 0.66921669 | 0.35255312 | 1.89820102 | 0.05766961 | 0.36897087 |
| Fgd3 | 209.532075 | -0.3394925 | 0.45816873 | -0.7409771 | 0.45870734 | 0.82986925 |
| Ano10 | 642.829239 | 0.3868394 | 0.34664294 | 1.11595926 | 0.26443957 | 0.70110302 |
| Wdr20 | 758.938816 | 0.33140068 | 0.27183313 | 1.2191328 | 0.22279379 | 0.65613364 |
| Nsrp1 | 544.753775 | -0.2497232 | 0.39165514 | -0.63761 | 0.52372758 | 0.85927568 |
| Card19 | 423.299052 | 0.10776168 | 0.22586572 | 0.47710505 | 0.63328732 | 0.90434445 |
| Zc3h7a | 1397.96659 | 0.08683348 | 0.26577186 | 0.32672187 | 0.74387826 | 0.93880767 |
| Ninj1 | 2862.8889 | 0.23696378 | 0.20243757 | 1.17055238 | 0.24177875 | 0.67708722 |
| 1110032A03I | 258.942917 | 0.48562554 | 0.3369502 | 1.4412383 | 0.14951737 | 0.57685486 |
| Snn | 33.1575398 | 1.71600375 | 1.0842349 | 1.58268633 | 0.11349298 | 0.51325677 |
| Rmi2 | 261.006143 | -0.5324662 | 0.4570784 | -1.1649341 | 0.24404567 | 0.67911981 |
| Rara | 198.053666 | 0.21113266 | 0.34478878 | 0.61235363 | 0.54030381 | 0.86741192 |
| Dhx38 | 3155.68682 | -0.4817301 | 0.33698006 | -1.4295508 | 0.15284599 | 0.58121386 |
| Igsf9 | 46.3129889 | 1.60801369 | 0.80972329 | 1.9858805 | 0.04704659 | 0.3347601 |
| Parp11 | 371.91211 | -0.0778729 | 0.40817125 | -0.190785 | 0.84869406 | 0.96765585 |
| Arap2 | 59.1867157 | -0.3650698 | 0.51148568 | -0.713744 | 0.47538547 | 0.83626553 |
| Acd | 1172.1135 | -0.1526267 | 0.20581065 | -0.7415878 | 0.45833714 | 0.82986925 |
| Cramp1l | 148.210782 | -0.4786572 | 0.76632946 | -0.6246102 | 0.53222693 | 0.86442675 |
| Hpf1 | 3041.6532 | -0.0681581 | 0.26091424 | -0.2612281 | 0.79391657 | 0.95310033 |
| Acer2 | 70.4811431 | 1.02963517 | 0.54235377 | 1.89845672 | 0.05763594 | 0.36897087 |
| Ccdc138 | 195.191164 | 0.46984451 | 0.36935431 | 1.27206993 | 0.20334827 | 0.63809363 |
| Wipf2 | 244.482956 | -0.2662886 | 0.61147856 | -0.4354831 | 0.66321181 | 0.91395288 |
| Fam120a | 2568.71589 | -0.2412034 | 0.21863876 | -1.103205 | 0.26993814 | 0.7070992 |
| Mindy4 | 122.449411 | 0.85097343 | 0.51151474 | 1.66363422 | 0.09618555 | 0.47560138 |
| Atp6v0a2 | 2251.85145 | -0.1861739 | 0.26480435 | -0.7030619 | 0.48201716 | 0.84038934 |
| Dennd4c | 151.333137 | 0.7596914 | 0.81227395 | 0.935265 | 0.34965175 | 0.76203275 |
| Phf2 | 73.2463068 | 0.13764638 | 0.56155686 | 0.24511566 | 0.80636688 | 0.95741157 |
| Tigar | 512.206496 | -0.447298 | 0.37324786 | -1.198394 | 0.23076368 | 0.6653874 |
| Igsf8 | 1146.9881 | -0.3326476 | 0.27586119 | -1.2058514 | 0.22787481 | 0.66182159 |
| Socs1 | 29.5513024 | -0.4078296 | 0.68770594 | -0.5930291 | 0.55316167 | 0.87136821 |
| Gcc2 | 444.095396 | -0.2918284 | 0.43107776 | -0.6769739 | 0.49842251 | 0.84870979 |
| Sult6b1 | 10.8355728 | 2.54905893 | 1.91649529 | 1.33006272 | 0.18349761 | 0.61524496 |
| Mrm3 | 889.725103 | -0.7553486 | 0.31866284 | -2.3703692 | 0.01777033 | 0.18756206 |
| Haus6 | 1049.72649 | -0.0159153 | 0.37344737 | -0.0426173 | 0.96600661 | 0.99443087 |

| | | | | | | |
|---|---|---|---|---|---|---|
| Dexi | 13.1023475 | -0.5452652 | 1.56694003 | -0.3479809 | 0.72785453 | 0.93576253 |
| Kmt2c | 430.514882 | 0.14685088 | 0.46720655 | 0.31431683 | 0.75328042 | 0.94018119 |
| Nod1 | 183.923445 | 0.679456 | 0.49336801 | 1.37717886 | 0.16845695 | 0.60312169 |
| Smim3 | 817.412068 | -0.9016342 | 0.35121335 | -2.567198 | 0.0102524 | 0.13050744 |
| Mturn | 13.9272029 | 3.03064163 | 2.01756107 | 1.5021313 | 0.1330632 | 0.55290835 |
| Csf3 | 27.7059384 | 2.48472246 | 1.83373976 | 1.35500277 | 0.17541667 | 0.60763832 |
| Rnf144b | 270.121662 | -0.5741343 | 0.43054912 | -1.3334932 | 0.18237001 | 0.61363505 |
| Cdkn2aip | 887.328619 | -0.4570227 | 0.35948859 | -1.2713134 | 0.20361718 | 0.63809363 |
| Cntln | 59.4716816 | 1.25299243 | 0.65421153 | 1.91527107 | 0.05545794 | 0.3627292 |
| Galnt11 | 763.040874 | 0.15736286 | 0.19793915 | 0.79500627 | 0.42660985 | 0.81021713 |
| Fkbp14 | 164.852928 | 2.43460108 | 0.46199831 | 5.26971853 | 1.366331165 | 1.265524296 |
| Tmem237 | 411.794929 | -0.3413109 | 0.25280915 | -1.3500735 | 0.1769924 | 0.60784615 |
| Kdm1b | 875.956499 | -0.1669571 | 0.27071729 | -0.6167212 | 0.53741867 | 0.86634214 |
| Opa1 | 1998.99197 | -0.3204863 | 0.29075429 | -1.1022581 | 0.27034947 | 0.70733226 |
| Cnbd2 | 146.216945 | 0.25419822 | 0.3388583 | 0.75016082 | 0.45315785 | 0.82610332 |
| Sbno1 | 2767.3608 | -0.0774155 | 0.31537575 | -0.2454706 | 0.80609205 | 0.95741157 |
| Trappc11 | 1125.70768 | 0.12071144 | 0.34084758 | 0.35415078 | 0.72322589 | 0.93433544 |
| Phf20 | 1645.73347 | 0.7343777 | 0.31575764 | 2.32576383 | 0.02003115 | 0.20003355 |
| Fam210a | 1595.97383 | -0.0417558 | 0.32790703 | -0.1273405 | 0.89867094 | 0.97989497 |
| Tbc1d32 | 69.0386832 | 0.4303493 | 0.74450808 | 0.57803175 | 0.56324268 | 0.8760902 |
| Mphosph9 | 266.818535 | -0.3258789 | 0.35704282 | -0.9127165 | 0.36139166 | 0.76887042 |
| Ccdc50 | 5031.06389 | 0.29785889 | 0.18552037 | 1.60553198 | 0.10837678 | 0.50301538 |
| Tmem181a | 1162.51179 | 0.33280569 | 0.22206643 | 1.4986763 | 0.13395763 | 0.55441542 |
| Stox2 | 12.7313749 | 2.32691688 | 1.70750197 | 1.36276088 | 0.17295788 | 0.60537734 |
| Snrk | 823.413825 | -0.1069757 | 0.41522978 | -0.2576302 | 0.79669232 | 0.95380086 |
| Cd84 | 3087.78121 | 0.66017698 | 0.35973025 | 1.83520005 | 0.06647605 | 0.39699582 |
| Ormdl3 | 376.442183 | -0.1914925 | 0.46641398 | -0.4105635 | 0.68139265 | 0.92061965 |
| Prdm1 | 256.884762 | -0.2123309 | 0.525348 | -0.4041719 | 0.68608632 | 0.92361549 |
| Gstp2 | 380.288027 | 2.33185883 | 0.40549442 | 5.75065581 | 8.889793187 | 1.316607998 |
| Spon1 | 10.5625855 | 1.73989613 | 2.59127706 | 0.67144349 | 0.50193804 | 0.85006796 |
| Atg5 | 3026.81039 | 0.51285728 | 0.30615393 | 1.6751615 | 0.09390255 | 0.46847874 |
| Pde4dip | 161.041256 | 1.250521 | 0.38183645 | 3.27501731 | 0.00105655 | 0.02426677 |
| Ttc39b | 543.834564 | 0.53598978 | 0.29402037 | 1.82296818 | 0.06830821 | 0.40309423 |
| Enpp6 | 59.3539119 | 0.71439061 | 1.27381906 | 0.56082582 | 0.57491629 | 0.88059206 |
| Fam126b | 393.898197 | 0.35012136 | 0.31807654 | 1.10074564 | 0.27100737 | 0.70830941 |
| Mylip | 597.998464 | 0.57522286 | 0.33610616 | 1.71143205 | 0.08700138 | 0.45109601 |
| Slc43a2 | 2201.9133 | 0.46693849 | 0.30952217 | 1.5085785 | 0.13140653 | 0.54970237 |
| Spag4 | 18.9935052 | -2.3787036 | 1.33339302 | -1.7839478 | 0.07443214 | 0.42056738 |
| Chpf2 | 634.118727 | 0.16097012 | 0.23787777 | 0.67669258 | 0.49860103 | 0.84870979 |
| Btbd10 | 1192.06058 | -0.4263061 | 0.23293691 | -1.8301352 | 0.06722972 | 0.39923356 |
| Scarf1 | 57.4274626 | 2.80141061 | 1.61707075 | 1.7323983 | 0.08320267 | 0.44330064 |
| Rilp | 20.6862635 | 1.88149176 | 1.42881375 | 1.31682087 | 0.18789868 | 0.61935434 |
| Asb10 | 44.9428533 | 0.22663918 | 0.85572618 | 0.26485012 | 0.79112493 | 0.95268772 |
| Prkab2 | 180.873143 | 0.39129937 | 0.49510584 | 0.79033479 | 0.42933227 | 0.81155572 |
| Fbxo8 | 620.763707 | 0.23623584 | 0.28125913 | 0.83992239 | 0.4009519 | 0.79698916 |
| Pgap3 | 29.8257409 | -0.3334181 | 0.84095559 | -0.3964753 | 0.69175444 | 0.92504756 |

| | | | | | | |
|---|---|---|---|---|---|---|
| Mfsd14b | 3504.74927 | 0.05671475 | 0.18240448 | 0.3109285 | 0.75585498 | 0.94102069 |
| Tapbpl | 515.107356 | 0.04743265 | 0.37579973 | 0.1262179 | 0.89955944 | 0.98033948 |
| Bend3 | 189.867931 | -0.5353206 | 0.39080111 | -1.3698032 | 0.17074833 | 0.60430173 |
| Cep44 | 551.551945 | 0.15066181 | 0.36450826 | 0.41332894 | 0.67936563 | 0.92044299 |
| Tlcd2 | 485.573586 | 0.34942542 | 0.28122294 | 1.24252102 | 0.21404439 | 0.64772533 |
| Primpol | 560.744743 | 0.27724697 | 0.2930994 | 0.94591447 | 0.34419221 | 0.75962035 |
| Hoxa9 | 17.8629097 | 3.45269523 | 1.76086451 | 1.96079551 | 0.04990288 | 0.34225988 |
| F11r | 694.263529 | -0.3924793 | 0.37394303 | -1.0495698 | 0.29391594 | 0.72683372 |
| Pdss2 | 523.93038 | 0.05800582 | 0.36524409 | 0.15881385 | 0.87381553 | 0.97350615 |
| Cep250 | 555.867224 | 0.08167326 | 0.387511 | 0.21076371 | 0.83307166 | 0.96565375 |
| Mical2 | 37.5411681 | 0.01390879 | 0.96095992 | 0.01447385 | 0.98845194 | 0.99635239 |
| Usp38 | 1206.07236 | -0.0075132 | 0.28894972 | -0.0260019 | 0.97925582 | 0.99483019 |
| Ncapd2 | 3852.77527 | -0.2075859 | 0.28050625 | -0.7400402 | 0.45927562 | 0.82987212 |
| Bcl9 | 171.834941 | 0.56621775 | 0.62063142 | 0.9123253 | 0.3615975 | 0.76887042 |
| Trpm4 | 74.5440603 | 0.53684761 | 0.73513742 | 0.73026838 | 0.46522615 | 0.83339036 |
| Slc22a23 | 16.2699246 | -0.0240774 | 1.29347138 | -0.0186145 | 0.98514861 | 0.99564261 |
| Ovca2 | 939.063763 | 0.41333051 | 0.39792145 | 1.03872387 | 0.29893318 | 0.73093388 |
| Iffo1 | 299.987092 | -0.1946634 | 0.52107309 | -0.3735818 | 0.70871544 | 0.92997777 |
| Fau | 19992.1408 | 0.42873248 | 0.44471098 | 0.96406992 | 0.33501087 | 0.75359284 |
| Nop2 | 6561.22457 | -0.5776898 | 0.22737809 | -2.5406574 | 0.01106443 | 0.13658035 |
| Ostm1 | 2571.75292 | 0.13163197 | 0.17913572 | 0.73481698 | 0.46245097 | 0.83108016 |
| Bphl | 1603.065 | -0.2029876 | 0.23544845 | -0.8621319 | 0.38861497 | 0.78942657 |
| Smg6 | 329.88638 | -0.2398866 | 0.36420133 | -0.6586649 | 0.51011096 | 0.85335831 |
| Snx25 | 80.8153763 | -0.6064993 | 0.78505103 | -0.7725604 | 0.43978262 | 0.81963526 |
| Atg9b | 84.5528417 | -0.5230541 | 0.88515622 | -0.5909173 | 0.55457586 | 0.87165957 |
| Galnt18 | 11.4082911 | -0.9448107 | 1.6572671 | -0.5701016 | 0.56860878 | 0.87936678 |
| Wdr36 | 4208.90342 | -0.0862177 | 0.1470004 | -0.5865134 | 0.55753057 | 0.87283439 |
| Snx10 | 1053.71046 | -0.6060697 | 0.2115438 | -2.8649843 | 0.0041703 | 0.06959792 |
| Afg1l | 225.378375 | 0.6067054 | 0.35161725 | 1.72547111 | 0.08444261 | 0.44526833 |
| Spats2l | 43.5129836 | 3.1918625 | 1.21562488 | 2.62569692 | 0.00864718 | 0.11762411 |
| Edem2 | 2796.24837 | 0.04920252 | 0.18419878 | 0.26711644 | 0.78937951 | 0.95255415 |
| 1700066M21 | 540.963798 | -0.295704 | 0.33731944 | -0.876629 | 0.38068819 | 0.78323054 |
| Trpc4ap | 3111.49697 | -0.073473 | 0.37790637 | -0.1944211 | 0.8458462 | 0.96765585 |
| Sesn1 | 8292.11136 | 0.99141418 | 0.18157669 | 5.46003009 | 4.760539035 | 5.153599471 |
| Tsr1 | 4129.75803 | -0.4790167 | 0.25089348 | -1.9092433 | 0.05623071 | 0.36472442 |
| Mlxip | 1343.65166 | -0.2476772 | 0.4518602 | -0.548128 | 0.58360403 | 0.88400546 |
| Txlng | 298.625923 | -0.1599748 | 0.39020596 | -0.4099754 | 0.681824 | 0.92061965 |
| Zfp384 | 96.3856246 | 0.15884489 | 0.64295865 | 0.24705304 | 0.80486718 | 0.95725854 |
| Sgsm2 | 26.824151 | -0.6997906 | 1.32635893 | -0.5276027 | 0.5977751 | 0.88928406 |
| Arl5c | 416.442506 | 0.41586152 | 0.62030174 | 0.67041811 | 0.50259129 | 0.85012672 |
| Fbxo25 | 262.861979 | 0.4809183 | 0.30225623 | 1.59109477 | 0.11158825 | 0.50906326 |
| Lasp1 | 4716.704 | 0.01393411 | 0.31195728 | 0.04466672 | 0.96437296 | 0.99406401 |
| Focad | 319.439045 | 0.42501478 | 0.44735768 | 0.95005585 | 0.34208388 | 0.75878732 |
| Ncoa6 | 225.214783 | -0.4206661 | 0.54947967 | -0.7655717 | 0.44393117 | 0.82068545 |
| Pcp4l1 | 19.2012093 | 0.16118921 | 1.19208265 | 0.13521647 | 0.89244072 | 0.97941957 |
| Sbf2 | 416.193148 | 0.92026928 | 0.45397624 | 2.02713095 | 0.04264902 | 0.31638902 |

| | | | | | | |
|---|---|---|---|---|---|---|
| Gmds | 1270.36616 | -0.1723004 | 0.18571369 | -0.9277745 | 0.35352455 | 0.76383312 |
| Rbm8a | 5684.63929 | -0.3250569 | 0.18440048 | -1.7627767 | 0.07793817 | 0.43160596 |
| Trp53inp2 | 286.727989 | 0.59562399 | 0.41105986 | 1.44899577 | 0.14733876 | 0.57410946 |
| Ttk | 1257.0322 | -0.2094283 | 0.26467568 | -0.7912639 | 0.42878999 | 0.81136783 |
| Pigu | 2327.41135 | -0.0721914 | 0.19790225 | -0.3647833 | 0.7152732 | 0.93235217 |
| Setd1b | 152.308865 | 0.10079426 | 0.38271573 | 0.26336587 | 0.79226859 | 0.95297175 |
| Rras | 19.137309 | 2.63896112 | 1.75000447 | 1.50797393 | 0.1315612 | 0.54984222 |
| Mpp6 | 903.540987 | -0.1673854 | 0.18925867 | -0.8844268 | 0.37646588 | 0.77905031 |
| Txnip | 5341.50991 | 0.17570176 | 0.2736574 | 0.6420501 | 0.52084066 | 0.8576517 |
| Upf3a | 777.200159 | -0.0530688 | 0.27651251 | -0.1919218 | 0.84780348 | 0.96765585 |
| Pmepa1 | 17.3892977 | 0.36084599 | 0.88221214 | 0.40902406 | 0.68252201 | 0.92104707 |
| Scaf1 | 297.133991 | -0.3079885 | 0.46018005 | -0.6692782 | 0.50331803 | 0.85012672 |
| Higd1a | 2333.94464 | -0.1216365 | 0.30892525 | -0.3937409 | 0.69377237 | 0.92609254 |
| Cdc16 | 8186.54576 | 0.31247053 | 0.25417984 | 1.22932855 | 0.21894865 | 0.65247231 |
| Fig4 | 2799.8687 | 0.05034158 | 0.1770334 | 0.28436208 | 0.77613292 | 0.94776024 |
| Egr1 | 9579.95212 | 0.69059935 | 0.37902827 | 1.82202597 | 0.06845105 | 0.40335339 |
| Hdhd3 | 105.218728 | 1.24064579 | 0.61906069 | 2.00407779 | 0.04506173 | 0.32675763 |
| Poli | 286.634553 | -0.0761472 | 0.47355851 | -0.1607979 | 0.87225259 | 0.97312047 |
| Usp5 | 6362.74007 | 0.08145488 | 0.15606283 | 0.52193642 | 0.6017146 | 0.89103464 |
| Mllt6 | 49.2098319 | 0.3376527 | 0.71056205 | 0.47519102 | 0.63465083 | 0.9050692 |
| Cdc40 | 1919.85933 | -0.295899 | 0.17382234 | -1.7023072 | 0.0886978 | 0.45521604 |
| Spsb2 | 363.276058 | 0.57447191 | 0.28613344 | 2.00770629 | 0.04467451 | 0.32480971 |
| Abhd17c | 193.47907 | -0.2197285 | 0.35053499 | -0.6268377 | 0.53076562 | 0.86375626 |
| Uqcrfs1 | 8882.92707 | -0.0142853 | 0.27217576 | -0.0524855 | 0.95814184 | 0.99246834 |
| Olfml2b | 70.9880213 | -0.2696041 | 0.63022726 | -0.4277886 | 0.66880502 | 0.91618049 |
| Chmp4b | 51.5813027 | -0.4186905 | 0.58810001 | -0.7119376 | 0.47650342 | 0.83654799 |
| Nos1ap | 24.4122796 | -0.3777309 | 1.61386784 | -0.2340532 | 0.8149437 | 0.95996209 |
| Cdk19 | 334.854313 | 0.05316778 | 0.28341434 | 0.18759735 | 0.8511923 | 0.9679302 |
| Tfdp1 | 5449.11478 | -0.3334539 | 0.19772763 | -1.6864302 | 0.09171296 | 0.46341374 |
| Socs7 | 157.734723 | -1.0710056 | 0.75145044 | -1.4252512 | 0.15408461 | 0.58303286 |
| Sv2a | 38.003644 | -0.003859 | 1.39109602 | -0.0027741 | 0.9977866 | 0.99967651 |
| Polr2l | 1864.47797 | -0.2708879 | 0.23355248 | -1.1598587 | 0.24610633 | 0.68117517 |
| Otud7b | 468.127617 | 0.06459028 | 0.49990287 | 0.12920567 | 0.89719491 | 0.97984623 |
| Tmco3 | 513.828324 | 0.66375172 | 0.42424392 | 1.56455211 | 0.11768797 | 0.52409061 |
| Prr3 | 591.773462 | -0.4027459 | 0.30611371 | -1.315674 | 0.18828349 | 0.61935434 |
| Ptov1 | 362.14546 | 0.43680427 | 0.30943185 | 1.41163318 | 0.158058 | 0.58698345 |
| Mesd | 6118.85393 | 0.41577387 | 0.28685917 | 1.44940068 | 0.14722572 | 0.57410946 |
| Dcun1d2 | 818.835925 | -0.1159427 | 0.26013568 | -0.4457008 | 0.65581338 | 0.91165371 |
| Parp12 | 507.945839 | 0.79205817 | 0.39285184 | 2.01617529 | 0.04378165 | 0.32051521 |
| Gdf15 | 22.2919266 | 2.54860466 | 1.3769229 | 1.85094217 | 0.06417787 | 0.3915482 |
| Rpf2 | 3580.73773 | -0.3032471 | 0.30633995 | -0.9899038 | 0.32222116 | 0.7445747 |
| Grtp1 | 147.812493 | 1.39584899 | 0.42295503 | 3.30023026 | 0.00096606 | 0.02277518 |
| Tbkbp1 | 364.859379 | 0.01269084 | 0.29025353 | 0.04372331 | 0.96512496 | 0.99430301 |
| Jarid2 | 1249.45757 | -0.4725834 | 0.36009655 | -1.3123796 | 0.1893921 | 0.61997152 |
| Tbc1d17 | 1251.69038 | 0.15079584 | 0.3400054 | 0.44351013 | 0.65739679 | 0.91258818 |
| Fchsd1 | 67.7122698 | 1.1593208 | 0.92071792 | 1.25914873 | 0.20797662 | 0.6418276 |

| | | | | | | |
|---|---|---|---|---|---|---|
| Armc10 | 1207.37753 | 0.13781023 | 0.19793983 | 0.69622287 | 0.48628927 | 0.84183502 |
| Car14 | 26.7656515 | 1.60146514 | 1.50418707 | 1.06467152 | 0.2870246 | 0.72261205 |
| C1rl | 204.753871 | 0.19895405 | 0.44272901 | 0.44938109 | 0.65315677 | 0.9107565 |
| Cbfa2t2 | 1571.48526 | -0.073328 | 0.40004162 | -0.1833008 | 0.85456198 | 0.96859981 |
| Osbpl7 | 363.459148 | 0.05904746 | 0.41528571 | 0.14218516 | 0.88693375 | 0.97827644 |
| Zfp280d | 373.16727 | -0.0236354 | 0.42961267 | -0.0550157 | 0.95612596 | 0.99215259 |
| Ubn2 | 281.645446 | -0.0368124 | 0.52175687 | -0.0705547 | 0.94375214 | 0.98992676 |
| Atf5 | 38.4388954 | -0.2036654 | 0.64020525 | -0.3181252 | 0.75038998 | 0.9399191 |
| Tmc3 | 21.7801258 | 1.60827159 | 1.51129654 | 1.0641668 | 0.28725315 | 0.72278333 |
| Pcid2 | 1984.30602 | -0.3692814 | 0.24301406 | -1.5195885 | 0.12861442 | 0.54422008 |
| BC028528 | 1134.48615 | 0.68138202 | 0.45673187 | 1.49186441 | 0.1357347 | 0.55758574 |
| Inip | 3011.00783 | -0.4085882 | 0.20613498 | -1.982139 | 0.04746369 | 0.33634712 |
| Cul7 | 391.075609 | -0.1384947 | 0.38046212 | -0.3640171 | 0.71584523 | 0.93235217 |
| Ranbp9 | 289.027903 | -0.0935396 | 0.49072748 | -0.1906141 | 0.84882798 | 0.96765585 |
| Ciart | 119.024957 | -0.4057513 | 0.52329363 | -0.7753798 | 0.43811527 | 0.81873879 |
| Fndc4 | 91.6666383 | 1.20818293 | 0.68998743 | 1.75102165 | 0.07994218 | 0.43723638 |
| Efl1 | 870.808378 | -0.7637373 | 0.29049065 | -2.6291287 | 0.00856039 | 0.11670212 |
| Ift172 | 106.863391 | 0.94696649 | 0.67043907 | 1.4124572 | 0.15781538 | 0.58658126 |
| Rad9b | 17.7800118 | -1.1204603 | 1.92920599 | -0.5807883 | 0.56138313 | 0.8752292 |
| Saxo2 | 15.371649 | 1.8040541 | 1.07937613 | 1.67138596 | 0.09464547 | 0.47133886 |
| Susd1 | 786.775899 | -0.3798338 | 0.34957504 | -1.0865587 | 0.27723188 | 0.71247429 |
| Pptc7 | 274.02586 | -0.4744716 | 0.4801379 | -0.9881987 | 0.32305534 | 0.7456483 |
| Akap12 | 39.1457095 | 1.82088685 | 1.3000656 | 1.40061152 | 0.16133028 | 0.59141542 |
| Tctn1 | 102.119215 | 0.43592193 | 0.52138583 | 0.8360832 | 0.4031081 | 0.79806139 |
| Cep85l | 342.292761 | -0.8422929 | 0.51283196 | -1.6424345 | 0.1005 | 0.48478567 |
| Ripor1 | 763.672157 | 0.25355892 | 0.36313464 | 0.69825042 | 0.4850206 | 0.84171939 |
| Samd10 | 221.151546 | 0.49347504 | 0.45402231 | 1.08689604 | 0.27708276 | 0.71247429 |
| Gng10 | 4400.41946 | 0.02300176 | 0.4589675 | 0.05011632 | 0.9600297 | 0.99306979 |
| Dock10 | 684.172846 | -0.728273 | 0.347824 | -2.0937974 | 0.03627803 | 0.2857625 |
| Phrf1 | 1609.4504 | -0.0460803 | 0.32909004 | -0.1400235 | 0.88864142 | 0.97846539 |
| Mcl1 | 2308.27622 | 0.07529269 | 0.26366913 | 0.28555747 | 0.77521708 | 0.94737568 |
| Nfe2l1 | 1329.07266 | -0.3412905 | 0.27035066 | -1.2623994 | 0.20680509 | 0.64174209 |
| Rassf7 | 257.754824 | 0.59950176 | 0.43948245 | 1.36410853 | 0.17253341 | 0.60492813 |
| Ensa | 3836.00886 | 0.17636361 | 0.20858428 | 0.84552688 | 0.39781673 | 0.79531492 |
| Med30 | 1151.5185 | -0.032532 | 0.33602484 | -0.0968143 | 0.92287387 | 0.98465273 |
| Tm6sf1 | 1913.14345 | -0.042551 | 0.22316262 | -0.1906726 | 0.8487821 | 0.96765585 |
| Polr3k | 3636.18827 | 0.09663511 | 0.29548849 | 0.3270351 | 0.74364134 | 0.93870235 |
| Degs1 | 6198.46876 | 0.15204554 | 0.15189496 | 1.00099135 | 0.31683099 | 0.74186126 |
| Lrrc56 | 114.04079 | 0.33981094 | 0.51512424 | 0.65966794 | 0.50946694 | 0.85335831 |
| Ctss | 16420.7283 | -0.129078 | 0.18392967 | -0.7017792 | 0.48281686 | 0.84106286 |
| Pold1 | 4468.60138 | -0.2622569 | 0.21675511 | -1.2099225 | 0.22630862 | 0.65977917 |
| Ramac | 2347.91611 | -0.1352636 | 0.1630885 | -0.8293877 | 0.40688508 | 0.79992362 |
| Creb3l2 | 327.209991 | 0.30221608 | 0.32832163 | 0.92048787 | 0.35731787 | 0.7664509 |
| Rnh1 | 9582.50908 | 0.19236127 | 0.27331645 | 0.70380424 | 0.48155469 | 0.84011103 |
| Ric1 | 545.112651 | 0.02571774 | 0.31952966 | 0.08048625 | 0.93585053 | 0.98794094 |
| Herc1 | 639.587347 | 0.31507432 | 0.27630802 | 1.14030101 | 0.25416092 | 0.68801737 |

| | | | | | | |
|---|---|---|---|---|---|---|
| Lpar1 | 52.4123146 | 1.87106437 | 1.20674544 | 1.55050461 | 0.12102045 | 0.52910763 |
| Arfrp1 | 1852.41206 | 0.22334217 | 0.25922183 | 0.86158704 | 0.38891481 | 0.78942657 |
| Trps1 | 459.330927 | -0.8199915 | 0.4028168 | -2.0356438 | 0.04178613 | 0.31168889 |
| Pak1ip1 | 3829.92165 | -0.2068784 | 0.33248419 | -0.6222203 | 0.533797 | 0.86500711 |
| Rtel1 | 1288.36526 | -0.0356206 | 0.34576907 | -0.1030184 | 0.91794838 | 0.98390186 |
| Atp5j2 | 4461.98464 | 0.45165387 | 0.42300701 | 1.06772196 | 0.28564595 | 0.721488 |
| Josd2 | 615.507461 | -0.066928 | 0.26036434 | -0.2570553 | 0.79713611 | 0.95397994 |
| Mapkap1 | 1309.4533 | 0.06226967 | 0.4088127 | 0.15231834 | 0.87893586 | 0.97535404 |
| Taf5l | 1248.17651 | -0.2223448 | 0.22410129 | -0.992162 | 0.32111852 | 0.74426731 |
| Dsel | 61.6906578 | -0.2761699 | 0.80949658 | -0.3411625 | 0.73298128 | 0.93728136 |
| Gmeb2 | 188.055147 | 0.06364264 | 0.49265494 | 0.12918299 | 0.89721285 | 0.97984623 |
| Golga4 | 287.412076 | 0.02831718 | 0.64544334 | 0.04387245 | 0.96500608 | 0.99428295 |
| Mindy1 | 1036.81489 | 0.30686087 | 0.30034982 | 1.0216782 | 0.30693323 | 0.73595594 |
| Atp5l | 6182.42377 | 0.52617701 | 0.44821789 | 1.17393129 | 0.24042255 | 0.67564175 |
| Pbx3 | 264.482372 | -0.0241552 | 0.48927188 | -0.0493697 | 0.96062465 | 0.99325523 |
| Bud31 | 5601.64457 | 0.40717165 | 0.40003936 | 1.01782896 | 0.30875925 | 0.73867239 |
| Mboat1 | 1190.67168 | 0.36441706 | 0.23789474 | 1.53184161 | 0.12556153 | 0.53724874 |
| Wdr26 | 2575.19423 | -0.1167141 | 0.29663344 | -0.3934624 | 0.69397802 | 0.92609254 |
| Nudcd1 | 2022.12472 | -0.0962014 | 0.2888624 | -0.3330353 | 0.73910765 | 0.93800224 |
| Mvb12b | 113.024882 | 0.58256751 | 0.3963032 | 1.47000456 | 0.14156052 | 0.56628837 |
| Angptl6 | 130.021412 | -0.2454292 | 0.62970123 | -0.389755 | 0.69671775 | 0.92646244 |
| Nup205 | 1272.9061 | -0.6435446 | 0.23245599 | -2.768458 | 0.00563223 | 0.08754282 |
| Abcf1 | 4101.08756 | -0.4558093 | 0.22830202 | -1.996519 | 0.04587746 | 0.33110261 |
| Alpk3 | 10.0457293 | 2.36061017 | 1.55548034 | 1.51760849 | 0.12911312 | 0.54519878 |
| Gabpb2 | 597.090861 | -0.247085 | 0.29741533 | -0.8307743 | 0.40610115 | 0.79992362 |
| Kdm3b | 405.422151 | -0.4248921 | 0.28148719 | -1.5094543 | 0.13118272 | 0.54970237 |
| Ascc3 | 580.917957 | -0.2345553 | 0.3577322 | -0.6556729 | 0.5120346 | 0.85398564 |
| Vill | 3353.45753 | 0.04197393 | 0.19023327 | 0.22064453 | 0.82536923 | 0.96324573 |
| Ephx1 | 32.0333416 | 1.23932457 | 0.75132729 | 1.64951358 | 0.09904246 | 0.48073092 |
| Smurf1 | 63.859049 | -0.2580473 | 0.87875884 | -0.2936497 | 0.76902556 | 0.94401697 |
| Stap2 | 11.3996901 | 1.50877765 | 2.06142504 | 0.73191002 | 0.46422349 | 0.83255328 |
| 1700028J19F | 24.1471098 | 0.74564835 | 1.35504611 | 0.55027526 | 0.58213059 | 0.88321857 |
| Cnot4 | 919.949431 | -0.4964249 | 0.30046498 | -1.6521889 | 0.09849604 | 0.48023199 |
| Lefty1 | 16.2331299 | 2.80471246 | 2.07838165 | 1.34946941 | 0.17718624 | 0.60816966 |
| Ost4 | 6519.89206 | 0.29121934 | 0.31198257 | 0.93344746 | 0.35058898 | 0.76238338 |
| Sde2 | 3692.90496 | 0.08232862 | 0.39490421 | 0.20847744 | 0.83485619 | 0.96641606 |
| Rap1gap2 | 238.387145 | -0.8211725 | 0.5081823 | -1.6159014 | 0.10611563 | 0.49777747 |
| Gngt2 | 1353.47317 | 0.36435263 | 0.40238763 | 0.90547671 | 0.36521289 | 0.77134829 |
| Trmt112 | 3160.2681 | -0.1248847 | 0.16650564 | -0.7500327 | 0.45323499 | 0.82610332 |
| Ctnnal1 | 107.394746 | -0.3131512 | 0.5312976 | -0.5894083 | 0.5555874 | 0.87235279 |
| Hace1 | 632.461594 | 0.31286926 | 0.26021408 | 1.20235333 | 0.22922666 | 0.6635423 |
| Abitram | 929.308611 | -0.4322037 | 0.23166186 | -1.8656664 | 0.06208805 | 0.3847635 |
| Tmem214 | 3273.67034 | 0.09897221 | 0.31110604 | 0.31813014 | 0.75038622 | 0.9399191 |
| Ralgps1 | 170.820701 | -1.3365674 | 0.46290853 | -2.8873251 | 0.00388533 | 0.06537715 |
| Agbl3 | 53.1614999 | 1.35424509 | 0.71380711 | 1.89721436 | 0.05779965 | 0.36898259 |
| Vars2 | 219.730625 | -0.0666936 | 0.46123696 | -0.1445973 | 0.88502884 | 0.97747903 |

| | | | | | | |
|---|---|---|---|---|---|---|
| Gcnt1 | 603.839062 | 0.07481308 | 0.27715348 | 0.26993375 | 0.78721122 | 0.95225386 |
| Kif16b | 269.809934 | 0.81631923 | 0.67547876 | 1.20850467 | 0.2268532 | 0.66015092 |
| Phb | 12151.8445 | 0.3444417 | 0.41418411 | 0.83161497 | 0.40562631 | 0.79982698 |
| Ythdf1 | 375.867877 | -0.4049414 | 0.33156139 | -1.2213164 | 0.22196624 | 0.65525654 |
| Itpkb | 84.4871492 | -0.4782354 | 0.74039659 | -0.6459179 | 0.51833252 | 0.8570838 |
| Baiap2l1 | 16.8277835 | 2.27000611 | 1.19814495 | 1.89460058 | 0.05814534 | 0.37027169 |
| Garnl3 | 19.6763481 | -0.2504847 | 1.41839638 | -0.1765971 | 0.85982485 | 0.96954641 |
| Pi4kb | 543.253061 | 0.0151248 | 0.25053752 | 0.06036942 | 0.95186142 | 0.99124208 |
| Zcchc2 | 181.597428 | -0.745772 | 0.46928087 | -1.5891804 | 0.11201967 | 0.50987629 |
| Bpgm | 503.017859 | 0.62440967 | 0.48345883 | 1.29154675 | 0.19651416 | 0.62897898 |
| Zfhx3 | 409.597043 | 0.32136007 | 0.49314396 | 0.65165571 | 0.5146233 | 0.85526675 |
| Rnf146 | 427.517693 | 0.00811684 | 0.53625711 | 0.0151361 | 0.9879236 | 0.9962454 |
| Mrps34 | 2450.29206 | -0.1016591 | 0.29501393 | -0.3445907 | 0.7304021 | 0.93653187 |
| A230050P20I | 216.17085 | 0.25375107 | 0.32720732 | 0.7755055 | 0.43804102 | 0.81873879 |
| Man2a2 | 644.533414 | -0.621045 | 0.49682389 | -1.2500304 | 0.21128844 | 0.64504405 |
| Ctu1 | 384.843227 | -0.4864461 | 0.36596669 | -1.3292087 | 0.18377912 | 0.61576828 |
| Fam117a | 72.7833753 | -1.6841388 | 1.1963361 | -1.4077472 | 0.15920594 | 0.58838861 |
| Irs2 | 87.5290219 | 0.16059929 | 0.51887842 | 0.30951236 | 0.75693181 | 0.9411065 |
| Zfp653 | 37.3558904 | 0.20323286 | 0.99280029 | 0.20470668 | 0.8378013 | 0.96683245 |
| Rpl12 | 1189.39846 | 0.36494391 | 0.4273127 | 0.85404414 | 0.39308053 | 0.79137753 |
| Pogz | 316.070722 | 0.27235873 | 0.30513044 | 0.89259771 | 0.37207265 | 0.77719221 |
| Kat7 | 4328.93522 | -0.2248729 | 0.2624263 | -0.8568992 | 0.39150062 | 0.79018736 |
| Plcl2 | 452.420803 | 0.40435229 | 0.31179273 | 1.29686248 | 0.19467851 | 0.62671111 |
| Dido1 | 690.973617 | -0.3520388 | 0.3618385 | -0.9729168 | 0.33059464 | 0.75003046 |
| Rccd1 | 72.2348588 | 0.58405371 | 0.61042156 | 0.95680388 | 0.33866626 | 0.7564609 |
| Sccpdh | 1634.14645 | 0.88575948 | 0.28410601 | 3.11770767 | 0.00182263 | 0.03753585 |
| Prc1 | 4868.30131 | -0.2041425 | 0.18319456 | -1.114348 | 0.2651299 | 0.70112124 |
| Cnst | 426.236377 | 0.22736465 | 0.50259199 | 0.45238415 | 0.65099226 | 0.91021183 |
| Supt3 | 559.670327 | 0.03991803 | 0.27638643 | 0.14442832 | 0.88516226 | 0.97751512 |
| Edc3 | 2079.1182 | -0.0081093 | 0.16004553 | -0.0506686 | 0.9595896 | 0.99286641 |
| Slco4a1 | 165.281976 | -2.728799 | 0.81445297 | -3.3504684 | 0.00080675 | 0.0197775 |
| Ube2l3 | 11668.2834 | 0.00781194 | 0.17482956 | 0.04468316 | 0.96435985 | 0.99406401 |
| Pdk2 | 50.2671256 | 0.51190453 | 1.30278993 | 0.39292945 | 0.69437159 | 0.92609254 |
| Lmtk2 | 59.7834873 | -0.7280503 | 1.22155654 | -0.5960021 | 0.5511738 | 0.87093412 |
| Rabggtb | 2038.37544 | -0.2675018 | 0.2346425 | -1.1400398 | 0.25426973 | 0.68815964 |
| Ppp1r9b | 374.279653 | 0.00454729 | 0.27216673 | 0.01670774 | 0.98666978 | 0.99611549 |
| Bloc1s5 | 1310.05312 | 0.75769014 | 0.3506882 | 2.16058068 | 0.03072775 | 0.26139952 |
| Cables2 | 48.2956531 | 0.6231637 | 0.62236508 | 1.00128321 | 0.31668991 | 0.74186126 |
| Txndc5 | 9857.56833 | -0.3764992 | 0.21813861 | -1.7259631 | 0.08435406 | 0.44526833 |
| Ube3c | 404.158826 | 0.1103371 | 0.30230143 | 0.36499034 | 0.71511862 | 0.93235217 |
| Rps21 | 2670.43393 | 0.30992321 | 0.44613424 | 0.69468599 | 0.48725211 | 0.84260037 |
| Tlr4 | 2839.83435 | 0.68799144 | 0.27789757 | 2.47570156 | 0.01329747 | 0.15581219 |
| Cpq | 231.316093 | 1.13560742 | 0.59814625 | 1.89854475 | 0.05762436 | 0.36897087 |
| Siglecf | 209.8919 | -1.8363741 | 0.51010382 | -3.6000006 | 0.00031822 | 0.00926599 |
| Timm8b | 2131.35694 | 0.76705375 | 0.51652952 | 1.48501436 | 0.13754002 | 0.5600695 |
| Mtg1 | 972.431991 | -0.0338608 | 0.19535217 | -0.1733319 | 0.86239056 | 0.9705375 |

| | | | | | | |
|---|---|---|---|---|---|---|
| Arhgap18 | 2859.52899 | 0.65532093 | 0.20130731 | 3.2553261 | 0.00113262 | 0.02575347 |
| Tasp1 | 646.268164 | 0.16067069 | 0.36689803 | 0.43791648 | 0.66144683 | 0.91376977 |
| Adrm1 | 4871.93098 | -0.2162505 | 0.23630848 | -0.9151195 | 0.36012888 | 0.76857232 |
| Arpin | 21.4231069 | 1.23703296 | 1.27820766 | 0.96778716 | 0.33315068 | 0.75248011 |
| Usp6nl | 113.56995 | 0.28496746 | 0.50290554 | 0.56664213 | 0.57095735 | 0.88035336 |
| Pigk | 1882.94643 | 0.14751301 | 0.26091763 | 0.56536239 | 0.5718273 | 0.88035336 |
| Foxred1 | 2511.02254 | -0.1631868 | 0.1581623 | -1.031768 | 0.30218082 | 0.73372823 |
| Osbpl2 | 1116.8827 | -0.2224005 | 0.24545673 | -0.906068 | 0.36489986 | 0.77108653 |
| Eme1 | 1344.02138 | -0.1332787 | 0.20115032 | -0.6625825 | 0.50759797 | 0.85197408 |
| Anpep | 50.1503845 | 1.15859591 | 0.74687988 | 1.55124798 | 0.12084227 | 0.52910763 |
| Atpsckmt | 699.613996 | 0.28020289 | 0.38918283 | 0.71997753 | 0.47153883 | 0.83393588 |
| Psmd7 | 10071.5082 | -0.0336358 | 0.19041648 | -0.1766432 | 0.85978863 | 0.96954641 |
| Zzz3 | 1236.73229 | 0.16290114 | 0.44853031 | 0.36318869 | 0.71646393 | 0.93235217 |
| Mtg2 | 1129.06216 | 0.00800272 | 0.29020641 | 0.02757595 | 0.97800036 | 0.99483019 |
| Chad | 6.64361664 | -6.1341487 | 2.22740316 | -2.7539463 | 0.00588814 | 0.09037597 |
| Ss18l1 | 74.7894461 | -0.6355995 | 0.77280801 | -0.8224546 | 0.4108182 | 0.80139357 |
| Rreb1 | 508.07211 | -0.0556507 | 0.5421053 | -0.1026565 | 0.91823556 | 0.98390186 |
| L3mbtl3 | 223.501424 | -0.007539 | 0.33427189 | -0.0225536 | 0.98200632 | 0.99523682 |
| Rsad1 | 151.627374 | -0.5893202 | 0.43642479 | -1.3503363 | 0.17690814 | 0.60784615 |
| March6 | 596.672013 | 0.32092163 | 0.40160646 | 0.79909478 | 0.42423545 | 0.80936873 |
| Atp6v1g1 | 9639.44784 | 0.67607119 | 0.32660579 | 2.06999141 | 0.03845315 | 0.29640666 |
| Lsm14b | 34.0020722 | 0.08224665 | 0.76968339 | 0.10685777 | 0.91490182 | 0.98347559 |
| F13a1 | 35664.6595 | 1.00072209 | 0.19114543 | 5.23539626 | 1.646312759 | 1.502936118 |
| Mycbpap | 13.5772357 | 0.7384824 | 2.07537053 | 0.35583159 | 0.72196669 | 0.93398714 |
| Nrn1 | 10.2473365 | 0.90171984 | 1.70629923 | 0.52846524 | 0.59717647 | 0.88891958 |
| Itga9 | 277.397777 | 1.22535301 | 0.39022829 | 3.14009269 | 0.00168894 | 0.03525742 |
| Taf4 | 116.764086 | 0.57605618 | 0.61482232 | 0.9369474 | 0.34878562 | 0.76163771 |
| Prune2 | 1060.44327 | 1.69609955 | 0.34104944 | 4.97317793 | 6.586415695 | 5.099469822 |
| Cdc123 | 5726.23505 | -0.0464275 | 0.20792838 | -0.2232858 | 0.82331308 | 0.9628698 |
| Zc3hc1 | 2548.14755 | 0.04963284 | 0.19543292 | 0.25396355 | 0.79952372 | 0.95515043 |
| Camk1d | 1331.46834 | -0.4492233 | 0.30809642 | -1.4580607 | 0.14482381 | 0.57067981 |
| Sart1 | 530.573909 | -0.2699332 | 0.3685405 | -0.7324384 | 0.46390104 | 0.83221901 |
| Runx2 | 50.3099934 | 1.07187941 | 0.86294345 | 1.24212011 | 0.21419225 | 0.64776608 |
| Stim2 | 314.521178 | -0.1470772 | 0.8479003 | -0.1734605 | 0.86228944 | 0.9705375 |
| Fam102a | 215.308072 | 1.24083686 | 0.80411195 | 1.54311456 | 0.12280298 | 0.53283544 |
| Akna | 1676.54954 | -0.3449063 | 0.36464606 | -0.9458659 | 0.34421701 | 0.75962035 |
| Ube2h | 1707.12186 | -0.0600773 | 0.27928175 | -0.2151136 | 0.82967876 | 0.96462215 |
| Cmc1 | 347.006906 | 0.12536133 | 0.32367336 | 0.38730815 | 0.69852809 | 0.92669304 |
| Naif1 | 444.090287 | -0.4890467 | 0.29346362 | -1.6664644 | 0.09562096 | 0.47434356 |
| Akap7 | 135.494926 | 0.6093565 | 0.45258546 | 1.34638991 | 0.1781768 | 0.60928307 |
| Adgrl4 | 98.1610513 | 0.56181287 | 0.72542502 | 0.77446029 | 0.43865864 | 0.8193372 |
| Dap | 1383.83017 | 0.34326211 | 0.19671931 | 1.74493343 | 0.08099646 | 0.4392955 |
| Polg | 1415.08792 | 0.00606017 | 0.2556475 | 0.02370517 | 0.98108778 | 0.99522729 |
| Tbc1d19 | 620.264754 | 0.55055057 | 0.25564495 | 2.15357497 | 0.03127352 | 0.26457051 |
| AW209491 | 964.236318 | 0.14636565 | 0.26086435 | 0.56107952 | 0.57474333 | 0.88059206 |
| Nubp2 | 7300.14689 | -0.1859517 | 0.25911615 | -0.7176382 | 0.47298038 | 0.83483892 |

| | | | | | | |
|---|---|---|---|---|---|---|
| Fanci | 932.955483 | 0.22288334 | 0.25598888 | 0.87067584 | 0.38393117 | 0.78704643 |
| Rbpj | 2877.18838 | 0.30077787 | 0.24954175 | 1.20532083 | 0.22807947 | 0.66211765 |
| Nlrc4 | 222.688668 | 0.31464736 | 0.38989323 | 0.80700905 | 0.41966127 | 0.80563709 |
| 1110008P14I | 188.773064 | -0.3129618 | 0.32789357 | -0.9544614 | 0.33985014 | 0.75726534 |
| Adk | 2901.99314 | -0.1493111 | 0.22039602 | -0.6774673 | 0.49810949 | 0.84860011 |
| Zdhhc1 | 86.0740367 | 1.04317634 | 1.19912322 | 0.86994924 | 0.38432814 | 0.78704643 |
| Atf7ip2 | 56.9081432 | 0.1407442 | 0.68530294 | 0.20537515 | 0.83727903 | 0.96683245 |
| Tbc1d25 | 379.812042 | -0.1837885 | 0.34001053 | -0.5405378 | 0.58882623 | 0.88690478 |
| Abhd2 | 320.902153 | -0.4999936 | 0.36478352 | -1.3706584 | 0.17048146 | 0.60430173 |
| Ciz1 | 962.050786 | -0.2975406 | 0.3451742 | -0.8620014 | 0.38868675 | 0.78942657 |
| Daglb | 6123.3344 | -0.0011715 | 0.17095153 | -0.0068529 | 0.99453224 | 0.99882461 |
| Metrnl | 32.1356059 | 1.50755406 | 0.98796637 | 1.52591637 | 0.12703069 | 0.54111158 |
| Rpl39l | 119.308284 | -0.4120384 | 0.57052229 | -0.7222126 | 0.47016377 | 0.83381643 |
| Gpatch2 | 405.050262 | 0.24381892 | 0.41767389 | 0.58375427 | 0.55938565 | 0.87397022 |
| Il18 | 524.198903 | 1.90688628 | 0.3892488 | 4.89888802 | 9.638054874 | 6.976468951 |
| Srrm2 | 3258.85279 | 0.1391586 | 0.41229738 | 0.33751998 | 0.73572496 | 0.9377627 |
| Arid4b | 607.827244 | 0.1057463 | 0.46154522 | 0.22911362 | 0.8187806 | 0.96105733 |
| Ppp1r10 | 1754.57351 | 0.22142658 | 0.24742773 | 0.89491416 | 0.37083298 | 0.77651486 |
| Rpl22l1 | 8185.3815 | 0.34366647 | 0.44793858 | 0.76721783 | 0.44295202 | 0.82038676 |
| Tbcd | 2150.02044 | -0.1455763 | 0.26274946 | -0.5540499 | 0.57954473 | 0.88286755 |
| Suv39h1 | 4015.16759 | -0.3317997 | 0.19280959 | -1.7208673 | 0.08527491 | 0.44656163 |
| Stx11 | 310.009891 | -0.2245327 | 0.5087742 | -0.441321 | 0.65898065 | 0.91280998 |
| Tbce | 879.282129 | -0.2569473 | 0.30419796 | -0.8446715 | 0.39829429 | 0.79555974 |
| Sec24d | 990.870113 | 0.97845648 | 0.34674263 | 2.82185223 | 0.00477472 | 0.0768541 |
| Isg20 | 56.0633626 | -0.9648712 | 1.00923553 | -0.9560416 | 0.33905121 | 0.75683748 |
| Tgfb2 | 45.5206114 | 3.55046841 | 1.79269992 | 1.98051462 | 0.04764573 | 0.33690765 |
| B3galnt2 | 906.778638 | -0.5592423 | 0.28614206 | -1.954422 | 0.05065133 | 0.34442343 |
| E130309D02 | 1754.10927 | -0.3222198 | 0.27419038 | -1.1751681 | 0.23992745 | 0.67563574 |
| Lyplal1 | 1638.69269 | 0.72241049 | 0.39871103 | 1.8118648 | 0.07000709 | 0.40878125 |
| Fn3krp | 391.387658 | 0.22125933 | 0.31365727 | 0.70541751 | 0.48055045 | 0.83935746 |
| Pomt1 | 999.711969 | -0.1373615 | 0.27712738 | -0.4956619 | 0.62013299 | 0.89896588 |
| Prrc2b | 546.409287 | 0.12660861 | 0.46170822 | 0.27421779 | 0.78391725 | 0.95094799 |
| Npepl1 | 1509.02242 | -0.1752287 | 0.22704776 | -0.7717703 | 0.44025048 | 0.81963526 |
| Megf9 | 256.736837 | -0.7709256 | 0.30269172 | -2.5469 | 0.01086846 | 0.13520102 |
| Foxk2 | 403.399462 | -0.1243706 | 0.34961725 | -0.3557336 | 0.72204009 | 0.93398714 |
| Azi2 | 2044.63442 | 0.06062145 | 0.17673077 | 0.3430158 | 0.73158657 | 0.93653187 |
| Fndc3b | 277.47468 | 0.03904891 | 0.41019635 | 0.09519564 | 0.92415943 | 0.9849488 |
| Cybc1 | 5561.14283 | 0.0736877 | 0.17203109 | 0.42833945 | 0.668404 | 0.91604147 |
| Cdk5rap2 | 374.285845 | -0.3136955 | 0.42150877 | -0.7442207 | 0.456743 | 0.82915295 |
| Hexdc | 53.5715367 | 0.83729981 | 0.6575252 | 1.27341099 | 0.20287224 | 0.63809363 |
| Ndst2 | 287.820143 | -0.2036572 | 0.40349581 | -0.5047318 | 0.61374722 | 0.89776035 |
| Rftn1 | 1098.51593 | 0.27602743 | 0.23829695 | 1.1583339 | 0.2467278 | 0.68177574 |
| Rab3gap2 | 1446.93584 | 0.32647364 | 0.31824844 | 1.02584523 | 0.30496454 | 0.73510332 |
| Igfbp2 | 84.8376921 | -0.7025314 | 1.41840846 | -0.4952955 | 0.62039153 | 0.89896588 |
| Rnf122 | 58.9450158 | 1.06123959 | 1.28808333 | 0.82389048 | 0.41000179 | 0.80117888 |
| Mettl22 | 419.836638 | -0.3590028 | 0.33821513 | -1.061463 | 0.28847954 | 0.72375492 |

| | | | | | | |
|---|---|---|---|---|---|---|
| Atp6v0e2 | 68.281923 | 0.65008195 | 0.97698464 | 0.66539629 | 0.50579706 | 0.85074333 |
| Smarcal1 | 961.878685 | 0.23927894 | 0.24918036 | 0.96026405 | 0.33692234 | 0.7553565 |
| Exosc2 | 2265.76173 | -0.1200697 | 0.23175968 | -0.5180783 | 0.60440366 | 0.89258167 |
| Fut11 | 1189.20397 | 0.21639505 | 0.19716609 | 1.09752673 | 0.2724112 | 0.71005689 |
| Picalm | 3402.53759 | 0.41608458 | 0.27458775 | 1.51530639 | 0.12969483 | 0.54639717 |
| Sec24c | 4927.55079 | -0.1590442 | 0.26086883 | -0.6096711 | 0.54207972 | 0.86853454 |
| Hlx | 552.721721 | 0.20620358 | 0.26496911 | 0.77821742 | 0.43644085 | 0.81735776 |
| Wdr45 | 377.138561 | 0.36019581 | 0.33818143 | 1.06509634 | 0.28683234 | 0.72260415 |
| Dusp10 | 233.908436 | 0.04507974 | 0.52870754 | 0.08526404 | 0.93205148 | 0.98730569 |
| Mreg | 52.8274832 | -2.127248 | 0.97540261 | -2.1808922 | 0.02919139 | 0.25470632 |
| Neil3 | 934.453371 | 0.05658673 | 0.21788703 | 0.25970675 | 0.79508999 | 0.95367917 |
| Prss23 | 19.9778812 | 2.8799818 | 2.08029534 | 1.38441006 | 0.16623293 | 0.59950691 |
| Prdm16 | 79.9195655 | 0.19880066 | 0.86594718 | 0.22957596 | 0.81842129 | 0.96105733 |
| Heatr5b | 259.953882 | -0.21022 | 0.48753774 | -0.4311871 | 0.66633236 | 0.91520501 |
| Alg1 | 1151.27644 | -0.1241104 | 0.21724031 | -0.5713047 | 0.56779313 | 0.87929304 |
| Tmem135 | 2035.51159 | 0.07903479 | 0.32913777 | 0.24012676 | 0.81023199 | 0.95884376 |
| Mtmr7 | 18.2459454 | -2.1527286 | 1.33206595 | -1.6160826 | 0.10607645 | 0.49777747 |
| Prpf18 | 1456.12085 | -0.2230496 | 0.22590898 | -0.9873428 | 0.32347461 | 0.74583177 |
| Dcxr | 580.491498 | 0.8130408 | 0.30075003 | 2.70337728 | 0.00686388 | 0.10086772 |
| Morc3 | 4108.88436 | 0.12429735 | 0.18405389 | 0.67533128 | 0.49946532 | 0.84872332 |
| Mtmr12 | 666.078138 | -0.3330643 | 0.36392842 | -0.9151918 | 0.36009095 | 0.76857232 |
| Tcta | 784.19029 | 0.66129927 | 0.32270097 | 2.04926335 | 0.04043637 | 0.30572195 |
| Slc9a8 | 694.600437 | -0.0846575 | 0.33070903 | -0.255988 | 0.79796013 | 0.95433165 |
| Zdhhc2 | 122.408982 | 1.24693152 | 0.6000135 | 2.07817244 | 0.03769348 | 0.29276466 |
| Ubn1 | 739.678697 | 0.20982457 | 0.35758366 | 0.58678456 | 0.55734841 | 0.87276537 |
| Wfs1 | 99.1712385 | 0.53858976 | 0.48727739 | 1.10530424 | 0.26902778 | 0.70592381 |
| Tnrc18 | 66.9854243 | 0.36290435 | 0.66022661 | 0.54966634 | 0.58254825 | 0.88343755 |
| Micu3 | 122.474147 | 0.94223755 | 0.38811402 | 2.42773385 | 0.01519349 | 0.17040121 |
| Nt5dc1 | 1090.23447 | 0.10219342 | 0.25990105 | 0.39320126 | 0.69417084 | 0.92609254 |
| Asb6 | 1167.31068 | -0.0275461 | 0.26853355 | -0.1025796 | 0.91829662 | 0.98390186 |
| Tspyl4 | 96.6190143 | 1.02301625 | 0.56767572 | 1.80211382 | 0.0715275 | 0.41122734 |
| Dse | 6811.66594 | 0.41458882 | 0.21178008 | 1.95763845 | 0.05027245 | 0.34299029 |
| Znfx1 | 234.42385 | -0.32564 | 0.48207243 | -0.6755001 | 0.49935811 | 0.84872332 |
| Nup133 | 2486.84141 | -0.2880844 | 0.16893381 | -1.7053095 | 0.08813671 | 0.45368281 |
| Uhrf1bp1 | 184.093429 | 0.59832265 | 0.44526621 | 1.34374142 | 0.17903202 | 0.60992118 |
| Ptpa | 5549.24987 | 0.0576126 | 0.16537864 | 0.34836786 | 0.72756394 | 0.93576253 |
| Cep104 | 582.793035 | -0.2530488 | 0.30601518 | -0.826916 | 0.40828467 | 0.80087574 |
| Tusc3 | 1633.51791 | 0.42402027 | 0.32993768 | 1.28515259 | 0.19873897 | 0.63148061 |
| Zup1 | 548.013027 | -0.2617692 | 0.33608163 | -0.7788858 | 0.43604698 | 0.81725842 |
| Stau1 | 2475.8212 | 0.23931375 | 0.31844687 | 0.7515029 | 0.45235005 | 0.82603347 |
| Ncam1 | 57.0363787 | 0.19737673 | 0.59678924 | 0.33073105 | 0.74084764 | 0.93800224 |
| Rsph4a | 8.57721041 | 1.32265766 | 2.67097371 | 0.49519681 | 0.6204612 | 0.89896588 |
| Ubald1 | 1319.01076 | 0.21207578 | 0.2617548 | 0.8102078 | 0.41782075 | 0.8051099 |
| Nphp4 | 7.88562796 | 2.38293635 | 2.6472469 | 0.90015644 | 0.368037 | 0.77410534 |
| Myo9a | 638.111177 | 0.44611607 | 0.32411722 | 1.37640349 | 0.16869675 | 0.60312169 |
| Fam149b | 140.546437 | -0.0688878 | 0.67400414 | -0.1022067 | 0.91859259 | 0.98390186 |

| | | | | | | |
|---|---|---|---|---|---|---|
| Tmem246 | 70.2040795 | 2.31993656 | 0.85067781 | 2.72716242 | 0.00638816 | 0.09644997 |
| Stub1 | 7989.63609 | -0.0642117 | 0.15004632 | -0.4279461 | 0.66869038 | 0.91618049 |
| Gm7488 | 2050.72923 | -0.2859598 | 0.22003973 | -1.2995827 | 0.19374402 | 0.62518945 |
| Prex1 | 1830.87313 | 0.01851107 | 0.28324295 | 0.06535405 | 0.9478921 | 0.99019648 |
| Ap5z1 | 1093.53381 | 0.08454371 | 0.19743479 | 0.42821078 | 0.66849767 | 0.91606721 |
| Hnrnpu | 9649.94412 | 0.41718613 | 0.55181896 | 0.75601993 | 0.44963725 | 0.82484234 |
| Lonrf1 | 48.2035715 | 0.14361505 | 0.69790008 | 0.20578168 | 0.83696146 | 0.96683245 |
| Zfp189 | 200.824419 | 1.09367823 | 0.53271909 | 2.05301115 | 0.0400715 | 0.30409095 |
| Coro7 | 3603.87249 | -0.0812078 | 0.23343478 | -0.3478822 | 0.72792869 | 0.93576253 |
| Mrpl12 | 7557.51432 | 0.16067639 | 0.36907319 | 0.43535101 | 0.66330767 | 0.91395288 |
| Vasn | 44.7636283 | 1.00278289 | 1.56696452 | 0.63995251 | 0.52220347 | 0.85816354 |
| Kyat1 | 245.043828 | -0.4837433 | 0.49226358 | -0.9826917 | 0.32575921 | 0.74723652 |
| Cpeb3 | 25.8159239 | 0.38443048 | 1.54441748 | 0.24891617 | 0.80342563 | 0.95672466 |
| Rxrb | 656.776492 | -0.6055816 | 0.23694178 | -2.5558243 | 0.01059366 | 0.13331258 |
| Spout1 | 2861.62673 | -0.1827677 | 0.19557369 | -0.9345208 | 0.35003532 | 0.76216093 |
| Icmt | 516.239048 | 0.49466808 | 0.32435717 | 1.52507213 | 0.12724111 | 0.54121713 |
| Oxld1 | 218.348242 | 0.55664314 | 0.67079994 | 0.8298199 | 0.40664062 | 0.79992362 |
| Zmynd8 | 2333.78554 | 0.16248198 | 0.21521886 | 0.75496162 | 0.45027201 | 0.82494795 |
| Tbc1d13 | 1080.50671 | 0.18533297 | 0.30746667 | 0.60277417 | 0.54665893 | 0.86994632 |
| Mrps6 | 1259.24213 | 0.21893999 | 0.35711318 | 0.61308291 | 0.53982152 | 0.86719392 |
| Lap3 | 4830.39154 | -0.1258209 | 0.18971615 | -0.6632063 | 0.50719846 | 0.85169216 |
| Zer1 | 49.8136779 | 0.18481522 | 0.58696048 | 0.31486826 | 0.75286168 | 0.94018119 |
| Msantd3 | 58.9564956 | 0.389965 | 0.68000238 | 0.57347593 | 0.56632249 | 0.87871931 |
| Ncoa7 | 472.86137 | -0.263177 | 0.33812162 | -0.77835 | 0.4363627 | 0.81733653 |
| Usp53 | 28.9983708 | -0.2491421 | 1.41160498 | -0.1764956 | 0.8599046 | 0.96954641 |
| Nploc4 | 557.46266 | -0.342671 | 0.44508992 | -0.7698916 | 0.4413642 | 0.81967485 |
| Lmbrd2 | 144.318799 | 0.2049394 | 0.58999627 | 0.34735712 | 0.72832304 | 0.93605089 |
| Ldb2 | 72.7228493 | 4.47454116 | 2.8435553 | 1.57357276 | 0.11558625 | 0.5202232 |
| Plekhg5 | 421.075965 | 0.08485058 | 0.28943042 | 0.29316402 | 0.76939678 | 0.94401697 |
| Wdr34 | 666.753494 | -0.4312785 | 0.28544762 | -1.5108849 | 0.13081779 | 0.54930207 |
| Dock3 | 20.7023015 | 0.13385261 | 1.09229616 | 0.12254241 | 0.90246946 | 0.98043702 |
| Trp53rka | 1044.69036 | 0.00622721 | 0.3075813 | 0.02024573 | 0.98384735 | 0.99564261 |
| Fnbp1l | 82.9319724 | 0.40749761 | 0.65403601 | 0.62305072 | 0.53325119 | 0.86492466 |
| Prkrip1 | 736.093143 | -0.0093528 | 0.37158881 | -0.0251699 | 0.97991946 | 0.99486654 |
| Slx4 | 259.19008 | 0.24863891 | 0.40554908 | 0.61309204 | 0.53981548 | 0.86719392 |
| Alg2 | 795.355276 | 0.08764135 | 0.34175646 | 0.25644387 | 0.79760812 | 0.95424353 |
| Bahcc1 | 72.2812704 | 1.62789971 | 0.72438447 | 2.24728688 | 0.0246217 | 0.22749037 |
| Htatip2 | 1089.17526 | 0.23669077 | 0.31925999 | 0.74137312 | 0.45846724 | 0.82986925 |
| Orai2 | 902.636403 | -0.4686466 | 0.38431838 | -1.2194229 | 0.22268373 | 0.65613364 |
| Exo1 | 2536.79627 | -0.0549242 | 0.17101591 | -0.3211641 | 0.748086 | 0.9399191 |
| Fbxl5 | 2851.50416 | 0.44425702 | 0.293443 | 1.51394655 | 0.1300394 | 0.54703299 |
| Alkbh4 | 504.016889 | 0.25232575 | 0.26551173 | 0.95033751 | 0.34194079 | 0.75874982 |
| Dnttip2 | 4727.79341 | -0.0304464 | 0.23603864 | -0.1289889 | 0.89736645 | 0.97984623 |
| Thap3 | 257.941753 | 0.07433323 | 0.31212064 | 0.23815544 | 0.81176054 | 0.95967622 |
| Dnajc28 | 62.7266625 | -0.3144121 | 0.9843784 | -0.3194017 | 0.74942195 | 0.9399191 |
| Cc2d2a | 37.6277296 | 1.92529029 | 0.6780157 | 2.83959545 | 0.00451708 | 0.07377492 |

| | | | | | | |
|---|---|---|---|---|---|---|
| Dnajc11 | 5757.84543 | -0.3496582 | 0.22188626 | -1.5758442 | 0.11506172 | 0.51881682 |
| Ypel5 | 2003.97094 | 0.4958398 | 0.17935698 | 2.76454142 | 0.00570029 | 0.08837973 |
| Polr2j | 2468.97638 | 0.25662864 | 0.35771033 | 0.71742026 | 0.47311482 | 0.83483892 |
| Galnt12 | 26.0605237 | -0.8564695 | 1.49807097 | -0.5717149 | 0.56751513 | 0.87929304 |
| Cep131 | 206.866324 | -0.5703677 | 0.4327537 | -1.3179962 | 0.18750493 | 0.61935434 |
| Cpeb2 | 125.83496 | -0.0744564 | 0.43543258 | -0.1709941 | 0.86422842 | 0.9705375 |
| Kmo | 186.99944 | -0.7682188 | 0.51506388 | -1.4915019 | 0.13582976 | 0.55758574 |
| Cercam | 464.232636 | -0.5212217 | 0.33234204 | -1.5683292 | 0.11680432 | 0.52252063 |
| Zfp597 | 147.571958 | -0.0900923 | 0.38782347 | -0.2323023 | 0.81630322 | 0.96079676 |
| Zfand1 | 1043.5142 | 0.38515929 | 0.387751 | 0.99331607 | 0.32055595 | 0.74413737 |
| Cplane1 | 79.2076807 | -0.5170667 | 0.88115366 | -0.5868065 | 0.55733367 | 0.87276537 |
| Ncoa5 | 366.154681 | -0.0342324 | 0.50979531 | -0.0671493 | 0.94646283 | 0.99019648 |
| Zc3h10 | 504.927344 | -0.0458594 | 0.33811578 | -0.1356323 | 0.892112 | 0.97937556 |
| Tbc1d2 | 114.355793 | -2.3216842 | 0.80534655 | -2.8828387 | 0.00394109 | 0.06622442 |
| Xkr5 | 221.084957 | -0.052526 | 0.48378142 | -0.1085737 | 0.9135406 | 0.98309098 |
| Trub2 | 852.012009 | 0.3004697 | 0.23910008 | 1.25666917 | 0.20887346 | 0.64228937 |
| Wdr70 | 1554.65854 | 0.29037352 | 0.24290778 | 1.19540645 | 0.23192827 | 0.66733333 |
| Olig2 | 408.568657 | -0.4789893 | 0.35580282 | -1.3462213 | 0.17823116 | 0.60928307 |
| Arhgap29 | 95.6097204 | 1.53137465 | 0.47469335 | 3.22602932 | 0.0012552 | 0.02751778 |
| Zfp335 | 497.190542 | -0.4807027 | 0.23779193 | -2.0215267 | 0.04322527 | 0.31923815 |
| Nhsl1 | 28.4343742 | 0.13104582 | 1.02884475 | 0.12737181 | 0.89864613 | 0.97989497 |
| Epg5 | 148.282318 | -0.1254719 | 0.86613875 | -0.1448635 | 0.88481864 | 0.97742337 |
| Zfp800 | 544.812354 | -0.437197 | 0.30235603 | -1.4459675 | 0.14818631 | 0.57506401 |
| Mcph1 | 1125.84055 | -0.3367714 | 0.29964649 | -1.1238957 | 0.26105729 | 0.69691361 |
| Rapgef1 | 501.118671 | -0.6719129 | 0.45704264 | -1.4701319 | 0.14152605 | 0.56628837 |
| Pcif1 | 1669.82551 | -0.1883508 | 0.20929784 | -0.8999176 | 0.36816412 | 0.77421783 |
| Endov | 260.081409 | 0.76390601 | 0.39827488 | 1.91803714 | 0.0551063 | 0.36145598 |
| Rere | 351.518139 | -0.4701931 | 0.57000709 | -0.8248899 | 0.40943408 | 0.80100711 |
| Trim14 | 660.62922 | -0.1474859 | 0.35460139 | -0.4159203 | 0.67746834 | 0.91958179 |
| Neurl2 | 324.798901 | -0.2316676 | 0.26870516 | -0.862163 | 0.38859784 | 0.78942657 |
| Heca | 84.9277365 | -0.5434471 | 0.60742885 | -0.8946679 | 0.37096467 | 0.7765614 |
| Tmem120a | 1475.28467 | 0.76411238 | 0.31179252 | 2.45070786 | 0.01425756 | 0.16269846 |
| Alg14 | 969.81833 | 0.29992855 | 0.19660189 | 1.52556291 | 0.12711876 | 0.54111158 |
| Txlnb | 21.2242705 | 4.02197045 | 1.312929 | 3.06335716 | 0.00218869 | 0.04271935 |
| Fgl2 | 416.577333 | -1.7079116 | 0.31880065 | -5.3573028 | 8.447348618 | 8.507181661 |
| Armh3 | 695.570735 | 0.36393003 | 0.24205408 | 1.50350711 | 0.13270833 | 0.55237188 |
| Slc26a11 | 526.861583 | 0.2851392 | 0.33461186 | 0.85214913 | 0.39413133 | 0.79233901 |
| Cited2 | 358.808161 | -0.1562127 | 0.37277411 | -0.4190547 | 0.67517615 | 0.91848602 |
| Spsb1 | 156.559083 | -0.6913493 | 0.68938717 | -1.0028462 | 0.31593507 | 0.74186126 |
| Coq10a | 42.2593187 | 0.80068897 | 0.85788459 | 0.93332947 | 0.35064988 | 0.76238338 |
| Rhbdd2 | 969.8148 | 0.09645196 | 0.22090831 | 0.43661537 | 0.66239032 | 0.91395288 |
| Urb1 | 232.632367 | -0.6612983 | 0.44776135 | -1.476899 | 0.1397027 | 0.56329174 |
| Gsap | 647.71207 | 0.2889021 | 0.43136226 | 0.66974358 | 0.50302127 | 0.85012672 |
| Pik3cd | 3706.11414 | -0.2931485 | 0.27714007 | -1.0577628 | 0.2901636 | 0.72418639 |
| Ptger4 | 276.472475 | -1.0840928 | 0.69943779 | -1.5499489 | 0.12115378 | 0.52912326 |
| Plcb4 | 109.660922 | -0.741364 | 0.74084762 | -1.000697 | 0.31697332 | 0.74186126 |

| | | | | | | |
|---|---|---|---|---|---|---|
| Dag1 | 884.426932 | 0.7135447 | 0.40961042 | 1.74200819 | 0.08150701 | 0.44011802 |
| Clstn1 | 109.032724 | 0.18358177 | 0.68348684 | 0.26859591 | 0.78824066 | 0.95225386 |
| Etfbkmt | 139.427148 | -0.012573 | 0.48595089 | -0.0258729 | 0.97935868 | 0.99483019 |
| Hip1 | 1289.82124 | 0.15961445 | 0.27961973 | 0.57082685 | 0.56811702 | 0.87929304 |
| Ccdc40 | 23.0620372 | -0.4697511 | 1.19791832 | -0.3921395 | 0.69495516 | 0.92609254 |
| Zfp292 | 427.33141 | -0.3513867 | 0.25258724 | -1.3911497 | 0.16418004 | 0.59585062 |
| Rsbn1l | 255.123899 | 0.25279024 | 0.44707837 | 0.56542713 | 0.57178327 | 0.88035336 |
| Tbc1d16 | 356.24664 | -0.2917895 | 0.47365348 | -0.6160399 | 0.53786815 | 0.86655355 |
| Zc3h12d | 137.650504 | -0.4629644 | 0.75217867 | -0.6154978 | 0.53822599 | 0.86663962 |
| Dtx4 | 1207.78949 | -0.0760877 | 0.20788465 | -0.3660091 | 0.71435828 | 0.93235217 |
| Ccdc32 | 860.370714 | 0.47440072 | 0.39041971 | 1.21510442 | 0.22432627 | 0.65793926 |
| Sinhcaf | 4179.38915 | -0.1718162 | 0.28476362 | -0.6033645 | 0.54626627 | 0.86994632 |
| Phtf2 | 444.658031 | -0.0735179 | 0.37021883 | -0.1985795 | 0.84259171 | 0.96765585 |
| Ankrd13c | 913.590393 | 0.03917533 | 0.2968354 | 0.13197662 | 0.89500279 | 0.9794437 |
| Cbx4 | 75.036766 | -1.0186763 | 0.85696714 | -1.1886994 | 0.23455798 | 0.67038085 |
| Edrf1 | 559.309061 | -0.2904878 | 0.29302833 | -0.99133 | 0.32152446 | 0.74447686 |
| Timeless | 2315.41832 | -0.1709303 | 0.24547393 | -0.6963277 | 0.4862236 | 0.84183502 |
| Ifi203 | 43.8052201 | -1.3320674 | 0.89378952 | -1.4903591 | 0.13612983 | 0.55833898 |
| Ginm1 | 612.286776 | -0.1449709 | 0.23377098 | -0.6201405 | 0.53516527 | 0.86539693 |
| Bahd1 | 254.872477 | -0.5527878 | 0.49024962 | -1.1275639 | 0.25950412 | 0.69591124 |
| Slc7a5 | 318.735801 | -1.175059 | 0.52480848 | -2.2390245 | 0.02515432 | 0.23084234 |
| Fkbp6 | 14.7747132 | -1.6033664 | 1.23315462 | -1.3002152 | 0.19352722 | 0.62506533 |
| Ptger3 | 570.432064 | -1.2291367 | 0.25426466 | -4.8340836 | 1.337603378 | 9.350229790 |
| Cox15 | 2030.27132 | -0.0431781 | 0.24483376 | -0.1763567 | 0.86001374 | 0.96954641 |
| Lats1 | 157.099511 | 0.15633113 | 0.4602409 | 0.3396724 | 0.73410325 | 0.93750666 |
| Rab11fip2 | 21.6093934 | -0.8363559 | 1.77239004 | -0.4718803 | 0.63701223 | 0.90565274 |
| Ythdf2 | 7600.15703 | -0.1240067 | 0.15039437 | -0.8245437 | 0.40963068 | 0.80100711 |
| Elavl1 | 8393.27099 | -0.3884542 | 0.18945793 | -2.0503452 | 0.04033076 | 0.3051541 |
| Ipo8 | 395.656528 | -0.3138861 | 0.32515409 | -0.9653457 | 0.33437171 | 0.75342957 |
| Stat2 | 606.718535 | -0.0667604 | 0.34660696 | -0.1926112 | 0.84726346 | 0.96765585 |
| Nup43 | 2566.87833 | 0.21613638 | 0.35055162 | 0.61656078 | 0.53752447 | 0.86634214 |
| Ift88 | 254.732399 | 0.50275101 | 0.40928597 | 1.22836122 | 0.2193114 | 0.65275823 |
| Rbms2 | 395.375105 | 0.49208109 | 0.41318578 | 1.19094393 | 0.2336756 | 0.66954422 |
| Orc3 | 4195.65602 | -0.1378232 | 0.16682159 | -0.8261714 | 0.40870686 | 0.80100711 |
| Ndufb10 | 4038.15335 | 0.03500586 | 0.29936151 | 0.11693508 | 0.9069115 | 0.98176897 |
| Baz2a | 456.000083 | -0.2292827 | 0.39943971 | -0.5740107 | 0.56596056 | 0.878492 |
| Plcb2 | 624.503569 | -0.3675672 | 0.31473625 | -1.1678577 | 0.24286416 | 0.67814218 |
| Ptges3-ps | 220.860876 | 0.27204028 | 0.71240201 | 0.38186343 | 0.70256267 | 0.92762537 |
| Bub1b | 6385.71627 | -0.2809971 | 0.25507418 | -1.1016289 | 0.27062303 | 0.70756376 |
| Bmf | 35.5089081 | -0.1594817 | 1.38890977 | -0.1148251 | 0.90858374 | 0.98195114 |
| Flywch1 | 742.987779 | 0.13538962 | 0.37301892 | 0.36295644 | 0.71663742 | 0.93235217 |
| Klhl42 | 277.588042 | -0.7087699 | 0.59977712 | -1.1817221 | 0.23731598 | 0.67356993 |
| Plpp6 | 65.2789089 | 0.59743431 | 0.65806361 | 0.9078671 | 0.36394844 | 0.77045439 |
| Gramd1b | 219.630345 | 0.62352096 | 0.47910054 | 1.30144075 | 0.19310763 | 0.624687 |
| Mrps35 | 1777.96342 | -0.054508 | 0.31722374 | -0.1718282 | 0.86357257 | 0.9705375 |
| Zmym5 | 438.972603 | 0.26313507 | 0.36632164 | 0.71831703 | 0.47256183 | 0.83483892 |

| | | | | | | |
|---|---|---|---|---|---|---|
| Gorab | 422.47165 | 0.43870258 | 0.34529847 | 1.27050255 | 0.20390568 | 0.63827486 |
| Pnrc1 | 1306.22305 | 0.58837131 | 0.65014922 | 0.90497887 | 0.36547658 | 0.77150562 |
| Gpr176 | 14.1179246 | -0.7835413 | 2.0329534 | -0.3854202 | 0.69992611 | 0.92669304 |
| 9430038I01F | 495.002872 | 0.68546293 | 0.38204646 | 1.79418738 | 0.07278326 | 0.41489174 |
| Maob | 50.0260935 | -1.7040271 | 1.24610404 | -1.3674838 | 0.17147371 | 0.60430173 |
| Hs2st1 | 213.279543 | -0.5120465 | 0.69812383 | -0.7334608 | 0.46327742 | 0.83179129 |
| Thbs1 | 866.008713 | -2.6718475 | 0.49926636 | -5.3515472 | 8.720537023 | 8.603091080 |
| Tax1bp3 | 3736.86971 | 0.85986387 | 0.19149274 | 4.49032095 | 7.111593271 | 0.00039012 |
| Ikzf5 | 639.00605 | 0.36036848 | 0.27538116 | 1.30861707 | 0.1906641 | 0.62164016 |
| Alkbh3 | 1963.6033 | 0.35561051 | 0.30966452 | 1.14837343 | 0.25081444 | 0.6842534 |
| 2310057M21 | 596.395542 | -0.3195769 | 0.24104314 | -1.3258079 | 0.18490333 | 0.61700012 |
| Fmo1 | 57.4209592 | 1.63069191 | 0.92256334 | 1.76756634 | 0.07713343 | 0.42909197 |
| Ankrd6 | 15.5588931 | -0.3361671 | 1.72606515 | -0.1947592 | 0.84558147 | 0.96765585 |
| Arntl2 | 38.4615668 | -0.2852265 | 0.94442025 | -0.3020122 | 0.76264274 | 0.94298237 |
| Scamp2 | 7540.34435 | 0.32357176 | 0.24083002 | 1.34356903 | 0.17908779 | 0.60992118 |
| Ccdc114 | 50.1744847 | 1.24680968 | 0.82870667 | 1.50452474 | 0.13244631 | 0.55219606 |
| Nemp1 | 399.949416 | 0.36747333 | 0.33227484 | 1.10593185 | 0.26875602 | 0.70566481 |
| Pclaf | 13856.1053 | 0.13260472 | 0.27435453 | 0.48333346 | 0.62885899 | 0.90276005 |
| Zfp704 | 161.40299 | 0.49698316 | 0.47048333 | 1.0563247 | 0.29081989 | 0.72470967 |
| Emp3 | 2636.86718 | -0.0095762 | 0.22641982 | -0.0422941 | 0.96626425 | 0.99443087 |
| Kyat3 | 667.158256 | -0.2247106 | 0.33603427 | -0.6687134 | 0.50367829 | 0.85012672 |
| Ttc12 | 64.1542738 | 0.09027497 | 0.70072179 | 0.12883141 | 0.89749105 | 0.97984623 |
| Gas8 | 563.285071 | 0.60149102 | 0.29080745 | 2.06834808 | 0.0386073 | 0.29684673 |
| Prrc2c | 3015.2747 | -0.2753966 | 0.24854338 | -1.1080424 | 0.26784351 | 0.70441538 |
| Gpr34 | 2469.82816 | 1.16087573 | 0.36378754 | 3.1910816 | 0.00141741 | 0.03015514 |
| Syngr4 | 15.0439496 | 0.93254624 | 0.95350896 | 0.97801519 | 0.32806681 | 0.74843736 |
| Tm7sf3 | 2452.93431 | -0.4909094 | 0.23269067 | -2.109708 | 0.03488351 | 0.28000657 |
| Trappc5 | 1789.03413 | -0.719773 | 0.20651463 | -3.4853367 | 0.00049152 | 0.01324392 |
| Fgfr1op2 | 3766.16916 | 0.13741434 | 0.26633453 | 0.51594639 | 0.60589185 | 0.89258167 |
| Tbc1d10c | 146.870814 | -1.0862943 | 0.57648982 | -1.8843252 | 0.059521 | 0.37661155 |
| Lrp1 | 1663.00914 | 0.39875736 | 0.29130842 | 1.36884941 | 0.17104635 | 0.60430173 |
| Ints13 | 1564.1669 | -0.4877296 | 0.26484947 | -1.8415351 | 0.06554319 | 0.39558985 |
| Gbp7 | 4045.91518 | 0.09371855 | 0.24006496 | 0.39038831 | 0.69624943 | 0.92638887 |
| Klhdc4 | 2652.55164 | -0.2323069 | 0.16961197 | -1.3696375 | 0.1708001 | 0.60430173 |
| Plekha1 | 909.962465 | 0.15989695 | 0.26289209 | 0.60822276 | 0.54303974 | 0.86919453 |
| Mrps28 | 2132.90403 | 0.20501893 | 0.35443285 | 0.57844223 | 0.56296559 | 0.87595498 |
| Bach2 | 50.0599027 | 2.02155914 | 1.56704562 | 1.29004486 | 0.19703508 | 0.62932233 |
| Accs | 465.24107 | 0.3655573 | 0.27122699 | 1.34779103 | 0.1777256 | 0.60906934 |
| Cdk6 | 3638.92422 | 0.01603554 | 0.26883393 | 0.05964848 | 0.95243561 | 0.9914597 |
| BC052040 | 631.569323 | 0.11755269 | 0.28677142 | 0.40991774 | 0.68186629 | 0.92061965 |
| Hey1 | 8.77313701 | 0.34118606 | 1.84493765 | 0.18493094 | 0.8532832 | 0.96811477 |
| Ddx58 | 179.683218 | 0.4714975 | 0.50727696 | 0.92946761 | 0.35264681 | 0.76366231 |
| Suco | 279.375577 | -0.1202184 | 0.49877705 | -0.2410263 | 0.8095347 | 0.95844313 |
| Rbm48 | 468.164999 | -0.3238218 | 0.45476764 | -0.7120598 | 0.47642772 | 0.83653513 |
| Cchcr1 | 187.069956 | 0.23137196 | 0.49363866 | 0.46870713 | 0.63927899 | 0.90632643 |
| Zfp770 | 62.05707 | -0.2159018 | 0.62821635 | -0.3436743 | 0.73109123 | 0.93653187 |

| | | | | | | |
|---|---|---|---|---|---|---|
| Slc25a24 | 1015.64575 | 0.47544428 | 0.22501393 | 2.1129549 | 0.03460463 | 0.27887708 |
| Gm15429 | 301.728332 | -0.4942756 | 0.33917546 | -1.4572859 | 0.14503748 | 0.57067981 |
| Dcaf1 | 373.616518 | -0.2412672 | 0.3115599 | -0.7743846 | 0.4387034 | 0.8193372 |
| Cul9 | 95.1401424 | 1.19507294 | 0.66255035 | 1.8037466 | 0.07127104 | 0.41122734 |
| Nsmce4a | 386.642337 | 0.2304078 | 0.42941715 | 0.53655938 | 0.59157201 | 0.88797037 |
| Fam102b | 812.366707 | 0.75634899 | 0.26943842 | 2.80713121 | 0.00499849 | 0.07982572 |
| Arhgap9 | 5092.82596 | -0.2214109 | 0.18660668 | -1.1865109 | 0.23542061 | 0.67164041 |
| Ankib1 | 234.100587 | -0.5628009 | 0.3155185 | -1.7837334 | 0.07446699 | 0.42057003 |
| Mars | 8206.3522 | -0.240325 | 0.15869812 | -1.5143532 | 0.12993629 | 0.5467873 |
| Skiv2l | 1043.92043 | -0.3867106 | 0.29718259 | -1.3012559 | 0.19317089 | 0.624687 |
| Ufl1 | 1996.84614 | -0.0180716 | 0.20044541 | -0.0901574 | 0.92816216 | 0.98666423 |
| Bcor | 266.073736 | 0.04344841 | 0.29160148 | 0.14899927 | 0.88155421 | 0.97582756 |
| Trim41 | 259.43534 | 0.48795195 | 0.26287928 | 1.85618264 | 0.06342753 | 0.3885373 |
| Etfrf1 | 407.496281 | -0.0634146 | 0.32212683 | -0.1968623 | 0.84393528 | 0.96765585 |
| Pex2 | 2224.39024 | 0.33100016 | 0.2433107 | 1.36040119 | 0.173703 | 0.60653633 |
| Aqr | 1529.9257 | -0.3444224 | 0.33507053 | -1.0279101 | 0.30399211 | 0.73436097 |
| Ppp1ca | 41582.8888 | 0.34709063 | 0.33144217 | 1.04721323 | 0.29500124 | 0.72771732 |
| Wdr47 | 258.791144 | 0.29982739 | 0.38614102 | 0.77647123 | 0.4374708 | 0.8185065 |
| Map3k10 | 19.3245277 | -0.6718118 | 1.40089246 | -0.4795598 | 0.63154041 | 0.90399255 |
| Abhd13 | 201.422673 | 0.01068561 | 0.57122015 | 0.01870663 | 0.98507514 | 0.99564261 |
| Akap9 | 534.417877 | -0.2392871 | 0.51208981 | -0.4672756 | 0.64030268 | 0.90677142 |
| Fbxl4 | 463.797487 | 0.42906494 | 0.3984581 | 1.07681319 | 0.28156373 | 0.7168302 |
| Timd2 | 113.253487 | -3.7190739 | 0.95601089 | -3.8902003 | 0.00010016 | 0.00361438 |
| Slc25a28 | 40.5993877 | 0.31105249 | 0.82119649 | 0.37877961 | 0.70485153 | 0.92874286 |
| Dtx3 | 65.1906625 | 0.02436091 | 0.55552919 | 0.04385172 | 0.9650226 | 0.99428295 |
| Rc3h1 | 1454.75897 | 0.20550382 | 0.47889431 | 0.42912146 | 0.66783484 | 0.91595755 |
| Plekha4 | 10.4849983 | 0.82975411 | 2.10997846 | 0.39325241 | 0.69413307 | 0.92609254 |
| Mterf1a | 124.058109 | -0.6732725 | 0.47401625 | -1.4203574 | 0.15550367 | 0.58452503 |
| Pitpnc1 | 762.184234 | 0.73383929 | 0.42470909 | 1.7278634 | 0.08401272 | 0.44471119 |
| Zbtb38 | 1270.15693 | 0.26663924 | 0.32334625 | 0.8246245 | 0.40958481 | 0.80100711 |
| Ppp1r15a | 3827.79433 | 0.64436817 | 0.27687628 | 2.32727831 | 0.01995046 | 0.19988038 |
| Slc26a10 | 22.5883655 | -2.4486026 | 1.03009689 | -2.3770605 | 0.01745123 | 0.18560042 |
| Rprd1a | 785.587715 | 0.41454816 | 0.3075465 | 1.34792029 | 0.17768402 | 0.60906934 |
| Spns2 | 35.8233598 | 1.98202121 | 1.5655883 | 1.26599133 | 0.20551619 | 0.6393895 |
| Sgms1 | 976.327239 | -0.3831559 | 0.38263337 | -1.0013655 | 0.31665012 | 0.74186126 |
| Usp45 | 220.481018 | -0.1362011 | 0.30884024 | -0.4410082 | 0.65920708 | 0.91280998 |
| Arglu1 | 2416.37001 | -0.1670299 | 0.25243552 | -0.6616736 | 0.50818043 | 0.85238774 |
| Os9 | 2283.83939 | -0.1759767 | 0.23885556 | -0.7367496 | 0.46127463 | 0.83018576 |
| Mybbp1a | 10682.5042 | -0.6110131 | 0.193923 | -3.1508028 | 0.00162822 | 0.03404797 |
| Gtpbp10 | 774.362457 | -0.0061316 | 0.27180949 | -0.0225585 | 0.98200243 | 0.99523682 |
| Blvrb | 1948.59923 | 2.20305055 | 0.41752056 | 5.27650798 | 1.316687802 | 1.239003222 |
| Rabggta | 1427.16859 | -0.351223 | 0.26407988 | -1.3299876 | 0.18352235 | 0.61524496 |
| Dgkz | 3344.57394 | 0.06171641 | 0.21143118 | 0.29189833 | 0.77036436 | 0.94454775 |
| Bptf | 682.295177 | -0.3064548 | 0.39502661 | -0.7757827 | 0.43787727 | 0.81866921 |
| Dxo | 908.167648 | 0.21520981 | 0.24158555 | 0.89082237 | 0.37302447 | 0.77748705 |
| Xaf1 | 225.842708 | 0.01824431 | 0.59954459 | 0.03043028 | 0.97572389 | 0.99458632 |

| | | | | | | |
|---|---|---|---|---|---|---|
| Ambra1 | 319.171108 | 0.11251887 | 0.34017043 | 0.33077205 | 0.74081667 | 0.93800224 |
| Pvr | 297.003474 | 0.01498485 | 0.48018969 | 0.0312061 | 0.97510518 | 0.99450266 |
| Manea | 295.045771 | 1.03255601 | 0.34822064 | 2.96523498 | 0.00302452 | 0.05433029 |
| Tsfm | 1707.63483 | -0.3219805 | 0.20728813 | -1.5532992 | 0.12035167 | 0.52825513 |
| Tlr8 | 176.13423 | -0.3798176 | 0.78596944 | -0.4832473 | 0.62892016 | 0.90276005 |
| Zfp609 | 49.5382971 | -0.5706638 | 1.22399904 | -0.4662289 | 0.64105165 | 0.90685007 |
| Milr1 | 5154.71326 | -0.1602322 | 0.32681488 | -0.4902843 | 0.62393273 | 0.90017327 |
| Abhd11 | 1910.19003 | -0.2255174 | 0.2050494 | -1.0998197 | 0.27141067 | 0.70898285 |
| Adam22 | 17.2995523 | 0.12300257 | 1.04019648 | 0.11824936 | 0.90587008 | 0.98144617 |
| Tex2 | 524.488189 | -0.1094473 | 0.2702145 | -0.4050385 | 0.68544923 | 0.92314423 |
| Ckap5 | 1341.35069 | 0.02008641 | 0.3433322 | 0.05850429 | 0.95334695 | 0.99149988 |
| Otud6b | 3604.93934 | -0.2020427 | 0.19808082 | -1.0200014 | 0.30772777 | 0.73711256 |
| C3ar1 | 19645.5915 | 0.8726152 | 0.25896415 | 3.36963711 | 0.00075267 | 0.01871431 |
| Aipl1 | 35.3908865 | 0.90898464 | 1.29301382 | 0.70299685 | 0.48205771 | 0.84038934 |
| Mettl27 | 88.2350383 | 1.19633545 | 0.68742993 | 1.74030166 | 0.08180606 | 0.44065767 |
| Wdr7 | 480.120384 | -0.4479068 | 0.31680666 | -1.4138175 | 0.15741548 | 0.58620126 |
| Gstm2 | 42.0771534 | 2.37004101 | 1.20623116 | 1.96483152 | 0.04943374 | 0.34140219 |
| Plppr2 | 20.2218231 | -0.4957559 | 1.08768106 | -0.4557916 | 0.64853986 | 0.90973357 |
| Apoc1 | 343.416002 | -0.2410856 | 0.51976195 | -0.4638385 | 0.64276346 | 0.90743513 |
| Btaf1 | 755.774489 | -0.3037928 | 0.4394584 | -0.6912891 | 0.48938387 | 0.84401986 |
| Ofd1 | 301.42401 | 0.33531515 | 0.42533541 | 0.78835465 | 0.43048929 | 0.81243065 |
| 1110051M20 | 405.999224 | 0.74454036 | 0.39676389 | 1.87653258 | 0.06058219 | 0.37940675 |
| Cd79b | 2997.98494 | 0.16351078 | 0.22208756 | 0.73624467 | 0.46158181 | 0.83025001 |
| Ranbp17 | 188.100303 | -0.0603114 | 0.45148712 | -0.133584 | 0.89373156 | 0.97941957 |
| Pogk | 183.418733 | 1.21297263 | 0.63698109 | 1.90425218 | 0.05687735 | 0.36677945 |
| Mis12 | 2994.09105 | 0.15286769 | 0.18430467 | 0.82942929 | 0.40686153 | 0.79992362 |
| Kazn | 19.175353 | -1.087295 | 1.34075048 | -0.81096 | 0.41738864 | 0.8049685 |
| Apobec1 | 3362.31441 | 0.228985 | 0.1693053 | 1.35249756 | 0.1762162 | 0.60763832 |
| Tmem51 | 611.902609 | 0.9518176 | 0.25787028 | 3.6910714 | 0.00022331 | 0.00710571 |
| Pck2 | 572.888364 | 0.13684563 | 0.35128004 | 0.38956278 | 0.69685988 | 0.92649569 |
| Dhx33 | 683.257595 | -0.2786186 | 0.33715824 | -0.8263735 | 0.40859226 | 0.80100711 |
| Gemin8 | 346.645987 | -0.0687051 | 0.40340908 | -0.1703112 | 0.86476543 | 0.9705375 |
| Plekhg1 | 22.5769954 | 3.01622687 | 1.90124389 | 1.58644921 | 0.11263744 | 0.51139203 |
| Dok4 | 163.43079 | 1.5796984 | 0.45707546 | 3.45609979 | 0.00054805 | 0.01441499 |
| Ppip5k2 | 694.764169 | -0.0313012 | 0.30439376 | -0.1028312 | 0.91809691 | 0.98390186 |
| Tasor | 194.949968 | 0.54294836 | 0.57105794 | 0.95077632 | 0.34171793 | 0.75866343 |
| Oaz2 | 3686.01839 | 0.3381214 | 0.20295103 | 1.66602456 | 0.09570852 | 0.47439317 |
| Ppp1r14c | 19.3207796 | 1.265694 | 1.61043055 | 0.78593517 | 0.43190547 | 0.81344118 |
| Dnph1 | 1106.30908 | 0.03099933 | 0.35265593 | 0.08790249 | 0.92995418 | 0.98699718 |
| Efhd2 | 5089.14337 | 0.41401514 | 0.40821809 | 1.01420088 | 0.31048691 | 0.7399039 |
| Rad54l2 | 252.720228 | 0.38885602 | 0.52055727 | 0.7469995 | 0.45506386 | 0.82777639 |
| Nup88 | 7510.69791 | -0.1645061 | 0.17817096 | -0.9233046 | 0.35584851 | 0.76499791 |
| Phc1 | 120.431441 | -0.0319682 | 0.51424837 | -0.0621648 | 0.95043158 | 0.9907652 |
| Mthfd1l | 3522.97817 | -0.3027894 | 0.21217418 | -1.4270794 | 0.15355702 | 0.58242478 |
| Hmgn1 | 16184.7017 | 0.0420291 | 0.21265339 | 0.19764132 | 0.84332571 | 0.96765585 |
| Madd | 723.167257 | -0.3464886 | 0.30779484 | -1.125713 | 0.26028704 | 0.69647146 |

| | | | | | | |
|---|---|---|---|---|---|---|
| Tbl3 | 5594.93007 | -0.3616852 | 0.17277676 | -2.0933672 | 0.03631638 | 0.28588052 |
| Apobec2 | 150.346664 | 0.73364086 | 0.41998065 | 1.74684445 | 0.08066432 | 0.43854735 |
| Dnajc16 | 487.287891 | 0.53169843 | 0.27552667 | 1.92975303 | 0.05363745 | 0.35874624 |
| Limd2 | 16840.3239 | -0.1938447 | 0.13542193 | -1.4314127 | 0.15231196 | 0.58026542 |
| Ap1g2 | 2664.33609 | -0.0464097 | 0.1848899 | -0.2510126 | 0.80180436 | 0.95629571 |
| St8sia4 | 1120.57337 | -0.8165136 | 0.23825424 | -3.4270684 | 0.00061014 | 0.01584668 |
| Sh3pxd2b | 185.210404 | -0.2436833 | 0.54751261 | -0.4450734 | 0.65626669 | 0.91176722 |
| Camta2 | 377.602918 | 0.42637769 | 0.40107246 | 1.06309391 | 0.28773936 | 0.72300867 |
| Creg1 | 6588.23594 | 0.01472194 | 0.36864367 | 0.03993543 | 0.9681446 | 0.99443087 |
| Rsc1a1 | 135.793294 | 0.06077532 | 0.51206252 | 0.11868731 | 0.90552309 | 0.98144617 |
| Virma | 970.68822 | -0.3554985 | 0.2091268 | -1.6999184 | 0.08914628 | 0.4566019 |
| Zfhx2 | 17.6667562 | 3.23092437 | 1.59820909 | 2.02159054 | 0.04321867 | 0.31923815 |
| Scamp5 | 2163.20237 | 0.57179508 | 0.21530621 | 2.65572963 | 0.0079137 | 0.1098846 |
| Rcsd1 | 3614.38822 | -0.0072014 | 0.21536791 | -0.0334378 | 0.97332545 | 0.99443087 |
| Hnrnpul1 | 681.276867 | -0.2043546 | 0.36292427 | -0.5630779 | 0.57338181 | 0.88046468 |
| Hesx1 | 15.4840754 | -0.1785517 | 1.34905378 | -0.1323533 | 0.89470485 | 0.97941957 |
| Eif4h | 17428.7022 | -0.3390813 | 0.20436203 | -1.6592187 | 0.09707174 | 0.47799662 |
| Erg | 10.8316406 | 2.0705517 | 2.14623435 | 0.964737 | 0.33467656 | 0.75351065 |
| Ints8 | 459.72192 | -1.2633448 | 0.54137438 | -2.3335881 | 0.0196173 | 0.19849335 |
| Rnf167 | 912.652852 | -0.1999075 | 0.26901571 | -0.7431071 | 0.45741682 | 0.82946634 |
| Cd53 | 18378.4745 | 0.0539733 | 0.28022278 | 0.19260855 | 0.84726555 | 0.96765585 |
| Siah1b | 736.515503 | -0.1181684 | 0.25922443 | -0.4558535 | 0.64849534 | 0.90973357 |
| Lat2 | 802.044652 | -1.7516701 | 0.47774588 | -3.6665311 | 0.00024586 | 0.00759506 |
| Appl1 | 1113.03303 | 0.18577767 | 0.2711781 | 0.68507623 | 0.49329582 | 0.84635172 |
| Spen | 84.1546466 | -0.3313369 | 0.6011116 | -0.5512069 | 0.58149184 | 0.88321857 |
| Snrnp25 | 2952.39874 | 0.7269533 | 0.48333967 | 1.50402159 | 0.13257581 | 0.55219606 |
| Oard1 | 1710.66366 | 0.14507406 | 0.30484045 | 0.47590159 | 0.63414449 | 0.9046635 |
| Cept1 | 4154.53772 | -0.083841 | 0.14958714 | -0.5604828 | 0.57515019 | 0.88059206 |
| Cop1 | 385.650182 | -0.2529323 | 0.38587423 | -0.6554786 | 0.51215963 | 0.85399887 |
| Ttc3 | 1555.2368 | 0.05342157 | 0.23626111 | 0.22611242 | 0.82111398 | 0.96205421 |
| Iqcc | 539.288324 | 0.15246392 | 0.41951032 | 0.36343307 | 0.7162814 | 0.93235217 |
| Eml2 | 21.8594757 | -0.1486493 | 0.94470544 | -0.1573499 | 0.87496908 | 0.97366476 |
| Tex264 | 1076.3894 | 0.56976174 | 0.20456142 | 2.78528442 | 0.00534808 | 0.08409132 |
| Dennd6a | 400.813217 | 0.32956724 | 0.34128109 | 0.96567683 | 0.33420592 | 0.75320272 |
| Hlcs | 178.903215 | 0.41617789 | 0.42593361 | 0.97709568 | 0.32852178 | 0.74843736 |
| 1700123O20 | 2019.15884 | 0.34035727 | 0.34516173 | 0.98608055 | 0.32409359 | 0.74593356 |
| Snrpd2 | 3450.7495 | 0.15720103 | 0.39848449 | 0.39449723 | 0.69321397 | 0.92577363 |
| Szrd1 | 1668.81804 | 0.14881847 | 0.31777301 | 0.46831689 | 0.63955799 | 0.90646657 |
| Tiprl | 2622.94425 | 0.25985764 | 0.19530903 | 1.3304947 | 0.18335533 | 0.61524496 |
| Sft2d2 | 728.882226 | 0.51724022 | 0.18771061 | 2.75551941 | 0.0058599 | 0.09016879 |
| Psme4 | 595.084902 | -0.4135174 | 0.35807892 | -1.1548219 | 0.24816337 | 0.68235696 |
| Reps2 | 137.651081 | 2.24616733 | 0.56846817 | 3.95126313 | 7.773979345 | 0.00289936 |
| Erf | 56.1465277 | 0.98089523 | 0.69054396 | 1.42046747 | 0.15547163 | 0.58452503 |
| Bsdc1 | 806.032318 | 0.06001863 | 0.30949442 | 0.19392477 | 0.84623479 | 0.96765585 |
| Crocc | 48.918885 | 0.02766457 | 1.05013239 | 0.02634389 | 0.97898305 | 0.99483019 |
| Ino80d | 183.779073 | 0.05534408 | 0.55752584 | 0.09926729 | 0.92092605 | 0.9839023 |

| | | | | | | |
|---|---|---|---|---|---|---|
| Wdr25 | 228.043737 | -0.3629044 | 0.42293088 | -0.8580702 | 0.39085368 | 0.7899826 |
| Tmem205 | 353.320257 | 0.49354868 | 0.36010382 | 1.3705733 | 0.17050801 | 0.60430173 |
| Gfer | 1220.0416 | 0.01933611 | 0.23709234 | 0.08155517 | 0.93500045 | 0.98789605 |
| Gm21988 | 643.89398 | 0.5269503 | 0.22595426 | 2.33211046 | 0.01969488 | 0.1989492 |
| Atp1a3 | 646.986457 | -1.9086564 | 0.33270698 | -5.7367491 | 9.651105144 | 1.377785152 |
| Fbxw4 | 179.83246 | -0.4825269 | 0.39681654 | -1.215995 | 0.22398681 | 0.65774578 |
| Slc19a2 | 592.23788 | 0.15999013 | 0.28110612 | 0.56914496 | 0.56925777 | 0.87961466 |
| S100pbp | 232.607646 | -0.3593215 | 0.32120337 | -1.1186728 | 0.26327975 | 0.69969611 |
| Rfx3 | 105.429188 | -1.3756487 | 0.70031866 | -1.9643183 | 0.04949319 | 0.34145768 |
| Ulk4 | 36.8929148 | 3.43539251 | 1.25688293 | 2.73326371 | 0.00627101 | 0.09487653 |
| Arhgef1 | 3711.82599 | -0.2985261 | 0.29599939 | -1.0085362 | 0.3131971 | 0.74065408 |
| Tet2 | 553.107899 | 0.18187842 | 0.42653992 | 0.42640422 | 0.66981332 | 0.91689368 |
| Rcc2 | 2479.19133 | -0.5148207 | 0.28214049 | -1.8246964 | 0.06804686 | 0.40193976 |
| Mgl2 | 107.896111 | -1.6239531 | 0.62645728 | -2.5922807 | 0.00953419 | 0.12534914 |
| Rps19 | 37229.3785 | 0.41809451 | 0.416149 | 1.00467503 | 0.31505336 | 0.74123109 |
| Cables1 | 140.078113 | 1.52784591 | 0.64201699 | 2.37975931 | 0.01732395 | 0.18521003 |
| Asgr2 | 6.58648103 | 3.18269476 | 2.0699311 | 1.53758488 | 0.12415015 | 0.53551789 |
| Arhgef10l | 2248.79164 | -0.3797557 | 0.30480643 | -1.2458914 | 0.21280428 | 0.64596443 |
| Gm11992 | 8.22651759 | 1.42674987 | 2.3309014 | 0.6121022 | 0.54047014 | 0.86741192 |
| Sh3kbp1 | 3577.56299 | -0.407667 | 0.28215031 | -1.4448575 | 0.14849792 | 0.5754118 |
| Abhd4 | 1050.55475 | 1.02432147 | 0.30014547 | 3.41275006 | 0.00064311 | 0.01659737 |
| Npnt | 15.3508963 | -0.7693505 | 2.0010322 | -0.3844768 | 0.70062505 | 0.92706828 |
| Trim62 | 46.7014185 | 0.76571198 | 0.73817919 | 1.03729826 | 0.29959688 | 0.73123875 |
| Cmtm8 | 82.1773429 | 0.37137262 | 0.66489204 | 0.55854574 | 0.57647178 | 0.88105119 |
| Iffo2 | 148.973029 | -0.2734692 | 0.40468447 | -0.675759 | 0.49919367 | 0.84872332 |
| Ghitm | 17067.206 | 0.29470024 | 0.26418477 | 1.11550807 | 0.26463276 | 0.70112124 |
| Gm17018 | 1073.65119 | -0.151314 | 0.20502709 | -0.7380193 | 0.46050272 | 0.82987212 |
| Irgq | 104.98262 | 0.94798216 | 0.68012559 | 1.39383399 | 0.16336774 | 0.59475807 |
| Fam117b | 262.983195 | -0.778734 | 0.29024315 | -2.6830401 | 0.00729562 | 0.10438985 |
| Slc7a13 | 54.8147532 | 3.97375259 | 1.56359941 | 2.54141346 | 0.01104053 | 0.13649787 |
| Wdr43 | 7951.95476 | -0.2819001 | 0.249812 | -1.1284489 | 0.25913037 | 0.69546977 |
| Wwp1 | 1287.4745 | 0.90934783 | 0.2964096 | 3.06787574 | 0.00215586 | 0.04233173 |
| Pif1 | 550.542459 | -0.2199179 | 0.2849718 | -0.771718 | 0.44028148 | 0.81963526 |
| Fzd7 | 151.569725 | 0.3161076 | 0.62917294 | 0.50241766 | 0.61537376 | 0.8987425 |
| Rwdd2b | 924.567381 | 0.10806659 | 0.33180483 | 0.32569325 | 0.74465645 | 0.93917909 |
| Ostc | 8608.1619 | 0.30727992 | 0.31460544 | 0.97671522 | 0.32871016 | 0.74843736 |
| Tspyl2 | 199.33834 | -0.1503626 | 0.43182015 | -0.3482066 | 0.72768507 | 0.93576253 |
| Elmo1 | 5726.32404 | 0.00664736 | 0.28806237 | 0.02307613 | 0.98158955 | 0.99523682 |
| Iqsec2 | 63.8808037 | 2.06479401 | 0.77622668 | 2.66004 | 0.00781314 | 0.10907646 |
| Ccdc8 | 166.594986 | 0.5397945 | 0.3648471 | 1.47950884 | 0.13900437 | 0.56272724 |
| Pde9a | 18.3143238 | 1.95365565 | 2.06190592 | 0.9474999 | 0.34338411 | 0.75947281 |
| Nbl1 | 7.47379255 | -1.5430766 | 2.73533626 | -0.5641268 | 0.57266782 | 0.88035336 |
| Msantd4 | 1097.45743 | 0.49991519 | 0.25536494 | 1.95765009 | 0.05027109 | 0.34299029 |
| H2afv | 1832.70871 | 0.55802905 | 0.37758991 | 1.47787066 | 0.1394424 | 0.56329174 |
| Zfp598 | 87.7656869 | -0.4046937 | 0.521344 | -0.7762507 | 0.43760095 | 0.8185065 |
| N4bp2l1 | 687.387702 | 0.08584602 | 0.26030334 | 0.32979223 | 0.74155696 | 0.93800224 |

| | | | | | | |
|---|---|---|---|---|---|---|
| Smc1a | 4833.58683 | -0.223087 | 0.25098916 | -0.8888313 | 0.37409374 | 0.77748705 |
| Ripk2 | 1055.48187 | -0.4811764 | 0.35381713 | -1.3599579 | 0.17384325 | 0.60653633 |
| Tmco4 | 259.855255 | 0.56145556 | 0.46637126 | 1.20388113 | 0.22863553 | 0.66258779 |
| Brca2 | 604.441183 | 0.29897812 | 0.3599714 | 0.83056076 | 0.40622181 | 0.79992362 |
| Osgin2 | 798.691612 | 0.18216953 | 0.30098672 | 0.60524109 | 0.54501882 | 0.86963259 |
| Otud3 | 24.3232542 | 1.17561418 | 1.08022569 | 1.08830422 | 0.27646083 | 0.7118544 |
| Zmiz2 | 333.230066 | -0.414206 | 0.40587533 | -1.0205252 | 0.30747943 | 0.73673815 |
| Lonp1 | 2915.54944 | -0.0489428 | 0.31624286 | -0.1547633 | 0.87700795 | 0.97458574 |
| Hectd2 | 14.9940062 | -0.7067479 | 1.44311725 | -0.489737 | 0.62432001 | 0.90046668 |
| Prkd2 | 322.3224 | -0.1594066 | 0.41691343 | -0.3823495 | 0.70220216 | 0.92762537 |
| Rpusd1 | 570.390094 | -0.0158681 | 0.23412553 | -0.0677759 | 0.94596405 | 0.99019648 |
| Trir | 1042.201 | -0.034367 | 0.36558267 | -0.0940061 | 0.92510432 | 0.98569821 |
| Yeats2 | 318.366727 | -0.0112553 | 0.3088943 | -0.0364374 | 0.97093357 | 0.99443087 |
| Arhgap11a | 1064.98791 | -0.0699343 | 0.22610187 | -0.3093043 | 0.75709002 | 0.94112534 |
| Elovl6 | 1143.92512 | 0.03926337 | 0.32006648 | 0.12267254 | 0.90236641 | 0.98043702 |
| Arhgap12 | 1384.54005 | 0.81093124 | 0.22723133 | 3.56874754 | 0.00035869 | 0.01025038 |
| Phf8 | 312.36992 | -0.9511483 | 0.35310987 | -2.6936327 | 0.0070678 | 0.10256275 |
| Ublcp1 | 713.383084 | 0.28935857 | 0.24390609 | 1.18635236 | 0.23548318 | 0.67164041 |
| Chd7 | 1024.91395 | -0.3559547 | 0.47830636 | -0.7441982 | 0.4567566 | 0.82915295 |
| Vps41 | 3671.70832 | 0.35150427 | 0.17429755 | 2.0166908 | 0.04372779 | 0.32046747 |
| Rbbp8 | 1754.59351 | 0.00010345 | 0.19757984 | 0.00052359 | 0.99958224 | 0.99976341 |
| Mul1 | 756.575836 | 0.41079223 | 0.34498829 | 1.19074254 | 0.23375467 | 0.66954422 |
| Zfp236 | 164.351101 | 0.49227793 | 0.45439111 | 1.08337932 | 0.27864009 | 0.71377266 |
| Car8 | 40.9545315 | 2.65785002 | 1.47454322 | 1.80249041 | NA | NA |
| Rusc1 | 228.246981 | 0.00635768 | 0.42126713 | 0.0150918 | 0.98795894 | 0.9962454 |
| Uspl1 | 2481.12279 | -0.1625032 | 0.19491303 | -0.8337217 | 0.40443784 | 0.79917613 |
| Dmxl2 | 68.6069551 | -0.1640769 | 1.32928109 | -0.1234328 | 0.90176435 | 0.98043702 |
| Tox | 30.6755166 | -4.7461355 | 1.35578647 | -3.5006511 | 0.00046412 | 0.0127016 |
| Ttc1 | 3167.19987 | 0.37039409 | 0.34464421 | 1.07471437 | 0.28250263 | 0.71787592 |
| Cdk13 | 114.898843 | -0.1927491 | 0.59978032 | -0.3213662 | 0.74793286 | 0.9399191 |
| Katnal1 | 121.396631 | 0.3492094 | 0.78031723 | 0.44752235 | 0.65449795 | 0.9108895 |
| Gtf3c3 | 1498.37725 | 0.06916235 | 0.16543055 | 0.41807481 | 0.67589242 | 0.91865406 |
| Sntb2 | 183.727303 | 0.5317741 | 0.82159932 | 0.64724263 | 0.51747493 | 0.85694744 |
| Slc7a1 | 863.955097 | -0.8101091 | 0.37627497 | -2.152971 | 0.03132096 | 0.26460586 |
| Thoc6 | 2276.44092 | 0.20726983 | 0.34513299 | 0.60055062 | 0.54813933 | 0.87059427 |
| Pcf11 | 358.402195 | 0.00924431 | 0.36065073 | 0.02563231 | 0.97955061 | 0.99483957 |
| Atg2b | 238.133824 | -1.0140284 | 0.49520275 | -2.0477035 | 0.04058906 | 0.3064975 |
| Ankrd42 | 35.7137379 | 2.08823334 | 1.78429812 | 1.17033881 | 0.24186465 | 0.67708722 |
| Wrap53 | 696.775565 | -0.1701791 | 0.19197929 | -0.8864451 | 0.37537771 | 0.77815049 |
| Tmem29 | 837.059178 | 0.25830208 | 0.35667578 | 0.72419293 | 0.4689473 | 0.83381643 |
| Rgl2 | 700.476063 | 0.08509634 | 0.2820709 | 0.30168423 | 0.76289279 | 0.94298237 |
| Ssr2 | 9221.41111 | 0.09045811 | 0.16781258 | 0.53904248 | 0.58985754 | 0.88722166 |
| Pum3 | 4818.04267 | -0.491942 | 0.2124456 | -2.315614 | 0.02057936 | 0.20384393 |
| Myzap | 57.2941742 | 1.21259863 | 1.36331002 | 0.88945185 | 0.37376029 | 0.77748705 |
| Shtn1 | 114.926065 | 0.14754609 | 0.47032743 | 0.3137093 | 0.75374184 | 0.94018119 |
| Ccdc9 | 72.5452037 | 1.31549364 | 0.56969102 | 2.30913529 | 0.02093607 | 0.20654113 |

| | | | | | | |
|---|---|---|---|---|---|---|
| Cldn5 | 385.161895 | 3.58786544 | 1.78889144 | 2.00563621 | 0.04489508 | 0.32574227 |
| Mdfic | 60.4219827 | 0.48662413 | 1.31263475 | 0.37072318 | 0.71084372 | 0.93142151 |
| Mettl18 | 195.344645 | 0.75757387 | 0.54413909 | 1.39224306 | 0.16384881 | 0.59565007 |
| BC055324 | 1563.92967 | -0.2022341 | 0.21515027 | -0.9399666 | 0.34723471 | 0.76101142 |
| Wapl | 1959.7127 | -0.1600717 | 0.2529155 | -0.6329058 | 0.52679514 | 0.86119003 |
| Dicer1 | 539.83381 | -0.5543141 | 0.4370964 | -1.2681735 | 0.20473601 | 0.63901202 |
| Pik3r1 | 1390.6052 | 0.000804 | 0.36290271 | 0.00221548 | 0.99823231 | 0.99967651 |
| Meis3 | 88.2252991 | 1.77308736 | 1.09927513 | 1.6129605 | 0.10675309 | 0.49920128 |
| Hibch | 1015.9915 | 0.00209782 | 0.38562822 | 0.00544 | 0.99565953 | 0.99884675 |
| Nthl1 | 241.695312 | -0.0227616 | 0.33096821 | -0.0687729 | 0.9451704 | 0.99019648 |
| Ccnb1 | 18844.957 | 0.05161776 | 0.18017842 | 0.28648136 | 0.77450947 | 0.94709254 |
| Utp4 | 5097.79216 | -0.369677 | 0.20636029 | -1.7914154 | 0.07322667 | 0.41625549 |
| Mfsd6 | 68.428163 | -0.5962377 | 0.97654104 | -0.6105609 | 0.54149034 | 0.86793126 |
| Gk5 | 74.2719086 | 0.10947944 | 0.65454535 | 0.16726028 | 0.86716526 | 0.97107585 |
| Arhgap32 | 18.1116263 | -0.0555669 | 1.605598 | -0.0346082 | 0.97239216 | 0.99443087 |
| Mmrn2 | 62.7254283 | 2.61061888 | 3.73098314 | 0.6997134 | 0.48410631 | 0.84128398 |
| Rpl21 | 15844.2428 | -0.105517 | 0.23835658 | -0.4426854 | 0.65799327 | 0.91258818 |
| Tardbp | 6630.14019 | -0.2137037 | 0.15426968 | -1.3852602 | 0.16597292 | 0.5992759 |
| Shld2 | 295.557582 | 0.59481919 | 0.35254919 | 1.68719487 | 0.09156589 | 0.46305312 |
| Dcp1b | 236.626755 | -0.5755394 | 0.36156782 | -1.5917882 | 0.11143229 | 0.50888686 |
| Serpina3g | 37.0426972 | 0.66309464 | 1.90855033 | 0.34743367 | NA | NA |
| Zfp281 | 920.896655 | -0.2393796 | 0.29832212 | -0.8024198 | 0.42231015 | 0.80821275 |
| Stx3 | 209.243552 | 0.13531935 | 0.39153167 | 0.34561534 | 0.72963182 | 0.93649408 |
| Cep78 | 1295.33968 | -0.1406747 | 0.26006407 | -0.5409233 | 0.58856047 | 0.88690478 |
| Kif14 | 233.803022 | -0.9907402 | 0.37804235 | -2.6207121 | 0.00877463 | 0.11893454 |
| Rrp9 | 2687.36935 | -0.5920593 | 0.24130253 | -2.4535977 | 0.01414351 | 0.16169868 |
| Irf8 | 9973.2806 | -0.4407655 | 0.138407 | -3.1845605 | 0.00144974 | 0.030736 |
| Rnf123 | 510.114024 | -0.4130819 | 0.37069712 | -1.114338 | 0.26513422 | 0.70112124 |
| Ago1 | 237.911028 | 0.82789159 | 0.44018458 | 1.88078278 | 0.06000147 | 0.37827102 |
| Nop53 | 4168.4169 | -0.1861911 | 0.1874857 | -0.9930952 | 0.32066357 | 0.74413737 |
| Camsap2 | 57.3568213 | 1.4999811 | 0.98475094 | 1.52320861 | 0.12770652 | 0.54225404 |
| Selenow | 1863.26757 | 0.38005435 | 0.35113102 | 1.08237189 | 0.27908731 | 0.71377266 |
| Tmtc4 | 352.12293 | 0.29438689 | 0.41218394 | 0.71421243 | 0.47509583 | 0.83587621 |
| Cdc42ep4 | 477.663121 | 0.32324523 | 0.23303557 | 1.38710682 | 0.16540919 | 0.5984068 |
| Mbp | 304.132421 | -0.2471924 | 0.27264905 | -0.9066323 | 0.3646013 | 0.77108653 |
| Bicdl1 | 75.9365578 | 0.43906712 | 0.52438284 | 0.83730261 | 0.40242249 | 0.79795242 |
| D11Wsu47e | 498.833267 | -0.7715377 | 0.35495129 | -2.1736438 | 0.0297319 | 0.25703906 |
| Ggact | 124.918239 | 0.48055496 | 0.4154428 | 1.15672955 | 0.24738287 | 0.68235696 |
| Fam104a | 1922.21902 | 0.19534317 | 0.30372103 | 0.64316644 | 0.52011611 | 0.8576517 |
| Mrps27 | 3543.48188 | -0.2007802 | 0.19273065 | -1.0417657 | 0.29752034 | 0.72979999 |
| Kctd12b | 139.538658 | 1.03242562 | 0.66754226 | 1.54660713 | 0.12195801 | 0.5306488 |
| Gcn1l1 | 1285.10636 | -0.7464315 | 0.34279141 | -2.1775095 | 0.02944258 | 0.2563495 |
| Kif21b | 368.680705 | -0.6499764 | 0.71894349 | -0.9040717 | 0.3659574 | 0.77212089 |
| Ddx24 | 4344.41324 | -0.2847729 | 0.18602629 | -1.5308206 | 0.12581374 | 0.5378361 |
| Klf8 | 69.8271526 | 4.59784281 | 1.23123182 | 3.73434373 | 0.00018821 | 0.00618902 |
| Pcca | 1256.73167 | 0.27148176 | 0.20643465 | 1.31509786 | 0.18847702 | 0.61935434 |

| | | | | | | |
|---|---|---|---|---|---|---|
| Slc39a11 | 1308.02914 | -1.2023676 | 0.27671472 | -4.3451523 | 1.391790669 | 0.00069808 |
| Pyroxd1 | 454.632572 | 0.23794618 | 0.37879886 | 0.6281597 | 0.52989933 | 0.86338019 |
| Lrrc29 | 81.401106 | 1.31687658 | 0.4872522 | 2.70265907 | 0.00687873 | 0.10086772 |
| Bivm | 449.434485 | -0.1077376 | 0.29297187 | -0.3677403 | 0.7130669 | 0.93235217 |
| Fcho2 | 372.856704 | 0.42597354 | 0.34330389 | 1.24080605 | 0.21467741 | 0.64854028 |
| Amot | 32.7269654 | 0.62752294 | 1.16904096 | 0.53678439 | 0.59141656 | 0.88792326 |
| Kcnj2 | 67.5480606 | -0.5774741 | 0.56698502 | -1.0184998 | 0.3084405 | 0.7383723 |
| Cox6a1 | 8632.38342 | 0.04370203 | 0.22001338 | 0.19863353 | 0.84254942 | 0.96765585 |
| Btbd7 | 193.143442 | 0.58685611 | 0.38991482 | 1.50508799 | 0.13230146 | 0.55203111 |
| Tmem273 | 2161.29132 | 0.21592968 | 0.27336439 | 0.78989687 | 0.429588 | 0.81155572 |
| Ubr7 | 2794.73722 | -0.3093654 | 0.18016623 | -1.7171111 | 0.08595887 | 0.44803361 |
| Gm20604 | 746.004727 | -0.0354738 | 0.25055071 | -0.1415831 | 0.8874093 | 0.97832097 |
| Alg13 | 659.637749 | 0.56751238 | 0.36438599 | 1.55744841 | 0.11936406 | 0.5265707 |
| Pi4ka | 556.058181 | -0.2884815 | 0.32768195 | -0.8803706 | 0.3786586 | 0.78126394 |
| Coro2b | 51.2886587 | 0.78264606 | 1.13716161 | 0.68824524 | 0.49129836 | 0.84493924 |
| Coq5 | 2392.29164 | -0.1273131 | 0.19103913 | -0.6664242 | 0.50514002 | 0.85034006 |
| Kirrel | 44.2080439 | 0.20191686 | 0.89580475 | 0.22540276 | 0.82166596 | 0.9623207 |
| Tspo | 13160.8386 | -0.6016504 | 0.39250104 | -1.5328633 | 0.12530955 | 0.53710993 |
| Rnf10 | 3869.42799 | -0.2427549 | 0.25199512 | -0.9633319 | 0.33538099 | 0.75359284 |
| Utp15 | 3077.53753 | -0.2761017 | 0.27661669 | -0.9981382 | 0.31821234 | 0.74336577 |
| Trem3 | 194.948436 | -2.7348493 | 0.40979028 | -6.6737777 | 2.493010927 | 5.647593087 |
| Gpr155 | 153.915013 | 0.29285653 | 0.59707013 | 0.49048934 | 0.62378767 | 0.90017327 |
| Tpp2 | 4146.5719 | -0.0240823 | 0.24821808 | -0.0970207 | 0.92270993 | 0.98465273 |
| Ubac2 | 2155.12215 | 0.12924551 | 0.17487181 | 0.73908716 | 0.45985408 | 0.82987212 |
| Ppp2r2d | 450.649701 | 0.15812813 | 0.30896981 | 0.51179153 | 0.60879692 | 0.89437372 |
| Enc1 | 90.2556485 | 0.03972999 | 0.61030403 | 0.06509869 | 0.94809543 | 0.99019648 |
| Ydjc | 395.530584 | -1.0268945 | 0.33600647 | -3.0561749 | 0.0022418 | 0.0433924 |
| Mapk1ip1 | 146.987815 | -0.6880833 | 0.59176895 | -1.1627567 | 0.24492823 | 0.68039867 |
| Cir1 | 342.622976 | 0.20458824 | 0.24870014 | 0.82263019 | 0.41071832 | 0.80139357 |
| Tram2 | 582.415897 | 0.13824427 | 0.36196903 | 0.38192293 | 0.70251853 | 0.92762537 |
| Cpsf2 | 3481.06002 | -0.2656371 | 0.18660608 | -1.4235177 | 0.15458612 | 0.58343244 |
| Abca9 | 663.020352 | 1.28840692 | 0.47433293 | 2.71625019 | 0.0066026 | 0.09849952 |
| Phlda3 | 57.5348691 | 0.7814831 | 0.83254181 | 0.93867129 | 0.34789954 | 0.76101142 |
| Poldip3 | 4969.77216 | 0.01801518 | 0.1865024 | 0.09659492 | 0.92304809 | 0.98465273 |
| Fam169a | 10.3360546 | 0.69589419 | 2.09944587 | 0.33146565 | 0.74029278 | 0.93800224 |
| Oasl1 | 54.8374985 | -0.6396991 | 0.62736686 | -1.0196571 | 0.30789113 | 0.73735947 |
| Ptpre | 1789.01813 | -0.5524757 | 0.31573062 | -1.7498327 | 0.08014719 | 0.43723638 |
| Pdcd7 | 209.165641 | -0.3144751 | 0.57136407 | -0.5503935 | 0.5820495 | 0.88321857 |
| Haus1 | 2317.24777 | 0.29940813 | 0.34252811 | 0.87411258 | 0.38205695 | 0.78483671 |
| Rpl37 | 5197.30664 | -0.1094473 | 0.28505412 | -0.3839526 | 0.70101361 | 0.92722415 |
| Rhod | 48.4904704 | 3.15669251 | 1.22542769 | 2.57599247 | 0.00999528 | 0.12925189 |
| Ppp4r3a | 2430.46966 | -0.0867968 | 0.45690279 | -0.1899678 | 0.84933436 | 0.96765585 |
| Card6 | 58.9444259 | -0.5352314 | 1.38757716 | -0.3857309 | 0.69969597 | 0.92669304 |
| Tcf20 | 653.346793 | -0.1406769 | 0.33081748 | -0.4252403 | 0.67066148 | 0.91711586 |
| Mcm3 | 22603.9718 | -0.3676771 | 0.2012948 | -1.8265603 | 0.06776591 | 0.40101338 |
| Ankrd13a | 930.204449 | -0.2050871 | 0.22625736 | -0.9064329 | 0.36470679 | 0.77108653 |

| | | | | | | |
|---|---|---|---|---|---|---|
| Ipo9 | 3996.63916 | 0.02372885 | 0.22512481 | 0.1054031 | 0.91605595 | 0.98385535 |
| Ndufa7 | 4790.21587 | 0.79052723 | 0.50510443 | 1.5650768 | 0.1175649 | 0.52373323 |
| Shisa4 | 30.3198616 | 3.11958504 | 1.26946546 | 2.45740048 | 0.01399466 | 0.16105047 |
| Git2 | 2108.88475 | 0.09883641 | 0.21190817 | 0.46641152 | 0.64092097 | 0.90685007 |
| Lman1 | 4137.98815 | -0.2467211 | 0.214568 | -1.1498504 | 0.2502055 | 0.6842534 |
| Wipi1 | 122.198543 | 0.48686011 | 0.47211538 | 1.03123119 | 0.30243242 | 0.73372823 |
| Gpr45 | 11.4728424 | 0.57952748 | 1.46703916 | 0.39503204 | 0.69281924 | 0.92550785 |
| Tdrkh | 260.563016 | 0.86865176 | 0.46412349 | 1.8715962 | 0.06126249 | 0.38220919 |
| Ammecr1l | 1559.24931 | -0.4474513 | 0.27880217 | -1.6049061 | 0.10851448 | 0.50301538 |
| Slc16a6 | 2455.40024 | -0.4091815 | 0.25243998 | -1.6209061 | 0.10503778 | 0.49581042 |
| Metap1d | 414.878495 | 0.07665295 | 0.29955807 | 0.25588678 | 0.79803826 | 0.95433165 |
| Rnpep | 17713.2595 | -0.0872625 | 0.22093354 | -0.3949719 | 0.69286364 | 0.92550785 |
| AW549877 | 307.91931 | 0.90660853 | 0.64050672 | 1.41545516 | 0.15693509 | 0.58620126 |
| Agrn | 19.9012029 | -0.6001444 | 1.59460792 | -0.3763586 | 0.70665033 | 0.92921138 |
| Mvk | 872.048122 | 0.55702892 | 0.28777607 | 1.93563324 | 0.05291264 | 0.3556485 |
| Mfsd9 | 136.628074 | 0.31546989 | 0.4057404 | 0.7775166 | 0.43685405 | 0.81788117 |
| Tango6 | 426.880129 | -0.1756378 | 0.49326789 | -0.3560698 | 0.72178832 | 0.93398714 |
| Pkp2 | 240.435388 | -0.6017697 | 0.3187001 | -1.8882006 | 0.05899903 | 0.3744938 |
| Pigs | 4703.02017 | -0.2636606 | 0.20067929 | -1.3138408 | 0.18889978 | 0.61935434 |
| S100a10 | 2739.19424 | 0.56916243 | 0.4059888 | 1.40191658 | 0.16094016 | 0.59141542 |
| Znrf3 | 619.405739 | -0.1522612 | 0.3760617 | -0.4048835 | 0.68556314 | 0.92314423 |
| Dcaf17 | 1153.74053 | 0.15885844 | 0.1913185 | 0.83033495 | 0.40634943 | 0.79992362 |
| Spidr | 2115.1524 | 0.09964041 | 0.17443627 | 0.57121382 | 0.56785471 | 0.87929304 |
| Mettl8 | 656.362298 | 0.34888914 | 0.26200873 | 1.33159358 | 0.18299378 | 0.61465208 |
| Arhgef11 | 91.4510097 | 0.16681794 | 0.46829965 | 0.35622051 | 0.72167544 | 0.93398714 |
| Rapgef5 | 141.762337 | -0.0724959 | 0.64943066 | -0.1116299 | 0.91111686 | 0.98289594 |
| Zbed3 | 687.919549 | -0.1032328 | 0.26943336 | -0.3831478 | 0.7016102 | 0.92738786 |
| Tlk1 | 1149.31696 | -0.1012647 | 0.31450538 | -0.3219809 | 0.74746715 | 0.9399191 |
| Wdr41 | 1872.61031 | 0.20596477 | 0.23256058 | 0.88563919 | 0.37581198 | 0.77828094 |
| Ncapg2 | 1158.92583 | 0.18146979 | 0.23588294 | 0.76932136 | 0.44170256 | 0.81967485 |
| Mat2b | 3633.27017 | 0.28410693 | 0.29156252 | 0.97442886 | 0.32984364 | 0.74916028 |
| Igsf3 | 70.29847 | -0.2336172 | 0.66519006 | -0.3512037 | 0.72543555 | 0.93544211 |
| Csgalnact2 | 408.603344 | -0.1624625 | 0.28992374 | -0.560363 | 0.57523184 | 0.88059206 |
| Tbca | 2255.38449 | 0.61608302 | 0.40421688 | 1.52413977 | 0.1274738 | 0.54145382 |
| Dstyk | 127.397438 | 0.69810561 | 0.57532576 | 1.21340927 | 0.22497338 | 0.65870021 |
| Wdr60 | 74.5162654 | 1.22971739 | 0.49316482 | 2.49352211 | 0.01264827 | 0.14978344 |
| Wdr11 | 1361.80518 | 0.02696492 | 0.31642213 | 0.08521818 | 0.93208793 | 0.98730569 |
| Zfp386 | 746.000058 | 0.43404215 | 0.36168566 | 1.20005354 | 0.23011855 | 0.66484742 |
| Tmcc2 | 19.4241898 | -0.629983 | 1.52648851 | -0.4127008 | 0.67982584 | 0.92054299 |
| Abhd14b | 198.634763 | -0.0928253 | 0.49506425 | -0.1875015 | 0.85126748 | 0.9679302 |
| Hnrnpf | 48928.6606 | -0.1075105 | 0.1476933 | -0.7279309 | 0.4666559 | 0.83345004 |
| Arsb | 1781.37303 | -0.224853 | 0.3422636 | -0.6569585 | 0.51120757 | 0.85373917 |
| Zfp239 | 150.173184 | -0.3199155 | 0.36874513 | -0.867579 | 0.38562487 | 0.78819532 |
| Kank3 | 253.759597 | -0.0794325 | 0.48323605 | -0.1643763 | 0.86943496 | 0.97228246 |
| Uggt2 | 39.8879095 | -0.7518077 | 1.04262605 | -0.7210712 | 0.47086568 | 0.83381643 |
| Inpp5f | 619.961437 | -0.1135551 | 0.30376472 | -0.373826 | 0.70853379 | 0.92988884 |

| | | | | | | |
|---|---|---|---|---|---|---|
| Inka1 | 2110.14542 | 0.17710025 | 0.33354502 | 0.53096355 | 0.59544404 | 0.88830085 |
| Ccdc115 | 5419.94803 | 0.18538978 | 0.30145011 | 0.61499326 | 0.53855917 | 0.86663962 |
| Ssh1 | 358.71935 | 0.82633274 | 0.40028497 | 2.06436115 | 0.03898349 | 0.29842616 |
| Rassf4 | 3145.55606 | -0.6890791 | 0.19677854 | -3.5018002 | 0.00046213 | 0.01267531 |
| Ppig | 1745.65903 | -0.2595437 | 0.24993022 | -1.0384648 | 0.29905371 | 0.73093388 |
| Msantd2 | 46.126378 | -0.1803103 | 1.12617561 | -0.1601085 | 0.8727956 | 0.97344599 |
| Cox10 | 1402.38681 | -0.451483 | 0.26093038 | -1.7302817 | 0.08357996 | 0.44386868 |
| Klhl23 | 778.937784 | -0.2441894 | 0.24237429 | -1.0074888 | 0.31369991 | 0.74065408 |
| Tent2 | 3587.47677 | 0.15268371 | 0.16298616 | 0.93678944 | 0.34886688 | 0.76167919 |
| Armc5 | 882.725777 | -0.1545072 | 0.28922719 | -0.5342069 | 0.59319842 | 0.88797037 |
| Bend6 | 41.7407405 | 0.71062159 | 1.3299594 | 0.53431826 | 0.59312136 | 0.88797037 |
| Nfrkb | 348.625529 | 0.23539501 | 0.41631507 | 0.56542515 | 0.57178462 | 0.88035336 |
| Cmklr1 | 60.9738918 | 1.05609769 | 0.61196772 | 1.72574086 | 0.08439404 | 0.44526833 |
| Slc35f2 | 21.2030277 | 1.35812467 | 1.37410659 | 0.98836922 | 0.32297185 | 0.74562749 |
| Zfp451 | 885.933276 | 0.11537934 | 0.34946389 | 0.33016098 | 0.74127833 | 0.93800224 |
| Chchd7 | 1290.07726 | 0.24453357 | 0.313223 | 0.78070119 | 0.43497826 | 0.81606911 |
| Slc35e2 | 406.780662 | 0.78010671 | 0.45948622 | 1.69778045 | 0.08954921 | 0.45701104 |
| Tbc1d22b | 274.347462 | 0.39958966 | 0.65902902 | 0.6063309 | 0.54429505 | 0.8695007 |
| Kdm5b | 486.267334 | 0.00198331 | 0.36900742 | 0.00537473 | 0.99571161 | 0.99884675 |
| 0610010F05I | 898.668734 | 0.07386238 | 0.27933791 | 0.26441948 | 0.7914567 | 0.95268772 |
| Abhd14a | 736.307025 | -0.2506012 | 0.22351558 | -1.1211798 | 0.26221133 | 0.69762406 |
| Fbxo38 | 1914.27397 | -0.1857583 | 0.2966422 | -0.6262032 | 0.53118165 | 0.86415301 |
| Zfand4 | 301.363 | -0.4045435 | 0.4459651 | -0.9071192 | 0.36434376 | 0.77081383 |
| Bag2 | 2115.91692 | -0.1043981 | 0.20841892 | -0.500905 | 0.61643797 | 0.89878056 |
| Sgsm1 | 68.0262841 | 1.19871186 | 0.83405714 | 1.43720593 | 0.15065949 | 0.57859667 |
| Ammecr1 | 259.734524 | -0.3691623 | 0.32112065 | -1.1496061 | 0.25030613 | 0.6842534 |
| Lyn | 12031.5957 | -0.0036239 | 0.22970946 | -0.0157759 | 0.98741318 | 0.9962376 |
| Rabif | 1994.19872 | -0.1007117 | 0.16870446 | -0.5969714 | 0.55052646 | 0.87093412 |
| Tmc7 | 23.8218675 | -0.9307239 | 0.88221857 | -1.0549811 | 0.29143395 | 0.72550093 |
| Grk3 | 32.282875 | -1.5210148 | 1.41493213 | -1.0749737 | 0.28238652 | 0.71787592 |
| Pm20d1 | 19.0191282 | -1.2291411 | 2.48263771 | -0.4950948 | 0.62053319 | 0.89896588 |
| Isl1 | 12.3610947 | 0.07503746 | 2.09485922 | 0.03581981 | 0.97142604 | 0.99443087 |
| Trem1 | 118.797358 | -3.4987828 | 0.89218182 | -3.9216029 | 8.796186581 | 0.00322167 |
| Nxt2 | 953.115765 | 0.71578112 | 0.29555798 | 2.42179593 | 0.01544402 | 0.17214842 |
| Sestd1 | 123.749359 | 0.33835108 | 0.69720916 | 0.48529351 | 0.62746816 | 0.9022352 |
| Pelo | 47.019349 | -0.1687256 | 0.92449235 | -0.1825062 | 0.85518545 | 0.96892504 |
| Stab1 | 8399.83959 | 1.28638616 | 0.34413735 | 3.73800213 | 0.00018549 | 0.00611612 |
| Hsd3b7 | 892.538313 | 1.1091903 | 0.41045327 | 2.70235467 | 0.00688503 | 0.10086772 |
| Mrtfa | 633.812372 | -0.1484428 | 0.26991006 | -0.5499714 | 0.58233896 | 0.88340185 |
| Gm5617 | 124.428869 | 0.91229851 | 0.68916431 | 1.32377504 | 0.18557777 | 0.61775377 |
| Ttc19 | 77.9822691 | -0.7674128 | 0.81731615 | -0.9389424 | 0.3477603 | 0.76101142 |
| Sgsm3 | 1345.5098 | -0.1151252 | 0.29584271 | -0.3891431 | 0.69717026 | 0.92650558 |
| Tmem183a | 3460.84954 | 0.21705034 | 0.17278702 | 1.25617274 | 0.20905335 | 0.64228937 |
| Setd1a | 107.029439 | 0.32327262 | 0.74818678 | 0.43207476 | 0.66568708 | 0.91520501 |
| S100a13 | 962.306501 | 0.43686514 | 0.44321749 | 0.98566765 | 0.32429623 | 0.74596967 |
| Prox2 | 73.487449 | 0.20140616 | 0.7364723 | 0.27347419 | 0.78448873 | 0.95094799 |

| | | | | | | |
|---|---|---|---|---|---|---|
| Pbrm1 | 1118.06207 | -0.1673605 | 0.29840508 | -0.5608502 | 0.57489966 | 0.88059206 |
| Hps4 | 643.362022 | -0.3018124 | 0.25398539 | -1.1883063 | 0.23471276 | 0.67038085 |
| Specc1 | 2070.59105 | -0.0749841 | 0.15616867 | -0.480148 | 0.63112217 | 0.90394609 |
| Tnfrsf14 | 118.064045 | -0.675075 | 0.52140616 | -1.2947201 | 0.1954168 | 0.62710749 |
| Ubash3a | 222.043841 | -3.5786787 | 0.52528593 | -6.8128205 | 9.570355452 | 2.341483164 |
| Arl15 | 100.740778 | -0.123956 | 0.63435983 | -0.1954034 | 0.84507718 | 0.96765585 |
| Ikbke | 628.966916 | -0.6342475 | 0.31216698 | -2.0317571 | 0.04217825 | 0.3140393 |
| Arel1 | 366.485197 | 0.11209755 | 0.29892694 | 0.37499981 | 0.70766061 | 0.92950127 |
| Grap2 | 19.5444883 | -7.6856286 | 1.78839238 | -4.2975069 | 1.727298959 | 0.00081867 |
| Frem3 | 6.77928991 | 1.52756991 | 2.76308912 | 0.55284858 | 0.5803671 | 0.88286755 |
| Gnl3 | 4333.19968 | -0.056831 | 0.24626045 | -0.2307759 | 0.81748893 | 0.96079676 |
| Lgalsl | 353.851431 | 0.82169683 | 0.37831849 | 2.17197114 | 0.02985784 | 0.25769767 |
| Snx18 | 2678.85584 | -0.284606 | 0.32843608 | -0.8665492 | 0.38618906 | 0.7886896 |
| Rbm45 | 152.532612 | -0.2676214 | 0.48675766 | -0.5498042 | 0.58245366 | 0.88343755 |
| Fam83g | 69.9377342 | 0.88321123 | 1.32045645 | 0.66886812 | 0.50357961 | 0.85012672 |
| Smim12 | 1350.83271 | 0.080661 | 0.33904817 | 0.23790424 | 0.81195537 | 0.95967622 |
| Tsen2 | 1161.76754 | 0.0289194 | 0.34385937 | 0.0841024 | 0.93297501 | 0.98748011 |
| Gatad2b | 507.560139 | -0.5821533 | 0.43669512 | -1.3330887 | 0.18250269 | 0.61383074 |
| Rbm7 | 6880.29224 | 0.57048944 | 0.33737603 | 1.69096022 | 0.0908444 | 0.46044203 |
| Dennd4b | 680.260017 | -0.2092667 | 0.37458269 | -0.5586661 | 0.5763896 | 0.88105119 |
| Atf4 | 6631.71387 | -0.2715838 | 0.17804048 | -1.525405 | 0.12715812 | 0.54111158 |
| Zmym6 | 73.3543485 | 2.24187823 | 0.95045884 | 2.35873257 | 0.01833747 | 0.19092573 |
| Agps | 1258.09093 | -0.2537127 | 0.23805231 | -1.0657855 | 0.28652061 | 0.72260415 |
| Nfkbil1 | 232.190132 | 0.07302278 | 0.28294536 | 0.25808086 | 0.7963445 | 0.9537323 |
| Fbrs | 289.304524 | -0.4530699 | 0.42722716 | -1.0604894 | 0.28892199 | 0.72395811 |
| Dhx29 | 1339.88257 | -0.3769333 | 0.3562451 | -1.0580729 | 0.2900222 | 0.72418639 |
| Zfp532 | 32.2815957 | 2.61865163 | 1.70308789 | 1.53759042 | 0.12414879 | 0.53551789 |
| Mindy2 | 177.261099 | 0.57479002 | 0.36233761 | 1.58633828 | 0.11266259 | 0.51139203 |
| Zmym4 | 176.073982 | -0.085204 | 0.57310979 | -0.1486696 | 0.88181434 | 0.97582756 |
| Mios | 1265.1546 | -0.1019409 | 0.23613212 | -0.4317114 | 0.66595115 | 0.91520501 |
| C1galt1 | 112.253889 | 0.24800768 | 0.43959404 | 0.56417435 | 0.57263548 | 0.88035336 |
| Dctpp1 | 2547.75625 | 0.05980676 | 0.39271819 | 0.15228926 | 0.87895879 | 0.97535404 |
| Zfp410 | 2412.84079 | 0.28294937 | 0.22247448 | 1.27182843 | 0.20343409 | 0.63809363 |
| Tbc1d8b | 46.5520845 | 0.39661893 | 0.68473948 | 0.57922603 | 0.56243667 | 0.8755831 |
| Leo1 | 1630.32842 | -0.0835514 | 0.20332556 | -0.4109243 | 0.68112801 | 0.92061965 |
| Clspn | 621.9715 | -0.1863156 | 0.39261269 | -0.4745532 | 0.63510543 | 0.90531997 |
| Tbc1d10b | 868.439703 | -0.1042891 | 0.20862215 | -0.4998947 | 0.61714921 | 0.89878056 |
| Prdm10 | 131.936364 | 0.44636312 | 0.87821752 | 0.50826033 | 0.61127079 | 0.89563727 |
| Radx | 252.738242 | 0.64921565 | 0.27323507 | 2.3760334 | 0.01749988 | 0.18560042 |
| Ago4 | 24.5848438 | -1.5761257 | 1.25608581 | -1.2547914 | 0.20955449 | 0.6434438 |
| Cd2bp2 | 1494.0384 | -0.4171058 | 0.21531628 | -1.9371773 | 0.05272367 | 0.35506328 |
| Sdhaf3 | 383.978768 | 0.06259652 | 0.35633505 | 0.1756676 | 0.86055509 | 0.96994723 |
| Usp22 | 3835.931 | 0.23158918 | 0.20400508 | 1.13521281 | 0.25628616 | 0.69126266 |
| Elmsan1 | 327.986595 | -0.1194366 | 0.36655922 | -0.3258317 | 0.74455172 | 0.93917909 |
| Dmtf1 | 1133.84032 | 0.09698277 | 0.33264444 | 0.29155086 | 0.77063005 | 0.94454775 |
| AA986860 | 16.6357849 | 2.77385136 | 1.490288 | 1.86128543 | 0.06270388 | 0.38662125 |

| | | | | | | |
|---|---|---|---|---|---|---|
| Pwwp3b | 267.811974 | 1.27992663 | 0.5209291 | 2.45700733 | 0.01400998 | 0.1610753 |
| Ubap2l | 9342.24221 | -0.1658068 | 0.26022614 | -0.6371642 | 0.52401791 | 0.8594073 |
| Dnal1 | 24.5078279 | 0.29610749 | 0.98569877 | 0.30040363 | 0.7638693 | 0.94300309 |
| Sun2 | 910.338674 | -0.2285158 | 0.28712472 | -0.7958765 | 0.42610383 | 0.81021713 |
| Gtpbp1 | 1231.66718 | -0.6555435 | 0.18314992 | -3.5792726 | 0.00034455 | 0.00991742 |
| Sem1 | 4903.36887 | 0.05699937 | 0.25591581 | 0.22272704 | 0.82374796 | 0.9628698 |
| Asxl1 | 447.663834 | 0.39596541 | 0.27893478 | 1.41956269 | 0.15573503 | 0.58481688 |
| Sin3a | 1134.30367 | -0.2861336 | 0.35296093 | -0.8106664 | 0.41755728 | 0.80503991 |
| Adprhl2 | 1856.73992 | -0.0547676 | 0.20408488 | -0.2683571 | 0.78842445 | 0.95225386 |
| Dhrs7b | 2658.98411 | 0.52355055 | 0.33566907 | 1.55972238 | 0.1188255 | 0.52515149 |
| Mier2 | 642.101329 | -0.367492 | 0.28750308 | -1.2782193 | 0.2011721 | 0.63508084 |
| Ube2q1 | 592.619657 | -0.2060351 | 0.28501014 | -0.7229045 | 0.46973855 | 0.83381643 |
| Ipo11 | 2794.70219 | 0.31220678 | 0.22726103 | 1.37378055 | 0.16950982 | 0.60382474 |
| Sh2b3 | 2814.31168 | -0.2226502 | 0.21360192 | -1.0423603 | 0.29724465 | 0.72954201 |
| Fam199x | 138.103735 | -0.4463909 | 0.8861021 | -0.5037692 | 0.61442358 | 0.89844745 |
| Kdm7a | 375.157832 | -0.5096431 | 0.47988781 | -1.0620046 | 0.2882336 | 0.72342257 |
| Atxn2 | 310.9893 | -0.4032246 | 0.37363028 | -1.0792074 | 0.28049529 | 0.7155865 |
| Hirip3 | 525.366032 | -0.2507539 | 0.3482501 | -0.7200398 | 0.47150048 | 0.83393588 |
| Asb4 | 345.917998 | 1.34740101 | 0.41465952 | 3.24941533 | 0.00115642 | 0.02624591 |
| Stk40 | 375.337774 | 0.05943561 | 0.37147335 | 0.15999966 | 0.87288134 | 0.97344599 |
| Pbxip1 | 785.459011 | -0.3118416 | 0.22566857 | -1.3818566 | 0.16701575 | 0.60073616 |
| Oscp1 | 140.280543 | 1.27050318 | 0.63525148 | 2.00000035 | 0.04550023 | 0.3293163 |
| Maff | 34.7743349 | 0.59842406 | 0.90762396 | 0.6593304 | 0.50968363 | 0.85335831 |
| Safb2 | 1143.77313 | -0.381459 | 0.50017876 | -0.7626453 | 0.44567495 | 0.82157047 |
| Shc1 | 3921.2722 | 0.11331185 | 0.18548576 | 0.61089245 | 0.54127078 | 0.86776854 |
| Zfyve1 | 255.299822 | 0.80329492 | 0.3504853 | 2.29195041 | 0.02190851 | 0.21320672 |
| Pla2g6 | 166.284493 | 0.83443499 | 0.63629069 | 1.31140531 | 0.18972087 | 0.62019167 |
| Rgsl1 | 93.9168379 | 1.31194273 | 1.35042762 | 0.9715017 | 0.33129851 | 0.75037486 |
| Flad1 | 1437.2065 | -0.1440143 | 0.21670732 | -0.6645567 | 0.50633404 | 0.85093891 |
| Itpr3 | 148.167603 | 0.48985805 | 0.61681108 | 0.79417842 | 0.42709157 | 0.81021713 |
| Acad12 | 126.154262 | 0.99809665 | 0.53662953 | 1.85993614 | 0.06289456 | 0.38701669 |
| Alkbh5 | 4304.86938 | -0.2173823 | 0.25368589 | -0.8568953 | 0.39150273 | 0.79018736 |
| Arrdc4 | 1658.16698 | -0.253416 | 0.29731337 | -0.8523532 | 0.39401808 | 0.79233901 |
| Wdr55 | 1971.45363 | -0.4443034 | 0.19306105 | -2.3013622 | 0.02137117 | 0.20964994 |
| Immp1l | 2038.05898 | 0.82697349 | 0.5256491 | 1.57324247 | 0.11566268 | 0.52030172 |
| Ypel3 | 106.643291 | 1.6968396 | 0.69120446 | 2.45490256 | 0.01409228 | 0.16167905 |
| Zc3h12a | 55.108378 | -0.6124603 | 0.79884103 | -0.7666861 | 0.44326817 | 0.82038676 |
| Myo15 | 27.1753447 | -0.211348 | 1.13022909 | -0.1869957 | 0.851664 | 0.96798344 |
| Selenok | 4639.42202 | 0.77357496 | 0.31347729 | 2.4677225 | 0.01359757 | 0.15766735 |
| Npl | 3492.0444 | 1.29272334 | 0.22586535 | 5.72342476 | 1.043978122 | 1.467929237 |
| Mapk6 | 2941.46306 | -0.0166644 | 0.36674062 | -0.0454392 | 0.96375724 | 0.99406401 |
| Stn1 | 1332.2171 | 0.23415561 | 0.25086381 | 0.93339733 | 0.35061486 | 0.76238338 |
| Dhx9 | 645.483464 | 0.21184013 | 0.43806664 | 0.48357969 | 0.62868419 | 0.90276005 |
| Sipa1l1 | 53.4159727 | 0.40832715 | 0.89121063 | 0.45817132 | 0.64682937 | 0.90927884 |
| Commd10 | 3103.90096 | 0.55894633 | 0.19298192 | 2.89636635 | 0.00377511 | 0.06422944 |
| Shcbp1l | 51.8672524 | -0.6979595 | 1.28971067 | -0.5411753 | 0.58838679 | 0.88690478 |

| | | | | | | |
|---|---|---|---|---|---|---|
| Atpaf2 | 2934.7754 | -0.1919012 | 0.1963435 | -0.977375 | 0.32838353 | 0.74843736 |
| Tceal9 | 3319.43585 | 0.95344164 | 0.28530036 | 3.34188728 | 0.00083211 | 0.02023694 |
| Naa25 | 2721.21492 | -0.6052395 | 0.35816268 | -1.6898453 | 0.09105755 | 0.46086347 |
| Trafd1 | 2049.69076 | -0.3717119 | 0.2718051 | -1.3675677 | 0.17144744 | 0.60430173 |
| Wdr74 | 2994.53464 | -0.4283052 | 0.24888759 | -1.720878 | 0.08527297 | 0.44656163 |
| Dpm3 | 323.713122 | 0.54479634 | 0.46874135 | 1.16225363 | 0.24513246 | 0.68039867 |
| Bmt2 | 400.350058 | 0.19604566 | 0.34144777 | 0.57416001 | 0.56585953 | 0.878492 |
| Sgtb | 57.8016769 | 0.98659899 | 0.714447 | 1.38092677 | 0.16730148 | 0.60123354 |
| Hectd4 | 412.341621 | -0.03582 | 0.37062623 | -0.0966471 | 0.92300663 | 0.98465273 |
| Id1 | 1784.39519 | -0.1345474 | 0.28034782 | -0.4799302 | 0.63127699 | 0.90394609 |
| Krtcap2 | 2480.86805 | -0.2414509 | 0.20720867 | -1.1652548 | 0.24391587 | 0.67911981 |
| Bex2 | 103.796037 | 0.25517836 | 0.65895728 | 0.38724568 | 0.69857434 | 0.92669304 |
| Apobr | 492.949573 | -0.428019 | 0.32099043 | -1.3334322 | 0.18239001 | 0.61363505 |
| Hebp1 | 102.121773 | 2.03459012 | 0.79241812 | 2.56757144 | 0.01024137 | 0.13050277 |
| Smg7 | 735.410699 | -0.1755961 | 0.34393461 | -0.5105508 | 0.60966562 | 0.89508835 |
| Exog | 1122.37885 | -0.1219057 | 0.20306756 | -0.6003211 | 0.54829223 | 0.87059427 |
| Rnf214 | 471.710688 | -0.4152575 | 0.51566237 | -0.8052895 | 0.42065265 | 0.80681152 |
| Aqp11 | 40.6052957 | 1.61022316 | 0.88692235 | 1.81551763 | 0.0694444 | 0.40704999 |
| Hecw2 | 35.1374088 | 1.34915878 | 0.90182095 | 1.49603841 | 0.13464365 | 0.55585235 |
| Krba1 | 159.67887 | -0.2433622 | 0.45934787 | -0.5297994 | 0.59625103 | 0.88873045 |
| Mcts2 | 599.423792 | -0.1283026 | 0.25848013 | -0.4963732 | 0.6196311 | 0.89896588 |
| Flt3 | 29.7108947 | -2.144991 | 1.44632831 | -1.4830595 | 0.1380586 | 0.56068131 |
| Snai1 | 13.3413132 | 0.85309243 | 1.56198595 | 0.54615883 | 0.58495676 | 0.88473987 |
| Fgf11 | 18.2613072 | 1.42550796 | 2.42565054 | 0.58768068 | 0.55674665 | 0.87276537 |
| Alkbh6 | 648.38311 | 0.30983151 | 0.32660204 | 0.94865148 | 0.3427979 | 0.75940723 |
| Nrep | 584.954654 | 2.27308522 | 0.61253356 | 3.71095623 | 0.00020648 | 0.00669985 |
| Serpinb6b | 427.642669 | 1.13429762 | 0.36443747 | 3.1124616 | 0.00185534 | 0.03795383 |
| Trp53rkb | 341.997454 | 0.16094244 | 0.41906334 | 0.38405278 | 0.70093933 | 0.92722415 |
| Tom1 | 2083.18551 | -0.0426431 | 0.15910228 | -0.2680232 | 0.78868148 | 0.95225386 |
| Aida | 746.570176 | 0.23437748 | 0.26294476 | 0.89135634 | 0.37273803 | 0.77748705 |
| Foxo4 | 13.0341315 | -2.114012 | 1.15297942 | -1.8335211 | 0.06672512 | 0.3975881 |
| Gm14117 | 359.758458 | 0.37779707 | 0.31493751 | 1.19959374 | 0.23029716 | 0.66491035 |
| Greb1l | 12.2728016 | 1.21655705 | 1.29014839 | 0.94295901 | 0.34570187 | 0.7602916 |
| Sbk1 | 14.9800911 | 0.23255447 | 1.83425252 | 0.12678433 | 0.8991111 | 0.98010294 |
| Borcs5 | 604.004837 | -0.6800239 | 0.27162434 | -2.5035457 | 0.01229558 | 0.14760729 |
| Nhlrc3 | 1044.57535 | 1.02767747 | 0.31726551 | 3.23917172 | 0.00119877 | 0.02685824 |
| Gng2 | 4085.26848 | 0.08576861 | 0.1721945 | 0.49809146 | 0.61841958 | 0.89878056 |
| Klhl6 | 8537.21641 | -0.0266194 | 0.13817543 | -0.192649 | 0.84723384 | 0.96765585 |
| Nemp2 | 327.00689 | 0.60143554 | 0.49319661 | 1.21946407 | 0.22266811 | 0.65613364 |
| Ptgir | 13.5609966 | 0.20310004 | 1.82552787 | 0.11125551 | 0.91141373 | 0.98292561 |
| Edem3 | 133.516383 | 0.08809561 | 0.63962646 | 0.13772977 | 0.89045399 | 0.97871887 |
| Disc1 | 103.079718 | 0.08590137 | 0.53237764 | 0.1613542 | 0.87181444 | 0.97298659 |
| Zfp513 | 77.2930079 | 1.61312103 | 0.8127985 | 1.9846506 | 0.04718336 | 0.33518817 |
| Tmem18 | 925.111936 | 0.92224731 | 0.31347851 | 2.94197941 | 0.00326122 | 0.05756776 |
| Spice1 | 261.487504 | 0.13348096 | 0.41273576 | 0.32340536 | 0.74638827 | 0.93960567 |
| Dpy19l1 | 1163.15383 | -0.44846 | 0.23233245 | -1.9302511 | 0.05357574 | 0.35852954 |

| | | | | | | |
|---|---|---|---|---|---|---|
| Synpo | 20.2451192 | 0.7579477 | 1.33573743 | 0.56743765 | 0.57041688 | 0.88027118 |
| Tmem82 | 17.4633574 | 0.88934522 | 1.4136843 | 0.62909747 | 0.52928525 | 0.86280549 |
| Zfp866 | 241.999296 | -0.0464106 | 0.40163812 | -0.1155533 | 0.90800656 | 0.9817854 |
| Tuba1c | 29083.5834 | 0.28799303 | 0.28508778 | 1.01019072 | 0.31240393 | 0.74051791 |
| Lrrn4 | 30.0017969 | 0.52951901 | 1.88003544 | 0.28165374 | 0.77820903 | 0.94837927 |
| Mob1a | 7179.01629 | -0.1134933 | 0.20911261 | -0.5427376 | 0.58731051 | 0.88632735 |
| Tmem186 | 987.760739 | 0.02484199 | 0.34066647 | 0.07292172 | 0.94186841 | 0.9896127 |
| Crppa | 151.917274 | 0.67578013 | 0.40151012 | 1.68309616 | 0.09235648 | 0.46597694 |
| Ppp2r3a | 86.6166424 | 0.6090701 | 0.49739452 | 1.22452113 | 0.2207557 | 0.65450908 |
| Hpdl | 13.8582962 | -2.6436999 | 2.46755677 | -1.0713836 | 0.28399698 | 0.71958057 |
| Arl11 | 847.760321 | 0.13287526 | 0.26626331 | 0.49903707 | 0.61775327 | 0.89878056 |
| Pyurf | 1126.76964 | -0.0969503 | 0.30495585 | -0.3179159 | 0.75054872 | 0.9399191 |
| Simc1 | 198.899349 | -0.5687121 | 0.54039181 | -1.0524069 | 0.2926129 | 0.72584701 |
| Rfesd | 299.695838 | 0.60060124 | 0.56839917 | 1.05665398 | 0.29066953 | 0.72461526 |
| Zmpste24 | 2676.19373 | 0.0112692 | 0.20326753 | 0.05544023 | 0.95578775 | 0.99215259 |
| Upf2 | 277.619256 | 0.19080259 | 0.35252537 | 0.54124499 | 0.58833873 | 0.88690478 |
| Fam129c | 17.4905492 | -0.3568176 | 1.61545396 | -0.2208776 | 0.82518772 | 0.96312579 |
| Exoc3l | 59.753909 | 0.47670536 | 1.02413531 | 0.46547107 | 0.64159415 | 0.90700632 |
| Tmem64 | 139.00156 | -0.0805209 | 0.43759191 | -0.1840091 | 0.8540063 | 0.96839628 |
| Pigv | 319.13217 | 0.41847642 | 0.40319809 | 1.03789287 | 0.29931993 | 0.73113229 |
| Uevld | 464.451255 | 0.52303512 | 0.26874742 | 1.94619588 | 0.05163121 | 0.35011344 |
| Ifi209 | 205.303492 | -0.1654442 | 0.45121721 | -0.3666619 | 0.71387122 | 0.93235217 |
| Trim56 | 89.6108154 | -0.7501827 | 0.73051914 | -1.0269173 | 0.30445942 | 0.73467909 |
| Tmem11 | 2652.94746 | -0.0470424 | 0.22601183 | -0.2081415 | 0.83511848 | 0.96641606 |
| Zfp784 | 65.3497475 | 0.53948011 | 0.77548562 | 0.69566745 | 0.48663712 | 0.84183502 |
| Smco3 | 29.889467 | -0.1028214 | 1.30250971 | -0.078941 | 0.93707958 | 0.98845735 |
| B3galnt1 | 711.958358 | 2.28814515 | 0.62841099 | 3.64116027 | 0.00027141 | 0.00823867 |
| D17H6S53E | 467.691486 | 0.42964365 | 0.2217807 | 1.93724549 | 0.05271534 | 0.35506328 |
| Fbrsl1 | 181.507476 | 0.46345508 | 0.39064694 | 1.18637837 | 0.23547291 | 0.67164041 |
| Filip1l | 1443.09479 | 0.21536239 | 0.51815348 | 0.41563436 | 0.67767758 | 0.91958179 |
| Gm6741 | 153.478778 | 0.38286157 | 0.41949005 | 0.91268332 | 0.36140912 | 0.76887042 |
| Hexim2 | 90.518498 | 0.76706092 | 0.61237503 | 1.25259992 | 0.21035135 | 0.64378984 |
| Gprasp1 | 279.696108 | 0.6104378 | 0.73926621 | 0.82573474 | 0.40895458 | 0.80100711 |
| 2510009E07I | 517.895239 | 0.70790815 | 0.47674894 | 1.4848657 | 0.1375794 | 0.5600695 |
| Usp48 | 2434.29781 | -0.1502757 | 0.27558306 | -0.545301 | 0.58554654 | 0.8848945 |
| Otud1 | 123.137306 | 0.01063449 | 0.53069301 | 0.02003888 | 0.98401236 | 0.99564261 |
| Rnf227 | 31.9214392 | -1.6394065 | 1.23239674 | -1.3302587 | 0.18343306 | 0.61524496 |
| Hilpda | 1061.89564 | 0.48579848 | 0.25379425 | 1.91414294 | 0.05560189 | 0.36276158 |
| Eif3j2 | 1951.00383 | -0.2637948 | 0.19495944 | -1.3530753 | 0.17603159 | 0.60763832 |
| Leng9 | 40.5753469 | 0.07872351 | 0.72757979 | 0.10819914 | 0.91383773 | 0.98319938 |
| Epop | 20.1717505 | -0.9313313 | 1.35209604 | -0.6888056 | 0.49094563 | 0.84477968 |
| Pgp | 3133.84733 | -0.3186053 | 0.25965561 | -1.2270304 | 0.21981117 | 0.65297477 |
| Zbtb37 | 229.556058 | -0.4975367 | 0.95380208 | -0.5216352 | 0.60192436 | 0.89103677 |
| Gm6863 | 70.1325385 | 0.86003469 | 0.88189283 | 0.97521452 | 0.32945386 | 0.74874249 |
| Acot6 | 10.8414554 | -1.3627636 | 1.61281294 | -0.8449608 | 0.39813275 | 0.79546892 |
| Gimap5 | 79.0019263 | 0.54925214 | 0.7133764 | 0.76993316 | 0.44133954 | 0.81967485 |

| | | | | | | |
|---|---|---|---|---|---|---|
| Hscb | 541.568198 | 0.60296434 | 0.46552825 | 1.29522612 | 0.19524224 | 0.62707955 |
| Setx | 528.222127 | -0.1361524 | 0.43089993 | -0.3159723 | 0.75202356 | 0.94018119 |
| Zc2hc1a | 401.998592 | 0.07563915 | 0.2585319 | 0.29257181 | 0.76984947 | 0.9442118 |
| Mdga1 | 72.1393755 | -0.1150353 | 0.85627302 | -0.1343442 | 0.89313038 | 0.97941957 |
| Pars2 | 409.800217 | 0.01956692 | 0.44854227 | 0.04362336 | 0.96520463 | 0.99430301 |
| Pxylp1 | 41.4013914 | -2.6393189 | 1.1798795 | -2.2369393 | 0.02529031 | 0.23122295 |
| Zfp3 | 60.6219166 | 1.14413518 | 0.59697911 | 1.91654139 | 0.05529622 | 0.3621192 |
| 1700019D03 | 10.528088 | -0.7196692 | 2.68541508 | -0.2679918 | 0.78870564 | 0.95225386 |
| Fam221b | 70.2433095 | 0.76367572 | 0.6672577 | 1.14449893 | 0.25241678 | 0.68582922 |
| Pld6 | 62.1236995 | 1.11990699 | 1.02937429 | 1.08794926 | 0.27661751 | 0.7118544 |
| Tmem221 | 14.8880055 | 1.91267042 | 1.45568369 | 1.31393271 | 0.18886885 | 0.61935434 |
| Dpy19l3 | 255.75893 | -1.3330585 | 0.69035918 | -1.9309637 | 0.05348755 | 0.35852954 |
| Fem1a | 780.61235 | -0.6711321 | 0.34160202 | -1.9646609 | 0.0494535 | 0.34140219 |
| Pde12 | 1817.69903 | -0.2178113 | 0.24108076 | -0.9034784 | 0.36627205 | 0.77230417 |
| Rpl7 | 34480.5498 | 0.10511206 | 0.34276802 | 0.30665655 | 0.7591048 | 0.94184564 |
| F830045P16I | 8.70573105 | 1.3459607 | 2.68732259 | 0.50085565 | 0.61647272 | 0.89878056 |
| Ptpn11 | 1007.61262 | -0.3902925 | 0.29158247 | -1.3385323 | 0.18072299 | 0.61122544 |
| B430306N03 | 427.230599 | -0.6475748 | 0.43579114 | -1.4859751 | 0.1372857 | 0.55981285 |
| Gm12481 | 545.075568 | 0.38397903 | 0.41419846 | 0.92704119 | 0.35390515 | 0.76401657 |
| Oaz1-ps | 8553.41329 | 0.35464666 | 0.35049247 | 1.01185242 | 0.31160862 | 0.74017637 |
| Ly6g5b | 10.4748335 | -2.1554818 | 1.94441794 | -1.1085486 | 0.26762496 | 0.70441538 |
| Lysmd4 | 315.5363 | 0.95252456 | 0.47716302 | 1.9962246 | 0.04590948 | 0.33113837 |
| Clec4a3 | 903.760524 | 1.74134638 | 0.30840679 | 5.64626478 | 1.639712515 | 2.031058461 |
| Taf10 | 243.560887 | -0.6754383 | 0.48992856 | -1.3786465 | 0.16800378 | 0.60251838 |
| Zmym1 | 811.093352 | -0.0516449 | 0.28923024 | -0.1785598 | 0.85828339 | 0.96947659 |
| Kbtbd7 | 155.176739 | 0.68647856 | 0.50171274 | 1.36827015 | 0.17122753 | 0.60430173 |
| Slc36a4 | 152.895451 | -0.2208757 | 0.59935612 | -0.3685216 | 0.71248432 | 0.93235217 |
| Gm8399 | 41.6277141 | -0.4263933 | 0.63126807 | -0.6754551 | 0.49938665 | 0.84872332 |
| S1pr2 | 1138.94115 | 0.57313595 | 0.22014037 | 2.60350224 | 0.00922767 | 0.12309928 |
| Trp53bp1 | 600.425795 | 0.16165076 | 0.33376118 | 0.48433063 | 0.62815124 | 0.90253396 |
| Ccdc84 | 187.130366 | 0.2986518 | 0.53564565 | 0.5575548 | 0.57714843 | 0.88121027 |
| Klhl15 | 213.377184 | 0.5511894 | 0.54098328 | 1.01886587 | 0.30826665 | 0.7381143 |
| A530064D06 | 28.8616219 | -0.9970582 | 1.93521775 | -0.5152176 | 0.606401 | 0.8926719 |
| Wdfy3 | 1698.71784 | 0.63350831 | 0.40766642 | 1.553987 | 0.12018752 | 0.52810845 |
| Ccrl2 | 1251.27737 | -0.1093683 | 0.52514497 | -0.2082631 | 0.83502355 | 0.96641606 |
| Thrap3 | 5280.72841 | -0.3250642 | 0.27801727 | -1.1692232 | 0.24231372 | 0.67753686 |
| Orai3 | 1702.07075 | 0.14861122 | 0.23978772 | 0.61976159 | 0.53541476 | 0.86539693 |
| Cep164 | 299.669325 | 0.64132031 | 0.665532 | 0.96362054 | 0.3352362 | 0.75359284 |
| Pura | 111.321376 | -0.053807 | 0.50526215 | -0.1064933 | 0.91519095 | 0.98347559 |
| Mgat2 | 6735.83967 | -0.0139422 | 0.14785882 | -0.0942942 | 0.92487542 | 0.98562601 |
| Gls2 | 10.518748 | -0.7280455 | 2.17346111 | -0.3349706 | 0.73764728 | 0.93800224 |
| Mrpl50 | 2930.31802 | 0.35739897 | 0.32715008 | 1.0924618 | 0.27463017 | 0.71147204 |
| Slc35g1 | 69.2938854 | 0.10191097 | 1.05058127 | 0.09700436 | 0.92272293 | 0.98465273 |
| Irf2bp1 | 147.543414 | -0.6789807 | 0.43161134 | -1.5731298 | 0.11568877 | 0.52030172 |
| Als2cl | 123.406649 | 0.32932192 | 0.70181023 | 0.4692464 | 0.63889352 | 0.90615794 |
| Fmn1 | 132.641036 | 0.1588672 | 0.48088266 | 0.33036583 | 0.74112356 | 0.93800224 |

| | | | | | | |
|---|---|---|---|---|---|---|
| Efcab9 | 11.7958226 | -2.2246139 | 1.46575311 | -1.5177276 | 0.12908309 | 0.54519878 |
| Cep68 | 1366.76721 | 0.31780819 | 0.27029829 | 1.17576842 | 0.23968743 | 0.67563574 |
| Zrsr1 | 39.4356664 | 0.69097868 | 1.01115837 | 0.68335356 | 0.49438346 | 0.84678075 |
| Tafa2 | 10.8077595 | -5.7669507 | 1.77981422 | -3.2401981 | 0.00119447 | 0.02681085 |
| Eml6 | 34.7936109 | 0.43060419 | 0.66419599 | 0.64830893 | 0.51678516 | 0.85673255 |
| S100a1 | 530.253736 | 1.13545747 | 0.44140186 | 2.57238942 | 0.01009992 | 0.12992799 |
| C130050O18 | 552.262339 | -0.243032 | 0.25234037 | -0.9631118 | 0.33549143 | 0.75359284 |
| Rsbn1 | 266.631119 | -0.5138663 | 0.30762583 | -1.6704265 | 0.09483502 | 0.47159219 |
| Proca1 | 15.1223222 | 0.79050473 | 1.23769172 | 0.63869275 | 0.52302283 | 0.85869525 |
| Pheta1 | 353.090732 | 1.05462693 | 0.38921537 | 2.709623 | 0.00673597 | 0.09963863 |
| 1810024B03I | 36.9878702 | 0.29360347 | 0.68086139 | 0.43122356 | 0.66630583 | 0.91520501 |
| Arf6 | 414.603281 | 0.13655165 | 0.32536555 | 0.41968686 | 0.67471423 | 0.91841317 |
| 1810030O07 | 2466.28497 | -0.0003988 | 0.24398136 | -0.0016345 | 0.99869587 | 0.99967651 |
| Nkrf | 63.3985343 | -1.267473 | 0.78475722 | -1.6151148 | 0.10628585 | 0.49777747 |
| Bclaf3 | 332.263483 | -0.064537 | 0.3226976 | -0.1999923 | 0.84148662 | 0.96765585 |
| Lsm8 | 1792.91678 | -0.3497904 | 0.21922633 | -1.5955676 | 0.11058536 | 0.50856795 |
| Tnip3 | 13.5327928 | -2.0241406 | 2.21552659 | -0.913616 | 0.36091864 | 0.76887042 |
| Foxo1 | 124.797709 | 0.29857908 | 0.37420317 | 0.79790635 | 0.42492484 | 0.80988459 |
| Gpr146 | 832.329028 | 0.7347113 | 0.42669478 | 1.72186616 | 0.08509377 | 0.44637738 |
| S1pr4 | 46.2556765 | -2.6167199 | 1.32731356 | -1.9714406 | 0.0486735 | 0.34046898 |
| Cdc25c | 1050.03861 | 0.0479574 | 0.21695588 | 0.22104678 | 0.82505601 | 0.96312579 |
| Grsf1 | 1622.02217 | 0.16312832 | 0.49453376 | 0.32986286 | 0.74150359 | 0.93800224 |
| Dnajc21 | 103.666977 | -1.014644 | 0.74927835 | -1.3541616 | 0.17568482 | 0.60763832 |
| Nxpe4 | 41.7548666 | -0.5516164 | 1.12241015 | -0.4914571 | 0.6231032 | 0.90017327 |
| Nhlrc1 | 55.6668789 | 0.25963016 | 0.81575907 | 0.31826819 | 0.75028151 | 0.9399191 |
| Il20rb | 37.1755722 | 1.70461905 | 1.24185855 | 1.37263544 | 0.16986571 | 0.60430173 |
| Pced1b | 80.8954239 | 0.21436657 | 0.70695781 | 0.303224 | 0.76171916 | 0.9428423 |
| Osbpl1a | 672.781322 | 0.1603673 | 0.3124075 | 0.51332732 | 0.60772238 | 0.89332706 |
| Ctla2a | 83.0650569 | 1.88742715 | 0.66536947 | 2.83666028 | 0.00455881 | 0.07415946 |
| Gm1821 | 9859.17152 | 0.04997009 | 0.18907352 | 0.26428918 | 0.7915571 | 0.95268772 |
| Ubr3 | 261.511333 | -0.0109002 | 0.29022341 | -0.037558 | 0.97004013 | 0.99443087 |
| Trp53i13 | 264.569994 | -0.2129468 | 0.31226428 | -0.6819443 | 0.4952742 | 0.84713217 |
| Aplnr | 554.224114 | 3.43833579 | 1.35200658 | 2.54313539 | 0.01098626 | 0.13620785 |
| Alkbh2 | 266.981023 | 0.04246845 | 0.40343363 | 0.1052675 | 0.91616355 | 0.98386323 |
| Phlpp1 | 32.7968413 | 0.30830526 | 0.9184415 | 0.33568306 | 0.73710988 | 0.93800224 |
| Slc25a53 | 137.700669 | -0.789054 | 0.39606721 | -1.9922226 | 0.04634664 | 0.33291745 |
| Snhg11 | 27.2913112 | 1.29185935 | 1.29028677 | 1.00121878 | 0.31672105 | 0.74186126 |
| Lacc1 | 347.187296 | 0.40213861 | 0.37193307 | 1.08121229 | 0.27960269 | 0.71436503 |
| Ccdc89 | 28.9748175 | 2.34524903 | 1.72512851 | 1.35946337 | 0.1739998 | 0.60653633 |
| Slc16a13 | 719.729976 | -0.0204994 | 0.26097048 | -0.0785505 | 0.93739015 | 0.98845735 |
| Tigd3 | 14.7279822 | -1.288143 | 1.98725836 | -0.6482011 | 0.51685492 | 0.85673255 |
| Sptssa | 4138.77123 | 0.50813861 | 0.39518551 | 1.28582299 | 0.19850484 | 0.63122167 |
| Gm9493 | 19.2881625 | -0.1917162 | 0.86220904 | -0.2223547 | 0.82403781 | 0.96290555 |
| N6amt1 | 526.062459 | 0.20011883 | 0.31825356 | 0.62880311 | 0.52947796 | 0.86283637 |
| Dock5 | 48.6921198 | -3.7575013 | 1.3012219 | -2.8876715 | 0.00388105 | 0.06537715 |
| Rin3 | 731.852108 | -0.918471 | 0.25326577 | -3.6265106 | 0.00028728 | 0.00861338 |

| | | | | | | |
|---|---|---|---|---|---|---|
| Fam160a2 | 416.904659 | -0.0698722 | 0.37651382 | -0.1855767 | 0.85277673 | 0.96798445 |
| Tent5c | 72.0999822 | -1.7414609 | 0.91946069 | -1.8940026 | 0.05822467 | 0.37058394 |
| Tnfaip8l1 | 334.781095 | 1.26883064 | 0.64372187 | 1.97108519 | 0.04871413 | 0.34046898 |
| Ascc1 | 893.137444 | 0.17631509 | 0.29564137 | 0.59638163 | 0.55092031 | 0.87093412 |
| Zfand3 | 219.762223 | 0.10418818 | 0.37968963 | 0.27440353 | 0.78377453 | 0.95094799 |
| 2510039O18 | 350.429973 | -0.6350018 | 0.34910484 | -1.818943 | 0.06892012 | 0.40475269 |
| Zfp758 | 276.968796 | 0.53705679 | 0.32403445 | 1.65740647 | 0.09743732 | 0.47887037 |
| Bod1 | 49.5509625 | -0.219888 | 0.56075827 | -0.3921262 | 0.69496498 | 0.92609254 |
| Tram1l1 | 17.6996645 | 0.87418417 | 1.87834511 | 0.46540125 | 0.64164414 | 0.90700632 |
| Rps2 | 65275.1138 | -0.2079554 | 0.21799336 | -0.9539527 | 0.34010759 | 0.75756302 |
| 9930012K11I | 286.124041 | 1.07155721 | 0.40012891 | 2.67802995 | 0.00740566 | 0.10532015 |
| Rasip1 | 39.4785264 | 1.75592915 | 1.33536234 | 1.31494584 | 0.18852811 | 0.61935434 |
| Cage1 | 15.163409 | 0.78626578 | 1.74462587 | 0.45067873 | 0.65222111 | 0.91037383 |
| Acp1 | 6765.80864 | 0.24064803 | 0.2775821 | 0.86694362 | 0.3859729 | 0.7886896 |
| Tlr7 | 1477.66401 | 0.71206429 | 0.31723735 | 2.244579 | 0.02479517 | 0.22874762 |
| Dnd1 | 35.3126009 | 0.9719759 | 1.30549936 | 0.74452422 | 0.45655939 | 0.82915295 |
| Smim7 | 4461.7894 | 0.67605654 | 0.30930755 | 2.18570974 | 0.02883685 | 0.25269424 |
| Zbtb39 | 20.1555595 | -0.9200331 | 0.97568029 | -0.9429657 | 0.34569844 | 0.7602916 |
| Liph | 45.0094205 | 2.56188586 | 1.19030857 | 2.15228716 | 0.03137474 | 0.26487733 |
| Swi5 | 4173.24888 | 0.26120015 | 0.35409851 | 0.73764826 | 0.46072822 | 0.82987212 |
| Cnrip1 | 466.402029 | 1.61500266 | 0.28689657 | 5.62921568 | 1.810309096 | 2.171128547 |
| Csrnp2 | 77.389494 | 0.26882302 | 0.82079854 | 0.32751401 | 0.74327915 | 0.93856621 |
| Plppr4 | 204.465257 | -1.3902439 | 0.5308399 | -2.6189514 | 0.00882005 | 0.11905611 |
| Fzd1 | 18.616769 | -1.4278192 | 1.93906328 | -0.7363448 | 0.46152089 | 0.83025001 |
| Zfp612 | 9.86476119 | 0.51403725 | 2.16186787 | 0.23777459 | 0.81205593 | 0.95967622 |
| Tmem201 | 876.411524 | -0.3187028 | 0.27063439 | -1.1776139 | 0.23895055 | 0.67507669 |
| Palb2 | 753.750486 | 0.2179414 | 0.22715763 | 0.95942805 | 0.33734315 | 0.75598717 |
| Phf11a | 55.6273512 | -0.9134751 | 0.56908279 | -1.6051709 | 0.10845621 | 0.50301538 |
| Gemin7 | 2510.41724 | 0.70759306 | 0.39758819 | 1.77971346 | 0.07512287 | 0.42213047 |
| Slc38a6 | 851.360709 | 0.90239324 | 0.2440838 | 3.69706324 | 0.00021811 | 0.00700293 |
| Gskip | 1671.45597 | 0.2009787 | 0.32605669 | 0.61639189 | 0.53763591 | 0.86640759 |
| 9930104L06F | 286.126572 | 0.21364191 | 0.3801809 | 0.56194803 | 0.57415143 | 0.88059206 |
| Serpinb1a | 911.689607 | -2.6916699 | 0.34160521 | -7.8794755 | 3.287581337 | 1.340566083 |
| Gm6430 | 13.9270871 | -0.7645112 | 1.17325123 | -0.6516176 | 0.51464786 | 0.85526675 |
| Trmt10c | 1667.67912 | -0.2039631 | 0.21314589 | -0.9569178 | 0.33860874 | 0.7564609 |
| D1Ertd622e | 2849.00629 | -0.1098111 | 0.15199733 | -0.7224544 | 0.47001516 | 0.83381643 |
| Hjurp | 4066.27965 | 0.22761145 | 0.1554447 | 1.46425997 | 0.14312293 | 0.56846902 |
| Zfp36 | 9559.00104 | 0.49980028 | 0.37688406 | 1.32613802 | 0.18479399 | 0.61700012 |
| Fads6 | 22.10951 | 0.17747411 | 1.29604761 | 0.13693487 | 0.89108228 | 0.97889088 |
| Setd2 | 925.675313 | -0.3715773 | 0.40842584 | -0.9097791 | 0.36293902 | 0.76990376 |
| Isca1 | 2791.11602 | -0.0589898 | 0.17720936 | -0.3328822 | 0.73922321 | 0.93800224 |
| Cyb5d1 | 309.866983 | 0.11831335 | 0.31313759 | 0.37783183 | 0.70555553 | 0.92896305 |
| Zfp354c | 21.4812748 | -1.9463624 | 1.56333966 | -1.2450029 | 0.2131307 | 0.64599096 |
| Cd300c2 | 1716.02176 | 1.30613325 | 0.45711859 | 2.85731814 | 0.00427237 | 0.07110741 |
| Shb | 12.9311666 | 1.53851076 | 1.54984109 | 0.99268936 | 0.32086136 | 0.74413737 |
| D630023F18 | 179.34501 | 0.03801774 | 0.5313538 | 0.07154883 | 0.94296096 | 0.98983253 |

| | | | | | | |
|---|---|---|---|---|---|---|
| Tlr1 | 1721.11712 | 0.86734687 | 0.29934992 | 2.89743476 | 0.00376228 | 0.06410023 |
| Lsm11 | 96.9101988 | -0.1465947 | 0.62293674 | -0.2353284 | 0.81395382 | 0.95988497 |
| 1700056E22I | 15.016034 | -0.3858402 | 1.27035757 | -0.3037256 | 0.76133692 | 0.94247845 |
| Lemd2 | 136.280032 | 0.27160869 | 0.43058684 | 0.63078725 | 0.52817964 | 0.86198732 |
| Ankrd50 | 109.105066 | 0.51641648 | 1.29919965 | 0.39748816 | 0.69100752 | 0.92470267 |
| Zfp444 | 864.45717 | 0.23779238 | 0.21972799 | 1.0822125 | 0.27915812 | 0.71377266 |
| Coa4 | 1198.05745 | 0.85010145 | 0.41972306 | 2.02538656 | 0.04282769 | 0.31713747 |
| Uqcrq | 4419.59746 | 0.50640093 | 0.43245508 | 1.17099082 | 0.24160247 | 0.67708722 |
| 4930503L19F | 801.42029 | 0.05371588 | 0.28186171 | 0.19057528 | 0.84885836 | 0.96765585 |
| Zfp367 | 1492.58647 | 0.02377375 | 0.32446329 | 0.07327099 | 0.94159047 | 0.98948137 |
| Ttc41 | 13.6872923 | 0.51300642 | 2.54234319 | 0.20178488 | 0.8400849 | 0.96765585 |
| Ubtd2 | 130.603806 | 0.32253904 | 0.38023401 | 0.84826458 | 0.39629064 | 0.79439542 |
| Pwwp2a | 102.703592 | 0.19701776 | 0.61933934 | 0.31810955 | 0.75040184 | 0.9399191 |
| Kctd21 | 157.633142 | 0.55401339 | 0.66183616 | 0.8370854 | 0.40254456 | 0.79795242 |
| Fbxo48 | 54.3054871 | 0.66333175 | 0.90369283 | 0.73402348 | 0.46293443 | 0.83156884 |
| Napepld | 195.930721 | 0.66879726 | 0.45634536 | 1.46555072 | 0.14277073 | 0.56834178 |
| Sft2d3 | 180.313242 | -0.7238636 | 0.35706508 | -2.0272595 | 0.04263588 | 0.31638902 |
| Tst | 97.8490935 | -0.2821805 | 0.53360526 | -0.5288189 | 0.59693111 | 0.88891958 |
| Shld1 | 443.60651 | 0.6434724 | 0.48180492 | 1.33554552 | 0.18169787 | 0.61290438 |
| Fzd5 | 33.9715479 | -0.0455521 | 1.12186192 | -0.0406041 | 0.96761155 | 0.99443087 |
| Cetn4 | 24.86981 | -0.2095808 | 0.86031858 | -0.2436084 | 0.80753412 | 0.95797316 |
| Prkce | 29.6573521 | 2.12226378 | 0.99452177 | 2.13395406 | 0.03284654 | 0.26964673 |
| Megf8 | 149.002478 | -0.0645921 | 0.45822819 | -0.1409605 | 0.88790112 | 0.97832097 |
| Rpsa-ps2 | 18.3049323 | -0.0338331 | 0.92446727 | -0.0365974 | 0.97080599 | 0.99443087 |
| E130308A19I | 142.641936 | -0.776596 | 0.80968692 | -0.9591312 | 0.33749267 | 0.75611716 |
| Rnf216 | 1452.71773 | 0.31307801 | 0.25458469 | 1.22975977 | 0.21878708 | 0.65230863 |
| S1pr1 | 725.810159 | -0.0175039 | 0.27443242 | -0.0637823 | 0.94914355 | 0.99028544 |
| Arhgef37 | 20.430079 | -1.6723213 | 1.45667708 | -1.1480385 | 0.25095268 | 0.6842534 |
| Magi1 | 54.2633058 | 0.46648255 | 0.59030811 | 0.79023572 | 0.42939012 | 0.81155572 |
| Kmt5b | 404.667918 | -0.4022398 | 0.40423155 | -0.9950728 | 0.31970087 | 0.7440831 |
| Slc25a26 | 397.500781 | 0.03563422 | 0.26993218 | 0.13201174 | 0.89497501 | 0.9794437 |
| Poln | 73.4904562 | 1.8608187 | 0.81909063 | 2.2718105 | 0.02309796 | 0.21802263 |
| Saysd1 | 466.505938 | 0.69030227 | 0.45612032 | 1.51342145 | 0.13017264 | 0.54740526 |
| Rpl18a | 34474.5043 | -0.1405276 | 0.20511772 | -0.685107 | 0.49327643 | 0.84635172 |
| Tubb2b | 232.325278 | 0.94945774 | 0.54264037 | 1.74969979 | 0.08017013 | 0.43723638 |
| Pigw | 94.5648779 | -1.1424175 | 0.60654137 | -1.8834947 | 0.05963336 | 0.37680521 |
| Bola3 | 1437.99521 | 0.26047106 | 0.41193347 | 0.63231343 | 0.52718209 | 0.86136217 |
| AI467606 | 1603.62078 | -0.054408 | 0.30664347 | -0.1774309 | 0.85916997 | 0.96954641 |
| Borcs6 | 229.645121 | 0.3251513 | 0.32179119 | 1.0104419 | 0.31228362 | 0.74051791 |
| Shroom2 | 19.5577198 | 1.36996899 | 1.91762407 | 0.71440957 | 0.47497396 | 0.83578199 |
| Cirbp | 537.228497 | -0.2332495 | 0.27796951 | -0.8391192 | 0.40140245 | 0.79737342 |
| Dpy19l4 | 475.58481 | 0.01624308 | 0.36111866 | 0.04497988 | 0.96412334 | 0.99406401 |
| Vcpip1 | 851.451284 | 0.3106643 | 0.26332309 | 1.17978372 | 0.23808624 | 0.67403588 |
| Nudt18 | 362.2824 | 0.13114711 | 0.28828755 | 0.4549177 | 0.64916845 | 0.90973357 |
| Hs6st1 | 109.981978 | -0.8752291 | 0.61848202 | -1.4151246 | 0.15703197 | 0.58620126 |
| 1110012L19F | 561.616411 | 0.50954414 | 0.42281206 | 1.20513153 | 0.22815253 | 0.66215656 |

| | | | | | | |
|---|---|---|---|---|---|---|
| Med26 | 480.258523 | -0.0946722 | 0.25071982 | -0.3776015 | 0.70572663 | 0.92896305 |
| Zfp688 | 264.143312 | 0.39798998 | 0.45733812 | 0.87023139 | 0.38417397 | 0.78704643 |
| Zfp574 | 759.666305 | -0.4133484 | 0.24985427 | -1.654358 | 0.09805479 | 0.47978865 |
| Morn2 | 173.866428 | 0.5139331 | 0.44032987 | 1.16715475 | 0.24314788 | 0.67814218 |
| Zfp691 | 103.797379 | -0.6475747 | 0.5225793 | -1.2391894 | 0.21527535 | 0.64854028 |
| Cenph | 1343.57343 | -0.0942738 | 0.245563 | -0.3839088 | 0.70104603 | 0.92722415 |
| Lca5l | 15.049948 | 1.48428149 | 1.93968644 | 0.76521723 | 0.4441422 | 0.82068545 |
| Tmem86b | 9.12985392 | -0.8762832 | 1.5448153 | -0.5672414 | 0.57055017 | 0.88035336 |
| Rtn4rl1 | 23.429316 | -0.9107483 | 1.21840361 | -0.7474931 | 0.45476599 | 0.82760373 |
| Insig1 | 3923.28681 | 0.56456778 | 0.25727692 | 2.1943973 | 0.02820686 | 0.24951635 |
| Preb | 4723.33491 | -0.3335287 | 0.2186573 | -1.5253491 | 0.12717206 | 0.54111158 |
| Lhfpl2 | 159.639382 | 1.85920231 | 1.80780559 | 1.02843045 | 0.30374739 | 0.73436097 |
| Fahd1 | 296.015973 | 0.44568705 | 0.27469723 | 1.62246647 | 0.10470351 | 0.49501875 |
| Tlr9 | 144.74469 | 0.17205103 | 0.56960866 | 0.30205129 | 0.76261296 | 0.94298237 |
| Fndc7 | 45.0946556 | 0.38720652 | 1.12535587 | 0.34407473 | 0.73079008 | 0.93653187 |
| 6330549D23 | 78.0990988 | -0.0219283 | 0.81144709 | -0.0270237 | 0.97844084 | 0.99483019 |
| Cenpe | 2455.72398 | -0.1989859 | 0.31403401 | -0.6336444 | 0.52631289 | 0.86109209 |
| Tnfrsf26 | 113.176349 | -0.4155264 | 0.54816352 | -0.7580337 | 0.44843081 | 0.82404297 |
| Wdr81 | 506.610473 | 0.32519759 | 0.41758336 | 0.7787609 | 0.43612057 | 0.81725842 |
| Tmem88 | 158.16174 | 1.06878823 | 0.46006623 | 2.32311821 | 0.0201728 | 0.20079243 |
| Pxt1 | 13.4056472 | -0.964249 | 1.65655054 | -0.5820825 | 0.56051114 | 0.87469256 |
| Olfr433 | 49.0982879 | -0.9682341 | 0.64975035 | -1.4901633 | 0.13618131 | 0.55833898 |
| Cxcr4 | 21.0814046 | 0.76013806 | 0.92285294 | 0.82368276 | 0.41011984 | 0.80117888 |
| Epcam | 15.5841054 | -0.6422363 | 2.01712059 | -0.3183926 | 0.75018712 | 0.9399191 |
| Kcnk13 | 343.241107 | 0.93170548 | 0.43324603 | 2.15052282 | 0.03151388 | 0.26539272 |
| Trim39 | 284.990979 | 0.16470903 | 0.51727889 | 0.31841437 | 0.75017064 | 0.9399191 |
| Akr1e1 | 1921.13686 | 0.77030913 | 0.32070315 | 2.40193812 | 0.01630847 | 0.17796742 |
| 2410002F23I | 1883.67868 | -0.3832313 | 0.21707232 | -1.7654545 | 0.0774874 | 0.43008322 |
| Dipk2a | 240.863156 | 0.65613035 | 0.4558729 | 1.43928352 | 0.15007021 | 0.57784353 |
| Hnrnph2 | 7069.08872 | -0.2490868 | 0.22151277 | -1.1244804 | 0.2608093 | 0.69691361 |
| Tmem60 | 934.851177 | 0.51616866 | 0.36040317 | 1.43219791 | 0.15208719 | 0.58009243 |
| Cox19 | 1400.4797 | 0.19265856 | 0.36582517 | 0.52664105 | 0.59844288 | 0.889626 |
| Gm9797 | 461.556151 | 0.49520344 | 0.51272281 | 0.96583072 | 0.33412889 | 0.75320272 |
| Zfp956 | 189.74749 | -0.652532 | 0.39653692 | -1.6455769 | 0.09985089 | 0.48298772 |
| Ttll13 | 13.6501099 | -0.2663941 | 1.68592258 | -0.1580109 | 0.87444825 | 0.97364644 |
| Trrap | 656.860549 | -0.0506986 | 0.26971671 | -0.1879699 | 0.85090025 | 0.9679302 |
| Hcar2 | 729.46079 | -1.7756931 | 0.49124932 | -3.6146475 | 0.00030076 | 0.00892999 |
| Sys1 | 3367.82901 | 0.21175964 | 0.24299291 | 0.87146426 | 0.38350071 | 0.78674564 |
| Gpr150 | 15.6441371 | 0.49024498 | 0.97378393 | 0.50344329 | 0.61465264 | 0.89844745 |
| Zfp560 | 301.360401 | 0.88941326 | 0.51643141 | 1.72222921 | 0.08502801 | 0.44637738 |
| Ddx28 | 814.475113 | -0.1565288 | 0.22802578 | -0.6864521 | 0.49242804 | 0.84569314 |
| Penk | 219.175627 | -1.2183236 | 0.35163808 | -3.464709 | 0.00053081 | 0.01408534 |
| St7l | 972.393565 | 0.08292735 | 0.32519255 | 0.25500998 | 0.79871539 | 0.9547279 |
| Glb1 | 3513.49816 | -0.065124 | 0.19695496 | -0.3306544 | 0.74090553 | 0.93800224 |
| Zfp553 | 283.442437 | -0.7448456 | 0.46408482 | -1.6049774 | 0.10849878 | 0.50301538 |
| Esf1 | 882.080607 | -0.2830463 | 0.37131881 | -0.762273 | 0.44589708 | 0.82166369 |

| | | | | | | |
|---|---|---|---|---|---|---|
| Pigz | 30.3995707 | 2.965218 | 1.15443776 | 2.56853864 | 0.01021283 | 0.13041084 |
| Mtus1 | 11798.9682 | -0.6329314 | 0.18672997 | -3.3895546 | 0.00070006 | 0.01773057 |
| Zfp629 | 110.341374 | -1.1753829 | 0.55753399 | -2.1081817 | 0.03501528 | 0.28075507 |
| Pid1 | 1090.79723 | -0.1527278 | 0.31759053 | -0.4808955 | 0.6305908 | 0.9036296 |
| Henmt1 | 16.3243589 | -1.9692883 | 1.70978638 | -1.1517745 | 0.24941377 | 0.68409835 |
| Cdc42ep2 | 114.796186 | 1.90513063 | 1.02385978 | 1.86073393 | 0.06278176 | 0.38690643 |
| Mfsd5 | 1219.99177 | 0.20257312 | 0.30844077 | 0.65676507 | 0.51133199 | 0.85373917 |
| Smtnl2 | 28.4270517 | -0.4132384 | 0.93707898 | -0.4409857 | 0.65922338 | 0.91280998 |
| Spred2 | 186.046823 | -0.2190892 | 0.46508201 | -0.4710766 | 0.63758606 | 0.90599527 |
| Pqlc3 | 514.69054 | -0.7153884 | 0.26551685 | -2.6943241 | 0.00705316 | 0.10256275 |
| Wdr89 | 580.989301 | -0.2003258 | 0.292166 | -0.6856574 | 0.49292917 | 0.84609645 |
| Thtpa | 393.682488 | 0.11596738 | 0.36993792 | 0.31347794 | 0.75391758 | 0.94018119 |
| Gm21411 | 20.8628923 | 0.3259793 | 0.92726667 | 0.3515486 | 0.72517682 | 0.93542036 |
| Smkr-ps | 34.6740227 | -0.1395102 | 0.86015193 | -0.1621925 | 0.87115425 | 0.97268995 |
| Adrb2 | 5278.90133 | 0.6370372 | 0.31048945 | 2.05171929 | 0.04019695 | 0.30448619 |
| Mms22l | 1314.5189 | -0.0149012 | 0.1835707 | -0.0811739 | 0.93530364 | 0.98794094 |
| Tssc4 | 2227.05435 | 0.11465953 | 0.27676558 | 0.41428392 | 0.67866619 | 0.92010678 |
| Zfp764 | 276.62464 | 0.63658985 | 0.45634826 | 1.394965 | 0.1630264 | 0.59424967 |
| Basp1 | 109.208837 | 1.12641254 | 0.53759075 | 2.09529746 | 0.03614456 | 0.28489457 |
| B230219D22 | 749.271759 | 0.32642815 | 0.3517828 | 0.92792527 | 0.35344634 | 0.76383312 |
| Olfr624 | 23.3807933 | 0.31212047 | 0.99219648 | 0.31457527 | 0.75308416 | 0.94018119 |
| Whamm | 44.2327331 | 0.60394801 | 0.65904387 | 0.91640032 | 0.35945697 | 0.76857232 |
| Gm9800 | 7.40190807 | -0.1119872 | 1.34813111 | -0.0830684 | 0.93379712 | 0.98756396 |
| Zfp36l2 | 946.072309 | -0.2354407 | 0.25143044 | -0.936405 | 0.34906468 | 0.7617085 |
| Zswim3 | 237.539079 | 0.7828348 | 0.54070983 | 1.44779094 | 0.14767553 | 0.57472133 |
| Ptprcap | 104.684319 | -3.0596655 | 0.68957016 | -4.437062 | 9.119502824 | 0.00047879 |
| Serpinb9 | 514.079741 | 1.12001269 | 0.39305675 | 2.8494936 | 0.00437889 | 0.07238776 |
| Lyrm2 | 700.858813 | 0.51994906 | 0.39354404 | 1.32119662 | 0.18643581 | 0.61862792 |
| Cradd | 307.613732 | 0.30177221 | 0.2709664 | 1.11368868 | 0.26541277 | 0.70125149 |
| Gvin1 | 372.907272 | -0.0243984 | 0.34349981 | -0.0710289 | 0.94337472 | 0.98989307 |
| Gm9803 | 223.663416 | 0.58438428 | 0.69827189 | 0.83690076 | 0.40264835 | 0.79795242 |
| Paip2b | 201.247832 | 0.10283953 | 0.38750352 | 0.26538992 | 0.7907091 | 0.95261385 |
| Tmem268 | 2398.97647 | 0.02381604 | 0.18609702 | 0.12797649 | 0.89816758 | 0.97989497 |
| Clec14a | 140.969442 | 4.43973199 | 2.03772997 | 2.17876365 | NA | NA |
| Ifit2 | 105.187024 | 0.75582229 | 0.523534 | 1.44369285 | 0.14882539 | 0.57595098 |
| Mtmr11 | 37.0403477 | 1.34582274 | 0.8514892 | 1.58055174 | 0.11398057 | 0.51527135 |
| BC049762 | 19.2599384 | 0.7046116 | 1.28491595 | 0.54837174 | 0.58343667 | 0.8838614 |
| Mrps12 | 2125.82452 | -0.1800163 | 0.2442885 | -0.7369005 | 0.46118284 | 0.83014269 |
| Wnk1 | 1619.0453 | 0.1670497 | 0.4530249 | 0.36874287 | 0.71231939 | 0.93235217 |
| Ing1 | 651.815085 | -0.0890386 | 0.19806112 | -0.4495513 | 0.65303399 | 0.9106891 |
| Slc25a51 | 1821.65357 | 0.14866167 | 0.20389149 | 0.72912154 | 0.46592732 | 0.83345004 |
| C2cd2 | 455.147926 | -0.2646603 | 0.5375711 | -0.4923262 | 0.62248879 | 0.89978794 |
| Tmem104 | 1631.10077 | 0.56495822 | 0.36215337 | 1.55999717 | 0.11876055 | 0.52515149 |
| Eif4g1 | 5518.55923 | -0.4247794 | 0.31651951 | -1.3420322 | 0.17958556 | 0.60992118 |
| Polr2k | 1142.23874 | 0.44767181 | 0.42987942 | 1.04138927 | 0.29769492 | 0.72979999 |
| Gapt | 422.156942 | -2.6045295 | 0.3286108 | -7.9258793 | 2.265374847 | 9.897260896 |

| | | | | | | |
|---|---|---|---|---|---|---|
| Zfp830 | 842.389524 | -0.3041826 | 0.21600743 | -1.4082044 | 0.15907055 | 0.58838861 |
| Pofut1 | 364.34597 | 0.01419678 | 0.38796589 | 0.03659285 | 0.97080964 | 0.99443087 |
| Stard5 | 1777.15831 | 0.76256024 | 0.19272893 | 3.95664637 | 7.600932243 | 0.00284349 |
| Calhm6 | 143.058376 | 0.35574689 | 0.51384429 | 0.69232431 | 0.48873369 | 0.8436846 |
| Snx12 | 1263.05088 | 0.23450024 | 0.19960355 | 1.17483001 | 0.24006273 | 0.67563574 |
| Otulin | 228.179829 | -0.5471046 | 0.34530727 | -1.5843992 | 0.1131029 | 0.51263346 |
| Sbsn | 8.63505014 | -0.3818659 | 1.40443853 | -0.2718993 | 0.78569942 | 0.95168899 |
| Ppp1r15b | 1670.47069 | -0.2089754 | 0.20743103 | -1.0074454 | 0.31372079 | 0.74065408 |
| Lrrc8d | 730.801695 | -0.2062203 | 0.34625427 | -0.5955749 | 0.55145928 | 0.87093412 |
| Mosmo | 59.5979632 | -0.4846723 | 0.71282487 | -0.6799318 | 0.49654764 | 0.84788767 |
| Mcmdc2 | 11.6816898 | 1.65042851 | 1.45742765 | 1.13242569 | 0.25745551 | 0.69274795 |
| Cep295 | 351.132 | -0.5626 | 0.42483614 | -1.3242753 | 0.18541164 | 0.61768533 |
| 9930021J03F | 73.7126433 | -1.5897245 | 0.89568395 | -1.7748722 | 0.075919 | 0.42465346 |
| Patl1 | 311.474161 | -0.2461288 | 0.57634334 | -0.4270524 | 0.66934119 | 0.91650445 |
| Fut10 | 454.352828 | 0.21958566 | 0.28373292 | 0.77391676 | 0.43898002 | 0.81963526 |
| Tmem229b | 2803.5859 | -0.7083086 | 0.22331189 | -3.1718355 | 0.00151479 | 0.03194896 |
| Olig1 | 6.08340443 | 0.40309785 | 2.32946022 | 0.17304346 | 0.86261727 | 0.9705375 |
| Adamts6 | 11.9568516 | 2.01648987 | 2.5022997 | 0.80585466 | 0.42032666 | 0.80643915 |
| E2f8 | 1053.29012 | -0.5345081 | 0.32300171 | -1.6548151 | 0.097962 | 0.47953948 |
| 4930550L24F | 23.5915047 | -1.188775 | 1.54543825 | -0.7692155 | 0.44176541 | 0.81967485 |
| Zfp84 | 620.288548 | 0.14338499 | 0.39987399 | 0.35857543 | 0.71991273 | 0.93328572 |
| Cd109 | 14.5241258 | 0.97206115 | 1.76653249 | 0.55026508 | 0.58213757 | 0.88321857 |
| Scaf8 | 1122.62949 | 0.15164168 | 0.29807801 | 0.50873153 | 0.61094042 | 0.89526044 |
| Pnma2 | 35.8501519 | 1.99859591 | 1.71462368 | 1.16561782 | 0.243769 | 0.67897148 |
| Pik3r6 | 1296.07018 | 0.26404159 | 0.20053234 | 1.31670324 | 0.18793812 | 0.61935434 |
| Plaur | 1412.27921 | 0.41488639 | 0.55886696 | 0.74237058 | 0.45786284 | 0.82986925 |
| Scand1 | 509.542012 | -0.0391054 | 0.25354342 | -0.1542357 | 0.87742393 | 0.97458574 |
| Vps13a | 278.255062 | -0.6751624 | 0.36980035 | -1.8257484 | 0.06788817 | 0.40139003 |
| Pilra | 1234.7578 | -1.0677224 | 0.33653012 | -3.1727394 | 0.00151008 | 0.03190468 |
| Ankle1 | 446.813997 | -0.1941858 | 0.27332658 | -0.7104533 | 0.47742307 | 0.836949 |
| Zfp62 | 869.953736 | 0.11970636 | 0.31454167 | 0.38057394 | 0.70351942 | 0.9283876 |
| Ermp1 | 781.923746 | 0.02810492 | 0.23465363 | 0.11977193 | 0.90466382 | 0.98132496 |
| Slc25a23 | 24.7822153 | 2.63627516 | 1.36574127 | 1.93028885 | 0.05357106 | 0.35852954 |
| Rpl37a | 9995.99098 | 0.08340482 | 0.34542707 | 0.2414542 | 0.80920311 | 0.95844313 |
| Zfp322a | 931.562746 | 0.2547474 | 0.32311128 | 0.78842001 | 0.43045107 | 0.81243065 |
| Rpl27a | 5554.06519 | 0.05222081 | 0.34157037 | 0.15288449 | 0.87848937 | 0.97518697 |
| Jrk | 63.1334375 | -0.1436778 | 0.70594674 | -0.203525 | 0.83872469 | 0.96744446 |
| Rbp1 | 73.6779882 | 0.84317523 | 0.83703569 | 1.00733486 | 0.31377388 | 0.74065408 |
| Yod1 | 726.294718 | 0.0359902 | 0.32900851 | 0.10938989 | 0.91289326 | 0.98292561 |
| Kcnk6 | 545.711015 | 0.1432003 | 0.30790059 | 0.46508615 | 0.64186976 | 0.90702012 |
| Lrrc75a | 102.539212 | 0.53905149 | 0.67840458 | 0.79458705 | 0.42685375 | 0.81021713 |
| Bex3 | 3965.36655 | 0.58358381 | 0.25492667 | 2.28922228 | 0.02206644 | 0.21355916 |
| Hnrnpa1 | 26064.861 | -0.5484741 | 0.16802731 | -3.2641959 | 0.00109775 | 0.02514758 |
| Cmtr2 | 251.57869 | 0.28654852 | 0.41516494 | 0.69020404 | 0.49006588 | 0.84448327 |
| Ppm1e | 11.6062427 | 0.09789276 | 1.05875005 | 0.09246069 | 0.92633202 | 0.98624627 |
| Sox18 | 73.2185344 | 2.65540174 | 1.3907764 | 1.9092945 | 0.05622411 | 0.36472442 |

| | | | | | | |
|---|---|---|---|---|---|---|
| Cox17 | 1496.99869 | -0.069334 | 0.36572735 | -0.1895783 | 0.84963963 | 0.96766052 |
| Golph3l | 1574.1106 | 0.12541239 | 0.20108333 | 0.62368367 | 0.53283535 | 0.8646427 |
| Kctd4 | 25.7260829 | 2.54719342 | 1.70523704 | 1.49374741 | 0.13524165 | 0.55685329 |
| Zfp526 | 23.6462644 | 0.40928785 | 1.1260896 | 0.3634594 | 0.71626174 | 0.93235217 |
| Fam43a | 347.643429 | 0.22065776 | 0.54377205 | 0.40579092 | 0.68489623 | 0.92314423 |
| Spin2c | 41.2336222 | 0.71772453 | 0.71339714 | 1.00606588 | 0.31438388 | 0.74065408 |
| Zfp319 | 192.138884 | 0.93157415 | 0.53172633 | 1.75198046 | 0.07977717 | 0.43686807 |
| Unc119b | 1112.23841 | 0.0200181 | 0.23821508 | 0.08403373 | 0.93302962 | 0.98748011 |
| 4930430F08I | 189.22576 | 0.16971131 | 0.32199824 | 0.52705663 | 0.59815427 | 0.88951331 |
| Zfp518b | 55.9794794 | -0.6998306 | 1.22581343 | -0.5709112 | 0.56805985 | 0.87929304 |
| Lyrm4 | 1976.39903 | 0.08151527 | 0.29986952 | 0.27183578 | 0.78574829 | 0.95168899 |
| Prr12 | 75.7675547 | -0.221624 | 0.76746055 | -0.2887758 | 0.77275296 | 0.94625031 |
| Ticrr | 457.730846 | -0.8000395 | 0.23976626 | -3.3367477 | 0.00084765 | 0.02053323 |
| Bdh1 | 11.3758524 | -3.2732379 | 1.35859609 | -2.4092796 | 0.01598405 | 0.17615771 |
| Tcaim | 114.736468 | -0.2068083 | 0.4867147 | -0.4249066 | 0.67090474 | 0.91711586 |
| B3gntl1 | 279.917935 | 0.17496228 | 0.26872575 | 0.65108117 | 0.51499409 | 0.85538665 |
| Zfp316 | 32.8109427 | 0.43480489 | 0.91745095 | 0.47392713 | 0.63555187 | 0.90531997 |
| Rbm12b1 | 165.75735 | 0.36607277 | 0.49033806 | 0.74657221 | 0.45532183 | 0.82788614 |
| Cxxc5 | 169.726977 | 0.20044199 | 0.39363153 | 0.50921222 | 0.61060348 | 0.89508835 |
| Mtfr1l | 3692.81546 | -0.0446082 | 0.20113043 | -0.2217876 | 0.82447923 | 0.962942 |
| Tmem251 | 691.43806 | 0.1631761 | 0.30658683 | 0.53223453 | 0.59456357 | 0.88828726 |
| C87436 | 600.285 | 0.20942674 | 0.30978699 | 0.67603466 | 0.49901864 | 0.84872332 |
| Tifa | 3300.55093 | -0.0275224 | 0.28825183 | -0.0954804 | 0.92393322 | 0.9849488 |
| Chtf8 | 1429.13355 | -0.1456723 | 0.2277902 | -0.6395022 | 0.5224963 | 0.85816354 |
| Csnk2a2 | 704.828974 | -0.6671006 | 0.27014023 | -2.4694604 | 0.0135317 | 0.1574243 |
| Hmga1 | 18951.4526 | -0.5380875 | 0.20742235 | -2.5941635 | 0.00948214 | 0.12499462 |
| Bst2 | 2163.1559 | 0.15157647 | 0.28231107 | 0.53691295 | 0.59132774 | 0.88792087 |
| Rpl14-ps1 | 12.1707923 | 2.01185181 | 1.19415041 | 1.68475579 | 0.0920357 | 0.46485248 |
| Cdc42se1 | 5535.09079 | 0.19451476 | 0.14333399 | 1.35707347 | 0.17475787 | 0.60698835 |
| Kctd11 | 131.490164 | 1.0741165 | 0.69987064 | 1.53473575 | 0.12484875 | 0.53626132 |
| Gprc5a | 29.3248226 | 0.14600658 | 1.63696695 | 0.08919336 | 0.92892824 | 0.98669576 |
| Ccdc66 | 374.718603 | 0.12729156 | 0.24255542 | 0.52479372 | 0.5997266 | 0.8902352 |
| Mrps7 | 4419.36211 | 0.00111654 | 0.29352138 | 0.00380394 | 0.99696491 | 0.99942546 |
| Fam83h | 12.8401989 | 6.18511737 | 1.72905512 | 3.5771661 | 0.00034734 | 0.00995083 |
| Rhoj | 132.975395 | 1.25338696 | 0.46781916 | 2.67921256 | 0.00737955 | 0.10509205 |
| Epm2aip1 | 240.241203 | 0.71760384 | 0.35539422 | 2.01917702 | 0.04346882 | 0.31959733 |
| Riox1 | 593.294571 | -0.4675046 | 0.46154957 | -1.0129022 | 0.31110686 | 0.74011268 |
| Zfp787 | 364.233232 | 0.580803 | 0.33387876 | 1.73956257 | 0.08193586 | 0.44107633 |
| Ppp1r3b | 576.717335 | 0.54387097 | 0.4388546 | 1.23929649 | 0.21523572 | 0.64854028 |
| Cldn12 | 224.78594 | -0.6009988 | 0.5091484 | -1.1804 | 0.23784117 | 0.67403588 |
| Mpeg1 | 660.217369 | -0.299772 | 0.24311992 | -1.2330211 | 0.2175679 | 0.65150232 |
| Cyren | 1077.51862 | -0.1110664 | 0.23048001 | -0.4818915 | 0.62988304 | 0.90321876 |
| Atp10d | 58.6132409 | -1.6466242 | 0.77681521 | -2.1197116 | 0.03403037 | 0.27569109 |
| Gchfr | 24.4249143 | 1.99178784 | 0.97210245 | 2.04894847 | 0.04046715 | 0.30576571 |
| Slc39a3 | 650.150442 | 0.09643638 | 0.3445227 | 0.279913 | 0.77954425 | 0.9488468 |
| Brox | 2788.87838 | 0.41867135 | 0.20136246 | 2.07919267 | 0.03759965 | 0.29222138 |

| | | | | | | |
|---|---|---|---|---|---|---|
| Ckap4 | 643.698646 | -0.0075578 | 0.32449541 | -0.0232908 | 0.9814183 | 0.99523682 |
| Hectd3 | 964.29911 | 0.02618501 | 0.27326509 | 0.09582275 | 0.92366135 | 0.98493282 |
| Pramef8 | 1308.32219 | 0.48985194 | 0.22870833 | 2.14181942 | 0.03220802 | 0.2665769 |
| Fbl | 3305.70736 | -0.5090504 | 0.15732829 | -3.2355937 | 0.0012139 | 0.02711313 |
| Mbtps2 | 525.074897 | -0.1026451 | 0.3583667 | -0.2864246 | 0.77455291 | 0.94709254 |
| Atxn1 | 199.205985 | 1.00248096 | 0.67015371 | 1.49589706 | 0.13468048 | 0.55585235 |
| Irgm1 | 1072.79134 | 0.13387538 | 0.27919925 | 0.47949763 | 0.63158466 | 0.90399255 |
| Zfp740 | 365.252047 | -0.1616501 | 0.45314863 | -0.3567265 | 0.72129653 | 0.93398714 |
| Ltb4r1 | 187.641008 | -2.5997388 | 0.50365963 | -5.1616977 | 2.447202559 | 2.169320935 |
| Tefm | 986.491361 | 0.43229512 | 0.53070698 | 0.8145646 | 0.41532158 | 0.80395688 |
| Adck2 | 492.167417 | 0.0054907 | 0.28530368 | 0.01924511 | 0.98464557 | 0.99564261 |
| Nqo2 | 2717.34141 | 0.43840833 | 0.33692801 | 1.30119287 | 0.19319245 | 0.624687 |
| Zbtb21 | 1010.61739 | 0.41618258 | 0.29511553 | 1.41023609 | 0.15846998 | 0.58731982 |
| Tshz1 | 124.045313 | -0.8196366 | 0.92067547 | -0.8902557 | 0.37332862 | 0.77748705 |
| Tapt1 | 89.3405605 | -0.8265392 | 0.55709052 | -1.4836712 | 0.13789617 | 0.56068131 |
| Wdr27 | 11.7803263 | 2.06993195 | 1.2225772 | 1.69308894 | 0.09043854 | 0.45983389 |
| Mars2 | 829.954369 | -0.3174557 | 0.3485346 | -0.9108297 | 0.36238512 | 0.76969598 |
| Spsb4 | 28.4862376 | -2.0480274 | 1.37032635 | -1.4945545 | 0.13503075 | 0.55654288 |
| Zfp41 | 327.46632 | 0.12245518 | 0.42451199 | 0.28846107 | 0.77299383 | 0.94625874 |
| Mipol1 | 163.203066 | -0.5755531 | 0.46396541 | -1.2405087 | 0.21478731 | 0.64854028 |
| Spata2 | 611.1548 | -0.2047649 | 0.3968687 | -0.5159512 | 0.60588851 | 0.89258167 |
| Zfp445 | 626.764912 | -0.0162207 | 0.49179559 | -0.0329827 | 0.97368842 | 0.99443087 |
| Nipa1 | 39.8774253 | -0.1360484 | 0.79083677 | -0.172031 | 0.86341316 | 0.9705375 |
| D030056L22I | 575.694206 | -0.1392508 | 0.40882879 | -0.340609 | 0.73339795 | 0.93728136 |
| Tmem164 | 2393.80828 | 0.04339866 | 0.2460414 | 0.17638763 | 0.85998942 | 0.96954641 |
| Dusp28 | 61.1343647 | -0.6213379 | 0.58480826 | -1.0624644 | 0.28802492 | 0.72304716 |
| Ngrn | 867.075148 | 0.11060556 | 0.37546388 | 0.29458375 | 0.76831188 | 0.94401697 |
| Tmem198b | 325.612433 | 0.62553017 | 0.39625328 | 1.57861196 | 0.11442509 | 0.51670806 |
| Rnf31 | 573.322129 | -0.2860556 | 0.32868241 | -0.8703101 | 0.38413096 | 0.78704643 |
| Dnajb7 | 7.41524258 | -0.0449801 | 2.79708418 | -0.0160811 | 0.98716972 | 0.99620914 |
| Fam221a | 63.2455731 | 0.51166258 | 0.77587667 | 0.65946381 | 0.50959798 | 0.85335831 |
| Ticam1 | 602.715042 | -0.0832891 | 0.4006757 | -0.2078716 | 0.8353292 | 0.96641606 |
| Cltc | 6493.83082 | 0.15806911 | 0.28038662 | 0.56375411 | 0.57292149 | 0.88035336 |
| Cd24a | 153.060708 | 0.76816391 | 1.06349815 | 0.72229924 | 0.47011052 | 0.83381643 |
| Zfp654 | 134.810238 | 0.73800264 | 0.5085675 | 1.45113999 | 0.14674088 | 0.57365079 |
| Tet1 | 52.9454541 | -0.8994627 | 0.89671325 | -1.0030661 | 0.31582896 | 0.74186126 |
| Khnyn | 440.466418 | -0.1651821 | 0.27344249 | -0.6040836 | 0.54578806 | 0.86990153 |
| Neurl3 | 1342.19069 | -0.2214855 | 0.27867701 | -0.7947749 | 0.42674446 | 0.81021713 |
| Rab2a | 950.578109 | 0.11462629 | 0.27442041 | 0.41770324 | 0.6761641 | 0.91885309 |
| Dync2h1 | 127.293562 | -0.7329214 | 0.92104148 | -0.7957529 | 0.4261757 | 0.81021713 |
| Dusp18 | 415.768046 | -0.1827578 | 0.40211287 | -0.4544938 | 0.64947345 | 0.90973357 |
| Ythdf3 | 1574.18246 | 0.16278483 | 0.28368813 | 0.57381616 | 0.56609221 | 0.87858116 |
| Rpl9 | 29811.2298 | 0.59278709 | 0.49236449 | 1.20395989 | 0.22860508 | 0.66258779 |
| Fam185a | 147.507677 | 0.06163847 | 0.57225571 | 0.10771142 | 0.91422461 | 0.98344264 |
| Taf9b | 309.590903 | -0.5174618 | 0.31210994 | -1.6579473 | 0.09732811 | 0.47854291 |
| Hist1h2be | 48.5338263 | 0.98083353 | 0.87463546 | 1.1214198 | 0.26210921 | 0.69762406 |

| | | | | | | |
|---|---|---|---|---|---|---|
| C2cd3 | 381.191501 | -0.7632376 | 0.53238404 | -1.4336222 | 0.15168009 | 0.57985599 |
| Ptgs1 | 14970.9319 | -0.3925741 | 0.18735104 | -2.0953934 | 0.03613604 | 0.28489457 |
| Emc6 | 3883.31006 | -0.121589 | 0.25933541 | -0.4688485 | 0.63917792 | 0.90632643 |
| Gap43 | 167.227376 | 2.97244025 | 0.73604447 | 4.03839766 | 5.381756350 | 0.00213549 |
| Zfp358 | 201.970246 | 0.70646476 | 0.42276085 | 1.67107424 | 0.09470701 | 0.47133886 |
| Sfn | 10.6937694 | -1.5692835 | 1.66047689 | -0.94508 | 0.34461802 | 0.75972469 |
| Neurl4 | 185.560525 | -0.2239838 | 0.54236585 | -0.4129754 | 0.67962462 | 0.92051478 |
| Gpr15 | 16.0766904 | -1.9462703 | 1.7986244 | -1.0820882 | 0.27921332 | 0.71377266 |
| Zfp286 | 30.1409301 | 0.43589422 | 0.78592885 | 0.55462301 | 0.57915255 | 0.88286755 |
| Abhd17b | 171.911763 | 0.04028578 | 0.38395877 | 0.10492216 | 0.91643758 | 0.98386323 |
| Dnah14 | 172.893984 | -0.6566855 | 0.37980235 | -1.729019 | 0.08380569 | 0.44417933 |
| Zfp768 | 144.370242 | -0.6418232 | 0.44034696 | -1.4575398 | 0.14496744 | 0.57067981 |
| B4gat1 | 883.249921 | 0.13627094 | 0.23655164 | 0.57607266 | 0.56456607 | 0.87721793 |
| Atmin | 574.92768 | 0.12749204 | 0.30232797 | 0.42170112 | 0.67324319 | 0.91786342 |
| Tgif1 | 2195.57386 | 0.07399248 | 0.54412623 | 0.13598402 | 0.89183391 | 0.97927786 |
| Ctdspl | 229.893585 | -0.5868356 | 0.46700015 | -1.2566069 | 0.20889601 | 0.64228937 |
| Zbtb44 | 256.239364 | 0.25912439 | 0.43269896 | 0.59885604 | 0.54926889 | 0.87093412 |
| Flrt2 | 66.7229424 | 2.90973671 | 1.52594376 | 1.90684401 | 0.05654079 | 0.36524665 |
| Rexo1 | 844.622362 | -0.1879584 | 0.32078932 | -0.5859248 | 0.55792605 | 0.87291829 |
| AI837181 | 87.3852731 | -0.7109643 | 0.59798714 | -1.1889291 | 0.23446757 | 0.67038085 |
| Xxylt1 | 204.900841 | -0.0693177 | 0.39285417 | -0.1764465 | 0.85994318 | 0.96954641 |
| Arl4a | 685.65765 | 0.86489768 | 0.38893108 | 2.22378133 | 0.02616316 | 0.23690151 |
| Gphn | 658.104016 | -0.0564067 | 0.30626018 | -0.1841792 | 0.85387289 | 0.96839628 |
| Dynlrb1 | 3993.9533 | -0.0552108 | 0.20720959 | -0.2664492 | 0.78989326 | 0.95255415 |
| 8030462N17 | 1011.74618 | 0.08738317 | 0.22658984 | 0.38564471 | 0.69975982 | 0.92669304 |
| Zfp30 | 37.8093675 | 1.37887802 | 0.73488667 | 1.87631383 | 0.0606122 | 0.37940675 |
| Adamts12 | 10.8932287 | 1.90513369 | 2.20835016 | 0.86269547 | 0.38830492 | 0.78942657 |
| Tspyl1 | 3803.02138 | -0.152467 | 0.22422916 | -0.6799606 | 0.49652939 | 0.84788767 |
| BC049715 | 139.016809 | -0.6391824 | 0.69114784 | -0.9248128 | 0.35506328 | 0.76456418 |
| Mis18bp1 | 1777.25059 | -0.1345312 | 0.21572005 | -0.6236381 | 0.53286531 | 0.8646427 |
| Fbxo28 | 689.602099 | 0.1228406 | 0.53884147 | 0.2279717 | 0.81966824 | 0.96136161 |
| Cltb | 1179.00838 | 0.11301591 | 0.20405583 | 0.55384798 | 0.57968289 | 0.88286755 |
| Tmem41b | 2974.34493 | 0.66388118 | 0.19483407 | 3.40741832 | 0.00065581 | 0.0168894 |
| Lxn | 1472.4165 | 0.48240749 | 0.21426195 | 2.25148461 | 0.02435486 | 0.22553596 |
| Tyw3 | 234.081616 | 0.22007104 | 0.32679872 | 0.67341463 | 0.50068354 | 0.84968141 |
| Zfp235 | 178.45842 | 0.40194449 | 0.32901861 | 1.22164667 | 0.22184127 | 0.65525654 |
| Frat2 | 9.75515231 | -1.2909219 | 1.58952202 | -0.8121447 | 0.4167086 | 0.8047849 |
| A430005L14F | 3079.06087 | 0.5408415 | 0.39530301 | 1.36816943 | 0.17125905 | 0.60430173 |
| Paxx | 160.466362 | 0.21706978 | 0.48578784 | 0.44684071 | 0.65499008 | 0.91113186 |
| Fgfbp3 | 49.374343 | -0.3130198 | 0.67855992 | -0.4613002 | 0.64458323 | 0.908327 |
| 2810006K23I | 856.21219 | 0.19261255 | 0.23025094 | 0.83653316 | 0.40285503 | 0.79795242 |
| Fbxo30 | 1308.36768 | 0.28434477 | 0.2472188 | 1.15017456 | 0.25007198 | 0.6842534 |
| Cd3eap | 703.135394 | -0.3993542 | 0.23291967 | -1.7145575 | 0.08642638 | 0.44913081 |
| Tssk6 | 66.3442978 | -0.0361249 | 0.62548033 | -0.0577554 | 0.95394347 | 0.99171765 |
| Trpt1 | 33.132397 | -0.1361756 | 0.85440678 | -0.1593803 | 0.87336927 | 0.97344599 |
| Msl3l2 | 65.5805202 | -0.4701104 | 0.57950938 | -0.8112214 | 0.41723856 | 0.8049685 |

| | | | | | | |
|---|---|---|---|---|---|---|
| Rps8 | 22647.4647 | 0.08210021 | 0.38766101 | 0.21178351 | 0.83227593 | 0.96550322 |
| Rpsa-ps10 | 49.0210157 | 0.25792183 | 0.70665676 | 0.36498885 | 0.71511973 | 0.93235217 |
| Yipf6 | 681.578628 | 0.02826931 | 0.31886567 | 0.08865586 | 0.92935542 | 0.9867 |
| Olfr1388 | 22.0465278 | -2.0486148 | 0.88077849 | -2.3259138 | 0.02002315 | 0.20003355 |
| Champ1 | 1387.68998 | -0.381685 | 0.33287324 | -1.1466378 | 0.25153134 | 0.68499173 |
| Ust | 76.1362142 | 1.67180601 | 1.16909132 | 1.43000464 | 0.15271569 | 0.58090075 |
| Ppp1r2 | 2924.16535 | -0.0569561 | 0.17667405 | -0.3223794 | 0.74716526 | 0.9399191 |
| Ubiad1 | 1090.91276 | -0.1552714 | 0.22999538 | -0.6751066 | 0.49960807 | 0.84873011 |
| Bola2 | 1856.06772 | 0.23862782 | 0.43332787 | 0.55068654 | 0.58184857 | 0.88321857 |
| Wbp1l | 1919.84608 | 0.06502579 | 0.30483451 | 0.21331505 | 0.83108124 | 0.96466657 |
| Samd9l | 63.2459047 | -1.2634871 | 0.72903009 | -1.7331069 | 0.08307667 | 0.4429983 |
| Rnf150 | 90.4994358 | 1.31305455 | 0.8828378 | 1.48731121 | 0.13693263 | 0.55967154 |
| Zc3hav1l | 140.113137 | 0.0715057 | 0.36779344 | 0.19441811 | 0.84584851 | 0.96765585 |
| Fancb | 752.269179 | 0.40259331 | 0.32369321 | 1.24374963 | 0.21359172 | 0.64658934 |
| Lrrc49 | 158.967272 | 0.02386477 | 0.38495857 | 0.0619931 | 0.95056833 | 0.99082331 |
| Atg16l2 | 401.322816 | -0.1105809 | 0.31776899 | -0.3479916 | 0.72784649 | 0.93576253 |
| Phf13 | 535.910803 | 0.09069465 | 0.41339048 | 0.21939221 | 0.82634454 | 0.9636485 |
| Lix1 | 82.63983 | 1.40533822 | 0.71484939 | 1.9659221 | 0.04930761 | 0.34139551 |
| Slc38a9 | 792.675472 | 0.05460001 | 0.34361514 | 0.15889872 | 0.87374867 | 0.97350615 |
| Cd300lf | 353.046215 | -1.6631545 | 0.49568966 | -3.3552334 | 0.00079298 | 0.01959703 |
| Akap10 | 1496.14477 | 0.22511043 | 0.22579615 | 0.9969631 | 0.31878242 | 0.74398524 |
| Ccdc88b | 643.622395 | -0.3039539 | 0.61313575 | -0.4957368 | 0.62008015 | 0.89896588 |
| Trim16 | 35.207685 | 1.27631389 | 1.26015969 | 1.01281916 | 0.31114655 | 0.74011268 |
| Pygo2 | 1002.06034 | -0.1016253 | 0.39060413 | -0.2601746 | 0.7947291 | 0.95367917 |
| Cdca4 | 3965.23469 | -0.351639 | 0.18795202 | -1.8708976 | 0.06135928 | 0.38244534 |
| Bri3 | 146.597396 | 0.47574089 | 0.36666479 | 1.29748181 | 0.19446546 | 0.62651174 |
| Bex4 | 139.150359 | 0.26758636 | 0.50194634 | 0.53309754 | 0.59396606 | 0.88797037 |
| Lonp2 | 2941.02877 | 0.08228074 | 0.14735182 | 0.55839648 | 0.57657368 | 0.88105119 |
| Gimap6 | 661.979225 | 1.09091364 | 0.27531154 | 3.96246966 | 7.417843717 | 0.00281809 |
| Gpr157 | 317.391174 | 1.91784763 | 0.39953022 | 4.80025678 | 1.584623168 | 0.00010651 |
| Usp14 | 9796.31058 | -0.2310652 | 0.16360608 | -1.4123267 | 0.1578538 | 0.58658126 |
| Cxcr5 | 68.2394439 | 1.74813864 | 0.76600034 | 2.28216431 | 0.02247964 | 0.21507833 |
| Rell1 | 357.684682 | 0.28227581 | 0.44736922 | 0.63096833 | 0.52806123 | 0.86198732 |
| Tnrc6b | 358.339688 | -0.2708075 | 0.45124638 | -0.6001321 | 0.54841817 | 0.87059427 |
| Ankrd16 | 353.547277 | 0.35815902 | 0.32396069 | 1.1055632 | 0.26891563 | 0.70592381 |
| Trappc9 | 561.357043 | 0.38214297 | 0.40125838 | 0.95236134 | 0.34091376 | 0.75839207 |
| Marcksl1 | 4527.42881 | 0.69594435 | 0.24053878 | 2.89327294 | 0.0038125 | 0.06468556 |
| Kcna3 | 15.3380848 | -0.4698917 | 0.91099894 | -0.5157983 | 0.6059953 | 0.89258167 |
| Stbd1 | 86.2441551 | -0.5724748 | 0.54352313 | -1.0532666 | 0.29221881 | 0.72584701 |
| Rpl9-ps7 | 41.5184636 | 0.82095778 | 0.94044989 | 0.87294155 | 0.38269494 | 0.7856196 |
| Ino80c | 2283.44699 | 0.09359602 | 0.20124845 | 0.46507698 | 0.64187633 | 0.90702012 |
| Prrg1 | 71.7379017 | -0.97064 | 0.74787481 | -1.2978642 | 0.19433399 | 0.62642628 |
| Gigyf2 | 841.875655 | -0.0814236 | 0.33976777 | -0.2396447 | 0.81060568 | 0.95905029 |
| Timm8a1 | 4213.18153 | -0.3756734 | 0.23157268 | -1.6222698 | 0.10474558 | 0.49501875 |
| Zfp473 | 95.1147496 | 0.68604497 | 0.72074071 | 0.95186099 | 0.34116748 | 0.75846782 |
| Rgmb | 396.497306 | 0.83852827 | 0.70269799 | 1.19329823 | 0.2327526 | 0.66805784 |

| | | | | | | |
|---|---|---|---|---|---|---|
| Isg20l2 | 1661.04848 | -0.6187779 | 0.21162183 | -2.9239794 | 0.00345588 | 0.06005075 |
| Arxes2 | 517.761494 | -0.1615283 | 0.4065384 | -0.3973261 | 0.69112699 | 0.92470267 |
| Zbtb33 | 71.9854806 | 0.10926045 | 0.76858757 | 0.14215746 | 0.88695564 | 0.97827644 |
| Ldlrad3 | 307.04657 | -0.9939829 | 0.4798456 | -2.0714641 | 0.03831545 | 0.29553145 |
| Arf1 | 38939.3054 | -0.0561528 | 0.13253379 | -0.423687 | 0.67179409 | 0.91719387 |
| Gm4737 | 15.1527897 | -0.6021708 | 1.01030968 | -0.596026 | 0.55115786 | 0.87093412 |
| Taf13 | 1277.32841 | 0.45674388 | 0.19943095 | 2.29023563 | 0.02200766 | 0.21347378 |
| 4632415L05I | 142.608599 | 0.15995446 | 0.69328354 | 0.23072012 | 0.81753224 | 0.96079676 |
| Tmem72 | 20.8714057 | -0.0764598 | 0.92424917 | -0.0827264 | 0.93406912 | 0.98756396 |
| Rbm15 | 1469.05456 | 0.00598565 | 0.29721577 | 0.02013908 | 0.98393243 | 0.99564261 |
| Arid4a | 959.950805 | -0.0623247 | 0.41630196 | -0.1497103 | 0.88099321 | 0.97582756 |
| Entpd1 | 4628.37968 | 0.84434698 | 0.15103301 | 5.59047983 | 2.264429883 | 2.663535650 |
| Dmrt2 | 23.4040623 | 4.28121578 | 1.56207515 | 2.74072331 | 0.00613041 | 0.09327527 |
| Nat8l | 47.9624998 | 0.34020094 | 0.93581245 | 0.36353538 | 0.71620499 | 0.93235217 |
| Kmt2d | 1502.53936 | -0.088062 | 0.42955585 | -0.2050072 | 0.83756654 | 0.96683245 |
| Selplg | 2319.08866 | -1.594319 | 0.23158769 | -6.8842994 | 5.807261104 | 1.480004689 |
| Mcmbp | 2970.41422 | -0.2613171 | 0.24038766 | -1.0870653 | 0.27700795 | 0.71247429 |
| Tmem81 | 30.0215514 | -0.4993813 | 1.06891094 | -0.467187 | 0.64036609 | 0.90677142 |
| Asb8 | 1857.34516 | 0.33596778 | 0.21286039 | 1.57834806 | 0.11448567 | 0.51679085 |
| Cracr2b | 167.72034 | 1.48909022 | 0.57305804 | 2.59849809 | 0.00936326 | 0.12401786 |
| Gpr85 | 30.469646 | 0.90157623 | 0.89818206 | 1.00377892 | 0.31548519 | 0.74186126 |
| Amigo2 | 16.0996673 | 0.00551772 | 2.24162262 | 0.00246148 | 0.99803602 | 0.99967651 |
| Mfap1b | 2798.59181 | 0.15834235 | 0.17015767 | 0.93056255 | 0.35207989 | 0.76348958 |
| Fbxo10 | 64.2728957 | 0.85000347 | 1.31239232 | 0.64767483 | 0.51719529 | 0.85694744 |
| Rnf149 | 823.150095 | 0.52101388 | 0.33564921 | 1.5522571 | 0.12060072 | 0.52878444 |
| Gng7 | 16.6295811 | -2.8082465 | 2.34991924 | -1.1950396 | 0.23207157 | 0.66733333 |
| Crebrf | 142.173189 | 1.05065541 | 0.58490486 | 1.79628429 | 0.07244931 | 0.41395258 |
| Dip2c | 53.593418 | 1.12688902 | 0.82831674 | 1.36045666 | 0.17368546 | 0.60653633 |
| Rbm33 | 434.80601 | -0.3754386 | 0.30820835 | -1.2181325 | 0.22317362 | 0.6565856 |
| Syngr2 | 7827.99519 | -0.4736891 | 0.20858747 | -2.2709376 | 0.02315075 | 0.21835248 |
| Sacs | 97.572628 | -0.545865 | 1.27236148 | -0.4290172 | 0.6679107 | 0.91595755 |
| Zfp738 | 147.090354 | 1.04139419 | 0.53818729 | 1.93500331 | 0.05298989 | 0.35597217 |
| Frmd6 | 95.3062443 | 0.26938523 | 0.50696856 | 0.53136477 | 0.59516603 | 0.88830085 |
| Ankrd46 | 1465.96299 | 0.38180347 | 0.22026267 | 1.73340072 | 0.08302447 | 0.4429983 |
| Pskh1 | 982.487759 | 0.11740876 | 0.31964239 | 0.36731285 | 0.71338567 | 0.93235217 |
| Ckap2l | 4121.81335 | -0.1124016 | 0.18393746 | -0.6110859 | 0.5411427 | 0.86776854 |
| Ric3 | 36.1287143 | -0.2434282 | 0.8232387 | -0.2956957 | 0.76746247 | 0.94401697 |
| Lhfp | 58.277701 | 2.78269371 | 1.63342275 | 1.70359676 | 0.08845645 | 0.45427696 |
| Coa7 | 1352.23939 | 0.13632997 | 0.22498642 | 0.60594757 | 0.54454957 | 0.86960875 |
| Arxes1 | 25.4825482 | 0.90561207 | 1.33614424 | 0.67778017 | 0.49791109 | 0.84860011 |
| Omd | 10.9696308 | 1.14495469 | 1.34991512 | 0.84816791 | 0.39634447 | 0.79439542 |
| Pdp2 | 1607.17008 | -0.2502159 | 0.28230647 | -0.8863272 | 0.37544122 | 0.77815049 |
| F2r | 223.527416 | 1.357456 | 0.70713421 | 1.91965822 | 0.05490108 | 0.36084578 |
| Socs4 | 695.986425 | -0.1458997 | 0.24994501 | -0.5837273 | 0.55940381 | 0.87397022 |
| Zfp407 | 179.250231 | -0.5614219 | 0.45679334 | -1.22905 | 0.21905306 | 0.65257977 |
| Mlf1 | 113.468488 | 0.2715092 | 0.58313602 | 0.46560184 | 0.64150053 | 0.90700632 |

| | | | | | | |
|---|---|---|---|---|---|---|
| Timm29 | 2606.12522 | -0.2267581 | 0.1769126 | -1.2817519 | 0.1999297 | 0.63247751 |
| Nupl2 | 778.970444 | -0.3457979 | 0.25942323 | -1.332949 | 0.18254854 | 0.61383074 |
| Cyp4f16 | 59.717658 | 1.0340999 | 0.98887588 | 1.04573276 | 0.29568443 | 0.72852118 |
| Ccdc57 | 77.3865662 | 0.25695476 | 0.71393799 | 0.35991188 | 0.71891303 | 0.93310883 |
| Spata33 | 7.45096216 | -0.080675 | 2.25368378 | -0.0357969 | 0.97144427 | 0.99443087 |
| Mypop | 45.608877 | -0.0471331 | 0.58486477 | -0.080588 | 0.93576962 | 0.98794094 |
| Fitm2 | 248.162878 | -0.9661324 | 0.67883078 | -1.4232301 | 0.15466945 | 0.58343244 |
| Depp1 | 29.6710615 | 0.05178205 | 1.40044071 | 0.03697554 | 0.97050451 | 0.99443087 |
| Nrip1 | 414.155946 | 0.30139878 | 0.36074369 | 0.83549289 | 0.40344025 | 0.79841492 |
| Tyw5 | 737.12988 | -0.2646601 | 0.38113275 | -0.6944039 | 0.48742894 | 0.84278703 |
| Mmgt2 | 806.402221 | -0.1217493 | 0.26600062 | -0.4577031 | 0.64716575 | 0.90950305 |
| Cd300e | 51.4631026 | -0.4504711 | 0.7777137 | -0.5792249 | 0.56243745 | 0.8755831 |
| Phldb1 | 21.1877297 | 1.43476165 | 1.52365166 | 0.94165988 | 0.3463668 | 0.7602916 |
| Tob2 | 799.831342 | -0.4437647 | 0.43671469 | -1.0161433 | 0.30956114 | 0.73898722 |
| Thnsl1 | 369.477592 | 0.40077368 | 0.54518149 | 0.73511975 | 0.46226657 | 0.83108016 |
| Mlec | 6684.37741 | 0.21550974 | 0.37010664 | 0.58229094 | 0.56037073 | 0.87469256 |
| E130311K13l | 127.322322 | 1.1846653 | 0.5815253 | 2.03716899 | 0.04163311 | 0.31092664 |
| Igf2 | 468.656519 | 1.79514679 | 0.79992007 | 2.2441577 | 0.02482226 | 0.22882492 |
| Myof | 18.5986979 | 0.71430499 | 1.00258528 | 0.71246307 | 0.47617807 | 0.83644936 |
| Gm6377 | 775.123016 | 0.9041671 | 0.4566203 | 1.980129 | 0.04768903 | 0.33701904 |
| Ctxn1 | 24.103555 | 1.51567716 | 1.38844551 | 1.09163604 | 0.27499311 | 0.71147204 |
| Exd1 | 39.7666346 | -1.7918239 | 0.81246579 | -2.2054147 | 0.02742501 | 0.24574023 |
| Lemd3 | 83.4852307 | -1.0211549 | 0.947702 | -1.0775063 | 0.28125414 | 0.71634017 |
| Rhno1 | 1332.94786 | -0.3658684 | 0.20493955 | -1.7852501 | 0.07422076 | 0.41976075 |
| Tpcn2 | 998.392963 | 0.68941153 | 0.21251231 | 3.24410166 | 0.00117822 | 0.02652579 |
| Mex3d | 15.426509 | 0.94834503 | 1.19945313 | 0.79064784 | 0.42914952 | 0.81140481 |
| Ccdc6 | 1590.85933 | 0.0143352 | 0.31386837 | 0.04567264 | 0.96357117 | 0.99401755 |
| Lurap1l | 12.3740086 | 2.40224337 | 1.7747902 | 1.35353653 | 0.17588429 | 0.60763832 |
| Tprn | 310.38723 | -0.138343 | 0.40467929 | -0.3418585 | 0.73245741 | 0.93725434 |
| Ggnbp1 | 12.8345067 | 0.27051785 | 1.05155963 | 0.25725393 | 0.79698275 | 0.95397994 |
| Klhl11 | 95.9637109 | -0.0665149 | 0.54619228 | -0.1217792 | 0.90307389 | 0.98043702 |
| Prss50 | 8.81144059 | 6.6218444 | 2.0414052 | 3.24376777 | 0.0011796 | 0.02652579 |
| Mcat | 192.468065 | 0.36098356 | 0.51280455 | 0.70393986 | 0.48147023 | 0.84011103 |
| Foxo3 | 572.331947 | 0.19918092 | 0.35050747 | 0.56826442 | 0.56985543 | 0.87984873 |
| Rpl29 | 11895.418 | -0.2468805 | 0.19717927 | -1.2520612 | 0.21054757 | 0.64401792 |
| Tmem53 | 96.4058158 | 1.34077976 | 0.53015079 | 2.52905361 | 0.01143706 | 0.14023051 |
| P2ry6 | 6384.03102 | 0.3982864 | 0.22454961 | 1.77371226 | 0.07611076 | 0.42514288 |
| Dcun1d3 | 82.4793053 | -0.3602113 | 0.61912258 | -0.5818093 | 0.56069512 | 0.87469256 |
| Cyb561d1 | 190.122511 | -0.1188077 | 0.38948281 | -0.3050396 | 0.76033601 | 0.94215089 |
| Cep120 | 957.067851 | 0.25963898 | 0.29306224 | 0.88595169 | 0.37564356 | 0.77819605 |
| Ifnb1 | 25.3995324 | 0.59130046 | 1.18697872 | 0.49815591 | 0.61837416 | 0.89878056 |
| Slc35e4 | 446.835299 | 0.19116911 | 0.40953442 | 0.46679621 | 0.64064569 | 0.90685007 |
| Pkd1l3 | 37.899671 | 0.06862734 | 0.78955162 | 0.08691938 | 0.93073559 | 0.98716777 |
| Vps37c | 1306.53247 | -0.0180373 | 0.3457894 | -0.0521626 | 0.95839916 | 0.99249408 |
| Slc39a9 | 1125.74173 | -0.1727913 | 0.28791412 | -0.6001488 | 0.54840704 | 0.87059427 |
| Slc25a47 | 28.1301551 | -2.2771687 | 1.21165489 | -1.8793872 | 0.06019164 | 0.37837837 |

| | | | | | | |
|---|---|---|---|---|---|---|
| Arhgap30 | 3581.08865 | -0.2516788 | 0.24581233 | -1.0238655 | 0.30589882 | 0.73554152 |
| Phf3 | 420.090733 | -0.3215058 | 0.3126911 | -1.0281896 | 0.30386063 | 0.73436097 |
| Hexim1 | 180.282728 | -0.4352703 | 0.57689725 | -0.7545023 | 0.45054768 | 0.82520583 |
| Zfp710 | 1495.8773 | -0.2305777 | 0.31906003 | -0.7226781 | 0.46987769 | 0.83381643 |
| 4930539E08I | 12.6960385 | 0.64635976 | 2.60925647 | 0.24771799 | 0.80435261 | 0.9570372 |
| Rnf24 | 105.108338 | -0.4606812 | 0.4805592 | -0.9586356 | 0.33774234 | 0.75611716 |
| Efna5 | 16.2786237 | -0.6007538 | 1.47690291 | -0.406766 | 0.68417988 | 0.92277536 |
| Fkrp | 974.178489 | -0.0152633 | 0.22526403 | -0.0677575 | 0.94597868 | 0.99019648 |
| Zfp689 | 148.165541 | -0.9624196 | 0.60447098 | -1.5921684 | 0.11134687 | 0.50888686 |
| Cdca2 | 2584.0501 | -0.0524691 | 0.20449647 | -0.2565768 | 0.79750545 | 0.95421403 |
| Ccdc125 | 108.306851 | -0.2629943 | 0.52793744 | -0.4981542 | 0.61837536 | 0.89878056 |
| Tada3 | 1172.65942 | -0.2410121 | 0.1763635 | -1.3665647 | 0.1717618 | 0.60430173 |
| Prex2 | 19.1643747 | 1.22130892 | 1.99209434 | 0.61307785 | 0.53982486 | 0.86719392 |
| Yjefn3 | 11.8356697 | 0.51563616 | 1.32221788 | 0.38997821 | 0.69655266 | 0.92646244 |
| C1galt1c1 | 2819.49846 | -1.7899096 | 0.30625716 | -5.8444661 | 5.081954581 | 8.073707844 |
| Atxn7l2 | 109.798106 | -0.3612749 | 0.55045356 | -0.6563223 | 0.51161678 | 0.85393154 |
| Clec4a1 | 2387.9672 | 0.67460016 | 0.32879964 | 2.05170593 | 0.04019825 | 0.30448619 |
| Mterf2 | 38.934104 | 0.34061565 | 0.9406387 | 0.36211103 | 0.71726906 | 0.93264801 |
| Armcx3 | 1736.56065 | 0.85944312 | 0.23422333 | 3.6693318 | 0.00024319 | 0.00756659 |
| Acap2 | 1491.59688 | -0.1192362 | 0.23748251 | -0.5020843 | 0.61560821 | 0.89876301 |
| Bmyc | 803.369281 | -0.207421 | 0.24682282 | -0.8403639 | 0.40070437 | 0.79698916 |
| Zadh2 | 396.831073 | 0.39991997 | 0.29620926 | 1.35012652 | 0.1769754 | 0.60784615 |
| Sephs2 | 4526.49889 | -0.2736629 | 0.19487099 | -1.4043286 | 0.16022103 | 0.5898236 |
| Ccr2 | 644.325359 | -4.7553599 | 0.40671258 | -11.692188 | 1.397393138 | 2.442044323 |
| Dcaf5 | 272.82211 | 0.25167129 | 0.34988973 | 0.71928745 | 0.47196382 | 0.83435479 |
| Fam110b | 89.0577268 | 0.37737339 | 0.63314703 | 0.59602805 | 0.55115649 | 0.87093412 |
| Gm8186 | 198.786374 | 0.74125784 | 0.62872248 | 1.17899052 | 0.23840195 | 0.67446139 |
| Ivl | 42.2151178 | -0.5574214 | 0.94921521 | -0.5872445 | 0.55703952 | 0.87276537 |
| C5ar1 | 9763.7548 | 1.38481831 | 0.29137318 | 4.75273087 | 2.006874050 | 0.00013332 |
| Nrap | 105.281397 | 2.12443012 | 0.57561619 | 3.69070596 | 0.00022363 | 0.00710571 |
| Zfp518a | 274.303676 | -0.3526264 | 0.39297845 | -0.8973174 | 0.36954956 | 0.77502139 |
| Purg | 8.73229899 | -0.4354449 | 2.23234703 | -0.1950615 | 0.84534484 | 0.96765585 |
| Pdp1 | 308.312381 | 1.07628378 | 0.4469077 | 2.40829095 | 0.0160274 | 0.17615771 |
| Tigd2 | 334.793368 | 0.41398101 | 0.30274142 | 1.3674409 | 0.17148715 | 0.60430173 |
| Gm7324 | 22.2966706 | -0.2037869 | 0.90907139 | -0.2241704 | 0.82262472 | 0.96252207 |
| Mblac1 | 149.3785 | 0.39676696 | 0.60708535 | 0.65356042 | 0.51339505 | 0.85497712 |
| Iba57 | 258.066103 | 0.20884203 | 0.47530373 | 0.43938647 | 0.66038153 | 0.91302324 |
| Lix1l | 38.3674904 | 0.04860094 | 1.19934811 | 0.0405228 | 0.96767633 | 0.99443087 |
| Zfp219 | 181.878934 | -0.1954494 | 0.43570846 | -0.4485784 | 0.65373581 | 0.91076467 |
| Trappc1 | 3143.02973 | 0.00120018 | 0.17422504 | 0.0068887 | 0.99450366 | 0.99882461 |
| Prmt6 | 745.605879 | -0.4543697 | 0.36364563 | -1.2494849 | 0.21148779 | 0.64533053 |
| Ccdc71 | 1177.61088 | 0.024057 | 0.27164863 | 0.08855924 | 0.9294322 | 0.9867 |
| Fut4 | 220.188365 | -0.0602961 | 0.53153554 | -0.1134376 | 0.90968361 | 0.98261895 |
| Sorl1 | 637.849042 | -3.160721 | 0.55691197 | -5.675441 | 1.383318530 | 1.819584471 |
| Zfp2 | 186.740105 | 1.99250569 | 0.81180519 | 2.45441358 | 0.01411146 | 0.16167905 |
| Smcr8 | 830.819727 | -0.0263032 | 0.34904565 | -0.0753575 | 0.93993029 | 0.98934006 |

| | | | | | | |
|---|---|---|---|---|---|---|
| Kmt5a | 728.183054 | -0.6079344 | 0.24290157 | -2.5028013 | 0.01232147 | 0.1477731 |
| Retreg2 | 481.079982 | 0.37559031 | 0.38456011 | 0.97667518 | 0.32872999 | 0.74843736 |
| Rd3 | 60.3741476 | 2.73982403 | 0.58623007 | 4.67363275 | 2.959182759 | 0.00018564 |
| Dcaf7 | 3603.84472 | -0.0621749 | 0.23607421 | -0.26337 | 0.79226541 | 0.95297175 |
| Krt8 | 80.3462151 | -1.1002461 | 0.75335036 | -1.4604707 | 0.14416076 | 0.57027792 |
| Gemin4 | 3259.00654 | -0.1979579 | 0.26814517 | -0.7382491 | 0.46036308 | 0.82987212 |
| Ogfr | 2339.22051 | -0.219409 | 0.19761099 | -1.1103075 | 0.26686653 | 0.7035729 |
| Rarres1 | 30.0931804 | 0.51082104 | 0.78322616 | 0.6522012 | 0.51427139 | 0.85523136 |
| Prokr1 | 31.1899756 | -0.1851875 | 0.78266612 | -0.2366111 | 0.81295852 | 0.95974923 |
| Tmem241 | 606.868814 | 1.005326 | 0.25397481 | 3.958369 | 7.546331025 | 0.00283173 |
| Zfp260 | 982.596591 | 0.5816813 | 0.33583676 | 1.73203579 | 0.08326719 | 0.44341423 |
| Chchd10 | 142.934913 | -0.0122034 | 0.38128302 | -0.032006 | 0.97446725 | 0.99446549 |
| Cyp20a1 | 2179.02266 | 0.07158098 | 0.189558 | 0.37762048 | 0.70571255 | 0.92896305 |
| Aff4 | 599.749713 | 1.88159942 | 0.44734377 | 4.2061599 | 2.597464933 | 0.00113481 |
| Ctu2 | 1810.24908 | -0.2770201 | 0.17440891 | -1.5883369 | 0.11221017 | 0.51028512 |
| Tmem67 | 196.684347 | -0.3101848 | 0.59829661 | -0.5184465 | 0.60414678 | 0.89257579 |
| Ccnq | 1045.35533 | 0.02102813 | 0.35404041 | 0.05939471 | 0.95263772 | 0.9914597 |
| Dtx3l | 700.311619 | -0.3702193 | 0.21155149 | -1.7500198 | 0.0801149 | 0.43723638 |
| Proser1 | 452.947122 | 0.29225243 | 0.51810968 | 0.56407443 | 0.57270348 | 0.88035336 |
| Spty2d1 | 972.932858 | -0.0471577 | 0.28765474 | -0.1639385 | 0.86977953 | 0.97239152 |
| Rps23 | 10745.851 | 0.03952369 | 0.35443559 | 0.11151164 | 0.91121064 | 0.98289594 |
| Cdc42ep1 | 45.0880824 | 1.66791296 | 1.38582405 | 1.2035532 | 0.22876232 | 0.66258779 |
| Tmem202 | 20.014688 | 0.9252134 | 1.49710495 | 0.6180017 | 0.53657422 | 0.86583728 |
| Sall2 | 18.5104676 | 0.3877497 | 1.26754546 | 0.30590595 | 0.75967625 | 0.94210735 |
| Tceal1 | 72.8067066 | 1.07899044 | 0.62119533 | 1.73695839 | 0.08239452 | 0.44183004 |
| Hist1h1a | 14.4846874 | -0.145371 | 1.11841284 | -0.1299797 | 0.89658246 | 0.97984623 |
| Clip1 | 1331.40664 | 0.06851136 | 0.33527486 | 0.20434387 | 0.83808479 | 0.96683245 |
| Polr1a | 1312.76866 | -0.3582192 | 0.27610342 | -1.2974095 | 0.19449031 | 0.62651174 |
| Ap5b1 | 444.469993 | -0.5498129 | 0.37850605 | -1.4525868 | 0.14633852 | 0.57340745 |
| Zfpm1 | 8.48500944 | 0.3299387 | 2.22886804 | 0.14802971 | 0.88231932 | 0.97598447 |
| Tsku | 26.2034044 | 2.63755247 | 1.21784616 | 2.16575177 | 0.03033016 | 0.26001357 |
| Ccdc69 | 186.023342 | -0.3051928 | 0.38835697 | -0.7858564 | 0.43195165 | 0.81344118 |
| Zbtb45 | 303.225097 | -0.2064015 | 0.27964973 | -0.7380716 | 0.46047095 | 0.82987212 |
| Zfp644 | 332.844089 | -0.3269018 | 0.27642719 | -1.1825964 | 0.23696914 | 0.6732629 |
| Gpr55 | 12.1375827 | -1.4908598 | 1.9441455 | -0.7668458 | 0.44317321 | 0.82038676 |
| Slc17a5 | 1084.0481 | 0.65707555 | 0.30136671 | 2.18031895 | 0.02923383 | 0.2548948 |
| Tifab | 5314.54359 | -0.795849 | 0.21777798 | -3.6544054 | 0.00025778 | 0.00790327 |
| 2310022A10I | 1054.10305 | -0.3543635 | 0.24420336 | -1.4510999 | 0.14675205 | 0.57365079 |
| Zbtb5 | 370.044734 | -0.1682107 | 0.39350566 | -0.4274671 | 0.66903911 | 0.91639855 |
| Bdp1 | 528.471078 | -0.2573501 | 0.50768044 | -0.5069136 | 0.61221543 | 0.89650055 |
| Aftph | 1320.74903 | -0.0943492 | 0.28991206 | -0.3254409 | 0.74484744 | 0.93917909 |
| Morn4 | 112.203015 | -0.3523644 | 0.56383882 | -0.6249381 | 0.53201166 | 0.8643494 |
| Zbtb14 | 655.61855 | 0.00563503 | 0.33967338 | 0.01658955 | 0.98676406 | 0.99612846 |
| Catsperg1 | 10.068184 | -0.4428763 | 2.67202716 | -0.1657454 | 0.86835729 | 0.97186901 |
| Urgcp | 819.419418 | 0.1952479 | 0.26833702 | 0.72762195 | 0.46684504 | 0.8334534 |
| Orai1 | 298.02088 | 0.19431248 | 0.39248052 | 0.49508822 | 0.62053785 | 0.89896588 |

| | | | | | | |
|---|---|---|---|---|---|---|
| Pheta2 | 379.272927 | -0.5577406 | 0.28884233 | -1.9309518 | 0.05348901 | 0.35852954 |
| Nlrp10 | 10.1734124 | -1.6188469 | 2.60122476 | -0.6223403 | 0.53371816 | 0.86500711 |
| Lig4 | 85.8564007 | 0.32519839 | 1.25717746 | 0.25867342 | 0.79588723 | 0.9537323 |
| Gal3st1 | 23.1458277 | 0.86940927 | 1.32651218 | 0.65540994 | 0.51220385 | 0.85399887 |
| Mmp12 | 50.9654922 | 1.24644229 | 1.00950105 | 1.23471123 | 0.21693799 | 0.65065857 |
| Zfp668 | 416.838605 | -0.2156603 | 0.37172855 | -0.5801553 | 0.56180993 | 0.87540592 |
| Trex1 | 1382.9637 | -0.2252061 | 0.25262692 | -0.8914571 | 0.37268401 | 0.77748705 |
| Zfp646 | 233.012138 | 0.34888903 | 0.45163561 | 0.77250115 | 0.43981767 | 0.81963526 |
| Arhgap15 | 1737.58307 | -0.9231636 | 0.3401825 | -2.7137303 | 0.00665303 | 0.09889005 |
| Rpl36al | 10850.7199 | 0.31073831 | 0.37545946 | 0.82762146 | 0.40788491 | 0.80064439 |
| Zfp672 | 569.740243 | 0.15971494 | 0.3460452 | 0.46154358 | 0.64440866 | 0.90829026 |
| Micos13 | 2706.10238 | 0.42832204 | 0.43042296 | 0.99511894 | 0.31967842 | 0.7440831 |
| Zfp280b | 350.249296 | 0.09116263 | 0.5624005 | 0.16209557 | 0.87123059 | 0.97268995 |
| Tmsb4x | 278777.089 | 0.65688618 | 0.38079242 | 1.72505058 | 0.08451836 | 0.44526833 |
| Fzd4 | 72.1749911 | -0.7091391 | 0.62819925 | -1.1288442 | 0.25896356 | 0.69546977 |
| Bag5 | 1441.80586 | -0.154197 | 0.19989336 | -0.7713964 | 0.44047201 | 0.81963526 |
| Sertad2 | 576.739107 | 0.0620876 | 0.29887322 | 0.20773892 | 0.83543283 | 0.96641606 |
| Armcx4 | 9.04857998 | -1.626444 | 2.2109776 | -0.7356221 | 0.46196071 | 0.83080938 |
| Arhgap23 | 59.2871752 | 0.8230777 | 1.03387215 | 0.79611169 | 0.42596712 | 0.81021713 |
| Fam161a | 9.20443275 | -0.2458141 | 1.73311166 | -0.141834 | 0.88721114 | 0.97827644 |
| Suox | 62.7085806 | -0.0465694 | 0.52946439 | -0.0879557 | 0.92991191 | 0.98699718 |
| Arl4c | 205.029295 | -0.2771604 | 0.45791967 | -0.6052598 | 0.54500639 | 0.86963259 |
| Nlrc3 | 210.89583 | 0.3086863 | 0.41075136 | 0.7515162 | 0.45234205 | 0.82603347 |
| Rlf | 802.045146 | 0.50942125 | 0.21412698 | 2.37906153 | 0.01735678 | 0.18521003 |
| 2810025M15 | 2599.09756 | 0.3941766 | 0.26600209 | 1.48185526 | 0.13837881 | 0.56089729 |
| Vcpkmt | 836.149493 | 0.17006323 | 0.34482766 | 0.49318325 | 0.62188311 | 0.89912494 |
| Rasd1 | 10.0021352 | -0.0530661 | 2.14286546 | -0.0247641 | 0.98024315 | 0.99504725 |
| Tmem17 | 277.927062 | 0.41313602 | 0.52436139 | 0.78788415 | 0.43076448 | 0.81254302 |
| Rasl11b | 124.954068 | 0.92589641 | 0.77388343 | 1.19642877 | 0.23152928 | 0.66689373 |
| 2610318N02 | 921.636053 | -0.0558149 | 0.20897438 | -0.2670896 | 0.78940021 | 0.95255415 |
| Slc35c1 | 943.214549 | -0.9164832 | 0.40092466 | -2.2859237 | 0.02225872 | 0.21435947 |
| Lpar4 | 126.211617 | 1.77663879 | 0.66042733 | 2.69013519 | 0.00714231 | 0.10299472 |
| H2afx | 4662.31757 | -0.3321288 | 0.15265302 | -2.1757103 | 0.02957693 | 0.25669202 |
| Lrrc4 | 21.8950169 | 0.97887471 | 1.12155272 | 0.87278528 | 0.38278012 | 0.78566263 |
| Pgrmc2 | 89.5607599 | -0.7016973 | 0.46030021 | -1.5244341 | 0.12740031 | 0.54145382 |
| Rpp38 | 539.678772 | 0.28505169 | 0.35710614 | 0.79822679 | 0.42473889 | 0.80982401 |
| Ccdc137 | 885.039378 | -0.5801499 | 0.26061983 | -2.226039 | 0.02601157 | 0.23622832 |
| Mrps16 | 2316.50331 | 0.44568461 | 0.4301641 | 1.03608042 | 0.30016462 | 0.73145694 |
| Plekhf2 | 2458.08082 | 0.00450531 | 0.2769984 | 0.01626474 | 0.98702319 | 0.99616298 |
| Ankrd55 | 20.1655529 | 0.38918106 | 0.89645756 | 0.43413216 | 0.66419247 | 0.9141614 |
| Lrrc25 | 2666.32713 | 0.39484533 | 0.24153032 | 1.63476508 | 0.1020983 | 0.48845074 |
| Ppp1r3d | 70.7451338 | -0.3623571 | 0.61116478 | -0.592896 | 0.55325079 | 0.87136821 |
| Idnk | 677.242604 | 0.28678031 | 0.43021245 | 0.66660162 | 0.50502663 | 0.85034006 |
| Shisa3 | 45.7810938 | 4.61186139 | 3.58085192 | 1.28792295 | 0.19777278 | 0.6306972 |
| Pitpnb | 5889.73191 | -0.0234038 | 0.15702952 | -0.1490406 | 0.88152157 | 0.97582756 |
| Amz1 | 405.687514 | -1.2675075 | 0.28156127 | -4.5017112 | 6.740854656 | 0.00037313 |

| | | | | | | |
|---|---|---|---|---|---|---|
| Rap2c | 2301.45451 | 0.20391823 | 0.19164143 | 1.0640613 | 0.28730093 | 0.72278333 |
| Fhl4 | 14.1841618 | -0.081119 | 2.56938142 | -0.0315714 | 0.97481383 | 0.9944856 |
| Tmx2 | 5726.16295 | 0.17686637 | 0.18646627 | 0.94851668 | 0.34286649 | 0.75942164 |
| Bex1 | 173.834275 | 0.29388557 | 0.43804588 | 0.67090134 | 0.50228338 | 0.85012672 |
| Gpr171 | 268.895335 | -1.8315158 | 0.40061586 | -4.5717506 | 4.836664198 | 0.00028276 |
| Rspry1 | 2005.55259 | -0.0800052 | 0.23095112 | -0.3464162 | 0.72902993 | 0.93617556 |
| 1600012H06 | 1493.13807 | 0.47212779 | 0.28782324 | 1.64033939 | 0.10093462 | 0.48566935 |
| Agmo | 95.7166932 | 1.2741072 | 0.98105531 | 1.29871088 | 0.19404317 | 0.62565369 |
| Tmc8 | 300.094035 | 0.49234372 | 0.52882159 | 0.93102048 | 0.35184297 | 0.76348958 |
| Haspin | 957.054686 | -0.2658352 | 0.26724856 | -0.9947116 | 0.31987655 | 0.74413737 |
| Sarm1 | 29.7526403 | 0.92670487 | 1.39794854 | 0.66290342 | 0.50739241 | 0.85189835 |
| Kcnk12 | 72.5291581 | -2.1148415 | 0.79598881 | -2.6568734 | 0.00788691 | 0.10963695 |
| Slc25a44 | 1129.28483 | -0.3458348 | 0.42798127 | -0.8080607 | 0.41905567 | 0.80529433 |
| F2rl3 | 35.6028129 | -0.287813 | 0.66685511 | -0.4315975 | 0.66603395 | 0.91520501 |
| Ubqln2 | 183.508343 | -0.1358027 | 0.40067092 | -0.3389381 | 0.73465634 | 0.93750666 |
| Nudt6 | 274.558582 | 0.00849753 | 0.3812932 | 0.02228609 | 0.98221975 | 0.99523682 |
| Lsm10 | 1419.78932 | 0.53637636 | 0.29967186 | 1.78987899 | 0.07347337 | 0.41734043 |
| Eif5a2 | 202.106977 | 0.15523454 | 0.3750037 | 0.41395469 | 0.67890729 | 0.92012773 |
| Lgr4 | 268.123922 | 0.15467386 | 0.39455486 | 0.39202118 | 0.69504256 | 0.92609254 |
| Eva1b | 191.853141 | 0.3869901 | 0.35899983 | 1.07796737 | 0.28104832 | 0.71611415 |
| Snip1 | 1467.91207 | -0.062514 | 0.20704475 | -0.3019347 | 0.76270187 | 0.94298237 |
| Pigm | 990.752923 | 0.03236495 | 0.22266645 | 0.14535174 | 0.88443317 | 0.9772623 |
| Cxcr3 | 189.062613 | 1.46226959 | 0.57140541 | 2.55907552 | 0.01049509 | 0.1323572 |
| Hic2 | 296.553488 | -0.1912531 | 0.42276289 | -0.4523886 | 0.65098908 | 0.91021183 |
| Heatr1 | 1758.8659 | -0.8246022 | 0.27057023 | -3.0476457 | 0.00230642 | 0.04429261 |
| Evc2 | 64.5309267 | 0.49758773 | 0.69021239 | 0.72091974 | 0.4709589 | 0.83381643 |
| Tmem220 | 50.5795192 | 0.59138174 | 0.63245058 | 0.93506397 | 0.34975533 | 0.76212273 |
| Prag1 | 523.244476 | 0.98743775 | 0.25560722 | 3.8631059 | 0.00011195 | 0.00396968 |
| Slc25a2 | 6.54716005 | -1.2862396 | 2.80645507 | -0.4583147 | 0.64672638 | 0.90925224 |
| Rictor | 161.253637 | 0.18318182 | 0.46645105 | 0.39271392 | 0.69453079 | 0.92609254 |
| Nsun3 | 544.477926 | 0.03844225 | 0.35950253 | 0.10693178 | 0.9148431 | 0.98347559 |
| Ndufaf6 | 1057.11737 | 0.0740139 | 0.45641429 | 0.16216385 | 0.87117683 | 0.97268995 |
| Amer1 | 239.51425 | 0.48162632 | 0.48035058 | 1.00265585 | 0.31602694 | 0.74186126 |
| Lgals3 | 1570.50693 | -0.4715872 | 0.23769137 | -1.9840315 | 0.04725233 | 0.33529328 |
| 4930486L24F | 11.3776141 | 6.99137388 | 1.88571443 | 3.70754647 | 0.00020928 | 0.00677271 |
| Ch25h | 27.6274333 | 0.7067467 | 1.1984192 | 0.58973246 | 0.55537003 | 0.8722354 |
| Snx21 | 163.934485 | 0.44262523 | 0.46986816 | 0.94202004 | 0.34618238 | 0.7602916 |
| Sept6 | 1220.17331 | -1.0592575 | 0.28578738 | -3.706453 | 0.00021018 | 0.00678406 |
| Kif7 | 56.6117161 | 0.49395376 | 0.61539682 | 0.80265894 | 0.42217189 | 0.80821275 |
| C77080 | 23.9784822 | -1.7935174 | 0.9811005 | -1.8280669 | 0.06753951 | 0.40045479 |
| Armcx6 | 37.6938542 | 1.37122966 | 1.00304987 | 1.3670603 | 0.17160641 | 0.60430173 |
| Tcf19 | 1496.3501 | 0.08965905 | 0.17901039 | 0.50085949 | 0.61647002 | 0.89878056 |
| Fbxo46 | 401.964474 | 0.10316103 | 0.37214765 | 0.27720457 | 0.78162303 | 0.95004728 |
| Lypd6 | 13.0660244 | -2.8450966 | 2.4467296 | -1.1628161 | 0.24490411 | 0.68039867 |
| Fam118b | 1708.42693 | 0.06140782 | 0.1824252 | 0.33661918 | 0.736404 | 0.93788965 |
| Fbxl22 | 13.0100579 | 0.39930302 | 2.01230667 | 0.1984305 | 0.84270826 | 0.96765585 |

| | | | | | | |
|---|---|---|---|---|---|---|
| Fam171a1 | 101.405664 | 0.58162643 | 0.50904097 | 1.14259257 | 0.25320779 | 0.68647781 |
| B230217C12 | 50.4886563 | -1.4375967 | 0.82292064 | -1.7469445 | 0.08064696 | 0.43854735 |
| Fam228b | 9.74015033 | -1.7041832 | 1.74327331 | -0.9775766 | 0.32828378 | 0.74843736 |
| Fam241a | 19.082258 | -0.9836623 | 1.59943823 | -0.6150049 | 0.53855149 | 0.86663962 |
| Lamtor4 | 858.581168 | -0.688463 | 0.31641388 | -2.1758305 | 0.02956793 | 0.25669202 |
| Hyls1 | 286.002046 | -0.151944 | 0.3242077 | -0.4686624 | 0.63931094 | 0.90632643 |
| Kcnb1 | 24.5425505 | 0.77129664 | 1.35675903 | 0.56848462 | 0.56970595 | 0.87972897 |
| Tor1aip2 | 3251.71096 | 0.08964237 | 0.23664201 | 0.37881005 | 0.70482893 | 0.92874286 |
| Maml1 | 154.12468 | 0.49250289 | 0.39063878 | 1.26076292 | 0.20739428 | 0.6418276 |
| Mmp13 | 52.5569648 | 3.76536145 | 1.4865348 | 2.53297901 | 0.01130977 | 0.13890808 |
| Fam78a | 710.071323 | -0.1171325 | 0.32141238 | -0.3644306 | 0.71553649 | 0.93235217 |
| Zfp61 | 132.763491 | -0.0807862 | 0.43165073 | -0.1871564 | 0.85153804 | 0.9679302 |
| Micos10 | 4764.98431 | 0.38637663 | 0.37263521 | 1.03687633 | 0.2997935 | 0.73128092 |
| Zscan29 | 491.109012 | 0.06998831 | 0.37213149 | 0.18807413 | 0.85081854 | 0.9679302 |
| Rps27rt | 4792.17449 | 0.62035741 | 0.49930422 | 1.24244377 | 0.21407288 | 0.64772533 |
| Gpd1l | 2342.16235 | -0.6312734 | 0.16560552 | -3.8119106 | 0.0001379 | 0.00475181 |
| Ubald2 | 323.492936 | 0.18050469 | 0.34689481 | 0.52034417 | 0.60282372 | 0.89168472 |
| Gpatch11 | 367.170681 | 0.02274863 | 0.3288029 | 0.06918623 | 0.94484138 | 0.99019648 |
| Ccdc96 | 21.2109059 | 0.35797256 | 1.95561094 | 0.18304897 | 0.8547596 | 0.96862197 |
| Prkaa1 | 973.401156 | -0.2644244 | 0.24669038 | -1.0718878 | 0.28377045 | 0.71951357 |
| 2310061I04R | 506.004109 | -0.6206821 | 0.3954843 | -1.5694227 | 0.11654947 | 0.52206138 |
| Ftl1 | 258900.943 | 0.42233517 | 0.13492198 | 3.13021761 | 0.00174677 | 0.03615604 |
| Zbtb26 | 39.6386767 | -0.0276633 | 0.78445465 | -0.0352644 | 0.97186893 | 0.99443087 |
| Plekho2 | 3939.17374 | 0.05979772 | 0.35143507 | 0.17015297 | 0.86488984 | 0.9705375 |
| Arhgap42 | 34.0738021 | 1.68579081 | 0.77507318 | 2.17500858 | 0.02962947 | 0.25669202 |
| Vamp8 | 7200.48878 | 0.4244883 | 0.27542503 | 1.54121175 | 0.12326526 | 0.53358242 |
| Ptges | 33.5191004 | -0.1600525 | 1.46097834 | -0.1095516 | 0.91276498 | 0.98292561 |
| Tmem37 | 3209.55616 | 1.75908542 | 0.29629355 | 5.93696833 | 2.903407178 | 4.865394523 |
| Ccdc126 | 177.878954 | 0.81248074 | 0.55960693 | 1.45187755 | 0.14653566 | 0.57365079 |
| B3galt6 | 468.406757 | -0.3194069 | 0.39890076 | -0.8007177 | 0.42329511 | 0.80870984 |
| Ecpas | 1138.76103 | -0.4886976 | 0.35748372 | -1.3670486 | 0.17161006 | 0.60430173 |
| Zfp623 | 450.416665 | -0.3993753 | 0.30351953 | -1.3158141 | 0.18823643 | 0.61935434 |
| Zfp940 | 57.1661666 | 1.15795229 | 0.78466023 | 1.47573719 | 0.14001446 | 0.56396008 |
| Atp5k | 1353.31062 | 0.35496863 | 0.42636707 | 0.8325423 | 0.40510291 | 0.79929418 |
| Clrn3 | 11.3878028 | 6.00843182 | 1.77489209 | 3.38523782 | 0.00071117 | 0.01793752 |
| Pdik1l | 537.656362 | 0.05614235 | 0.43953037 | 0.12773259 | 0.8983606 | 0.97989497 |
| Tatdn1 | 569.446684 | -0.1975373 | 0.38930246 | -0.5074136 | 0.61186467 | 0.8961854 |
| Tvp23a | 13.6707906 | -0.2887397 | 2.09830658 | -0.1376061 | 0.89055176 | 0.97871887 |
| Cdr2l | 26.4666095 | -2.5431526 | 1.1216759 | -2.2672793 | 0.02337317 | 0.22004437 |
| Tmem123 | 6611.28218 | -0.2509408 | 0.17411351 | -1.4412485 | 0.14951448 | 0.57685486 |
| Ankrd37 | 324.08518 | 0.32698555 | 0.4002728 | 0.81690676 | 0.41398171 | 0.80395688 |
| Zfp366 | 32.1463751 | 4.22410109 | 1.56730072 | 2.69514398 | 0.00703582 | 0.10256275 |
| Map10 | 133.631122 | 1.42858242 | 0.57384068 | 2.48951052 | 0.01279191 | 0.15120563 |
| Sgms2 | 48.6793127 | -4.9789071 | 1.11124992 | -4.4804567 | 7.448349788 | 0.00040496 |
| Zfp438 | 145.486094 | 0.26090401 | 0.36071272 | 0.72330138 | 0.46949474 | 0.83381643 |
| Gja1 | 1660.59754 | 1.85512671 | 0.31351545 | 5.91717804 | 3.275121194 | 5.341941009 |

| | | | | | | |
|---|---|---|---|---|---|---|
| Zfp169 | 54.92468 | 0.29188786 | 0.97669708 | 0.29885198 | 0.76505299 | 0.94334172 |
| Insl6 | 32.1738222 | 1.023243 | 0.91550214 | 1.117685 | 0.26370155 | 0.70008189 |
| Prkca | 177.526653 | -0.0642718 | 0.57347288 | -0.1120747 | 0.91076415 | 0.98288717 |
| Lin28a | 113.180229 | -1.355575 | 0.54251495 | -2.4986869 | 0.01246544 | 0.14871281 |
| Creg2 | 9.48400098 | 2.2651327 | 1.98051471 | 1.1437091 | 0.2527443 | 0.68630877 |
| Gdpgp1 | 465.841896 | -0.5076802 | 0.34125117 | -1.4877026 | 0.13682934 | 0.55946682 |
| Selenon | 376.312138 | 0.29113201 | 0.40327568 | 0.7219181 | 0.47034483 | 0.83381643 |
| Adgb | 102.989373 | 0.56080904 | 0.57400927 | 0.97700345 | 0.32856745 | 0.74843736 |
| Gatd1 | 1582.34657 | 0.19911061 | 0.22391407 | 0.88922778 | 0.37388068 | 0.77748705 |
| Hs3st1 | 105.047758 | 0.59734951 | 0.54787391 | 1.09030471 | 0.27557894 | 0.71181528 |
| Zfp11 | 78.6530424 | 0.89350809 | 1.30317195 | 0.68564097 | 0.49293954 | 0.84609645 |
| Zfp455 | 155.227076 | -0.2638748 | 0.58818353 | -0.4486266 | 0.65370106 | 0.91076467 |
| Gprc5c | 373.013103 | 2.56045897 | 0.49342159 | 5.18919115 | 2.112094997 | 1.913870970 |
| Mblac2 | 120.642213 | 0.30594454 | 0.39760999 | 0.76945888 | 0.44162095 | 0.81967485 |
| Fam71e1 | 8.20498999 | 0.58064783 | 1.2009101 | 0.48350649 | 0.62873615 | 0.90276005 |
| Olfr77 | 124.654535 | -0.1259268 | 0.40465819 | -0.3111931 | 0.75565385 | 0.94095211 |
| Gimap9 | 147.603895 | 0.091793 | 0.46553224 | 0.19717861 | 0.84368777 | 0.96765585 |
| Nat2 | 126.979271 | 0.47553937 | 0.59943776 | 0.793309 | 0.42759782 | 0.81062404 |
| Commd3 | 4062.36446 | 0.69603836 | 0.37847273 | 1.83907135 | 0.06590469 | 0.39558985 |
| Eml5 | 36.2120669 | -0.8278798 | 1.51729819 | -0.5456276 | 0.58532192 | 0.8848945 |
| Rpusd3 | 324.20105 | -0.3241319 | 0.36244559 | -0.8942913 | 0.37116607 | 0.7765614 |
| Zfp524 | 330.818091 | -0.0950157 | 0.31642365 | -0.3002799 | 0.76396369 | 0.94300309 |
| Fam174a | 1663.84773 | 0.22927398 | 0.26378888 | 0.86915711 | 0.3847612 | 0.78748265 |
| Gpr183 | 5063.75461 | -0.5849809 | 0.1828419 | -3.1993811 | 0.00137723 | 0.02950552 |
| Ercc6l | 1630.65649 | -0.1307721 | 0.23656756 | -0.5527895 | 0.58040754 | 0.88286755 |
| Bzw1 | 27478.1233 | 0.08816079 | 0.21271201 | 0.41446079 | 0.67853668 | 0.92003316 |
| Tceanc | 328.670999 | -0.2079445 | 0.33264053 | -0.6251327 | 0.53188395 | 0.8643494 |
| Tmem199 | 2094.64048 | 0.46039769 | 0.34408023 | 1.33805331 | 0.18087906 | 0.61136731 |
| Rnf7 | 5691.4403 | 0.56021895 | 0.36327828 | 1.5421207 | 0.12304426 | 0.5331037 |
| Gen1 | 482.638762 | -0.200466 | 0.32367194 | -0.6193494 | 0.53568619 | 0.86539693 |
| Msrb3 | 17.0219983 | -2.2178675 | 1.33928263 | -1.6560116 | 0.09771948 | 0.47892724 |
| Swsap1 | 37.02056 | -1.8305524 | 0.75409417 | -2.4274852 | 0.01520391 | 0.17040121 |
| Jagn1 | 2326.71277 | 0.31308247 | 0.33834607 | 0.92533206 | 0.35479321 | 0.76419569 |
| Trap1a | 18.0471237 | 1.5415783 | 2.0102489 | 0.76685942 | 0.4431651 | 0.82038676 |
| Zgrf1 | 218.171259 | -0.3927398 | 0.59368454 | -0.6615294 | 0.50827284 | 0.85238774 |
| Pcmtd1 | 450.761555 | 0.00062712 | 0.30362945 | 0.00206541 | 0.99835204 | 0.99967651 |
| Usp42 | 294.546767 | -0.4895736 | 0.37446018 | -1.3074115 | 0.19107298 | 0.62228856 |
| Taf7 | 1196.31396 | 0.26119444 | 0.25789295 | 1.01280181 | 0.31115484 | 0.74011268 |
| Mtln | 219.256117 | 0.29701935 | 0.57586817 | 0.51577664 | 0.60601042 | 0.89258167 |
| Nup160 | 971.3358 | -0.1978459 | 0.34740316 | -0.5694995 | 0.56901718 | 0.87961466 |
| Gfod1 | 18.6178855 | 0.3528093 | 1.81402117 | 0.19449018 | 0.84579209 | 0.96765585 |
| 2900026A02I | 511.933121 | -0.0577038 | 0.43259638 | -0.1333896 | 0.89388529 | 0.97941957 |
| Zfp52 | 441.041477 | 0.32342219 | 0.40025039 | 0.80804966 | 0.419062 | 0.80529433 |
| Rab11fip5 | 70.6761962 | 1.38831928 | 0.53260768 | 2.60664527 | 0.0091434 | 0.12230094 |
| Plekhm3 | 369.316409 | 0.1305992 | 0.32996054 | 0.39580247 | 0.69225075 | 0.92539651 |
| Spryd4 | 1073.97539 | -0.2910574 | 0.34247257 | -0.8498706 | 0.39539703 | 0.79332326 |

| | | | | | | |
|---|---|---|---|---|---|---|
| Zfp46 | 364.91058 | -0.039655 | 0.44590713 | -0.0889311 | 0.92913669 | 0.98669576 |
| Commd1 | 4851.9664 | 0.41892115 | 0.34132378 | 1.22734242 | 0.21969392 | 0.65289787 |
| Plpp7 | 152.673269 | 0.76169391 | 0.63436394 | 1.2007207 | 0.22985956 | 0.66443101 |
| Kif18b | 380.156332 | -0.4297705 | 0.3570288 | -1.2037417 | 0.22868942 | 0.66258779 |
| Flrt3 | 287.697987 | 0.20282663 | 0.51032225 | 0.39744814 | 0.69103702 | 0.92470267 |
| Zbtb22 | 253.132316 | -0.464725 | 0.28354794 | -1.6389644 | 0.10122068 | 0.48631215 |
| Ywhag | 4116.78385 | -0.3843692 | 0.27225768 | -1.4117845 | 0.15801344 | 0.58698345 |
| Gm45902 | 262.841189 | -0.634339 | 0.3920496 | -1.6180069 | 0.1056611 | 0.49732676 |
| Tacstd2 | 17.6252803 | -5.7113627 | 1.72451145 | -3.3118729 | 0.00092674 | 0.02201314 |
| Ppp1r37 | 26.2695801 | 0.20246784 | 1.11163481 | 0.18213521 | 0.85547662 | 0.96915653 |
| Vamp7 | 2405.26834 | 0.75821677 | 0.35198438 | 2.1541205 | 0.03123072 | 0.26439131 |
| Plagl2 | 1504.47679 | -0.268973 | 0.34295786 | -0.7842741 | 0.43287932 | 0.81435873 |
| Ccdc157 | 17.0846336 | 0.08261042 | 2.00767816 | 0.04114724 | 0.96717851 | 0.99443087 |
| Fhad1 | 38.2306183 | -0.0910177 | 0.74892248 | -0.1215316 | 0.90327 | 0.98043702 |
| Cd14 | 2568.73284 | 0.13470206 | 0.25469329 | 0.5288795 | 0.59688904 | 0.88891958 |
| Bbs12 | 152.375926 | 0.44763067 | 1.12825309 | 0.39674669 | 0.69155427 | 0.92497085 |
| Crebzf | 104.430039 | -1.0016698 | 0.64823307 | -1.545231 | 0.1222904 | 0.53143108 |
| Hspb3 | 24.4133088 | -1.48292 | 1.07097337 | -1.3846469 | 0.16616046 | 0.59942225 |
| Spn | 49.9917737 | -6.4084062 | 1.15539363 | -5.5465134 | 2.914223652 | 3.395209327 |
| Zfp24 | 1318.78068 | 0.05441455 | 0.25834375 | 0.21062847 | 0.83317719 | 0.96565375 |
| Cbr1 | 718.182135 | 1.42398708 | 0.42839104 | 3.3240356 | 0.00088725 | 0.02124014 |
| Irf2bp2 | 859.56131 | -0.0290032 | 0.27472424 | -0.1055722 | 0.91592177 | 0.98379761 |
| Tlr6 | 361.885259 | 0.91251004 | 0.5442971 | 1.67649257 | 0.09364175 | 0.46774991 |
| Ufsp1 | 43.1844695 | -0.5907146 | 0.79933779 | -0.739005 | 0.45990399 | 0.82987212 |
| Gm6583 | 35.510543 | 1.25284515 | 1.52302901 | 0.82260098 | 0.41073494 | 0.80139357 |
| Siglech | 429.919899 | 0.69143248 | 0.35279732 | 1.95985753 | 0.05001245 | 0.34252555 |
| Wdfy4 | 1241.55825 | -0.0637996 | 0.25503316 | -0.2501619 | 0.80246211 | 0.95658926 |
| Mafg | 1562.71602 | -0.5627625 | 0.27847645 | -2.020862 | 0.04329405 | 0.31923815 |
| Fam181b | 11.3331641 | -0.5222901 | 1.17962658 | -0.4427589 | 0.65794016 | 0.91258818 |
| Arhgef39 | 1244.41847 | -0.0805491 | 0.19427492 | -0.4146139 | 0.67842454 | 0.91998308 |
| Rps19bp1 | 1185.15522 | 0.00681615 | 0.21588383 | 0.03157323 | 0.97481239 | 0.9944856 |
| Pusl1 | 1411.09134 | -0.1226163 | 0.27517871 | -0.445588 | 0.65589487 | 0.91165371 |
| Tceal8 | 2850.75542 | 0.70847717 | 0.21908695 | 3.23377173 | 0.00122167 | 0.0271397 |
| Mical3 | 85.5228668 | -0.1689159 | 0.612975 | -0.2755674 | 0.78288036 | 0.95075701 |
| Rap2a | 256.044389 | 0.20435583 | 0.2988393 | 0.68383187 | 0.49408134 | 0.84678075 |
| Hist1h1e | 232.501648 | -0.2534355 | 0.38173576 | -0.6639031 | 0.50675235 | 0.85129106 |
| Gm5812 | 15.3959055 | 1.98618091 | 1.62073247 | 1.22548351 | 0.2203931 | 0.65381019 |
| B3gnt2 | 939.091604 | -0.0717105 | 0.37764946 | -0.1898863 | 0.84939823 | 0.96765585 |
| Coa6 | 641.735837 | 0.35637523 | 0.4043161 | 0.88142725 | 0.37808661 | 0.78087685 |
| Dcun1d4 | 406.705322 | -0.24083 | 0.44898982 | -0.5363818 | 0.59169472 | 0.88797037 |
| Trim32 | 899.731116 | 0.23782965 | 0.26033021 | 0.91356916 | 0.36094328 | 0.76887042 |
| Treml4 | 50.1092924 | 0.81883314 | 1.43946684 | 0.56884474 | 0.56946151 | 0.87961466 |
| Pcbp1 | 8427.01943 | -0.0821031 | 0.34633581 | -0.2370621 | 0.81260858 | 0.95967622 |
| Senp8 | 278.465429 | 0.08351938 | 0.29454855 | 0.28355047 | 0.7767549 | 0.94800948 |
| Apon | 18.834083 | -1.5168266 | 1.4489146 | -1.0468709 | 0.29515911 | 0.72795995 |
| Wdcp | 597.797532 | -0.2832049 | 0.42812615 | -0.6614988 | 0.50829249 | 0.85238774 |

| | | | | | | |
|---|---|---|---|---|---|---|
| Kctd14 | 38.8982757 | -1.4319435 | 1.0134065 | -1.4130001 | 0.1576557 | 0.58620126 |
| Mettl5 | 638.606923 | -0.1572366 | 0.33767118 | -0.4656501 | 0.64146596 | 0.90700632 |
| Rinl | 3381.26332 | -0.287545 | 0.16561791 | -1.736195 | 0.08252936 | 0.44183004 |
| Fam229b | 55.6792784 | 0.8655638 | 0.89051936 | 0.9719764 | 0.33106229 | 0.75011761 |
| Wfdc21 | 10.4152601 | -3.7193679 | 1.61203153 | -2.3072551 | 0.0210406 | 0.2072381 |
| Xrcc1 | 1916.03369 | -0.3049854 | 0.20899981 | -1.4592617 | 0.14449308 | 0.57027792 |
| Tubgcp6 | 238.266754 | -0.5502975 | 0.68608755 | -0.8020805 | 0.42250638 | 0.80821275 |
| Cox6b2 | 21.6537538 | -0.9030045 | 0.89447967 | -1.0095305 | 0.31272028 | 0.74065408 |
| Sox12 | 24.5754526 | -0.1796915 | 0.85998128 | -0.2089482 | 0.83448867 | 0.96641606 |
| Rtl8c | 40.6425778 | 0.33043768 | 0.77825303 | 0.42458901 | 0.67113631 | 0.91711586 |
| Arf3 | 1287.11082 | -0.2552705 | 0.23928825 | -1.066791 | 0.28606625 | 0.72202931 |
| Mest | 2601.49376 | 2.44997073 | 1.14032824 | 2.14847852 | 0.03167576 | 0.26539272 |
| Tbc1d22a | 1618.11682 | 0.13530226 | 0.20124793 | 0.67231627 | 0.50138238 | 0.84968141 |
| Klhdc1 | 106.741432 | -0.3350927 | 0.52216169 | -0.6417413 | 0.52104117 | 0.85774412 |
| Spats2 | 491.123571 | 0.36676372 | 0.24586641 | 1.49171948 | 0.1357727 | 0.55758574 |
| B3glct | 594.682102 | 0.93204023 | 0.25461313 | 3.66061337 | 0.00025161 | 0.00775308 |
| Prdm9 | 84.7598069 | -1.2906588 | 0.92092885 | -1.401475 | 0.16107207 | 0.59141542 |
| Erich1 | 290.786822 | 0.08986759 | 0.37389847 | 0.24035293 | 0.81005666 | 0.95882179 |
| Sec31b | 12.2007098 | 1.27702441 | 1.28161093 | 0.99642129 | 0.31904549 | 0.74398524 |
| Smim11 | 1017.07058 | 0.32225567 | 0.34152886 | 0.9435679 | 0.3453905 | 0.7602916 |
| Btla | 161.520029 | -4.3774319 | 0.90847718 | -4.818428 | 1.446937113 | 9.833545395 |
| Tagap1 | 563.355751 | 0.16987002 | 0.3170957 | 0.53570586 | 0.59216185 | 0.88797037 |
| Pfdn4 | 585.721837 | -0.2441238 | 0.30422303 | -0.8024502 | 0.42229259 | 0.80821275 |
| Klf13 | 3014.6879 | -0.2734188 | 0.21973721 | -1.244299 | 0.21338954 | 0.64625661 |
| Zfp217 | 291.06202 | -0.7938983 | 0.49749094 | -1.5958046 | 0.11053243 | 0.50851567 |
| Dock8 | 646.848748 | 0.3789358 | 0.32135377 | 1.17918579 | 0.2383242 | 0.67446139 |
| Rgs14 | 921.41065 | -0.2505095 | 0.23737514 | -1.0553317 | 0.29127365 | 0.72536876 |
| Gnpda1 | 6944.72775 | 0.84056109 | 0.21147228 | 3.97480501 | 7.043694172 | 0.00270961 |
| Rbm12b2 | 294.633812 | 0.57270441 | 0.4338224 | 1.32013564 | 0.18678973 | 0.61862792 |
| Babam2 | 2237.69788 | -0.2555693 | 0.21613606 | -1.1824462 | 0.23702869 | 0.6732629 |
| Rasal3 | 1749.09608 | -1.0357871 | 0.36890572 | -2.8077284 | 0.00498923 | 0.07978199 |
| Ppp4r2 | 843.164847 | -0.3211834 | 0.21889814 | -1.4672735 | 0.14230167 | 0.56795311 |
| Rps10 | 11062.216 | -0.0099445 | 0.3455721 | -0.0287768 | 0.97704256 | 0.99483019 |
| Plpp2 | 552.735727 | 0.34802266 | 0.26794844 | 1.2988419 | 0.19399819 | 0.62565369 |
| Acvr2a | 127.864542 | 0.92559755 | 0.52546246 | 1.7614913 | 0.07815528 | 0.43258005 |
| Pld4 | 13744.5244 | -0.5493172 | 0.16681786 | -3.2929158 | 0.00099154 | 0.02319222 |
| Hbb-y | 1006.09214 | 0.22988682 | 1.20479264 | 0.19081028 | 0.84867423 | 0.96765585 |
| Opa3 | 2580.63354 | 0.0279491 | 0.1704612 | 0.16396168 | 0.86976131 | 0.97239152 |
| Hbb-bh1 | 932.933011 | 1.58910674 | 1.35969576 | 1.16872229 | 0.24251554 | 0.67794621 |
| Zfp622 | 2518.07816 | 0.29423087 | 0.43935402 | 0.66968973 | 0.5030556 | 0.85012672 |
| Taf9 | 186.709244 | -0.4589464 | 0.45638117 | -1.0056209 | 0.31459799 | 0.74066151 |
| Ppp6r1 | 327.711316 | -0.1703438 | 0.38375256 | -0.4438897 | 0.65712233 | 0.91251438 |
| Cdc42se2 | 8213.16202 | 0.35990811 | 0.15325948 | 2.34835793 | 0.01885639 | 0.19384053 |
| Ltn1 | 818.198495 | -0.3588554 | 0.27862123 | -1.2879686 | 0.1977569 | 0.6306972 |
| Tbc1d30 | 7.00117766 | -0.5530711 | 2.33050802 | -0.2373178 | 0.81241024 | 0.95967622 |
| Slc39a1 | 7009.19811 | 0.64249594 | 0.16217803 | 3.96167062 | 7.442716726 | 0.00281878 |

| | | | | | | |
|---|---|---|---|---|---|---|
| Ankrd44 | 479.012932 | -0.2488552 | 0.3105189 | -0.8014172 | 0.42289018 | 0.80844471 |
| Cx3cr1 | 45399.1043 | -0.0502505 | 0.1918531 | -0.2619217 | 0.79338182 | 0.95310033 |
| Immt | 12974.3554 | 0.07906199 | 0.14303578 | 0.55274272 | 0.5804396 | 0.88286755 |
| Tmem106c | 1336.52116 | 0.20336375 | 0.20520475 | 0.99102848 | 0.32167167 | 0.74447686 |
| Nrros | 18465.5808 | 0.18702428 | 0.17095401 | 1.09400347 | 0.27395346 | 0.7109191 |
| Acot4 | 13.0233421 | 5.28376292 | 1.72336789 | 3.06595181 | 0.00216978 | 0.04253678 |
| Rft1 | 966.398434 | -0.1031387 | 0.26152491 | -0.3943743 | 0.69330471 | 0.92579374 |
| Mogat2 | 100.957108 | 3.53048286 | 1.3569362 | 2.60180461 | 0.00927347 | 0.12344105 |
| Ezr | 3006.40219 | -0.3602674 | 0.25208358 | -1.4291585 | 0.15295867 | 0.58128098 |
| Rexo4 | 2249.86973 | -0.1394886 | 0.1575094 | -0.8855892 | 0.37583891 | 0.77828094 |
| Ccdc171 | 32.9235402 | 0.89610176 | 1.33207412 | 0.67271164 | 0.50113076 | 0.84968141 |
| 2610001J05F | 4314.30485 | 0.47179155 | 0.26752575 | 1.76353696 | 0.07780997 | 0.4314473 |
| B4galt3 | 1119.84308 | -0.026423 | 0.25748544 | -0.1026194 | 0.91826505 | 0.98390186 |
| Tmco1 | 5772.01463 | 0.08962516 | 0.27053631 | 0.33128697 | 0.74042773 | 0.93800224 |
| Cebpe | 51.6964561 | -3.8514453 | 0.91125162 | -4.2265443 | 2.373076088 | 0.00104424 |
| Zfp961 | 579.418485 | 0.38912713 | 0.40834844 | 0.95292916 | 0.34062597 | 0.75821364 |
| Asna1 | 1289.66297 | 0.1075051 | 0.22353985 | 0.4809214 | 0.63057236 | 0.9036296 |
| Atp6v1a | 10573.9406 | 0.34192975 | 0.15040336 | 2.27341836 | 0.02300098 | 0.2172749 |
| Cherp | 1752.81968 | -0.2230719 | 0.28315857 | -0.7877986 | 0.43081452 | 0.81254302 |
| Nav2 | 31.8095744 | 1.81112469 | 1.58001616 | 1.14626972 | 0.25168357 | 0.68505999 |
| Spdya | 109.230617 | 0.13333069 | 0.44689498 | 0.29834904 | 0.76543678 | 0.94347227 |
| Nup188 | 885.788986 | 0.5313428 | 0.29360028 | 1.80974898 | 0.07033472 | 0.40971651 |
| Pbx1 | 114.743707 | 0.38487528 | 0.45442597 | 0.84694824 | 0.39702397 | 0.79502279 |
| Magi3 | 74.6617169 | -0.321684 | 1.00273369 | -0.320807 | 0.74835667 | 0.9399191 |
| St6galnac3 | 132.627161 | -0.2365494 | 0.68903988 | -0.3433029 | 0.73137063 | 0.93653187 |
| Gan | 111.691869 | -0.6290298 | 0.46556327 | -1.3511156 | 0.1766584 | 0.60784615 |
| Cpne8 | 117.824005 | 2.86788631 | 0.82819904 | 3.4627984 | 0.00053459 | 0.01415503 |
| Hist1h1d | 30.6305581 | 0.56962617 | 0.71971935 | 0.79145597 | 0.42867796 | 0.81136783 |
| Hook2 | 259.919965 | 0.29296199 | 0.43284367 | 0.67683095 | 0.49851322 | 0.84870979 |
| Adam17 | 5880.87294 | 0.32250616 | 0.23789147 | 1.35568609 | 0.17519907 | 0.60743838 |
| Isoc2b | 187.371173 | 0.64019069 | 0.48286017 | 1.32583039 | 0.18489589 | 0.61700012 |
| Plekhg3 | 92.9341698 | 0.88219147 | 0.71161455 | 1.23970411 | 0.21508486 | 0.64854028 |
| Asap2 | 25.9680025 | 0.69447064 | 0.898227 | 0.77315716 | 0.43942938 | 0.81963526 |
| Rnf103 | 165.357734 | 0.02161553 | 0.48766229 | 0.0443248 | 0.96464551 | 0.99406401 |
| Zmat1 | 32.1550136 | 0.39285159 | 1.87880345 | 0.20909669 | NA | NA |
| Rap1b | 8960.53348 | 0.76699148 | 0.22289828 | 3.44099329 | 0.00057958 | 0.01518209 |
| Jun | 26098.8692 | 0.22136941 | 0.25718226 | 0.86074917 | 0.38937621 | 0.78942657 |
| Rab7b | 2451.11326 | 0.06945073 | 0.29288167 | 0.23712897 | 0.81255674 | 0.95967622 |
| Tln2 | 1866.80454 | 1.97594577 | 0.36096062 | 5.47413114 | 4.396634004 | 4.822428606 |
| Tnrc6a | 615.270586 | 0.08840129 | 0.34658914 | 0.25506077 | 0.79867616 | 0.9547279 |
| BC004004 | 2404.6802 | 0.16484406 | 0.25248623 | 0.65288337 | 0.51383147 | 0.85497712 |
| Zfp608 | 141.794264 | -0.3256601 | 0.55068929 | -0.5913682 | 0.55427375 | 0.87165957 |
| Kcnt2 | 9.59212342 | 0.43413593 | 1.79860217 | 0.24137407 | 0.80926521 | 0.95844313 |
| Map1b | 212.220155 | 2.05070094 | 0.74298847 | 2.76007101 | 0.00577888 | 0.08924119 |
| Suclg1 | 7585.3332 | -0.0199959 | 0.27293292 | -0.0732631 | 0.94159672 | 0.98948137 |
| Swt1 | 882.352521 | 0.22705537 | 0.27492769 | 0.82587303 | 0.40887612 | 0.80100711 |

| | | | | | | |
|---|---|---|---|---|---|---|
| Trim30b | 25.4042559 | 0.29461114 | 1.54217665 | 0.19103592 | 0.84849745 | 0.96765585 |
| Repin1 | 525.029556 | 0.20370984 | 0.34871554 | 0.58417194 | 0.55910464 | 0.87394928 |
| Traf7 | 3842.32289 | -0.3958583 | 0.29178504 | -1.3566777 | 0.17488364 | 0.60708046 |
| A630001G21 | 954.738412 | 0.12387825 | 0.18838517 | 0.65757964 | 0.51080828 | 0.85359566 |
| Zfp212 | 509.395497 | -0.0368275 | 0.25867478 | -0.14237 | 0.88678774 | 0.97827644 |
| Oas1a | 494.005096 | -0.2798919 | 0.36299675 | -0.7710589 | 0.44067204 | 0.81963526 |
| Grk4 | 17.4547739 | -0.3835347 | 0.97930031 | -0.3916415 | 0.69532309 | 0.92616369 |
| 1700030K09l | 384.529094 | 0.04163951 | 0.30080815 | 0.13842546 | 0.88990418 | 0.97871887 |
| Nup107 | 4893.37973 | -0.3087382 | 0.17269484 | -1.7877674 | 0.07381354 | 0.41842493 |
| Atad2b | 142.930887 | 0.40541661 | 0.43683629 | 0.92807448 | 0.35336894 | 0.76383312 |
| Cysltr1 | 1230.20839 | 0.36010628 | 0.20245265 | 1.77871853 | 0.07528592 | 0.4226584 |
| Sae1 | 14250.9572 | -0.260866 | 0.18434873 | -1.4150681 | 0.15704852 | 0.58620126 |
| Junb | 9426.4334 | 0.69501199 | 0.47962751 | 1.44906615 | 0.14731911 | 0.57410946 |
| Reep1 | 108.066471 | 0.05066422 | 0.59887636 | 0.0845988 | 0.93258035 | 0.98738166 |
| Prkcb | 310.136632 | 0.2428429 | 0.34516417 | 0.70355767 | 0.48170828 | 0.84022786 |
| Ubxn8 | 1198.09016 | -0.0585101 | 0.19233767 | -0.304205 | 0.76097173 | 0.94244354 |
| Gm9894 | 32.1299716 | 0.64815371 | 0.88990727 | 0.72833848 | 0.46640641 | 0.83345004 |
| Lamb2 | 61.7428442 | 1.04461672 | 1.26597793 | 0.82514608 | 0.40928866 | 0.80100711 |
| Smarca5-ps | 18.1819792 | -0.8225949 | 1.32715718 | -0.6198173 | 0.5353781 | 0.86539693 |
| Cyp2j6 | 82.9212149 | 4.84937427 | 1.51740732 | 3.19582898 | 0.0013943 | 0.02971505 |
| Msl1 | 517.07125 | -0.0526809 | 0.38590518 | -0.1365126 | 0.8914161 | 0.97905307 |
| Senp7 | 211.701444 | 0.29473966 | 0.43249625 | 0.68148489 | 0.49556473 | 0.84727369 |
| Rnaseh2a | 2628.62475 | 0.37185065 | 0.35222679 | 1.05571371 | 0.29109903 | 0.72510985 |
| Ctif | 12.3390735 | 1.57549775 | 2.01221014 | 0.78296879 | 0.43364544 | 0.81483708 |
| Fbxo31 | 242.144436 | 0.50350838 | 0.67519504 | 0.74572287 | 0.45583484 | 0.8284397 |
| Glis3 | 17.6893563 | 7.62774504 | 1.71132034 | 4.45722806 | 8.302626110 | 0.00044352 |
| Rnf157 | 16.3009397 | 0.9169429 | 1.48507117 | 0.61744038 | 0.53694429 | 0.86596304 |
| Gas1 | 58.4810457 | 0.52258835 | 0.83554459 | 0.6254464 | 0.53167812 | 0.8643494 |
| Mrpl35 | 2702.63622 | -0.0773151 | 0.19301363 | -0.400568 | 0.6887382 | 0.92470267 |
| Uba2 | 11935.4579 | -0.1339367 | 0.19039258 | -0.7034762 | 0.48175903 | 0.84022786 |
| Creb5 | 113.289941 | 0.8505017 | 0.70002362 | 1.21496143 | 0.2243808 | 0.65793926 |
| Krcc1 | 1200.3123 | 0.46380376 | 0.28038651 | 1.65415859 | 0.09809529 | 0.47978865 |
| Sv2b | 59.7654428 | -0.4211797 | 0.90657243 | -0.4645847 | 0.6422289 | 0.90705802 |
| Gm6180 | 13.918149 | 0.37656944 | 1.10287005 | 0.34144497 | 0.73276862 | 0.93728136 |
| Aph1c | 348.181745 | 0.73785895 | 0.25924867 | 2.84614364 | 0.00442522 | 0.07285835 |
| Clec12a | 2290.98864 | -1.6970654 | 0.3637606 | -4.6653359 | 3.081132944 | 0.00019036 |
| Cfap300 | 25.6725162 | 0.84170484 | 1.40672237 | 0.59834468 | 0.54960997 | 0.87093412 |
| 2700081O15 | 30.3637029 | 0.28061042 | 0.95618958 | 0.29346735 | 0.76916495 | 0.94401697 |
| Lins1 | 306.407986 | 0.13800843 | 0.57922953 | 0.23826207 | 0.81167784 | 0.95967622 |
| Tmem248 | 2553.97209 | 0.16220114 | 0.4430684 | 0.366086 | 0.7143009 | 0.93235217 |
| Gpr141 | 572.57693 | -3.5459549 | 0.42834343 | -8.278299 | 1.249471777 | 6.368661773 |
| Socs3 | 1140.19627 | 0.13714166 | 0.43745359 | 0.31349991 | 0.75390089 | 0.94018119 |
| Chmp3 | 3789.41555 | 0.33015496 | 0.18837247 | 1.75267093 | 0.07965851 | 0.43676068 |
| Rnf26 | 1054.7349 | 0.07763059 | 0.4310871 | 0.18008098 | 0.85708899 | 0.96936093 |
| Supt7l | 1026.40293 | 0.08896315 | 0.26666604 | 0.3336126 | 0.73867191 | 0.93800224 |
| Mapk11 | 39.2769817 | 2.4723489 | 1.66757157 | 1.48260437 | 0.13817955 | 0.56068131 |

| | | | | | | |
|---|---|---|---|---|---|---|
| Fes | 8215.1768 | -0.3674807 | 0.21362685 | -1.7201991 | 0.08539626 | 0.44669728 |
| Rpl18-ps2 | 12.8711726 | 0.31982516 | 1.06883922 | 0.29922663 | 0.76476713 | 0.94317938 |
| Bcl3 | 127.03052 | 0.36431433 | 0.77490438 | 0.47014101 | 0.63825428 | 0.90599527 |
| Mterf1b | 38.1291787 | 1.36437223 | 1.26913404 | 1.07504187 | 0.28235598 | 0.71787592 |
| Mllt11 | 354.971544 | 0.42741184 | 0.45939717 | 0.93037544 | 0.35217673 | 0.76348958 |
| Prx | 29.3551292 | 1.0103805 | 0.70131656 | 1.44069107 | 0.14967198 | 0.57685486 |
| Styx | 800.058677 | -0.0438419 | 0.19950697 | -0.2197513 | 0.82606488 | 0.96345784 |
| Raet1e | 16.3776305 | 0.55828276 | 1.11928591 | 0.49878476 | 0.61793102 | 0.89878056 |
| Dand5 | 65.0231467 | -0.1575601 | 0.49322105 | -0.3194512 | 0.74938438 | 0.9399191 |
| Ndfip2 | 737.244992 | 0.26817632 | 0.19967702 | 1.34305054 | 0.17925561 | 0.60992118 |
| Trmt1l | 1309.21493 | -0.1518738 | 0.26585951 | -0.5712557 | 0.56782632 | 0.87929304 |
| Ddx10 | 1874.01469 | -0.3452434 | 0.2655426 | -1.3001433 | 0.19355184 | 0.62506533 |
| Rab4b | 1509.37042 | -0.0114275 | 0.2105174 | -0.0542828 | 0.95670988 | 0.99215259 |
| Pom121 | 581.089299 | -0.5124704 | 0.33484123 | -1.5304877 | 0.12589605 | 0.53789768 |
| AI854703 | 10.4882698 | -3.0282344 | 1.43713213 | -2.1071371 | 0.0351057 | 0.28086853 |
| Sec61b | 5124.48488 | 0.28340545 | 0.3867813 | 0.73272791 | 0.4637244 | 0.83202414 |
| Slamf8 | 92.2613786 | -0.7240313 | 0.63789629 | -1.1350297 | 0.25636286 | 0.69126266 |
| Gatd3a | 300.625143 | -0.0250517 | 0.35067979 | -0.0714375 | 0.9430496 | 0.98983253 |
| Dis3l2 | 824.875367 | 0.04912395 | 0.25707513 | 0.19108792 | 0.84845671 | 0.96765585 |
| Ficd | 158.179511 | 0.10660871 | 0.63723575 | 0.16729869 | 0.86713504 | 0.97107585 |
| Tarm1 | 179.203381 | -4.0487245 | 0.73174319 | -5.5329856 | 3.148253454 | 3.633262689 |
| Zfp943 | 330.670409 | 0.09355582 | 0.28934432 | 0.32333734 | 0.74643978 | 0.93960567 |
| Zfp952 | 486.331598 | 0.34618814 | 0.25842512 | 1.33960716 | 0.1803731 | 0.61122544 |
| Phgdh | 34844.1549 | -0.096511 | 0.29169931 | -0.3308577 | 0.74075196 | 0.93800224 |
| Cbx7 | 62.9521638 | -0.2633786 | 0.82765963 | -0.3182209 | 0.75031735 | 0.9399191 |
| Hunk | 15.7267161 | -0.8335857 | 1.5464351 | -0.539037 | 0.58986134 | 0.88722166 |
| Mapk14 | 1237.2887 | -0.1897047 | 0.27609354 | -0.6871031 | 0.49201778 | 0.84522588 |
| Thoc7 | 3059.18565 | 0.1880965 | 0.38929742 | 0.48316914 | 0.62897564 | 0.90276005 |
| Ggcx | 631.694629 | 0.43729898 | 0.32890612 | 1.32955562 | 0.18366473 | 0.61555359 |
| Kdm3a | 727.767827 | 0.46721037 | 0.27450142 | 1.70203266 | 0.08874925 | 0.45521604 |
| Tnfaip6 | 68.2725123 | -0.9980401 | 0.83105909 | -1.2009255 | 0.2297801 | 0.6643583 |
| Tcf4 | 1936.64825 | 0.39225429 | 0.21137133 | 1.85575918 | 0.06348789 | 0.3887124 |
| Usp21 | 1082.95413 | 0.41553586 | 0.26497846 | 1.56818734 | 0.11683742 | 0.52252063 |
| Nrd1 | 4078.78434 | -0.1887423 | 0.28046531 | -0.6729613 | 0.50097189 | 0.84968141 |
| Cstf2t | 1806.16494 | -0.0158965 | 0.21211667 | -0.0749424 | 0.94026052 | 0.98934006 |
| 3110082I17R | 1975.77852 | 0.13573033 | 0.3471705 | 0.3909616 | 0.69582562 | 0.92623267 |
| Smagp | 748.943493 | 1.49324449 | 0.29357423 | 5.08642901 | 3.648682095 | 2.995592488 |
| Ier2 | 9306.57344 | 0.14059941 | 0.26834012 | 0.52395971 | 0.60030656 | 0.8902352 |
| Eif3k | 6817.48169 | -0.0581016 | 0.27362609 | -0.2123395 | 0.83184215 | 0.96518306 |
| Tanc2 | 182.254721 | 1.26329022 | 0.61120869 | 2.06687214 | 0.0387462 | 0.29754068 |
| Zfand2a | 1781.729 | 0.09793802 | 0.17845132 | 0.54882203 | 0.58312759 | 0.88350258 |
| Zfp472 | 637.529861 | -0.0996367 | 0.33528136 | -0.2971734 | 0.7663341 | 0.94360659 |
| Rpia | 28.6495876 | -0.5224339 | 0.94763656 | -0.551302 | 0.58142669 | 0.88321857 |
| Sh3pxd2a | 59.5720506 | 1.2923803 | 0.90900193 | 1.42175748 | 0.15509667 | 0.58414333 |
| Dennd4a | 306.255185 | 0.5523731 | 0.41278663 | 1.33815647 | 0.18084544 | 0.61136731 |
| Aldh7a1 | 2192.56733 | 0.65823698 | 0.19039351 | 3.45724495 | 0.00054573 | 0.01441499 |

| | | | | | | |
|---|---|---|---|---|---|---|
| BC048403 | 71.5341203 | -0.5624107 | 0.53574163 | -1.0497797 | 0.2938194 | 0.72683372 |
| Dpep2 | 15.1450758 | 0.72754094 | 1.50580827 | 0.48315642 | 0.62898467 | 0.90276005 |
| Dusp7 | 438.545808 | -0.2364075 | 0.28948889 | -0.8166374 | 0.41413567 | 0.80395688 |
| Tmem39b | 164.901975 | -0.8342196 | 0.46193297 | -1.8059321 | 0.07092894 | 0.41102498 |
| Ptrh1 | 110.322543 | 0.10385512 | 0.45215115 | 0.22969116 | 0.81833176 | 0.96105733 |
| Chd8 | 423.53722 | 0.53030098 | 0.40209468 | 1.31884603 | 0.18722059 | 0.61935434 |
| Chchd3 | 3559.38086 | -0.0886342 | 0.20980407 | -0.4224619 | 0.67268789 | 0.91780571 |
| Lysmd1 | 326.473979 | 0.2694519 | 0.34448478 | 0.78218812 | 0.43410403 | 0.81510277 |
| Ubxn7 | 1083.4641 | -0.1274184 | 0.25612933 | -0.4974766 | 0.61885299 | 0.89878056 |
| Exoc6 | 2205.28132 | -0.3356809 | 0.21639865 | -1.5512152 | 0.12085012 | 0.52910763 |
| Grwd1 | 4305.2233 | -0.3541965 | 0.19240292 | -1.8409101 | 0.06563474 | 0.39558985 |
| Camk2d | 4486.43774 | 0.14707926 | 0.22417823 | 0.65608178 | 0.5117715 | 0.85393154 |
| H2-T24 | 190.210243 | -0.5941676 | 0.42135552 | -1.4101337 | 0.15850021 | 0.58731982 |
| Nudcd3 | 2529.71337 | -0.4849051 | 0.20617032 | -2.3519636 | 0.0186746 | 0.1927818 |
| Txlna | 3666.6793 | -0.2512367 | 0.23233258 | -1.0813665 | 0.27953411 | 0.714339 |
| Fpgt | 347.125905 | 1.12317863 | 0.46743085 | 2.40287658 | 0.01626668 | 0.17782865 |
| Srcap | 405.893975 | -0.1680196 | 0.25877253 | -0.6492944 | 0.51614808 | 0.85627177 |
| Slc39a8 | 1383.42925 | 1.74854014 | 0.30410269 | 5.74983445 | 8.933087863 | 1.316607998 |
| Ech1 | 4956.11375 | 0.64378583 | 0.27850944 | 2.31154045 | 0.02080302 | 0.20556007 |
| Mat2a | 14278.4584 | -0.1776032 | 0.13182154 | -1.3473006 | 0.17788345 | 0.60906934 |
| Nanp | 1017.28806 | 0.28982137 | 0.33780032 | 0.85796653 | 0.39091095 | 0.7899826 |
| Cyhr1 | 317.752213 | 0.69555114 | 0.3755548 | 1.85206297 | 0.06401678 | 0.39097218 |
| Cnn3 | 2149.40865 | 0.2661591 | 0.28670658 | 0.92833275 | 0.35323499 | 0.76376846 |
| Adnp2 | 630.863785 | -0.2179695 | 0.52484754 | -0.4153007 | 0.67792182 | 0.91965617 |
| Lgals4 | 24.1403955 | -0.3464087 | 0.85940106 | -0.4030815 | 0.68688824 | 0.92426981 |
| Pde5a | 27.999916 | -1.7453354 | 1.87918337 | -0.9287733 | 0.35300657 | 0.76376537 |
| Zfp14 | 59.3185212 | 2.70976104 | 1.27807512 | 2.12018917 | 0.0339901 | 0.27569109 |
| Ndst1 | 493.841217 | -0.325769 | 0.25014678 | -1.3023112 | 0.19281001 | 0.62447573 |
| Sirt5 | 218.920945 | -0.6008356 | 0.40725543 | -1.4753287 | 0.14012419 | 0.56396008 |
| Nt5dc3 | 39.1166361 | -1.3986233 | 0.71743947 | -1.9494652 | 0.05123989 | 0.34803863 |
| Ercc6 | 71.6649799 | -1.5743951 | 1.03859304 | -1.5158922 | 0.12954662 | 0.54639717 |
| A930004D18 | 29.6188429 | -0.0982967 | 0.75780217 | -0.1297129 | 0.89679362 | 0.97984623 |
| Pkp3 | 32.7308367 | -1.9721476 | 1.35966353 | -1.4504674 | 0.14692823 | 0.57367728 |
| Iigp1 | 22.1652752 | -0.0247691 | 1.67233727 | -0.0148111 | 0.98818292 | 0.9962454 |
| Skida1 | 27.7137358 | 1.72081212 | 1.19656599 | 1.43812555 | 0.15039844 | 0.57837915 |
| Utp18 | 2726.06002 | -0.0481767 | 0.2526105 | -0.1907153 | 0.84874865 | 0.96765585 |
| 1810037I17R | 2556.89659 | 0.42154768 | 0.41451458 | 1.01696708 | 0.30916909 | 0.73882897 |
| Slc25a40 | 206.428238 | 0.29146262 | 0.40780704 | 0.71470718 | 0.47479 | 0.83578199 |
| Skp2 | 2295.30457 | 0.51862404 | 0.21591327 | 2.40200162 | 0.01630564 | 0.17796742 |
| Syne3 | 76.3125958 | -0.7284827 | 0.53828584 | -1.3533381 | 0.17594764 | 0.60763832 |
| Klf1 | 17.4359478 | -0.9855724 | 1.22480623 | -0.8046762 | 0.42100655 | 0.80708017 |
| Cthrc1 | 13.7031338 | -0.5777162 | 1.6114987 | -0.3584962 | 0.71997199 | 0.93328572 |
| Gon4l | 480.411074 | -0.6685834 | 0.41626275 | -1.6061571 | 0.1082394 | 0.50301538 |
| Ifi205 | 112.176012 | -0.0122675 | 0.90174934 | -0.0136041 | 0.9891458 | 0.99650865 |
| Gzmm | 10.6833293 | 0.61612667 | 1.21872605 | 0.50554977 | 0.61317275 | 0.89739894 |
| Tprkb | 924.692891 | -0.0389321 | 0.3345726 | -0.1163637 | 0.9073643 | 0.98176897 |

| | | | | | | |
|---|---|---|---|---|---|---|
| Fra10ac1 | 496.05686 | -0.4639349 | 0.27885703 | -1.6637018 | 0.09617203 | 0.47560138 |
| Lifr | 1392.82499 | 1.05132721 | 0.28400574 | 3.70178149 | 0.00021409 | 0.00689204 |
| Arfgap3 | 497.868018 | 0.53156833 | 0.26053008 | 2.04033384 | 0.04131709 | 0.30948419 |
| Prr14l | 1510.83801 | 0.30848591 | 0.39761719 | 0.77583646 | 0.43784555 | 0.81866921 |
| P2ry10b | 459.880677 | 0.68729506 | 0.49045984 | 1.40132789 | 0.16111604 | 0.59141542 |
| Eapp | 1758.24401 | 0.40950826 | 0.30968382 | 1.3223431 | 0.18605393 | 0.61790851 |
| D130007C19 | 11.661536 | 1.16085059 | 1.62368358 | 0.71494878 | 0.4746407 | 0.83578199 |
| Cpsf3 | 8814.22928 | -0.280322 | 0.15238715 | -1.8395383 | 0.06583605 | 0.39558985 |
| Mrps21 | 1834.63945 | 0.08954125 | 0.27926724 | 0.32062927 | 0.74849136 | 0.9399191 |
| Taf4b | 358.704804 | -0.1537529 | 0.41698978 | -0.3687211 | 0.71233566 | 0.93235217 |
| Kcnn4 | 34.076294 | -1.4568241 | 0.86168222 | -1.6906744 | 0.090899 | 0.46044203 |
| Rhob | 2576.70899 | 0.90967907 | 0.37398425 | 2.4323994 | 0.01499916 | 0.16926632 |
| Zfp747 | 368.176969 | 0.20741723 | 0.30393457 | 0.68244041 | 0.49496051 | 0.84681245 |
| Pnma1 | 8.72936318 | -0.7007495 | 2.15644394 | -0.324956 | 0.74521435 | 0.93917909 |
| Mdm4 | 898.891743 | -0.3090557 | 0.28453089 | -1.086194 | 0.27739318 | 0.71260013 |
| Cklf | 865.033429 | 0.39647833 | 0.39538273 | 1.00277097 | 0.31597138 | 0.74186126 |
| Slfn5 | 90.4126877 | 0.23239772 | 0.58031847 | 0.40046584 | 0.68881344 | 0.92470267 |
| Dnajc8 | 5406.33107 | -0.0388169 | 0.23166497 | -0.1675563 | 0.86693234 | 0.9709928 |
| Spcs3 | 11567.3837 | 0.01659048 | 0.16929275 | 0.09799876 | 0.92193328 | 0.98437722 |
| Slc30a7 | 1976.81573 | -0.1554515 | 0.2616328 | -0.5941592 | 0.55240561 | 0.87093412 |
| Atpif1 | 12072.2903 | 0.45666823 | 0.37632149 | 1.21350558 | 0.22493658 | 0.65870021 |
| Tmem120b | 202.812106 | 0.11996964 | 0.51753651 | 0.23180903 | 0.81668634 | 0.96079676 |
| Gimap4 | 30.3433752 | 1.80276451 | 1.37026624 | 1.31563083 | 0.18829798 | 0.61935434 |
| Aes | 851.761397 | -0.171337 | 0.22614532 | -0.757641 | 0.44866595 | 0.82417561 |
| Vapb | 246.133713 | -0.0544595 | 0.28649439 | -0.1900893 | 0.84923918 | 0.96765585 |
| Lclat1 | 241.558349 | 0.17391546 | 0.49936137 | 0.34827576 | 0.72763309 | 0.93576253 |
| Thnsl2 | 185.47306 | 1.11336563 | 0.47925267 | 2.3231287 | 0.02017224 | 0.20079243 |
| Tmem62 | 132.362088 | -0.516148 | 0.79363215 | -0.6503618 | 0.51545858 | 0.85587477 |
| Dedd2 | 401.371784 | -0.1802054 | 0.29897177 | -0.6027504 | 0.54667472 | 0.86994632 |
| Parp4 | 45.0714683 | -0.0247155 | 0.74792308 | -0.0330455 | 0.97363834 | 0.99443087 |
| Trim65 | 282.640449 | 0.60489965 | 0.432905 | 1.39730346 | 0.16232234 | 0.59324932 |
| Zfp867 | 127.041556 | 2.79132242 | 0.64834401 | 4.30531072 | 1.667515122 | 0.00079372 |
| Sh3bp2 | 818.570975 | 0.06911872 | 0.27138686 | 0.25468705 | 0.79896482 | 0.95475225 |
| Ugt1a6a | 61.4428946 | -2.1732099 | 1.44661197 | -1.5022756 | 0.13302595 | 0.55290835 |
| 9130230L23R | 11.0919013 | -0.1450854 | 1.61369834 | -0.0899086 | 0.92835984 | 0.98666423 |
| Cggbp1 | 2174.90154 | 0.00870732 | 0.26438076 | 0.03293476 | 0.97372661 | 0.99443087 |
| Kdm2a | 742.154096 | -0.1410658 | 0.42470199 | -0.3321524 | 0.73977418 | 0.93800224 |
| Mgmt | 440.000692 | 0.60142234 | 0.50329287 | 1.19497488 | 0.23209684 | 0.66733333 |
| Mettl7a1 | 45.106574 | 1.04359679 | 0.82702391 | 1.26187016 | 0.20699551 | 0.64174209 |
| Xlr | 222.816918 | 0.72785033 | 0.49647744 | 1.46602901 | 0.14264038 | 0.56834178 |
| Slc8a1 | 269.690677 | -0.4718706 | 0.72696481 | -0.6490968 | 0.51627579 | 0.85627177 |
| Zfp869 | 1832.84491 | 0.20145088 | 0.28870986 | 0.69776237 | 0.48532582 | 0.84183502 |
| Tmem119 | 667.770187 | -0.3902655 | 0.31009355 | -1.2585413 | 0.20819606 | 0.6418276 |
| 1600014C10 | 1869.60167 | -1.1024806 | 0.21297093 | -5.1766718 | 2.258790082 | 2.016918180 |
| Emcn | 209.599279 | 3.26590068 | 2.27328784 | 1.43664195 | 0.15081976 | 0.57859667 |
| Adam10 | 4731.80689 | -0.0121462 | 0.26353921 | -0.0460886 | 0.96323962 | 0.99401755 |

| | | | | | | |
|---|---|---|---|---|---|---|
| Ap1s3 | 12.9945976 | -2.6762117 | 2.4385708 | -1.0974509 | 0.27244434 | 0.71005689 |
| Ankrd24 | 81.6863303 | 0.64841167 | 0.76986872 | 0.84223667 | 0.39965549 | 0.79612475 |
| Zscan22 | 499.776011 | -0.3544194 | 0.36177474 | -0.9796687 | 0.32724966 | 0.74811039 |
| Zfp771 | 479.177137 | -0.1390481 | 0.25726801 | -0.5404794 | 0.58886647 | 0.88690478 |
| Hmgb2 | 23632.0938 | 0.02944523 | 0.33052946 | 0.08908505 | 0.92901432 | 0.98669576 |
| Lrrc8c | 414.455194 | -0.0193076 | 0.44918416 | -0.0429838 | 0.96571444 | 0.99443087 |
| Vmac | 274.187153 | 0.23643314 | 0.32491161 | 0.72768449 | 0.46680675 | 0.8334534 |
| Phactr1 | 193.376233 | -0.1609429 | 0.3745598 | -0.4296854 | 0.66742453 | 0.91595755 |
| Msra | 847.415575 | -0.2859854 | 0.23461613 | -1.2189502 | 0.2228631 | 0.65613364 |
| Zfp182 | 122.955475 | 0.63300627 | 0.58411012 | 1.0837105 | 0.27849318 | 0.71377266 |
| Set | 35163.8915 | -0.5577958 | 0.16500339 | -3.380511 | 0.00072351 | 0.01813672 |
| Kctd18 | 222.308756 | -0.3556338 | 0.54089897 | -0.6574865 | 0.51086813 | 0.85359566 |
| Klhl18 | 347.012733 | 0.32535521 | 0.47961626 | 0.67836567 | 0.49753987 | 0.84827947 |
| Actn4 | 8214.93514 | -0.2396156 | 0.23944158 | -1.0007267 | 0.31695896 | 0.74186126 |
| Usp46 | 694.804585 | 0.205331 | 0.33073819 | 0.62082641 | 0.53471385 | 0.8652846 |
| Nsd3 | 1331.0985 | -0.1969271 | 0.20897857 | -0.9423315 | 0.34602296 | 0.7602916 |
| Elp6 | 848.857459 | -0.0970413 | 0.22042484 | -0.4402467 | 0.65975846 | 0.91280998 |
| Atrnl1 | 78.8623628 | 0.19128736 | 0.63705939 | 0.30026614 | 0.76397416 | 0.94300309 |
| Smim10l2a | 196.798596 | 0.89154625 | 0.56103706 | 1.58910404 | 0.11203691 | 0.50987629 |
| Rnd1 | 44.1913667 | -0.4531783 | 0.98068707 | -0.4621028 | 0.64400758 | 0.90823885 |
| Pcnx3 | 497.397505 | 0.07207222 | 0.36428619 | 0.19784505 | 0.8431663 | 0.96765585 |
| Zfp667 | 137.41199 | 0.69376643 | 0.52269718 | 1.32728176 | 0.1844155 | 0.61658486 |
| Dmac2l | 371.367443 | -0.1446613 | 0.48878054 | -0.2959637 | 0.76725778 | 0.94401697 |
| Stfa3 | 33.9568151 | -5.0802512 | 1.12679912 | -4.5085687 | 6.526645009 | 0.00036333 |
| Klhl5 | 262.755855 | -0.439802 | 0.4297263 | -1.0234468 | 0.30609663 | 0.73554152 |
| Zkscan4 | 15.8916396 | 1.14686912 | 1.70150002 | 0.67403415 | 0.5002896 | 0.84938937 |
| Afp | 251.134451 | 1.58067196 | 0.69115401 | 2.28700397 | 0.02219559 | 0.21429774 |
| Miga1 | 343.285535 | -0.1306607 | 0.42410546 | -0.3080853 | 0.75801739 | 0.94129085 |
| 9130008F23I | 28.5420826 | -0.8875394 | 1.10587011 | -0.8025711 | 0.42222266 | 0.80821275 |
| Zfp647 | 61.9826511 | -1.2698003 | 1.16447298 | -1.0904507 | 0.27551467 | 0.71181528 |
| Agap1 | 35.9892517 | -0.4987114 | 1.34123657 | -0.3718295 | 0.71001977 | 0.93077435 |
| Ep300 | 683.853649 | -0.0780784 | 0.36324631 | -0.2149461 | 0.82980931 | 0.96464393 |
| Commd5 | 1090.9044 | 0.43890428 | 0.41460389 | 1.05861108 | 0.28977694 | 0.72395811 |
| Pdlim1 | 126.232281 | 0.27407668 | 0.38555639 | 0.71086016 | 0.4771709 | 0.83683779 |
| Nfic | 649.419921 | 0.29737104 | 0.33940895 | 0.87614379 | 0.38095187 | 0.7832891 |
| Ddx17 | 807.333846 | -0.3391496 | 0.24107113 | -1.4068444 | 0.15947354 | 0.5886662 |
| Smyd3 | 587.047815 | 0.39637596 | 0.28348346 | 1.39823309 | 0.16204308 | 0.59315429 |
| Rab39 | 378.546293 | 0.40847365 | 0.3514912 | 1.16211627 | 0.24518824 | 0.68039867 |
| Gm8430 | 14.0715668 | 1.36637668 | 1.266601 | 1.07877435 | 0.28068833 | 0.7155865 |
| Arntl | 145.520164 | 0.38576136 | 0.49469476 | 0.77979673 | 0.43551053 | 0.81649473 |
| Cgrrf1 | 1161.35881 | 0.61189497 | 0.31783461 | 1.9251993 | 0.05420443 | 0.35994983 |
| Sugct | 40.4900258 | -0.2195583 | 0.60567343 | -0.3625028 | 0.71697635 | 0.93256478 |
| Klf2 | 900.096514 | 0.36586475 | 0.44999435 | 0.81304298 | 0.41619342 | 0.80440685 |
| Zfp78 | 32.8225998 | 0.81474347 | 1.10865087 | 0.73489634 | 0.46240263 | 0.83108016 |
| C1ra | 81.3892881 | 1.10016336 | 0.81334297 | 1.35264384 | 0.17616944 | 0.60763832 |
| Fam72a | 468.643653 | -0.6066474 | 0.33947709 | -1.7870055 | 0.07393661 | 0.41892844 |

| | | | | | | |
|---|---|---|---|---|---|---|
| Sertad3 | 479.908143 | 0.5517354 | 0.52909249 | 1.04279575 | 0.2970429 | 0.72954201 |
| Ankrd17 | 1186.79812 | -0.2771945 | 0.34308751 | -0.8079411 | 0.4191245 | 0.80529433 |
| Zfp935 | 6.73496682 | -0.8617119 | 2.80942063 | -0.3067223 | 0.75905476 | 0.94184564 |
| Wdr86 | 18.4841317 | -3.3957829 | 1.4350656 | -2.366291 | 0.01796731 | 0.18931452 |
| Kcmf1 | 403.927824 | -0.3843048 | 0.28169015 | -1.3642819 | 0.17247886 | 0.60492813 |
| Zfp101 | 321.525645 | 0.524733 | 0.39752178 | 1.3200107 | 0.18683144 | 0.61862792 |
| Tmem245 | 467.837447 | 0.90448504 | 0.57534161 | 1.57208348 | 0.11593119 | 0.52081758 |
| Mrfap1 | 14032.0715 | -0.1791648 | 0.17172684 | -1.0433128 | 0.29680342 | 0.72954201 |
| Zfp93 | 76.3136265 | -0.2809452 | 1.12467682 | -0.2498009 | 0.80274136 | 0.95672466 |
| Pgbd1 | 14.76899 | 0.91310791 | 2.17261117 | 0.42028133 | 0.67427995 | 0.91822108 |
| Sec23ip | 2601.06301 | -0.3768851 | 0.20495892 | -1.8388324 | 0.06593984 | 0.39560669 |
| Tns1 | 13.5565397 | 1.39006426 | 1.8367805 | 0.75679389 | 0.44917335 | 0.82484234 |
| Snupn | 724.861656 | 0.00042616 | 0.20999113 | 0.00202941 | 0.99838076 | 0.99967651 |
| Stam2 | 2100.87035 | 0.23783947 | 0.24052587 | 0.98883117 | 0.32274575 | 0.74549637 |
| Fbxo6 | 540.659905 | 0.97404335 | 0.39373399 | 2.4738615 | 0.01336615 | 0.15631751 |
| Maf | 3591.20235 | 0.32951167 | 0.22712382 | 1.45080192 | 0.14683503 | 0.57365079 |
| Srsf11 | 5672.16387 | -0.1446348 | 0.13984143 | -1.0342773 | 0.30100656 | 0.73263296 |
| Cd47 | 4506.70746 | -0.0307301 | 0.25001668 | -0.1229123 | 0.90217657 | 0.98043702 |
| Zfp458 | 39.4991212 | -1.1083137 | 0.86269791 | -1.2847066 | 0.19889484 | 0.63148061 |
| Soga1 | 71.4393848 | 0.81556069 | 0.63661104 | 1.28109731 | 0.20015949 | 0.63253708 |
| Pprc1 | 1233.97006 | 0.0856072 | 0.34127613 | 0.25084438 | 0.80193443 | 0.95632784 |
| Cpsf6 | 5146.16994 | -0.053019 | 0.20519209 | -0.2583871 | 0.79610814 | 0.9537323 |
| Zcchc24 | 108.463188 | 1.96251014 | 1.26358924 | 1.5531235 | 0.12039363 | 0.52825513 |
| Lair1 | 2047.29624 | 0.20879839 | 0.37833087 | 0.55189359 | 0.58102126 | 0.88314588 |
| Timd4 | 598.10452 | 4.04168763 | 0.46200453 | 8.74815578 | 2.168677546 | 1.473857356 |
| Kxd1 | 2293.91219 | -0.2827312 | 0.20816802 | -1.3581874 | 0.17440423 | 0.60656845 |
| Hba-x | 525.484117 | -0.6321983 | 0.79439968 | -0.7958189 | 0.42613729 | 0.81021713 |
| Cdca7 | 6392.92839 | -0.7013225 | 0.22545068 | -3.1107582 | 0.00186608 | 0.03810972 |
| B4galnt4 | 9.63978795 | 0.01165283 | 2.11828239 | 0.00550108 | 0.9956108 | 0.99884675 |
| Zfp580 | 30.3610653 | -0.0256858 | 0.80047532 | -0.0320882 | 0.97440171 | 0.99446549 |
| Klhl25 | 813.946936 | 0.15223266 | 0.26805177 | 0.5679226 | 0.57008753 | 0.88006507 |
| Gpc3 | 578.560228 | -0.4430548 | 0.57567985 | -0.7696201 | 0.44152531 | 0.81967485 |
| Mettl4 | 823.183441 | -0.2246547 | 0.22668358 | -0.9910497 | 0.32166133 | 0.74447686 |
| Zzef1 | 615.316098 | -0.1191577 | 0.3547063 | -0.3359333 | 0.73692117 | 0.93800224 |
| Cope | 7666.19969 | 0.15385167 | 0.28007353 | 0.54932601 | 0.58278174 | 0.88343755 |
| Tmem191c | 37.903732 | 1.42289619 | 0.88954864 | 1.59957098 | 0.10969379 | 0.50605049 |
| Klhl26 | 218.688083 | 0.34833903 | 0.52359854 | 0.66527885 | 0.50587215 | 0.85074333 |
| Ubl7 | 1363.35022 | -0.2729089 | 0.22504824 | -1.2126683 | 0.22525665 | 0.65875319 |
| Rras2 | 254.931684 | -0.0348659 | 0.42137707 | -0.0827428 | 0.93405609 | 0.98756396 |
| Nap1l3 | 11.8688216 | 0.25616185 | 2.15260109 | 0.11900108 | 0.9052745 | 0.9814201 |
| Ghr | 93.324213 | -0.5882912 | 0.66253189 | -0.887944 | 0.37457089 | 0.77797919 |
| Rtl6 | 168.402461 | -0.1833974 | 0.59227312 | -0.3096501 | 0.75682704 | 0.94107188 |
| Gemin6 | 2217.09356 | -0.1965655 | 0.29112574 | -0.675191 | 0.49955445 | 0.84873011 |
| Eef1d | 13717.1145 | -0.1922672 | 0.23703919 | -0.8111197 | 0.41729693 | 0.8049685 |
| Abcd2 | 2450.68602 | -0.2019863 | 0.18787357 | -1.0751181 | 0.28232187 | 0.71787592 |
| Fmnl1 | 951.284648 | -0.905749 | 0.32140156 | -2.8181226 | 0.00483054 | 0.07744685 |

| | | | | | | |
|---|---|---|---|---|---|---|
| Mta3 | 632.813064 | -0.2922272 | 0.28595554 | -1.0219322 | 0.30681299 | 0.73595594 |
| Zfp1 | 199.44071 | 0.05377328 | 0.42518855 | 0.12646926 | 0.89936048 | 0.98020997 |
| Elob | 3585.49271 | 0.03259757 | 0.23828525 | 0.13680061 | 0.8911884 | 0.97889088 |
| Rnf181 | 4667.91636 | 0.22312287 | 0.14299136 | 1.56039409 | 0.11866678 | 0.52515149 |
| Izumo4 | 62.7736341 | 1.00225699 | 0.64380178 | 1.55677885 | 0.119523 | 0.52689185 |
| Per2 | 43.347635 | 0.76405037 | 1.30600232 | 0.58502987 | 0.55852763 | 0.87327051 |
| Fancm | 257.42874 | -0.4048716 | 0.45604528 | -0.8877881 | 0.37465474 | 0.77799209 |
| Ppp4r1l-ps | 141.528308 | 0.2544146 | 0.43313366 | 0.58738128 | 0.55694767 | 0.87276537 |
| Tmem69 | 610.39414 | -0.4240094 | 0.24687133 | -1.7175322 | 0.08588198 | 0.44801003 |
| Tmem150a | 71.5437977 | 1.034892 | 0.75263009 | 1.37503406 | 0.16912089 | 0.60316496 |
| Zfp277 | 2448.7539 | 0.33404967 | 0.31864484 | 1.04834485 | 0.29447975 | 0.72738877 |
| Aasdh | 265.007793 | -0.3354172 | 0.48153923 | -0.6965521 | 0.48608311 | 0.84183502 |
| Fto | 1285.47392 | 0.04237607 | 0.23434264 | 0.18082954 | 0.85650138 | 0.96924897 |
| Emc7 | 3923.12138 | 0.04729627 | 0.30479586 | 0.15517361 | 0.87668445 | 0.9745677 |
| Triqk | 72.4205987 | -0.2504959 | 0.55490264 | -0.4514232 | 0.65168455 | 0.91036394 |
| Zkscan5 | 330.345992 | 0.64761085 | 0.45901353 | 1.4108753 | 0.15828139 | 0.5871574 |
| Nod2 | 18.2781286 | 1.89459455 | 1.06056532 | 1.78640062 | 0.07403441 | 0.41913963 |
| A430033K04I | 161.751084 | -0.787319 | 0.5038952 | -1.5624658 | 0.1181783 | 0.52512718 |
| Zfp709 | 258.507348 | 0.28575197 | 0.32607152 | 0.87634753 | 0.38084114 | 0.7832891 |
| Mia3 | 1386.46687 | -0.0732068 | 0.21721059 | -0.3370315 | 0.73609312 | 0.9377627 |
| S100a8 | 1392.72769 | -5.8455545 | 1.20987074 | -4.8315529 | 1.354721934 | 9.416087173 |
| Sag | 89.4189395 | -0.3179401 | 0.55266953 | -0.5752807 | 0.56510148 | 0.87793833 |
| Otulinl | 7866.12326 | 0.16804548 | 0.18691811 | 0.89903267 | 0.36863526 | 0.77456461 |
| S100a9 | 402.346111 | -5.1707223 | 1.97570639 | -2.6171512 | 0.00886671 | 0.1194564 |
| Eif3b | 2835.45003 | 0.02122929 | 0.45083577 | 0.04708874 | 0.9624425 | 0.99401755 |
| St3gal5 | 1488.11273 | 0.33357958 | 0.40317895 | 0.8273735 | 0.40802539 | 0.80066966 |
| H2-T22 | 458.355231 | 0.63096746 | 0.31908292 | 1.97744039 | 0.04799187 | 0.33778401 |
| Fez2 | 4554.75624 | 0.27451149 | 0.22440418 | 1.22329041 | 0.22122003 | 0.65469944 |
| B4galt6 | 160.75245 | 0.84443301 | 0.4863669 | 1.73620576 | 0.08252746 | 0.44183004 |
| Ticam2 | 2640.98149 | 0.10408577 | 0.21357861 | 0.48734176 | 0.62601616 | 0.90169029 |
| Pgm3 | 1630.33695 | -0.0060733 | 0.19157201 | -0.0317024 | 0.97470939 | 0.9944856 |
| Trim34a | 413.058825 | 0.80434365 | 0.36917741 | 2.17874559 | 0.02935057 | 0.25573046 |
| Socs6 | 1247.77708 | -0.1948391 | 0.33002482 | -0.5903771 | 0.55493788 | 0.87189251 |
| Cnot10 | 2771.36974 | -0.0772107 | 0.19861379 | -0.3887478 | 0.69746272 | 0.92667877 |
| Snx32 | 105.7654 | 0.47929951 | 0.43056204 | 1.113195 | 0.26562469 | 0.70163359 |
| Cfl1 | 66271.0183 | -0.1636495 | 0.14827084 | -1.1037201 | 0.26971457 | 0.70681628 |
| Pgpep1 | 821.546294 | 0.37975349 | 0.2185448 | 1.73764599 | 0.08227321 | 0.44161834 |
| Npm3 | 98.5733316 | -0.5762898 | 0.51568538 | -1.1175221 | 0.26377115 | 0.70008189 |
| R3hdm1 | 806.085785 | -0.5877506 | 0.39401264 | -1.491705 | 0.13577649 | 0.55758574 |
| Pard6g | 67.4929701 | 0.1436495 | 0.63507415 | 0.22619328 | 0.8210511 | 0.96205421 |
| Cebpg | 450.59667 | -0.2308706 | 0.27119525 | -0.8513076 | 0.39459853 | 0.79263117 |
| Pla2g4a | 8942.22555 | 0.06173778 | 0.24503286 | 0.25195713 | 0.8010742 | 0.95605275 |
| Cars2 | 1483.67304 | -0.1682715 | 0.28630447 | -0.587736 | 0.55670948 | 0.87276537 |
| Ncoa4 | 10751.4121 | 0.31600251 | 0.24242311 | 1.30351645 | 0.1923985 | 0.62396896 |
| Lrif1 | 407.040285 | 0.29058134 | 0.23971603 | 1.21218987 | 0.22543971 | 0.65900564 |
| Cep70 | 552.964056 | -0.0444334 | 0.34846854 | -0.1275106 | 0.89853628 | 0.97989497 |

| | | | | | | |
|---|---|---|---|---|---|---|
| Dennd1b | 200.29767 | -0.3296811 | 0.4075173 | -0.8089991 | 0.41851568 | 0.8052381 |
| Ms4a4b | 14.2371794 | 0.79241588 | 1.9193993 | 0.41284577 | 0.67971961 | 0.92051478 |
| Zfp981 | 16.5169469 | -0.0642935 | 0.94632109 | -0.0679404 | 0.94583305 | 0.99019648 |
| Usp39 | 4898.5636 | -0.2180585 | 0.14616862 | -1.4918287 | 0.13574405 | 0.55758574 |
| Tyw1 | 2173.39638 | -0.2112894 | 0.22452454 | -0.9410526 | 0.34667792 | 0.76056509 |
| Tcim | 110.462729 | 1.29640775 | 0.57929775 | 2.23789537 | 0.02522788 | 0.23122295 |
| Usp34 | 625.596355 | -0.6563649 | 0.5586271 | -1.1749606 | 0.24001047 | 0.67563574 |
| Actr3b | 28.3010183 | -0.6792276 | 0.84046275 | -0.8081591 | 0.41899901 | 0.80529433 |
| AI987944 | 585.267311 | 0.58946997 | 0.29496331 | 1.99845184 | 0.0456677 | 0.32978331 |
| Lig1 | 6621.26729 | -0.0559917 | 0.18192173 | -0.3077788 | 0.75825064 | 0.94131116 |
| Adap1 | 485.36139 | -0.6656063 | 0.25972405 | -2.5627442 | 0.01038485 | 0.13164548 |
| Tgoln1 | 2477.17477 | 0.12819728 | 0.29189985 | 0.43918242 | 0.66052936 | 0.91302324 |
| Mok | 25.3957949 | 1.06436704 | 0.85334433 | 1.24728905 | 0.21229153 | 0.64580271 |
| Zbtb25 | 426.172493 | 0.10582948 | 0.33572658 | 0.31522522 | 0.75259066 | 0.94018119 |
| Med12l | 128.867484 | 0.13201943 | 0.42840459 | 0.30816529 | 0.75795656 | 0.94129085 |
| Chn1 | 14.6747894 | 0.32890738 | 1.76809589 | 0.1860235 | 0.85242633 | 0.96798445 |
| Foxk1 | 149.777454 | -0.6129663 | 0.67348661 | -0.9101388 | 0.36274932 | 0.76990376 |
| Tmem154 | 240.25642 | -1.8495604 | 0.37140085 | -4.9799572 | 6.359833876 | 4.955404319 |
| Cebpb | 215.207197 | -0.497273 | 0.67165091 | -0.7403742 | 0.45907295 | 0.82987212 |
| Rab31 | 9401.34794 | 0.59773695 | 0.15234257 | 3.9236371 | 8.722209245 | 0.00320417 |
| Ptafr | 4811.99089 | 0.4119497 | 0.2893995 | 1.42346378 | 0.15460175 | 0.58343244 |
| Ccdc18 | 159.968152 | 0.36155623 | 0.63092447 | 0.57305787 | 0.56660551 | 0.87883863 |
| Pign | 885.662425 | 0.10464181 | 0.3127278 | 0.33460988 | 0.73791939 | 0.93800224 |
| Rlim | 6546.37684 | 0.0099879 | 0.25228051 | 0.03959044 | 0.96841965 | 0.99443087 |
| Zfp658 | 420.904443 | -0.1396408 | 0.36812535 | -0.3793295 | 0.70444322 | 0.9285501 |
| Drc3 | 23.022563 | 0.06316042 | 1.58109699 | 0.03994721 | 0.96813521 | 0.99443087 |
| Fry | 183.641168 | -0.1505599 | 0.39655997 | -0.3796648 | 0.70419427 | 0.9285501 |
| Chd9 | 1854.76615 | -0.0251558 | 0.20052382 | -0.1254506 | 0.90016684 | 0.98043702 |
| Ppp1r14b | 367.308651 | -0.1790274 | 0.30191784 | -0.5929673 | 0.55320304 | 0.87136821 |
| Fkbp2 | 2373.08236 | 0.08942896 | 0.18458169 | 0.48449528 | 0.62803442 | 0.9024721 |
| Retsat | 1268.63804 | 0.04574229 | 0.27207885 | 0.16812146 | 0.86648773 | 0.97060021 |
| Prelid2 | 715.377121 | 0.53057718 | 0.31780074 | 1.66952782 | 0.09501282 | 0.47206881 |
| Kdm5d | 116.255466 | 0.57067976 | 0.66828345 | 0.85394867 | 0.39313343 | 0.79137753 |
| Ilrun | 747.953561 | -0.1063889 | 0.29577134 | -0.3596998 | 0.71907165 | 0.93310883 |
| Elmod3 | 396.030548 | 0.15881038 | 0.29003154 | 0.54756246 | 0.58399237 | 0.88404713 |
| Ier5 | 504.102856 | -0.0375129 | 0.50836027 | -0.0737919 | 0.94117598 | 0.98935054 |
| Nbeal2 | 253.812464 | -0.6644901 | 0.60616403 | -1.0962216 | 0.27298182 | 0.71005689 |
| Capg | 13111.2475 | 0.30170093 | 0.21429881 | 1.40785163 | 0.15917501 | 0.58838861 |
| Nfil3 | 1997.04863 | -1.0823517 | 0.1885488 | -5.7404327 | 9.443496367 | 1.375265369 |
| Hmga2 | 138.288882 | -1.0406092 | 0.54058626 | -1.9249641 | 0.05423384 | 0.35994983 |
| Cspp1 | 410.833851 | 0.33080592 | 0.32683257 | 1.01215715 | 0.31146292 | 0.74017637 |
| Setd3 | 3040.10916 | -0.2876796 | 0.22447037 | -1.2815928 | 0.19998553 | 0.63247751 |
| Tsnax | 4946.04412 | -0.2333076 | 0.14813684 | -1.5749463 | 0.11526884 | 0.51917664 |
| Ttc26 | 52.5278909 | 1.3404604 | 0.9069978 | 1.47790921 | 0.13943208 | 0.56329174 |
| Pcbp2 | 3920.71338 | -0.2259791 | 0.17763612 | -1.2721459 | 0.20332127 | 0.63809363 |
| Gulp1 | 45.1226588 | 1.10046487 | 0.99245631 | 1.10882953 | 0.26750373 | 0.70441538 |

| | | | | | | |
|---|---|---|---|---|---|---|
| Gadl1 | 10.4763544 | 0.47906182 | 1.15690041 | 0.4140908 | 0.67880761 | 0.92012773 |
| Glipr1 | 3013.97452 | -1.5593235 | 0.33909295 | -4.5985135 | 4.255162004 | 0.00025436 |
| Hist3h2ba | 15.7773711 | 1.47791151 | 1.26780401 | 1.16572554 | 0.24372543 | 0.67897148 |
| Immp2l | 171.045428 | 0.05546581 | 0.59952324 | 0.09251653 | 0.92628765 | 0.98624627 |
| Gm5620 | 49.4045241 | 0.04086509 | 0.54735203 | 0.07465962 | 0.94048554 | 0.98935054 |
| Sipa1 | 4987.42404 | -0.1237874 | 0.26203647 | -0.4724051 | 0.6366377 | 0.90542378 |
| Cep162 | 301.639978 | 0.50043717 | 0.37997271 | 1.31703452 | 0.18782705 | 0.61935434 |
| Acbd4 | 103.298813 | 0.3995843 | 0.46640194 | 0.85673807 | 0.39158966 | 0.79018736 |
| Commd7 | 4631.41412 | 0.36055929 | 0.32696904 | 1.10273218 | 0.27014349 | 0.70727945 |
| Tatdn2 | 39.9233378 | 0.39937912 | 0.96925142 | 0.41204904 | 0.68030348 | 0.92061965 |
| Jmjd6 | 2926.85842 | -0.2568236 | 0.205884 | -1.2474188 | 0.21224398 | 0.64580271 |
| Ide | 4065.52713 | -0.0532645 | 0.19150063 | -0.2781429 | 0.78090264 | 0.94986398 |
| Gm7536 | 5635.36627 | 0.05621102 | 0.36614995 | 0.15351912 | 0.87798891 | 0.97498533 |
| Ero1lb | 444.60231 | -0.6141598 | 0.26420884 | -2.3245239 | 0.02009743 | 0.20036827 |
| Zfp607b | 50.4229939 | 0.49582785 | 0.96151942 | 0.51567117 | 0.60608409 | 0.89258167 |
| Zfp180 | 787.875689 | -0.2412305 | 0.28947799 | -0.8333295 | 0.40465895 | 0.79917613 |
| Nat8f1 | 79.5939197 | 1.26604246 | 0.81138095 | 1.56035517 | 0.11867597 | 0.52515149 |
| Cntrl | 1109.78841 | -0.132642 | 0.29371725 | -0.4515975 | 0.65155901 | 0.91036394 |
| Npm1 | 54097.3584 | -0.2916331 | 0.19252021 | -1.5148181 | 0.12981848 | 0.54647951 |
| Txnl4a | 3619.14293 | -0.09499 | 0.28621114 | -0.3318879 | 0.73997393 | 0.93800224 |
| Rpgrip1 | 4158.79181 | -0.4831011 | 0.29514105 | -1.6368483 | 0.10166218 | 0.48717118 |
| Chd6 | 195.338541 | -0.320097 | 0.60561346 | -0.52855 | 0.59711763 | 0.88891958 |
| Ado | 3745.80678 | 0.1755822 | 0.26236247 | 0.66923519 | 0.50334546 | 0.85012672 |
| Scimp | 105.444929 | 0.8725589 | 0.54721144 | 1.59455528 | 0.11081172 | 0.50865282 |
| Tmem140 | 483.292692 | 2.12866918 | 0.53596869 | 3.97162971 | 7.138261599 | 0.00272425 |
| Trim12c | 163.758016 | 0.53517092 | 0.56777531 | 0.94257518 | 0.34589824 | 0.7602916 |
| Dph6 | 1179.58874 | -0.1814708 | 0.23603602 | -0.7688266 | 0.44199626 | 0.81985447 |
| Homez | 288.138428 | 0.05741796 | 0.57375632 | 0.10007377 | 0.92028576 | 0.9839023 |
| Ccdc189 | 37.9897878 | 0.64292949 | 0.86294652 | 0.74503979 | 0.45624767 | 0.82894467 |
| Gsk3a | 34.5882176 | -0.9928185 | 0.73872021 | -1.3439709 | 0.1789578 | 0.60992118 |
| 5730455P16I | 820.504118 | 0.3702662 | 0.27984351 | 1.32311874 | 0.18579589 | 0.61790851 |
| AB124611 | 1068.99987 | 0.2972915 | 0.26467242 | 1.12324321 | 0.26133421 | 0.69710018 |
| Slc44a2 | 3428.64306 | -0.1414834 | 0.23162812 | -0.6108213 | 0.54131789 | 0.86776854 |
| Armc7 | 763.314207 | 0.4193584 | 0.24934589 | 1.68183401 | 0.09260103 | 0.46597694 |
| Dmac2 | 812.213292 | -0.2995592 | 0.22452182 | -1.3342097 | 0.18213514 | 0.61344341 |
| Aak1 | 353.758091 | 0.38778396 | 0.87824467 | 0.44154433 | 0.65881897 | 0.91280998 |
| Mettl15 | 338.743702 | 0.23517492 | 0.25499757 | 0.92226339 | 0.35639121 | 0.76553707 |
| Rbbp4 | 8704.80391 | 0.21978391 | 0.29270085 | 0.7508824 | 0.45272343 | 0.82610332 |
| Gm6139 | 15.3854054 | 0.19578241 | 0.90737835 | 0.21576712 | 0.82916928 | 0.96462215 |
| Snrpg | 6349.28379 | 0.32984625 | 0.45159904 | 0.73039627 | 0.465148 | 0.83339036 |
| St6galnac2 | 54.6118285 | 0.11919149 | 0.86343874 | 0.13804279 | 0.8902066 | 0.97871887 |
| Arhgap24 | 334.409729 | -0.6587475 | 0.47630945 | -1.3830242 | 0.16665744 | 0.59997659 |
| Rpl38 | 4923.45572 | 0.11059019 | 0.36259567 | 0.30499589 | 0.76036928 | 0.94215089 |
| Bcl2 | 53.5114115 | -0.8500112 | 0.94657303 | -0.897988 | 0.36919196 | 0.77480275 |
| Cep170 | 719.435511 | 0.20790784 | 0.28973461 | 0.71758028 | 0.47301612 | 0.83483892 |
| Sphk2 | 1018.41225 | -0.0679185 | 0.27429048 | -0.2476152 | 0.80443213 | 0.9570372 |

| | | | | | | |
|---|---|---|---|---|---|---|
| Uxs1 | 706.987033 | 0.32992158 | 0.25470557 | 1.29530572 | 0.19521479 | 0.62707955 |
| Birc2 | 504.806002 | 0.17132874 | 0.35522393 | 0.48231193 | 0.62958437 | 0.90310807 |
| Yipf1 | 2023.69849 | 0.1602788 | 0.29502735 | 0.54326759 | 0.58694557 | 0.88615705 |
| Mrpl18 | 6727.17892 | -0.0867262 | 0.25158994 | -0.3447123 | 0.73031064 | 0.93653187 |
| Zfp759 | 59.2202699 | 1.39921027 | 1.10179576 | 1.26993616 | 0.20410737 | 0.63846817 |
| Nsd2 | 1682.40654 | -0.1155332 | 0.32385512 | -0.3567436 | 0.72128374 | 0.93398714 |
| Zfp53 | 805.037176 | 0.6348039 | 0.39121334 | 1.62265402 | 0.10466339 | 0.49501875 |
| Fam173a | 1685.44368 | 0.37334569 | 0.26609198 | 1.40307005 | 0.16059595 | 0.5908482 |
| Las1l | 3586.96444 | -0.4967202 | 0.25614847 | -1.9391887 | 0.05247836 | 0.35389626 |
| Mpp7 | 143.075188 | -0.1361018 | 0.41037983 | -0.3316484 | 0.74015474 | 0.93800224 |
| E2f6 | 258.211481 | 0.1939713 | 0.30978768 | 0.62614271 | 0.53122134 | 0.86415301 |
| Fam136a | 7614.09205 | 0.00902583 | 0.32987809 | 0.02736112 | 0.97817171 | 0.99483019 |
| Bloc1s2 | 1861.46795 | 0.39811285 | 0.42920573 | 0.92755716 | 0.35363733 | 0.76391408 |
| Spop | 5115.10768 | -0.2145463 | 0.15921044 | -1.3475646 | 0.17779848 | 0.60906934 |
| Ece1 | 1049.72497 | -0.4238636 | 0.25185338 | -1.6829774 | 0.09237946 | 0.46597694 |
| Dtnbp1 | 517.602189 | 0.37164707 | 0.25020134 | 1.48539199 | 0.13744002 | 0.55999057 |
| Pus7 | 2227.17926 | -0.521317 | 0.21605138 | -2.4129306 | 0.01582483 | 0.17485494 |
| Zfp317 | 932.517351 | 0.511862 | 0.28175207 | 1.81671073 | 0.06926142 | 0.40636687 |
| Lgals8 | 2392.15397 | 0.46571611 | 0.21287071 | 2.18778859 | 0.02868501 | 0.25226719 |
| Eif1a | 12064.3591 | 0.03115413 | 0.15089715 | 0.20645937 | 0.8364321 | 0.96683245 |
| Zbtb8os | 1143.59439 | -0.2425792 | 0.19016152 | -1.2756483 | 0.20207986 | 0.63663222 |
| Arl16 | 845.212781 | 0.2021579 | 0.30252645 | 0.66823215 | 0.50398542 | 0.85012672 |
| Trim30d | 252.344842 | -0.2216266 | 0.45875936 | -0.4830999 | 0.62902483 | 0.90276005 |
| Gnai1 | 18.9645971 | 1.45462534 | 1.92658717 | 0.755027 | 0.45023278 | 0.82494795 |
| Prdm2 | 296.456929 | -0.0666653 | 0.41732032 | -0.1597461 | 0.87308109 | 0.97344599 |
| Brd9 | 1134.84459 | -0.3555793 | 0.26655104 | -1.3340007 | 0.18220363 | 0.61344341 |
| Gapdh | 101916.939 | -0.2995398 | 0.16568304 | -1.8079087 | 0.07062072 | 0.41021047 |
| Bloc1s3 | 128.373108 | -0.8390861 | 0.43317595 | -1.937056 | 0.05273849 | 0.35506328 |
| Pkn1 | 2466.53286 | 0.17147818 | 0.18017313 | 0.95174111 | 0.34122829 | 0.75846782 |
| Zfp746 | 137.283753 | -0.0351717 | 0.49630118 | -0.0708676 | 0.94350314 | 0.98989307 |
| Mex3b | 34.300626 | 1.10345662 | 0.68705838 | 1.60605946 | 0.10826085 | 0.50301538 |
| Prtn3 | 4004.14768 | -3.3717839 | 0.34129742 | -9.8793125 | 5.118286776 | 4.816307856 |
| Sptan1 | 568.274191 | -0.0451085 | 0.2847045 | -0.1584397 | 0.87411032 | 0.97350615 |
| Ankrd29 | 13.5737247 | 2.38392758 | 1.75507357 | 1.35830635 | 0.1743665 | 0.60656845 |
| Cyb5d2 | 208.987974 | -0.5902513 | 0.32752361 | -1.802164 | 0.07151961 | 0.41122734 |
| Ddx49 | 2308.71519 | -0.3349555 | 0.16642609 | -2.0126382 | 0.04415271 | 0.32226735 |
| Bak1 | 35.6124056 | -0.1236969 | 0.79973679 | -0.1546721 | 0.87707986 | 0.97458574 |
| Zfp119a | 136.659556 | -0.5856348 | 0.48176207 | -1.21561 | 0.22413353 | 0.65793926 |
| Xlr3a | 41.5325058 | 0.13503676 | 1.04001133 | 0.12984162 | 0.89669173 | 0.97984623 |
| Rpl32 | 24446.9265 | 0.43886832 | 0.44869912 | 0.97809045 | 0.32802959 | 0.74843736 |
| Zfp595 | 120.711845 | -0.0934112 | 0.48788284 | -0.1914624 | 0.8481633 | 0.96765585 |
| Fam204a | 587.222221 | -0.1501859 | 0.34772761 | -0.4319066 | 0.66580926 | 0.91520501 |
| Rpl36 | 6050.76135 | 0.45355414 | 0.48148912 | 0.94198211 | 0.3462018 | 0.7602916 |
| Zfp329 | 87.3825067 | -0.8580776 | 0.94369425 | -0.909275 | 0.36320496 | 0.77000799 |
| Zfp105 | 399.161881 | 0.54036482 | 0.39338096 | 1.37364254 | 0.16955268 | 0.60382474 |
| Cacnb2 | 14.9079959 | -1.3995311 | 1.17774048 | -1.1883187 | 0.23470786 | 0.67038085 |

| | | | | | | |
|---|---|---|---|---|---|---|
| Unc13d | 427.627596 | -0.9811737 | 0.37510302 | -2.6157446 | 0.00890331 | 0.1197646 |
| Itpk1 | 840.495128 | -0.2174931 | 0.26198903 | -0.8301611 | 0.40644771 | 0.79992362 |
| Zfp809 | 366.479004 | -0.2141357 | 0.31463672 | -0.6805808 | 0.49613679 | 0.84777781 |
| E030024N20 | 17.6096593 | -0.0160759 | 0.91339055 | -0.0176003 | 0.98595772 | 0.99572532 |
| Mdn1 | 376.048006 | -0.02144 | 0.39157033 | -0.054754 | 0.95633446 | 0.99215259 |
| Sept11 | 3088.31514 | -0.2108034 | 0.22272349 | -0.9464801 | 0.34390375 | 0.75962035 |
| Adtrp | 14.8200938 | 1.14331213 | 1.5405386 | 0.74215091 | 0.45799591 | 0.82986925 |
| 4933430I17R | 13.5318502 | 3.02943061 | 1.76798464 | 1.71349374 | 0.08662174 | 0.44976392 |
| Palld | 602.613612 | 0.95658987 | 0.39722608 | 2.40817487 | 0.0160325 | 0.17615771 |
| Eml1 | 23.676104 | 2.55493439 | 1.20918531 | 2.11293867 | 0.03460602 | 0.27887708 |
| Sdhc | 4726.91239 | 0.35143633 | 0.21886653 | 1.60571071 | 0.10833749 | 0.50301538 |
| Zfp729b | 416.030776 | 0.17578394 | 0.47938308 | 0.36668783 | 0.71385188 | 0.93235217 |
| Nfam1 | 7521.83116 | -0.2393988 | 0.22840189 | -1.0481473 | 0.29457075 | 0.72738877 |
| Tpm3-rs7 | 59.2496517 | 0.31709234 | 0.59487494 | 0.53304033 | 0.59400566 | 0.88797037 |
| Gstm1 | 450.492457 | 0.12076983 | 0.40883531 | 0.29539971 | 0.76768856 | 0.94401697 |
| Gm5431 | 225.030404 | 0.95865203 | 0.4248682 | 2.25635159 | 0.02404862 | 0.22439877 |
| Smco4 | 17.1016112 | 0.87283892 | 1.06499647 | 0.81956978 | 0.41246141 | 0.80251302 |
| Gm5148 | 1903.10594 | 0.19991441 | 0.40312314 | 0.49591401 | 0.61995508 | 0.89896588 |
| Zfp980 | 22.0352005 | 0.07689535 | 0.89302269 | 0.08610683 | 0.93138149 | 0.98730569 |
| Zfp846 | 188.183806 | -0.0089588 | 0.5344865 | -0.0167615 | 0.98662685 | 0.99611549 |
| Arhgap35 | 71.4737471 | 0.91986777 | 0.53771335 | 1.71070287 | 0.08713598 | 0.45109601 |
| Usf2 | 462.189123 | -0.1457197 | 0.26935764 | -0.5409897 | 0.5885147 | 0.88690478 |
| Cryzl1 | 1747.96614 | 0.85521183 | 0.35502216 | 2.40889706 | 0.01600081 | 0.17615771 |
| Tspan7 | 121.849174 | 1.32174925 | 0.44343442 | 2.98070962 | 0.00287581 | 0.0525072 |
| Idi1 | 8352.3337 | 0.55908047 | 0.29079251 | 1.92260961 | 0.0545291 | 0.36033644 |
| Mrps14 | 4242.31123 | -0.0560813 | 0.21939786 | -0.2556145 | 0.79824856 | 0.95435639 |
| Espl1 | 513.704493 | -0.0971064 | 0.42457199 | -0.228716 | 0.81908964 | 0.96105733 |
| Zfp68 | 2001.49724 | 0.48273895 | 0.35041188 | 1.37763293 | 0.16831665 | 0.60296969 |
| Mcm9 | 263.972019 | 0.30045058 | 0.28994599 | 1.03622947 | 0.3000951 | 0.73145694 |
| Upf1 | 204.01599 | -0.6016615 | 0.44556892 | -1.350322 | 0.17691271 | 0.60784615 |
| Ube2e2 | 56.2348579 | 0.60659576 | 0.63722533 | 0.95193291 | 0.34113101 | 0.75846782 |
| Phf21a | 165.523261 | -0.0924756 | 0.4043363 | -0.2287097 | 0.81909456 | 0.96105733 |
| Dock1 | 627.65327 | 0.54991846 | 0.39588574 | 1.38908378 | 0.16480728 | 0.59718229 |
| Zfp85 | 143.83797 | 0.9207175 | 0.7684364 | 1.19817008 | 0.23085081 | 0.6653874 |
| Smim4 | 620.370421 | 0.24815935 | 0.41522164 | 0.59765516 | 0.55007005 | 0.87093412 |
| Abce1 | 9841.08099 | -0.2649168 | 0.2045519 | -1.295108 | 0.19528296 | 0.62707955 |
| Hist1h2bg | 43.2294238 | 0.04559843 | 0.71319252 | 0.06393565 | 0.94902146 | 0.99028544 |
| Phtf1 | 150.152412 | -0.6070189 | 0.58868973 | -1.0311356 | 0.30247724 | 0.73372823 |
| Rrp1b | 1988.38884 | -0.8694164 | 0.21076974 | -4.1249585 | 3.708016357 | 0.00156956 |
| Zfp420 | 141.17402 | 0.77488912 | 0.56407762 | 1.37372783 | 0.16952619 | 0.60382474 |
| Txndc9 | 3405.50042 | 0.30589957 | 0.28677269 | 1.06669702 | 0.28610867 | 0.72202931 |
| Cxcl2 | 815.696763 | 1.79734854 | 0.75453307 | 2.3820673 | 0.01721575 | 0.18456841 |
| Nrf1 | 3224.86458 | -0.3293117 | 0.28265782 | -1.1650544 | 0.24399696 | 0.67911981 |
| Rpl10-ps3 | 34545.3795 | -0.0139132 | 0.26219451 | -0.0530643 | 0.95768067 | 0.99242502 |
| Map2k5 | 1022.66777 | -0.130648 | 0.19588311 | -0.6669694 | 0.50479167 | 0.85034006 |
| Znrf2 | 391.284653 | -1.1926788 | 0.46749503 | -2.5512117 | 0.01073491 | 0.13400016 |

| | | | | | | |
|---|---|---|---|---|---|---|
| Gm26920 | 14.3898957 | 1.00240789 | 1.66401261 | 0.60240402 | 0.54690523 | 0.86994632 |
| Dhcr7 | 1858.19217 | 0.39902147 | 0.21766594 | 1.83318285 | 0.06677538 | 0.39767663 |
| Gm8369 | 9.59770782 | 0.36579536 | 1.61467733 | 0.22654394 | 0.82077839 | 0.96205421 |
| Wdr91 | 705.711983 | 0.52123656 | 0.21365701 | 2.43959496 | 0.01470374 | 0.16670141 |
| Scp2-ps2 | 10.3851967 | -0.1516556 | 1.21113278 | -0.125218 | 0.90035097 | 0.98043702 |
| Fam133b | 418.394276 | 0.17393121 | 0.28561521 | 0.60897042 | 0.54254405 | 0.86903891 |
| Gm15590 | 71.8409936 | 0.52568002 | 0.5803843 | 0.90574473 | 0.36507098 | 0.77118172 |
| Rpl23a | 13638.0551 | -0.0702738 | 0.23817914 | -0.2950458 | 0.76795889 | 0.94401697 |
| Rpl5 | 3520.33216 | 0.03809952 | 0.35992032 | 0.10585543 | 0.91569706 | 0.98369452 |
| Tmed9 | 1548.37136 | -0.2154282 | 0.22032498 | -0.9777747 | 0.32818577 | 0.74843736 |
| Serhl | 1231.37373 | 0.18350726 | 0.4304307 | 0.42633404 | 0.66986445 | 0.91689368 |
| Tmod3 | 6318.20245 | 0.16120211 | 0.16106448 | 1.00085452 | 0.31689715 | 0.74186126 |
| Fbh1 | 3648.75127 | 0.18385184 | 0.22471247 | 0.81816481 | 0.41326309 | 0.80359996 |
| Rpl30 | 9221.86674 | -0.1422772 | 0.23333502 | -0.6097552 | 0.54202401 | 0.86853454 |
| Rpl28-ps1 | 2618.00028 | -0.2214892 | 0.2346379 | -0.9439617 | 0.34518924 | 0.76016201 |
| Gda | 25.8722551 | -5.2322468 | 1.61843982 | -3.2328955 | 0.00122542 | 0.0271397 |
| Gm17383 | 29.0585211 | 0.17449932 | 0.71839061 | 0.24290312 | 0.80808044 | 0.95809696 |
| Zfp110 | 2099.26253 | 0.23888055 | 0.21626677 | 1.1045643 | 0.26934843 | 0.70619525 |
| Eif4b | 8056.80282 | -0.4921689 | 0.2162015 | -2.2764361 | 0.02281992 | 0.2163471 |
| Tubb2a | 13796.0001 | 0.20673703 | 0.18268455 | 1.13166125 | 0.25777688 | 0.69309071 |
| Ccser2 | 361.686307 | 0.13935857 | 0.35024221 | 0.39789198 | 0.69070981 | 0.92470267 |
| Memo1 | 3158.61385 | 0.25892249 | 0.32507271 | 0.79650637 | 0.42573777 | 0.81021713 |
| 0610030E20l | 1071.48385 | -0.0599775 | 0.24124577 | -0.2486156 | 0.80365817 | 0.95680296 |
| Egln2 | 1920.17031 | 0.19356499 | 0.15612086 | 1.23984068 | 0.21503433 | 0.64854028 |
| Fcer1g | 18781.6417 | 0.24825585 | 0.16112346 | 1.54078026 | 0.12337028 | 0.5338481 |
| Cd300c | 14.1605187 | -1.3027837 | 1.50571499 | -0.8652259 | 0.38691479 | 0.78911204 |
| Lin9 | 674.038982 | -0.4404059 | 0.36428545 | -1.2089582 | 0.22667889 | 0.66015092 |
| Kcnj14 | 33.7090266 | 2.91711281 | 1.59063269 | 1.83393239 | 0.06666403 | 0.39741769 |
| Zfp958 | 355.52188 | 1.22677861 | 0.52283313 | 2.34640563 | 0.01895547 | 0.19469539 |
| Osm | 295.576766 | -0.533127 | 0.57561209 | -0.9261915 | 0.35434649 | 0.76419569 |
| Thra | 52.2714174 | 0.13495885 | 0.55310936 | 0.2440003 | 0.80723059 | 0.95787117 |
| Rnf169 | 357.053388 | 0.53163318 | 0.34084303 | 1.55975958 | 0.11881671 | 0.52515149 |
| Hist1h1b | 157.659551 | 0.2909428 | 0.50019389 | 0.58166004 | 0.5607957 | 0.87469256 |
| Cds2 | 821.049948 | 0.19782644 | 0.30925551 | 0.63968607 | 0.52237671 | 0.85816354 |
| Nfe2 | 387.667225 | -2.5900289 | 0.40871402 | -6.33702 | 2.342516476 | 4.761813280 |
| Nap1l1 | 13703.6809 | -0.3167007 | 0.18955806 | -1.6707318 | 0.09477466 | 0.47148369 |
| Hspd1-ps3 | 10.5531757 | -0.8776468 | 1.13778709 | -0.7713629 | 0.44049184 | 0.81963526 |
| Pirb | 558.719645 | -0.5169538 | 0.34658095 | -1.4915818 | 0.13580881 | 0.55758574 |
| Rex1bd | 796.251339 | -0.0948849 | 0.24343925 | -0.3897682 | 0.69670799 | 0.92646244 |
| Abi1 | 4287.11025 | 0.06225694 | 0.1743371 | 0.35710667 | 0.72101195 | 0.93398714 |
| Rps27a-ps2 | 30.4516515 | 1.17978654 | 0.93724942 | 1.25877542 | 0.20811146 | 0.6418276 |
| Zfp516 | 144.487794 | -0.3296334 | 0.65840954 | -0.500651 | 0.61661676 | 0.89878056 |
| Zfp708 | 26.4006915 | -1.3089404 | 1.38236268 | -0.9468864 | 0.34369668 | 0.75947281 |
| Deaf1 | 564.095325 | 0.1898617 | 0.36978853 | 0.51343318 | 0.60764834 | 0.89332558 |
| Rsl1 | 244.613234 | -0.2504282 | 0.55380705 | -0.4521939 | 0.65112931 | 0.91021183 |
| C1qtnf3 | 26.4807611 | 0.54518477 | 1.99453426 | 0.27333939 | 0.78459234 | 0.95094799 |

| | | | | | | |
|---|---|---|---|---|---|---|
| Gm10053 | 6993.68104 | -0.1503531 | 0.24003549 | -0.6263786 | 0.53106666 | 0.86413122 |
| Hdhd5 | 509.883213 | 0.0311195 | 0.2783323 | 0.111807 | 0.91097644 | 0.98288717 |
| Vwa8 | 382.479113 | -0.3137275 | 0.3761622 | -0.8340218 | 0.40426872 | 0.79917613 |
| Hnrnpa3 | 3453.20699 | -0.5231847 | 0.18204916 | -2.8738652 | 0.00405482 | 0.06785584 |
| Sh2d3c | 1279.94747 | -0.5803371 | 0.21001049 | -2.7633722 | 0.00572075 | 0.08847273 |
| Kcp | 7.31330843 | 0.50396074 | 2.7825063 | 0.18111755 | 0.85627531 | 0.96924897 |
| Eno1b | 233.129475 | -0.1017257 | 0.3948415 | -0.2576368 | 0.79668724 | 0.95380086 |
| Rad51b | 304.471294 | 0.66374575 | 0.5208939 | 1.27424366 | 0.20257708 | 0.63759103 |
| Rpl18 | 18545.2091 | -0.145506 | 0.28436763 | -0.5116827 | 0.60887308 | 0.89437372 |
| Fcgr4 | 1315.07158 | 0.67399728 | 0.40421299 | 1.66743102 | 0.09542873 | 0.47377421 |
| Ifitm6 | 174.618943 | -1.9093046 | 0.53738205 | -3.5529743 | 0.0003809 | 0.01080469 |
| Nap1l4 | 8965.44978 | -0.0077985 | 0.15272364 | -0.051063 | 0.95927528 | 0.99271349 |
| Zfp945 | 224.291087 | 0.98043512 | 0.40377593 | 2.42816631 | 0.01517538 | 0.17040121 |
| Mfsd4a | 14.6894091 | -1.062644 | 1.15833933 | -0.9173857 | 0.35894055 | 0.76764331 |
| Skap2 | 7551.8175 | 0.30200519 | 0.29081757 | 1.03846956 | 0.2990515 | 0.73093388 |
| Mtfmt | 652.14741 | 0.12090433 | 0.25221142 | 0.47937692 | 0.63167051 | 0.90399255 |
| Gm12715 | 34.4075685 | -0.5833397 | 0.81986135 | -0.7115102 | 0.47676814 | 0.83666308 |
| Hnrnpm | 13126.2423 | -0.3178816 | 0.2412466 | -1.3176627 | 0.18761659 | 0.61935434 |
| Sept9 | 9735.3686 | 0.06840251 | 0.26600372 | 0.25714869 | 0.79706399 | 0.95397994 |
| Usp47 | 1005.27949 | -0.1631598 | 0.36030058 | -0.4528436 | 0.65066134 | 0.91021183 |
| Zc3h4 | 478.804837 | -0.3341012 | 0.25266038 | -1.3223332 | 0.18605723 | 0.61790851 |
| Naa38 | 1575.74402 | 0.36409589 | 0.42114768 | 0.86453257 | 0.3872954 | 0.78911204 |
| Cdyl | 288.382786 | -0.2737439 | 0.38190769 | -0.7167802 | 0.47350975 | 0.83483892 |
| Rpl11 | 12638.2154 | -0.0084444 | 0.32669812 | -0.0258478 | 0.97937873 | 0.99483019 |
| Vpreb1 | 14.3472944 | -1.9676838 | 1.39946038 | -1.4060304 | 0.1597151 | 0.58902465 |
| Slc27a4 | 229.686334 | 0.84395575 | 0.33057124 | 2.55302234 | 0.01067926 | 0.13344172 |
| Tonsl | 769.901895 | -0.2768623 | 0.34804296 | -0.7954832 | 0.42633247 | 0.81021713 |
| Csf2ra | 1540.15554 | -0.9326586 | 0.30822592 | -3.0258927 | 0.002479 | 0.04672671 |
| Slc14a1 | 149.491733 | 0.90117792 | 0.5026259 | 1.79293967 | 0.07298257 | 0.41525386 |
| Wdr83os | 1403.48093 | -0.0496444 | 0.19779756 | -0.2509859 | 0.80182499 | 0.95629571 |
| Fxn | 718.199013 | 0.35318392 | 0.37615604 | 0.93892927 | 0.34776706 | 0.76101142 |
| Nkapl | 16.7282579 | 0.24046445 | 1.15753013 | 0.20773926 | 0.83543256 | 0.96641606 |
| Ppp2r5d | 4292.30901 | -0.0606372 | 0.30834423 | -0.1966541 | 0.84409824 | 0.96765585 |
| Fxyd2 | 46.7301305 | 2.93627532 | 0.6916848 | 4.24510601 | 2.184900965 | 0.00097352 |
| Gm8116 | 156.892327 | -0.07713 | 0.45578796 | -0.1692234 | 0.86562091 | 0.9705375 |
| Zfp933 | 54.2313225 | 0.92017184 | 0.61096474 | 1.50609647 | 0.13204241 | 0.55148116 |
| Max | 229.828227 | -0.0839728 | 0.36382638 | -0.2308045 | 0.8174667 | 0.96079676 |
| Bcas3 | 527.33259 | -0.2113888 | 0.39168496 | -0.5396909 | 0.58941025 | 0.88722166 |
| Hadhb | 9179.98198 | 0.286852 | 0.23912255 | 1.19960249 | 0.23029376 | 0.66491035 |
| Ptk2b | 1010.35592 | -1.1164803 | 0.34080906 | -3.2759702 | 0.001053 | 0.02425859 |
| Gm7331 | 14.7280637 | 0.86021172 | 1.01506833 | 0.84744218 | 0.3967487 | 0.79486192 |
| Mbtd1 | 371.284696 | -0.1871526 | 0.30262407 | -0.6184326 | 0.53629022 | 0.86549318 |
| Zfp426 | 598.28918 | 0.44382558 | 0.2469662 | 1.7971106 | 0.07231806 | 0.41349974 |
| B3gnt8 | 169.992656 | -0.3304981 | 0.56650019 | -0.5834034 | 0.5596218 | 0.87419914 |
| 2610301B20I | 516.63547 | 0.11603986 | 0.35422376 | 0.32758916 | 0.74322232 | 0.93856621 |
| Kbtbd2 | 1575.65915 | 0.14906416 | 0.28283044 | 0.52704425 | 0.59816286 | 0.88951331 |

| | | | | | | |
|---|---|---|---|---|---|---|
| Arhgef12 | 16.0933519 | 3.80037393 | 1.6234241 | 2.34096188 | 0.01923413 | 0.19585467 |
| Fcgr3 | 29393.2249 | 0.88458996 | 0.2680438 | 3.30016941 | 0.00096626 | 0.02277518 |
| Znhit1 | 820.580179 | 0.32184403 | 0.31729548 | 1.01433536 | 0.31042276 | 0.7399039 |
| Uqcr10 | 5272.8349 | 0.65216512 | 0.45754419 | 1.42535984 | 0.15405321 | 0.58303286 |
| Tcea2 | 112.246891 | 0.01548383 | 0.44411829 | 0.03486421 | 0.97218802 | 0.99443087 |
| Trp53 | 9283.28405 | -0.265034 | 0.19268788 | -1.3754576 | 0.16898961 | 0.60312169 |
| Ccdc28a | 84.0565595 | 0.25484268 | 0.44606101 | 0.57131799 | 0.56778411 | 0.87929304 |
| Tor4a | 2067.18289 | 0.19075524 | 0.23145359 | 0.82416197 | 0.40984753 | 0.80116089 |
| Nsmce2 | 3052.24826 | 0.47559561 | 0.32877247 | 1.44657981 | 0.14801464 | 0.57481369 |
| Calcrl | 680.820877 | 1.05900832 | 0.44829693 | 2.36229215 | 0.01816232 | 0.19071219 |
| Stfa2l1 | 13.2426005 | -6.1511261 | 1.7924748 | -3.4316388 | 0.00059995 | 0.01561519 |
| Taf1b | 2085.31811 | 0.04519276 | 0.35941121 | 0.12574111 | 0.89993686 | 0.98043702 |
| Zfp637 | 686.960505 | 0.07235578 | 0.35622074 | 0.20312062 | 0.83904075 | 0.96757028 |
| Akap17b | 41.2070094 | -0.4138386 | 1.24359808 | -0.3327752 | 0.73930398 | 0.93800224 |
| Rcan3 | 221.811115 | 0.69453783 | 0.49759038 | 1.39580236 | 0.16277402 | 0.59421503 |
| Flot1 | 2653.41037 | -0.1722065 | 0.18199995 | -0.9461901 | 0.34405165 | 0.75962035 |
| Ndufs8 | 7094.64523 | 0.3263029 | 0.35171435 | 0.92774972 | 0.35353741 | 0.76383312 |
| Fdps | 8920.26342 | 0.22289566 | 0.21641224 | 1.02995864 | 0.30302942 | 0.73428908 |
| Rps3a3 | 9.79602104 | -0.0989195 | 1.13501433 | -0.0871526 | 0.93055018 | 0.98716777 |
| Slx1b | 390.571266 | -0.269747 | 0.36369135 | -0.7416921 | 0.4582739 | 0.82986925 |
| Rps26-ps1 | 15.2252253 | 0.26068745 | 0.93022501 | 0.28024128 | 0.7792924 | 0.9488468 |
| Rpl13-ps6 | 23.2835276 | 0.34493965 | 0.89055717 | 0.38733015 | 0.69851181 | 0.92669304 |
| Nrm | 2503.95078 | -0.3456674 | 0.1879032 | -1.8396038 | 0.06582642 | 0.39558985 |
| Eif4a1 | 46498.748 | -0.076414 | 0.17485268 | -0.4370191 | 0.66209751 | 0.91395288 |
| Rgs3 | 612.927103 | 0.68346352 | 0.41050906 | 1.66491702 | 0.09592931 | 0.47482666 |
| Atl2 | 1628.79446 | -0.0548048 | 0.17399527 | -0.3149789 | 0.75277765 | 0.94018119 |
| Nkapd1 | 520.275479 | -0.0273831 | 0.25722293 | -0.1064567 | 0.91521999 | 0.98347559 |
| Dbp | 22.6987936 | 0.47647792 | 1.23438868 | 0.38600315 | 0.69949434 | 0.92669304 |
| Sclt1 | 235.699226 | -0.0517775 | 0.39086333 | -0.1324696 | 0.89461291 | 0.97941957 |
| Zfp874b | 114.344285 | 0.59733436 | 0.64177663 | 0.93075119 | 0.35198228 | 0.76348958 |
| Zfp341 | 27.2128393 | -1.2814444 | 1.21728265 | -1.052709 | 0.29247438 | 0.72584701 |
| Kmt5c | 1448.07205 | 0.09365136 | 0.22299671 | 0.41996747 | 0.67450922 | 0.91832667 |
| Tnip2 | 907.891067 | -0.2492462 | 0.24424933 | -1.0204581 | 0.30751124 | 0.73673815 |
| Zfp422 | 3181.44618 | -0.1178703 | 0.16491626 | -0.714728 | 0.47477716 | 0.83578199 |
| Irak4 | 1562.13165 | 0.0813465 | 0.19828884 | 0.41024244 | 0.68162811 | 0.92061965 |
| Ube4a | 1346.23003 | -0.0409462 | 0.30592561 | -0.1338437 | 0.89352618 | 0.97941957 |
| Ptp4a3 | 2016.74701 | -0.1747073 | 0.19583326 | -0.8921228 | 0.37232714 | 0.77738145 |
| Zfp930 | 888.156809 | -0.3954416 | 0.49277712 | -0.8024756 | 0.42227792 | 0.80821275 |
| Tmem40 | 62.9077656 | 0.41677133 | 0.63925891 | 0.65196013 | 0.51442689 | 0.85525743 |
| 4930453N24 | 1042.20791 | -0.2782045 | 0.24011313 | -1.1586391 | 0.24660334 | 0.68166056 |
| Grb2 | 766.288278 | -0.1986229 | 0.23189435 | -0.8565233 | 0.39170841 | 0.79019938 |
| 9430015G10 | 861.743775 | -0.2079176 | 0.32116233 | -0.6473911 | 0.51737888 | 0.85694744 |
| Serpinb12 | 46.7817803 | -0.5785303 | 0.62779198 | -0.9215319 | 0.35677279 | 0.76595323 |
| Hspa2 | 149.249169 | 0.01480851 | 0.46105637 | 0.03211865 | 0.97437743 | 0.99446549 |
| Zfp74 | 190.242825 | 0.48791296 | 0.68942766 | 0.70770725 | 0.47912707 | 0.83869135 |
| Taok2 | 430.153628 | 0.33303492 | 0.32947514 | 1.0108044 | 0.31211006 | 0.74050472 |

| | | | | | | |
|---|---|---|---|---|---|---|
| Fcrl1 | 440.418432 | -0.0743724 | 0.26190132 | -0.2839712 | 0.77643243 | 0.94800857 |
| Atxn7l3 | 201.947951 | -0.5875979 | 0.50316675 | -1.1677995 | 0.24288765 | 0.67814218 |
| Chpt1 | 90.5558058 | 0.76792698 | 0.71996366 | 1.06661908 | 0.28614388 | 0.72202931 |
| Kif13b | 202.775417 | 0.61749352 | 0.55894056 | 1.10475704 | 0.26926488 | 0.70619525 |
| H2afj | 1733.14942 | 0.14775855 | 0.32919957 | 0.44884186 | 0.65354574 | 0.91076467 |
| Ctf2 | 102.893466 | 0.24735663 | 0.80064016 | 0.30894856 | 0.75736066 | 0.94116141 |
| Rpl3 | 68605.0117 | 0.08288853 | 0.31153376 | 0.26606597 | 0.79018841 | 0.95261385 |
| Dhps | 1771.90396 | -0.102667 | 0.30273238 | -0.3391344 | 0.7345085 | 0.93750666 |
| Alox5ap | 5486.89501 | 0.25063044 | 0.25902001 | 0.96761036 | 0.333239 | 0.75254065 |
| Psma3 | 19655.5551 | 0.41091892 | 0.41406512 | 0.99240167 | 0.32100162 | 0.74413737 |
| Rp2 | 2536.55587 | 0.47791259 | 0.20445068 | 2.33754463 | 0.01941088 | 0.19738432 |
| Amd-ps3 | 54.2991004 | 1.05191224 | 3.72884109 | 0.28210165 | 0.77786557 | 0.94837927 |
| Prmt7 | 6999.84922 | 0.12688496 | 0.25125539 | 0.50500393 | 0.61355608 | 0.89769543 |
| Gemin2 | 960.284559 | 0.22963168 | 0.35406896 | 0.64855072 | 0.51662882 | 0.85670602 |
| Tpt1 | 11065.5783 | 0.28108389 | 0.33894846 | 0.82928211 | 0.40694479 | 0.79992362 |
| Atp8b4 | 253.048012 | -2.2946855 | 0.36864242 | -6.2246921 | 4.825025902 | 9.080698749 |
| Serpinb6a | 3417.22392 | 2.19321271 | 0.38352161 | 5.71861573 | 1.073953627 | 1.492917582 |
| Pop5 | 2226.06725 | 0.35633809 | 0.38139383 | 0.9343048 | 0.35014669 | 0.76216093 |
| Zdhhc8 | 33.2098652 | -0.3920758 | 0.7876292 | -0.4977923 | 0.61863042 | 0.89878056 |
| Slc35e3 | 165.93485 | -0.3838943 | 0.39934429 | -0.9613117 | 0.33639547 | 0.75493042 |
| Gm11353 | 16.8952973 | 0.21438913 | 0.89915033 | 0.23843524 | 0.81154353 | 0.95967622 |
| Arrb2 | 1200.77032 | 0.19973293 | 0.22378312 | 0.89252903 | 0.37210944 | 0.77719221 |
| Pyroxd2 | 284.038693 | 1.05979982 | 0.34956311 | 3.03178397 | 0.00243113 | 0.0460372 |
| Casc4 | 187.474327 | 0.00220935 | 0.32656219 | 0.00676547 | 0.99460197 | 0.99882461 |
| Alyref2 | 176.042467 | -0.118409 | 0.32434857 | -0.3650671 | 0.71506132 | 0.93235217 |
| Pwwp2b | 230.013427 | -0.4206659 | 0.46366414 | -0.9072642 | 0.36426709 | 0.77081383 |
| Gtf2i | 2337.50383 | -0.0486094 | 0.22174396 | -0.2192142 | 0.82648322 | 0.96370073 |
| Ap2a1 | 758.539305 | -0.115188 | 0.28587938 | -0.4029252 | 0.68700324 | 0.92426981 |
| Sp7 | 7.13522592 | -0.7938197 | 2.20815618 | -0.3594943 | 0.71922531 | 0.93310883 |
| Ppih | 2616.85244 | 0.33448976 | 0.44152659 | 0.75757558 | 0.44870511 | 0.82417561 |
| 2610008E11I | 211.879959 | 0.4014423 | 0.41943627 | 0.95709964 | 0.33851696 | 0.7564609 |
| Zfp937 | 142.40384 | -0.107622 | 0.63384433 | -0.1697926 | 0.86517328 | 0.9705375 |
| Hnrnpc | 24190.0382 | 0.04885903 | 0.27379131 | 0.17845356 | 0.85836679 | 0.96947659 |
| Bckdha | 1292.33863 | -0.2663533 | 0.24996834 | -1.0655481 | 0.28662799 | 0.72260415 |
| Zfp128 | 42.1025864 | 1.18336746 | 1.37811329 | 0.85868663 | 0.39051343 | 0.7899826 |
| Rps16-ps2 | 3210.86179 | 0.12116833 | 0.38984235 | 0.31081366 | 0.75594228 | 0.94102069 |
| Zfp868 | 837.15756 | 0.44594889 | 0.25931486 | 1.71971978 | 0.08548339 | 0.44669728 |
| Sntb1 | 66.9007163 | 0.93765325 | 0.7291255 | 1.28599707 | 0.19844408 | 0.63122167 |
| Trim5 | 135.445704 | 0.16005297 | 0.427666 | 0.37424759 | 0.70822012 | 0.92967665 |
| Rnf14 | 2035.543 | 0.60299923 | 0.24059124 | 2.50632245 | 0.01219943 | 0.14688547 |
| Adgrg3 | 169.648343 | -4.1444829 | 0.84879492 | -4.8827848 | 1.045980019 | 7.526749166 |
| Wtap | 7167.88225 | 0.11117762 | 0.19613603 | 0.56683935 | 0.57082333 | 0.88035336 |
| Irak2 | 795.898405 | -0.1606303 | 0.43047002 | -0.3731509 | 0.7090361 | 0.93004918 |
| Xcr1 | 33.3158224 | -0.7564843 | 1.11902334 | -0.6760219 | 0.49902672 | 0.84872332 |
| Zfp266 | 1571.52116 | 0.1043376 | 0.32632819 | 0.31973211 | 0.74917142 | 0.9399191 |
| 0610040J01F | 875.039106 | -0.1837199 | 0.2226657 | -0.8250931 | 0.40931873 | 0.80100711 |

| | | | | | | |
|---|---|---|---|---|---|---|
| Tor3a | 2562.3119 | -0.5752893 | 0.19325115 | -2.9769 | 0.00291179 | 0.05299195 |
| Tmem219 | 917.292531 | 0.263306 | 0.34315911 | 0.76730004 | 0.44290314 | 0.82038676 |
| Tnfrsf19 | 7.85784514 | -1.6254207 | 2.27452748 | -0.7146191 | 0.47484447 | 0.83578199 |
| Fam78b | 11.2027659 | -0.414301 | 1.4564012 | -0.284469 | 0.776051 | 0.94776024 |
| Mfap2 | 75.1026743 | 1.27809458 | 0.90777221 | 1.40794636 | 0.15914696 | 0.58838861 |
| Fhit | 26.5627608 | 1.28202287 | 1.33824051 | 0.95799138 | 0.3380671 | 0.75632313 |
| Ifitm2 | 4709.76042 | 0.87789431 | 0.20204443 | 4.34505585 | 1.392402314 | 0.00069808 |
| Layn | 403.876299 | 0.46383997 | 0.35248655 | 1.31590827 | 0.18820483 | 0.61935434 |
| Eno3 | 476.616469 | 0.737396 | 0.36745271 | 2.00677797 | 0.04477331 | 0.3252446 |
| Nr1h2 | 1582.88575 | -0.178395 | 0.19619122 | -0.9092913 | 0.36319636 | 0.77000799 |
| Rpl35a | 11692.5713 | 0.28402536 | 0.4243427 | 0.66933013 | 0.50328491 | 0.85012672 |
| Hist1h4i | 991.883759 | 0.0549496 | 0.29318888 | 0.18742047 | 0.85133097 | 0.9679302 |
| Marf1 | 386.784761 | 0.46507784 | 0.27804782 | 1.67265413 | 0.0943954 | 0.47055378 |
| Atp8b2 | 96.9305778 | 0.23807525 | 0.65884786 | 0.36135088 | 0.71783717 | 0.93285856 |
| Pla2g16 | 532.14205 | 1.35097316 | 0.50150764 | 2.6938237 | 0.00706375 | 0.10256275 |
| Hist1h4c | 22.2112196 | 0.45279432 | 1.01579584 | 0.44575327 | 0.65577548 | 0.91165371 |
| Mrps9 | 2559.23151 | 0.05112275 | 0.25853049 | 0.19774362 | 0.84324566 | 0.96765585 |
| Slc9a6 | 442.967112 | 0.73323998 | 0.31110874 | 2.35686074 | 0.01843016 | 0.19120986 |
| Cd302 | 2849.89896 | 0.58706097 | 0.31906838 | 1.83992214 | 0.06577967 | 0.39558985 |
| Bloc1s4 | 843.61478 | 0.19368575 | 0.22568629 | 0.85820786 | 0.3907777 | 0.7899826 |
| Ipmk | 1294.46103 | 0.22933475 | 0.36047177 | 0.63620723 | 0.52464136 | 0.85973714 |
| Nsa2 | 3274.58173 | -0.1384841 | 0.2459703 | -0.5630115 | 0.57342708 | 0.88046468 |
| H3f3a | 30992.4307 | 0.3347065 | 0.38706923 | 0.86471998 | 0.3871925 | 0.78911204 |
| Tsga10 | 337.357972 | -0.0856202 | 0.30218724 | -0.283335 | 0.77692003 | 0.94800948 |
| Gmfg | 3187.45734 | 0.49918413 | 0.41049324 | 1.21605932 | 0.22396233 | 0.65774578 |
| Intu | 150.816896 | -0.4042676 | 0.38403716 | -1.0526782 | 0.29248849 | 0.72584701 |
| B2m | 30572.7608 | -0.1329323 | 0.17231455 | -0.7714515 | 0.44043934 | 0.81963526 |
| Gstp1 | 7790.98481 | 0.87016596 | 0.22752744 | 3.82444406 | 0.00013107 | 0.00454206 |
| Ube2s | 375.320181 | -0.2921891 | 0.29153414 | -1.0022465 | 0.31622454 | 0.74186126 |
| Zbtb40 | 157.631734 | -0.6879556 | 0.61926962 | -1.1109145 | 0.26660517 | 0.70333859 |
| Arl1 | 9613.15762 | 0.77970527 | 0.37048234 | 2.10456797 | 0.03532893 | 0.28154973 |
| Acyp2 | 30.4605937 | 1.10893923 | 0.9837618 | 1.12724364 | 0.25963949 | 0.69596146 |
| Tmem263 | 1783.83137 | -0.0605694 | 0.23867063 | -0.2537782 | 0.79966693 | 0.95518266 |
| Rpl26 | 15124.8905 | 0.25157814 | 0.4073756 | 0.61755821 | 0.5368666 | 0.86596304 |
| Trmt61a | 917.236871 | -0.2109607 | 0.31067836 | -0.6790326 | 0.49711722 | 0.84814011 |
| Tdrd5 | 26.8970858 | -7.1823101 | 1.56199852 | -4.5981542 | 4.262504378 | 0.00025436 |
| Copz1 | 12395.0454 | -0.3128702 | 0.16800982 | -1.862214 | 0.06257293 | 0.38618662 |
| Rrs1 | 3112.69571 | -0.4192682 | 0.213554 | -1.9632888 | 0.04961263 | 0.34192183 |
| Clasrp | 488.115888 | -0.208691 | 0.35078187 | -0.5949309 | 0.55188966 | 0.87093412 |
| Rrp1 | 8906.99656 | -0.2261038 | 0.17921292 | -1.2616488 | 0.20707519 | 0.64179145 |
| Haghl | 256.018833 | 0.08720443 | 0.35611677 | 0.24487595 | 0.80655248 | 0.95741157 |
| Cdh3 | 22.4787469 | -1.2277842 | 1.94308491 | -0.6318737 | 0.52746942 | 0.86160147 |
| Zfp143 | 1240.59144 | 0.14770221 | 0.21637409 | 0.68262432 | 0.49484426 | 0.84678075 |
| Plac1 | 92.2126325 | 5.51226476 | 3.74840576 | 1.47056245 | NA | NA |
| Sap18b | 5758.9148 | -0.0381125 | 0.23043109 | -0.1653965 | 0.86863187 | 0.9719395 |
| Mcrip1 | 3982.55501 | 0.45180241 | 0.28977137 | 1.55916855 | 0.11895649 | 0.52515149 |

| | | | | | | |
|---|---|---|---|---|---|---|
| Dnajc30 | 692.535034 | -0.0780999 | 0.20710619 | -0.3771006 | 0.70609888 | 0.92896305 |
| Prcp | 2983.40389 | 0.09204981 | 0.26101088 | 0.35266656 | 0.72433843 | 0.93488415 |
| Ppm1b | 156.811009 | -0.5596622 | 0.43585082 | -1.2840682 | 0.1991181 | 0.63169391 |
| Blnk | 388.322982 | -0.9938459 | 0.27265901 | -3.645014 | 0.00026738 | 0.00815667 |
| Prpf40a | 5562.72654 | -0.2558501 | 0.17681192 | -1.4470182 | 0.14789183 | 0.57472133 |
| Maml3 | 140.030342 | -0.514138 | 0.63867689 | -0.8050049 | 0.42081685 | 0.80699993 |
| Rpl15-ps3 | 11390.8895 | 0.24129092 | 0.34964993 | 0.69009285 | 0.4901358 | 0.84448327 |
| Fnip2 | 325.980571 | 0.37111477 | 0.48625078 | 0.76321682 | 0.4453341 | 0.82118815 |
| Sfmbt2 | 172.451007 | -0.729716 | 0.42196838 | -1.7293145 | 0.08375282 | 0.44417933 |
| Stk19 | 669.676458 | 0.18632538 | 0.19853002 | 0.93852498 | 0.34797469 | 0.76101142 |
| H2-K1 | 5054.89768 | 0.2237005 | 0.19497255 | 1.14734353 | 0.25123967 | 0.6844769 |
| Exoc5 | 2075.88766 | -0.0094645 | 0.25261052 | -0.0374669 | 0.97011269 | 0.99443087 |
| Mmgt1 | 971.740831 | 0.07899917 | 0.2777871 | 0.28438745 | 0.77611347 | 0.94776024 |
| Exosc5 | 2126.54186 | -0.429327 | 0.27501807 | -1.5610864 | 0.11850336 | 0.52515149 |
| Taok3 | 3751.26003 | -0.3066461 | 0.30374866 | -1.0095389 | 0.31271625 | 0.74065408 |
| Slc38a10 | 5106.40868 | 0.20813673 | 0.23051127 | 0.90293516 | 0.36656031 | 0.77259343 |
| Ddhd2 | 1137.57589 | 0.24000764 | 0.23874048 | 1.0053077 | 0.31474871 | 0.74087376 |
| Naca | 23033.2995 | 0.26788627 | 0.37566365 | 0.71310139 | 0.47578302 | 0.83639059 |
| Dnaic1 | 14.488798 | 1.09534808 | 1.7461177 | 0.62730484 | 0.53045943 | 0.86356925 |
| Gm17132 | 19.0399986 | -0.1281086 | 1.0766467 | -0.1189885 | 0.90528443 | 0.9814201 |
| Cxcl12 | 93.3490271 | 1.99100225 | 0.98561474 | 2.02006137 | 0.04337702 | 0.31946485 |
| Phf5a | 4934.15843 | -0.2257973 | 0.18494978 | -1.2208576 | 0.22213993 | 0.65543602 |
| Zfp873 | 211.416853 | -0.0427888 | 0.42570302 | -0.1005132 | 0.91993691 | 0.9839023 |
| Fiz1 | 1819.62047 | -0.1182894 | 0.33314381 | -0.3550701 | 0.72253707 | 0.93423486 |
| Acvr2b | 35.7068425 | -0.6332738 | 0.9364748 | -0.6762316 | 0.49889362 | 0.84872332 |
| Zcchc14 | 34.1445722 | 2.46328143 | 1.71866065 | 1.43325644 | 0.15178457 | 0.57988151 |
| Nol4l | 216.290265 | 0.12356805 | 0.36225865 | 0.34110449 | 0.73302492 | 0.93728136 |
| Cracr2a | 154.826448 | -0.2141963 | 0.56216368 | -0.3810213 | 0.70318743 | 0.92804961 |
| Hipk2 | 655.760465 | 0.0113953 | 0.32909463 | 0.0346262 | 0.97237781 | 0.99443087 |
| Stx17 | 879.86222 | 0.46024546 | 0.19116466 | 2.40758649 | 0.01605836 | 0.17618107 |
| Nol10 | 349.570532 | -0.8294176 | 0.35110342 | -2.362317 | 0.0181611 | 0.19071219 |
| Smim20 | 2195.43788 | 0.30292101 | 0.38906814 | 0.77858086 | 0.43622666 | 0.81729305 |
| Mrps36 | 1720.66099 | 0.16024564 | 0.28966223 | 0.55321552 | 0.58011584 | 0.88286755 |
| Rps7 | 21683.5725 | 0.28567269 | 0.40190998 | 0.71078776 | 0.47721576 | 0.83683779 |
| Snrpa | 5971.62984 | -0.3336592 | 0.1786365 | -1.8678112 | 0.06178839 | 0.38368396 |
| Hist1h4d | 218.597398 | -0.2980523 | 0.38060635 | -0.7830986 | 0.43356921 | 0.81483708 |
| Cox5b | 6912.8847 | 0.15238742 | 0.32012603 | 0.47602322 | 0.63405784 | 0.90465549 |
| Tmem236 | 40.7544329 | 2.02354824 | 1.17885875 | 1.71653155 | 0.0860648 | 0.44836429 |
| Cep128 | 831.805583 | 0.50977831 | 0.31489797 | 1.61886818 | 0.10547563 | 0.49702749 |
| Sec22c | 161.476938 | -0.3472691 | 0.69607391 | -0.4988969 | 0.61785203 | 0.89878056 |
| Zfp229 | 47.7104231 | 1.75515452 | 1.25569823 | 1.39775185 | 0.1621876 | 0.59320305 |
| Wdr61 | 5509.53127 | 0.16882832 | 0.3663857 | 0.46079398 | 0.64494642 | 0.90835328 |
| Adgrg5 | 100.637688 | -2.217447 | 1.05590969 | -2.1000347 | 0.03572579 | 0.28378802 |
| Dot1l | 501.787185 | 0.02720327 | 0.41358383 | 0.06577449 | 0.94755736 | 0.99019648 |
| Mdc1 | 1059.29673 | -0.5355998 | 0.25274005 | -2.1191727 | 0.03407588 | 0.27569457 |
| U2af1 | 4064.85249 | -0.317085 | 0.1843431 | -1.7200806 | 0.0854178 | 0.44669728 |

| | | | | | | |
|---|---|---|---|---|---|---|
| Hist1h2ab | 103.751142 | -0.0715344 | 0.57472144 | -0.1244679 | 0.90094483 | 0.98043702 |
| Med9 | 1835.9512 | 0.09091371 | 0.15973066 | 0.56916879 | 0.5692416 | 0.87961466 |
| Cd2ap | 667.336132 | -0.0913089 | 0.3727011 | -0.2449924 | 0.80646234 | 0.95741157 |
| Gdpd1 | 117.243582 | 3.13661526 | 1.08973242 | 2.87833527 | 0.0039978 | 0.06699327 |
| Dlgap4 | 1110.88816 | 0.19006501 | 0.2208797 | 0.86049105 | 0.38951842 | 0.78942657 |
| Tmem91 | 35.4730631 | 0.26168988 | 0.75091641 | 0.34849402 | 0.72746921 | 0.93576253 |
| Ext1 | 169.674503 | 0.43712437 | 0.43324712 | 1.00894928 | 0.31299896 | 0.74065408 |
| Kalrn | 51.7531391 | 1.24358566 | 0.99709952 | 1.24720315 | 0.21232302 | 0.64580271 |
| Bod1l | 409.04789 | -0.0760581 | 0.41800452 | -0.1819552 | 0.85561788 | 0.96915653 |
| Akr1b10 | 2147.8372 | 0.38477989 | 0.23663136 | 1.62607314 | 0.1039341 | 0.49356592 |
| Armt1 | 1075.46342 | -0.3024748 | 0.18289776 | -1.653792 | 0.09816979 | 0.47978865 |
| Mospd2 | 697.699179 | 0.98365301 | 0.24546121 | 4.00736633 | 6.139959048 | 0.00239968 |
| Rps17 | 14000.415 | 0.16877634 | 0.36502863 | 0.46236465 | 0.64381984 | 0.90823885 |
| Ttr | 27.2485984 | 2.38286982 | 1.21049463 | 1.96850921 | 0.04900948 | 0.34122535 |
| Gm6311 | 14.6524007 | 0.64515151 | 0.96366419 | 0.66947752 | 0.50319092 | 0.85012672 |
| Suclg2 | 4627.74454 | -0.2187458 | 0.15426853 | -1.4179545 | 0.15620404 | 0.58525083 |
| Patj | 29.3279586 | -1.6416532 | 1.36324683 | -1.204223 | 0.22850339 | 0.66258779 |
| Ccdc62 | 163.673523 | 0.38660327 | 0.50112408 | 0.77147215 | 0.44042713 | 0.81963526 |
| Ssbp3 | 381.225498 | -0.3039955 | 0.31269364 | -0.9721831 | 0.33095945 | 0.75011761 |
| Zscan20 | 69.6215036 | -0.2100244 | 0.63870689 | -0.3288276 | 0.74228603 | 0.93840202 |
| Rbak | 98.7567018 | 0.36764757 | 0.66652061 | 0.55159221 | 0.58122778 | 0.88314588 |
| Slc25a3 | 23334.1516 | -0.0753256 | 0.13939033 | -0.5403936 | 0.58892565 | 0.88690478 |
| Ppp4r1 | 2233.84608 | 0.13144874 | 0.26180266 | 0.50209093 | 0.61560356 | 0.89876301 |
| Rcc1l | 1957.96041 | -0.1258571 | 0.20739797 | -0.6068385 | 0.54395807 | 0.86947682 |
| Flot2 | 1830.75623 | -0.1043377 | 0.21228916 | -0.4914886 | 0.6230809 | 0.90017327 |
| Rps12 | 782.603203 | 0.0347744 | 0.45102033 | 0.07710162 | 0.93854271 | 0.9890914 |
| Hist1h2af | 13.1508357 | 0.09889498 | 1.07757705 | 0.09177533 | 0.92687655 | 0.98630274 |
| Rpl34 | 11913.7507 | 0.52753255 | 0.47035871 | 1.12155371 | 0.26205224 | 0.69762406 |
| Zfp13 | 41.0306917 | 3.15251567 | 1.29912676 | 2.42664208 | 0.01523928 | 0.17056005 |
| Gmfb | 4168.7906 | 0.3841038 | 0.14946247 | 2.5699013 | 0.01017275 | 0.13017076 |
| Abca14 | 11.5551 | 0.14091576 | 1.10470355 | 0.12755979 | 0.89849735 | 0.97989497 |
| Pgghg | 310.211246 | 0.18643474 | 0.42700857 | 0.43660656 | 0.66239671 | 0.91395288 |
| Gm10108 | 587.072922 | -0.1012898 | 0.34204221 | -0.2961323 | 0.76712904 | 0.94399854 |
| Zfp27 | 181.703637 | 0.60653056 | 0.44661158 | 1.35807173 | 0.17444093 | 0.60656845 |
| Iah1 | 978.571314 | 0.04855309 | 0.39553847 | 0.12275187 | 0.90230358 | 0.98043702 |
| Pgk1 | 42991.2835 | 0.17875964 | 0.27755617 | 0.64404851 | 0.51954398 | 0.8576517 |
| Ccn6 | 12.1755301 | -1.6029149 | 2.06201786 | -0.7773526 | 0.4369508 | 0.81793712 |
| Lmnb2 | 1694.12798 | -0.2566359 | 0.24636564 | -1.0416872 | 0.29755674 | 0.72979999 |
| Qk | 11199.3888 | 0.5681717 | 0.25793557 | 2.202766 | 0.02761125 | 0.24636643 |
| Cd200r4 | 46.9817052 | -1.9699997 | 1.20725775 | -1.6317971 | 0.10272224 | 0.48958378 |
| Btbd3 | 200.027143 | 0.59078969 | 0.45652057 | 1.29411406 | 0.19562603 | 0.62744971 |
| Zfp119b | 264.579265 | 0.78456506 | 0.56361861 | 1.39201412 | 0.16391812 | 0.59572501 |
| Scfd2 | 896.283822 | -0.1508955 | 0.21517312 | -0.7012747 | 0.48313159 | 0.84106286 |
| Rai1 | 191.021071 | -0.3208554 | 0.33942608 | -0.945288 | 0.34451184 | 0.75972469 |
| Zfp954 | 671.7375 | 0.02245331 | 0.35218147 | 0.06375494 | 0.94916536 | 0.99028544 |
| Cttnbp2nl | 1556.56295 | 0.56704049 | 0.30523821 | 1.85769823 | 0.06321188 | 0.38764352 |

| | | | | | | |
|---|---|---|---|---|---|---|
| Cnih4 | 3477.23039 | 0.50773507 | 0.41912444 | 1.21141843 | 0.22573508 | 0.65900564 |
| Fmr1nb | 93.5339975 | 0.38726408 | 0.71242338 | 0.54358699 | 0.58672571 | 0.88599131 |
| Tgif2 | 572.081083 | 0.07261969 | 0.27714275 | 0.26202991 | 0.79329838 | 0.95310033 |
| Lancl2 | 1048.15111 | -0.1544612 | 0.21195035 | -0.7287613 | 0.46614767 | 0.83345004 |
| 2700097O09 | 566.468749 | 0.23597139 | 0.38494538 | 0.61299967 | 0.53987656 | 0.86719392 |
| Btbd9 | 1643.94633 | -0.0347344 | 0.215542 | -0.1611492 | 0.87197592 | 0.97298927 |
| Gspt1 | 2300.5405 | -0.1154258 | 0.34968747 | -0.3300828 | 0.74133739 | 0.93800224 |
| Tnfaip8 | 7173.68503 | 0.35035713 | 0.20109103 | 1.74228126 | 0.08145924 | 0.44011802 |
| Rapgef2 | 237.024861 | 1.08979294 | 0.54790673 | 1.98901179 | 0.0466999 | 0.33444287 |
| Gak | 2922.91254 | -0.0481303 | 0.20423602 | -0.23566 | 0.81369649 | 0.95988497 |
| Cks2 | 4425.16526 | -0.0541084 | 0.33879787 | -0.1597071 | 0.87311179 | 0.97344599 |
| Morf4l1 | 12774.4371 | 0.38613431 | 0.28306312 | 1.36412795 | 0.1725273 | 0.60492813 |
| Nectin2 | 474.861783 | 0.16543025 | 0.27458365 | 0.60247671 | 0.54685686 | 0.86994632 |
| Glrp1 | 24.4063511 | 3.26673232 | 1.69615806 | 1.92595984 | 0.05410939 | 0.35989907 |
| Rpl17 | 13310.7311 | 0.50651646 | 0.47766374 | 1.06040383 | 0.28896092 | 0.72395811 |
| Serpinb2 | 51.4201961 | 0.82907644 | 1.057653 | 0.78388323 | 0.43310863 | 0.81435873 |
| Itgb1bp1 | 2727.66478 | 0.49415287 | 0.41159596 | 1.20057756 | 0.22991511 | 0.66443457 |
| Gm15772 | 12.8823156 | 0.11321937 | 1.13161948 | 0.10005074 | 0.92030404 | 0.9839023 |
| Tmem65 | 70.3242086 | -0.0007832 | 0.70993563 | -0.0011032 | 0.99911981 | 0.99969186 |
| Borcs7 | 963.194577 | 0.41747445 | 0.3115355 | 1.34005419 | 0.18022773 | 0.61122544 |
| Tubb3 | 52.4772758 | 2.72370615 | 1.68598069 | 1.61550258 | 0.1062019 | 0.49777747 |
| Ftl1-ps1 | 2443.20554 | 0.52970808 | 0.16404926 | 3.22895737 | 0.00124242 | 0.02738482 |
| Zfp706 | 15909.5536 | -0.2335457 | 0.16873803 | -1.3840727 | 0.1663362 | 0.59960123 |
| Arf2 | 4467.43746 | 0.38393266 | 0.16492304 | 2.32795043 | 0.01991474 | 0.19988038 |
| Rpl9-ps6 | 179.926483 | 0.49080616 | 0.59027684 | 0.83148471 | 0.40569986 | 0.79982698 |
| Gm8623 | 9.21993516 | -0.2208801 | 1.26530048 | -0.1745673 | 0.86141965 | 0.97007287 |
| Nsl1 | 1482.16746 | -0.0014956 | 0.22348617 | -0.0066922 | 0.99466046 | 0.99882461 |
| Fabp4 | 27.911776 | -1.4127318 | 1.0311768 | -1.370019 | 0.17068096 | 0.60430173 |
| Zfp398 | 275.668164 | 0.45035731 | 0.67224087 | 0.66993443 | 0.50289959 | 0.85012672 |
| Mppe1 | 982.042391 | -0.1287166 | 0.29133514 | -0.4418162 | 0.65862225 | 0.91265586 |
| Tlr12 | 41.9978262 | 2.34996514 | 1.24721089 | 1.88417626 | 0.05954114 | 0.37661155 |
| Timm17a | 4114.77993 | 0.04402668 | 0.25791791 | 0.17070035 | 0.86445939 | 0.9705375 |
| Cnr2 | 208.461561 | -1.0035479 | 0.50438396 | -1.9896507 | 0.04662943 | 0.33435982 |
| Armc9 | 241.3221 | 0.13757061 | 0.49034327 | 0.28055981 | 0.77904805 | 0.94883461 |
| Tubb4a | 31.2887337 | -0.0749203 | 1.10974489 | -0.0675113 | 0.94617471 | 0.99019648 |
| Gm49339 | 1673.18412 | 0.87740029 | 0.23541081 | 3.72710273 | 0.00019369 | 0.00635242 |
| Srpk2 | 1388.22614 | 0.01337882 | 0.18423432 | 0.07261848 | 0.94210972 | 0.98962444 |
| 2310039H08 | 535.463609 | 0.09499304 | 0.36977998 | 0.2568907 | 0.79726315 | 0.95401743 |
| Mysm1 | 997.504139 | -0.0760688 | 0.37752514 | -0.2014934 | 0.84031278 | 0.96765585 |
| Ganc | 316.270354 | -0.1738542 | 0.36847961 | -0.4718149 | 0.63705892 | 0.90565274 |
| Rpl7a | 23492.7848 | -0.0342729 | 0.30322572 | -0.1130277 | 0.91000861 | 0.98262295 |
| Ncs1 | 170.180696 | 0.86628809 | 0.66274885 | 1.30711368 | 0.19117411 | 0.62230783 |
| Atp5g2 | 2429.40131 | 0.01495718 | 0.28856532 | 0.05183292 | 0.95866182 | 0.99249408 |
| Cebpzos | 1773.97917 | 1.13806198 | 0.4986959 | 2.28207606 | 0.02248485 | 0.21507833 |
| Hist1h2bk | 43.8984012 | -0.5496713 | 0.7551676 | -0.7278799 | 0.46668711 | 0.83345004 |
| Ppox | 761.249712 | 0.16156497 | 0.253789 | 0.63661139 | 0.524378 | 0.85953586 |

| | | | | | | |
|---|---|---|---|---|---|---|
| Zfp677 | 66.0434296 | 0.93303958 | 0.6504162 | 1.43452697 | 0.15142194 | 0.57958216 |
| AI413582 | 1882.26919 | 0.64401919 | 0.50030874 | 1.28724354 | 0.19800942 | 0.63077867 |
| Gm16477 | 15.4113869 | -0.2957739 | 0.99376991 | -0.2976282 | 0.76598697 | 0.94347227 |
| Zfp512 | 1260.64816 | -0.1562681 | 0.21402037 | -0.7301551 | 0.46529538 | 0.83339036 |
| Ei24 | 400.740411 | 0.27667911 | 0.34507018 | 0.80180532 | 0.42266558 | 0.80826451 |
| Zfp599 | 22.6038035 | 1.12501592 | 1.55715916 | 0.72247973 | 0.46999959 | 0.83381643 |
| Hikeshi | 2501.39469 | 0.17444979 | 0.26168182 | 0.66664851 | 0.50499667 | 0.85034006 |
| 4833420G17 | 718.620345 | -0.1274948 | 0.38850024 | -0.3281717 | 0.74278187 | 0.93850397 |
| Actg1 | 151489.985 | -0.2926104 | 0.20106178 | -1.4553259 | 0.14557905 | 0.57242843 |
| Zfp28 | 163.655479 | 0.7706499 | 0.41036202 | 1.8779757 | 0.06038449 | 0.37881203 |
| Phactr2 | 1427.08532 | 0.18686195 | 0.28333965 | 0.65949807 | 0.50957598 | 0.85335831 |
| Impdh2 | 9987.64818 | -0.6004286 | 0.14739036 | -4.0737301 | 4.626608921 | 0.00190563 |
| Klhl24 | 587.390104 | 0.84450504 | 0.47275419 | 1.78635125 | 0.0740424 | 0.41913963 |
| Hdac10 | 533.378245 | 1.08805611 | 0.35090137 | 3.10074629 | 0.00193034 | 0.03908648 |
| Acadm | 2811.35876 | 0.3728779 | 0.17984364 | 2.07334494 | 0.03814019 | 0.29511002 |
| Cfl2 | 402.984442 | 0.5282039 | 0.32133723 | 1.64376816 | 0.1002241 | 0.4838364 |
| Zfp938 | 317.840746 | 0.17255708 | 0.50599683 | 0.34102403 | 0.73308549 | 0.93728136 |
| Gm10123 | 28346.7304 | 0.3444473 | 0.36365903 | 0.94717102 | 0.34355164 | 0.75947281 |
| Mtap | 5082.18959 | -0.2446333 | 0.15864417 | -1.542025 | 0.12306751 | 0.5331037 |
| Stat4 | 225.952241 | -0.266657 | 0.42939293 | -0.6210094 | 0.53459344 | 0.8652846 |
| 9130023H24 | 171.658783 | 0.97595808 | 0.5808622 | 1.68018865 | 0.09292062 | 0.46643328 |
| Atp11c | 288.516719 | -0.715931 | 0.33461705 | -2.1395531 | 0.0323909 | 0.26700664 |
| Kdr | 86.9928336 | 2.20019198 | 1.01656033 | 2.16434964 | 0.03043753 | 0.2603792 |
| Ufc1 | 2721.42639 | 0.47065458 | 0.37317655 | 1.26121158 | 0.20723263 | 0.6418276 |
| Cped1 | 120.874882 | -1.3479612 | 0.43639897 | -3.0888277 | 0.00200948 | 0.04023234 |
| Mrpl42 | 7530.63865 | 0.18352237 | 0.30201877 | 0.60765221 | 0.54341817 | 0.86919906 |
| Nrg1 | 172.510195 | -4.6407103 | 0.90894412 | -5.1056058 | 3.297365141 | 2.781839156 |
| Ica1 | 508.864107 | 0.29275649 | 0.31844844 | 0.91932147 | 0.35792745 | 0.76674184 |
| Rpl35 | 12972.5328 | 0.52793782 | 0.45996113 | 1.1477879 | 0.25105613 | 0.6842534 |
| Ccni | 891.943796 | -0.1175887 | 0.26277121 | -0.4474946 | 0.654518 | 0.9108895 |
| Manbal | 874.006735 | 0.20251775 | 0.20910516 | 0.96849713 | 0.33279615 | 0.75195702 |
| Zfp780b | 323.326215 | 0.0905123 | 0.28410954 | 0.3185824 | 0.7500432 | 0.9399191 |
| Ing2 | 108.243968 | 0.70447955 | 0.40683744 | 1.7315996 | 0.08334488 | 0.44341423 |
| Lrrc40 | 3258.57311 | -0.0194184 | 0.19790388 | -0.0981206 | 0.92183654 | 0.98437722 |
| Sox7 | 38.037234 | 3.33040703 | 1.72857403 | 1.92667885 | 0.05401966 | 0.35989907 |
| Mapk3 | 995.309835 | -0.5027165 | 0.25638688 | -1.9607729 | 0.04990551 | 0.34225988 |
| Kif1b | 738.694169 | 0.48677966 | 0.39081205 | 1.24555949 | 0.21292616 | 0.64596443 |
| Zfp26 | 263.357607 | 1.03838014 | 0.41350438 | 2.51117083 | 0.01203314 | 0.14545599 |
| Bbs5 | 225.717338 | -0.2241026 | 0.37756838 | -0.5935418 | 0.55281863 | 0.8711362 |
| Clip2 | 11.6177467 | -0.5117689 | 1.35818278 | -0.3768041 | 0.70631916 | 0.92897562 |
| Numbl | 31.5941629 | -0.3086391 | 0.91240637 | -0.3382694 | 0.7351602 | 0.93750666 |
| Rps4l | 2867.68456 | -0.0040409 | 0.25928869 | -0.0155846 | 0.98756578 | 0.9962376 |
| Hspb11 | 154.457331 | 0.94033754 | 0.500875 | 1.87738963 | 0.06046471 | 0.37912087 |
| Pstk | 829.333383 | 0.89697622 | 0.50873343 | 1.76315563 | 0.07787424 | 0.4314473 |
| Cd300lb | 591.157344 | -1.4391806 | 0.31159677 | -4.6187275 | 3.861004951 | 0.00023382 |
| Nol7 | 1560.10487 | -0.1762718 | 0.22064049 | -0.7989098 | 0.42434271 | 0.80944713 |

| | | | | | | |
|---|---|---|---|---|---|---|
| Ldha | 199149.607 | 0.00867872 | 0.26626194 | 0.03259467 | 0.97399782 | 0.99444457 |
| Gpr84 | 3028.48763 | 0.46047481 | 0.44832329 | 1.02710437 | 0.30437132 | 0.73467909 |
| 1110038F14I | 846.030594 | -0.077904 | 0.26954446 | -0.2890211 | 0.77256523 | 0.94611977 |
| Zfp993 | 637.373528 | 0.42923992 | 0.31029411 | 1.38333245 | 0.16656295 | 0.59997655 |
| Scoc | 2110.97725 | 0.51732958 | 0.17948363 | 2.88232175 | 0.00394756 | 0.06624218 |
| Parp10 | 483.410391 | 0.21219188 | 0.32545099 | 0.65199334 | 0.51440547 | 0.85525743 |
| Naa15 | 4148.41377 | -0.0690411 | 0.28426553 | -0.2428755 | 0.80810186 | 0.95809696 |
| Hacd1 | 1473.15932 | -0.3824035 | 0.21654995 | -1.7658904 | 0.07741423 | 0.43008322 |
| Zfp35 | 1515.0943 | -0.143032 | 0.2763088 | -0.5176528 | 0.60470052 | 0.89258167 |
| Gm8995 | 754.265347 | -0.1869107 | 0.43429571 | -0.4303765 | 0.66692179 | 0.91565144 |
| Rpl27 | 9607.42216 | 0.08030208 | 0.38184154 | 0.21030211 | 0.83343189 | 0.96565375 |
| Usp31 | 208.750246 | -0.04139 | 0.74303536 | -0.055704 | 0.95557763 | 0.99215259 |
| 1190007I07R | 327.441233 | 0.04268014 | 0.43520738 | 0.09806852 | 0.92187789 | 0.98437722 |
| Krr1 | 2599.32274 | -0.5255758 | 0.19920764 | -2.6383313 | 0.00833152 | 0.11464503 |
| Slc39a4 | 50.6057412 | -0.132102 | 0.84073649 | -0.1571266 | 0.87514511 | 0.97370497 |
| Mapk1 | 7744.08496 | -0.1143337 | 0.21765559 | -0.5252963 | 0.59937721 | 0.8902352 |
| Alg11 | 1284.53438 | 0.2228894 | 0.19992226 | 1.11488037 | 0.26490169 | 0.70112124 |
| 3300002I08R | 38.3697455 | 0.89296619 | 0.78593431 | 1.13618426 | 0.25587946 | 0.6905302 |
| Bcl9l | 64.3105546 | -0.0497047 | 0.68274626 | -0.0728011 | 0.94196439 | 0.9896127 |
| Zfp947 | 184.036969 | 0.1647306 | 0.59135409 | 0.27856508 | 0.78057861 | 0.94965869 |
| Tmed5 | 4314.02332 | -0.295447 | 0.22255182 | -1.3275423 | 0.18432935 | 0.61658486 |
| Stk24 | 488.800972 | -0.1803779 | 0.26127111 | -0.690386 | 0.48995146 | 0.84448327 |
| Gm10131 | 2857.28793 | -0.0387496 | 0.37875569 | -0.1023077 | 0.91851248 | 0.98390186 |
| B9d2 | 1113.70797 | 0.04044452 | 0.40833972 | 0.09904626 | 0.92110154 | 0.98391854 |
| Nmral1 | 2499.16293 | 0.12933838 | 0.34117705 | 0.37909462 | 0.7046176 | 0.92863468 |
| Syne2 | 59.4221708 | 0.73170224 | 0.954414 | 0.76665078 | 0.44328917 | 0.82038676 |
| D630045J12I | 23.5437997 | -2.2841351 | 1.58322597 | -1.4427095 | 0.14910234 | 0.57645724 |
| Rps15 | 12493.5675 | 0.03691458 | 0.32755103 | 0.1126987 | 0.91026943 | 0.98275189 |
| Lrmda | 144.566634 | -0.3795398 | 0.5560226 | -0.6825977 | 0.49486108 | 0.84678075 |
| Snu13 | 12045.7005 | -0.2513399 | 0.24535918 | -1.0243752 | 0.3056581 | 0.73532263 |
| Zkscan7 | 24.3998397 | -0.2035529 | 0.86449705 | -0.2354582 | 0.81385315 | 0.95988497 |
| Arhgap22 | 102.581826 | 0.99617736 | 0.58599109 | 1.69998722 | 0.08913333 | 0.4566019 |
| Snrnp70 | 4553.89505 | -0.3388558 | 0.21755208 | -1.5575848 | 0.11933171 | 0.5265707 |
| Eno1 | 137978.281 | -0.1805989 | 0.16102597 | -1.1215513 | 0.26205329 | 0.69762406 |
| Zfp773 | 79.8308451 | -0.755646 | 0.8220196 | -0.9192555 | 0.35796196 | 0.76674184 |
| Nup98 | 2536.08332 | 0.03169799 | 0.22140491 | 0.14316751 | 0.88615789 | 0.97793139 |
| Jazf1 | 90.4050267 | -0.2808363 | 0.71018983 | -0.3954384 | 0.69251939 | 0.92549717 |
| Klhdc3 | 4597.75276 | -0.1898488 | 0.18841814 | -1.007593 | 0.31364987 | 0.74065408 |
| Ccdc102a | 459.35017 | -1.5343476 | 0.34568045 | -4.4386301 | 9.053326353 | 0.00047737 |
| Sox11 | 135.919669 | 1.14703855 | 1.18160514 | 0.97074607 | 0.33167474 | 0.75068746 |
| Slc22a21 | 109.765474 | -0.0672223 | 0.86948964 | -0.0773124 | 0.93837505 | 0.98906962 |
| Zbtb18 | 44.6696742 | 0.05273719 | 0.77499589 | 0.06804835 | 0.94574715 | 0.99019648 |
| Brwd3 | 239.0113 | -0.1726273 | 0.41050339 | -0.420526 | 0.67410126 | 0.91822108 |
| Cycs | 10241.6306 | -0.1940079 | 0.22980195 | -0.8442393 | 0.39853567 | 0.79557554 |
| Gm8730 | 18.0170985 | -0.081701 | 0.94264729 | -0.0866719 | 0.93093233 | 0.98726443 |
| Sfxn4 | 128.488402 | 0.61873138 | 0.58498211 | 1.05769281 | 0.29019551 | 0.72418639 |

| | | | | | | |
|---|---|---|---|---|---|---|
| Rnf217 | 25.8949623 | -2.7484098 | 1.66432349 | -1.6513676 | 0.09866354 | 0.48025164 |
| Utp14a | 1513.16358 | -0.4359187 | 0.18726822 | -2.3277771 | 0.01992394 | 0.19988038 |
| Chchd1 | 2350.81667 | 0.39391554 | 0.41847523 | 0.94131148 | 0.34654526 | 0.76054685 |
| Prpf38a | 2813.61918 | 0.09286957 | 0.28908147 | 0.32125743 | 0.74801531 | 0.9399191 |
| Ap3s2 | 2025.67825 | 0.02701286 | 0.31786082 | 0.0849833 | 0.93227467 | 0.98730569 |
| Hspbp1 | 3177.74741 | -0.0680098 | 0.1892398 | -0.3593843 | 0.71930765 | 0.93311669 |
| Lin28b | 858.440768 | -0.287531 | 0.2202969 | -1.3051976 | 0.19182558 | 0.62286958 |
| Gpatch1 | 499.920196 | -0.3789445 | 0.31616791 | -1.1985545 | 0.23070123 | 0.6653874 |
| Alms1 | 70.7618767 | 1.1432752 | 0.6556494 | 1.74372949 | 0.08120627 | 0.43985341 |
| Ppcdc | 1102.84559 | -0.2134733 | 0.32031982 | -0.6664379 | 0.50513125 | 0.85034006 |
| Gpx1 | 55329.3196 | 0.14627046 | 0.40303152 | 0.3629256 | 0.71666046 | 0.93235217 |
| Chd4 | 1353.92707 | -0.1797509 | 0.3443971 | -0.5219291 | 0.6017197 | 0.89103464 |
| Slc24a3 | 19.5767516 | -0.9033956 | 1.34282078 | -0.6727596 | 0.50110023 | 0.84968141 |
| Rps6-ps1 | 12.5830503 | 0.10951426 | 1.00616003 | 0.10884378 | 0.9133264 | 0.98308155 |
| Uqcrh | 4429.62974 | 0.6033293 | 0.45935998 | 1.31341283 | 0.18904388 | 0.61961804 |
| Ptcd3 | 5779.80169 | 0.01537387 | 0.22510589 | 0.06829617 | 0.94554987 | 0.99019648 |
| Rpl7l1 | 11499.0931 | 0.10491521 | 0.24322926 | 0.43134289 | 0.66621907 | 0.91520501 |
| Crem | 1126.09333 | 0.30638775 | 0.22830417 | 1.34201556 | 0.17959095 | 0.60992118 |
| Zkscan8 | 91.2078577 | 0.11549711 | 0.78615089 | 0.14691468 | 0.88319936 | 0.97651643 |
| CAAA011183 | 113.324283 | 0.44777328 | 0.6381456 | 0.70167887 | 0.48287945 | 0.84106286 |
| Gm7964 | 13.9966401 | 0.62093985 | 0.98662831 | 0.6293554 | 0.52911641 | 0.86278569 |
| Dpp3 | 8435.80006 | -0.0607502 | 0.16717154 | -0.3634005 | 0.71630572 | 0.93235217 |
| Pepd | 8473.71142 | 0.5944758 | 0.14496421 | 4.1008454 | 4.116435122 | 0.00171865 |
| Brpf3 | 97.5134376 | -0.6202573 | 0.49607065 | -1.2503406 | 0.21117514 | 0.6450201 |
| Amd2 | 23.8323413 | -0.7412302 | 0.83479143 | -0.8879226 | 0.37458239 | 0.77797919 |
| Nr6a1 | 65.2818492 | 0.4314307 | 0.77942419 | 0.5535249 | 0.57990403 | 0.88286755 |
| Klk8 | 2953.23715 | -0.4042527 | 0.36784341 | -1.0989804 | 0.27177663 | 0.70940613 |
| Pym1 | 300.455986 | 0.01170976 | 0.28630905 | 0.04089903 | 0.96737639 | 0.99443087 |
| Gpn1 | 3983.90888 | 0.04170522 | 0.24201957 | 0.17232167 | 0.86318465 | 0.9705375 |
| Dzip3 | 520.032608 | 0.10186018 | 0.24535862 | 0.41514816 | 0.67803345 | 0.91965663 |
| Ipcef1 | 69.7195835 | -0.7538812 | 0.90828632 | -0.8300039 | 0.40653656 | 0.79992362 |
| Mtx1 | 1939.07863 | 0.11966143 | 0.18920636 | 0.63243874 | 0.52710022 | 0.86134345 |
| Vrk2 | 3152.32434 | 0.15985942 | 0.2832937 | 0.56428864 | 0.57255771 | 0.88035336 |
| Cnnm2 | 175.340402 | 0.5016881 | 0.53442227 | 0.9387485 | 0.34785989 | 0.76101142 |
| Hcst | 92.0543324 | 0.15408084 | 0.51877234 | 0.29701051 | 0.76645848 | 0.94363877 |
| Mocs1 | 619.546017 | -0.3820177 | 0.31780569 | -1.2020482 | 0.22934487 | 0.66356408 |
| Med14 | 464.356582 | -0.2845908 | 0.40433008 | -0.7038575 | 0.4815215 | 0.84011103 |
| Cenpj | 372.423756 | 0.11133128 | 0.34118573 | 0.32630695 | 0.74419213 | 0.93906039 |
| Fam172a | 1679.38893 | 0.02293087 | 0.20932718 | 0.10954558 | 0.91276977 | 0.98292561 |
| Zfp69 | 137.988036 | 0.14731054 | 0.47045645 | 0.31312258 | 0.75418754 | 0.94018119 |
| Arih2 | 2571.12194 | -0.2246003 | 0.23463089 | -0.9572496 | 0.3384413 | 0.7564609 |
| Rab44 | 100.648889 | -3.8416527 | 0.9627728 | -3.9901966 | 6.601855426 | 0.00255207 |
| Tnnt1 | 444.370369 | 0.3688491 | 0.36072266 | 1.02252823 | 0.30653096 | 0.7358307 |
| Rab3ip | 324.173944 | -0.2956135 | 0.28677782 | -1.0308101 | 0.30262988 | 0.73372823 |
| Gm4735 | 26.5256738 | -1.1327245 | 0.90918315 | -1.2458706 | 0.21281192 | 0.64596443 |
| Ano6 | 1417.27783 | 0.31617633 | 0.36509176 | 0.86601881 | 0.38647985 | 0.78902002 |

| | | | | | | |
|---|---|---|---|---|---|---|
| Ifi27 | 253.520169 | 0.87336044 | 0.55532599 | 1.57269868 | 0.11578861 | 0.52055936 |
| Hist2h2aa1 | 55.1241179 | -0.1521929 | 0.73234556 | -0.2078156 | 0.83537293 | 0.96641606 |
| Plcxd1 | 21.0565286 | 0.2847095 | 0.88827824 | 0.32051838 | 0.7485754 | 0.9399191 |
| Ethe1 | 2341.55061 | -0.5806875 | 0.17961934 | -3.2328787 | 0.0012255 | 0.0271397 |
| Zfp428 | 151.391607 | -0.2737012 | 0.3413371 | -0.8018501 | 0.42263965 | 0.80826451 |
| Hvcn1 | 85.8785646 | 0.70459018 | 0.51226154 | 1.3754501 | 0.16899195 | 0.60312169 |
| Rpl19-ps1 | 23.295567 | 0.2173008 | 0.97356785 | 0.22320047 | 0.82337948 | 0.9628698 |
| Cdpf1 | 1170.81597 | 0.32916573 | 0.36177324 | 0.90986755 | 0.36289237 | 0.76990376 |
| Tank | 1365.96822 | -0.0910622 | 0.17417791 | -0.5228117 | 0.60110533 | 0.89086204 |
| Clasp1 | 849.083516 | -0.2711868 | 0.23579903 | -1.150076 | 0.25011259 | 0.6842534 |
| Lrrc51 | 98.5724701 | 0.4454415 | 0.48306147 | 0.92212178 | 0.35646506 | 0.76556128 |
| Siva1 | 3778.28299 | -0.1155362 | 0.37093043 | -0.3114768 | 0.7554382 | 0.94087513 |
| mt-Nd1 | 43847.2928 | -0.1381177 | 0.26740016 | -0.5165206 | 0.60549086 | 0.89258167 |
| mt-Nd2 | 3564.56155 | -0.2703903 | 0.26036859 | -1.0384905 | 0.29904177 | 0.73093388 |
| mt-Co1 | 67145.4694 | -0.1000442 | 0.33561782 | -0.2980896 | 0.7656348 | 0.94347227 |
| mt-Co2 | 25168.7342 | -0.1998288 | 0.31494674 | -0.6344844 | 0.52576473 | 0.86056156 |
| mt-Atp8 | 1451.90934 | -0.0168702 | 0.34021011 | -0.0495875 | 0.96045112 | 0.99325523 |
| mt-Atp6 | 66788.3681 | -0.1291979 | 0.22898682 | -0.5642153 | 0.5726076 | 0.88035336 |
| mt-Co3 | 11371.5784 | -0.1043944 | 0.23132744 | -0.451284 | 0.6517849 | 0.91037383 |
| mt-Nd3 | 2861.26129 | 0.19692821 | 0.27634191 | 0.7126252 | 0.47607771 | 0.83644936 |
| mt-Nd4 | 17018.8048 | -0.1232749 | 0.31812456 | -0.3875053 | 0.69838219 | 0.92669304 |
| mt-Nd5 | 9902.10758 | -0.2273466 | 0.39313059 | -0.578298 | 0.56306292 | 0.875995 |
| mt-Nd6 | 266.600486 | -0.2466706 | 0.40483608 | -0.6093098 | 0.54231911 | 0.8688043 |
| mt-Cytb | 52094.8196 | -0.1237156 | 0.279638 | -0.4424135 | 0.65819004 | 0.91258818 |
| Selenop | 64345.2721 | 1.73857416 | 0.39138422 | 4.44211616 | 8.907844774 | 0.00047173 |
| mt-Nd4l | 82.5451179 | 3.68944683 | 2.40427452 | 1.53453643 | 0.12489774 | 0.53626132 |
| C330021F23 | 69.1194867 | 0.24418877 | 0.57824897 | 0.42229002 | 0.67281334 | 0.91780571 |
| Tacc1 | 8119.45294 | -0.0253049 | 0.29286338 | -0.0864051 | 0.9311444 | 0.98730569 |
| Cpped1 | 937.469912 | 0.26292756 | 0.23342924 | 1.12636941 | 0.26000919 | 0.69625824 |
| Aurkaip1 | 3955.63837 | -0.0795643 | 0.3025903 | -0.2629441 | 0.7925937 | 0.95297175 |
| Zfp979 | 1291.80811 | 0.22613041 | 0.3733714 | 0.60564469 | 0.54475072 | 0.86963259 |
| Zfp600 | 29.0602897 | 0.17148753 | 0.87111188 | 0.19686051 | 0.84393669 | 0.96765585 |
| Zfp987 | 20.2041819 | -3.5297192 | 1.36820727 | -2.5798132 | 0.00988538 | 0.12828558 |
| Dhrs3 | 5795.36815 | 0.4949506 | 0.15984315 | 3.09647675 | 0.00195835 | 0.03946709 |
| Ubr4 | 864.852955 | 0.70473461 | 0.34512446 | 2.04197236 | 0.04115427 | 0.30904863 |
| Hnrnpr | 4240.48738 | -0.2058735 | 0.1763807 | -1.1672112 | 0.24312509 | 0.67814218 |
| Med18 | 466.7246 | 0.26138641 | 0.47049608 | 0.55555493 | 0.57851515 | 0.88230592 |
| Phactr4 | 134.916461 | -0.5285777 | 0.48423935 | -1.0915629 | 0.27502526 | 0.71147204 |
| Prpf4 | 2162.91501 | -0.2873399 | 0.2061648 | -1.3937391 | 0.1633964 | 0.59475807 |
| Cdc26 | 2569.99402 | 0.46924709 | 0.39826053 | 1.17824153 | 0.23870033 | 0.67452555 |
| Slc31a1 | 6197.57449 | 0.4156372 | 0.14899801 | 2.7895487 | 0.00527816 | 0.08320577 |
| Fkbp15 | 2991.06577 | 0.08583659 | 0.23163869 | 0.37056243 | 0.71096347 | 0.93147865 |
| Slc31a2 | 1591.80102 | 0.38221416 | 0.25408926 | 1.50425154 | 0.13251661 | 0.55219606 |
| Anks6 | 12.518914 | -1.1259493 | 2.02478699 | -0.5560828 | 0.57815423 | 0.8820854 |
| Ipo7 | 2236.18474 | -0.3196778 | 0.18712777 | -1.7083399 | 0.0875733 | 0.45207544 |
| Tmem42 | 198.316951 | 0.51559379 | 0.5481091 | 0.9406773 | 0.34687025 | 0.7607142 |

| | | | | | | |
|---|---|---|---|---|---|---|
| Pomgnt2 | 117.142628 | -0.1003513 | 0.67512622 | -0.1486408 | 0.88183709 | 0.97582756 |
| Trim12a | 354.38634 | -0.0818008 | 0.35283439 | -0.2318392 | 0.8166629 | 0.96079676 |
| Vps37b | 1502.37855 | -0.181371 | 0.29588781 | -0.6129721 | 0.53989481 | 0.86719392 |
| Numa1 | 1224.2824 | -0.3491172 | 0.30203773 | -1.1558727 | 0.24773322 | 0.68235696 |
| Impad1 | 339.697375 | -0.0644084 | 0.31157254 | -0.2067203 | 0.83622832 | 0.96683245 |
| Wdr6 | 1661.03521 | 0.21057047 | 0.18431034 | 1.14247776 | 0.25325549 | 0.68647781 |
| Rps13-ps1 | 41.2828399 | 0.36665832 | 0.66760458 | 0.5492148 | 0.58285805 | 0.88343755 |
| Serpina3f | 44.1588597 | -2.9922343 | 1.28393936 | -2.3305106 | 0.01977918 | 0.19950359 |
| Iqcf5 | 33.6741191 | -0.730179 | 0.84780904 | -0.861254 | 0.38909814 | 0.78942657 |
| Akap13 | 2600.56624 | -0.293306 | 0.38242019 | -0.7669731 | 0.44309749 | 0.82038676 |
| Msl2 | 251.03747 | -0.6852513 | 0.45713026 | -1.4990286 | 0.13386622 | 0.55436206 |
| Zfyve26 | 282.23357 | -0.1966783 | 0.60517431 | -0.3249944 | 0.74518532 | 0.93917909 |
| Rdh11 | 1176.24039 | 0.29736324 | 0.28088968 | 1.05864778 | 0.28976022 | 0.72395811 |
| Mthfs | 1225.89188 | 0.38728331 | 0.44065944 | 0.87887215 | 0.3794706 | 0.78211296 |
| Gm10163 | 27.7014019 | 1.35921729 | 1.94726501 | 0.69801351 | 0.48516874 | 0.8417344 |
| Hmgn3 | 41.413794 | -0.4994918 | 0.66631019 | -0.7496385 | 0.45347246 | 0.82635611 |
| Gm5786 | 19.3495016 | -0.0331256 | 0.85190399 | -0.0388842 | 0.96898275 | 0.99443087 |
| Rpl17-ps9 | 17.8434292 | 0.78395512 | 1.08747709 | 0.72089346 | 0.47097507 | 0.83381643 |
| Hmgb1 | 39001.5664 | -0.0707677 | 0.18922901 | -0.373979 | 0.70841993 | 0.92983917 |
| Lsm14a | 873.667146 | -0.2484896 | 0.23974757 | -1.0364635 | 0.29998597 | 0.73145694 |
| 4931406P16I | 305.028014 | 0.53801939 | 0.44522219 | 1.20842896 | 0.22688231 | 0.66015092 |
| Flvcr1 | 211.220237 | -0.451267 | 0.35238276 | -1.280616 | 0.20032856 | 0.63290788 |
| Zfp932 | 1044.48685 | 0.51784839 | 0.4067682 | 1.27307982 | 0.20298972 | 0.63809363 |
| Tecpr1 | 440.363575 | 0.2487702 | 0.49733212 | 0.50020941 | 0.61692763 | 0.89878056 |
| Gm5121 | 10.5845636 | 0.09608888 | 1.25305496 | 0.07668369 | 0.93887518 | 0.98910786 |
| Ttc32 | 697.588034 | -0.0272075 | 0.33941091 | -0.080161 | 0.93610922 | 0.98804349 |
| Fbxl18 | 264.825293 | -0.6188046 | 0.72978256 | -0.8479301 | 0.39647688 | 0.79444744 |
| Wdr35 | 131.026952 | 0.95693949 | 0.49657281 | 1.92708797 | 0.05396867 | 0.35989907 |
| Gm5113 | 57.3875378 | 0.62068453 | 0.60240896 | 1.03033747 | 0.30285162 | 0.73405663 |
| Ifi208 | 27.1347541 | 1.32400082 | 1.34239846 | 0.98629495 | 0.3239884 | 0.74583177 |
| Pilrb2 | 74.7764988 | -2.200363 | 1.00295114 | -2.1938885 | 0.02824342 | 0.24951635 |
| Pilrb1 | 100.286623 | -1.1649586 | 0.68555435 | -1.6992943 | 0.08926375 | 0.4566019 |
| Zbtb16 | 29.0465621 | -3.8954971 | 1.37310785 | -2.8369928 | 0.00455406 | 0.07415946 |
| Fxyd6 | 31.6117576 | 1.3572627 | 1.37437319 | 0.98755033 | 0.32337292 | 0.74583177 |
| Gm10175 | 3150.17121 | 0.0164297 | 0.24290669 | 0.0676379 | 0.94607389 | 0.99019648 |
| Vkorc1l1 | 271.945636 | 0.13584433 | 0.3722409 | 0.36493659 | 0.71515874 | 0.93235217 |
| Zbtb6 | 113.500421 | -0.191208 | 0.5072917 | -0.3769192 | 0.70623366 | 0.92896305 |
| Rnasel | 2550.98692 | 0.99323734 | 0.30486651 | 3.25794175 | 0.00112223 | 0.02561249 |
| Zfp810 | 191.110581 | 1.35266502 | 0.5844937 | 2.3142508 | 0.02065397 | 0.20441748 |
| Zfp772 | 165.430405 | 0.10215581 | 0.49867358 | 0.20485506 | 0.83768537 | 0.96683245 |
| Ecsit | 2084.08406 | -0.3732632 | 0.24970971 | -1.4947886 | 0.13496962 | 0.55648241 |
| Oas1g | 44.2378805 | -1.2702856 | 0.95779672 | -1.326258 | 0.18475426 | 0.61700012 |
| Nck2 | 24.0992049 | 1.39832265 | 1.00082865 | 1.39716489 | 0.16236399 | 0.59324932 |
| Gm10184 | 165.581659 | -0.6318091 | 0.40979988 | -1.5417504 | 0.12313426 | 0.53320405 |
| Zfp617 | 722.061265 | 0.5367387 | 0.30591017 | 1.75456314 | 0.07933406 | 0.43617688 |
| Fbxl12 | 283.496829 | 0.0963068 | 0.33397248 | 0.28836747 | 0.77306547 | 0.94625874 |

| | | | | | | |
|---|---|---|---|---|---|---|
| Vsig10 | 27.0431132 | 0.47517416 | 1.19058099 | 0.39911116 | 0.6898113 | 0.92470267 |
| Suds3 | 168.084815 | -0.6116016 | 0.430452 | -1.4208357 | 0.15536453 | 0.58452503 |
| Myo1h | 18.1938588 | -1.4705048 | 2.1396253 | -0.6872721 | 0.49191127 | 0.84516159 |
| Cryba4 | 650.022035 | 2.01519485 | 0.46312395 | 4.35130776 | 1.353279460 | 0.00068692 |
| Bub3 | 9936.77026 | -0.1967417 | 0.16522901 | -1.1907212 | 0.23376305 | 0.66954422 |
| Gm3608 | 210.772058 | 0.15096343 | 0.35572519 | 0.42438216 | 0.67128713 | 0.91711586 |
| Rps12-ps3 | 2743.05337 | 0.34207014 | 0.48506917 | 0.70519869 | 0.4806866 | 0.83943456 |
| Hes6 | 1791.59942 | -0.4709014 | 0.24737604 | -1.9035852 | 0.05696423 | 0.36695286 |
| Gm5529 | 13.5873549 | 0.11197668 | 1.05489185 | 0.10614991 | 0.91546341 | 0.98356437 |
| Polr1c | 2104.05768 | 0.06450176 | 0.35051963 | 0.18401755 | 0.85399969 | 0.96839628 |
| Xpo5 | 1935.73192 | -0.4451787 | 0.1975505 | -2.2534931 | 0.02422807 | 0.22470299 |
| Gm5560 | 14.3960671 | 0.08564386 | 0.96468054 | 0.0887795 | 0.92925715 | 0.98669576 |
| Eif1ax | 6174.7951 | 0.24027326 | 0.3358435 | 0.71543222 | 0.47434201 | 0.83575195 |
| Frat1 | 126.068768 | -0.8959941 | 0.40765862 | -2.1979031 | 0.02795601 | 0.24817555 |
| H2-K2 | 86.6201355 | 2.12486708 | 1.07098767 | 1.98402572 | 0.04725297 | 0.33529328 |
| Lrrc66 | 9.2282006 | 0.5601168 | 2.23882803 | 0.25018304 | 0.8024458 | 0.95658926 |
| H2-T23 | 599.848477 | -0.2886015 | 0.43952823 | -0.6566165 | 0.51142756 | 0.85375182 |
| Gm3695 | 10.5762196 | -0.2145419 | 1.10807615 | -0.1936165 | 0.84647614 | 0.96765585 |
| H2-Q10 | 22.7537954 | 1.82696532 | 1.01151042 | 1.80617548 | 0.07089093 | 0.41099943 |
| Rplp0 | 95760.0426 | 0.00575748 | 0.30966646 | 0.01859253 | 0.98516616 | 0.99564261 |
| Capn6 | 43.455986 | 1.72094768 | 1.04272185 | 1.65043793 | 0.0988534 | 0.48025164 |
| Rps28 | 7082.19338 | -0.0962086 | 0.29583092 | -0.3252147 | 0.74501858 | 0.93917909 |
| Bmpr2 | 430.088089 | 0.06164287 | 0.44398267 | 0.13884072 | 0.88957602 | 0.97870754 |
| Tmem128 | 1298.48076 | 0.31793807 | 0.33599409 | 0.9462609 | 0.34401553 | 0.75962035 |
| Lyar | 2742.52697 | -0.8486088 | 0.28148405 | -3.0147668 | 0.00257177 | 0.04780338 |
| Trmt2b | 868.441068 | -0.1664903 | 0.1943395 | -0.856698 | 0.39161179 | 0.79018736 |
| B3galt4 | 159.369572 | -0.1691238 | 0.3501026 | -0.4830692 | 0.62904657 | 0.90276005 |
| Tspan6 | 246.208004 | 1.38295347 | 0.83778023 | 1.65073539 | 0.09879262 | 0.48025164 |
| Zfp563 | 114.1664 | 0.3310882 | 0.92073565 | 0.35959094 | 0.71915306 | 0.93310883 |
| Zfp763 | 242.173838 | 0.3336112 | 0.58547902 | 0.56980896 | 0.56880727 | 0.87956256 |
| Gm7666 | 863.572794 | 0.7804149 | 0.66339261 | 1.17639974 | 0.23943518 | 0.67562272 |
| Hdac8 | 303.160086 | -0.3766311 | 0.28836291 | -1.306101 | 0.1915182 | 0.62259956 |
| S1pr3 | 22.2397567 | 2.37598555 | 1.86922839 | 1.271105 | 0.2036913 | 0.63809363 |
| Pcna-ps2 | 9.57988882 | -0.563827 | 1.29096118 | -0.4367498 | 0.66229283 | 0.91395288 |
| Ankrd23 | 30.3194053 | 0.56558333 | 1.84787657 | 0.30607203 | 0.75954981 | 0.94210735 |
| Prkag1 | 5272.99587 | 0.20371308 | 0.23817245 | 0.85531755 | 0.39237537 | 0.79076243 |
| Lpar5 | 195.903943 | 0.43688812 | 0.45487524 | 0.96045702 | 0.33682525 | 0.75534066 |
| BC003965 | 1551.80847 | 0.41030664 | 0.35325701 | 1.16149609 | 0.24544021 | 0.68067787 |
| Gm10222 | 1592.61724 | -0.4102385 | 0.37989567 | -1.0798715 | 0.28019942 | 0.71532524 |
| Xlr4b | 151.453349 | 0.46965726 | 0.65097479 | 0.72146767 | 0.47062183 | 0.83381643 |
| Nnat | 113.775072 | 1.00618519 | 0.52764439 | 1.90693809 | 0.05652861 | 0.36524665 |
| Blcap | 786.644544 | 0.76201141 | 0.20368958 | 3.74104264 | 0.00018326 | 0.00607898 |
| Myl9 | 59.2451046 | 0.75402003 | 0.78467502 | 0.96093289 | 0.33658591 | 0.75494233 |
| Pex26 | 38.5482739 | -0.81137 | 0.82242828 | -0.9865541 | 0.32386129 | 0.74583177 |
| Romo1 | 2501.19982 | 0.30501186 | 0.37442909 | 0.81460515 | 0.41529836 | 0.80395688 |
| Arfgef1 | 346.784716 | -0.9205078 | 0.32675224 | -2.817143 | 0.00484529 | 0.07758179 |

| | | | | | | |
|---|---|---|---|---|---|---|
| Rpl31-ps8 | 2568.69904 | 0.11714795 | 0.37303087 | 0.31404357 | 0.75348795 | 0.94018119 |
| Htatsf1 | 2008.98468 | -0.2559921 | 0.18002563 | -1.4219758 | 0.15503327 | 0.58408439 |
| Map7d3 | 164.485722 | -0.4769981 | 0.33309655 | -1.4320115 | 0.15214052 | 0.58009243 |
| Zfp991 | 1122.94788 | 0.388071 | 0.28505029 | 1.36141238 | 0.1733834 | 0.60651964 |
| Rtl8b | 110.61634 | 0.59495445 | 0.39261425 | 1.51536643 | 0.12967963 | 0.54639717 |
| Rtl8a | 97.9755401 | 0.12189886 | 0.43033891 | 0.28326247 | 0.77697564 | 0.94800948 |
| Zfp760 | 160.4728 | 1.45088096 | 0.725674 | 1.99935641 | 0.0455698 | 0.32946538 |
| Zfp948 | 503.271265 | 2.0429351 | 0.33097905 | 6.17240002 | 6.726106318 | 1.228066546 |
| Zfp160 | 299.361881 | 0.22860459 | 0.39050732 | 0.58540412 | 0.55827602 | 0.87321194 |
| Gtf2f2 | 1891.90753 | -0.0388133 | 0.27945954 | -0.1388869 | 0.88953954 | 0.97870754 |
| Lrch1 | 710.812994 | 0.23209945 | 0.35621993 | 0.65156222 | 0.51468362 | 0.85526675 |
| Afdn | 11.2263403 | 0.62650537 | 1.33542935 | 0.46914153 | 0.63896847 | 0.90615794 |
| Tcp1 | 29947.7562 | -0.2350101 | 0.18147858 | -1.2949742 | 0.19532912 | 0.62707955 |
| Tm9sf4 | 3828.83712 | -0.1867479 | 0.2919977 | -0.6395527 | 0.52246342 | 0.85816354 |
| Cenpm | 705.833269 | -0.2449974 | 0.27446223 | -0.8926454 | 0.3720471 | 0.77719221 |
| Ccdc134 | 992.913783 | -0.2524155 | 0.30968091 | -0.8150825 | 0.41502509 | 0.80395688 |
| Ninl | 368.183549 | 0.61321618 | 0.42977466 | 1.42683188 | 0.15362836 | 0.58242478 |
| Cst7 | 76.8846156 | -2.4800791 | 0.58306767 | -4.2535013 | 2.104536770 | 0.00094999 |
| Zfp442 | 102.046796 | 0.33617419 | 0.50302296 | 0.66830784 | 0.50393711 | 0.85012672 |
| Zfp120 | 1527.15711 | 0.38036585 | 0.33454482 | 1.1369653 | 0.25555279 | 0.69025543 |
| Gm10233 | 407.34327 | -0.0363918 | 0.61394978 | -0.0592749 | 0.95273319 | 0.9914747 |
| Ndufaf2 | 793.256064 | -0.079285 | 0.4085589 | -0.1940601 | 0.84612879 | 0.96765585 |
| Macrod2 | 118.928274 | 0.11627229 | 0.49992577 | 0.23257911 | 0.81608825 | 0.96075523 |
| Pick1 | 278.595607 | 0.1910943 | 0.31698239 | 0.60285463 | 0.5466054 | 0.86994632 |
| Lgals1 | 19250.5852 | 1.36626943 | 0.32314822 | 4.22799622 | 2.357817544 | 0.00104127 |
| Gm11808 | 4127.7968 | -0.0968503 | 0.2345884 | -0.4128521 | 0.67971496 | 0.92051478 |
| Phf11d | 86.1098902 | 1.03318824 | 0.59131508 | 1.74727192 | 0.08059018 | 0.43854735 |
| Amn1 | 172.783514 | 0.72175985 | 0.40501305 | 1.78206567 | 0.07473852 | 0.42093241 |
| Gm5879 | 25.7331399 | -0.559181 | 0.8372562 | -0.6678733 | 0.50421449 | 0.85012672 |
| Ap5s1 | 423.221107 | 0.00454792 | 0.23706687 | 0.01918414 | 0.98469421 | 0.99564261 |
| Cenpb | 320.478094 | 0.15576432 | 0.31976756 | 0.48711733 | 0.62617518 | 0.90181316 |
| Usf3 | 91.3824841 | -0.2265142 | 0.67072103 | -0.3377175 | 0.7355761 | 0.9377627 |
| Ddrgk1 | 1128.28398 | -0.0669553 | 0.25872086 | -0.2587936 | 0.79579446 | 0.9537323 |
| Nat8f4 | 30.8076139 | 0.64483704 | 1.43758469 | 0.44855586 | 0.65375208 | 0.91076467 |
| Slc4a5 | 104.500327 | -0.4736634 | 0.63995879 | -0.7401467 | 0.45921101 | 0.82987212 |
| Aup1 | 4437.64361 | -0.1206043 | 0.16643767 | -0.7246216 | 0.46868422 | 0.83381643 |
| Htra2 | 3653.18825 | -0.0666632 | 0.19001309 | -0.3508348 | 0.72571225 | 0.93544211 |
| Dok1 | 542.501666 | 0.22447681 | 0.21208001 | 1.05845341 | 0.28984878 | 0.72395811 |
| Chrac1 | 2349.73888 | 0.25283536 | 0.33571003 | 0.75313616 | 0.45136809 | 0.82559598 |
| Cep152 | 172.552826 | -0.3927477 | 0.57707211 | -0.6805869 | 0.49613293 | 0.84777781 |
| Rnase12 | 19.0334068 | -0.6709643 | 0.93392405 | -0.7184356 | 0.47248871 | 0.83483892 |
| Pnp2 | 37.5794426 | 0.10407242 | 0.7266914 | 0.14321405 | 0.88612113 | 0.97793139 |
| Uty | 141.789885 | 1.57252854 | 0.7952454 | 1.97741294 | 0.04799497 | 0.33778401 |
| Mfap1a | 1630.60988 | -0.1871548 | 0.16585481 | -1.1284256 | 0.25914024 | 0.69546977 |
| Gng5 | 6771.48931 | -0.0183096 | 0.22977787 | -0.079684 | 0.93648861 | 0.98827339 |
| Zfp467 | 363.08416 | 0.17467819 | 0.37937223 | 0.4604401 | 0.64520036 | 0.90835953 |

| | | | | | | |
|---|---|---|---|---|---|---|
| Myadm | 4456.56056 | -0.3580075 | 0.22370051 | -1.6003875 | 0.10951265 | 0.50553519 |
| Zfyve19 | 2568.18163 | 0.07161953 | 0.27376645 | 0.26160814 | 0.79362357 | 0.95310033 |
| Gm8225 | 7.97042133 | -0.0568964 | 1.31401024 | -0.0432999 | 0.96546251 | 0.99443087 |
| Actc1 | 71.944242 | -0.0536833 | 0.88354075 | -0.0607592 | 0.95155096 | 0.99116839 |
| Clec16a | 372.88594 | -0.2444206 | 0.26383175 | -0.9264259 | 0.35422469 | 0.76419569 |
| Cd59b | 15.2235849 | 0.74624986 | 0.94095713 | 0.79307531 | 0.42773395 | 0.81062404 |
| Myoz1 | 87.7977155 | 1.00836279 | 0.4594721 | 2.19461158 | 0.02819147 | 0.24951635 |
| Tmem167b | 1094.67887 | 0.04372031 | 0.21692637 | 0.20154445 | 0.84027288 | 0.96765585 |
| Trp53i11 | 40.5280955 | 0.80604958 | 1.01037505 | 0.79777265 | 0.42500243 | 0.80988459 |
| Sars | 5500.73065 | -0.0096445 | 0.16528384 | -0.058351 | 0.95346901 | 0.99149988 |
| Cry2 | 469.276107 | 0.04459581 | 0.43076884 | 0.10352608 | 0.91754545 | 0.98390186 |
| Psrc1 | 313.005103 | 0.07426967 | 0.32017202 | 0.23196803 | 0.81656284 | 0.96079676 |
| Sort1 | 510.575259 | -0.225988 | 0.25183561 | -0.8973632 | 0.36952513 | 0.77502139 |
| Psma5 | 19200.1641 | 0.43356513 | 0.38453028 | 1.12751883 | 0.25952318 | 0.69591124 |
| Il3ra | 104.182209 | -1.2468885 | 0.55180306 | -2.2596621 | 0.02384223 | 0.2230773 |
| Rap1a | 4903.713 | 0.27466428 | 0.22000938 | 1.24842078 | 0.211877 | 0.64580271 |
| Csde1 | 17201.5423 | -0.0385197 | 0.15976723 | -0.2410986 | 0.80947868 | 0.95844313 |
| Hist2h2be | 80.3213161 | 0.7483835 | 0.56256457 | 1.33030685 | 0.18341719 | 0.61524496 |
| Hist2h2ac | 56.8645806 | 0.24518571 | 0.67936269 | 0.36090547 | 0.71817011 | 0.93286094 |
| Sf3b4 | 3509.28757 | -0.344465 | 0.22695426 | -1.517773 | 0.12907164 | 0.54519878 |
| Gm128 | 59.3276102 | -0.5821665 | 0.83326423 | -0.6986578 | 0.48476593 | 0.84163505 |
| Selenbp1 | 685.174536 | -0.2487062 | 0.23452345 | -1.0604746 | 0.28892872 | 0.72395811 |
| Ssb | 8786.3391 | -0.1860767 | 0.1849929 | -1.0058586 | 0.31448361 | 0.74066151 |
| Clk2 | 135.012594 | -0.5837399 | 0.5804404 | -1.0056845 | 0.31456734 | 0.74066151 |
| Dap3 | 3759.66379 | -0.1451767 | 0.27736018 | -0.5234231 | 0.60067984 | 0.89046491 |
| Msto1 | 2533.23459 | -0.2762711 | 0.16272045 | -1.6978264 | 0.08954053 | 0.45701104 |
| Syt11 | 293.293467 | -0.0471837 | 0.45330279 | -0.1040887 | 0.91709894 | 0.98390186 |
| Zfp619 | 33.6752873 | 1.12221192 | 1.20826636 | 0.92877858 | 0.35300385 | 0.76376537 |
| Zfp114 | 43.5749816 | -1.2931156 | 1.87341052 | -0.6902468 | 0.49003901 | 0.84448327 |
| Gm6361 | 256.883413 | 0.1618082 | 0.47432129 | 0.34113629 | 0.73300099 | 0.93728136 |
| Zbtb34 | 261.675936 | -0.0942647 | 0.38305904 | -0.246084 | 0.80561721 | 0.95737871 |
| Gm5641 | 2336.8639 | -0.2039452 | 0.23923586 | -0.852486 | 0.39394438 | 0.79233901 |
| Urm1 | 469.799983 | 0.63832103 | 0.40105384 | 1.59160934 | 0.11147251 | 0.50888686 |
| Slc25a31 | 19.2805155 | 1.00657772 | 1.56392054 | 0.64362459 | 0.5198189 | 0.8576517 |
| Ddx3y | 794.071991 | 0.23307646 | 0.32100837 | 0.72607594 | 0.46779222 | 0.83381643 |
| Eif2s3y | 1058.18667 | -0.048639 | 0.32113847 | -0.1514579 | 0.87961454 | 0.97541902 |
| Cdk7 | 2742.56383 | 0.37493796 | 0.35225071 | 1.06440654 | 0.28714457 | 0.72276534 |
| Pde7a | 242.130266 | 0.71567768 | 0.40253126 | 1.7779431 | 0.0754132 | 0.42279087 |
| Zbtb10 | 74.4933377 | 1.05716106 | 0.74761893 | 1.41403731 | 0.15735095 | 0.58620126 |
| Gm10260 | 30.4337448 | 0.41872925 | 0.90582834 | 0.46226115 | 0.64389405 | 0.90823885 |
| Fgfr1op | 1429.56148 | -0.2292172 | 0.28116406 | -0.8152436 | 0.41493289 | 0.80395688 |
| Zfp72 | 58.2305047 | -0.2092717 | 0.5166139 | -0.4050834 | 0.6854162 | 0.92314423 |
| Zfp874a | 425.010065 | 0.99899661 | 0.58592098 | 1.70500227 | 0.088194 | 0.453691 |
| Zfp825 | 422.790897 | 0.35145081 | 0.32448586 | 1.08310053 | 0.2787638 | 0.71377266 |
| Fam8a1 | 66.2458036 | 0.96267586 | 0.54883641 | 1.75403061 | 0.07942526 | 0.43628614 |
| Dusp22 | 1744.3985 | 0.5988999 | 0.30086591 | 1.99058745 | 0.04652626 | 0.33401159 |

| | | | | | | |
|---|---|---|---|---|---|---|
| Hist1h3b | 13.0045058 | -0.7629143 | 1.05954735 | -0.7200379 | 0.47150163 | 0.83393588 |
| Hist1h2ac | 38.6978489 | 1.12271285 | 0.74765381 | 1.50164801 | 0.13318804 | 0.55309812 |
| Hist1h2ae | 1634.52936 | -0.2216368 | 0.32818421 | -0.6753426 | 0.49945811 | 0.84872332 |
| Hist1h3e | 18.1213045 | 0.29487723 | 0.99476402 | 0.29642933 | 0.76690223 | 0.94381438 |
| Hist1h2bj | 24.4829931 | -0.4056398 | 0.94083759 | -0.4311475 | 0.6663611 | 0.91520501 |
| Hist1h2ag | 133.977156 | -0.025993 | 0.48267656 | -0.0538519 | 0.95705318 | 0.99217217 |
| Hist1h4n | 35.5691915 | -0.1710211 | 0.86875979 | -0.1968566 | 0.84393974 | 0.96765585 |
| Hist1h2bq | 10.7317882 | -0.3664728 | 1.19940652 | -0.3055451 | 0.75995104 | 0.94215089 |
| Hist1h2bp | 12.6874176 | 1.35881738 | 1.34284992 | 1.01189072 | 0.31159031 | 0.74017637 |
| Hist1h2an | 30.1366892 | -0.3421443 | 0.7764785 | -0.4406359 | 0.65947658 | 0.91280998 |
| Hist1h3c | 51.3093425 | -0.1951968 | 0.67114091 | -0.2908433 | 0.77117119 | 0.94488554 |
| Epc2 | 147.423995 | 0.67367302 | 0.51038461 | 1.31993208 | 0.18685769 | 0.61862792 |
| Lyz2 | 59465.9917 | -0.3004819 | 0.42578968 | -0.705705 | 0.48037162 | 0.83935746 |
| Tmem19 | 1009.43912 | 0.19890523 | 0.2079933 | 0.95630595 | 0.33891769 | 0.75683748 |
| Scyl2 | 343.055319 | 0.39064641 | 0.49352115 | 0.79154947 | 0.42862342 | 0.81136783 |
| Dazap1 | 277.901628 | -0.7310139 | 0.42126674 | -1.7352756 | 0.082692 | 0.44231364 |
| Strada | 1694.0817 | -0.2296791 | 0.2196093 | -1.0458533 | 0.29562876 | 0.72852118 |
| Pex11g | 105.860271 | -0.1338459 | 0.47704035 | -0.2805756 | 0.77903592 | 0.94883461 |
| Marcks | 4878.24514 | 0.27033277 | 0.2234769 | 1.2096676 | 0.22640647 | 0.65990715 |
| Pcgf1 | 163.281722 | 0.21153067 | 0.47797988 | 0.44255142 | 0.65809023 | 0.91258818 |
| Gm10275 | 10990.006 | 0.34282656 | 0.42424359 | 0.80808897 | 0.41903937 | 0.80529433 |
| Zfp975 | 224.744567 | 0.78073642 | 0.44495487 | 1.7546418 | 0.0793206 | 0.43617688 |
| Arid1b | 278.821369 | 0.08212049 | 0.36480816 | 0.22510594 | 0.82189686 | 0.96232782 |
| Zfp820 | 256.266822 | 0.47949928 | 0.52409016 | 0.91491754 | 0.36023492 | 0.76864294 |
| Psmb3 | 11645.7819 | 0.00823369 | 0.31811107 | 0.02588307 | 0.97935061 | 0.99483019 |
| Zfp125 | 380.066301 | 0.27256042 | 0.30472753 | 0.89443975 | 0.37108665 | 0.7765614 |
| Msi2 | 118.311203 | -0.539961 | 0.55460993 | -0.9735871 | 0.33026162 | 0.74978051 |
| Wfdc17 | 495.825848 | 0.8759364 | 0.53504855 | 1.63711575 | 0.10160629 | 0.48717118 |
| Slfn9 | 709.097631 | 0.27599896 | 0.27628511 | 0.99896429 | 0.31781199 | 0.74307991 |
| Fam57a | 382.645037 | 0.47970276 | 0.25919526 | 1.85073894 | 0.06420712 | 0.3915482 |
| Nlrp1a | 192.388838 | 0.03985597 | 0.43430459 | 0.09176962 | 0.92688108 | 0.98630274 |
| Ahnak | 609.948517 | 1.24380319 | 0.27003117 | 4.60614675 | 4.101989578 | 0.00024719 |
| Sat2 | 112.051421 | 0.79695296 | 0.50625309 | 1.57421848 | 0.11543695 | 0.51974244 |
| Sco1 | 1040.27766 | -0.4308001 | 0.27042203 | -1.593066 | 0.11114538 | 0.50888686 |
| Rps12-ps9 | 9.99180515 | 0.82274566 | 1.20394312 | 0.68337586 | 0.49436937 | 0.84678075 |
| Irgm2 | 589.636047 | -0.042254 | 0.38195656 | -0.1106251 | 0.91191363 | 0.98292561 |
| 9930111J21F | 1383.68124 | 0.11692763 | 0.32276893 | 0.36226419 | 0.71715461 | 0.93259832 |
| 9930111J21F | 699.583361 | 0.57066397 | 0.4007688 | 1.42392315 | 0.15446872 | 0.58343244 |
| Atxn1l | 234.866158 | -0.4236461 | 0.58341875 | -0.7261441 | 0.46775046 | 0.83381643 |
| Gm12166 | 12.780622 | 1.33387887 | 1.12363011 | 1.18711563 | 0.23518201 | 0.6714076 |
| Spdl1 | 1637.19906 | -0.2562211 | 0.17033076 | -1.5042563 | 0.13251539 | 0.55219606 |
| Hba-a2 | 270.276692 | 0.30062881 | 0.86343152 | 0.34817909 | 0.72770569 | 0.93576253 |
| Hba-a1 | 314.065216 | -0.0509296 | 0.85458058 | -0.0595961 | 0.95247736 | 0.9914597 |
| B3gnt9 | 21.9056103 | 0.49113707 | 1.4500117 | 0.33871249 | 0.73482633 | 0.93750666 |
| Gm12070 | 257.454398 | -0.5020147 | 0.28632309 | -1.7533154 | 0.07954788 | 0.4364971 |
| Rps13-ps2 | 1422.47067 | 0.00494087 | 0.37803474 | 0.01306989 | 0.98957204 | 0.99650865 |

| | | | | | | |
|---|---|---|---|---|---|---|
| Fcho1 | 78.0154144 | -3.4265486 | 0.68092561 | -5.0321923 | 4.849024823 | 3.877001350 |
| Ell | 666.268429 | 0.00321761 | 0.44907025 | 0.00716505 | 0.99428317 | 0.99880383 |
| Ssbp4 | 1705.22226 | -0.025996 | 0.25547621 | -0.101755 | 0.91895117 | 0.9839023 |
| Gm10284 | 41.9158399 | -0.2153235 | 0.62171522 | -0.3463378 | 0.7290889 | 0.93617556 |
| Sp140 | 1275.8175 | -0.403993 | 0.17131338 | -2.3582104 | 0.01836328 | 0.19092573 |
| Sp110 | 1453.26244 | -0.3365666 | 0.23255144 | -1.4472782 | 0.14781902 | 0.57472133 |
| Fam149a | 136.511905 | 0.91452492 | 0.35664584 | 2.56423827 | 0.01034025 | 0.1312161 |
| Fat1 | 19.5104001 | 0.46059508 | 1.61807009 | 0.28465706 | 0.77590689 | 0.94776024 |
| Mfhas1 | 62.0958488 | 0.87336838 | 1.156323 | 0.75529794 | 0.45007024 | 0.82484234 |
| 3000002C10 | 84.2798173 | 0.55242923 | 0.48370748 | 1.14207295 | 0.2534237 | 0.68662948 |
| Ndufaf3 | 685.199677 | -0.5035455 | 0.27662013 | -1.82035 | 0.06870573 | 0.40407556 |
| Gmppb | 1320.23156 | -0.0708696 | 0.33598416 | -0.2109314 | 0.8329408 | 0.96565375 |
| Slc35g2 | 50.3585432 | 3.04942064 | 1.63399385 | 1.86623753 | 0.06200814 | 0.38446304 |
| Mpzl3 | 32.8816853 | 0.43485308 | 1.0005635 | 0.43460818 | 0.66384686 | 0.91399629 |
| Eif3g | 7383.78521 | -0.3462065 | 0.18872964 | -1.8344045 | 0.06659398 | 0.39719362 |
| Rnf213 | 293.635465 | -0.3524595 | 0.42732083 | -0.8248123 | 0.40947817 | 0.80100711 |
| Fbxo47 | 22.7577049 | 0.18429532 | 1.44647061 | 0.12741035 | 0.89861563 | 0.97989497 |
| Gm10288 | 603.172454 | 0.40836797 | 0.53849804 | 0.75834624 | 0.44824373 | 0.82382295 |
| Ccnd1 | 1313.25051 | 0.69132771 | 0.31074941 | 2.22471125 | 0.02610063 | 0.23668568 |
| Evi2 | 303.237759 | -0.1272637 | 0.34835505 | -0.3653277 | 0.7148668 | 0.93235217 |
| Prss36 | 80.9581174 | -0.2404206 | 0.62176138 | -0.3866767 | 0.69899558 | 0.92669304 |
| Capza1 | 30028.784 | 0.18546342 | 0.2037625 | 0.91019406 | 0.36272017 | 0.76990376 |
| Nlrp1b | 14.1846687 | 0.51008854 | 1.60473147 | 0.31786535 | 0.75058707 | 0.9399191 |
| Tmem256 | 2101.06374 | 0.97025334 | 0.52489625 | 1.84846691 | 0.06453483 | 0.39212252 |
| Hs3st3b1 | 52.5208668 | -0.1130199 | 1.26115062 | -0.0896165 | 0.92859195 | 0.98666423 |
| Xntrpc | 8.72095834 | -1.2816924 | 2.13967323 | -0.5990131 | 0.54916412 | 0.87093412 |
| Rnf121 | 1209.48544 | 0.64642326 | 0.29888277 | 2.16279864 | 0.03055667 | 0.26085118 |
| Il18bp | 10.3477161 | 0.30430204 | 2.20564432 | 0.13796514 | 0.89026797 | 0.97871887 |
| Serpinh1 | 761.585062 | 3.17600938 | 0.47379008 | 6.70341052 | 2.036104026 | 4.699558596 |
| Gm10291 | 16.4612481 | -0.717924 | 0.9736525 | -0.7373514 | 0.46090869 | 0.82987212 |
| Tlnrd1 | 269.919278 | 0.23388781 | 0.67514241 | 0.34642737 | 0.72902156 | 0.93617556 |
| Fam217b | 174.689333 | 0.16321822 | 0.42587226 | 0.38325628 | 0.70152976 | 0.92738786 |
| Gm10293 | 30.6123066 | -0.490711 | 0.67349398 | -0.728605 | 0.46624334 | 0.83345004 |
| Chchd2 | 19184.7801 | 0.21666561 | 0.32107144 | 0.67482057 | 0.49978977 | 0.84892089 |
| Nsmce3 | 1009.89703 | -0.1192914 | 0.17232236 | -0.6922574 | 0.48877567 | 0.8436846 |
| Gm6505 | 38.2278098 | 0.50628291 | 0.70031211 | 0.72293896 | 0.46971739 | 0.83381643 |
| Peg12 | 197.114776 | -0.0506242 | 0.41504494 | -0.1219728 | 0.90292059 | 0.98043702 |
| Mkrn3 | 15.2791891 | 2.1645962 | 1.89145935 | 1.14440535 | 0.25245557 | 0.68582922 |
| Top1 | 1712.55372 | -0.2538741 | 0.32920675 | -0.7711691 | 0.44060671 | 0.81963526 |
| Rasal2 | 33.3439509 | -0.2894431 | 0.89841277 | -0.3221716 | 0.7473227 | 0.9399191 |
| Trmt112-ps2 | 37.2812687 | -0.562417 | 1.27158494 | -0.442296 | 0.65827501 | 0.91258818 |
| Mn1 | 276.10685 | -0.9556635 | 0.50564767 | -1.889979 | 0.05876077 | 0.37321937 |
| Fv1 | 38.9974667 | -0.8072065 | 0.66971057 | -1.2053065 | 0.228085 | 0.66211765 |
| Vsig10l | 108.042984 | -0.0045716 | 0.86687376 | -0.0052736 | 0.99579227 | 0.99884675 |
| Zfp992 | 962.87524 | 0.2933608 | 0.31213041 | 0.93986613 | 0.34728623 | 0.76101142 |
| Lrrc8b | 74.7184393 | 0.62871656 | 0.57716747 | 1.08931392 | 0.27601548 | 0.7118544 |

| | | | | | | |
|---|---|---|---|---|---|---|
| Sox13 | 22.2220371 | 0.80391676 | 1.51948581 | 0.52907158 | 0.59675579 | 0.88891958 |
| Runx3 | 20.4091651 | -1.7948083 | 0.93528096 | -1.9190044 | 0.05498378 | 0.36084578 |
| Utp3 | 2646.11441 | -0.2090353 | 0.19249583 | -1.0859212 | 0.27751386 | 0.71260013 |
| Sars2 | 828.583348 | -0.3144837 | 0.21492558 | -1.4632214 | 0.1434068 | 0.56846902 |
| Eid2b | 83.8364488 | 0.1421813 | 0.58691709 | 0.24225108 | 0.80858562 | 0.95844313 |
| Zfp974 | 47.7447071 | 1.04807532 | 0.68059025 | 1.53995055 | 0.12357241 | 0.53434474 |
| Gm10282 | 314.609542 | -0.3450638 | 0.35706941 | -0.9663774 | 0.33385539 | 0.75320272 |
| Rmdn3 | 1826.42781 | -0.1201932 | 0.18114533 | -0.6635183 | 0.50699863 | 0.85158785 |
| Rbm44 | 86.6406225 | -0.6998655 | 0.51323196 | -1.3636437 | 0.17267974 | 0.60492813 |
| Fryl | 481.530431 | -0.0065574 | 0.53055038 | -0.0123596 | 0.99013872 | 0.99657454 |
| Tmem35b | 381.542161 | 0.88417081 | 0.32445016 | 2.72513596 | 0.0064275 | 0.09680087 |
| Dgkd | 1586.1923 | -0.0661399 | 0.26999998 | -0.2449628 | 0.80648524 | 0.95741157 |
| Rbm47 | 1967.80854 | 0.40258647 | 0.32801794 | 1.22733065 | 0.21969834 | 0.65289787 |
| Cited4 | 13.1843805 | -0.6983397 | 1.51764368 | -0.4601473 | 0.64541049 | 0.90847224 |
| Bicra | 82.678007 | -0.4227667 | 0.82251115 | -0.5139952 | 0.60725538 | 0.89332558 |
| Zswim9 | 112.499746 | -0.0135465 | 0.45405439 | -0.0298345 | 0.97619904 | 0.994746 |
| Zscan18 | 72.155412 | -0.3840809 | 1.30378586 | -0.2945889 | 0.76830792 | 0.94401697 |
| Gm1673 | 69.8437086 | 0.53267314 | 0.59776973 | 0.89110089 | 0.37287505 | 0.77748705 |
| Ccnyl1 | 148.761724 | 0.16985999 | 0.49388293 | 0.34392763 | 0.73090071 | 0.93653187 |
| Lilra5 | 46.0141901 | 1.75578062 | 1.19801855 | 1.46557048 | 0.14276534 | 0.56834178 |
| Ccdc173 | 16.7150393 | 1.02467882 | 0.98075578 | 1.0447849 | 0.2961224 | 0.72871963 |
| Klhl9 | 3449.56765 | 0.5080851 | 0.21114138 | 2.40637393 | 0.01611176 | 0.17660861 |
| Rraga | 794.605584 | 0.34995944 | 0.36493653 | 0.95895975 | 0.33757902 | 0.75611716 |
| Tgfbrap1 | 527.227312 | 0.42941718 | 0.31200437 | 1.37631785 | 0.16872325 | 0.60312169 |
| Il1rl2 | 754.66075 | 0.24264746 | 0.38327275 | 0.63309343 | 0.52667263 | 0.86115578 |
| Rabepk | 1540.22666 | 0.30836831 | 0.36745896 | 0.83919117 | 0.40136204 | 0.79737342 |
| Gm19680 | 1750.23747 | 0.89219482 | 0.20607285 | 4.32951177 | 1.494402839 | 0.00074616 |
| Dnajc25 | 147.356131 | 0.03193812 | 0.47929077 | 0.06663622 | 0.94687131 | 0.99019648 |
| Ndufb6 | 4254.39222 | 0.26284481 | 0.39407724 | 0.66698804 | 0.50477977 | 0.85034006 |
| Camkmt | 139.366093 | -0.5875659 | 0.57894567 | -1.0148895 | 0.3101585 | 0.73989254 |
| Rasgrp3 | 1273.54004 | 0.9218325 | 0.32544277 | 2.83254875 | 0.00461785 | 0.07492067 |
| Safb | 2076.19251 | -0.3202883 | 0.37042879 | -0.864642 | 0.38723531 | 0.78911204 |
| Treml2 | 491.58936 | -0.6118304 | 0.33005984 | -1.8536953 | 0.06378277 | 0.39032248 |
| Ptges3 | 15279.4543 | -0.2312244 | 0.13483078 | -1.7149226 | 0.08635941 | 0.44906118 |
| Yipf3 | 2056.92029 | -0.1124037 | 0.20978021 | -0.5358165 | 0.59208537 | 0.88797037 |
| Jund | 56.6580107 | 0.28790391 | 0.54634675 | 0.52696188 | 0.59822006 | 0.88951331 |
| Nr2c2ap | 1776.37963 | -0.0579581 | 0.26220887 | -0.2210381 | 0.82506277 | 0.96312579 |
| Srsf3 | 20476.0011 | 0.0390669 | 0.30090221 | 0.12983254 | 0.89669892 | 0.97984623 |
| Arhgef10 | 9.48686467 | 1.93688876 | 2.03551987 | 0.951545 | 0.34132778 | 0.75846782 |
| Smim15 | 3802.63723 | 0.77578014 | 0.31075139 | 2.49646556 | 0.01254379 | 0.1492686 |
| Naip5 | 180.756275 | -0.8431921 | 0.36202789 | -2.3290806 | 0.01985479 | 0.19957576 |
| Cecr2 | 32.32531 | 0.00277585 | 1.56029357 | 0.00177906 | 0.99858052 | 0.99967651 |
| 2210408I21R | 143.354791 | 0.58147161 | 0.44237117 | 1.31444281 | 0.18869723 | 0.61935434 |
| Slc25a16 | 176.030143 | 0.5310786 | 0.41413672 | 1.28237507 | 0.19971111 | 0.63247751 |
| Zfp213 | 416.657404 | 0.02591811 | 0.25229629 | 0.10272885 | 0.91817817 | 0.98390186 |
| Zfp946 | 529.719571 | -0.0853792 | 0.25303989 | -0.337414 | 0.73580483 | 0.9377627 |

| | | | | | | |
|---|---|---|---|---|---|---|
| Zfp942 | 461.995343 | 0.0001515 | 0.30217698 | 0.00050137 | 0.99959996 | 0.99976341 |
| Zfp65 | 160.095078 | 0.54001259 | 0.63885036 | 0.84528807 | 0.39795002 | 0.79531492 |
| Zfp58 | 215.908916 | 0.41229587 | 0.46358658 | 0.88936111 | 0.37380904 | 0.77748705 |
| 2610044O15 | 295.906001 | 0.72636708 | 0.48884258 | 1.48589161 | 0.13730779 | 0.55981285 |
| Rps8-ps1 | 9.36952717 | -0.334647 | 1.66879838 | -0.2005317 | 0.84106473 | 0.96765585 |
| Gpr31b | 20.6784389 | -0.8294303 | 1.13690052 | -0.729554 | 0.46566288 | 0.83339036 |
| Armc2 | 12.4560776 | -1.8141335 | 1.78311373 | -1.0173964 | 0.30896488 | 0.73867239 |
| Mfsd4b3-ps | 27.8850541 | 0.09125398 | 0.87164432 | 0.10469176 | 0.9166204 | 0.98389485 |
| Tia1 | 1772.17824 | -0.0345733 | 0.32438171 | -0.106582 | 0.91512058 | 0.98347559 |
| Egr4 | 9.34286015 | 0.16444141 | 1.92487349 | 0.08542972 | 0.93191976 | 0.98730569 |
| Gm10327 | 20.0431733 | -0.642936 | 0.87666722 | -0.7333867 | 0.46332264 | 0.83179129 |
| Setdb2 | 580.188728 | -0.1657143 | 0.20723537 | -0.7996429 | 0.42391773 | 0.80917688 |
| Tbpl1 | 2946.59197 | 0.10899625 | 0.21078477 | 0.51709738 | 0.60508818 | 0.89258167 |
| Map3k5 | 289.371838 | -0.3951136 | 0.3924187 | -1.0068674 | 0.31399849 | 0.74065408 |
| Hpcal1 | 1629.68594 | -0.4655442 | 0.18027214 | -2.5824524 | 0.00981009 | 0.1278028 |
| Rpl23 | 19360.7974 | -0.185005 | 0.18896893 | -0.9790233 | 0.32756846 | 0.74829972 |
| Psmg4 | 1076.05953 | 0.31780256 | 0.42956732 | 0.73982016 | 0.45940912 | 0.82987212 |
| Dtnb | 162.771209 | -0.3181204 | 0.47440038 | -0.6705738 | 0.5024921 | 0.85012672 |
| Zfp777 | 227.871289 | -0.9121955 | 0.44176117 | -2.0649066 | 0.03893184 | 0.29821741 |
| Hist1h2ad | 15.6584213 | 0.16588623 | 1.17019778 | 0.14175914 | 0.88727026 | 0.97827644 |
| Nutf2-ps1 | 8027.4351 | 0.09409202 | 0.23601584 | 0.39866824 | 0.69013767 | 0.92470267 |
| Hist1h2ai | 11.5893685 | 0.41346732 | 1.59576991 | 0.25910209 | 0.79555647 | 0.9537323 |
| Atp5md | 4864.46119 | 0.5784163 | 0.48964117 | 1.1813065 | 0.23748099 | 0.67388191 |
| Pcnp | 7226.05407 | 0.10811977 | 0.19187869 | 0.56347984 | 0.57310818 | 0.88035336 |
| Klrg2 | 52.1988682 | -0.5311412 | 0.70463 | -0.7537874 | 0.45097692 | 0.82549763 |
| Nt5dc2 | 1712.48221 | -0.7550802 | 0.1644415 | -4.5917861 | 4.394686956 | 0.00025971 |
| Cfap44 | 20.7069197 | -1.858238 | 1.69138268 | -1.0986502 | 0.27192065 | 0.70940613 |
| Cstdc5 | 417.812261 | -4.3830963 | 0.51486477 | -8.5131019 | 1.693410778 | 1.035774702 |
| Stfa1 | 1663.47689 | -4.0949869 | 0.38710835 | -10.5784 | 3.753130866 | 5.101338877 |
| Rnls | 92.0531633 | -0.0570082 | 0.50938189 | -0.1119164 | 0.91088973 | 0.98288717 |
| 2510002D24 | 718.313532 | 0.59093398 | 0.34544 | 1.71067042 | 0.08714197 | 0.45109601 |
| Cebpd | 965.077786 | -0.0850544 | 0.22776633 | -0.3734284 | 0.70882962 | 0.92997777 |
| Eef1g | 43280.7031 | -0.0230165 | 0.22727425 | -0.1012719 | 0.91933466 | 0.9839023 |
| Tut1 | 1072.38343 | -0.1258876 | 0.23677366 | -0.5316792 | 0.59494818 | 0.88830085 |
| Mta2 | 8543.55098 | -0.2762912 | 0.20494167 | -1.3481456 | 0.17761157 | 0.60906934 |
| Eml3 | 1173.77005 | -0.3517264 | 0.25031209 | -1.4051517 | 0.15997618 | 0.58927692 |
| Rom1 | 85.4764949 | 0.33874766 | 0.75542897 | 0.44841762 | 0.65385183 | 0.91076467 |
| B3gat3 | 472.593792 | 0.71558994 | 0.26394817 | 2.71110021 | 0.00670604 | 0.09943628 |
| Ganab | 6342.69219 | -0.1937387 | 0.195199 | -0.9925187 | 0.32094456 | 0.74413737 |
| Ints5 | 1344.90232 | -0.2118039 | 0.26952229 | -0.7858495 | 0.43195569 | 0.81344118 |
| 1810009A15I | 3361.04062 | -0.0182993 | 0.33520523 | -0.0545914 | 0.95646396 | 0.99215259 |
| Uqcc3 | 879.207307 | 0.15085474 | 0.35195856 | 0.42861507 | 0.66820338 | 0.91597175 |
| Ubxn1 | 2895.27736 | 0.04458659 | 0.19295538 | 0.23107203 | 0.81725884 | 0.96079676 |
| Bscl2 | 1779.12081 | -0.2997725 | 0.17705769 | -1.6930779 | 0.09044064 | 0.45983389 |
| Hnrnpul2 | 319.288377 | -1.0252214 | 0.37894147 | -2.7054874 | 0.00682042 | 0.10051233 |
| Ttc9c | 1944.433 | -0.1023644 | 0.21370572 | -0.4789968 | 0.63194089 | 0.90405469 |

| | | | | | | |
|---|---|---|---|---|---|---|
| Zbtb3 | 63.7743653 | -0.946878 | 0.73631414 | -1.2859702 | 0.19845347 | 0.63122167 |
| Polr2g | 3096.1185 | 0.121084 | 0.25263384 | 0.47928656 | 0.63173478 | 0.90399255 |
| Snx29 | 134.693281 | 1.93224989 | 0.95156893 | 2.03059372 | 0.04229623 | 0.31453479 |
| Sms | 2712.62415 | 0.00541423 | 0.37175639 | 0.01456391 | 0.98838009 | 0.99635239 |
| Csf2rb | 3447.92333 | -0.8340904 | 0.29745925 | -2.8040491 | 0.00504652 | 0.0804639 |
| Csf2rb2 | 180.908238 | -0.1665964 | 0.6161362 | -0.270389 | 0.78686102 | 0.95218824 |
| Ncf4 | 4135.19752 | -0.8066237 | 0.25491262 | -3.1643145 | 0.00155449 | 0.03267361 |
| Spin4 | 21.9904703 | 0.73733014 | 0.82026317 | 0.8988946 | 0.3687088 | 0.77458609 |
| Gspt2 | 319.855274 | -0.0930389 | 0.31145 | -0.2987283 | 0.76514737 | 0.94336301 |
| Smpd5 | 39.7306512 | 0.99131302 | 0.68351733 | 1.45031147 | 0.14697168 | 0.57367728 |
| Gm14698 | 154.242666 | -0.0163795 | 0.395421 | -0.0414228 | 0.9669588 | 0.99443087 |
| Zhx2 | 24.458207 | -0.865258 | 0.78067046 | -1.1083525 | 0.26770962 | 0.70441538 |
| 2610005L07F | 743.324421 | 0.14263679 | 0.36895855 | 0.386593 | 0.69905754 | 0.92669304 |
| 6820431F20I | 1078.20352 | 0.00344514 | 0.1923906 | 0.017907 | 0.98571305 | 0.99564261 |
| Apcdd1 | 17.9865694 | 2.32689382 | 1.40578305 | 1.65522968 | 0.09787792 | 0.47939805 |
| Ccdc112 | 32.6599009 | -0.0737805 | 0.91419411 | -0.0807055 | 0.93567614 | 0.98794094 |
| Mcc | 42.8757404 | 0.17948444 | 0.74249174 | 0.24173258 | 0.80898738 | 0.95844313 |
| Ppia | 94548.4988 | 0.26762476 | 0.3538653 | 0.75628993 | 0.44947539 | 0.82484234 |
| 6720489N17 | 495.86086 | 0.59037045 | 0.57728962 | 1.02265904 | 0.30646908 | 0.73582655 |
| Ccnf | 5348.93155 | -0.2510055 | 0.16845822 | -1.4900166 | 0.13621988 | 0.55833898 |
| Ang | 1027.39349 | 0.90407313 | 0.2934079 | 3.08128422 | 0.0020611 | 0.04112033 |
| Tuba1a | 5348.80339 | 1.10353195 | 0.44229664 | 2.49500414 | 0.01259557 | 0.14937799 |
| Trim6 | 216.649039 | 1.42027957 | 0.40214243 | 3.53178239 | 0.00041277 | 0.01152831 |
| Taf1a | 295.259521 | -0.8136249 | 0.34739084 | -2.3421023 | 0.01917546 | 0.19562439 |
| Klf12 | 17.6434794 | 0.52903949 | 0.98206779 | 0.53869956 | 0.59009417 | 0.88731984 |
| Nanos1 | 71.2724536 | -0.1711813 | 1.2069319 | -0.1418318 | 0.88721288 | 0.97827644 |
| Ppp1r3e | 12.5156208 | -1.4622758 | 1.50718106 | -0.9702058 | 0.33194392 | 0.75105716 |
| Phf20l1 | 1030.99835 | -0.078571 | 0.38297515 | -0.2051595 | 0.83744755 | 0.96683245 |
| Fam84b | 350.511036 | -0.2012957 | 0.360959 | -0.5576693 | 0.57707023 | 0.88121027 |
| Ptrh2 | 1766.69497 | 0.19800965 | 0.32609328 | 0.60721782 | 0.54370636 | 0.86947682 |
| Ear2 | 10.6239273 | -6.8116645 | 1.9711182 | -3.4557362 | 0.00054879 | 0.01441499 |
| Slfn2 | 5829.25454 | 0.34913997 | 0.18703052 | 1.86675401 | 0.06193595 | 0.38421017 |
| Slfn10-ps | 102.909761 | 0.87458509 | 0.83099328 | 1.05245747 | 0.29258971 | 0.72584701 |
| Zfp9 | 30.9269494 | 5.53553997 | 1.21101282 | 4.5710003 | 4.854015962 | 0.00028276 |
| Lyrm9 | 250.041178 | 1.33480795 | 0.71315784 | 1.87168656 | 0.06124998 | 0.38220919 |
| Zfp783 | 150.181288 | -0.6426923 | 0.39231433 | -1.6382075 | 0.10137841 | 0.48633809 |
| Spef2 | 20.3707361 | -0.1190728 | 1.40148112 | -0.0849621 | 0.93229153 | 0.98730569 |
| Tmem254a | 28.2098893 | -0.8429881 | 1.9512586 | -0.4320228 | 0.66572488 | 0.91520501 |
| Tmem254c | 347.471204 | -0.3674102 | 0.45312878 | -0.8108296 | 0.41746355 | 0.80498606 |
| Rpl37rt | 198.687138 | 0.44036836 | 0.64604644 | 0.68163576 | 0.4954693 | 0.84722895 |
| Gm10401 | 181.520655 | 0.02824316 | 0.32404107 | 0.08715919 | 0.93054498 | 0.98716777 |
| 1500011B03I | 1125.67846 | 0.26928628 | 0.35249803 | 0.76393698 | 0.4449048 | 0.82118815 |
| Smim10l1 | 1532.83104 | 0.1330525 | 0.22030027 | 0.60395977 | 0.54587039 | 0.86990153 |
| Gm10044 | 13.7464341 | 0.57116473 | 1.61085049 | 0.3545734 | 0.72290921 | 0.93433544 |
| 4930522L14F | 550.451941 | 1.05853485 | 0.53865308 | 1.96515139 | 0.04939672 | 0.34139551 |
| 5430403G16 | 67.4434372 | -0.876451 | 0.69614843 | -1.2590001 | 0.20803029 | 0.6418276 |

| | | | | | | |
|---|---|---|---|---|---|---|
| Acrbp | 25.0940276 | -0.4520265 | 1.49379067 | -0.3026037 | 0.76219193 | 0.94298237 |
| Grcc10 | 1259.74876 | 0.36707608 | 0.35711059 | 1.0279059 | 0.3039941 | 0.73436097 |
| Zfp951 | 450.681601 | 0.54895512 | 0.36802371 | 1.49162976 | 0.13579623 | 0.55758574 |
| Gm10420 | 14.2154705 | -0.8475797 | 0.97502094 | -0.8692939 | 0.38468641 | 0.78748265 |
| Rybp | 212.516802 | -0.1037914 | 0.43660359 | -0.2377245 | 0.81209478 | 0.95967622 |
| Nfxl1 | 450.038588 | 1.1436993 | 0.4613754 | 2.47889093 | 0.01317916 | 0.15501987 |
| Gm10434 | 8.7736582 | -0.4328654 | 1.40355057 | -0.3084074 | 0.75777237 | 0.94129085 |
| Noxred1 | 61.1537455 | -0.9869255 | 0.61103627 | -1.6151668 | 0.10627457 | 0.49777747 |
| Nup62cl | 27.8994925 | -0.3859832 | 1.7759702 | -0.2173365 | 0.8279461 | 0.96432112 |
| Ptgr2 | 2264.29283 | 0.23434624 | 0.19964791 | 1.1737976 | 0.2404761 | 0.67564175 |
| Acot1 | 70.5994889 | 0.64676718 | 0.66499247 | 0.97259323 | 0.33075551 | 0.75011761 |
| Tmsb15l | 35.6908005 | 0.42348606 | 0.77675087 | 0.54520191 | 0.58561466 | 0.8848945 |
| Bhlhb9 | 1659.0855 | -0.1978181 | 0.19629942 | -1.0077366 | 0.31358093 | 0.74065408 |
| Gprasp2 | 86.272974 | 0.26728179 | 0.70954974 | 0.37669211 | 0.70640242 | 0.92898526 |
| Armcx5 | 184.810739 | -0.003039 | 0.47495268 | -0.0063984 | 0.99489482 | 0.99884675 |
| Adam4 | 15.6411119 | 1.53375605 | 1.47147512 | 1.04232551 | 0.29726079 | 0.72954201 |
| Oip5 | 1320.2167 | 0.20694461 | 0.24427247 | 0.84718763 | 0.39689055 | 0.79492355 |
| Vamp5 | 165.754017 | 1.67244076 | 0.64698369 | 2.58498134 | 0.00973843 | 0.12700445 |
| Gm5127 | 13.6623652 | 0.40889016 | 1.74694449 | 0.23406019 | 0.81493825 | 0.95996209 |
| Uprt | 113.802201 | 1.56772283 | 0.47713578 | 3.28569539 | 0.00101731 | 0.02370428 |
| Zxdb | 54.0975949 | -1.6496151 | 0.87732069 | -1.8802875 | 0.06006891 | 0.37831609 |
| Srp54a | 5170.6489 | -0.0270779 | 0.16600591 | -0.1631141 | 0.8704286 | 0.97265735 |
| Lrrc61 | 351.914343 | 0.05666079 | 0.40223057 | 0.14086646 | 0.88797544 | 0.97832097 |
| Xlr3b | 242.201177 | 0.53601539 | 0.58200183 | 0.92098575 | 0.35705786 | 0.76632459 |
| Vma21 | 2105.24457 | 0.34717907 | 0.25030755 | 1.38700999 | 0.16543872 | 0.5984068 |
| Tmem185a | 1625.39912 | -0.1298222 | 0.19747429 | -0.6574131 | 0.51091534 | 0.85359566 |
| 9030624G23 | 34.8550123 | 0.95145879 | 0.70812001 | 1.34364058 | 0.17906464 | 0.60992118 |
| Zfp449 | 205.79826 | 0.8061137 | 0.82111286 | 0.98173314 | 0.32623134 | 0.74761857 |
| 5730507C01 | 94.6572462 | 0.41089357 | 0.72632509 | 0.56571579 | 0.571587 | 0.88035336 |
| Nudt11 | 12.2453353 | 2.0336531 | 1.62497275 | 1.25149982 | 0.21075218 | 0.64401792 |
| H2-D1 | 15192.2476 | 0.00266285 | 0.17184577 | 0.01549559 | 0.9876368 | 0.9962376 |
| Lst1 | 3432.17039 | 0.07714169 | 0.22014422 | 0.35041434 | 0.72602777 | 0.93558387 |
| H2-Ab1 | 95.4980575 | -0.1525276 | 0.59341799 | -0.2570323 | 0.79715384 | 0.95397994 |
| H2-Ke6 | 362.339112 | -0.1385117 | 0.35083512 | -0.3948055 | 0.6929864 | 0.92557077 |
| Zfp414 | 884.469942 | -0.2026093 | 0.20552304 | -0.9858228 | 0.32422006 | 0.74596967 |
| Gm4924 | 91.5182629 | 0.25751997 | 0.79666435 | 0.32324777 | 0.74650761 | 0.93960567 |
| Arhgdig | 7.95754493 | -1.1275443 | 1.81531774 | -0.6211278 | 0.53451554 | 0.8652846 |
| Wdr90 | 195.849822 | -0.5547728 | 0.41837237 | -1.3260263 | 0.18483099 | 0.61700012 |
| Nme3 | 584.953062 | 0.34130274 | 0.55071218 | 0.61974794 | 0.53542375 | 0.86539693 |
| Eme2 | 439.475876 | -0.3503425 | 0.33598771 | -1.042724 | 0.29707614 | 0.72954201 |
| Gm10509 | 51.1579497 | 0.39855403 | 0.563047 | 0.70785214 | 0.47903708 | 0.83869135 |
| Pnldc1 | 12.8908351 | -0.4216792 | 1.69304671 | -0.2490653 | 0.80331028 | 0.95672466 |
| Sft2d1 | 3104.07581 | 0.59893457 | 0.30219342 | 1.9819577 | 0.04748398 | 0.33634712 |
| Rsph3a | 726.408524 | -0.048732 | 0.41452297 | -0.1175617 | 0.90641499 | 0.98159944 |
| Marc2 | 3024.31034 | 0.29909794 | 0.24434165 | 1.22409727 | 0.22091554 | 0.65452328 |
| Ifi204 | 3176.43912 | -0.0047372 | 0.26316882 | -0.0180004 | 0.9856385 | 0.99564261 |

| | | | | | | |
|---|---|---|---|---|---|---|
| Ifi207 | 2982.42859 | 1.6537633 | 0.4701226 | 3.51772772 | 0.00043526 | 0.01207374 |
| Cep76 | 506.467217 | -0.9282546 | 0.44720346 | -2.0756873 | 0.03792288 | 0.29398642 |
| Gm4951 | 24.3214444 | 2.95280652 | 1.66337245 | 1.77519264 | 0.07586609 | 0.42455161 |
| Ppp1r12b | 40.6975062 | -0.2464617 | 1.35508797 | -0.1818788 | 0.85567787 | 0.96915653 |
| Csnk1g3 | 206.975912 | 0.27902092 | 0.43221866 | 0.64555501 | 0.51856758 | 0.8570838 |
| Ecscr | 154.450752 | 2.52489163 | 0.87338533 | 2.89092516 | 0.0038411 | 0.06499049 |
| D2hgdh | 409.892635 | 0.67112825 | 0.40312328 | 1.66482137 | 0.0959484 | 0.47482666 |
| Cops9 | 3097.63832 | 0.7436458 | 0.51900958 | 1.43281711 | 0.15191011 | 0.57999887 |
| Fbxo36 | 163.954077 | 0.26647682 | 0.35458072 | 0.75152654 | 0.45233584 | 0.82603347 |
| Rab18 | 4738.34952 | 0.13260104 | 0.15993674 | 0.82908432 | 0.4070567 | 0.79992362 |
| Rpl27-ps3 | 2154.36097 | 0.36731356 | 0.43623738 | 0.84200387 | 0.39978579 | 0.79612475 |
| Wdfy1 | 2762.61504 | -0.259147 | 0.16527072 | -1.5680154 | 0.11687753 | 0.52252063 |
| Gm10557 | 141.366386 | 0.2517331 | 0.39515108 | 0.63705532 | 0.52408881 | 0.8594073 |
| Catip | 17.0421971 | 1.45927903 | 1.51555532 | 0.96286754 | 0.335614 | 0.75372977 |
| Nbeal1 | 83.5164718 | -0.6036544 | 0.6549756 | -0.9216441 | 0.35671425 | 0.76595323 |
| Hspe1 | 10599.1584 | 0.23135492 | 0.40258791 | 0.57466931 | 0.56551497 | 0.87835762 |
| Pgap1 | 441.177391 | 0.49049534 | 0.45274316 | 1.08338544 | 0.27863738 | 0.71377266 |
| Faap20 | 1148.85093 | -0.2228695 | 0.16942228 | -1.3154673 | 0.1883529 | 0.61935434 |
| Klhl21 | 420.721997 | -0.194778 | 0.43406222 | -0.4487329 | 0.65362431 | 0.91076467 |
| Rpl31 | 8348.49033 | 0.0970712 | 0.35279934 | 0.27514564 | 0.78320435 | 0.95081654 |
| Cenps | 931.228795 | 0.11910749 | 0.34307139 | 0.34717989 | 0.72845618 | 0.93605089 |
| Lmbrd1 | 1484.52654 | 0.62996098 | 0.26900457 | 2.34182255 | 0.01918984 | 0.19562439 |
| Cplane2 | 56.3404327 | -0.1830476 | 0.63107909 | -0.290055 | 0.77177418 | 0.94534029 |
| Gm10566 | 10.1123452 | -0.1782086 | 1.13434931 | -0.1571021 | 0.87516441 | 0.97370497 |
| 5730409E04I | 42.4831601 | 1.43261279 | 0.82257185 | 1.74162633 | 0.08157385 | 0.44018215 |
| Btbd19 | 229.056208 | -1.0677561 | 0.438242 | -2.4364531 | 0.01483209 | 0.16769038 |
| Kti12 | 2016.45266 | -0.4706418 | 0.18665579 | -2.5214423 | 0.01168749 | 0.1424034 |
| Efcab7 | 282.151914 | 0.28046757 | 0.34876161 | 0.80418132 | 0.42129227 | 0.80727887 |
| Alg6 | 937.990201 | -0.1597336 | 0.26821928 | -0.5955334 | 0.55148695 | 0.87093412 |
| 6430531B16I | 23.1048045 | -0.2670294 | 1.38637518 | -0.1926098 | 0.84726457 | 0.96765585 |
| Tufm | 6868.2492 | -0.4737335 | 0.19772804 | -2.3958841 | 0.01658033 | 0.17972015 |
| Iqck | 32.6437482 | 1.34710451 | 1.34071708 | 1.00476419 | 0.31501041 | 0.74123109 |
| Ccl27a | 20.9451 | 1.25348033 | 1.07173412 | 1.16958143 | 0.24216945 | 0.67749347 |
| Il11ra1 | 1316.58425 | 0.69522078 | 0.19942458 | 3.48613378 | 0.00049006 | 0.01323718 |
| Gm1966 | 450.994533 | -2.1508123 | 0.47253704 | -4.5516268 | 5.323269716 | 0.0003043 |
| Mob3b | 364.062592 | -0.4216968 | 0.46409905 | -0.9086354 | 0.36354263 | 0.77021423 |
| Hbb-bt | 115.951695 | 2.39715567 | 0.79806152 | 3.00372291 | 0.00266698 | 0.0491343 |
| Trim68 | 161.205344 | 0.11201147 | 0.52675378 | 0.21264483 | 0.831604 | 0.96508982 |
| Rhog | 4878.37341 | -0.1288085 | 0.21313214 | -0.6043597 | 0.54560454 | 0.86990153 |
| Ggh | 1579.39165 | 0.23846815 | 0.35296674 | 0.67561082 | 0.49928777 | 0.84872332 |
| Ttpa | 93.6797764 | -1.063819 | 0.61812038 | -1.7210547 | 0.08524091 | 0.44656163 |
| Exoc8 | 199.038352 | 0.35924102 | 0.48016681 | 0.74815879 | 0.45436438 | 0.82736521 |
| Gm5921 | 22.609029 | -0.5263129 | 0.76948025 | -0.683985 | 0.49398465 | 0.84678075 |
| Osgin1 | 310.747615 | 0.14940371 | 0.35942264 | 0.41567696 | 0.6776464 | 0.91958179 |
| Mlycd | 445.76779 | 0.91140083 | 0.44407702 | 2.05234853 | 0.0401358 | 0.30439012 |
| Snrnp40 | 6181.74375 | -0.3887731 | 0.16084846 | -2.4170149 | 0.01564838 | 0.17370835 |

| | | | | | | |
|---|---|---|---|---|---|---|
| Svip | 226.162411 | -2.8010077 | 0.73634068 | -3.8039562 | 0.0001424 | 0.00486598 |
| Rbm15b | 208.635396 | -0.6523827 | 0.42282218 | -1.5429244 | 0.12284911 | 0.53283544 |
| Rpl13a | 22807.6972 | 0.64578954 | 0.47206814 | 1.36800069 | 0.17131186 | 0.60430173 |
| Il4i1 | 23.2891678 | 1.60461559 | 1.46427431 | 1.09584357 | 0.27314724 | 0.71005689 |
| Nlrc5 | 13.2620013 | -0.5957435 | 1.52345662 | -0.3910472 | 0.69576231 | 0.92623267 |
| Zfp976 | 40.993674 | 0.31059692 | 0.91482117 | 0.33951655 | 0.73422063 | 0.93750666 |
| Zfp788 | 262.941751 | 0.40418764 | 0.30918192 | 1.30728094 | 0.19111731 | 0.62228856 |
| AW146154 | 204.254685 | 0.04656361 | 0.51548075 | 0.09033046 | 0.92802461 | 0.98666423 |
| Plekhf1 | 151.270817 | 0.52944783 | 0.59875872 | 0.88424236 | 0.3765654 | 0.77905031 |
| Znhit6 | 1037.00641 | -0.3670643 | 0.24510277 | -1.4975932 | 0.13423896 | 0.55496628 |
| Zfp791 | 29.1600853 | 1.27460104 | 1.19961719 | 1.06250649 | 0.28800582 | 0.72304716 |
| G430095P16 | 13.7163186 | -1.8679707 | 1.61692386 | -1.155262 | 0.24798314 | 0.68235696 |
| Adh1 | 40.7635889 | 1.6899311 | 1.72995024 | 0.97686688 | 0.32863506 | 0.74843736 |
| Sdhaf1 | 650.243515 | 0.18549483 | 0.38845555 | 0.47751881 | 0.63299273 | 0.90430301 |
| Dnajb14 | 836.628958 | 0.18212558 | 0.26182067 | 0.69561193 | 0.48667189 | 0.84183502 |
| Gm10643 | 23.1481342 | 0.60018103 | 1.46431769 | 0.40987078 | 0.68190074 | 0.92061965 |
| Cox7a1 | 108.108734 | 1.38921855 | 0.73180686 | 1.89834042 | 0.05765125 | 0.36897087 |
| Zfp382 | 113.345611 | 1.03749844 | 0.55463575 | 1.87059423 | 0.06140135 | 0.38244534 |
| Zfp568 | 552.566063 | -0.0033074 | 0.38487264 | -0.0085934 | 0.99314353 | 0.99809447 |
| Spint2 | 41.5798021 | -1.8681878 | 0.78286891 | -2.3863354 | 0.01701722 | 0.18292766 |
| Ap1ar | 392.957787 | -0.20861 | 0.27815371 | -0.7499809 | 0.4532662 | 0.82610332 |
| Dda1 | 3834.56024 | -0.1642461 | 0.19072705 | -0.8611579 | 0.3891511 | 0.78942657 |
| Rec114 | 54.7653438 | 0.40637688 | 0.78425388 | 0.51817006 | 0.60433962 | 0.89258167 |
| Ceacam1 | 513.772244 | -0.0873691 | 0.35399845 | -0.2468063 | 0.80505811 | 0.95725854 |
| Zfp94 | 84.3367366 | 0.83767058 | 0.84158487 | 0.99534891 | 0.3195666 | 0.7440831 |
| Zfp109 | 63.5879875 | -0.954848 | 0.83085995 | -1.1492285 | 0.25046175 | 0.6842534 |
| Peak1 | 323.436311 | 0.60340938 | 0.37481026 | 1.60990624 | 0.10741833 | 0.50135383 |
| Ovgp1 | 13.5914459 | -1.0700636 | 1.52443065 | -0.7019431 | 0.48271465 | 0.84106286 |
| I830077J02R | 1735.89876 | -0.324089 | 0.22932381 | -1.4132377 | 0.15758584 | 0.58620126 |
| Tmigd3 | 43.0096667 | -2.3721874 | 1.10374015 | -2.1492264 | 0.03161646 | 0.26539272 |
| Ccdc61 | 433.484027 | -0.018107 | 0.34022365 | -0.0532208 | 0.95755595 | 0.99242502 |
| C5ar2 | 301.442057 | 0.42841044 | 0.38049294 | 1.12593533 | 0.2601929 | 0.69647146 |
| Ehd2 | 40.3745455 | 2.7388872 | 1.24530501 | 2.19937058 | 0.02785158 | 0.24777117 |
| Hist2h3b | 148.743958 | -0.5828496 | 0.40988002 | -1.4220006 | 0.15502608 | 0.58408439 |
| Zfp865 | 89.4475216 | -0.4734763 | 0.53907446 | -0.8783134 | 0.37977366 | 0.78237978 |
| Zfp628 | 16.4143223 | -0.8497357 | 1.41622349 | -0.6000011 | 0.54850549 | 0.87061992 |
| Gm10689 | 9.54907122 | -0.0009469 | 1.12187768 | -0.000844 | 0.9993266 | 0.99973522 |
| Gm14548 | 68.3909678 | -3.2595067 | 1.12749254 | -2.8909342 | 0.00384099 | 0.06499049 |
| Pate2 | 9.08105113 | 0.19056507 | 2.22721685 | 0.08556198 | 0.93181463 | 0.98730569 |
| S100a16 | 72.4105175 | 3.73662615 | 2.60395359 | 1.43498186 | 0.15129227 | 0.57926708 |
| Spc24 | 3991.77255 | -0.0833859 | 0.28598395 | -0.2915756 | 0.77061114 | 0.94454775 |
| Mex3a | 67.8326251 | 0.21819807 | 0.5674414 | 0.38452969 | 0.70058589 | 0.92706828 |
| Clec4g | 16.8579712 | -7.4772532 | 1.83899115 | -4.0659539 | 4.783639974 | 0.00194413 |
| Zfp558 | 72.8293298 | -1.0561978 | 0.78672383 | -1.3425268 | 0.17942526 | 0.60992118 |
| Arfip1 | 1375.43574 | -0.0791249 | 0.21932255 | -0.3607696 | 0.71827166 | 0.93286094 |
| Zfp971 | 226.069906 | 0.57677588 | 0.60234999 | 0.95754278 | 0.33829337 | 0.75644564 |

| | | | | | | |
|---|---|---|---|---|---|---|
| Gm14327 | 186.951694 | 0.50041536 | 0.55936525 | 0.89461287 | 0.37099407 | 0.7765614 |
| Gm14296 | 197.041533 | 0.36445355 | 0.50064947 | 0.72796151 | 0.46663715 | 0.83345004 |
| Cass4 | 516.761874 | 0.39637183 | 0.29445077 | 1.34613957 | 0.17825751 | 0.60928307 |
| Mocs3 | 18.7847823 | 0.04884748 | 0.8785335 | 0.05560115 | 0.95565955 | 0.99215259 |
| Lekr1 | 131.592363 | 1.44494352 | 0.48867668 | 2.95684972 | 0.003108 | 0.0554229 |
| 4931440P22I | 20.1937058 | 0.94354992 | 0.88786291 | 1.06272029 | 0.28790882 | 0.72304716 |
| Arfgef2 | 501.186206 | 0.21942971 | 0.33195284 | 0.66102677 | 0.50859515 | 0.85251363 |
| Mgst2 | 94.4055741 | -1.3989182 | 0.72423688 | -1.9315754 | 0.05341193 | 0.35841367 |
| Tox2 | 13.0684593 | 1.0702345 | 1.47942955 | 0.72341025 | 0.46942788 | 0.83381643 |
| Mafb | 951.973828 | -1.4049794 | 0.24857864 | -5.652052 | 1.585435982 | 1.999447255 |
| Cpne1 | 2991.77373 | -0.3441381 | 0.14645816 | -2.3497368 | 0.01878669 | 0.19344914 |
| 6430550D23 | 18.5277093 | 2.94711949 | 1.44424688 | 2.0405926 | 0.04129134 | 0.30948419 |
| BC029722 | 430.411018 | -0.3787407 | 0.25252296 | -1.4998269 | 0.13365925 | 0.55425547 |
| Eif2s2 | 8862.77435 | 0.06383394 | 0.28797212 | 0.22166707 | 0.82457307 | 0.96295965 |
| Tspyl3 | 427.738474 | 0.14018963 | 0.35957213 | 0.38987903 | 0.696626 | 0.92646244 |
| Zcchc3 | 1135.89688 | -0.3136979 | 0.24674428 | -1.2713482 | 0.2036048 | 0.63809363 |
| Csnk2a1 | 2864.81814 | -0.3622587 | 0.19612068 | -1.8471212 | 0.06472958 | 0.39236898 |
| Zfp950 | 312.740255 | 0.65094453 | 0.49421455 | 1.31712943 | 0.18779524 | 0.61935434 |
| Thbd | 383.901412 | -0.6091747 | 0.44654162 | -1.3642058 | 0.17250279 | 0.60492813 |
| Pdzd8 | 253.678525 | -0.4345647 | 0.36149289 | -1.202139 | 0.22930968 | 0.66356408 |
| Atxn7l3b | 2817.65791 | -0.0527133 | 0.16899955 | -0.3119141 | 0.75510582 | 0.94084431 |
| Kiz | 470.00349 | 0.17780233 | 0.31257139 | 0.5688375 | 0.56946642 | 0.87961466 |
| Smim26 | 271.827303 | 0.32500572 | 0.53597746 | 0.60637946 | 0.54426281 | 0.8695007 |
| Ube2n | 13299.8625 | 0.09704737 | 0.25881109 | 0.37497374 | 0.70767999 | 0.92950127 |
| Plxnc1 | 446.552837 | -0.5185492 | 0.53160484 | -0.975441 | 0.32934155 | 0.74874249 |
| Hspa12b | 8.85849911 | 0.56261356 | 2.70317007 | 0.20813102 | 0.83512666 | 0.96641606 |
| Arrdc3 | 1346.52858 | 0.0953242 | 0.21471784 | 0.44395099 | 0.65707802 | 0.91251438 |
| Itpa | 3937.06127 | 0.00647563 | 0.21045415 | 0.03076977 | 0.97545315 | 0.99450266 |
| Gas2l3 | 1080.51838 | 0.63856283 | 0.34963133 | 1.82638902 | 0.06779168 | 0.40101338 |
| Hps6 | 339.915097 | -0.7108541 | 0.42382785 | -1.6772236 | 0.09349877 | 0.46750793 |
| Rslcan18 | 23.8838315 | -0.802916 | 1.52294697 | -0.5272121 | 0.59804633 | 0.88951331 |
| Itpripl1 | 4292.85509 | 0.04682904 | 0.21570426 | 0.21709835 | 0.8281317 | 0.96434151 |
| Gm10767 | 15.1263178 | -0.0501796 | 1.53494315 | -0.0326915 | 0.97392059 | 0.99444457 |
| 2010315B03I | 402.139598 | 0.46768208 | 0.26802268 | 1.74493474 | 0.08099623 | 0.4392955 |
| Platr25 | 765.741845 | 0.13550611 | 0.33944139 | 0.39920326 | 0.68974344 | 0.92470267 |
| Zfp934 | 489.004492 | 0.13894678 | 0.33240359 | 0.41800625 | 0.67594255 | 0.91865406 |
| Zfp808 | 440.4363 | 0.97350488 | 0.50543327 | 1.92607991 | 0.0540944 | 0.35989907 |
| Ctla2b | 528.568695 | 2.66405346 | 0.52580653 | 5.06660397 | 4.049756055 | 3.302711055 |
| Serf2 | 9002.47969 | 0.43280624 | 0.33228521 | 1.30251432 | 0.19274063 | 0.62446288 |
| Grk6 | 349.647523 | -0.7227956 | 0.30238697 | -2.3903 | 0.01683462 | 0.18128333 |
| Lcmt2 | 423.694204 | 0.38708083 | 0.33340475 | 1.16099373 | 0.24564445 | 0.68093554 |
| Ifit3 | 89.4207519 | 1.79291475 | 1.21895758 | 1.47085901 | 0.14132925 | 0.5661037 |
| Ranbp6 | 161.444851 | 0.25811276 | 0.46030723 | 0.56074018 | 0.57497467 | 0.88059206 |
| Chst14 | 613.914556 | 0.19642218 | 0.22095183 | 0.88898189 | 0.37401281 | 0.77748705 |
| Inafm2 | 654.879964 | -0.5285811 | 0.44218533 | -1.1953837 | 0.23193717 | 0.66733333 |
| Fam122a | 48.6100146 | -0.2374146 | 0.69669902 | -0.3407707 | 0.73327624 | 0.93728136 |

| | | | | | | |
|---|---|---|---|---|---|---|
| Ptar1 | 436.521462 | -0.0745343 | 0.27504927 | -0.2709854 | 0.78640228 | 0.9519156 |
| Actg-ps1 | 11.7203838 | 0.24500443 | 1.08396654 | 0.22602583 | 0.82118133 | 0.96205421 |
| Qser1 | 175.153426 | -0.1891064 | 0.51121532 | -0.3699153 | 0.71144561 | 0.93191071 |
| Nrbf2 | 1218.50577 | -0.1379468 | 0.24601946 | -0.5607148 | 0.57499195 | 0.88059206 |
| Gm10800 | 16.5038331 | -0.0657911 | 1.01869297 | -0.0645838 | 0.94850537 | 0.99019648 |
| Prdm11 | 27.7880445 | -0.5570482 | 0.89562207 | -0.621968 | 0.53396294 | 0.86516141 |
| Hist1h2bb | 20.3191025 | -0.1955982 | 0.91468699 | -0.2138417 | 0.83067049 | 0.96466657 |
| Nxpe3 | 26.9868819 | -2.0823781 | 0.89707881 | -2.3212878 | 0.02027132 | 0.20128169 |
| Zfp408 | 110.526601 | 0.10380677 | 0.51497605 | 0.20157592 | 0.84024827 | 0.96765585 |
| Yae1d1 | 849.241921 | 0.34680534 | 0.22143377 | 1.56618087 | 0.11730628 | 0.52353438 |
| Cd80 | 84.5432577 | 0.28144593 | 0.49927056 | 0.56371425 | 0.57294862 | 0.88035336 |
| Ccdc162 | 52.6153501 | 0.7006655 | 0.72604254 | 0.96504745 | 0.33452105 | 0.75348848 |
| Znhit2 | 643.078856 | -0.5387934 | 0.24778285 | -2.1744578 | 0.02967077 | 0.25669202 |
| Ccdc58 | 1168.62292 | 0.34836609 | 0.38080407 | 0.91481714 | 0.36028764 | 0.76864294 |
| Amd1 | 6331.42572 | -0.317489 | 0.18161258 | -1.7481663 | 0.08043523 | 0.43809626 |
| Heg1 | 115.373894 | 1.23187852 | 0.54570477 | 2.25740837 | 0.02398257 | 0.22412435 |
| Cenpw | 1569.55313 | 0.15449912 | 0.34968842 | 0.44181937 | 0.65861992 | 0.91265586 |
| Bex6 | 132.239232 | -1.4242738 | 0.47850232 | -2.9765244 | 0.00291536 | 0.05299195 |
| Ttc30a1 | 422.581844 | 0.60485117 | 0.59171837 | 1.02219435 | 0.30668892 | 0.73594057 |
| Ttc30a2 | 52.4921665 | 1.17799951 | 1.38208513 | 0.85233499 | 0.39402819 | 0.79233901 |
| Ttc30b | 167.984575 | 0.54842583 | 0.48812364 | 1.12353874 | 0.26120876 | 0.69693483 |
| Mrpl23-ps1 | 2505.4514 | 0.12935506 | 0.28619172 | 0.45198744 | 0.65127803 | 0.91021183 |
| Wipf1 | 505.375496 | -0.354285 | 0.32939537 | -1.0755617 | 0.28212331 | 0.71778152 |
| Carns1 | 41.2531241 | -0.350939 | 0.975509 | -0.3597496 | 0.71903438 | 0.93310883 |
| H60b | 44.9191547 | 0.45856456 | 1.10309578 | 0.41570693 | 0.67762447 | 0.91958179 |
| Klhl41 | 10.6107734 | -0.7587208 | 2.65912405 | -0.2853273 | 0.77539337 | 0.94749646 |
| Zbtb2 | 725.448217 | -0.1204434 | 0.45349798 | -0.2655875 | 0.79055689 | 0.95261385 |
| Rc3h2 | 156.214919 | 0.8889053 | 0.47692991 | 1.86380697 | 0.06234882 | 0.38520862 |
| Fnbp1 | 3738.13115 | 0.10479992 | 0.17798282 | 0.58882042 | 0.55598174 | 0.87263596 |
| Dolk | 1361.90826 | 0.14935947 | 0.2173197 | 0.68727994 | 0.49190634 | 0.84516159 |
| Dnlz | 947.304582 | 0.26261562 | 0.2587589 | 1.01490467 | 0.31015127 | 0.73989254 |
| Alg10b | 421.196136 | -0.6371496 | 0.46718235 | -1.3638135 | 0.17262628 | 0.60492813 |
| Commd6 | 1990.52786 | 0.74277501 | 0.39003676 | 1.90437181 | 0.05686178 | 0.36677945 |
| Kbtbd6 | 125.459585 | -0.669538 | 0.53574081 | -1.2497423 | 0.21139369 | 0.64520435 |
| Aarsd1 | 2238.99375 | -0.2638564 | 0.15828228 | -1.6669993 | 0.09551455 | 0.47400792 |
| Nynrin | 105.828654 | 0.32313027 | 0.71772286 | 0.45021593 | 0.65255475 | 0.91051509 |
| Gal3st4 | 133.420218 | 0.27738261 | 0.45782228 | 0.60587399 | 0.54459843 | 0.86960875 |
| Zfp652 | 739.905966 | -0.1362492 | 0.53182138 | -0.2561935 | 0.79780146 | 0.95433165 |
| Zc3h3 | 213.550073 | -0.9136468 | 0.70641432 | -1.2933583 | 0.19588716 | 0.62779346 |
| Selenot | 1180.85743 | 0.0994383 | 0.32855908 | 0.30264967 | 0.76215686 | 0.94298237 |
| Selenos | 3627.90113 | 0.13592467 | 0.24297525 | 0.55941778 | 0.57587663 | 0.88080372 |
| Selenom | 118.528119 | -0.0020711 | 0.51033921 | -0.0040582 | 0.99676202 | 0.99937627 |
| Selenoi | 704.15322 | -0.6975022 | 0.40375702 | -1.7275296 | 0.08407259 | 0.44483565 |
| Txnrd2 | 2039.64282 | 0.26009135 | 0.16333154 | 1.59241347 | 0.11129183 | 0.50888686 |
| Msrb1 | 196.417446 | -0.3862465 | 0.35249928 | -1.0957369 | 0.27319393 | 0.71005689 |
| Gpx4 | 9366.9604 | 0.59020979 | 0.42687773 | 1.38262025 | 0.16678134 | 0.60024598 |

| | | | | | | |
|---|---|---|---|---|---|---|
| Sox4 | 139.64028 | 0.05492943 | 0.44513485 | 0.12339953 | 0.90179073 | 0.98043702 |
| Ywhaq | 26724.9178 | 0.35793294 | 0.28990408 | 1.23465989 | 0.21695711 | 0.65065857 |
| Acsf2 | 31.1858879 | -0.3664378 | 1.43881378 | -0.2546805 | 0.7989699 | 0.95475225 |
| Selenoh | 6603.4071 | 0.17824685 | 0.36449911 | 0.48901862 | 0.62482852 | 0.90093438 |
| Ass1 | 55.9531005 | -0.7339258 | 1.01827831 | -0.7207517 | 0.47106232 | 0.83381643 |
| Ighm | 13799.678 | 0.11272205 | 0.15406268 | 0.73166357 | 0.46437394 | 0.83257898 |
| Rab11b | 4605.98013 | -0.1436167 | 0.17559117 | -0.8179037 | 0.4134122 | 0.80364811 |
| Rps12l1 | 2120.93155 | 0.22354015 | 0.45576313 | 0.49047441 | 0.62379823 | 0.90017327 |
| Gm12355 | 12.7119817 | -0.0787005 | 1.05776759 | -0.0744024 | 0.94069018 | 0.98935054 |
| Psme2b | 161.092867 | 0.19669843 | 0.32927487 | 0.59736848 | 0.55026139 | 0.87093412 |
| Rbm8a2 | 10.8022463 | -0.6778507 | 1.08863628 | -0.6226604 | 0.53350769 | 0.86492515 |
| Chml | 168.125642 | 0.02470765 | 0.47654485 | 0.05184747 | 0.95865023 | 0.99249408 |
| Gm2000 | 6276.39067 | 0.59473623 | 0.46150846 | 1.28867893 | 0.19750973 | 0.63018689 |
| Tmem203 | 193.260014 | 0.06092561 | 0.32320968 | 0.1885018 | 0.8504833 | 0.9679302 |
| Gm3550 | 14.180654 | 0.48865539 | 1.24723769 | 0.3917901 | 0.69521331 | 0.92616369 |
| Hmga1b | 12.6742743 | -1.262592 | 1.0699284 | -1.1800715 | 0.23797178 | 0.67403588 |
| Gm2606 | 179.513737 | -0.2590062 | 0.33810286 | -0.7660574 | 0.44364214 | 0.82068545 |
| F8a | 200.52498 | 0.0005704 | 0.33275961 | 0.00171415 | 0.99863231 | 0.99967651 |
| Sf3b5 | 3513.89721 | -0.0344857 | 0.30157422 | -0.1143524 | 0.90895847 | 0.98209583 |
| Smim1 | 194.71143 | 2.93169497 | 0.51442218 | 5.69900579 | 1.205081350 | 1.610222881 |
| Gm6316 | 11.2879564 | -0.1142383 | 1.03936949 | -0.1099112 | 0.91247981 | 0.98292561 |
| Gm4294 | 29.5477385 | 0.1833732 | 1.92612881 | 0.09520298 | NA | NA |
| Sarnp | 5905.79657 | -0.090557 | 0.29071843 | -0.3114939 | 0.75542519 | 0.94087513 |
| Ctdsp2 | 604.442542 | 0.11708901 | 0.35964497 | 0.32556834 | 0.74475097 | 0.93917909 |
| AU041133 | 131.236622 | 1.27277676 | 0.66718211 | 1.90769019 | 0.05643127 | 0.36517938 |
| Smim24 | 143.046038 | 0.30295901 | 0.40823652 | 0.74211638 | 0.45801683 | 0.82986925 |
| Dohh | 4384.82236 | -0.1021956 | 0.20532359 | -0.4977293 | 0.61867485 | 0.89878056 |
| Raet1d | 209.105755 | 0.59158375 | 0.52943122 | 1.1173949 | 0.26382551 | 0.70008189 |
| Abracl | 2970.48349 | -0.0082835 | 0.25205389 | -0.0328641 | 0.97378299 | 0.99443087 |
| Plekhn1 | 63.7139398 | -0.4941362 | 0.58070725 | -0.8509213 | 0.39481307 | 0.79293192 |
| Cfap74 | 8.18605792 | 1.29319608 | 2.70711219 | 0.47770317 | 0.63286149 | 0.90430301 |
| Zfp984 | 1230.6666 | 0.3295426 | 0.24948176 | 1.32090859 | 0.18653184 | 0.61862792 |
| Zfp982 | 111.702227 | 0.28319548 | 0.43528582 | 0.65059661 | 0.51530692 | 0.85579005 |
| Gm13212 | 368.959852 | -0.1488019 | 0.3419228 | -0.4351914 | 0.66342349 | 0.91395288 |
| Ddi2 | 2157.68018 | -0.2847303 | 0.37790442 | -0.7534454 | 0.45118234 | 0.82550977 |
| Emc1 | 1522.58464 | -0.0347435 | 0.30025606 | -0.115713 | 0.90788004 | 0.9817854 |
| Aunip | 525.916857 | -0.2690115 | 0.2977663 | -0.9034317 | 0.36629681 | 0.77230417 |
| Zfp995 | 243.511967 | 1.08106158 | 0.45051807 | 2.3995965 | 0.01641315 | 0.17863176 |
| Bnip3 | 6736.79046 | 0.43123964 | 0.33635377 | 1.28210139 | 0.19980708 | 0.63247751 |
| 1110065P20I | 241.208396 | 0.71756159 | 0.54356709 | 1.32009756 | 0.18680245 | 0.61862792 |
| Ndufaf8 | 1027.46821 | 0.13429803 | 0.31752082 | 0.42295818 | 0.67232577 | 0.91761254 |
| Ube2d3 | 33760.4736 | 0.18644848 | 0.26771573 | 0.696442 | 0.48615207 | 0.84183502 |
| E430018J23F | 102.270516 | 0.6059415 | 0.91246877 | 0.6640682 | 0.50664666 | 0.85123041 |
| AU022252 | 606.533552 | 0.34815568 | 0.41955839 | 0.8298146 | 0.40664361 | 0.79992362 |
| Gm4609 | 31.3094344 | -0.9291506 | 0.82977556 | -1.1197613 | 0.26281548 | 0.69888997 |
| Gm4070 | 279.884247 | 0.03981389 | 0.36774349 | 0.10826539 | 0.91378517 | 0.98319938 |

| | | | | | | |
|---|---|---|---|---|---|---|
| Trim30c | 188.984104 | 0.31161525 | 0.43693852 | 0.71317872 | 0.47573517 | 0.83639059 |
| Smarcd2 | 601.363331 | 0.1435213 | 0.4515095 | 0.31786995 | 0.75058359 | 0.9399191 |
| Ccdc47 | 3874.64471 | -0.183063 | 0.22227052 | -0.8236046 | 0.41016424 | 0.80117888 |
| Tomt | 23.7871032 | -0.9825079 | 1.08012002 | -0.9096285 | 0.36301847 | 0.76990376 |
| Gm7336 | 249.507564 | 0.15045457 | 0.28690157 | 0.52441181 | 0.59999215 | 0.8902352 |
| Aoc2 | 8.70023163 | -0.7143638 | 1.40036933 | -0.5101253 | 0.60996368 | 0.89508835 |
| Psme3 | 2648.84423 | -0.1639498 | 0.88155825 | -0.1859772 | 0.85246261 | 0.96798445 |
| Cntd1 | 25.1158991 | 0.06716657 | 1.23605208 | 0.05433959 | 0.95666461 | 0.99215259 |
| Vps25 | 5603.30807 | -0.1373161 | 0.14384887 | -0.9545857 | 0.33978723 | 0.75726534 |
| Fam174b | 26.6200651 | -1.2434892 | 1.30515693 | -0.9527507 | 0.34071638 | 0.75821364 |
| Chd2 | 507.566295 | -0.2751081 | 0.34731588 | -0.7920976 | 0.42830374 | 0.81130995 |
| Casc3 | 457.023327 | -0.3933252 | 0.48553087 | -0.8100932 | 0.41788662 | 0.8051099 |
| Tm2d3 | 847.480474 | -0.0046825 | 0.23532595 | -0.0198978 | 0.98412493 | 0.99564261 |
| 5830417I10F | 492.07818 | 0.04670804 | 0.31874915 | 0.14653542 | 0.88349873 | 0.97667737 |
| Cisd3 | 500.738375 | 0.60761672 | 0.25151948 | 2.41578395 | 0.01570137 | 0.17413864 |
| Tomm5 | 4506.24326 | 0.46869484 | 0.48694114 | 0.96252875 | 0.33578407 | 0.7539733 |
| Haus5 | 2489.9981 | -0.1452847 | 0.22577478 | -0.6434939 | 0.51990366 | 0.8576517 |
| U2af1l4 | 1041.90017 | 0.17867235 | 0.30654164 | 0.58286487 | 0.55998427 | 0.87454201 |
| Zfp566 | 152.346132 | 0.14069585 | 0.65787365 | 0.21386454 | 0.83065269 | 0.96466657 |
| Evi2a | 5375.49727 | 0.26486759 | 0.27405068 | 0.96649126 | 0.33379842 | 0.75320272 |
| Rad54b | 783.543195 | 0.07650423 | 0.19356315 | 0.39524171 | 0.69266451 | 0.92549717 |
| Zfp59 | 153.586619 | -0.1370628 | 0.637022 | -0.2151618 | 0.8296412 | 0.96462215 |
| 1810022K09I | 2340.30064 | 0.17974225 | 0.36068865 | 0.49833078 | 0.61825092 | 0.89878056 |
| BC024978 | 224.795163 | 0.07828495 | 0.41618992 | 0.18809911 | 0.85079896 | 0.9679302 |
| Dph1 | 1384.74789 | 0.01231425 | 0.18595325 | 0.06622231 | 0.94720084 | 0.99019648 |
| Eif5a | 81631.615 | -0.2165089 | 0.20643046 | -1.0488226 | 0.29425979 | 0.72720808 |
| Leng1 | 557.662232 | -0.2160715 | 0.34528121 | -0.6257841 | 0.53145658 | 0.86432064 |
| Hist3h2a | 548.455258 | 0.47653555 | 0.24781119 | 1.92297833 | 0.05448277 | 0.36033644 |
| Igtp | 793.565297 | 0.0067093 | 0.20346795 | 0.03297473 | 0.97369474 | 0.99443087 |
| Zfp931 | 394.700531 | 0.87943886 | 0.63233723 | 1.39077508 | 0.16429364 | 0.59585062 |
| Gm14326 | 499.779086 | 0.77335925 | 0.51971698 | 1.48803922 | 0.13674055 | 0.55946682 |
| Gm14322 | 81.4465368 | 0.55563786 | 0.71651882 | 0.77546862 | 0.4380628 | 0.81873879 |
| Zfp970 | 36.8695768 | 0.01913051 | 0.67669944 | 0.02827032 | 0.97744655 | 0.99483019 |
| Gm14418 | 159.006681 | 0.6713193 | 0.50970628 | 1.31707089 | 0.18781486 | 0.61935434 |
| Gm14410 | 480.95279 | 1.08322699 | 0.58087142 | 1.86483092 | 0.06220511 | 0.38511032 |
| Gm14419 | 162.056702 | 0.66063709 | 0.7499354 | 0.88092532 | 0.37835824 | 0.78108777 |
| Gm14295 | 388.109319 | 0.93221546 | 0.55842784 | 1.66935708 | 0.09504663 | 0.47206881 |
| Gm14305 | 988.223343 | 1.31555561 | 0.57555909 | 2.28570032 | 0.0222718 | 0.21435947 |
| Zfp973 | 39.0572126 | 0.55662672 | 0.69391688 | 0.80215188 | 0.42246511 | 0.80821275 |
| Gm14308 | 45.7885467 | 1.21982407 | 0.68888644 | 1.77071865 | 0.0766075 | 0.42713744 |
| Gm14434 | 1200.83171 | 0.76007268 | 0.50325417 | 1.51031571 | 0.13096288 | 0.54940637 |
| Gm2026 | 67.5639792 | 0.22961259 | 0.74139653 | 0.30970282 | 0.75678696 | 0.94107188 |
| Gm6710 | 462.318708 | 0.77339713 | 0.49524481 | 1.56164611 | 0.11837138 | 0.52515149 |
| Gm14288 | 300.369695 | 1.70096798 | 0.81512699 | 2.08675214 | 0.03691055 | 0.28869994 |
| 2210418O10 | 977.133641 | 0.68660596 | 0.40492948 | 1.69561863 | 0.08995813 | 0.45827038 |
| Zfp965 | 465.208922 | 0.99839464 | 0.56308655 | 1.77307491 | 0.07621629 | 0.42553808 |

| | | | | | | |
|---|---|---|---|---|---|---|
| Gm4724 | 1040.08177 | 0.84577919 | 0.4950391 | 1.70850987 | 0.08754178 | 0.45207544 |
| Zfp968 | 108.980573 | 0.61982571 | 0.73802459 | 0.83984425 | 0.40099572 | 0.79698916 |
| Gm4631 | 304.609619 | -0.265433 | 0.29178244 | -0.9096951 | 0.36298335 | 0.76990376 |
| Gm14391 | 252.126717 | 0.6555349 | 0.5511641 | 1.18936431 | 0.23429634 | 0.67028698 |
| Mon1b | 653.853456 | -0.1328309 | 0.40854757 | -0.3251296 | 0.74508296 | 0.93917909 |
| Dpm1 | 1164.23112 | 0.00937955 | 0.19573712 | 0.04791909 | 0.96178072 | 0.99369539 |
| Ifi47 | 359.523099 | 1.23255045 | 0.43393826 | 2.84038204 | 0.00450595 | 0.07369162 |
| Ube2v1 | 12911.6971 | -0.1662638 | 0.17872008 | -0.9303027 | 0.35221438 | 0.76348958 |
| Pdf | 1836.83602 | -0.0607942 | 0.35569687 | -0.1709158 | 0.86428994 | 0.9705375 |
| Ak6 | 10850.2545 | 0.22098451 | 0.3050562 | 0.72440591 | 0.46881657 | 0.83381643 |
| Naip6 | 71.4800942 | -0.392655 | 1.01602583 | -0.3864616 | 0.6991548 | 0.92669304 |
| Naip2 | 567.730572 | -0.9200128 | 0.27724876 | -3.3183659 | 0.00090546 | 0.02163373 |
| Gm12033 | 29.3310329 | -0.987524 | 0.9076968 | -1.0879448 | 0.27661949 | 0.7118544 |
| Gm3272 | 41.9002011 | -0.2642186 | 0.69148814 | -0.3821014 | 0.70238615 | 0.92762537 |
| Wdr92 | 1136.27362 | 0.03402702 | 0.24607118 | 0.13828122 | 0.89001817 | 0.97871887 |
| Sec61g | 827.84338 | 0.57434788 | 0.49677028 | 1.15616393 | 0.24761411 | 0.68235696 |
| Zfp429 | 495.922159 | 0.78406111 | 0.31620442 | 2.47960196 | 0.01315291 | 0.15486003 |
| Zfp456 | 160.330491 | 0.72457698 | 0.42144051 | 1.7192865 | 0.08556222 | 0.44691828 |
| Samd1 | 103.096257 | -0.4356223 | 0.53763835 | -0.8102516 | 0.41779558 | 0.8051099 |
| Ifi27l2a | 57.1229743 | 0.13054341 | 0.86413291 | 0.15106867 | 0.87992155 | 0.97541902 |
| Slc45a4 | 448.342034 | 0.25622254 | 0.25958496 | 0.98704696 | 0.32361962 | 0.74583177 |
| Alkbh1 | 1210.63293 | -0.2421754 | 0.36030949 | -0.6721317 | 0.50149984 | 0.84968141 |
| Prnp | 69.2902756 | 0.89336312 | 0.67983135 | 1.31409521 | 0.18881417 | 0.61935434 |
| D130040H23 | 59.9963087 | -0.6239998 | 0.7395957 | -0.8437039 | 0.3988349 | 0.7957029 |
| Fastkd5 | 989.711257 | -0.5142238 | 0.23643424 | -2.1749126 | 0.02963667 | 0.25669202 |
| Kcnip3 | 179.592975 | -0.3176246 | 0.40214352 | -0.789829 | 0.42962763 | 0.81155572 |
| Cyp4v3 | 319.860511 | 0.73291289 | 0.30807159 | 2.37903433 | 0.01735806 | 0.18521003 |
| BC005561 | 48.6697876 | -0.6911557 | 1.23415736 | -0.5600224 | 0.57546418 | 0.88072731 |
| Jrkl | 79.2159309 | -0.4067541 | 0.52949168 | -0.7681974 | 0.44236994 | 0.82019298 |
| Ccdc82 | 607.767994 | -0.3062986 | 0.37088835 | -0.8258513 | 0.40888843 | 0.80100711 |
| Prps1l3 | 1713.16363 | -0.0699394 | 0.17439197 | -0.4010472 | 0.6883854 | 0.92470267 |
| Srp54c | 2836.60462 | -0.0659226 | 0.16287221 | -0.4047507 | 0.68566079 | 0.92314423 |
| Pms2 | 1354.75415 | 0.13981105 | 0.36189741 | 0.38632785 | 0.69925388 | 0.92669304 |
| Capn3 | 72.3626875 | -0.8940244 | 0.80690695 | -1.1079646 | 0.26787708 | 0.70441538 |
| Kdelr2 | 7324.08846 | -0.0692686 | 0.16857445 | -0.4109081 | 0.6811399 | 0.92061965 |
| Gm8818 | 230.615459 | 0.75413444 | 0.65818356 | 1.14578135 | 0.25188564 | 0.68505999 |
| A130010J15F | 594.95827 | 0.11446037 | 0.2929078 | 0.39077269 | 0.69596527 | 0.92623267 |
| Tlr5 | 47.115183 | 2.20335689 | 1.34806155 | 1.63446313 | 0.10216164 | 0.48856269 |
| Sap25 | 46.1385851 | 0.05452543 | 0.7331394 | 0.07437253 | 0.94071397 | 0.98935054 |
| Zan | 64.6630816 | 0.14100243 | 0.73755724 | 0.1911749 | 0.84838857 | 0.96765585 |
| Mphosph8 | 520.470772 | -0.0367361 | 0.27585496 | -0.1331719 | 0.89405743 | 0.97941957 |
| Psme2 | 8112.20091 | 0.24535995 | 0.38337942 | 0.63999248 | 0.52217749 | 0.85816354 |
| Zfp664 | 410.094236 | 0.0378163 | 0.37511866 | 0.10081157 | 0.91970004 | 0.9839023 |
| Ccr5 | 1683.0172 | -0.0944491 | 0.36586887 | -0.2581502 | 0.79629097 | 0.9537323 |
| Xirp1 | 13.2127959 | 4.7210759 | 1.62689999 | 2.90188453 | 0.00370925 | 0.06346194 |
| Trim71 | 104.31177 | 0.47895428 | 0.44827384 | 1.06844129 | 0.2853215 | 0.72114419 |

| | | | | | | |
|---|---|---|---|---|---|---|
| Tmppe | 533.847505 | 0.02362779 | 0.56888683 | 0.04153337 | 0.96687069 | 0.99443087 |
| 2310009B15I | 186.767139 | 0.38137036 | 0.53205606 | 0.71678604 | 0.47350613 | 0.83483892 |
| Clec7a | 6696.97002 | -0.6557257 | 0.41360197 | -1.5854028 | 0.11287484 | 0.5119755 |
| Gm3222 | 32.2301114 | -0.2305946 | 0.72113882 | -0.3197644 | 0.7491469 | 0.9399191 |
| Rab9 | 1297.26015 | 0.54031944 | 0.33876046 | 1.5949897 | 0.11071454 | 0.50858841 |
| Trappc2 | 656.562566 | 0.36181874 | 0.28318473 | 1.27767746 | 0.20136318 | 0.63552005 |
| Gm20661 | 7.70201335 | -0.1049064 | 1.9235007 | -0.0545393 | 0.95650548 | 0.99215259 |
| Naa80 | 301.748127 | 0.06994914 | 0.47008102 | 0.1488023 | 0.88170964 | 0.97582756 |
| Ifit1bl1 | 20.7214651 | 3.38854445 | 1.8422124 | 1.83938858 | 0.06585805 | 0.39558985 |
| Gbp4 | 100.812913 | 1.83358327 | 1.32346038 | 1.38544629 | 0.16591605 | 0.5992759 |
| Cntf | 16.2443594 | -0.9032204 | 1.53584563 | -0.5880932 | 0.55646973 | 0.87276537 |
| Atg4a | 1193.10253 | 0.40397933 | 0.32825363 | 1.23069263 | 0.21843785 | 0.65178055 |
| Ms4a6c | 6976.96282 | 1.10890147 | 0.3537491 | 3.13471176 | 0.00172023 | 0.03578286 |
| Arpc4 | 24185.3307 | 0.12138985 | 0.21616243 | 0.56156775 | 0.57441056 | 0.88059206 |
| Mthfsl | 1073.07769 | 0.43234406 | 0.45115892 | 0.95829658 | 0.33791323 | 0.75611716 |
| Mroh2a | 44.3524782 | -0.1518717 | 0.86742805 | -0.1750827 | 0.86101463 | 0.96996131 |
| Rpl36a | 6666.97457 | 0.05286612 | 0.3296688 | 0.16036131 | 0.87259647 | 0.97341534 |
| St6galnac4 | 4801.64688 | -0.2568361 | 0.18623369 | -1.3791067 | 0.16786185 | 0.60218592 |
| B3gnt7 | 205.350329 | 0.03834416 | 0.51941287 | 0.07382212 | 0.94115192 | 0.98935054 |
| Cldn34c1 | 74.5566242 | 0.74121675 | 0.58396047 | 1.26929268 | 0.2043367 | 0.63848041 |
| Pigb | 574.529735 | -0.123735 | 0.47405445 | -0.2610143 | 0.79408147 | 0.95310033 |
| Utp14b | 112.456721 | -0.1177967 | 0.5559873 | -0.2118695 | 0.83220886 | 0.96550322 |
| Rab7 | 15024.8347 | 0.16841557 | 0.14734632 | 1.14299137 | 0.25304217 | 0.68647781 |
| Sssca1 | 912.616714 | -0.4456886 | 0.27514024 | -1.6198599 | 0.10526238 | 0.49640504 |
| Pin4 | 630.141261 | -0.0870167 | 0.25638984 | -0.3393922 | 0.73431427 | 0.93750666 |
| Nhsl2 | 483.315666 | 0.16984448 | 0.57646944 | 0.29462877 | 0.76827748 | 0.94401697 |
| Phyhd1 | 536.735998 | 1.78410837 | 0.34621706 | 5.15314974 | 2.561472689 | 2.238178243 |
| Med12 | 248.560818 | -0.1506372 | 0.32769217 | -0.4596911 | 0.64573797 | 0.90847224 |
| Gm5138 | 44.4475677 | -0.9758895 | 0.68527783 | -1.4240785 | 0.15442374 | 0.58343244 |
| Apoo | 1125.17751 | 0.34027062 | 0.33759852 | 1.00791503 | 0.31349525 | 0.74065408 |
| Zfx | 951.877725 | 0.09307755 | 0.2827174 | 0.32922469 | 0.74198586 | 0.93825853 |
| Gm42688 | 47.239727 | 0.23595565 | 0.66731341 | 0.35359045 | 0.72364583 | 0.93453787 |
| Tmsb10 | 4142.67712 | -0.0043439 | 0.36448602 | -0.011918 | 0.99049104 | 0.9966603 |
| Gm5640 | 18.7784566 | -1.9734392 | 1.37832499 | -1.4317663 | 0.15221073 | 0.58009243 |
| H2-DMb1 | 108.89503 | -2.0614507 | 0.72008653 | -2.8627819 | 0.0041994 | 0.06998802 |
| Kifc1 | 142.474553 | 0.04028335 | 0.51274558 | 0.07856401 | 0.93737941 | 0.98845735 |
| Haus3 | 2225.32738 | -0.127831 | 0.2003845 | -0.6379286 | 0.52352018 | 0.85905063 |
| March2 | 242.70853 | 0.45699449 | 0.38915554 | 1.1743235 | 0.24026548 | 0.67564175 |
| Colca2 | 72.1598843 | 2.06642056 | 0.67963947 | 3.04046581 | 0.00236213 | 0.04507937 |
| Maea | 4032.49023 | -0.0749517 | 0.20844304 | -0.3595786 | 0.71916226 | 0.93310883 |
| Cstdc4 | 76.5895495 | -4.5840156 | 0.90800234 | -5.0484623 | 4.453803841 | 3.584433052 |
| Zbtb9 | 651.034978 | 0.06313475 | 0.36057213 | 0.17509603 | 0.86100417 | 0.96996131 |
| Stard6 | 59.0420066 | -0.3783317 | 0.78172173 | -0.4839723 | 0.62840551 | 0.90276005 |
| Ankrd39 | 372.912532 | -0.0164886 | 0.46546729 | -0.0354237 | 0.9717419 | 0.99443087 |
| Seh1l | 5355.81438 | -0.2865553 | 0.16987114 | -1.6868982 | 0.09162293 | 0.46315012 |
| Rpl39 | 6498.10741 | 0.14280379 | 0.40106977 | 0.35605723 | 0.72179771 | 0.93398714 |

| | | | | | | |
|---|---|---|---|---|---|---|
| Eloc | 4304.08155 | 0.37750582 | 0.2898379 | 1.30247227 | 0.19275499 | 0.62446288 |
| Tmem243 | 127.232282 | -0.5800721 | 0.55234917 | -1.050191 | 0.29363031 | 0.7265331 |
| Fdx1l | 1347.7611 | -0.1234065 | 0.25266676 | -0.4884161 | 0.62525517 | 0.9011746 |
| Ulbp1 | 443.363952 | 0.84951883 | 0.46192241 | 1.83909421 | 0.06590133 | 0.39558985 |
| Tcte3 | 6.67699268 | -1.1923876 | 2.82367516 | -0.4222821 | 0.6728191 | 0.91780571 |
| Gm3448 | 17.1279152 | 1.08612792 | 1.8411221 | 0.58992715 | 0.55523949 | 0.8722354 |
| 3110001I22R | 390.779455 | 0.20874601 | 0.31098253 | 0.67124674 | 0.50206336 | 0.85006796 |
| AC168977.1 | 7.92175166 | 0.66395102 | 1.40853906 | 0.47137566 | 0.6373725 | 0.90599323 |
| Tmlhe | 66.7463474 | 1.34800541 | 0.83248564 | 1.61925363 | 0.1053927 | 0.4968281 |
| Xlr4a | 44.223459 | 0.53191718 | 1.01855243 | 0.52222857 | 0.60151119 | 0.89094157 |
| Brms1 | 3135.85066 | 0.01362119 | 0.18940327 | 0.07191632 | 0.9426685 | 0.98983253 |
| B230307C23 | 286.008807 | 0.71894219 | 0.39702357 | 1.81082997 | 0.07016718 | 0.40928741 |
| Gm12174 | 12.1720526 | 0.66248256 | 1.16082944 | 0.57069759 | 0.56820465 | 0.8792976 |
| Gm15760 | 42.1653888 | 0.03686458 | 1.56048677 | 0.02362377 | 0.98115271 | 0.99522729 |
| Gm9385 | 10.2359923 | -0.1112575 | 1.74774568 | -0.0636577 | 0.94924279 | 0.99028544 |
| Rpl10-ps6 | 67.6882326 | -0.1262123 | 0.6089009 | -0.2072789 | 0.83579202 | 0.96674014 |
| Gm14387 | 13.7897029 | 0.4893993 | 1.37826309 | 0.3550841 | 0.72252659 | 0.93423486 |
| Ywhaq-ps3 | 18.1882754 | 0.44423287 | 1.19516122 | 0.37169284 | 0.71012155 | 0.93077435 |
| Gm14048 | 12.6999672 | 2.05898327 | 1.29491917 | 1.59004772 | 0.11182405 | 0.50962308 |
| Gm13464 | 25.7407645 | 0.28992139 | 0.78175722 | 0.37085861 | 0.71074284 | 0.93141199 |
| Psmb7-ps2 | 11.4268538 | 0.83054064 | 1.55875038 | 0.53282466 | 0.59415496 | 0.88797037 |
| Gm15427 | 65.3414925 | 0.02843952 | 0.5412634 | 0.05254285 | 0.95809615 | 0.99246834 |
| Hist2h3c2 | 211.223523 | -0.6373303 | 0.34813258 | -1.8307116 | 0.0671436 | 0.39891584 |
| Rpl19-ps11 | 31.6282725 | -0.1509222 | 0.77967581 | -0.1935705 | 0.84651221 | 0.96765585 |
| BC022960 | 45.435109 | -0.1063922 | 0.7914816 | -0.1344216 | 0.89306918 | 0.97941957 |
| Gm12922 | 13.4200822 | 0.07391982 | 1.10034393 | 0.06717883 | 0.94643934 | 0.99019648 |
| Gm12933 | 15.5467621 | 0.162945 | 0.94393993 | 0.17262222 | 0.86294839 | 0.9705375 |
| Rpl35a-ps2 | 12.6402688 | -0.11959 | 1.18566862 | -0.1008629 | 0.9196593 | 0.9839023 |
| Gm13509 | 13.8582108 | 0.59483021 | 0.960002 | 0.61961351 | 0.53551227 | 0.86539693 |
| Gm12397 | 82.2064802 | -0.6729909 | 0.53375964 | -1.2608502 | 0.20736283 | 0.6418276 |
| Rps6-ps4 | 4662.01056 | 0.15744856 | 0.33827925 | 0.46543961 | 0.64161668 | 0.90700632 |
| Hmgb1-ps3 | 27.8796337 | -0.3085044 | 0.71863784 | -0.4292905 | 0.66771186 | 0.91595755 |
| Slc48a1 | 3755.2803 | 0.56210139 | 0.17688938 | 3.17770008 | 0.00148448 | 0.03141812 |
| Gm12411 | 11.4352613 | -0.1699937 | 1.23900285 | -0.137202 | 0.89087111 | 0.97889088 |
| Gm12286 | 33.3951932 | -0.7552316 | 0.67300891 | -1.1221718 | 0.2617894 | 0.6974044 |
| Gm14633 | 12.7465083 | -0.4612856 | 1.08855163 | -0.4237609 | 0.6717402 | 0.91719387 |
| Gm15922 | 39.4644894 | -1.940066 | 1.05665974 | -1.8360366 | 0.06635224 | 0.39699582 |
| Rps2-ps13 | 17.6373598 | -0.2078384 | 0.90785178 | -0.2289343 | 0.81891995 | 0.96105733 |
| Gm15931 | 27.3691201 | -0.172249 | 0.78924644 | -0.2182449 | 0.82723828 | 0.9639556 |
| Sms-ps | 1874.37006 | 0.00661665 | 0.39361725 | 0.01680987 | 0.9865883 | 0.99611549 |
| Gm15599 | 9.30772524 | 0.04648597 | 1.25628584 | 0.0370027 | 0.97048285 | 0.99443087 |
| Gm15539 | 71.9227513 | 0.7912578 | 0.64295411 | 1.23065984 | 0.21845011 | 0.65178055 |
| Gm15484 | 6.94477623 | -0.366989 | 1.40446088 | -0.2613024 | 0.79385933 | 0.95310033 |
| Gm12435 | 58.8995795 | -0.0556876 | 0.62095749 | -0.0896803 | 0.92854131 | 0.98666423 |
| Dnajb3 | 20.6796002 | 2.49305856 | 1.15794927 | 2.15299463 | 0.0313191 | 0.26460586 |
| Gm13461 | 32.0026534 | -0.0179928 | 0.85599341 | -0.0210197 | 0.98322992 | 0.99564261 |

| | | | | | | |
|---|---|---|---|---|---|---|
| Pgam1-ps2 | 1850.87169 | -0.0187742 | 0.38536808 | -0.0487175 | 0.96114446 | 0.99338983 |
| Rpl17-ps8 | 20.2343929 | 1.35951184 | 1.09184028 | 1.24515634 | 0.21307429 | 0.64599096 |
| Ftl2-ps | 18.4176422 | 1.03672643 | 0.84887034 | 1.22130128 | 0.22197197 | 0.65525654 |
| Gm13230 | 9.20604531 | -0.0601909 | 1.31266676 | -0.0458539 | 0.96342669 | 0.99401755 |
| Gm14276 | 114.760342 | 0.23901923 | 0.48695399 | 0.49084562 | 0.62353564 | 0.90017327 |
| Rpl27-ps2 | 22.1209599 | 2.00522292 | 1.12403282 | 1.78395407 | 0.07443113 | 0.42056738 |
| Rpl5-ps1 | 14.2425585 | 0.80171994 | 1.15808125 | 0.69228298 | 0.48875964 | 0.8436846 |
| Pisd-ps1 | 23.1097749 | 1.42771902 | 0.84035305 | 1.69895144 | 0.08932833 | 0.4566019 |
| Gm12250 | 112.894911 | 1.80603555 | 0.97535432 | 1.85167124 | 0.06407305 | 0.39112055 |
| Gm8822 | 33.5855865 | -0.0445571 | 0.72092371 | -0.0618056 | 0.95071768 | 0.99089456 |
| Gm11425 | 6.27207576 | 1.02044574 | 2.32777143 | 0.43837884 | 0.66111168 | 0.91351849 |
| Gm13292 | 36.8781803 | -0.2712191 | 0.62430944 | -0.4344306 | 0.66397576 | 0.91399629 |
| Gm11688 | 8.09890969 | 1.38254143 | 1.92436832 | 0.71843909 | 0.47248659 | 0.83483892 |
| Rps6-ps3 | 11.6824018 | -0.0244459 | 1.05363466 | -0.0232015 | 0.98148951 | 0.99523682 |
| Gm11687 | 14.2686291 | -0.3591514 | 1.02802689 | -0.3493599 | 0.72681915 | 0.93576253 |
| Gm13456 | 50.9859837 | -0.0700742 | 0.64141626 | -0.1092491 | 0.91300489 | 0.98292561 |
| Gm15387 | 7373.72917 | -0.1286997 | 0.21175594 | -0.6077737 | 0.54333758 | 0.86919453 |
| Gm15443 | 62.9468449 | 0.33336052 | 0.63543805 | 0.52461529 | 0.59985065 | 0.8902352 |
| Dynlt1-ps1 | 586.162163 | 0.98880075 | 0.50182808 | 1.9703974 | 0.04879284 | 0.34068656 |
| Gm12366 | 33.3102493 | -0.0642095 | 0.74886651 | -0.0857423 | 0.93167127 | 0.98730569 |
| Gm14150 | 64.1970998 | -0.0444706 | 0.54346868 | -0.0818273 | 0.93478405 | 0.98775272 |
| Gm11953 | 8.81311709 | -0.7561362 | 1.27227074 | -0.5943202 | 0.55229798 | 0.87093412 |
| Rpsa-ps9 | 10.729814 | -0.3793604 | 1.40196783 | -0.2705913 | 0.78670536 | 0.95209405 |
| Gm5844 | 41.8357638 | -0.450146 | 0.65086958 | -0.6916071 | 0.48918413 | 0.84401986 |
| Gm12191 | 78.4006525 | 0.12119296 | 0.48145522 | 0.25172218 | 0.80125581 | 0.95615712 |
| Gm11249 | 27.7267367 | -0.0322324 | 0.69483921 | -0.0463883 | 0.96300072 | 0.99401755 |
| Gm14494 | 146.858165 | 0.53495177 | 0.61494579 | 0.86991696 | 0.38434579 | 0.78704643 |
| Gm14421 | 11.8163818 | -0.3688555 | 1.0316352 | -0.3575445 | 0.7206842 | 0.93398714 |
| Gm6341 | 11.9728811 | 0.60385199 | 1.08023509 | 0.55900053 | 0.57616136 | 0.88091263 |
| Gm11427 | 97.5104822 | -0.7911315 | 0.57834606 | -1.3679206 | 0.17133693 | 0.60430173 |
| Ctsf | 1195.31961 | 1.36730493 | 0.21298973 | 6.41958149 | 1.366494930 | 2.932689910 |
| AA414768 | 27.1678936 | -0.3974426 | 0.92906029 | -0.4277899 | 0.66880409 | 0.91618049 |
| Vcp-rs | 22.3963838 | -0.8922501 | 0.84780575 | -1.0524229 | 0.29260558 | 0.72584701 |
| Llph-ps2 | 263.850341 | -0.2765207 | 0.31268187 | -0.8843514 | 0.37650652 | 0.77905031 |
| Ndufb4c | 931.806883 | 0.65413758 | 0.47618877 | 1.37369383 | 0.16953675 | 0.60382474 |
| Gm15542 | 25.9765037 | -0.1476062 | 1.10825347 | -0.1331881 | 0.89404461 | 0.97941957 |
| Gm15470 | 25.0776177 | -0.3794843 | 1.14938385 | -0.3301633 | 0.7412766 | 0.93800224 |
| Gm13340 | 14.2506851 | -0.6866056 | 1.16979426 | -0.5869456 | 0.55724022 | 0.87276537 |
| Gm4459 | 15.5328294 | 0.55467614 | 1.55918593 | 0.35574728 | 0.72202984 | 0.93398714 |
| Rasl2-9 | 30.0051539 | 2.19331008 | 1.10319151 | 1.9881499 | 0.04679511 | 0.33444287 |
| Gm10169 | 10.8201379 | -0.0449992 | 1.18708374 | -0.0379074 | 0.96976153 | 0.99443087 |
| Zfp133-ps | 16.6961382 | -1.0483565 | 1.53241626 | -0.6841199 | 0.49389944 | 0.84678075 |
| Gm8624 | 12.3630843 | 0.44008209 | 1.07084 | 0.41096904 | 0.68109523 | 0.92061965 |
| Gm5883 | 9.89214237 | -0.4384526 | 1.13476101 | -0.3863832 | 0.6992129 | 0.92669304 |
| Ccnb2-ps | 9.44298825 | -0.6631042 | 1.85435193 | -0.3575935 | 0.72064754 | 0.93398714 |
| Gm7803 | 17.8815133 | -0.2030369 | 0.96629331 | -0.2101193 | 0.83357456 | 0.96572759 |

| | | | | | | |
|---|---|---|---|---|---|---|
| Rps15a-ps4 | 176.226414 | 0.4283316 | 0.52749759 | 0.81200674 | 0.41678776 | 0.8047849 |
| Gm13394 | 36.2640171 | -0.7726076 | 0.64678332 | -1.1945386 | 0.23226736 | 0.6674267 |
| Gm13033 | 18.4845698 | -0.6889631 | 1.38414202 | -0.4977546 | 0.61865702 | 0.89878056 |
| Hmgb1-ps5 | 14.6766498 | -1.2791161 | 1.01369219 | -1.2618388 | 0.20700681 | 0.64174209 |
| Ube2d-ps | 596.087107 | 0.79031012 | 0.40753617 | 1.9392392 | 0.05247222 | 0.35389626 |
| Gm13341 | 9.48974094 | -0.3896354 | 1.1853142 | -0.3287191 | 0.74236804 | 0.93840202 |
| Gm12346 | 46.110489 | -0.0605845 | 0.61413297 | -0.0986505 | 0.92141575 | 0.98416824 |
| Gm4342 | 9.9922383 | 0.94038786 | 1.1993809 | 0.78406106 | 0.43300428 | 0.81435873 |
| Gm6136 | 22.5573527 | 0.09478643 | 0.95299448 | 0.09946167 | 0.92077172 | 0.9839023 |
| Gm15710 | 18.844754 | 0.14720511 | 0.98809164 | 0.14897921 | 0.88157004 | 0.97582756 |
| Rpl3-ps2 | 22.386842 | 0.45489815 | 0.84857006 | 0.53607613 | 0.59190593 | 0.88797037 |
| Gm15421 | 12.9580235 | 0.08206715 | 1.25064565 | 0.06561982 | 0.94768051 | 0.99019648 |
| Gm6977 | 25.507177 | 0.38459327 | 0.83738582 | 0.45927846 | 0.64603422 | 0.90869686 |
| Tpt1-ps3 | 4283.17847 | 0.3252715 | 0.35713943 | 0.91076893 | 0.36241714 | 0.76969598 |
| Gm14438 | 13.660447 | 0.00823166 | 1.56070845 | 0.00527431 | 0.99579173 | 0.99884675 |
| Rpl3-ps1 | 21.1840658 | 0.01160746 | 0.89518563 | 0.01296654 | 0.98965449 | 0.99650865 |
| Znf41-ps | 586.638788 | 0.46363549 | 0.31428312 | 1.47521603 | 0.14015447 | 0.56396008 |
| Rps2-ps4 | 18.3554587 | 0.17009868 | 0.91573427 | 0.18575114 | 0.85263993 | 0.96798445 |
| Ubl5 | 1949.66432 | 0.09083392 | 0.28748934 | 0.31595578 | 0.75203607 | 0.94018119 |
| Ccdc85c | 273.868609 | 0.22194976 | 0.37151523 | 0.59741766 | 0.55022856 | 0.87093412 |
| Bbip1 | 1525.22245 | 0.61073678 | 0.4073971 | 1.49911912 | 0.13384273 | 0.55436206 |
| Slc2a4rg-ps | 37.9987085 | 0.52345303 | 0.92043855 | 0.5686996 | 0.56956002 | 0.87961466 |
| Abhd11os | 204.800126 | 0.79658555 | 0.41940646 | 1.89931637 | 0.05752289 | 0.36897087 |
| Gm3362 | 15.8450982 | -0.4588219 | 0.96366236 | -0.4761231 | 0.6339867 | 0.90465549 |
| Gm11346 | 17.8812551 | 2.37856718 | 1.46140845 | 1.62758548 | 0.10361281 | 0.49263248 |
| 3110053B16I | 18.2669554 | -0.3162127 | 1.03712809 | -0.3048926 | 0.76044794 | 0.94215089 |
| Gm13092 | 75.0802266 | 0.04066006 | 0.77484562 | 0.05247504 | 0.95815018 | 0.99246834 |
| Lin52 | 806.085526 | -0.149592 | 0.29830403 | -0.5014748 | 0.61603698 | 0.89878056 |
| Zfp703 | 163.127494 | 0.65216612 | 0.59947533 | 1.08789485 | 0.27664153 | 0.7118544 |
| Gm7805 | 8.91120174 | 0.44777432 | 2.15544293 | 0.20774121 | 0.83543104 | 0.96641606 |
| Ccpg1os | 49.4356498 | 0.50034789 | 0.57884134 | 0.86439558 | 0.38737062 | 0.78911204 |
| Nbdy | 720.062662 | -0.0074759 | 0.28759097 | -0.0259949 | 0.97926142 | 0.99483019 |
| E130218I03R | 13.414957 | 0.23559426 | 1.23585451 | 0.19063269 | 0.84881338 | 0.96765585 |
| Cd101 | 133.554947 | -0.0795441 | 0.49201155 | -0.1616713 | 0.87156474 | 0.97288543 |
| Gm15500 | 62441.6866 | 0.32880307 | 0.39217779 | 0.83840307 | 0.40180437 | 0.79752354 |
| Isoc2a | 6.57959423 | -1.1257956 | 1.76005855 | -0.6396353 | 0.52240972 | 0.85816354 |
| Gm13835 | 1298.35955 | 0.8809359 | 0.5956285 | 1.47900227 | 0.13913971 | 0.56286245 |
| Rtl10 | 12.2390397 | -1.6070509 | 1.69103919 | -0.9503333 | 0.34194292 | 0.75874982 |
| Gm8093 | 28.493764 | 0.72202761 | 1.1366729 | 0.63521142 | 0.52529053 | 0.86045516 |
| Plcxd2 | 33.6817955 | 0.80311855 | 0.95815901 | 0.83818921 | 0.40192444 | 0.79752501 |
| BC064078 | 16.3991683 | -0.7101868 | 0.97291829 | -0.7299553 | 0.46541752 | 0.83339036 |
| Gm6483 | 25.8182172 | -0.4226707 | 0.78201801 | -0.5404871 | 0.58886113 | 0.88690478 |
| Gm13147 | 13.6614574 | 0.18952532 | 1.16299872 | 0.16296262 | 0.87054786 | 0.97265735 |
| Lamtor5 | 6681.05196 | 0.51351891 | 0.41243615 | 1.24508706 | 0.21309975 | 0.64599096 |
| Tmem170b | 3063.70805 | -0.0943711 | 0.34017151 | -0.277422 | 0.78145606 | 0.95004728 |
| Frg2f1 | 11.3197424 | 3.10389298 | 1.56001629 | 1.9896542 | 0.04662904 | 0.33435982 |

| | | | | | | |
|---|---|---|---|---|---|---|
| Kantr | 180.212863 | -0.8942282 | 0.50150827 | -1.7830777 | 0.07457365 | 0.42093241 |
| Gm15501 | 1874.39052 | -0.0131082 | 0.36413105 | -0.0359985 | 0.97128356 | 0.99443087 |
| Zfp111 | 421.912454 | -0.0213153 | 0.36451001 | -0.0584765 | 0.95336905 | 0.99149988 |
| 1500009L16I | 12.7569218 | 1.31165566 | 2.11779045 | 0.61935101 | 0.53568514 | 0.86539693 |
| Tmem250-ps | 3691.71267 | 0.44453547 | 0.19523336 | 2.27694418 | 0.02278956 | 0.2163471 |
| Pet100 | 472.415064 | 0.68053796 | 0.5744571 | 1.18466281 | 0.23615079 | 0.67182855 |
| Gm13493 | 22.6218414 | 0.01847636 | 0.95997519 | 0.01924671 | 0.9846443 | 0.99564261 |
| Fcor | 134.945241 | -1.0356504 | 0.61030668 | -1.6969344 | 0.08970905 | 0.45763589 |
| Bcl2l2 | 19.2510073 | 0.45637756 | 1.31003649 | 0.34837011 | 0.72756225 | 0.93576253 |
| Rab42 | 25.487273 | 1.83794453 | 0.84449346 | 2.17638694 | 0.02952634 | 0.25669202 |
| Galnt2 | 609.582546 | 1.00894526 | 0.54563064 | 1.849136 | 0.06443818 | 0.39212252 |
| Cbx6 | 195.023737 | -0.9908646 | 0.41914426 | -2.3640182 | 0.01807793 | 0.19031606 |
| Clec2f | 15.9958112 | -1.9169892 | 1.22470491 | -1.5652662 | 0.1175205 | 0.52372617 |
| Zfp966 | 10.3292679 | 0.63193395 | 1.64984608 | 0.383026 | 0.70170048 | 0.92738786 |
| Gm16580 | 9.02962248 | 0.93252667 | 1.38645331 | 0.67259868 | 0.50120264 | 0.84968141 |
| Tmsb15b1 | 27.0555831 | 0.53142737 | 0.99680591 | 0.53313024 | 0.59394343 | 0.88797037 |
| Slc5a3 | 206.036157 | 0.98470487 | 0.64116849 | 1.53579735 | 0.12458809 | 0.53617731 |
| Rasgef1b | 4990.95927 | 0.13071491 | 0.46378319 | 0.28184486 | 0.77806247 | 0.94837927 |
| Rbm12 | 1900.91101 | 0.17355402 | 0.28513868 | 0.60866529 | 0.54274632 | 0.86903891 |
| Shkbp1 | 1259.53387 | -0.1919316 | 0.26363332 | -0.7280247 | 0.46659849 | 0.83345004 |
| Timm10b | 1755.29603 | 0.36349625 | 0.38415911 | 0.94621274 | 0.34404009 | 0.75962035 |
| Zfp882 | 60.039552 | -0.005812 | 0.62480897 | -0.009302 | 0.99257818 | 0.99796243 |
| Umad1 | 1133.83304 | -0.0059711 | 0.23023204 | -0.0259349 | 0.97930923 | 0.99483019 |
| Rps6kc1 | 709.777967 | 0.19383649 | 0.32672539 | 0.59327036 | 0.55300023 | 0.87131012 |
| Tmem102 | 9.20803267 | 2.63063381 | 1.65096239 | 1.59339415 | 0.1110718 | 0.50888686 |
| Gm8113 | 27.6549884 | 1.63825222 | 1.15581068 | 1.41740533 | 0.15636446 | 0.5856725 |
| Mfsd14a | 3671.96242 | 0.02071402 | 0.17476878 | 0.11852245 | 0.90565371 | 0.98144617 |
| Uckl1 | 1225.29181 | 0.06690261 | 0.23524412 | 0.28439652 | 0.77610652 | 0.94776024 |
| Bcl2a1b | 1024.24955 | 0.09832282 | 0.4193317 | 0.23447504 | 0.81461621 | 0.95994071 |
| Pira2 | 174.823181 | -0.3308011 | 0.56652036 | -0.5839174 | 0.55927588 | 0.87397022 |
| Pakap | 16.0388377 | 2.18084595 | 1.33391815 | 1.63491736 | 0.10206637 | 0.48845074 |
| Gm14435 | 559.358037 | 0.47766283 | 0.50301948 | 0.94959111 | 0.34232006 | 0.75889838 |
| Fbxo24 | 7.47099617 | 1.3368431 | 2.35871416 | 0.56676774 | 0.57087199 | 0.88035336 |
| Ier3ip1 | 2166.4661 | -0.033909 | 0.27709452 | -0.1223735 | 0.90260322 | 0.98043702 |
| Gm15446 | 540.824512 | -0.0345137 | 0.43641394 | -0.0790849 | 0.93696512 | 0.98843518 |
| Gimap1 | 290.663054 | 0.0044489 | 0.49741387 | 0.00894407 | 0.99286376 | 0.99800348 |
| Galnt4 | 316.013573 | -0.1103819 | 0.45799062 | -0.2410134 | 0.80954475 | 0.95844313 |
| Lime1 | 33.4841254 | -0.8735986 | 0.73289905 | -1.1919766 | 0.2332704 | 0.66891108 |
| Rnf8 | 1531.06798 | -0.1639462 | 0.17946899 | -0.9135069 | 0.360976 | 0.76887042 |
| Gm14399 | 60.9609809 | 0.73337766 | 0.86613483 | 0.84672459 | 0.39714865 | 0.79505223 |
| Ttbk2 | 43.9669117 | -0.3492313 | 1.05601214 | -0.3307076 | 0.74086535 | 0.93800224 |
| Cmc4 | 148.291427 | 1.09213281 | 0.54957597 | 1.9872281 | 0.04689713 | 0.3345146 |
| Shprh | 193.801744 | 0.3682206 | 0.39097847 | 0.94179253 | 0.34629887 | 0.7602916 |
| Usp49 | 42.9263257 | 0.54335369 | 0.96697146 | 0.56191285 | 0.5741754 | 0.88059206 |
| Ugt1a7c | 2826.99043 | 0.6941645 | 0.19936849 | 3.48181656 | 0.00049802 | 0.01336039 |
| Gm10177 | 3413.47926 | 0.71492562 | 0.47570111 | 1.50288827 | 0.13286786 | 0.55270061 |

| | | | | | | |
|---|---|---|---|---|---|---|
| Uba52 | 9787.75534 | -0.0381318 | 0.26811501 | -0.1422218 | 0.88690481 | 0.97827644 |
| Acad11 | 450.769241 | -0.0971836 | 0.38000614 | -0.2557423 | 0.79814985 | 0.95433165 |
| BC035044 | 8365.52572 | -0.1769169 | 0.39338958 | -0.4497245 | 0.65290907 | 0.9106891 |
| Fbxw10 | 17.3183141 | -0.1631731 | 2.00937777 | -0.0812058 | 0.93527829 | 0.98794094 |
| Tmem189 | 322.988251 | -0.4481979 | 0.35422009 | -1.265309 | 0.20576057 | 0.63991024 |
| Cfb | 650.860468 | 0.07227148 | 0.45535035 | 0.1587162 | 0.87389247 | 0.97350615 |
| Bloc1s1 | 1583.41386 | 0.57894629 | 0.4123828 | 1.40390505 | 0.16034713 | 0.59011022 |
| Churc1 | 3573.13974 | 0.95039759 | 0.49265603 | 1.92913015 | 0.05371471 | 0.35906667 |
| Mettl23 | 1325.64564 | 0.3815927 | 0.46296443 | 0.82423762 | 0.40980456 | 0.80116089 |
| Mndal | 344.352116 | -0.3307702 | 0.4141534 | -0.7986658 | 0.42448421 | 0.80959079 |
| Tarbp1 | 76.9621343 | -0.0871175 | 0.46671817 | -0.1866597 | 0.85192744 | 0.96798445 |
| Gm6158 | 66.1920298 | -0.8352719 | 1.20060031 | -0.6957118 | 0.4866093 | 0.84183502 |
| 4930523C07 | 110.375971 | -2.2528689 | 0.58843064 | -3.8286056 | 0.00012887 | 0.0044914 |
| Gypc | 1549.57024 | 0.43082894 | 0.22627607 | 1.90399694 | 0.05691058 | 0.3668004 |
| Gm6576 | 12.9331156 | -0.4799753 | 1.14603759 | -0.4188129 | 0.6753529 | 0.91848602 |
| Snrpe | 7062.02238 | 0.19966253 | 0.42829769 | 0.466177 | 0.64108881 | 0.90685007 |
| A930033H14 | 9.09492847 | 1.57277798 | 2.25310968 | 0.69804768 | 0.48514738 | 0.8417344 |
| Tex9 | 92.9253983 | 0.22349023 | 0.54366032 | 0.41108431 | 0.68101071 | 0.92061965 |
| Gm20425 | 19.6056426 | -1.0471066 | 1.95031225 | -0.5368918 | 0.59134238 | 0.88792087 |
| Zfp712 | 28.9722674 | 1.38704163 | 1.22474145 | 1.13251792 | 0.25741676 | 0.69274795 |
| Zfp493 | 82.3008066 | 0.36624876 | 0.51874233 | 0.70603214 | 0.48016815 | 0.83935746 |
| Gad1-ps | 20.9298163 | -1.3293249 | 1.34343389 | -0.9894978 | 0.32241965 | 0.74488377 |
| Olfr111 | 21.5840176 | 3.88912883 | 1.57317091 | 2.47215913 | 0.01342997 | 0.1567552 |
| Apold1 | 60.3192705 | 0.80371334 | 1.38345384 | 0.58094698 | 0.56127619 | 0.8752292 |
| Rps27 | 5934.5193 | 0.45775438 | 0.42765957 | 1.07037095 | 0.28445237 | 0.71998879 |
| Klhl33 | 47.9493853 | 0.44069488 | 1.30149566 | 0.33860649 | 0.73490619 | 0.93750666 |
| Myl6 | 15663.3383 | 0.32186688 | 0.3320566 | 0.9693133 | 0.3323889 | 0.75145322 |
| Rps13 | 10062.3156 | 0.06629319 | 0.38532793 | 0.17204357 | 0.86340328 | 0.9705375 |
| Hspa1b | 890.302457 | -0.3017747 | 0.40810207 | -0.7394588 | 0.45962844 | 0.82987212 |
| Olfr110 | 17.7195283 | 0.35546211 | 1.53734143 | 0.23121871 | 0.81714489 | 0.96079676 |
| Pabpc4l | 195.692724 | 1.33745498 | 0.63625587 | 2.10207096 | 0.03554706 | 0.2827355 |
| Synj2bp | 1340.61615 | -0.0606184 | 0.22594749 | -0.2682851 | 0.78847984 | 0.95225386 |
| Gm17147 | 22.5132035 | 1.35665003 | 0.93936665 | 1.44421779 | 0.14867772 | 0.57574377 |
| Ccdc71l | 107.417552 | -0.3751753 | 0.61621209 | -0.6088412 | 0.54262969 | 0.86903891 |
| Lrrc32 | 21.438846 | 0.72042581 | 1.58714779 | 0.45391224 | 0.64989199 | 0.90973357 |
| Gm17655 | 61.18347 | 1.26450373 | 0.86294888 | 1.46532867 | 0.14283127 | 0.56839782 |
| 1700030C10 | 34.0901088 | -0.0235308 | 0.70841017 | -0.0332163 | 0.97350209 | 0.99443087 |
| Rpl6l | 9.74725218 | -0.2169646 | 1.19844922 | -0.1810378 | 0.85633794 | 0.96924897 |
| Ccdc152 | 141.774713 | 1.55714284 | 0.51566445 | 3.01968238 | 0.0025304 | 0.04723614 |
| Phf11c | 16.5942607 | -0.4779653 | 1.95231067 | -0.2448203 | 0.80659557 | 0.95741157 |
| Gm5141 | 205.926123 | -0.3389393 | 0.51674942 | -0.6559066 | 0.51188424 | 0.85393154 |
| Gm20390 | 3750.97537 | 0.15340211 | 0.31563697 | 0.48600807 | 0.62696144 | 0.90205481 |
| Smim13 | 301.983188 | 0.09151464 | 0.28957779 | 0.31602784 | 0.75198137 | 0.94018119 |
| Gm6682 | 33.7600564 | 0.10270667 | 0.67857541 | 0.15135631 | 0.87969465 | 0.97541902 |
| Eid1 | 301.004049 | -0.566646 | 0.38597126 | -1.4681041 | 0.14207594 | 0.5673724 |
| Gm4202 | 374.191338 | 0.9041956 | 0.47107538 | 1.91942868 | 0.0549301 | 0.36084578 |

| | | | | | | |
|---|---|---|---|---|---|---|
| Gm17084 | 17.8690489 | -0.3146258 | 0.94783852 | -0.3319403 | 0.73993433 | 0.93800224 |
| Gm3739 | 11.6179385 | -0.139866 | 2.20572314 | -0.0634105 | 0.94943962 | 0.99040631 |
| 2610021A01I | 323.735108 | 0.62235134 | 0.37010732 | 1.68154293 | 0.0926575 | 0.46597694 |
| Gm10039 | 23.8011592 | -0.1893037 | 1.25985954 | -0.1502578 | 0.88056123 | 0.97571608 |
| Lamtor3 | 4292.93321 | 0.43569779 | 0.28459688 | 1.53092961 | 0.12578679 | 0.5378361 |
| Tma7 | 7235.94864 | 0.37135816 | 0.38867054 | 0.95545746 | 0.33934641 | 0.75683748 |
| Gm6548 | 37.7867422 | 0.60243559 | 0.92274717 | 0.65287178 | 0.51383894 | 0.85497712 |
| Cyp4f17 | 18.0626197 | 1.24865335 | 1.79906043 | 0.69405859 | 0.48764547 | 0.84304225 |
| Lsm5 | 1827.70725 | 0.18517335 | 0.37231271 | 0.49735974 | 0.61893539 | 0.89879352 |
| Gm3756 | 40.9369633 | 0.24600594 | 0.6419262 | 0.38323087 | 0.70154859 | 0.92738786 |
| Phf11b | 438.844337 | -0.7940543 | 0.3100813 | -2.560794 | 0.01044333 | 0.13211292 |
| Yy2 | 105.697539 | -0.7562522 | 1.24367328 | -0.6080794 | 0.54313479 | 0.86919453 |
| Zfp964 | 8.727786 | -0.1249252 | 2.73946311 | -0.0456021 | 0.96362741 | 0.99401755 |
| Sco2 | 331.211649 | -0.1222305 | 0.26919251 | -0.4540635 | 0.64978314 | 0.90973357 |
| Cox16 | 1261.86975 | 0.30738632 | 0.40658765 | 0.75601489 | 0.44964027 | 0.82484234 |
| Inafm1 | 200.385092 | -0.1213936 | 0.52128775 | -0.2328726 | 0.81586035 | 0.96063519 |
| Gm8251 | 18.8469331 | 1.53043595 | 1.45115371 | 1.05463393 | 0.29159278 | 0.72554658 |
| Rpl36-ps12 | 33.695021 | 0.23418376 | 0.82885216 | 0.28253984 | 0.7775296 | 0.94837927 |
| Ube2d2a | 3094.00033 | 0.03020012 | 0.20038537 | 0.1507102 | 0.88020433 | 0.97549733 |
| Gon7 | 624.455471 | -0.4131562 | 0.21772311 | -1.8976219 | 0.0577459 | 0.36898259 |
| Rps2-ps10 | 232.49665 | -0.1435182 | 0.34998039 | -0.4100751 | 0.68175086 | 0.92061965 |
| Hspa1a | 52308.8659 | 0.02879161 | 0.30208642 | 0.09530917 | 0.92406926 | 0.9849488 |
| BC049352 | 58.3500073 | 1.17067495 | 0.75491372 | 1.55074007 | 0.12096399 | 0.52910763 |
| Gm9027 | 16.2801331 | 0.32092845 | 1.01770053 | 0.31534664 | 0.75249847 | 0.94018119 |
| Peg10 | 23.3751748 | 1.57020688 | 1.56517814 | 1.00321288 | 0.31575816 | 0.74186126 |
| Bend4 | 2741.84287 | 0.30980812 | 0.33150594 | 0.93454772 | 0.35002143 | 0.76216093 |
| Dynlt1a | 1591.21526 | 0.70107729 | 0.35728657 | 1.96222682 | 0.04973608 | 0.34225366 |
| Fancf | 63.1179603 | 0.90065299 | 0.56761236 | 1.58673956 | 0.11257164 | 0.51139203 |
| B930094E09I | 10.9109814 | -0.0401195 | 1.41741131 | -0.0283048 | 0.97741906 | 0.99483019 |
| Gm45351 | 83.8693021 | 0.97771183 | 1.26466333 | 0.77310049 | 0.43946291 | 0.81963526 |
| Dnaaf4 | 17.7197247 | 0.20270612 | 1.08991395 | 0.1859836 | 0.85245762 | 0.96798445 |
| Stk19-ps1 | 30.6119608 | 2.32737878 | 0.77209656 | 3.01436234 | 0.0025752 | 0.04780338 |
| Gm2308 | 15.0159917 | 0.05547209 | 1.01479121 | 0.05466355 | 0.95640651 | 0.99215259 |
| Zfp963 | 218.204045 | 0.37489229 | 0.41332007 | 0.90702658 | 0.36439274 | 0.77081383 |
| Hmgb1-ps7 | 10.2155256 | 0.23737362 | 1.15022424 | 0.2063716 | 0.83650065 | 0.96683245 |
| Gm20388 | 1502.32519 | 0.55432646 | 0.25708705 | 2.15618194 | 0.03106946 | 0.26339065 |
| Platr17 | 74.5188808 | 0.20903543 | 0.5021821 | 0.41625423 | 0.67722398 | 0.91958179 |
| Gm20541 | 32.4012307 | 0.62063404 | 0.88441851 | 0.70174248 | 0.48283977 | 0.84106286 |
| Zfp141 | 456.550578 | 0.43276609 | 0.33290016 | 1.29998763 | 0.19360521 | 0.62506533 |
| Gpank1 | 631.344091 | 0.07666942 | 0.39516701 | 0.19401776 | 0.84616198 | 0.96765585 |
| Gm20482 | 25.0082613 | 2.15865037 | 1.60185005 | 1.34759829 | 0.17778762 | 0.60906934 |
| Gm17907 | 14.298392 | 1.03653089 | 1.7386062 | 0.59618497 | 0.55105166 | 0.87093412 |
| Nlrp1c-ps | 28.1926937 | -0.2379705 | 1.39544838 | -0.1705334 | 0.8645907 | 0.9705375 |
| Gm20422 | 10.5342086 | 1.45218441 | 1.68352037 | 0.86258797 | 0.38836405 | 0.78942657 |
| Med20 | 1406.43358 | 0.0089537 | 0.27052695 | 0.03309726 | 0.97359703 | 0.99443087 |
| Serpinb10 | 146.425008 | -1.6278507 | 0.58324615 | -2.7910183 | 0.00525425 | 0.08309185 |

| | | | | | | |
|---|---|---|---|---|---|---|
| 2810047C21 | 24.3940321 | 0.19459465 | 2.01064988 | 0.09678197 | 0.92289955 | 0.98465273 |
| Scnm1 | 839.215048 | 0.40022711 | 0.22133785 | 1.80821815 | 0.07057256 | 0.41021047 |
| Gm6061 | 47.6620731 | 0.22870642 | 0.74599132 | 0.30658054 | 0.75916267 | 0.94184564 |
| Lrch4 | 4315.87538 | -0.2910892 | 0.22877491 | -1.2723824 | 0.2032373 | 0.63809363 |
| Gm20695 | 75.8222501 | 0.64845218 | 0.53532823 | 1.21131699 | 0.22577394 | 0.65900564 |
| Eif4e3 | 676.128503 | -1.0561655 | 0.37171874 | -2.8413028 | 0.00449296 | 0.07360533 |
| Rpl41 | 24828.4832 | 0.62909618 | 0.52063492 | 1.208325 | 0.22692228 | 0.66015092 |
| Gm20716 | 372.011076 | 0.128888 | 0.39048519 | 0.33007143 | 0.74134599 | 0.93800224 |
| Gm8355 | 66.7979073 | 0.10719392 | 0.50044464 | 0.21419735 | 0.83039316 | 0.96466657 |
| Ppp2r3d | 253.562383 | -0.3151135 | 0.3404923 | -0.9254644 | 0.35472438 | 0.76419569 |
| Tomm20 | 10851.9326 | 0.35225904 | 0.43225597 | 0.81493157 | 0.41511148 | 0.80395688 |
| Hmgcs1 | 5970.27957 | 0.85188793 | 0.17356338 | 4.90822376 | 9.190496290 | 6.732176115 |
| Evi2b | 4032.45291 | -0.5453342 | 0.24976562 | -2.1833839 | 0.02900755 | 0.25346382 |
| Mrgpra2a | 35.8970561 | -1.0820606 | 0.92544409 | -1.1692339 | 0.2423094 | 0.67753686 |
| Gm2237 | 30.1922159 | -1.1966303 | 1.39257961 | -0.8592904 | 0.39018031 | 0.78985201 |
| Rnasek | 2864.50398 | 0.2633238 | 0.17890816 | 1.47183784 | 0.14106467 | 0.5654142 |
| Hist1h2ao | 9.18554208 | -0.3038237 | 1.36706803 | -0.2222448 | 0.82412335 | 0.96290555 |
| Gm11942 | 9.30846197 | -0.4178111 | 1.27661989 | -0.3272791 | 0.74345676 | 0.93856621 |
| Gm38394 | 34.7315956 | -0.4450652 | 1.43474294 | -0.3102055 | 0.75640468 | 0.94102069 |
| Gm9830 | 79.0110178 | 0.66683659 | 0.58068255 | 1.14836685 | 0.25081715 | 0.6842534 |
| Zfp955a | 410.961479 | -0.2352115 | 0.3830655 | -0.6140243 | 0.53919924 | 0.86719392 |
| 9430069I07R | 49.5442566 | -0.7637765 | 0.90659018 | -0.8424716 | 0.39952402 | 0.79612475 |
| Purb | 919.388327 | -0.1353045 | 0.2401111 | -0.563508 | 0.57308902 | 0.88035336 |
| Fsbp | 9.61777786 | 0.85393839 | 1.20848925 | 0.70661646 | 0.47980486 | 0.83933259 |
| Gm3055 | 216.572223 | 1.33316175 | 1.83071357 | 0.72821973 | 0.46647909 | 0.83345004 |
| Gm10359 | 8036.5916 | -0.7046103 | 0.36390922 | -1.9362255 | 0.05284009 | 0.35543223 |
| Rnaset2b | 7495.16243 | 0.08221488 | 0.26018372 | 0.31598781 | 0.75201176 | 0.94018119 |
| Hist1h2ap | 16754.0328 | -0.3464754 | 0.30454565 | -1.1376798 | 0.25525422 | 0.68975587 |
| Gm14403 | 93.4840928 | 0.62455893 | 0.47773381 | 1.30733667 | 0.1910984 | 0.62228856 |
| BC147527 | 260.735036 | 0.42000869 | 0.60960777 | 0.68898185 | 0.49083469 | 0.84477968 |
| Zfp131 | 778.814111 | -0.2842054 | 0.32263631 | -0.8808846 | 0.37838029 | 0.78108777 |
| Rbm4 | 1466.62838 | 0.08969115 | 0.24851821 | 0.36090374 | 0.7181714 | 0.93286094 |
| Gm3604 | 316.536951 | 0.26724435 | 0.42469819 | 0.6292571 | 0.52918075 | 0.86278569 |
| AC149090.1 | 529.792542 | 0.17808829 | 0.3197226 | 0.55700877 | 0.57752144 | 0.88150999 |
| Gm12537 | 43.2821276 | -1.0207691 | 0.66285655 | -1.5399548 | 0.12357138 | 0.53434474 |
| Ccdc85b | 345.579369 | 0.01360121 | 0.25701303 | 0.05292031 | 0.9577954 | 0.99242502 |
| Gm13285 | 13.1776592 | 1.89722957 | 1.73663821 | 1.09247255 | 0.27462545 | 0.71147204 |
| Itpripl2 | 209.343592 | -0.0800387 | 0.43346144 | -0.1846501 | 0.85350351 | 0.96827492 |
| Mid1-ps1 | 35.3141854 | 0.08169601 | 1.15096196 | 0.07098063 | 0.94341317 | 0.98989307 |
| Zfp967 | 114.022131 | 0.52883176 | 0.55485125 | 0.95310545 | 0.34053664 | 0.75821364 |
| Zfp799 | 141.668545 | 1.45020896 | 0.6698184 | 2.16507781 | 0.03038173 | 0.26008374 |
| Gm3488 | 24.0635449 | 1.55311394 | 1.3390951 | 1.15982349 | 0.24612068 | 0.68117517 |
| Gm10130 | 290.560485 | -0.1235252 | 0.47000253 | -0.2628183 | 0.79269066 | 0.95297175 |
| Zfp870 | 258.076259 | -0.258911 | 0.46748571 | -0.5538373 | 0.5796902 | 0.88286755 |
| Gm14325 | 150.82804 | 0.99143133 | 0.53750539 | 1.84450491 | 0.06510959 | 0.39352056 |
| Gm21092 | 50.1823351 | 0.04399766 | 0.74932543 | 0.05871636 | 0.95317803 | 0.99149988 |

| | | | | | | |
|---|---|---|---|---|---|---|
| Rps2-ps6 | 35.9759612 | -0.2232187 | 0.67316667 | -0.331595 | 0.74019513 | 0.93800224 |
| Zfp748 | 347.408104 | -0.3161883 | 0.30095548 | -1.0506147 | 0.29343558 | 0.72646828 |
| Entpd4 | 476.725596 | 0.33399976 | 0.39136402 | 0.8534248 | 0.39342377 | 0.79183169 |
| Noc2l | 657.649821 | -0.7797741 | 0.33579439 | -2.3221772 | 0.0202234 | 0.20113236 |
| Fam177a | 1946.33137 | 0.01373001 | 0.20935106 | 0.06558368 | 0.94770928 | 0.99019648 |
| Gm6472 | 110.934386 | -0.5828401 | 0.96450544 | -0.6042891 | 0.54565145 | 0.86990153 |
| Csta2 | 21.5343828 | 0.47089286 | 1.03164249 | 0.45644965 | 0.64806667 | 0.90973357 |
| Gm2004 | 58.1242661 | -0.9313871 | 0.84119285 | -1.1072218 | 0.26819804 | 0.7046535 |
| Dynlt1f | 6052.84307 | 0.74004055 | 0.37730772 | 1.96137134 | 0.04983572 | 0.34225988 |
| Rnaset2a | 2891.17214 | 0.05642586 | 0.27338426 | 0.20639763 | 0.83648032 | 0.96683245 |
| Rab11b-ps2 | 316.712344 | 0.05198259 | 0.32195106 | 0.16146115 | 0.87173021 | 0.97298136 |
| Nupr1l | 13.0091157 | -2.2400588 | 2.25474465 | -0.9934867 | 0.32047283 | 0.74413737 |
| Gm13237 | 10.7689212 | -0.6128773 | 1.08625772 | -0.5642098 | 0.57261133 | 0.88035336 |
| Nim1k | 68.6020348 | 0.68432735 | 0.72263739 | 0.94698581 | 0.34364601 | 0.75947281 |
| Zfp97 | 553.729091 | 0.22733319 | 0.4729603 | 0.4806602 | 0.63075802 | 0.90373189 |
| 2610528A11l | 69.9513379 | 0.47124086 | 0.51888985 | 0.90817129 | 0.36378773 | 0.77045439 |
| Gm21596 | 317.586736 | -0.2125861 | 0.24995902 | -0.8504838 | 0.39505615 | 0.79305241 |
| Hist4h4 | 15.7944862 | -0.1243634 | 1.2976903 | -0.0958344 | 0.92365209 | 0.98493282 |
| D830030K20 | 15.1829246 | 1.71724042 | 1.99804445 | 0.85946057 | 0.39008647 | 0.78985201 |
| Vkorc1 | 1095.51109 | 0.38424811 | 0.34861872 | 1.10220159 | 0.27037404 | 0.70733226 |
| Cmtm4 | 25.1911796 | -0.8196733 | 1.33167775 | -0.6155193 | 0.53821185 | 0.86663962 |
| Ptrhd1 | 1264.70103 | 0.32382795 | 0.36158626 | 0.89557593 | 0.37047929 | 0.77617284 |
| H1f0 | 2201.88297 | 0.25135158 | 0.22067898 | 1.13899194 | 0.25470652 | 0.68873227 |
| Dynlt1b | 2220.80511 | 0.84951091 | 0.41063586 | 2.06876944 | 0.03856773 | 0.29684673 |
| Gm21992 | 141.941775 | -0.4822405 | 0.50823133 | -0.9488603 | 0.34269168 | 0.75932198 |
| Gm21811 | 97.0713756 | 0.34751642 | 0.45738169 | 0.75979521 | 0.447377 | 0.82297186 |
| Zfp994 | 273.567697 | 1.51079814 | 0.51415103 | 2.93843257 | 0.00329876 | 0.05814666 |
| Moap1 | 381.329098 | -0.0235534 | 0.29046173 | -0.0810896 | 0.93537068 | 0.98794094 |
| Cdkn2d | 1284.24372 | -0.0960989 | 0.32322998 | -0.2973082 | 0.76623124 | 0.94360659 |
| Gm4617 | 30.5342456 | 0.20088214 | 0.68727885 | 0.29228622 | 0.77006779 | 0.94429023 |
| Gm15801 | 412.626836 | 0.62709235 | 0.28195101 | 2.22411813 | 0.0261405 | 0.23687163 |
| Tpbgl | 210.62431 | 0.77309578 | 0.67188235 | 1.15064159 | 0.24987972 | 0.6842534 |
| Gm5559 | 42.241509 | -1.0637945 | 0.64931306 | -1.6383383 | 0.10135114 | 0.48633809 |
| Mfsd4b4 | 85.7489595 | 0.09876881 | 0.75447924 | 0.13090992 | 0.89584657 | 0.97984623 |
| Zfp960 | 254.196966 | 0.59996498 | 0.62224646 | 0.96419188 | 0.33494973 | 0.75359284 |
| Zfp781 | 136.883946 | 0.84653461 | 0.68445484 | 1.23680127 | 0.21616087 | 0.64982959 |
| Mrgpra2b | 12.1992913 | -3.7601774 | 1.22508925 | -3.0693089 | 0.00214555 | 0.04220001 |
| Psmb9 | 1428.7029 | 0.20043292 | 0.35492513 | 0.56471886 | 0.572265 | 0.88035336 |
| Lbhd1 | 1127.92527 | -0.0823686 | 0.33242474 | -0.2477812 | 0.80430372 | 0.9570372 |
| Tmem181b-p | 97.0363771 | 1.04345803 | 0.69166489 | 1.50861789 | 0.13139646 | 0.54970237 |
| Zfp433 | 241.933191 | 0.34850174 | 0.38493663 | 0.90534833 | 0.36528088 | 0.77135871 |
| Gm10481 | 24.4526499 | -0.2943349 | 0.7683806 | -0.3830587 | 0.70167625 | 0.92738786 |
| Gm21748 | 27.6783224 | 1.29163044 | 0.71714621 | 1.80106987 | 0.07169187 | 0.41193359 |
| Gm10778 | 347.875078 | 0.04834408 | 0.3777583 | 0.12797623 | 0.89816778 | 0.97989497 |
| Zfp955b | 379.197963 | -0.5029363 | 0.39959019 | -1.2586304 | 0.20816387 | 0.6418276 |
| Zfp850 | 198.933173 | -0.6201932 | 0.31012055 | -1.9998457 | 0.04551693 | 0.3293163 |

| | | | | | | |
|---|---|---|---|---|---|---|
| Rps19-ps6 | 11.533653 | 0.34055201 | 1.2173966 | 0.27973793 | 0.77967857 | 0.9488468 |
| Cdiptos | 30.2245684 | -0.028767 | 0.78807582 | -0.0365028 | 0.97088141 | 0.99443087 |
| Gm3839 | 38.8630204 | -0.3953629 | 0.61732151 | -0.6404489 | 0.52188081 | 0.85816354 |
| Gm27029 | 313.272642 | -0.5658336 | 0.32129412 | -1.7611078 | 0.07822017 | 0.43258005 |
| Rhox5 | 128.169545 | 0.38041256 | 0.66390526 | 0.57299224 | 0.56664995 | 0.87883863 |
| E030030I06R | 135.730013 | 0.59269999 | 0.44115609 | 1.34351538 | 0.17910515 | 0.60992118 |
| Zfp87 | 485.910616 | -0.0616716 | 0.25885046 | -0.238252 | 0.81168565 | 0.95967622 |
| C920009B18 | 501.741807 | -0.0147929 | 0.25372783 | -0.0583023 | 0.9535078 | 0.99149988 |
| Gm3200 | 22.6413821 | -0.1399207 | 0.77678454 | -0.1801281 | 0.85705198 | 0.96936093 |
| Tmem134 | 1050.61202 | 0.23382108 | 0.22078029 | 1.0590668 | 0.28956936 | 0.72395811 |
| 1810014B01I | 436.788353 | -0.2372719 | 0.43925446 | -0.5401695 | 0.58908014 | 0.88691905 |
| Ptges3l | 311.910436 | 0.44402903 | 0.47106517 | 0.94260636 | 0.34588229 | 0.7602916 |
| Gm4349 | 55.5455265 | 1.49134209 | 0.82928311 | 1.79835096 | 0.07212141 | 0.41303124 |
| Gm5523 | 9.10277342 | -0.0152676 | 1.15186411 | -0.0132547 | 0.9894246 | 0.99650865 |
| Gpr137b-ps | 369.124552 | 1.33048928 | 0.44900152 | 2.96321774 | 0.00304441 | 0.05460745 |
| Myadm.1 | 382.182358 | -0.175519 | 0.46389436 | -0.3783598 | 0.70516334 | 0.92879471 |
| Gm3650 | 20.9657018 | -0.3520216 | 0.82391996 | -0.4272522 | 0.66919566 | 0.91643685 |
| Gm9625 | 17.3140744 | -0.3351849 | 1.03726539 | -0.3231429 | 0.74658705 | 0.93960898 |
| Gm4691 | 34.1065859 | -0.1926853 | 0.65330019 | -0.2949414 | 0.76803864 | 0.94401697 |
| Gm4335 | 16.6858942 | 0.01815587 | 0.90703157 | 0.02001681 | 0.98402997 | 0.99564261 |
| Gm8565 | 8.6903167 | -0.3426395 | 1.21884523 | -0.2811182 | 0.77861976 | 0.94853305 |
| Gm2546 | 17.1451891 | -0.4628955 | 0.87890953 | -0.5266703 | 0.5984226 | 0.889626 |
| Zfp82 | 106.881579 | -0.3278638 | 1.19869673 | -0.2735169 | 0.78445589 | 0.95094799 |
| Gm6981 | 15.1137087 | -0.3343129 | 1.14586438 | -0.2917561 | 0.77047313 | 0.94454775 |
| Gm26992 | 11.93062 | 0.73981891 | 1.24174803 | 0.59578827 | 0.55131668 | 0.87093412 |
| 2700099C18 | 171.539446 | 0.04464369 | 0.40712274 | 0.10965658 | 0.91268174 | 0.98292561 |
| Gm16470 | 17.3751048 | 0.12643947 | 0.88549108 | 0.14279023 | 0.88645586 | 0.97817197 |
| Gm4654 | 27.424312 | -0.6918372 | 0.72604624 | -0.9528832 | 0.34064926 | 0.75821364 |
| Bin2 | 14232.574 | -0.6563285 | 0.19457564 | -3.3731277 | 0.00074319 | 0.01855409 |
| Gm2445 | 10.7455336 | -0.0941148 | 1.1439709 | -0.0822703 | 0.93443178 | 0.98756396 |
| Gm4518 | 55.4825793 | -0.1436952 | 0.56538355 | -0.2541553 | 0.79937562 | 0.95514377 |
| Rnf113a2 | 408.709953 | -0.1692206 | 0.2739954 | -0.6176038 | 0.53683656 | 0.86596304 |
| Ccdc166 | 100.932764 | -0.1052818 | 0.50966706 | -0.2065697 | 0.83634594 | 0.96683245 |
| Sowahc | 430.851947 | -0.3818622 | 0.38960772 | -0.9801196 | 0.32702708 | 0.74811039 |
| Gm3534 | 8.83797314 | 0.03298942 | 1.28324172 | 0.02570787 | 0.97949034 | 0.99483957 |
| Rpl24 | 12556.305 | 0.13008589 | 0.35607157 | 0.36533637 | 0.71486033 | 0.93235217 |
| Gm28043 | 409.778299 | 0.232662 | 0.32773484 | 0.70990925 | 0.47776041 | 0.83731277 |
| Kctd12 | 2079.87371 | 0.37543424 | 0.2530763 | 1.48348242 | 0.13794628 | 0.56068131 |
| Gm28048 | 460.105123 | 1.16872622 | 0.81927848 | 1.42653109 | 0.1537151 | 0.58242478 |
| Gm28041 | 13.8311036 | -0.6339317 | 1.44326956 | -0.4392331 | 0.66049267 | 0.91302324 |
| Rps2-ps5 | 14.7243744 | -0.3198009 | 1.05688509 | -0.3025881 | 0.76220376 | 0.94298237 |
| Gm28044 | 12.5406048 | 1.04916022 | 1.61642005 | 0.6490641 | 0.51629695 | 0.85627177 |
| Jmjd7 | 633.145708 | 0.53365757 | 0.34834319 | 1.53198797 | 0.1255254 | 0.53724874 |
| Zfp953 | 62.4488781 | 0.12464392 | 0.70182379 | 0.17760003 | 0.8590371 | 0.96954641 |
| Rpl15-ps2 | 12.4552388 | 0.6254702 | 0.9973681 | 0.62712072 | 0.53058011 | 0.86356925 |
| Tmem185b | 1158.76556 | -0.3235947 | 0.30939161 | -1.0459066 | 0.29560414 | 0.72852118 |

| | | | | | | |
|---|---|---|---|---|---|---|
| Gm18853 | 50.4128255 | -0.2020108 | 0.99756609 | -0.2025036 | 0.83952301 | 0.96757914 |
| Gm28036 | 1180.23234 | -0.0384766 | 0.34850999 | -0.1104031 | 0.91208966 | 0.98292561 |
| 2810039B14l | 64.4577037 | 0.75970921 | 0.63891414 | 1.18906308 | 0.23441485 | 0.67038085 |
| Atf7 | 459.325612 | 0.62262166 | 0.36470597 | 1.707188 | 0.08778711 | 0.45274021 |
| Gm18852 | 31.9571728 | -0.8140229 | 1.4112289 | -0.5768185 | 0.56406209 | 0.87688035 |
| Gm18180 | 49.6810076 | 0.72477543 | 0.68021748 | 1.06550545 | 0.28664726 | 0.72260415 |
| Ms4a14 | 18.7856099 | 2.03651378 | 1.90474903 | 1.06917697 | 0.28498994 | 0.72045503 |
| Xndc1 | 342.578931 | -0.5958524 | 0.40337731 | -1.4771589 | 0.13963304 | 0.56329174 |
| Hist1h3g | 16.1075396 | -0.2687522 | 1.09218177 | -0.2460691 | 0.80562874 | 0.95737871 |
| Hist1h3d | 37.6639409 | 0.02651468 | 0.79924673 | 0.03317458 | 0.97353537 | 0.99443087 |
| 2310034G01 | 11.1600706 | -0.6680737 | 1.39396587 | -0.4792612 | 0.63175283 | 0.90399255 |
| Zfp383 | 30.8008186 | 0.84567981 | 1.37467382 | 0.6151858 | 0.53843202 | 0.86663962 |
| Bcl2a1d | 205.839922 | -0.9458552 | 0.61718123 | -1.5325404 | 0.12538914 | 0.53724874 |
| Gm8337 | 25.9583045 | 0.04102014 | 0.74824458 | 0.05482183 | 0.95628041 | 0.99215259 |
| Gm6170 | 31.199576 | 0.23246614 | 0.78723556 | 0.29529426 | 0.7677691 | 0.94401697 |
| Ppp1ccb | 61.347425 | -0.9155081 | 3.72835328 | -0.2455529 | 0.80602832 | 0.95741157 |
| Hist1h3f | 16.2366864 | -1.3043484 | 1.24495735 | -1.0477053 | 0.29477441 | 0.72759794 |
| Gm28557 | 80.9374441 | 0.56796 | 0.55491609 | 1.02350611 | 0.30606861 | 0.73554152 |
| Gm13283 | 10.4402932 | -1.2926272 | 1.75286172 | -0.7374382 | 0.46085589 | 0.82987212 |
| Rps23-ps1 | 9162.11404 | 0.16159126 | 0.38314682 | 0.42174763 | 0.67320924 | 0.91786342 |
| Gm15459 | 420.145752 | 0.21807727 | 0.30319642 | 0.71926069 | 0.47198031 | 0.83435479 |
| Gm29094 | 12.9888791 | -0.3612288 | 1.35920023 | -0.2657657 | 0.79041966 | 0.95261385 |
| Gm9320 | 22.8498859 | 1.01482601 | 1.10782953 | 0.91604889 | 0.35964126 | 0.76857232 |
| Gm29666 | 342.322026 | 0.49368815 | 0.50123521 | 0.98494308 | 0.32465203 | 0.74601961 |
| Gm28437 | 53726.3136 | 0.05630075 | 0.24640475 | 0.22848888 | 0.8192662 | 0.96116653 |
| Gm17971 | 14.1267839 | 0.96044721 | 1.2792614 | 0.75078261 | 0.4527835 | 0.82610332 |
| Eif4a-ps4 | 344.539292 | 0.88413561 | 0.69751436 | 1.26755184 | 0.20495804 | 0.63911591 |
| Gm29216 | 5940.50657 | -0.1005244 | 0.31091686 | -0.3233161 | 0.74645585 | 0.93960567 |
| Ms4a4a | 2085.23204 | 1.80268637 | 0.25402917 | 7.09637551 | 1.280712039 | 3.730226279 |
| Gm7901 | 11.8590103 | 0.27337643 | 1.12466082 | 0.24307456 | 0.80794764 | 0.95809696 |
| Gm29609 | 59.9783507 | 0.15120231 | 0.77128496 | 0.19603949 | 0.84457925 | 0.96765585 |
| 9130401M01 | 1051.68007 | -0.3430132 | 0.27380214 | -1.2527775 | 0.21028668 | 0.64378984 |
| Gm28438 | 391.819533 | -0.7575654 | 2.00239007 | -0.3783306 | 0.70518502 | 0.92879471 |
| Bcl2a1a | 78.8237926 | -0.6105234 | 0.51800368 | -1.1786082 | 0.23855421 | 0.67452555 |
| Zbed6 | 19.0063798 | -0.542134 | 1.92424893 | -0.281738 | 0.77814445 | 0.94837927 |
| Gm20257 | 18.3557405 | 1.57953139 | 1.3712691 | 1.15187557 | 0.24937221 | 0.68409835 |
| Gm28661 | 3111.4037 | 0.37296717 | 0.42780923 | 0.87180721 | 0.38331356 | 0.7864936 |
| Snrpn | 629.155709 | -0.1944447 | 0.23917806 | -0.8129706 | 0.41623492 | 0.80440685 |
| Cd28.1 | 29.9807346 | -0.7518263 | 0.94595618 | -0.7947791 | 0.42674198 | 0.81021713 |
| Sh2d1b1 | 120.015471 | -1.8161228 | 0.58465737 | -3.1063027 | 0.00189443 | 0.0386242 |
| Pcdhga12 | 20.0429493 | 1.17725647 | 1.75059871 | 0.67248791 | 0.50127313 | 0.84968141 |
| Rpl21-ps1 | 21.5132363 | 0.32899507 | 0.94448075 | 0.34833433 | 0.72758912 | 0.93576253 |
| Thap6 | 266.448753 | -0.8738315 | 0.52549304 | -1.6628792 | 0.09633662 | 0.47596359 |
| Pcdhga11 | 19.9943972 | -0.4292573 | 1.43505538 | -0.2991224 | 0.76484663 | 0.94318234 |
| Pcdhgb2 | 17.3493752 | 3.00713013 | 2.05171143 | 1.46566914 | 0.14273845 | 0.56834178 |
| Gm7694 | 26.0882827 | -1.2810446 | 1.51727247 | -0.8443076 | 0.39849755 | 0.79557554 |

| | | | | | | |
|---|---|---|---|---|---|---|
| Gm8242 | 12.7891469 | 1.02889097 | 1.1023205 | 0.93338641 | 0.3506205 | 0.76238338 |
| Pcdhgc3 | 76.3677124 | 2.3515749 | 0.67813322 | 3.46771819 | 0.0005249 | 0.01398926 |
| Zc3h11a | 943.695962 | -0.2668659 | 0.2962788 | -0.9007254 | 0.36773432 | 0.77386646 |
| Gm8797 | 50.752447 | 0.10289089 | 0.59245574 | 0.17366849 | 0.862126 | 0.9705375 |
| Gm38158 | 58.1887971 | -0.3688022 | 0.58542568 | -0.6299727 | 0.52871242 | 0.8623652 |
| Pcdhga5 | 20.6781024 | -0.629943 | 1.81757532 | -0.3465843 | 0.72890368 | 0.93617556 |
| Gm18445 | 6.49009994 | -5.1054972 | 2.02738907 | -2.5182622 | 0.01179355 | 0.14331975 |
| Tstd1 | 27.4771956 | 0.42061976 | 0.82469266 | 0.51003213 | 0.61002895 | 0.89508835 |
| Tigd5 | 15.7814042 | 0.48533579 | 1.72490546 | 0.2813695 | 0.77842701 | 0.94853305 |
| Gm17771 | 12.8296584 | -0.1922761 | 0.97674922 | -0.1968531 | 0.84394252 | 0.96765585 |
| Pcdhgb7 | 22.1927399 | 4.2034988 | 1.49925528 | 2.80372452 | 0.0050516 | 0.0804639 |
| Ighd | 63.9286483 | 0.84397685 | 0.55576186 | 1.51859441 | 0.12886462 | 0.54471351 |
| Gm37988 | 13.9679544 | -0.3550041 | 1.47646469 | -0.240442 | 0.80998761 | 0.95882179 |
| Rpl18-ps1 | 9.93308456 | 0.05575285 | 1.16568313 | 0.04782847 | 0.96185294 | 0.99369539 |
| Gm37164 | 22.9116718 | 0.19357942 | 0.90978807 | 0.21277419 | 0.8315031 | 0.96506427 |
| Gm5869 | 37.4470006 | 0.05667699 | 0.63329489 | 0.08949542 | 0.92868819 | 0.986679 |
| Gm6560 | 43.2980157 | -0.1034596 | 0.57865644 | -0.1787928 | 0.85810041 | 0.96944425 |
| Gm43064 | 13.1471429 | -0.3767861 | 1.5514456 | -0.2428613 | 0.80811285 | 0.95809696 |
| Gm7285 | 9.79006433 | 1.34396124 | 1.7826276 | 0.75392148 | 0.45089637 | 0.82549763 |
| Gbp5 | 16.6999419 | 0.36322763 | 1.30502063 | 0.27833095 | 0.78075832 | 0.94978287 |
| Gm6204 | 9.73060813 | 0.47704964 | 1.39136806 | 0.34286373 | 0.73170098 | 0.93658032 |
| Gm4332 | 133.884844 | 0.00917669 | 0.49070735 | 0.01870094 | 0.98507968 | 0.99564261 |
| Gm20768 | 57.0048067 | 0.10568411 | 0.69653273 | 0.15172885 | 0.87940081 | 0.97541902 |
| Gm2451 | 52.0643925 | 0.32949112 | 0.71212326 | 0.46268833 | 0.64358779 | 0.90823885 |
| Tmem265 | 25.8993776 | -1.8188688 | 2.32355792 | -0.7827947 | 0.43374769 | 0.81483708 |
| Ube2n-ps1 | 1982.80341 | 0.20483979 | 0.28077328 | 0.72955585 | 0.46566171 | 0.83339036 |
| Gtf3c2 | 3821.32016 | -0.1232295 | 0.24394178 | -0.5051595 | 0.6134468 | 0.89764291 |
| Mrpl33 | 2429.28238 | 0.12709742 | 0.46853048 | 0.2712682 | 0.78618476 | 0.95184068 |
| Rpl7-ps7 | 9.51343637 | 0.81416159 | 1.32594896 | 0.61402182 | 0.53920091 | 0.86719392 |
| 0610012G03 | 3341.51672 | 0.48098947 | 0.2150175 | 2.2369782 | 0.02528776 | 0.23122295 |
| Gm42715 | 608.247332 | -0.0911797 | 0.27260138 | -0.3344801 | 0.73801728 | 0.93800224 |
| Gm42742 | 1483.59626 | -0.0524862 | 0.19680812 | -0.266687 | 0.78971014 | 0.95255415 |
| Gm9794 | 26.466775 | 0.5532692 | 0.91852261 | 0.60234685 | 0.54694328 | 0.86994632 |
| Mpv17 | 2133.80248 | 0.74104585 | 0.29534935 | 2.50904851 | 0.01210569 | 0.14604423 |
| Gm4366 | 38.8301262 | -0.4250523 | 0.77624664 | -0.5475737 | 0.58398463 | 0.88404713 |
| Zfp862-ps | 31.5781265 | -0.1403829 | 1.35953044 | -0.1032583 | 0.91775794 | 0.98390186 |
| Gm45062 | 67.3487723 | 1.37738977 | 1.44817671 | 0.95111996 | 0.34154348 | 0.75855146 |
| Gm16042 | 11.2789864 | -1.2662093 | 1.51161755 | -0.8376519 | 0.40222624 | 0.79786503 |
| Gm2011 | 23.5591314 | 0.23205062 | 1.20012295 | 0.1933557 | 0.84668041 | 0.96765585 |
| Gm44126 | 11.2952248 | -1.5214714 | 1.99727753 | -0.7617727 | 0.44619569 | 0.82166369 |
| Gm18584 | 10.6008762 | -0.2107225 | 1.06098448 | -0.1986103 | 0.84256757 | 0.96765585 |
| Gm45140 | 240.803196 | 0.88273612 | 0.32372037 | 2.72684765 | 0.00639426 | 0.09644997 |
| Gm6210 | 151.024908 | -0.0450489 | 0.38398302 | -0.11732 | 0.90660648 | 0.98163543 |
| Gm4045 | 11.88863 | 0.37313394 | 1.06154509 | 0.35150079 | 0.72521268 | 0.93542036 |
| Prkcz2 | 13.9824239 | -0.9820941 | 1.75168308 | -0.5606574 | 0.57503111 | 0.88059206 |
| Gm5586 | 10.526242 | 0.9716466 | 1.20138803 | 0.80877 | 0.41864746 | 0.8052381 |

| | | | | | | |
|---|---|---|---|---|---|---|
| Gm49368 | 35.2188352 | -2.8910673 | 1.79386003 | -1.611646 | 0.107039 | 0.49996493 |
| Rpl19-ps9 | 8.98688045 | -0.0403034 | 1.23852498 | -0.0325414 | 0.97404029 | 0.99444457 |
| Gm9354 | 10.9000005 | -0.1630723 | 1.12665462 | -0.1447403 | 0.88491593 | 0.97744258 |
| Gm44973 | 27.7338186 | 0.57743609 | 0.78522077 | 0.73538056 | 0.46210776 | 0.83095167 |
| 4930431P19I | 18.3884627 | 1.39882478 | 0.94136375 | 1.48595564 | 0.13729085 | 0.55981285 |
| Prmt1 | 3305.90552 | -0.2243908 | 0.26986624 | -0.8314888 | 0.40569756 | 0.79982698 |
| Samd4b | 290.297388 | -0.1205485 | 0.42686167 | -0.2824066 | 0.77763176 | 0.94837927 |
| Gm49396 | 38.1424871 | 0.4335783 | 1.6082429 | 0.26959752 | 0.78746991 | 0.95225386 |
| Nup62 | 7563.19308 | -0.2197851 | 0.2179011 | -1.008646 | 0.31314444 | 0.74065408 |
| Gdf1 | 27.7337457 | 0.76634692 | 1.40727125 | 0.54456234 | 0.58605457 | 0.88518071 |
| Nxpe1-ps | 27.2565098 | -0.3083869 | 0.96311137 | -0.3201986 | 0.74881779 | 0.9399191 |
| Pvrig | 90.4050278 | 0.27636842 | 0.6493268 | 0.425623 | 0.67038258 | 0.91711586 |
| Gm18194 | 27.1690372 | 0.44164274 | 0.94046296 | 0.46960141 | 0.63863981 | 0.90615794 |
| Gm35315 | 180.723561 | 0.5579967 | 0.67762922 | 0.8234543 | 0.41024969 | 0.80117888 |
| Hspa14 | 6463.49201 | -0.1131515 | 0.19956242 | -0.5669978 | 0.57071569 | 0.88035336 |
| Chmp1b | 1229.65111 | -0.1345136 | 0.21114412 | -0.6370702 | 0.52407913 | 0.8594073 |
| Gm45808 | 11.3403602 | -0.1377591 | 2.08124158 | -0.0661908 | 0.94722592 | 0.99019648 |
| Brd3os | 50.0466317 | 0.51822353 | 0.8139308 | 0.63669237 | 0.52432524 | 0.85953586 |
| Gm19196 | 12.4480916 | 1.11624014 | 1.18134172 | 0.94489183 | 0.3447141 | 0.75979957 |
| Gm45844 | 29.7585119 | -0.0011929 | 0.95204437 | -0.001253 | 0.99900027 | 0.99969186 |
| Gm4032 | 9.38358309 | 0.4286633 | 1.63806904 | 0.26168817 | 0.79356186 | 0.95310033 |
| Pde2a | 86.7405018 | 2.06499445 | 1.53040182 | 1.34931521 | 0.17723574 | 0.60816966 |
| Flt3l | 84.6914291 | -0.0780885 | 0.49531837 | -0.157653 | 0.8747302 | 0.97366465 |
| Gm45799 | 74.380212 | 0.02597571 | 0.75118211 | 0.03457977 | 0.97241483 | 0.99443087 |
| Gm45871 | 231.548527 | 0.03832056 | 0.64465311 | 0.05944369 | 0.95259871 | 0.9914597 |
| Gm45716 | 296.866303 | 0.95421743 | 0.36812816 | 2.59207945 | 0.00953977 | 0.12534914 |
| Gm9619 | 10.0070541 | -0.5004996 | 1.10700311 | -0.4521212 | 0.65118168 | 0.91021183 |
| Gm10033 | 191.764534 | -0.0704264 | 0.4905067 | -0.1435789 | 0.885833 | 0.97792574 |
| Gm10358 | 30.3895355 | -0.6703288 | 0.86919833 | -0.7712035 | 0.44058633 | 0.81963526 |
| Gm45855 | 10.229699 | -0.2446505 | 1.27883964 | -0.1913066 | 0.84828539 | 0.96765585 |
| Gm46223 | 170.88506 | -0.6169403 | 0.43627585 | -1.414106 | 0.15733077 | 0.58620126 |
| Ifi30 | 7951.63647 | -0.0144389 | 0.20698439 | -0.0697582 | 0.94438609 | 0.99011613 |
| Ptpmt1 | 873.803909 | 0.83166798 | 0.41145256 | 2.02129739 | 0.04324899 | 0.31923815 |
| Nudt8 | 216.857024 | 0.36569806 | 0.54225948 | 0.6743968 | 0.50005907 | 0.84914912 |
| Gm2981 | 41.7558872 | 1.77881677 | 3.73125724 | 0.47673389 | 0.63355163 | 0.90455392 |
| Btbd8 | 29.2157554 | 0.71069509 | 0.90719497 | 0.7833984 | 0.43339319 | 0.81476855 |
| Gm38431 | 437.280486 | -0.2815251 | 0.35699835 | -0.7885894 | 0.43035205 | 0.81242232 |
| AEKR010771! | 117.2311 | 0.00513414 | 0.40891804 | 0.01255544 | 0.98998247 | 0.99650865 |
| Gm3365 | 27.2400144 | -0.4413071 | 0.93865347 | -0.4701491 | 0.6382485 | 0.90599527 |
| Gm4977 | 14.2404062 | 1.00776942 | 0.99775755 | 1.01003437 | 0.31247882 | 0.74051791 |
| Gm49373 | 7.54600055 | 0.39003259 | 1.43374867 | 0.27203693 | 0.78559362 | 0.95168899 |
| Gm6477 | 122.932932 | -0.3445808 | 0.37221493 | -0.9257577 | 0.35457194 | 0.76419569 |
| Gm2436 | 21.1553593 | -1.1982637 | 1.90090157 | -0.630366 | 0.52845517 | 0.86217134 |
| Lilr4b | 193.335575 | 1.01431438 | 0.37651557 | 2.69395073 | 0.00706106 | 0.10256275 |
| Gm47924 | 7.71568827 | -1.7654341 | 2.10049252 | -0.8404858 | 0.40063608 | 0.79698916 |
| Pbld1 | 26.963345 | 0.30815709 | 0.85057921 | 0.36229088 | 0.71713467 | 0.93259832 |

| | | | | | | |
|---|---|---|---|---|---|---|
| Lilrb4a | 964.972162 | 0.71957311 | 0.33343069 | 2.15808901 | 0.03092091 | 0.26267744 |
| BC024063 | 125.099016 | 0.56273839 | 0.54926954 | 1.0245214 | 0.30558907 | 0.73530116 |
| Srp54b | 3543.80104 | -0.1267781 | 0.24192384 | -0.5240415 | 0.60024967 | 0.8902352 |
| Gm7476 | 10.7695252 | 0.15866644 | 1.1144296 | 0.14237457 | 0.88678415 | 0.97827644 |
| Gm32687 | 265.933377 | 1.46612225 | 0.47369076 | 3.09510415 | 0.00196744 | 0.03958498 |
| Gm17794 | 8.05464774 | -0.4857986 | 1.22182391 | -0.3976011 | 0.69092423 | 0.92470267 |
| Gm4739 | 8160.9426 | 0.0539237 | 0.31804438 | 0.16954773 | 0.86536584 | 0.9705375 |
| Gm9118 | 9.89376852 | -1.3941667 | 1.11212096 | -1.2536107 | 0.20998354 | 0.64350283 |
| Gm7172 | 36.3186562 | -0.0783685 | 0.62859376 | -0.1246727 | 0.90078266 | 0.98043702 |
| Gm11361 | 73.4163078 | 0.55718752 | 0.68367074 | 0.81499395 | 0.41507577 | 0.80395688 |
| Gm49383 | 299.221069 | -0.2219668 | 0.2794892 | -0.7941875 | 0.42708631 | 0.81021713 |
| Gm9973 | 20.2414681 | 0.69084075 | 1.20670034 | 0.57250399 | 0.56698059 | 0.87918285 |
| Tes3-ps | 18.8750493 | 0.16451377 | 0.83127467 | 0.19790543 | 0.84311906 | 0.96765585 |
| Gm48551 | 82.930659 | -0.0346618 | 0.48150101 | -0.071987 | 0.94261225 | 0.98983253 |
| Gm49328 | 8.97319786 | -0.0456153 | 1.15909901 | -0.0393541 | 0.96860809 | 0.99443087 |
| Gm49359 | 539.915837 | 0.4377864 | 0.4906218 | 0.89230931 | 0.37222716 | 0.77730538 |
| Gm49327 | 9.04790874 | 0.17542576 | 1.3699619 | 0.12805156 | 0.89810817 | 0.97989497 |
| Gm48035 | 24.8490739 | 2.80250177 | 1.28218348 | 2.18572599 | 0.02883566 | 0.25269424 |
| Gm3379 | 21.9556438 | -0.0574098 | 0.86896468 | -0.0660668 | 0.9473246 | 0.99019648 |
| Gm7240 | 200.289301 | -0.0674958 | 0.50545246 | -0.1335354 | 0.89376995 | 0.97941957 |
| Adat3 | 233.034813 | -0.7417488 | 0.36266024 | -2.0452996 | 0.04082533 | 0.30771181 |
| Gm49686 | 23.0454335 | 0.30433787 | 1.19391815 | 0.25490681 | 0.79879508 | 0.95472986 |
| Gm38699 | 62.7320198 | 0.02171219 | 0.55669851 | 0.03900171 | 0.96888903 | 0.99443087 |
| Gm49384 | 351.509425 | -0.4292693 | 0.27247424 | -1.5754492 | 0.11515279 | 0.51884494 |
| Gm3325 | 391.689197 | 0.12151155 | 0.40924097 | 0.29691931 | 0.76652811 | 0.94363877 |
| Rpl17-ps3 | 22.5291031 | 1.0040905 | 1.02100499 | 0.98343348 | 0.32539414 | 0.74695937 |
| Scamp4 | 4926.22244 | 0.02078218 | 0.23400015 | 0.08881267 | 0.92923078 | 0.98669576 |
| Gm49331 | 67.7330623 | 0.14999854 | 0.47564438 | 0.31535858 | 0.75248941 | 0.94018119 |
| Gm20075 | 205.81412 | -1.0388907 | 0.38720977 | -2.6830178 | 0.00729611 | 0.10438985 |
| Gm36298 | 66.6472699 | 0.04224237 | 0.92073671 | 0.04587888 | 0.96340679 | 0.99401755 |
| Gm7143 | 25.2715025 | -0.0217418 | 0.76556221 | -0.0283998 | 0.97734327 | 0.99483019 |
| Hist1h2bm | 17.8914641 | 0.28316849 | 1.04342886 | 0.27138266 | 0.78609674 | 0.95184068 |
| Gm49391 | 43.2102627 | 3.39077828 | 1.31900542 | 2.57070838 | 0.01014907 | 0.13017076 |
| Hist1h2bh | 17.3887618 | 1.22044592 | 1.42754874 | 0.85492417 | 0.39259313 | 0.79094067 |
| Gm3226 | 51.126905 | 0.05802461 | 0.57537712 | 0.10084623 | 0.91967253 | 0.9839023 |
| Gm49392 | 13.1050722 | 0.81442593 | 2.03469763 | 0.40026878 | 0.68895856 | 0.92470267 |
| Gm49336 | 2993.42678 | -0.19222 | 0.17564226 | -1.0943836 | 0.27378679 | 0.71078816 |
| Gm49345 | 67.691685 | -0.6575612 | 0.54281165 | -1.2113985 | 0.22574273 | 0.65900564 |
| AL732309.1 | 48.2023318 | -0.5360391 | 1.27538033 | -0.4202975 | 0.67426817 | 0.91822108 |
| Gm48909 | 9.97346507 | 0.01612957 | 1.24065044 | 0.0130009 | 0.98962707 | 0.99650865 |
| Gm16374 | 41.6177556 | -0.0731405 | 0.66192493 | -0.1104966 | 0.91201553 | 0.98292561 |
| Eef1akmt4 | 2331.51424 | 0.13420398 | 0.32211666 | 0.41663161 | 0.67694789 | 0.91950697 |
| Gm49273 | 17.8395821 | -1.2241052 | 1.04621948 | -1.1700272 | 0.24199004 | 0.677251 |
| Pnp | 9818.13416 | 0.30915794 | 0.20375792 | 1.51728065 | 0.12919584 | 0.54535981 |
| AL731706.1 | 901.147978 | 0.81050073 | 0.47509386 | 1.70598021 | 0.08801176 | 0.45351639 |
| Gm3219 | 9.89714602 | -0.0386198 | 1.12972419 | -0.0341851 | 0.97272954 | 0.99443087 |

| | | | | | | |
|---|---|---|---|---|---|---|
| Gm2387 | 11.3768333 | -0.6197697 | 1.12574943 | -0.5505397 | 0.58194929 | 0.88321857 |
| Gm4824 | 20.2918412 | 0.67995478 | 0.96784087 | 0.70254811 | 0.48233741 | 0.84075713 |
| Gm7232 | 23.3982528 | 0.2822675 | 0.7700524 | 0.36655622 | 0.71395007 | 0.93235217 |
| LO018689.1 | 82.3152385 | 0.20611211 | 0.52092025 | 0.39566921 | 0.69234907 | 0.92542681 |
| AC119928.2 | 12.1675646 | 0.08704561 | 1.36693652 | 0.06367933 | 0.94922556 | 0.99028544 |
| Vps28 | 3457.77696 | 0.07990044 | 0.26068228 | 0.30650506 | 0.75922013 | 0.94184564 |
| Sept2.1 | 30683.7643 | 0.35621628 | 0.20136195 | 1.76903467 | 0.07688809 | 0.42792174 |
| Gm49510 | 52.7551091 | -0.3513053 | 0.70608114 | -0.4975424 | 0.61880662 | 0.89878056 |
| C030006K11 | 150.714852 | -1.1381278 | 0.39277629 | -2.8976489 | 0.00375971 | 0.06410023 |
| Pdxp | 324.841461 | -0.9484288 | 0.54067965 | -1.7541419 | 0.0794062 | 0.43628614 |
| Nnt.1 | 2661.28256 | 0.06397617 | 0.21307735 | 0.30024857 | 0.76398756 | 0.94300309 |
| Trerf1 | 58.8826515 | -1.1439516 | 0.65359564 | -1.7502436 | 0.08007628 | 0.43723638 |
| Lmbrd2.1 | 28.0002612 | 1.80137352 | 1.78102934 | 1.01142271 | 0.31181416 | 0.74030066 |
| Zc3h11a.1 | 1046.71098 | -0.0516265 | 0.2547561 | -0.2026508 | 0.83940795 | 0.96757914 |
| Riok2 | 1790.93015 | -0.0098985 | 0.24858527 | -0.0398194 | 0.96823709 | 0.99443087 |
| Gm4828 | 14.7005122 | 0.18686118 | 1.05795484 | 0.17662491 | 0.85980303 | 0.96954641 |
| Magef1 | 97.3065024 | -1.0542236 | 0.67233722 | -1.5679982 | 0.11688155 | 0.52252063 |
| Gm8670 | 16.0548098 | 0.36451194 | 1.51269965 | 0.24096782 | 0.80958006 | 0.95844313 |
| A630089N07 | 449.228473 | 0.42441572 | 0.51256808 | 0.82801824 | 0.40766017 | 0.8005951 |
| Gm6553 | 17.6044858 | 0.42739682 | 1.09995803 | 0.38855739 | 0.69760359 | 0.92667877 |
| Gm49760 | 234.507904 | 0.15161566 | 0.42991079 | 0.35266772 | 0.72433756 | 0.93488415 |
| Rps2-ps7 | 7.69730364 | 0.43083035 | 1.33027246 | 0.32386624 | 0.7460393 | 0.93959466 |
| Gm49594 | 12.525062 | 0.72645247 | 1.28784377 | 0.56408432 | 0.57269675 | 0.88035336 |
| Morf4l1-ps1 | 20.2231321 | 1.63340467 | 1.38875908 | 1.17616129 | 0.23953043 | 0.67562272 |
| Gm3435 | 386.834581 | 0.47540517 | 0.4223275 | 1.12567894 | 0.26030144 | 0.69647146 |
| 9030025P20I | 447.297189 | 0.49296482 | 0.35422942 | 1.3916541 | 0.16402717 | 0.59585062 |
| Gm21926 | 83.1535353 | 0.00397222 | 0.49967826 | 0.00794955 | 0.99365725 | 0.9984729 |
| Gm30531 | 22.8405367 | 0.40732092 | 0.8993235 | 0.45291925 | 0.65060689 | 0.91021183 |
| CT010433.1 | 32.09015 | 1.29940843 | 1.10617797 | 1.17468298 | 0.24012157 | 0.67563574 |
| Gm9805 | 26.436548 | 0.75242298 | 1.17188071 | 0.64206448 | 0.52083132 | 0.8576517 |
| Gm49909 | 2906.05877 | 0.14287565 | 0.19709857 | 0.7248944 | 0.46851681 | 0.83381643 |
| Gm20008 | 74.61827 | 0.72180719 | 0.77114848 | 0.93601584 | 0.34926503 | 0.76186861 |
| Gpr31c | 19.6854961 | -1.4729892 | 0.91097919 | -1.6169296 | 0.10589348 | 0.49777747 |
| Gm7072 | 693.97027 | 0.3120532 | 0.30861895 | 1.01112778 | 0.31195528 | 0.7404247 |
| Ptp4a1.1 | 12710.6989 | 0.12801644 | 0.15779587 | 0.81127877 | 0.4172056 | 0.8049685 |
| Gm50045 | 10.420427 | -0.2507201 | 1.12453966 | -0.2229535 | 0.82357167 | 0.9628698 |
| Apbb3 | 234.035073 | -0.1577657 | 0.4431427 | -0.3560156 | 0.72182886 | 0.93398714 |
| Gm28285 | 102.467512 | -0.6607462 | 0.45828952 | -1.4417659 | 0.14936843 | 0.57685486 |
| Chtf8.1 | 190.486036 | -0.3745392 | 0.66157486 | -0.5661327 | 0.57130355 | 0.88035336 |
| AC124502.3 | 69.5027023 | -0.2198216 | 0.51742984 | -0.4248336 | 0.67095802 | 0.91711586 |
| AC166779.2 | 16.5147368 | 0.62788578 | 1.01821747 | 0.61665194 | 0.53746433 | 0.86634214 |
| AC133509.1 | 22.1206056 | -0.1272641 | 0.77817791 | -0.1635412 | 0.87009238 | 0.97248575 |
| AC124512.1 | 24.7674957 | 0.11124381 | 0.80283322 | 0.13856404 | 0.88979466 | 0.97871887 |
| Tmem223 | 944.071164 | 0.06444637 | 0.29168414 | 0.22094577 | 0.82513466 | 0.96312579 |
| AC115631.1 | 157.998532 | 0.43471305 | 0.56236915 | 0.77300302 | 0.43952059 | 0.81963526 |
| Itprip | 2249.27008 | -0.3029649 | 0.32425356 | -0.9343455 | 0.35012568 | 0.76216093 |

| | | | | | | |
|---|---|---|---|---|---|---|
| AC137605.2 | 17.0182655 | -0.3245513 | 1.44736127 | -0.2242365 | 0.82257325 | 0.96252207 |
| Sssca1.1 | 334.837232 | 0.03058934 | 0.35052665 | 0.08726681 | 0.93045943 | 0.98716777 |
| AC090881.3 | 28.5842048 | -0.2451449 | 0.69862299 | -0.3508972 | 0.72566544 | 0.93544211 |
| AC102106.1 | 12.26281 | -0.0486344 | 1.31676125 | -0.0369348 | 0.97053697 | 0.99443087 |
| Fam220a | 210.343005 | 0.32908644 | 0.41703518 | 0.78910954 | 0.43004799 | 0.81197362 |
| Tmem179b | 1681.45534 | -0.2364436 | 0.17181398 | -1.3761603 | 0.16877202 | 0.60312169 |
| AC154328.1 | 107.423945 | 0.75608897 | 0.59785094 | 1.26467807 | 0.20598676 | 0.64036492 |

| significant | GeneID | Entrez |
|---|---|---|
| NA | ENSMUSG00 | 14679 |
| Significant | ENSMUSG00 | 12544 |
| NA | ENSMUSG00 | 107815 |
| NA | ENSMUSG00 | 67608 |
| NA | ENSMUSG00 | 12390 |
| NA | ENSMUSG00 | 23849 |
| NA | ENSMUSG00 | 29871 |
| NA | ENSMUSG00 | 12858 |
| NA | ENSMUSG00 | 14158 |
| NA | ENSMUSG00 | 74204 |
| NA | ENSMUSG00 | 209446 |
| NA | ENSMUSG00 | 231841 |
| NA | ENSMUSG00 | 14673 |
| NA | ENSMUSG00 | 18400 |
| NA | ENSMUSG00 | 72614 |
| NA | ENSMUSG00 | 235339 |
| NA | ENSMUSG00 | 66925 |
| Significant | ENSMUSG00 | 12444 |
| NA | ENSMUSG00 | 277463 |
| NA | ENSMUSG00 | 27027 |
| NA | ENSMUSG00 | 66355 |
| NA | ENSMUSG00 | 217069 |
| NA | ENSMUSG00 | 56077 |
| Significant | ENSMUSG00 | 74617 |
| NA | ENSMUSG00 | 17428 |
| Significant | ENSMUSG00 | 16414 |
| NA | ENSMUSG00 | 69692 |
| NA | ENSMUSG00 | 21987 |
| NA | ENSMUSG00 | 18618 |
| NA | ENSMUSG00 | 12029 |
| Significant | ENSMUSG00 | 17312 |
| NA | ENSMUSG00 | NA |
| Significant | ENSMUSG00 | 12846 |
| NA | ENSMUSG00 | 66368 |
| NA | ENSMUSG00 | 13171 |
| NA | ENSMUSG00 | 23994 |
| NA | ENSMUSG00 | 68995 |
| NA | ENSMUSG00 | 19700 |
| NA | ENSMUSG00 | 30054 |
| NA | ENSMUSG00 | 216131 |
| NA | ENSMUSG00 | 216527 |
| NA | ENSMUSG00 | 21379 |
| Significant | ENSMUSG00 | 17857 |
| NA | ENSMUSG00 | 14089 |
| NA | ENSMUSG00 | 66108 |

| | | |
|---|---|---|
| NA | ENSMUSG00 | 16818 |
| NA | ENSMUSG00 | 14423 |
| NA | ENSMUSG00 | 67027 |
| NA | ENSMUSG00 | 19016 |
| NA | ENSMUSG00 | 110157 |
| NA | ENSMUSG00 | NA |
| NA | ENSMUSG00 | 18591 |
| NA | ENSMUSG00 | 11482 |
| NA | ENSMUSG00 | 56149 |
| NA | ENSMUSG00 | 11479 |
| NA | ENSMUSG00 | 216810 |
| NA | ENSMUSG00 | 29813 |
| NA | ENSMUSG00 | 16402 |
| Significant | ENSMUSG00 | 70465 |
| Significant | ENSMUSG00 | NA |
| NA | ENSMUSG00 | 11950 |
| NA | ENSMUSG00 | 20682 |
| NA | ENSMUSG00 | 11991 |
| NA | ENSMUSG00 | 100040563 |
| NA | ENSMUSG00 | 57316 |
| NA | ENSMUSG00 | 14667 |
| NA | ENSMUSG00 | 79264 |
| NA | ENSMUSG00 | 12727 |
| NA | ENSMUSG00 | 15277 |
| NA | ENSMUSG00 | 360013 |
| Significant | ENSMUSG00 | 107766 |
| NA | ENSMUSG00 | 23833 |
| NA | ENSMUSG00 | 67477 |
| Significant | ENSMUSG00 | 16950 |
| NA | ENSMUSG00 | 18519 |
| NA | ENSMUSG00 | 19344 |
| NA | ENSMUSG00 | 50723 |
| NA | ENSMUSG00 | 234847 |
| NA | ENSMUSG00 | 270106 |
| NA | ENSMUSG00 | 234852 |
| NA | ENSMUSG00 | 68275 |
| NA | ENSMUSG00 | 20317 |
| NA | ENSMUSG00 | 259279 |
| NA | ENSMUSG00 | 67065 |
| NA | ENSMUSG00 | 13205 |
| NA | ENSMUSG00 | 16161 |
| NA | ENSMUSG00 | 237898 |
| NA | ENSMUSG00 | 232223 |
| NA | ENSMUSG00 | 269401 |
| NA | ENSMUSG00 | 13002 |
| NA | ENSMUSG00 | 66314 |

| | | |
|---|---|---|
| NA | ENSMUSG00 | 14265 |
| NA | ENSMUSG00 | 14025 |
| NA | ENSMUSG00 | 59038 |
| NA | ENSMUSG00 | 13972 |
| NA | ENSMUSG00 | 17385 |
| NA | ENSMUSG00 | 20587 |
| NA | ENSMUSG00 | 29816 |
| NA | ENSMUSG00 | 100609 |
| NA | ENSMUSG00 | 72960 |
| NA | ENSMUSG00 | 17387 |
| Significant | ENSMUSG00 | 20540 |
| NA | ENSMUSG00 | 69089 |
| NA | ENSMUSG00 | 217026 |
| NA | ENSMUSG00 | 20302 |
| NA | ENSMUSG00 | 67781 |
| NA | ENSMUSG00 | 66511 |
| NA | ENSMUSG00 | 20615 |
| NA | ENSMUSG00 | 20198 |
| NA | ENSMUSG00 | 20200 |
| NA | ENSMUSG00 | 15896 |
| NA | ENSMUSG00 | 23939 |
| NA | ENSMUSG00 | 13855 |
| NA | ENSMUSG00 | 27078 |
| NA | ENSMUSG00 | 99683 |
| NA | ENSMUSG00 | 212706 |
| NA | ENSMUSG00 | 66089 |
| NA | ENSMUSG00 | 52530 |
| NA | ENSMUSG00 | 72325 |
| NA | ENSMUSG00 | 57247 |
| NA | ENSMUSG00 | 68294 |
| NA | ENSMUSG00 | 269593 |
| NA | ENSMUSG00 | 20500 |
| NA | ENSMUSG00 | 330171 |
| NA | ENSMUSG00 | 67811 |
| NA | ENSMUSG00 | 55978 |
| NA | ENSMUSG00 | 59093 |
| NA | ENSMUSG00 | 16859 |
| NA | ENSMUSG00 | 11836 |
| NA | ENSMUSG00 | 18636 |
| NA | ENSMUSG00 | 22294 |
| NA | ENSMUSG00 | 94218 |
| NA | ENSMUSG00 | 214895 |
| NA | ENSMUSG00 | 54387 |
| NA | ENSMUSG00 | 18541 |
| NA | ENSMUSG00 | 17119 |
| NA | ENSMUSG00 | 23885 |

| | | |
|---|---|---|
| NA | ENSMUSG00 | 66618 |
| NA | ENSMUSG00 | 320634 |
| NA | ENSMUSG00 | 12313 |
| Significant | ENSMUSG00 | 28169 |
| NA | ENSMUSG00 | 20359 |
| NA | ENSMUSG00 | 18140 |
| NA | ENSMUSG00 | 224897 |
| NA | ENSMUSG00 | 54409 |
| NA | ENSMUSG00 | 54135 |
| NA | ENSMUSG00 | 52857 |
| Significant | ENSMUSG00 | 15451 |
| NA | ENSMUSG00 | 12709 |
| NA | ENSMUSG00 | 20683 |
| Significant | ENSMUSG00 | 16421 |
| NA | ENSMUSG00 | 60315 |
| NA | ENSMUSG00 | 19411 |
| NA | ENSMUSG00 | 56612 |
| NA | ENSMUSG00 | 67223 |
| NA | ENSMUSG00 | 11858 |
| NA | ENSMUSG00 | 27364 |
| NA | ENSMUSG00 | 11433 |
| NA | ENSMUSG00 | 71538 |
| NA | ENSMUSG00 | 73288 |
| NA | ENSMUSG00 | 15115 |
| NA | ENSMUSG00 | 66492 |
| NA | ENSMUSG00 | 68612 |
| NA | ENSMUSG00 | 229512 |
| NA | ENSMUSG00 | 12462 |
| NA | ENSMUSG00 | 56700 |
| NA | ENSMUSG00 | 17261 |
| NA | ENSMUSG00 | 71913 |
| NA | ENSMUSG00 | 56463 |
| NA | ENSMUSG00 | 12822 |
| NA | ENSMUSG00 | 20509 |
| NA | ENSMUSG00 | 16211 |
| NA | ENSMUSG00 | 19155 |
| NA | ENSMUSG00 | 107732 |
| NA | ENSMUSG00 | 13121 |
| NA | ENSMUSG00 | 66855 |
| NA | ENSMUSG00 | 67951 |
| NA | ENSMUSG00 | 23854 |
| NA | ENSMUSG00 | 17285 |
| NA | ENSMUSG00 | 12842 |
| NA | ENSMUSG00 | 14235 |
| NA | ENSMUSG00 | 101142 |
| NA | ENSMUSG00 | 60345 |

| | | |
|---|---|---|
| NA | ENSMUSG00 | 22158 |
| NA | ENSMUSG00 | 14885 |
| NA | ENSMUSG00 | 22154 |
| NA | ENSMUSG00 | 192657 |
| NA | ENSMUSG00 | 16480 |
| NA | ENSMUSG00 | 14230 |
| NA | ENSMUSG00 | 433864 |
| NA | ENSMUSG00 | 67458 |
| Significant | ENSMUSG00 | 216859 |
| NA | ENSMUSG00 | 21401 |
| NA | ENSMUSG00 | 15982 |
| NA | ENSMUSG00 | 232533 |
| NA | ENSMUSG00 | 78783 |
| NA | ENSMUSG00 | 11677 |
| NA | ENSMUSG00 | 103140 |
| NA | ENSMUSG00 | 13202 |
| NA | ENSMUSG00 | 66942 |
| NA | ENSMUSG00 | 24109 |
| NA | ENSMUSG00 | 107022 |
| NA | ENSMUSG00 | 66143 |
| NA | ENSMUSG00 | 11651 |
| Significant | ENSMUSG00 | 60363 |
| NA | ENSMUSG00 | 16170 |
| NA | ENSMUSG00 | 27060 |
| Significant | ENSMUSG00 | 27419 |
| NA | ENSMUSG00 | 71743 |
| NA | ENSMUSG00 | 319757 |
| NA | ENSMUSG00 | 232670 |
| NA | ENSMUSG00 | 66877 |
| Significant | ENSMUSG00 NA | |
| NA | ENSMUSG00 | 66917 |
| NA | ENSMUSG00 | 28088 |
| NA | ENSMUSG00 | 103136 |
| NA | ENSMUSG00 | 69754 |
| NA | ENSMUSG00 | 12336 |
| NA | ENSMUSG00 | 20480 |
| NA | ENSMUSG00 | 235072 |
| NA | ENSMUSG00 | 72881 |
| NA | ENSMUSG00 | 19353 |
| NA | ENSMUSG00 | 227720 |
| NA | ENSMUSG00 | 108897 |
| NA | ENSMUSG00 | 216558 |
| NA | ENSMUSG00 | 212528 |
| NA | ENSMUSG00 | 66830 |
| NA | ENSMUSG00 | 20514 |
| NA | ENSMUSG00 | 22201 |

| | | |
|---|---|---|
| NA | ENSMUSG00 | 22371 |
| NA | ENSMUSG00 | 22619 |
| NA | ENSMUSG00 | 69524 |
| NA | ENSMUSG00 | 20686 |
| NA | ENSMUSG00 | 108160 |
| NA | ENSMUSG00 | 13726 |
| NA | ENSMUSG00 | 70207 |
| Significant | ENSMUSG00 | 53623 |
| NA | ENSMUSG00 | 244668 |
| NA | ENSMUSG00 | 108011 |
| Significant | ENSMUSG00 | 109778 |
| NA | ENSMUSG00 | 56504 |
| NA | ENSMUSG00 | 15929 |
| Significant | ENSMUSG00 | 93843 |
| NA | ENSMUSG00 | 20832 |
| NA | ENSMUSG00 | 27061 |
| NA | ENSMUSG00 | 72722 |
| NA | ENSMUSG00 | 214162 |
| NA | ENSMUSG00 | 76568 |
| NA | ENSMUSG00 | 67091 |
| NA | ENSMUSG00 | 20926 |
| NA | ENSMUSG00 | 54141 |
| NA | ENSMUSG00 | 22248 |
| NA | ENSMUSG00 | 19376 |
| NA | ENSMUSG00 | 20316 |
| NA | ENSMUSG00 | 12447 |
| NA | ENSMUSG00 | 170770 |
| NA | ENSMUSG00 | 19182 |
| NA | ENSMUSG00 | 11432 |
| NA | ENSMUSG00 | 19400 |
| NA | ENSMUSG00 | 68427 |
| NA | ENSMUSG00 | 14007 |
| Significant | ENSMUSG00 | 22259 |
| NA | ENSMUSG00 | 107986 |
| NA | ENSMUSG00 | 20375 |
| NA | ENSMUSG00 | 67465 |
| NA | ENSMUSG00 | 20852 |
| NA | ENSMUSG00 | 12745 |
| Significant | ENSMUSG00 | 16541 |
| NA | ENSMUSG00 | 101568 |
| NA | ENSMUSG00 | 71997 |
| NA | ENSMUSG00 | 55982 |
| NA | ENSMUSG00 | 68477 |
| NA | ENSMUSG00 | 17454 |
| NA | ENSMUSG00 | 71887 |
| Significant | ENSMUSG00 | 11853 |

| | | |
|---|---|---|
| NA | ENSMUSG00 | NA |
| Significant | ENSMUSG00 | 23853 |
| NA | ENSMUSG00 | 18616 |
| NA | ENSMUSG00 | 70083 |
| NA | ENSMUSG00 | 76483 |
| NA | ENSMUSG00 | 67563 |
| NA | ENSMUSG00 | 57875 |
| NA | ENSMUSG00 | 27214 |
| NA | ENSMUSG00 | 13163 |
| NA | ENSMUSG00 | 54219 |
| NA | ENSMUSG00 | 75751 |
| NA | ENSMUSG00 | 74140 |
| NA | ENSMUSG00 | 16391 |
| NA | ENSMUSG00 | 105446 |
| NA | ENSMUSG00 | 67881 |
| NA | ENSMUSG00 | 52585 |
| NA | ENSMUSG00 | 234371 |
| NA | ENSMUSG00 | 76813 |
| NA | ENSMUSG00 | 72368 |
| NA | ENSMUSG00 | 73095 |
| NA | ENSMUSG00 | 66616 |
| NA | ENSMUSG00 | 71810 |
| NA | ENSMUSG00 | 69875 |
| NA | ENSMUSG00 | 13864 |
| NA | ENSMUSG00 | 67023 |
| NA | ENSMUSG00 | 77090 |
| NA | ENSMUSG00 | 13549 |
| NA | ENSMUSG00 | 109880 |
| NA | ENSMUSG00 | 68198 |
| NA | ENSMUSG00 | 20585 |
| NA | ENSMUSG00 | 68879 |
| NA | ENSMUSG00 | 56470 |
| NA | ENSMUSG00 | 20641 |
| NA | ENSMUSG00 | 77832 |
| NA | ENSMUSG00 | 21844 |
| NA | ENSMUSG00 | 22084 |
| Significant | ENSMUSG00 | 65962 |
| NA | ENSMUSG00 | 67959 |
| NA | ENSMUSG00 | 99412 |
| NA | ENSMUSG00 | 22245 |
| NA | ENSMUSG00 | 103742 |
| NA | ENSMUSG00 | 26362 |
| NA | ENSMUSG00 | 21803 |
| NA | ENSMUSG00 | 52132 |
| NA | ENSMUSG00 | 22700 |
| NA | ENSMUSG00 | 54194 |

| | | |
|---|---|---|
| NA | ENSMUSG00 | 68079 |
| NA | ENSMUSG00 | 98053 |
| NA | ENSMUSG00 | 53895 |
| NA | ENSMUSG00 | 66400 |
| NA | ENSMUSG00 | 70785 |
| NA | ENSMUSG00 | 69792 |
| NA | ENSMUSG00 | 16822 |
| NA | ENSMUSG00 | 110750 |
| NA | ENSMUSG00 | 67877 |
| NA | ENSMUSG00 | 23992 |
| NA | ENSMUSG00 | 14231 |
| NA | ENSMUSG00 | 83435 |
| NA | ENSMUSG00 | 56418 |
| NA | ENSMUSG00 | 22385 |
| NA | ENSMUSG00 | 224824 |
| NA | ENSMUSG00 | 27398 |
| NA | ENSMUSG00 | 17256 |
| NA | ENSMUSG00 | 68137 |
| NA | ENSMUSG00 | 70209 |
| NA | ENSMUSG00 | 110175 |
| NA | ENSMUSG00 | 16450 |
| NA | ENSMUSG00 | 399566 |
| NA | ENSMUSG00 | 66174 |
| NA | ENSMUSG00 | 14248 |
| NA | ENSMUSG00 | 21975 |
| NA | ENSMUSG00 | 235040 |
| NA | ENSMUSG00 | 60507 |
| NA | ENSMUSG00 | 15193 |
| NA | ENSMUSG00 | 27221 |
| NA | ENSMUSG00 | 11544 |
| NA | ENSMUSG00 | 67846 |
| NA | ENSMUSG00 | 76916 |
| NA | ENSMUSG00 | 85031 |
| NA | ENSMUSG00 | 17216 |
| NA | ENSMUSG00 | 24100 |
| NA | ENSMUSG00 | 17936 |
| NA | ENSMUSG00 | 26364 |
| NA | ENSMUSG00 | 16172 |
| NA | ENSMUSG00 | 16777 |
| NA | ENSMUSG00 | 70807 |
| Significant | ENSMUSG00 | 12491 |
| NA | ENSMUSG00 | 26400 |
| NA | ENSMUSG00 | 21856 |
| NA | ENSMUSG00 | 11772 |
| NA | ENSMUSG00 | 59047 |
| NA | ENSMUSG00 | 75613 |

| | | |
|---|---|---|
| NA | ENSMUSG00 | 56457 |
| NA | ENSMUSG00 | 19698 |
| NA | ENSMUSG00 | 53333 |
| Significant | ENSMUSG00 | 11816 |
| NA | ENSMUSG00 | 11813 |
| NA | ENSMUSG00 | 73389 |
| NA | ENSMUSG00 | 19088 |
| NA | ENSMUSG00 | 12576 |
| NA | ENSMUSG00 | 16600 |
| NA | ENSMUSG00 | 11767 |
| NA | ENSMUSG00 | 17274 |
| NA | ENSMUSG00 | 15331 |
| NA | ENSMUSG00 | 67922 |
| NA | ENSMUSG00 | 76205 |
| NA | ENSMUSG00 | 20869 |
| NA | ENSMUSG00 | 66043 |
| NA | ENSMUSG00 | 19060 |
| NA | ENSMUSG00 | 69131 |
| NA | ENSMUSG00 | 16890 |
| NA | ENSMUSG00 | 18475 |
| NA | ENSMUSG00 | 54610 |
| NA | ENSMUSG00 | 52846 |
| NA | ENSMUSG00 | 20527 |
| NA | ENSMUSG00 | 60611 |
| NA | ENSMUSG00 | 109552 |
| NA | ENSMUSG00 | 13356 |
| NA | ENSMUSG00 | 54131 |
| NA | ENSMUSG00 | 75736 |
| NA | ENSMUSG00 | 224893 |
| NA | ENSMUSG00 | 68047 |
| Significant | ENSMUSG00 | 20405 |
| NA | ENSMUSG00 | 50498 |
| NA | ENSMUSG00 | 72886 |
| NA | ENSMUSG00 | 19386 |
| NA | ENSMUSG00 | 14773 |
| NA | ENSMUSG00 | 13544 |
| NA | ENSMUSG00 | 27406 |
| NA | ENSMUSG00 | 224045 |
| NA | ENSMUSG00 | 19158 |
| NA | ENSMUSG00 | 56711 |
| NA | ENSMUSG00 | 15162 |
| NA | ENSMUSG00 | 66163 |
| NA | ENSMUSG00 | 50868 |
| NA | ENSMUSG00 | 11768 |
| NA | ENSMUSG00 | 20340 |
| NA | ENSMUSG00 | 208198 |

| | | |
|---|---|---|
| NA | ENSMUSG00 | 103236 |
| NA | ENSMUSG00 | 216169 |
| NA | ENSMUSG00 | 208228 |
| NA | ENSMUSG00 | 66120 |
| NA | ENSMUSG00 | 74351 |
| NA | ENSMUSG00 | 18807 |
| NA | ENSMUSG00 | 14470 |
| NA | ENSMUSG00 | 27049 |
| NA | ENSMUSG00 | 19089 |
| NA | ENSMUSG00 | 241230 |
| Significant | ENSMUSG00 | 14132 |
| NA | ENSMUSG00 | 66394 |
| NA | ENSMUSG00 | 68845 |
| NA | ENSMUSG00 | 27207 |
| NA | ENSMUSG00 | 20924 |
| NA | ENSMUSG00 | 54624 |
| NA | ENSMUSG00 | 66525 |
| NA | ENSMUSG00 | 67224 |
| NA | ENSMUSG00 | 59287 |
| NA | ENSMUSG00 | 19298 |
| Significant | ENSMUSG00 | 72054 |
| NA | ENSMUSG00 | 23917 |
| NA | ENSMUSG00 | 72349 |
| NA | ENSMUSG00 | 19125 |
| NA | ENSMUSG00 | 27886 |
| NA | ENSMUSG00 | 13358 |
| NA | ENSMUSG00 | 13353 |
| Significant | ENSMUSG00 | 15937 |
| NA | ENSMUSG00 | 14282 |
| NA | ENSMUSG00 | 74764 |
| NA | ENSMUSG00 | 13870 |
| NA | ENSMUSG00 | 57344 |
| NA | ENSMUSG00 | 26558 |
| NA | ENSMUSG00 | 382056 |
| NA | ENSMUSG00 | 71673 |
| NA | ENSMUSG00 | 74268 |
| NA | ENSMUSG00 | 74114 |
| Significant | ENSMUSG00 | 20111 |
| NA | ENSMUSG00 | 320632 |
| NA | ENSMUSG00 | 26371 |
| NA | ENSMUSG00 | NA |
| NA | ENSMUSG00 | 72999 |
| NA | ENSMUSG00 | 16650 |
| NA | ENSMUSG00 | 17155 |
| NA | ENSMUSG00 | 233011 |
| NA | ENSMUSG00 | 76889 |

| | | |
|---|---|---|
| NA | ENSMUSG00 | 78656 |
| NA | ENSMUSG00 | 19348 |
| NA | ENSMUSG00 | 66590 |
| NA | ENSMUSG00 | 270076 |
| NA | ENSMUSG00 | 17776 |
| NA | ENSMUSG00 | NA |
| NA | ENSMUSG00 | 19358 |
| NA | ENSMUSG00 | 12317 |
| NA | ENSMUSG00 | 71846 |
| NA | ENSMUSG00 | 54446 |
| NA | ENSMUSG00 | 67619 |
| NA | ENSMUSG00 | 18104 |
| NA | ENSMUSG00 | 14936 |
| NA | ENSMUSG00 | 20174 |
| NA | ENSMUSG00 | 12028 |
| NA | ENSMUSG00 | 21780 |
| NA | ENSMUSG00 | 224694 |
| Significant | ENSMUSG00 | 67468 |
| NA | ENSMUSG00 | 70186 |
| NA | ENSMUSG00 | 26961 |
| Significant | ENSMUSG00 | 66970 |
| NA | ENSMUSG00 | 67030 |
| NA | ENSMUSG00 | 14866 |
| NA | ENSMUSG00 | 68312 |
| NA | ENSMUSG00 | 20848 |
| NA | ENSMUSG00 | 20850 |
| NA | ENSMUSG00 | 19285 |
| NA | ENSMUSG00 | 26403 |
| NA | ENSMUSG00 | 11652 |
| NA | ENSMUSG00 | 83945 |
| NA | ENSMUSG00 | 15369 |
| NA | ENSMUSG00 | 66626 |
| NA | ENSMUSG00 | 65964 |
| NA | ENSMUSG00 | 66070 |
| NA | ENSMUSG00 | 13433 |
| NA | ENSMUSG00 | 235036 |
| NA | ENSMUSG00 | 26360 |
| NA | ENSMUSG00 | 108943 |
| Significant | ENSMUSG00 | 14009 |
| NA | ENSMUSG00 | 16581 |
| NA | ENSMUSG00 | 19156 |
| NA | ENSMUSG00 | 16151 |
| NA | ENSMUSG00 | 70560 |
| NA | ENSMUSG00 | 13498 |
| NA | ENSMUSG00 | 12034 |
| NA | ENSMUSG00 | 15170 |

| | | |
|---|---|---|
| NA | ENSMUSG00 | 14791 |
| NA | ENSMUSG00 | 14792 |
| NA | ENSMUSG00 | 66144 |
| NA | ENSMUSG00 | 12728 |
| NA | ENSMUSG00 | 12725 |
| NA | ENSMUSG00 | 53417 |
| NA | ENSMUSG00 | 70683 |
| NA | ENSMUSG00 | 20728 |
| Significant | ENSMUSG00 | 26554 |
| NA | ENSMUSG00 | 16795 |
| NA | ENSMUSG00 | 52513 |
| NA | ENSMUSG00 | 103694 |
| NA | ENSMUSG00 | 12122 |
| NA | ENSMUSG00 | 64143 |
| NA | ENSMUSG00 NA | |
| NA | ENSMUSG00 | 67838 |
| NA | ENSMUSG00 | 70573 |
| NA | ENSMUSG00 | 243834 |
| NA | ENSMUSG00 | 14389 |
| NA | ENSMUSG00 | 23790 |
| NA | ENSMUSG00 | 52440 |
| Significant | ENSMUSG00 | 13034 |
| NA | ENSMUSG00 | 29811 |
| NA | ENSMUSG00 | 52535 |
| NA | ENSMUSG00 | 268739 |
| NA | ENSMUSG00 | 50767 |
| NA | ENSMUSG00 | 94178 |
| NA | ENSMUSG00 | 102098 |
| NA | ENSMUSG00 | 109333 |
| Significant | ENSMUSG00 | 12489 |
| NA | ENSMUSG00 | 110826 |
| Significant | ENSMUSG00 | 72310 |
| NA | ENSMUSG00 | 20911 |
| Significant | ENSMUSG00 | 20392 |
| Significant | ENSMUSG00 | 69993 |
| NA | ENSMUSG00 | 80707 |
| NA | ENSMUSG00 | 20492 |
| NA | ENSMUSG00 | 56380 |
| Significant | ENSMUSG00 | 12798 |
| NA | ENSMUSG00 | 66420 |
| NA | ENSMUSG00 | 17925 |
| NA | ENSMUSG00 | 79221 |
| NA | ENSMUSG00 | 17085 |
| NA | ENSMUSG00 | 18106 |
| Significant | ENSMUSG00 | 13733 |
| NA | ENSMUSG00 | 67900 |

| | | |
|---|---|---|
| NA | ENSMUSG00 | 19335 |
| Significant | ENSMUSG00 | 53869 |
| NA | ENSMUSG00 | 217715 |
| NA | ENSMUSG00 | 78920 |
| NA | ENSMUSG00 | 18654 |
| NA | ENSMUSG00 | 29869 |
| Significant | ENSMUSG00 | 56221 |
| NA | ENSMUSG00 | 110524 |
| Significant | ENSMUSG00 | 71520 |
| NA | ENSMUSG00 | 68942 |
| Significant | ENSMUSG00 | 26433 |
| NA | ENSMUSG00 | 11769 |
| NA | ENSMUSG00 | 26415 |
| NA | ENSMUSG00 | 20815 |
| Significant | ENSMUSG00 | 98386 |
| NA | ENSMUSG00 | 12904 |
| NA | ENSMUSG00 | 18008 |
| NA | ENSMUSG00 | 94315 |
| NA | ENSMUSG00 | 229503 |
| NA | ENSMUSG00 | 15191 |
| NA | ENSMUSG00 | 71803 |
| Significant | ENSMUSG00 | 50492 |
| NA | ENSMUSG00 | 57267 |
| NA | ENSMUSG00 | 17179 |
| NA | ENSMUSG00 | 59004 |
| NA | ENSMUSG00 | 26395 |
| NA | ENSMUSG00 | 52551 |
| NA | ENSMUSG00 | 70544 |
| NA | ENSMUSG00 | 74198 |
| NA | ENSMUSG00 | 15507 |
| NA | ENSMUSG00 | 54153 |
| NA | ENSMUSG00 | 53379 |
| NA | ENSMUSG00 | 108017 |
| NA | ENSMUSG00 | 67736 |
| NA | ENSMUSG00 | 67873 |
| NA | ENSMUSG00 | 18749 |
| NA | ENSMUSG00 | 27029 |
| Significant | ENSMUSG00 | 13014 |
| Significant | ENSMUSG00 | 23921 |
| NA | ENSMUSG00 | 19305 |
| NA | ENSMUSG00 | 104771 |
| NA | ENSMUSG00 | 12505 |
| NA | ENSMUSG00 | 27103 |
| NA | ENSMUSG00 | 22388 |
| NA | ENSMUSG00 | 117591 |
| NA | ENSMUSG00 | 17988 |

| | | |
|---|---|---|
| NA | ENSMUSG00 | 17159 |
| NA | ENSMUSG00 | 12224 |
| NA | ENSMUSG00 | 67836 |
| NA | ENSMUSG00 | 21672 |
| NA | ENSMUSG00 | 20020 |
| NA | ENSMUSG00 | 80886 |
| Significant | ENSMUSG00 | 231999 |
| NA | ENSMUSG00 | 66442 |
| NA | ENSMUSG00 | 327954 |
| NA | ENSMUSG00 | 22230 |
| NA | ENSMUSG00 | 170740 |
| NA | ENSMUSG00 | 56384 |
| NA | ENSMUSG00 | 56085 |
| NA | ENSMUSG00 | 56551 |
| NA | ENSMUSG00 | 20512 |
| NA | ENSMUSG00 | 58799 |
| NA | ENSMUSG00 | 17688 |
| NA | ENSMUSG00 | 225055 |
| NA | ENSMUSG00 | 58805 |
| NA | ENSMUSG00 | 27368 |
| NA | ENSMUSG00 | 66138 |
| NA | ENSMUSG00 | 18073 |
| NA | ENSMUSG00 | 17218 |
| Significant | ENSMUSG00 | 15368 |
| NA | ENSMUSG00 | 26936 |
| NA | ENSMUSG00 | 71722 |
| NA | ENSMUSG00 | 18476 |
| NA | ENSMUSG00 | 18747 |
| NA | ENSMUSG00 | 66929 |
| NA | ENSMUSG00 | 68278 |
| NA | ENSMUSG00 | 81489 |
| NA | ENSMUSG00 | 227334 |
| NA | ENSMUSG00 | 67136 |
| NA | ENSMUSG00 | 13046 |
| NA | ENSMUSG00 | 68349 |
| NA | ENSMUSG00 | 18984 |
| NA | ENSMUSG00 | 16001 |
| NA | ENSMUSG00 | 16337 |
| NA | ENSMUSG00 | 11944 |
| NA | ENSMUSG00 | 21849 |
| NA | ENSMUSG00 | 22192 |
| NA | ENSMUSG00 | 11515 |
| NA | ENSMUSG00 | 17260 |
| NA | ENSMUSG00 | 22083 |
| NA | ENSMUSG00 | 13690 |
| Significant | ENSMUSG00 | 17540 |

| | | |
|---|---|---|
| NA | ENSMUSG00 | 13026 |
| NA | ENSMUSG00 | 233410 |
| NA | ENSMUSG00 | 19185 |
| Significant | ENSMUSG00 | 72183 |
| Significant | ENSMUSG00 | 17768 |
| NA | ENSMUSG00 | 16590 |
| NA | ENSMUSG00 | 641376 |
| NA | ENSMUSG00 | 11807 |
| NA | ENSMUSG00 | 103135 |
| NA | ENSMUSG00 | 12974 |
| NA | ENSMUSG00 | 11717 |
| NA | ENSMUSG00 | 68183 |
| NA | ENSMUSG00 | 13018 |
| NA | ENSMUSG00 | 56513 |
| NA | ENSMUSG00 | 83383 |
| NA | ENSMUSG00 | 19385 |
| NA | ENSMUSG00 | 53970 |
| NA | ENSMUSG00 | 19172 |
| Significant | ENSMUSG00 | 17394 |
| NA | ENSMUSG00 | 22785 |
| Significant | ENSMUSG00 | 59010 |
| NA | ENSMUSG00 | 18457 |
| NA | ENSMUSG00 | 75624 |
| NA | ENSMUSG00 | 78308 |
| NA | ENSMUSG00 | 50930 |
| NA | ENSMUSG00 | 66409 |
| NA | ENSMUSG00 | 11789 |
| NA | ENSMUSG00 | 13476 |
| NA | ENSMUSG00 | 66366 |
| NA | ENSMUSG00 | 56046 |
| NA | ENSMUSG00 | 17978 |
| NA | ENSMUSG00 | 22026 |
| NA | ENSMUSG00 | 22025 |
| NA | ENSMUSG00 | 77626 |
| NA | ENSMUSG00 | 71382 |
| NA | ENSMUSG00 | 66989 |
| NA | ENSMUSG00 | 83429 |
| NA | ENSMUSG00 | 74637 |
| NA | ENSMUSG00 | 22156 |
| NA | ENSMUSG00 | 19672 |
| NA | ENSMUSG00 | 13419 |
| NA | ENSMUSG00 | 68015 |
| NA | ENSMUSG00 | 74763 |
| NA | ENSMUSG00 | NA |
| NA | ENSMUSG00 | 68423 |
| NA | ENSMUSG00 | 108989 |

| | | |
|---|---|---|
| NA | ENSMUSG00 | 226499 |
| NA | ENSMUSG00 | 71723 |
| NA | ENSMUSG00 | 70394 |
| NA | ENSMUSG00 | 108124 |
| NA | ENSMUSG00 | 24068 |
| NA | ENSMUSG00 | 11951 |
| NA | ENSMUSG00 | 27681 |
| NA | ENSMUSG00 | 66411 |
| NA | ENSMUSG00 | 19062 |
| NA | ENSMUSG00 | 12929 |
| NA | ENSMUSG00 | 67425 |
| NA | ENSMUSG00 | 216705 |
| NA | ENSMUSG00 | 54373 |
| NA | ENSMUSG00 | 68916 |
| NA | ENSMUSG00 | 22642 |
| NA | ENSMUSG00 | 74202 |
| Significant | ENSMUSG00 | 13857 |
| NA | ENSMUSG00 | 67119 |
| NA | ENSMUSG00 | 68473 |
| NA | ENSMUSG00 | 11966 |
| NA | ENSMUSG00 | 13859 |
| NA | ENSMUSG00 | 21745 |
| NA | ENSMUSG00 | 219022 |
| NA | ENSMUSG00 | 66246 |
| NA | ENSMUSG00 | 227290 |
| NA | ENSMUSG00 | 69660 |
| NA | ENSMUSG00 | 76709 |
| NA | ENSMUSG00 | 75410 |
| NA | ENSMUSG00 | 69804 |
| NA | ENSMUSG00 | 76846 |
| NA | ENSMUSG00 | 69714 |
| NA | ENSMUSG00 | 23887 |
| NA | ENSMUSG00 | 14598 |
| NA | ENSMUSG00 | 68337 |
| Significant | ENSMUSG00 | 12925 |
| NA | ENSMUSG00 | 12398 |
| NA | ENSMUSG00 | 14114 |
| NA | ENSMUSG00 | 53328 |
| NA | ENSMUSG00 | 26912 |
| NA | ENSMUSG00 | 21687 |
| NA | ENSMUSG00 | 54325 |
| NA | ENSMUSG00 | 80509 |
| NA | ENSMUSG00 | 68180 |
| NA | ENSMUSG00 | 107995 |
| NA | ENSMUSG00 | 71740 |
| NA | ENSMUSG00 | 18637 |

| | | |
|---|---|---|
| NA | ENSMUSG00 | 81018 |
| NA | ENSMUSG00 | 77644 |
| NA | ENSMUSG00 | 18011 |
| Significant | ENSMUSG00 | 20810 |
| NA | ENSMUSG00 | 13836 |
| NA | ENSMUSG00 | 56275 |
| NA | ENSMUSG00 | 70605 |
| NA | ENSMUSG00 | 52028 |
| NA | ENSMUSG00 | 78797 |
| NA | ENSMUSG00 | 56876 |
| NA | ENSMUSG00 | 228026 |
| NA | ENSMUSG00 | 19205 |
| NA | ENSMUSG00 | 192156 |
| NA | ENSMUSG00 | 13057 |
| NA | ENSMUSG00 | 69179 |
| NA | ENSMUSG00 | 54650 |
| NA | ENSMUSG00 | 51799 |
| NA | ENSMUSG00 | 67177 |
| NA | ENSMUSG00 | 12395 |
| NA | ENSMUSG00 | 11821 |
| NA | ENSMUSG00 | 14884 |
| Significant | ENSMUSG00 | 15216 |
| NA | ENSMUSG00 | 57742 |
| NA | ENSMUSG00 | 330064 |
| NA | ENSMUSG00 | 54631 |
| NA | ENSMUSG00 | 69232 |
| NA | ENSMUSG00 | 74443 |
| NA | ENSMUSG00 | 71472 |
| NA | ENSMUSG00 | 18968 |
| NA | ENSMUSG00 | 12540 |
| NA | ENSMUSG00 | 18771 |
| NA | ENSMUSG00 | 57441 |
| NA | ENSMUSG00 | 66834 |
| NA | ENSMUSG00 | 193452 |
| NA | ENSMUSG00 | 12567 |
| Significant | ENSMUSG00 | 14421 |
| NA | ENSMUSG00 | 17299 |
| NA | ENSMUSG00 | 67125 |
| NA | ENSMUSG00 | 16573 |
| NA | ENSMUSG00 | 78935 |
| NA | ENSMUSG00 | 12799 |
| NA | ENSMUSG00 | 74407 |
| NA | ENSMUSG00 | 72043 |
| NA | ENSMUSG00 | 20656 |
| NA | ENSMUSG00 | 71983 |
| NA | ENSMUSG00 | 70527 |

| | | |
|---|---|---|
| NA | ENSMUSG00 | 14055 |
| NA | ENSMUSG00 | 66180 |
| NA | ENSMUSG00 | 12387 |
| NA | ENSMUSG00 | 68969 |
| NA | ENSMUSG00 | 21762 |
| NA | ENSMUSG00 | 22321 |
| NA | ENSMUSG00 | 27762 |
| NA | ENSMUSG00 | 15482 |
| NA | ENSMUSG00 | 70129 |
| NA | ENSMUSG00 | 17687 |
| NA | ENSMUSG00 | 193742 |
| NA | ENSMUSG00 | 18010 |
| Significant | ENSMUSG00 | 51793 |
| NA | ENSMUSG00 | 114584 |
| NA | ENSMUSG00 | 27756 |
| NA | ENSMUSG00 | 18973 |
| NA | ENSMUSG00 | 20907 |
| NA | ENSMUSG00 | 243372 |
| NA | ENSMUSG00 | 18120 |
| NA | ENSMUSG00 | 329727 |
| NA | ENSMUSG00 | 17169 |
| NA | ENSMUSG00 | 67210 |
| NA | ENSMUSG00 | 17113 |
| NA | ENSMUSG00 | 241296 |
| NA | ENSMUSG00 | 51792 |
| NA | ENSMUSG00 | 72775 |
| NA | ENSMUSG00 | 28113 |
| NA | ENSMUSG00 | 224907 |
| NA | ENSMUSG00 | 70359 |
| NA | ENSMUSG00 | 21812 |
| NA | ENSMUSG00 | 26556 |
| NA | ENSMUSG00 | 114714 |
| NA | ENSMUSG00 | 12389 |
| NA | ENSMUSG00 | 59046 |
| NA | ENSMUSG00 | 12048 |
| NA | ENSMUSG00 | 16549 |
| NA | ENSMUSG00 | 234388 |
| NA | ENSMUSG00 | 12464 |
| NA | ENSMUSG00 | 66050 |
| NA | ENSMUSG00 | 78070 |
| NA | ENSMUSG00 | 72611 |
| NA | ENSMUSG00 | 11848 |
| NA | ENSMUSG00 | 328365 |
| NA | ENSMUSG00 | 232339 |
| NA | ENSMUSG00 | 69748 |
| NA | ENSMUSG00 | 77134 |

| | | |
|---|---|---|
| NA | ENSMUSG00 | 65116 |
| NA | ENSMUSG00 | 59013 |
| NA | ENSMUSG00 | 76411 |
| NA | ENSMUSG00 | 15903 |
| NA | ENSMUSG00 | 93760 |
| Significant | ENSMUSG00 | 13033 |
| NA | ENSMUSG00 | 56040 |
| NA | ENSMUSG00 | 64296 |
| NA | ENSMUSG00 | 67286 |
| NA | ENSMUSG00 | 14365 |
| NA | ENSMUSG00 | 68041 |
| NA | ENSMUSG00 | 232910 |
| NA | ENSMUSG00 | 116701 |
| NA | ENSMUSG00 | 74666 |
| NA | ENSMUSG00 | 14200 |
| NA | ENSMUSG00 | 69683 |
| NA | ENSMUSG00 | 232370 |
| NA | ENSMUSG00 | 68628 |
| NA | ENSMUSG00 | 55935 |
| NA | ENSMUSG00 | 67260 |
| NA | ENSMUSG00 | 74616 |
| NA | ENSMUSG00 | 56698 |
| NA | ENSMUSG00 | 21885 |
| NA | ENSMUSG00 | 320100 |
| NA | ENSMUSG00 | 20639 |
| NA | ENSMUSG00 | 22190 |
| NA | ENSMUSG00 | 68988 |
| NA | ENSMUSG00 | 55942 |
| Significant | ENSMUSG00 | 52502 |
| NA | ENSMUSG00 | 13713 |
| NA | ENSMUSG00 | 80517 |
| NA | ENSMUSG00 | 108841 |
| NA | ENSMUSG00 | 68051 |
| NA | ENSMUSG00 | 67771 |
| NA | ENSMUSG00 | 18987 |
| NA | ENSMUSG00 | 56615 |
| NA | ENSMUSG00 | 94232 |
| NA | ENSMUSG00 | 20084 |
| NA | ENSMUSG00 | 110954 |
| NA | ENSMUSG00 | 267019 |
| NA | ENSMUSG00 | 52683 |
| NA | ENSMUSG00 | 15257 |
| NA | ENSMUSG00 | 17156 |
| NA | ENSMUSG00 | 66204 |
| NA | ENSMUSG00 | 15184 |
| NA | ENSMUSG00 | 56044 |

| | | |
|---|---|---|
| NA | ENSMUSG00 | 22335 |
| NA | ENSMUSG00 | 21427 |
| NA | ENSMUSG00 | 14390 |
| NA | ENSMUSG00 | 56455 |
| NA | ENSMUSG00 | 67466 |
| NA | ENSMUSG00 | 223693 |
| NA | ENSMUSG00 | 18016 |
| NA | ENSMUSG00 | 67178 |
| NA | ENSMUSG00 | 14030 |
| NA | ENSMUSG00 | 11764 |
| NA | ENSMUSG00 | 14686 |
| NA | ENSMUSG00 | 94044 |
| NA | ENSMUSG00 | 93838 |
| NA | ENSMUSG00 | 69605 |
| NA | ENSMUSG00 | 17173 |
| NA | ENSMUSG00 | 108705 |
| NA | ENSMUSG00 | 28240 |
| NA | ENSMUSG00 | 22213 |
| Significant | ENSMUSG00 | 17523 |
| Significant | ENSMUSG00 | 17295 |
| NA | ENSMUSG00 | 240638 |
| NA | ENSMUSG00 | 13712 |
| NA | ENSMUSG00 | 215690 |
| NA | ENSMUSG00 | 238799 |
| NA | ENSMUSG00 | 67897 |
| NA | ENSMUSG00 | 20813 |
| NA | ENSMUSG00 | 107951 |
| NA | ENSMUSG00 | 30933 |
| NA | ENSMUSG00 | 14287 |
| NA | ENSMUSG00 | 239719 |
| NA | ENSMUSG00 | 12419 |
| NA | ENSMUSG00 | 80287 |
| NA | ENSMUSG00 | 22325 |
| NA | ENSMUSG00 | 19053 |
| NA | ENSMUSG00 | 14373 |
| NA | ENSMUSG00 | 66233 |
| NA | ENSMUSG00 | 215999 |
| NA | ENSMUSG00 | 18302 |
| NA | ENSMUSG00 | 110279 |
| Significant | ENSMUSG00 | 18301 |
| NA | ENSMUSG00 | 16506 |
| NA | ENSMUSG00 | 21422 |
| NA | ENSMUSG00 | 22221 |
| Significant | ENSMUSG00 | 74137 |
| NA | ENSMUSG00 | 56542 |
| NA | ENSMUSG00 | 67680 |

| | | |
|---|---|---|
| Significant | ENSMUSG00 | 84682 |
| NA | ENSMUSG00 | 67826 |
| NA | ENSMUSG00 | 70750 |
| NA | ENSMUSG00 | 20479 |
| NA | ENSMUSG00 | 75617 |
| NA | ENSMUSG00 | 83454 |
| NA | ENSMUSG00 | 66826 |
| NA | ENSMUSG00 | 286942 |
| NA | ENSMUSG00 | 11671 |
| NA | ENSMUSG00 | 56395 |
| NA | ENSMUSG00 | 15587 |
| NA | ENSMUSG00 | 15983 |
| NA | ENSMUSG00 | 15586 |
| NA | ENSMUSG00 | 80385 |
| NA | ENSMUSG00 | 56032 |
| NA | ENSMUSG00 | 56289 |
| NA | ENSMUSG00 | 17254 |
| NA | ENSMUSG00 | 53319 |
| NA | ENSMUSG00 | 56389 |
| NA | ENSMUSG00 | 57916 |
| NA | ENSMUSG00 | 19130 |
| NA | ENSMUSG00 | 71766 |
| NA | ENSMUSG00 | 72503 |
| NA | ENSMUSG00 | 98404 |
| Significant | ENSMUSG00 | 67893 |
| NA | ENSMUSG00 | 70110 |
| NA | ENSMUSG00 | 66599 |
| NA | ENSMUSG00 | 18002 |
| NA | ENSMUSG00 | 53334 |
| NA | ENSMUSG00 | 68836 |
| NA | ENSMUSG00 | 226976 |
| NA | ENSMUSG00 | 14084 |
| NA | ENSMUSG00 | 76927 |
| NA | ENSMUSG00 | 67493 |
| NA | ENSMUSG00 | 66268 |
| NA | ENSMUSG00 | 67039 |
| NA | ENSMUSG00 | 19165 |
| NA | ENSMUSG00 | 235674 |
| NA | ENSMUSG00 | 18799 |
| Significant | ENSMUSG00 | 76267 |
| NA | ENSMUSG00 | 56873 |
| NA | ENSMUSG00 | 79202 |
| NA | ENSMUSG00 | 27267 |
| NA | ENSMUSG00 | 56369 |
| NA | ENSMUSG00 | 27402 |
| NA | ENSMUSG00 | 234729 |

| | | |
|---|---|---|
| Significant | ENSMUSG00 | 68279 |
| NA | ENSMUSG00 | 67605 |
| NA | ENSMUSG00 | 21376 |
| Significant | ENSMUSG00 | 11565 |
| NA | ENSMUSG00 | 72308 |
| NA | ENSMUSG00 | 22355 |
| NA | ENSMUSG00 | 18263 |
| NA | ENSMUSG00 | 66628 |
| NA | ENSMUSG00 | 11492 |
| Significant | ENSMUSG00 | 230721 |
| NA | ENSMUSG00 | 74463 |
| NA | ENSMUSG00 | 63872 |
| NA | ENSMUSG00 | 70616 |
| NA | ENSMUSG00 | 71755 |
| NA | ENSMUSG00 | 233056 |
| NA | ENSMUSG00 | 70300 |
| NA | ENSMUSG00 | 80297 |
| NA | ENSMUSG00 | 18648 |
| NA | ENSMUSG00 | 14020 |
| NA | ENSMUSG00 | 213027 |
| NA | ENSMUSG00 | 102209 |
| NA | ENSMUSG00 | 216963 |
| NA | ENSMUSG00 | 56356 |
| NA | ENSMUSG00 | 12175 |
| NA | ENSMUSG00 | 12455 |
| NA | ENSMUSG00 | 52592 |
| NA | ENSMUSG00 | 94112 |
| NA | ENSMUSG00 | 67422 |
| NA | ENSMUSG00 | 230806 |
| NA | ENSMUSG00 | 67586 |
| NA | ENSMUSG00 | 74094 |
| NA | ENSMUSG00 | 66480 |
| NA | ENSMUSG00 | 66074 |
| NA | ENSMUSG00 | 117167 |
| NA | ENSMUSG00 | 66308 |
| NA | ENSMUSG00 | 16551 |
| NA | ENSMUSG00 | 68240 |
| Significant | ENSMUSG00 | 74568 |
| NA | ENSMUSG00 | 320938 |
| NA | ENSMUSG00 | 320244 |
| NA | ENSMUSG00 | 69930 |
| NA | ENSMUSG00 | 57264 |
| NA | ENSMUSG00 | 22295 |
| NA | ENSMUSG00 | 20103 |
| NA | ENSMUSG00 | 330814 |
| NA | ENSMUSG00 | 75723 |

| | | |
|---|---|---|
| NA | ENSMUSG00 | 104156 |
| NA | ENSMUSG00 | 70575 |
| NA | ENSMUSG00 | 102124 |
| NA | ENSMUSG00 | 11972 |
| NA | ENSMUSG00 | 19280 |
| NA | ENSMUSG00 | 98396 |
| NA | ENSMUSG00 | 140486 |
| NA | ENSMUSG00 | 58202 |
| NA | ENSMUSG00 | 72392 |
| NA | ENSMUSG00 | 27883 |
| NA | ENSMUSG00 | 19378 |
| NA | ENSMUSG00 | 226646 |
| NA | ENSMUSG00 | 381629 |
| Significant | ENSMUSG00 | 69719 |
| NA | ENSMUSG00 | 79566 |
| NA | ENSMUSG00 | 67979 |
| NA | ENSMUSG00 | 19211 |
| Significant | ENSMUSG00 | 18611 |
| NA | ENSMUSG00 | 53600 |
| NA | ENSMUSG00 | 69769 |
| NA | ENSMUSG00 | 70047 |
| NA | ENSMUSG00 | 110147 |
| NA | ENSMUSG00 | 66126 |
| NA | ENSMUSG00 | 216154 |
| NA | ENSMUSG00 | 20442 |
| NA | ENSMUSG00 | 216157 |
| NA | ENSMUSG00 | 77733 |
| NA | ENSMUSG00 | NA |
| NA | ENSMUSG00 | 21945 |
| Significant | ENSMUSG00 | 69189 |
| NA | ENSMUSG00 | 27045 |
| NA | ENSMUSG00 | 114604 |
| NA | ENSMUSG00 | 70238 |
| NA | ENSMUSG00 | 66061 |
| NA | ENSMUSG00 | 56398 |
| NA | ENSMUSG00 | 63873 |
| NA | ENSMUSG00 | 100503085 |
| NA | ENSMUSG00 | 67510 |
| NA | ENSMUSG00 | 56354 |
| NA | ENSMUSG00 | 12301 |
| NA | ENSMUSG00 | 76779 |
| NA | ENSMUSG00 | 69747 |
| NA | ENSMUSG00 | 327942 |
| NA | ENSMUSG00 | 17991 |
| NA | ENSMUSG00 | 66449 |
| NA | ENSMUSG00 | 83396 |

| | | |
|---|---|---|
| NA | ENSMUSG00 | 12864 |
| NA | ENSMUSG00 | 268470 |
| Significant | ENSMUSG00 | 72477 |
| NA | ENSMUSG00 | 17222 |
| NA | ENSMUSG00 | 17289 |
| NA | ENSMUSG00 | 22088 |
| NA | ENSMUSG00 | 246694 |
| NA | ENSMUSG00 | 26407 |
| NA | ENSMUSG00 | 234839 |
| NA | ENSMUSG00 | 68718 |
| NA | ENSMUSG00 | 105522 |
| NA | ENSMUSG00 | 237615 |
| NA | ENSMUSG00 | 53871 |
| NA | ENSMUSG00 | 66384 |
| NA | ENSMUSG00 | 268752 |
| NA | ENSMUSG00 | 78287 |
| NA | ENSMUSG00 | 64658 |
| NA | ENSMUSG00 | 27369 |
| NA | ENSMUSG00 | 100072 |
| NA | ENSMUSG00 | 12977 |
| NA | ENSMUSG00 | 229707 |
| NA | ENSMUSG00 | 67863 |
| NA | ENSMUSG00 | 66531 |
| NA | ENSMUSG00 | 231600 |
| NA | ENSMUSG00 | 21766 |
| NA | ENSMUSG00 | 67544 |
| NA | ENSMUSG00 | 21374 |
| NA | ENSMUSG00 | 19170 |
| NA | ENSMUSG00 | 18567 |
| NA | ENSMUSG00 | 234686 |
| NA | ENSMUSG00 | 234683 |
| NA | ENSMUSG00 | 20855 |
| NA | ENSMUSG00 | 74356 |
| NA | ENSMUSG00 | 67971 |
| NA | ENSMUSG00 | 17686 |
| NA | ENSMUSG00 | 66320 |
| NA | ENSMUSG00 | 104394 |
| NA | ENSMUSG00 | 20932 |
| NA | ENSMUSG00 | 20931 |
| NA | ENSMUSG00 | 27362 |
| NA | ENSMUSG00 | 234344 |
| NA | ENSMUSG00 | 22612 |
| NA | ENSMUSG00 | 19046 |
| NA | ENSMUSG00 | 70231 |
| NA | ENSMUSG00 | 66637 |
| NA | ENSMUSG00 | 76740 |

| | | |
|---|---|---|
| NA | ENSMUSG00 | 59005 |
| NA | ENSMUSG00 | 257633 |
| NA | ENSMUSG00 | 13680 |
| NA | ENSMUSG00 | 50917 |
| NA | ENSMUSG00 | 14461 |
| NA | ENSMUSG00 | 69379 |
| NA | ENSMUSG00 | 227624 |
| NA | ENSMUSG00 | 59022 |
| NA | ENSMUSG00 | 30839 |
| NA | ENSMUSG00 | 50799 |
| NA | ENSMUSG00 | 22196 |
| NA | ENSMUSG00 | 68327 |
| NA | ENSMUSG00 | 74154 |
| NA | ENSMUSG00 | 233328 |
| NA | ENSMUSG00 | 56847 |
| NA | ENSMUSG00 | 109711 |
| NA | ENSMUSG00 | 64383 |
| NA | ENSMUSG00 | 15388 |
| NA | ENSMUSG00 | 70769 |
| NA | ENSMUSG00 | 213819 |
| NA | ENSMUSG00 | 215748 |
| NA | ENSMUSG00 | 53332 |
| NA | ENSMUSG00 | 15354 |
| NA | ENSMUSG00 | 11303 |
| NA | ENSMUSG00 | 66536 |
| NA | ENSMUSG00 | 66192 |
| NA | ENSMUSG00 NA | |
| NA | ENSMUSG00 | 14567 |
| NA | ENSMUSG00 | 70097 |
| NA | ENSMUSG00 | 17873 |
| NA | ENSMUSG00 | 30925 |
| NA | ENSMUSG00 | 27218 |
| NA | ENSMUSG00 | 66220 |
| NA | ENSMUSG00 | 13804 |
| Significant | ENSMUSG00 | 13058 |
| NA | ENSMUSG00 NA | |
| NA | ENSMUSG00 | 22439 |
| NA | ENSMUSG00 | 12506 |
| NA | ENSMUSG00 | 270166 |
| NA | ENSMUSG00 | 67976 |
| NA | ENSMUSG00 | 69440 |
| NA | ENSMUSG00 | 12522 |
| NA | ENSMUSG00 | 12915 |
| NA | ENSMUSG00 | 18132 |
| NA | ENSMUSG00 | 54397 |
| NA | ENSMUSG00 | 54197 |

| | | |
|---|---|---|
| NA | ENSMUSG00 | 381356 |
| NA | ENSMUSG00 | 15273 |
| NA | ENSMUSG00 | 11863 |
| NA | ENSMUSG00 | 17434 |
| NA | ENSMUSG00 | 66176 |
| NA | ENSMUSG00 | 16956 |
| NA | ENSMUSG00 | 11974 |
| NA | ENSMUSG00 | 57908 |
| NA | ENSMUSG00 | 20807 |
| NA | ENSMUSG00 | 14462 |
| Significant | ENSMUSG00 | 70358 |
| NA | ENSMUSG00 | 15481 |
| NA | ENSMUSG00 | 321007 |
| NA | ENSMUSG00 | 72621 |
| NA | ENSMUSG00 | 19166 |
| NA | ENSMUSG00 | 75398 |
| NA | ENSMUSG00 | 84505 |
| NA | ENSMUSG00 | 71790 |
| NA | ENSMUSG00 | 229589 |
| NA | ENSMUSG00 | 76893 |
| NA | ENSMUSG00 | 12343 |
| NA | ENSMUSG00 | 67220 |
| NA | ENSMUSG00 | 22365 |
| NA | ENSMUSG00 | 70767 |
| NA | ENSMUSG00 | 66471 |
| NA | ENSMUSG00 | 226548 |
| NA | ENSMUSG00 | 68652 |
| NA | ENSMUSG00 | 67418 |
| NA | ENSMUSG00 | 12793 |
| Significant | ENSMUSG00 | 13860 |
| NA | ENSMUSG00 | 20933 |
| NA | ENSMUSG00 | 20930 |
| NA | ENSMUSG00 | 66999 |
| NA | ENSMUSG00 | 110391 |
| NA | ENSMUSG00 | 18412 |
| NA | ENSMUSG00 | 18024 |
| NA | ENSMUSG00 | 20181 |
| NA | ENSMUSG00 | 229595 |
| Significant | ENSMUSG00 | 80891 |
| NA | ENSMUSG00 | 11801 |
| NA | ENSMUSG00 | 67763 |
| NA | ENSMUSG00 | 54392 |
| NA | ENSMUSG00 | 209707 |
| NA | ENSMUSG00 | 16993 |
| NA | ENSMUSG00 | 56431 |
| Significant | ENSMUSG00 | 26914 |

| | | |
|---|---|---|
| NA | ENSMUSG00 | 114674 |
| NA | ENSMUSG00 | 69168 |
| NA | ENSMUSG00 | 80909 |
| NA | ENSMUSG00 | 14129 |
| NA | ENSMUSG00 | 17969 |
| NA | ENSMUSG00 | 11566 |
| NA | ENSMUSG00 | 12289 |
| NA | ENSMUSG00 | 56249 |
| NA | ENSMUSG00 | 76612 |
| NA | ENSMUSG00 | 14272 |
| NA | ENSMUSG00 | 72198 |
| NA | ENSMUSG00 | 16803 |
| NA | ENSMUSG00 | 14247 |
| NA | ENSMUSG00 | 215193 |
| Significant | ENSMUSG00 | 15483 |
| NA | ENSMUSG00 | 329324 |
| Significant | ENSMUSG00 | 14991 |
| NA | ENSMUSG00 | 74365 |
| NA | ENSMUSG00 | 67126 |
| NA | ENSMUSG00 | 57314 |
| NA | ENSMUSG00 | 64138 |
| NA | ENSMUSG00 | 66390 |
| NA | ENSMUSG00 | 22209 |
| NA | ENSMUSG00 | 11740 |
| NA | ENSMUSG00 | 66496 |
| NA | ENSMUSG00 | 102866 |
| NA | ENSMUSG00 | 77015 |
| NA | ENSMUSG00 | 67050 |
| NA | ENSMUSG00 | 54405 |
| NA | ENSMUSG00 | 13557 |
| NA | ENSMUSG00 | 12946 |
| NA | ENSMUSG00 | 67533 |
| Significant | ENSMUSG00 | 12490 |
| NA | ENSMUSG00 | 67759 |
| NA | ENSMUSG00 | 60533 |
| NA | ENSMUSG00 | 66596 |
| NA | ENSMUSG00 | 76366 |
| NA | ENSMUSG00 | 140887 |
| Significant | ENSMUSG00 | 226419 |
| NA | ENSMUSG00 | 17164 |
| NA | ENSMUSG00 | 16153 |
| NA | ENSMUSG00 | 16784 |
| NA | ENSMUSG00 | 54138 |
| NA | ENSMUSG00 | 239554 |
| NA | ENSMUSG00 | 55944 |
| NA | ENSMUSG00 | 15081 |

| | | |
|---|---|---|
| NA | ENSMUSG00 | 18141 |
| NA | ENSMUSG00 | 67042 |
| NA | ENSMUSG00 | 23970 |
| NA | ENSMUSG00 | 12763 |
| NA | ENSMUSG00 | 223723 |
| NA | ENSMUSG00 | 268741 |
| NA | ENSMUSG00 | 68735 |
| NA | ENSMUSG00 | 67996 |
| NA | ENSMUSG00 | 18803 |
| NA | ENSMUSG00 | 70382 |
| NA | ENSMUSG00 | 69259 |
| NA | ENSMUSG00 | 68145 |
| Significant | ENSMUSG00 | 20568 |
| Significant | ENSMUSG00 | 20971 |
| NA | ENSMUSG00 | 16164 |
| NA | ENSMUSG00 | 17966 |
| NA | ENSMUSG00 | 19157 |
| NA | ENSMUSG00 | 74194 |
| NA | ENSMUSG00 | 12189 |
| NA | ENSMUSG00 | 68106 |
| NA | ENSMUSG00 | 52469 |
| NA | ENSMUSG00 | 23989 |
| NA | ENSMUSG00 | 22123 |
| NA | ENSMUSG00 | 50912 |
| NA | ENSMUSG00 | 67201 |
| NA | ENSMUSG00 | 68299 |
| NA | ENSMUSG00 | 216965 |
| NA | ENSMUSG00 | 76233 |
| NA | ENSMUSG00 | 170789 |
| NA | ENSMUSG00 | 52685 |
| NA | ENSMUSG00 | 18099 |
| NA | ENSMUSG00 | 22032 |
| NA | ENSMUSG00 | 11676 |
| NA | ENSMUSG00 | 19921 |
| NA | ENSMUSG00 | 140859 |
| NA | ENSMUSG00 | 12298 |
| NA | ENSMUSG00 | 75869 |
| NA | ENSMUSG00 | 22680 |
| NA | ENSMUSG00 | 69077 |
| Significant | ENSMUSG00 | 21858 |
| NA | ENSMUSG00 | 76014 |
| NA | ENSMUSG00 | 21974 |
| NA | ENSMUSG00 | 218772 |
| Significant | ENSMUSG00 | 16010 |
| NA | ENSMUSG00 | 23834 |
| NA | ENSMUSG00 | 52615 |

| | | |
|---|---|---|
| NA | ENSMUSG00 | 237877 |
| NA | ENSMUSG00 | 54394 |
| NA | ENSMUSG00 | 217169 |
| NA | ENSMUSG00 | 21927 |
| NA | ENSMUSG00 | 109934 |
| NA | ENSMUSG00 | 21939 |
| NA | ENSMUSG00 | 228875 |
| NA | ENSMUSG00 | 228876 |
| NA | ENSMUSG00 | 140579 |
| NA | ENSMUSG00 | 78889 |
| NA | ENSMUSG00 | 76080 |
| NA | ENSMUSG00 | 59040 |
| NA | ENSMUSG00 | 11486 |
| NA | ENSMUSG00 | 26943 |
| NA | ENSMUSG00 | 71776 |
| NA | ENSMUSG00 | 74451 |
| NA | ENSMUSG00 | 11799 |
| NA | ENSMUSG00 | 71562 |
| NA | ENSMUSG00 | 78928 |
| NA | ENSMUSG00 | 52840 |
| Significant | ENSMUSG00 | 17395 |
| NA | ENSMUSG00 | 80860 |
| NA | ENSMUSG00 | 18830 |
| NA | ENSMUSG00 | 20499 |
| NA | ENSMUSG00 | 19025 |
| NA | ENSMUSG00 | 71971 |
| NA | ENSMUSG00 | 20498 |
| NA | ENSMUSG00 | 17913 |
| NA | ENSMUSG00 | 12928 |
| NA | ENSMUSG00 | 12867 |
| NA | ENSMUSG00 | 18738 |
| NA | ENSMUSG00 | 21428 |
| NA | ENSMUSG00 | 67998 |
| NA | ENSMUSG00 | 80861 |
| NA | ENSMUSG00 | 271457 |
| NA | ENSMUSG00 | 71966 |
| NA | ENSMUSG00 | 26931 |
| NA | ENSMUSG00 | 245866 |
| NA | ENSMUSG00 | 17865 |
| NA | ENSMUSG00 | 56336 |
| NA | ENSMUSG00 | 228858 |
| NA | ENSMUSG00 | 228889 |
| NA | ENSMUSG00 | 19159 |
| NA | ENSMUSG00 | 72318 |
| NA | ENSMUSG00 | 170758 |
| NA | ENSMUSG00 | 74778 |

| | | |
|---|---|---|
| NA | ENSMUSG00 | 109754 |
| NA | ENSMUSG00 | 70422 |
| NA | ENSMUSG00 | 76199 |
| NA | ENSMUSG00 | 68024 |
| NA | ENSMUSG00 | 17135 |
| NA | ENSMUSG00 | 19014 |
| NA | ENSMUSG00 | 13867 |
| NA | ENSMUSG00 | 59045 |
| NA | ENSMUSG00 | 17305 |
| NA | ENSMUSG00 | 75909 |
| NA | ENSMUSG00 | 56207 |
| NA | ENSMUSG00 | 69367 |
| NA | ENSMUSG00 | 20822 |
| NA | ENSMUSG00 | 58231 |
| Significant | ENSMUSG00 | 18858 |
| NA | ENSMUSG00 | 14566 |
| NA | ENSMUSG00 | 67966 |
| NA | ENSMUSG00 | 19175 |
| NA | ENSMUSG00 | 216873 |
| NA | ENSMUSG00 | 18643 |
| NA | ENSMUSG00 | 67145 |
| NA | ENSMUSG00 | 54401 |
| NA | ENSMUSG00 | 16706 |
| Significant | ENSMUSG00 | 14778 |
| NA | ENSMUSG00 | 11749 |
| NA | ENSMUSG00 | 16162 |
| NA | ENSMUSG00 | 52712 |
| NA | ENSMUSG00 | 16647 |
| NA | ENSMUSG00 | 66313 |
| NA | ENSMUSG00 | 320162 |
| NA | ENSMUSG00 | 22344 |
| NA | ENSMUSG00 | 110809 |
| Significant | ENSMUSG00 | 66610 |
| NA | ENSMUSG00 | 71774 |
| Significant | ENSMUSG00 | 16568 |
| Significant | ENSMUSG00 | 20362 |
| NA | ENSMUSG00 | 170749 |
| NA | ENSMUSG00 | 217038 |
| NA | ENSMUSG00 | 76719 |
| NA | ENSMUSG00 | 66771 |
| NA | ENSMUSG00 | 17912 |
| NA | ENSMUSG00 | 67487 |
| NA | ENSMUSG00 | 77864 |
| NA | ENSMUSG00 | 11640 |
| NA | ENSMUSG00 | 68979 |
| NA | ENSMUSG00 | 116891 |

| | | |
|---|---|---|
| Significant | ENSMUSG00 | 12261 |
| NA | ENSMUSG00 | 69723 |
| NA | ENSMUSG00 | 103712 |
| NA | ENSMUSG00 | 114644 |
| NA | ENSMUSG00 | 216848 |
| NA | ENSMUSG00 | 216850 |
| NA | ENSMUSG00 | 66884 |
| NA | ENSMUSG00 | 11541 |
| NA | ENSMUSG00 | 20185 |
| NA | ENSMUSG00 | 22368 |
| NA | ENSMUSG00 | 73139 |
| NA | ENSMUSG00 | 22658 |
| NA | ENSMUSG00 | 67480 |
| NA | ENSMUSG00 | 108083 |
| NA | ENSMUSG00 | 68729 |
| NA | ENSMUSG00 | 67181 |
| NA | ENSMUSG00 | 54351 |
| NA | ENSMUSG00 | 56486 |
| NA | ENSMUSG00 | 70419 |
| NA | ENSMUSG00 | 78246 |
| NA | ENSMUSG00 | 11370 |
| NA | ENSMUSG00 | 27041 |
| NA | ENSMUSG00 | 11927 |
| NA | ENSMUSG00 | 20692 |
| NA | ENSMUSG00 | 237781 |
| NA | ENSMUSG00 | 21386 |
| NA | ENSMUSG00 | 56405 |
| NA | ENSMUSG00 | 217031 |
| NA | ENSMUSG00 | 22778 |
| NA | ENSMUSG00 | 103711 |
| NA | ENSMUSG00 | 16834 |
| NA | ENSMUSG00 | 12412 |
| NA | ENSMUSG00 | 80280 |
| Significant | ENSMUSG00 | 56358 |
| NA | ENSMUSG00 | 68066 |
| NA | ENSMUSG00 | 78912 |
| NA | ENSMUSG00 | 56321 |
| NA | ENSMUSG00 | 13424 |
| NA | ENSMUSG00 | 103737 |
| NA | ENSMUSG00 | 83431 |
| NA | ENSMUSG00 | 71998 |
| NA | ENSMUSG00 | 75580 |
| NA | ENSMUSG00 | 24070 |
| NA | ENSMUSG00 | 23879 |
| NA | ENSMUSG00 | 228033 |
| NA | ENSMUSG00 | 12514 |

| | | |
|---|---|---|
| NA | ENSMUSG00 | 14081 |
| NA | ENSMUSG00 | 319822 |
| Significant | ENSMUSG00 | 16985 |
| NA | ENSMUSG00 | 319740 |
| NA | ENSMUSG00 | 69534 |
| NA | ENSMUSG00 | 19364 |
| NA | ENSMUSG00 | 211347 |
| NA | ENSMUSG00 | 104458 |
| NA | ENSMUSG00 | 68572 |
| NA | ENSMUSG00 | 14149 |
| NA | ENSMUSG00 | 67036 |
| NA | ENSMUSG00 | 16362 |
| NA | ENSMUSG00 | 20520 |
| Significant | ENSMUSG00 | 18452 |
| NA | ENSMUSG00 | 109689 |
| NA | ENSMUSG00 | 66102 |
| NA | ENSMUSG00 | 75273 |
| NA | ENSMUSG00 | 66172 |
| Significant | ENSMUSG00 | 20305 |
| NA | ENSMUSG00 | 20303 |
| NA | ENSMUSG00 | 24083 |
| NA | ENSMUSG00 | 26397 |
| NA | ENSMUSG00 | 22629 |
| NA | ENSMUSG00 | 53890 |
| NA | ENSMUSG00 | 242705 |
| NA | ENSMUSG00 | 20557 |
| Significant | ENSMUSG00 | 244141 |
| NA | ENSMUSG00 | 58246 |
| NA | ENSMUSG00 | 67789 |
| NA | ENSMUSG00 | 66437 |
| Significant | ENSMUSG00 | 18822 |
| NA | ENSMUSG00 | 19340 |
| NA | ENSMUSG00 | 69572 |
| NA | ENSMUSG00 | 68267 |
| NA | ENSMUSG00 | 54411 |
| Significant | ENSMUSG00 | 69537 |
| Significant | ENSMUSG00 | 20308 |
| NA | ENSMUSG00 | 69938 |
| NA | ENSMUSG00 | 79555 |
| NA | ENSMUSG00 | 71780 |
| NA | ENSMUSG00 | 70791 |
| NA | ENSMUSG00 | 69094 |
| NA | ENSMUSG00 | 19345 |
| NA | ENSMUSG00 | 17448 |
| NA | ENSMUSG00 | 14950 |
| NA | ENSMUSG00 | 74315 |

| | | |
|---|---|---|
| NA | ENSMUSG00 | 20266 |
| NA | ENSMUSG00 | 11973 |
| NA | ENSMUSG00 | 214901 |
| NA | ENSMUSG00 | 238323 |
| NA | ENSMUSG00 | 232807 |
| NA | ENSMUSG00 | 11622 |
| NA | ENSMUSG00 | 270058 |
| NA | ENSMUSG00 | 68366 |
| NA | ENSMUSG00 | 67842 |
| NA | ENSMUSG00 | 11975 |
| NA | ENSMUSG00 | 19183 |
| NA | ENSMUSG00 | 78266 |
| NA | ENSMUSG00 | 71584 |
| NA | ENSMUSG00 | 101966 |
| NA | ENSMUSG00 | 12315 |
| NA | ENSMUSG00 | 26572 |
| NA | ENSMUSG00 | 14232 |
| NA | ENSMUSG00 | 53817 |
| NA | ENSMUSG00 | 67903 |
| NA | ENSMUSG00 | 68385 |
| NA | ENSMUSG00 | 70310 |
| NA | ENSMUSG00 | 52666 |
| NA | ENSMUSG00 | 67439 |
| NA | ENSMUSG00 | 12539 |
| NA | ENSMUSG00 | 19341 |
| NA | ENSMUSG00 | 106628 |
| NA | ENSMUSG00 | 26893 |
| NA | ENSMUSG00 | 22187 |
| NA | ENSMUSG00 | 11781 |
| NA | ENSMUSG00 | 27357 |
| Significant | ENSMUSG00 | 52377 |
| NA | ENSMUSG00 | 13496 |
| Significant | ENSMUSG00 | 27273 |
| NA | ENSMUSG00 | 66530 |
| NA | ENSMUSG00 | 28106 |
| NA | ENSMUSG00 | 20358 |
| NA | ENSMUSG00 | 72654 |
| NA | ENSMUSG00 | 66117 |
| NA | ENSMUSG00 | 23797 |
| NA | ENSMUSG00 | 67707 |
| NA | ENSMUSG00 | 74412 |
| NA | ENSMUSG00 | 67217 |
| NA | ENSMUSG00 | 17101 |
| NA | ENSMUSG00 | 270066 |
| NA | ENSMUSG00 | 73316 |
| NA | ENSMUSG00 | 353499 |

| | | |
|---|---|---|
| NA | ENSMUSG00 | 69920 |
| NA | ENSMUSG00 | 16661 |
| NA | ENSMUSG00 | 66084 |
| NA | ENSMUSG00 | 13982 |
| NA | ENSMUSG00 | 67141 |
| NA | ENSMUSG00 | 108853 |
| NA | ENSMUSG00 | 15182 |
| NA | ENSMUSG00 | 14302 |
| NA | ENSMUSG00 | 66521 |
| NA | ENSMUSG00 | 78808 |
| NA | ENSMUSG00 | 66847 |
| NA | ENSMUSG00 | 73681 |
| NA | ENSMUSG00 | 23924 |
| NA | ENSMUSG00 | 18537 |
| NA | ENSMUSG00 | 237253 |
| NA | ENSMUSG00 | 67851 |
| NA | ENSMUSG00 | 140740 |
| Significant | ENSMUSG00 | 54198 |
| NA | ENSMUSG00 | 66253 |
| NA | ENSMUSG00 | 66757 |
| NA | ENSMUSG00 | 56535 |
| NA | ENSMUSG00 | 66848 |
| NA | ENSMUSG00 | 103268 |
| NA | ENSMUSG00 | 353258 |
| NA | ENSMUSG00 | 22634 |
| NA | ENSMUSG00 | 53599 |
| NA | ENSMUSG00 | 22288 |
| NA | ENSMUSG00 | 20598 |
| NA | ENSMUSG00 | 171580 |
| NA | ENSMUSG00 | 268294 |
| NA | ENSMUSG00 | 83767 |
| NA | ENSMUSG00 | 67844 |
| NA | ENSMUSG00 | NA |
| NA | ENSMUSG00 | 67371 |
| NA | ENSMUSG00 | 72472 |
| NA | ENSMUSG00 | 19714 |
| NA | ENSMUSG00 | 103213 |
| NA | ENSMUSG00 | 14360 |
| NA | ENSMUSG00 | 71924 |
| NA | ENSMUSG00 | 19072 |
| NA | ENSMUSG00 | 21929 |
| Significant | ENSMUSG00 | 64058 |
| NA | ENSMUSG00 | 56016 |
| NA | ENSMUSG00 | 19707 |
| NA | ENSMUSG00 | 66403 |
| NA | ENSMUSG00 | 94221 |

| | | |
|---|---|---|
| NA | ENSMUSG00 | 76563 |
| NA | ENSMUSG00 | 170728 |
| NA | ENSMUSG00 | 11630 |
| NA | ENSMUSG00 NA | |
| Significant | ENSMUSG00 | 57319 |
| NA | ENSMUSG00 | 28193 |
| NA | ENSMUSG00 | 18768 |
| NA | ENSMUSG00 | 56442 |
| NA | ENSMUSG00 | 15500 |
| NA | ENSMUSG00 | 52665 |
| Significant | ENSMUSG00 | 19272 |
| NA | ENSMUSG00 | 66686 |
| NA | ENSMUSG00 | 52713 |
| NA | ENSMUSG00 | 17931 |
| NA | ENSMUSG00 | 18451 |
| NA | ENSMUSG00 | 103080 |
| NA | ENSMUSG00 | 110829 |
| NA | ENSMUSG00 | 52696 |
| NA | ENSMUSG00 | 216080 |
| NA | ENSMUSG00 | 12534 |
| NA | ENSMUSG00 | 67972 |
| NA | ENSMUSG00 | 69288 |
| NA | ENSMUSG00 | 71371 |
| NA | ENSMUSG00 | 67019 |
| NA | ENSMUSG00 | 75089 |
| NA | ENSMUSG00 | 382406 |
| NA | ENSMUSG00 | 67603 |
| NA | ENSMUSG00 | 21917 |
| NA | ENSMUSG00 | 17311 |
| NA | ENSMUSG00 | 19164 |
| NA | ENSMUSG00 | 20393 |
| NA | ENSMUSG00 | 216274 |
| NA | ENSMUSG00 | 67454 |
| NA | ENSMUSG00 | 56422 |
| NA | ENSMUSG00 | 13822 |
| NA | ENSMUSG00 | 11783 |
| NA | ENSMUSG00 | 17863 |
| NA | ENSMUSG00 | 70208 |
| NA | ENSMUSG00 | 52906 |
| NA | ENSMUSG00 | 11846 |
| NA | ENSMUSG00 | 17997 |
| NA | ENSMUSG00 | 71804 |
| NA | ENSMUSG00 | 17761 |
| NA | ENSMUSG00 | 53331 |
| NA | ENSMUSG00 | 18634 |
| NA | ENSMUSG00 | 15979 |

| | | |
|---|---|---|
| NA | ENSMUSG00 | 237459 |
| NA | ENSMUSG00 | 15109 |
| NA | ENSMUSG00 | 69878 |
| NA | ENSMUSG00 | 327799 |
| Significant | ENSMUSG00 | 13998 |
| NA | ENSMUSG00 | 66414 |
| NA | ENSMUSG00 | 319880 |
| NA | ENSMUSG00 | 77048 |
| NA | ENSMUSG00 | 216233 |
| NA | ENSMUSG00 | 71207 |
| NA | ENSMUSG00 | 77976 |
| NA | ENSMUSG00 | 216198 |
| NA | ENSMUSG00 | 12952 |
| NA | ENSMUSG00 | 74007 |
| NA | ENSMUSG00 | 21859 |
| NA | ENSMUSG00 | 22027 |
| Significant | ENSMUSG00 | 16000 |
| NA | ENSMUSG00 | 67282 |
| NA | ENSMUSG00 | 71712 |
| NA | ENSMUSG00 | 93759 |
| NA | ENSMUSG00 | 67345 |
| NA | ENSMUSG00 | 432467 |
| NA | ENSMUSG00 | 70432 |
| NA | ENSMUSG00 | 67307 |
| NA | ENSMUSG00 | 67500 |
| NA | ENSMUSG00 | 56200 |
| NA | ENSMUSG00 | 94213 |
| Significant | ENSMUSG00 | 19073 |
| NA | ENSMUSG00 | 30930 |
| NA | ENSMUSG00 | 338359 |
| NA | ENSMUSG00 | 71361 |
| NA | ENSMUSG00 | 404634 |
| NA | ENSMUSG00 | 71767 |
| NA | ENSMUSG00 | 20224 |
| NA | ENSMUSG00 | 67895 |
| NA | ENSMUSG00 | 13688 |
| NA | ENSMUSG00 | 27355 |
| NA | ENSMUSG00 | 20397 |
| NA | ENSMUSG00 | 13180 |
| NA | ENSMUSG00 | 71279 |
| NA | ENSMUSG00 | 74048 |
| NA | ENSMUSG00 | 20503 |
| NA | ENSMUSG00 | 52717 |
| NA | ENSMUSG00 | 74747 |
| NA | ENSMUSG00 | 56709 |
| NA | ENSMUSG00 | 216001 |

| | | |
|---|---|---|
| NA | ENSMUSG00 | 71902 |
| NA | ENSMUSG00 | 56480 |
| NA | ENSMUSG00 | 66249 |
| NA | ENSMUSG00 | 56193 |
| NA | ENSMUSG00 | 14479 |
| Significant | ENSMUSG00 | 50701 |
| NA | ENSMUSG00 | 245944 |
| NA | ENSMUSG00 | 66687 |
| NA | ENSMUSG00 | 18551 |
| NA | ENSMUSG00 | 216344 |
| NA | ENSMUSG00 | 66374 |
| NA | ENSMUSG00 | 67245 |
| NA | ENSMUSG00 | 66816 |
| NA | ENSMUSG00 | 55963 |
| NA | ENSMUSG00 | 94176 |
| NA | ENSMUSG00 | 19324 |
| NA | ENSMUSG00 | 14431 |
| NA | ENSMUSG00 | 66713 |
| NA | ENSMUSG00 | 75406 |
| NA | ENSMUSG00 | 68114 |
| NA | ENSMUSG00 | 17268 |
| NA | ENSMUSG00 | 66594 |
| NA | ENSMUSG00 | 72068 |
| NA | ENSMUSG00 | 21423 |
| NA | ENSMUSG00 | 327826 |
| NA | ENSMUSG00 | 64050 |
| NA | ENSMUSG00 | 76877 |
| NA | ENSMUSG00 | 14783 |
| NA | ENSMUSG00 | 67332 |
| NA | ENSMUSG00 | 70574 |
| NA | ENSMUSG00 | 17246 |
| NA | ENSMUSG00 | 52679 |
| NA | ENSMUSG00 | 13008 |
| NA | ENSMUSG00 | 237542 |
| NA | ENSMUSG00 | 17347 |
| NA | ENSMUSG00 | 53604 |
| NA | ENSMUSG00 | 104248 |
| NA | ENSMUSG00 | 11776 |
| NA | ENSMUSG00 | 21664 |
| NA | ENSMUSG00 | 20222 |
| NA | ENSMUSG00 | 17245 |
| NA | ENSMUSG00 | 30055 |
| NA | ENSMUSG00 | 230895 |
| NA | ENSMUSG00 | 66225 |
| NA | ENSMUSG00 | 68212 |
| NA | ENSMUSG00 | 73914 |

| | | |
|---|---|---|
| NA | ENSMUSG00 | 117599 |
| Significant | ENSMUSG00 | 15468 |
| NA | ENSMUSG00 | 64451 |
| NA | ENSMUSG00 | 15353 |
| NA | ENSMUSG00 | 56371 |
| NA | ENSMUSG00 | 103425 |
| NA | ENSMUSG00 | 67933 |
| NA | ENSMUSG00 | 18045 |
| NA | ENSMUSG00 | 50493 |
| NA | ENSMUSG00 | 67905 |
| NA | ENSMUSG00 | 28109 |
| NA | ENSMUSG00 | 216188 |
| NA | ENSMUSG00 | 77305 |
| NA | ENSMUSG00 | 235582 |
| NA | ENSMUSG00 | 80294 |
| NA | ENSMUSG00 | 215335 |
| Significant | ENSMUSG00 | 216190 |
| NA | ENSMUSG00 | 20610 |
| NA | ENSMUSG00 | 15254 |
| NA | ENSMUSG00 | 75530 |
| NA | ENSMUSG00 | 103583 |
| NA | ENSMUSG00 | 20868 |
| NA | ENSMUSG00 | 216578 |
| NA | ENSMUSG00 | 19696 |
| NA | ENSMUSG00 | 18641 |
| NA | ENSMUSG00 | 74467 |
| NA | ENSMUSG00 | 13650 |
| NA | ENSMUSG00 | 72129 |
| NA | ENSMUSG00 | 67884 |
| NA | ENSMUSG00 | 268395 |
| NA | ENSMUSG00 | 268390 |
| NA | ENSMUSG00 | 17168 |
| NA | ENSMUSG00 | 103573 |
| NA | ENSMUSG00 | 67579 |
| NA | ENSMUSG00 | 20856 |
| NA | ENSMUSG00 | 65257 |
| NA | ENSMUSG00 | 216150 |
| NA | ENSMUSG00 | 110012 |
| NA | ENSMUSG00 | 68044 |
| NA | ENSMUSG00 | 66753 |
| NA | ENSMUSG00 | 20742 |
| NA | ENSMUSG00 | 216560 |
| NA | ENSMUSG00 | 17449 |
| Significant | ENSMUSG00 | 73106 |
| NA | ENSMUSG00 | 12450 |
| NA | ENSMUSG00 | 52653 |

| | | |
|---|---|---|
| NA | ENSMUSG00 | 216151 |
| NA | ENSMUSG00 | 15366 |
| NA | ENSMUSG00 | 30805 |
| NA | ENSMUSG00 | 76884 |
| NA | ENSMUSG00 | 17308 |
| NA | ENSMUSG00 | 19052 |
| NA | ENSMUSG00 | 14257 |
| NA | ENSMUSG00 | 15384 |
| NA | ENSMUSG00 | 72947 |
| NA | ENSMUSG00 | 15525 |
| NA | ENSMUSG00 | 104625 |
| NA | ENSMUSG00 | 14584 |
| NA | ENSMUSG00 | 21408 |
| NA | ENSMUSG00 | 26420 |
| NA | ENSMUSG00 | 12330 |
| NA | ENSMUSG00 | 14694 |
| NA | ENSMUSG00 | 216724 |
| NA | ENSMUSG00 | 59044 |
| Significant | ENSMUSG00 | 17001 |
| NA | ENSMUSG00 | 19360 |
| NA | ENSMUSG00 | 68067 |
| NA | ENSMUSG00 | 12750 |
| NA | ENSMUSG00 | 66397 |
| Significant | ENSMUSG00 | 30794 |
| NA | ENSMUSG00 | 213084 |
| NA | ENSMUSG00 | 22210 |
| NA | ENSMUSG00 | 52626 |
| NA | ENSMUSG00 | 84035 |
| NA | ENSMUSG00 | 66213 |
| NA | ENSMUSG00 | 171285 |
| NA | ENSMUSG00 | 57783 |
| NA | ENSMUSG00 | 22333 |
| NA | ENSMUSG00 | 22271 |
| NA | ENSMUSG00 | 193116 |
| NA | ENSMUSG00 | 75452 |
| NA | ENSMUSG00 | 15574 |
| NA | ENSMUSG00 | 30939 |
| NA | ENSMUSG00 | 75828 |
| NA | ENSMUSG00 | 545938 |
| NA | ENSMUSG00 | 319939 |
| NA | ENSMUSG00 | 12227 |
| NA | ENSMUSG00 | 71962 |
| NA | ENSMUSG00 | 16009 |
| NA | ENSMUSG00 | 64934 |
| Significant | ENSMUSG00 | 21452 |
| Significant | ENSMUSG00 | 246177 |

| | | |
|---|---|---|
| NA | ENSMUSG00 | 29856 |
| NA | ENSMUSG00 | 11844 |
| NA | ENSMUSG00 | 67862 |
| NA | ENSMUSG00 | 14923 |
| NA | ENSMUSG00 | 193670 |
| NA | ENSMUSG00 | 16886 |
| NA | ENSMUSG00 | 56218 |
| NA | ENSMUSG00 | 74203 |
| NA | ENSMUSG00 | 94091 |
| NA | ENSMUSG00 | 18293 |
| NA | ENSMUSG00 | 13494 |
| NA | ENSMUSG00 | 68585 |
| NA | ENSMUSG00 | 76784 |
| NA | ENSMUSG00 | 78294 |
| NA | ENSMUSG00 | 216618 |
| NA | ENSMUSG00 | 104570 |
| NA | ENSMUSG00 | 71701 |
| NA | ENSMUSG00 | 18972 |
| NA | ENSMUSG00 | 268417 |
| NA | ENSMUSG00 | 54125 |
| NA | ENSMUSG00 | 13169 |
| NA | ENSMUSG00 | 64660 |
| NA | ENSMUSG00 | 104479 |
| NA | ENSMUSG00 | 68097 |
| NA | ENSMUSG00 | 22433 |
| NA | ENSMUSG00 | 20922 |
| NA | ENSMUSG00 | 69944 |
| NA | ENSMUSG00 | 66140 |
| NA | ENSMUSG00 | 270906 |
| NA | ENSMUSG00 | 74133 |
| NA | ENSMUSG00 | 108660 |
| NA | ENSMUSG00 | 56427 |
| NA | ENSMUSG00 | 216767 |
| NA | ENSMUSG00 | 69125 |
| NA | ENSMUSG00 | 72508 |
| NA | ENSMUSG00 | 50724 |
| NA | ENSMUSG00 | 171212 |
| NA | ENSMUSG00 | 76892 |
| NA | ENSMUSG00 | 216760 |
| NA | ENSMUSG00 | 67726 |
| NA | ENSMUSG00 | 53892 |
| NA | ENSMUSG00 | 448850 |
| NA | ENSMUSG00 | 66196 |
| NA | ENSMUSG00 | 212627 |
| NA | ENSMUSG00 | 217039 |
| NA | ENSMUSG00 | 107476 |

| | | |
|---|---|---|
| NA | ENSMUSG00 | 20425 |
| NA | ENSMUSG00 | 16897 |
| NA | ENSMUSG00 | 13495 |
| NA | ENSMUSG00 | 20787 |
| NA | ENSMUSG00 | 71943 |
| NA | ENSMUSG00 | 69802 |
| NA | ENSMUSG00 | 20913 |
| NA | ENSMUSG00 | 66912 |
| NA | ENSMUSG00 | 68626 |
| NA | ENSMUSG00 | 18559 |
| NA | ENSMUSG00 | 28071 |
| NA | ENSMUSG00 | 380753 |
| NA | ENSMUSG00 | 19027 |
| NA | ENSMUSG00 | 71853 |
| NA | ENSMUSG00 | 59027 |
| NA | ENSMUSG00 | 30955 |
| NA | ENSMUSG00 | 71169 |
| NA | ENSMUSG00 | 66109 |
| NA | ENSMUSG00 | 19878 |
| NA | ENSMUSG00 | 17182 |
| NA | ENSMUSG00 | 17775 |
| NA | ENSMUSG00 | 76820 |
| NA | ENSMUSG00 | 217463 |
| NA | ENSMUSG00 | 20969 |
| NA | ENSMUSG00 | 14245 |
| NA | ENSMUSG00 | 80913 |
| NA | ENSMUSG00 | 319504 |
| NA | ENSMUSG00 | 217410 |
| NA | ENSMUSG00 | 74008 |
| NA | ENSMUSG00 | 58240 |
| NA | ENSMUSG00 | 105005 |
| NA | ENSMUSG00 | 67241 |
| NA | ENSMUSG00 | 13929 |
| NA | ENSMUSG00 | 14674 |
| NA | ENSMUSG00 | 19084 |
| NA | ENSMUSG00 | 105014 |
| Significant | ENSMUSG00 | 26399 |
| NA | ENSMUSG00 | 217449 |
| NA | ENSMUSG00 | 104923 |
| NA | ENSMUSG00 | 19819 |
| NA | ENSMUSG00 | 217379 |
| NA | ENSMUSG00 | 14226 |
| NA | ENSMUSG00 | 22169 |
| NA | ENSMUSG00 | 382562 |
| NA | ENSMUSG00 | 20403 |
| NA | ENSMUSG00 | 58185 |

| | | |
|---|---|---|
| NA | ENSMUSG00 | 108089 |
| NA | ENSMUSG00 | 15902 |
| NA | ENSMUSG00 | 67216 |
| NA | ENSMUSG00 | 17977 |
| NA | ENSMUSG00 | 71916 |
| NA | ENSMUSG00 | 20135 |
| NA | ENSMUSG00 | 12033 |
| NA | ENSMUSG00 | 52504 |
| NA | ENSMUSG00 | 194655 |
| NA | ENSMUSG00 | 104111 |
| NA | ENSMUSG00 | 195733 |
| NA | ENSMUSG00 | 217378 |
| NA | ENSMUSG00 | 668212 |
| NA | ENSMUSG00 | 104836 |
| NA | ENSMUSG00 | 18976 |
| NA | ENSMUSG00 | 13435 |
| NA | ENSMUSG00 | 13382 |
| NA | ENSMUSG00 | 16570 |
| NA | ENSMUSG00 | 24057 |
| NA | ENSMUSG00 | 19325 |
| NA | ENSMUSG00 | 69675 |
| NA | ENSMUSG00 | 78394 |
| NA | ENSMUSG00 | 21410 |
| NA | ENSMUSG00 | 70439 |
| NA | ENSMUSG00 | 118453 |
| NA | ENSMUSG00 | 217232 |
| NA | ENSMUSG00 | 16416 |
| NA | ENSMUSG00 | 52686 |
| NA | ENSMUSG00 | 217011 |
| NA | ENSMUSG00 | 24086 |
| NA | ENSMUSG00 | 67338 |
| NA | ENSMUSG00 | 16882 |
| NA | ENSMUSG00 | 26406 |
| NA | ENSMUSG00 | 72047 |
| NA | ENSMUSG00 | 56095 |
| NA | ENSMUSG00 | 71956 |
| NA | ENSMUSG00 | 19184 |
| NA | ENSMUSG00 | 216991 |
| NA | ENSMUSG00 | 75870 |
| NA | ENSMUSG00 | 78943 |
| NA | ENSMUSG00 | 18015 |
| NA | ENSMUSG00 | 18613 |
| NA | ENSMUSG00 | 50776 |
| NA | ENSMUSG00 | 13207 |
| NA | ENSMUSG00 | 66997 |
| NA | ENSMUSG00 | 78455 |

| | | |
|---|---|---|
| NA | ENSMUSG00 | 76380 |
| NA | ENSMUSG00 | 26941 |
| NA | ENSMUSG00 | 50773 |
| NA | ENSMUSG00 | 15374 |
| NA | ENSMUSG00 | 170930 |
| NA | ENSMUSG00 | 445007 |
| NA | ENSMUSG00 | 260302 |
| NA | ENSMUSG00 | 74148 |
| NA | ENSMUSG00 | 69674 |
| NA | ENSMUSG00 | 67283 |
| NA | ENSMUSG00 | 18472 |
| NA | ENSMUSG00 | 71947 |
| NA | ENSMUSG00 | 170472 |
| NA | ENSMUSG00 | 57230 |
| NA | ENSMUSG00 | 14635 |
| NA | ENSMUSG00 | 217331 |
| Significant | ENSMUSG00 | 217333 |
| Significant | ENSMUSG00 | 11484 |
| NA | ENSMUSG00 | 60441 |
| NA | ENSMUSG00 | 217335 |
| NA | ENSMUSG00 | 11430 |
| NA | ENSMUSG00 | 69535 |
| NA | ENSMUSG00 | 217337 |
| NA | ENSMUSG00 | 76265 |
| NA | ENSMUSG00 | 217325 |
| NA | ENSMUSG00 | 66874 |
| Significant | ENSMUSG00 | 55984 |
| Significant | ENSMUSG00 | 18436 |
| Significant | ENSMUSG00 | 53313 |
| NA | ENSMUSG00 | 11736 |
| NA | ENSMUSG00 | 53413 |
| NA | ENSMUSG00 | 67128 |
| Significant | ENSMUSG00 | 77577 |
| NA | ENSMUSG00 | 67279 |
| NA | ENSMUSG00 | 217342 |
| NA | ENSMUSG00 | 52700 |
| NA | ENSMUSG00 | 217344 |
| NA | ENSMUSG00 | 74477 |
| NA | ENSMUSG00 | 109212 |
| Significant | ENSMUSG00 | 67622 |
| NA | ENSMUSG00 | 54189 |
| Significant | ENSMUSG00 | 69900 |
| NA | ENSMUSG00 | 74136 |
| NA | ENSMUSG00 | 50932 |
| Significant | ENSMUSG00 | 18806 |
| Significant | ENSMUSG00 | 52466 |

| | | |
|---|---|---|
| NA | ENSMUSG00 | 104457 |
| NA | ENSMUSG00 | 57837 |
| NA | ENSMUSG00 | 70451 |
| NA | ENSMUSG00 | 216961 |
| NA | ENSMUSG00 | 104184 |
| NA | ENSMUSG00 | 12874 |
| NA | ENSMUSG00 | 56322 |
| NA | ENSMUSG00 | 18230 |
| NA | ENSMUSG00 | 76566 |
| NA | ENSMUSG00 | 22627 |
| NA | ENSMUSG00 | 192159 |
| NA | ENSMUSG00 | 18103 |
| NA | ENSMUSG00 | 70834 |
| NA | ENSMUSG00 | 67684 |
| NA | ENSMUSG00 | 71452 |
| NA | ENSMUSG00 | 76408 |
| NA | ENSMUSG00 | 217119 |
| NA | ENSMUSG00 | 98238 |
| NA | ENSMUSG00 | 110172 |
| NA | ENSMUSG00 | 74479 |
| NA | ENSMUSG00 | 217140 |
| NA | ENSMUSG00 | 72194 |
| NA | ENSMUSG00 | 13385 |
| NA | ENSMUSG00 | 13543 |
| NA | ENSMUSG00 | 217166 |
| NA | ENSMUSG00 | 18626 |
| NA | ENSMUSG00 | 22318 |
| NA | ENSMUSG00 | 66910 |
| NA | ENSMUSG00 | 20877 |
| NA | ENSMUSG00 | 68964 |
| NA | ENSMUSG00 | 237823 |
| NA | ENSMUSG00 | 77579 |
| NA | ENSMUSG00 | 320207 |
| NA | ENSMUSG00 | 55943 |
| NA | ENSMUSG00 | 66358 |
| NA | ENSMUSG00 | 21973 |
| NA | ENSMUSG00 | 104112 |
| NA | ENSMUSG00 | 14534 |
| NA | ENSMUSG00 | 20851 |
| NA | ENSMUSG00 | 76547 |
| NA | ENSMUSG00 | 268490 |
| NA | ENSMUSG00 | 21429 |
| NA | ENSMUSG00 | 52715 |
| NA | ENSMUSG00 | 20624 |
| NA | ENSMUSG00 | 68087 |
| NA | ENSMUSG00 | 18107 |

| | | |
|---|---|---|
| NA | ENSMUSG00 | 72469 |
| NA | ENSMUSG00 | 53859 |
| NA | ENSMUSG00 | 56494 |
| NA | ENSMUSG00 | 66689 |
| NA | ENSMUSG00 | 30795 |
| NA | ENSMUSG00 | 76983 |
| NA | ENSMUSG00 | 94186 |
| NA | ENSMUSG00 | 11782 |
| NA | ENSMUSG00 | 328092 |
| NA | ENSMUSG00 | 108800 |
| NA | ENSMUSG00 | 83602 |
| NA | ENSMUSG00 | 20338 |
| NA | ENSMUSG00 | 109065 |
| NA | ENSMUSG00 | 18974 |
| NA | ENSMUSG00 | 69554 |
| NA | ENSMUSG00 | 66244 |
| NA | ENSMUSG00 | 20334 |
| NA | ENSMUSG00 | 217666 |
| NA | ENSMUSG00 | 78232 |
| NA | ENSMUSG00 | 18949 |
| NA | ENSMUSG00 | 338320 |
| NA | ENSMUSG00 | 14420 |
| NA | ENSMUSG00 | 104871 |
| NA | ENSMUSG00 | 24000 |
| NA | ENSMUSG00 | 75553 |
| NA | ENSMUSG00 | 76260 |
| NA | ENSMUSG00 | 245841 |
| NA | ENSMUSG00 | 59032 |
| NA | ENSMUSG00 | 66132 |
| NA | ENSMUSG00 | 26443 |
| NA | ENSMUSG00 | 18035 |
| NA | ENSMUSG00 | 56784 |
| NA | ENSMUSG00 | 217588 |
| NA | ENSMUSG00 | 14874 |
| NA | ENSMUSG00 | 20773 |
| NA | ENSMUSG00 | 217737 |
| NA | ENSMUSG00 | 104799 |
| NA | ENSMUSG00 | 66354 |
| NA | ENSMUSG00 | 380773 |
| NA | ENSMUSG00 | 72113 |
| NA | ENSMUSG00 | 664883 |
| NA | ENSMUSG00 | 108156 |
| NA | ENSMUSG00 | 26932 |
| NA | ENSMUSG00 | 81535 |
| NA | ENSMUSG00 | 238276 |
| NA | ENSMUSG00 | 20741 |

| | | |
|---|---|---|
| NA | ENSMUSG00 | 53618 |
| NA | ENSMUSG00 | 73991 |
| NA | ENSMUSG00 | 64010 |
| NA | ENSMUSG00 | 18080 |
| NA | ENSMUSG00 | 110095 |
| NA | ENSMUSG00 | 72736 |
| NA | ENSMUSG00 | 56444 |
| NA | ENSMUSG00 | 30056 |
| NA | ENSMUSG00 | 104001 |
| NA | ENSMUSG00 | 78257 |
| NA | ENSMUSG00 | 66375 |
| NA | ENSMUSG00 | 19042 |
| NA | ENSMUSG00 | 75801 |
| NA | ENSMUSG00 | 73046 |
| NA | ENSMUSG00 | 17420 |
| NA | ENSMUSG00 | 18755 |
| NA | ENSMUSG00 | 15251 |
| NA | ENSMUSG00 | 18789 |
| NA | ENSMUSG00 | 56217 |
| NA | ENSMUSG00 | 75627 |
| NA | ENSMUSG00 | 73834 |
| NA | ENSMUSG00 | 22367 |
| NA | ENSMUSG00 | 13665 |
| NA | ENSMUSG00 | 110417 |
| NA | ENSMUSG00 | 77974 |
| NA | ENSMUSG00 | 53612 |
| NA | ENSMUSG00 | 11847 |
| NA | ENSMUSG00 | 12192 |
| NA | ENSMUSG00 | 13877 |
| NA | ENSMUSG00 | 217684 |
| NA | ENSMUSG00 | 20384 |
| NA | ENSMUSG00 | 64075 |
| NA | ENSMUSG00 | 105940408 |
| NA | ENSMUSG00 | 54604 |
| NA | ENSMUSG00 | 217893 |
| NA | ENSMUSG00 | 116870 |
| NA | ENSMUSG00 | 207615 |
| NA | ENSMUSG00 | 69237 |
| NA | ENSMUSG00 | 66505 |
| NA | ENSMUSG00 | 52635 |
| NA | ENSMUSG00 | 217946 |
| NA | ENSMUSG00 | 78767 |
| NA | ENSMUSG00 | 104884 |
| NA | ENSMUSG00 | 19179 |
| NA | ENSMUSG00 | 217827 |
| NA | ENSMUSG00 | 73086 |

| | | |
|---|---|---|
| NA | ENSMUSG00 | 68339 |
| NA | ENSMUSG00 | 217830 |
| NA | ENSMUSG00 | 23876 |
| NA | ENSMUSG00 | 109181 |
| NA | ENSMUSG00 | 110616 |
| Significant | ENSMUSG00 | 19141 |
| NA | ENSMUSG00 | 27277 |
| NA | ENSMUSG00 | 69617 |
| NA | ENSMUSG00 | 56421 |
| Significant | ENSMUSG00 | 65256 |
| NA | ENSMUSG00 | 68149 |
| NA | ENSMUSG00 | 622402 |
| NA | ENSMUSG00 | 27384 |
| NA | ENSMUSG00 | 56349 |
| NA | ENSMUSG00 | 243931 |
| NA | ENSMUSG00 | 14569 |
| NA | ENSMUSG00 | 73828 |
| NA | ENSMUSG00 | 18222 |
| NA | ENSMUSG00 | 171210 |
| NA | ENSMUSG00 | 217705 |
| NA | ENSMUSG00 | 217707 |
| NA | ENSMUSG00 | 12499 |
| NA | ENSMUSG00 | 104776 |
| NA | ENSMUSG00 | 19300 |
| NA | ENSMUSG00 | 74316 |
| NA | ENSMUSG00 | 67963 |
| NA | ENSMUSG00 | 73736 |
| NA | ENSMUSG00 | 56531 |
| NA | ENSMUSG00 | 217716 |
| NA | ENSMUSG00 | 68581 |
| NA | ENSMUSG00 | 14281 |
| NA | ENSMUSG00 | 58520 |
| NA | ENSMUSG00 | 21809 |
| NA | ENSMUSG00 | 70373 |
| NA | ENSMUSG00 | 68737 |
| NA | ENSMUSG00 | 12454 |
| NA | ENSMUSG00 | 14026 |
| NA | ENSMUSG00 | 22632 |
| NA | ENSMUSG00 | 214663 |
| NA | ENSMUSG00 | 22375 |
| NA | ENSMUSG00 | 15519 |
| NA | ENSMUSG00 | 72805 |
| NA | ENSMUSG00 | 14137 |
| NA | ENSMUSG00 | 104859 |
| NA | ENSMUSG00 | 67236 |
| NA | ENSMUSG00 | 22031 |

| | | |
|---|---|---|
| NA | ENSMUSG00 | 217866 |
| NA | ENSMUSG00 | 21928 |
| NA | ENSMUSG00 | 217869 |
| NA | ENSMUSG00 | 21981 |
| NA | ENSMUSG00 | 68520 |
| NA | ENSMUSG00 | 74335 |
| NA | ENSMUSG00 | 16593 |
| NA | ENSMUSG00 | 70257 |
| NA | ENSMUSG00 | 56696 |
| NA | ENSMUSG00 | 14593 |
| NA | ENSMUSG00 | 83924 |
| NA | ENSMUSG00 | 238505 |
| NA | ENSMUSG00 | 14634 |
| NA | ENSMUSG00 | 27052 |
| NA | ENSMUSG00 | 19720 |
| NA | ENSMUSG00 | 72739 |
| NA | ENSMUSG00 | 68732 |
| NA | ENSMUSG00 | 380836 |
| NA | ENSMUSG00 | 14756 |
| NA | ENSMUSG00 | 16364 |
| NA | ENSMUSG00 | 66482 |
| NA | ENSMUSG00 | 14538 |
| NA | ENSMUSG00 | 66154 |
| NA | ENSMUSG00 | 107889 |
| NA | ENSMUSG00 | 54326 |
| Significant | ENSMUSG00 | 18003 |
| NA | ENSMUSG00 | 110521 |
| NA | ENSMUSG00 | 13614 |
| NA | ENSMUSG00 | 67046 |
| NA | ENSMUSG00 | 76137 |
| NA | ENSMUSG00 | 218210 |
| NA | ENSMUSG00 | 16553 |
| NA | ENSMUSG00 | 22017 |
| NA | ENSMUSG00 | 110052 |
| NA | ENSMUSG00 | 66329 |
| NA | ENSMUSG00 | 75678 |
| NA | ENSMUSG00 | 66336 |
| NA | ENSMUSG00 | 70930 |
| NA | ENSMUSG00 | 20729 |
| NA | ENSMUSG00 | 78903 |
| NA | ENSMUSG00 | 19766 |
| Significant | ENSMUSG00 | 66895 |
| NA | ENSMUSG00 | 19134 |
| NA | ENSMUSG00 | 23986 |
| NA | ENSMUSG00 | 208366 |
| NA | ENSMUSG00 | 69955 |

| | | |
|---|---|---|
| NA | ENSMUSG00 | 17084 |
| NA | ENSMUSG00 | 107513 |
| NA | ENSMUSG00 | 71340 |
| NA | ENSMUSG00 | 67797 |
| NA | ENSMUSG00 | 108652 |
| NA | ENSMUSG00 | 20354 |
| NA | ENSMUSG00 | 23882 |
| NA | ENSMUSG00 | 14120 |
| NA | ENSMUSG00 | 20963 |
| NA | ENSMUSG00 | 72061 |
| NA | ENSMUSG00 | 11992 |
| NA | ENSMUSG00 | 14088 |
| NA | ENSMUSG00 | 19206 |
| NA | ENSMUSG00 | 268656 |
| NA | ENSMUSG00 | 76251 |
| NA | ENSMUSG00 | 14057 |
| NA | ENSMUSG00 | 56541 |
| Significant | ENSMUSG00 | 13039 |
| NA | ENSMUSG00 | 26919 |
| NA | ENSMUSG00 | 66129 |
| NA | ENSMUSG00 | 105278 |
| NA | ENSMUSG00 | 66890 |
| NA | ENSMUSG00 | 17121 |
| NA | ENSMUSG00 | 66494 |
| NA | ENSMUSG00 | 18193 |
| NA | ENSMUSG00 | 67399 |
| NA | ENSMUSG00 | 72935 |
| NA | ENSMUSG00 | 212483 |
| NA | ENSMUSG00 | 72562 |
| NA | ENSMUSG00 | 69672 |
| NA | ENSMUSG00 | 212880 |
| NA | ENSMUSG00 | 12328 |
| NA | ENSMUSG00 | 218271 |
| NA | ENSMUSG00 | 212281 |
| NA | ENSMUSG00 | 170936 |
| NA | ENSMUSG00 | 19210 |
| NA | ENSMUSG00 | 66410 |
| NA | ENSMUSG00 | 67530 |
| NA | ENSMUSG00 | 69577 |
| NA | ENSMUSG00 | 12626 |
| NA | ENSMUSG00 | 17129 |
| NA | ENSMUSG00 | 15387 |
| NA | ENSMUSG00 | 66671 |
| NA | ENSMUSG00 | 218397 |
| NA | ENSMUSG00 | 70153 |
| NA | ENSMUSG00 | 56278 |

| | | |
|---|---|---|
| NA | ENSMUSG00 | 78689 |
| NA | ENSMUSG00 | 105348 |
| NA | ENSMUSG00 NA | |
| Significant | ENSMUSG00 | 69635 |
| NA | ENSMUSG00 | 69716 |
| NA | ENSMUSG00 | 74470 |
| NA | ENSMUSG00 | 72948 |
| NA | ENSMUSG00 | 11624 |
| NA | ENSMUSG00 | 18570 |
| NA | ENSMUSG00 | 66945 |
| NA | ENSMUSG00 | 67433 |
| NA | ENSMUSG00 | 80898 |
| NA | ENSMUSG00 | 12380 |
| NA | ENSMUSG00 | 73296 |
| NA | ENSMUSG00 | 93692 |
| NA | ENSMUSG00 | 77041 |
| NA | ENSMUSG00 | 78925 |
| NA | ENSMUSG00 | 28114 |
| NA | ENSMUSG00 | 78771 |
| NA | ENSMUSG00 | 105377 |
| NA | ENSMUSG00 | 28077 |
| NA | ENSMUSG00 | 407785 |
| NA | ENSMUSG00 | 94066 |
| NA | ENSMUSG00 | 210992 |
| NA | ENSMUSG00 | 218335 |
| NA | ENSMUSG00 | 21752 |
| NA | ENSMUSG00 | 12950 |
| Significant | ENSMUSG00 | 13003 |
| NA | ENSMUSG00 | 108138 |
| NA | ENSMUSG00 | 66795 |
| NA | ENSMUSG00 | 69085 |
| NA | ENSMUSG00 | 17079 |
| NA | ENSMUSG00 | 69048 |
| NA | ENSMUSG00 | 19356 |
| NA | ENSMUSG00 | 218518 |
| NA | ENSMUSG00 | 18260 |
| NA | ENSMUSG00 | 23894 |
| NA | ENSMUSG00 | 20365 |
| NA | ENSMUSG00 | 20595 |
| NA | ENSMUSG00 | 78038 |
| NA | ENSMUSG00 | 68927 |
| NA | ENSMUSG00 | 218490 |
| NA | ENSMUSG00 | 68558 |
| NA | ENSMUSG00 | 110596 |
| NA | ENSMUSG00 | 15212 |
| NA | ENSMUSG00 | 320806 |

| | | |
|---|---|---|
| NA | ENSMUSG00 | 27015 |
| NA | ENSMUSG00 | 68018 |
| NA | ENSMUSG00 | 15357 |
| NA | ENSMUSG00 | 67463 |
| NA | ENSMUSG00 | 544963 |
| NA | ENSMUSG00 | 66549 |
| NA | ENSMUSG00 | 218461 |
| NA | ENSMUSG00 | 11774 |
| NA | ENSMUSG00 | 107767 |
| NA | ENSMUSG00 | 57748 |
| NA | ENSMUSG00 | 66254 |
| NA | ENSMUSG00 | 16563 |
| NA | ENSMUSG00 | 71991 |
| Significant | ENSMUSG00 | 74559 |
| NA | ENSMUSG00 | 218581 |
| NA | ENSMUSG00 | 238871 |
| NA | ENSMUSG00 | 20620 |
| NA | ENSMUSG00 | 218442 |
| NA | ENSMUSG00 | 382793 |
| NA | ENSMUSG00 | 218441 |
| NA | ENSMUSG00 | 13361 |
| NA | ENSMUSG00 | 59079 |
| NA | ENSMUSG00 | 75805 |
| NA | ENSMUSG00 | 66975 |
| NA | ENSMUSG00 | 81003 |
| NA | ENSMUSG00 | 238831 |
| NA | ENSMUSG00 | 60411 |
| NA | ENSMUSG00 | 67285 |
| NA | ENSMUSG00 | 67288 |
| Significant | ENSMUSG00 | 52882 |
| Significant | ENSMUSG00 | 71816 |
| Significant | ENSMUSG00 | 52552 |
| Significant | ENSMUSG00 | 13723 |
| NA | ENSMUSG00 | 59054 |
| NA | ENSMUSG00 | 218756 |
| NA | ENSMUSG00 | 66413 |
| NA | ENSMUSG00 | 246103 |
| NA | ENSMUSG00 NA | |
| NA | ENSMUSG00 | 68263 |
| NA | ENSMUSG00 | 71393 |
| NA | ENSMUSG00 | 26401 |
| NA | ENSMUSG00 | 16195 |
| NA | ENSMUSG00 | 13206 |
| NA | ENSMUSG00 | 19012 |
| NA | ENSMUSG00 | 69590 |
| NA | ENSMUSG00 | 17993 |

| | | |
|---|---|---|
| NA | ENSMUSG00 | 54169 |
| NA | ENSMUSG00 | 67630 |
| NA | ENSMUSG00 | 22334 |
| NA | ENSMUSG00 | 69721 |
| NA | ENSMUSG00 | 69156 |
| NA | ENSMUSG00 | 22194 |
| NA | ENSMUSG00 | 353187 |
| NA | ENSMUSG00 | 21834 |
| NA | ENSMUSG00 | 59007 |
| NA | ENSMUSG00 | 71147 |
| NA | ENSMUSG00 | 70564 |
| NA | ENSMUSG00 | 14661 |
| NA | ENSMUSG00 | 12166 |
| Significant | ENSMUSG00 | 18074 |
| NA | ENSMUSG00 | 68045 |
| NA | ENSMUSG00 | 67725 |
| NA | ENSMUSG00 | 70601 |
| NA | ENSMUSG00 | 108671 |
| NA | ENSMUSG00 | 11750 |
| NA | ENSMUSG00 | 19056 |
| NA | ENSMUSG00 | 268721 |
| NA | ENSMUSG00 | 12325 |
| Significant | ENSMUSG00 | 18792 |
| NA | ENSMUSG00 | 22330 |
| NA | ENSMUSG00 | 55946 |
| NA | ENSMUSG00 | 70561 |
| NA | ENSMUSG00 | 50527 |
| NA | ENSMUSG00 | 67089 |
| NA | ENSMUSG00 | 12159 |
| NA | ENSMUSG00 | 74480 |
| NA | ENSMUSG00 | 218975 |
| NA | ENSMUSG00 | 16709 |
| NA | ENSMUSG00 | NA |
| NA | ENSMUSG00 | 11744 |
| NA | ENSMUSG00 | NA |
| NA | ENSMUSG00 | 105675 |
| NA | ENSMUSG00 | 83997 |
| NA | ENSMUSG00 | 75019 |
| Significant | ENSMUSG00 | 58809 |
| NA | ENSMUSG00 | 11843 |
| Significant | ENSMUSG00 | 78416 |
| NA | ENSMUSG00 | 56794 |
| NA | ENSMUSG00 | 14744 |
| NA | ENSMUSG00 | 74427 |
| NA | ENSMUSG00 | 67011 |
| Significant | ENSMUSG00 | 24056 |

| | | |
|---|---|---|
| NA | ENSMUSG00 | 12339 |
| NA | ENSMUSG00 | 71704 |
| NA | ENSMUSG00 | 26363 |
| NA | ENSMUSG00 | 104416 |
| NA | ENSMUSG00 | 71838 |
| NA | ENSMUSG00 | 105638 |
| NA | ENSMUSG00 | 218885 |
| NA | ENSMUSG00 | 64652 |
| NA | ENSMUSG00 | 26430 |
| NA | ENSMUSG00 | 239017 |
| NA | ENSMUSG00 | 76485 |
| NA | ENSMUSG00 | 69019 |
| NA | ENSMUSG00 | 23955 |
| NA | ENSMUSG00 | 68177 |
| NA | ENSMUSG00 | 16648 |
| NA | ENSMUSG00 | 66674 |
| NA | ENSMUSG00 | 67153 |
| NA | ENSMUSG00 | 26419 |
| NA | ENSMUSG00 | 66645 |
| Significant | ENSMUSG00 | 13030 |
| NA | ENSMUSG00 | 76007 |
| NA | ENSMUSG00 | 68631 |
| NA | ENSMUSG00 | 18753 |
| NA | ENSMUSG00 | 68043 |
| NA | ENSMUSG00 | 57258 |
| NA | ENSMUSG00 | 21881 |
| NA | ENSMUSG00 | 72400 |
| NA | ENSMUSG00 | 50523 |
| NA | ENSMUSG00 | 75901 |
| NA | ENSMUSG00 | 20220 |
| NA | ENSMUSG00 | 219114 |
| NA | ENSMUSG00 | 67840 |
| NA | ENSMUSG00 | 75965 |
| NA | ENSMUSG00 | 219150 |
| NA | ENSMUSG00 | 68514 |
| NA | ENSMUSG00 | 210925 |
| NA | ENSMUSG00 | 54616 |
| NA | ENSMUSG00 | 69008 |
| NA | ENSMUSG00 | 71891 |
| Significant | ENSMUSG00 NA | |
| NA | ENSMUSG00 | 219135 |
| Significant | ENSMUSG00 | 219140 |
| NA | ENSMUSG00 | 70478 |
| NA | ENSMUSG00 | 13885 |
| NA | ENSMUSG00 | 18826 |
| NA | ENSMUSG00 | 67302 |

| | | |
|---|---|---|
| NA | ENSMUSG00 | 67554 |
| NA | ENSMUSG00 | 67467 |
| NA | ENSMUSG00 | 27275 |
| NA | ENSMUSG00 | 21807 |
| NA | ENSMUSG00 | 66148 |
| NA | ENSMUSG00 | 108670 |
| NA | ENSMUSG00 | 219181 |
| Significant | ENSMUSG00 | 66214 |
| NA | ENSMUSG00 | 219249 |
| NA | ENSMUSG00 | 66897 |
| NA | ENSMUSG00 | 56419 |
| NA | ENSMUSG00 | 211253 |
| NA | ENSMUSG00 | 22380 |
| NA | ENSMUSG00 | 67955 |
| Significant | ENSMUSG00 | 380924 |
| NA | ENSMUSG00 | 74195 |
| NA | ENSMUSG00 | 52033 |
| NA | ENSMUSG00 | 71988 |
| NA | ENSMUSG00 | 67179 |
| NA | ENSMUSG00 | 12759 |
| NA | ENSMUSG00 | 66854 |
| NA | ENSMUSG00 | 12934 |
| NA | ENSMUSG00 | 12177 |
| NA | ENSMUSG00 | 71978 |
| NA | ENSMUSG00 | 18040 |
| NA | ENSMUSG00 | 18039 |
| NA | ENSMUSG00 | 52023 |
| NA | ENSMUSG00 | 77744 |
| NA | ENSMUSG00 | 21933 |
| NA | ENSMUSG00 | 246710 |
| NA | ENSMUSG00 | 57784 |
| NA | ENSMUSG00 | 213019 |
| NA | ENSMUSG00 | 20410 |
| NA | ENSMUSG00 | 19057 |
| NA | ENSMUSG00 | 213053 |
| NA | ENSMUSG00 | 239170 |
| NA | ENSMUSG00 | 12153 |
| NA | ENSMUSG00 | 65246 |
| NA | ENSMUSG00 | 13449 |
| Significant | ENSMUSG00 | 19645 |
| NA | ENSMUSG00 | 105670 |
| Significant | ENSMUSG00 | 16432 |
| NA | ENSMUSG00 | 67381 |
| NA | ENSMUSG00 | 20916 |
| NA | ENSMUSG00 | 50933 |
| NA | ENSMUSG00 | 24064 |

| | | |
|---|---|---|
| NA | ENSMUSG00 | 74213 |
| NA | ENSMUSG00 | 72486 |
| Significant | ENSMUSG00 | 13618 |
| NA | ENSMUSG00 | 64929 |
| NA | ENSMUSG00 | 50789 |
| NA | ENSMUSG00 | 211286 |
| NA | ENSMUSG00 | 16365 |
| NA | ENSMUSG00 | 13190 |
| NA | ENSMUSG00 | 76355 |
| NA | ENSMUSG00 | 58245 |
| NA | ENSMUSG00 | 100037258 |
| NA | ENSMUSG00 | 105559 |
| NA | ENSMUSG00 | 71175 |
| NA | ENSMUSG00 | 170762 |
| NA | ENSMUSG00 | 23880 |
| Significant | ENSMUSG00 | 13132 |
| NA | ENSMUSG00 | 67515 |
| NA | ENSMUSG00 | 76338 |
| NA | ENSMUSG00 | 56335 |
| NA | ENSMUSG00 | 13135 |
| NA | ENSMUSG00 | 65107 |
| NA | ENSMUSG00 | 219072 |
| NA | ENSMUSG00 | 16475 |
| NA | ENSMUSG00 | 74359 |
| NA | ENSMUSG00 | 50934 |
| NA | ENSMUSG00 | 106052 |
| NA | ENSMUSG00 | 56215 |
| NA | ENSMUSG00 | 67041 |
| NA | ENSMUSG00 | 14000 |
| NA | ENSMUSG00 | 19173 |
| NA | ENSMUSG00 | 54196 |
| NA | ENSMUSG00 | 77877 |
| NA | ENSMUSG00 | 68070 |
| NA | ENSMUSG00 | 59049 |
| NA | ENSMUSG00 | 66629 |
| NA | ENSMUSG00 | 22763 |
| NA | ENSMUSG00 | 68966 |
| NA | ENSMUSG00 | 20024 |
| NA | ENSMUSG00 | 28200 |
| NA | ENSMUSG00 | 28199 |
| NA | ENSMUSG00 | 68680 |
| NA | ENSMUSG00 | 19186 |
| NA | ENSMUSG00 | 85308 |
| NA | ENSMUSG00 | 21816 |
| NA | ENSMUSG00 | 12684 |
| NA | ENSMUSG00 | 104110 |

| | | |
|---|---|---|
| NA | ENSMUSG00 | 56532 |
| NA | ENSMUSG00 | 654795 |
| NA | ENSMUSG00 | 432731 |
| NA | ENSMUSG00 | 12465 |
| Significant | ENSMUSG00 | 69574 |
| NA | ENSMUSG00 | 252967 |
| NA | ENSMUSG00 | 67434 |
| NA | ENSMUSG00 | 110960 |
| NA | ENSMUSG00 | 17117 |
| NA | ENSMUSG00 | 75646 |
| NA | ENSMUSG00 | 67832 |
| NA | ENSMUSG00 | 19355 |
| NA | ENSMUSG00 | 68646 |
| NA | ENSMUSG00 | 67154 |
| Significant | ENSMUSG00 | 114128 |
| NA | ENSMUSG00 | 15529 |
| NA | ENSMUSG00 | 223435 |
| NA | ENSMUSG00 | 11732 |
| NA | ENSMUSG00 | 66270 |
| NA | ENSMUSG00 | 17909 |
| NA | ENSMUSG00 | 30945 |
| Significant | ENSMUSG00 | 18458 |
| NA | ENSMUSG00 | 22631 |
| NA | ENSMUSG00 | 252973 |
| NA | ENSMUSG00 | 382985 |
| NA | ENSMUSG00 | 66335 |
| NA | ENSMUSG00 | 69906 |
| NA | ENSMUSG00 | 223499 |
| NA | ENSMUSG00 | 75766 |
| NA | ENSMUSG00 | 239393 |
| NA | ENSMUSG00 | 170719 |
| NA | ENSMUSG00 | 11600 |
| NA | ENSMUSG00 | 68135 |
| NA | ENSMUSG00 | 78581 |
| NA | ENSMUSG00 | 19357 |
| NA | ENSMUSG00 | 20419 |
| NA | ENSMUSG00 | 15473 |
| NA | ENSMUSG00 | 17181 |
| NA | ENSMUSG00 | 67724 |
| NA | ENSMUSG00 | 56274 |
| NA | ENSMUSG00 | 13992 |
| NA | ENSMUSG00 | 380993 |
| NA | ENSMUSG00 | 16341 |
| NA | ENSMUSG00 | 66736 |
| NA | ENSMUSG00 | 223527 |
| NA | ENSMUSG00 | 55960 |

| | | |
|---|---|---|
| NA | ENSMUSG00 | 17869 |
| NA | ENSMUSG00 | 223593 |
| NA | ENSMUSG00 | 20775 |
| Significant | ENSMUSG00 | 211401 |
| NA | ENSMUSG00 | 66218 |
| NA | ENSMUSG00 | 67731 |
| NA | ENSMUSG00 | 76773 |
| NA | ENSMUSG00 | 70472 |
| NA | ENSMUSG00 | 22770 |
| NA | ENSMUSG00 | NA |
| NA | ENSMUSG00 | 210544 |
| NA | ENSMUSG00 | 67819 |
| NA | ENSMUSG00 | 105837 |
| NA | ENSMUSG00 | 68537 |
| NA | ENSMUSG00 | 12818 |
| NA | ENSMUSG00 | 20491 |
| NA | ENSMUSG00 | 13196 |
| NA | ENSMUSG00 | 223601 |
| NA | ENSMUSG00 | 29870 |
| NA | ENSMUSG00 | 72026 |
| NA | ENSMUSG00 | 223770 |
| NA | ENSMUSG00 | 21685 |
| NA | ENSMUSG00 | 20286 |
| NA | ENSMUSG00 | 19387 |
| NA | ENSMUSG00 | 214669 |
| NA | ENSMUSG00 | 56438 |
| NA | ENSMUSG00 | 321003 |
| NA | ENSMUSG00 | 70356 |
| NA | ENSMUSG00 | 20524 |
| NA | ENSMUSG00 | 11564 |
| NA | ENSMUSG00 | 213956 |
| NA | ENSMUSG00 | 239555 |
| NA | ENSMUSG00 | 66513 |
| NA | ENSMUSG00 | 20972 |
| NA | ENSMUSG00 | 97998 |
| NA | ENSMUSG00 | 54152 |
| NA | ENSMUSG00 | 72107 |
| NA | ENSMUSG00 | 18606 |
| NA | ENSMUSG00 | 74158 |
| NA | ENSMUSG00 | 223696 |
| NA | ENSMUSG00 | 73739 |
| NA | ENSMUSG00 | 13404 |
| NA | ENSMUSG00 | 27373 |
| NA | ENSMUSG00 | 73225 |
| NA | ENSMUSG00 | 20401 |
| NA | ENSMUSG00 | 68653 |

| | | |
|---|---|---|
| NA | ENSMUSG00 | 170736 |
| Significant | ENSMUSG00 | 64099 |
| NA | ENSMUSG00 | 72709 |
| NA | ENSMUSG00 | 319953 |
| NA | ENSMUSG00 | 17886 |
| NA | ENSMUSG00 | 67130 |
| Significant | ENSMUSG00 | 19230 |
| NA | ENSMUSG00 | 69029 |
| NA | ENSMUSG00 | 17939 |
| NA | ENSMUSG00 | 67760 |
| NA | ENSMUSG00 | 20788 |
| Significant | ENSMUSG00 | 69354 |
| NA | ENSMUSG00 | 71919 |
| NA | ENSMUSG00 | 223864 |
| NA | ENSMUSG00 | 14375 |
| NA | ENSMUSG00 | 28075 |
| NA | ENSMUSG00 | 29858 |
| NA | ENSMUSG00 | 56233 |
| NA | ENSMUSG00 | 78929 |
| NA | ENSMUSG00 | 11429 |
| Significant | ENSMUSG00 | 22337 |
| NA | ENSMUSG00 | 105855 |
| NA | ENSMUSG00 | 18574 |
| NA | ENSMUSG00 | 106200 |
| NA | ENSMUSG00 | 56722 |
| NA | ENSMUSG00 | 26425 |
| NA | ENSMUSG00 | 69053 |
| NA | ENSMUSG00 | 12053 |
| NA | ENSMUSG00 | 16180 |
| NA | ENSMUSG00 | 72615 |
| NA | ENSMUSG00 | 66911 |
| NA | ENSMUSG00 | 17237 |
| NA | ENSMUSG00 | 106393 |
| NA | ENSMUSG00 | 12914 |
| NA | ENSMUSG00 | 71591 |
| Significant | ENSMUSG00 | 15205 |
| NA | ENSMUSG00 | 74120 |
| NA | ENSMUSG00 | 224088 |
| NA | ENSMUSG00 | 54483 |
| NA | ENSMUSG00 | 74022 |
| Significant | ENSMUSG00 | 224090 |
| NA | ENSMUSG00 | 224092 |
| Significant | ENSMUSG00 | 66049 |
| NA | ENSMUSG00 NA | |
| NA | ENSMUSG00 | 70511 |
| NA | ENSMUSG00 | 50505 |

| | | |
|---|---|---|
| NA | ENSMUSG00 | 76282 |
| NA | ENSMUSG00 | 268822 |
| NA | ENSMUSG00 | 66445 |
| NA | ENSMUSG00 | 106025 |
| NA | ENSMUSG00 | 68877 |
| NA | ENSMUSG00 | 59053 |
| NA | ENSMUSG00 | 13350 |
| NA | ENSMUSG00 | 15499 |
| Significant | ENSMUSG00 | 12181 |
| NA | ENSMUSG00 | 223658 |
| NA | ENSMUSG00 | 30840 |
| NA | ENSMUSG00 | 52710 |
| NA | ENSMUSG00 | 14731 |
| Significant | ENSMUSG00 | 75475 |
| NA | ENSMUSG00 | 66168 |
| NA | ENSMUSG00 | 18810 |
| NA | ENSMUSG00 | 105782 |
| NA | ENSMUSG00 | 22122 |
| NA | ENSMUSG00 | 66194 |
| NA | ENSMUSG00 | 223646 |
| NA | ENSMUSG00 | 69146 |
| Significant | ENSMUSG00 | 100041546 |
| Significant | ENSMUSG00 | 17069 |
| NA | ENSMUSG00 | 271377 |
| NA | ENSMUSG00 | 11838 |
| NA | ENSMUSG00 | 74201 |
| NA | ENSMUSG00 | 14083 |
| NA | ENSMUSG00 | 105847 |
| NA | ENSMUSG00 | 72962 |
| NA | ENSMUSG00 | 12651 |
| NA | ENSMUSG00 | 11883 |
| NA | ENSMUSG00 | 68708 |
| NA | ENSMUSG00 | 16564 |
| NA | ENSMUSG00 | 67057 |
| NA | ENSMUSG00 | 67197 |
| NA | ENSMUSG00 | 11658 |
| NA | ENSMUSG00 | 208650 |
| NA | ENSMUSG00 | 70508 |
| NA | ENSMUSG00 | 58998 |
| NA | ENSMUSG00 | 67841 |
| NA | ENSMUSG00 | 74102 |
| NA | ENSMUSG00 | 67896 |
| NA | ENSMUSG00 | 57781 |
| NA | ENSMUSG00 | 66067 |
| NA | ENSMUSG00 | 72083 |
| NA | ENSMUSG00 | 19090 |

| | | |
|---|---|---|
| NA | ENSMUSG00 | 17217 |
| NA | ENSMUSG00 | 70620 |
| NA | ENSMUSG00 | 20583 |
| NA | ENSMUSG00 | 66086 |
| NA | ENSMUSG00 | 67203 |
| NA | ENSMUSG00 | 93734 |
| NA | ENSMUSG00 | 94184 |
| NA | ENSMUSG00 | 18203 |
| NA | ENSMUSG00 | 106298 |
| NA | ENSMUSG00 | 67118 |
| NA | ENSMUSG00 | 74108 |
| NA | ENSMUSG00 | 108105 |
| NA | ENSMUSG00 | 72117 |
| NA | ENSMUSG00 | 212153 |
| NA | ENSMUSG00 | 15260 |
| NA | ENSMUSG00 | 106248 |
| NA | ENSMUSG00 | 18100 |
| NA | ENSMUSG00 | 74185 |
| NA | ENSMUSG00 | 56490 |
| NA | ENSMUSG00 | 252870 |
| NA | ENSMUSG00 | 54128 |
| NA | ENSMUSG00 | 94223 |
| NA | ENSMUSG00 | 15547 |
| NA | ENSMUSG00 | 56297 |
| NA | ENSMUSG00 | 224273 |
| NA | ENSMUSG00 | 67014 |
| NA | ENSMUSG00 | 12892 |
| NA | ENSMUSG00 | 224250 |
| Significant | ENSMUSG00 | 54613 |
| NA | ENSMUSG00 | 66497 |
| NA | ENSMUSG00 | 67581 |
| NA | ENSMUSG00 | 224023 |
| NA | ENSMUSG00 | 52633 |
| NA | ENSMUSG00 | 28185 |
| NA | ENSMUSG00 | 21787 |
| NA | ENSMUSG00 | 69009 |
| NA | ENSMUSG00 | 66863 |
| NA | ENSMUSG00 | 67474 |
| NA | ENSMUSG00 | 64136 |
| NA | ENSMUSG00 | 13383 |
| NA | ENSMUSG00 | 66053 |
| NA | ENSMUSG00 | 320213 |
| NA | ENSMUSG00 | 106369 |
| NA | ENSMUSG00 | 68092 |
| NA | ENSMUSG00 | 21976 |
| NA | ENSMUSG00 | 224105 |

| | | |
|---|---|---|
| NA | ENSMUSG00 | 68980 |
| NA | ENSMUSG00 | 224014 |
| NA | ENSMUSG00 | 74006 |
| NA | ENSMUSG00 | 51789 |
| NA | ENSMUSG00 | 70120 |
| NA | ENSMUSG00 | 56375 |
| NA | ENSMUSG00 | 22042 |
| NA | ENSMUSG00 | 12549 |
| NA | ENSMUSG00 | 69823 |
| NA | ENSMUSG00 | 70144 |
| NA | ENSMUSG00 | 239833 |
| Significant | ENSMUSG00 | 106326 |
| NA | ENSMUSG00 | 69150 |
| NA | ENSMUSG00 | 22661 |
| NA | ENSMUSG00 | 56637 |
| NA | ENSMUSG00 | 22247 |
| NA | ENSMUSG00 | 14314 |
| NA | ENSMUSG00 | 16419 |
| NA | ENSMUSG00 NA | |
| NA | ENSMUSG00 | 27416 |
| NA | ENSMUSG00 | 67657 |
| NA | ENSMUSG00 | 74197 |
| NA | ENSMUSG00 | 15163 |
| NA | ENSMUSG00 | 239839 |
| NA | ENSMUSG00 | 107589 |
| NA | ENSMUSG00 | 320299 |
| Significant | ENSMUSG00 | 11773 |
| NA | ENSMUSG00 | 72599 |
| NA | ENSMUSG00 | 224132 |
| NA | ENSMUSG00 | 66667 |
| NA | ENSMUSG00 | 74147 |
| NA | ENSMUSG00 | 75826 |
| NA | ENSMUSG00 | 66664 |
| NA | ENSMUSG00 | 20462 |
| NA | ENSMUSG00 | 110197 |
| NA | ENSMUSG00 | 12228 |
| NA | ENSMUSG00 | 67102 |
| NA | ENSMUSG00 | 30940 |
| NA | ENSMUSG00 | 67742 |
| NA | ENSMUSG00 | 106344 |
| NA | ENSMUSG00 | 13682 |
| Significant | ENSMUSG00 | 20440 |
| NA | ENSMUSG00 | 27393 |
| NA | ENSMUSG00 | 11957 |
| NA | ENSMUSG00 | 11820 |
| Significant | ENSMUSG00 | 11504 |

| | | |
|---|---|---|
| NA | ENSMUSG00 | 23872 |
| NA | ENSMUSG00 | 13548 |
| NA | ENSMUSG00 | 13185 |
| NA | ENSMUSG00 | 106347 |
| Significant | ENSMUSG00 | 12524 |
| NA | ENSMUSG00 | 16646 |
| NA | ENSMUSG00 | 80285 |
| NA | ENSMUSG00 | 68146 |
| NA | ENSMUSG00 | 19128 |
| NA | ENSMUSG00 | 56088 |
| NA | ENSMUSG00 | 93871 |
| NA | ENSMUSG00 | 56176 |
| NA | ENSMUSG00 | 110749 |
| NA | ENSMUSG00 | 70028 |
| NA | ENSMUSG00 | 109857 |
| NA | ENSMUSG00 | 224440 |
| NA | ENSMUSG00 | 209195 |
| Significant | ENSMUSG00 | 54720 |
| NA | ENSMUSG00 | 12394 |
| NA | ENSMUSG00 | 28080 |
| Significant | ENSMUSG00 | 16443 |
| NA | ENSMUSG00 | 60364 |
| NA | ENSMUSG00 | 20658 |
| NA | ENSMUSG00 | 14450 |
| NA | ENSMUSG00 | 77975 |
| NA | ENSMUSG00 | 15980 |
| NA | ENSMUSG00 | 15975 |
| NA | ENSMUSG00 | 16155 |
| Significant | ENSMUSG00 | 15976 |
| NA | ENSMUSG00 | 68001 |
| NA | ENSMUSG00 | 104015 |
| NA | ENSMUSG00 | 67367 |
| NA | ENSMUSG00 | 66578 |
| NA | ENSMUSG00 | 20655 |
| NA | ENSMUSG00 | 224432 |
| NA | ENSMUSG00 | 69612 |
| NA | ENSMUSG00 | 11512 |
| Significant | ENSMUSG00 | 13800 |
| NA | ENSMUSG00 | 74775 |
| NA | ENSMUSG00 | 22143 |
| NA | ENSMUSG00 | 54614 |
| NA | ENSMUSG00 | 22379 |
| NA | ENSMUSG00 | 380969 |
| NA | ENSMUSG00 | 110213 |
| NA | ENSMUSG00 | 26934 |
| NA | ENSMUSG00 | 11419 |

| | | |
|---|---|---|
| NA | ENSMUSG00 | 83797 |
| NA | ENSMUSG00 | 14555 |
| NA | ENSMUSG00 | 66379 |
| NA | ENSMUSG00 | 71949 |
| NA | ENSMUSG00 | 65970 |
| NA | ENSMUSG00 | 207214 |
| NA | ENSMUSG00 | 239667 |
| NA | ENSMUSG00 | 11908 |
| Significant | ENSMUSG00 | 18174 |
| Significant | ENSMUSG00 | 109901 |
| NA | ENSMUSG00 | 59033 |
| NA | ENSMUSG00 | 15370 |
| NA | ENSMUSG00 | 93707 |
| NA | ENSMUSG00 | 16668 |
| NA | ENSMUSG00 | 246277 |
| NA | ENSMUSG00 | 223920 |
| NA | ENSMUSG00 | 66151 |
| NA | ENSMUSG00 | 26404 |
| NA | ENSMUSG00 | 21357 |
| NA | ENSMUSG00 | 67488 |
| NA | ENSMUSG00 | 246102 |
| NA | ENSMUSG00 | 12575 |
| NA | ENSMUSG00 | 52014 |
| NA | ENSMUSG00 | 72140 |
| NA | ENSMUSG00 | 70380 |
| NA | ENSMUSG00 | 68050 |
| NA | ENSMUSG00 | 57080 |
| NA | ENSMUSG00 | 12457 |
| NA | ENSMUSG00 | 17250 |
| NA | ENSMUSG00 | 68202 |
| NA | ENSMUSG00 | 14199 |
| NA | ENSMUSG00 | 76467 |
| NA | ENSMUSG00 | 19718 |
| NA | ENSMUSG00 | 68184 |
| NA | ENSMUSG00 | 27374 |
| NA | ENSMUSG00 | 68188 |
| NA | ENSMUSG00 | 27426 |
| NA | ENSMUSG00 | 71446 |
| NA | ENSMUSG00 | 117198 |
| NA | ENSMUSG00 NA | |
| NA | ENSMUSG00 | 105727 |
| NA | ENSMUSG00 | 56310 |
| NA | ENSMUSG00 | 12215 |
| NA | ENSMUSG00 | 67776 |
| NA | ENSMUSG00 | 16169 |
| NA | ENSMUSG00 | 12258 |

| | | |
|---|---|---|
| NA | ENSMUSG00 | 20300 |
| NA | ENSMUSG00 | 16529 |
| NA | ENSMUSG00 | 235587 |
| NA | ENSMUSG00 | 171429 |
| NA | ENSMUSG00 | 109652 |
| NA | ENSMUSG00 | 76737 |
| NA | ENSMUSG00 | 23999 |
| NA | ENSMUSG00 | 675812 |
| NA | ENSMUSG00 | 140499 |
| NA | ENSMUSG00 | 69104 |
| NA | ENSMUSG00 | 69185 |
| NA | ENSMUSG00 | 17858 |
| NA | ENSMUSG00 | 70235 |
| NA | ENSMUSG00 | 22051 |
| NA | ENSMUSG00 | 56620 |
| NA | ENSMUSG00 | 213990 |
| Significant | ENSMUSG00 | 66058 |
| NA | ENSMUSG00 | 228993 |
| NA | ENSMUSG00 | 320951 |
| NA | ENSMUSG00 | 21991 |
| NA | ENSMUSG00 | 19328 |
| NA | ENSMUSG00 | 56734 |
| Significant | ENSMUSG00 | 59092 |
| NA | ENSMUSG00 | 14793 |
| NA | ENSMUSG00 | 67389 |
| NA | ENSMUSG00 | 17151 |
| NA | ENSMUSG00 | 66627 |
| NA | ENSMUSG00 | 64656 |
| NA | ENSMUSG00 | 69159 |
| NA | ENSMUSG00 | 78887 |
| NA | ENSMUSG00 | 84653 |
| NA | ENSMUSG00 | 72084 |
| NA | ENSMUSG00 | NA |
| NA | ENSMUSG00 | 24001 |
| NA | ENSMUSG00 | 20975 |
| NA | ENSMUSG00 | 100037282 |
| NA | ENSMUSG00 | NA |
| NA | ENSMUSG00 | 50873 |
| NA | ENSMUSG00 | 68262 |
| NA | ENSMUSG00 | 16004 |
| NA | ENSMUSG00 | 110460 |
| NA | ENSMUSG00 | 240028 |
| NA | ENSMUSG00 | 12648 |
| NA | ENSMUSG00 | 55951 |
| NA | ENSMUSG00 | 66931 |
| NA | ENSMUSG00 | 22712 |

| | | |
|---|---|---|
| NA | ENSMUSG00 | 72057 |
| NA | ENSMUSG00 | 22709 |
| NA | ENSMUSG00 | 353502 |
| NA | ENSMUSG00 | 27279 |
| NA | ENSMUSG00 | 54419 |
| NA | ENSMUSG00 | 268930 |
| Significant | ENSMUSG00 | 76498 |
| NA | ENSMUSG00 | 27226 |
| NA | ENSMUSG00 | 94185 |
| NA | ENSMUSG00 | 83815 |
| NA | ENSMUSG00 | 17850 |
| NA | ENSMUSG00 | 72238 |
| NA | ENSMUSG00 | 20230 |
| NA | ENSMUSG00 | 71702 |
| NA | ENSMUSG00 | 71020 |
| NA | ENSMUSG00 | 224805 |
| NA | ENSMUSG00 | 68463 |
| NA | ENSMUSG00 | 72415 |
| NA | ENSMUSG00 | 63959 |
| NA | ENSMUSG00 | 15516 |
| NA | ENSMUSG00 | 18037 |
| NA | ENSMUSG00 | 56055 |
| NA | ENSMUSG00 | 80905 |
| NA | ENSMUSG00 | 224794 |
| NA | ENSMUSG00 | 56050 |
| NA | ENSMUSG00 | 50758 |
| NA | ENSMUSG00 | 75564 |
| NA | ENSMUSG00 | 68565 |
| NA | ENSMUSG00 | 224823 |
| NA | ENSMUSG00 | 71461 |
| NA | ENSMUSG00 | 72029 |
| NA | ENSMUSG00 | 224826 |
| NA | ENSMUSG00 | 63856 |
| NA | ENSMUSG00 | 14913 |
| NA | ENSMUSG00 | 112694759 |
| NA | ENSMUSG00 | 53414 |
| NA | ENSMUSG00 | 21425 |
| NA | ENSMUSG00 | 74123 |
| NA | ENSMUSG00 | 83433 |
| NA | ENSMUSG00 | 18044 |
| NA | ENSMUSG00 | 57261 |
| NA | ENSMUSG00 | 106504 |
| NA | ENSMUSG00 | 68816 |
| NA | ENSMUSG00 | 56462 |
| NA | ENSMUSG00 | 26382 |
| NA | ENSMUSG00 | 18712 |

| | | |
|---|---|---|
| NA | ENSMUSG00 | 68597 |
| NA | ENSMUSG00 | 74157 |
| NA | ENSMUSG00 | 109801 |
| NA | ENSMUSG00 | 11307 |
| NA | ENSMUSG00 | 224674 |
| NA | ENSMUSG00 | 57773 |
| NA | ENSMUSG00 | 78330 |
| NA | ENSMUSG00 | 12411 |
| NA | ENSMUSG00 | 17691 |
| NA | ENSMUSG00 | 268970 |
| Significant | ENSMUSG00 | 13823 |
| NA | ENSMUSG00 | 56399 |
| NA | ENSMUSG00 | 67268 |
| NA | ENSMUSG00 | 17929 |
| NA | ENSMUSG00 | 22404 |
| NA | ENSMUSG00 | 64898 |
| Significant | ENSMUSG00 | 246707 |
| NA | ENSMUSG00 | 74355 |
| NA | ENSMUSG00 | 170716 |
| NA | ENSMUSG00 | 67052 |
| NA | ENSMUSG00 | 77889 |
| Significant | ENSMUSG00 | 22436 |
| NA | ENSMUSG00 | 66310 |
| NA | ENSMUSG00 | 50850 |
| NA | ENSMUSG00 | 210148 |
| NA | ENSMUSG00 | 75292 |
| NA | ENSMUSG00 | 67864 |
| NA | ENSMUSG00 | 12211 |
| NA | ENSMUSG00 | 50766 |
| NA | ENSMUSG00 | 268980 |
| NA | ENSMUSG00 | 74196 |
| NA | ENSMUSG00 | 19106 |
| NA | ENSMUSG00 | 12607 |
| NA | ENSMUSG00 | 73694 |
| NA | ENSMUSG00 | 224938 |
| Significant | ENSMUSG00 | 70536 |
| NA | ENSMUSG00 | 17158 |
| NA | ENSMUSG00 | 30960 |
| NA | ENSMUSG00 | 72692 |
| NA | ENSMUSG00 | 19765 |
| NA | ENSMUSG00 | 225027 |
| NA | ENSMUSG00 | 65960 |
| NA | ENSMUSG00 | 72900 |
| NA | ENSMUSG00 | 68767 |
| NA | ENSMUSG00 | 28006 |
| NA | ENSMUSG00 | 72016 |

| | | |
|---|---|---|
| NA | ENSMUSG00 | 72416 |
| NA | ENSMUSG00 | 11984 |
| NA | ENSMUSG00 | 18607 |
| NA | ENSMUSG00 | 213760 |
| NA | ENSMUSG00 | 27410 |
| NA | ENSMUSG00 | 13177 |
| NA | ENSMUSG00 | 78586 |
| NA | ENSMUSG00 | 66705 |
| NA | ENSMUSG00 | 13560 |
| NA | ENSMUSG00 | 56716 |
| NA | ENSMUSG00 | 104215 |
| NA | ENSMUSG00 | 18701 |
| NA | ENSMUSG00 | 56724 |
| NA | ENSMUSG00 | 193813 |
| NA | ENSMUSG00 | 17685 |
| NA | ENSMUSG00 | 14651 |
| NA | ENSMUSG00 | 79043 |
| NA | ENSMUSG00 | 30957 |
| Significant | ENSMUSG00 | 12266 |
| NA | ENSMUSG00 | 52009 |
| NA | ENSMUSG00 | 407831 |
| NA | ENSMUSG00 | 106633 |
| NA | ENSMUSG00 | 71718 |
| NA | ENSMUSG00 | 72836 |
| Significant | ENSMUSG00 | 56520 |
| NA | ENSMUSG00 | 60455 |
| NA | ENSMUSG00 | 68611 |
| NA | ENSMUSG00 | 12005 |
| Significant | ENSMUSG00 | 106581 |
| NA | ENSMUSG00 | 66978 |
| NA | ENSMUSG00 | 19252 |
| NA | ENSMUSG00 | 224630 |
| NA | ENSMUSG00 | 21652 |
| NA | ENSMUSG00 | 67675 |
| NA | ENSMUSG00 | 66905 |
| NA | ENSMUSG00 | 193796 |
| NA | ENSMUSG00 | 67267 |
| NA | ENSMUSG00 | 64144 |
| NA | ENSMUSG00 | 56409 |
| NA | ENSMUSG00 | 20630 |
| NA | ENSMUSG00 | 68776 |
| NA | ENSMUSG00 | 224650 |
| NA | ENSMUSG00 | 224656 |
| NA | ENSMUSG00 | 14229 |
| NA | ENSMUSG00 | 67993 |
| NA | ENSMUSG00 | 71745 |

| | | |
|---|---|---|
| NA | ENSMUSG00 | 68010 |
| NA | ENSMUSG00 | 67440 |
| NA | ENSMUSG00 | 26410 |
| NA | ENSMUSG00 | 225115 |
| NA | ENSMUSG00 | 21417 |
| NA | ENSMUSG00 | 13831 |
| NA | ENSMUSG00 | 20662 |
| NA | ENSMUSG00 | 225028 |
| NA | ENSMUSG00 | 72167 |
| NA | ENSMUSG00 | 106522 |
| NA | ENSMUSG00 | 20463 |
| NA | ENSMUSG00 | 240174 |
| NA | ENSMUSG00 | 213575 |
| NA | ENSMUSG00 | 69241 |
| NA | ENSMUSG00 | 67453 |
| NA | ENSMUSG00 | 269003 |
| NA | ENSMUSG00 | 66648 |
| NA | ENSMUSG00 | 106957 |
| NA | ENSMUSG00 | 58523 |
| NA | ENSMUSG00 | 68046 |
| NA | ENSMUSG00 NA | |
| NA | ENSMUSG00 | 69256 |
| NA | ENSMUSG00 | 212307 |
| NA | ENSMUSG00 | 225131 |
| NA | ENSMUSG00 | 67974 |
| NA | ENSMUSG00 | 225160 |
| NA | ENSMUSG00 | 19877 |
| NA | ENSMUSG00 | 77805 |
| NA | ENSMUSG00 | 225164 |
| NA | ENSMUSG00 NA | |
| NA | ENSMUSG00 | 208292 |
| NA | ENSMUSG00 | 224697 |
| NA | ENSMUSG00 | 17916 |
| NA | ENSMUSG00 | 16580 |
| NA | ENSMUSG00 | 12558 |
| NA | ENSMUSG00 | 21356 |
| NA | ENSMUSG00 | 14976 |
| Significant | ENSMUSG00 | 57315 |
| NA | ENSMUSG00 | 56515 |
| NA | ENSMUSG00 | 224705 |
| NA | ENSMUSG00 | 19763 |
| NA | ENSMUSG00 | 14977 |
| NA | ENSMUSG00 | 15001 |
| NA | ENSMUSG00 | 14312 |
| NA | ENSMUSG00 | 16913 |
| NA | ENSMUSG00 | 21355 |

| | | |
|---|---|---|
| Significant | ENSMUSG00 | 547431 |
| NA | ENSMUSG00 | 67199 |
| NA | ENSMUSG00 | 72512 |
| NA | ENSMUSG00 | 76594 |
| NA | ENSMUSG00 | 71242 |
| NA | ENSMUSG00 | 81500 |
| NA | ENSMUSG00 | 15526 |
| NA | ENSMUSG00 | 225363 |
| NA | ENSMUSG00 | 27632 |
| NA | ENSMUSG00 | 52563 |
| NA | ENSMUSG00 | 12263 |
| NA | ENSMUSG00 | 13824 |
| Significant | ENSMUSG00 | 170459 |
| Significant | ENSMUSG00 | 30948 |
| NA | ENSMUSG00 | 13872 |
| NA | ENSMUSG00 | 26405 |
| NA | ENSMUSG00 | 73473 |
| NA | ENSMUSG00 | 13001 |
| NA | ENSMUSG00 | 224727 |
| NA | ENSMUSG00 | 53761 |
| NA | ENSMUSG00 | 11629 |
| NA | ENSMUSG00 | 16994 |
| NA | ENSMUSG00 | 74320 |
| NA | ENSMUSG00 | 21926 |
| NA | ENSMUSG00 | 66237 |
| NA | ENSMUSG00 | 66878 |
| NA | ENSMUSG00 | 76482 |
| NA | ENSMUSG00 | 18145 |
| NA | ENSMUSG00 | 94064 |
| NA | ENSMUSG00 | 225207 |
| NA | ENSMUSG00 | 69192 |
| NA | ENSMUSG00 | 16210 |
| NA | ENSMUSG00 | 72747 |
| NA | ENSMUSG00 | 65113 |
| NA | ENSMUSG00 | 73242 |
| NA | ENSMUSG00 | 24066 |
| NA | ENSMUSG00 | 14670 |
| NA | ENSMUSG00 | 14815 |
| NA | ENSMUSG00 | 66973 |
| NA | ENSMUSG00 | 53601 |
| NA | ENSMUSG00 | 66839 |
| NA | ENSMUSG00 | 67676 |
| NA | ENSMUSG00 | 106952 |
| NA | ENSMUSG00 | 15183 |
| NA | ENSMUSG00 | 13367 |
| NA | ENSMUSG00 | 22670 |

| | | |
|---|---|---|
| NA | ENSMUSG00 | 70640 |
| NA | ENSMUSG00 | 24010 |
| NA | ENSMUSG00 | 225467 |
| NA | ENSMUSG00 | 11777 |
| NA | ENSMUSG00 | 108857 |
| NA | ENSMUSG00 | 15200 |
| NA | ENSMUSG00 | 67180 |
| NA | ENSMUSG00 | 225432 |
| NA | ENSMUSG00 | 107045 |
| NA | ENSMUSG00 | 56070 |
| NA | ENSMUSG00 | 22240 |
| NA | ENSMUSG00 | 68857 |
| NA | ENSMUSG00 | 15488 |
| NA | ENSMUSG00 | 17191 |
| NA | ENSMUSG00 | 17128 |
| NA | ENSMUSG00 | 66286 |
| NA | ENSMUSG00 | 58801 |
| NA | ENSMUSG00 | 14680 |
| NA | ENSMUSG00 | 114663 |
| NA | ENSMUSG00 | 69597 |
| NA | ENSMUSG00 | 67222 |
| NA | ENSMUSG00 | 225655 |
| NA | ENSMUSG00 | 68166 |
| NA | ENSMUSG00 | 67847 |
| Significant | ENSMUSG00 | 69226 |
| NA | ENSMUSG00 | 107047 |
| NA | ENSMUSG00 | 19038 |
| NA | ENSMUSG00 | 19255 |
| NA | ENSMUSG00 | 70799 |
| NA | ENSMUSG00 | 52662 |
| Significant | ENSMUSG00 | 107029 |
| NA | ENSMUSG00 | 74322 |
| NA | ENSMUSG00 | 17190 |
| NA | ENSMUSG00 | 17126 |
| NA | ENSMUSG00 | 20689 |
| NA | ENSMUSG00 | 50771 |
| NA | ENSMUSG00 | 68731 |
| NA | ENSMUSG00 NA | |
| NA | ENSMUSG00 | 93687 |
| Significant | ENSMUSG00 | 240334 |
| NA | ENSMUSG00 | 17714 |
| NA | ENSMUSG00 | 108123 |
| NA | ENSMUSG00 | 53382 |
| NA | ENSMUSG00 | 70223 |
| NA | ENSMUSG00 | 14151 |
| NA | ENSMUSG00 | 83814 |

| | | |
|---|---|---|
| NA | ENSMUSG00 | 16906 |
| NA | ENSMUSG00 | 77422 |
| NA | ENSMUSG00 | 73137 |
| NA | ENSMUSG00 | 20496 |
| NA | ENSMUSG00 | 14119 |
| NA | ENSMUSG00 | 66307 |
| NA | ENSMUSG00 | 67665 |
| NA | ENSMUSG00 | 66810 |
| NA | ENSMUSG00 | 20044 |
| NA | ENSMUSG00 | 16149 |
| NA | ENSMUSG00 | 21453 |
| Significant | ENSMUSG00 | 67988 |
| NA | ENSMUSG00 | 12322 |
| NA | ENSMUSG00 | 18596 |
| NA | ENSMUSG00 | 12978 |
| NA | ENSMUSG00 | 106894 |
| NA | ENSMUSG00 | 14682 |
| NA | ENSMUSG00 | 107272 |
| NA | ENSMUSG00 | 21888 |
| Significant | ENSMUSG00 | 66054 |
| NA | ENSMUSG00 | 67105 |
| Significant | ENSMUSG00 | 109672 |
| NA | ENSMUSG00 | 66514 |
| Significant | ENSMUSG00 | 16952 |
| NA | ENSMUSG00 | 16319 |
| Significant | ENSMUSG00 | 14319 |
| NA | ENSMUSG00 | 74760 |
| NA | ENSMUSG00 | 60527 |
| NA | ENSMUSG00 | 56473 |
| NA | ENSMUSG00 | 72982 |
| NA | ENSMUSG00 | 68642 |
| NA | ENSMUSG00 | 66072 |
| NA | ENSMUSG00 | 12507 |
| Significant | ENSMUSG00 | 109225 |
| NA | ENSMUSG00 | 64380 |
| Significant | ENSMUSG00 | 69774 |
| Significant | ENSMUSG00 | 68774 |
| Significant | ENSMUSG00 | 170813 |
| NA | ENSMUSG00 | 94063 |
| NA | ENSMUSG00 | 76303 |
| NA | ENSMUSG00 | 107373 |
| NA | ENSMUSG00 | 109910 |
| NA | ENSMUSG00 | 107321 |
| NA | ENSMUSG00 | 54391 |
| NA | ENSMUSG00 | 20409 |
| NA | ENSMUSG00 | 67383 |

| | | |
|---|---|---|
| NA | ENSMUSG00 | 108673 |
| NA | ENSMUSG00 | 28000 |
| NA | ENSMUSG00 | 98170 |
| NA | ENSMUSG00 | 65221 |
| NA | ENSMUSG00 | 13194 |
| NA | ENSMUSG00 | 14156 |
| NA | ENSMUSG00 | 22682 |
| NA | ENSMUSG00 | 83921 |
| NA | ENSMUSG00 | 20168 |
| NA | ENSMUSG00 | 109168 |
| NA | ENSMUSG00 | 70999 |
| NA | ENSMUSG00 | 381236 |
| NA | ENSMUSG00 | 107260 |
| Significant | ENSMUSG00 | 13660 |
| NA | ENSMUSG00 | 329015 |
| NA | ENSMUSG00 | 76630 |
| NA | ENSMUSG00 | 225849 |
| NA | ENSMUSG00 | 14102 |
| NA | ENSMUSG00 | 67072 |
| NA | ENSMUSG00 | 16889 |
| NA | ENSMUSG00 | 56248 |
| NA | ENSMUSG00 | 59028 |
| NA | ENSMUSG00 | 69024 |
| NA | ENSMUSG00 | 16452 |
| NA | ENSMUSG00 | 66406 |
| NA | ENSMUSG00 | 67849 |
| NA | ENSMUSG00 | 81909 |
| NA | ENSMUSG00 | 85030 |
| NA | ENSMUSG00 | 240641 |
| NA | ENSMUSG00 | 68505 |
| NA | ENSMUSG00 | 73166 |
| NA | ENSMUSG00 | 54364 |
| NA | ENSMUSG00 | 76073 |
| NA | ENSMUSG00 | 74126 |
| NA | ENSMUSG00 | 74493 |
| NA | ENSMUSG00 | 21873 |
| NA | ENSMUSG00 | 67457 |
| NA | ENSMUSG00 | 109113 |
| NA | ENSMUSG00 | 107375 |
| NA | ENSMUSG00 | 19367 |
| NA | ENSMUSG00 | 19708 |
| NA | ENSMUSG00 | 353242 |
| NA | ENSMUSG00 | 58988 |
| NA | ENSMUSG00 | 20589 |
| NA | ENSMUSG00 | 18969 |
| NA | ENSMUSG00 | 23789 |

| | | |
|---|---|---|
| NA | ENSMUSG00 | 69860 |
| NA | ENSMUSG00 | 12660 |
| NA | ENSMUSG00 | 23825 |
| NA | ENSMUSG00 | 11632 |
| NA | ENSMUSG00 | 18739 |
| NA | ENSMUSG00 | 319322 |
| NA | ENSMUSG00 | 69745 |
| NA | ENSMUSG00 | 107975 |
| NA | ENSMUSG00 | 52004 |
| NA | ENSMUSG00 | 110355 |
| NA | ENSMUSG00 | 71670 |
| NA | ENSMUSG00 | 76308 |
| NA | ENSMUSG00 | 68090 |
| NA | ENSMUSG00 | 226043 |
| NA | ENSMUSG00 | 67689 |
| NA | ENSMUSG00 | 54447 |
| NA | ENSMUSG00 | 19671 |
| NA | ENSMUSG00 | 13340 |
| NA | ENSMUSG00 | 18563 |
| NA | ENSMUSG00 | 17330 |
| NA | ENSMUSG00 | 319924 |
| NA | ENSMUSG00 | 23972 |
| NA | ENSMUSG00 | 12894 |
| NA | ENSMUSG00 | 66419 |
| NA | ENSMUSG00 | 17771 |
| NA | ENSMUSG00 | 71711 |
| NA | ENSMUSG00 | 52036 |
| Significant | ENSMUSG00 | 58859 |
| NA | ENSMUSG00 | 58249 |
| NA | ENSMUSG00 | 14283 |
| NA | ENSMUSG00 | 16973 |
| NA | ENSMUSG00 | 66556 |
| NA | ENSMUSG00 | 67155 |
| NA | ENSMUSG00 | 68209 |
| NA | ENSMUSG00 | 81601 |
| NA | ENSMUSG00 | 19697 |
| NA | ENSMUSG00 | 114601 |
| NA | ENSMUSG00 | 17826 |
| NA | ENSMUSG00 | 78891 |
| NA | ENSMUSG00 | 12333 |
| NA | ENSMUSG00 | 226026 |
| NA | ENSMUSG00 | 56327 |
| NA | ENSMUSG00 | 17283 |
| NA | ENSMUSG00 | 26412 |
| NA | ENSMUSG00 | 22668 |
| NA | ENSMUSG00 | NA |

| | | |
|---|---|---|
| NA | ENSMUSG00 | 54683 |
| NA | ENSMUSG00 | 26379 |
| NA | ENSMUSG00 | 107173 |
| NA | ENSMUSG00 | 12015 |
| NA | ENSMUSG00 | 18797 |
| NA | ENSMUSG00 | 22340 |
| NA | ENSMUSG00 | 57431 |
| NA | ENSMUSG00 | 108101 |
| NA | ENSMUSG00 | 20867 |
| NA | ENSMUSG00 | 104383 |
| NA | ENSMUSG00 | 13728 |
| NA | ENSMUSG00 | 68229 |
| NA | ENSMUSG00 | 13006 |
| NA | ENSMUSG00 | 18569 |
| NA | ENSMUSG00 | 56392 |
| NA | ENSMUSG00 | 14732 |
| NA | ENSMUSG00 | 433256 |
| NA | ENSMUSG00 | 66980 |
| NA | ENSMUSG00 | 53611 |
| NA | ENSMUSG00 | 21416 |
| NA | ENSMUSG00 | 15242 |
| NA | ENSMUSG00 | 74107 |
| NA | ENSMUSG00 | 19662 |
| NA | ENSMUSG00 | 13669 |
| NA | ENSMUSG00 | 67894 |
| NA | ENSMUSG00 | 11757 |
| NA | ENSMUSG00 | 57753 |
| NA | ENSMUSG00 | 15201 |
| NA | ENSMUSG00 | 20411 |
| NA | ENSMUSG00 | 56454 |
| NA | ENSMUSG00 | 67590 |
| NA | ENSMUSG00 | 240665 |
| NA | ENSMUSG00 | 107358 |
| NA | ENSMUSG00 | 83490 |
| NA | ENSMUSG00 | 212391 |
| NA | ENSMUSG00 | 76479 |
| NA | ENSMUSG00 | 17859 |
| NA | ENSMUSG00 | 27360 |
| NA | ENSMUSG00 | 170750 |
| NA | ENSMUSG00 | 93679 |
| NA | ENSMUSG00 | 56350 |
| NA | ENSMUSG00 | 94279 |
| NA | ENSMUSG00 | 17161 |
| NA | ENSMUSG00 | 72018 |
| NA | ENSMUSG00 | 76952 |
| Significant | ENSMUSG00 | 20288 |

| | | |
|---|---|---|
| NA | ENSMUSG00 | 18572 |
| NA | ENSMUSG00 | 226182 |
| NA | ENSMUSG00 | 71041 |
| NA | ENSMUSG00 | 71398 |
| NA | ENSMUSG00 | 14933 |
| NA | ENSMUSG00 | 20874 |
| NA | ENSMUSG00 | 67788 |
| NA | ENSMUSG00 | 14873 |
| NA | ENSMUSG00 | 68214 |
| NA | ENSMUSG00 | 12369 |
| NA | ENSMUSG00 | 55947 |
| NA | ENSMUSG00 | 66866 |
| NA | ENSMUSG00 | 226251 |
| NA | ENSMUSG00 | 72133 |
| NA | ENSMUSG00 | 73442 |
| NA | ENSMUSG00 | 214084 |
| NA | ENSMUSG00 | 67290 |
| NA | ENSMUSG00 | 83962 |
| NA | ENSMUSG00 | 29877 |
| NA | ENSMUSG00 | 18453 |
| NA | ENSMUSG00 | 192662 |
| NA | ENSMUSG00 | 101861 |
| NA | ENSMUSG00 | 21681 |
| NA | ENSMUSG00 | 66156 |
| NA | ENSMUSG00 | 68671 |
| NA | ENSMUSG00 | 209011 |
| NA | ENSMUSG00 | 54473 |
| NA | ENSMUSG00 | 209027 |
| NA | ENSMUSG00 | 68515 |
| NA | ENSMUSG00 | 68938 |
| NA | ENSMUSG00 | 20892 |
| NA | ENSMUSG00 | 217366 |
| NA | ENSMUSG00 | 101513 |
| NA | ENSMUSG00 | 12409 |
| Significant | ENSMUSG00 | 94275 |
| NA | ENSMUSG00 | 14104 |
| Significant | ENSMUSG00 | 71085 |
| NA | ENSMUSG00 | 68730 |
| NA | ENSMUSG00 | 209318 |
| NA | ENSMUSG00 | 74168 |
| NA | ENSMUSG00 | 19719 |
| NA | ENSMUSG00 | 72199 |
| NA | ENSMUSG00 | 80879 |
| NA | ENSMUSG00 | 104318 |
| NA | ENSMUSG00 | 66179 |
| NA | ENSMUSG00 | 226122 |

| | | |
|---|---|---|
| NA | ENSMUSG00 | 66840 |
| NA | ENSMUSG00 | 84095 |
| NA | ENSMUSG00 | 52013 |
| NA | ENSMUSG00 | 192236 |
| NA | ENSMUSG00 | 14718 |
| NA | ENSMUSG00 | 93685 |
| NA | ENSMUSG00 | 66388 |
| NA | ENSMUSG00 | 71972 |
| NA | ENSMUSG00 | 93721 |
| NA | ENSMUSG00 | 226144 |
| NA | ENSMUSG00 | 12675 |
| NA | ENSMUSG00 | 72502 |
| NA | ENSMUSG00 | 20250 |
| NA | ENSMUSG00 | 67264 |
| NA | ENSMUSG00 | 94067 |
| NA | ENSMUSG00 | 226153 |
| NA | ENSMUSG00 | 94280 |
| NA | ENSMUSG00 | 12234 |
| NA | ENSMUSG00 | 56626 |
| NA | ENSMUSG00 | 76055 |
| NA | ENSMUSG00 | 16825 |
| NA | ENSMUSG00 | 107338 |
| NA | ENSMUSG00 | 18034 |
| NA | ENSMUSG00 | 68431 |
| NA | ENSMUSG00 | 75146 |
| NA | ENSMUSG00 | 54130 |
| NA | ENSMUSG00 | 24069 |
| NA | ENSMUSG00 | 15211 |
| NA | ENSMUSG00 | 23806 |
| NA | ENSMUSG00 | 102774 |
| NA | ENSMUSG00 | 72141 |
| NA | ENSMUSG00 | 67287 |
| NA | ENSMUSG00 | 29806 |
| Significant | ENSMUSG00 | 83493 |
| NA | ENSMUSG00 | 17281 |
| NA | ENSMUSG00 | 93730 |
| NA | ENSMUSG00 | 21372 |
| NA | ENSMUSG00 | 66611 |
| NA | ENSMUSG00 | 15108 |
| NA | ENSMUSG00 | 59026 |
| NA | ENSMUSG00 | 207375 |
| NA | ENSMUSG00 | 69499 |
| NA | ENSMUSG00 | 14163 |
| NA | ENSMUSG00 | 237107 |
| Significant | ENSMUSG00 | 80884 |
| NA | ENSMUSG00 | 77622 |

| | | |
|---|---|---|
| NA | ENSMUSG00 | 18639 |
| NA | ENSMUSG00 | 66082 |
| NA | ENSMUSG00 | 286940 |
| NA | ENSMUSG00 | 13421 |
| NA | ENSMUSG00 | 218832 |
| NA | ENSMUSG00 | 20229 |
| NA | ENSMUSG00 | 56360 |
| NA | ENSMUSG00 | 53381 |
| NA | ENSMUSG00 | 20088 |
| NA | ENSMUSG00 | 19271 |
| NA | ENSMUSG00 | 53325 |
| NA | ENSMUSG00 | 20688 |
| NA | ENSMUSG00 | 11982 |
| NA | ENSMUSG00 | 22215 |
| NA | ENSMUSG00 | 20591 |
| NA | ENSMUSG00 | 66711 |
| NA | ENSMUSG00 | 56715 |
| NA | ENSMUSG00 | 19682 |
| NA | ENSMUSG00 | 12512 |
| NA | ENSMUSG00 | 14561 |
| NA | ENSMUSG00 | 66844 |
| NA | ENSMUSG00 | 74330 |
| NA | ENSMUSG00 | 58223 |
| NA | ENSMUSG00 | 13139 |
| NA | ENSMUSG00 | 12566 |
| NA | ENSMUSG00 | 20431 |
| NA | ENSMUSG00 | 27370 |
| Significant | ENSMUSG00 | 18813 |
| Significant | ENSMUSG00 | 23943 |
| NA | ENSMUSG00 | 68094 |
| NA | ENSMUSG00 | 208092 |
| NA | ENSMUSG00 | 108100 |
| NA | ENSMUSG00 | 67588 |
| NA | ENSMUSG00 | 69917 |
| NA | ENSMUSG00 | 11302 |
| NA | ENSMUSG00 | 78777 |
| NA | ENSMUSG00 | 56530 |
| NA | ENSMUSG00 | 71885 |
| NA | ENSMUSG00 | 11947 |
| NA | ENSMUSG00 | 19075 |
| NA | ENSMUSG00 | 17937 |
| NA | ENSMUSG00 | 108037 |
| NA | ENSMUSG00 | 71750 |
| NA | ENSMUSG00 | 13198 |
| NA | ENSMUSG00 | 110962 |
| NA | ENSMUSG00 | 69654 |

| | | |
|---|---|---|
| NA | ENSMUSG00 | 72354 |
| NA | ENSMUSG00 | 117150 |
| NA | ENSMUSG00 | 71206 |
| NA | ENSMUSG00 | 76987 |
| NA | ENSMUSG00 | 17344 |
| NA | ENSMUSG00 | 11946 |
| NA | ENSMUSG00 | 19201 |
| NA | ENSMUSG00 | 68876 |
| NA | ENSMUSG00 | 170822 |
| NA | ENSMUSG00 | 12729 |
| NA | ENSMUSG00 | 218693 |
| NA | ENSMUSG00 | 18115 |
| Significant | ENSMUSG00 | 212503 |
| NA | ENSMUSG00 | 93747 |
| NA | ENSMUSG00 | 69064 |
| NA | ENSMUSG00 | 69752 |
| Significant | ENSMUSG00 | 11501 |
| NA | ENSMUSG00 | 74237 |
| NA | ENSMUSG00 | 212111 |
| NA | ENSMUSG00 | 54399 |
| NA | ENSMUSG00 | 101489 |
| NA | ENSMUSG00 | 64384 |
| NA | ENSMUSG00 | 23997 |
| NA | ENSMUSG00 | 73835 |
| Significant | ENSMUSG00 | 68713 |
| Significant | ENSMUSG00 | 66141 |
| NA | ENSMUSG00 | 24058 |
| NA | ENSMUSG00 | 27388 |
| NA | ENSMUSG00 | 54123 |
| NA | ENSMUSG00 | 15461 |
| NA | ENSMUSG00 | 21351 |
| NA | ENSMUSG00 | 71448 |
| NA | ENSMUSG00 | 57913 |
| NA | ENSMUSG00 | 67186 |
| NA | ENSMUSG00 | 66853 |
| NA | ENSMUSG00 | 12476 |
| NA | ENSMUSG00 | 64540 |
| NA | ENSMUSG00 | 68038 |
| NA | ENSMUSG00 | 73067 |
| NA | ENSMUSG00 | 68117 |
| NA | ENSMUSG00 | 245595 |
| NA | ENSMUSG00 | 12662 |
| NA | ENSMUSG00 | 12909 |
| NA | ENSMUSG00 | 109900 |
| NA | ENSMUSG00 | 110006 |
| NA | ENSMUSG00 | 67902 |

| | | |
|---|---|---|
| NA | ENSMUSG00 | 68059 |
| NA | ENSMUSG00 | 69634 |
| NA | ENSMUSG00 | 223254 |
| NA | ENSMUSG00 | 105445 |
| NA | ENSMUSG00 | 217351 |
| NA | ENSMUSG00 | 217353 |
| NA | ENSMUSG00 | 21877 |
| NA | ENSMUSG00 | 76025 |
| NA | ENSMUSG00 | 52897 |
| NA | ENSMUSG00 | 12416 |
| NA | ENSMUSG00 | 30951 |
| NA | ENSMUSG00 | 14387 |
| NA | ENSMUSG00 | 192170 |
| NA | ENSMUSG00 | 74370 |
| NA | ENSMUSG00 | 66282 |
| NA | ENSMUSG00 | 93837 |
| NA | ENSMUSG00 | 237010 |
| NA | ENSMUSG00 | 80902 |
| NA | ENSMUSG00 | 54160 |
| NA | ENSMUSG00 | 27418 |
| NA | ENSMUSG00 | 12013 |
| NA | ENSMUSG00 | 12469 |
| NA | ENSMUSG00 | 74112 |
| NA | ENSMUSG00 | 70998 |
| NA | ENSMUSG00 | 15452 |
| NA | ENSMUSG00 | 66658 |
| NA | ENSMUSG00 | NA |
| NA | ENSMUSG00 | 66940 |
| NA | ENSMUSG00 | 270198 |
| NA | ENSMUSG00 | 22273 |
| NA | ENSMUSG00 | 236915 |
| NA | ENSMUSG00 | 67071 |
| NA | ENSMUSG00 | 192216 |
| NA | ENSMUSG00 | 11689 |
| NA | ENSMUSG00 | 71779 |
| NA | ENSMUSG00 | 71968 |
| NA | ENSMUSG00 | 68039 |
| NA | ENSMUSG00 | 56529 |
| NA | ENSMUSG00 | 14755 |
| NA | ENSMUSG00 | 224624 |
| NA | ENSMUSG00 | 68347 |
| NA | ENSMUSG00 | 68241 |
| NA | ENSMUSG00 | 214952 |
| NA | ENSMUSG00 | 72106 |
| NA | ENSMUSG00 | 268933 |
| NA | ENSMUSG00 | 110639 |

| | | |
|---|---|---|
| NA | ENSMUSG00 | 20970 |
| NA | ENSMUSG00 | 97212 |
| NA | ENSMUSG00 | 16193 |
| NA | ENSMUSG00 | 22171 |
| NA | ENSMUSG00 | 18415 |
| NA | ENSMUSG00 | 20873 |
| NA | ENSMUSG00 | 72175 |
| NA | ENSMUSG00 | 214048 |
| NA | ENSMUSG00 | 269424 |
| NA | ENSMUSG00 | 73852 |
| NA | ENSMUSG00 | 17087 |
| Significant | ENSMUSG00 | 209378 |
| NA | ENSMUSG00 | 11949 |
| NA | ENSMUSG00 | 209361 |
| NA | ENSMUSG00 | 66446 |
| NA | ENSMUSG00 | 69035 |
| NA | ENSMUSG00 | 108116 |
| NA | ENSMUSG00 | 72157 |
| NA | ENSMUSG00 | 27376 |
| NA | ENSMUSG00 | 15239 |
| NA | ENSMUSG00 | 67115 |
| NA | ENSMUSG00 | 192678 |
| NA | ENSMUSG00 | 12768 |
| NA | ENSMUSG00 | 16412 |
| NA | ENSMUSG00 | 18186 |
| NA | ENSMUSG00 | 93742 |
| NA | ENSMUSG00 | 209692 |
| NA | ENSMUSG00 | 57743 |
| NA | ENSMUSG00 | 53893 |
| NA | ENSMUSG00 | 101095 |
| NA | ENSMUSG00 | 12304 |
| NA | ENSMUSG00 | 66383 |
| NA | ENSMUSG00 | 80752 |
| NA | ENSMUSG00 | 19085 |
| NA | ENSMUSG00 | 18590 |
| NA | ENSMUSG00 | 433956 |
| NA | ENSMUSG00 | 67604 |
| NA | ENSMUSG00 | 11798 |
| NA | ENSMUSG00 | 20843 |
| NA | ENSMUSG00 | 67044 |
| NA | ENSMUSG00 | 28126 |
| NA | ENSMUSG00 | 56795 |
| NA | ENSMUSG00 | 97820 |
| NA | ENSMUSG00 | 73666 |
| NA | ENSMUSG00 | 76577 |
| NA | ENSMUSG00 | 74257 |

| | | |
|---|---|---|
| Significant | ENSMUSG00 | 212032 |
| NA | ENSMUSG00 | 20184 |
| NA | ENSMUSG00 | 17131 |
| NA | ENSMUSG00 | 12362 |
| NA | ENSMUSG00 | 69149 |
| NA | ENSMUSG00 | 67618 |
| NA | ENSMUSG00 | 244672 |
| NA | ENSMUSG00 | 67667 |
| NA | ENSMUSG00 | 20671 |
| NA | ENSMUSG00 | 18777 |
| NA | ENSMUSG00 | 12421 |
| NA | ENSMUSG00 | 76187 |
| NA | ENSMUSG00 | 170755 |
| NA | ENSMUSG00 | 26754 |
| NA | ENSMUSG00 | 29819 |
| NA | ENSMUSG00 | 98711 |
| NA | ENSMUSG00 | 21749 |
| NA | ENSMUSG00 | 74229 |
| NA | ENSMUSG00 | 75712 |
| NA | ENSMUSG00 | 72265 |
| NA | ENSMUSG00 | 212442 |
| NA | ENSMUSG00 | 66799 |
| NA | ENSMUSG00 | 70397 |
| NA | ENSMUSG00 | 18711 |
| Significant | ENSMUSG00 | 15926 |
| NA | ENSMUSG00 | 67099 |
| NA | ENSMUSG00 | 12912 |
| NA | ENSMUSG00 | 93691 |
| NA | ENSMUSG00 | 75619 |
| NA | ENSMUSG00 | 55949 |
| NA | ENSMUSG00 | 227197 |
| Significant | ENSMUSG00 | 18187 |
| NA | ENSMUSG00 | 68115 |
| NA | ENSMUSG00 | 74013 |
| NA | ENSMUSG00 | 19070 |
| NA | ENSMUSG00 | 15510 |
| NA | ENSMUSG00 | 67876 |
| NA | ENSMUSG00 | 81898 |
| NA | ENSMUSG00 | 227059 |
| Significant | ENSMUSG00 | 53945 |
| NA | ENSMUSG00 | 73674 |
| NA | ENSMUSG00 | 22779 |
| NA | ENSMUSG00 | 14768 |
| NA | ENSMUSG00 | 11363 |
| NA | ENSMUSG00 | 68691 |
| NA | ENSMUSG00 | 66646 |

| | | |
|---|---|---|
| NA | ENSMUSG00 | 54167 |
| NA | ENSMUSG00 | 12487 |
| NA | ENSMUSG00 | 77300 |
| NA | ENSMUSG00 | 241066 |
| NA | ENSMUSG00 | 70375 |
| NA | ENSMUSG00 | 57750 |
| NA | ENSMUSG00 | 55989 |
| NA | ENSMUSG00 | 22218 |
| NA | ENSMUSG00 | 74018 |
| NA | ENSMUSG00 | 227154 |
| NA | ENSMUSG00 | 70827 |
| NA | ENSMUSG00 | 12370 |
| NA | ENSMUSG00 | 12633 |
| NA | ENSMUSG00 | 66495 |
| NA | ENSMUSG00 | 12747 |
| NA | ENSMUSG00 | 70225 |
| NA | ENSMUSG00 | 65102 |
| NA | ENSMUSG00 | 18393 |
| NA | ENSMUSG00 | 68549 |
| NA | ENSMUSG00 | 12832 |
| NA | ENSMUSG00 | 12825 |
| NA | ENSMUSG00 | 72050 |
| NA | ENSMUSG00 | 22592 |
| NA | ENSMUSG00 | 75623 |
| NA | ENSMUSG00 | 19243 |
| Significant | ENSMUSG00 | 16174 |
| NA | ENSMUSG00 | 17082 |
| Significant | ENSMUSG00 | 16178 |
| NA | ENSMUSG00 | 26921 |
| NA | ENSMUSG00 | 68833 |
| NA | ENSMUSG00 | 98388 |
| NA | ENSMUSG00 | 56210 |
| NA | ENSMUSG00 | 226982 |
| NA | ENSMUSG00 | 107734 |
| NA | ENSMUSG00 | 69028 |
| NA | ENSMUSG00 | 98267 |
| NA | ENSMUSG00 | 70396 |
| NA | ENSMUSG00 | 72085 |
| NA | ENSMUSG00 | 227102 |
| NA | ENSMUSG00 | 227099 |
| NA | ENSMUSG00 | 16329 |
| NA | ENSMUSG00 | 14660 |
| NA | ENSMUSG00 | 20846 |
| Significant | ENSMUSG00 | 109019 |
| NA | ENSMUSG00 | 269181 |
| NA | ENSMUSG00 | 67387 |

| | | |
|---|---|---|
| NA | ENSMUSG00 | 76178 |
| NA | ENSMUSG00 | 269180 |
| NA | ENSMUSG00 | 56030 |
| NA | ENSMUSG00 | 20353 |
| Significant | ENSMUSG00 | 226971 |
| NA | ENSMUSG00 | 19253 |
| NA | ENSMUSG00 | 27993 |
| Significant | ENSMUSG00 | 13518 |
| NA | ENSMUSG00 | 19076 |
| NA | ENSMUSG00 | 77264 |
| NA | ENSMUSG00 | 76867 |
| NA | ENSMUSG00 | 12839 |
| NA | ENSMUSG00 | 75734 |
| NA | ENSMUSG00 | 68187 |
| NA | ENSMUSG00 | 68002 |
| NA | ENSMUSG00 | 98366 |
| Significant | ENSMUSG00 | 70155 |
| NA | ENSMUSG00 | 15463 |
| NA | ENSMUSG00 | 75570 |
| NA | ENSMUSG00 | 104086 |
| NA | ENSMUSG00 | 57751 |
| NA | ENSMUSG00 | 66821 |
| NA | ENSMUSG00 | 58184 |
| NA | ENSMUSG00 | 227292 |
| NA | ENSMUSG00 | 18173 |
| NA | ENSMUSG00 | 56695 |
| Significant | ENSMUSG00 | 12765 |
| NA | ENSMUSG00 | 68606 |
| NA | ENSMUSG00 | 16011 |
| NA | ENSMUSG00 | 22596 |
| NA | ENSMUSG00 | 111175 |
| Significant | ENSMUSG00 | 108147 |
| NA | ENSMUSG00 | 14268 |
| NA | ENSMUSG00 | 12021 |
| NA | ENSMUSG00 | 68818 |
| NA | ENSMUSG00 | 74104 |
| NA | ENSMUSG00 | 52231 |
| NA | ENSMUSG00 | 74577 |
| NA | ENSMUSG00 | 20872 |
| Significant | ENSMUSG00 | 22145 |
| NA | ENSMUSG00 | 56812 |
| Significant | ENSMUSG00 | 11790 |
| NA | ENSMUSG00 | 13437 |
| NA | ENSMUSG00 | 98733 |
| NA | ENSMUSG00 | 71728 |
| NA | ENSMUSG00 | 14897 |

| | | |
|---|---|---|
| Significant | ENSMUSG00 | 20684 |
| NA | ENSMUSG00 | 64294 |
| NA | ENSMUSG00 | 70247 |
| NA | ENSMUSG00 | 17975 |
| NA | ENSMUSG00 | 13838 |
| NA | ENSMUSG00 | 19231 |
| NA | ENSMUSG00 | 18582 |
| NA | ENSMUSG00 | 26895 |
| Significant | ENSMUSG00 | 23874 |
| NA | ENSMUSG00 | 69163 |
| NA | ENSMUSG00 | 20720 |
| NA | ENSMUSG00 | 26987 |
| NA | ENSMUSG00 | 67273 |
| NA | ENSMUSG00 | 108657 |
| NA | ENSMUSG00 | 23830 |
| NA | ENSMUSG00 | 64095 |
| NA | ENSMUSG00 | 69821 |
| NA | ENSMUSG00 | 269224 |
| NA | ENSMUSG00 | 66385 |
| NA | ENSMUSG00 | 18000 |
| NA | ENSMUSG00 | 59041 |
| NA | ENSMUSG00 | 51800 |
| NA | ENSMUSG00 | 67026 |
| NA | ENSMUSG00 | 66615 |
| NA | ENSMUSG00 | 21915 |
| NA | ENSMUSG00 | 66262 |
| NA | ENSMUSG00 | 18566 |
| NA | ENSMUSG00 | 16331 |
| NA | ENSMUSG00 | 77040 |
| NA | ENSMUSG00 | 16978 |
| NA | ENSMUSG00 | 50880 |
| NA | ENSMUSG00 | 67444 |
| NA | ENSMUSG00 | 65247 |
| NA | ENSMUSG00 | 208727 |
| Significant | ENSMUSG00 | 20725 |
| NA | ENSMUSG00 | 26889 |
| NA | ENSMUSG00 | 227446 |
| NA | ENSMUSG00 | 21934 |
| NA | ENSMUSG00 | 252876 |
| NA | ENSMUSG00 | 18484 |
| NA | ENSMUSG00 | 70829 |
| NA | ENSMUSG00 | 74117 |
| NA | ENSMUSG00 | 74150 |
| NA | ENSMUSG00 | 72949 |
| NA | ENSMUSG00 | 67812 |
| NA | ENSMUSG00 | 17219 |

| | | |
|---|---|---|
| NA | ENSMUSG00 | 226414 |
| Significant | ENSMUSG00 | 64214 |
| NA | ENSMUSG00 | 50778 |
| NA | ENSMUSG00 | 19735 |
| NA | ENSMUSG00 | 214498 |
| NA | ENSMUSG00 | 12628 |
| NA | ENSMUSG00 | 22099 |
| NA | ENSMUSG00 | 67949 |
| NA | ENSMUSG00 | 226352 |
| NA | ENSMUSG00 | 19258 |
| NA | ENSMUSG00 | 13167 |
| NA | ENSMUSG00 | 73103 |
| NA | ENSMUSG00 | 68428 |
| NA | ENSMUSG00 | 59125 |
| NA | ENSMUSG00 | 19264 |
| NA | ENSMUSG00 | 13136 |
| NA | ENSMUSG00 | 67997 |
| NA | ENSMUSG00 | 18640 |
| NA | ENSMUSG00 | 66241 |
| NA | ENSMUSG00 | 21952 |
| NA | ENSMUSG00 | 13007 |
| NA | ENSMUSG00 | 14270 |
| NA | ENSMUSG00 | 68724 |
| NA | ENSMUSG00 | 16865 |
| NA | ENSMUSG00 | 67196 |
| NA | ENSMUSG00 | 54354 |
| NA | ENSMUSG00 | 226422 |
| NA | ENSMUSG00 | 98415 |
| NA | ENSMUSG00 | 212980 |
| NA | ENSMUSG00 | 13714 |
| NA | ENSMUSG00 | 18557 |
| NA | ENSMUSG00 | 213464 |
| NA | ENSMUSG00 | 240752 |
| NA | ENSMUSG00 | 240756 |
| NA | ENSMUSG00 | 72017 |
| NA | ENSMUSG00 | 72674 |
| NA | ENSMUSG00 | 68507 |
| NA | ENSMUSG00 | 17928 |
| NA | ENSMUSG00 | 381290 |
| NA | ENSMUSG00 | 208263 |
| NA | ENSMUSG00 | 19775 |
| NA | ENSMUSG00 | 58244 |
| NA | ENSMUSG00 | 15064 |
| NA | ENSMUSG00 | 14645 |
| NA | ENSMUSG00 | 19734 |
| NA | ENSMUSG00 | 226519 |

| | | |
|---|---|---|
| NA | ENSMUSG00 | 17970 |
| NA | ENSMUSG00 | 19731 |
| NA | ENSMUSG00 | 63913 |
| NA | ENSMUSG00 | 19821 |
| NA | ENSMUSG00 | 67426 |
| NA | ENSMUSG00 | 226751 |
| NA | ENSMUSG00 | 226747 |
| NA | ENSMUSG00 | 15278 |
| NA | ENSMUSG00 | 68226 |
| NA | ENSMUSG00 | 11545 |
| NA | ENSMUSG00 | 170760 |
| NA | ENSMUSG00 | 66359 |
| NA | ENSMUSG00 | 78825 |
| NA | ENSMUSG00 | 76816 |
| NA | ENSMUSG00 | 12334 |
| NA | ENSMUSG00 | 209456 |
| NA | ENSMUSG00 | 27058 |
| NA | ENSMUSG00 | 67459 |
| NA | ENSMUSG00 | 208795 |
| NA | ENSMUSG00 | 69051 |
| NA | ENSMUSG00 | 13603 |
| NA | ENSMUSG00 | 14194 |
| Significant | ENSMUSG00 | 381308 |
| NA | ENSMUSG00 | 68440 |
| NA | ENSMUSG00 | 71870 |
| NA | ENSMUSG00 | 21346 |
| Significant | ENSMUSG00 | 98365 |
| NA | ENSMUSG00 | 12847 |
| NA | ENSMUSG00 | 98193 |
| NA | ENSMUSG00 | 93840 |
| NA | ENSMUSG00 | 80914 |
| NA | ENSMUSG00 | 27878 |
| NA | ENSMUSG00 | 240892 |
| NA | ENSMUSG00 | 18986 |
| NA | ENSMUSG00 | 68481 |
| NA | ENSMUSG00 | 70456 |
| NA | ENSMUSG00 | 74106 |
| NA | ENSMUSG00 | 171567 |
| NA | ENSMUSG00 | 11931 |
| NA | ENSMUSG00 | 66352 |
| NA | ENSMUSG00 | 74895 |
| Significant | ENSMUSG00 | 14067 |
| Significant | ENSMUSG00 | 20343 |
| NA | ENSMUSG00 | 240880 |
| NA | ENSMUSG00 | 16579 |
| NA | ENSMUSG00 | 18933 |

| | | |
|---|---|---|
| NA | ENSMUSG00 | 78255 |
| NA | ENSMUSG00 | 11352 |
| NA | ENSMUSG00 | 20652 |
| NA | ENSMUSG00 | 226830 |
| NA | ENSMUSG00 | 108000 |
| NA | ENSMUSG00 | 226823 |
| NA | ENSMUSG00 | 107508 |
| NA | ENSMUSG00 | 23827 |
| NA | ENSMUSG00 | 381314 |
| NA | ENSMUSG00 | 18005 |
| NA | ENSMUSG00 | 226856 |
| NA | ENSMUSG00 | 226849 |
| Significant | ENSMUSG00 | 66950 |
| NA | ENSMUSG00 | 11910 |
| NA | ENSMUSG00 | 381319 |
| NA | ENSMUSG00 | 68972 |
| NA | ENSMUSG00 | 52477 |
| NA | ENSMUSG00 | 22033 |
| NA | ENSMUSG00 | 18845 |
| NA | ENSMUSG00 | 22278 |
| NA | ENSMUSG00 | 18108 |
| NA | ENSMUSG00 | 64707 |
| NA | ENSMUSG00 | 227525 |
| NA | ENSMUSG00 | 75472 |
| NA | ENSMUSG00 | 104362 |
| Significant | ENSMUSG00 | 66540 |
| Significant | ENSMUSG00 | 14130 |
| NA | ENSMUSG00 | 209630 |
| NA | ENSMUSG00 | 80915 |
| NA | ENSMUSG00 | 109079 |
| NA | ENSMUSG00 | 226641 |
| NA | ENSMUSG00 | 16922 |
| NA | ENSMUSG00 | 16589 |
| NA | ENSMUSG00 | 70024 |
| NA | ENSMUSG00 | 107652 |
| NA | ENSMUSG00 | 71648 |
| NA | ENSMUSG00 | 15490 |
| NA | ENSMUSG00 | 71233 |
| NA | ENSMUSG00 | 66977 |
| NA | ENSMUSG00 | 56752 |
| NA | ENSMUSG00 | 66447 |
| NA | ENSMUSG00 | 71449 |
| NA | ENSMUSG00 | 53330 |
| NA | ENSMUSG00 | 67292 |
| Significant | ENSMUSG00 | 11758 |
| NA | ENSMUSG00 | 226541 |

| | | |
|---|---|---|
| NA | ENSMUSG00 | 74455 |
| NA | ENSMUSG00 | 70454 |
| NA | ENSMUSG00 | 226539 |
| Significant | ENSMUSG00 | 17533 |
| NA | ENSMUSG00 | 11905 |
| NA | ENSMUSG00 | 20844 |
| NA | ENSMUSG00 | 29809 |
| NA | ENSMUSG00 | 13434 |
| NA | ENSMUSG00 | 20163 |
| NA | ENSMUSG00 | 22352 |
| NA | ENSMUSG00 | 19212 |
| NA | ENSMUSG00 | 18718 |
| NA | ENSMUSG00 | 12151 |
| NA | ENSMUSG00 | 13418 |
| NA | ENSMUSG00 | 17354 |
| NA | ENSMUSG00 | 67448 |
| Significant | ENSMUSG00 | 59126 |
| NA | ENSMUSG00 | 19177 |
| NA | ENSMUSG00 | 67857 |
| NA | ENSMUSG00 | 76899 |
| NA | ENSMUSG00 | 74192 |
| NA | ENSMUSG00 | 26428 |
| NA | ENSMUSG00 | 16574 |
| NA | ENSMUSG00 | 109129 |
| NA | ENSMUSG00 | 66960 |
| NA | ENSMUSG00 | 241226 |
| NA | ENSMUSG00 | 76857 |
| NA | ENSMUSG00 | 170768 |
| NA | ENSMUSG00 | 27377 |
| NA | ENSMUSG00 | 18761 |
| NA | ENSMUSG00 | 67121 |
| NA | ENSMUSG00 | 74159 |
| NA | ENSMUSG00 | 329165 |
| NA | ENSMUSG00 | 56075 |
| NA | ENSMUSG00 | 263803 |
| NA | ENSMUSG00 | 54519 |
| NA | ENSMUSG00 | 71834 |
| NA | ENSMUSG00 | 18286 |
| Significant | ENSMUSG00 | 56017 |
| NA | ENSMUSG00 | 227738 |
| NA | ENSMUSG00 | 227737 |
| NA | ENSMUSG00 | 20910 |
| NA | ENSMUSG00 | 227683 |
| NA | ENSMUSG00 | 68975 |
| NA | ENSMUSG00 | 22130 |
| NA | ENSMUSG00 | 227674 |

| | | |
|---|---|---|
| NA | ENSMUSG00 | 68870 |
| NA | ENSMUSG00 | 69987 |
| NA | ENSMUSG00 | 13481 |
| NA | ENSMUSG00 | 50935 |
| NA | ENSMUSG00 | 64930 |
| NA | ENSMUSG00 | 13805 |
| NA | ENSMUSG00 | 14582 |
| NA | ENSMUSG00 | 70239 |
| NA | ENSMUSG00 | 11636 |
| NA | ENSMUSG00 | 227731 |
| NA | ENSMUSG00 | 96979 |
| NA | ENSMUSG00 | 19730 |
| NA | ENSMUSG00 | 18227 |
| Significant | ENSMUSG00 | 14571 |
| Significant | ENSMUSG00 | 227929 |
| Significant | ENSMUSG00 | 14134 |
| NA | ENSMUSG00 | 11477 |
| NA | ENSMUSG00 | 11350 |
| NA | ENSMUSG00 | 320267 |
| NA | ENSMUSG00 | 30934 |
| NA | ENSMUSG00 | 30931 |
| NA | ENSMUSG00 | 227707 |
| NA | ENSMUSG00 | 12908 |
| NA | ENSMUSG00 | 74270 |
| NA | ENSMUSG00 | 57170 |
| NA | ENSMUSG00 | 66617 |
| NA | ENSMUSG00 | 108958 |
| NA | ENSMUSG00 | 227700 |
| NA | ENSMUSG00 | 14828 |
| NA | ENSMUSG00 | 70789 |
| NA | ENSMUSG00 | 66691 |
| NA | ENSMUSG00 | 66998 |
| NA | ENSMUSG00 | 24136 |
| NA | ENSMUSG00 | 74016 |
| NA | ENSMUSG00 | 22029 |
| NA | ENSMUSG00 | 68365 |
| NA | ENSMUSG00 | 227753 |
| Significant | ENSMUSG00 | 13830 |
| NA | ENSMUSG00 | 69601 |
| NA | ENSMUSG00 | 67871 |
| NA | ENSMUSG00 | 67889 |
| NA | ENSMUSG00 | 227960 |
| NA | ENSMUSG00 | 75495 |
| NA | ENSMUSG00 | 68375 |
| NA | ENSMUSG00 | 71586 |
| NA | ENSMUSG00 | 59029 |

| | | |
|---|---|---|
| NA | ENSMUSG00 | 20744 |
| NA | ENSMUSG00 | 140858 |
| NA | ENSMUSG00 | 67382 |
| NA | ENSMUSG00 | 353156 |
| NA | ENSMUSG00 | 67512 |
| NA | ENSMUSG00 | 18128 |
| NA | ENSMUSG00 | 227648 |
| NA | ENSMUSG00 | 64436 |
| NA | ENSMUSG00 | 66865 |
| NA | ENSMUSG00 | 68112 |
| NA | ENSMUSG00 | 332579 |
| NA | ENSMUSG00 | 67839 |
| NA | ENSMUSG00 | 67991 |
| NA | ENSMUSG00 | 227634 |
| Significant | ENSMUSG00 | 14133 |
| Significant | ENSMUSG00 | 51875 |
| NA | ENSMUSG00 | 381352 |
| NA | ENSMUSG00 | 22030 |
| NA | ENSMUSG00 | 11305 |
| NA | ENSMUSG00 | 64685 |
| NA | ENSMUSG00 | 72080 |
| Significant | ENSMUSG00 | 227620 |
| Significant | ENSMUSG00 | 83768 |
| NA | ENSMUSG00 | 65103 |
| NA | ENSMUSG00 | 99152 |
| NA | ENSMUSG00 | 68475 |
| NA | ENSMUSG00 | 56878 |
| NA | ENSMUSG00 | 215705 |
| NA | ENSMUSG00 | 67187 |
| NA | ENSMUSG00 | 67228 |
| NA | ENSMUSG00 | 18510 |
| NA | ENSMUSG00 | 57438 |
| NA | ENSMUSG00 | 215632 |
| NA | ENSMUSG00 | 17076 |
| NA | ENSMUSG00 | 16181 |
| Significant | ENSMUSG00 | 140483 |
| NA | ENSMUSG00 | 407823 |
| NA | ENSMUSG00 | 72137 |
| NA | ENSMUSG00 | 227937 |
| NA | ENSMUSG00 | 14425 |
| NA | ENSMUSG00 | 69482 |
| NA | ENSMUSG00 | 68082 |
| NA | ENSMUSG00 | 50884 |
| NA | ENSMUSG00 | 66861 |
| NA | ENSMUSG00 | 70599 |
| NA | ENSMUSG00 | 16401 |

| | | |
|---|---|---|
| Significant | ENSMUSG00 | 78830 |
| NA | ENSMUSG00 | 22193 |
| NA | ENSMUSG00 | 13427 |
| NA | ENSMUSG00 | 80744 |
| NA | ENSMUSG00 | 107435 |
| Significant | ENSMUSG00 | 53416 |
| NA | ENSMUSG00 | 241447 |
| NA | ENSMUSG00 | 20833 |
| NA | ENSMUSG00 | 58207 |
| NA | ENSMUSG00 | 72401 |
| NA | ENSMUSG00 | 30059 |
| NA | ENSMUSG00 | 56791 |
| NA | ENSMUSG00 | 98985 |
| NA | ENSMUSG00 | 381379 |
| NA | ENSMUSG00 | 21788 |
| NA | ENSMUSG00 | 320720 |
| NA | ENSMUSG00 | 16410 |
| NA | ENSMUSG00 | 73373 |
| NA | ENSMUSG00 | 69082 |
| NA | ENSMUSG00 | 53375 |
| NA | ENSMUSG00 | 11909 |
| NA | ENSMUSG00 | 11435 |
| NA | ENSMUSG00 | 67059 |
| NA | ENSMUSG00 | 20687 |
| NA | ENSMUSG00 | 16403 |
| NA | ENSMUSG00 | 228421 |
| NA | ENSMUSG00 | 212772 |
| NA | ENSMUSG00 | 107723 |
| NA | ENSMUSG00 | 68032 |
| NA | ENSMUSG00 | 72425 |
| NA | ENSMUSG00 | 66181 |
| Significant | ENSMUSG00 | 99010 |
| NA | ENSMUSG00 | 68201 |
| NA | ENSMUSG00 | 22343 |
| NA | ENSMUSG00 | 76501 |
| NA | ENSMUSG00 | 22034 |
| NA | ENSMUSG00 | 68170 |
| NA | ENSMUSG00 | 99349 |
| NA | ENSMUSG00 | 77766 |
| NA | ENSMUSG00 | 98221 |
| NA | ENSMUSG00 | 211896 |
| NA | ENSMUSG00 | 320554 |
| NA | ENSMUSG00 | 228410 |
| NA | ENSMUSG00 | 15259 |
| NA | ENSMUSG00 | 57443 |
| NA | ENSMUSG00 | 53872 |

| | | |
|---|---|---|
| NA | ENSMUSG00 | 98956 |
| NA | ENSMUSG00 | 12359 |
| Significant | ENSMUSG00 | 80985 |
| NA | ENSMUSG00 | 11800 |
| NA | ENSMUSG00 | 74569 |
| NA | ENSMUSG00 | 56348 |
| NA | ENSMUSG00 | 14043 |
| Significant | ENSMUSG00 | 67092 |
| Significant | ENSMUSG00 | 214968 |
| NA | ENSMUSG00 | 17876 |
| NA | ENSMUSG00 | 110074 |
| NA | ENSMUSG00 | 12848 |
| NA | ENSMUSG00 | 69976 |
| NA | ENSMUSG00 | 17536 |
| NA | ENSMUSG00 | 12521 |
| Significant | ENSMUSG00 | 241556 |
| NA | ENSMUSG00 | 76969 |
| NA | ENSMUSG00 | 18633 |
| NA | ENSMUSG00 | 20322 |
| NA | ENSMUSG00 | 26427 |
| NA | ENSMUSG00 | 78655 |
| NA | ENSMUSG00 | 17242 |
| NA | ENSMUSG00 | 241627 |
| NA | ENSMUSG00 | 241547 |
| NA | ENSMUSG00 | 51897 |
| NA | ENSMUSG00 | 67693 |
| NA | ENSMUSG00 | 269344 |
| NA | ENSMUSG00 | 228359 |
| NA | ENSMUSG00 | 14827 |
| NA | ENSMUSG00 | 228357 |
| NA | ENSMUSG00 | 77038 |
| NA | ENSMUSG00 | 80708 |
| NA | ENSMUSG00 | 75894 |
| NA | ENSMUSG00 | 51885 |
| NA | ENSMUSG00 | 22222 |
| NA | ENSMUSG00 | 59030 |
| NA | ENSMUSG00 | 16449 |
| NA | ENSMUSG00 | 74243 |
| NA | ENSMUSG00 | 56428 |
| NA | ENSMUSG00 | 68968 |
| NA | ENSMUSG00 | 66296 |
| NA | ENSMUSG00 | 66606 |
| NA | ENSMUSG00 | 20619 |
| NA | ENSMUSG00 | 20402 |
| NA | ENSMUSG00 | 269338 |
| NA | ENSMUSG00 | 98878 |

| | | |
|---|---|---|
| NA | ENSMUSG00 | 22174 |
| NA | ENSMUSG00 | 140629 |
| NA | ENSMUSG00 | 19262 |
| NA | ENSMUSG00 | 76246 |
| NA | ENSMUSG00 | 69702 |
| NA | ENSMUSG00 | 108907 |
| NA | ENSMUSG00 | 228602 |
| Significant | ENSMUSG00 | 11990 |
| NA | ENSMUSG00 | 69065 |
| NA | ENSMUSG00 | 20732 |
| NA | ENSMUSG00 | 20612 |
| NA | ENSMUSG00 | 19361 |
| NA | ENSMUSG00 | 271842 |
| NA | ENSMUSG00 | 76464 |
| NA | ENSMUSG00 | 67326 |
| NA | ENSMUSG00 | 12531 |
| NA | ENSMUSG00 | 51944 |
| NA | ENSMUSG00 | 56357 |
| NA | ENSMUSG00 | 228608 |
| NA | ENSMUSG00 | 215653 |
| NA | ENSMUSG00 | 54338 |
| NA | ENSMUSG00 | 70612 |
| NA | ENSMUSG00 | 18538 |
| NA | ENSMUSG00 | 74182 |
| NA | ENSMUSG00 | 68215 |
| NA | ENSMUSG00 | 114715 |
| NA | ENSMUSG00 | 66634 |
| NA | ENSMUSG00 | 66586 |
| NA | ENSMUSG00 | 12156 |
| Significant | ENSMUSG00 | 15186 |
| NA | ENSMUSG00 | 14391 |
| NA | ENSMUSG00 | 84092 |
| NA | ENSMUSG00 | 75083 |
| NA | ENSMUSG00 | 58800 |
| NA | ENSMUSG00 | 66552 |
| NA | ENSMUSG00 | 99138 |
| NA | ENSMUSG00 | 13537 |
| NA | ENSMUSG00 | 68126 |
| NA | ENSMUSG00 | 77721 |
| NA | ENSMUSG00 | 53885 |
| NA | ENSMUSG00 | 12235 |
| NA | ENSMUSG00 | 12125 |
| NA | ENSMUSG00 | 69487 |
| NA | ENSMUSG00 | 57432 |
| NA | ENSMUSG00 | 69737 |
| NA | ENSMUSG00 | 20017 |

| | | |
|---|---|---|
| NA | ENSMUSG00 | 20515 |
| NA | ENSMUSG00 | 16176 |
| NA | ENSMUSG00 | 16175 |
| NA | ENSMUSG00 | 20638 |
| NA | ENSMUSG00 | 67134 |
| NA | ENSMUSG00 | 170718 |
| NA | ENSMUSG00 | 56264 |
| NA | ENSMUSG00 | 80743 |
| NA | ENSMUSG00 | 81910 |
| NA | ENSMUSG00 | 69178 |
| NA | ENSMUSG00 | 74528 |
| NA | ENSMUSG00 | 228714 |
| NA | ENSMUSG00 | 70408 |
| NA | ENSMUSG00 | 26450 |
| NA | ENSMUSG00 | 27054 |
| NA | ENSMUSG00 | 66044 |
| NA | ENSMUSG00 | 24128 |
| Significant | ENSMUSG00 | 17064 |
| NA | ENSMUSG00 | 17957 |
| NA | ENSMUSG00 | 74533 |
| NA | ENSMUSG00 | 13010 |
| Significant | ENSMUSG00 | 68738 |
| NA | ENSMUSG00 | 69270 |
| NA | ENSMUSG00 | 386649 |
| NA | ENSMUSG00 | 73847 |
| NA | ENSMUSG00 | 11602 |
| NA | ENSMUSG00 | 69698 |
| NA | ENSMUSG00 | 67231 |
| NA | ENSMUSG00 | 24105 |
| NA | ENSMUSG00 | 72119 |
| NA | ENSMUSG00 | 68559 |
| NA | ENSMUSG00 | 16569 |
| NA | ENSMUSG00 | 13436 |
| NA | ENSMUSG00 | 13589 |
| NA | ENSMUSG00 | 66971 |
| Significant | ENSMUSG00 | 20648 |
| NA | ENSMUSG00 | 13555 |
| NA | ENSMUSG00 | 67017 |
| NA | ENSMUSG00 | 20878 |
| NA | ENSMUSG00 | 67337 |
| NA | ENSMUSG00 | 18767 |
| NA | ENSMUSG00 | 66404 |
| NA | ENSMUSG00 | 21985 |
| NA | ENSMUSG00 | 94212 |
| NA | ENSMUSG00 | 66679 |
| NA | ENSMUSG00 | 56190 |

| | | |
|---|---|---|
| NA | ENSMUSG00 | 58203 |
| NA | ENSMUSG00 | 19334 |
| NA | ENSMUSG00 | 228960 |
| NA | ENSMUSG00 | 14683 |
| NA | ENSMUSG00 | 55980 |
| NA | ENSMUSG00 | 16592 |
| NA | ENSMUSG00 | 74718 |
| NA | ENSMUSG00 | 66371 |
| NA | ENSMUSG00 | 19246 |
| Significant | ENSMUSG00 | 18019 |
| NA | ENSMUSG00 | 11981 |
| NA | ENSMUSG00 | 99377 |
| NA | ENSMUSG00 | 71710 |
| NA | ENSMUSG00 | 22722 |
| NA | ENSMUSG00 | 13559 |
| Significant | ENSMUSG00 | 71934 |
| NA | ENSMUSG00 | 12350 |
| NA | ENSMUSG00 | 12349 |
| NA | ENSMUSG00 | 26444 |
| NA | ENSMUSG00 | 73247 |
| NA | ENSMUSG00 | 76425 |
| NA | ENSMUSG00 | 228998 |
| NA | ENSMUSG00 | 229003 |
| NA | ENSMUSG00 | 229007 |
| NA | ENSMUSG00 | 72147 |
| NA | ENSMUSG00 | 245867 |
| NA | ENSMUSG00 | 19383 |
| NA | ENSMUSG00 | 269378 |
| NA | ENSMUSG00 | 16396 |
| NA | ENSMUSG00 | 74252 |
| NA | ENSMUSG00 | 67472 |
| NA | ENSMUSG00 | 66734 |
| NA | ENSMUSG00 | 60525 |
| NA | ENSMUSG00 | 14854 |
| NA | ENSMUSG00 | 19124 |
| NA | ENSMUSG00 | 16418 |
| NA | ENSMUSG00 | 12807 |
| NA | ENSMUSG00 | 18041 |
| NA | ENSMUSG00 | 170791 |
| NA | ENSMUSG00 | 13821 |
| NA | ENSMUSG00 | 68295 |
| NA | ENSMUSG00 | 81004 |
| NA | ENSMUSG00 | 29812 |
| NA | ENSMUSG00 | 66934 |
| NA | ENSMUSG00 | 67388 |
| NA | ENSMUSG00 | 56045 |

| | | |
|---|---|---|
| NA | ENSMUSG00 | 19650 |
| NA | ENSMUSG00 | 20014 |
| NA | ENSMUSG00 | 20779 |
| NA | ENSMUSG00 | 66642 |
| NA | ENSMUSG00 | 75425 |
| NA | ENSMUSG00 | 70470 |
| NA | ENSMUSG00 | 228850 |
| NA | ENSMUSG00 | 71878 |
| NA | ENSMUSG00 | 71715 |
| NA | ENSMUSG00 | 20482 |
| NA | ENSMUSG00 | 22401 |
| NA | ENSMUSG00 | 18706 |
| NA | ENSMUSG00 | 67778 |
| NA | ENSMUSG00 | 67414 |
| NA | ENSMUSG00 | 14696 |
| NA | ENSMUSG00 | 74614 |
| NA | ENSMUSG00 | 56456 |
| NA | ENSMUSG00 | 66046 |
| NA | ENSMUSG00 | 67120 |
| NA | ENSMUSG00 | 17979 |
| NA | ENSMUSG00 | 67713 |
| NA | ENSMUSG00 | 14359 |
| NA | ENSMUSG00 | 665113 |
| NA | ENSMUSG00 | 18805 |
| NA | ENSMUSG00 | 320024 |
| NA | ENSMUSG00 | 13605 |
| NA | ENSMUSG00 | 71827 |
| NA | ENSMUSG00 | 69276 |
| NA | ENSMUSG00 | 114893 |
| NA | ENSMUSG00 | 72039 |
| NA | ENSMUSG00 | 229211 |
| Significant | ENSMUSG00 | 11747 |
| NA | ENSMUSG00 | 50911 |
| NA | ENSMUSG00 | 12428 |
| NA | ENSMUSG00 | 57815 |
| NA | ENSMUSG00 | 26570 |
| NA | ENSMUSG00 | 19338 |
| NA | ENSMUSG00 | 67542 |
| NA | ENSMUSG00 | 67890 |
| NA | ENSMUSG00 | 50706 |
| NA | ENSMUSG00 | 56790 |
| NA | ENSMUSG00 | 69639 |
| NA | ENSMUSG00 | 56758 |
| NA | ENSMUSG00 | 18441 |
| NA | ENSMUSG00 | 72162 |
| NA | ENSMUSG00 | 28030 |

| | | |
|---|---|---|
| NA | ENSMUSG00 | 66868 |
| NA | ENSMUSG00 | 30953 |
| NA | ENSMUSG00 | 68259 |
| NA | ENSMUSG00 | 16649 |
| NA | ENSMUSG00 | 97112 |
| NA | ENSMUSG00 | 26422 |
| NA | ENSMUSG00 | 97064 |
| Significant | ENSMUSG00 | 67738 |
| NA | ENSMUSG00 | 18645 |
| NA | ENSMUSG00 | 72033 |
| NA | ENSMUSG00 | 28146 |
| NA | ENSMUSG00 | 66841 |
| NA | ENSMUSG00 | 229317 |
| NA | ENSMUSG00 | 75939 |
| NA | ENSMUSG00 | 11416 |
| NA | ENSMUSG00 | 229363 |
| NA | ENSMUSG00 | 16497 |
| NA | ENSMUSG00 | 67437 |
| NA | ENSMUSG00 | 56706 |
| NA | ENSMUSG00 | 20713 |
| NA | ENSMUSG00 | 56426 |
| NA | ENSMUSG00 | 19260 |
| NA | ENSMUSG00 | 140917 |
| NA | ENSMUSG00 | 99543 |
| NA | ENSMUSG00 | 18176 |
| NA | ENSMUSG00 | 66641 |
| NA | ENSMUSG00 | 70747 |
| NA | ENSMUSG00 | 229658 |
| NA | ENSMUSG00 | 14547 |
| NA | ENSMUSG00 | 18129 |
| NA | ENSMUSG00 | 20333 |
| NA | ENSMUSG00 | 66921 |
| NA | ENSMUSG00 | 20912 |
| NA | ENSMUSG00 | 76123 |
| NA | ENSMUSG00 | 229725 |
| NA | ENSMUSG00 | 109674 |
| NA | ENSMUSG00 | 14865 |
| NA | ENSMUSG00 | 229709 |
| NA | ENSMUSG00 | 67171 |
| NA | ENSMUSG00 | 72121 |
| NA | ENSMUSG00 | 53975 |
| Significant | ENSMUSG00 | 20195 |
| NA | ENSMUSG00 | 18160 |
| NA | ENSMUSG00 | 26568 |
| NA | ENSMUSG00 | 229543 |
| NA | ENSMUSG00 | 68328 |

| | | |
|---|---|---|
| NA | ENSMUSG00 | 74343 |
| NA | ENSMUSG00 | 23922 |
| NA | ENSMUSG00 | 59069 |
| NA | ENSMUSG00 | 99650 |
| NA | ENSMUSG00 | 23897 |
| NA | ENSMUSG00 | 16194 |
| NA | ENSMUSG00 | 11444 |
| NA | ENSMUSG00 | 56417 |
| NA | ENSMUSG00 | 68603 |
| NA | ENSMUSG00 | 19729 |
| NA | ENSMUSG00 | 13636 |
| NA | ENSMUSG00 | 68659 |
| Significant | ENSMUSG00 | 70652 |
| NA | ENSMUSG00 | 229782 |
| NA | ENSMUSG00 | 72776 |
| NA | ENSMUSG00 | 109245 |
| NA | ENSMUSG00 | 22329 |
| NA | ENSMUSG00 | 58193 |
| NA | ENSMUSG00 | 28036 |
| NA | ENSMUSG00 | 67225 |
| NA | ENSMUSG00 | 15107 |
| NA | ENSMUSG00 | 80890 |
| NA | ENSMUSG00 | 66815 |
| NA | ENSMUSG00 | 24088 |
| NA | ENSMUSG00 | 20319 |
| NA | ENSMUSG00 | 12368 |
| NA | ENSMUSG00 | 53317 |
| NA | ENSMUSG00 | 66350 |
| NA | ENSMUSG00 | 76561 |
| NA | ENSMUSG00 | 68147 |
| NA | ENSMUSG00 | 74776 |
| Significant | ENSMUSG00 | 229445 |
| NA | ENSMUSG00 | 71793 |
| NA | ENSMUSG00 | 13645 |
| NA | ENSMUSG00 | 67553 |
| NA | ENSMUSG00 | 71481 |
| NA | ENSMUSG00 | 13722 |
| NA | ENSMUSG00 | 271981 |
| NA | ENSMUSG00 | 23971 |
| Significant | ENSMUSG00 | 51886 |
| NA | ENSMUSG00 | 67035 |
| NA | ENSMUSG00 | 13639 |
| NA | ENSMUSG00 | 11490 |
| NA | ENSMUSG00 | 22724 |
| NA | ENSMUSG00 | 54124 |
| NA | ENSMUSG00 | 14466 |

| | | |
|---|---|---|
| NA | ENSMUSG00 | 24045 |
| NA | ENSMUSG00 | 192195 |
| NA | ENSMUSG00 | 19769 |
| NA | ENSMUSG00 | 16800 |
| NA | ENSMUSG00 | 74200 |
| NA | ENSMUSG00 | 83409 |
| NA | ENSMUSG00 | 16905 |
| Significant | ENSMUSG00 | 20351 |
| NA | ENSMUSG00 | 67037 |
| NA | ENSMUSG00 | 404710 |
| NA | ENSMUSG00 | 66614 |
| NA | ENSMUSG00 | 246703 |
| NA | ENSMUSG00 | 73182 |
| NA | ENSMUSG00 | 12479 |
| NA | ENSMUSG00 | 80877 |
| NA | ENSMUSG00 | 20091 |
| NA | ENSMUSG00 | 27059 |
| NA | ENSMUSG00 | 229487 |
| NA | ENSMUSG00 | 50754 |
| NA | ENSMUSG00 | 14263 |
| NA | ENSMUSG00 | 68058 |
| NA | ENSMUSG00 | 66659 |
| NA | ENSMUSG00 | 67549 |
| NA | ENSMUSG00 | 67845 |
| NA | ENSMUSG00 | 74414 |
| NA | ENSMUSG00 | 78373 |
| NA | ENSMUSG00 | 229615 |
| NA | ENSMUSG00 | 18632 |
| NA | ENSMUSG00 | 69870 |
| NA | ENSMUSG00 | 75137 |
| NA | ENSMUSG00 | 71807 |
| NA | ENSMUSG00 | 13601 |
| NA | ENSMUSG00 | 210529 |
| NA | ENSMUSG00 | 29815 |
| NA | ENSMUSG00 | 14630 |
| NA | ENSMUSG00 | 18720 |
| NA | ENSMUSG00 | 19299 |
| NA | ENSMUSG00 | 14066 |
| NA | ENSMUSG00 | 66568 |
| NA | ENSMUSG00 | 56195 |
| NA | ENSMUSG00 | 76742 |
| NA | ENSMUSG00 | 11532 |
| NA | ENSMUSG00 | 78523 |
| NA | ENSMUSG00 | 75778 |
| NA | ENSMUSG00 | 229877 |
| NA | ENSMUSG00 | 56224 |

| | | |
|---|---|---|
| NA | ENSMUSG00 | 13684 |
| NA | ENSMUSG00 | 17777 |
| NA | ENSMUSG00 | 26377 |
| NA | ENSMUSG00 | 19055 |
| NA | ENSMUSG00 | 18033 |
| NA | ENSMUSG00 | 110173 |
| NA | ENSMUSG00 | 67006 |
| Significant | ENSMUSG00 | 68151 |
| NA | ENSMUSG00 | 76131 |
| NA | ENSMUSG00 | 107869 |
| NA | ENSMUSG00 | 53861 |
| NA | ENSMUSG00 | 99633 |
| NA | ENSMUSG00 | 70285 |
| NA | ENSMUSG00 | 70951 |
| NA | ENSMUSG00 | 74245 |
| NA | ENSMUSG00 | 12042 |
| Significant | ENSMUSG00 | 69219 |
| Significant | ENSMUSG00 | 16007 |
| NA | ENSMUSG00 | 12972 |
| NA | ENSMUSG00 | 65973 |
| NA | ENSMUSG00 | 60599 |
| NA | ENSMUSG00 | 12448 |
| NA | ENSMUSG00 | 14579 |
| NA | ENSMUSG00 | 68099 |
| NA | ENSMUSG00 | 72519 |
| NA | ENSMUSG00 | 67460 |
| NA | ENSMUSG00 | 27354 |
| NA | ENSMUSG00 | 70568 |
| NA | ENSMUSG00 | 66302 |
| NA | ENSMUSG00 | 72098 |
| NA | ENSMUSG00 | 116940 |
| NA | ENSMUSG00 | 67427 |
| NA | ENSMUSG00 | 242341 |
| NA | ENSMUSG00 | 68053 |
| NA | ENSMUSG00 | 18201 |
| NA | ENSMUSG00 | 230027 |
| NA | ENSMUSG00 | 66625 |
| Significant | ENSMUSG00 | 53378 |
| NA | ENSMUSG00 | 77032 |
| NA | ENSMUSG00 | 51813 |
| NA | ENSMUSG00 | 52184 |
| NA | ENSMUSG00 | 68493 |
| NA | ENSMUSG00 | 16911 |
| NA | ENSMUSG00 | 55932 |
| NA | ENSMUSG00 | 14469 |
| NA | ENSMUSG00 | 229906 |

| | | |
|---|---|---|
| NA | ENSMUSG00 | 56376 |
| NA | ENSMUSG00 | 24018 |
| NA | ENSMUSG00 | 56228 |
| NA | ENSMUSG00 | 26885 |
| NA | ENSMUSG00 | 26409 |
| NA | ENSMUSG00 | 433693 |
| NA | ENSMUSG00 | 109093 |
| NA | ENSMUSG00 | 24060 |
| NA | ENSMUSG00 | 66291 |
| Significant | ENSMUSG00 | 73205 |
| NA | ENSMUSG00 | 109331 |
| NA | ENSMUSG00 | 14211 |
| NA | ENSMUSG00 | 64424 |
| NA | ENSMUSG00 | 66362 |
| NA | ENSMUSG00 | 21916 |
| NA | ENSMUSG00 | 22590 |
| NA | ENSMUSG00 | 433702 |
| NA | ENSMUSG00 | 74753 |
| NA | ENSMUSG00 | 67628 |
| NA | ENSMUSG00 | 94181 |
| Significant | ENSMUSG00 | 107684 |
| NA | ENSMUSG00 | 12819 |
| NA | ENSMUSG00 | 18124 |
| NA | ENSMUSG00 | 76299 |
| NA | ENSMUSG00 | 16348 |
| NA | ENSMUSG00 | 269536 |
| NA | ENSMUSG00 | 230157 |
| NA | ENSMUSG00 | 22166 |
| NA | ENSMUSG00 | 67103 |
| NA | ENSMUSG00 | 22234 |
| NA | ENSMUSG00 | 230257 |
| NA | ENSMUSG00 | 72479 |
| NA | ENSMUSG00 | 209131 |
| NA | ENSMUSG00 | 22696 |
| NA | ENSMUSG00 | 71354 |
| NA | ENSMUSG00 | 17025 |
| NA | ENSMUSG00 | 59001 |
| NA | ENSMUSG00 | 76804 |
| NA | ENSMUSG00 | 66928 |
| NA | ENSMUSG00 | 17475 |
| NA | ENSMUSG00 | 68268 |
| NA | ENSMUSG00 | 11428 |
| NA | ENSMUSG00 | 100504309 |
| NA | ENSMUSG00 | 74255 |
| NA | ENSMUSG00 | 15502 |
| NA | ENSMUSG00 | 66408 |

| | | |
|---|---|---|
| NA | ENSMUSG00 | 100434 |
| NA | ENSMUSG00 | 14595 |
| NA | ENSMUSG00 | 246179 |
| NA | ENSMUSG00 | 12017 |
| NA | ENSMUSG00 | 76959 |
| NA | ENSMUSG00 | 52076 |
| NA | ENSMUSG00 | 74164 |
| NA | ENSMUSG00 | 19359 |
| NA | ENSMUSG00 | 230082 |
| NA | ENSMUSG00 | 230233 |
| NA | ENSMUSG00 | 68926 |
| NA | ENSMUSG00 | 68970 |
| NA | ENSMUSG00 | 67123 |
| NA | ENSMUSG00 | 109242 |
| NA | ENSMUSG00 | 71901 |
| NA | ENSMUSG00 | 66401 |
| NA | ENSMUSG00 | 69638 |
| NA | ENSMUSG00 | 53598 |
| NA | ENSMUSG00 | 269523 |
| NA | ENSMUSG00 | 60534 |
| NA | ENSMUSG00 | 56703 |
| NA | ENSMUSG00 | 66592 |
| NA | ENSMUSG00 | 22249 |
| NA | ENSMUSG00 | 21754 |
| NA | ENSMUSG00 | 12517 |
| NA | ENSMUSG00 | 622404 |
| NA | ENSMUSG00 | 230099 |
| NA | ENSMUSG00 | 22004 |
| NA | ENSMUSG00 | 21894 |
| NA | ENSMUSG00 | 12913 |
| NA | ENSMUSG00 | 230101 |
| NA | ENSMUSG00 | 242406 |
| NA | ENSMUSG00 | 68917 |
| NA | ENSMUSG00 | 53614 |
| Significant | ENSMUSG00 | 12757 |
| NA | ENSMUSG00 | 50798 |
| NA | ENSMUSG00 | 384009 |
| NA | ENSMUSG00 | 77634 |
| NA | ENSMUSG00 | 101739 |
| NA | ENSMUSG00 | 11520 |
| NA | ENSMUSG00 | 20104 |
| NA | ENSMUSG00 | 70122 |
| Significant | ENSMUSG00 | 66775 |
| NA | ENSMUSG00 | 329908 |
| NA | ENSMUSG00 | 67916 |
| NA | ENSMUSG00 | 242585 |

| | | |
|---|---|---|
| NA | ENSMUSG00 | 71148 |
| NA | ENSMUSG00 | 73094 |
| NA | ENSMUSG00 | 18578 |
| NA | ENSMUSG00 | 11639 |
| NA | ENSMUSG00 | 16451 |
| NA | ENSMUSG00 | 320508 |
| NA | ENSMUSG00 | 20441 |
| NA | ENSMUSG00 | 67728 |
| NA | ENSMUSG00 | 53418 |
| NA | ENSMUSG00 | 67621 |
| NA | ENSMUSG00 | 15572 |
| NA | ENSMUSG00 | 67733 |
| NA | ENSMUSG00 | 242557 |
| NA | ENSMUSG00 | 12580 |
| NA | ENSMUSG00 | 13858 |
| NA | ENSMUSG00 | 230603 |
| NA | ENSMUSG00 | 67299 |
| NA | ENSMUSG00 | 29864 |
| Significant | ENSMUSG00 | 100273 |
| NA | ENSMUSG00 | 230484 |
| NA | ENSMUSG00 | 94043 |
| NA | ENSMUSG00 | 18027 |
| NA | ENSMUSG00 | 66073 |
| NA | ENSMUSG00 | 70533 |
| NA | ENSMUSG00 | 75578 |
| NA | ENSMUSG00 | 67694 |
| NA | ENSMUSG00 | 18786 |
| NA | ENSMUSG00 | 67770 |
| NA | ENSMUSG00 | 80912 |
| NA | ENSMUSG00 | 16792 |
| NA | ENSMUSG00 | 319965 |
| NA | ENSMUSG00 | 14726 |
| NA | ENSMUSG00 | 18392 |
| NA | ENSMUSG00 | 67305 |
| NA | ENSMUSG00 | 21938 |
| NA | ENSMUSG00 | 52430 |
| NA | ENSMUSG00 | 21941 |
| NA | ENSMUSG00 | 20280 |
| NA | ENSMUSG00 | 12896 |
| NA | ENSMUSG00 | 74098 |
| NA | ENSMUSG00 | 17149 |
| NA | ENSMUSG00 | 16975 |
| NA | ENSMUSG00 | 72787 |
| NA | ENSMUSG00 | 77809 |
| NA | ENSMUSG00 | 56374 |
| NA | ENSMUSG00 | 66526 |

| | | |
|---|---|---|
| NA | ENSMUSG00 | 230582 |
| NA | ENSMUSG00 | 56280 |
| NA | ENSMUSG00 | 73172 |
| NA | ENSMUSG00 | 69181 |
| NA | ENSMUSG00 | 51797 |
| NA | ENSMUSG00 | 16656 |
| NA | ENSMUSG00 | 106564 |
| NA | ENSMUSG00 | 22608 |
| NA | ENSMUSG00 | 56401 |
| NA | ENSMUSG00 | 69216 |
| NA | ENSMUSG00 | 20525 |
| NA | ENSMUSG00 | 54170 |
| NA | ENSMUSG00 | 56309 |
| NA | ENSMUSG00 | 595136 |
| NA | ENSMUSG00 | 11426 |
| NA | ENSMUSG00 | 56031 |
| NA | ENSMUSG00 | 66966 |
| NA | ENSMUSG00 | 16918 |
| NA | ENSMUSG00 | 76574 |
| NA | ENSMUSG00 | 12331 |
| NA | ENSMUSG00 | 19063 |
| NA | ENSMUSG00 | 27224 |
| NA | ENSMUSG00 | 66193 |
| NA | ENSMUSG00 | 26394 |
| NA | ENSMUSG00 | 74246 |
| NA | ENSMUSG00 | 15356 |
| NA | ENSMUSG00 | 71665 |
| NA | ENSMUSG00 | 52830 |
| NA | ENSMUSG00 | 14105 |
| NA | ENSMUSG00 | 66743 |
| NA | ENSMUSG00 | 73804 |
| NA | ENSMUSG00 | 12795 |
| NA | ENSMUSG00 | 108067 |
| NA | ENSMUSG00 | 22275 |
| NA | ENSMUSG00 | 70603 |
| NA | ENSMUSG00 | 68276 |
| NA | ENSMUSG00 | 68394 |
| NA | ENSMUSG00 | 67096 |
| NA | ENSMUSG00 | 18477 |
| NA | ENSMUSG00 | 58810 |
| NA | ENSMUSG00 | 50927 |
| NA | ENSMUSG00 | 16351 |
| NA | ENSMUSG00 | 18710 |
| NA | ENSMUSG00 NA | |
| NA | ENSMUSG00 | 19366 |
| NA | ENSMUSG00 | 230654 |

| | | |
|---|---|---|
| NA | ENSMUSG00 | 72181 |
| NA | ENSMUSG00 | 17346 |
| NA | ENSMUSG00 | 100465 |
| NA | ENSMUSG00 | 230649 |
| NA | ENSMUSG00 | 21349 |
| NA | ENSMUSG00 | 20460 |
| NA | ENSMUSG00 | 66588 |
| NA | ENSMUSG00 | 69072 |
| NA | ENSMUSG00 | 212647 |
| NA | ENSMUSG00 | 69902 |
| NA | ENSMUSG00 | 110198 |
| Significant | ENSMUSG00 | 212555 |
| NA | ENSMUSG00 | 12345 |
| NA | ENSMUSG00 | 68943 |
| NA | ENSMUSG00 | 13200 |
| NA | ENSMUSG00 | 15441 |
| NA | ENSMUSG00 | 230861 |
| NA | ENSMUSG00 | 15530 |
| NA | ENSMUSG00 | 19248 |
| NA | ENSMUSG00 | 619605 |
| NA | ENSMUSG00 | 14077 |
| NA | ENSMUSG00 | 94242 |
| NA | ENSMUSG00 | 67898 |
| NA | ENSMUSG00 | 19244 |
| NA | ENSMUSG00 | 20218 |
| NA | ENSMUSG00 | 11637 |
| NA | ENSMUSG00 | 75234 |
| NA | ENSMUSG00 | 66264 |
| NA | ENSMUSG00 | 54383 |
| NA | ENSMUSG00 | 76799 |
| NA | ENSMUSG00 | 54709 |
| NA | ENSMUSG00 | 230761 |
| NA | ENSMUSG00 | 433759 |
| NA | ENSMUSG00 | 74552 |
| NA | ENSMUSG00 | 73680 |
| NA | ENSMUSG00 | 51796 |
| NA | ENSMUSG00 | 107271 |
| NA | ENSMUSG00 | 71514 |
| NA | ENSMUSG00 | 68592 |
| NA | ENSMUSG00 | 71817 |
| NA | ENSMUSG00 | 66146 |
| NA | ENSMUSG00 | 100317 |
| NA | ENSMUSG00 | 16765 |
| NA | ENSMUSG00 | 26562 |
| NA | ENSMUSG00 | 26445 |
| NA | ENSMUSG00 | 68040 |

| | | |
|---|---|---|
| NA | ENSMUSG00 | 214150 |
| NA | ENSMUSG00 | 73723 |
| NA | ENSMUSG00 | 27096 |
| NA | ENSMUSG00 | 100210 |
| NA | ENSMUSG00 | 245877 |
| NA | ENSMUSG00 | 242691 |
| NA | ENSMUSG00 | 18221 |
| NA | ENSMUSG00 | 20544 |
| NA | ENSMUSG00 | 52174 |
| Significant | ENSMUSG00 | 12986 |
| NA | ENSMUSG00 | 66407 |
| NA | ENSMUSG00 | 53608 |
| NA | ENSMUSG00 | 70088 |
| NA | ENSMUSG00 | 15234 |
| NA | ENSMUSG00 | 242687 |
| NA | ENSMUSG00 | 230737 |
| NA | ENSMUSG00 | 192199 |
| NA | ENSMUSG00 | 52276 |
| Significant | ENSMUSG00 | 14191 |
| NA | ENSMUSG00 | 230789 |
| NA | ENSMUSG00 | 100226 |
| NA | ENSMUSG00 | 100336 |
| NA | ENSMUSG00 | 20346 |
| NA | ENSMUSG00 | 19891 |
| NA | ENSMUSG00 | 100340 |
| NA | ENSMUSG00 | 14050 |
| NA | ENSMUSG00 | 230734 |
| NA | ENSMUSG00 | 17764 |
| NA | ENSMUSG00 | 230784 |
| NA | ENSMUSG00 | 16330 |
| Significant | ENSMUSG00 | 100088 |
| NA | ENSMUSG00 | 71787 |
| NA | ENSMUSG00 | 66464 |
| NA | ENSMUSG00 | 56809 |
| NA | ENSMUSG00 | 75062 |
| NA | ENSMUSG00 | 269587 |
| NA | ENSMUSG00 | 67205 |
| NA | ENSMUSG00 | 26922 |
| NA | ENSMUSG00 | 57317 |
| NA | ENSMUSG00 | 12371 |
| NA | ENSMUSG00 | 69582 |
| NA | ENSMUSG00 | 213499 |
| NA | ENSMUSG00 | 66147 |
| Significant | ENSMUSG00 | 18647 |
| NA | ENSMUSG00 | 18600 |
| NA | ENSMUSG00 | 16498 |

| | | |
|---|---|---|
| NA | ENSMUSG00 | 19181 |
| NA | ENSMUSG00 | 57434 |
| NA | ENSMUSG00 | 19934 |
| Significant | ENSMUSG00 | 70025 |
| NA | ENSMUSG00 | 108099 |
| NA | ENSMUSG00 | 19744 |
| NA | ENSMUSG00 | 74035 |
| NA | ENSMUSG00 | 100090 |
| NA | ENSMUSG00 | 27407 |
| NA | ENSMUSG00 | 53312 |
| Significant | ENSMUSG00 | 22319 |
| NA | ENSMUSG00 | 18628 |
| NA | ENSMUSG00 | 64295 |
| NA | ENSMUSG00 | 66587 |
| NA | ENSMUSG00 | 63958 |
| NA | ENSMUSG00 | 110208 |
| NA | ENSMUSG00 | 20535 |
| NA | ENSMUSG00 | 57320 |
| NA | ENSMUSG00 | 74155 |
| NA | ENSMUSG00 | 12568 |
| NA | ENSMUSG00 | 18669 |
| NA | ENSMUSG00 | 12353 |
| NA | ENSMUSG00 | 74610 |
| NA | ENSMUSG00 | 13347 |
| NA | ENSMUSG00 | 56273 |
| NA | ENSMUSG00 | 56485 |
| NA | ENSMUSG00 | 69743 |
| NA | ENSMUSG00 | 18127 |
| NA | ENSMUSG00 | 100198 |
| NA | ENSMUSG00 | 70556 |
| NA | ENSMUSG00 | 52323 |
| NA | ENSMUSG00 | 67087 |
| NA | ENSMUSG00 | 69151 |
| NA | ENSMUSG00 | 56717 |
| NA | ENSMUSG00 | 66454 |
| Significant | ENSMUSG00 | 84652 |
| NA | ENSMUSG00 | 66169 |
| NA | ENSMUSG00 | 72772 |
| NA | ENSMUSG00 | 71890 |
| NA | ENSMUSG00 | 69188 |
| NA | ENSMUSG00 | 11610 |
| NA | ENSMUSG00 | 17769 |
| NA | ENSMUSG00 | 26429 |
| NA | ENSMUSG00 | 22791 |
| NA | ENSMUSG00 | 26372 |
| NA | ENSMUSG00 | 73078 |

| | | |
|---|---|---|
| NA | ENSMUSG00 | 170731 |
| NA | ENSMUSG00 | 28010 |
| NA | ENSMUSG00 | 13368 |
| NA | ENSMUSG00 | 72946 |
| NA | ENSMUSG00 | 59002 |
| NA | ENSMUSG00 | 67808 |
| NA | ENSMUSG00 | 140500 |
| NA | ENSMUSG00 | 71957 |
| NA | ENSMUSG00 | 108888 |
| NA | ENSMUSG00 | 68991 |
| NA | ENSMUSG00 | 668173 |
| NA | ENSMUSG00 | 67830 |
| NA | ENSMUSG00 | 76866 |
| NA | ENSMUSG00 | 20481 |
| NA | ENSMUSG00 | 269614 |
| NA | ENSMUSG00 | 66469 |
| NA | ENSMUSG00 | 76580 |
| NA | ENSMUSG00 | 12537 |
| NA | ENSMUSG00 | 192185 |
| NA | ENSMUSG00 | 14688 |
| NA | ENSMUSG00 | 66448 |
| NA | ENSMUSG00 | 56036 |
| Significant | ENSMUSG00 | 74761 |
| NA | ENSMUSG00 | 13542 |
| NA | ENSMUSG00 | 79554 |
| NA | ENSMUSG00 | 20318 |
| NA | ENSMUSG00 | 12182 |
| Significant | ENSMUSG00 | 12494 |
| NA | ENSMUSG00 | 19126 |
| NA | ENSMUSG00 NA | |
| NA | ENSMUSG00 | 70693 |
| NA | ENSMUSG00 | 70292 |
| NA | ENSMUSG00 | 78558 |
| NA | ENSMUSG00 | 78890 |
| NA | ENSMUSG00 | 80911 |
| NA | ENSMUSG00 | 71729 |
| NA | ENSMUSG00 | 16976 |
| NA | ENSMUSG00 | 15194 |
| NA | ENSMUSG00 | 11518 |
| NA | ENSMUSG00 | 54216 |
| NA | ENSMUSG00 | 19822 |
| NA | ENSMUSG00 | 24116 |
| NA | ENSMUSG00 | 17160 |
| NA | ENSMUSG00 | 12933 |
| NA | ENSMUSG00 | 59056 |
| NA | ENSMUSG00 | 71116 |

| | | |
|---|---|---|
| NA | ENSMUSG00 | 18196 |
| NA | ENSMUSG00 | 75079 |
| NA | ENSMUSG00 | 100972 |
| NA | ENSMUSG00 | 23950 |
| NA | ENSMUSG00 | 14284 |
| NA | ENSMUSG00 | 71336 |
| NA | ENSMUSG00 | 20534 |
| NA | ENSMUSG00 | 13667 |
| NA | ENSMUSG00 | 266781 |
| NA | ENSMUSG00 | 14208 |
| NA | ENSMUSG00 | 192292 |
| NA | ENSMUSG00 | 68095 |
| NA | ENSMUSG00 | 433904 |
| NA | ENSMUSG00 | 24051 |
| NA | ENSMUSG00 | 16548 |
| NA | ENSMUSG00 | 100952 |
| NA | ENSMUSG00 | 231093 |
| NA | ENSMUSG00 | 100732 |
| NA | ENSMUSG00 | 13204 |
| NA | ENSMUSG00 | 66681 |
| NA | ENSMUSG00 | 211006 |
| NA | ENSMUSG00 | 57915 |
| NA | ENSMUSG00 | 74919 |
| NA | ENSMUSG00 | 52206 |
| NA | ENSMUSG00 | 12615 |
| NA | ENSMUSG00 | 16599 |
| NA | ENSMUSG00 | 78796 |
| NA | ENSMUSG00 | 68303 |
| NA | ENSMUSG00 | 67073 |
| NA | ENSMUSG00 | 320661 |
| NA | ENSMUSG00 | 19687 |
| NA | ENSMUSG00 | 100855 |
| NA | ENSMUSG00 | 231151 |
| NA | ENSMUSG00 | 17713 |
| NA | ENSMUSG00 | 79464 |
| NA | ENSMUSG00 | 22235 |
| NA | ENSMUSG00 | 71521 |
| NA | ENSMUSG00 | 53323 |
| NA | ENSMUSG00 | 74734 |
| NA | ENSMUSG00 | 11787 |
| NA | ENSMUSG00 | 231279 |
| NA | ENSMUSG00 | 67980 |
| NA | ENSMUSG00 | 27784 |
| NA | ENSMUSG00 | 21682 |
| NA | ENSMUSG00 | 109108 |
| NA | ENSMUSG00 | 22223 |

| | | |
|---|---|---|
| NA | ENSMUSG00 | 66899 |
| NA | ENSMUSG00 | 74277 |
| NA | ENSMUSG00 | 57357 |
| NA | ENSMUSG00 | 21982 |
| NA | ENSMUSG00 | 12753 |
| NA | ENSMUSG00 | 231327 |
| NA | ENSMUSG00 | 67054 |
| NA | ENSMUSG00 | 19712 |
| NA | ENSMUSG00 | 231329 |
| NA | ENSMUSG00 | 12617 |
| NA | ENSMUSG00 | 56792 |
| NA | ENSMUSG00 | 433931 |
| NA | ENSMUSG00 | 13486 |
| NA | ENSMUSG00 | 17765 |
| NA | ENSMUSG00 | 67266 |
| NA | ENSMUSG00 | 170823 |
| NA | ENSMUSG00 | 12545 |
| NA | ENSMUSG00 | 21814 |
| NA | ENSMUSG00 | 54367 |
| NA | ENSMUSG00 | 52822 |
| NA | ENSMUSG00 | 236573 |
| NA | ENSMUSG00 | 27405 |
| NA | ENSMUSG00 | 20750 |
| NA | ENSMUSG00 | 13602 |
| NA | ENSMUSG00 | 74167 |
| NA | ENSMUSG00 | 114664 |
| NA | ENSMUSG00 | 246293 |
| NA | ENSMUSG00 | 17355 |
| NA | ENSMUSG00 | 231510 |
| NA | ENSMUSG00 | 71883 |
| Significant | ENSMUSG00 | 231507 |
| NA | ENSMUSG00 | 67870 |
| NA | ENSMUSG00 | 50926 |
| NA | ENSMUSG00 | 74596 |
| NA | ENSMUSG00 | 71914 |
| Significant | ENSMUSG00 | 12960 |
| NA | ENSMUSG00 | 22022 |
| NA | ENSMUSG00 | 54723 |
| NA | ENSMUSG00 | 70118 |
| NA | ENSMUSG00 | 72151 |
| NA | ENSMUSG00 | 59043 |
| NA | ENSMUSG00 | 13178 |
| NA | ENSMUSG00 | 57349 |
| Significant | ENSMUSG00 | 56744 |
| NA | ENSMUSG00 | 665563 |
| NA | ENSMUSG00 | 330122 |

| | | |
|---|---|---|
| NA | ENSMUSG00 | 14825 |
| NA | ENSMUSG00 | 12452 |
| NA | ENSMUSG00 | 209357 |
| NA | ENSMUSG00 | 209354 |
| NA | ENSMUSG00 | 67848 |
| NA | ENSMUSG00 | 56334 |
| NA | ENSMUSG00 | 75695 |
| NA | ENSMUSG00 | 13445 |
| NA | ENSMUSG00 | 68098 |
| NA | ENSMUSG00 | 80291 |
| NA | ENSMUSG00 | 76167 |
| NA | ENSMUSG00 | 53886 |
| NA | ENSMUSG00 | 65105 |
| NA | ENSMUSG00 | 23881 |
| Significant | ENSMUSG00 | 19679 |
| NA | ENSMUSG00 | 56041 |
| NA | ENSMUSG00 | 56325 |
| NA | ENSMUSG00 | 67111 |
| NA | ENSMUSG00 | 208628 |
| NA | ENSMUSG00 | 231452 |
| NA | ENSMUSG00 | 100561 |
| NA | ENSMUSG00 | 208606 |
| NA | ENSMUSG00 | 12492 |
| NA | ENSMUSG00 | 70650 |
| NA | ENSMUSG00 | 13852 |
| NA | ENSMUSG00 | 19384 |
| NA | ENSMUSG00 | 14467 |
| NA | ENSMUSG00 | 66593 |
| NA | ENSMUSG00 | 77573 |
| NA | ENSMUSG00 | 77045 |
| NA | ENSMUSG00 | 231769 |
| NA | ENSMUSG00 | 67151 |
| NA | ENSMUSG00 | 100678 |
| NA | ENSMUSG00 | 12466 |
| NA | ENSMUSG00 | 23912 |
| NA | ENSMUSG00 | 77462 |
| NA | ENSMUSG00 | 17165 |
| NA | ENSMUSG00 | 11669 |
| NA | ENSMUSG00 | 71985 |
| NA | ENSMUSG00 | 72399 |
| NA | ENSMUSG00 | 319604 |
| NA | ENSMUSG00 | 56433 |
| NA | ENSMUSG00 | 68948 |
| NA | ENSMUSG00 | 68080 |
| NA | ENSMUSG00 | 56378 |
| NA | ENSMUSG00 | 56317 |

| | | |
|---|---|---|
| NA | ENSMUSG00 | 11938 |
| NA | ENSMUSG00 | 18439 |
| NA | ENSMUSG00 | 12589 |
| Significant | ENSMUSG00 | 18438 |
| NA | ENSMUSG00 | 207565 |
| NA | ENSMUSG00 | 59008 |
| NA | ENSMUSG00 | 80751 |
| NA | ENSMUSG00 | 30841 |
| NA | ENSMUSG00 | 20602 |
| Significant | ENSMUSG00 | 208144 |
| NA | ENSMUSG00 | 78894 |
| NA | ENSMUSG00 | 11745 |
| NA | ENSMUSG00 | 94061 |
| NA | ENSMUSG00 | 19301 |
| NA | ENSMUSG00 | 72542 |
| NA | ENSMUSG00 | 71782 |
| NA | ENSMUSG00 | 269682 |
| NA | ENSMUSG00 | 69663 |
| NA | ENSMUSG00 | 75560 |
| NA | ENSMUSG00 | 56361 |
| NA | ENSMUSG00 | 22241 |
| NA | ENSMUSG00 | 19079 |
| NA | ENSMUSG00 | 12704 |
| NA | ENSMUSG00 | 77407 |
| NA | ENSMUSG00 | 50883 |
| NA | ENSMUSG00 | 75387 |
| NA | ENSMUSG00 | 19303 |
| NA | ENSMUSG00 | 64213 |
| NA | ENSMUSG00 | 69076 |
| NA | ENSMUSG00 | 384281 |
| NA | ENSMUSG00 | 108014 |
| NA | ENSMUSG00 | 11409 |
| NA | ENSMUSG00 | 68510 |
| NA | ENSMUSG00 | 74585 |
| NA | ENSMUSG00 | 66506 |
| NA | ENSMUSG00 | 21753 |
| NA | ENSMUSG00 | 21426 |
| NA | ENSMUSG00 | 17120 |
| NA | ENSMUSG00 | 68017 |
| NA | ENSMUSG00 | 72357 |
| Significant | ENSMUSG00 | 231834 |
| NA | ENSMUSG00 | 23962 |
| NA | ENSMUSG00 | 114142 |
| NA | ENSMUSG00 | 101118 |
| NA | ENSMUSG00 | 16848 |
| NA | ENSMUSG00 | 71900 |

| | | |
|---|---|---|
| NA | ENSMUSG00 | 77697 |
| NA | ENSMUSG00 | 117146 |
| NA | ENSMUSG00 | 74781 |
| NA | ENSMUSG00 | 11461 |
| Significant | ENSMUSG00 | 14086 |
| NA | ENSMUSG00 | 231866 |
| NA | ENSMUSG00 | 22256 |
| NA | ENSMUSG00 NA | |
| NA | ENSMUSG00 | 74111 |
| NA | ENSMUSG00 | 71772 |
| NA | ENSMUSG00 | 71990 |
| NA | ENSMUSG00 | 100764 |
| NA | ENSMUSG00 | 19415 |
| NA | ENSMUSG00 | 231872 |
| NA | ENSMUSG00 | 15467 |
| NA | ENSMUSG00 | 19988 |
| NA | ENSMUSG00 | 67397 |
| NA | ENSMUSG00 | 231874 |
| NA | ENSMUSG00 | 56443 |
| NA | ENSMUSG00 | 11867 |
| NA | ENSMUSG00 | 231887 |
| NA | ENSMUSG00 | 71799 |
| NA | ENSMUSG00 | 54188 |
| NA | ENSMUSG00 | 67235 |
| NA | ENSMUSG00 | 75725 |
| NA | ENSMUSG00 | 17992 |
| NA | ENSMUSG00 | 74132 |
| NA | ENSMUSG00 | 264064 |
| NA | ENSMUSG00 | 170772 |
| NA | ENSMUSG00 | 22217 |
| NA | ENSMUSG00 | 68895 |
| NA | ENSMUSG00 | 20018 |
| NA | ENSMUSG00 | 72587 |
| NA | ENSMUSG00 | 14254 |
| NA | ENSMUSG00 | 66537 |
| NA | ENSMUSG00 | 71706 |
| NA | ENSMUSG00 | 77521 |
| NA | ENSMUSG00 | 381695 |
| NA | ENSMUSG00 | 15505 |
| NA | ENSMUSG00 | 12843 |
| NA | ENSMUSG00 | 269831 |
| NA | ENSMUSG00 | 71777 |
| NA | ENSMUSG00 | 27999 |
| NA | ENSMUSG00 | 319974 |
| NA | ENSMUSG00 | 16885 |
| NA | ENSMUSG00 | 101185 |

| | | |
|---|---|---|
| NA | ENSMUSG00 | 12054 |
| NA | ENSMUSG00 | 73178 |
| NA | ENSMUSG00 | 26965 |
| NA | ENSMUSG00 | 14056 |
| NA | ENSMUSG00 | 109267 |
| NA | ENSMUSG00 | 68272 |
| NA | ENSMUSG00 | 71735 |
| NA | ENSMUSG00 | 13047 |
| NA | ENSMUSG00 | 74375 |
| NA | ENSMUSG00 | 13846 |
| NA | ENSMUSG00 | 14693 |
| NA | ENSMUSG00 | 57330 |
| NA | ENSMUSG00 | 74097 |
| NA | ENSMUSG00 | 18542 |
| NA | ENSMUSG00 | NA |
| NA | ENSMUSG00 | 231801 |
| NA | ENSMUSG00 | NA |
| NA | ENSMUSG00 | 69871 |
| NA | ENSMUSG00 | 231803 |
| NA | ENSMUSG00 | 74570 |
| NA | ENSMUSG00 | 17220 |
| NA | ENSMUSG00 | 29807 |
| NA | ENSMUSG00 | 27053 |
| Significant | ENSMUSG00 | 269823 |
| NA | ENSMUSG00 | 109624 |
| Significant | ENSMUSG00 | 14187 |
| NA | ENSMUSG00 | 20336 |
| NA | ENSMUSG00 | 12321 |
| NA | ENSMUSG00 | 27056 |
| NA | ENSMUSG00 | 74340 |
| NA | ENSMUSG00 | 76788 |
| NA | ENSMUSG00 | 58875 |
| Significant | ENSMUSG00 | 353172 |
| NA | ENSMUSG00 | 107569 |
| Significant | ENSMUSG00 | 27055 |
| NA | ENSMUSG00 | 72649 |
| NA | ENSMUSG00 | 78937 |
| NA | ENSMUSG00 | 83922 |
| NA | ENSMUSG00 | 67138 |
| NA | ENSMUSG00 | 26357 |
| NA | ENSMUSG00 | 73998 |
| Significant | ENSMUSG00 | 65963 |
| NA | ENSMUSG00 | 140488 |
| NA | ENSMUSG00 | 75593 |
| NA | ENSMUSG00 | 93695 |
| NA | ENSMUSG00 | 101214 |

| | | |
|---|---|---|
| NA | ENSMUSG00 | 109648 |
| NA | ENSMUSG00 | 54722 |
| NA | ENSMUSG00 | 71720 |
| NA | ENSMUSG00 | 192196 |
| NA | ENSMUSG00 | 78781 |
| NA | ENSMUSG00 | 21848 |
| NA | ENSMUSG00 | 12417 |
| NA | ENSMUSG00 | 19242 |
| NA | ENSMUSG00 | 14489 |
| NA | ENSMUSG00 | 22793 |
| NA | ENSMUSG00 | 12366 |
| NA | ENSMUSG00 | 76263 |
| NA | ENSMUSG00 | 13848 |
| NA | ENSMUSG00 | 56150 |
| NA | ENSMUSG00 | 381760 |
| NA | ENSMUSG00 | 70779 |
| Significant | ENSMUSG00 | 23845 |
| NA | ENSMUSG00 | 69923 |
| NA | ENSMUSG00 | 14548 |
| Significant | ENSMUSG00 | 54486 |
| NA | ENSMUSG00 | 13990 |
| NA | ENSMUSG00 | 54484 |
| NA | ENSMUSG00 | 19331 |
| NA | ENSMUSG00 | 72144 |
| Significant | ENSMUSG00 | 21391 |
| NA | ENSMUSG00 | 14583 |
| NA | ENSMUSG00 | 56748 |
| NA | ENSMUSG00 | 11746 |
| NA | ENSMUSG00 | 66881 |
| NA | ENSMUSG00 | 72102 |
| NA | ENSMUSG00 | 12468 |
| NA | ENSMUSG00 | 73327 |
| NA | ENSMUSG00 | 18139 |
| NA | ENSMUSG00 | 101358 |
| NA | ENSMUSG00 | 243574 |
| NA | ENSMUSG00 | 75659 |
| NA | ENSMUSG00 | 70020 |
| NA | ENSMUSG00 | 22377 |
| NA | ENSMUSG00 | 57377 |
| NA | ENSMUSG00 | 68499 |
| NA | ENSMUSG00 | 110958 |
| NA | ENSMUSG00 | 66979 |
| NA | ENSMUSG00 | 56284 |
| NA | ENSMUSG00 | 232201 |
| NA | ENSMUSG00 | 72103 |
| NA | ENSMUSG00 | 54368 |

| | | |
|---|---|---|
| NA | ENSMUSG00 | 69834 |
| NA | ENSMUSG00 | 57905 |
| NA | ENSMUSG00 | 12785 |
| NA | ENSMUSG00 | 54161 |
| NA | ENSMUSG00 | 232286 |
| NA | ENSMUSG00 | 232210 |
| NA | ENSMUSG00 | 22200 |
| NA | ENSMUSG00 | 103963 |
| NA | ENSMUSG00 | 232288 |
| NA | ENSMUSG00 | 108655 |
| NA | ENSMUSG00 | NA |
| NA | ENSMUSG00 | 56505 |
| NA | ENSMUSG00 | 53421 |
| NA | ENSMUSG00 | 80283 |
| NA | ENSMUSG00 | 18844 |
| NA | ENSMUSG00 | 66098 |
| NA | ENSMUSG00 | 66277 |
| NA | ENSMUSG00 | 107747 |
| NA | ENSMUSG00 | 71699 |
| Significant | ENSMUSG00 | 54563 |
| NA | ENSMUSG00 | 22591 |
| NA | ENSMUSG00 | 74122 |
| NA | ENSMUSG00 | 21366 |
| NA | ENSMUSG00 | 58911 |
| Significant | ENSMUSG00 | 16438 |
| NA | ENSMUSG00 | 20893 |
| NA | ENSMUSG00 | 192193 |
| NA | ENSMUSG00 | 67166 |
| NA | ENSMUSG00 | 24110 |
| NA | ENSMUSG00 | 14412 |
| NA | ENSMUSG00 | 14562 |
| NA | ENSMUSG00 | 30853 |
| NA | ENSMUSG00 | 69202 |
| Significant | ENSMUSG00 | 67784 |
| NA | ENSMUSG00 | 16768 |
| NA | ENSMUSG00 | 330401 |
| NA | ENSMUSG00 | 26894 |
| Significant | ENSMUSG00 | 53857 |
| NA | ENSMUSG00 | 213895 |
| NA | ENSMUSG00 | 56619 |
| NA | ENSMUSG00 | 17474 |
| NA | ENSMUSG00 | 72720 |
| NA | ENSMUSG00 | 26888 |
| Significant | ENSMUSG00 | 27007 |
| NA | ENSMUSG00 | 232408 |
| Significant | ENSMUSG00 | 12515 |

| | | |
|---|---|---|
| NA | ENSMUSG00 | 93694 |
| Significant | ENSMUSG00 | 56760 |
| NA | ENSMUSG00 | 57436 |
| Significant | ENSMUSG00 | 108078 |
| NA | ENSMUSG00 | 19365 |
| NA | ENSMUSG00 | 68465 |
| NA | ENSMUSG00 | 111173 |
| NA | ENSMUSG00 | 67200 |
| NA | ENSMUSG00 | 214899 |
| NA | ENSMUSG00 | 66441 |
| NA | ENSMUSG00 | 56449 |
| Significant | ENSMUSG00 | 14011 |
| NA | ENSMUSG00 | 66813 |
| NA | ENSMUSG00 | 16974 |
| NA | ENSMUSG00 | 70686 |
| NA | ENSMUSG00 | 67755 |
| NA | ENSMUSG00 | 14840 |
| NA | ENSMUSG00 | 74525 |
| NA | ENSMUSG00 | 13730 |
| NA | ENSMUSG00 | 54343 |
| Significant | ENSMUSG00 | 66857 |
| NA | ENSMUSG00 | 60321 |
| Significant | ENSMUSG00 | 69187 |
| NA | ENSMUSG00 | 11857 |
| NA | ENSMUSG00 | 19277 |
| NA | ENSMUSG00 | 20901 |
| NA | ENSMUSG00 | 232449 |
| NA | ENSMUSG00 | 109135 |
| NA | ENSMUSG00 | 11569 |
| NA | ENSMUSG00 | 19691 |
| NA | ENSMUSG00 | 66964 |
| NA | ENSMUSG00 | 16832 |
| NA | ENSMUSG00 | 20928 |
| NA | ENSMUSG00 | 58186 |
| NA | ENSMUSG00 | 79362 |
| NA | ENSMUSG00 | 259302 |
| NA | ENSMUSG00 | 71323 |
| NA | ENSMUSG00 | 16970 |
| NA | ENSMUSG00 | 14911 |
| NA | ENSMUSG00 | 16653 |
| Significant | ENSMUSG00 | 12035 |
| NA | ENSMUSG00 | 97287 |
| NA | ENSMUSG00 | 18294 |
| Significant | ENSMUSG00 | 52163 |
| NA | ENSMUSG00 | 75320 |
| NA | ENSMUSG00 | 101100 |

| | | |
|---|---|---|
| NA | ENSMUSG00 | 74741 |
| NA | ENSMUSG00 | 171095 |
| NA | ENSMUSG00 | 12764 |
| NA | ENSMUSG00 | 171508 |
| NA | ENSMUSG00 | 66087 |
| NA | ENSMUSG00 | 16439 |
| NA | ENSMUSG00 | 108098 |
| NA | ENSMUSG00 | 110379 |
| NA | ENSMUSG00 | 67015 |
| NA | ENSMUSG00 | 330450 |
| NA | ENSMUSG00 | 67456 |
| Significant | ENSMUSG00 | 232560 |
| NA | ENSMUSG00 | 320560 |
| NA | ENSMUSG00 | 74244 |
| NA | ENSMUSG00 | 232334 |
| NA | ENSMUSG00 | 68971 |
| NA | ENSMUSG00 | 67088 |
| NA | ENSMUSG00 | 212516 |
| NA | ENSMUSG00 | 17193 |
| NA | ENSMUSG00 | 81896 |
| NA | ENSMUSG00 | 67602 |
| NA | ENSMUSG00 | 28019 |
| NA | ENSMUSG00 | 66493 |
| NA | ENSMUSG00 | NA |
| NA | ENSMUSG00 | 17000 |
| NA | ENSMUSG00 | 21937 |
| NA | ENSMUSG00 | 12527 |
| NA | ENSMUSG00 | 101320 |
| NA | ENSMUSG00 | 19362 |
| NA | ENSMUSG00 | 28040 |
| NA | ENSMUSG00 | 109246 |
| NA | ENSMUSG00 | 14228 |
| NA | ENSMUSG00 | 93675 |
| NA | ENSMUSG00 | 97387 |
| NA | ENSMUSG00 | 109889 |
| NA | ENSMUSG00 | 67370 |
| NA | ENSMUSG00 | 22775 |
| NA | ENSMUSG00 | 232944 |
| NA | ENSMUSG00 | 12715 |
| NA | ENSMUSG00 | 13871 |
| Significant | ENSMUSG00 | 22323 |
| NA | ENSMUSG00 | 67369 |
| NA | ENSMUSG00 | 13400 |
| NA | ENSMUSG00 | 13401 |
| Significant | ENSMUSG00 | 21946 |
| NA | ENSMUSG00 | 56330 |

| | | |
|---|---|---|
| NA | ENSMUSG00 | 19777 |
| NA | ENSMUSG00 | 66161 |
| NA | ENSMUSG00 | NA |
| NA | ENSMUSG00 | 18726 |
| NA | ENSMUSG00 | 664968 |
| NA | ENSMUSG00 | 19943 |
| NA | ENSMUSG00 | 22185 |
| NA | ENSMUSG00 | 213011 |
| NA | ENSMUSG00 | 212569 |
| NA | ENSMUSG00 | 20430 |
| NA | ENSMUSG00 | 15204 |
| NA | ENSMUSG00 | 93790 |
| NA | ENSMUSG00 | 234353 |
| NA | ENSMUSG00 | 243958 |
| NA | ENSMUSG00 | 210105 |
| NA | ENSMUSG00 | 243983 |
| NA | ENSMUSG00 | 83382 |
| NA | ENSMUSG00 | 54678 |
| NA | ENSMUSG00 | 101831 |
| NA | ENSMUSG00 | 14453 |
| NA | ENSMUSG00 | 71974 |
| NA | ENSMUSG00 | 117589 |
| NA | ENSMUSG00 | 545975 |
| NA | ENSMUSG00 | 68981 |
| NA | ENSMUSG00 | 272396 |
| NA | ENSMUSG00 | 21872 |
| NA | ENSMUSG00 | 67973 |
| NA | ENSMUSG00 | 233315 |
| NA | ENSMUSG00 | 70461 |
| NA | ENSMUSG00 | 12144 |
| NA | ENSMUSG00 | 18550 |
| NA | ENSMUSG00 | 68695 |
| NA | ENSMUSG00 | 101869 |
| NA | ENSMUSG00 | 233405 |
| NA | ENSMUSG00 | 29875 |
| NA | ENSMUSG00 | 23991 |
| NA | ENSMUSG00 | 20352 |
| NA | ENSMUSG00 | 269951 |
| NA | ENSMUSG00 | 18631 |
| NA | ENSMUSG00 | 11819 |
| NA | ENSMUSG00 | 67009 |
| NA | ENSMUSG00 | 67867 |
| NA | ENSMUSG00 | 17258 |
| NA | ENSMUSG00 | 72433 |
| Significant | ENSMUSG00 | 13032 |
| NA | ENSMUSG00 | 22177 |

| | | |
|---|---|---|
| NA | ENSMUSG00 | 74206 |
| NA | ENSMUSG00 | 29861 |
| NA | ENSMUSG00 | 77254 |
| NA | ENSMUSG00 | 233046 |
| NA | ENSMUSG00 | 73833 |
| NA | ENSMUSG00 | 57296 |
| NA | ENSMUSG00 | 20190 |
| NA | ENSMUSG00 | 18036 |
| NA | ENSMUSG00 | 50760 |
| NA | ENSMUSG00 | 70584 |
| NA | ENSMUSG00 | 23996 |
| NA | ENSMUSG00 | 71163 |
| NA | ENSMUSG00 | 17304 |
| NA | ENSMUSG00 | 68048 |
| NA | ENSMUSG00 | 76375 |
| NA | ENSMUSG00 | 67994 |
| NA | ENSMUSG00 | 67308 |
| NA | ENSMUSG00 | 66365 |
| NA | ENSMUSG00 | 68472 |
| NA | ENSMUSG00 | 66271 |
| NA | ENSMUSG00 | 13626 |
| NA | ENSMUSG00 | 65098 |
| NA | ENSMUSG00 | 14085 |
| NA | ENSMUSG00 | 74041 |
| NA | ENSMUSG00 | 75985 |
| NA | ENSMUSG00 | 68197 |
| NA | ENSMUSG00 | 75430 |
| NA | ENSMUSG00 | 12850 |
| NA | ENSMUSG00 | 54208 |
| NA | ENSMUSG00 | 233789 |
| NA | ENSMUSG00 | 53322 |
| NA | ENSMUSG00 | 18704 |
| NA | ENSMUSG00 | 70572 |
| NA | ENSMUSG00 | 56372 |
| NA | ENSMUSG00 | 244209 |
| NA | ENSMUSG00 | 18576 |
| NA | ENSMUSG00 | 17907 |
| NA | ENSMUSG00 | 67375 |
| NA | ENSMUSG00 | 110033 |
| NA | ENSMUSG00 | 17188 |
| NA | ENSMUSG00 NA | |
| NA | ENSMUSG00 | 78388 |
| NA | ENSMUSG00 | 52858 |
| NA | ENSMUSG00 | 233877 |
| NA | ENSMUSG00 | 56018 |
| NA | ENSMUSG00 | 233875 |

| | | |
|---|---|---|
| NA | ENSMUSG00 | 207278 |
| NA | ENSMUSG00 | 11674 |
| NA | ENSMUSG00 | 56420 |
| NA | ENSMUSG00 | 68616 |
| NA | ENSMUSG00 | 19346 |
| NA | ENSMUSG00 | 52443 |
| NA | ENSMUSG00 | 12721 |
| NA | ENSMUSG00 | 69387 |
| NA | ENSMUSG00 | 20887 |
| NA | ENSMUSG00 | 75565 |
| NA | ENSMUSG00 | 56312 |
| NA | ENSMUSG00 | 72590 |
| Significant | ENSMUSG00 | 12752 |
| NA | ENSMUSG00 | 18020 |
| NA | ENSMUSG00 | 67164 |
| NA | ENSMUSG00 | 67967 |
| NA | ENSMUSG00 | 70314 |
| NA | ENSMUSG00 | 70974 |
| NA | ENSMUSG00 | 20399 |
| NA | ENSMUSG00 | 101488 |
| NA | ENSMUSG00 | 56347 |
| NA | ENSMUSG00 | 73658 |
| NA | ENSMUSG00 | 16797 |
| NA | ENSMUSG00 | 27050 |
| NA | ENSMUSG00 | 60504 |
| Significant | ENSMUSG00 | 67800 |
| NA | ENSMUSG00 | 16190 |
| NA | ENSMUSG00 | 67711 |
| NA | ENSMUSG00 | 26440 |
| NA | ENSMUSG00 | 77035 |
| NA | ENSMUSG00 | 72981 |
| NA | ENSMUSG00 | 70349 |
| NA | ENSMUSG00 | 67420 |
| NA | ENSMUSG00 | 66190 |
| NA | ENSMUSG00 | 17921 |
| NA | ENSMUSG00 | 30949 |
| NA | ENSMUSG00 | 70497 |
| NA | ENSMUSG00 | 68897 |
| NA | ENSMUSG00 | 233836 |
| NA | ENSMUSG00 | 57342 |
| NA | ENSMUSG00 | 18479 |
| NA | ENSMUSG00 | 19647 |
| NA | ENSMUSG00 | 233913 |
| Significant | ENSMUSG00 | 16409 |
| Significant | ENSMUSG00 | 114332 |
| Significant | ENSMUSG00 | 67150 |

| | | |
|---|---|---|
| NA | ENSMUSG00 | 11535 |
| NA | ENSMUSG00 | 66824 |
| NA | ENSMUSG00 | 233908 |
| NA | ENSMUSG00 | 21677 |
| Significant | ENSMUSG00 | 12493 |
| NA | ENSMUSG00 | 67773 |
| NA | ENSMUSG00 | 12041 |
| NA | ENSMUSG00 | 20909 |
| NA | ENSMUSG00 | 233902 |
| NA | ENSMUSG00 | 12055 |
| NA | ENSMUSG00 | 68961 |
| NA | ENSMUSG00 | 233900 |
| NA | ENSMUSG00 | 233895 |
| NA | ENSMUSG00 | 78921 |
| NA | ENSMUSG00 | 18220 |
| NA | ENSMUSG00 | 12036 |
| NA | ENSMUSG00 | 16408 |
| Significant | ENSMUSG00 | 211548 |
| NA | ENSMUSG00 | 27414 |
| NA | ENSMUSG00 | 66508 |
| NA | ENSMUSG00 | 67865 |
| NA | ENSMUSG00 | 21843 |
| NA | ENSMUSG00 | 29810 |
| NA | ENSMUSG00 | 11907 |
| NA | ENSMUSG00 | 57752 |
| NA | ENSMUSG00 | 66885 |
| NA | ENSMUSG00 | 18817 |
| NA | ENSMUSG00 | 59288 |
| NA | ENSMUSG00 | 70316 |
| NA | ENSMUSG00 | 28018 |
| NA | ENSMUSG00 | 67417 |
| NA | ENSMUSG00 | 74105 |
| NA | ENSMUSG00 | 59052 |
| NA | ENSMUSG00 | 12585 |
| NA | ENSMUSG00 | 27397 |
| NA | ENSMUSG00 | 26939 |
| NA | ENSMUSG00 | 76932 |
| NA | ENSMUSG00 | 67003 |
| NA | ENSMUSG00 | 101867 |
| NA | ENSMUSG00 | 16202 |
| Significant | ENSMUSG00 | 12751 |
| NA | ENSMUSG00 | 233806 |
| NA | ENSMUSG00 | 20128 |
| NA | ENSMUSG00 | 73919 |
| NA | ENSMUSG00 | 434234 |
| NA | ENSMUSG00 | 71151 |

| | | |
|---|---|---|
| Significant | ENSMUSG00 | 77590 |
| NA | ENSMUSG00 | 18242 |
| NA | ENSMUSG00 | 233802 |
| NA | ENSMUSG00 | 76429 |
| NA | ENSMUSG00 | 77938 |
| NA | ENSMUSG00 | 72096 |
| NA | ENSMUSG00 | 109359 |
| NA | ENSMUSG00 | 360216 |
| NA | ENSMUSG00 | 13017 |
| NA | ENSMUSG00 | 20133 |
| NA | ENSMUSG00 | 22276 |
| NA | ENSMUSG00 | 66356 |
| NA | ENSMUSG00 | 71517 |
| NA | ENSMUSG00 | 66165 |
| Significant | ENSMUSG00 | 101437 |
| NA | ENSMUSG00 | 20866 |
| NA | ENSMUSG00 | 233575 |
| NA | ENSMUSG00 | 17345 |
| Significant | ENSMUSG00 | 70495 |
| NA | ENSMUSG00 | 22284 |
| NA | ENSMUSG00 | 12361 |
| NA | ENSMUSG00 | 20947 |
| NA | ENSMUSG00 | 22390 |
| NA | ENSMUSG00 | 56786 |
| NA | ENSMUSG00 | 57373 |
| NA | ENSMUSG00 | 76954 |
| NA | ENSMUSG00 | 66085 |
| NA | ENSMUSG00 | 104130 |
| NA | ENSMUSG00 | 236732 |
| NA | ENSMUSG00 | 18555 |
| NA | ENSMUSG00 | 236733 |
| NA | ENSMUSG00 | 30926 |
| NA | ENSMUSG00 | 72284 |
| NA | ENSMUSG00 | 14082 |
| Significant | ENSMUSG00 | 13043 |
| NA | ENSMUSG00 | 78914 |
| NA | ENSMUSG00 | 75974 |
| NA | ENSMUSG00 | 72584 |
| NA | ENSMUSG00 | 21953 |
| NA | ENSMUSG00 | 74131 |
| NA | ENSMUSG00 | 56501 |
| NA | ENSMUSG00 | 19337 |
| NA | ENSMUSG00 | 20523 |
| NA | ENSMUSG00 | 209003 |
| NA | ENSMUSG00 | 209224 |
| Significant | ENSMUSG00 | 70415 |

| | | |
|---|---|---|
| NA | ENSMUSG00 | 14735 |
| NA | ENSMUSG00 | 70691 |
| Significant | ENSMUSG00 | 331004 |
| NA | ENSMUSG00 | 73341 |
| NA | ENSMUSG00 | 19655 |
| NA | ENSMUSG00 | 14168 |
| NA | ENSMUSG00 | 14071 |
| NA | ENSMUSG00 | 54652 |
| NA | ENSMUSG00 | 54638 |
| NA | ENSMUSG00 | 20977 |
| NA | ENSMUSG00 | 54630 |
| NA | ENSMUSG00 | 18824 |
| NA | ENSMUSG00 | 209416 |
| NA | ENSMUSG00 | 54637 |
| NA | ENSMUSG00 | 54648 |
| NA | ENSMUSG00 | 54645 |
| NA | ENSMUSG00 | 54644 |
| NA | ENSMUSG00 | 18715 |
| NA | ENSMUSG00 | 22232 |
| NA | ENSMUSG00 | 54633 |
| NA | ENSMUSG00 | 21855 |
| NA | ENSMUSG00 | 15185 |
| NA | ENSMUSG00 | 14460 |
| NA | ENSMUSG00 | 22376 |
| NA | ENSMUSG00 | 73447 |
| NA | ENSMUSG00 | NA |
| NA | ENSMUSG00 | 13595 |
| NA | ENSMUSG00 | 54632 |
| NA | ENSMUSG00 | 19893 |
| NA | ENSMUSG00 | 67117 |
| NA | ENSMUSG00 | 22327 |
| NA | ENSMUSG00 | 67391 |
| NA | ENSMUSG00 | 17763 |
| NA | ENSMUSG00 | 210766 |
| NA | ENSMUSG00 | 17698 |
| NA | ENSMUSG00 | 76206 |
| Significant | ENSMUSG00 | 94190 |
| Significant | ENSMUSG00 | 236920 |
| NA | ENSMUSG00 | 13641 |
| NA | ENSMUSG00 | 245532 |
| NA | ENSMUSG00 | 18518 |
| NA | ENSMUSG00 | 272790 |
| NA | ENSMUSG00 | 67683 |
| NA | ENSMUSG00 | 107528 |
| NA | ENSMUSG00 | 22589 |
| NA | ENSMUSG00 | 66142 |

| | | |
|---|---|---|
| NA | ENSMUSG00 | 67075 |
| NA | ENSMUSG00 | 16431 |
| NA | ENSMUSG00 | 67028 |
| NA | ENSMUSG00 | 50887 |
| NA | ENSMUSG00 | 56726 |
| NA | ENSMUSG00 | 27359 |
| NA | ENSMUSG00 | 108062 |
| NA | ENSMUSG00 | 102920 |
| NA | ENSMUSG00 | 12229 |
| NA | ENSMUSG00 | 11605 |
| NA | ENSMUSG00 | 12830 |
| NA | ENSMUSG00 | 50790 |
| NA | ENSMUSG00 | 382253 |
| NA | ENSMUSG00 | 110094 |
| NA | ENSMUSG00 | 11989 |
| NA | ENSMUSG00 | 18597 |
| NA | ENSMUSG00 | 270672 |
| NA | ENSMUSG00 | 16186 |
| NA | ENSMUSG00 | 110651 |
| NA | ENSMUSG00 | 56364 |
| NA | ENSMUSG00 | 53610 |
| NA | ENSMUSG00 | 270627 |
| NA | ENSMUSG00 | 20102 |
| NA | ENSMUSG00 | 192176 |
| NA | ENSMUSG00 | 11306 |
| NA | ENSMUSG00 | 17772 |
| NA | ENSMUSG00 | 14758 |
| NA | ENSMUSG00 | 26370 |
| NA | ENSMUSG00 | 18194 |
| NA | ENSMUSG00 | 15159 |
| NA | ENSMUSG00 | 245688 |
| Significant | ENSMUSG00 | 11856 |
| NA | ENSMUSG00 | 67043 |
| NA | ENSMUSG00 | 17692 |
| NA | ENSMUSG00 | 55936 |
| NA | ENSMUSG00 | 72891 |
| NA | ENSMUSG00 | 27081 |
| NA | ENSMUSG00 | 108012 |
| NA | ENSMUSG00 | 22184 |
| NA | ENSMUSG00 | 73738 |
| NA | ENSMUSG00 | 56078 |
| NA | ENSMUSG00 | 12111 |
| NA | ENSMUSG00 | 11666 |
| NA | ENSMUSG00 | 69656 |
| NA | ENSMUSG00 | 18700 |
| NA | ENSMUSG00 | 75590 |

| | | |
|---|---|---|
| NA | ENSMUSG00 | 140571 |
| NA | ENSMUSG00 | 15161 |
| Significant | ENSMUSG00 | 19703 |
| NA | ENSMUSG00 | 56292 |
| NA | ENSMUSG00 | 171207 |
| NA | ENSMUSG00 | 16179 |
| NA | ENSMUSG00 | 17257 |
| NA | ENSMUSG00 | 18846 |
| NA | ENSMUSG00 | 66294 |
| Significant | ENSMUSG00 | 14381 |
| Significant | ENSMUSG00 | 17524 |
| NA | ENSMUSG00 | 245474 |
| NA | ENSMUSG00 | 56397 |
| NA | ENSMUSG00 | 53380 |
| NA | ENSMUSG00 | 14605 |
| NA | ENSMUSG00 | 19139 |
| NA | ENSMUSG00 | 237073 |
| NA | ENSMUSG00 | 75746 |
| Significant | ENSMUSG00 | 66889 |
| NA | ENSMUSG00 | 50770 |
| Significant | ENSMUSG00 | 14058 |
| NA | ENSMUSG00 | 66901 |
| NA | ENSMUSG00 | 99375 |
| NA | ENSMUSG00 | 16783 |
| Significant | ENSMUSG00 | 14456 |
| NA | ENSMUSG00 | 19414 |
| Significant | ENSMUSG00 | 66423 |
| NA | ENSMUSG00 | 11601 |
| NA | ENSMUSG00 | 52123 |
| NA | ENSMUSG00 | 23954 |
| NA | ENSMUSG00 | 70160 |
| NA | ENSMUSG00 | 56229 |
| NA | ENSMUSG00 | 18408 |
| NA | ENSMUSG00 | 244373 |
| NA | ENSMUSG00 | 114863 |
| NA | ENSMUSG00 | 78560 |
| NA | ENSMUSG00 | 66653 |
| NA | ENSMUSG00 | 75767 |
| NA | ENSMUSG00 | 13685 |
| NA | ENSMUSG00 | 243897 |
| Significant | ENSMUSG00 | 170786 |
| NA | ENSMUSG00 | 24099 |
| NA | ENSMUSG00 | 12826 |
| NA | ENSMUSG00 | 12827 |
| NA | ENSMUSG00 | 19332 |
| NA | ENSMUSG00 | 69225 |

| | | |
|---|---|---|
| NA | ENSMUSG00 | 320139 |
| NA | ENSMUSG00 | 102334 |
| NA | ENSMUSG00 | 54126 |
| NA | ENSMUSG00 | 68192 |
| NA | ENSMUSG00 | 22428 |
| NA | ENSMUSG00 | 11593 |
| NA | ENSMUSG00 | 50768 |
| NA | ENSMUSG00 | 67276 |
| NA | ENSMUSG00 | 21951 |
| NA | ENSMUSG00 | 319520 |
| NA | ENSMUSG00 | 67887 |
| NA | ENSMUSG00 | 57312 |
| NA | ENSMUSG00 | 102032 |
| NA | ENSMUSG00 | 18970 |
| NA | ENSMUSG00 | 16150 |
| NA | ENSMUSG00 | 18791 |
| NA | ENSMUSG00 | 64933 |
| NA | ENSMUSG00 | 244349 |
| NA | ENSMUSG00 | 102247 |
| NA | ENSMUSG00 | 109145 |
| NA | ENSMUSG00 | 20377 |
| NA | ENSMUSG00 | 11497 |
| NA | ENSMUSG00 | 11502 |
| NA | ENSMUSG00 | 69742 |
| NA | ENSMUSG00 | 83436 |
| Significant | ENSMUSG00 | 320685 |
| NA | ENSMUSG00 | 52357 |
| NA | ENSMUSG00 | 14182 |
| NA | ENSMUSG00 | 192174 |
| NA | ENSMUSG00 | 71910 |
| NA | ENSMUSG00 | 20845 |
| NA | ENSMUSG00 | 23808 |
| NA | ENSMUSG00 | 234138 |
| NA | ENSMUSG00 | 67920 |
| NA | ENSMUSG00 | 22427 |
| Significant | ENSMUSG00 | 14782 |
| NA | ENSMUSG00 | 68153 |
| NA | ENSMUSG00 | 19663 |
| NA | ENSMUSG00 | 14300 |
| NA | ENSMUSG00 | 11886 |
| NA | ENSMUSG00 | 18536 |
| NA | ENSMUSG00 | 234199 |
| NA | ENSMUSG00 | 68797 |
| NA | ENSMUSG00 | 11988 |
| NA | ENSMUSG00 | 52348 |
| NA | ENSMUSG00 | 18983 |

| | | |
|---|---|---|
| Significant | ENSMUSG00 | 66234 |
| NA | ENSMUSG00 | 77113 |
| NA | ENSMUSG00 | 108150 |
| NA | ENSMUSG00 | 60406 |
| NA | ENSMUSG00 | 15446 |
| NA | ENSMUSG00 | 234463 |
| NA | ENSMUSG00 | 20467 |
| NA | ENSMUSG00 | 16363 |
| NA | ENSMUSG00 | 12367 |
| NA | ENSMUSG00 | 71876 |
| NA | ENSMUSG00 | 66756 |
| NA | ENSMUSG00 | 11739 |
| NA | ENSMUSG00 | 192169 |
| NA | ENSMUSG00 | 53318 |
| NA | ENSMUSG00 | 67620 |
| NA | ENSMUSG00 | 142980 |
| NA | ENSMUSG00 | 234309 |
| NA | ENSMUSG00 | 59009 |
| NA | ENSMUSG00 | 18004 |
| NA | ENSMUSG00 | 80750 |
| Significant | ENSMUSG00 | 234549 |
| NA | ENSMUSG00 | 11513 |
| NA | ENSMUSG00 | 26992 |
| NA | ENSMUSG00 | 93960 |
| NA | ENSMUSG00 | 71607 |
| NA | ENSMUSG00 | 19651 |
| NA | ENSMUSG00 | 14339 |
| NA | ENSMUSG00 | 13666 |
| NA | ENSMUSG00 | 78833 |
| NA | ENSMUSG00 | 66083 |
| NA | ENSMUSG00 | 14719 |
| NA | ENSMUSG00 | 12552 |
| NA | ENSMUSG00 | 17125 |
| NA | ENSMUSG00 | 78651 |
| NA | ENSMUSG00 | 76775 |
| NA | ENSMUSG00 | 212999 |
| NA | ENSMUSG00 | 65114 |
| NA | ENSMUSG00 | 56452 |
| Significant | ENSMUSG00 | 108682 |
| NA | ENSMUSG00 | 56445 |
| NA | ENSMUSG00 | 71927 |
| NA | ENSMUSG00 | 19724 |
| NA | ENSMUSG00 | 106529 |
| NA | ENSMUSG00 | 71310 |
| NA | ENSMUSG00 | 30932 |
| NA | ENSMUSG00 | 16168 |

| | | |
|---|---|---|
| Significant | ENSMUSG00 | 14388 |
| NA | ENSMUSG00 | 93762 |
| Significant | ENSMUSG00 | 15439 |
| NA | ENSMUSG00 | 234723 |
| NA | ENSMUSG00 | 75871 |
| NA | ENSMUSG00 | 71955 |
| NA | ENSMUSG00 | 56749 |
| NA | ENSMUSG00 | 11765 |
| NA | ENSMUSG00 | 244650 |
| Significant | ENSMUSG00 | 17390 |
| NA | ENSMUSG00 | 20444 |
| NA | ENSMUSG00 | 23802 |
| NA | ENSMUSG00 | 102339 |
| NA | ENSMUSG00 | 68219 |
| NA | ENSMUSG00 | 67378 |
| NA | ENSMUSG00 | 72155 |
| NA | ENSMUSG00 | 75796 |
| NA | ENSMUSG00 | 17750 |
| NA | ENSMUSG00 | 17748 |
| NA | ENSMUSG00 | 67528 |
| NA | ENSMUSG00 | 64209 |
| NA | ENSMUSG00 | 102122 |
| NA | ENSMUSG00 | 107566 |
| NA | ENSMUSG00 | 20312 |
| NA | ENSMUSG00 | 109006 |
| NA | ENSMUSG00 | 67914 |
| NA | ENSMUSG00 | 20021 |
| Significant | ENSMUSG00 | 14766 |
| NA | ENSMUSG00 | 74187 |
| NA | ENSMUSG00 | 16582 |
| Significant | ENSMUSG00 | 74166 |
| NA | ENSMUSG00 | 101985 |
| NA | ENSMUSG00 | 14894 |
| NA | ENSMUSG00 | 326618 |
| NA | ENSMUSG00 | 72297 |
| NA | ENSMUSG00 | 16453 |
| NA | ENSMUSG00 | 66171 |
| NA | ENSMUSG00 | 26457 |
| NA | ENSMUSG00 | 67443 |
| NA | ENSMUSG00 | 73711 |
| NA | ENSMUSG00 | 234814 |
| NA | ENSMUSG00 | 12857 |
| NA | ENSMUSG00 | 18117 |
| NA | ENSMUSG00 | 68251 |
| NA | ENSMUSG00 | 272551 |
| NA | ENSMUSG00 | 382034 |

| | | |
|---|---|---|
| NA | ENSMUSG00 | 102193 |
| NA | ENSMUSG00 | 234797 |
| NA | ENSMUSG00 | 78892 |
| NA | ENSMUSG00 | 22224 |
| NA | ENSMUSG00 | 72042 |
| NA | ENSMUSG00 | 234796 |
| NA | ENSMUSG00 | 21341 |
| NA | ENSMUSG00 | 546071 |
| NA | ENSMUSG00 | 18709 |
| NA | ENSMUSG00 | 56453 |
| NA | ENSMUSG00 | 68196 |
| NA | ENSMUSG00 | 19339 |
| NA | ENSMUSG00 | 68533 |
| NA | ENSMUSG00 | 69528 |
| NA | ENSMUSG00 | 50783 |
| Significant | ENSMUSG00 | 66566 |
| NA | ENSMUSG00 | 74549 |
| NA | ENSMUSG00 | 170759 |
| NA | ENSMUSG00 | 70885 |
| NA | ENSMUSG00 | 13191 |
| NA | ENSMUSG00 | 12562 |
| Significant | ENSMUSG00 | 68119 |
| NA | ENSMUSG00 | 234664 |
| NA | ENSMUSG00 | 68523 |
| NA | ENSMUSG00 | 56437 |
| NA | ENSMUSG00 | 12354 |
| NA | ENSMUSG00 | 12400 |
| NA | ENSMUSG00 | 71609 |
| NA | ENSMUSG00 | 234678 |
| NA | ENSMUSG00 | 19171 |
| NA | ENSMUSG00 | 66369 |
| NA | ENSMUSG00 | 18021 |
| NA | ENSMUSG00 | 192654 |
| NA | ENSMUSG00 | 330836 |
| NA | ENSMUSG00 | 58994 |
| NA | ENSMUSG00 | 22751 |
| NA | ENSMUSG00 | 116733 |
| NA | ENSMUSG00 | 97484 |
| NA | ENSMUSG00 | 66164 |
| NA | ENSMUSG00 | 77116 |
| NA | ENSMUSG00 | 21750 |
| NA | ENSMUSG00 | 74360 |
| NA | ENSMUSG00 | 66427 |
| Significant | ENSMUSG00 | 270118 |
| NA | ENSMUSG00 | 17535 |
| NA | ENSMUSG00 | 66894 |

| | | |
|---|---|---|
| NA | ENSMUSG00 | 56503 |
| NA | ENSMUSG00 | 55991 |
| NA | ENSMUSG00 | 234959 |
| NA | ENSMUSG00 | 70984 |
| NA | ENSMUSG00 | 75316 |
| NA | ENSMUSG00 | 85305 |
| NA | ENSMUSG00 | 30947 |
| NA | ENSMUSG00 | 93739 |
| NA | ENSMUSG00 | 234740 |
| NA | ENSMUSG00 | 66817 |
| NA | ENSMUSG00 | 23837 |
| NA | ENSMUSG00 | 12927 |
| NA | ENSMUSG00 | 319481 |
| NA | ENSMUSG00 | 234734 |
| NA | ENSMUSG00 | 114896 |
| NA | ENSMUSG00 | 66948 |
| NA | ENSMUSG00 | 73420 |
| NA | ENSMUSG00 | 56199 |
| NA | ENSMUSG00 | 382038 |
| NA | ENSMUSG00 | 76332 |
| NA | ENSMUSG00 | 68865 |
| NA | ENSMUSG00 | 69551 |
| NA | ENSMUSG00 | 66523 |
| NA | ENSMUSG00 | 14712 |
| NA | ENSMUSG00 | 244666 |
| NA | ENSMUSG00 | 112405 |
| NA | ENSMUSG00 | 69091 |
| NA | ENSMUSG00 | 83964 |
| NA | ENSMUSG00 | 102607 |
| NA | ENSMUSG00 | 19143 |
| NA | ENSMUSG00 | 11804 |
| NA | ENSMUSG00 | 11796 |
| NA | ENSMUSG00 | 76863 |
| NA | ENSMUSG00 | 75747 |
| NA | ENSMUSG00 | 53376 |
| NA | ENSMUSG00 | 102644 |
| NA | ENSMUSG00 | 235293 |
| NA | ENSMUSG00 | 72828 |
| NA | ENSMUSG00 | 71566 |
| NA | ENSMUSG00 | 104444 |
| NA | ENSMUSG00 | 75717 |
| NA | ENSMUSG00 | 23871 |
| NA | ENSMUSG00 | 20443 |
| NA | ENSMUSG00 | 69305 |
| NA | ENSMUSG00 | 117149 |
| NA | ENSMUSG00 | 67398 |

| | | |
|---|---|---|
| NA | ENSMUSG00 | 71989 |
| Significant | ENSMUSG00 | 76192 |
| NA | ENSMUSG00 | 110446 |
| NA | ENSMUSG00 | 19684 |
| NA | ENSMUSG00 | 14148 |
| NA | ENSMUSG00 | 73699 |
| NA | ENSMUSG00 | 102580 |
| NA | ENSMUSG00 | 12955 |
| NA | ENSMUSG00 | 330938 |
| Significant | ENSMUSG00 | 170752 |
| NA | ENSMUSG00 | 19286 |
| Significant | ENSMUSG00 | 54725 |
| NA | ENSMUSG00 | 215051 |
| NA | ENSMUSG00 | 22687 |
| NA | ENSMUSG00 | 11806 |
| NA | ENSMUSG00 | 21345 |
| NA | ENSMUSG00 | 23821 |
| NA | ENSMUSG00 | 16154 |
| NA | ENSMUSG00 | 213827 |
| NA | ENSMUSG00 | 13209 |
| NA | ENSMUSG00 | 30959 |
| NA | ENSMUSG00 | 67049 |
| NA | ENSMUSG00 | 270151 |
| NA | ENSMUSG00 | 60409 |
| NA | ENSMUSG00 | 12649 |
| NA | ENSMUSG00 | 14385 |
| NA | ENSMUSG00 | 12282 |
| NA | ENSMUSG00 | 16430 |
| NA | ENSMUSG00 | 102423 |
| NA | ENSMUSG00 | 71764 |
| NA | ENSMUSG00 | 66279 |
| Significant | ENSMUSG00 | 56857 |
| NA | ENSMUSG00 | 13478 |
| NA | ENSMUSG00 | 74144 |
| NA | ENSMUSG00 | 15288 |
| NA | ENSMUSG00 | 71732 |
| NA | ENSMUSG00 | 84004 |
| NA | ENSMUSG00 | 23988 |
| NA | ENSMUSG00 | 54721 |
| NA | ENSMUSG00 | 18577 |
| NA | ENSMUSG00 | 16201 |
| NA | ENSMUSG00 | 17083 |
| NA | ENSMUSG00 | 74766 |
| NA | ENSMUSG00 | 70082 |
| NA | ENSMUSG00 | 59035 |
| NA | ENSMUSG00 | 50876 |

| | | |
|---|---|---|
| NA | ENSMUSG00 | 20586 |
| NA | ENSMUSG00 | 14697 |
| NA | ENSMUSG00 | 16835 |
| NA | ENSMUSG00 | 235041 |
| NA | ENSMUSG00 | 319899 |
| NA | ENSMUSG00 | 28015 |
| Significant | ENSMUSG00 | 11891 |
| NA | ENSMUSG00 | 64008 |
| NA | ENSMUSG00 | 66660 |
| NA | ENSMUSG00 | 225215 |
| NA | ENSMUSG00 | 17999 |
| NA | ENSMUSG00 | 93836 |
| NA | ENSMUSG00 | 12442 |
| NA | ENSMUSG00 | 71602 |
| NA | ENSMUSG00 | 17427 |
| NA | ENSMUSG00 | 21406 |
| Significant | ENSMUSG00 | 12306 |
| NA | ENSMUSG00 | 68178 |
| NA | ENSMUSG00 | 93697 |
| NA | ENSMUSG00 | 19883 |
| NA | ENSMUSG00 | 55934 |
| NA | ENSMUSG00 | 14155 |
| NA | ENSMUSG00 | 76524 |
| NA | ENSMUSG00 | 75600 |
| NA | ENSMUSG00 | 11737 |
| NA | ENSMUSG00 | 65099 |
| NA | ENSMUSG00 | 93683 |
| NA | ENSMUSG00 | 83946 |
| NA | ENSMUSG00 | 71819 |
| NA | ENSMUSG00 | 75782 |
| NA | ENSMUSG00 | 212531 |
| NA | ENSMUSG00 | 12040 |
| NA | ENSMUSG00 | 26951 |
| NA | ENSMUSG00 | 212943 |
| NA | ENSMUSG00 | 235323 |
| NA | ENSMUSG00 | 67834 |
| NA | ENSMUSG00 | 21887 |
| NA | ENSMUSG00 | 58233 |
| NA | ENSMUSG00 | 102462 |
| NA | ENSMUSG00 | 56294 |
| NA | ENSMUSG00 | 12903 |
| NA | ENSMUSG00 | 64602 |
| NA | ENSMUSG00 | 18746 |
| NA | ENSMUSG00 | 73744 |
| NA | ENSMUSG00 | 72774 |
| NA | ENSMUSG00 | 66199 |

| | | |
|---|---|---|
| NA | ENSMUSG00 | 74211 |
| NA | ENSMUSG00 | 26441 |
| NA | ENSMUSG00 | 78323 |
| NA | ENSMUSG00 | 110119 |
| NA | ENSMUSG00 | 109161 |
| NA | ENSMUSG00 | 71742 |
| NA | ENSMUSG00 | 71999 |
| NA | ENSMUSG00 | 83961 |
| NA | ENSMUSG00 | 12988 |
| NA | ENSMUSG00 | 110842 |
| NA | ENSMUSG00 | 102414 |
| NA | ENSMUSG00 | 26611 |
| NA | ENSMUSG00 | 19200 |
| NA | ENSMUSG00 | 56434 |
| NA | ENSMUSG00 | 69981 |
| NA | ENSMUSG00 | 66867 |
| NA | ENSMUSG00 | 12866 |
| NA | ENSMUSG00 | 69106 |
| NA | ENSMUSG00 | 20320 |
| NA | ENSMUSG00 | 18007 |
| NA | ENSMUSG00 | 68291 |
| NA | ENSMUSG00 | 214763 |
| NA | ENSMUSG00 | 14860 |
| NA | ENSMUSG00 | 68801 |
| NA | ENSMUSG00 | 14629 |
| NA | ENSMUSG00 | 214345 |
| NA | ENSMUSG00 | 66111 |
| NA | ENSMUSG00 | 13036 |
| NA | ENSMUSG00 | 22003 |
| NA | ENSMUSG00 | 22038 |
| NA | ENSMUSG00 | 80907 |
| NA | ENSMUSG00 | 76459 |
| NA | ENSMUSG00 | 26432 |
| NA | ENSMUSG00 | 208117 |
| NA | ENSMUSG00 | 235441 |
| NA | ENSMUSG00 | 13143 |
| NA | ENSMUSG00 | 68250 |
| NA | ENSMUSG00 | 56440 |
| NA | ENSMUSG00 | 19035 |
| NA | ENSMUSG00 | 214897 |
| NA | ENSMUSG00 | 56404 |
| NA | ENSMUSG00 | 71973 |
| NA | ENSMUSG00 | 27965 |
| Significant | ENSMUSG00 | 214424 |
| NA | ENSMUSG00 | 74388 |
| NA | ENSMUSG00 | 213550 |

| | | |
|---|---|---|
| NA | ENSMUSG00 | 66131 |
| NA | ENSMUSG00 | 330959 |
| NA | ENSMUSG00 | 67891 |
| NA | ENSMUSG00 | 68014 |
| NA | ENSMUSG00 | 17127 |
| NA | ENSMUSG00 | 69478 |
| NA | ENSMUSG00 | 56469 |
| NA | ENSMUSG00 | 67958 |
| NA | ENSMUSG00 | 245000 |
| NA | ENSMUSG00 | 24127 |
| NA | ENSMUSG00 | 211586 |
| NA | ENSMUSG00 | 11933 |
| NA | ENSMUSG00 | 114713 |
| NA | ENSMUSG00 | 70348 |
| NA | ENSMUSG00 | 69519 |
| NA | ENSMUSG00 | 17436 |
| Significant | ENSMUSG00 | 23959 |
| NA | ENSMUSG00 | 244962 |
| NA | ENSMUSG00 | 56403 |
| NA | ENSMUSG00 | 71640 |
| NA | ENSMUSG00 | 56693 |
| NA | ENSMUSG00 | 67213 |
| NA | ENSMUSG00 | 235661 |
| NA | ENSMUSG00 | 102545 |
| NA | ENSMUSG00 | 68292 |
| NA | ENSMUSG00 | 21813 |
| NA | ENSMUSG00 | 192287 |
| NA | ENSMUSG00 | 74080 |
| NA | ENSMUSG00 | 50797 |
| NA | ENSMUSG00 | 64655 |
| NA | ENSMUSG00 | 74769 |
| NA | ENSMUSG00 | 23873 |
| NA | ENSMUSG00 | 74125 |
| NA | ENSMUSG00 | 83703 |
| NA | ENSMUSG00 | 17973 |
| NA | ENSMUSG00 | 12530 |
| NA | ENSMUSG00 | 54369 |
| NA | ENSMUSG00 | 17758 |
| NA | ENSMUSG00 | 72831 |
| NA | ENSMUSG00 | 20588 |
| NA | ENSMUSG00 | 235623 |
| NA | ENSMUSG00 | 19225 |
| NA | ENSMUSG00 | 16578 |
| NA | ENSMUSG00 | 67169 |
| NA | ENSMUSG00 | 71268 |
| NA | ENSMUSG00 | 17350 |

| | | |
|---|---|---|
| NA | ENSMUSG00 | 211770 |
| NA | ENSMUSG00 | 18571 |
| NA | ENSMUSG00 | 72179 |
| NA | ENSMUSG00 | 17874 |
| NA | ENSMUSG00 | 67561 |
| NA | ENSMUSG00 | 74498 |
| NA | ENSMUSG00 | 215418 |
| NA | ENSMUSG00 | 16785 |
| NA | ENSMUSG00 | 208638 |
| NA | ENSMUSG00 | 18087 |
| NA | ENSMUSG00 | 26901 |
| NA | ENSMUSG00 | 66904 |
| NA | ENSMUSG00 | 56332 |
| NA | ENSMUSG00 | 28135 |
| NA | ENSMUSG00 | 67095 |
| NA | ENSMUSG00 | 67469 |
| NA | ENSMUSG00 | 20187 |
| NA | ENSMUSG00 | 270192 |
| NA | ENSMUSG00 | 66202 |
| NA | ENSMUSG00 | 20818 |
| Significant | ENSMUSG00 | 22041 |
| NA | ENSMUSG00 | 235559 |
| NA | ENSMUSG00 | 66663 |
| NA | ENSMUSG00 | 74025 |
| NA | ENSMUSG00 | 235567 |
| NA | ENSMUSG00 | 56318 |
| NA | ENSMUSG00 | 14678 |
| NA | ENSMUSG00 | 94062 |
| NA | ENSMUSG00 | 75686 |
| NA | ENSMUSG00 | 66595 |
| NA | ENSMUSG00 | 235574 |
| NA | ENSMUSG00 | 75669 |
| NA | ENSMUSG00 | 74840 |
| Significant | ENSMUSG00 | 102626 |
| NA | ENSMUSG00 | 12700 |
| NA | ENSMUSG00 | 69536 |
| NA | ENSMUSG00 | 83486 |
| NA | ENSMUSG00 | 19654 |
| NA | ENSMUSG00 | 72825 |
| NA | ENSMUSG00 | 22036 |
| Significant | ENSMUSG00 | 12217 |
| NA | ENSMUSG00 | 235606 |
| NA | ENSMUSG00 | 320844 |
| NA | ENSMUSG00 | 27399 |
| NA | ENSMUSG00 | 69398 |
| NA | ENSMUSG00 | 74153 |

| | | |
|---|---|---|
| NA | ENSMUSG00 | 80987 |
| NA | ENSMUSG00 | 76500 |
| NA | ENSMUSG00 | 19087 |
| NA | ENSMUSG00 | 57279 |
| NA | ENSMUSG00 | 97541 |
| NA | ENSMUSG00 | 66257 |
| NA | ENSMUSG00 | 434437 |
| NA | ENSMUSG00 | 78267 |
| NA | ENSMUSG00 | 22258 |
| NA | ENSMUSG00 | 103850 |
| NA | ENSMUSG00 | 218543 |
| NA | ENSMUSG00 | 78798 |
| NA | ENSMUSG00 | 216805 |
| NA | ENSMUSG00 | 233871 |
| NA | ENSMUSG00 | 269941 |
| NA | ENSMUSG00 | 14760 |
| NA | ENSMUSG00 | 14201 |
| NA | ENSMUSG00 | 19309 |
| NA | ENSMUSG00 | 232430 |
| NA | ENSMUSG00 | 68521 |
| NA | ENSMUSG00 | 69399 |
| Significant | ENSMUSG00 | 330260 |
| NA | ENSMUSG00 | 67939 |
| NA | ENSMUSG00 | 12509 |
| NA | ENSMUSG00 | 228778 |
| NA | ENSMUSG00 | 240354 |
| NA | ENSMUSG00 | 246728 |
| NA | ENSMUSG00 | 216799 |
| NA | ENSMUSG00 | 16909 |
| NA | ENSMUSG00 | 97827 |
| NA | ENSMUSG00 | 67246 |
| NA | ENSMUSG00 | 71709 |
| NA | ENSMUSG00 | 228775 |
| NA | ENSMUSG00 | 17240 |
| NA | ENSMUSG00 | 67729 |
| NA | ENSMUSG00 | 99382 |
| Significant | ENSMUSG00 | 14276 |
| NA | ENSMUSG00 | 218613 |
| NA | ENSMUSG00 | 235406 |
| NA | ENSMUSG00 | 16332 |
| Significant | ENSMUSG00 | 378460 |
| NA | ENSMUSG00 | 108686 |
| NA | ENSMUSG00 | 252972 |
| NA | ENSMUSG00 | 233865 |
| NA | ENSMUSG00 | 73274 |
| NA | ENSMUSG00 NA | |

| | | |
|---|---|---|
| NA | ENSMUSG00 | 170756 |
| NA | ENSMUSG00 | 12068 |
| NA | ENSMUSG00 | 216136 |
| NA | ENSMUSG00 | 66066 |
| Significant | ENSMUSG00 | 277328 |
| NA | ENSMUSG00 | 233863 |
| NA | ENSMUSG00 | 216846 |
| NA | ENSMUSG00 | 78733 |
| NA | ENSMUSG00 | 11655 |
| NA | ENSMUSG00 | 216134 |
| NA | ENSMUSG00 | 76650 |
| NA | ENSMUSG00 | 321022 |
| NA | ENSMUSG00 | 214601 |
| NA | ENSMUSG00 | 69710 |
| NA | ENSMUSG00 | 14087 |
| NA | ENSMUSG00 | 109676 |
| NA | ENSMUSG00 | 243725 |
| NA | ENSMUSG00 | 110816 |
| NA | ENSMUSG00 | 76792 |
| NA | ENSMUSG00 | 72446 |
| NA | ENSMUSG00 | 224814 |
| NA | ENSMUSG00 | 239273 |
| NA | ENSMUSG00 | 18763 |
| NA | ENSMUSG00 | 18442 |
| NA | ENSMUSG00 | 231672 |
| NA | ENSMUSG00 | 228769 |
| NA | ENSMUSG00 | 69780 |
| NA | ENSMUSG00 NA | |
| NA | ENSMUSG00 | 74205 |
| NA | ENSMUSG00 | 242662 |
| NA | ENSMUSG00 | 57785 |
| NA | ENSMUSG00 | 18046 |
| NA | ENSMUSG00 | 231670 |
| NA | ENSMUSG00 | 20501 |
| NA | ENSMUSG00 | 67526 |
| NA | ENSMUSG00 | 269473 |
| Significant | ENSMUSG00 | 52614 |
| NA | ENSMUSG00 | 110920 |
| NA | ENSMUSG00 | 71805 |
| NA | ENSMUSG00 | 19395 |
| NA | ENSMUSG00 | 67489 |
| NA | ENSMUSG00 | 23980 |
| NA | ENSMUSG00 | 73916 |
| NA | ENSMUSG00 | 14225 |
| NA | ENSMUSG00 | 108707 |
| NA | ENSMUSG00 | 74241 |

| | | |
|---|---|---|
| NA | ENSMUSG00 | 230700 |
| NA | ENSMUSG00 | 105689 |
| NA | ENSMUSG00 | 270086 |
| NA | ENSMUSG00 | 94093 |
| NA | ENSMUSG00 | 18018 |
| NA | ENSMUSG00 | 69833 |
| NA | ENSMUSG00 | 69080 |
| NA | ENSMUSG00 | 12583 |
| NA | ENSMUSG00 | 224171 |
| NA | ENSMUSG00 | 106877 |
| NA | ENSMUSG00 | 72691 |
| NA | ENSMUSG00 | 665989 |
| NA | ENSMUSG00 | 27008 |
| NA | ENSMUSG00 | 223691 |
| NA | ENSMUSG00 | 244879 |
| NA | ENSMUSG00 | 223690 |
| NA | ENSMUSG00 | 110078 |
| NA | ENSMUSG00 | 18642 |
| NA | ENSMUSG00 | 14457 |
| NA | ENSMUSG00 | 12497 |
| NA | ENSMUSG00 | 223870 |
| NA | ENSMUSG00 | 210789 |
| NA | ENSMUSG00 | 110253 |
| NA | ENSMUSG00 | 71881 |
| NA | ENSMUSG00 | 97961 |
| NA | ENSMUSG00 | 218294 |
| NA | ENSMUSG00 | 16987 |
| NA | ENSMUSG00 | 66432 |
| NA | ENSMUSG00 | 67664 |
| NA | ENSMUSG00 | 218734 |
| NA | ENSMUSG00 | 70484 |
| NA | ENSMUSG00 | 245860 |
| NA | ENSMUSG00 | 216119 |
| NA | ENSMUSG00 | 106039 |
| NA | ENSMUSG00 | 242126 |
| NA | ENSMUSG00 | 208177 |
| NA | ENSMUSG00 | 319655 |
| NA | ENSMUSG00 | 213012 |
| NA | ENSMUSG00 | 69171 |
| NA | ENSMUSG00 | 11928 |
| NA | ENSMUSG00 | 72662 |
| NA | ENSMUSG00 | 105844 |
| NA | ENSMUSG00 | 23945 |
| NA | ENSMUSG00 | 66676 |
| NA | ENSMUSG00 | 76789 |
| NA | ENSMUSG00 | 21846 |

| | | |
|---|---|---|
| NA | ENSMUSG00 | 270084 |
| NA | ENSMUSG00 | 209683 |
| NA | ENSMUSG00 | 65967 |
| NA | ENSMUSG00 | 19354 |
| NA | ENSMUSG00 | 74044 |
| NA | ENSMUSG00 | 22420 |
| NA | ENSMUSG00 | 72193 |
| NA | ENSMUSG00 | 229644 |
| NA | ENSMUSG00 | 77044 |
| NA | ENSMUSG00 | 230676 |
| NA | ENSMUSG00 | 333669 |
| NA | ENSMUSG00 | 67534 |
| NA | ENSMUSG00 | 67843 |
| NA | ENSMUSG00 | 269209 |
| NA | ENSMUSG00 | 19274 |
| NA | ENSMUSG00 | 244585 |
| Significant | ENSMUSG00 | 269470 |
| NA | ENSMUSG00 | 100608 |
| NA | ENSMUSG00 | 19268 |
| NA | ENSMUSG00 | 210126 |
| NA | ENSMUSG00 | 17319 |
| NA | ENSMUSG00 | 50788 |
| NA | ENSMUSG00 | 14872 |
| NA | ENSMUSG00 | 240263 |
| NA | ENSMUSG00 | 67655 |
| NA | ENSMUSG00 | 230674 |
| NA | ENSMUSG00 | 13430 |
| NA | ENSMUSG00 | 54371 |
| NA | ENSMUSG00 | 26398 |
| NA | ENSMUSG00 | 67775 |
| NA | ENSMUSG00 | 78895 |
| NA | ENSMUSG00 | 319651 |
| NA | ENSMUSG00 | 230673 |
| NA | ENSMUSG00 | 110606 |
| NA | ENSMUSG00 | 114143 |
| NA | ENSMUSG00 | 75964 |
| NA | ENSMUSG00 | 20321 |
| NA | ENSMUSG00 | 76499 |
| NA | ENSMUSG00 | 214585 |
| NA | ENSMUSG00 | 77559 |
| NA | ENSMUSG00 | 214254 |
| NA | ENSMUSG00 | 329506 |
| NA | ENSMUSG00 | 68778 |
| NA | ENSMUSG00 | 78832 |
| NA | ENSMUSG00 | 140546 |
| NA | ENSMUSG00 | 73724 |

| | | |
|---|---|---|
| NA | ENSMUSG00 | 57321 |
| NA | ENSMUSG00 | 107999 |
| NA | ENSMUSG00 | 67416 |
| NA | ENSMUSG00 | 229780 |
| NA | ENSMUSG00 | 74392 |
| NA | ENSMUSG00 | 67168 |
| NA | ENSMUSG00 | 72536 |
| NA | ENSMUSG00 | 268564 |
| NA | ENSMUSG00 | 330554 |
| NA | ENSMUSG00 | 78248 |
| NA | ENSMUSG00 | 57914 |
| NA | ENSMUSG00 | 66119 |
| NA | ENSMUSG00 | 226252 |
| NA | ENSMUSG00 | 212670 |
| NA | ENSMUSG00 | 319448 |
| NA | ENSMUSG00 | 226527 |
| NA | ENSMUSG00 | 217980 |
| NA | ENSMUSG00 | 229776 |
| NA | ENSMUSG00 | 327655 |
| NA | ENSMUSG00 | 104718 |
| NA | ENSMUSG00 | 12363 |
| NA | ENSMUSG00 | 15932 |
| NA | ENSMUSG00 | 54324 |
| NA | ENSMUSG00 | 235459 |
| NA | ENSMUSG00 | 170737 |
| NA | ENSMUSG00 | 69740 |
| NA | ENSMUSG00 | 215015 |
| NA | ENSMUSG00 | 93686 |
| NA | ENSMUSG00 | 17920 |
| NA | ENSMUSG00 | 319765 |
| NA | ENSMUSG00 | 17984 |
| NA | ENSMUSG00 | 72549 |
| NA | ENSMUSG00 | 78779 |
| NA | ENSMUSG00 | 234736 |
| NA | ENSMUSG00 | 75735 |
| NA | ENSMUSG00 | 71751 |
| NA | ENSMUSG00 | 69587 |
| NA | ENSMUSG00 | 319518 |
| NA | ENSMUSG00 | 225326 |
| NA | ENSMUSG00 | 57874 |
| Significant | ENSMUSG00 | 225289 |
| NA | ENSMUSG00 | 57746 |
| NA | ENSMUSG00 | 209018 |
| NA | ENSMUSG00 | 234733 |
| NA | ENSMUSG00 | 223669 |
| NA | ENSMUSG00 | 74081 |

| | | |
|---|---|---|
| NA | ENSMUSG00 | 104009 |
| NA | ENSMUSG00 | 22228 |
| NA | ENSMUSG00 | 223666 |
| NA | ENSMUSG00 | 72482 |
| NA | ENSMUSG00 | 234730 |
| NA | ENSMUSG00 | 668880 |
| NA | ENSMUSG00 | 232187 |
| NA | ENSMUSG00 | 219158 |
| NA | ENSMUSG00 | 71375 |
| NA | ENSMUSG00 | 94282 |
| NA | ENSMUSG00 | 57257 |
| NA | ENSMUSG00 | 223664 |
| NA | ENSMUSG00 | 13655 |
| NA | ENSMUSG00 | 101943 |
| NA | ENSMUSG00 | 20751 |
| NA | ENSMUSG00 | 333564 |
| NA | ENSMUSG00 | 56299 |
| NA | ENSMUSG00 | 102060 |
| NA | ENSMUSG00 | 76915 |
| NA | ENSMUSG00 | 66704 |
| NA | ENSMUSG00 | 79456 |
| NA | ENSMUSG00 | 244654 |
| NA | ENSMUSG00 | 229473 |
| NA | ENSMUSG00 | 75914 |
| NA | ENSMUSG00 | 231571 |
| NA | ENSMUSG00 | 279572 |
| NA | ENSMUSG00 | 142688 |
| NA | ENSMUSG00 | 233276 |
| NA | ENSMUSG00 | 11977 |
| NA | ENSMUSG00 | 108664 |
| NA | ENSMUSG00 | 105203 |
| NA | ENSMUSG00 | 211499 |
| NA | ENSMUSG00 | 208624 |
| NA | ENSMUSG00 | 21399 |
| NA | ENSMUSG00 | 73062 |
| NA | ENSMUSG00 | 13417 |
| NA | ENSMUSG00 | 27395 |
| NA | ENSMUSG00 | 232889 |
| NA | ENSMUSG00 | 102238433 |
| NA | ENSMUSG00 | 217364 |
| NA | ENSMUSG00 | 234854 |
| NA | ENSMUSG00 | 16601 |
| NA | ENSMUSG00 | 170826 |
| NA | ENSMUSG00 | 19039 |
| NA | ENSMUSG00 | 241944 |
| NA | ENSMUSG00 | 218699 |

| | | |
|---|---|---|
| NA | ENSMUSG00 | 213582 |
| NA | ENSMUSG00 | 26390 |
| NA | ENSMUSG00 | 101565 |
| NA | ENSMUSG00 | 108672 |
| NA | ENSMUSG00 | 72344 |
| NA | ENSMUSG00 | 68953 |
| NA | ENSMUSG00 | 56209 |
| NA | ENSMUSG00 | 381038 |
| NA | ENSMUSG00 | 52202 |
| NA | ENSMUSG00 | 22346 |
| NA | ENSMUSG00 | 66916 |
| NA | ENSMUSG00 | 101314 |
| NA | ENSMUSG00 | 29808 |
| NA | ENSMUSG00 | 74464 |
| NA | ENSMUSG00 | 28105 |
| NA | ENSMUSG00 | 12316 |
| NA | ENSMUSG00 | 19058 |
| NA | ENSMUSG00 | 240185 |
| NA | ENSMUSG00 | 269870 |
| NA | ENSMUSG00 | 226470 |
| NA | ENSMUSG00 | 20502 |
| NA | ENSMUSG00 | 381845 |
| NA | ENSMUSG00 | 69263 |
| NA | ENSMUSG00 | 319615 |
| NA | ENSMUSG00 | 12812 |
| Significant | ENSMUSG00 | 230661 |
| NA | ENSMUSG00 | 218343 |
| NA | ENSMUSG00 | 66943 |
| NA | ENSMUSG00 | 244891 |
| NA | ENSMUSG00 | 100710 |
| NA | ENSMUSG00 | 94230 |
| NA | ENSMUSG00 | 211651 |
| Significant | ENSMUSG00 | 12461 |
| NA | ENSMUSG00 | 68925 |
| NA | ENSMUSG00 | 622665 |
| NA | ENSMUSG00 | 232237 |
| NA | ENSMUSG00 | 17095 |
| NA | ENSMUSG00 | 77110 |
| NA | ENSMUSG00 | 18679 |
| NA | ENSMUSG00 | 224143 |
| NA | ENSMUSG00 | 227377 |
| NA | ENSMUSG00 | 619331 |
| NA | ENSMUSG00 | 228136 |
| NA | ENSMUSG00 | 232236 |
| NA | ENSMUSG00 | 110611 |
| NA | ENSMUSG00 | 12388 |

| | | |
|---|---|---|
| NA | ENSMUSG00 | 74347 |
| NA | ENSMUSG00 | 12460 |
| NA | ENSMUSG00 | 73124 |
| NA | ENSMUSG00 | 212919 |
| NA | ENSMUSG00 | 382620 |
| NA | ENSMUSG00 | 22324 |
| NA | ENSMUSG00 | 22021 |
| NA | ENSMUSG00 | 20382 |
| NA | ENSMUSG00 | 380718 |
| NA | ENSMUSG00 | 217734 |
| NA | ENSMUSG00 | 216350 |
| NA | ENSMUSG00 | 70661 |
| NA | ENSMUSG00 | 211446 |
| NA | ENSMUSG00 | 68142 |
| NA | ENSMUSG00 | 207777 |
| NA | ENSMUSG00 | 217732 |
| NA | ENSMUSG00 | 320184 |
| NA | ENSMUSG00 | 108155 |
| NA | ENSMUSG00 | 216345 |
| Significant | ENSMUSG00 | 140703 |
| Significant | ENSMUSG00 | 12445 |
| NA | ENSMUSG00 | 238330 |
| NA | ENSMUSG00 | 71970 |
| NA | ENSMUSG00 | 279766 |
| Significant | ENSMUSG00 | 207742 |
| NA | ENSMUSG00 | 18195 |
| NA | ENSMUSG00 | 72552 |
| NA | ENSMUSG00 | 105513 |
| NA | ENSMUSG00 | 67678 |
| NA | ENSMUSG00 | 71843 |
| NA | ENSMUSG00 | 78926 |
| NA | ENSMUSG00 | 72170 |
| NA | ENSMUSG00 | 94352 |
| NA | ENSMUSG00 | 77782 |
| NA | ENSMUSG00 | 230648 |
| NA | ENSMUSG00 | 66258 |
| NA | ENSMUSG00 | 228545 |
| NA | ENSMUSG00 | 11920 |
| NA | ENSMUSG00 | 14733 |
| NA | ENSMUSG00 | 228543 |
| NA | ENSMUSG00 | 78787 |
| NA | ENSMUSG00 | 224139 |
| NA | ENSMUSG00 | 232232 |
| NA | ENSMUSG00 | 353047 |
| NA | ENSMUSG00 | 67712 |
| NA | ENSMUSG00 | 215351 |

| | | |
|---|---|---|
| NA | ENSMUSG00 | 55979 |
| NA | ENSMUSG00 | 544817 |
| NA | ENSMUSG00 | 109075 |
| NA | ENSMUSG00 | 69882 |
| NA | ENSMUSG00 | 224454 |
| NA | ENSMUSG00 | 53314 |
| NA | ENSMUSG00 | 72895 |
| NA | ENSMUSG00 | 107829 |
| NA | ENSMUSG00 | 235086 |
| NA | ENSMUSG00 | 69408 |
| NA | ENSMUSG00 | 18082 |
| NA | ENSMUSG00 | 217718 |
| NA | ENSMUSG00 | 74019 |
| NA | ENSMUSG00 | 327987 |
| NA | ENSMUSG00 | 66306 |
| NA | ENSMUSG00 | 245902 |
| NA | ENSMUSG00 | 74032 |
| NA | ENSMUSG00 | 16571 |
| NA | ENSMUSG00 | 232227 |
| NA | ENSMUSG00 | 66949 |
| NA | ENSMUSG00 | 13521 |
| NA | ENSMUSG00 | 66583 |
| NA | ENSMUSG00 | 105440 |
| NA | ENSMUSG00 | 237911 |
| NA | ENSMUSG00 | 234779 |
| NA | ENSMUSG00 | 223773 |
| NA | ENSMUSG00 | 22378 |
| NA | ENSMUSG00 | 12402 |
| NA | ENSMUSG00 | 67921 |
| NA | ENSMUSG00 | 66467 |
| NA | ENSMUSG00 | 70099 |
| Significant | ENSMUSG00 | 51801 |
| NA | ENSMUSG00 | 74302 |
| NA | ENSMUSG00 | 234577 |
| NA | ENSMUSG00 | 225913 |
| NA | ENSMUSG00 | 68842 |
| NA | ENSMUSG00 | 69544 |
| NA | ENSMUSG00 | 74440 |
| NA | ENSMUSG00 | 16878 |
| Significant | ENSMUSG00 | 67946 |
| NA | ENSMUSG00 | 18744 |
| NA | ENSMUSG00 | 103724 |
| NA | ENSMUSG00 | 76645 |
| NA | ENSMUSG00 | 547253 |
| NA | ENSMUSG00 | 68133 |
| NA | ENSMUSG00 | 217328 |

| | | |
|---|---|---|
| NA | ENSMUSG00 | 69020 |
| NA | ENSMUSG00 | 80292 |
| NA | ENSMUSG00 | 108679 |
| NA | ENSMUSG00 | 76074 |
| NA | ENSMUSG00 | 76357 |
| NA | ENSMUSG00 | 225912 |
| NA | ENSMUSG00 | 192970 |
| NA | ENSMUSG00 | 70428 |
| NA | ENSMUSG00 | 245527 |
| NA | ENSMUSG00 | 15957 |
| NA | ENSMUSG00 | 18764 |
| NA | ENSMUSG00 | 140721 |
| NA | ENSMUSG00 | 317717 |
| NA | ENSMUSG00 | 54004 |
| Significant | ENSMUSG00 | 67804 |
| NA | ENSMUSG00 | 72565 |
| Significant | ENSMUSG00 | 68304 |
| NA | ENSMUSG00 | 67708 |
| NA | ENSMUSG00 | 66591 |
| NA | ENSMUSG00 | 70823 |
| NA | ENSMUSG00 | 380912 |
| NA | ENSMUSG00 | 218333 |
| NA | ENSMUSG00 | 232854 |
| NA | ENSMUSG00 | 74522 |
| NA | ENSMUSG00 | 66880 |
| NA | ENSMUSG00 | 245596 |
| NA | ENSMUSG00 | 230597 |
| NA | ENSMUSG00 | 319277 |
| NA | ENSMUSG00 | 72278 |
| NA | ENSMUSG00 | 71679 |
| NA | ENSMUSG00 | 19249 |
| NA | ENSMUSG00 | 208846 |
| NA | ENSMUSG00 | 210106 |
| NA | ENSMUSG00 | 217310 |
| NA | ENSMUSG00 | 17918 |
| NA | ENSMUSG00 | 76448 |
| NA | ENSMUSG00 | 76967 |
| NA | ENSMUSG00 | 67074 |
| NA | ENSMUSG00 | 230594 |
| NA | ENSMUSG00 | 58250 |
| NA | ENSMUSG00 | 319468 |
| NA | ENSMUSG00 | 216505 |
| NA | ENSMUSG00 | 245857 |
| NA | ENSMUSG00 | 210009 |
| NA | ENSMUSG00 | 216395 |
| NA | ENSMUSG00 | 237943 |

| | | |
|---|---|---|
| NA | ENSMUSG00 | 414872 |
| NA | ENSMUSG00 | 74729 |
| NA | ENSMUSG00 | 99929 |
| Significant | ENSMUSG00 | 217305 |
| NA | ENSMUSG00 | 106585 |
| Significant | ENSMUSG00 | 217303 |
| NA | ENSMUSG00 | 240255 |
| NA | ENSMUSG00 | 12286 |
| NA | ENSMUSG00 | 68539 |
| NA | ENSMUSG00 | 140780 |
| Significant | ENSMUSG00 | 16399 |
| NA | ENSMUSG00 | 73192 |
| NA | ENSMUSG00 | 18515 |
| NA | ENSMUSG00 | 21665 |
| NA | ENSMUSG00 | 56320 |
| NA | ENSMUSG00 | 19826 |
| NA | ENSMUSG00 | 75612 |
| Significant | ENSMUSG00 | 14824 |
| NA | ENSMUSG00 | 73825 |
| NA | ENSMUSG00 | 117160 |
| NA | ENSMUSG00 | 52837 |
| NA | ENSMUSG00 | 231464 |
| NA | ENSMUSG00 | 64657 |
| NA | ENSMUSG00 | 380921 |
| NA | ENSMUSG00 | 329416 |
| NA | ENSMUSG00 | 56174 |
| NA | ENSMUSG00 | 50721 |
| NA | ENSMUSG00 | 328329 |
| NA | ENSMUSG00 | 72053 |
| NA | ENSMUSG00 | 114606 |
| NA | ENSMUSG00 | 399510 |
| NA | ENSMUSG00 | 240672 |
| NA | ENSMUSG00 | 217216 |
| NA | ENSMUSG00 | 14672 |
| NA | ENSMUSG00 | 106512 |
| NA | ENSMUSG00 | 19336 |
| NA | ENSMUSG00 | 14676 |
| NA | ENSMUSG00 | 68401 |
| NA | ENSMUSG00 | 108811 |
| NA | ENSMUSG00 | 382018 |
| NA | ENSMUSG00 | 72180 |
| NA | ENSMUSG00 | 20663 |
| NA | ENSMUSG00 | 234407 |
| NA | ENSMUSG00 | 269061 |
| NA | ENSMUSG00 | 269113 |
| NA | ENSMUSG00 | 194388 |

| | | |
|---|---|---|
| Significant | ENSMUSG00 | 67596 |
| NA | ENSMUSG00 | 14685 |
| NA | ENSMUSG00 | 84094 |
| NA | ENSMUSG00 | 73668 |
| NA | ENSMUSG00 | 69470 |
| NA | ENSMUSG00 | 329910 |
| NA | ENSMUSG00 | 73822 |
| NA | ENSMUSG00 | 15945 |
| NA | ENSMUSG00 | 235493 |
| NA | ENSMUSG00 | 382014 |
| NA | ENSMUSG00 | 245886 |
| Significant | ENSMUSG00 | 67938 |
| NA | ENSMUSG00 | 110959 |
| NA | ENSMUSG00 | 94065 |
| NA | ENSMUSG00 | 69706 |
| NA | ENSMUSG00 | 70312 |
| NA | ENSMUSG00 | 20090 |
| NA | ENSMUSG00 | 338337 |
| NA | ENSMUSG00 | 18717 |
| NA | ENSMUSG00 | 319876 |
| NA | ENSMUSG00 | 215387 |
| NA | ENSMUSG00 | 214597 |
| NA | ENSMUSG00 | 234395 |
| NA | ENSMUSG00 | 27375 |
| NA | ENSMUSG00 | 74754 |
| NA | ENSMUSG00 | 105239 |
| NA | ENSMUSG00 | 20166 |
| NA | ENSMUSG00 | 217207 |
| NA | ENSMUSG00 | 66047 |
| NA | ENSMUSG00 | 80981 |
| NA | ENSMUSG00 | 217030 |
| Significant | ENSMUSG00 | 217203 |
| NA | ENSMUSG00 | 233824 |
| NA | ENSMUSG00 | 12606 |
| NA | ENSMUSG00 | 271221 |
| NA | ENSMUSG00 | 16815 |
| NA | ENSMUSG00 | 320615 |
| NA | ENSMUSG00 | 13144 |
| NA | ENSMUSG00 | 15466 |
| NA | ENSMUSG00 | 26949 |
| NA | ENSMUSG00 | 13629 |
| NA | ENSMUSG00 | 14236 |
| NA | ENSMUSG00 | 80719 |
| NA | ENSMUSG00 | 217201 |
| NA | ENSMUSG00 | 16969 |
| NA | ENSMUSG00 | 217578 |

| | | |
|---|---|---|
| NA | ENSMUSG00 | 78658 |
| NA | ENSMUSG00 | 26396 |
| NA | ENSMUSG00 | 208677 |
| NA | ENSMUSG00 | 547176 |
| NA | ENSMUSG00 | 215194 |
| NA | ENSMUSG00 | 69010 |
| NA | ENSMUSG00 | 107094 |
| NA | ENSMUSG00 | 106794 |
| NA | ENSMUSG00 | 245522 |
| NA | ENSMUSG00 | 13631 |
| NA | ENSMUSG00 | 67013 |
| NA | ENSMUSG00 | 210376 |
| NA | ENSMUSG00 | 56208 |
| NA | ENSMUSG00 | 70354 |
| NA | ENSMUSG00 | 232146 |
| Significant | ENSMUSG00 | 112407 |
| NA | ENSMUSG00 | 73379 |
| NA | ENSMUSG00 | 271849 |
| NA | ENSMUSG00 | 330361 |
| NA | ENSMUSG00 | 242584 |
| NA | ENSMUSG00 | 11855 |
| NA | ENSMUSG00 | 75420 |
| NA | ENSMUSG00 | 76826 |
| NA | ENSMUSG00 | 26905 |
| NA | ENSMUSG00 | 244548 |
| NA | ENSMUSG00 | 71770 |
| NA | ENSMUSG00 | 17342 |
| NA | ENSMUSG00 | 18130 |
| NA | ENSMUSG00 | 244871 |
| NA | ENSMUSG00 | 57442 |
| NA | ENSMUSG00 | 66860 |
| NA | ENSMUSG00 | NA |
| NA | ENSMUSG00 | 213993 |
| NA | ENSMUSG00 | 320487 |
| NA | ENSMUSG00 | 317750 |
| NA | ENSMUSG00 | 103733 |
| NA | ENSMUSG00 | 65106 |
| NA | ENSMUSG00 | 108015 |
| NA | ENSMUSG00 | 102436 |
| NA | ENSMUSG00 | 13854 |
| NA | ENSMUSG00 | 73218 |
| NA | ENSMUSG00 | 276950 |
| NA | ENSMUSG00 | 230514 |
| NA | ENSMUSG00 | 66094 |
| NA | ENSMUSG00 | 66624 |
| NA | ENSMUSG00 | 232821 |

| | | |
|---|---|---|
| NA | ENSMUSG00 | 236900 |
| NA | ENSMUSG00 | 70681 |
| NA | ENSMUSG00 | 66597 |
| NA | ENSMUSG00 | 320271 |
| NA | ENSMUSG00 | 50877 |
| NA | ENSMUSG00 | 18245 |
| NA | ENSMUSG00 | 101351 |
| NA | ENSMUSG00 | 236899 |
| NA | ENSMUSG00 | 207304 |
| NA | ENSMUSG00 | 214290 |
| NA | ENSMUSG00 | 191578 |
| NA | ENSMUSG00 | 18769 |
| Significant | ENSMUSG00 | 15442 |
| NA | ENSMUSG00 | 78670 |
| NA | ENSMUSG00 | 11554 |
| NA | ENSMUSG00 | 320528 |
| NA | ENSMUSG00 | 217558 |
| NA | ENSMUSG00 | 76646 |
| NA | ENSMUSG00 | 26891 |
| NA | ENSMUSG00 | 17318 |
| NA | ENSMUSG00 | NA |
| NA | ENSMUSG00 | 432486 |
| NA | ENSMUSG00 | 233552 |
| NA | ENSMUSG00 | 69162 |
| NA | ENSMUSG00 | 70611 |
| NA | ENSMUSG00 | 226154 |
| NA | ENSMUSG00 | 69736 |
| Significant | ENSMUSG00 | 20293 |
| NA | ENSMUSG00 | 78610 |
| NA | ENSMUSG00 | 80859 |
| NA | ENSMUSG00 | 75317 |
| NA | ENSMUSG00 | 74386 |
| NA | ENSMUSG00 | NA |
| NA | ENSMUSG00 | 72056 |
| Significant | ENSMUSG00 | 20306 |
| NA | ENSMUSG00 | 70757 |
| NA | ENSMUSG00 | 72171 |
| NA | ENSMUSG00 | 18554 |
| Significant | ENSMUSG00 | 20296 |
| NA | ENSMUSG00 | 227801 |
| NA | ENSMUSG00 | 118445 |
| NA | ENSMUSG00 | 66680 |
| NA | ENSMUSG00 | 233545 |
| NA | ENSMUSG00 | 103743 |
| NA | ENSMUSG00 | 227800 |
| NA | ENSMUSG00 | 76478 |

| | | |
|---|---|---|
| NA | ENSMUSG00 | 338367 |
| NA | ENSMUSG00 | 77862 |
| NA | ENSMUSG00 | 12771 |
| NA | ENSMUSG00 | 60530 |
| NA | ENSMUSG00 | 71330 |
| NA | ENSMUSG00 | 319625 |
| NA | ENSMUSG00 | 66724 |
| NA | ENSMUSG00 | 17192 |
| Significant | ENSMUSG00 | 21810 |
| NA | ENSMUSG00 | 272027 |
| NA | ENSMUSG00 | 70335 |
| NA | ENSMUSG00 | 231430 |
| NA | ENSMUSG00 | 213311 |
| NA | ENSMUSG00 | 100121 |
| NA | ENSMUSG00 | 214505 |
| NA | ENSMUSG00 | 72843 |
| NA | ENSMUSG00 | 20918 |
| NA | ENSMUSG00 | 381522 |
| NA | ENSMUSG00 | 232798 |
| NA | ENSMUSG00 | 12337 |
| NA | ENSMUSG00 | 234384 |
| NA | ENSMUSG00 | 72535 |
| NA | ENSMUSG00 | 77087 |
| NA | ENSMUSG00 | 242418 |
| NA | ENSMUSG00 | 216987 |
| NA | ENSMUSG00 | 69707 |
| NA | ENSMUSG00 | 66078 |
| NA | ENSMUSG00 | 69770 |
| NA | ENSMUSG00 | 77582 |
| NA | ENSMUSG00 | 328110 |
| NA | ENSMUSG00 | 69934 |
| NA | ENSMUSG00 | 328108 |
| NA | ENSMUSG00 | 237422 |
| NA | ENSMUSG00 | 59090 |
| NA | ENSMUSG00 | 233532 |
| NA | ENSMUSG00 | 100502698 |
| NA | ENSMUSG00 | 232791 |
| NA | ENSMUSG00 | 76238 |
| NA | ENSMUSG00 | 66273 |
| NA | ENSMUSG00 | 319885 |
| NA | ENSMUSG00 | 269252 |
| NA | ENSMUSG00 | 212168 |
| NA | ENSMUSG00 | 216161 |
| NA | ENSMUSG00 | 66091 |
| NA | ENSMUSG00 | 21950 |
| NA | ENSMUSG00 | 17279 |

| | | |
|---|---|---|
| NA | ENSMUSG00 | 100038882 |
| NA | ENSMUSG00 | 73469 |
| NA | ENSMUSG00 | 70719 |
| NA | ENSMUSG00 | 381903 |
| NA | ENSMUSG00 | 27261 |
| NA | ENSMUSG00 | 244144 |
| NA | ENSMUSG00 | 27403 |
| NA | ENSMUSG00 | 19108 |
| NA | ENSMUSG00 | 114741 |
| NA | ENSMUSG00 | 269060 |
| NA | ENSMUSG00 | 216156 |
| NA | ENSMUSG00 | 223776 |
| NA | ENSMUSG00 | 71769 |
| NA | ENSMUSG00 | 72745 |
| NA | ENSMUSG00 | 268882 |
| NA | ENSMUSG00 | 69190 |
| NA | ENSMUSG00 | 102448 |
| NA | ENSMUSG00 | 234663 |
| NA | ENSMUSG00 | 118451 |
| NA | ENSMUSG00 | 171486 |
| NA | ENSMUSG00 | 14594 |
| NA | ENSMUSG00 | 109284 |
| NA | ENSMUSG00 | 11475 |
| NA | ENSMUSG00 | 66994 |
| NA | ENSMUSG00 | 320150 |
| NA | ENSMUSG00 | 57813 |
| NA | ENSMUSG00 | 223775 |
| NA | ENSMUSG00 | 67486 |
| Significant | ENSMUSG00 | 216152 |
| NA | ENSMUSG00 | 80289 |
| NA | ENSMUSG00 | 320209 |
| NA | ENSMUSG00 | 223774 |
| NA | ENSMUSG00 | 15931 |
| NA | ENSMUSG00 | 231386 |
| Significant | ENSMUSG00 | 18483 |
| NA | ENSMUSG00 | 73229 |
| NA | ENSMUSG00 | 114774 |
| NA | ENSMUSG00 | 320799 |
| NA | ENSMUSG00 | 235386 |
| NA | ENSMUSG00 | 12868 |
| NA | ENSMUSG00 | 70294 |
| NA | ENSMUSG00 | 223753 |
| NA | ENSMUSG00 | 231380 |
| NA | ENSMUSG00 | 223752 |
| NA | ENSMUSG00 | 19347 |
| NA | ENSMUSG00 | 102657 |

| | | |
|---|---|---|
| NA | ENSMUSG00 | 319845 |
| NA | ENSMUSG00 | 238123 |
| NA | ENSMUSG00 | 208076 |
| NA | ENSMUSG00 | 214579 |
| NA | ENSMUSG00 | 108837 |
| NA | ENSMUSG00 | 239570 |
| NA | ENSMUSG00 | 30050 |
| NA | ENSMUSG00 | 219024 |
| NA | ENSMUSG00 | 238130 |
| NA | ENSMUSG00 | 56196 |
| NA | ENSMUSG00 | 11792 |
| Significant | ENSMUSG00 | 236790 |
| NA | ENSMUSG00 | 216742 |
| NA | ENSMUSG00 | 230088 |
| NA | ENSMUSG00 | 193385 |
| NA | ENSMUSG00 | 216292 |
| NA | ENSMUSG00 | 278279 |
| NA | ENSMUSG00 | 78755 |
| NA | ENSMUSG00 | 11546 |
| NA | ENSMUSG00 | 224807 |
| NA | ENSMUSG00 | 235472 |
| NA | ENSMUSG00 | 22715 |
| NA | ENSMUSG00 | 223739 |
| NA | ENSMUSG00 | 56323 |
| NA | ENSMUSG00 | 71409 |
| NA | ENSMUSG00 | 234373 |
| NA | ENSMUSG00 | 104831 |
| NA | ENSMUSG00 | 71726 |
| Significant | ENSMUSG00 | 227659 |
| NA | ENSMUSG00 | 14430 |
| NA | ENSMUSG00 | 18391 |
| NA | ENSMUSG00 | 226409 |
| NA | ENSMUSG00 | 75991 |
| NA | ENSMUSG00 | 109685 |
| NA | ENSMUSG00 | 75423 |
| NA | ENSMUSG00 | 226151 |
| NA | ENSMUSG00 | 215008 |
| Significant | ENSMUSG00 | 140792 |
| NA | ENSMUSG00 | 226407 |
| NA | ENSMUSG00 | 109270 |
| Significant | ENSMUSG00 | 171170 |
| NA | ENSMUSG00 | 109594 |
| NA | ENSMUSG00 | 56307 |
| NA | ENSMUSG00 | 69961 |
| NA | ENSMUSG00 | 19727 |
| NA | ENSMUSG00 | 113868 |

| | | |
|---|---|---|
| NA | ENSMUSG00 | 107895 |
| NA | ENSMUSG00 | 106042 |
| NA | ENSMUSG00 | 20935 |
| NA | ENSMUSG00 | 223828 |
| NA | ENSMUSG00 | 237465 |
| NA | ENSMUSG00 | 234366 |
| NA | ENSMUSG00 | 50708 |
| NA | ENSMUSG00 | 217473 |
| NA | ENSMUSG00 | 223827 |
| NA | ENSMUSG00 | 67184 |
| NA | ENSMUSG00 | 51869 |
| NA | ENSMUSG00 | 98402 |
| NA | ENSMUSG00 | 212547 |
| NA | ENSMUSG00 | 76416 |
| NA | ENSMUSG00 | 66468 |
| NA | ENSMUSG00 | 67615 |
| NA | ENSMUSG00 | 67419 |
| NA | ENSMUSG00 | 78816 |
| NA | ENSMUSG00 | 71684 |
| NA | ENSMUSG00 | 67452 |
| NA | ENSMUSG00 | 234699 |
| NA | ENSMUSG00 | 66725 |
| NA | ENSMUSG00 | 213673 |
| NA | ENSMUSG00 | 227644 |
| NA | ENSMUSG00 | 70646 |
| NA | ENSMUSG00 | 56412 |
| NA | ENSMUSG00 | 74385 |
| NA | ENSMUSG00 | 207798 |
| NA | ENSMUSG00 | 407819 |
| NA | ENSMUSG00 | 332175 |
| NA | ENSMUSG00 | 211134 |
| NA | ENSMUSG00 | 21402 |
| NA | ENSMUSG00 | 66136 |
| NA | ENSMUSG00 | 66661 |
| NA | ENSMUSG00 | 227638 |
| NA | ENSMUSG00 | 110877 |
| NA | ENSMUSG00 | 77480 |
| NA | ENSMUSG00 | 242050 |
| NA | ENSMUSG00 | 218989 |
| NA | ENSMUSG00 | 98766 |
| Significant | ENSMUSG00 | 70839 |
| NA | ENSMUSG00 | 74191 |
| NA | ENSMUSG00 | 381110 |
| NA | ENSMUSG00 | 66870 |
| NA | ENSMUSG00 | 69038 |
| NA | ENSMUSG00 | 30946 |

| | | |
|---|---|---|
| NA | ENSMUSG00 | 320827 |
| Significant | ENSMUSG00 | NA |
| NA | ENSMUSG00 | 13197 |
| NA | ENSMUSG00 | 77371 |
| NA | ENSMUSG00 | 244757 |
| NA | ENSMUSG00 | 76497 |
| Significant | ENSMUSG00 | 14701 |
| NA | ENSMUSG00 | 381644 |
| NA | ENSMUSG00 | 68283 |
| NA | ENSMUSG00 | 14751 |
| NA | ENSMUSG00 | 72726 |
| NA | ENSMUSG00 | 20439 |
| NA | ENSMUSG00 | 69940 |
| NA | ENSMUSG00 | 12314 |
| NA | ENSMUSG00 | 59016 |
| NA | ENSMUSG00 | 17022 |
| NA | ENSMUSG00 | 319594 |
| Significant | ENSMUSG00 | 71302 |
| NA | ENSMUSG00 | 13709 |
| NA | ENSMUSG00 | 72925 |
| NA | ENSMUSG00 | 224617 |
| NA | ENSMUSG00 | 12226 |
| NA | ENSMUSG00 | 216238 |
| NA | ENSMUSG00 | 225358 |
| NA | ENSMUSG00 | 245386 |
| NA | ENSMUSG00 | 24017 |
| NA | ENSMUSG00 | 75454 |
| NA | ENSMUSG00 | 52245 |
| NA | ENSMUSG00 | 268527 |
| NA | ENSMUSG00 | 108723 |
| NA | ENSMUSG00 | 19024 |
| NA | ENSMUSG00 | 77980 |
| Significant | ENSMUSG00 | 260409 |
| NA | ENSMUSG00 | 234595 |
| NA | ENSMUSG00 | 69942 |
| NA | ENSMUSG00 | 234594 |
| NA | ENSMUSG00 | 381062 |
| NA | ENSMUSG00 | 231147 |
| NA | ENSMUSG00 | 74239 |
| NA | ENSMUSG00 | 243914 |
| NA | ENSMUSG00 | 71474 |
| NA | ENSMUSG00 | 234593 |
| NA | ENSMUSG00 | 78339 |
| NA | ENSMUSG00 | 210982 |
| NA | ENSMUSG00 | 68134 |
| NA | ENSMUSG00 | 229285 |

| | | |
|---|---|---|
| NA | ENSMUSG00 | 71435 |
| NA | ENSMUSG00 | 59031 |
| NA | ENSMUSG00 | 140570 |
| NA | ENSMUSG00 | 67484 |
| NA | ENSMUSG00 | 380752 |
| NA | ENSMUSG00 | 170767 |
| NA | ENSMUSG00 | 103534 |
| NA | ENSMUSG00 | 74772 |
| NA | ENSMUSG00 | 66248 |
| NA | ENSMUSG00 | 17136 |
| NA | ENSMUSG00 | 26373 |
| NA | ENSMUSG00 | 17766 |
| NA | ENSMUSG00 | 76795 |
| NA | ENSMUSG00 | 227619 |
| NA | ENSMUSG00 | 105841 |
| Significant | ENSMUSG00 | 77574 |
| NA | ENSMUSG00 | 320394 |
| NA | ENSMUSG00 | 237500 |
| NA | ENSMUSG00 | 223921 |
| NA | ENSMUSG00 | 212139 |
| NA | ENSMUSG00 | 75416 |
| NA | ENSMUSG00 | 239528 |
| NA | ENSMUSG00 | 12283 |
| NA | ENSMUSG00 | 74256 |
| NA | ENSMUSG00 | 231830 |
| NA | ENSMUSG00 | 22758 |
| NA | ENSMUSG00 | 68035 |
| NA | ENSMUSG00 | 108737 |
| NA | ENSMUSG00 | 73677 |
| NA | ENSMUSG00 | 67116 |
| NA | ENSMUSG00 | 110323 |
| NA | ENSMUSG00 | 227613 |
| NA | ENSMUSG00 | 30045 |
| NA | ENSMUSG00 | 227325 |
| NA | ENSMUSG00 | 209737 |
| NA | ENSMUSG00 | 72404 |
| NA | ENSMUSG00 | 26378 |
| NA | ENSMUSG00 | 68743 |
| NA | ENSMUSG00 | 214627 |
| NA | ENSMUSG00 | 67941 |
| Significant | ENSMUSG00 | 67455 |
| NA | ENSMUSG00 | 109241 |
| NA | ENSMUSG00 | 382030 |
| NA | ENSMUSG00 | 77053 |
| NA | ENSMUSG00 | 194952 |
| NA | ENSMUSG00 | 245847 |

| | | |
|---|---|---|
| NA | ENSMUSG00 | 106021 |
| NA | ENSMUSG00 | 99167 |
| NA | ENSMUSG00 | 241274 |
| NA | ENSMUSG00 | 66340 |
| NA | ENSMUSG00 | 20437 |
| NA | ENSMUSG00 | 75660 |
| NA | ENSMUSG00 | 107733 |
| NA | ENSMUSG00 | 171166 |
| NA | ENSMUSG00 | 243912 |
| NA | ENSMUSG00 | 19208 |
| NA | ENSMUSG00 | 67212 |
| NA | ENSMUSG00 | 233651 |
| NA | ENSMUSG00 | 333193 |
| NA | ENSMUSG00 | 66421 |
| Significant | ENSMUSG00 | 327762 |
| NA | ENSMUSG00 | 102093 |
| NA | ENSMUSG00 | 52538 |
| NA | ENSMUSG00 | 233071 |
| Significant | ENSMUSG00 | 12259 |
| NA | ENSMUSG00 | 227835 |
| NA | ENSMUSG00 | 77683 |
| NA | ENSMUSG00 | 74012 |
| Significant | ENSMUSG00 | 12262 |
| NA | ENSMUSG00 | 72154 |
| Significant | ENSMUSG00 | 12260 |
| NA | ENSMUSG00 | 54445 |
| NA | ENSMUSG00 | 208968 |
| NA | ENSMUSG00 | 225049 |
| NA | ENSMUSG00 | 50878 |
| NA | ENSMUSG00 | 243910 |
| NA | ENSMUSG00 | 26926 |
| NA | ENSMUSG00 | 66714 |
| NA | ENSMUSG00 | 99982 |
| NA | ENSMUSG00 | 114615 |
| NA | ENSMUSG00 | 235442 |
| NA | ENSMUSG00 | 213068 |
| NA | ENSMUSG00 | 231807 |
| NA | ENSMUSG00 | 277468 |
| NA | ENSMUSG00 | 72320 |
| NA | ENSMUSG00 | 320376 |
| NA | ENSMUSG00 | 56631 |
| NA | ENSMUSG00 | 223918 |
| NA | ENSMUSG00 | 66455 |
| NA | ENSMUSG00 | 66343 |
| NA | ENSMUSG00 | 68999 |
| NA | ENSMUSG00 | 21343 |

| | | |
|---|---|---|
| NA | ENSMUSG00 | 224481 |
| NA | ENSMUSG00 | 208884 |
| NA | ENSMUSG00 | 18854 |
| NA | ENSMUSG00 | 55992 |
| NA | ENSMUSG00 | 73945 |
| NA | ENSMUSG00 | 56488 |
| NA | ENSMUSG00 | 22698 |
| NA | ENSMUSG00 | 56314 |
| NA | ENSMUSG00 | 15275 |
| NA | ENSMUSG00 | 268996 |
| NA | ENSMUSG00 | 22697 |
| NA | ENSMUSG00 | 233064 |
| NA | ENSMUSG00 | 109136 |
| NA | ENSMUSG00 | 26465 |
| Significant | ENSMUSG00 | 70423 |
| NA | ENSMUSG00 | 11785 |
| NA | ENSMUSG00 | 20597 |
| NA | ENSMUSG00 | 67869 |
| NA | ENSMUSG00 | 109042 |
| NA | ENSMUSG00 | 54673 |
| NA | ENSMUSG00 | 19656 |
| NA | ENSMUSG00 | 20249 |
| NA | ENSMUSG00 | 93684 |
| NA | ENSMUSG00 | 75841 |
| NA | ENSMUSG00 | 68260 |
| NA | ENSMUSG00 | 73836 |
| NA | ENSMUSG00 NA | |
| NA | ENSMUSG00 | 215445 |
| NA | ENSMUSG00 | 212123 |
| NA | ENSMUSG00 | 56468 |
| NA | ENSMUSG00 | 381678 |
| NA | ENSMUSG00 | 241694 |
| NA | ENSMUSG00 | 73251 |
| NA | ENSMUSG00 | 235344 |
| NA | ENSMUSG00 | 210998 |
| NA | ENSMUSG00 | 73728 |
| NA | ENSMUSG00 | 102182 |
| NA | ENSMUSG00 | 78514 |
| NA | ENSMUSG00 | 104721 |
| NA | ENSMUSG00 | 216011 |
| NA | ENSMUSG00 | 66377 |
| NA | ENSMUSG00 | 67749 |
| NA | ENSMUSG00 | 18109 |
| NA | ENSMUSG00 | 18119 |
| NA | ENSMUSG00 | 243780 |
| NA | ENSMUSG00 | 69257 |

| | | |
|---|---|---|
| NA | ENSMUSG00 | 74127 |
| NA | ENSMUSG00 | 56368 |
| NA | ENSMUSG00 | 76938 |
| NA | ENSMUSG00 | 68118 |
| NA | ENSMUSG00 | 231128 |
| NA | ENSMUSG00 | 24063 |
| NA | ENSMUSG00 | 73754 |
| NA | ENSMUSG00 | 623661 |
| NA | ENSMUSG00 | 68929 |
| NA | ENSMUSG00 | 231125 |
| NA | ENSMUSG00 | 320191 |
| NA | ENSMUSG00 | 17122 |
| NA | ENSMUSG00 | 17184 |
| NA | ENSMUSG00 | 29876 |
| NA | ENSMUSG00 | 103836 |
| NA | ENSMUSG00 | 74653 |
| NA | ENSMUSG00 | 240396 |
| NA | ENSMUSG00 | 66939 |
| Significant | ENSMUSG00 | 52120 |
| NA | ENSMUSG00 | 16588 |
| NA | ENSMUSG00 | 27981 |
| NA | ENSMUSG00 | 229227 |
| NA | ENSMUSG00 | 216766 |
| NA | ENSMUSG00 | 69071 |
| Significant | ENSMUSG00 | 207839 |
| NA | ENSMUSG00 | 20842 |
| NA | ENSMUSG00 | 272589 |
| NA | ENSMUSG00 | 100017 |
| NA | ENSMUSG00 | 67207 |
| NA | ENSMUSG00 | 234875 |
| NA | ENSMUSG00 | 230815 |
| NA | ENSMUSG00 | 21335 |
| NA | ENSMUSG00 | 382207 |
| NA | ENSMUSG00 | 67384 |
| NA | ENSMUSG00 | 215243 |
| NA | ENSMUSG00 | 21354 |
| NA | ENSMUSG00 | 71492 |
| NA | ENSMUSG00 | 50817 |
| NA | ENSMUSG00 | 73158 |
| NA | ENSMUSG00 | 15110 |
| Significant | ENSMUSG00 | 26411 |
| NA | ENSMUSG00 | 74504 |
| NA | ENSMUSG00 | 236727 |
| NA | ENSMUSG00 | 319944 |
| NA | ENSMUSG00 | 83704 |
| NA | ENSMUSG00 | 60322 |

| | | |
|---|---|---|
| NA | ENSMUSG00 | 71904 |
| NA | ENSMUSG00 | 68323 |
| NA | ENSMUSG00 | 226977 |
| NA | ENSMUSG00 | 68614 |
| NA | ENSMUSG00 | 71101 |
| NA | ENSMUSG00 | 66055 |
| NA | ENSMUSG00 | 270035 |
| NA | ENSMUSG00 | 83701 |
| NA | ENSMUSG00 | 100163 |
| NA | ENSMUSG00 | 22289 |
| NA | ENSMUSG00 | 18605 |
| NA | ENSMUSG00 | 13016 |
| NA | ENSMUSG00 | 226861 |
| NA | ENSMUSG00 | 66926 |
| NA | ENSMUSG00 | 116838 |
| NA | ENSMUSG00 | 214742 |
| NA | ENSMUSG00 | 76295 |
| NA | ENSMUSG00 | 15894 |
| NA | ENSMUSG00 | 330723 |
| NA | ENSMUSG00 | 94220 |
| NA | ENSMUSG00 | 67016 |
| NA | ENSMUSG00 | 18787 |
| NA | ENSMUSG00 | 74334 |
| NA | ENSMUSG00 | 240283 |
| NA | ENSMUSG00 | 24115 |
| NA | ENSMUSG00 | 71946 |
| NA | ENSMUSG00 | 277854 |
| NA | ENSMUSG00 | 100534273 |
| NA | ENSMUSG00 | 22782 |
| NA | ENSMUSG00 | 70012 |
| NA | ENSMUSG00 | 214855 |
| NA | ENSMUSG00 | 76306 |
| NA | ENSMUSG00 | 54375 |
| NA | ENSMUSG00 | 77065 |
| NA | ENSMUSG00 | 21847 |
| NA | ENSMUSG00 | 104732 |
| NA | ENSMUSG00 | 320011 |
| NA | ENSMUSG00 | 76843 |
| NA | ENSMUSG00 | 331401 |
| NA | ENSMUSG00 | 75302 |
| NA | ENSMUSG00 | 70790 |
| NA | ENSMUSG00 | 56506 |
| NA | ENSMUSG00 | 66208 |
| NA | ENSMUSG00 | 214469 |
| NA | ENSMUSG00 | 226970 |
| NA | ENSMUSG00 | 74450 |

| | | |
|---|---|---|
| NA | ENSMUSG00 | 233977 |
| NA | ENSMUSG00 | 228607 |
| NA | ENSMUSG00 | 75284 |
| NA | ENSMUSG00 | 100504663 |
| NA | ENSMUSG00 | 74600 |
| NA | ENSMUSG00 | 192786 |
| NA | ENSMUSG00 | 78938 |
| NA | ENSMUSG00 | 218977 |
| NA | ENSMUSG00 | 15000 |
| NA | ENSMUSG00 | 101497 |
| NA | ENSMUSG00 | 503610 |
| NA | ENSMUSG00 | 20055 |
| NA | ENSMUSG00 | 226841 |
| NA | ENSMUSG00 | 51812 |
| NA | ENSMUSG00 | 218973 |
| NA | ENSMUSG00 | 22057 |
| NA | ENSMUSG00 | 14528 |
| NA | ENSMUSG00 | 23957 |
| NA | ENSMUSG00 | 217887 |
| NA | ENSMUSG00 | 18102 |
| NA | ENSMUSG00 | 319387 |
| NA | ENSMUSG00 | 72567 |
| NA | ENSMUSG00 | 79201 |
| NA | ENSMUSG00 | 26942 |
| NA | ENSMUSG00 | 230796 |
| NA | ENSMUSG00 | 72391 |
| NA | ENSMUSG00 | 194744 |
| NA | ENSMUSG00 | 382639 |
| NA | ENSMUSG00 | 22718 |
| NA | ENSMUSG00 | 18759 |
| NA | ENSMUSG00 | 666173 |
| Significant | ENSMUSG00 | 14998 |
| NA | ENSMUSG00 | 241915 |
| NA | ENSMUSG00 | 20516 |
| NA | ENSMUSG00 | 71862 |
| NA | ENSMUSG00 | 12577 |
| NA | ENSMUSG00 | 68832 |
| NA | ENSMUSG00 | 319758 |
| NA | ENSMUSG00 | 70435 |
| NA | ENSMUSG00 | 70882 |
| NA | ENSMUSG00 | 11980 |
| NA | ENSMUSG00 | 230793 |
| NA | ENSMUSG00 | 114874 |
| NA | ENSMUSG00 | 12520 |
| NA | ENSMUSG00 | 58909 |
| NA | ENSMUSG00 | 52637 |

| | | |
|---|---|---|
| NA | ENSMUSG00 | 218952 |
| NA | ENSMUSG00 | 67878 |
| NA | ENSMUSG00 | 54342 |
| NA | ENSMUSG00 | 80986 |
| NA | ENSMUSG00 | 80732 |
| NA | ENSMUSG00 | 230787 |
| NA | ENSMUSG00 | 77569 |
| NA | ENSMUSG00 | 99045 |
| NA | ENSMUSG00 | 13627 |
| NA | ENSMUSG00 | 216971 |
| NA | ENSMUSG00 | 381560 |
| NA | ENSMUSG00 | 228852 |
| Significant | ENSMUSG00 | 19217 |
| NA | ENSMUSG00 | 109275 |
| NA | ENSMUSG00 | 19935 |
| NA | ENSMUSG00 | 319513 |
| NA | ENSMUSG00 | 72507 |
| NA | ENSMUSG00 | 68020 |
| NA | ENSMUSG00 | 232023 |
| NA | ENSMUSG00 | 268448 |
| NA | ENSMUSG00 | 333789 |
| NA | ENSMUSG00 | 19896 |
| NA | ENSMUSG00 | 72826 |
| NA | ENSMUSG00 | 12385 |
| NA | ENSMUSG00 | 109082 |
| NA | ENSMUSG00 | 269423 |
| Significant | ENSMUSG00 | 21817 |
| NA | ENSMUSG00 | 68552 |
| NA | ENSMUSG00 | 52588 |
| NA | ENSMUSG00 | 243382 |
| NA | ENSMUSG00 | 382137 |
| NA | ENSMUSG00 | 170799 |
| Significant | ENSMUSG00 | 225994 |
| NA | ENSMUSG00 | 105148 |
| NA | ENSMUSG00 | 12876 |
| NA | ENSMUSG00 | 68564 |
| NA | ENSMUSG00 | 383619 |
| NA | ENSMUSG00 | 13654 |
| NA | ENSMUSG00 | 108829 |
| NA | ENSMUSG00 | 67333 |
| NA | ENSMUSG00 | 18218 |
| NA | ENSMUSG00 | 213081 |
| NA | ENSMUSG00 | 73173 |
| NA | ENSMUSG00 | 51788 |
| NA | ENSMUSG00 | 217864 |
| NA | ENSMUSG00 | 19261 |

| | | |
|---|---|---|
| NA | ENSMUSG00 | 74251 |
| Significant | ENSMUSG00 | 76809 |
| NA | ENSMUSG00 | 268445 |
| NA | ENSMUSG00 | 243025 |
| NA | ENSMUSG00 | 17995 |
| NA | ENSMUSG00 | 234311 |
| NA | ENSMUSG00 | 242248 |
| NA | ENSMUSG00 | 237860 |
| NA | ENSMUSG00 | 76895 |
| NA | ENSMUSG00 | 57376 |
| NA | ENSMUSG00 | 20778 |
| NA | ENSMUSG00 | 66170 |
| NA | ENSMUSG00 | 234515 |
| NA | ENSMUSG00 | 30938 |
| NA | ENSMUSG00 | 102566 |
| NA | ENSMUSG00 | 69641 |
| NA | ENSMUSG00 | 237859 |
| NA | ENSMUSG00 | 68480 |
| NA | ENSMUSG00 | 106205 |
| NA | ENSMUSG00 | 18081 |
| NA | ENSMUSG00 | 68721 |
| NA | ENSMUSG00 | 20621 |
| NA | ENSMUSG00 | 223970 |
| NA | ENSMUSG00 | 19401 |
| NA | ENSMUSG00 | 64340 |
| NA | ENSMUSG00 | 93842 |
| NA | ENSMUSG00 | 101187 |
| NA | ENSMUSG00 | 212285 |
| NA | ENSMUSG00 | 497652 |
| NA | ENSMUSG00 | 57354 |
| NA | ENSMUSG00 | 72612 |
| NA | ENSMUSG00 | 230379 |
| NA | ENSMUSG00 | 76138 |
| NA | ENSMUSG00 | 68524 |
| NA | ENSMUSG00 | 218236 |
| NA | ENSMUSG00 | 330323 |
| NA | ENSMUSG00 | 21871 |
| NA | ENSMUSG00 | 329877 |
| NA | ENSMUSG00 | 18676 |
| NA | ENSMUSG00 | 319801 |
| NA | ENSMUSG00 | 140559 |
| NA | ENSMUSG00 | 12703 |
| NA | ENSMUSG00 | 70297 |
| NA | ENSMUSG00 | 73671 |
| NA | ENSMUSG00 | 67390 |
| NA | ENSMUSG00 | 230376 |

| | | |
|---|---|---|
| NA | ENSMUSG00 | 58239 |
| NA | ENSMUSG00 | 231051 |
| NA | ENSMUSG00 | 107607 |
| NA | ENSMUSG00 | 106878 |
| NA | ENSMUSG00 | 68235 |
| NA | ENSMUSG00 | 12985 |
| NA | ENSMUSG00 | 218215 |
| NA | ENSMUSG00 | 70925 |
| NA | ENSMUSG00 | 338349 |
| NA | ENSMUSG00 | 231050 |
| Significant | ENSMUSG00 | 231997 |
| NA | ENSMUSG00 | 381259 |
| NA | ENSMUSG00 | 218214 |
| NA | ENSMUSG00 | 74143 |
| NA | ENSMUSG00 | 70873 |
| NA | ENSMUSG00 | 243272 |
| NA | ENSMUSG00 | 320714 |
| NA | ENSMUSG00 | 228829 |
| NA | ENSMUSG00 | 108654 |
| NA | ENSMUSG00 | 544696 |
| NA | ENSMUSG00 | 269702 |
| NA | ENSMUSG00 | 67501 |
| NA | ENSMUSG00 | 77106 |
| NA | ENSMUSG00 | 71069 |
| NA | ENSMUSG00 | 20623 |
| NA | ENSMUSG00 | 12523 |
| NA | ENSMUSG00 | 66612 |
| NA | ENSMUSG00 | 12142 |
| Significant | ENSMUSG00 | 14869 |
| NA | ENSMUSG00 | 233744 |
| NA | ENSMUSG00 | 11793 |
| Significant | ENSMUSG00 | 83679 |
| NA | ENSMUSG00 | 69863 |
| NA | ENSMUSG00 | 320981 |
| NA | ENSMUSG00 | 213056 |
| NA | ENSMUSG00 | 218203 |
| NA | ENSMUSG00 | 215113 |
| NA | ENSMUSG00 | 245865 |
| NA | ENSMUSG00 | 100910 |
| NA | ENSMUSG00 | 68815 |
| NA | ENSMUSG00 | 380713 |
| NA | ENSMUSG00 | 280408 |
| NA | ENSMUSG00 | 117590 |
| NA | ENSMUSG00 | 108097 |
| NA | ENSMUSG00 | 50753 |
| NA | ENSMUSG00 | 320655 |

| | | |
|---|---|---|
| NA | ENSMUSG00 | 66631 |
| NA | ENSMUSG00 | 213233 |
| NA | ENSMUSG00 | 331623 |
| NA | ENSMUSG00 | 382010 |
| NA | ENSMUSG00 | 380712 |
| NA | ENSMUSG00 | 408022 |
| NA | ENSMUSG00 | 15405 |
| NA | ENSMUSG00 | 16456 |
| NA | ENSMUSG00 | 71365 |
| NA | ENSMUSG00 | 16328 |
| NA | ENSMUSG00 | 320878 |
| NA | ENSMUSG00 | 74841 |
| NA | ENSMUSG00 | 68298 |
| NA | ENSMUSG00 | 77578 |
| NA | ENSMUSG00 | 68667 |
| NA | ENSMUSG00 | 73102 |
| NA | ENSMUSG00 | 246257 |
| NA | ENSMUSG00 | 320678 |
| NA | ENSMUSG00 | 14109 |
| NA | ENSMUSG00 | 110109 |
| NA | ENSMUSG00 | 14628 |
| NA | ENSMUSG00 | 68021 |
| NA | ENSMUSG00 | 103677 |
| NA | ENSMUSG00 | 102141 |
| NA | ENSMUSG00 | 213948 |
| NA | ENSMUSG00 | 233733 |
| NA | ENSMUSG00 | 225348 |
| NA | ENSMUSG00 | 71982 |
| NA | ENSMUSG00 | 215951 |
| NA | ENSMUSG00 | 67198 |
| NA | ENSMUSG00 | 108687 |
| NA | ENSMUSG00 | 73467 |
| NA | ENSMUSG00 | 56407 |
| Significant | ENSMUSG00 | 140742 |
| NA | ENSMUSG00 | 104662 |
| NA | ENSMUSG00 | 208104 |
| NA | ENSMUSG00 | 353170 |
| NA | ENSMUSG00 | 269800 |
| NA | ENSMUSG00 | 97761 |
| NA | ENSMUSG00 | 217151 |
| NA | ENSMUSG00 | 66822 |
| NA | ENSMUSG00 | 16796 |
| NA | ENSMUSG00 | 230393 |
| NA | ENSMUSG00 | 56406 |
| NA | ENSMUSG00 | 66425 |
| NA | ENSMUSG00 | 319934 |

| | | |
|---|---|---|
| NA | ENSMUSG00 | 218138 |
| NA | ENSMUSG00 | 60365 |
| NA | ENSMUSG00 | 68728 |
| NA | ENSMUSG00 | 22137 |
| NA | ENSMUSG00 | 228812 |
| NA | ENSMUSG00 | 208043 |
| NA | ENSMUSG00 | 20130 |
| NA | ENSMUSG00 | 56524 |
| NA | ENSMUSG00 | 56338 |
| NA | ENSMUSG00 | 67031 |
| NA | ENSMUSG00 | 65112 |
| NA | ENSMUSG00 | 233208 |
| NA | ENSMUSG00 | 56295 |
| NA | ENSMUSG00 | 69957 |
| NA | ENSMUSG00 | 103199 |
| NA | ENSMUSG00 | 13653 |
| NA | ENSMUSG00 | 72748 |
| NA | ENSMUSG00 | 26447 |
| NA | ENSMUSG00 | 22225 |
| NA | ENSMUSG00 | 246198 |
| NA | ENSMUSG00 | 71713 |
| NA | ENSMUSG00 | 14794 |
| NA | ENSMUSG00 | 70178 |
| NA | ENSMUSG00 | 66694 |
| NA | ENSMUSG00 | 320078 |
| NA | ENSMUSG00 | 75608 |
| NA | ENSMUSG00 | 70729 |
| NA | ENSMUSG00 | 78334 |
| NA | ENSMUSG00 | 21781 |
| NA | ENSMUSG00 | 192157 |
| NA | ENSMUSG00 | 64051 |
| NA | ENSMUSG00 | 66491 |
| NA | ENSMUSG00 | 229603 |
| NA | ENSMUSG00 | 234076 |
| NA | ENSMUSG00 | 75210 |
| NA | ENSMUSG00 | 84113 |
| NA | ENSMUSG00 | 67943 |
| NA | ENSMUSG00 | 102323 |
| NA | ENSMUSG00 | 243771 |
| NA | ENSMUSG00 | 23886 |
| NA | ENSMUSG00 | 67239 |
| Significant | ENSMUSG00 | 66790 |
| NA | ENSMUSG00 | 73174 |
| NA | ENSMUSG00 | 16468 |
| NA | ENSMUSG00 | 233204 |
| NA | ENSMUSG00 | 319262 |

| | | |
|---|---|---|
| NA | ENSMUSG00 | 67211 |
| NA | ENSMUSG00 | 23831 |
| NA | ENSMUSG00 | 232371 |
| NA | ENSMUSG00 | 12396 |
| NA | ENSMUSG00 | 71240 |
| NA | ENSMUSG00 | 235469 |
| NA | ENSMUSG00 | 320538 |
| NA | ENSMUSG00 | 107503 |
| NA | ENSMUSG00 | 233424 |
| NA | ENSMUSG00 | 234069 |
| NA | ENSMUSG00 | 229600 |
| NA | ENSMUSG00 | 66209 |
| NA | ENSMUSG00 | 66515 |
| NA | ENSMUSG00 | 56705 |
| NA | ENSMUSG00 | 229599 |
| NA | ENSMUSG00 | 64339 |
| NA | ENSMUSG00 | 101592 |
| NA | ENSMUSG00 | 67661 |
| NA | ENSMUSG00 | 231724 |
| NA | ENSMUSG00 | 330577 |
| NA | ENSMUSG00 | 634731 |
| NA | ENSMUSG00 | 320717 |
| NA | ENSMUSG00 | 83397 |
| NA | ENSMUSG00 | 654470 |
| NA | ENSMUSG00 | 100038725 |
| NA | ENSMUSG00 | 75687 |
| NA | ENSMUSG00 | 229011 |
| NA | ENSMUSG00 | 14700 |
| NA | ENSMUSG00 | 210293 |
| NA | ENSMUSG00 | 101471 |
| NA | ENSMUSG00 | 17210 |
| NA | ENSMUSG00 | 18023 |
| NA | ENSMUSG00 | 66985 |
| NA | ENSMUSG00 | 56205 |
| NA | ENSMUSG00 | 69790 |
| NA | ENSMUSG00 | 107769 |
| NA | ENSMUSG00 | 67005 |
| NA | ENSMUSG00 | 13244 |
| NA | ENSMUSG00 | 70552 |
| NA | ENSMUSG00 | 13040 |
| NA | ENSMUSG00 | 18971 |
| NA | ENSMUSG00 | 67148 |
| NA | ENSMUSG00 | 208647 |
| NA | ENSMUSG00 | 107702 |
| NA | ENSMUSG00 | 226089 |
| NA | ENSMUSG00 | 235439 |

| | | |
|---|---|---|
| NA | ENSMUSG00 | 14745 |
| NA | ENSMUSG00 | 76688 |
| NA | ENSMUSG00 | 83925 |
| NA | ENSMUSG00 | 68083 |
| NA | ENSMUSG00 | 269400 |
| NA | ENSMUSG00 | 57423 |
| NA | ENSMUSG00 | 66124 |
| NA | ENSMUSG00 | 227743 |
| NA | ENSMUSG00 | 102162 |
| NA | ENSMUSG00 | 319901 |
| NA | ENSMUSG00 | 229004 |
| NA | ENSMUSG00 | 54214 |
| NA | ENSMUSG00 | 75007 |
| NA | ENSMUSG00 | 27425 |
| NA | ENSMUSG00 | 18516 |
| NA | ENSMUSG00 | 231889 |
| NA | ENSMUSG00 | 218121 |
| NA | ENSMUSG00 | 226757 |
| NA | ENSMUSG00 | 67429 |
| NA | ENSMUSG00 | 72543 |
| NA | ENSMUSG00 | 70726 |
| NA | ENSMUSG00 | 70699 |
| NA | ENSMUSG00 | 224742 |
| NA | ENSMUSG00 | 116904 |
| NA | ENSMUSG00 | 213054 |
| NA | ENSMUSG00 | 277250 |
| NA | ENSMUSG00 | 77987 |
| NA | ENSMUSG00 | 22351 |
| NA | ENSMUSG00 | 13849 |
| NA | ENSMUSG00 | 75788 |
| NA | ENSMUSG00 | 106766 |
| NA | ENSMUSG00 | 70004 |
| NA | ENSMUSG00 | 53621 |
| NA | ENSMUSG00 | 13590 |
| NA | ENSMUSG00 | 67695 |
| NA | ENSMUSG00 | 208768 |
| NA | ENSMUSG00 | 380711 |
| NA | ENSMUSG00 | 14710 |
| NA | ENSMUSG00 | 67674 |
| NA | ENSMUSG00 | 54366 |
| NA | ENSMUSG00 | 209462 |
| NA | ENSMUSG00 | 230234 |
| NA | ENSMUSG00 | 68796 |
| NA | ENSMUSG00 | 241308 |
| NA | ENSMUSG00 | 76223 |
| NA | ENSMUSG00 | 68915 |

| | | |
|---|---|---|
| NA | ENSMUSG00 | 14537 |
| NA | ENSMUSG00 | 16558 |
| NA | ENSMUSG00 | 18673 |
| NA | ENSMUSG00 | 228994 |
| NA | ENSMUSG00 | 320404 |
| NA | ENSMUSG00 | 66898 |
| NA | ENSMUSG00 | 99326 |
| NA | ENSMUSG00 | 107650 |
| NA | ENSMUSG00 | 227449 |
| NA | ENSMUSG00 | 12183 |
| NA | ENSMUSG00 | 11906 |
| NA | ENSMUSG00 | 68031 |
| NA | ENSMUSG00 | 79044 |
| NA | ENSMUSG00 | 319278 |
| NA | ENSMUSG00 | 140481 |
| NA | ENSMUSG00 | 233189 |
| NA | ENSMUSG00 | 215512 |
| NA | ENSMUSG00 | 384783 |
| NA | ENSMUSG00 | 319601 |
| NA | ENSMUSG00 | 269261 |
| NA | ENSMUSG00 | 229584 |
| NA | ENSMUSG00 | 217127 |
| NA | ENSMUSG00 | 224860 |
| NA | ENSMUSG00 | 23856 |
| NA | ENSMUSG00 | 269955 |
| Significant | ENSMUSG00 | 109232 |
| NA | ENSMUSG00 | 233406 |
| NA | ENSMUSG00 | 226744 |
| NA | ENSMUSG00 | 109115 |
| NA | ENSMUSG00 | 353190 |
| Significant | ENSMUSG00 | 108115 |
| NA | ENSMUSG00 | 22195 |
| NA | ENSMUSG00 | 18604 |
| NA | ENSMUSG00 | 231876 |
| NA | ENSMUSG00 | 19352 |
| NA | ENSMUSG00 | 217124 |
| NA | ENSMUSG00 | 17828 |
| NA | ENSMUSG00 | 252966 |
| NA | ENSMUSG00 | 105245 |
| NA | ENSMUSG00 | 100763 |
| NA | ENSMUSG00 | 66481 |
| NA | ENSMUSG00 | 21898 |
| NA | ENSMUSG00 | 54381 |
| Significant | ENSMUSG00 | 233186 |
| NA | ENSMUSG00 | 30057 |
| NA | ENSMUSG00 | 212508 |

| | | |
|---|---|---|
| Significant | ENSMUSG00 | 73910 |
| NA | ENSMUSG00 | 75812 |
| NA | ENSMUSG00 | 56436 |
| NA | ENSMUSG00 | 70420 |
| NA | ENSMUSG00 | 98910 |
| NA | ENSMUSG00 | 329777 |
| NA | ENSMUSG00 | 235169 |
| NA | ENSMUSG00 | 228983 |
| NA | ENSMUSG00 | 268465 |
| NA | ENSMUSG00 | 16790 |
| NA | ENSMUSG00 | 68073 |
| NA | ENSMUSG00 | 17463 |
| NA | ENSMUSG00 | 108946 |
| NA | ENSMUSG00 | 52856 |
| NA | ENSMUSG00 | 12643 |
| NA | ENSMUSG00 | 269397 |
| NA | ENSMUSG00 | 68750 |
| NA | ENSMUSG00 | 237339 |
| NA | ENSMUSG00 | 237926 |
| NA | ENSMUSG00 | 223455 |
| NA | ENSMUSG00 | 66290 |
| NA | ENSMUSG00 | 241846 |
| Significant | ENSMUSG00 | 74145 |
| NA | ENSMUSG00 | 104601 |
| NA | ENSMUSG00 | 68404 |
| Significant | ENSMUSG00 | 104099 |
| NA | ENSMUSG00 | 228980 |
| Significant | ENSMUSG00 | 353211 |
| NA | ENSMUSG00 | 98828 |
| NA | ENSMUSG00 | 232679 |
| NA | ENSMUSG00 | 227541 |
| NA | ENSMUSG00 | 20227 |
| NA | ENSMUSG00 | 12393 |
| NA | ENSMUSG00 | 116873 |
| NA | ENSMUSG00 | 98952 |
| NA | ENSMUSG00 | 100182 |
| NA | ENSMUSG00 | 22214 |
| NA | ENSMUSG00 | 67899 |
| NA | ENSMUSG00 | 71254 |
| NA | ENSMUSG00 | 432442 |
| NA | ENSMUSG00 | 170757 |
| NA | ENSMUSG00 | 223453 |
| NA | ENSMUSG00 | 18975 |
| NA | ENSMUSG00 | 67249 |
| NA | ENSMUSG00 | 105351 |
| NA | ENSMUSG00 | 26426 |

| | | |
|---|---|---|
| NA | ENSMUSG00 | 208836 |
| NA | ENSMUSG00 | 19664 |
| NA | ENSMUSG00 | 268973 |
| NA | ENSMUSG00 | 73737 |
| NA | ENSMUSG00 | 11534 |
| NA | ENSMUSG00 | 70796 |
| NA | ENSMUSG00 | 75329 |
| NA | ENSMUSG00 | 209815 |
| NA | ENSMUSG00 | 54608 |
| NA | ENSMUSG00 | 68379 |
| NA | ENSMUSG00 | 231871 |
| NA | ENSMUSG00 | 210029 |
| NA | ENSMUSG00 | 68172 |
| NA | ENSMUSG00 | 67769 |
| Significant | ENSMUSG00 | 16173 |
| NA | ENSMUSG00 | 75956 |
| NA | ENSMUSG00 | 94246 |
| NA | ENSMUSG00 | 52040 |
| NA | ENSMUSG00 | 68028 |
| NA | ENSMUSG00 | 108903 |
| NA | ENSMUSG00 | 20937 |
| NA | ENSMUSG00 | 74732 |
| NA | ENSMUSG00 | 70430 |
| NA | ENSMUSG00 | 69608 |
| NA | ENSMUSG00 | 57444 |
| NA | ENSMUSG00 | 21808 |
| NA | ENSMUSG00 | 97884 |
| NA | ENSMUSG00 | 231868 |
| NA | ENSMUSG00 | 226791 |
| NA | ENSMUSG00 | 238024 |
| NA | ENSMUSG00 | 99011 |
| NA | ENSMUSG00 | 227723 |
| NA | ENSMUSG00 | 228961 |
| NA | ENSMUSG00 | 230316 |
| NA | ENSMUSG00 | 68837 |
| NA | ENSMUSG00 | 27215 |
| NA | ENSMUSG00 | 72007 |
| NA | ENSMUSG00 | 217370 |
| NA | ENSMUSG00 | 214444 |
| NA | ENSMUSG00 | 238023 |
| NA | ENSMUSG00 | 17423 |
| NA | ENSMUSG00 | 76438 |
| NA | ENSMUSG00 | 98732 |
| NA | ENSMUSG00 | 16008 |
| NA | ENSMUSG00 | 68867 |
| NA | ENSMUSG00 | 239706 |

| | | |
|---|---|---|
| NA | ENSMUSG00 | 76252 |
| NA | ENSMUSG00 | 54380 |
| NA | ENSMUSG00 | 227715 |
| NA | ENSMUSG00 | 73068 |
| NA | ENSMUSG00 | 233489 |
| NA | ENSMUSG00 | 218811 |
| NA | ENSMUSG00 | 15284 |
| NA | ENSMUSG00 | 54636 |
| NA | ENSMUSG00 | 63953 |
| NA | ENSMUSG00 | 381269 |
| NA | ENSMUSG00 | 234258 |
| NA | ENSMUSG00 | 76453 |
| NA | ENSMUSG00 | 70673 |
| NA | ENSMUSG00 | 320473 |
| NA | ENSMUSG00 | 208211 |
| NA | ENSMUSG00 | 72759 |
| NA | ENSMUSG00 | 54384 |
| NA | ENSMUSG00 | 67229 |
| NA | ENSMUSG00 | 67880 |
| NA | ENSMUSG00 | 338467 |
| NA | ENSMUSG00 | 268783 |
| NA | ENSMUSG00 | 102791 |
| NA | ENSMUSG00 | 77031 |
| NA | ENSMUSG00 | 70546 |
| NA | ENSMUSG00 | 170644 |
| NA | ENSMUSG00 | 22393 |
| NA | ENSMUSG00 | 231861 |
| NA | ENSMUSG00 | 78506 |
| NA | ENSMUSG00 | 319638 |
| NA | ENSMUSG00 | 72323 |
| NA | ENSMUSG00 | 72480 |
| NA | ENSMUSG00 | 212898 |
| NA | ENSMUSG00 | 98999 |
| NA | ENSMUSG00 | 234865 |
| NA | ENSMUSG00 | 224648 |
| NA | ENSMUSG00 | 110854 |
| NA | ENSMUSG00 | 230967 |
| NA | ENSMUSG00 | NA |
| NA | ENSMUSG00 | 72580 |
| NA | ENSMUSG00 | 20853 |
| NA | ENSMUSG00 | 17967 |
| NA | ENSMUSG00 | 212892 |
| NA | ENSMUSG00 | 207740 |
| NA | ENSMUSG00 | 260305 |
| NA | ENSMUSG00 | 270163 |
| NA | ENSMUSG00 | 105428 |

| | | |
|---|---|---|
| NA | ENSMUSG00 | 67063 |
| NA | ENSMUSG00 | 56424 |
| NA | ENSMUSG00 NA | |
| NA | ENSMUSG00 | 277360 |
| NA | ENSMUSG00 | 231855 |
| NA | ENSMUSG00 | 51810 |
| NA | ENSMUSG00 | 244421 |
| NA | ENSMUSG00 | 230162 |
| NA | ENSMUSG00 | 78885 |
| NA | ENSMUSG00 | 56282 |
| NA | ENSMUSG00 | 246154 |
| NA | ENSMUSG00 | 70266 |
| NA | ENSMUSG00 | 208922 |
| NA | ENSMUSG00 | 20182 |
| NA | ENSMUSG00 | 227695 |
| NA | ENSMUSG00 | 57295 |
| NA | ENSMUSG00 | 66431 |
| NA | ENSMUSG00 | 228880 |
| NA | ENSMUSG00 | 70296 |
| NA | ENSMUSG00 | 121022 |
| NA | ENSMUSG00 | 66988 |
| NA | ENSMUSG00 | 227693 |
| NA | ENSMUSG00 | 66665 |
| NA | ENSMUSG00 | 211329 |
| NA | ENSMUSG00 | 99526 |
| NA | ENSMUSG00 | 217365 |
| NA | ENSMUSG00 | 320506 |
| NA | ENSMUSG00 | 16826 |
| NA | ENSMUSG00 | 269608 |
| NA | ENSMUSG00 | 71820 |
| NA | ENSMUSG00 | 208869 |
| NA | ENSMUSG00 | 381406 |
| NA | ENSMUSG00 | 214459 |
| NA | ENSMUSG00 | 66801 |
| NA | ENSMUSG00 | 52864 |
| NA | ENSMUSG00 | 56737 |
| NA | ENSMUSG00 | 268515 |
| NA | ENSMUSG00 | 53415 |
| NA | ENSMUSG00 | 269717 |
| NA | ENSMUSG00 | 26909 |
| NA | ENSMUSG00 | 242960 |
| NA | ENSMUSG00 | 72041 |
| NA | ENSMUSG00 | 99480 |
| NA | ENSMUSG00 | 69876 |
| NA | ENSMUSG00 | 246738 |
| NA | ENSMUSG00 | 231214 |

| | | |
|---|---|---|
| NA | ENSMUSG00 | 230935 |
| NA | ENSMUSG00 | 383295 |
| NA | ENSMUSG00 | 20022 |
| NA | ENSMUSG00 | 230145 |
| NA | ENSMUSG00 | 12009 |
| NA | ENSMUSG00 | 231207 |
| NA | ENSMUSG00 | 98256 |
| NA | ENSMUSG00 | 99151 |
| NA | ENSMUSG00 | 71063 |
| NA | ENSMUSG00 | 66361 |
| NA | ENSMUSG00 | 73692 |
| NA | ENSMUSG00 | 228869 |
| NA | ENSMUSG00 | 103284 |
| NA | ENSMUSG00 | 381605 |
| NA | ENSMUSG00 | 319581 |
| NA | ENSMUSG00 | 227682 |
| NA | ENSMUSG00 | 545085 |
| NA | ENSMUSG00 | 50913 |
| Significant | ENSMUSG00 | 214137 |
| NA | ENSMUSG00 | 329559 |
| NA | ENSMUSG00 | 215819 |
| NA | ENSMUSG00 | 100502841 |
| NA | ENSMUSG00 | 627049 |
| NA | ENSMUSG00 | 244329 |
| NA | ENSMUSG00 | 107746 |
| NA | ENSMUSG00 | 228866 |
| NA | ENSMUSG00 | 338371 |
| NA | ENSMUSG00 | 68703 |
| NA | ENSMUSG00 | 74735 |
| NA | ENSMUSG00 | 415115 |
| NA | ENSMUSG00 | 380629 |
| NA | ENSMUSG00 | 215210 |
| NA | ENSMUSG00 | 66789 |
| Significant | ENSMUSG00 | 378431 |
| Significant | ENSMUSG00 | 14190 |
| NA | ENSMUSG00 | 71617 |
| NA | ENSMUSG00 | 268512 |
| NA | ENSMUSG00 | 17684 |
| NA | ENSMUSG00 | 74646 |
| NA | ENSMUSG00 | 210582 |
| NA | ENSMUSG00 | 215160 |
| NA | ENSMUSG00 | 207932 |
| NA | ENSMUSG00 | 212167 |
| NA | ENSMUSG00 | 18707 |
| NA | ENSMUSG00 | 19219 |
| NA | ENSMUSG00 | 18798 |

| | | |
|---|---|---|
| NA | ENSMUSG00 | 13138 |
| NA | ENSMUSG00 | 65945 |
| NA | ENSMUSG00 | 320204 |
| NA | ENSMUSG00 | 215114 |
| NA | ENSMUSG00 | 207607 |
| NA | ENSMUSG00 | 30046 |
| NA | ENSMUSG00 | 242860 |
| NA | ENSMUSG00 | 207592 |
| NA | ENSMUSG00 | 237256 |
| NA | ENSMUSG00 | 207521 |
| NA | ENSMUSG00 | 269336 |
| NA | ENSMUSG00 | 56306 |
| NA | ENSMUSG00 | 68770 |
| NA | ENSMUSG00 | 433667 |
| NA | ENSMUSG00 | 12418 |
| NA | ENSMUSG00 | 214764 |
| NA | ENSMUSG00 | 21853 |
| NA | ENSMUSG00 | 15950 |
| NA | ENSMUSG00 | 215751 |
| NA | ENSMUSG00 | 228536 |
| NA | ENSMUSG00 | 20539 |
| NA | ENSMUSG00 | 94244 |
| Significant | ENSMUSG00 | 19218 |
| NA | ENSMUSG00 | 226139 |
| NA | ENSMUSG00 | 16798 |
| NA | ENSMUSG00 | 74998 |
| NA | ENSMUSG00 | 213541 |
| NA | ENSMUSG00 | 15568 |
| NA | ENSMUSG00 | 320727 |
| NA | ENSMUSG00 | 20847 |
| NA | ENSMUSG00 | 69912 |
| NA | ENSMUSG00 | 21821 |
| NA | ENSMUSG00 | 56516 |
| NA | ENSMUSG00 | 50793 |
| NA | ENSMUSG00 | 68342 |
| NA | ENSMUSG00 | 116848 |
| NA | ENSMUSG00 | 18796 |
| NA | ENSMUSG00 NA | |
| NA | ENSMUSG00 | 12236 |
| NA | ENSMUSG00 | 171543 |
| NA | ENSMUSG00 | 224613 |
| NA | ENSMUSG00 | 232539 |
| NA | ENSMUSG00 | 74411 |
| NA | ENSMUSG00 | 235283 |
| NA | ENSMUSG00 | 232536 |
| NA | ENSMUSG00 | 219105 |

| | | |
|---|---|---|
| NA | ENSMUSG00 | 98376 |
| NA | ENSMUSG00 | 108767 |
| NA | ENSMUSG00 | 381413 |
| NA | ENSMUSG00 | 77252 |
| NA | ENSMUSG00 | 109731 |
| NA | ENSMUSG00 | 23908 |
| Significant | ENSMUSG00 | 21825 |
| Significant | ENSMUSG00 | 76281 |
| NA | ENSMUSG00 | 67143 |
| NA | ENSMUSG00 | 69113 |
| NA | ENSMUSG00 | 68277 |
| NA | ENSMUSG00 | 14261 |
| NA | ENSMUSG00 | 140577 |
| NA | ENSMUSG00 | 272322 |
| NA | ENSMUSG00 | 24044 |
| NA | ENSMUSG00 | 211535 |
| NA | ENSMUSG00 | NA |
| NA | ENSMUSG00 | 68026 |
| NA | ENSMUSG00 | 170753 |
| NA | ENSMUSG00 | 13732 |
| NA | ENSMUSG00 | 229905 |
| NA | ENSMUSG00 | 235330 |
| NA | ENSMUSG00 | 104346 |
| NA | ENSMUSG00 | 226562 |
| Significant | ENSMUSG00 | 23890 |
| NA | ENSMUSG00 | 58867 |
| NA | ENSMUSG00 | 67623 |
| Significant | ENSMUSG00 | 66682 |
| NA | ENSMUSG00 | 67529 |
| NA | ENSMUSG00 | 108995 |
| NA | ENSMUSG00 | 16971 |
| NA | ENSMUSG00 | 71177 |
| NA | ENSMUSG00 | 229900 |
| NA | ENSMUSG00 | 234825 |
| NA | ENSMUSG00 | 101476 |
| NA | ENSMUSG00 | 66230 |
| NA | ENSMUSG00 | 12014 |
| NA | ENSMUSG00 | 329470 |
| NA | ENSMUSG00 | 12571 |
| NA | ENSMUSG00 | 399568 |
| NA | ENSMUSG00 | 15213 |
| NA | ENSMUSG00 | 230073 |
| NA | ENSMUSG00 | 226551 |
| NA | ENSMUSG00 | 269623 |
| NA | ENSMUSG00 | 240084 |
| NA | ENSMUSG00 | 228491 |

| | | |
|---|---|---|
| NA | ENSMUSG00 | 229731 |
| NA | ENSMUSG00 NA | |
| NA | ENSMUSG00 | 321006 |
| NA | ENSMUSG00 | 78309 |
| NA | ENSMUSG00 | 67872 |
| NA | ENSMUSG00 | 329739 |
| NA | ENSMUSG00 | 216445 |
| NA | ENSMUSG00 | 70797 |
| NA | ENSMUSG00 | 216443 |
| NA | ENSMUSG00 | 108077 |
| NA | ENSMUSG00 | 67490 |
| NA | ENSMUSG00 | 71458 |
| NA | ENSMUSG00 | 211007 |
| NA | ENSMUSG00 | 67636 |
| NA | ENSMUSG00 | 19302 |
| NA | ENSMUSG00 | 11834 |
| NA | ENSMUSG00 | 19045 |
| NA | ENSMUSG00 | 99512 |
| NA | ENSMUSG00 | 269881 |
| NA | ENSMUSG00 | 68904 |
| NA | ENSMUSG00 | 100986 |
| NA | ENSMUSG00 | 269514 |
| Significant | ENSMUSG00 | 171284 |
| NA | ENSMUSG00 | 246696 |
| NA | ENSMUSG00 | 80904 |
| NA | ENSMUSG00 | 381305 |
| NA | ENSMUSG00 | 69217 |
| NA | ENSMUSG00 | 545725 |
| NA | ENSMUSG00 | 71795 |
| NA | ENSMUSG00 | 245007 |
| NA | ENSMUSG00 | 17872 |
| NA | ENSMUSG00 | 216441 |
| NA | ENSMUSG00 | 225283 |
| NA | ENSMUSG00 | 216892 |
| NA | ENSMUSG00 | 208449 |
| NA | ENSMUSG00 | 77593 |
| NA | ENSMUSG00 | 234023 |
| NA | ENSMUSG00 | 216440 |
| Significant | ENSMUSG00 | 18432 |
| NA | ENSMUSG00 | 207704 |
| Significant | ENSMUSG00 | 233016 |
| NA | ENSMUSG00 | 56187 |
| NA | ENSMUSG00 | 104418 |
| NA | ENSMUSG00 | 207165 |
| NA | ENSMUSG00 | 112403 |
| NA | ENSMUSG00 | 327959 |

| | | |
|---|---|---|
| NA | ENSMUSG00 | 228361 |
| NA | ENSMUSG00 | 52118 |
| NA | ENSMUSG00 | 242362 |
| NA | ENSMUSG00 | 66399 |
| NA | ENSMUSG00 | 170744 |
| NA | ENSMUSG00 | 214812 |
| NA | ENSMUSG00 | 380732 |
| NA | ENSMUSG00 | 68758 |
| NA | ENSMUSG00 | 11496 |
| NA | ENSMUSG00 | 21763 |
| NA | ENSMUSG00 | 75786 |
| NA | ENSMUSG00 | 72201 |
| Significant | ENSMUSG00 | 12267 |
| NA | ENSMUSG00 | 114230 |
| NA | ENSMUSG00 | 79565 |
| NA | ENSMUSG00 | 104082 |
| NA | ENSMUSG00 | 14863 |
| NA | ENSMUSG00 | 235044 |
| NA | ENSMUSG00 | 11812 |
| NA | ENSMUSG00 | 107182 |
| NA | ENSMUSG00 | 237222 |
| NA | ENSMUSG00 | 228356 |
| NA | ENSMUSG00 | 15985 |
| NA | ENSMUSG00 | 66011 |
| NA | ENSMUSG00 | 71592 |
| NA | ENSMUSG00 | 67139 |
| NA | ENSMUSG00 | 71529 |
| NA | ENSMUSG00 | 11810 |
| Significant | ENSMUSG00 | 214359 |
| NA | ENSMUSG00 | 74551 |
| NA | ENSMUSG00 | 216877 |
| NA | ENSMUSG00 | 237221 |
| NA | ENSMUSG00 | 213783 |
| Significant | ENSMUSG00 | 114255 |
| NA | ENSMUSG00 | 227399 |
| NA | ENSMUSG00 | 218850 |
| NA | ENSMUSG00 | 18247 |
| NA | ENSMUSG00 | 76142 |
| NA | ENSMUSG00 | 381101 |
| NA | ENSMUSG00 | 27984 |
| NA | ENSMUSG00 | 81000 |
| NA | ENSMUSG00 | 19069 |
| NA | ENSMUSG00 | 13619 |
| NA | ENSMUSG00 | 270685 |
| NA | ENSMUSG00 | 15312 |
| NA | ENSMUSG00 | 228355 |

| | | |
|---|---|---|
| NA | ENSMUSG00 | 213773 |
| NA | ENSMUSG00 | 11811 |
| NA | ENSMUSG00 | 214063 |
| NA | ENSMUSG00 | 67803 |
| NA | ENSMUSG00 | 11766 |
| Significant | ENSMUSG00 | 20452 |
| NA | ENSMUSG00 | 268396 |
| NA | ENSMUSG00 | 216874 |
| NA | ENSMUSG00 | 433375 |
| NA | ENSMUSG00 | 69994 |
| NA | ENSMUSG00 | 66185 |
| NA | ENSMUSG00 | 239102 |
| NA | ENSMUSG00 | 56807 |
| NA | ENSMUSG00 | 226594 |
| NA | ENSMUSG00 | 232989 |
| NA | ENSMUSG00 | 15209 |
| NA | ENSMUSG00 | 22384 |
| NA | ENSMUSG00 | 13876 |
| NA | ENSMUSG00 | 72656 |
| NA | ENSMUSG00 | 70510 |
| NA | ENSMUSG00 | 12508 |
| NA | ENSMUSG00 | 20438 |
| Significant | ENSMUSG00 | 56743 |
| NA | ENSMUSG00 | 72993 |
| NA | ENSMUSG00 | 56381 |
| NA | ENSMUSG00 | 78372 |
| NA | ENSMUSG00 | 106821 |
| NA | ENSMUSG00 | 99712 |
| NA | ENSMUSG00 | 26374 |
| NA | ENSMUSG00 | 22129 |
| NA | ENSMUSG00 | 230767 |
| NA | ENSMUSG00 | 72205 |
| NA | ENSMUSG00 | 21767 |
| NA | ENSMUSG00 | 211922 |
| NA | ENSMUSG00 | 110948 |
| NA | ENSMUSG00 | 58248 |
| NA | ENSMUSG00 | 107686 |
| NA | ENSMUSG00 | 213491 |
| NA | ENSMUSG00 | 226591 |
| NA | ENSMUSG00 | 108735 |
| NA | ENSMUSG00 | 103554 |
| Significant | ENSMUSG00 | 194590 |
| NA | ENSMUSG00 | 13875 |
| NA | ENSMUSG00 | 100383 |
| NA | ENSMUSG00 | 230872 |
| NA | ENSMUSG00 | 227195 |

| | | |
|---|---|---|
| NA | ENSMUSG00 | 212198 |
| NA | ENSMUSG00 | 235043 |
| NA | ENSMUSG00 | 11692 |
| NA | ENSMUSG00 | NA |
| Significant | ENSMUSG00 | 232975 |
| NA | ENSMUSG00 | 30838 |
| NA | ENSMUSG00 | 116914 |
| NA | ENSMUSG00 | 74648 |
| NA | ENSMUSG00 | 19726 |
| NA | ENSMUSG00 | 209012 |
| NA | ENSMUSG00 | 16801 |
| NA | ENSMUSG00 | 214133 |
| NA | ENSMUSG00 | 108911 |
| NA | ENSMUSG00 | 216864 |
| NA | ENSMUSG00 | 20085 |
| NA | ENSMUSG00 | 63955 |
| NA | ENSMUSG00 | 11890 |
| NA | ENSMUSG00 | 72754 |
| NA | ENSMUSG00 | 626870 |
| NA | ENSMUSG00 | 58194 |
| Significant | ENSMUSG00 | 105501 |
| NA | ENSMUSG00 | 114249 |
| NA | ENSMUSG00 | 67525 |
| NA | ENSMUSG00 | 70031 |
| NA | ENSMUSG00 | 212632 |
| NA | ENSMUSG00 | 66092 |
| NA | ENSMUSG00 | 226162 |
| NA | ENSMUSG00 | 210146 |
| NA | ENSMUSG00 | 72750 |
| NA | ENSMUSG00 | 74087 |
| NA | ENSMUSG00 | 72515 |
| Significant | ENSMUSG00 | 107568 |
| NA | ENSMUSG00 | 208084 |
| NA | ENSMUSG00 | 14369 |
| NA | ENSMUSG00 | 53858 |
| NA | ENSMUSG00 | 66357 |
| NA | ENSMUSG00 | 52808 |
| NA | ENSMUSG00 | 140580 |
| NA | ENSMUSG00 | 245666 |
| NA | ENSMUSG00 | 434130 |
| NA | ENSMUSG00 | 18585 |
| NA | ENSMUSG00 | 17965 |
| NA | ENSMUSG00 | 78100 |
| NA | ENSMUSG00 | 77605 |
| NA | ENSMUSG00 | 213753 |
| NA | ENSMUSG00 | 100637 |

| | | |
|---|---|---|
| NA | ENSMUSG00 | 24061 |
| NA | ENSMUSG00 | 192656 |
| NA | ENSMUSG00 | 77056 |
| NA | ENSMUSG00 | 12190 |
| NA | ENSMUSG00 | 209212 |
| NA | ENSMUSG00 | 73162 |
| NA | ENSMUSG00 | 52915 |
| NA | ENSMUSG00 | 74142 |
| NA | ENSMUSG00 | 226098 |
| NA | ENSMUSG00 | 101540 |
| NA | ENSMUSG00 | 106707 |
| NA | ENSMUSG00 | 68544 |
| NA | ENSMUSG00 | 208146 |
| NA | ENSMUSG00 | 228482 |
| NA | ENSMUSG00 | 170439 |
| Significant | ENSMUSG00 | 75415 |
| NA | ENSMUSG00 | 320595 |
| NA | ENSMUSG00 | 79560 |
| NA | ENSMUSG00 | 320790 |
| NA | ENSMUSG00 | 218035 |
| NA | ENSMUSG00 | 225182 |
| NA | ENSMUSG00 | 68350 |
| NA | ENSMUSG00 | 329002 |
| NA | ENSMUSG00 | 12319 |
| NA | ENSMUSG00 | 72296 |
| NA | ENSMUSG00 | 231915 |
| NA | ENSMUSG00 | 235380 |
| Significant | ENSMUSG00 | 252838 |
| NA | ENSMUSG00 | 66827 |
| NA | ENSMUSG00 | 69562 |
| NA | ENSMUSG00 | 231912 |
| NA | ENSMUSG00 | 98488 |
| NA | ENSMUSG00 | 20650 |
| NA | ENSMUSG00 | 11987 |
| NA | ENSMUSG00 | 386612 |
| NA | ENSMUSG00 | 74737 |
| NA | ENSMUSG00 | 76559 |
| NA | ENSMUSG00 | 73845 |
| NA | ENSMUSG00 | 216853 |
| NA | ENSMUSG00 | 382245 |
| NA | ENSMUSG00 | 19732 |
| NA | ENSMUSG00 | 66256 |
| NA | ENSMUSG00 | 52874 |
| NA | ENSMUSG00 | 102371 |
| NA | ENSMUSG00 | 71653 |
| NA | ENSMUSG00 | 243846 |

| | | |
|---|---|---|
| NA | ENSMUSG00 | 12741 |
| NA | ENSMUSG00 | 16543 |
| NA | ENSMUSG00 | 69962 |
| NA | ENSMUSG00 | 381306 |
| NA | ENSMUSG00 | 218914 |
| NA | ENSMUSG00 | 192119 |
| NA | ENSMUSG00 | 18708 |
| NA | ENSMUSG00 | 17537 |
| NA | ENSMUSG00 | 227095 |
| NA | ENSMUSG00 | 18207 |
| NA | ENSMUSG00 | 268697 |
| NA | ENSMUSG00 | 21771 |
| NA | ENSMUSG00 | 98682 |
| NA | ENSMUSG00 | 235533 |
| NA | ENSMUSG00 | 330914 |
| NA | ENSMUSG00 | 105450 |
| NA | ENSMUSG00 | 19933 |
| NA | ENSMUSG00 | 230908 |
| NA | ENSMUSG00 | 75698 |
| NA | ENSMUSG00 | 319618 |
| NA | ENSMUSG00 | 20715 |
| NA | ENSMUSG00 | 226442 |
| NA | ENSMUSG00 | 20908 |
| NA | ENSMUSG00 | 208518 |
| NA | ENSMUSG00 | 381293 |
| NA | ENSMUSG00 | 27966 |
| Significant | ENSMUSG00 | 15900 |
| NA | ENSMUSG00 | 84585 |
| NA | ENSMUSG00 | 236511 |
| NA | ENSMUSG00 | 68077 |
| NA | ENSMUSG00 | 67886 |
| NA | ENSMUSG00 | 20364 |
| NA | ENSMUSG00 | 70551 |
| NA | ENSMUSG00 | 56699 |
| NA | ENSMUSG00 | 17196 |
| NA | ENSMUSG00 | 75665 |
| NA | ENSMUSG00 | 276852 |
| NA | ENSMUSG00 | 223267 |
| NA | ENSMUSG00 | 28081 |
| NA | ENSMUSG00 | 218506 |
| NA | ENSMUSG00 | 207474 |
| NA | ENSMUSG00 | 231659 |
| NA | ENSMUSG00 | 16565 |
| NA | ENSMUSG00 | 27225 |
| Significant | ENSMUSG00 | 245671 |
| NA | ENSMUSG00 | 110821 |

| | | |
|---|---|---|
| Significant | ENSMUSG00 | 69806 |
| NA | ENSMUSG00 | 232491 |
| NA | ENSMUSG00 | 234684 |
| NA | ENSMUSG00 | 246229 |
| NA | ENSMUSG00 | 218503 |
| NA | ENSMUSG00 | 27494 |
| NA | ENSMUSG00 | 16518 |
| NA | ENSMUSG00 | 12861 |
| NA | ENSMUSG00 | 238386 |
| NA | ENSMUSG00 | 69069 |
| NA | ENSMUSG00 | 66622 |
| NA | ENSMUSG00 | 100859931 |
| NA | ENSMUSG00 | 67574 |
| NA | ENSMUSG00 | 224020 |
| NA | ENSMUSG00 | 235431 |
| NA | ENSMUSG00 | 52064 |
| NA | ENSMUSG00 | 170643 |
| NA | ENSMUSG00 | 12257 |
| NA | ENSMUSG00 | 50849 |
| NA | ENSMUSG00 | 105372 |
| Significant | ENSMUSG00 | 58218 |
| NA | ENSMUSG00 | 68526 |
| NA | ENSMUSG00 | 22019 |
| NA | ENSMUSG00 | 68889 |
| NA | ENSMUSG00 | 52432 |
| NA | ENSMUSG00 | 13803 |
| Significant | ENSMUSG00 | 69101 |
| NA | ENSMUSG00 | 69546 |
| NA | ENSMUSG00 | 66935 |
| NA | ENSMUSG00 | 170829 |
| NA | ENSMUSG00 | 51786 |
| NA | ENSMUSG00 | 217262 |
| NA | ENSMUSG00 | 27280 |
| NA | ENSMUSG00 | 73826 |
| NA | ENSMUSG00 | 320557 |
| NA | ENSMUSG00 | 231655 |
| NA | ENSMUSG00 | 19267 |
| NA | ENSMUSG00 | 50996 |
| NA | ENSMUSG00 | 225745 |
| NA | ENSMUSG00 | NA |
| NA | ENSMUSG00 | 11854 |
| NA | ENSMUSG00 | 68734 |
| NA | ENSMUSG00 | 239319 |
| NA | ENSMUSG00 | 21411 |
| NA | ENSMUSG00 | 17215 |
| NA | ENSMUSG00 | 68420 |

| | | |
|---|---|---|
| NA | ENSMUSG00 | 226432 |
| NA | ENSMUSG00 | 66416 |
| NA | ENSMUSG00 | 77552 |
| NA | ENSMUSG00 | 26431 |
| NA | ENSMUSG00 | 70361 |
| NA | ENSMUSG00 | 52639 |
| NA | ENSMUSG00 | 93690 |
| NA | ENSMUSG00 | 72634 |
| NA | ENSMUSG00 | 225339 |
| NA | ENSMUSG00 | 104681 |
| NA | ENSMUSG00 | 66559 |
| NA | ENSMUSG00 | 215615 |
| NA | ENSMUSG00 | 106064 |
| NA | ENSMUSG00 | 11603 |
| NA | ENSMUSG00 | 17855 |
| NA | ENSMUSG00 | 211798 |
| NA | ENSMUSG00 | 272538 |
| NA | ENSMUSG00 | 67451 |
| NA | ENSMUSG00 | 276846 |
| NA | ENSMUSG00 | 20194 |
| NA | ENSMUSG00 | 407821 |
| NA | ENSMUSG00 | 75763 |
| NA | ENSMUSG00 | 224008 |
| NA | ENSMUSG00 | 228019 |
| NA | ENSMUSG00 | 213498 |
| NA | ENSMUSG00 | 217944 |
| NA | ENSMUSG00 | 72114 |
| NA | ENSMUSG00 | 228012 |
| NA | ENSMUSG00 | 218460 |
| NA | ENSMUSG00 | 76044 |
| NA | ENSMUSG00 | 108645 |
| NA | ENSMUSG00 | 78908 |
| NA | ENSMUSG00 | 78752 |
| NA | ENSMUSG00 | 21371 |
| NA | ENSMUSG00 | 213452 |
| NA | ENSMUSG00 | 217935 |
| NA | ENSMUSG00 | 207425 |
| NA | ENSMUSG00 | 56220 |
| NA | ENSMUSG00 | 68875 |
| NA | ENSMUSG00 | 76491 |
| NA | ENSMUSG00 | 98758 |
| NA | ENSMUSG00 | 11881 |
| NA | ENSMUSG00 | 22685 |
| NA | ENSMUSG00 | 80880 |
| NA | ENSMUSG00 | 66435 |
| NA | ENSMUSG00 | 101490 |

| | | |
|---|---|---|
| NA | ENSMUSG00 | 68176 |
| NA | ENSMUSG00 | 69668 |
| NA | ENSMUSG00 | 231637 |
| Significant | ENSMUSG00 | 213391 |
| NA | ENSMUSG00 | 228005 |
| NA | ENSMUSG00 | 235184 |
| NA | ENSMUSG00 | 70383 |
| NA | ENSMUSG00 | 277396 |
| NA | ENSMUSG00 | 100715 |
| NA | ENSMUSG00 | 233912 |
| NA | ENSMUSG00 | 320705 |
| NA | ENSMUSG00 | 235134 |
| NA | ENSMUSG00 | 14747 |
| NA | ENSMUSG00 | 72022 |
| NA | ENSMUSG00 | 98403 |
| NA | ENSMUSG00 | 66433 |
| NA | ENSMUSG00 | 320541 |
| NA | ENSMUSG00 | 381085 |
| NA | ENSMUSG00 | 75605 |
| NA | ENSMUSG00 | 71675 |
| NA | ENSMUSG00 | 68644 |
| NA | ENSMUSG00 | 107035 |
| NA | ENSMUSG00 | 67492 |
| NA | ENSMUSG00 | 213539 |
| NA | ENSMUSG00 | 52850 |
| NA | ENSMUSG00 | 56068 |
| NA | ENSMUSG00 | 17096 |
| NA | ENSMUSG00 | 98710 |
| NA | ENSMUSG00 | 209760 |
| NA | ENSMUSG00 | 320129 |
| NA | ENSMUSG00 | 212933 |
| NA | ENSMUSG00 | 16392 |
| Significant | ENSMUSG00 | 58217 |
| NA | ENSMUSG00 | 237082 |
| NA | ENSMUSG00 | 228071 |
| NA | ENSMUSG00 | 105083 |
| Significant | ENSMUSG00 | 192187 |
| NA | ENSMUSG00 | 101502 |
| NA | ENSMUSG00 | 223701 |
| NA | ENSMUSG00 | 434402 |
| NA | ENSMUSG00 | 72795 |
| NA | ENSMUSG00 | 105835 |
| NA | ENSMUSG00 | 57439 |
| NA | ENSMUSG00 | 233904 |
| NA | ENSMUSG00 | 20196 |
| NA | ENSMUSG00 | 73422 |

| | | |
|---|---|---|
| NA | ENSMUSG00 | 66923 |
| NA | ENSMUSG00 | 192232 |
| NA | ENSMUSG00 | 432572 |
| NA | ENSMUSG00 | 230979 |
| Significant | ENSMUSG00 | 328795 |
| NA | ENSMUSG00 | 218639 |
| NA | ENSMUSG00 | 56489 |
| NA | ENSMUSG00 | 68497 |
| Significant | ENSMUSG00 | 17444 |
| NA | ENSMUSG00 | 333315 |
| NA | ENSMUSG00 | 30877 |
| NA | ENSMUSG00 | 216551 |
| NA | ENSMUSG00 | 170625 |
| NA | ENSMUSG00 | 241490 |
| NA | ENSMUSG00 | 69640 |
| NA | ENSMUSG00 | 80284 |
| NA | ENSMUSG00 | 381802 |
| NA | ENSMUSG00 | 229542 |
| NA | ENSMUSG00 | 67010 |
| NA | ENSMUSG00 | 229541 |
| NA | ENSMUSG00 | 11911 |
| NA | ENSMUSG00 | 100177 |
| NA | ENSMUSG00 | 228061 |
| NA | ENSMUSG00 | 18038 |
| NA | ENSMUSG00 | 14123 |
| NA | ENSMUSG00 | 218629 |
| NA | ENSMUSG00 | 328977 |
| NA | ENSMUSG00 | 235461 |
| NA | ENSMUSG00 | 67785 |
| NA | ENSMUSG00 | 252875 |
| NA | ENSMUSG00 | 94192 |
| NA | ENSMUSG00 | 66422 |
| NA | ENSMUSG00 | 52708 |
| NA | ENSMUSG00 | 245638 |
| NA | ENSMUSG00 | 235497 |
| NA | ENSMUSG00 | 269582 |
| NA | ENSMUSG00 | 68449 |
| NA | ENSMUSG00 | 382066 |
| NA | ENSMUSG00 | 102871 |
| NA | ENSMUSG00 | 76850 |
| NA | ENSMUSG00 | 70233 |
| NA | ENSMUSG00 | 71238 |
| NA | ENSMUSG00 | 216825 |
| NA | ENSMUSG00 | 238317 |
| NA | ENSMUSG00 | 23857 |
| NA | ENSMUSG00 | 212439 |

| | | |
|---|---|---|
| NA | ENSMUSG00 | 245631 |
| NA | ENSMUSG00 | 74383 |
| NA | ENSMUSG00 | 105000 |
| NA | ENSMUSG00 | 223697 |
| Significant | ENSMUSG00 | 14904 |
| NA | ENSMUSG00 | 20422 |
| NA | ENSMUSG00 | 228790 |
| NA | ENSMUSG00 | 20466 |
| NA | ENSMUSG00 | 100206 |
| NA | ENSMUSG00 | 216820 |
| NA | ENSMUSG00 | 70427 |
| NA | ENSMUSG00 | 70093 |
| NA | ENSMUSG00 NA | |
| NA | ENSMUSG00 | 16923 |
| NA | ENSMUSG00 | 245622 |
| NA | ENSMUSG00 | 338523 |
| NA | ENSMUSG00 | 20239 |
| NA | ENSMUSG00 | 233876 |
| Significant | ENSMUSG00 | 65255 |
| NA | ENSMUSG00 | 74178 |
| NA | ENSMUSG00 | 229534 |
| NA | ENSMUSG00 | 230751 |
| NA | ENSMUSG00 | 17133 |
| NA | ENSMUSG00 | 224902 |
| NA | ENSMUSG00 | 20416 |
| NA | ENSMUSG00 | 217695 |
| NA | ENSMUSG00 | 53357 |
| NA | ENSMUSG00 | 240816 |
| NA | ENSMUSG00 | 319945 |
| NA | ENSMUSG00 | 16440 |
| NA | ENSMUSG00 | 338350 |
| NA | ENSMUSG00 | 268420 |
| NA | ENSMUSG00 | 66412 |
| NA | ENSMUSG00 | 67936 |
| NA | ENSMUSG00 | 66541 |
| NA | ENSMUSG00 | 66090 |
| NA | ENSMUSG00 | 230738 |
| NA | ENSMUSG00 | 17910 |
| NA | ENSMUSG00 | 80795 |
| Significant | ENSMUSG00 | 74091 |
| NA | ENSMUSG00 | 50772 |
| NA | ENSMUSG00 | 108689 |
| NA | ENSMUSG00 | 13211 |
| NA | ENSMUSG00 | 217692 |
| NA | ENSMUSG00 | 69456 |
| NA | ENSMUSG00 | 71836 |

| | | |
|---|---|---|
| NA | ENSMUSG00 | 246782 |
| Significant | ENSMUSG00 | 22381 |
| NA | ENSMUSG00 | 231713 |
| NA | ENSMUSG00 | 231712 |
| NA | ENSMUSG00 | 107071 |
| NA | ENSMUSG00 | 68563 |
| NA | ENSMUSG00 | 101148 |
| NA | ENSMUSG00 | 218544 |
| NA | ENSMUSG00 | 269700 |
| NA | ENSMUSG00 | 15901 |
| NA | ENSMUSG00 | 66059 |
| NA | ENSMUSG00 | 12069 |
| NA | ENSMUSG00 | 171504 |
| NA | ENSMUSG00 | 15199 |
| NA | ENSMUSG00 | 226517 |
| NA | ENSMUSG00 | 208194 |
| NA | ENSMUSG00 | 235315 |
| NA | ENSMUSG00 | 66333 |
| NA | ENSMUSG00 | 329152 |
| NA | ENSMUSG00 | 77827 |
| NA | ENSMUSG00 | 66405 |
| NA | ENSMUSG00 | 14255 |
| NA | ENSMUSG00 | 20613 |
| NA | ENSMUSG00 | 14166 |
| NA | ENSMUSG00 | 233065 |
| Significant | ENSMUSG00 | 27528 |
| Significant | ENSMUSG00 | 20708 |
| NA | ENSMUSG00 | 76367 |
| NA | ENSMUSG00 | 21968 |
| NA | ENSMUSG00 | 108909 |
| NA | ENSMUSG00 | 54601 |
| NA | ENSMUSG00 | 100418259 |
| NA | ENSMUSG00 | 381157 |
| NA | ENSMUSG00 | 104175 |
| NA | ENSMUSG00 | 67774 |
| Significant | ENSMUSG00 | 212114 |
| NA | ENSMUSG00 | 14702 |
| NA | ENSMUSG00 | 239743 |
| NA | ENSMUSG00 | 227094 |
| NA | ENSMUSG00 | 19222 |
| NA | ENSMUSG00 | 66967 |
| NA | ENSMUSG00 | 244667 |
| NA | ENSMUSG00 | 101023 |
| NA | ENSMUSG00 | 211986 |
| NA | ENSMUSG00 | 212514 |
| NA | ENSMUSG00 | 244745 |

| | | |
|---|---|---|
| NA | ENSMUSG00 | 104027 |
| NA | ENSMUSG00 | 213989 |
| NA | ENSMUSG00 | 330788 |
| NA | ENSMUSG00 | 22146 |
| NA | ENSMUSG00 | 320974 |
| NA | ENSMUSG00 | 232157 |
| NA | ENSMUSG00 | 66690 |
| NA | ENSMUSG00 | 75847 |
| NA | ENSMUSG00 | 235542 |
| NA | ENSMUSG00 | 242642 |
| NA | ENSMUSG00 | 219144 |
| NA | ENSMUSG00 | 66459 |
| NA | ENSMUSG00 | 319719 |
| NA | ENSMUSG00 | 218341 |
| NA | ENSMUSG00 | 230709 |
| NA | ENSMUSG00 | 326622 |
| NA | ENSMUSG00 | 100037278 |
| NA | ENSMUSG00 | 277978 |
| NA | ENSMUSG00 | 100201 |
| NA | ENSMUSG00 | 230801 |
| NA | ENSMUSG00 | 54122 |
| NA | ENSMUSG00 | 236312 |
| NA | ENSMUSG00 | 384309 |
| NA | ENSMUSG00 | 216821 |
| NA | ENSMUSG00 | 654801 |
| NA | ENSMUSG00 | 654818 |
| Significant | ENSMUSG00 | 26879 |
| NA | ENSMUSG00 | 114585 |
| NA | ENSMUSG00 | 381668 |
| NA | ENSMUSG00 | 78749 |
| NA | ENSMUSG00 | NA |
| NA | ENSMUSG00 | 71059 |
| NA | ENSMUSG00 | 67298 |
| NA | ENSMUSG00 | 72190 |
| NA | ENSMUSG00 | 170707 |
| NA | ENSMUSG00 | 71198 |
| NA | ENSMUSG00 | 80515 |
| NA | ENSMUSG00 | 69573 |
| NA | ENSMUSG00 | 100042807 |
| NA | ENSMUSG00 | 243813 |
| NA | ENSMUSG00 | 103551 |
| NA | ENSMUSG00 | 67078 |
| NA | ENSMUSG00 | 240869 |
| NA | ENSMUSG00 | 628277 |
| NA | ENSMUSG00 | 217700 |
| NA | ENSMUSG00 | 317757 |

| | | |
|---|---|---|
| NA | ENSMUSG00 | 100900 |
| NA | ENSMUSG00 | 269254 |
| NA | ENSMUSG00 | 67306 |
| NA | ENSMUSG00 | 74762 |
| NA | ENSMUSG00 | 230577 |
| NA | ENSMUSG00 | 235534 |
| NA | ENSMUSG00 | 193043 |
| NA | ENSMUSG00 | 67080 |
| NA | ENSMUSG00 | 242408 |
| NA | ENSMUSG00 | 194908 |
| NA | ENSMUSG00 | 434325 |
| NA | ENSMUSG00 | 233115 |
| NA | ENSMUSG00 | 14154 |
| NA | ENSMUSG00 | 211948 |
| NA | ENSMUSG00 | 19989 |
| NA | ENSMUSG00 | 228592 |
| NA | ENSMUSG00 | 19247 |
| NA | ENSMUSG00 | 320148 |
| NA | ENSMUSG00 | NA |
| NA | ENSMUSG00 | NA |
| NA | ENSMUSG00 | 266614 |
| NA | ENSMUSG00 | 75099 |
| Significant | ENSMUSG00 | 73149 |
| NA | ENSMUSG00 | 24075 |
| NA | ENSMUSG00 | 68310 |
| NA | ENSMUSG00 | 211255 |
| NA | ENSMUSG00 | 234967 |
| NA | ENSMUSG00 | NA |
| NA | ENSMUSG00 | 14739 |
| NA | ENSMUSG00 | 27223 |
| NA | ENSMUSG00 | 382073 |
| NA | ENSMUSG00 | 236904 |
| NA | ENSMUSG00 | 328830 |
| NA | ENSMUSG00 | 72145 |
| NA | ENSMUSG00 | 54199 |
| NA | ENSMUSG00 | 230753 |
| NA | ENSMUSG00 | 269999 |
| NA | ENSMUSG00 | 214552 |
| NA | ENSMUSG00 | 19290 |
| NA | ENSMUSG00 | 217664 |
| NA | ENSMUSG00 | 216456 |
| NA | ENSMUSG00 | 28028 |
| NA | ENSMUSG00 | 240660 |
| NA | ENSMUSG00 | 272359 |
| NA | ENSMUSG00 | 235633 |
| NA | ENSMUSG00 | 14260 |

| | | |
|---|---|---|
| NA | ENSMUSG00 | 69306 |
| NA | ENSMUSG00 | 216543 |
| NA | ENSMUSG00 | NA |
| Significant | ENSMUSG00 | 268354 |
| NA | ENSMUSG00 | 237711 |
| NA | ENSMUSG00 | 20193 |
| NA | ENSMUSG00 | 319772 |
| NA | ENSMUSG00 | 229675 |
| NA | ENSMUSG00 | 216974 |
| NA | ENSMUSG00 | 231717 |
| NA | ENSMUSG00 | 329509 |
| NA | ENSMUSG00 | 11845 |
| NA | ENSMUSG00 | 69155 |
| NA | ENSMUSG00 | 77286 |
| NA | ENSMUSG00 | 382252 |
| NA | ENSMUSG00 | 76522 |
| NA | ENSMUSG00 | 414084 |
| NA | ENSMUSG00 | 56458 |
| NA | ENSMUSG00 | 80290 |
| NA | ENSMUSG00 | 13611 |
| NA | ENSMUSG00 | 12532 |
| NA | ENSMUSG00 | 231413 |
| NA | ENSMUSG00 | 78244 |
| NA | ENSMUSG00 | 244853 |
| NA | ENSMUSG00 | 105193 |
| NA | ENSMUSG00 | 213208 |
| NA | ENSMUSG00 | 239647 |
| NA | ENSMUSG00 | 64291 |
| NA | ENSMUSG00 | 13024 |
| NA | ENSMUSG00 | NA |
| NA | ENSMUSG00 | 68795 |
| NA | ENSMUSG00 | 216964 |
| NA | ENSMUSG00 | 23796 |
| NA | ENSMUSG00 | 231642 |
| NA | ENSMUSG00 | 98432 |
| NA | ENSMUSG00 | 67062 |
| NA | ENSMUSG00 | 319317 |
| NA | ENSMUSG00 | 210808 |
| NA | ENSMUSG00 | 70054 |
| NA | ENSMUSG00 | 69309 |
| NA | ENSMUSG00 | 332359 |
| NA | ENSMUSG00 | 104725 |
| NA | ENSMUSG00 | 102638448 |
| NA | ENSMUSG00 | 67768 |
| NA | ENSMUSG00 | 68813 |
| Significant | ENSMUSG00 | 217835 |

| | | |
|---|---|---|
| NA | ENSMUSG00 | 74349 |
| NA | ENSMUSG00 | 74645 |
| NA | ENSMUSG00 | 66443 |
| NA | ENSMUSG00 | 69090 |
| NA | ENSMUSG00 | 21769 |
| NA | ENSMUSG00 | 77034 |
| NA | ENSMUSG00 | 224598 |
| NA | ENSMUSG00 | 69556 |
| NA | ENSMUSG00 | 229801 |
| NA | ENSMUSG00 | 16898 |
| NA | ENSMUSG00 | 268759 |
| NA | ENSMUSG00 | 69903 |
| NA | ENSMUSG00 | 71213 |
| NA | ENSMUSG00 | 11431 |
| NA | ENSMUSG00 | 170743 |
| NA | ENSMUSG00 | 213236 |
| NA | ENSMUSG00 | 66818 |
| NA | ENSMUSG00 | 320080 |
| NA | ENSMUSG00 | 239759 |
| NA | ENSMUSG00 | 72931 |
| Significant | ENSMUSG00 | 380686 |
| NA | ENSMUSG00 | 207785 |
| NA | ENSMUSG00 | 229791 |
| NA | ENSMUSG00 | 14362 |
| NA | ENSMUSG00 | 234725 |
| NA | ENSMUSG00 | 230917 |
| NA | ENSMUSG00 | 233826 |
| NA | ENSMUSG00 | 219131 |
| NA | ENSMUSG00 | 69731 |
| Significant | ENSMUSG00 | 625098 |
| NA | ENSMUSG00 | 66787 |
| NA | ENSMUSG00 | 194268 |
| Significant | ENSMUSG00 | 66222 |
| NA | ENSMUSG00 NA | |
| NA | ENSMUSG00 | 52575 |
| NA | ENSMUSG00 | 52392 |
| NA | ENSMUSG00 NA | |
| NA | ENSMUSG00 | 22695 |
| NA | ENSMUSG00 | 328035 |
| NA | ENSMUSG00 | 235626 |
| NA | ENSMUSG00 | 69046 |
| NA | ENSMUSG00 | 327951 |
| NA | ENSMUSG00 | 30944 |
| NA | ENSMUSG00 | 140497 |
| NA | ENSMUSG00 | 230126 |
| NA | ENSMUSG00 | 98303 |

| | | |
|---|---|---|
| NA | ENSMUSG00 | 21897 |
| NA | ENSMUSG00 | 72290 |
| NA | ENSMUSG00 | 73363 |
| NA | ENSMUSG00 | 224640 |
| NA | ENSMUSG00 | 99696 |
| NA | ENSMUSG00 | 72667 |
| NA | ENSMUSG00 | 68185 |
| NA | ENSMUSG00 | 22272 |
| NA | ENSMUSG00 | 269033 |
| NA | ENSMUSG00 | 238673 |
| NA | ENSMUSG00 | 103220 |
| NA | ENSMUSG00 | 327900 |
| NA | ENSMUSG00 | 70802 |
| NA | ENSMUSG00 | 622320 |
| NA | ENSMUSG00 | 319701 |
| NA | ENSMUSG00 | 242864 |
| NA | ENSMUSG00 | 67158 |
| NA | ENSMUSG00 | 22117 |
| NA | ENSMUSG00 | 73747 |
| NA | ENSMUSG00 | 14367 |
| NA | ENSMUSG00 | 207175 |
| NA | ENSMUSG00 | 18754 |
| NA | ENSMUSG00 | 269878 |
| NA | ENSMUSG00 | 622655 |
| NA | ENSMUSG00 | 230259 |
| NA | ENSMUSG00 | 108086 |
| NA | ENSMUSG00 | 13609 |
| NA | ENSMUSG00 | 328967 |
| NA | ENSMUSG00 NA | |
| NA | ENSMUSG00 | 225888 |
| NA | ENSMUSG00 | 67582 |
| NA | ENSMUSG00 | 272158 |
| NA | ENSMUSG00 | 67509 |
| NA | ENSMUSG00 | 76808 |
| NA | ENSMUSG00 | 73710 |
| NA | ENSMUSG00 | 70325 |
| NA | ENSMUSG00 | 78653 |
| NA | ENSMUSG00 | 101602 |
| NA | ENSMUSG00 | 71923 |
| NA | ENSMUSG00 | 110380 |
| NA | ENSMUSG00 | 12696 |
| NA | ENSMUSG00 | 381510 |
| NA | ENSMUSG00 | 70675 |
| NA | ENSMUSG00 | 213484 |
| NA | ENSMUSG00 | 50785 |
| NA | ENSMUSG00 | 68618 |

| | | |
|---|---|---|
| NA | ENSMUSG00 | 70625 |
| NA | ENSMUSG00 | 69234 |
| NA | ENSMUSG00 | 232976 |
| NA | ENSMUSG00 | 378462 |
| NA | ENSMUSG00 | 195522 |
| NA | ENSMUSG00 | 26886 |
| NA | ENSMUSG00 | 385668 |
| NA | ENSMUSG00 | 68255 |
| NA | ENSMUSG00 | 237847 |
| NA | ENSMUSG00 | 231070 |
| NA | ENSMUSG00 | 50907 |
| NA | ENSMUSG00 | 218454 |
| NA | ENSMUSG00 | 68636 |
| NA | ENSMUSG00 | 81897 |
| NA | ENSMUSG00 | 320181 |
| NA | ENSMUSG00 NA | |
| NA | ENSMUSG00 | 229841 |
| NA | ENSMUSG00 | 244237 |
| NA | ENSMUSG00 | 192652 |
| NA | ENSMUSG00 | 67020 |
| NA | ENSMUSG00 | 69307 |
| NA | ENSMUSG00 | 258712 |
| NA | ENSMUSG00 | 12767 |
| NA | ENSMUSG00 | 17075 |
| NA | ENSMUSG00 | 217826 |
| NA | ENSMUSG00 | 79263 |
| NA | ENSMUSG00 | 56043 |
| NA | ENSMUSG00 | 668661 |
| NA | ENSMUSG00 | 68861 |
| NA | ENSMUSG00 | 56258 |
| NA | ENSMUSG00 | 212090 |
| NA | ENSMUSG00 | 68033 |
| NA | ENSMUSG00 NA | |
| NA | ENSMUSG00 | 101197 |
| NA | ENSMUSG00 | 269954 |
| NA | ENSMUSG00 | 100683 |
| Significant | ENSMUSG00 | 80885 |
| NA | ENSMUSG00 | 66460 |
| NA | ENSMUSG00 | 238725 |
| NA | ENSMUSG00 | 434377 |
| NA | ENSMUSG00 | 71986 |
| Significant | ENSMUSG00 | 18619 |
| NA | ENSMUSG00 | 229681 |
| NA | ENSMUSG00 | 12091 |
| NA | ENSMUSG00 | 233887 |
| NA | ENSMUSG00 | 66580 |

| | | |
|---|---|---|
| NA | ENSMUSG00 | 239827 |
| Significant | ENSMUSG00 | 102103 |
| NA | ENSMUSG00 | 320683 |
| NA | ENSMUSG00 | 98496 |
| NA | ENSMUSG00 | 66715 |
| NA | ENSMUSG00 | 104252 |
| NA | ENSMUSG00 | 106073 |
| NA | ENSMUSG00 | 276829 |
| NA | ENSMUSG00 | 114716 |
| NA | ENSMUSG00 | 217430 |
| NA | ENSMUSG00 | 72338 |
| NA | ENSMUSG00 | 105663 |
| NA | ENSMUSG00 | 115489953 |
| NA | ENSMUSG00 | NA |
| NA | ENSMUSG00 | 11555 |
| NA | ENSMUSG00 | 212377 |
| NA | ENSMUSG00 | 56844 |
| NA | ENSMUSG00 | 233893 |
| NA | ENSMUSG00 | 70350 |
| NA | ENSMUSG00 | 78521 |
| NA | ENSMUSG00 | 258189 |
| NA | ENSMUSG00 | 434204 |
| NA | ENSMUSG00 | NA |
| NA | ENSMUSG00 | 12193 |
| NA | ENSMUSG00 | 67538 |
| Significant | ENSMUSG00 | 19265 |
| NA | ENSMUSG00 | 20723 |
| NA | ENSMUSG00 | 108755 |
| NA | ENSMUSG00 | 12905 |
| NA | ENSMUSG00 | 74558 |
| NA | ENSMUSG00 | 100042179 |
| NA | ENSMUSG00 | 232164 |
| NA | ENSMUSG00 | 230279 |
| NA | ENSMUSG00 | 66864 |
| NA | ENSMUSG00 | 15958 |
| NA | ENSMUSG00 | 194126 |
| NA | ENSMUSG00 | 193286 |
| NA | ENSMUSG00 | 24030 |
| NA | ENSMUSG00 | 232341 |
| NA | ENSMUSG00 | 26356 |
| NA | ENSMUSG00 | 230125 |
| NA | ENSMUSG00 | 207781 |
| NA | ENSMUSG00 | 320534 |
| NA | ENSMUSG00 | 208643 |
| NA | ENSMUSG00 | 17749 |
| Significant | ENSMUSG00 | 238875 |

| | | |
|---|---|---|
| NA | ENSMUSG00 | 66983 |
| NA | ENSMUSG00 | 140484 |
| Significant | ENSMUSG00 | 170460 |
| NA | ENSMUSG00 | 215900 |
| NA | ENSMUSG00 | 55988 |
| NA | ENSMUSG00 | 432940 |
| NA | ENSMUSG00 | 282619 |
| NA | ENSMUSG00 | 108954 |
| NA | ENSMUSG00 | 231549 |
| NA | ENSMUSG00 | 233812 |
| NA | ENSMUSG00 | 240697 |
| NA | ENSMUSG00 | 319675 |
| NA | ENSMUSG00 | 240613 |
| NA | ENSMUSG00 | 225929 |
| NA | ENSMUSG00 | 171167 |
| Significant | ENSMUSG00 | 268567 |
| NA | ENSMUSG00 | 50914 |
| NA | ENSMUSG00 | 108154 |
| NA | ENSMUSG00 | 108961 |
| NA | ENSMUSG00 | 75352 |
| NA | ENSMUSG00 | 74352 |
| NA | ENSMUSG00 | 235505 |
| NA | ENSMUSG00 | 106583 |
| NA | ENSMUSG00 | 239157 |
| NA | ENSMUSG00 | 104709 |
| NA | ENSMUSG00 | 18793 |
| NA | ENSMUSG00 | 19018 |
| NA | ENSMUSG00 | 271564 |
| Significant | ENSMUSG00 | 231805 |
| NA | ENSMUSG00 | 234396 |
| NA | ENSMUSG00 | 22720 |
| NA | ENSMUSG00 | 226090 |
| NA | ENSMUSG00 | 66972 |
| NA | ENSMUSG00 | 19981 |
| NA | ENSMUSG00 | 218100 |
| NA | ENSMUSG00 | 26451 |
| NA | ENSMUSG00 | 16469 |
| NA | ENSMUSG00 | 19659 |
| NA | ENSMUSG00 | 226418 |
| NA | ENSMUSG00 | 52150 |
| NA | ENSMUSG00 | 192976 |
| NA | ENSMUSG00 | 12070 |
| Significant | ENSMUSG00 | 15382 |
| NA | ENSMUSG00 | 234728 |
| NA | ENSMUSG00 | 320472 |
| NA | ENSMUSG00 | 20672 |

| | | |
|---|---|---|
| NA | ENSMUSG00 | 12856 |
| NA | ENSMUSG00 | 229593 |
| NA | ENSMUSG00 | 67516 |
| NA | ENSMUSG00 | 210172 |
| NA | ENSMUSG00 | 224093 |
| NA | ENSMUSG00 | 278240 |
| NA | ENSMUSG00 | 79233 |
| NA | ENSMUSG00 | 106840 |
| NA | ENSMUSG00 | 68281 |
| NA | ENSMUSG00 | 100515 |
| NA | ENSMUSG00 | 380840 |
| NA | ENSMUSG00 | 233210 |
| Significant | ENSMUSG00 | 77011 |
| NA | ENSMUSG00 | 71911 |
| NA | ENSMUSG00 | 382117 |
| NA | ENSMUSG00 | 210004 |
| NA | ENSMUSG00 | 54201 |
| NA | ENSMUSG00 | 72397 |
| NA | ENSMUSG00 | 67393 |
| NA | ENSMUSG00 | 76824 |
| NA | ENSMUSG00 | 320351 |
| NA | ENSMUSG00 | 232196 |
| NA | ENSMUSG00 | 211550 |
| NA | ENSMUSG00 | 214987 |
| NA | ENSMUSG00 | 13000 |
| NA | ENSMUSG00 | 15361 |
| NA | ENSMUSG00 | 69550 |
| NA | ENSMUSG00 | 100040970 |
| NA | ENSMUSG00 | 57912 |
| NA | ENSMUSG00 | 216858 |
| NA | ENSMUSG00 | 232431 |
| NA | ENSMUSG00 | 320234 |
| NA | ENSMUSG00 | 50529 |
| Significant | ENSMUSG00 | 105732 |
| NA | ENSMUSG00 | 80837 |
| NA | ENSMUSG00 | 77781 |
| NA | ENSMUSG00 | 71952 |
| NA | ENSMUSG00 | 67109 |
| NA | ENSMUSG00 | 244416 |
| NA | ENSMUSG00 | 64945 |
| NA | ENSMUSG00 | 17476 |
| NA | ENSMUSG00 | 78412 |
| NA | ENSMUSG00 | 231287 |
| NA | ENSMUSG00 | 320415 |
| NA | ENSMUSG00 | 106947 |
| NA | ENSMUSG00 | 71678 |

| | | |
|---|---|---|
| NA | ENSMUSG00 | 216197 |
| NA | ENSMUSG00 | 76608 |
| NA | ENSMUSG00 | 242736 |
| Significant | ENSMUSG00 | 14113 |
| NA | ENSMUSG00 | 270669 |
| NA | ENSMUSG00 | 20238 |
| NA | ENSMUSG00 | 15944 |
| NA | ENSMUSG00 | 68744 |
| Significant | ENSMUSG00 | 16995 |
| NA | ENSMUSG00 | 68550 |
| NA | ENSMUSG00 | 57869 |
| NA | ENSMUSG00 | 18105 |
| NA | ENSMUSG00 | 114565 |
| NA | ENSMUSG00 | 110796 |
| NA | ENSMUSG00 | 231225 |
| NA | ENSMUSG00 | 71682 |
| NA | ENSMUSG00 | 212679 |
| NA | ENSMUSG00 | 211949 |
| NA | ENSMUSG00 | 22701 |
| NA | ENSMUSG00 | 73490 |
| NA | ENSMUSG00 | 263876 |
| NA | ENSMUSG00 | 235682 |
| NA | ENSMUSG00 | 233280 |
| NA | ENSMUSG00 | 225995 |
| NA | ENSMUSG00 | 209497 |
| NA | ENSMUSG00 | 67446 |
| NA | ENSMUSG00 | 83485 |
| NA | ENSMUSG00 | 73827 |
| NA | ENSMUSG00 | 268749 |
| NA | ENSMUSG00 | 57755 |
| NA | ENSMUSG00 | 231946 |
| NA | ENSMUSG00 | 106759 |
| NA | ENSMUSG00 | 67300 |
| NA | ENSMUSG00 | 12484 |
| NA | ENSMUSG00 | 72020 |
| NA | ENSMUSG00 | 52463 |
| NA | ENSMUSG00 | 219094 |
| NA | ENSMUSG00 | 214854 |
| NA | ENSMUSG00 | 59021 |
| NA | ENSMUSG00 | 110350 |
| NA | ENSMUSG00 | 75219 |
| NA | ENSMUSG00 | 229096 |
| NA | ENSMUSG00 | 20005 |
| NA | ENSMUSG00 | 330050 |
| NA | ENSMUSG00 | 407786 |
| NA | ENSMUSG00 | 319179 |

| | | |
|---|---|---|
| NA | ENSMUSG00 | 277939 |
| NA | ENSMUSG00 | 19224 |
| NA | ENSMUSG00 | 66048 |
| Significant | ENSMUSG00 | 14432 |
| NA | ENSMUSG00 | 140482 |
| NA | ENSMUSG00 | 55948 |
| NA | ENSMUSG00 | 216860 |
| NA | ENSMUSG00 | 71223 |
| NA | ENSMUSG00 | 192651 |
| NA | ENSMUSG00 | 226016 |
| NA | ENSMUSG00 | 240960 |
| NA | ENSMUSG00 | 233890 |
| NA | ENSMUSG00 | 108902 |
| NA | ENSMUSG00 | 234776 |
| NA | ENSMUSG00 | 21815 |
| NA | ENSMUSG00 | 69274 |
| NA | ENSMUSG00 | 235132 |
| NA | ENSMUSG00 | 399558 |
| NA | ENSMUSG00 | 66932 |
| NA | ENSMUSG00 | 107242 |
| NA | ENSMUSG00 | 268880 |
| NA | ENSMUSG00 | 11861 |
| NA | ENSMUSG00 | 268566 |
| NA | ENSMUSG00 | 67068 |
| NA | ENSMUSG00 | 212163 |
| NA | ENSMUSG00 | 22693 |
| NA | ENSMUSG00 | 239337 |
| NA | ENSMUSG00 | 22110 |
| NA | ENSMUSG00 | 320135 |
| NA | ENSMUSG00 | 217653 |
| NA | ENSMUSG00 | 67948 |
| NA | ENSMUSG00 | 74325 |
| Significant | ENSMUSG00 | 233724 |
| NA | ENSMUSG00 | 17035 |
| NA | ENSMUSG00 | 209584 |
| NA | ENSMUSG00 | 56525 |
| NA | ENSMUSG00 | 212398 |
| NA | ENSMUSG00 | 97159 |
| NA | ENSMUSG00 | 227622 |
| NA | ENSMUSG00 | 72514 |
| NA | ENSMUSG00 | 72650 |
| NA | ENSMUSG00 | 71865 |
| NA | ENSMUSG00 | 70333 |
| NA | ENSMUSG00 | 83984 |
| NA | ENSMUSG00 | 107328 |
| NA | ENSMUSG00 | 73390 |

| | | |
|---|---|---|
| NA | ENSMUSG00 | NA |
| NA | ENSMUSG00 | 100040633 |
| NA | ENSMUSG00 | 77929 |
| NA | ENSMUSG00 | 258459 |
| NA | ENSMUSG00 | 101994 |
| NA | ENSMUSG00 | 338362 |
| NA | ENSMUSG00 | 66849 |
| NA | ENSMUSG00 | 71707 |
| NA | ENSMUSG00 | 66162 |
| NA | ENSMUSG00 | 226178 |
| NA | ENSMUSG00 | 209086 |
| NA | ENSMUSG00 | 330812 |
| NA | ENSMUSG00 | 209032 |
| NA | ENSMUSG00 | 237211 |
| NA | ENSMUSG00 | 102747 |
| NA | ENSMUSG00 | 73683 |
| NA | ENSMUSG00 | 230936 |
| NA | ENSMUSG00 | 66643 |
| NA | ENSMUSG00 | 268706 |
| Significant | ENSMUSG00 | 246746 |
| NA | ENSMUSG00 | 56697 |
| NA | ENSMUSG00 | 78317 |
| NA | ENSMUSG00 | 94092 |
| NA | ENSMUSG00 | 68911 |
| NA | ENSMUSG00 | 71963 |
| NA | ENSMUSG00 | 55950 |
| NA | ENSMUSG00 | 406217 |
| NA | ENSMUSG00 | 66887 |
| Significant | ENSMUSG00 | 231931 |
| Significant | ENSMUSG00 | 269604 |
| NA | ENSMUSG00 | 59025 |
| NA | ENSMUSG00 | 12145 |
| NA | ENSMUSG00 | 100532 |
| NA | ENSMUSG00 | 213988 |
| NA | ENSMUSG00 | 320816 |
| NA | ENSMUSG00 | 76510 |
| NA | ENSMUSG00 | 17357 |
| NA | ENSMUSG00 | 16491 |
| NA | ENSMUSG00 | 52331 |
| NA | ENSMUSG00 | NA |
| NA | ENSMUSG00 | 225280 |
| NA | ENSMUSG00 | 546336 |
| NA | ENSMUSG00 | 227331 |
| NA | ENSMUSG00 | 30058 |
| NA | ENSMUSG00 | 243963 |
| NA | ENSMUSG00 | 68799 |

| | | |
|---|---|---|
| NA | ENSMUSG00 | 229504 |
| NA | ENSMUSG00 | 76976 |
| NA | ENSMUSG00 | 56805 |
| NA | ENSMUSG00 | 241576 |
| NA | ENSMUSG00 | 11840 |
| NA | ENSMUSG00 | 11615 |
| NA | ENSMUSG00 | 99730 |
| NA | ENSMUSG00 | NA |
| NA | ENSMUSG00 | 319776 |
| NA | ENSMUSG00 | 229700 |
| NA | ENSMUSG00 | 238247 |
| Significant | ENSMUSG00 | 12495 |
| NA | ENSMUSG00 | 226049 |
| NA | ENSMUSG00 | 269642 |
| NA | ENSMUSG00 | 381022 |
| Significant | ENSMUSG00 | 20345 |
| NA | ENSMUSG00 | 210711 |
| NA | ENSMUSG00 | 74626 |
| NA | ENSMUSG00 | 78541 |
| NA | ENSMUSG00 | 213573 |
| NA | ENSMUSG00 | 64450 |
| NA | ENSMUSG00 | 105827 |
| NA | ENSMUSG00 | 100034361 |
| NA | ENSMUSG00 | 269529 |
| NA | ENSMUSG00 | 67702 |
| NA | ENSMUSG00 | 14708 |
| NA | ENSMUSG00 | 77128 |
| NA | ENSMUSG00 | 208440 |
| NA | ENSMUSG00 | 381626 |
| NA | ENSMUSG00 | 20973 |
| NA | ENSMUSG00 | 50720 |
| NA | ENSMUSG00 | 408068 |
| NA | ENSMUSG00 | 319710 |
| NA | ENSMUSG00 | 68839 |
| NA | ENSMUSG00 | 244631 |
| NA | ENSMUSG00 | 70466 |
| NA | ENSMUSG00 | 320360 |
| NA | ENSMUSG00 | 108927 |
| NA | ENSMUSG00 | 69893 |
| NA | ENSMUSG00 | 76219 |
| NA | ENSMUSG00 | 27047 |
| NA | ENSMUSG00 | 382051 |
| NA | ENSMUSG00 | 14062 |
| NA | ENSMUSG00 | 67296 |
| NA | ENSMUSG00 | 240476 |
| NA | ENSMUSG00 | 17349 |

| | | |
|---|---|---|
| NA | ENSMUSG00 | 69773 |
| NA | ENSMUSG00 | 231042 |
| NA | ENSMUSG00 | 70101 |
| NA | ENSMUSG00 | 71276 |
| NA | ENSMUSG00 | 320869 |
| NA | ENSMUSG00 | 232934 |
| NA | ENSMUSG00 | 228859 |
| NA | ENSMUSG00 | 213393 |
| NA | ENSMUSG00 | 268903 |
| NA | ENSMUSG00 | 68736 |
| NA | ENSMUSG00 | 216829 |
| NA | ENSMUSG00 | 217306 |
| NA | ENSMUSG00 | 102693 |
| NA | ENSMUSG00 | 57259 |
| NA | ENSMUSG00 | 208967 |
| NA | ENSMUSG00 | 109154 |
| NA | ENSMUSG00 | 329659 |
| NA | ENSMUSG00 | 16002 |
| NA | ENSMUSG00 | 226101 |
| NA | ENSMUSG00 | 622976 |
| NA | ENSMUSG00 | 330695 |
| NA | ENSMUSG00 | 241624 |
| NA | ENSMUSG00 | 380664 |
| NA | ENSMUSG00 | 72440 |
| Significant | ENSMUSG00 | 233979 |
| NA | ENSMUSG00 | 237400 |
| NA | ENSMUSG00 | 76551 |
| NA | ENSMUSG00 | 52829 |
| NA | ENSMUSG00 | 97031 |
| NA | ENSMUSG00 | 70772 |
| NA | ENSMUSG00 | 217194 |
| Significant | ENSMUSG00 | 235631 |
| NA | ENSMUSG00 | 223722 |
| NA | ENSMUSG00 | 56484 |
| NA | ENSMUSG00 | 19944 |
| NA | ENSMUSG00 | 68777 |
| NA | ENSMUSG00 | 233571 |
| NA | ENSMUSG00 | 233805 |
| NA | ENSMUSG00 | 72023 |
| NA | ENSMUSG00 | 225523 |
| NA | ENSMUSG00 | 15977 |
| NA | ENSMUSG00 | 103710 |
| NA | ENSMUSG00 | 244646 |
| NA | ENSMUSG00 | 107305 |
| NA | ENSMUSG00 | 328133 |
| NA | ENSMUSG00 | 104910 |

| | | |
|---|---|---|
| NA | ENSMUSG00 | 226652 |
| NA | ENSMUSG00 | 213109 |
| NA | ENSMUSG00 | 192231 |
| NA | ENSMUSG00 | 209225 |
| NA | ENSMUSG00 | 207819 |
| NA | ENSMUSG00 | 51902 |
| NA | ENSMUSG00 | 13640 |
| NA | ENSMUSG00 | 243853 |
| NA | ENSMUSG00 | 71131 |
| NA | ENSMUSG00 | 108912 |
| NA | ENSMUSG00 | 76041 |
| NA | ENSMUSG00 | 101206 |
| NA | ENSMUSG00 | 109294 |
| NA | ENSMUSG00 | 234365 |
| Significant | ENSMUSG00 | 59048 |
| NA | ENSMUSG00 | 72522 |
| NA | ENSMUSG00 | 269799 |
| NA | ENSMUSG00 | 74238 |
| Significant | ENSMUSG00 | 71703 |
| NA | ENSMUSG00 | 78618 |
| NA | ENSMUSG00 | 107771 |
| NA | ENSMUSG00 | 225791 |
| NA | ENSMUSG00 | 20768 |
| Significant | ENSMUSG00 | 12772 |
| NA | ENSMUSG00 | 320808 |
| NA | ENSMUSG00 | 242297 |
| NA | ENSMUSG00 | NA |
| NA | ENSMUSG00 | 16447 |
| Significant | ENSMUSG00 | 12273 |
| Significant | ENSMUSG00 | 18175 |
| NA | ENSMUSG00 | 72672 |
| NA | ENSMUSG00 | 75029 |
| NA | ENSMUSG00 | 381511 |
| NA | ENSMUSG00 | 68140 |
| NA | ENSMUSG00 | NA |
| NA | ENSMUSG00 | 330216 |
| NA | ENSMUSG00 | 216792 |
| NA | ENSMUSG00 | 280411 |
| NA | ENSMUSG00 | 69890 |
| NA | ENSMUSG00 | 245828 |
| NA | ENSMUSG00 | 99890 |
| NA | ENSMUSG00 | 72454 |
| NA | ENSMUSG00 | 14345 |
| Significant | ENSMUSG00 | 20660 |
| NA | ENSMUSG00 | 22678 |
| NA | ENSMUSG00 | 237782 |

| | | |
|---|---|---|
| NA | ENSMUSG00 | 67956 |
| NA | ENSMUSG00 | 227298 |
| Significant | ENSMUSG00 | 74023 |
| NA | ENSMUSG00 | 71833 |
| NA | ENSMUSG00 | 16691 |
| NA | ENSMUSG00 | 276919 |
| NA | ENSMUSG00 | 72075 |
| NA | ENSMUSG00 | 109222 |
| NA | ENSMUSG00 | 58182 |
| Significant | ENSMUSG00 | 338363 |
| NA | ENSMUSG00 | 26466 |
| NA | ENSMUSG00 | 103172 |
| NA | ENSMUSG00 | 77951 |
| Significant | ENSMUSG00 | 93736 |
| NA | ENSMUSG00 | 66965 |
| NA | ENSMUSG00 | 329795 |
| NA | ENSMUSG00 | 69109 |
| NA | ENSMUSG00 | 209200 |
| NA | ENSMUSG00 | 212127 |
| NA | ENSMUSG00 | 101685 |
| NA | ENSMUSG00 | 66475 |
| NA | ENSMUSG00 | 104445 |
| NA | ENSMUSG00 | 73893 |
| NA | ENSMUSG00 | 50524 |
| NA | ENSMUSG00 | 237052 |
| NA | ENSMUSG00 | 80838 |
| NA | ENSMUSG00 | 56430 |
| NA | ENSMUSG00 | 20019 |
| NA | ENSMUSG00 | 381201 |
| NA | ENSMUSG00 | 22761 |
| NA | ENSMUSG00 | 244152 |
| NA | ENSMUSG00 | 52570 |
| NA | ENSMUSG00 | 232879 |
| NA | ENSMUSG00 | 52397 |
| NA | ENSMUSG00 | 227326 |
| NA | ENSMUSG00 | 235504 |
| Significant | ENSMUSG00 | 212937 |
| NA | ENSMUSG00 | 66367 |
| NA | ENSMUSG00 | 230119 |
| NA | ENSMUSG00 | 544971 |
| NA | ENSMUSG00 | 216549 |
| NA | ENSMUSG00 | 226123 |
| NA | ENSMUSG00 | 22666 |
| NA | ENSMUSG00 | 320225 |
| NA | ENSMUSG00 | 72046 |
| NA | ENSMUSG00 | 109305 |

| | | |
|---|---|---|
| NA | ENSMUSG00 | 338368 |
| NA | ENSMUSG00 | 244202 |
| NA | ENSMUSG00 | 319583 |
| NA | ENSMUSG00 | 53897 |
| NA | ENSMUSG00 | 17381 |
| NA | ENSMUSG00 | 244219 |
| NA | ENSMUSG00 | 22040 |
| NA | ENSMUSG00 | 233905 |
| NA | ENSMUSG00 | 76117 |
| NA | ENSMUSG00 | 66483 |
| NA | ENSMUSG00 | 319475 |
| NA | ENSMUSG00 | 224904 |
| NA | ENSMUSG00 | 64453 |
| NA | ENSMUSG00 | 19241 |
| NA | ENSMUSG00 | 14366 |
| NA | ENSMUSG00 | 70369 |
| NA | ENSMUSG00 | 58172 |
| NA | ENSMUSG00 | 100503043 |
| NA | ENSMUSG00 | 58996 |
| NA | ENSMUSG00 | 73873 |
| NA | ENSMUSG00 | 211389 |
| NA | ENSMUSG00 | 320982 |
| NA | ENSMUSG00 | 268857 |
| NA | ENSMUSG00 | 109263 |
| NA | ENSMUSG00 | NA |
| NA | ENSMUSG00 | 207965 |
| NA | ENSMUSG00 | 19416 |
| NA | ENSMUSG00 | 103765 |
| NA | ENSMUSG00 | 68939 |
| NA | ENSMUSG00 | 70458 |
| NA | ENSMUSG00 | 228368 |
| NA | ENSMUSG00 | 78134 |
| NA | ENSMUSG00 | 15270 |
| NA | ENSMUSG00 | 192198 |
| NA | ENSMUSG00 | 70804 |
| NA | ENSMUSG00 | 227522 |
| NA | ENSMUSG00 | 67291 |
| NA | ENSMUSG00 | 66242 |
| NA | ENSMUSG00 | 71801 |
| NA | ENSMUSG00 | 77318 |
| NA | ENSMUSG00 | 211228 |
| NA | ENSMUSG00 | 228966 |
| NA | ENSMUSG00 | 75731 |
| NA | ENSMUSG00 | 330096 |
| NA | ENSMUSG00 | 56305 |
| Significant | ENSMUSG00 | 231842 |

| | | |
|---|---|---|
| NA | ENSMUSG00 | 72065 |
| NA | ENSMUSG00 | 14202 |
| NA | ENSMUSG00 | 66958 |
| NA | ENSMUSG00 | 19716 |
| Significant | ENSMUSG00 | 229323 |
| NA | ENSMUSG00 | 67610 |
| NA | ENSMUSG00 | 67912 |
| NA | ENSMUSG00 | 319660 |
| NA | ENSMUSG00 | 217356 |
| NA | ENSMUSG00 | 14841 |
| NA | ENSMUSG00 | 237868 |
| NA | ENSMUSG00 | 210741 |
| NA | ENSMUSG00 | 229517 |
| NA | ENSMUSG00 | 14065 |
| NA | ENSMUSG00 | 54609 |
| NA | ENSMUSG00 | 229228 |
| NA | ENSMUSG00 | 116748 |
| NA | ENSMUSG00 | 208691 |
| NA | ENSMUSG00 | 107515 |
| NA | ENSMUSG00 | 230752 |
| NA | ENSMUSG00 | 76793 |
| NA | ENSMUSG00 | 67556 |
| NA | ENSMUSG00 | 12766 |
| NA | ENSMUSG00 | 58180 |
| Significant | ENSMUSG00 | 217995 |
| NA | ENSMUSG00 | 68525 |
| NA | ENSMUSG00 | 338369 |
| Significant | ENSMUSG00 | 244418 |
| NA | ENSMUSG00 | 83885 |
| NA | ENSMUSG00 | 78757 |
| NA | ENSMUSG00 | 106338 |
| NA | ENSMUSG00 | 76947 |
| NA | ENSMUSG00 | 72345 |
| NA | ENSMUSG00 | 16854 |
| Significant | ENSMUSG00 | 214639 |
| NA | ENSMUSG00 | 12642 |
| NA | ENSMUSG00 | 101113 |
| Significant | ENSMUSG00 | 56526 |
| NA | ENSMUSG00 | 16576 |
| NA | ENSMUSG00 | 97130 |
| NA | ENSMUSG00 | 278097 |
| NA | ENSMUSG00 | 106795 |
| NA | ENSMUSG00 | 243867 |
| NA | ENSMUSG00 | 320343 |
| NA | ENSMUSG00 | 109229 |
| NA | ENSMUSG00 | 74165 |

| | | |
|---|---|---|
| NA | ENSMUSG00 | 269233 |
| NA | ENSMUSG00 | NA |
| NA | ENSMUSG00 | 207921 |
| NA | ENSMUSG00 | 70617 |
| NA | ENSMUSG00 | 66096 |
| NA | ENSMUSG00 | 76832 |
| NA | ENSMUSG00 | 16500 |
| NA | ENSMUSG00 | 240832 |
| NA | ENSMUSG00 | 103806 |
| NA | ENSMUSG00 | 17386 |
| NA | ENSMUSG00 | 241303 |
| NA | ENSMUSG00 | 22719 |
| NA | ENSMUSG00 | 433771 |
| NA | ENSMUSG00 | 99334 |
| NA | ENSMUSG00 | 100043813 |
| Significant | ENSMUSG00 | 333433 |
| NA | ENSMUSG00 | 319370 |
| NA | ENSMUSG00 | 53951 |
| NA | ENSMUSG00 | 66717 |
| NA | ENSMUSG00 | 105787 |
| NA | ENSMUSG00 | 69662 |
| Significant | ENSMUSG00 | 14325 |
| NA | ENSMUSG00 | 320633 |
| NA | ENSMUSG00 | 102595 |
| NA | ENSMUSG00 | 71544 |
| NA | ENSMUSG00 | 22320 |
| NA | ENSMUSG00 | 64292 |
| Significant | ENSMUSG00 | 170706 |
| NA | ENSMUSG00 | 57895 |
| NA | ENSMUSG00 | 117592 |
| NA | ENSMUSG00 | 230249 |
| NA | ENSMUSG00 | 78834 |
| NA | ENSMUSG00 | 233057 |
| NA | ENSMUSG00 | 11958 |
| Significant | ENSMUSG00 | 212070 |
| NA | ENSMUSG00 | 230809 |
| NA | ENSMUSG00 | 69694 |
| NA | ENSMUSG00 | 383103 |
| NA | ENSMUSG00 | 237988 |
| NA | ENSMUSG00 | 71929 |
| NA | ENSMUSG00 | 654824 |
| NA | ENSMUSG00 | 238803 |
| NA | ENSMUSG00 | 74393 |
| Significant | ENSMUSG00 | 74442 |
| NA | ENSMUSG00 | 240186 |
| Significant | ENSMUSG00 | 14609 |

| | | |
|---|---|---|
| NA | ENSMUSG00 | 67911 |
| NA | ENSMUSG00 | 27356 |
| NA | ENSMUSG00 | 18750 |
| NA | ENSMUSG00 | 83557 |
| NA | ENSMUSG00 | 263764 |
| NA | ENSMUSG00 | 269952 |
| NA | ENSMUSG00 | 74777 |
| NA | ENSMUSG00 | 215772 |
| NA | ENSMUSG00 | 213350 |
| NA | ENSMUSG00 | 15476 |
| NA | ENSMUSG00 | 22648 |
| NA | ENSMUSG00 | 218311 |
| Significant | ENSMUSG00 | 70355 |
| NA | ENSMUSG00 | 72852 |
| NA | ENSMUSG00 | 75538 |
| NA | ENSMUSG00 | 258336 |
| NA | ENSMUSG00 | 317758 |
| NA | ENSMUSG00 | 17961 |
| NA | ENSMUSG00 | 12238 |
| NA | ENSMUSG00 | 319670 |
| NA | ENSMUSG00 | 101122 |
| NA | ENSMUSG00 | 66056 |
| NA | ENSMUSG00 | 67698 |
| Significant | ENSMUSG00 | 321019 |
| NA | ENSMUSG00 | 236930 |
| NA | ENSMUSG00 | 66882 |
| NA | ENSMUSG00 | 245695 |
| NA | ENSMUSG00 | 195040 |
| NA | ENSMUSG00 | 19823 |
| NA | ENSMUSG00 | 209334 |
| NA | ENSMUSG00 | 320183 |
| NA | ENSMUSG00 | 66962 |
| NA | ENSMUSG00 | 67767 |
| NA | ENSMUSG00 | 22037 |
| NA | ENSMUSG00 | 71643 |
| NA | ENSMUSG00 | 319263 |
| NA | ENSMUSG00 | 76800 |
| NA | ENSMUSG00 | 24074 |
| NA | ENSMUSG00 | 67885 |
| NA | ENSMUSG00 | 59015 |
| NA | ENSMUSG00 | 328232 |
| NA | ENSMUSG00 | 243219 |
| NA | ENSMUSG00 | 22710 |
| NA | ENSMUSG00 | 52055 |
| NA | ENSMUSG00 | 241075 |
| NA | ENSMUSG00 | 66701 |

| | | |
|---|---|---|
| NA | ENSMUSG00 | 22704 |
| NA | ENSMUSG00 | 17846 |
| NA | ENSMUSG00 | 227721 |
| NA | ENSMUSG00 | 70218 |
| NA | ENSMUSG00 | 71436 |
| NA | ENSMUSG00 | 81630 |
| NA | ENSMUSG00 | 22628 |
| NA | ENSMUSG00 | 119065974 |
| Significant | ENSMUSG00 | 56753 |
| NA | ENSMUSG00 | 232947 |
| NA | ENSMUSG00 | NA |
| NA | ENSMUSG00 | 54711 |
| NA | ENSMUSG00 | 216516 |
| NA | ENSMUSG00 | 329977 |
| NA | ENSMUSG00 | 12475 |
| NA | ENSMUSG00 | 241950 |
| NA | ENSMUSG00 | 233490 |
| NA | ENSMUSG00 | 56534 |
| Significant | ENSMUSG00 | 20737 |
| NA | ENSMUSG00 | 59057 |
| Significant | ENSMUSG00 | 12408 |
| NA | ENSMUSG00 | 270110 |
| NA | ENSMUSG00 | 21899 |
| NA | ENSMUSG00 | 70240 |
| NA | ENSMUSG00 | 625424 |
| NA | ENSMUSG00 | 233274 |
| NA | ENSMUSG00 | 545030 |
| NA | ENSMUSG00 | 17134 |
| NA | ENSMUSG00 | 58238 |
| NA | ENSMUSG00 | 230098 |
| NA | ENSMUSG00 | 66538 |
| NA | ENSMUSG00 | 433813 |
| Significant | ENSMUSG00 | 66684 |
| NA | ENSMUSG00 | 194401 |
| NA | ENSMUSG00 | 76108 |
| NA | ENSMUSG00 | 50709 |
| NA | ENSMUSG00 | 545184 |
| NA | ENSMUSG00 | 53625 |
| NA | ENSMUSG00 | 67892 |
| NA | ENSMUSG00 | 100737 |
| NA | ENSMUSG00 | 69807 |
| NA | ENSMUSG00 | 224840 |
| NA | ENSMUSG00 | 23983 |
| NA | ENSMUSG00 | 71599 |
| NA | ENSMUSG00 | 28194 |
| NA | ENSMUSG00 | 238037 |

| | | |
|---|---|---|
| NA | ENSMUSG00 | 233529 |
| NA | ENSMUSG00 | 75422 |
| NA | ENSMUSG00 | 320435 |
| NA | ENSMUSG00 | 66337 |
| NA | ENSMUSG00 | 66107 |
| NA | ENSMUSG00 | 22594 |
| NA | ENSMUSG00 | 328580 |
| NA | ENSMUSG00 | 333182 |
| NA | ENSMUSG00 | 20667 |
| NA | ENSMUSG00 | 72865 |
| NA | ENSMUSG00 | 11842 |
| NA | ENSMUSG00 | 17294 |
| NA | ENSMUSG00 | 223754 |
| NA | ENSMUSG00 | 271005 |
| NA | ENSMUSG00 | 72572 |
| Significant | ENSMUSG00 | 381694 |
| NA | ENSMUSG00 | 213389 |
| NA | ENSMUSG00 | 234086 |
| NA | ENSMUSG00 | 240667 |
| NA | ENSMUSG00 | 68936 |
| Significant | ENSMUSG00 | 208154 |
| NA | ENSMUSG00 | 380608 |
| NA | ENSMUSG00 | 109054 |
| NA | ENSMUSG00 | 50794 |
| NA | ENSMUSG00 | 228913 |
| NA | ENSMUSG00 | 76088 |
| NA | ENSMUSG00 | 51791 |
| Significant | ENSMUSG00 | 26384 |
| NA | ENSMUSG00 | 77604 |
| NA | ENSMUSG00 | 107976 |
| NA | ENSMUSG00 | 320484 |
| NA | ENSMUSG00 | 232314 |
| NA | ENSMUSG00 | 67097 |
| NA | ENSMUSG00 | 50784 |
| NA | ENSMUSG00 | 11480 |
| Significant | ENSMUSG00 | 104759 |
| NA | ENSMUSG00 | 15135 |
| NA | ENSMUSG00 | 403187 |
| NA | ENSMUSG00 | 15132 |
| NA | ENSMUSG00 | 52521 |
| NA | ENSMUSG00 | 108143 |
| NA | ENSMUSG00 | 243819 |
| NA | ENSMUSG00 | 72729 |
| NA | ENSMUSG00 | 78913 |
| NA | ENSMUSG00 | 74694 |
| Significant | ENSMUSG00 | 30791 |

| | | |
|---|---|---|
| NA | ENSMUSG00 | 329154 |
| NA | ENSMUSG00 | 13051 |
| NA | ENSMUSG00 | 76614 |
| NA | ENSMUSG00 | 380967 |
| NA | ENSMUSG00 | 224109 |
| Significant | ENSMUSG00 | 171282 |
| NA | ENSMUSG00 | 328370 |
| NA | ENSMUSG00 | 233549 |
| NA | ENSMUSG00 | 22350 |
| NA | ENSMUSG00 | 227656 |
| NA | ENSMUSG00 | 320226 |
| NA | ENSMUSG00 | 66520 |
| NA | ENSMUSG00 | 57370 |
| NA | ENSMUSG00 | 68944 |
| Significant | ENSMUSG00 | 110794 |
| NA | ENSMUSG00 | 234413 |
| NA | ENSMUSG00 | 56495 |
| NA | ENSMUSG00 | 11964 |
| NA | ENSMUSG00 | 27967 |
| NA | ENSMUSG00 | 78286 |
| NA | ENSMUSG00 | 70891 |
| NA | ENSMUSG00 | 227699 |
| NA | ENSMUSG00 | 18514 |
| NA | ENSMUSG00 | 99470 |
| NA | ENSMUSG00 | 20447 |
| NA | ENSMUSG00 | 209239 |
| Significant | ENSMUSG00 | 66871 |
| NA | ENSMUSG00 | 14957 |
| NA | ENSMUSG00 | 170833 |
| NA | ENSMUSG00 | 11491 |
| NA | ENSMUSG00 | 67441 |
| NA | ENSMUSG00 | 263406 |
| NA | ENSMUSG00 | 211914 |
| NA | ENSMUSG00 | 22644 |
| NA | ENSMUSG00 | 215693 |
| Significant | ENSMUSG00 | 215449 |
| NA | ENSMUSG00 | 16476 |
| NA | ENSMUSG00 | 226421 |
| Significant | ENSMUSG00 | 70549 |
| NA | ENSMUSG00 | 233833 |
| NA | ENSMUSG00 | 80748 |
| NA | ENSMUSG00 | 269023 |
| NA | ENSMUSG00 | 240776 |
| NA | ENSMUSG00 | 17755 |
| NA | ENSMUSG00 | 56451 |
| NA | ENSMUSG00 | 66875 |

| | | |
|---|---|---|
| NA | ENSMUSG00 | NA |
| NA | ENSMUSG00 | 58887 |
| NA | ENSMUSG00 | 224619 |
| NA | ENSMUSG00 | 319997 |
| NA | ENSMUSG00 | 232784 |
| NA | ENSMUSG00 | 246730 |
| NA | ENSMUSG00 | 14772 |
| NA | ENSMUSG00 | 72254 |
| NA | ENSMUSG00 | 103468 |
| NA | ENSMUSG00 | 320817 |
| NA | ENSMUSG00 | 58861 |
| NA | ENSMUSG00 | 56459 |
| NA | ENSMUSG00 | 16477 |
| NA | ENSMUSG00 | 52250 |
| NA | ENSMUSG00 | 18751 |
| NA | ENSMUSG00 | 108159 |
| NA | ENSMUSG00 | NA |
| NA | ENSMUSG00 | 16779 |
| NA | ENSMUSG00 | 545700 |
| Significant | ENSMUSG00 | 13110 |
| NA | ENSMUSG00 | 74026 |
| NA | ENSMUSG00 | 66315 |
| NA | ENSMUSG00 | 69724 |
| NA | ENSMUSG00 | 269037 |
| NA | ENSMUSG00 | 76454 |
| Significant | ENSMUSG00 | 226075 |
| NA | ENSMUSG00 | 217340 |
| NA | ENSMUSG00 | 14451 |
| NA | ENSMUSG00 | 66223 |
| NA | ENSMUSG00 | 50995 |
| NA | ENSMUSG00 | 231991 |
| NA | ENSMUSG00 | 57896 |
| NA | ENSMUSG00 | 64176 |
| NA | ENSMUSG00 | NA |
| NA | ENSMUSG00 | 68318 |
| Significant | ENSMUSG00 | 232413 |
| NA | ENSMUSG00 | 234912 |
| NA | ENSMUSG00 | 108899 |
| NA | ENSMUSG00 | 72635 |
| NA | ENSMUSG00 | 71667 |
| Significant | ENSMUSG00 | 353346 |
| NA | ENSMUSG00 | 12702 |
| NA | ENSMUSG00 | 66700 |
| NA | ENSMUSG00 | 213211 |
| NA | ENSMUSG00 | 72195 |
| NA | ENSMUSG00 | 19094 |

| | | |
|---|---|---|
| NA | ENSMUSG00 | 14159 |
| NA | ENSMUSG00 | 652986 |
| NA | ENSMUSG00 | 12051 |
| NA | ENSMUSG00 | 208595 |
| NA | ENSMUSG00 | 56772 |
| NA | ENSMUSG00 | 19153 |
| NA | ENSMUSG00 | 56291 |
| NA | ENSMUSG00 | 379043 |
| NA | ENSMUSG00 | 23863 |
| NA | ENSMUSG00 | 76273 |
| NA | ENSMUSG00 | 98685 |
| NA | ENSMUSG00 | 77591 |
| NA | ENSMUSG00 | 19342 |
| NA | ENSMUSG00 | 107939 |
| NA | ENSMUSG00 | 243373 |
| NA | ENSMUSG00 | 66212 |
| NA | ENSMUSG00 | 74748 |
| NA | ENSMUSG00 | 28295 |
| NA | ENSMUSG00 | 208718 |
| NA | ENSMUSG00 | 231630 |
| Significant | ENSMUSG00 | 245126 |
| NA | ENSMUSG00 | 74670 |
| NA | ENSMUSG00 | 240067 |
| NA | ENSMUSG00 | 236539 |
| NA | ENSMUSG00 | 52609 |
| NA | ENSMUSG00 | 26559 |
| NA | ENSMUSG00 | 26416 |
| NA | ENSMUSG00 | 66231 |
| NA | ENSMUSG00 | 56316 |
| NA | ENSMUSG00 | 104263 |
| NA | ENSMUSG00 | 21930 |
| NA | ENSMUSG00 | 21413 |
| NA | ENSMUSG00 | 30941 |
| NA | ENSMUSG00 | 230598 |
| NA | ENSMUSG00 | 83410 |
| NA | ENSMUSG00 | 73212 |
| Significant | ENSMUSG00 | 207818 |
| NA | ENSMUSG00 | 15936 |
| NA | ENSMUSG00 | 73830 |
| NA | ENSMUSG00 | 77097 |
| NA | ENSMUSG00 | 100494 |
| NA | ENSMUSG00 | 224691 |
| NA | ENSMUSG00 | 19895 |
| NA | ENSMUSG00 | 14218 |
| NA | ENSMUSG00 | 102442 |
| Significant | ENSMUSG00 | 110695 |

| | | |
|---|---|---|
| NA | ENSMUSG00 | 270802 |
| NA | ENSMUSG00 | 319446 |
| NA | ENSMUSG00 | 235584 |
| NA | ENSMUSG00 | 230770 |
| NA | ENSMUSG00 | 329384 |
| NA | ENSMUSG00 | 67772 |
| NA | ENSMUSG00 | 66075 |
| NA | ENSMUSG00 | 217779 |
| NA | ENSMUSG00 | 224111 |
| NA | ENSMUSG00 | 107371 |
| NA | ENSMUSG00 | 101612 |
| NA | ENSMUSG00 | 108058 |
| NA | ENSMUSG00 | 15042 |
| NA | ENSMUSG00 | 209586 |
| NA | ENSMUSG00 | 109658 |
| NA | ENSMUSG00 | 75540 |
| NA | ENSMUSG00 | 100043597 |
| Significant | ENSMUSG00 | 67547 |
| NA | ENSMUSG00 | 51798 |
| NA | ENSMUSG00 | 232087 |
| NA | ENSMUSG00 | 67311 |
| NA | ENSMUSG00 | 54151 |
| NA | ENSMUSG00 | 71994 |
| NA | ENSMUSG00 | 240442 |
| NA | ENSMUSG00 | 16855 |
| NA | ENSMUSG00 | 242202 |
| NA | ENSMUSG00 | 243906 |
| NA | ENSMUSG00 | 15531 |
| NA | ENSMUSG00 | 68346 |
| NA | ENSMUSG00 | 103466 |
| NA | ENSMUSG00 | 319955 |
| NA | ENSMUSG00 | 77940 |
| NA | ENSMUSG00 | 56460 |
| NA | ENSMUSG00 | 60440 |
| NA | ENSMUSG00 | 72668 |
| NA | ENSMUSG00 | 217109 |
| NA | ENSMUSG00 | 67704 |
| NA | ENSMUSG00 | 319653 |
| NA | ENSMUSG00 | 27401 |
| NA | ENSMUSG00 | 212073 |
| NA | ENSMUSG00 | 16596 |
| NA | ENSMUSG00 | 68588 |
| NA | ENSMUSG00 | 76022 |
| NA | ENSMUSG00 | 226695 |
| NA | ENSMUSG00 | 16904 |
| NA | ENSMUSG00 | 69786 |

| | | |
|---|---|---|
| NA | ENSMUSG00 | 70567 |
| Significant | ENSMUSG00 | 16880 |
| NA | ENSMUSG00 | 66251 |
| NA | ENSMUSG00 | 215476 |
| NA | ENSMUSG00 | 213438 |
| NA | ENSMUSG00 | 66266 |
| NA | ENSMUSG00 NA | |
| NA | ENSMUSG00 | 54451 |
| NA | ENSMUSG00 | 66292 |
| NA | ENSMUSG00 | 72504 |
| NA | ENSMUSG00 | 16534 |
| NA | ENSMUSG00 | 11852 |
| NA | ENSMUSG00 | 269997 |
| NA | ENSMUSG00 | 70481 |
| NA | ENSMUSG00 | 17248 |
| NA | ENSMUSG00 | 75458 |
| NA | ENSMUSG00 | 327978 |
| NA | ENSMUSG00 | 68598 |
| NA | ENSMUSG00 | 76687 |
| NA | ENSMUSG00 | 66500 |
| NA | ENSMUSG00 | 11983 |
| NA | ENSMUSG00 | 330189 |
| NA | ENSMUSG00 | 107526 |
| NA | ENSMUSG00 | 14797 |
| NA | ENSMUSG00 | 56491 |
| NA | ENSMUSG00 | 225010 |
| NA | ENSMUSG00 | 232078 |
| NA | ENSMUSG00 | 96957 |
| NA | ENSMUSG00 | 67379 |
| NA | ENSMUSG00 | 328417 |
| NA | ENSMUSG00 | 338364 |
| Significant | ENSMUSG00 | 237775 |
| NA | ENSMUSG00 | 24055 |
| NA | ENSMUSG00 | 94284 |
| NA | ENSMUSG00 | 231253 |
| NA | ENSMUSG00 | 106143 |
| NA | ENSMUSG00 | 225876 |
| NA | ENSMUSG00 | 17314 |
| NA | ENSMUSG00 | 70152 |
| NA | ENSMUSG00 | 22441 |
| NA | ENSMUSG00 | 20541 |
| NA | ENSMUSG00 | 66869 |
| NA | ENSMUSG00 | 231633 |
| Significant | ENSMUSG00 | 72244 |
| NA | ENSMUSG00 | 59308 |
| NA | ENSMUSG00 | 11487 |

| | | |
|---|---|---|
| NA | ENSMUSG00 | 252903 |
| NA | ENSMUSG00 | 70615 |
| NA | ENSMUSG00 | 232878 |
| NA | ENSMUSG00 | 244216 |
| NA | ENSMUSG00 | 97165 |
| NA | ENSMUSG00 | 100604 |
| NA | ENSMUSG00 | 106639 |
| NA | ENSMUSG00 | 218194 |
| NA | ENSMUSG00 | 110265 |
| NA | ENSMUSG00 | 319535 |
| Significant | ENSMUSG00 | 56086 |
| NA | ENSMUSG00 | 51960 |
| NA | ENSMUSG00 | 270201 |
| NA | ENSMUSG00 | 60595 |
| NA | ENSMUSG00 | 69727 |
| NA | ENSMUSG00 | 234135 |
| NA | ENSMUSG00 | 72341 |
| NA | ENSMUSG00 | 226255 |
| NA | ENSMUSG00 | 320237 |
| NA | ENSMUSG00 | 223881 |
| NA | ENSMUSG00 | 104401 |
| NA | ENSMUSG00 | 384763 |
| NA | ENSMUSG00 | 68055 |
| Significant | ENSMUSG00 | 20863 |
| NA | ENSMUSG00 | 71778 |
| NA | ENSMUSG00 | 544922 |
| NA | ENSMUSG00 | 11576 |
| NA | ENSMUSG00 | 215708 |
| NA | ENSMUSG00 | 71583 |
| NA | ENSMUSG00 | 239546 |
| NA | ENSMUSG00 | 347722 |
| NA | ENSMUSG00 | 328572 |
| NA | ENSMUSG00 | 66398 |
| NA | ENSMUSG00 | 54132 |
| NA | ENSMUSG00 | 18029 |
| NA | ENSMUSG00 | 67040 |
| NA | ENSMUSG00 | 69726 |
| NA | ENSMUSG00 | 270160 |
| NA | ENSMUSG00 | 667035 |
| NA | ENSMUSG00 | 11865 |
| NA | ENSMUSG00 | 68755 |
| NA | ENSMUSG00 | 192136 |
| NA | ENSMUSG00 | 16598 |
| NA | ENSMUSG00 | 330463 |
| NA | ENSMUSG00 | 50909 |
| NA | ENSMUSG00 | 108900 |

| | | |
|---|---|---|
| NA | ENSMUSG00 | 170742 |
| NA | ENSMUSG00 | 81702 |
| NA | ENSMUSG00 | NA |
| NA | ENSMUSG00 | 269633 |
| NA | ENSMUSG00 | 74287 |
| NA | ENSMUSG00 | 22643 |
| NA | ENSMUSG00 | 242474 |
| NA | ENSMUSG00 | 67568 |
| NA | ENSMUSG00 | 22755 |
| NA | ENSMUSG00 | 319207 |
| NA | ENSMUSG00 | 207352 |
| NA | ENSMUSG00 | 21961 |
| NA | ENSMUSG00 | 66069 |
| NA | ENSMUSG00 | 56324 |
| NA | ENSMUSG00 | 50762 |
| NA | ENSMUSG00 | 17132 |
| NA | ENSMUSG00 | 69207 |
| NA | ENSMUSG00 | 16423 |
| NA | ENSMUSG00 | 238690 |
| NA | ENSMUSG00 | 320706 |
| NA | ENSMUSG00 | 226169 |
| NA | ENSMUSG00 | 432508 |
| NA | ENSMUSG00 | 71918 |
| NA | ENSMUSG00 | 52855 |
| Significant | ENSMUSG00 | 276891 |
| NA | ENSMUSG00 | 75620 |
| NA | ENSMUSG00 | 15126 |
| Significant | ENSMUSG00 | 66953 |
| NA | ENSMUSG00 | 330671 |
| NA | ENSMUSG00 | 68992 |
| NA | ENSMUSG00 | 207952 |
| NA | ENSMUSG00 | 14734 |
| NA | ENSMUSG00 | 76781 |
| NA | ENSMUSG00 | 195018 |
| NA | ENSMUSG00 | 59042 |
| NA | ENSMUSG00 | 224019 |
| NA | ENSMUSG00 | 234378 |
| NA | ENSMUSG00 | 69459 |
| NA | ENSMUSG00 | 66922 |
| NA | ENSMUSG00 | 54561 |
| NA | ENSMUSG00 | 14600 |
| NA | ENSMUSG00 | 223732 |
| NA | ENSMUSG00 | 67242 |
| NA | ENSMUSG00 | 66656 |
| NA | ENSMUSG00 | 26874 |
| NA | ENSMUSG00 | 57778 |

| | | |
|---|---|---|
| NA | ENSMUSG00 | 116871 |
| NA | ENSMUSG00 | 22640 |
| NA | ENSMUSG00 | 67673 |
| NA | ENSMUSG00 | 66510 |
| NA | ENSMUSG00 | 71564 |
| NA | ENSMUSG00 | 18627 |
| NA | ENSMUSG00 | 104806 |
| NA | ENSMUSG00 NA | |
| NA | ENSMUSG00 | 230657 |
| NA | ENSMUSG00 | 232086 |
| NA | ENSMUSG00 | 246196 |
| NA | ENSMUSG00 | 231326 |
| NA | ENSMUSG00 | 26383 |
| NA | ENSMUSG00 | 73024 |
| NA | ENSMUSG00 | 208820 |
| NA | ENSMUSG00 | 22757 |
| NA | ENSMUSG00 | 257632 |
| NA | ENSMUSG00 | 243308 |
| NA | ENSMUSG00 | 236193 |
| NA | ENSMUSG00 | 338366 |
| Significant | ENSMUSG00 | 20201 |
| NA | ENSMUSG00 | 20215 |
| NA | ENSMUSG00 | 223433 |
| NA | ENSMUSG00 | 20202 |
| NA | ENSMUSG00 | 27979 |
| NA | ENSMUSG00 | 20454 |
| NA | ENSMUSG00 | 15039 |
| NA | ENSMUSG00 | 225020 |
| NA | ENSMUSG00 | 56386 |
| NA | ENSMUSG00 | 225471 |
| NA | ENSMUSG00 | 109785 |
| NA | ENSMUSG00 | 94094 |
| NA | ENSMUSG00 | 54607 |
| NA | ENSMUSG00 | 78893 |
| NA | ENSMUSG00 | 225861 |
| NA | ENSMUSG00 | 12631 |
| NA | ENSMUSG00 | 66522 |
| NA | ENSMUSG00 | 18150 |
| NA | ENSMUSG00 | 226412 |
| NA | ENSMUSG00 | 93737 |
| NA | ENSMUSG00 | 12611 |
| NA | ENSMUSG00 | 18783 |
| NA | ENSMUSG00 | 71941 |
| NA | ENSMUSG00 | 27057 |
| NA | ENSMUSG00 | 321000 |
| NA | ENSMUSG00 | 68121 |

| | | |
|---|---|---|
| NA | ENSMUSG00 | 329260 |
| NA | ENSMUSG00 | 60361 |
| NA | ENSMUSG00 | 100041433 |
| NA | ENSMUSG00 | 28035 |
| NA | ENSMUSG00 | 100929 |
| NA | ENSMUSG00 | 69068 |
| NA | ENSMUSG00 | 17847 |
| NA | ENSMUSG00 | 242894 |
| NA | ENSMUSG00 | 233168 |
| NA | ENSMUSG00 | 16881 |
| NA | ENSMUSG00 | 231821 |
| NA | ENSMUSG00 | 22134 |
| NA | ENSMUSG00 | 26448 |
| NA | ENSMUSG00 | 109929 |
| NA | ENSMUSG00 | 329650 |
| NA | ENSMUSG00 | 108699 |
| NA | ENSMUSG00 | 17425 |
| Significant | ENSMUSG00 | 320782 |
| NA | ENSMUSG00 | 12608 |
| Significant | ENSMUSG00 | 106572 |
| NA | ENSMUSG00 | 19204 |
| NA | ENSMUSG00 | 73254 |
| NA | ENSMUSG00 | 27392 |
| NA | ENSMUSG00 | 19820 |
| NA | ENSMUSG00 | 210104 |
| NA | ENSMUSG00 | 74665 |
| NA | ENSMUSG00 | 320365 |
| NA | ENSMUSG00 | 109151 |
| NA | ENSMUSG00 | 18938 |
| NA | ENSMUSG00 | NA |
| NA | ENSMUSG00 | 67442 |
| NA | ENSMUSG00 | 77619 |
| NA | ENSMUSG00 | 20592 |
| NA | ENSMUSG00 | 224647 |
| NA | ENSMUSG00 | 232089 |
| NA | ENSMUSG00 | 15939 |
| NA | ENSMUSG00 | 235627 |
| NA | ENSMUSG00 | 12332 |
| Significant | ENSMUSG00 | 18030 |
| NA | ENSMUSG00 | 15364 |
| NA | ENSMUSG00 | 211660 |
| NA | ENSMUSG00 | 52690 |
| NA | ENSMUSG00 | 53424 |
| NA | ENSMUSG00 | 264134 |
| NA | ENSMUSG00 | 18521 |
| NA | ENSMUSG00 | 70676 |

| | | |
|---|---|---|
| NA | ENSMUSG00 | 73748 |
| Significant | ENSMUSG00 | 73690 |
| NA | ENSMUSG00 | 78303 |
| NA | ENSMUSG00 | 93757 |
| NA | ENSMUSG00 | 434428 |
| NA | ENSMUSG00 | 20469 |
| NA | ENSMUSG00 | 382090 |
| NA | ENSMUSG00 | 67131 |
| NA | ENSMUSG00 | 99311 |
| NA | ENSMUSG00 | 381801 |
| NA | ENSMUSG00 | 107817 |
| NA | ENSMUSG00 | 15925 |
| NA | ENSMUSG00 | 665189 |
| NA | ENSMUSG00 | 67475 |
| NA | ENSMUSG00 | 112415 |
| NA | ENSMUSG00 | 210135 |
| NA | ENSMUSG00 | 66116 |
| NA | ENSMUSG00 | 26920 |
| NA | ENSMUSG00 | 18148 |
| NA | ENSMUSG00 | 27366 |
| NA | ENSMUSG00 | 77945 |
| NA | ENSMUSG00 | 71389 |
| NA | ENSMUSG00 | 211488 |
| NA | ENSMUSG00 | 327957 |
| Significant | ENSMUSG00 | 68487 |
| NA | ENSMUSG00 | 319236 |
| NA | ENSMUSG00 | 66632 |
| NA | ENSMUSG00 | 239099 |
| NA | ENSMUSG00 | 233899 |
| NA | ENSMUSG00 | 606496 |
| NA | ENSMUSG00 | 70591 |
| NA | ENSMUSG00 | 382062 |
| NA | ENSMUSG00 | 68682 |
| NA | ENSMUSG00 | 276905 |
| NA | ENSMUSG00 | 66349 |
| NA | ENSMUSG00 | 269774 |
| NA | ENSMUSG00 | 76894 |
| NA | ENSMUSG00 | 19646 |
| NA | ENSMUSG00 | 108169072 |
| NA | ENSMUSG00 | 68011 |
| NA | ENSMUSG00 | 20446 |
| NA | ENSMUSG00 | 231532 |
| NA | ENSMUSG00 | 67671 |
| NA | ENSMUSG00 | 12043 |
| NA | ENSMUSG00 | 545389 |
| NA | ENSMUSG00 | 56632 |

| | | |
|---|---|---|
| NA | ENSMUSG00 | 67883 |
| NA | ENSMUSG00 | 11797 |
| NA | ENSMUSG00 | 230584 |
| NA | ENSMUSG00 | 67681 |
| NA | ENSMUSG00 | 268670 |
| NA | ENSMUSG00 | 107823 |
| NA | ENSMUSG00 | 24132 |
| NA | ENSMUSG00 | 214917 |
| NA | ENSMUSG00 | 76130 |
| NA | ENSMUSG00 | 75739 |
| NA | ENSMUSG00 | 50496 |
| NA | ENSMUSG00 NA | |
| NA | ENSMUSG00 | 73689 |
| NA | ENSMUSG00 | 20747 |
| NA | ENSMUSG00 | 230857 |
| NA | ENSMUSG00 | 94245 |
| NA | ENSMUSG00 | 78697 |
| NA | ENSMUSG00 | 244713 |
| NA | ENSMUSG00 | 56048 |
| NA | ENSMUSG00 | 13664 |
| NA | ENSMUSG00 | 67106 |
| NA | ENSMUSG00 | 70317 |
| NA | ENSMUSG00 | 209387 |
| NA | ENSMUSG00 | 14677 |
| NA | ENSMUSG00 | 110593 |
| NA | ENSMUSG00 | 105246 |
| NA | ENSMUSG00 | 14433 |
| NA | ENSMUSG00 | 232946 |
| NA | ENSMUSG00 | 320795 |
| NA | ENSMUSG00 | 69228 |
| NA | ENSMUSG00 | 108797 |
| Significant | ENSMUSG00 | 19152 |
| NA | ENSMUSG00 | 20740 |
| NA | ENSMUSG00 | 225187 |
| NA | ENSMUSG00 | 192986 |
| NA | ENSMUSG00 | 234374 |
| NA | ENSMUSG00 | 12018 |
| NA | ENSMUSG00 | 104349 |
| NA | ENSMUSG00 | 22445 |
| NA | ENSMUSG00 | 19951 |
| NA | ENSMUSG00 | 218314 |
| NA | ENSMUSG00 | 76539 |
| NA | ENSMUSG00 | 54217 |
| NA | ENSMUSG00 | 67230 |
| NA | ENSMUSG00 | 22646 |
| NA | ENSMUSG00 | 12296 |

| | | |
|---|---|---|
| NA | ENSMUSG00 | 70450 |
| NA | ENSMUSG00 | 217837 |
| NA | ENSMUSG00 | 235047 |
| NA | ENSMUSG00 | NA |
| NA | ENSMUSG00 | 100019 |
| NA | ENSMUSG00 | 52398 |
| NA | ENSMUSG00 | 109254 |
| NA | ENSMUSG00 | 214106 |
| NA | ENSMUSG00 | 72333 |
| NA | ENSMUSG00 | 68519 |
| NA | ENSMUSG00 | 66052 |
| NA | ENSMUSG00 | 100416706 |
| NA | ENSMUSG00 | 74039 |
| NA | ENSMUSG00 | 621054 |
| NA | ENSMUSG00 | 14862 |
| NA | ENSMUSG00 | 432555 |
| NA | ENSMUSG00 | 170748 |
| NA | ENSMUSG00 | 381438 |
| NA | ENSMUSG00 | 100041379 |
| NA | ENSMUSG00 | 244721 |
| NA | ENSMUSG00 | 232906 |
| NA | ENSMUSG00 | 22282 |
| NA | ENSMUSG00 | 66609 |
| NA | ENSMUSG00 | 21912 |
| NA | ENSMUSG00 | 319554 |
| NA | ENSMUSG00 | 64659 |
| NA | ENSMUSG00 | 105988 |
| NA | ENSMUSG00 | 24135 |
| NA | ENSMUSG00 | 71567 |
| NA | ENSMUSG00 | 19704 |
| NA | ENSMUSG00 | 218793 |
| NA | ENSMUSG00 | 192285 |
| NA | ENSMUSG00 | 330662 |
| NA | ENSMUSG00 | 22746 |
| NA | ENSMUSG00 | 66487 |
| NA | ENSMUSG00 | 24015 |
| NA | ENSMUSG00 | 319181 |
| NA | ENSMUSG00 | 18685 |
| Significant | ENSMUSG00 | 72462 |
| NA | ENSMUSG00 | 233058 |
| NA | ENSMUSG00 | 98258 |
| NA | ENSMUSG00 | 20310 |
| NA | ENSMUSG00 | 18181 |
| NA | ENSMUSG00 | 100043346 |
| NA | ENSMUSG00 | 23938 |
| NA | ENSMUSG00 | 387524 |

| | | |
|---|---|---|
| NA | ENSMUSG00 | NA |
| NA | ENSMUSG00 | 13360 |
| NA | ENSMUSG00 | 666926 |
| NA | ENSMUSG00 | 101240 |
| NA | ENSMUSG00 | 20282 |
| NA | ENSMUSG00 | 68152 |
| NA | ENSMUSG00 | NA |
| NA | ENSMUSG00 | NA |
| NA | ENSMUSG00 | 100503670 |
| NA | ENSMUSG00 | 67511 |
| NA | ENSMUSG00 | 68607 |
| NA | ENSMUSG00 | 50875 |
| NA | ENSMUSG00 | 50755 |
| NA | ENSMUSG00 | 19946 |
| NA | ENSMUSG00 | 100042670 |
| Significant | ENSMUSG00 | 14544 |
| NA | ENSMUSG00 | 100191011 |
| NA | ENSMUSG00 | 65020 |
| NA | ENSMUSG00 | 75705 |
| NA | ENSMUSG00 | 22151 |
| NA | ENSMUSG00 | 72972 |
| NA | ENSMUSG00 | 76890 |
| NA | ENSMUSG00 | 68364 |
| NA | ENSMUSG00 | 112406 |
| NA | ENSMUSG00 | 14127 |
| NA | ENSMUSG00 | 387565 |
| NA | ENSMUSG00 | 72568 |
| NA | ENSMUSG00 | 211480 |
| NA | ENSMUSG00 | 233987 |
| NA | ENSMUSG00 | 18413 |
| NA | ENSMUSG00 | 21833 |
| NA | ENSMUSG00 | 108937 |
| NA | ENSMUSG00 | 56702 |
| NA | ENSMUSG00 | 110911 |
| Significant | ENSMUSG00 | 18022 |
| NA | ENSMUSG00 | 53605 |
| NA | ENSMUSG00 | NA |
| NA | ENSMUSG00 | 18733 |
| NA | ENSMUSG00 | 66462 |
| NA | ENSMUSG00 | 11308 |
| NA | ENSMUSG00 | 619900 |
| NA | ENSMUSG00 | 329003 |
| NA | ENSMUSG00 | 432769 |
| NA | ENSMUSG00 | 54006 |
| NA | ENSMUSG00 | 380855 |
| NA | ENSMUSG00 | 81799 |

| | | |
|---|---|---|
| NA | ENSMUSG00 | NA |
| NA | ENSMUSG00 | 214932 |
| NA | ENSMUSG00 | 219189 |
| NA | ENSMUSG00 | 229279 |
| NA | ENSMUSG00 | 27387 |
| NA | ENSMUSG00 | 333088 |
| NA | ENSMUSG00 | 433182 |
| NA | ENSMUSG00 | 19363 |
| NA | ENSMUSG00 | 19899 |
| NA | ENSMUSG00 | 246256 |
| Significant | ENSMUSG00 | 213002 |
| NA | ENSMUSG00 | 17955 |
| NA | ENSMUSG00 | 240041 |
| NA | ENSMUSG00 | 213006 |
| NA | ENSMUSG00 | 54353 |
| NA | ENSMUSG00 | 69606 |
| NA | ENSMUSG00 | NA |
| NA | ENSMUSG00 | 76936 |
| NA | ENSMUSG00 | 53860 |
| NA | ENSMUSG00 | 74996 |
| NA | ENSMUSG00 | 330474 |
| NA | ENSMUSG00 | 78304 |
| NA | ENSMUSG00 | 12593 |
| NA | ENSMUSG00 | NA |
| NA | ENSMUSG00 | 22362 |
| NA | ENSMUSG00 | 26569 |
| NA | ENSMUSG00 | 72749 |
| Significant | ENSMUSG00 | 12982 |
| NA | ENSMUSG00 | 108052 |
| NA | ENSMUSG00 | 414077 |
| NA | ENSMUSG00 | 14297 |
| NA | ENSMUSG00 | 66707 |
| NA | ENSMUSG00 | 21770 |
| Significant | ENSMUSG00 | 11936 |
| NA | ENSMUSG00 | NA |
| NA | ENSMUSG00 | 242747 |
| NA | ENSMUSG00 | 17187 |
| NA | ENSMUSG00 | 192197 |
| NA | ENSMUSG00 | 231086 |
| Significant | ENSMUSG00 | 19229 |
| NA | ENSMUSG00 | NA |
| NA | ENSMUSG00 | 103537 |
| NA | ENSMUSG00 | 235028 |
| NA | ENSMUSG00 | 232984 |
| NA | ENSMUSG00 | 67157 |
| NA | ENSMUSG00 | 210973 |

| | | |
|---|---|---|
| NA | ENSMUSG00 | 69632 |
| Significant | ENSMUSG00 | 14131 |
| NA | ENSMUSG00 | 70103 |
| NA | ENSMUSG00 | 66152 |
| NA | ENSMUSG00 | 21400 |
| NA | ENSMUSG00 | 22059 |
| NA | ENSMUSG00 | 215814 |
| NA | ENSMUSG00 | 227612 |
| NA | ENSMUSG00 | 68501 |
| NA | ENSMUSG00 | 54598 |
| Significant | ENSMUSG00 | 268885 |
| NA | ENSMUSG00 | 21340 |
| NA | ENSMUSG00 | 232337 |
| NA | ENSMUSG00 | 338351 |
| NA | ENSMUSG00 | 53902 |
| NA | ENSMUSG00 | 14251 |
| NA | ENSMUSG00 | 225887 |
| NA | ENSMUSG00 | 110196 |
| NA | ENSMUSG00 | 668144 |
| NA | ENSMUSG00 | 75764 |
| NA | ENSMUSG00 | NA |
| NA | ENSMUSG00 | 100040416 |
| NA | ENSMUSG00 | 106582 |
| NA | ENSMUSG00 | 13681 |
| NA | ENSMUSG00 | 50780 |
| NA | ENSMUSG00 | 56298 |
| NA | ENSMUSG00 | 270156 |
| NA | ENSMUSG00 | 13170 |
| NA | ENSMUSG00 | 67161 |
| NA | ENSMUSG00 | 408067 |
| NA | ENSMUSG00 | 228807 |
| NA | ENSMUSG00 | 232811 |
| NA | ENSMUSG00 | 231130 |
| NA | ENSMUSG00 | 67255 |
| NA | ENSMUSG00 | 266632 |
| NA | ENSMUSG00 | 140630 |
| NA | ENSMUSG00 | 19245 |
| NA | ENSMUSG00 | 234358 |
| NA | ENSMUSG00 | 94346 |
| NA | ENSMUSG00 | 67609 |
| NA | ENSMUSG00 | 14784 |
| NA | ENSMUSG00 | 230996 |
| NA | ENSMUSG00 | 71869 |
| NA | ENSMUSG00 | 15512 |
| NA | ENSMUSG00 | 72723 |
| NA | ENSMUSG00 | 381921 |

| | | |
|---|---|---|
| NA | ENSMUSG00 | 229499 |
| NA | ENSMUSG00 | 217218 |
| NA | ENSMUSG00 | 212862 |
| NA | ENSMUSG00 | 16554 |
| NA | ENSMUSG00 | 232440 |
| NA | ENSMUSG00 | 244218 |
| NA | ENSMUSG00 | 27367 |
| NA | ENSMUSG00 | 330817 |
| NA | ENSMUSG00 | 11690 |
| NA | ENSMUSG00 | 19167 |
| NA | ENSMUSG00 | 19889 |
| NA | ENSMUSG00 | NA |
| NA | ENSMUSG00 | 214572 |
| NA | ENSMUSG00 | 66603 |
| NA | ENSMUSG00 | 22070 |
| Significant | ENSMUSG00 | 241633 |
| Significant | ENSMUSG00 | 20719 |
| NA | ENSMUSG00 | 117109 |
| NA | ENSMUSG00 | 27801 |
| NA | ENSMUSG00 | 215436 |
| NA | ENSMUSG00 | 100042180 |
| NA | ENSMUSG00 | 216869 |
| Significant | ENSMUSG00 | 74580 |
| NA | ENSMUSG00 | 319996 |
| NA | ENSMUSG00 | 56009 |
| NA | ENSMUSG00 | 101631 |
| NA | ENSMUSG00 | 14886 |
| NA | ENSMUSG00 | 11771 |
| NA | ENSMUSG00 | 170574 |
| NA | ENSMUSG00 | 66101 |
| NA | ENSMUSG00 | 72128 |
| NA | ENSMUSG00 | 245174 |
| NA | ENSMUSG00 | 15381 |
| NA | ENSMUSG00 | 12039 |
| NA | ENSMUSG00 | 243833 |
| NA | ENSMUSG00 | NA |
| NA | ENSMUSG00 | 234362 |
| NA | ENSMUSG00 | 20649 |
| NA | ENSMUSG00 | 667823 |
| NA | ENSMUSG00 | 56736 |
| Significant | ENSMUSG00 | 54672 |
| NA | ENSMUSG00 | 60532 |
| NA | ENSMUSG00 | 108960 |
| NA | ENSMUSG00 | 23832 |
| NA | ENSMUSG00 | 77519 |
| NA | ENSMUSG00 | 76261 |

| | | |
|---|---|---|
| NA | ENSMUSG00 | 30935 |
| NA | ENSMUSG00 | 68742 |
| NA | ENSMUSG00 | 29820 |
| NA | ENSMUSG00 | 226610 |
| NA | ENSMUSG00 | 17150 |
| NA | ENSMUSG00 | 14198 |
| Significant | ENSMUSG00 | 80876 |
| NA | ENSMUSG00 | 244864 |
| NA | ENSMUSG00 | 13808 |
| NA | ENSMUSG00 | 22260 |
| NA | ENSMUSG00 | 57808 |
| NA | ENSMUSG00 | 319158 |
| NA | ENSMUSG00 | 223989 |
| NA | ENSMUSG00 | 54667 |
| NA | ENSMUSG00 | 225845 |
| NA | ENSMUSG00 | 319155 |
| NA | ENSMUSG00 | 69527 |
| NA | ENSMUSG00 | 236794 |
| NA | ENSMUSG00 | 66205 |
| NA | ENSMUSG00 | 117197 |
| NA | ENSMUSG00 | 69718 |
| NA | ENSMUSG00 | 59050 |
| NA | ENSMUSG00 | 15078 |
| NA | ENSMUSG00 | 211484 |
| NA | ENSMUSG00 | 63986 |
| NA | ENSMUSG00 | 380614 |
| NA | ENSMUSG00 | 12010 |
| Significant | ENSMUSG00 | 14870 |
| NA | ENSMUSG00 | 77891 |
| NA | ENSMUSG00 | 230848 |
| NA | ENSMUSG00 | 104303 |
| NA | ENSMUSG00 | 75572 |
| NA | ENSMUSG00 | 103266 |
| NA | ENSMUSG00 | 19941 |
| NA | ENSMUSG00 | 328162 |
| Significant | ENSMUSG00 | 214575 |
| NA | ENSMUSG00 | 56447 |
| NA | ENSMUSG00 | 59014 |
| NA | ENSMUSG00 | 53609 |
| NA | ENSMUSG00 | 18114 |
| NA | ENSMUSG00 | 68977 |
| NA | ENSMUSG00 | 12560 |
| NA | ENSMUSG00 | 20841 |
| NA | ENSMUSG00 | 56096 |
| NA | ENSMUSG00 | 100041953 |
| NA | ENSMUSG00 | 192173 |

| | | |
|---|---|---|
| NA | ENSMUSG00 | 66114 |
| NA | ENSMUSG00 | 72461 |
| NA | ENSMUSG00 | 19043 |
| Significant | ENSMUSG00 | 17060 |
| NA | ENSMUSG00 | 56194 |
| NA | ENSMUSG00 | 433586 |
| NA | ENSMUSG00 | 100034724 |
| NA | ENSMUSG00 | 329679 |
| NA | ENSMUSG00 | 353282 |
| NA | ENSMUSG00 | 54402 |
| NA | ENSMUSG00 | 14972 |
| NA | ENSMUSG00 | 105504 |
| NA | ENSMUSG00 | 236792 |
| NA | ENSMUSG00 | 27998 |
| NA | ENSMUSG00 | 330177 |
| NA | ENSMUSG00 | 72055 |
| NA | ENSMUSG00 | 72108 |
| NA | ENSMUSG00 | 17938 |
| NA | ENSMUSG00 | 68922 |
| NA | ENSMUSG00 | NA |
| NA | ENSMUSG00 | 20315 |
| NA | ENSMUSG00 | 68479 |
| NA | ENSMUSG00 | 408062 |
| NA | ENSMUSG00 | 23877 |
| NA | ENSMUSG00 | 11481 |
| NA | ENSMUSG00 | 142682 |
| NA | ENSMUSG00 | 329540 |
| NA | ENSMUSG00 | 381812 |
| NA | ENSMUSG00 | 15258 |
| NA | ENSMUSG00 | 67727 |
| NA | ENSMUSG00 | 217431 |
| NA | ENSMUSG00 | 66278 |
| NA | ENSMUSG00 | 66128 |
| NA | ENSMUSG00 | 20115 |
| NA | ENSMUSG00 | 53607 |
| NA | ENSMUSG00 | 319156 |
| NA | ENSMUSG00 | 12859 |
| NA | ENSMUSG00 | 625286 |
| NA | ENSMUSG00 | 75216 |
| NA | ENSMUSG00 | 215474 |
| NA | ENSMUSG00 | 381067 |
| NA | ENSMUSG00 | 66317 |
| NA | ENSMUSG00 | 382045 |
| NA | ENSMUSG00 | 208266 |
| NA | ENSMUSG00 | 240087 |
| NA | ENSMUSG00 | 108121 |

| | | |
|---|---|---|
| NA | ENSMUSG00 | 319172 |
| NA | ENSMUSG00 | 192191 |
| NA | ENSMUSG00 | 12488 |
| NA | ENSMUSG00 | 66569 |
| NA | ENSMUSG00 | 228836 |
| NA | ENSMUSG00 | 320208 |
| NA | ENSMUSG00 | 14042 |
| NA | ENSMUSG00 | 545156 |
| NA | ENSMUSG00 | 665775 |
| NA | ENSMUSG00 | 67861 |
| NA | ENSMUSG00 | 73419 |
| Significant | ENSMUSG00 | 76763 |
| NA | ENSMUSG00 | 20068 |
| NA | ENSMUSG00 | 22139 |
| NA | ENSMUSG00 | 622313 |
| NA | ENSMUSG00 | 20917 |
| NA | ENSMUSG00 | 12695 |
| NA | ENSMUSG00 | 208908 |
| NA | ENSMUSG00 | 72475 |
| NA | ENSMUSG00 | 269585 |
| NA | ENSMUSG00 | 57782 |
| NA | ENSMUSG00 | 18674 |
| NA | ENSMUSG00 | 70351 |
| NA | ENSMUSG00 | 94254 |
| NA | ENSMUSG00 | 14252 |
| NA | ENSMUSG00 | 20042 |
| NA | ENSMUSG00 | 319173 |
| NA | ENSMUSG00 | NA |
| NA | ENSMUSG00 | 22654 |
| NA | ENSMUSG00 | 63985 |
| NA | ENSMUSG00 | 67928 |
| NA | ENSMUSG00 | 212974 |
| NA | ENSMUSG00 | NA |
| NA | ENSMUSG00 | 22689 |
| NA | ENSMUSG00 | 67732 |
| NA | ENSMUSG00 | 18655 |
| NA | ENSMUSG00 | 327743 |
| NA | ENSMUSG00 | 16907 |
| NA | ENSMUSG00 | 19317 |
| NA | ENSMUSG00 | 239849 |
| NA | ENSMUSG00 | 228662 |
| NA | ENSMUSG00 | 240120 |
| NA | ENSMUSG00 | 212986 |
| NA | ENSMUSG00 | 19377 |
| NA | ENSMUSG00 | 232853 |
| NA | ENSMUSG00 | 80281 |

| | | |
|---|---|---|
| NA | ENSMUSG00 | 98417 |
| NA | ENSMUSG00 | 207854 |
| NA | ENSMUSG00 | 228839 |
| NA | ENSMUSG00 | 71835 |
| NA | ENSMUSG00 | 72658 |
| NA | ENSMUSG00 | 224671 |
| NA | ENSMUSG00 | 14852 |
| NA | ENSMUSG00 | 106869 |
| NA | ENSMUSG00 | 76089 |
| NA | ENSMUSG00 | 231580 |
| NA | ENSMUSG00 | 66197 |
| NA | ENSMUSG00 | NA |
| NA | ENSMUSG00 | 19294 |
| NA | ENSMUSG00 | 14659 |
| NA | ENSMUSG00 | 319195 |
| NA | ENSMUSG00 | 18788 |
| NA | ENSMUSG00 | NA |
| NA | ENSMUSG00 | 100034726 |
| NA | ENSMUSG00 | 74868 |
| NA | ENSMUSG00 | 66439 |
| NA | ENSMUSG00 | 22152 |
| Significant | ENSMUSG00 | 434624 |
| NA | ENSMUSG00 | 68036 |
| NA | ENSMUSG00 | 11841 |
| NA | ENSMUSG00 | 100042823 |
| NA | ENSMUSG00 | 667422 |
| NA | ENSMUSG00 | 381318 |
| NA | ENSMUSG00 | 11770 |
| NA | ENSMUSG00 | 272347 |
| NA | ENSMUSG00 | 225651 |
| NA | ENSMUSG00 | 384059 |
| NA | ENSMUSG00 | 21854 |
| NA | ENSMUSG00 | 12802 |
| NA | ENSMUSG00 | 78795 |
| NA | ENSMUSG00 | 22153 |
| Significant | ENSMUSG00 | 14728 |
| NA | ENSMUSG00 | 20817 |
| NA | ENSMUSG00 | 67101 |
| NA | ENSMUSG00 | 320713 |
| NA | ENSMUSG00 | 76051 |
| NA | ENSMUSG00 | 27176 |
| NA | ENSMUSG00 | 14299 |
| NA | ENSMUSG00 | 67942 |
| NA | ENSMUSG00 | 68554 |
| NA | ENSMUSG00 | 319184 |
| NA | ENSMUSG00 | 19044 |

| | | |
|---|---|---|
| NA | ENSMUSG00 | 210503 |
| NA | ENSMUSG00 | 106672 |
| NA | ENSMUSG00 | 677073 |
| NA | ENSMUSG00 | 269639 |
| NA | ENSMUSG00 | 13663 |
| NA | ENSMUSG00 | 235048 |
| NA | ENSMUSG00 | 67669 |
| NA | ENSMUSG00 | 67392 |
| NA | ENSMUSG00 | 11465 |
| NA | ENSMUSG00 | 22690 |
| NA | ENSMUSG00 | 215789 |
| Significant | ENSMUSG00 | 23918 |
| NA | ENSMUSG00 | 75785 |
| Significant | ENSMUSG00 | 170787 |
| NA | ENSMUSG00 | 11364 |
| NA | ENSMUSG00 | 12632 |
| NA | ENSMUSG00 | 237411 |
| NA | ENSMUSG00 | NA |
| NA | ENSMUSG00 | 66902 |
| NA | ENSMUSG00 | 20849 |
| NA | ENSMUSG00 | 100043133 |
| NA | ENSMUSG00 | 320940 |
| NA | ENSMUSG00 | 16542 |
| NA | ENSMUSG00 | 66155 |
| Significant | ENSMUSG00 | 214642 |
| NA | ENSMUSG00 | 67270 |
| Significant | ENSMUSG00 | 211323 |
| NA | ENSMUSG00 | 15893 |
| NA | ENSMUSG00 | 66489 |
| NA | ENSMUSG00 | 12453 |
| NA | ENSMUSG00 | 69161 |
| NA | ENSMUSG00 | 338354 |
| NA | ENSMUSG00 | 69260 |
| NA | ENSMUSG00 | 67144 |
| NA | ENSMUSG00 | 20680 |
| NA | ENSMUSG00 | 26417 |
| NA | ENSMUSG00 | 16561 |
| NA | ENSMUSG00 | 22688 |
| NA | ENSMUSG00 | 72569 |
| NA | ENSMUSG00 | 269713 |
| NA | ENSMUSG00 | 18223 |
| NA | ENSMUSG00 | NA |
| NA | ENSMUSG00 | 72938 |
| NA | ENSMUSG00 | 214580 |
| Significant | ENSMUSG00 | 217304 |
| NA | ENSMUSG00 | 70078 |

| | | |
|---|---|---|
| NA | ENSMUSG00 | 16828 |
| NA | ENSMUSG00 | 80910 |
| NA | ENSMUSG00 | 117171 |
| NA | ENSMUSG00 | NA |
| NA | ENSMUSG00 | 56367 |
| NA | ENSMUSG00 | 671535 |
| NA | ENSMUSG00 | 74838 |
| NA | ENSMUSG00 | 30963 |
| NA | ENSMUSG00 | 22694 |
| NA | ENSMUSG00 | NA |
| NA | ENSMUSG00 | 19942 |
| NA | ENSMUSG00 | 76179 |
| NA | ENSMUSG00 | 544717 |
| NA | ENSMUSG00 | 52705 |
| NA | ENSMUSG00 | 72027 |
| NA | ENSMUSG00 | 26413 |
| NA | ENSMUSG00 | 207958 |
| NA | ENSMUSG00 | 69277 |
| NA | ENSMUSG00 | 80288 |
| NA | ENSMUSG00 | 210853 |
| NA | ENSMUSG00 | 73130 |
| NA | ENSMUSG00 | 223255 |
| NA | ENSMUSG00 | 100042777 |
| NA | ENSMUSG00 | 232987 |
| NA | ENSMUSG00 | 67824 |
| NA | ENSMUSG00 | 319565 |
| NA | ENSMUSG00 | 330286 |
| NA | ENSMUSG00 | 20054 |
| NA | ENSMUSG00 | 76633 |
| NA | ENSMUSG00 | 20826 |
| NA | ENSMUSG00 | 382118 |
| NA | ENSMUSG00 | 239027 |
| NA | ENSMUSG00 | 20637 |
| NA | ENSMUSG00 | 13806 |
| NA | ENSMUSG00 | 76373 |
| NA | ENSMUSG00 | 269966 |
| NA | ENSMUSG00 | 231986 |
| NA | ENSMUSG00 | 71765 |
| Significant | ENSMUSG00 | 234582 |
| NA | ENSMUSG00 | 20666 |
| NA | ENSMUSG00 | 56517 |
| NA | ENSMUSG00 | 30928 |
| NA | ENSMUSG00 | 382236 |
| NA | ENSMUSG00 | 13063 |
| NA | ENSMUSG00 | 667618 |
| NA | ENSMUSG00 | 94281 |

| | | |
|---|---|---|
| NA | ENSMUSG00 | 268291 |
| NA | ENSMUSG00 | 72554 |
| NA | ENSMUSG00 | 66121 |
| NA | ENSMUSG00 | 230596 |
| NA | ENSMUSG00 | 11778 |
| NA | ENSMUSG00 | 66245 |
| NA | ENSMUSG00 | 380669 |
| NA | ENSMUSG00 | 67471 |
| NA | ENSMUSG00 | 236266 |
| NA | ENSMUSG00 | 66812 |
| NA | ENSMUSG00 | 14775 |
| NA | ENSMUSG00 | 107932 |
| NA | ENSMUSG00 | 94249 |
| NA | ENSMUSG00 | 626259 |
| NA | ENSMUSG00 | 66576 |
| NA | ENSMUSG00 | 69956 |
| NA | ENSMUSG00 | 66229 |
| NA | ENSMUSG00 | 12916 |
| NA | ENSMUSG00 | 93681 |
| NA | ENSMUSG00 | NA |
| NA | ENSMUSG00 | NA |
| NA | ENSMUSG00 | 75221 |
| Significant | ENSMUSG00 | 18624 |
| NA | ENSMUSG00 | 268936 |
| NA | ENSMUSG00 | 100041585 |
| NA | ENSMUSG00 | 14536 |
| NA | ENSMUSG00 | 259277 |
| NA | ENSMUSG00 | 78428 |
| NA | ENSMUSG00 | 74254 |
| NA | ENSMUSG00 | 224170 |
| NA | ENSMUSG00 | 320495 |
| NA | ENSMUSG00 | 17827 |
| NA | ENSMUSG00 | 69922 |
| NA | ENSMUSG00 | 94219 |
| NA | ENSMUSG00 | 23900 |
| NA | ENSMUSG00 | 56738 |
| NA | ENSMUSG00 | 26896 |
| NA | ENSMUSG00 | 219103 |
| NA | ENSMUSG00 | 68675 |
| NA | ENSMUSG00 | 381549 |
| NA | ENSMUSG00 | 23807 |
| Significant | ENSMUSG00 | 442827 |
| NA | ENSMUSG00 | 21955 |
| NA | ENSMUSG00 | 216363 |
| NA | ENSMUSG00 | NA |
| NA | ENSMUSG00 | 105722 |

| | | |
|---|---|---|
| NA | ENSMUSG00 | 52668 |
| NA | ENSMUSG00 | 15267 |
| NA | ENSMUSG00 | 403178 |
| Significant | ENSMUSG00 | 66071 |
| NA | ENSMUSG00 | 232969 |
| NA | ENSMUSG00 | 74096 |
| NA | ENSMUSG00 | 100039695 |
| NA | ENSMUSG00 | 72355 |
| NA | ENSMUSG00 | 21353 |
| NA | ENSMUSG00 | 76707 |
| NA | ENSMUSG00 | 69358 |
| NA | ENSMUSG00 | 30954 |
| NA | ENSMUSG00 | 17716 |
| NA | ENSMUSG00 | 17717 |
| NA | ENSMUSG00 | 17708 |
| NA | ENSMUSG00 | 17709 |
| NA | ENSMUSG00 | 17706 |
| NA | ENSMUSG00 | 17705 |
| NA | ENSMUSG00 | 17710 |
| NA | ENSMUSG00 | 17718 |
| NA | ENSMUSG00 | 17719 |
| NA | ENSMUSG00 | 17721 |
| NA | ENSMUSG00 | 17722 |
| NA | ENSMUSG00 | 17711 |
| Significant | ENSMUSG00 | 20363 |
| NA | ENSMUSG00 | 17720 |
| NA | ENSMUSG00 | 546049 |
| NA | ENSMUSG00 | 320165 |
| NA | ENSMUSG00 | 223978 |
| NA | ENSMUSG00 | 66077 |
| NA | ENSMUSG00 | 112422 |
| NA | ENSMUSG00 | 667666 |
| NA | ENSMUSG00 | 626316 |
| Significant | ENSMUSG00 | 20148 |
| NA | ENSMUSG00 | 69116 |
| NA | ENSMUSG00 | 74326 |
| NA | ENSMUSG00 | 67219 |
| NA | ENSMUSG00 | 100169 |
| NA | ENSMUSG00 | 70052 |
| NA | ENSMUSG00 | 66440 |
| NA | ENSMUSG00 | 20529 |
| NA | ENSMUSG00 | 338355 |
| NA | ENSMUSG00 | 20530 |
| NA | ENSMUSG00 | 75691 |
| NA | ENSMUSG00 | 233726 |
| NA | ENSMUSG00 | 66079 |

| | | |
|---|---|---|
| NA | ENSMUSG00 | 215494 |
| NA | ENSMUSG00 | 76681 |
| NA | ENSMUSG00 | 330192 |
| NA | ENSMUSG00 | 101706 |
| NA | ENSMUSG00 | 242291 |
| NA | ENSMUSG00 | 83669 |
| NA | ENSMUSG00 | 625298 |
| NA | ENSMUSG00 | 238393 |
| NA | ENSMUSG00 | 75470 |
| NA | ENSMUSG00 | 75547 |
| NA | ENSMUSG00 | 77853 |
| NA | ENSMUSG00 | 211978 |
| NA | ENSMUSG00 | 17252 |
| NA | ENSMUSG00 | 107885 |
| NA | ENSMUSG00 | 100039601 |
| NA | ENSMUSG00 | 94353 |
| NA | ENSMUSG00 | 544872 |
| NA | ENSMUSG00 | 100043872 |
| NA | ENSMUSG00 | 15289 |
| NA | ENSMUSG00 | 67070 |
| NA | ENSMUSG00 | 233103 |
| NA | ENSMUSG00 | 226844 |
| NA | ENSMUSG00 | 69504 |
| NA | ENSMUSG00 | 70381 |
| NA | ENSMUSG00 | 101154638 |
| NA | ENSMUSG00 | 75516 |
| NA | ENSMUSG00 | 231863 |
| NA | ENSMUSG00 | 74682 |
| NA | ENSMUSG00 | NA |
| NA | ENSMUSG00 | 100033459 |
| NA | ENSMUSG00 | 545812 |
| NA | ENSMUSG00 | 170741 |
| NA | ENSMUSG00 | 235320 |
| NA | ENSMUSG00 | 59095 |
| NA | ENSMUSG00 | NA |
| NA | ENSMUSG00 | 69568 |
| NA | ENSMUSG00 | 241322 |
| Significant | ENSMUSG00 | 24014 |
| NA | ENSMUSG00 | 235050 |
| NA | ENSMUSG00 | 232855 |
| NA | ENSMUSG00 | 26940 |
| NA | ENSMUSG00 | 23960 |
| NA | ENSMUSG00 | 17974 |
| NA | ENSMUSG00 | 100043906 |
| NA | ENSMUSG00 | 170938 |
| NA | ENSMUSG00 | 30843 |

| | | |
|---|---|---|
| NA | ENSMUSG00 | 231668 |
| NA | ENSMUSG00 | 71954 |
| NA | ENSMUSG00 | 231646 |
| Significant | ENSMUSG00 | 12959 |
| NA | ENSMUSG00 | 12237 |
| NA | ENSMUSG00 | NA |
| NA | ENSMUSG00 | 100034727 |
| NA | ENSMUSG00 | 55927 |
| NA | ENSMUSG00 | NA |
| NA | ENSMUSG00 | 20016 |
| NA | ENSMUSG00 | 72322 |
| NA | ENSMUSG00 | NA |
| NA | ENSMUSG00 | 66235 |
| NA | ENSMUSG00 | 14296 |
| NA | ENSMUSG00 | NA |
| NA | ENSMUSG00 | 231296 |
| NA | ENSMUSG00 | 15040 |
| NA | ENSMUSG00 | 100042148 |
| NA | ENSMUSG00 | 15007 |
| NA | ENSMUSG00 | 11837 |
| NA | ENSMUSG00 | 12338 |
| NA | ENSMUSG00 | 54127 |
| NA | ENSMUSG00 | 12168 |
| NA | ENSMUSG00 | 66309 |
| Significant | ENSMUSG00 | 17089 |
| NA | ENSMUSG00 | 215201 |
| NA | ENSMUSG00 | 54218 |
| NA | ENSMUSG00 | 56496 |
| NA | ENSMUSG00 | 240068 |
| NA | ENSMUSG00 | 73451 |
| NA | ENSMUSG00 | 665511 |
| NA | ENSMUSG00 | 70315 |
| NA | ENSMUSG00 | 13610 |
| NA | ENSMUSG00 | NA |
| NA | ENSMUSG00 | 78321 |
| NA | ENSMUSG00 | 19082 |
| NA | ENSMUSG00 | 381810 |
| NA | ENSMUSG00 | 214489 |
| NA | ENSMUSG00 | 100142658 |
| NA | ENSMUSG00 | 27083 |
| NA | ENSMUSG00 | 18111 |
| Significant | ENSMUSG00 | 53619 |
| NA | ENSMUSG00 | 98932 |
| NA | ENSMUSG00 | 74043 |
| NA | ENSMUSG00 | 67067 |
| NA | ENSMUSG00 | 211673 |

| | | |
|---|---|---|
| NA | ENSMUSG00 | NA |
| NA | ENSMUSG00 | 72459 |
| NA | ENSMUSG00 | 320923 |
| NA | ENSMUSG00 | 666532 |
| NA | ENSMUSG00 | 553127 |
| NA | ENSMUSG00 | 66158 |
| NA | ENSMUSG00 | 240034 |
| Significant | ENSMUSG00 | 381066 |
| NA | ENSMUSG00 | 224585 |
| NA | ENSMUSG00 | 68705 |
| NA | ENSMUSG00 | 380916 |
| NA | ENSMUSG00 | 17356 |
| NA | ENSMUSG00 | 21454 |
| NA | ENSMUSG00 | 99237 |
| NA | ENSMUSG00 | 66570 |
| NA | ENSMUSG00 | 76457 |
| NA | ENSMUSG00 | 78177 |
| Significant | ENSMUSG00 | 13011 |
| NA | ENSMUSG00 | 668923 |
| NA | ENSMUSG00 | 104348 |
| NA | ENSMUSG00 | NA |
| NA | ENSMUSG00 | 75597 |
| NA | ENSMUSG00 | 72899 |
| NA | ENSMUSG00 | 18693 |
| Significant | ENSMUSG00 | 16852 |
| NA | ENSMUSG00 | 666586 |
| NA | ENSMUSG00 | 219132 |
| NA | ENSMUSG00 | 232566 |
| NA | ENSMUSG00 | 545864 |
| NA | ENSMUSG00 | 69596 |
| NA | ENSMUSG00 | 12616 |
| NA | ENSMUSG00 | 207806 |
| NA | ENSMUSG00 | 77006 |
| NA | ENSMUSG00 | 75541 |
| NA | ENSMUSG00 | 232156 |
| NA | ENSMUSG00 | 11993 |
| NA | ENSMUSG00 | 64704 |
| NA | ENSMUSG00 | 13448 |
| NA | ENSMUSG00 | 93696 |
| NA | ENSMUSG00 | 99100 |
| NA | ENSMUSG00 | 497106 |
| NA | ENSMUSG00 | 667034 |
| NA | ENSMUSG00 | 22290 |
| NA | ENSMUSG00 | 67532 |
| NA | ENSMUSG00 | 14707 |
| NA | ENSMUSG00 | 68910 |

| | | |
|---|---|---|
| NA | ENSMUSG00 | 50918 |
| NA | ENSMUSG00 | 72008 |
| NA | ENSMUSG00 | 666668 |
| NA | ENSMUSG00 | 11464 |
| NA | ENSMUSG00 | 74374 |
| NA | ENSMUSG00 | 333883 |
| NA | ENSMUSG00 | 59011 |
| NA | ENSMUSG00 | 67495 |
| NA | ENSMUSG00 | 277414 |
| NA | ENSMUSG00 | 20226 |
| NA | ENSMUSG00 | 12953 |
| NA | ENSMUSG00 | 56742 |
| NA | ENSMUSG00 | 20661 |
| NA | ENSMUSG00 | 26442 |
| NA | ENSMUSG00 | 16188 |
| NA | ENSMUSG00 | 109905 |
| NA | ENSMUSG00 | 229663 |
| NA | ENSMUSG00 | 319190 |
| NA | ENSMUSG00 | 319176 |
| NA | ENSMUSG00 | 107701 |
| NA | ENSMUSG00 | 229588 |
| NA | ENSMUSG00 | 20341 |
| NA | ENSMUSG00 | 20823 |
| NA | ENSMUSG00 | 12748 |
| NA | ENSMUSG00 | 65111 |
| NA | ENSMUSG00 | 229524 |
| NA | ENSMUSG00 | 229521 |
| NA | ENSMUSG00 | 70227 |
| NA | ENSMUSG00 | 232966 |
| NA | ENSMUSG00 | NA |
| NA | ENSMUSG00 | 241311 |
| NA | ENSMUSG00 | NA |
| NA | ENSMUSG00 | 68205 |
| NA | ENSMUSG00 | 73333 |
| NA | ENSMUSG00 | 26900 |
| NA | ENSMUSG00 | 26908 |
| NA | ENSMUSG00 | 12572 |
| NA | ENSMUSG00 | 18583 |
| NA | ENSMUSG00 | 229055 |
| NA | ENSMUSG00 | 100039740 |
| NA | ENSMUSG00 | 75296 |
| NA | ENSMUSG00 | 238722 |
| NA | ENSMUSG00 | 238692 |
| NA | ENSMUSG00 | 235956 |
| NA | ENSMUSG00 | 97863 |
| NA | ENSMUSG00 | 105352 |

| | | |
|---|---|---|
| NA | ENSMUSG00 | 319150 |
| NA | ENSMUSG00 | 319164 |
| NA | ENSMUSG00 | 319166 |
| NA | ENSMUSG00 | 319151 |
| NA | ENSMUSG00 | 319183 |
| NA | ENSMUSG00 | 319167 |
| NA | ENSMUSG00 | 319161 |
| NA | ENSMUSG00 | 665596 |
| NA | ENSMUSG00 | 319188 |
| NA | ENSMUSG00 | 319170 |
| NA | ENSMUSG00 | 319148 |
| NA | ENSMUSG00 | 227867 |
| NA | ENSMUSG00 | 17105 |
| NA | ENSMUSG00 | 67226 |
| NA | ENSMUSG00 | 213326 |
| NA | ENSMUSG00 | 70248 |
| NA | ENSMUSG00 | 72149 |
| NA | ENSMUSG00 | 69129 |
| NA | ENSMUSG00 | 17118 |
| NA | ENSMUSG00 | 69837 |
| NA | ENSMUSG00 | NA |
| NA | ENSMUSG00 | 434179 |
| NA | ENSMUSG00 | 239985 |
| NA | ENSMUSG00 | 75424 |
| NA | ENSMUSG00 | 26446 |
| NA | ENSMUSG00 | NA |
| NA | ENSMUSG00 | 76626 |
| NA | ENSMUSG00 | 100034251 |
| NA | ENSMUSG00 | 237886 |
| NA | ENSMUSG00 | 116972 |
| NA | ENSMUSG00 | 195046 |
| Significant | ENSMUSG00 | 66395 |
| NA | ENSMUSG00 | 69215 |
| NA | ENSMUSG00 | 52892 |
| NA | ENSMUSG00 | NA |
| NA | ENSMUSG00 | 54396 |
| NA | ENSMUSG00 | 245240 |
| NA | ENSMUSG00 | 667214 |
| NA | ENSMUSG00 | 52335 |
| NA | ENSMUSG00 | 100039624 |
| NA | ENSMUSG00 | 70385 |
| NA | ENSMUSG00 | 110257 |
| NA | ENSMUSG00 | 15122 |
| NA | ENSMUSG00 | 97440 |
| NA | ENSMUSG00 | NA |
| NA | ENSMUSG00 | 100039924 |

| | | |
|---|---|---|
| Significant | ENSMUSG00 | 74015 |
| NA | ENSMUSG00 | 13716 |
| NA | ENSMUSG00 | 76900 |
| NA | ENSMUSG00 | 100040634 |
| NA | ENSMUSG00 | 434484 |
| NA | ENSMUSG00 | 109032 |
| NA | ENSMUSG00 | 212326 |
| NA | ENSMUSG00 | 14107 |
| NA | ENSMUSG00 | 52065 |
| NA | ENSMUSG00 | NA |
| NA | ENSMUSG00 | 66706 |
| NA | ENSMUSG00 | 331026 |
| NA | ENSMUSG00 | 245020 |
| NA | ENSMUSG00 | 319742 |
| NA | ENSMUSG00 | 53356 |
| NA | ENSMUSG00 | 672511 |
| NA | ENSMUSG00 | 72973 |
| NA | ENSMUSG00 | 100040540 |
| NA | ENSMUSG00 | 12443 |
| NA | ENSMUSG00 | NA |
| NA | ENSMUSG00 | 77613 |
| NA | ENSMUSG00 | 12340 |
| NA | ENSMUSG00 | 637515 |
| NA | ENSMUSG00 | 69186 |
| NA | ENSMUSG00 | 54710 |
| NA | ENSMUSG00 | 102443351 |
| NA | ENSMUSG00 | 75212 |
| NA | ENSMUSG00 | 16068 |
| Significant | ENSMUSG00 | 12406 |
| NA | ENSMUSG00 | 100041748 |
| NA | ENSMUSG00 | 80889 |
| NA | ENSMUSG00 | 71532 |
| NA | ENSMUSG00 | 100039762 |
| NA | ENSMUSG00 | 14004 |
| NA | ENSMUSG00 | 66647 |
| NA | ENSMUSG00 | NA |
| NA | ENSMUSG00 | 27412 |
| NA | ENSMUSG00 | 22652 |
| NA | ENSMUSG00 | 21969 |
| NA | ENSMUSG00 | 226525 |
| NA | ENSMUSG00 | NA |
| NA | ENSMUSG00 | 433938 |
| NA | ENSMUSG00 | 14349 |
| NA | ENSMUSG00 | 75690 |
| NA | ENSMUSG00 | 433791 |
| NA | ENSMUSG00 | 433926 |

| | | |
|---|---|---|
| NA | ENSMUSG00 | 20668 |
| NA | ENSMUSG00 | 12399 |
| NA | ENSMUSG00 | 65961 |
| NA | ENSMUSG00 | 71984 |
| NA | ENSMUSG00 | 434156 |
| NA | ENSMUSG00 | 73430 |
| NA | ENSMUSG00 | 100042405 |
| NA | ENSMUSG00 | 67809 |
| NA | ENSMUSG00 | 329207 |
| NA | ENSMUSG00 | 72313 |
| NA | ENSMUSG00 | 100039968 |
| NA | ENSMUSG00 | 227333 |
| NA | ENSMUSG00 | 245945 |
| NA | ENSMUSG00 | 56222 |
| NA | ENSMUSG00 | 243842 |
| NA | ENSMUSG00 | 321008 |
| NA | ENSMUSG00 | 232875 |
| NA | ENSMUSG00 | 381633 |
| NA | ENSMUSG00 | 227210 |
| NA | ENSMUSG00 | 232801 |
| NA | ENSMUSG00 | 75051 |
| NA | ENSMUSG00 | 242521 |
| NA | ENSMUSG00 | 68441 |
| NA | ENSMUSG00 | 73122 |
| NA | ENSMUSG00 | 107527 |
| NA | ENSMUSG00 | 227746 |
| Significant | ENSMUSG00 | 100503415 |
| NA | ENSMUSG00 | 72429 |
| NA | ENSMUSG00 | 230075 |
| NA | ENSMUSG00 | 73582 |
| NA | ENSMUSG00 | 240168 |
| NA | ENSMUSG00 | 224903 |
| NA | ENSMUSG00 | 328833 |
| NA | ENSMUSG00 | 56351 |
| NA | ENSMUSG00 | 28064 |
| NA | ENSMUSG00 | 16478 |
| NA | ENSMUSG00 | 75692 |
| NA | ENSMUSG00 | 20383 |
| NA | ENSMUSG00 | 234094 |
| NA | ENSMUSG00 | 75616 |
| NA | ENSMUSG00 | 17951 |
| NA | ENSMUSG00 | 330409 |
| NA | ENSMUSG00 | 72371 |
| NA | ENSMUSG00 | 73132 |
| NA | ENSMUSG00 | 449521 |
| NA | ENSMUSG00 | 74149 |

| | | |
|---|---|---|
| NA | ENSMUSG00 | 73233 |
| NA | ENSMUSG00 | 235907 |
| NA | ENSMUSG00 | 238693 |
| NA | ENSMUSG00 | 72139 |
| NA | ENSMUSG00 | 629732 |
| NA | ENSMUSG00 | 436440 |
| NA | ENSMUSG00 | 213402 |
| NA | ENSMUSG00 | NA |
| NA | ENSMUSG00 | 21841 |
| NA | ENSMUSG00 | 13656 |
| NA | ENSMUSG00 | 100041399 |
| NA | ENSMUSG00 | 239122 |
| NA | ENSMUSG00 | 237336 |
| NA | ENSMUSG00 | 26408 |
| NA | ENSMUSG00 | 53602 |
| NA | ENSMUSG00 | 65019 |
| NA | ENSMUSG00 | 69666 |
| NA | ENSMUSG00 | 13528 |
| NA | ENSMUSG00 | 72306 |
| NA | ENSMUSG00 | 319165 |
| NA | ENSMUSG00 | 100043462 |
| NA | ENSMUSG00 | 319191 |
| NA | ENSMUSG00 | NA |
| NA | ENSMUSG00 | 76302 |
| NA | ENSMUSG00 | 74253 |
| Significant | ENSMUSG00 | 70021 |
| NA | ENSMUSG00 | 212517 |
| Significant | ENSMUSG00 | 100034684 |
| Significant | ENSMUSG00 | 20861 |
| NA | ENSMUSG00 | 67795 |
| NA | ENSMUSG00 | 72307 |
| NA | ENSMUSG00 | 12609 |
| NA | ENSMUSG00 | 67160 |
| NA | ENSMUSG00 | 70044 |
| NA | ENSMUSG00 | 23942 |
| NA | ENSMUSG00 | 225898 |
| NA | ENSMUSG00 | 19881 |
| NA | ENSMUSG00 | 72727 |
| NA | ENSMUSG00 | 14376 |
| NA | ENSMUSG00 | 109077 |
| NA | ENSMUSG00 | 66276 |
| NA | ENSMUSG00 | 107197 |
| NA | ENSMUSG00 | 225896 |
| NA | ENSMUSG00 | 14705 |
| NA | ENSMUSG00 | 68693 |
| NA | ENSMUSG00 | 70387 |

| | | |
|---|---|---|
| NA | ENSMUSG00 | 75291 |
| NA | ENSMUSG00 | 67710 |
| NA | ENSMUSG00 | 74478 |
| NA | ENSMUSG00 | 20603 |
| NA | ENSMUSG00 | 12983 |
| NA | ENSMUSG00 | 12984 |
| Significant | ENSMUSG00 | 17972 |
| NA | ENSMUSG00 | 270624 |
| NA | ENSMUSG00 | 14853 |
| NA | ENSMUSG00 | 100503915 |
| NA | ENSMUSG00 | 100040244 |
| NA | ENSMUSG00 | 387609 |
| NA | ENSMUSG00 | NA |
| NA | ENSMUSG00 | NA |
| NA | ENSMUSG00 | 494504 |
| NA | ENSMUSG00 | 240261 |
| NA | ENSMUSG00 | 328949 |
| NA | ENSMUSG00 | 268373 |
| NA | ENSMUSG00 | 211378 |
| NA | ENSMUSG00 | 12449 |
| Significant | ENSMUSG00 | 11727 |
| NA | ENSMUSG00 | 22142 |
| Significant | ENSMUSG00 | 94088 |
| NA | ENSMUSG00 | 21339 |
| NA | ENSMUSG00 | 16597 |
| NA | ENSMUSG00 | 332397 |
| NA | ENSMUSG00 | 105651 |
| NA | ENSMUSG00 | 239510 |
| NA | ENSMUSG00 | 399603 |
| NA | ENSMUSG00 | 217057 |
| Significant | ENSMUSG00 | 13587 |
| NA | ENSMUSG00 | 20556 |
| NA | ENSMUSG00 | NA |
| Significant | ENSMUSG00 | 22750 |
| NA | ENSMUSG00 | 66274 |
| NA | ENSMUSG00 | NA |
| NA | ENSMUSG00 | 320277 |
| NA | ENSMUSG00 | 66039 |
| NA | ENSMUSG00 | NA |
| NA | ENSMUSG00 | NA |
| NA | ENSMUSG00 | NA |
| NA | ENSMUSG00 | NA |
| NA | ENSMUSG00 | 381820 |
| NA | ENSMUSG00 | NA |
| NA | ENSMUSG00 | 100041734 |
| NA | ENSMUSG00 | 77200 |

| | | |
|---|---|---|
| NA | ENSMUSG00 | 54137 |
| NA | ENSMUSG00 | 14790 |
| NA | ENSMUSG00 | 626391 |
| NA | ENSMUSG00 | 108169197 |
| NA | ENSMUSG00 | NA |
| NA | ENSMUSG00 | 100978 |
| NA | ENSMUSG00 | NA |
| NA | ENSMUSG00 | 71275 |
| NA | ENSMUSG00 | 279706 |
| NA | ENSMUSG00 | 77219 |
| NA | ENSMUSG00 | 26897 |
| NA | ENSMUSG00 | 399591 |
| NA | ENSMUSG00 | 70237 |
| NA | ENSMUSG00 | 245607 |
| NA | ENSMUSG00 | 494468 |
| NA | ENSMUSG00 | 11498 |
| NA | ENSMUSG00 | 70645 |
| NA | ENSMUSG00 | 53620 |
| NA | ENSMUSG00 | 331493 |
| Significant | ENSMUSG00 | 331487 |
| NA | ENSMUSG00 | 668166 |
| NA | ENSMUSG00 | 24067 |
| NA | ENSMUSG00 | 243371 |
| NA | ENSMUSG00 | 574437 |
| NA | ENSMUSG00 | 67048 |
| NA | ENSMUSG00 | 236848 |
| NA | ENSMUSG00 | 66808 |
| NA | ENSMUSG00 | 78619 |
| NA | ENSMUSG00 | 236366 |
| NA | ENSMUSG00 | 58242 |
| NA | ENSMUSG00 | 14964 |
| NA | ENSMUSG00 | 16988 |
| NA | ENSMUSG00 | 14961 |
| NA | ENSMUSG00 | 14979 |
| NA | ENSMUSG00 | 328801 |
| NA | ENSMUSG00 | 237412 |
| NA | ENSMUSG00 | 14570 |
| NA | ENSMUSG00 | 106618 |
| NA | ENSMUSG00 | 79059 |
| NA | ENSMUSG00 | 193838 |
| NA | ENSMUSG00 | 100310809 |
| NA | ENSMUSG00 | 240023 |
| NA | ENSMUSG00 | 106489 |
| NA | ENSMUSG00 | 66832 |
| NA | ENSMUSG00 | 67247 |
| NA | ENSMUSG00 | 15951 |

| | | |
|---|---|---|
| Significant | ENSMUSG00 | 226691 |
| NA | ENSMUSG00 | 225659 |
| NA | ENSMUSG00 | 240327 |
| NA | ENSMUSG00 | 329251 |
| NA | ENSMUSG00 | 70425 |
| NA | ENSMUSG00 | 68545 |
| NA | ENSMUSG00 | 98314 |
| NA | ENSMUSG00 | 66915 |
| NA | ENSMUSG00 | 66153 |
| NA | ENSMUSG00 | 19330 |
| NA | ENSMUSG00 | 621100 |
| NA | ENSMUSG00 | 69368 |
| NA | ENSMUSG00 | NA |
| NA | ENSMUSG00 | 241112 |
| NA | ENSMUSG00 | 269198 |
| NA | ENSMUSG00 | 15528 |
| NA | ENSMUSG00 | 241062 |
| NA | ENSMUSG00 | 67513 |
| NA | ENSMUSG00 | 242785 |
| NA | ENSMUSG00 | 114641 |
| NA | ENSMUSG00 | 69928 |
| NA | ENSMUSG00 | 68421 |
| NA | ENSMUSG00 | 76166 |
| NA | ENSMUSG00 | 100039959 |
| NA | ENSMUSG00 | 230757 |
| NA | ENSMUSG00 | 78611 |
| NA | ENSMUSG00 | 100087 |
| NA | ENSMUSG00 | 230500 |
| NA | ENSMUSG00 | 320438 |
| NA | ENSMUSG00 | 381933 |
| NA | ENSMUSG00 | 233870 |
| NA | ENSMUSG00 | 434232 |
| NA | ENSMUSG00 | NA |
| Significant | ENSMUSG00 | 16157 |
| Significant | ENSMUSG00 | NA |
| NA | ENSMUSG00 | 214944 |
| Significant | ENSMUSG00 | 101488143 |
| NA | ENSMUSG00 | 101700 |
| NA | ENSMUSG00 | 56212 |
| NA | ENSMUSG00 | 14590 |
| NA | ENSMUSG00 | 50500 |
| NA | ENSMUSG00 | 102058 |
| NA | ENSMUSG00 | 546164 |
| NA | ENSMUSG00 | 71839 |
| NA | ENSMUSG00 | 56690 |
| NA | ENSMUSG00 | 66585 |

| | | |
|---|---|---|
| Significant | ENSMUSG00 | 75744 |
| NA | ENSMUSG00 | 109095 |
| NA | ENSMUSG00 | 22121 |
| NA | ENSMUSG00 | NA |
| NA | ENSMUSG00 | 434341 |
| NA | ENSMUSG00 | 208111 |
| NA | ENSMUSG00 | 67607 |
| NA | ENSMUSG00 | 101835 |
| NA | ENSMUSG00 | 72287 |
| NA | ENSMUSG00 | 229937 |
| NA | ENSMUSG00 | 244556 |
| NA | ENSMUSG00 | 619292 |
| NA | ENSMUSG00 | 11522 |
| NA | ENSMUSG00 | 68332 |
| NA | ENSMUSG00 | 70604 |
| NA | ENSMUSG00 | NA |
| NA | ENSMUSG00 | 12865 |
| NA | ENSMUSG00 | 233060 |
| NA | ENSMUSG00 | 243905 |
| NA | ENSMUSG00 | 20733 |
| NA | ENSMUSG00 | 211556 |
| NA | ENSMUSG00 | 66498 |
| NA | ENSMUSG00 | 73673 |
| NA | ENSMUSG00 | 26365 |
| NA | ENSMUSG00 | 22756 |
| NA | ENSMUSG00 | 56869 |
| NA | ENSMUSG00 | 244895 |
| NA | ENSMUSG00 | 12659 |
| NA | ENSMUSG00 | 433638 |
| NA | ENSMUSG00 | 69296 |
| NA | ENSMUSG00 | 232933 |
| NA | ENSMUSG00 | 319430 |
| NA | ENSMUSG00 | 259300 |
| NA | ENSMUSG00 | 319154 |
| NA | ENSMUSG00 | 319748 |
| NA | ENSMUSG00 | 232816 |
| NA | ENSMUSG00 | 100113396 |
| NA | ENSMUSG00 | NA |
| NA | ENSMUSG00 | 330921 |
| NA | ENSMUSG00 | 67860 |
| NA | ENSMUSG00 | 67629 |
| NA | ENSMUSG00 | 72640 |
| Significant | ENSMUSG00 | 75863 |
| NA | ENSMUSG00 | 72230 |
| NA | ENSMUSG00 | 99889 |
| NA | ENSMUSG00 | 626848 |

| | | |
|---|---|---|
| NA | ENSMUSG00 | NA |
| NA | ENSMUSG00 | 102639598 |
| NA | ENSMUSG00 | 320664 |
| NA | ENSMUSG00 | 69372 |
| NA | ENSMUSG00 | 624866 |
| NA | ENSMUSG00 | NA |
| NA | ENSMUSG00 | 99371 |
| NA | ENSMUSG00 | 211666 |
| NA | ENSMUSG00 | 269389 |
| Significant | ENSMUSG00 | 16658 |
| NA | ENSMUSG00 | 266692 |
| NA | ENSMUSG00 | 320095 |
| NA | ENSMUSG00 | 613262 |
| NA | ENSMUSG00 | 67204 |
| NA | ENSMUSG00 | 241732 |
| NA | ENSMUSG00 | 67917 |
| NA | ENSMUSG00 | 12995 |
| NA | ENSMUSG00 | 414758 |
| NA | ENSMUSG00 | 21824 |
| NA | ENSMUSG00 | 107368 |
| NA | ENSMUSG00 | 382423 |
| NA | ENSMUSG00 | 228730 |
| NA | ENSMUSG00 | 228715 |
| NA | ENSMUSG00 | 93765 |
| NA | ENSMUSG00 | 54712 |
| NA | ENSMUSG00 | 72630 |
| NA | ENSMUSG00 | 105171 |
| NA | ENSMUSG00 | 16434 |
| NA | ENSMUSG00 | 237436 |
| NA | ENSMUSG00 | 20170 |
| NA | ENSMUSG00 | 432770 |
| NA | ENSMUSG00 | 73338 |
| NA | ENSMUSG00 | 100038538 |
| NA | ENSMUSG00 | 630836 |
| NA | ENSMUSG00 | 435366 |
| NA | ENSMUSG00 | 77117 |
| NA | ENSMUSG00 | 630579 |
| Significant | ENSMUSG00 | 13025 |
| NA | ENSMUSG00 | 378702 |
| NA | ENSMUSG00 | 26385 |
| NA | ENSMUSG00 | 329504 |
| NA | ENSMUSG00 | 15959 |
| NA | ENSMUSG00 | 240614 |
| NA | ENSMUSG00 | 72136 |
| NA | ENSMUSG00 | 100043272 |
| NA | ENSMUSG00 | 68034 |

| | | |
|---|---|---|
| NA | ENSMUSG00 | 72351 |
| NA | ENSMUSG00 | 621269 |
| NA | ENSMUSG00 | 99003 |
| NA | ENSMUSG00 | 641340 |
| NA | ENSMUSG00 | NA |
| NA | ENSMUSG00 | 100042784 |
| NA | ENSMUSG00 | 319178 |
| NA | ENSMUSG00 | 385658 |
| NA | ENSMUSG00 | 381410 |
| NA | ENSMUSG00 | 67008 |
| NA | ENSMUSG00 | 12519 |
| NA | ENSMUSG00 | 75973 |
| NA | ENSMUSG00 | 29805 |
| NA | ENSMUSG00 | 381045 |
| NA | ENSMUSG00 | 11702 |
| NA | ENSMUSG00 | 77446 |
| NA | ENSMUSG00 | 66311 |
| NA | ENSMUSG00 | 328660 |
| NA | ENSMUSG00 | 78802 |
| NA | ENSMUSG00 | 620631 |
| NA | ENSMUSG00 | 72421 |
| NA | ENSMUSG00 | 100040519 |
| NA | ENSMUSG00 | 215280 |
| NA | ENSMUSG00 | 107239 |
| NA | ENSMUSG00 | 667281 |
| NA | ENSMUSG00 | 228003 |
| NA | ENSMUSG00 | 381990 |
| NA | ENSMUSG00 | 319817 |
| NA | ENSMUSG00 | 14269 |
| NA | ENSMUSG00 | 227697 |
| NA | ENSMUSG00 | 52838 |
| NA | ENSMUSG00 | 380959 |
| NA | ENSMUSG00 | 66200 |
| NA | ENSMUSG00 | 432879 |
| NA | ENSMUSG00 | 69684 |
| NA | ENSMUSG00 | 277154 |
| NA | ENSMUSG00 | 330217 |
| NA | ENSMUSG00 | 268469 |
| NA | ENSMUSG00 | 223642 |
| NA | ENSMUSG00 | 69227 |
| NA | ENSMUSG00 | 109815 |
| NA | ENSMUSG00 | 114679 |
| NA | ENSMUSG00 | 28042 |
| NA | ENSMUSG00 | 26462 |
| NA | ENSMUSG00 | 27361 |
| NA | ENSMUSG00 | 625249 |

| | | |
|---|---|---|
| NA | ENSMUSG00 | 20677 |
| NA | ENSMUSG00 | 22630 |
| NA | ENSMUSG00 | 264895 |
| NA | ENSMUSG00 | 72657 |
| NA | ENSMUSG00 | 11898 |
| NA | ENSMUSG00 | 16019 |
| NA | ENSMUSG00 | 19326 |
| NA | ENSMUSG00 | 672959 |
| NA | ENSMUSG00 | 100502754 |
| NA | ENSMUSG00 | 621823 |
| NA | ENSMUSG00 | NA |
| NA | ENSMUSG00 | 12663 |
| NA | ENSMUSG00 | 100038991 |
| NA | ENSMUSG00 | 227615 |
| NA | ENSMUSG00 | NA |
| NA | ENSMUSG00 | 111241 |
| NA | ENSMUSG00 | 100040109 |
| NA | ENSMUSG00 | 14070 |
| NA | ENSMUSG00 | 66125 |
| Significant | ENSMUSG00 | 68859 |
| NA | ENSMUSG00 | 622339 |
| NA | ENSMUSG00 | 100043209 |
| NA | ENSMUSG00 | 66118 |
| NA | ENSMUSG00 | 52468 |
| NA | ENSMUSG00 | 216177 |
| NA | ENSMUSG00 | 72273 |
| NA | ENSMUSG00 | 102115 |
| NA | ENSMUSG00 | NA |
| NA | ENSMUSG00 | 73112 |
| NA | ENSMUSG00 | 231002 |
| NA | ENSMUSG00 | 544678 |
| NA | ENSMUSG00 | 100041677 |
| NA | ENSMUSG00 | 195531 |
| NA | ENSMUSG00 | 433801 |
| NA | ENSMUSG00 | 68817 |
| NA | ENSMUSG00 | 230866 |
| NA | ENSMUSG00 | 69885 |
| NA | ENSMUSG00 | 70081 |
| NA | ENSMUSG00 | 12176 |
| NA | ENSMUSG00 | 68920 |
| NA | ENSMUSG00 | 208501 |
| NA | ENSMUSG00 | 66105 |
| NA | ENSMUSG00 | 101604 |
| NA | ENSMUSG00 | 230696 |
| NA | ENSMUSG00 | NA |
| NA | ENSMUSG00 | 100042856 |

| | | |
|---|---|---|
| NA | ENSMUSG00 | 434219 |
| NA | ENSMUSG00 | 83796 |
| NA | ENSMUSG00 | 67163 |
| NA | ENSMUSG00 | 791260 |
| NA | ENSMUSG00 | 654473 |
| NA | ENSMUSG00 | 237940 |
| NA | ENSMUSG00 | 19192 |
| NA | ENSMUSG00 | 68107 |
| NA | ENSMUSG00 | 28084 |
| NA | ENSMUSG00 | 100038347 |
| NA | ENSMUSG00 | 244059 |
| NA | ENSMUSG00 | 192160 |
| NA | ENSMUSG00 | 68634 |
| NA | ENSMUSG00 | NA |
| NA | ENSMUSG00 | 217149 |
| NA | ENSMUSG00 | 68512 |
| NA | ENSMUSG00 | 71909 |
| NA | ENSMUSG00 | 233073 |
| NA | ENSMUSG00 | 72556 |
| NA | ENSMUSG00 | 14017 |
| NA | ENSMUSG00 | 623474 |
| NA | ENSMUSG00 | 22717 |
| NA | ENSMUSG00 | 69126 |
| NA | ENSMUSG00 | 414069 |
| NA | ENSMUSG00 | 116905 |
| NA | ENSMUSG00 | 276770 |
| NA | ENSMUSG00 | 69757 |
| NA | ENSMUSG00 | 319162 |
| NA | ENSMUSG00 | 16145 |
| NA | ENSMUSG00 | 353208 |
| NA | ENSMUSG00 | 665211 |
| NA | ENSMUSG00 | 626802 |
| NA | ENSMUSG00 | 628308 |
| NA | ENSMUSG00 | 668066 |
| NA | ENSMUSG00 | 100043403 |
| NA | ENSMUSG00 | 627901 |
| NA | ENSMUSG00 | 100039123 |
| NA | ENSMUSG00 | 100043387 |
| NA | ENSMUSG00 | 545490 |
| NA | ENSMUSG00 | 100043381 |
| NA | ENSMUSG00 | 668039 |
| NA | ENSMUSG00 | 100039060 |
| NA | ENSMUSG00 | 626832 |
| NA | ENSMUSG00 | 13999 |
| NA | ENSMUSG00 | 100504263 |
| NA | ENSMUSG00 | 100503949 |

| | | |
|---|---|---|
| NA | ENSMUSG00 | 100043915 |
| NA | ENSMUSG00 | 100043914 |
| NA | ENSMUSG00 | NA |
| NA | ENSMUSG00 | 665001 |
| NA | ENSMUSG00 | 270096 |
| NA | ENSMUSG00 | 13480 |
| NA | ENSMUSG00 | 15953 |
| NA | ENSMUSG00 | 66589 |
| NA | ENSMUSG00 | 68023 |
| NA | ENSMUSG00 | NA |
| NA | ENSMUSG00 | 17952 |
| Significant | ENSMUSG00 | 17948 |
| NA | ENSMUSG00 | 100042746 |
| NA | ENSMUSG00 | 100041325 |
| NA | ENSMUSG00 | 103784 |
| NA | ENSMUSG00 | 20335 |
| NA | ENSMUSG00 | 72807 |
| NA | ENSMUSG00 | 408065 |
| NA | ENSMUSG00 | 666704 |
| NA | ENSMUSG00 | 76933 |
| NA | ENSMUSG00 | 106068 |
| NA | ENSMUSG00 | NA |
| NA | ENSMUSG00 | 19122 |
| NA | ENSMUSG00 | 211135 |
| NA | ENSMUSG00 | 380601 |
| NA | ENSMUSG00 | 56461 |
| NA | ENSMUSG00 | 102294 |
| NA | ENSMUSG00 | NA |
| NA | ENSMUSG00 | 77532 |
| NA | ENSMUSG00 | 66396 |
| NA | ENSMUSG00 | 328099 |
| NA | ENSMUSG00 | 100101806 |
| NA | ENSMUSG00 | 18861 |
| NA | ENSMUSG00 | 12335 |
| NA | ENSMUSG00 | 66913 |
| NA | ENSMUSG00 | NA |
| NA | ENSMUSG00 | 319266 |
| NA | ENSMUSG00 | 53791 |
| NA | ENSMUSG00 | 751865 |
| NA | ENSMUSG00 | 22635 |
| NA | ENSMUSG00 | 75339 |
| NA | ENSMUSG00 | 19188 |
| NA | ENSMUSG00 | 269704 |
| NA | ENSMUSG00 | 12774 |
| NA | ENSMUSG00 | 22437 |
| NA | ENSMUSG00 | 636931 |

| | | |
|---|---|---|
| NA | ENSMUSG00 | 100504715 |
| NA | ENSMUSG00 | 69549 |
| NA | ENSMUSG00 | 56644 |
| NA | ENSMUSG00 | 100041236 |
| NA | ENSMUSG00 | 56382 |
| NA | ENSMUSG00 | 66226 |
| NA | ENSMUSG00 | NA |
| NA | ENSMUSG00 | 56441 |
| NA | ENSMUSG00 | 667373 |
| NA | ENSMUSG00 | 17472 |
| NA | ENSMUSG00 | 12803 |
| NA | ENSMUSG00 | 666468 |
| Significant | ENSMUSG00 | 73656 |
| NA | ENSMUSG00 | 68089 |
| NA | ENSMUSG00 | 100039707 |
| NA | ENSMUSG00 | 100040766 |
| NA | ENSMUSG00 | 19982 |
| NA | ENSMUSG00 | 20448 |
| NA | ENSMUSG00 | 227327 |
| NA | ENSMUSG00 | 73061 |
| NA | ENSMUSG00 | 55981 |
| NA | ENSMUSG00 | 195434 |
| NA | ENSMUSG00 | 19349 |
| NA | ENSMUSG00 | 56390 |
| NA | ENSMUSG00 | 69713 |
| NA | ENSMUSG00 | 100042480 |
| Significant | ENSMUSG00 | 227696 |
| NA | ENSMUSG00 | 59024 |
| NA | ENSMUSG00 | 380687 |
| NA | ENSMUSG00 | 68316 |
| NA | ENSMUSG00 | 22764 |
| NA | ENSMUSG00 | NA |
| NA | ENSMUSG00 | 19240 |
| NA | ENSMUSG00 | 434797 |
| NA | ENSMUSG00 | 14999 |
| NA | ENSMUSG00 | 100502766 |
| NA | ENSMUSG00 | 231123 |
| NA | ENSMUSG00 | 224703 |
| Significant | ENSMUSG00 | 100502940 |
| NA | ENSMUSG00 | 59003 |
| Significant | ENSMUSG00 | 433016 |
| NA | ENSMUSG00 | 474156 |
| NA | ENSMUSG00 | 170461 |
| NA | ENSMUSG00 | 109346 |
| NA | ENSMUSG00 | 72124 |
| NA | ENSMUSG00 | 67248 |

| | | |
|---|---|---|
| NA | ENSMUSG00 | 67923 |
| NA | ENSMUSG00 | 652925 |
| NA | ENSMUSG00 | 68165 |
| NA | ENSMUSG00 | 77777 |
| NA | ENSMUSG00 | 21647 |
| NA | ENSMUSG00 | 100041639 |
| NA | ENSMUSG00 | NA |
| NA | ENSMUSG00 | 102638047 |
| NA | ENSMUSG00 | NA |
| NA | ENSMUSG00 | 434794 |
| NA | ENSMUSG00 | 107392 |
| NA | ENSMUSG00 | 245305 |
| NA | ENSMUSG00 | 620678 |
| NA | ENSMUSG00 | NA |
| NA | ENSMUSG00 | NA |
| NA | ENSMUSG00 | 100043010 |
| NA | ENSMUSG00 | NA |
| NA | ENSMUSG00 | NA |
| NA | ENSMUSG00 | 100418329 |
| NA | ENSMUSG00 | 100041245 |
| NA | ENSMUSG00 | 111283 |
| NA | ENSMUSG00 | 100041686 |
| NA | ENSMUSG00 | 97114 |
| NA | ENSMUSG00 | 100041511 |
| NA | ENSMUSG00 | NA |
| NA | ENSMUSG00 | 100039110 |
| NA | ENSMUSG00 | 620877 |
| NA | ENSMUSG00 | NA |
| NA | ENSMUSG00 | NA |
| NA | ENSMUSG00 | NA |
| NA | ENSMUSG00 | 105244208 |
| NA | ENSMUSG00 | NA |
| Significant | ENSMUSG00 | 67739 |
| NA | ENSMUSG00 | 100040140 |
| NA | ENSMUSG00 | NA |
| NA | ENSMUSG00 | 102639974 |
| NA | ENSMUSG00 | 18722 |
| NA | ENSMUSG00 | 546298 |
| NA | ENSMUSG00 | 112136069 |
| NA | ENSMUSG00 | NA |
| NA | ENSMUSG00 | 100416143 |
| NA | ENSMUSG00 | 100416290 |
| NA | ENSMUSG00 | NA |
| NA | ENSMUSG00 | 100417846 |
| NA | ENSMUSG00 | 15504 |
| NA | ENSMUSG00 | NA |

| | | |
|---|---|---|
| NA | ENSMUSG00 | NA |
| NA | ENSMUSG00 | NA |
| NA | ENSMUSG00 | NA |
| NA | ENSMUSG00 | NA |
| NA | ENSMUSG00 | 668943 |
| NA | ENSMUSG00 | 194960 |
| NA | ENSMUSG00 | 665407 |
| NA | ENSMUSG00 | NA |
| NA | ENSMUSG00 | NA |
| NA | ENSMUSG00 | 667799 |
| NA | ENSMUSG00 | 102635048 |
| NA | ENSMUSG00 | 100040898 |
| NA | ENSMUSG00 | NA |
| NA | ENSMUSG00 | 434404 |
| NA | ENSMUSG00 | 627160 |
| NA | ENSMUSG00 | NA |
| NA | ENSMUSG00 | NA |
| NA | ENSMUSG00 | NA |
| NA | ENSMUSG00 | NA |
| NA | ENSMUSG00 | 620559 |
| NA | ENSMUSG00 | 100043840 |
| NA | ENSMUSG00 | 100041720 |
| NA | ENSMUSG00 | 665412 |
| NA | ENSMUSG00 | NA |
| NA | ENSMUSG00 | NA |
| NA | ENSMUSG00 | 433715 |
| NA | ENSMUSG00 | NA |
| NA | ENSMUSG00 | NA |
| NA | ENSMUSG00 | NA |
| NA | ENSMUSG00 | NA |
| Significant | ENSMUSG00 | 56464 |
| NA | ENSMUSG00 | 245350 |
| NA | ENSMUSG00 | NA |
| NA | ENSMUSG00 | NA |
| NA | ENSMUSG00 | 100041273 |
| NA | ENSMUSG00 | 100043546 |
| NA | ENSMUSG00 | NA |
| NA | ENSMUSG00 | NA |
| NA | ENSMUSG00 | NA |
| NA | ENSMUSG00 | 19428 |
| NA | ENSMUSG00 | NA |
| NA | ENSMUSG00 | NA |
| NA | ENSMUSG00 | NA |
| NA | ENSMUSG00 | NA |
| NA | ENSMUSG00 | NA |
| NA | ENSMUSG00 | 665816 |

| | | |
|---|---|---|
| NA | ENSMUSG00 | NA |
| NA | ENSMUSG00 | 100033452 |
| NA | ENSMUSG00 | 100038703 |
| NA | ENSMUSG00 | 100043312 |
| NA | ENSMUSG00 | NA |
| NA | ENSMUSG00 | NA |
| NA | ENSMUSG00 | 544800 |
| NA | ENSMUSG00 | 100043296 |
| NA | ENSMUSG00 | 620213 |
| NA | ENSMUSG00 | 100040605 |
| NA | ENSMUSG00 | 628388 |
| NA | ENSMUSG00 | NA |
| NA | ENSMUSG00 | NA |
| NA | ENSMUSG00 | NA |
| NA | ENSMUSG00 | NA |
| NA | ENSMUSG00 | 100043000 |
| NA | ENSMUSG00 | NA |
| NA | ENSMUSG00 | 667731 |
| NA | ENSMUSG00 | 66177 |
| NA | ENSMUSG00 | 668158 |
| NA | ENSMUSG00 | 100503572 |
| NA | ENSMUSG00 | NA |
| NA | ENSMUSG00 | NA |
| NA | ENSMUSG00 | 108168233 |
| NA | ENSMUSG00 | NA |
| NA | ENSMUSG00 | NA |
| NA | ENSMUSG00 | NA |
| NA | ENSMUSG00 | 217708 |
| NA | ENSMUSG00 | 353310 |
| NA | ENSMUSG00 | 108168352 |
| NA | ENSMUSG00 | 546143 |
| NA | ENSMUSG00 | 70123 |
| NA | ENSMUSG00 | 77490 |
| NA | ENSMUSG00 | 630146 |
| NA | ENSMUSG00 | 100043295 |
| NA | ENSMUSG00 | 664994 |
| NA | ENSMUSG00 | NA |
| NA | ENSMUSG00 | NA |
| NA | ENSMUSG00 | NA |
| NA | ENSMUSG00 | 433022 |
| NA | ENSMUSG00 | NA |
| NA | ENSMUSG00 | 624198 |
| NA | ENSMUSG00 | NA |
| NA | ENSMUSG00 | 68576 |
| NA | ENSMUSG00 | 621976 |
| NA | ENSMUSG00 | 433752 |

| | | |
|---|---|---|
| NA | ENSMUSG00 | 72997 |
| NA | ENSMUSG00 | 100040298 |
| NA | ENSMUSG00 | 56707 |
| NA | ENSMUSG00 | 69784 |
| NA | ENSMUSG00 | 109299 |
| NA | ENSMUSG00 | 100503890 |
| NA | ENSMUSG00 | NA |
| NA | ENSMUSG00 | 100503924 |
| NA | ENSMUSG00 | 12050 |
| NA | ENSMUSG00 | 242681 |
| NA | ENSMUSG00 | 108148 |
| NA | ENSMUSG00 | 494448 |
| NA | ENSMUSG00 | 435921 |
| NA | ENSMUSG00 | 667962 |
| NA | ENSMUSG00 | 100417756 |
| NA | ENSMUSG00 | 666244 |
| NA | ENSMUSG00 | 53881 |
| NA | ENSMUSG00 | NA |
| NA | ENSMUSG00 | 75710 |
| NA | ENSMUSG00 | 192192 |
| NA | ENSMUSG00 | 14356 |
| NA | ENSMUSG00 | 382019 |
| NA | ENSMUSG00 | 100036521 |
| NA | ENSMUSG00 | 320119 |
| NA | ENSMUSG00 | 380705 |
| NA | ENSMUSG00 | NA |
| NA | ENSMUSG00 | 15247 |
| NA | ENSMUSG00 | 68556 |
| NA | ENSMUSG00 | 12045 |
| NA | ENSMUSG00 | 18725 |
| NA | ENSMUSG00 | 677884 |
| NA | ENSMUSG00 | NA |
| NA | ENSMUSG00 | 71176 |
| NA | ENSMUSG00 | 66191 |
| NA | ENSMUSG00 | 100642166 |
| NA | ENSMUSG00 | 16205 |
| NA | ENSMUSG00 | 14426 |
| NA | ENSMUSG00 | 72699 |
| NA | ENSMUSG00 | 58230 |
| NA | ENSMUSG00 | 100043761 |
| NA | ENSMUSG00 | 140810 |
| NA | ENSMUSG00 | 105886298 |
| NA | ENSMUSG00 | 268281 |
| NA | ENSMUSG00 | 224836 |
| Significant | ENSMUSG00 | 394432 |
| NA | ENSMUSG00 | NA |

| | | |
|---|---|---|
| NA | ENSMUSG00 | 22186 |
| NA | ENSMUSG00 | 102632 |
| NA | ENSMUSG00 | 232406 |
| NA | ENSMUSG00 | 213980 |
| NA | ENSMUSG00 | 407243 |
| NA | ENSMUSG00 | 14962 |
| NA | ENSMUSG00 | 14533 |
| NA | ENSMUSG00 | 211151 |
| NA | ENSMUSG00 | 74319 |
| NA | ENSMUSG00 | 100040462 |
| NA | ENSMUSG00 | 212728 |
| NA | ENSMUSG00 | NA |
| Significant | ENSMUSG00 | 67647 |
| NA | ENSMUSG00 | 71683 |
| NA | ENSMUSG00 | 625311 |
| NA | ENSMUSG00 | 20643 |
| NA | ENSMUSG00 | 320700 |
| NA | ENSMUSG00 | 21778 |
| NA | ENSMUSG00 | NA |
| NA | ENSMUSG00 | 78251 |
| NA | ENSMUSG00 | 72958 |
| NA | ENSMUSG00 | NA |
| NA | ENSMUSG00 | 545205 |
| NA | ENSMUSG00 | 381823 |
| NA | ENSMUSG00 | 57294 |
| NA | ENSMUSG00 | 546611 |
| NA | ENSMUSG00 | 17904 |
| NA | ENSMUSG00 | 68052 |
| NA | ENSMUSG00 | 15511 |
| NA | ENSMUSG00 | 258325 |
| NA | ENSMUSG00 | 241989 |
| NA | ENSMUSG00 | 24071 |
| NA | ENSMUSG00 | NA |
| NA | ENSMUSG00 | 72123 |
| NA | ENSMUSG00 | 434215 |
| NA | ENSMUSG00 | NA |
| NA | ENSMUSG00 | NA |
| NA | ENSMUSG00 | 432502 |
| Significant | ENSMUSG00 | 100039139 |
| NA | ENSMUSG00 | 628705 |
| NA | ENSMUSG00 | 380850 |
| NA | ENSMUSG00 | NA |
| NA | ENSMUSG00 | 108934 |
| NA | ENSMUSG00 | 626534 |
| NA | ENSMUSG00 | 58521 |
| NA | ENSMUSG00 | 100418460 |

| | | |
|---|---|---|
| NA | ENSMUSG00 | NA |
| NA | ENSMUSG00 | 100042235 |
| NA | ENSMUSG00 | NA |
| NA | ENSMUSG00 | NA |
| NA | ENSMUSG00 | 56692 |
| NA | ENSMUSG00 | NA |
| NA | ENSMUSG00 | NA |
| NA | ENSMUSG00 | 208285 |
| NA | ENSMUSG00 | 66373 |
| NA | ENSMUSG00 | 100042266 |
| NA | ENSMUSG00 | 236451 |
| NA | ENSMUSG00 | 100073351 |
| NA | ENSMUSG00 | 636741 |
| NA | ENSMUSG00 | 100126824 |
| NA | ENSMUSG00 | 66272 |
| NA | ENSMUSG00 | 66300 |
| NA | ENSMUSG00 | 102636082 |
| NA | ENSMUSG00 | 100043718 |
| NA | ENSMUSG00 | 56550 |
| NA | ENSMUSG00 | 100233175 |
| NA | ENSMUSG00 | 667279 |
| NA | ENSMUSG00 | 193740 |
| NA | ENSMUSG00 | 408059 |
| NA | ENSMUSG00 | NA |
| NA | ENSMUSG00 | 170676 |
| NA | ENSMUSG00 | 666938 |
| NA | ENSMUSG00 | 100310872 |
| NA | ENSMUSG00 | 100040608 |
| NA | ENSMUSG00 | NA |
| NA | ENSMUSG00 | 72716 |
| NA | ENSMUSG00 | 67685 |
| Significant | ENSMUSG00 | 114241 |
| NA | ENSMUSG00 | 100039556 |
| NA | ENSMUSG00 | 620419 |
| NA | ENSMUSG00 | NA |
| NA | ENSMUSG00 | NA |
| NA | ENSMUSG00 | 474160 |
| NA | ENSMUSG00 | 118568338 |
| NA | ENSMUSG00 | 434178 |
| NA | ENSMUSG00 | 81845 |
| NA | ENSMUSG00 | 102632136 |
| NA | ENSMUSG00 | 100416076 |
| NA | ENSMUSG00 | NA |
| NA | ENSMUSG00 | NA |
| NA | ENSMUSG00 | 56771 |
| NA | ENSMUSG00 | 241197 |

| | | |
|---|---|---|
| NA | ENSMUSG00 | NA |
| NA | ENSMUSG00 | NA |
| NA | ENSMUSG00 | NA |
| NA | ENSMUSG00 | 231798 |
| NA | ENSMUSG00 | NA |
| NA | ENSMUSG00 | 66892 |
| NA | ENSMUSG00 | 67945 |
| NA | ENSMUSG00 | NA |
| NA | ENSMUSG00 | 666904 |
| NA | ENSMUSG00 | 19054 |
| NA | ENSMUSG00 | 67952 |
| Significant | ENSMUSG00 | 208715 |
| NA | ENSMUSG00 | NA |
| NA | ENSMUSG00 | 668727 |
| NA | ENSMUSG00 | 100039441 |
| NA | ENSMUSG00 | 52898 |
| NA | ENSMUSG00 | 665433 |
| NA | ENSMUSG00 | NA |
| NA | ENSMUSG00 | 667118 |
| NA | ENSMUSG00 | NA |
| NA | ENSMUSG00 | 77652 |
| NA | ENSMUSG00 | 77358 |
| NA | ENSMUSG00 | 19291 |
| NA | ENSMUSG00 | 100503583 |
| NA | ENSMUSG00 | 100040944 |
| NA | ENSMUSG00 | 100042349 |
| NA | ENSMUSG00 | 68195 |
| NA | ENSMUSG00 | 319171 |
| NA | ENSMUSG00 | NA |
| NA | ENSMUSG00 | 625360 |
| NA | ENSMUSG00 | 72465 |
| NA | ENSMUSG00 | 19653 |
| NA | ENSMUSG00 | 100041979 |
| NA | ENSMUSG00 | NA |
| NA | ENSMUSG00 | 100041342 |
| NA | ENSMUSG00 | 240514 |
| NA | ENSMUSG00 | 545654 |
| NA | ENSMUSG00 | 319622 |
| NA | ENSMUSG00 | 107988026 |
| NA | ENSMUSG00 | 100303732 |
| NA | ENSMUSG00 | 240064 |
| NA | ENSMUSG00 | 100041735 |
| NA | ENSMUSG00 | 102633750 |
| NA | ENSMUSG00 | 240066 |
| NA | ENSMUSG00 | 329575 |
| NA | ENSMUSG00 | 100861634 |

| | | |
|---|---|---|
| NA | ENSMUSG00 | 100043752 |
| NA | ENSMUSG00 | 212276 |
| NA | ENSMUSG00 | 67464 |
| NA | ENSMUSG00 | 57741 |
| NA | ENSMUSG00 | 73385 |
| NA | ENSMUSG00 | NA |
| NA | ENSMUSG00 | 76770 |
| NA | ENSMUSG00 | 628147 |
| NA | ENSMUSG00 | 100040531 |
| NA | ENSMUSG00 | 100037283 |
| NA | ENSMUSG00 | NA |
| NA | ENSMUSG00 | 69034 |
| NA | ENSMUSG00 | NA |
| NA | ENSMUSG00 | 245269 |
| NA | ENSMUSG00 | 22759 |
| NA | ENSMUSG00 | 70045 |
| NA | ENSMUSG00 | NA |
| NA | ENSMUSG00 | 320332 |
| NA | ENSMUSG00 | 320333 |
| NA | ENSMUSG00 | 27973 |
| NA | ENSMUSG00 | 97487 |
| NA | ENSMUSG00 | 69709 |
| NA | ENSMUSG00 | 14958 |
| NA | ENSMUSG00 | 21648 |
| NA | ENSMUSG00 | 102902673 |
| NA | ENSMUSG00 | 101056094 |
| NA | ENSMUSG00 | 240038 |
| NA | ENSMUSG00 | 64113 |
| NA | ENSMUSG00 | 12581 |
| NA | ENSMUSG00 | NA |
| NA | ENSMUSG00 | NA |
| NA | ENSMUSG00 | 100503386 |
| NA | ENSMUSG00 | 433921 |
| NA | ENSMUSG00 | 213332 |
| NA | ENSMUSG00 | 449000 |
| NA | ENSMUSG00 | 331188 |
| Significant | ENSMUSG00 | 235712 |
| NA | ENSMUSG00 | 16912 |
| NA | ENSMUSG00 | 102308570 |
| NA | ENSMUSG00 | NA |
| NA | ENSMUSG00 | 73610 |
| NA | ENSMUSG00 | 433845 |
| NA | ENSMUSG00 | NA |
| NA | ENSMUSG00 | 100233208 |
| NA | ENSMUSG00 | 100043468 |
| NA | ENSMUSG00 | 100043772 |

| | | |
|---|---|---|
| NA | ENSMUSG00 | 100503302 |
| NA | ENSMUSG00 | 381922 |
| NA | ENSMUSG00 | NA |
| NA | ENSMUSG00 | NA |
| NA | ENSMUSG00 | NA |
| NA | ENSMUSG00 | NA |
| NA | ENSMUSG00 | 170763 |
| NA | ENSMUSG00 | NA |
| NA | ENSMUSG00 | 100041204 |
| NA | ENSMUSG00 | NA |
| NA | ENSMUSG00 | NA |
| NA | ENSMUSG00 | 73635 |
| NA | ENSMUSG00 | NA |
| NA | ENSMUSG00 | 433273 |
| NA | ENSMUSG00 | NA |
| NA | ENSMUSG00 | NA |
| NA | ENSMUSG00 | 100042074 |
| NA | ENSMUSG00 | 674597 |
| NA | ENSMUSG00 | 100043853 |
| NA | ENSMUSG00 | 100043283 |
| NA | ENSMUSG00 | 667309 |
| NA | ENSMUSG00 | 100040001 |
| NA | ENSMUSG00 | 330502 |
| NA | ENSMUSG00 | NA |
| NA | ENSMUSG00 | NA |
| NA | ENSMUSG00 | NA |
| NA | ENSMUSG00 | 676923 |
| NA | ENSMUSG00 | 100043797 |
| Significant | ENSMUSG00 | 668218 |
| NA | ENSMUSG00 | 100039827 |
| NA | ENSMUSG00 | NA |
| NA | ENSMUSG00 | 66381 |
| NA | ENSMUSG00 | 223648 |
| NA | ENSMUSG00 | 268301 |
| NA | ENSMUSG00 | 100041831 |
| NA | ENSMUSG00 | 68193 |
| NA | ENSMUSG00 | NA |
| NA | ENSMUSG00 | 239217 |
| NA | ENSMUSG00 | NA |
| NA | ENSMUSG00 | NA |
| NA | ENSMUSG00 | 545808 |
| NA | ENSMUSG00 | NA |
| NA | ENSMUSG00 | 433466 |
| NA | ENSMUSG00 | 629016 |
| NA | ENSMUSG00 | NA |
| NA | ENSMUSG00 | 226351 |

| | | |
|---|---|---|
| NA | ENSMUSG00 | 100417831 |
| NA | ENSMUSG00 | NA |
| NA | ENSMUSG00 | NA |
| NA | ENSMUSG00 | 223922 |
| NA | ENSMUSG00 | 100417829 |
| NA | ENSMUSG00 | 100416669 |
| NA | ENSMUSG00 | 383435 |
| NA | ENSMUSG00 | 102443350 |
| NA | ENSMUSG00 | 97908 |
| NA | ENSMUSG00 | 319149 |
| NA | ENSMUSG00 | NA |
| NA | ENSMUSG00 | 73729 |
| NA | ENSMUSG00 | 12047 |
| NA | ENSMUSG00 | 666869 |
| NA | ENSMUSG00 | NA |
| NA | ENSMUSG00 | NA |
| NA | ENSMUSG00 | 260423 |
| NA | ENSMUSG00 | NA |
| NA | ENSMUSG00 | 545645 |
| NA | ENSMUSG00 | 100038875 |
| NA | ENSMUSG00 | 727711 |
| NA | ENSMUSG00 | NA |
| NA | ENSMUSG00 | 668724 |
| NA | ENSMUSG00 | NA |
| NA | ENSMUSG00 | NA |
| NA | ENSMUSG00 | 100416214 |
| NA | ENSMUSG00 | NA |
| NA | ENSMUSG00 | NA |
| Significant | ENSMUSG00 | 666907 |
| NA | ENSMUSG00 | 666036 |
| NA | ENSMUSG00 | NA |
| NA | ENSMUSG00 | 75758 |
| NA | ENSMUSG00 | NA |
| NA | ENSMUSG00 | 12044 |
| NA | ENSMUSG00 | NA |
| NA | ENSMUSG00 | NA |
| NA | ENSMUSG00 | NA |
| NA | ENSMUSG00 | 20646 |
| NA | ENSMUSG00 | NA |
| Significant | ENSMUSG00 | 26904 |
| NA | ENSMUSG00 | 93724 |
| NA | ENSMUSG00 | 433387 |
| NA | ENSMUSG00 | 381650 |
| NA | ENSMUSG00 | 93723 |
| NA | ENSMUSG00 | 93700 |
| NA | ENSMUSG00 | 665574 |

| | | |
|---|---|---|
| NA | ENSMUSG00 | NA |
| Significant | ENSMUSG00 | 93706 |
| NA | ENSMUSG00 | 70579 |
| NA | ENSMUSG00 | 667759 |
| NA | ENSMUSG00 | NA |
| NA | ENSMUSG00 | 93713 |
| NA | ENSMUSG00 | NA |
| NA | ENSMUSG00 | 226654 |
| NA | ENSMUSG00 | 105734 |
| NA | ENSMUSG00 | 100046095 |
| NA | ENSMUSG00 | 93704 |
| NA | ENSMUSG00 | 380797 |
| NA | ENSMUSG00 | NA |
| NA | ENSMUSG00 | NA |
| NA | ENSMUSG00 | 115489679 |
| NA | ENSMUSG00 | 545767 |
| NA | ENSMUSG00 | NA |
| NA | ENSMUSG00 | NA |
| NA | ENSMUSG00 | NA |
| NA | ENSMUSG00 | 229898 |
| NA | ENSMUSG00 | 621155 |
| NA | ENSMUSG00 | 100043278 |
| NA | ENSMUSG00 | NA |
| NA | ENSMUSG00 | 100039840 |
| NA | ENSMUSG00 | 105180375 |
| NA | ENSMUSG00 | NA |
| NA | ENSMUSG00 | 71752 |
| NA | ENSMUSG00 | 66845 |
| NA | ENSMUSG00 | 433912 |
| NA | ENSMUSG00 | 106264 |
| NA | ENSMUSG00 | NA |
| NA | ENSMUSG00 | 67278 |
| NA | ENSMUSG00 | 100042008 |
| NA | ENSMUSG00 | 17527 |
| NA | ENSMUSG00 | 100043330 |
| NA | ENSMUSG00 | NA |
| NA | ENSMUSG00 | NA |
| NA | ENSMUSG00 | NA |
| NA | ENSMUSG00 | NA |
| NA | ENSMUSG00 | 118567384 |
| NA | ENSMUSG00 | NA |
| NA | ENSMUSG00 | NA |
| NA | ENSMUSG00 | 621306 |
| NA | ENSMUSG00 | NA |
| NA | ENSMUSG00 | NA |
| NA | ENSMUSG00 | 434127 |

| | | |
|---|---|---|
| NA | ENSMUSG00 | NA |
| NA | ENSMUSG00 | 668034 |
| NA | ENSMUSG00 | 668779 |
| NA | ENSMUSG00 | NA |
| NA | ENSMUSG00 | NA |
| NA | ENSMUSG00 | 15469 |
| NA | ENSMUSG00 | 233033 |
| NA | ENSMUSG00 | NA |
| NA | ENSMUSG00 | 18226 |
| NA | ENSMUSG00 | 14559 |
| NA | ENSMUSG00 | 640268 |
| NA | ENSMUSG00 | 102640920 |
| NA | ENSMUSG00 | NA |
| NA | ENSMUSG00 | 102638847 |
| NA | ENSMUSG00 | 50497 |
| NA | ENSMUSG00 | 67064 |
| NA | ENSMUSG00 | NA |
| NA | ENSMUSG00 | 100504116 |
| NA | ENSMUSG00 | 100418415 |
| NA | ENSMUSG00 | 76229 |
| NA | ENSMUSG00 | NA |
| NA | ENSMUSG00 | 207728 |
| NA | ENSMUSG00 | 14256 |
| NA | ENSMUSG00 | 14356 |
| NA | ENSMUSG00 | 108168395 |
| NA | ENSMUSG00 | 121811722 |
| NA | ENSMUSG00 | NA |
| NA | ENSMUSG00 | 378466 |
| NA | ENSMUSG00 | 100039229 |
| NA | ENSMUSG00 | NA |
| NA | ENSMUSG00 | NA |
| NA | ENSMUSG00 | NA |
| NA | ENSMUSG00 | NA |
| NA | ENSMUSG00 | 66387 |
| NA | ENSMUSG00 | 100040812 |
| NA | ENSMUSG00 | 100503185 |
| NA | ENSMUSG00 | NA |
| NA | ENSMUSG00 | NA |
| NA | ENSMUSG00 | NA |
| NA | ENSMUSG00 | NA |
| NA | ENSMUSG00 | NA |
| NA | ENSMUSG00 | NA |
| NA | ENSMUSG00 | 100039815 |
| NA | ENSMUSG00 | 14727 |
| NA | ENSMUSG00 | NA |
| NA | ENSMUSG00 | 68371 |

| | | |
|---|---|---|
| NA | ENSMUSG00 | 14728 |
| NA | ENSMUSG00 | 666584 |
| NA | ENSMUSG00 | 665155 |
| NA | ENSMUSG00 | 665071 |
| Significant | ENSMUSG00 | 102635315 |
| NA | ENSMUSG00 | 100126821 |
| NA | ENSMUSG00 | NA |
| NA | ENSMUSG00 | NA |
| NA | ENSMUSG00 | NA |
| NA | ENSMUSG00 | 100042388 |
| NA | ENSMUSG00 | NA |
| NA | ENSMUSG00 | NA |
| NA | ENSMUSG00 | 54339 |
| NA | ENSMUSG00 | 117934533 |
| NA | ENSMUSG00 | NA |
| NA | ENSMUSG00 | 71508 |
| NA | ENSMUSG00 | NA |
| NA | ENSMUSG00 | NA |
| NA | ENSMUSG00 | 100041509 |
| NA | ENSMUSG00 | NA |
| NA | ENSMUSG00 | 100113398 |
| NA | ENSMUSG00 | NA |
| NA | ENSMUSG00 | NA |
| NA | ENSMUSG00 | NA |
| NA | ENSMUSG00 | NA |
| NA | ENSMUSG00 | 100040745 |
| NA | ENSMUSG00 | 56214 |
| NA | ENSMUSG00 | NA |
| NA | ENSMUSG00 | 100504125 |
| NA | ENSMUSG00 | NA |
| NA | ENSMUSG00 | 634678 |
| NA | ENSMUSG00 | 319186 |
| NA | ENSMUSG00 | NA |
| NA | ENSMUSG00 | 319182 |
| NA | ENSMUSG00 | 100041240 |
| NA | ENSMUSG00 | NA |
| NA | ENSMUSG00 | 71844 |
| NA | ENSMUSG00 | NA |
| NA | ENSMUSG00 | NA |
| NA | ENSMUSG00 | 105245868 |
| NA | ENSMUSG00 | 100039214 |
| NA | ENSMUSG00 | 110599566 |
| NA | ENSMUSG00 | NA |
| NA | ENSMUSG00 | 18950 |
| NA | ENSMUSG00 | NA |
| NA | ENSMUSG00 | NA |

| | | |
|---|---|---|
| NA | ENSMUSG00 | 100039720 |
| NA | ENSMUSG00 | NA |
| NA | ENSMUSG00 | 638252 |
| NA | ENSMUSG00 | NA |
| NA | ENSMUSG00 | NA |
| NA | ENSMUSG00 | 66914 |
| NA | ENSMUSG00 | 18000 |
| NA | ENSMUSG00 | NA |
| NA | ENSMUSG00 | 223665 |
| NA | ENSMUSG00 | 57028 |
| NA | ENSMUSG00 | 18115 |
| NA | ENSMUSG00 | NA |
| NA | ENSMUSG00 | NA |
| NA | ENSMUSG00 | NA |
| NA | ENSMUSG00 | 67045 |
| NA | ENSMUSG00 | NA |
| NA | ENSMUSG00 | NA |
| NA | ENSMUSG00 | 667501 |
| NA | ENSMUSG00 | 320586 |
| NA | ENSMUSG00 | 625102 |
| NA | ENSMUSG00 | NA |
| NA | ENSMUSG00 | 622795 |
| NA | ENSMUSG00 | NA |
| NA | ENSMUSG00 | NA |
| NA | ENSMUSG00 | 100041621 |
| NA | ENSMUSG00 | 100041574 |
| NA | ENSMUSG00 | 100417722 |
| NA | ENSMUSG00 | NA |
| NA | ENSMUSG00 | NA |
| NA | ENSMUSG00 | NA |
| NA | ENSMUSG00 | NA |
| NA | ENSMUSG00 | 100504002 |
| NA | ENSMUSG00 | 667359 |
| NA | ENSMUSG00 | 631624 |
| NA | ENSMUSG00 | 19243 |
| NA | ENSMUSG00 | 115488794 |
| NA | ENSMUSG00 | 225372 |
| NA | ENSMUSG00 | NA |
| NA | ENSMUSG00 | 116621581 |
| NA | ENSMUSG00 | NA |
| NA | ENSMUSG00 | 668032 |
| NA | ENSMUSG00 | NA |
| NA | ENSMUSG00 | NA |
| NA | ENSMUSG00 | 66836 |
| NA | ENSMUSG00 | NA |
| NA | ENSMUSG00 | 414801 |

| | | |
|---|---|---|
| NA | ENSMUSG00 | 667490 |
| NA | ENSMUSG00 | NA |
| NA | ENSMUSG00 | 623242 |
| NA | ENSMUSG00 | 433193 |
| NA | ENSMUSG00 | NA |
| NA | ENSMUSG00 | 67706 |
| NA | ENSMUSG00 | NA |
